# Aerobic Adaptation and Metabolic Dynamics of *Propionibacterium freudenreichii* DSM 20271: Insights from Comparative Transcriptomics and Surfaceome Analysis

**DOI:** 10.1101/2024.04.30.591863

**Authors:** Iida Loivamaa, Annika Sillanpää, Paulina Deptula, Bhawani Chamlagain, Minnamari Edelmann, Petri Auvinen, Tuula A. Nyman, Kirsi Savijoki, Vieno Piironen, Pekka Varmanen

## Abstract

*Propionibacterium freudenreichii* (*PFR*) DSM 20271 is a bacterium known for its ability to thrive in diverse environments and to produce vitamin B12. Despite its anaerobic preference, recent studies have elucidated its ability to prosper in the presence of oxygen, prompting a deeper exploration of its physiology under aerobic conditions. Here, we investigated the response of DSM 20271 to aerobic growth by employing comparative transcriptomic and surfaceome analyses alongside metabolite profiling. Cultivation under controlled partial pressure of oxygen (pO2) conditions revealed significant increases in biomass formation and altered metabolite production, notably of B12 vitamin, pseudovitamin-B12, propionate and acetate, under aerobic conditions. Transcriptomic analysis identified differential expression of genes involved in lactate metabolism, TCA cycle, and electron transport chain, suggesting metabolic adjustments to aerobic environments. Moreover, surfaceome analysis unveiled growth environment-dependent changes in surface protein abundance, with implications for sensing and adaptation to atmospheric conditions. Supplementation experiments with key compounds highlighted the potential for enhancing aerobic growth, emphasizing the importance of iron and α-ketoglutarate availability. Furthermore, in liquid culture, FeSO_4_ supplementation led to increased heme production and reduced vitamin B12 production, highlighting the impact of oxygen and iron availability on the metabolic pathways. These findings deepen our understanding of *PFR*’s physiological responses to oxygen availability and offer insights for optimizing its growth in industrial applications.

**Importance:** The study of the response of *Propionibacterium freudenreichii* to aerobic growth is crucial for understanding how this bacterium adapts to different environments and produces essential compounds like vitamin B12. By investigating its physiological changes under aerobic conditions, we can gain insights into its metabolic adjustments and potential for enhanced growth. These findings not only deepen our understanding of *P. freudenreichii* responses to oxygen availability but also offer valuable information for optimizing its growth in industrial applications. This research sheds light on the adaptive mechanisms of this bacterium, providing a foundation for further exploration and potential applications in various fields.

## 1. Introduction

*Propionibacterium freudenreichii* (*PFR*) is a Gram-positive bacterium belonging to the Actinomycetota phylum. It is commonly found in dairy products, particularly Swiss-type cheeses, where it plays a crucial role in shaping the flavor compounds and texture characteristics of the cheese (1, 2). The bacterium metabolizes lactate to produce propionic acid and carbon dioxide, which leads to the formation of gas bubbles responsible for the characteristic eyes in Swiss-type cheeses (3). Besides its role in cheese production, *PFR* has garnered significant attention due to its potential health benefits, including its immunomodulatory properties and the production of vitamin B12 (hereafter B12) (2, 4, 5).

B12, also known as cobalamin, is an essential nutrient vital for various physiological processes in humans (6). Unlike many other organisms, humans cannot synthesize B12 and must obtain it from dietary sources or microbial production (7). *PFR* stands out as one of the microbial species capable of producing this essential vitamin. Its history of safe use in fermented foods and ability to produce vitamin B12 have attracted considerable interest from the food and pharmaceutical industries (8, 9). Given the prevalence of B12 deficiency, particularly among individuals following vegetarian or vegan diets, the microbial production of vitamin B12 by *PFR* offers a promising solution to address this nutritional deficiency (10).

Notably, the synthesis of cobalamin, as well as the human inactive pseudovitamin, by *PFR* exhibits intriguing dependencies on environmental conditions, particularly oxygen availability. While the first steps of cobalamin synthesis follow the anaerobic pathway where cobalt is inserted at an early stage (11), the biosynthesis of the lower ligand, 5,6-dimethylbenzimidazole (DMBI), requires oxygen. Absence of oxygen during biosynthesis leads to the production of pseudovitamin B12 (hereafter pseudo-B12) with adenine as the lower ligand (12).

*PFR* DSM 20271, the type strain of the species, has garnered significant attention due to its ability to produce the active form of B12 in various plant food matrices and under different applications (13–15). Furthermore, the response of DSM 20271 to oxygen and its ability to grow under aerobic conditions have attracted scientific interest. Recent studies have revealed that certain strains of *PFR*, including DSM 20271, while traditionally considered anaerobes, exhibit tolerance to oxygen. They can grow to high cell densities under mildly aerated conditions, which allow the consumption of oxygen and thereby keep levels of dissolved oxygen below the detection limit (16). The adaptation of DSM 20271 to these microaerobic conditions was shown to involve complex metabolic adjustments, affecting various cellular processes, including energy metabolism, redox balance, and gene expression (16, 17). *PFR* under aerobic conditions, including detectable oxygen concentrations, remains to be studied.

The genome sequence of DSM 20271 (18) indicates that the strain possesses enzymatic systems, such as superoxide dismutase and heme-containing catalase, which likely play crucial roles in protecting the bacterium against oxidative stress caused by oxygen exposure. Furthermore, like other *PFR* strains (19, 20), DSM 20271 is equipped with the genes required for aerobic respiration and for the complete pathway for heme synthesis. As heme iron is more easily absorbed and has a higher bioavailability than other forms of iron (21), heme biosynthesis through bioprocessing of food material potentially increases the nutritional value. Heme and cobalamin synthesis are interconnected as both pathways begin with the formation of uroporphyrinogen III precursor (22).

In this study, we cultivated the DSM 20271 strain in bioreactors under control of the partial pressure of oxygen (pO2), keeping it at approximately 20% by sparging with pure oxygen/air as well as under anaerobic conditions by nitrogen gas sparging. Employing a comparative transcriptomic approach alongside comparative surfaceome analyses and targeted metabolite profiling, we sought to elucidate the bacterium’s response to oxygen. The findings of this study contribute to a deeper understanding of the physiological responses of *PFR* to oxygen availability and offer valuable insights for optimizing its growth. Furthermore, the identification of key compounds and metabolites enhancing the aerobic growth and colony-forming ability of *PFR* under aerobic conditions can have significant implications for industrial applications.

## 2. Results and Discussion

In the present study, we used pH-controlled bioreactor cultivations to investigate the cellular responses of DSM 20271 during growth under aerobic and anaerobic conditions. Our focus was on monitoring the central growth characteristics, the production of B12 and pseudo-B12 vitamins, and the major metabolic end products **(Fig. 1A)** at specific time points of growth and conditions **(Fig. 1B)**. These findings were complemented with transcriptomic analyses and validated with label-free quantitative proteomics on cell surface proteins at the indicated timepoints **(Fig. 1B)**. Additional follow-up studies were conducted to confirm the most relevant metabolic processes.

**Figure 1.**
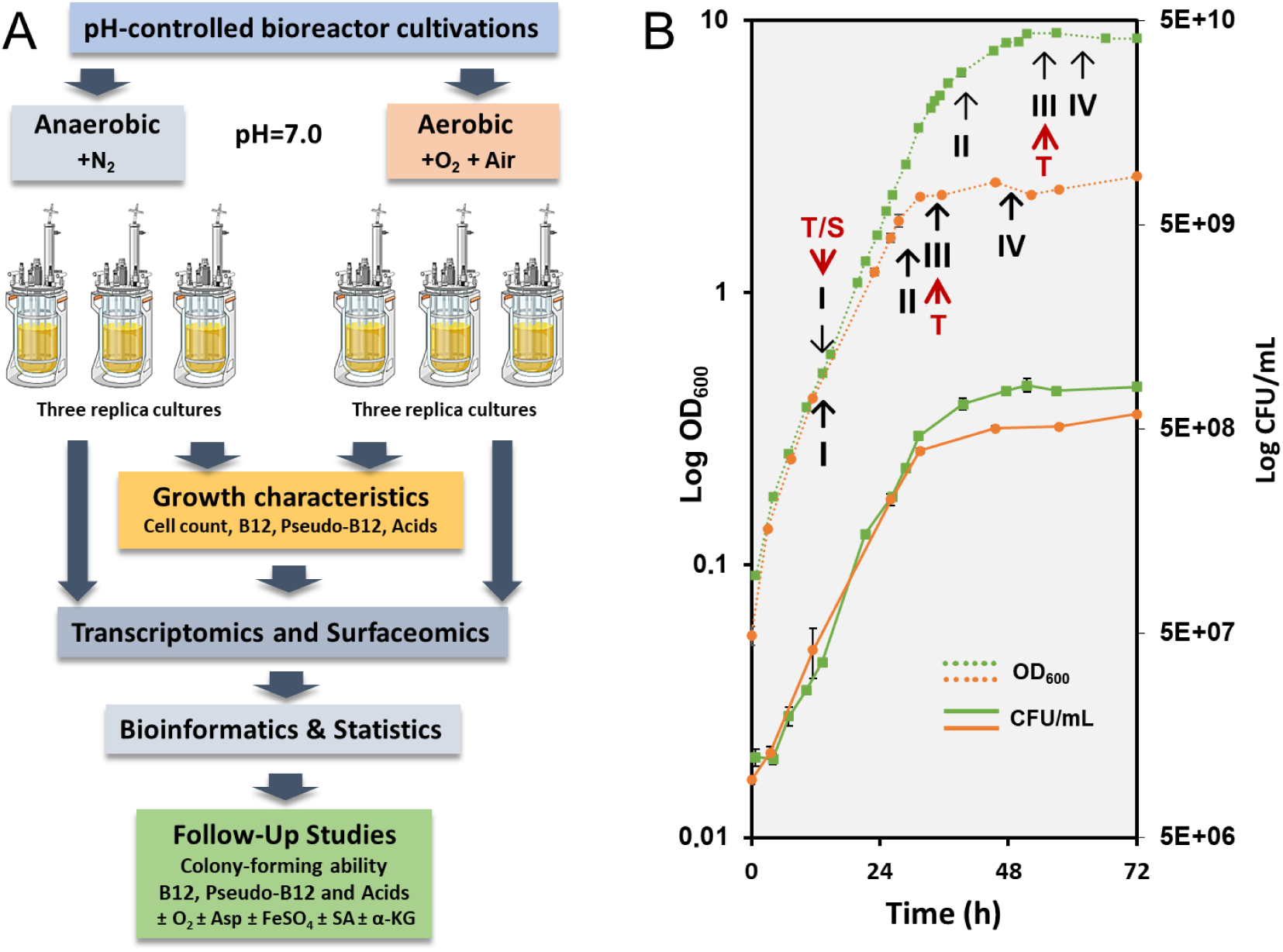
(**A**) A workflow illustrating an overview of experiments including bioreactor cultivations and parameters studied as well as follow-up growth experiments conducted in supplemented media. Asp, aspartate; SA, succinate; α-KG, α-ketoglutarate. **(B)** Cell densities (dotted line) and cell viability (solid line) in bioreactor-conducted experiments under aerobic (green) and anaerobic (orange) growth conditions. Sampling points for OD_600_ measurement and determining colony-forming units are marked with circles (anaerobic) and squares (aerobic). For the analysis of key metabolite production dynamics and transcriptomics over time, samples were taken at four growth stages: mid-logarithmic (I), late logarithmic (II), early stationary (III), and late stationary (IV) phases. RNA sequencing was performed on samples I and III. Arrows in red indicate the timepoint/growth stage of *PFR* at which the cells were withdrawn for transcriptomics (T) and surfaceomics (S).

### 2.1 Oxygen affects lactate metabolism, biomass, and formation of organic acids, B12, and pseudo-B12

#### Formation of biomass

To initiate the characterization of the response of DSM 20271 to aerobic growth, the strain was cultivated in bioreactors at a constant pO2 level of 20% and compared the growth characteristics to control cultivation under anaerobic control conditions (pO2 level below the detection limit). A pO2 level of 20% was selected based on preliminary experiments indicating that higher oxygen concentrations no longer increased cell density (Loivamaa et al., unpublished data). A significant increase in biomass formation was observed under aerobic conditions with a pO2 level of 20% compared to anaerobic conditions. Specifically, the cell densities determined at 600 nm and viable cell counts on agar (CFU/mL) were 3.2 and 1.4 times higher, respectively, under the maintained aerobic atmosphere over the 72-hour-cultivation **(Fig. 1B)**.

#### Secretion of organic acids

Lactate was found to be consumed faster under anaerobic conditions but was depleted from the media under both conditions in the early stationary phase sample (III) **(Fig. 2A)**. Under anaerobic conditions, the levels of propionate and acetate in the growth medium increased over time, especially during logarithmic growth **(Fig. 2B and 2C)**. In contrast, under aerobic conditions, only low levels of propionate were detected in logarithmic phase samples, with levels dropping below the limit of detection in stationary phase samples. Acetate levels followed a similar pattern under aerobic conditions, being fully depleted from the medium after an initial increase **(Fig. 2C)**. During logarithmic growth, the pyruvate levels in the media were higher under aerobic conditions but decreased to low levels in stationary phase samples **(Fig. 2D)**, whereas succinate was only detected in growth media in samples from anaerobic conditions **(Fig. 2E)**. Pyruvate in culture supernatants, at concentrations reflecting its intracellular levels in *PFR*, has been reported previously (23), but to the best of our knowledge, there are no reports of the effect of atmospheric conditions. As the elevated growth under aerobic conditions is accompanied by diminished lactate consumption and, on the other hand, increased pyruvate excretion, it is tempting to assume that ATP production is maintained by upregulation of the later stages of the aerobic respiration pathway. The accumulation of succinate during adaptation to anaerobic or oxygen-limiting conditions has been previously observed in *Mycobacterium tuberculosis* (24).

**Figure 2.**
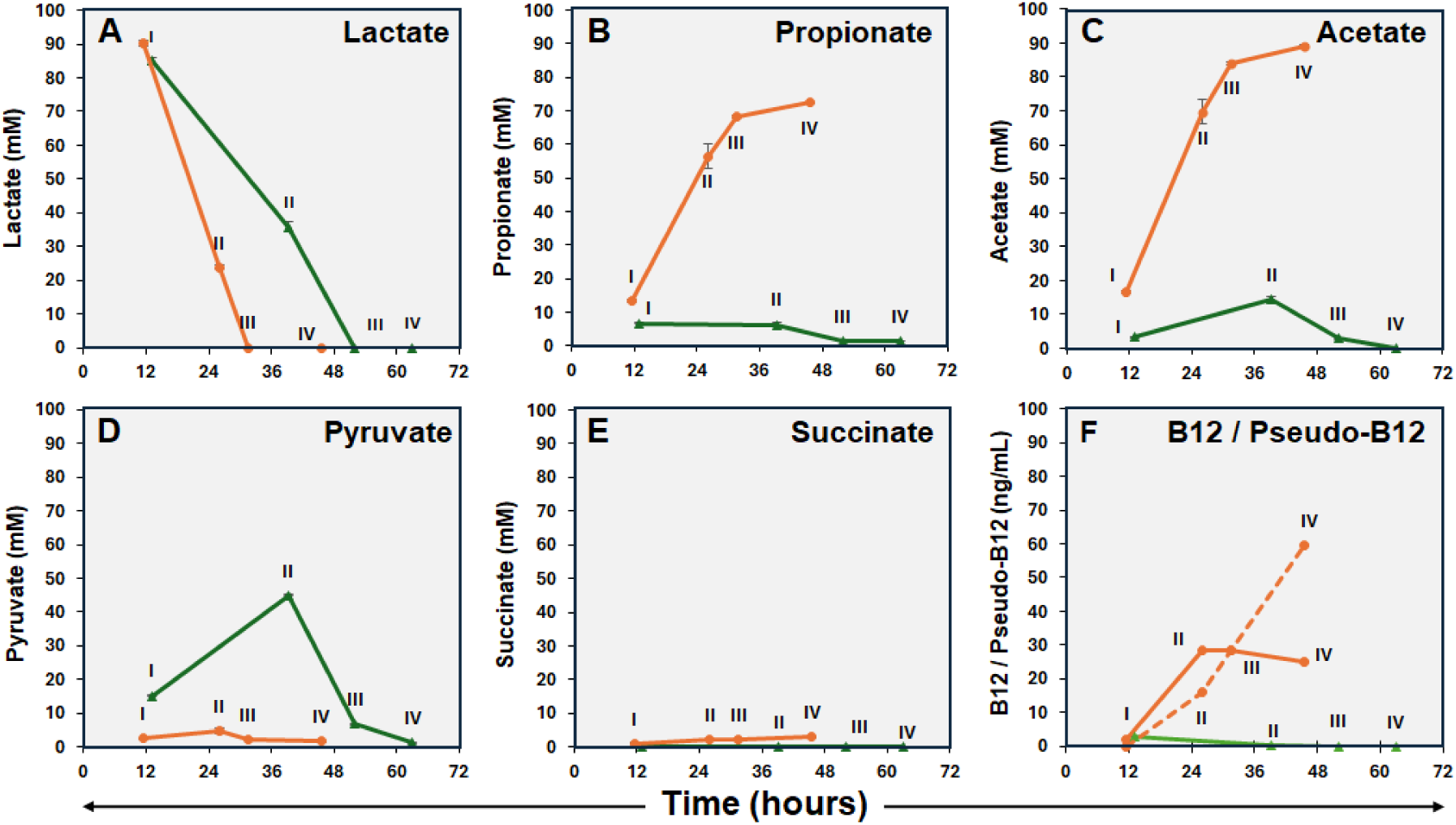
Lactate utilization (A), excreted propionate (B), acetate (C), pyruvate (D), succinate (E) and intracellular vitamin B12 and pseudovitamin (F; vitamin B12 - solid line, pseudovitamin – dashed line) of PFR DSM 20271 during cultivation under aerobic (green) and anaerobic (orange) fermentations performed in bioreactors. N = 3, except pseudovitamin samples III and IV: n=2. No pseudovitamin was detected under aerobic growth conditions.

#### Synthesis of B12 and pseudo-B12

B12 and pseudo-B12 levels were analyzed in cellular and supernatant (media) samples. However, no B12 or pseudo-B12 was detected in the media of either aerobic or anaerobic cultivations. Under aerobic conditions, the concentration of B12 in the cells was very low at each sampling point and below the detection limit at sampling points II-IV **(Fig. 2F)**. Under anaerobic conditions, B12 reached 28.5 ng/mL as fermentation progressed, while the concentration of pseudo-B12 increased as the fermentation process approached completion and reached 59.8 ng/mL. **(Fig. 2F)**. It is noteworthy that cultivation of this strain in our bioreactors under identical conditions but with no gas sparging, thus resembling microaerobic conditions, resulted in superior B12 production. The levels were 1.7 to 4.9 times higher compared to anaerobic conditions, along with a distinct metabolite production pattern **(Fig. S1)**.

In a previous study (16), the DSM 20271 strain was cultivated in bioreactors under conditions with an oxygen supply at a level that allowed its concentration to drop below the detection limit in the culture through consumption by cellular activity. Like those microaerobic conditions, aerobic conditions used here were found to increase biomass production compared to anaerobic conditions. However, the metabolite production profile under aerobic conditions seems to differ from those reported under microaerobic conditions. Firstly, lower oxygen concentrations promoted propionate production during the lactate consumption phase followed by its oxidation to acetate (16) **(Fig. S1)**. Under the more aerobic conditions used here, the consumption of lactate from the media was not accompanied by propionate accumulation. Furthermore, the B12 production appears to be drastically affected by increasing oxygen concentration, as microaerobic conditions increased B12 production compared to anaerobic conditions (17) **(Fig. S1)**, which is opposite to what was observed here under aerobic conditions **(Fig. 2F)**. The ratio of B12 to pseudovitamin in microaerobic conditions was not analyzed in previous studies due to the use of a microbiological assay that does not distinguish between them (17). Here, we observed that the active form of B12 is produced under microaerobic conditions, whereas very low levels of either B12 or pseudo-b12 are produced under aerobic conditions **(Fig. 1; Fig. S1).** Thus, the typical features associated with *PFR* as an efficient producer of B12 and short-chain fatty acids, such as propionate and acetate, do not apply under aerobic growth conditions.

### 2.2 Transcriptomics of DSM 20271 under anaerobic and aerobic conditions

In this study, we used gene expression data to uncover genes important for growth in both aerobic and anaerobic atmospheres, with focus on genes contributing to the growth of *PFR* under aerobic conditions as neither the physiology of this species in aerobic environments nor gene expression has yet been extensively studied (16). The gene expression of strain DSM 20271 grown under aerobic and anaerobic conditions was compared by isolating the total RNA from cells at the mid-logarithmic (sampling point I) and stationary phases (sampling point III) of growth for RNA sequencing.

To specifically identify differences between growth phases and conditions, pairwise differential expression analyses were utilized. A total of 1376 genes were found to be differentially expressed at sampling point I, with an adjusted p-value (padj) of < 0.05. Among these, 713 genes were downregulated, and 663 genes were upregulated under aerobic conditions. Additionally, 906 genes showed differential expressions at sampling point III. Out of the 906 genes, 388 genes were downregulated, and 518 genes were upregulated under aerobic conditions **(Fig. 3)**. A list of differentially expressed genes, annotation results, and expression levels of under anaerobic and aerobic atmospheres at sampling points I and III are shown in **Table S1**.

**Figure 3.**
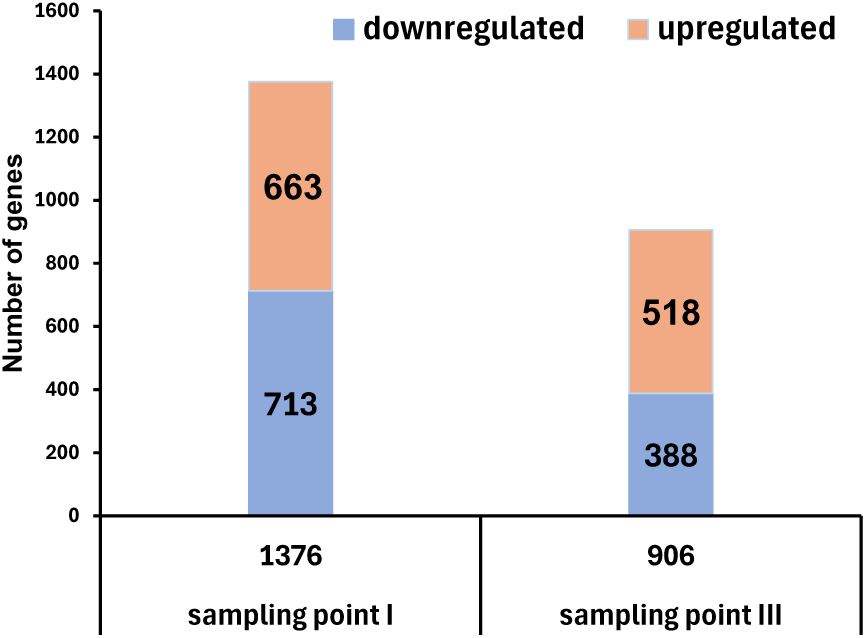
The number of genes exhibiting differential expression in *PFR* DSM 20271 cultivated under aerobic compared to anaerobic atmospheres. Pairwise analyses of differential expression were conducted to identify variances between growth environments. During sampling point I, 1376 genes showed significant differential expression (adjusted p-value < 0.05), with 713 genes downregulated and 663 genes upregulated. Similarly, at sampling point III, 906 genes displayed differential expression, with 388 genes downregulated and 518 genes upregulated.

#### Gene ontology (GO) enrichments

GO enrichments were conducted to obtain a comprehensive understanding of the functional groups represented by the differentially expressed genes (DEGs). The putative functions of the DEGs were further classified into various categories using the gene ontology (GO) framework. At sampling point I, the DEGs were organized into nine distinct GO categories, exhibiting statistical significance (p < 0.05). This categorization involved a comparison between the aerobic and anaerobic growth conditions. These categories encompassed transmembrane transport, transmembrane transporter activity, ATPase activity, zinc ion binding, magnesium ion binding, 4 iron, 4 sulfur cluster binding, ‘de novo’ IMP biosynthetic process, transferase activity, transferring glycosyl groups, and carbohydrate metabolic process **(Fig. 4A)**. Similarly, at sampling point III, the DEGs were categorized into 11 GO categories, with statistical significance observed (p < 0.05). These categories included structural constituent of ribosome, translation, rRNA binding, ribosome, plasma membrane, tRNA binding, glycolytic process, DNA recombination, peptidoglycan biosynthetic process, cytoplasm, and transferase activity; transferring acyl groups **(Fig. 4B**).

**Figure 4.**
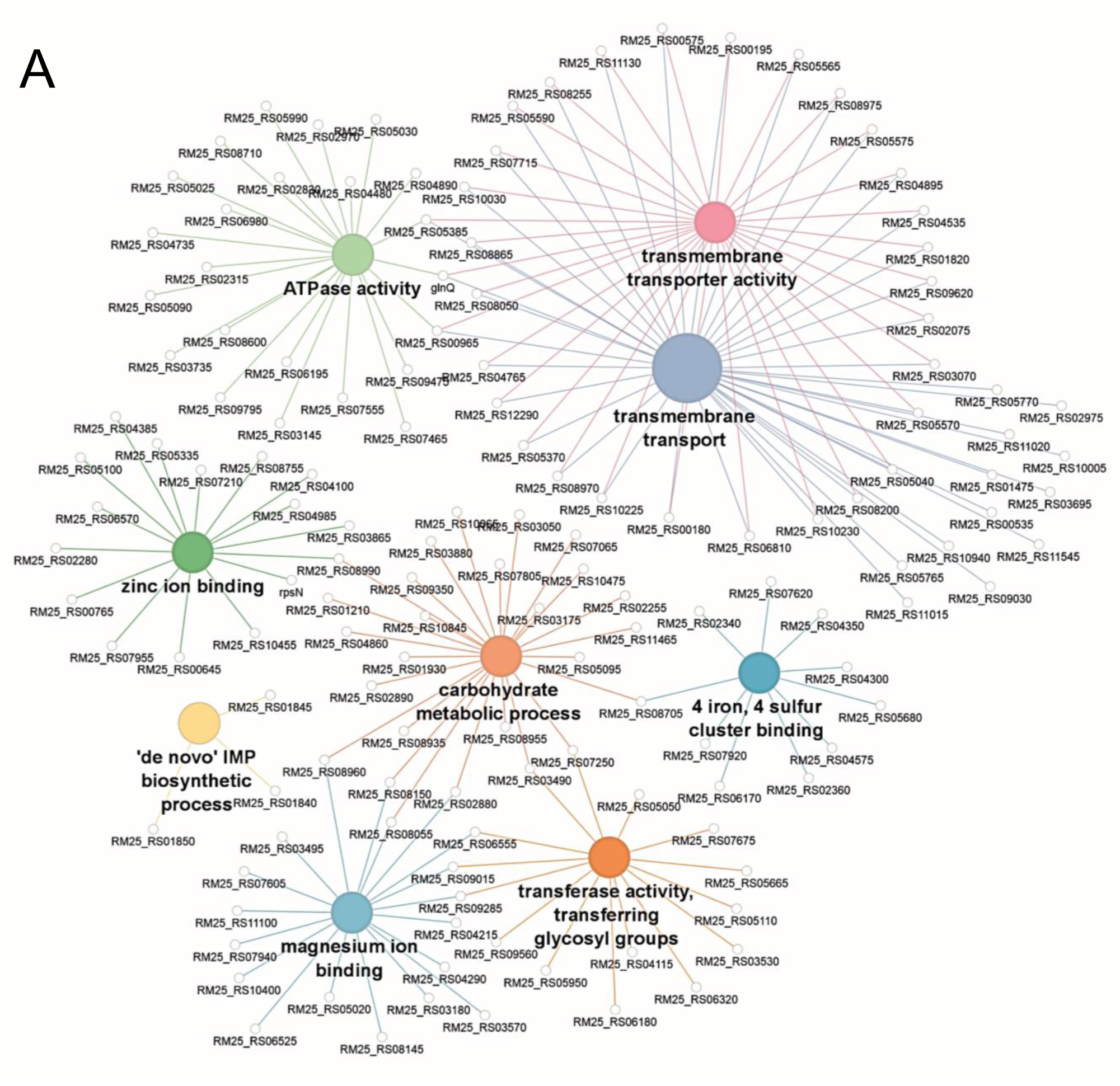
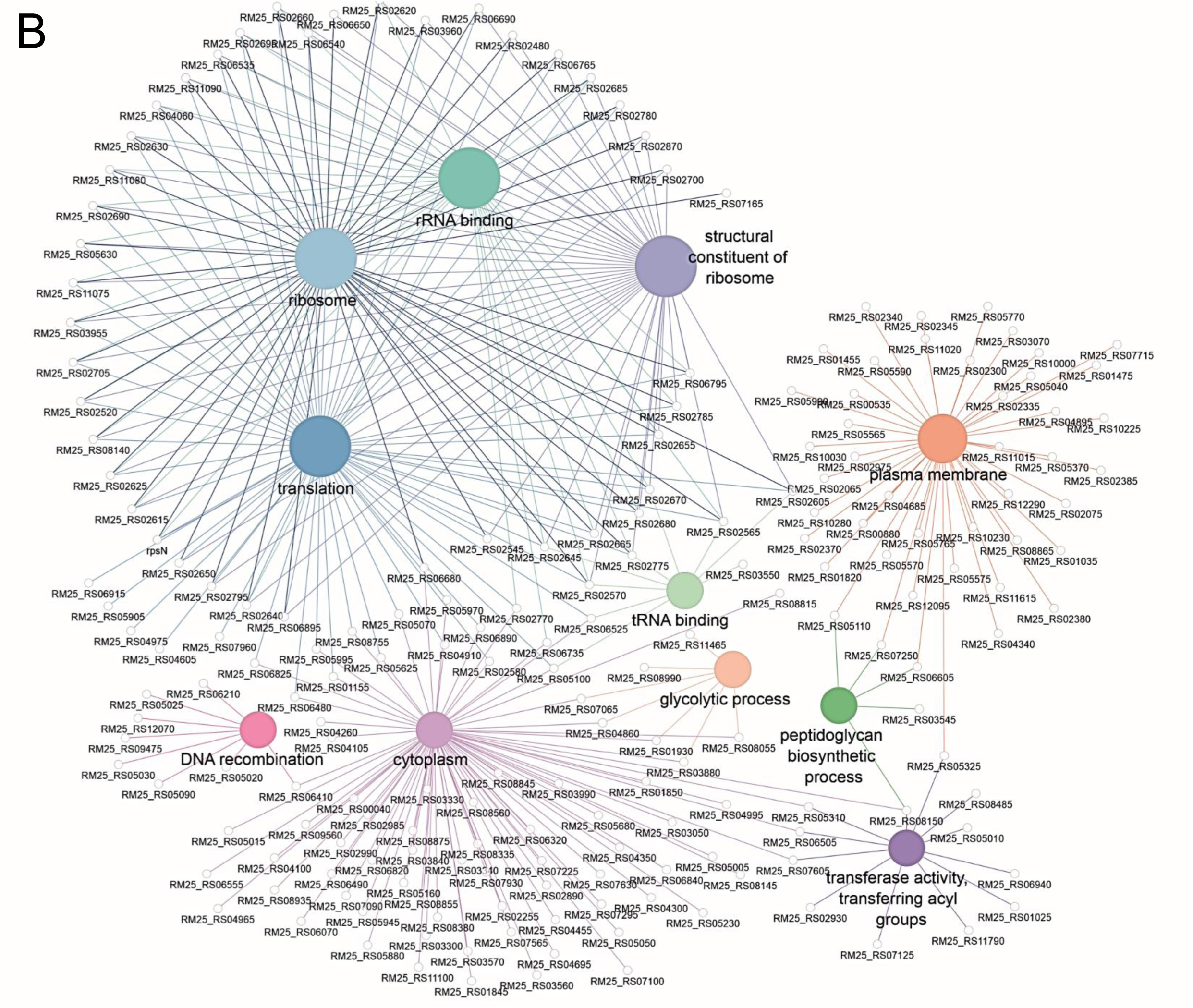
Gene Ontology (GO) classification of differentially expressed genes (DEGs) at sampling point I **(A**) and sampling point III **(B)** was conducted. At sampling point I, DEGs’ putative functions were categorized into nine distinct GO categories, all exhibiting statistical significance (p < 0.05). At sampling point III, DEGs were distributed across 11 distinct GO categories with statistical significance (p < 0.05). The size of the spheres corresponds to the number of genes within each gene ontology category. A straight line emerging from each sphere, representing a category, is connected to a gene. A single gene may be associated with multiple categories, depicted as multiple connections from one gene to several spheres.

#### Lactate metabolism

Analyses of organic acids from bioreactor cultivations revealed that lactate consumption was more rapid under anaerobic conditions compared to the aerobic atmosphere (**Figure 2 A**). The gene RM25_RS08745 encodes L-lactate permease and is the first gene within a predicted four-gene operon (RM25_RS08730 ̶ RM25_RS08745). This operon also contains genes encoding lactate utilization proteins A, B, and C. These proteins exhibit homology to the *lldEFG*-encoded proteins found in *Shewanella oneidensis* (25) in which together with lactate dehydrogenase (LDH), they form an enzyme complex responsible for catalyzing the oxidation of both D- and L-lactate stereoisomers to pyruvate (25). In our study, the expression levels of these operon genes were upregulated under anaerobic conditions in both sampling points (I and III). Specifically, in the III samples, the fold difference in expression ranged from 2.4 to 3.9, which correlates with the observed lactate consumption results. Regardless of the atmospheric conditions, the lactate utilization operon was downregulated upon shifting to the stationary phase.

The DSM20271 strain possesses two paralogs of LDH, namely RM25_RS04460 and RM25_RS05095, which encode NAD-dependent L-lactate dehydrogenases. Additionally, it carries one FAD-dependent D-lactate dehydrogenase encoded by RM25_RS01945. Under anaerobic conditions in the later sampling points (III), the expression level of D-lactate dehydrogenase was found to be 3.8-fold higher compared to aerobic conditions. These findings suggest that lactate metabolism is more active when oxygen is limited (anaerobic conditions). Multiple pathways are available for the conversion of pyruvate into acetyl-CoA. Two enzymes, the pyruvate dehydrogenase complex and pyruvate ferredoxin oxidoreductase, can perform this conversion. In this study, the gene cluster encoding pyruvate dehydrogenase (*pdh*), specifically genes RM25_RS01015, RM25_RS01020, and RM25_RS01025, showed significantly higher expression levels (2.0-8.6-fold) under aerobic conditions (at both sampling points, I and III). On the other hand, one of the two paralogs encoding pyruvate ferredoxin oxidoreductase, RM25_RS00920, only displayed higher expression (2.8-fold) under anaerobic conditions in later samples. These findings indicate that different pathways for pyruvate conversion are utilized under aerobic and anaerobic conditions.

#### Propionate metabolism

The gene cluster RM25_RS08835 to RM25_RS08850 encodes the key enzyme, methylmalonyl-CoA carboxytransferase (MMC), of the propionate-forming Wood-Werkman cycle. Interestingly, this gene cluster was found to be constitutively expressed under the conditions investigated in this study. This finding is consistent with a previous study by Brzuszkiewicz et al. (26), which also observed constitutive expression of the MMC genes at both logarithmic and stationary growth phases in closely related bacterium *Cutibacterium acnes*. Additionally, it aligns with the study conducted by Dank et al. (16), where proteins involved in the Wood-Werkman cycle were found to be as abundant in propionate consumption as in lactate consumption phases in *PFR*. The TCA cycle utilizes acetyl-CoA and converts NAD+ to NADH as part of its metabolic reactions. Certain genes encoding enzymes involved in the TCA cycle, such as citrate synthase (RM25_RS11475), α-ketoglutarate decarboxylase (RM25_RS05675), and isocitrate dehydrogenase (RM25_RS03180), exhibited higher expression levels under aerobic conditions in sampling point III. These results indicate that aerobic conditions promote increased expression of these TCA cycle enzymes, which contribute to the utilization of acetyl-CoA and the generation of NADH. The fermentation of pyruvate by propionibacteria typically produces propionic and acetic acids, as well as significant amounts of other products, such as succinate (27).

#### Expression of succinate dehydrogenase and other respiratory complexes

The primary function of the succinate dehydrogenase (Sdh) complex in bacteria is to catalyze the oxidation of succinate to fumarate while transferring electrons to the electron transport chain. The genome of DSM20271 harbors two gene clusters encoding the subunits of the Sdh complexes. These clusters consist of three genes each, namely RM25_RS06150, RM25_RS06155, RM25_RS06160, and RM25_RS06655, RM25_RS06660, RM25_RS06665. Here, the *sdh* gene clusters exhibited distinct patterns of gene expression under different atmospheric conditions. Specifically, the cluster RM25_RS06150 to RM25_RS06160 demonstrated a significantly higher expression level (2.8-3.4-fold increase) during anaerobic conditions (I and III samples). Conversely, at sampling point III, the RM25_RS06655 and RM25_RS06660 genes of the other cluster exhibited significantly higher expression levels (2.3 and 2.8-fold increase) under aerobic conditions. Based on the different expression patterns observed, we propose that the gene cluster RM25_RS06655 to RM25_RS06665 is responsible for encoding Sdh, catalyzing the conversion of succinate to fumarate, and is involved in both the TCA cycle and aerobic respiration. In contrast, the gene cluster RM25_RS06150 to RM25_RS06160 is responsible for encoding fumarate reductase, which reduces fumarate to succinate during anaerobic respiration. This is also consistent with the accumulation of succinate in the medium under anaerobic conditions and its absence from the medium under aerobic conditions. These two sets of succinate dehydrogenase/fumarate reductase complexes have been reported earlier in various studies on *Propionibacterium* strains (16, 19, 28), providing further support to our findings. Furthermore, it has been suggested, but not yet explored, that in *Acidipropionibacterium acidipropionici* the other succinate dehydrogenase complex could function in anaerobic fermentation as fumarate reductase, and another complex in aerobic fermentation as succinate dehydrogenase (28).

Expression of the NADH dehydrogenase complex, encoded by the gene cluster RM25_RS02335 to RM25_RS02400, and the expression of the cytochrome-bd oxidase, encoded by genes RM25_RS00880 to RM25_RS12185, were not significantly influenced by atmospheric conditions. However, the gene cluster RM25_RS05560 to RM25_RS05595, which encodes the ATP synthase enzyme complex, demonstrated higher expression levels under aerobic conditions, particularly in the later sampling point (III). This indicates that ATP synthesis is more efficient in the presence of oxygen which correlates with the results that ATP yields have been higher under aerobic growth conditions (Loivamaa et al., unpublished data). These findings align with the conducted study by Dank et al. (16), where they observed that the respiratory pathway remains consistently active under both anaerobic and microaerobic conditions.

The expression of the NADH dehydrogenase complex, the cytochrome-bd oxidase, and the ATP synthase complex were found to be impacted differently by the growth phase. The gene cluster of the NADH dehydrogenase complex (RM25_RS02335 to RM25_RS02400) and the ATP synthase enzyme complex (RM25_RS05560 to RM25_RS05595) showed downregulation at the onset of the stationary phase. In contrast, the cluster of cytochrome-bd oxidase (RM25_RS00880 to RM25_RS12185) was either constitutively expressed under both atmospheric conditions and, in the case of RM25_RS00880, exhibited a 2.5-fold increase in expression at the stationary phase under anaerobic conditions.

#### Synthesis of B12 and heme

B12 and heme are tetrapyrrols that play crucial roles in various biological processes. Vitamin B12 serves as an essential coenzyme for enzymatic reactions, particularly in the metabolic pathways involved in amino acid and nucleotide synthesis (29). Heme, however, is involved in important cellular oxidative metabolic pathways, such as oxygen transportation, response to oxidative stress, and oxidative phosphorylation. In our study, we observed a significant effect of atmospheric conditions on cobalamin production by DSM20271 in bioreactor cultivations. The cobalamin biosynthesis genes in *PFR* are mainly located in three clusters: RM25_RS02055 to RM25_RS02075, RM25_RS03590 to RM25_RS03625, and RM25_RS04745 to RM25_RS04770. Cluster RM25_RS02055 to RM25_RS02075 forms the locus for cobalt transport, while RM25_RS03590 to RM25_RS03625 and RM25_RS04745 to RM25_RS04770 encode the assembly and activation of the corrin ring (19, 30).

Additionally, there are single genes, such as RM25_RS02940 (*bluB*-*cobT2*), RM25_RS03660 (*cobQ2*), and RM25_RS05545 (*cobA*), involved in cobalamin biosynthesis. Under aerobic conditions during later sampling point (III), the genes RM25_RS02070 (*cbiN*) and RM25_RS02075 (*cbiM*) encoding cobalt transport proteins showed significant upregulation (4.7- and 3.2-fold, respectively), while the expression of all the other B12 biosynthetic genes was not significantly affected by the atmospheric conditions. The regulatory mechanism of the B12 biosynthetic pathway is known to involve cobalamin riboswitches, which were predicted as located upstream of RM25_RS03590 (*cbiL*), RM25_RS04770 (*cobD*) and RM25_RS02075 (*cbiM*) (31, 32). Cobalamin riboswitch controls gene expression through transcriptional and/or translational modifications (33, 34). Notably, the function of riboswitch upstream of *cbiM* was recently characterized in *PFR* strain P.UF1, which revealed control at the transcriptional level (31). Our results are in line with this and indicated significantly differential transcription of the *cbiM* and *cbiN* genes under the conditions used. On the other hand, differential B12 production and the constitutive expression of other B12 genes under studied conditions suggest regulation by means other than at the transcriptional level.

5-Aminolevulinic acid (ALA) is a precursor metabolite involved in the biosynthesis of tetrapyrrole compounds, which in *PFR* is synthesized via the C5 pathway (35–37). In this pathway, ALA is synthesized from glutamate through the coordinated actions of glutamyl-tRNA synthetase (encoded by *gltX*), glutamyl-tRNA reductase (encoded by *hemA*), and glutamate semialdehyde aminotransferase (HemL). HemB converts two ALA molecules into porphobilinogen (PBG), which is further polymerized into hydroxymethylbilane (HMB) by HemC. This HMB is then transformed into uroporphyrinogen III, a crucial intermediate in heme and B12 synthesis, by uroporphyrinogen III synthase. Our study identified significant upregulation (2.2–3.2-fold) of ALA and uroporphyrinogen III synthesis genes, specifically *hemA* (RM25_RS08900), *hemB* (RM25_RS08870), and *hemL2* (RM25_RS09040), under aerobic conditions during later sampling point (III). However, the expression of genes involved in later steps of heme biosynthesis, such as RM25_RS09440 encoding putative uroporphyrinogen III synthase, *hemH* (RM25_RS08875), *hemY* (RM25_RS08880), and *hemE* (RM25_RS12355), did not show significant differential expression in response to atmospheric conditions. Furthermore, the expression of RM25_RS09440 was found to be affected by the growth stage and downregulated during the shift to the stationary phase under both atmospheric conditions.

#### Transport of iron and α-ketoglutarate are increased under aerobic conditions

The results of the GO enrichment analysis revealed a significant impact on “transmembrane transport.” To delve deeper into this observation, genes from DSM 20271 were filtered based on annotations related to “transport,” “import,” or “export,” resulting in 208 genes potentially encoding transport functions. Within this pool of 208 genes, nine demonstrated more than a two-fold upregulation under aerobic conditions at sampling point I, and 24 exhibited more than a two-fold upregulation under aerobic conditions at sampling point III **(Table 1)**. Conversely, among these 208 genes, 74 displayed more than a two-fold upregulation under anaerobic conditions compared to aerobic conditions at sampling point I. Subsequently, at the later sampling point (III), 34 genes were observed to be upregulated under anaerobic conditions **(Table S2)**.

**Table 1.** List of transport-, import-, or export-associated genes expressed more than 2-fold in DSM 20271 at sampling point I (nine genes) and sampling point III (24 genes) under aerobic conditions compared to anaerobic conditions.

| Locus tag | Protein name | Fold-change<sup>a</sup> |
| --- | --- | --- |
| **Sampling point I** |  |  |
| RM25_RS07080 | ABC transporter, ATP-binding protein | 5.02 |
| RM25_RS01195 | Putative iron-siderophore transport system, periplasmic binding protein | 3.67 |
| RM25_RS02225 | Putative cation-transporting P-type ATPase | 3.45 |
| RM25_RS09250 | Ferrous iron transport protein A | 2.70 |
| RM25_RS12135 | MscS transporter, small conductance mechanosensitive ion channel | 2.67 |
| RM25_RS01240 | Glycerol-3-phosphate transporter | 2.60 |
| RM25_RS04570 | Alpha-ketoglutarate transporter, MFS super | 2.23 |
| RM25_RS09245 | Ferrous iron transport protein B | 2.18 |
| RM25_RS08970 | MFS transporter, sugar porter family protein | 2.16 |
| **Sampling point III** |  |  |
| RM25_RS02070 | Cobalt transport protein CbiN | 4.66 |
| RM25_RS09620 | D-serine/D-alanine/glycine transporter | 4.27 |
| RM25_RS01240 | Glycerol-3-phosphate transporter | 4.14 |
| RM25_RS09250 | Ferrous iron transport protein A | 4.07 |
| RM25_RS05305 | Putative sulfonate/nitrate/taurine transport system substrate-binding | 3.92 |
| RM25_RS03735 | ATP-binding cassette protein, ChvD family | 3.57 |
| RM25_RS07080 | ABC transporter, ATP-binding protein | 3.22 |
| RM25_RS02075 | Cobalt transport protein CbiM | 3.21 |
| RM25_RS02225 | Putative cation-transporting P-type ATPase | 3.17 |
| RM25_RS08710 | ATP binding protein of ABC transporter for sugars | 2.88 |
| RM25_RS03575 | LAO/AO transport system ATPase | 2.72 |
| RM25_RS08050 | Putative cationic amino acid transport integral membrane protein roce | 2.62 |
| RM25_RS02025 | Low affinity inorganic phosphate transporter membrane protein pitA | 2.61 |
| RM25_RS05380 | High-affinity branched-chain amino acid transport ATP-binding protein livG | 2.53 |
| RM25_RS05770 | ABC-type dipeptide transport system, periplasmic componen | 2.49 |
| RM25_RS04530 | Trk family potassium (K<sup>+</sup>) transporter, NAD<sup>+</sup> binding protein | 2.46 |
| RM25_RS01470 | Methionine import ATP-binding protein MetN 2 | 2.3 |
| RM25_RS01195 | Putative iron-siderophore transport system, periplasmic binding protein | 2.29 |
| RM25_RS01475 | ABC D-methionine transporter, permease component | 2.26 |
| RM25_RS11530 | ABC transporter domain-containing permease component | 2.19 |
| RM25_RS00345 | ABC-2 type transport system integral membrane protein | 2.16 |
| RM25_RS10625 | Putative potassium transport system protein kup | 2.09 |
| RM25_RS00035 | ABC-type antimicrobial peptide transport system | 2.02 |
| RM25_RS00535 | Putative polar amino acid ABC transporter substrate-binding protein | 2.01 |

a) The fold-changes were calculated by exponentiating the log2Fold values, which were derived from the differential expression analysis performed using DESeq2.

Among the transport genes that exhibited upregulation during aerobic conditioning at sampling point I; three genes (RM25_RS09245, RM25_RS09250, and RM25_RS01195) are predicted to be involved in either iron uptake systems or siderophore-mediated iron uptake systems. Iron plays a vital role as an essential component of electron transport chains and respiratory enzymes, such as cytochromes, as well as in the functioning of catalase and peroxidases, which protect reactive oxygen species (ROS). Additionally, the gene *glpT* (RM25_RS01240), encoding the glycerol-3-phosphate transporter, was upregulated by 2.6- and 4.1-fold under aerobic conditions at sampling points I and III, respectively. Furthermore, the gene RM25_RS04570, encoding the α-ketoglutarate permease, was upregulated under aerobic conditions. This permease facilitates the transport of α-ketoglutarate across the cell membrane, ensuring its availability for utilization in the TCA cycle, enabling subsequent energy production through oxidative phosphorylation. Four other predicted transport genes that were upregulated under aerobic conditions at sampling point I, are associated with the transportation of different molecules, including cations (RM25_RS02225), lipoproteins (RM25_RS07080), myo-inositol (RM25_RS08970), and ions (RM25_RS12135).

#### Fe-S cluster biogenesis is upregulated in response to aerobic conditions

The SUF (Sulfur mobilization) system is a crucial mechanism involved in the biogenesis of iron-sulfur (FeS) clusters, which play a critical role in aerobic growth by mobilizing and utilizing sulfur. The SUF system is responsible for assembling iron-sulfur clusters, which serve as vital cofactors in the proper functioning of numerous enzymes and proteins (38, 39). Notably, in *Mycobacterium tuberculosis*, the SUF system is the sole machinery responsible for the biogenesis of Fe-S clusters (40). In the DSM2071 strain, the SUF system is likely encoded by a gene cluster consisting of eight genes, namely RM25_RS06965 to RM25_RS07000. Among these genes, the last gene in the cluster (RM25_RS07000, sufR) is likely responsible for encoding the transcriptional regulator, while the function of the first gene (RM25_RS06965) remains unknown. Interestingly, the expression levels of the genes within this cluster are upregulated by 4.7-8.3-fold under aerobic conditions, specifically at sampling point III.

#### Transfer of amino groups from aspartate/alanine to α-ketoglutarate is upregulated under aerobic conditions

While the majority of the amino acid metabolism-associated genes showed increased expression under anaerobic conditions **(Table S1)**, RM25_RS06515 coding for a putative alanine or aspartate transferase was over 2 times more expressed under aerobic in comparison to the same gene under anaerobic conditions at both sampling points. This indicates that this pathway, involved in the production of pyruvate and glutamate or oxaloacetate and glutamate, is more active during aerobic growth.

### 2.3 The surfaceome of PFR is influenced by the atmosphere

As proteins exposed at the cell surface (surfaceome) of a bacterium play multiple roles in sensing, responding, and adapting to changing atmospheric conditions, we next complemented the transcriptome data with proteomics. To this end, we conducted an additional bioreactor cultivation analysis with DSM 20271, harvested the cells at the mid-logarithmic growth phase under both the anaerobic and aerobic conditions, and subjected the washed cells in triplicate to surfaceome shaving with trypsin and label-free quantitative (LFQ) proteomics.

#### Peptidoglycan-degrading enzymes were detected as most abundant on PFR cell surfaces

A total of 965 proteins were identified in the present study and these proteins along with their predicted subcellular locations and motifs are listed in **Table S3.** Eight proteins were exclusively identified in aerobic conditions, while 19 were exclusively identified in anaerobic conditions **(Fig. 5A, Table S4)**. Among these proteins, 938 were found in cells cultivated under both environmental conditions **(Figure 5A)**, with 85 proteins harboring a signal peptide (Sec/SPI, Sec/SPII, or Tat/SPI type) and 79 proteins with 2–14 transmembrane domains, directing the protein outside of the cells or into the cell membrane, respectively. Cytoplasmic proteins comprised the largest group (n, 634), suggesting that the proteins might have entered the cell surface in yet unknown fashion **(Fig. 5B)**.

**Figure 5.**
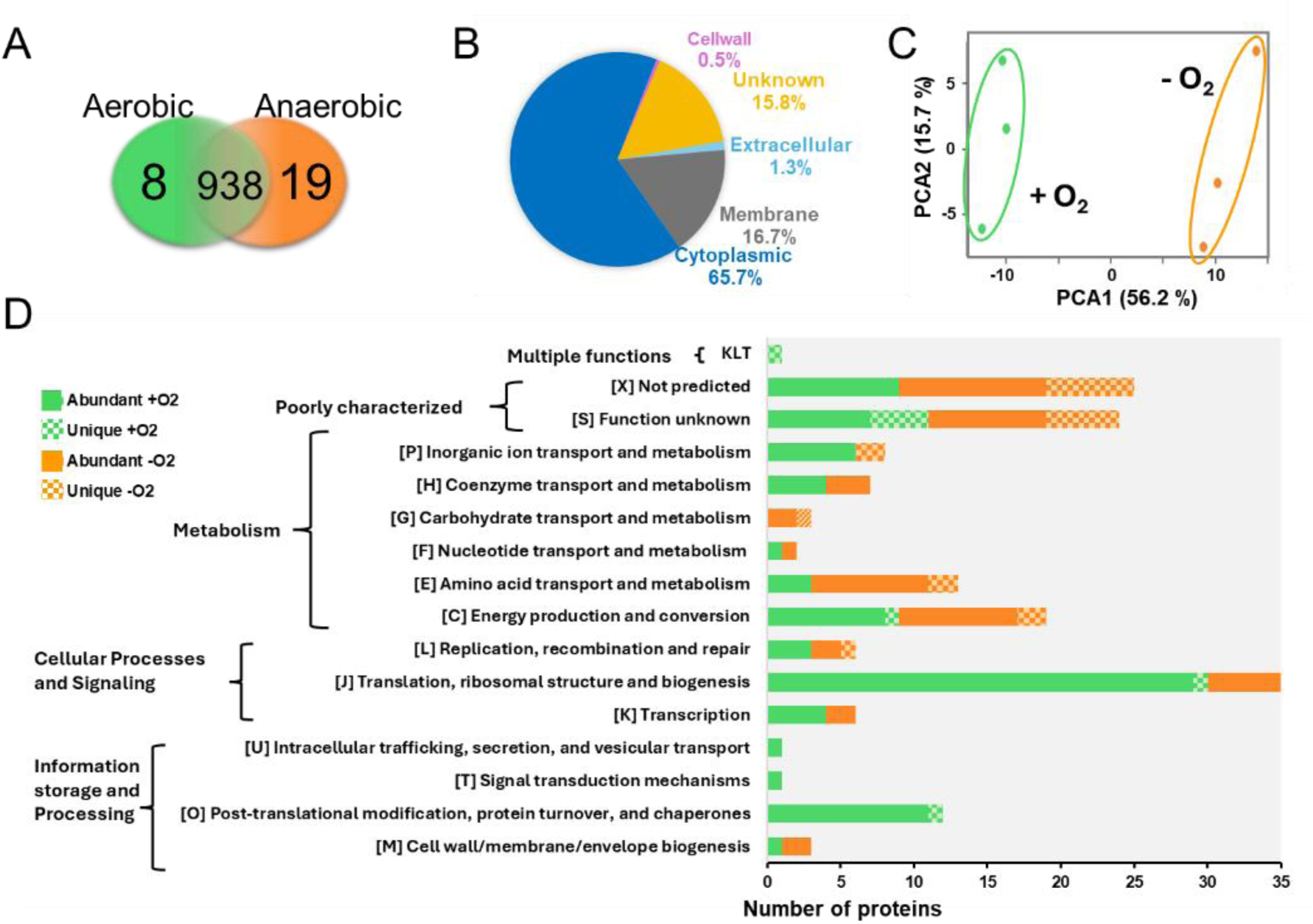
Proteomic analysis of *PFR* DSM 20271T surfaceomes. **(A)** The Venn diagram indicates uniquely and intersectively identified proteins in aerobic and anaerobic growth environments. **(B)** Subcellular localization of the identified surfaceomes (predicted by EggNOG 5.0.0). **(C)** The principal component analysis (PCA) plot of the LFQ intensities of the individual samples with 3 replicates. PCA identifies a correlation between the protein abundance profiles between aerobically (green) and anaerobically (orange) cultured bacterial cells. **D)** Functional categorization of proteins that were either uniquely identified from aerobic (green) or anaerobic (orange) culture or differentially abundant between the two growth environments (predicted by EggNOG 5.0.0).

Proteins identified with the highest intensity values, all of which have also been identified in previous studies from the cell surface or the extracellular milieu of *PFR* (Table 2) (4, 41) are likely to represent the most abundant surface proteins in DSM20271, five were found to be differentially abundant in response to atmospheric conditions after the LFQ analyses.

**Table 2.**
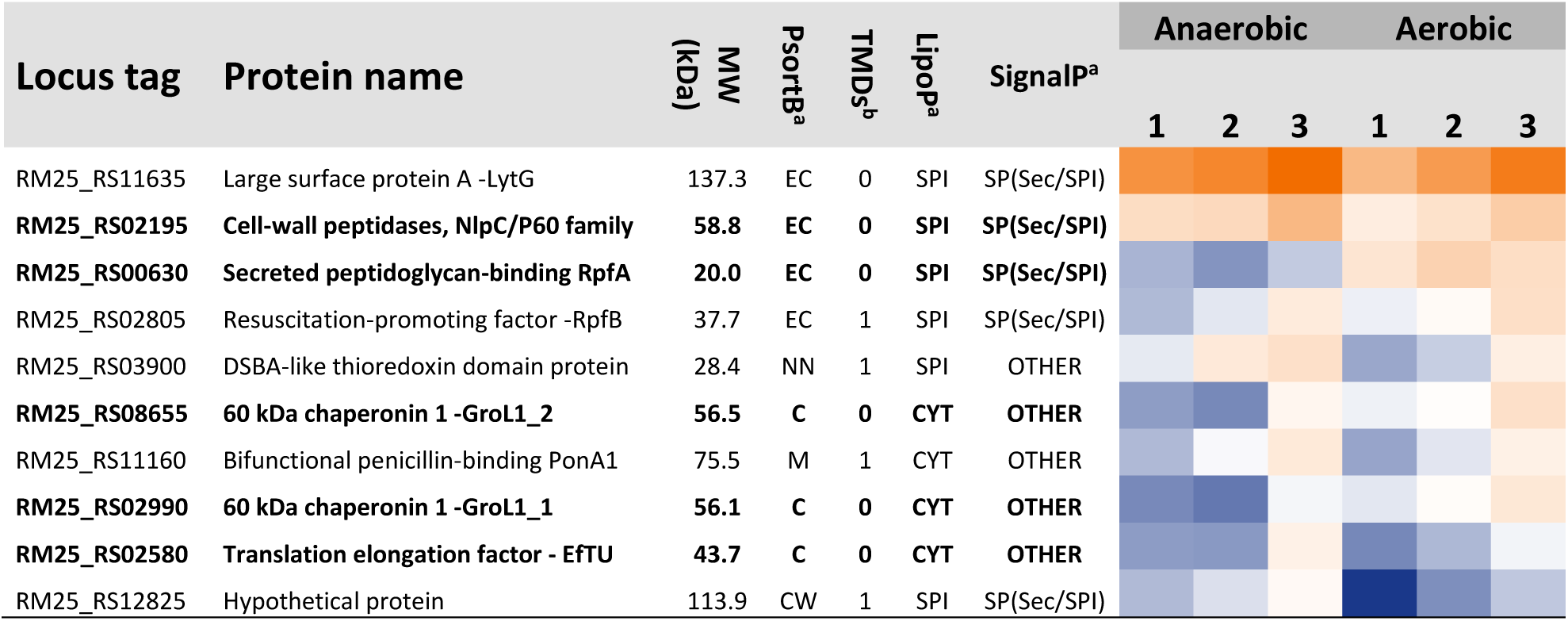
Top ten proteins identified with the highest raw intensity values (average of three independent replica samples). Color code, low (blue) and high (orange) protein abundances. Proteins in bold show statistically significant abundance change between the indicated conditions. ***a)*** *PSORTb v.3.0.3, and the presence of possible classical and non-classical signal peptide sequences were analyzed with LipoP 1.0 and SignalP 5.0. **b)** The number of transmembrane spanning domains (TMD) was predicted using TMHMM 2.0, subcellular localization of proteins was done with prediction tool*.

GroL1_1, GroL1_2, and RpfA were more abundant under aerobic conditions, whereas Ef-Tu and the NlpC/P60 family-secreted peptidase were more abundant during anaerobic growth. NlpC/P60 peptidases are a family of peptidoglycan hydrolases involved in bacterial cell wall turnover and division, as well as interaction with the environment (42). GroL and Ef-Tu belong to typical cytosolic moonlighting proteins, which exhibit a second function on the cell surface of various bacteria (43). The reported moonlighting surface functions of Ef-Tu include mediating attachment to human intestinal cells and mucins (44) and functioning as an environmental sensor (45). A large number of different biological moonlighting functions have been ascribed to chaperonin 60 (Hsp60) proteins (46). Notably, a recent study revealed antiadipogenic activity of surface-exposed Hsp60 in *P. freudenreichii* strain MJ2 (47). For *Helicobacter pylori* Hsp60, extracellular properties include iron binding (48), an ability that could be beneficial in aerobic conditions, thus explaining differential surface abundance in response to atmospheric conditions for *PFR*. However, the moonlighting functions of Ef-Tu and the two GroL proteins of *PFR* remain still to be studied. RpfA belongs to resuscitation-promoting factors that in *M. tuberculosis* have been shown to mediate resuscitation from dormancy via a mechanism involving peptidoglycan hydrolysis (49). The increased abundance of both RpfA and LytG with a predicted glucosaminidase domain suggests their potential roles in controlling the autolysis of the *PFR* cells.

#### Ribosomal moonlighters were more abundant within the aerobic-surfaceomes

The principal component analysis (PCA) of the LFQ intensities of each detected cell surface protein was applied to explore the statistically significant differences (p < 0.05) in protein abundances between the anaerobic and aerobic growth conditions. The PCA plot in **Fig. 5C** explains over 70% of the total variation with a clear clustering for each aerobic and anaerobic-associated surfaceome. Both surfaceomes demonstrate a strong negative (aerobic) and positive (anaerobic) correlation, respectively, indicating that *PFR* responded to oxygen level alterations by introducing specific protein abundance changes at the cell surface. These oxygen-dependent changes result from either gene expression alterations or changes in protein secretion and/or stability.

An additional two-sample t-test (FDR 0.05) showed that the LFQ intensities of 139 proteins were significantly differentially abundant on the cell surface between the two growth conditions **(Table S5)**. Among these, 88 proteins were more abundant in aerobically cultivated cells, while 51 proteins were more abundant in anaerobically cultivated *PFR* cells. The largest proportion of distinct or differentially abundant surface proteins mainly fell under the functional category of translation, ribosomal structure, and biogenesis **(Fig. 5D)**. This aligns with a previous report that the largest number of intracellular/surface moonlighting protein candidates identified in various bacterial surface proteomics studies belong to this functional category (50). Most of the proteins were ribosomal proteins (r-proteins) from both the small (Rps) and large (Rpl) ribosomal subunits, which are commonly associated with the cellular process of translation in the cytoplasm. Ribosomal proteins have been identified in surfaceome studies of various bacterial species (50–54). Here, the r-proteins were more abundant on the aerobic surfaceome compared to these proteins detected on anaerobic cell surfaces. While we did not observe significantly increased transcription of genes encoding r-proteins in log phase samples, the differential surface abundance is not likely the result of differential gene expression. A Similar finding has been demonstrated in *Lacticaseibacillus rhamnosus* GG, in which a ribosomal protein L2 (RPL2) bile-dependent abundance change occurred (53).

Interestingly, the majority of the differentially abundant ribosomal proteins identified here have previously been identified from the vesicles produced by *PFR* (5, 55, 56). In these studies, the proteins delivered by vesicles contributed to translation, ribosomal structure, and biogenesis. Proteomics data obtained from membrane vesicles of two *Limosilactobacillus reuterii* strains identified over 50 ribosome-associated proteins (57) and similar findings have been reported for *S. aureus, Bacillus anthracis*, *and Streptococcus pneumoniae* (58–60). These observations suggest that vesicles may serve as a means for transporting ribosomal proteins to the external surroundings and neighboring cells. Rodovalho et al., (5) also proposed that in response to environmental stress, *PFR* might exhibit an enhanced ability to produce vesicles. In *L. reuterii* DSM 17938 strain the oxygen stress influenced both the number of membrane vesicles and the protein concentration of *L. reuteri*-derived membrane vesicles (57). Consequently, further investigation is warranted to understand how the oxidative stress response affects the morphological, biochemical, and biological characteristics of *PFR*-secreted vesicles.

Taken together, the surfaceome data suggest that the release of cytoplasmic proteins depends on growth conditions rather than trypsin-triggered cell lysis, as several of these proteins were uniquely detected in one condition only, while expressed under both conditions. This is in line with many other studies reporting the presence of cytoplasmic proteins on the cell surface of various bacteria, and it is proposed that their controlled release or export involves phages, autolysins, hydrolytic enzymes, and/or vesicles (61–67). Thus, the increased abundance of RpfA and LytG on aerobic cell surfaces could be one of the factors explaining the oxygen-dependent increase in the abundance of r-proteins at the PFR cell surface. Additionally, changes in externally provided oxygen might have affected the physicochemical characteristics of the cell surface in such a way as to stimulate the binding of certain cytoplasmic proteins, as bacteria are known to modify their cell surface hydrophobicity and charge in response to environmental changes (68). Since r-proteins are typically negatively charged and their gene expression did not show significant atmospheric-dependent changes, the externally provided oxygen could have altered the negative net charge of the bacterial cell surface, as observed in some bacteria (69, 70). Therefore, aerobic conditions could have promoted the binding of these proteins to the cell surface, either directly or indirectly through oxygen-triggered gene expression changes affecting the charge and hydrophobicity of the cell surface.

### 2.4 Increasing availability of key compounds and metabolites can enhance aerobic growth of PFR

Omics data suggest that pathways involving iron and α-ketoglutarate transport and metabolism, as well as succinate metabolism, are more active under aerobic conditions. This data also suggest that alanine/aspartate transferase activity is necessary when *PFR* is exposed to oxygen. Aspartate aminotransferase catalyzes the transamination of aspartate and α-ketoglutarate, forming glutamate and oxaloacetate, both of which are intermediate metabolites in the TCA cycle, alongside α-ketoglutarate and succinate. Thus, it could be that increased availability of iron, aspartate, α-ketoglutarate, or succinate influences the aerobic growth of *PFR*. To test this, the colony formation of DSM 20271 together with another *PFR* type strain (DSM4902) on YEL agar with and without FeSO4, aspartate, α-ketoglutarate, or succinate both under aerobic and anaerobic conditions were compared **(Fig. 6)**. Earlier studies reported an inability of *PFR* to form colonies under aerobic conditions (71, 72), whereas a recent study demonstrated that DSM 20271 thrives in the presence of oxygen (16), implying strain-dependent differences in cells response to oxygen. Our results revealed that under anaerobic conditions, 4 days after incubation at 30 °C, colonies were formed equally by both strains. However, after 6 days of incubation under aerobic conditions, no visible separate colonies of DSM4902 formed without supplementation, whereas miniscule colonies of DSM 20271 were observed **(Fig. 6)**. Notably, addition of FeSO4 and α-ketoglutarate significantly improved the colony-forming ability of DSM4902, while aspartate and succinate had a lesser effect **(Fig. 6)**. Moreover, DSM 20271 formed small colonies under aerobic conditions even without supplementation. The addition of FeSO_4_ and α-ketoglutaric acid increased the size of DSM20271 colonies under aerobic conditions **(Fig. 6)**. α-ketoglutarate and succinate are important intermediates in the TCA cycle, while aspartic acid can be metabolized in various ways, including conversion to another TCA intermediate, oxaloacetate. Thus, it is tempting to suggest that the addition of these metabolites in YEL enhances the flow of carbon and energy through the TCA cycle, thereby promoting growth and colony-forming ability under aerobic conditions. On the other hand, iron is crucial for various cellular processes, including electron transport in the respiratory chain. Thus, iron sulfate can help improve the efficiency of energy production by supporting the function of iron-containing enzymes in the bacterium’s electron transport chain.

**Figure 6.**
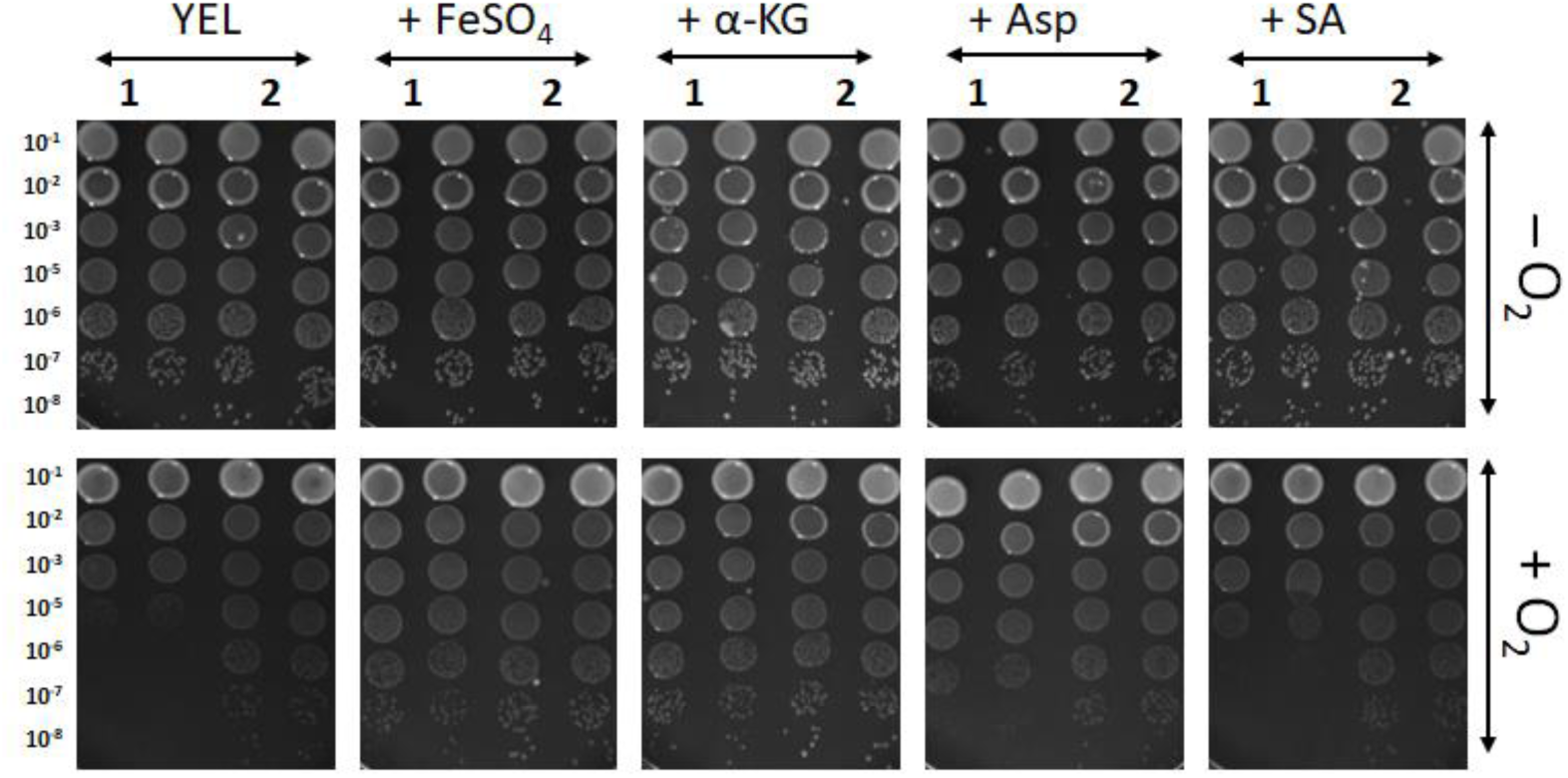
Colony formation of DSM 4902 (1) and DSM 20271 (2) under anaerobic (-O2) and aerobic (+O2) conditions on YEL agar plates with and without supplementation of FeSO4 (1 mM), aspartic acid (10 mM), α-ketoglutaric acid (1 mM), or succinic acid (1 mM) after four (anaerobic) or six (aerobic) days of incubation at 30°C. The images are representative of experiments repeated at least three times for each condition. YEL, yeast extract − lactate medium; α-KG, α-ketoglutarate; Asp, aspartate; SA, succinate.

#### FeSO_4_ controls the production of heme in the presence of oxygen

FeSO_4_ supplementation was found to enhance the colony formation ability under aerobic conditions, which potentially occurred through increased heme production and the activity of heme-dependent enzymes (e.g., cytochromes, catalase, peroxides). To study this further, DSM 20271 was cultured in triplicate for 96 hours under microaerobic conditions, with and without 1 mM FeSO_4_, while control cultures were propagated anaerobically.

Results revealed no impact on final cell densities with FeSO_4_ supplementation (data not shown). However, under microaerobic conditions, the addition of FeSO_4_ elevated heme production and reduced B12 vitamin production **(Fig. 7).** This suggests a shift in tetrapyrrole synthesis toward heme in the presence of excess oxygen and iron, at the expense of B12 production. Superior B12 production under microaerobic conditions aligns with a previous report by Dank et al. (17) as well as our bioreactor results **(Fig. S1BF)**. Notably, the cobamide detected under microaerobic conditions is the B12 form, while both pseudo-B12 and B12 forms are produced under anaerobic conditions **(Fig. 7)**. Acetate and succinate were present in all samples, while propionic acid was exclusive to anaerobic conditions, indicating potential differences in propionic acid metabolism under microaerobic conditions (16).

**Figure 7.**
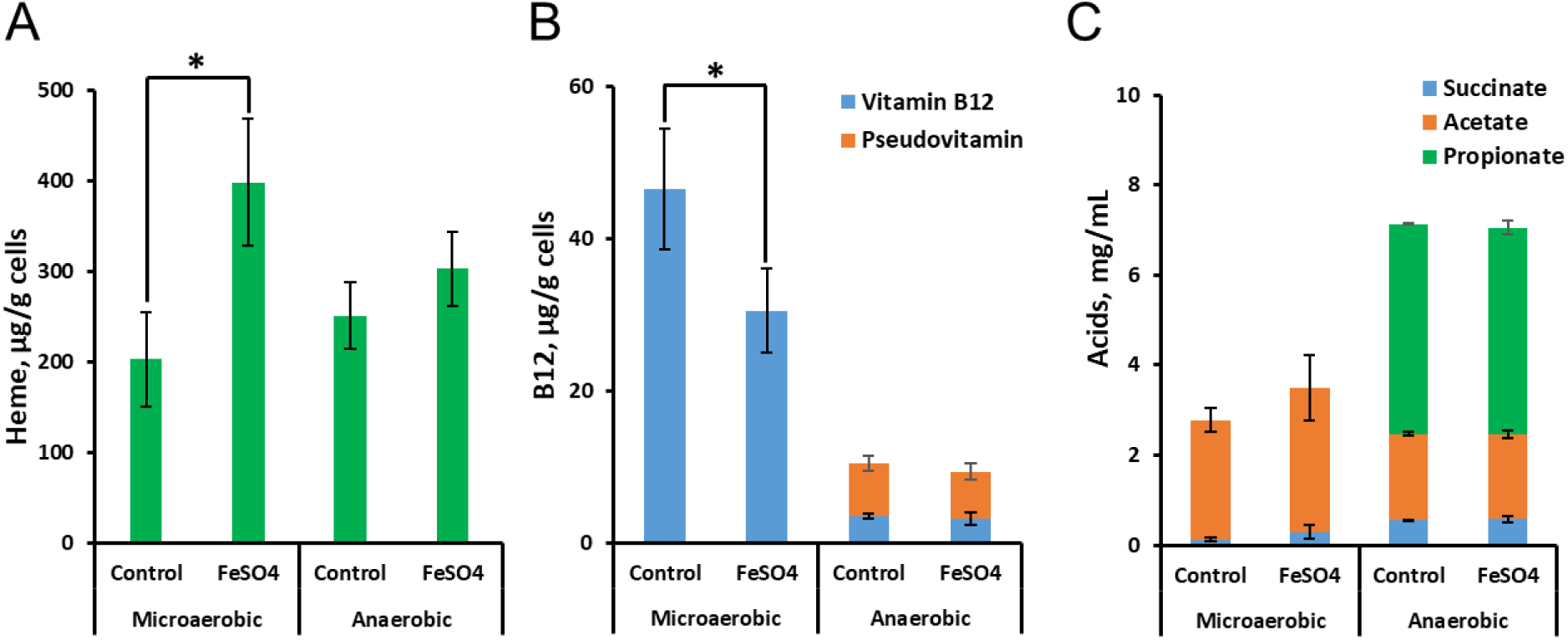
Effect of FeSO_4_ (1 mM) supplementation on the biosynthesis of heme **(A)**, vitamin B12 **(B)**, and accumulation of organic acids **(C)** when grown under microaerobic and anaerobic conditions in yeast extract-lactate medium supplemented with CoCl_2_ (5 mg/L). The values are means and standard deviations of three biological replicates. A significant difference between the treatments (p < 0.05) is indicated by an asterisk.

## 3. Conclusion

We investigated the metabolism and growth characteristics of *PFR* DSM 20271 under aerobic and anaerobic conditions. Our results showed enhanced biomass formation under aerobic conditions. Additionally, lactate consumption was faster under anaerobic conditions, while propionate and acetate levels increased over time under anaerobic conditions but remained low under aerobic conditions. Cobamide synthesis was notably impacted by oxygen, with minimal B12 or pseudo-B12 production under aerobic conditions compared to anaerobic or microaerobic conditions. Genes related to iron and α-ketoglutarate transport and metabolism, succinate metabolism, and alanine/aspartate transferase activity were found up-regulated under aerobic conditions, indicating energy production through respiration and classifying PFR as a facultative anaerobe. Surfaceome analyses revealed an oxygen-dependent increase in peptidoglycan hydrolyzing enzymes (RpfA and LytG) along with a several r-proteins and other cytoplasmic proteins. This suggests that PFR may utilize controlled cell lysis or vesiculation to release cytoplasmic moonlighters, influenced by changes in cell surface structures or gene expression triggered by oxygen. Moreover, our study demonstrated that the growth of DSM 20271 under aerobic conditions could be enhanced by supplementation with FeSO4, aspartate, α-ketoglutarate, or succinate. Interestingly, FeSO4 supplementation also increased heme production at the expense of B12 production in the presence of oxygen, suggesting a negative effect of externally available iron on B12 synthesis in *PFR*. In summary, our findings underscore the significant role of oxygen in the metabolism and growth of *PFR* DSM 20271, impacting lactate consumption, organic acid production, vitamin synthesis, and cell surface protein abundance.

## 4. Materials and Methods

### 4.1 Culture conditions

Strain DSM 20271 was grown in yeast extract-lactate-based medium (YEL) and routinely cultured at 30 °C starting by streaking from glycerol stock (15 %, −80 °C) onto a solid medium (supplemented with 1.5 % agar) and growing for four days in anaerobic jars (Anaerocult, Merck). Liquid cultures were obtained by inoculation of three separated colonies into 10 mL of liquid medium in 15 ml Greiner tubes and propagation for three days at 30 °C without shaking. Experimental cultures were routinely inoculated to 1 % (v/v) and 1.5 % (v/v) in test tubes and bioreactors, respectively. All experiments were performed with three biological replicates.

### 4.2 Cell density and viable cell count

Cell densities were assessed by measuring the optical density (OD) at a wavelength of 600 nm (OD600) by aseptically removing small aliquotes from the bioreactors. This measurement was routinely taken during the 72-h bioreactor cultivations. Microbial counts were determined using the plate counting technique. Dilution series were prepared in sterile Saline (0.9% NaCl) or PBS, and appropriate dilutions were either spread (100 μL) or spotted (10 μL) onto YEL agar plates. After incubation for 4-7 days in anaerobic conditions using Anaerocult (Merck) at a temperature of 30 °C, colonies were counted, and the CFU/mL (colony-forming units per milliliter) were determined.

### 4.3 Fermentation in bioreactors and sampling

To investigate the effect of available oxygen on gene expression, growth kinetics, and metabolic activity, batch fermentations were conducted in bioreactors under aerobic and anaerobic conditions.

Under aerobic conditions, the pO2 level at 20% was maintained throughout the cultivation process, recognizing the significant impact of increasing cell density and fluid viscosity on oxygen transfer efficiencies - a concern that can be mitigated by controlling oxygen partial pressure (pO2) in bioreactors conducted experiments (73). This decision aimed to ensure optimal oxygen availability, consistent with the reported oxygen conditions in a minimal lactate medium. During anaerobic growth, the pO2 level was verified to be at 0% to ensure the absence of oxygen in the experiment.

The fermentations were carried out in Sartorius A bioreactors with a growth volume of 750 mL. The temperature was maintained at 30 °C, and the pH was controlled at 7.0 using NaOH (5 M) and HCl (1 M). Stirring was set at 200 rpm for anaerobic experiments. In anaerobic conditions, continuous nitrogen gas flow at 20 cc was utilized, while in aerobic conditions, the oxygen saturation level was maintained at 20% through air and oxygen gas flow, with stirring alternating between 200-300 rpm.

Samples of 12 mL were taken aseptically in three biological replicates at four-time points (I-IV) for each atmospheric condition. For anaerobic fermentation, the sampling points were at 12 h (I), 26 h (II), 32 h (III), and 46 h (IV). In aerobic fermentation, samples were collected at 13 h (I), 40 h (II), 52 h (III), and 66 h (IV). Surfaceome analyses were performed on samples from the initial time point (I) in both aerobic and anaerobic fermentations. For gene expression and ddPCR analysis, samples from time points I and III were analyzed.

Cell harvesting was conducted by centrifugation at 3220 x g for 10 min at +4 °C. The supernatants were collected and stored at −80 °C for acid and vitamin analyses. Harvested cells for vitamin analysis were washed with 1 M Tris-HCl buffer (pH 8.0) and stored at −20 °C.

For RNA sequencing, the harvested cells were immediately resuspended in 1500 μL of RNA later (Invitrogen™) and incubated overnight at +4 °C. After incubation, the cells were harvested by centrifugation at 12,000 G for 5 min at +4 °C and stored at −80 °C.

### 4.4 RNA extraction and purification

Cells stored at −80 °C were thawed and resuspended in 50 μL of Tris-EDTA buffer (TE buffer). The cell suspension was then transferred to lysing matrix tubes containing glass beads. Cell lysis was performed using the FastPrep-24 homogenizer, with three cycles of 30 seconds each at a speed of 6.5 M/S. Between cycles, the samples were kept on ice for 60 seconds to maintain the temperature. After the final ice incubation, the lysates were suspended in 150 μL of TE buffer and thoroughly vortexed. Subsequently, the lysates were centrifuged at 12,000 g for 5 min at +4 °C to remove cell debris and glass beads. The supernatants were carefully collected and stored at −20 °C.

Total RNA extraction was carried out using the RNeasy Mini Kit (Qiagen) according to the manufacturer’s instructions, including DNase digestion to remove any remaining DNA. The RNA was eluted with 40 μL of nuclease-free water.

Following RNA extraction, the samples underwent DNAase treatment and were further purified using an RNA clean-up method. The concentration of purified single-stranded RNA was measured using a NanoDrop spectrophotometer. The RNA samples were divided into aliquots and stored at −80 °C. To deplete ribosomal RNA (rRNA), the Ribo-Zero Plus rRNA Depletion Kit (Illumina Inc., USA) was employed, following the protocol provided by the manufacturer.

### 4.5 RNA sequencing and data analysis

The RNA-seq libraries were generated using the QIAseq Stranded Total RNA Lib kit from Qiagen, and sequencing was performed on the Illumina NextSeq 500 platform.

To assess the quality of the sequencing reads, FastQC v0.11.7 was employed. Adapter sequences and low-quality reads (using parameters -m 30 -q 25) were removed using cutadapt v1.9.1. Reads that mapped to rRNA sequences were eliminated using sortmerna v2.1, while the remaining reads were aligned to the *PFR* DSM 20271 genome assembly GCF_000940845.1_ASM94084v1 using the mem algorithm of BWA v0.7.11. The number of reads mapped to each gene was counted using Htseq-count v0.1.11. Differential expression analysis was performed using DESeq2 (74), comparing atmospheric conditions (anaerobic versus aerobic) within growth phases (logarithmic and stationary). Additional detailed information on the validation of the RNAseq data with ddPCR is provided in the supplemental material.

The RNA-seq data can be accessed at the National Center for Biotechnology Information under the accession PRJNA872337 (https://www.ncbi.nlm.nih.gov/sra/PRJNA872337).

Genes were considered to be significantly differentially expressed if their adjusted p-value (padj) was ≤ 0.05 and fold changes were ≥ 2.0 (log2Fold ≥ 1). Predicted GO terms for the genes were obtained using PANNZER2 (75) with default parameters. The lists of differentially expressed genes from the various comparisons were analyzed for enrichment of GO terms in the Biological Process ontology using the R package clusterProfiler (v. 4.0) (76, 77). The reference set used for the analysis included all the genes annotated with GO terms in the genome. The annotated GO terms were visualized using Cytoscape 3.9.1 (78) with the ClueGO plug-in (79).

### 4.6 UHPLC Analysis of Vitamin B12 and Pseudovitamin

Analysis for the active vitamin B12 and pseudovitamin was based on ultra-high performance liquid chromatography (UHPLC) method described by Chamlagain et al. (80). The B12 was extracted in its cyano form from cellular pellets using established methods, as detailed by Chamlagain et al. (80, 81). Specifically, 0.1–0.2 g of the cell pellet underwent extraction with a pH 4.5 buffer (comprising 8.3 mM sodium hydroxide and 20.7 mM acetic acid) in the presence of sodium cyanide, yielding a 25 mL extract. The analysis was performed with Waters UPLCsystem (Waters, Milford, MA, USA) with a C18 column (Waters Acquity HSS T3, 2.1 x 100 mm, 1.8 µm) at a flow rate of 0.32 mL/min, a UV detection by a photodiode array (PDA) detector at 361 nm and an injection volume of 5-15 μL were used. Cyanocobalamin was identified based on retention time and absorption spectrum and quantified against its standard.

### 4.7 HPLC method to analyze acids and glucose

Glucose, acetic acid (AA), lactic acid (LA), propionic acid (PA), pyruvate and succinate were determined using a high-performance liquid chromatography (HPLC) method described in Chamlagain et al. (81) with modifications. Samples were centrifuged at 12,000 x g for 10 min, and the resulting supernatants were collected. After syringe filtration (Pall, MI, USA; 0.2 µm), the samples were stored at −20 °C. Analyses were conducted on the thawed samples. The analysis was performed on a Hi-Plex H column (Agilent, CA, USA; 300×6.5 mm), with a HiPlex H guard column (Agilent, CA, USA; 50 x 7.7 mm). The HPLC system was equipped with a Waters 515 pump, autosampler, ultraviolet (UV) detector (Waters 717), and refractive index detector (HP 1047A, HP, USA). The mobile phase was 10 mM H_2_SO_4_, and the flow rate was set at 0.5ml/min for 35 min with the column temperature maintained at 40 °C.

### 4.8 Quantitative heme analysis

The inoculated YEL was incubated at 30 °C under microaerobic and anaerobic conditions for 4 days. To create a microaerobic condition, the cultures (20 mL) were grown in 100-mL glass bottles under shaking conditions (200 rpm) and opened once every day in a sterile environment. The medium was supplemented with 5 mg/L CoCl2 and in the case of Fe-supplemented samples, FeSO_4_ was also added (1 mM). After incubation, the cultures were centrifuged (6800 × g, 10 min at room temperature). B12 and heme were measured in cells, while the supernatants were analyzed for the organic acid contents.

Samples for heme measurements were prepared as follows. A portion of the cell pellet (0.1 g) was suspended in 0.1 mL of 0.1 M NaOH (82) in lysing matrix tubes containing glass beads and the cells were lysed using the FastPrep-24 homogenizer (5 cycles of 30 seconds each at a speed of 4.5 M/s). To maintain the temperature, the samples were kept on ice for 60 seconds between the cycles. Another 0.1 mL of NaOH was added to the cell lysates, homogenized, and centrifuged (12,000 g for 10 min at room temperature). The supernatants were carefully collected and analyzed for the heme concentration.

Heme concentrations were measured using a heme assay kit (MAK316, Sigma-Aldrich, St. Louis, MO, USA) according to the manufacturer’s instructions. Absorbance was measured at 405 nm. The concentration of heme in the samples was calculated against the heme calibrant. The heme content is reported as µg heme/g cells.

### 4.9 Colony forming ability using spot-plating assay

*PFR* DSM 4902 and DSM 20271 cells were grown at 30°C in YEL liquid medium for 3d until OD_600_ reached approximately 1.2. Cells were pelleted by centrifugation, washed once with PBS, and resuspended in PBS to the same OD_600_, serially 10-fold diluted, and spotted (5 µl of dilutions) on YEL agar plates supplemented with appropriate chemicals. Plates were incubated under anaerobic or aerobic conditions at 30°C for 4 days (anaerobic) or 6 days (aerobic).

### 4.10 Surfaceome analyses

#### Preparing the cell samples for proteomics and enzymatic cell-surface shaving

The surfaceome samples were harvested from mid-log (I) growth phases of both aerobic and anaerobic fermentations by centrifugation (4 min, 4 °C, 3320× g). The cells were washed with 0.1 M sodium acetate (pH 5.0, Sigma-Aldrich) and resuspended into 50 mM TEAB (17% sucrose) (triethylammonium bicarbonate buffer, Sigma-Aldrich) after centrifugation (4 min, 4 °C, 3320× g). Enzymatic surface shaving was carried out with 0.05 µg/µL of trypsin (Promega, Madison, WI, USA) for 30 min at 37 °C. Afterward, the protein-cell suspensions were first time centrifuged (4 min, RT, 4000× g) to ensure cell-free supernatant and second time centrifuged (2 min, RT, 16 000× g) through Spin-X® filters to recover the peptides and the enzyme. Flow-troughs were left to incubate for 17 h at 37 °C before the tryptic digestions were terminated with the addition of trifluoroacetic acid (TFA, Sigma-Aldrich) at a final concentration of 0.6%. The peptide concentrations were measured spectrophotometrically with NanoDrop-1000 (Thermo Fisher Scientific, DE, USA). Samples were stored at −20 °C.

#### LC-MS/MS identification

The tryptic peptides were concentrated and purified using ZipTips C_18_ (Merck Millipore) according to the manufacturer’s instructions. An equal amount of each peptide sample was loaded into an Easy-nLC 1000 nano-LC system (Thermo Scientific, Waltham, MA, USA) coupled with a quadrupole Orbitrap mass spectrometer (Q ExactiveTM, ThermoElectron, Bremen, Germany) equipped with a nano-electrospray ion source (EASY-SprayTM, Thermo Scientific, Waltham, MA, USA), and analyzed as previously described (83). The acquired MS raw files, for protein identification and label-free quantification (LFQ), were searched against an in-house database of DSM 20271T (CP010341) (18) with MaxQuant (ver. 1.6.1.0) under the following settings: carbamidomethyl (C) was specified as a fixed modification; oxidation of methionine was set as a variable modification; a peptide tolerance was set at 20 ppm in the first search and the main search error of 4.5 ppm was applied. Additionally, trypsin without proline restriction enzyme option was used, allowing two miscleavages; the false discovery rate (FDR) filtering was set to 1 % for both peptide and protein identification, and the minimal number of unique and razor peptides were defined as 1.

#### Surfaceome bioinformatics and statistics

The identified proteins were submitted to SignalP 5.0 (84), SecretomeP 2.0 (85), LipoP 1.0 (86, 87) and TatP 1.0 (88) analysis tools to determine the presence of possible classical and non-classical signal peptide sequences. Additionally, the prediction of protein transmembrane helices was acquired by TMHMM Server v. 2.0 (89, 90), general protein function annotation, according to the COG, was accomplished with EggNOG 5.0.0 (91) and the isoelectric points (pIs) and molecular weights (MWs) were predicted by EMBOSS Pepsats (92).

The statistical comparisons of the MaxQuant-derived protein identifications were performed in Perseus version 1.6.15.0 (93), at which the known contaminant, reverse hits, and proteins only identified by site were excluded. The MS-raw intensities were filtered to contain minimally 2/3 valid values in at least one of the growth environments to obtain the total and unique identifications. Statistical comparisons of the normalized log2-transformed LFQ levels were also carried out in Perseus. The data were filtered to include proteins with valid LFQ-values in at least 70% of biological replicates in at least one group. Before a Student’s t-test with a permutation-based false discovery rate of 0.05, the missing values were imputed from a normal distribution using parameters: width 0.3 and downshift 1.8. A complete linkage hierarchical clustering was performed on normalized (Z-score) values. The mass spectrometry proteomics data have been deposited to the ProteomeXchange Consortium via the PRIDE (94) partner repository with the dataset identifier PXD051703.

## Data availability statement

The transcriptomics and proteomics data sets generated during and/or analysed during the current study are available in the SRA database under the accession PRJNA872337 and PRIDE repository with dataset identifiers PXD051703. The other data sets generated during and/or analysed during the current study and not included in the manuscript are available from the corresponding author on reasonable request.

## Supporting information

Supplemental experiment

## Acknowledgements

Funding for this study was provided by the University of Helsinki Doctoral School in Environmental, Food and Biological Sciences. The study was also funded by Academy of Finland (project number 325784), Novo Nordisk Foundation (project number NNF20OC0065096), core Facilities programme of the South-Eastern Norway Regional Health Authority (TN), Research Council of Norway INFRASTRUKTUR-programme (295910). We also thank Finnish Food and Drink Industries’ Federation (ETL) for their support in funding the research.

The following core facilities are acknowledged for their services and support, which were essential for this work: Institute of Biotechnology, DNA Sequencing and Genomics, University of Helsinki, Proteomics Core Facility at Oslo University Hospital.

## Author contributions

IL: Study design and experimental work – transcriptomics, acids, vitamin, cell counts, Writing – original draft

AS: Study design and experimental work – proteomics, Writing – original draft

PD: Experimental work – Bioinformatics, Writing – review

BC: Methodology – Acid, vitamin & heme, Supplementation experiment, Writing – original draft

ME: Methodology – Acid and vitamin

PA: Writing – review

TN: Methodology – proteomics, Writing – review, Funding acquisition

KS: Methodology – proteomics, Writing - review & editing

VP: Writing – review

PV: Conceptualization, Supervision, Methodology, Writing – original draft, review & editing, Funding acquisition

## Supplementary Materials

**Table S1.** Results of RNA sequencing of the strain *P. freudenreichii* DSM 20271 grown in bioreactors under anaerobic (nitrogen) and aerobic (oxygen) atmospheres at sampling points I and III. Columns A-I specify the order of annotated genes, locus tags of the current NCBI annotations, the previous version of the NCBI locus tags, product names, locations in the genome and strand orientation. Column J-AM: results of DEseq2 analyses with models comparing the effect of atmosphere (anaerobic vs. aerobic) at both sampling points (columns J-O); the effect of atmosphere (aerobic vs. anaerobic) at sampling point I (columns P-U); the effect of atmosphere (aerobic vs. anaerobic) at sampling point III (columns V-AA); the effect of growth phase under anaerobic atmosphere (columns AB-AG) and the effect of growth phase under aerobic atmosphere (AH-AM). Columns AN-BA list results from previously reported bioinformatics analyses (Deptula et al., 2017), including annotations with PROKKA, core genes identified through comparative analyses against other *PFR* strains with ROARY, IslandViewer, Prophinder, Phaster, REBASE and CRISPRFinder.

**Table S2.** List of 74 genes encoding transport proteins detected with more than two-fold higher expression under anaerobic conditions.

**Table S3.** List of all identified proteins with the detected raw intensity values for each. Reverse hits, potential contaminants, and proteins only identified by site were filtered out. Proteins were filtered to contain minimally 2/3 valid values in at least one of the growth environments. The number of transmembrane spanning domains (TMD) were predicted using TMHMM 2.0, subcellular localization of proteins were done with prediction tool PSORTb 3.0.3, and the presence of possible classical and non-classical signal peptide sequences were analysed with LipoP 1.0, SignalP 5.0, and SecretomeP 2.0. The COG categorization was accomplished with EggNOG 5.0.0.

**Table S4.** Proteins uniquely identified in at least two out of three replica samples in aerobic or anaerobic growth conditions. The colour gradient from pink to red indicates the log-2 transformed raw intensities for each biological replicate (grey colour indicates zero). The number of transmembrane spanning domains (TMD) were predicted using TMHMM 2.0, subcellular localization of proteins were done with prediction tool PSORTb 3.0.3 PSORTb v3.0.2, and the presence of possible classical and non-classical signal peptide sequences were analysed with LipoP 1.0, SignalP 5.0, and SecretomeP 2.0. The COG categorization was accomplished with EggNOG 5.0.0.

**Table_S5.** List of proteins showing statistically significant protein abundance change between the aerobic and anaerobic surfaceomes. For this purpose, the LFQ intensities after log2 conversion and missing value imputation were subjected to two-sample t-test with a permutation-based false discovery rate of 0.05 to confirm the statistical significance. The number of transmembrane spanning domains (TMD) were predicted using TMHMM 2.0, subcellular localization of proteins were done with prediction tool PSORTb 3.0.3 PSORTb v3.0.2, and the presence of possible classical and non-classical signal peptide sequences were analysed with LipoP 1.0, SignalP 5.0, and SecretomeP 2.0. The COG categorization was accomplished with EggNOG 5.0.0.

**Figure_S1.** (A) Cell densities (OD600 values - solid line) and cell viability (CFU/mL - dotted line) in bioreactors conducted experimented under aerobic (green) and microaerobic (blue) growth conditions. (B) Lactate utilization (A), excreted propionate (B), acetate (C), pyruvate (D), succinate (E) and intracellular vitamin B12 of P. freudenreichii DSM 20271 during cultivation under aerobic (green) and microaerobic (blue) fermentations performed in bioreactors. n = 3. In aerobic fermentation, samples were collected at 13 h (I), 40 h (II), 52 h (III), and 66 h (IV). For microaerobic fermentation, samples were obtained at 12 h (I), 26 h (II), 22 h (III), and 51 h (IV).

**Supplemental material.** Material, methods and results for validation of RNAseq data with ddPCR.

