## Supplementary figures and images for "Aerobic Adaptation and Metabolic Dynamics of *Propionibacterium freudenreichii* DSM 20271: Insights from Comparative Transcriptomics and Surfaceome Analysis"

### Supplemental Figure 1

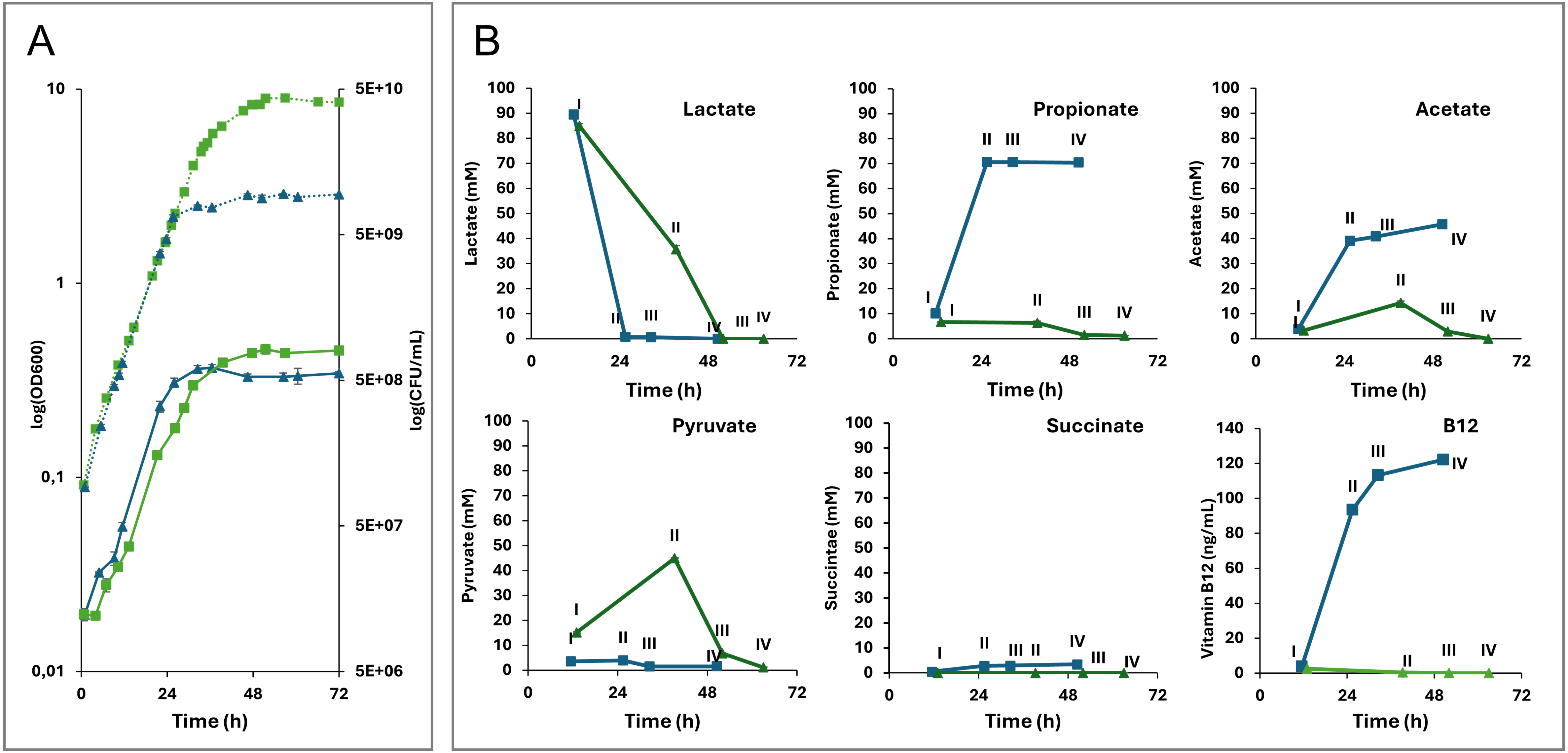
