## Supplemental Table 1 for "Aerobic Adaptation and Metabolic Dynamics of *Propionibacterium freudenreichii* DSM 20271: Insights from Comparative Transcriptomics and Surfaceome Analysis"

**Table S1.** Results of RNA sequencing of the strain *P. freudenreichii* DSM 20271 grown in bioreactors under anaerobic (nitrogen) and aerobic (oxygen) atmospheres at sampling points I and III. Columns A-I specify the order of annotated genes, locus tags of the current NCBI annotations, the previous version of the NCBI locus tags, product names, locations in the genome, and strand orientation. Column J-AM: results of DEseq2 analyses with models comparing the effect of atmosphere (anaerobic vs. aerobic) at both sampling points (columns J-O); the effect of atmosphere (aerobic vs. anaerobic) at sampling point I (columns P-U); the effect of atmosphere (aerobic vs. anaerobic) at sampling point III (columns V-AA); the effect of growth phase under anaerobic atmosphere (columns AB-AG) and the effect of growth phase under aerobic atmosphere (AH-AM). Columns AN-BA list results from previously reported bioinformatics analyses (Deptula et al., 2017), including annotations with PROKKA, core genes identified through comparative analyses against other *PFR* strains with ROARY, IslandViewer, Prophinder, Phaster, REBASE, and CRISPRFinder.

| | | | | | | | | | All_samples_atmospheres_ref:nitrogen | | | | | | Sampling_point_I_ref:oxygen | | | | | | Sampling_point_III_ref:oxygen | | | | | | Nitrogen_ref:sampling_point_III | | | | | | Oxygen_ref:sampling_point_III | | | | | |
|---|---|---|---|---|---|---|---|---|---|---|---|---|---|---|---|---|---|---|---|---|---|---|---|---|---|---|---|---|---|---|---|---|---|---|---|---|---|---|
| No. | NCBI locus tag | Old locus tag | Product / Feature name | Gene number | Feature type | Feaure start | Feature end | Strand | BaseMean | log2FoldChange | lfcSE | stat | pvalue | padj | baseMean2 | log2FoldChange3 | lfcSE4 | stat5 | pvalue6 | padj7 | baseMean | log2FoldChange9 | lfcSE10 | stat11 | pvalue12 | padj13 | BaseMean | log2FoldCl | lfcSE16 | stat17 | pvalue18 | padj19 | baseMean20 | log2FoldChange21 | lfcSE22 | stat23 | pvalue24 | padj25 |
| 1 | RM25_RS00005 | RM25_0001 | Chromosomal replication initiator protein DnaA | 1 | gene | 232 | 1707 | + | 2152,419 | -0,212429314 | 0,167861 | -1,26551 | 0,205689 | 0,309781 | 2436,889111 | -0,121023728 | 0,119205 | -1,01526 | 0,309982 | 0,37687 | 1867,949 | -0,331890892 | 0,215246 | -1,54191 | 0,123095 | 0,202044 | 2442,674 | -0,34943 | 0,177606 | -1,96747 | 0,04913 | 0,08379 | 1957,824386 | -0,501998071 | 0,1506 | -3,33333 | 0,000858 | 0,001854 |
| 2 | RM25_RS00010 | RM25_0002 | Drug resistance transporter, EmrB/QacA subfamily | 2 | gene | 1682 | 3151 | - | 612,7505 | -1,282039543 | 0,635872 | -2,01619 | 0,04378 | 0,114094 | 344,7756906 | -2,826772763 | 0,360442 | -7,84252 | 4,42E-15 | 7,49E-14 | 880,7253 | -0,849429197 | 0,484367 | -1,75369 | 0,079484 | 0,144667 | 912,0041 | 0,833005 | 0,403006 | 2,06698 | 0,038736 | 0,069115 | 353,0562168 | 2,891521869 | 0,473503 | 6,106657 | 1,02E-09 | 5,68E-09 |
| 3 | RM25_RS00015 | RM25_0003 | Transcriptional regulator, MerR family | 3 | gene | 3393 | 4010 | + | 269,4061 | -0,681246353 | 0,284055 | -2,39829 | 0,016472 | 0,064385 | 220,1706704 | -0,844687219 | 0,260622 | -3,24104 | 0,001191 | 0,003134 | 318,6415 | -0,576124436 | 0,362346 | -1,58998 | 0,111838 | 0,188209 | 349,5284 | 0,363962 | 0,26029 | 1,398294 | 0,162025 | 0,220875 | 203,5715424 | 0,69966854 | 0,378052 | 1,850723 | 0,064209 | 0,09262 |
| 4 | RM25_RS00020 | RM25_0004 | SqdX | 4 | gene | 4007 | 5188 | - | 477,8525 | -0,140906941 | 0,260693 | -0,54051 | 0,588846 | 0,680015 | 419,0627687 | 0,01763822 | 0,159685 | 0,110456 | 0,912048 | 0,933028 | 536,6421 | -0,267027435 | 0,423211 | -0,63096 | 0,528069 | 0,626717 | 526,5198 | 0,427464 | 0,264063 | 1,618793 | 0,105492 | 0,15543 | 447,7764879 | 0,212999139 | 0,367962 | 0,578861 | 0,562683 | 0,623528 |
| 5 | RM25_RS00025 | RM25_0005 | UDP-sulfoquinovose synthase | 5 | gene | 5202 | 6449 | - | 1237,366 | -0,523103819 | 0,155495 | -3,36411 | 0,000768 | 0,008405 | 1141,285778 | -0,514600623 | 0,176298 | -2,91892 | 0,003512 | 0,008085 | 1333,447 | -0,531335219 | 0,245171 | -2,1672 | 0,030219 | 0,069831 | 1536,77 | 0,167723 | 0,228074 | 0,735386 | 0,462104 | 0,534173 | 997,3940861 | 0,211888559 | 0,196379 | 1,078976 | 0,280598 | 0,341793 |
| 6 | RM25_RS00030 | RM25_0006 | Ectoine/hydroxyectoine ABC transporter, ATP-binding prote | 6 | gene | 6920 | 7864 | + | 1307,585 | 0,619610518 | 0,254615 | 2,43352 | 0,014953 | 0,060854 | 1621,415719 | 0,516387759 | 0,148773 | 3,470977 | 0,000519 | 0,001494 | 993,7551 | 0,792880058 | 0,165399 | 4,79374 | 1,64E-06 | 2,12E-05 | 1096,348 | -0,93719 | 0,172815 | -5,42306 | 5,86E-08 | 5,68E-07 | 1553,070094 | -0,607260773 | 0,129976 | -4,67208 | 2,98E-06 | 9,55E-06 |
| 7 | RM25_RS00035 | RM25_0007 | ABC-type antimicrobial peptide transport system | 7 | gene | 7866 | 10424 | + | 3502,376 | 0,769088273 | 0,323455 | 2,377729 | 0,01742 | 0,066339 | 4575,407361 | 0,645248099 | 0,140928 | 4,578576 | 4,68E-06 | 2,09E-05 | 2429,344 | 1,01120378 | 0,237826 | 4,251856 | 2,12E-05 | 0,000197 | 2754,775 | -1,20797 | 0,185747 | -6,50333 | 7,86E-11 | 1,45E-09 | 4325,557835 | -0,79378288 | 0,163407 | -4,85771 | 1,19E-06 | 4,08E-06 |
| 8 | NA | RM25_0008 | Hypothetical protein | 8 | gene | 11396 | 12172 | - | 374,241 | -1,317292134 | 0,873904 | -1,50736 | 0,131717 | 0,230139 | 187,809937 | -5,067215488 | 0,467361 | -10,8422 | 2,17E-27 | 1,65E-25 | 560,672 | -0,752635444 | 0,412507 | -1,82454 | 0,068071 | 0,128164 | 559,6815 | 0,877481 | 0,403392 | 2,175258 | 0,029611 | 0,055834 | 211,5060234 | 5,266024494 | 0,502607 | 10,47743 | 1,1E-25 | 4,54E-24 |
| 9 | RM25_RS00040 | RM25_0009 | DNA polymerase III, beta subunit | 9 | gene | 12300 | 13460 | + | 2498,819 | 0,036430198 | 0,233587 | 0,15596 | 0,876065 | 0,915692 | 3035,729768 | 0,025421005 | 0,143681 | 0,176926 | 0,859566 | 0,890157 | 1961,907 | 0,054778275 | 0,231083 | 0,23705 | 0,812618 | 0,865899 | 2619,428 | -0,69867 | 0,221453 | -3,15493 | 0,001605 | 0,005241 | 2479,106379 | -0,62736361 | 0,139864 | -4,48554 | 7,27E-06 | 2,18E-05 |
| 10 | RM25_RS00045 | RM25_0010 | DNA replication and repair protein RecF | 10 | gene | 13534 | 14835 | + | 919,4924 | -0,267467359 | 0,168646 | -1,58597 | 0,112746 | 0,207287 | 832,9486078 | -0,193841141 | 0,161345 | -1,2014 | 0,229594 | 0,292832 | 1006,036 | -0,329529628 | 0,269556 | -1,22249 | 0,221522 | 0,31876 | 1059,07 | 0,267976 | 0,192961 | 1,388759 | 0,164906 | 0,223638 | 820,2121065 | 0,19802723 | 0,249265 | 0,794445 | 0,426936 | 0,492314 |
| 11 | RM25_RS13060 | RM25_0011 | Hypothetical protein | 11 | gene | 14864 | 15610 | + | 454,9384 | -0,265352978 | 0,177623 | -1,49391 | 0,135199 | 0,234139 | 410,7881831 | -0,327777529 | 0,165769 | -1,97731 | 0,048007 | 0,076454 | 499,0886 | -0,217219939 | 0,287751 | -0,75489 | 0,450316 | 0,554112 | 523,6973 | 0,170068 | 0,1613 | 1,054361 | 0,291718 | 0,365733 | 406,1828768 | 0,341706919 | 0,283305 | 1,206146 | 0,227761 | 0,284039 |
| 12 | RM25_RS00055 | RM25_0012 | Transposase insI for insertion sequence element IS30B/C/D | 12 | gene | 17183 | 18217 | - | 38,69136 | -1,613476031 | 0,782992 | -2,06066 | 0,039336 | 0,10757 | 24,45828975 | -4,613011575 | 0,931358 | -4,95299 | 7,31E-07 | 3,76E-06 | 52,92444 | -0,946577645 | 0,374805 | -2,52552 | 0,011553 | 0,033995 | 61,49429 | 0,493891 | 0,505768 | 0,976517 | 0,328809 | 0,404173 | 18,68766841 | 4,227003981 | 0,925996 | 4,564819 | 5E-06 | 1,54E-05 |
| 13 | RM25_RS00065 | RM25_0013 | Hypothetical protein | 13 | gene | 18918 | 19901 | + | 354,2535 | -0,773174334 | 0,548174 | -1,41045 | 0,158406 | 0,26091 | 195,9353963 | -1,910127318 | 0,320365 | -5,96234 | 2,49E-09 | 2,01E-08 | 512,5716 | -0,409165742 | 0,354561 | -1,15401 | 0,248497 | 0,349051 | 468,3637 | 0,852143 | 0,325614 | 2,617034 | 0,00887 | 0,021464 | 257,4802776 | 2,419757375 | 0,360678 | 6,708912 | 1,96E-11 | 1,4E-10 |
| 14 | RM25_RS00070 | RM25_0014 | DNA gyrase, B subunit | 14 | gene | 20322 | 22364 | + | 3256,997 | 0,081668477 | 0,212679 | 0,383998 | 0,70098 | 0,77751 | 3038,192105 | -0,444461922 | 0,131273 | -3,38579 | 0,00071 | 0,001976 | 3475,802 | 0,544764171 | 0,255862 | 2,12913 | 0,033243 | 0,074695 | 3349,003 | -0,36692 | 0,225027 | -1,63055 | 0,102986 | 0,152329 | 3275,896217 | 0,668952253 | 0,148417 | 4,507262 | 6,57E-06 | 1,98E-05 |
| 15 | RM25_RS00075 | RM25_0015 | DNA gyrase, A subunit | 15 | gene | 22411 | 25146 | + | 3901,363 | 0,485618622 | 0,302881 | 1,603333 | 0,108861 | 0,203156 | 4377,323896 | -0,114788333 | 0,160477 | -0,71529 | 0,474427 | 0,541776 | 3425,402 | 1,329319477 | 0,259454 | 5,123536 | 3E-07 | 4,84E-06 | 3457,144 | -1,28055 | 0,227565 | -5,62718 | 1,83E-08 | 1,95E-07 | 4454,123569 | 0,209394863 | 0,169036 | 1,238757 | 0,215436 | 0,271396 |
| 16 | RM25_RS00080 | RM25_0016 | Hypothetical protein | 16 | gene | 25470 | 26252 | + | 390,6406 | 0,844816509 | 0,202793 | 4,165905 | 3,1E-05 | 0,000671 | 416,1133964 | 1,229376298 | 0,236715 | 5,193484 | 2,06E-07 | 1,18E-06 | 365,1677 | 0,440555602 | 0,229634 | 1,918515 | 0,055046 | 0,1083 | 294,2245 | 0,247566 | 0,269207 | 0,919613 | 0,357775 | 0,434007 | 493,2320976 | -0,476065389 | 0,216962 | -2,19423 | 0,028219 | 0,044759 |
| 17 | RM25_RS00085 | RM25_0017 | Hypothetical protein | 17 | gene | 26383 | 26658 | - | 74,78266 | -1,00400397 | 0,528418 | -1,90002 | 0,057431 | 0,13528 | 40,13467982 | -1,32231129 | 0,528363 | -2,50266 | 0,012326 | 0,023714 | 109,4306 | -0,899822806 | 0,380724 | -2,36345 | 0,018106 | 0,047681 | 104,8106 | 1,24873 | 0,492304 | 2,536503 | 0,011197 | 0,025873 | 48,65681879 | 1,71921251 | 0,401667 | 4,280195 | 1,87E-05 | 5,3E-05 |
| 18 | RM25_RS00090 | RM25_0018 | UPF0103 protein | 18 | gene | 26899 | 27696 | + | 696,2216 | -0,189316276 | 0,267855 | -0,70679 | 0,4797 | 0,580119 | 525,1389272 | -0,338785068 | 0,207178 | -1,63524 | 0,101999 | 0,144732 | 867,3043 | -0,101580768 | 0,210552 | -0,48245 | 0,629486 | 0,7136 | 780,2964 | 0,547793 | 0,238559 | 2,296258 | 0,021661 | 0,0436 | 638,3006747 | 0,846301494 | 0,190041 | 4,453252 | 8,46E-06 | 2,5E-05 |
| 19 | RM25_RS00095 | RM25_0019 | AMMECR1 domain protein | 19 | gene | 27761 | 28333 | + | 398,0309 | -0,208870812 | 0,45473 | -0,45933 | 0,645997 | 0,730973 | 226,1050808 | -0,830053351 | 0,286627 | -2,89594 | 0,00378 | 0,0086 | 569,9568 | 0,026767699 | 0,253462 | 0,105608 | 0,915893 | 0,941457 | 447,1227 | 0,899768 | 0,292052 | 3,080851 | 0,002064 | 0,006444 | 362,4457314 | 1,816704097 | 0,251017 | 7,237363 | 4,57E-13 | 4,16E-12 |
| 20 | RM25_RS00100 | RM25_0020 | Pyruvate-formate lyase-activating enzyme | 20 | gene | 28330 | 29418 | + | 532,3125 | -0,165373118 | 0,426453 | -0,38779 | 0,698173 | 0,774762 | 322,0104885 | -0,731264775 | 0,254556 | -2,87271 | 0,00407 | 0,009135 | 742,6145 | 0,071896274 | 0,309875 | 0,232017 | 0,816525 | 0,867695 | 590,4911 | 0,778137 | 0,296918 | 2,620716 | 0,008775 | 0,021277 | 493,1588941 | 1,650770324 | 0,278003 | 5,937968 | 2,89E-09 | 1,5E-08 |
| 21 | RM25_RS00105 | RM25_0021 | Putative membrane protein | 21 | gene | 29575 | 30093 | + | 1641,089 | 0,480949593 | 0,229644 | 2,094328 | 0,036231 | 0,102256 | 1865,616373 | 0,464530293 | 0,193755 | 2,39751 | 0,016507 | 0,030515 | 1416,562 | 0,503844042 | 0,333582 | 1,510405 | 0,13094 | 0,211968 | 1453,757 | -0,46959 | 0,338065 | -1,38906 | 0,164813 | 0,223638 | 1872,692605 | -0,392489429 | 0,183178 | -2,14267 | 0,032139 | 0,050101 |
| 23 | RM25_RS00115 | RM25_0023 | Hypothetical protein | 25 | gene | 30731 | 31411 | - | 543,0507 | -0,310976927 | 0,390204 | -0,79696 | 0,425475 | 0,533996 | 389,3957296 | -1,346516761 | 0,209826 | -6,41731 | 1,39E-10 | 1,33E-09 | 696,7058 | 0,215261565 | 0,210615 | 1,02206 | 0,306752 | 0,410806 | 634,9319 | 0,136049 | 0,245293 | 0,554639 | 0,579142 | 0,641465 | 473,7179382 | 1,756278956 | 0,162206 | 10,82749 | 2,55E-27 | 1,23E-25 |
| 24 | RM25_RS00120 | RM25_0024 | Hypothetical protein | 26 | gene | 31456 | 31740 | - | 367,6255 | -0,83065582 | 0,417847 | -1,98794 | 0,046818 | 0,119523 | 261,6508276 | -1,826366466 | 0,274429 | -6,65515 | 2,83E-11 | 3,05E-10 | 473,6001 | -0,369411416 | 0,225932 | -1,63506 | 0,102037 | 0,17491 | 495,5897 | 0,320078 | 0,276935 | 1,155788 | 0,247768 | 0,317041 | 259,1114483 | 1,838370956 | 0,235367 | 7,810653 | 5,69E-15 | 6,84E-14 |
| 25 | RM25_RS00125 | RM25_0025 | Hypothetical protein | 27 | gene | 31944 | 33524 | - | 598,2911 | -1,104247483 | 0,400559 | -2,75677 | 0,005838 | 0,032838 | 419,8888488 | -1,792041842 | 0,282137 | -6,35168 | 2,13E-10 | 1,99E-09 | 776,6933 | -0,782176832 | 0,269909 | -2,89792 | 0,003756 | 0,014444 | 859,8097 | 0,524853 | 0,31752 | 1,652978 | 0,098335 | 0,146891 | 373,3436365 | 1,601235895 | 0,269433 | 5,942982 | 2,8E-09 | 1,46E-08 |
| 26 | RM25_RS00130 | RM25_0026 | Hypothetical protein | 28 | gene | 33518 | 34039 | - | 253,042 | -1,438211027 | 0,623179 | -2,30786 | 0,021007 | 0,073505 | 150,2539544 | -3,122193302 | 0,402243 | -7,76197 | 8,36E-15 | 1,37E-13 | 355,83 | -0,960335418 | 0,387368 | -2,47913 | 0,01317 | 0,037764 | 387,7084 | 0,730793 | 0,410022 | 1,782327 | 0,074696 | 0,11706 | 134,3338433 | 2,96610374 | 0,415715 | 7,13495 | 9,68E-13 | 8,39E-12 |
| 27 | RM25_RS00135 | RM25_0027 | Hypothetical protein | 29 | gene | 34036 | 34338 | - | 124,8921 | -0,694034221 | 0,246325 | -2,81755 | 0,004839 | 0,028963 | 107,0141939 | -0,803494573 | 0,328309 | -2,44737 | 0,01439 | 0,027134 | 142,7699 | -0,618600818 | 0,285362 | -2,16777 | 0,030176 | 0,069798 | 162,5616 | 0,28312 | 0,234747 | 1,206065 | 0,227793 | 0,295654 | 93,64902232 | 0,530077756 | 0,39276 | 1,349621 | 0,177137 | 0,22893 |
| 28 | RM25_RS00140 | RM25_0028 | Hypothetical protein | 30 | gene | 34468 | 34689 | - | 45,20343 | -1,174407984 | 0,53495 | -2,19536 | 0,028138 | 0,086721 | 30,24546932 | -2,446747728 | 0,613018 | -3,99131 | 6,57E-05 | 0,000234 | 60,16138 | -0,697107089 | 0,278368 | -2,50426 | 0,012271 | 0,035662 | 66,28188 | 0,450743 | 0,366643 | 1,229379 | 0,21893 | 0,28525 | 27,08365154 | 2,27975083 | 0,606651 | 3,757928 | 0,000171 | 0,000418 |
| 29 | RM25_RS00145 | RM25_0029 | Hypothetical protein | 31 | gene | 34686 | 34970 | - | 182,1976 | -1,000158496 | 0,33261 | -3,007 | 0,002638 | 0,018959 | 138,3710938 | -1,401845213 | 0,304591 | -4,60238 | 4,18E-06 | 1,89E-05 | 226,0242 | -0,778327626 | 0,312367 | -2,49171 | 0,012713 | 0,036766 | 255,5555 | 0,442938 | 0,297735 | 1,487694 | 0,136832 | 0,191972 | 119,273826 | 1,130289088 | 0,344329 | 3,282586 | 0,001029 | 0,002172 |
| 30 | RM25_RS00150 | RM25_0030 | PhiRv2 phage integrase | 32 | gene | 35225 | 36421 | - | 879,6899 | -0,890995214 | 0,281013 | -3,17066 | 0,001521 | 0,012787 | 688,3104182 | -1,109410381 | 0,250207 | -4,43397 | 9,25E-06 | 3,91E-05 | 1071,069 | -0,757272519 | 0,248588 | -3,04629 | 0,002317 | 0,009717 | 1202,818 | 0,45011 | 0,294313 | 1,52936 | 0,126175 | 0,179803 | 605,2174843 | 0,866554828 | 0,226397 | 3,827592 | 0,000129 | 0,000323 |
| 31 | RM25_RS00155 | RM25_0031 | Hypothetical protein | 33 | gene | 36751 | 37197 | + | 162,2274 | -1,308029203 | 0,817785 | -1,59948 | 0,109714 | 0,20394 | 77,62814829 | -4,333703722 | 0,503353 | -8,60967 | 7,33E-18 | 1,71E-16 | 246,8267 | -0,810647458 | 0,446674 | -1,81485 | 0,069547 | 0,129787 | 242,1606 | 1,016481 | 0,334705 | 3,036949 | 0,00239 | 0,007335 | 92,22202213 | 4,61510182 | 0,610491 | 7,559656 | 4,04E-14 | 4,35E-13 |
| 32 | RM25_RS00160 | RM25_0032 | Hypothetical protein | 34 | gene | 37212 | 37670 | + | 132,1119 | -1,326922195 | 0,622093 | -2,133 | 0,032925 | 0,095934 | 79,04803692 | -3,036806343 | 0,464558 | -6,53698 | 6,28E-11 | 6,35E-10 | 185,1758 | -0,829222841 | 0,355407 | -2,33316 | 0,01964 | 0,050591 | 199,2088 | 0,672319 | 0,368282 | 1,825554 | 0,067917 | 0,109363 | 74,1087296 | 2,957599325 | 0,506586 | 5,838301 | 5,27E-09 | 2,61E-08 |
| 33 | RM25_RS00165 | RM25_0033 | Hypothetical protein | 35 | gene | 37707 | 38117 | - | 120,0254 | 0,374938043 | 0,322811 | 1,161477 | 0,245448 | 0,354703 | 85,5129946 | -0,167995856 | 0,282457 | -0,59477 | 0,551999 | 0,617391 | 154,5378 | 0,679151618 | 0,264578 | 2,566925 | 0,01026 | 0,030965 | 110,2388 | 0,326437 | 0,211533 | 1,543199 | 0,122782 | 0,175819 | 133,1195995 | 1,238507185 | 0,319856 | 3,872074 | 0,000108 | 0,000272 |
| 34 | RM25_RS00170 | RM25_0034 | Transcriptional regulator, LacI family | 36 | gene | 38390 | 39172 | + | 557,8658 | -0,695678349 | 0,311912 | -2,23037 | 0,025723 | 0,082377 | 396,0238509 | -0,618507153 | 0,166455 | -3,71576 | 0,000203 | 0,000646 | 719,7078 | -0,740271662 | 0,326164 | -2,26963 | 0,02323 | 0,057393 | 725,8026 | 0,847303 | 0,214396 | 3,952045 | 7,75E-05 | 0,000388 | 418,9172268 | 0,787588448 | 0,309631 | 2,543639 | 0,01097 | 0,019027 |
| 35 | RM25_RS13065 | RM25_0035 | Hypothetical protein | 37 | gene | 39307 | 39465 | + | 45,88494 | -0,802659895 | 0,646474 | -1,2416 | 0,214385 | 0,319608 | 26,16168794 | -2,137443948 | 0,685839 | -3,11654 | 0,00183 | 0,004545 | 65,60819 | -0,381052048 | 0,519172 | -0,73396 | 0,462973 | 0,565883 | 61,41239 | 0,715281 | 0,509723 | 1,403275 | 0,160535 | 0,21936 | 33,01020206 | 2,559704604 | 0,719051 | 3,559836 | 0,000371 | 0,000852 |
| 36 | NA | RM25_0036 | putative myo-inositol 2-dehydrogenase | 38 | gene | 39843 | 40226 | + | 198,3303 | -1,28038406 | 0,733254 | -1,74617 | 0,080782 | 0,168108 | 102,2652079 | -3,410409257 | 0,525 | -6,49601 | 8,25E-11 | 8,13E-10 | 294,3954 | -0,813240865 | 0,451645 | -1,80062 | 0,071763 | 0,13329 | 294,5425 | 0,932986 | 0,475206 | 1,963329 | 0,049608 | 0,084362 | 114,2285415 | 3,607294604 | 0,544351 | 6,626776 | 3,43E-11 | 2,38E-10 |
| 37 | RM25_RS00175 | RM25_0037 | Oxidoreductase, NAD-binding domain protein | 39 | gene | 40223 | 41032 | + | 285,4405 | -1,571455455 | 0,747821 | -2,10138 | 0,035608 | 0,1008 | 157,820989 | -4,065413414 | 0,445753 | -9,12033 | 7,49E-20 | 2,23E-18 | 413,06 | -1,046257807 | 0,452251 | -2,31344 | 0,020698 | 0,052628 | 448,5472 | 0,830414 | 0,408939 | 2,030655 | 0,04229 | 0,07407 | 142,0492322 | 3,927878118 | 0,527491 | 7,446342 | 9,6E-14 | 9,67E-13 |
| 38 | RM25_RS00180 | RM25_0038 | MFS transporter, sugar porter family protein | 40 | gene | 41049 | 42701 | + | 585,4724 | -1,413031639 | 0,556701 | -2,53822 | 0,011142 | 0,050223 | 376,3796709 | -2,761237185 | 0,353016 | -7,82184 | 5,21E-15 | 8,64E-14 | 794,5652 | -0,948660234 | 0,377969 | -2,50989 | 0,012077 | 0,035224 | 895,9076 | 0,603183 | 0,392307 | 1,537529 | 0,124164 | 0,177474 | 315,1855648 | 2,491553653 | 0,381973 | 6,522856 | 6,9E-11 | 4,61E-10 |
| 39 | RM25_RS00185 | RM25_0039 | Regulatory protein C | 41 | gene | 42915 | 43574 | - | 250,2384 | -0,436891373 | 0,306205 | -1,4268 | 0,153639 | 0,25466 | 196,0635283 | -0,575032756 | 0,300485 | -1,91368 | 0,055661 | 0,086197 | 304,4132 | -0,352454124 | 0,34825 | -1,01207 | 0,311504 | 0,41528 | 303,4759 | 0,46683 | 0,289713 | 1,611352 | 0,107103 | 0,157089 | 209,0892028 | 0,763162007 | 0,380868 | 2,003744 | 0,045097 | 0,067489 |
| 40 | RM25_RS00190 | RM25_0040 | Two-component system sensor kinase | 42 | gene | 43594 | 45072 | - | 825,9591 | -0,252814033 | 0,331559 | -0,7625 | 0,445762 | 0,553853 | 592,8522652 | -0,699148811 | 0,191258 | -3,65553 | 0,000257 | 0,000803 | 1059,066 | -0,011389056 | 0,305257 | -0,03731 | 0,970238 | 0,982746 | 944,5291 | 0,469127 | 0,242898 | 1,931378 | 0,053436 | 0,089644 | 740,962866 | 1,222065751 | 0,266116 | 4,592238 | 4,39E-06 | 1,37E-05 |
| 41 | RM25_RS00195 | RM25_0041 | Alpha-ketoglutarate transporter, MFS super | 43 | gene | 45274 | 46608 | - | 1163,499 | -0,08818964 | 0,269266 | -0,32752 | 0,743276 | 0,813325 | 1456,284648 | -0,286793784 | 0,167192 | -1,71536 | 0,086279 | 0,124693 | 870,7133 | 0,243572939 | 0,186979 | 1,302677 | 0,192685 | 0,288538 | 1276,289 | -1,06737 | 0,181611 | -5,87724 | 4,17E-09 | 4,99E-08 | 1107,203977 | -0,478847959 | 0,181761 | -2,63449 | 0,008426 | 0,014944 |
| 42 | RM25_RS00200 | RM25_0042 | Putative hippurate hydrolase | 44 | gene | 46627 | 47838 | - | 1377,924 | -0,51873708 | 0,306127 | -1,69452 | 0,090167 | 0,180895 | 1792,158243 | -0,692007869 | 0,185645 | -3,72759 | 0,000193 | 0,00062 | 963,6899 | -0,204159076 | 0,232097 | -0,87963 | 0,379061 | 0,484516 | 1727,899 | -1,16161 | 0,218396 | -5,31881 | 1,04E-07 | 9,73E-07 | 1112,133346 | -0,618384598 | 0,208349 | -2,96803 | 0,002997 | 0,005776 |
| 43 | RM25_RS00205 | RM25_0043 | Bifunctional ABC transporter | 45 | gene | 48375 | 50267 | + | 676,7796 | -0,209549232 | 0,388876 | -0,53886 | 0,589984 | 0,680945 | 441,2885826 | -0,502523812 | 0,197076 | -2,5499 | 0,010775 | 0,021182 | 912,2707 | -0,071217359 | 0,393062 | -0,18119 | 0,856221 | 0,895348 | 760,4763 | 0,789929 | 0,235915 | 3,348363 | 0,000813 | 0,002943 | 618,6206502 | 1,289210164 | 0,358169 | 3,599447 | 0,000319 | 0,000743 |
| 44 | RM25_RS00210 | RM25_0044 | ABC-type multidrug transport system ATPase and permease | 46 | gene | 50267 | 52234 | + | 766,7972 | -0,227956811 | 0,391397 | -0,58242 | 0,560285 | 0,654731 | 499,4890166 | -0,505483412 | 0,172751 | -2,92608 | 0,003433 | 0,007924 | 1034,105 | -0,097225377 | 0,411299 | -0,23639 | 0,813133 | 0,865899 | 866,4444 | 0,801987 | 0,244773 | 3,276447 | 0,001051 | 0,003671 | 696,1881288 | 1,279854485 | 0,361105 | 3,54427 | 0,000394 | 0,000896 |
| 45 | RM25_RS00215 | RM25_0045 | Alanine dehydrogenase | 47 | gene | 52458 | 53573 | + | 7343,994 | -5,555126534 | 0,981728 | -5,65852 | 1,53E-08 | 1,29E-06 | 328,0167051 | -2,636925454 | 0,335206 | -7,86658 | 3,64E-15 | 6,22E-14 | 14359,97 | -5,759271093 | 0,48891 | -11,7798 | 4,96E-32 | 2,92E-29 | 15009,28 | 5,59016 | 0,537811 | 10,39428 | 2,63E-25 | 4,98E-23 | 300,342025 | 2,516213828 | 0,282405 | 8,909944 | 5,11E-19 | 1,07E-17 |
| 46 | RM25_RS00220 | RM25_0046 | D-serine/D-alanine/glycine transporter | 48 | gene | 53941 | 55479 | + | 2559,453 | -3,204145749 | 0,639857 | -5,00759 | 5,51E-07 | 2,45E-05 | 668,007018 | -1,900248915 | 0,235482 | -8,06962 | 7,05E-16 | 1,27E-14 | 4450,9 | -3,507304541 | 0,397675 | -8,81954 | 1,15E-18 | 1,35E-16 | 4829,428 | 2,903496 | 0,405388 | 7,162272 | 7,94E-13 | 2,08E-11 | 492,5731865 | 1,349621481 | 0,264418 | 5,104115 | 3,32E-07 | 1,25E-06 |
| 47 | RM25_RS00225 | RM25_0047 | Putative conserved lipoprotein lppS | 49 | gene | 55696 | 56895 | - | 1411,08 | 0,971680661 | 0,252469 | 3,848708 | 0,000119 | 0,00193 | 1177,205001 | 0,246614807 | 0,128897 | 1,913272 | 0,055713 | 0,086222 | 1644,955 | 1,568534943 | 0,225604 | 6,952601 | 3,59E-12 | 1,8E-10 | 1008,99 | -0,43702 | 0,146458 | -2,98394 | 0,002846 | 0,008468 | 1826,030745 | 0,932661999 | 0,167262 | 5,576056 | 2,46E-08 | 1,11E-07 |
| 48 | NA | RM25_0048 | Hypothetical protein | 50 | gene | 56910 | 57047 | + | 56,13164 | -1,387788514 | 0,815826 | -1,70108 | 0,088927 | 0,179363 | 28,54456637 | -4,148794804 | 0,856675 | -4,8429 | 1,28E-06 | 6,38E-06 | 83,71872 | -0,877798651 | 0,374046 | -2,34676 | 0,018937 | 0,04943 | 85,22469 | 0,926902 | 0,561229 | 1,651557 | 0,098625 | 0,147125 | 30,50597048 | 4,267108384 | 0,757285 | 5,634747 | 1,75E-08 | 8,07E-08 |
| 49 | RM25_RS00230 | RM25_0049 | Putative acylphosphatase | 51 | gene | 57141 | 57479 | + | 208,0839 | -0,540467902 | 0,485109 | -1,11412 | 0,265229 | 0,372776 | 114,4892116 | -1,19560015 | 0,296511 | -4,03223 | 5,53E-05 | 0,000202 | 301,6786 | -0,312025717 | 0,307037 | -1,01625 | 0,309512 | 0,413537 | 258,6184 | 0,999063 | 0,237842 | 4,200528 | 2,66E-05 | 0,000149 | 166,6180395 | 1,949229727 | 0,35666 | 5,465232 | 4,62E-08 | 2,02E-07 |
| 50 | RM25_RS00235 | RM25_0050 | IMPACT family member yvyE | 52 | gene | 57526 | 58209 | - | 362,326 | -0,161365635 | 0,361987 | -0,44578 | 0,655758 | 0,738119 | 240,1972318 | -0,217264702 | 0,196172 | -1,10752 | 0,268068 | 0,333775 | 484,4548 | -0,136109274 | 0,360217 | -0,37785 | 0,705539 | 0,779998 | 400,9869 | 0,912523 | 0,186704 | 4,887529 | 1,02E-06 | 7,81E-06 | 336,9099422 | 1,058356143 | 0,354603 | 2,98462 | 0,002839 | 0,00549 |
| 51 | RM25_RS00240 | RM25_0051 | Nitroreductase | 53 | gene | 58340 | 58849 | + | 884,5537 | -0,073744335 | 0,356681 | -0,20675 | 0,836204 | 0,891018 | 607,2049427 | 0,530055565 | 0,156172 | 3,394058 | 0,000689 | 0,001923 | 1161,903 | -0,387059524 | 0,324198 | -1,1939 | 0,232518 | 0,330718 | 951,6335 | 1,352464 | 0,31988 | 4,228035 | 2,36E-05 | 0,000133 | 843,4643037 | 0,478641641 | 0,149491 | 3,201819 | 0,001366 | 0,002801 |
| 52 | RM25_RS00245 | RM25_0052 | Conserved transmembrane protein | 54 | gene | 59044 | 60735 | + | 1699,364 | 0,335792656 | 0,142965 | 2,348781 | 0,018835 | 0,069555 | 1869,992934 | 0,412140383 | 0,143943 | 2,863226 | 0,004194 | 0,00936 | 1528,735 | 0,243689744 | 0,205173 | 1,187726 | 0,234941 | 0,333589 | 1587,97 | -0,26036 | 0,179198 | -1,45294 | 0,146241 | 0,20264 | 1860,294388 | -0,372311894 | 0,155353 | -2,39655 | 0,01655 | 0,027676 |
| 53 | RM25_RS00250 | RM25_0053 | Hypothetical protein | 55 | gene | 60773 | 61351 | - | 221,5755 | -0,872865793 | 0,481189 | -1,81398 | 0,069681 | 0,15335 | 135,4393988 | -1,627079425 | 0,306313 | -5,31182 | 1,09E-07 | 6,54E-07 | 307,7116 | -0,581186371 | 0,400267 | -1,452 | 0,146502 | 0,230974 | 301,1371 | 0,78005 | 0,270653 | 2,882106 | 0,00395 | 0,010915 | 154,2675365 | 1,900505985 | 0,441916 | 4,3006 | 1,7E-05 | 4,85E-05 |
| 54 | RM25_RS00255 | RM25_0054 | Putative membrane protein | 56 | gene | 61324 | 62340 | - | 572,663 | -0,120771822 | 0,181155 | -0,66668 | 0,504978 | 0,60204 | 664,2255877 | -0,199077544 | 0,15538 | -1,28123 | 0,200114 | 0,260016 | 481,1004 | -0,010391155 | 0,181135 | -0,05737 | 0,954253 | 0,971566 | 632,7372 | -0,61672 | 0,178983 | -3,44569 | 0,00057 | 0,002197 | 538,1555524 | -0,372232663 | 0,15363 | -2,42292 | 0,015396 | 0,025903 |
| 55 | RM25_RS00260 | RM25_0055 | putative ATP-dependent serine protease | 57 | gene | 62337 | 64427 | - | 1728,037 | -0,25895614 | 0,124866 | -2,07386 | 0,038092 | 0,105876 | 1671,269173 | -0,440137105 | 0,122611 | -3,5897 | 0,000331 | 0,001011 | 1784,805 | -0,092481357 | 0,221329 | -0,41785 | 0,67606 | 0,754582 | 1986,633 | -0,12502 | 0,183298 | -0,68207 | 0,495197 | 0,563039 | 1543,817765 | 0,280610708 | 0,154056 | 1,82149 | 0,068532 | 0,098255 |
| 56 | RM25_RS00265 | RM25_0056 | Glucose-1-phosphate thymidylyltransferase | 58 | gene | 64483 | 65352 | - | 1096,526 | 0,119570261 | 0,112571 | 1,062181 | 0,288153 | 0,397645 | 1108,371618 | 0,0181882 | 0,150207 | 0,121088 | 0,903622 | 0,926822 | 1084,681 | 0,222264232 | 0,21015 | 1,057644 | 0,290218 | 0,394202 | 1113,64 | -0,19501 | 0,211532 | -0,92187 | 0,356595 | 0,433244 | 1117,639675 | 0,056043851 | 0,14224 | 0,394009 | 0,693574 | 0,743146 |
| 57 | RM25_RS00270 | RM25_0057 | Glycosyltransferase, group 2 family protein | 59 | gene | 65516 | 66262 | + | 1105,038 | 0,459066392 | 0,224364 | 2,046081 | 0,040748 | 0,109639 | 1323,693736 | 0,714866675 | 0,159116 | 4,492742 | 7,03E-06 | 3,02E-05 | 886,3822 | 0,090834551 | 0,187586 | 0,48423 | 0,628223 | 0,713257 | 985,2469 | -0,28257 | 0,175405 | -1,61096 | 0,107187 | 0,157115 | 1254,511735 | -0,858481961 | 0,162071 | -5,29694 | 1,18E-07 | 4,75E-07 |
| 58 | RM25_RS00275 | RM25_0058 | Hypothetical protein | 60 | gene | 66259 | 66684 | + | 486,8891 | 0,086577685 | 0,119139 | 0,726696 | 0,467412 | 0,569305 | 486,4981514 | 0,184865052 | 0,186375 | 0,9919 | 0,321246 | 0,389094 | 487,28 | -0,009001072 | 0,200489 | -0,0449 | 0,964191 | 0,979146 | 497,4366 | 0,047871 | 0,160243 | 0,298742 | 0,765137 | 0,808349 | 492,172662 | -0,095549461 | 0,20396 | -0,46847 | 0,639447 | 0,695135 |
| 59 | RM25_RS00280 | RM25_0059 | Alpha-1, 6-rhamnosyltransferase MigA | 61 | gene | 66823 | 67704 | + | 1560,941 | 0,232005688 | 0,154031 | 1,506223 | 0,13201 | 0,230139 | 1715,836545 | 0,356209106 | 0,152925 | 2,329307 | 0,019843 | 0,035458 | 1406,046 | 0,083043437 | 0,219082 | 0,379053 | 0,704649 | 0,779424 | 1519,133 | -0,19495 | 0,213378 | -0,91364 | 0,360907 | 0,437131 | 1651,927743 | -0,424647215 | 0,145728 | -2,91398 | 0,003569 | 0,006767 |
| 60 | RM25_RS00285 | RM25_0060 | Cps9H protein | 62 | gene | 67715 | 68506 | + | 849,3046 | 0,158670927 | 0,21754 | 0,729387 | 0,465765 | 0,568518 | 1024,857823 | 0,112510783 | 0,136461 | 0,824487 | 0,409663 | 0,478244 | 673,7514 | 0,229892666 | 0,186202 | 1,23464 | 0,216965 | 0,314312 | 850,4519 | -0,72586 | 0,149138 | -4,86707 | 1,13E-06 | 8,6E-06 | 878,0820617 | -0,560044218 | 0,127378 | -4,3967 | 1,1E-05 | 3,2E-05 |
| 61 | RM25_RS11630 | RM25_0061 | Family 2 glycosyl transferase | 63 | gene | 68521 | 70428 | + | 1539,766 | 0,23053443 | 0,128116 | 1,799416 | 0,071953 | 0,156596 | 1652,957686 | 0,119180843 | 0,130352 | 0,914297 | 0,360561 | 0,428996 | 1426,575 | 0,360518306 | 0,201991 | 1,784823 | 0,07429 | 0,136585 | 1498,213 | -0,40402 | 0,178364 | -2,26512 | 0,023505 | 0,046596 | 1630,982009 | -0,106102406 | 0,13313 | -0,79698 | 0,425461 | 0,491333 |
| 62 | RM25_RS00300 | RM25_0062 | Polysaccharide biosynthesis protein | 64 | gene | 70416 | 71693 | - | 1039,766 | -0,592815496 | 0,164927 | -3,5944 | 0,000325 | 0,004211 | 1001,692421 | -0,70777773 | 0,153081 | -4,62355 | 3,77E-06 | 1,73E-05 | 1077,839 | -0,488391997 | 0,295905 | -1,6505 | 0,098841 | 0,171174 | 1319,096 | -0,04355 | 0,17621 | -0,24717 | 0,80478 | 0,842678 | 814,9795204 | 0,239824292 | 0,268742 | 0,892396 | 0,372181 | 0,437085 |
| 63 | RM25_RS00305 | RM25_0063 | Putative membrane protein | 65 | gene | 71941 | 73131 | - | 1056,523 | 5,09885E-05 | 0,115965 | 0,00044 | 0,999649 | 0,999649 | 1014,173455 | -0,234534021 | 0,131956 | -1,77736 | 0,075509 | 0,111443 | 1098,873 | 0,214513605 | 0,169306 | 1,26702 | 0,205148 | 0,30202 | 1118,402 | -0,16615 | 0,168104 | -0,98839 | 0,322961 | 0,398336 | 1033,879963 | 0,331537384 | 0,117312 | 2,826107 | 0,004712 | 0,008758 |
| 64 | RM25_RS00310 | RM25_0064 | DTDP-4-keto-L-rhamnose reductase and dTDP-4-keto-6-deo | 66 | gene | 73627 | 75087 | - | 2609,45 | 0,277849494 | 0,276649 | 1,00434 | 0,315215 | 0,424792 | 3332,803356 | 0,231540819 | 0,158018 | 1,465285 | 0,142843 | 0,195508 | 1886,097 | 0,360909035 | 0,14741 | 2,448337 | 0,014352 | 0,040032 | 2504,092 | -0,94892 | 0,172885 | -5,4887 | 4,05E-08 | 4,04E-07 | 2804,093688 | -0,772940793 | 0,105979 | -7,29334 | 3,02E-13 | 2,83E-12 |
| 65 | RM25_RS00315 | RM25_0065 | DTDP-glucose 4, 6-dehydratase (Precursor) | 67 | gene | 75092 | 76090 | - | 2168,004 | 0,220640338 | 0,190759 | 1,156642 | 0,247419 | 0,355806 | 2477,555431 | 0,394903047 | 0,135262 | 2,919538 | 0,003506 | 0,008085 | 1858,452 | -0,007926413 | 0,244769 | -0,03238 | 0,974166 | 0,985032 | 2120,21 | -0,25575 | 0,203449 | -1,25706 | 0,208733 | 0,273414 | 2286,708018 | -0,613777648 | 0,175908 | -3,48921 | 0,000484 | 0,00109 |
| 66 | RM25_RS11635 | RM25_0066 | Large surface protein A | 68 | gene | 76221 | 80255 | - | 7136,037 | -0,096377484 | 0,404974 | -0,23798 | 0,811893 | 0,869438 | 10147,63233 | -0,167997065 | 0,158087 | -1,06269 | 0,287925 | 0,355681 | 4124,442 | 0,080114642 | 0,242202 | 0,330776 | 0,740814 | 0,808753 | 7857,476 | -1,47638 | 0,240638 | -6,13528 | 8,5E-10 | 1,22E-08 | 6763,710412 | -1,184694599 | 0,134237 | -8,8254 | 1,09E-18 | 2,2E-17 |
| 67 | RM25_RS00335 | RM25_0067 | UDP-galactopyranose mutase | 69 | gene | 80548 | 81735 | + | 5504,01 | 0,833296703 | 0,302131 | 2,75806 | 0,005815 | 0,032787 | 7011,749974 | 0,809235262 | 0,15624 | 5,179437 | 2,23E-07 | 1,27E-06 | 3996,269 | 0,876394203 | 0,311598 | 2,812583 | 0,004915 | 0,017626 | 4208,455 | -0,90944 | 0,227945 | -3,98975 | 6,61E-05 | 0,000339 | 6901,080391 | -0,804146808 | 0,234607 | -3,42764 | 0,000609 | 0,001349 |
| 68 | RM25_RS00340 | RM25_0068 | Bifunctional UDP-galactofuranosyl transferase glfT | 70 | gene | 81732 | 83744 | + | 5734,109 | 1,054519042 | 0,428646 | 2,460115 | 0,013889 | 0,057942 | 7912,073269 | 0,827333026 | 0,175229 | 4,721445 | 2,34E-06 | 1,11E-05 | 3556,145 | 1,615990526 | 0,29971 | 5,391848 | 6,97E-08 | 1,26E-06 | 3975,151 | -1,76002 | 0,254782 | -6,90796 | 4,92E-12 | 1,13E-10 | 7583,404977 | -0,930364157 | 0,2083 | -4,46646 | 7,95E-06 | 2,36E-05 |
| 69 | RM25_RS00345 | RM25_0069 | ABC-2 type transport system integral membrane protein | 71 | gene | 83749 | 84633 | + | 2050,226 | 0,926759911 | 0,430332 | 2,153595 | 0,031272 | 0,092882 | 2921,723389 | 0,855619835 | 0,18462 | 4,634501 | 3,58E-06 | 1,65E-05 | 1178,728 | 1,110550391 | 0,310108 | 3,581168 | 0,000342 | 0,002078 | 1506,977 | -1,53312 | 0,238485 | -6,42859 | 1,29E-10 | 2,23E-09 | 2631,270474 | -1,241699031 | 0,240342 | -5,16639 | 2,39E-07 | 9,24E-07 |
| 70 | RM25_RS00350 | RM25_0070 | O-antigen ABC transporter ATP-binding protein | 72 | gene | 84633 | 85403 | + | 1756,89 | 0,820149006 | 0,324853 | 2,524677 | 0,01158 | 0,051402 | 2343,029008 | 0,831893269 | 0,13547 | 6,140801 | 8,21E-10 | 7,06E-09 | 1170,75 | 0,79795491 | 0,223481 | 3,57057 | 0,000356 | 0,002147 | 1348,811 | -1,0384 | 0,166988 | -6,21841 | 5,02E-10 | 7,64E-09 | 2199,025878 | -1,024887666 | 0,151846 | -6,74952 | 1,48E-11 | 1,08E-10 |
| 71 | RM25_RS11640 | RM25_0071 | Glycosyltransferase | 73 | gene | 85516 | 87924 | - | 1673,966 | -0,108450417 | 0,105489 | -1,02807 | 0,303916 | 0,413586 | 1706,202182 | -0,077776176 | 0,115125 | -0,67558 | 0,499308 | 0,565147 | 1641,73 | -0,139712191 | 0,225632 | -0,6192 | 0,535782 | 0,633319 | 1832,742 | -0,08823 | 0,168099 | -0,5249 | 0,599656 | 0,660462 | 1581,959575 | -0,090750841 | 0,174515 | -0,52002 | 0,603052 | 0,662654 |
| 72 | RM25_RS00360 | RM25_0072 | Amylovoran biosynthesis glycosyltransferase AmsE | 74 | gene | 88100 | 88957 | - | 759,0792 | -0,327873025 | 0,151691 | -2,16146 | 0,03066 | 0,092058 | 681,727354 | -0,143015045 | 0,153823 | -0,92974 | 0,352506 | 0,421113 | 836,4311 | -0,478826005 | 0,209378 | -2,2869 | 0,022202 | 0,055669 | 891,4583 | 0,389017 | 0,218087 | 1,783774 | 0,07446 | 0,117002 | 660,321606 | 0,105696757 | 0,159318 | 0,663434 | 0,507052 | 0,568294 |
| 73 | RM25_RS00365 | RM25_0073 | DTDP-rhamnosyl transferase RfbF | 75 | gene | 88954 | 89916 | - | 493,2174 | -0,544771463 | 0,286269 | -1,90301 | 0,05704 | 0,134847 | 364,7365402 | -0,53519654 | 0,192813 | -2,77573 | 0,005508 | 0,011943 | 621,6982 | -0,551931366 | 0,289071 | -1,90933 | 0,05622 | 0,110058 | 614,8692 | 0,71491 | 0,18246 | 3,918174 | 8,92E-05 | 0,000439 | 394,4037315 | 0,759551391 | 0,291798 | 2,603 | 0,009241 | 0,016267 |
| 74 | RM25_RS00370 | RM25_0074 | ThiX protein | 76 | gene | 90140 | 90556 | - | 268,6311 | -0,30325012 | 0,34458 | -0,88006 | 0,378829 | 0,48926 | 202,2335561 | -0,332579998 | 0,353889 | -0,93979 | 0,347327 | 0,416468 | 335,0287 | -0,287458726 | 0,367419 | -0,78237 | 0,433996 | 0,538951 | 311,0007 | 0,641908 | 0,364424 | 1,761434 | 0,078165 | 0,121128 | 236,7046739 | 0,755522937 | 0,374259 | 2,018714 | 0,043517 | 0,065498 |
| 75 | RM25_RS00375 | RM25_0075 | Integral membrane protein | 77 | gene | 90561 | 91070 | - | 207,7511 | -0,035706164 | 0,393209 | -0,09081 | 0,927646 | 0,951049 | 139,5606465 | -0,122345906 | 0,328975 | -0,3719 | 0,709967 | 0,760288 | 275,9416 | 0,004280356 | 0,403518 | 0,010608 | 0,991537 | 0,995359 | 220,6408 | 0,847232 | 0,352948 | 2,400443 | 0,016375 | 0,035069 | 202,1881765 | 1,051052375 | 0,398576 | 2,637016 | 0,008364 | 0,014845 |
| 76 | RM25_RS00380 | RM25_0076 | Acyltransferase | 78 | gene | 92189 | 92983 | - | 804,5744 | 0,62470583 | 0,190509 | 3,279139 | 0,001041 | 0,010209 | 924,1327389 | 0,669248424 | 0,214004 | 3,127271 | 0,001764 | 0,004396 | 685,0161 | 0,566152882 | 0,179198 | 3,159369 | 0,001581 | 0,007098 | 669,4396 | -0,4267 | 0,234506 | -1,81956 | 0,068827 | 0,110357 | 957,7320698 | -0,476552582 | 0,163728 | -2,91063 | 0,003607 | 0,006823 |
| 77 | RM25_RS00385 | RM25_0077 | Family 2 glycosyl transferase | 79 | gene | 92983 | 93879 | - | 1539,128 | 0,39138802 | 0,199013 | 1,966647 | 0,049224 | 0,123341 | 1780,02194 | 0,417220988 | 0,190058 | 2,195232 | 0,028147 | 0,04829 | 1298,235 | 0,357374363 | 0,221225 | 1,615436 | 0,106216 | 0,180369 | 1412,483 | -0,47703 | 0,253275 | -1,88344 | 0,059641 | 0,098003 | 1710,575678 | -0,493686318 | 0,142326 | -3,46871 | 0,000523 | 0,001171 |
| 78 | RM25_RS00390 | RM25_0078 | Acetate kinase | 80 | gene | 93892 | 95082 | - | 2887,943 | 0,564013328 | 0,497456 | 1,133794 | 0,256881 | 0,364457 | 3751,917333 | -0,129909658 | 0,189754 | -0,68462 | 0,493584 | 0,559863 | 2023,969 | 2,160020019 | 0,296501 | 7,285028 | 3,22E-13 | 1,99E-11 | 2494,5 | -2,4666 | 0,230748 | -10,6896 | 1,14E-26 | 2,98E-24 | 3367,801229 | -0,13229285 | 0,228024 | -0,58017 | 0,561799 | 0,623135 |
| 79 | RM25_RS00395 | RM25_0079 | Phospho acetyltransferase | 81 | gene | 95243 | 96775 | - | 2080,277 | 0,027236287 | 0,1565 | 0,174034 | 0,861839 | 0,907665 | 1874,765555 | -0,203617716 | 0,167206 | -1,21776 | 0,223314 | 0,28606 | 2285,789 | 0,21583147 | 0,172399 | 1,251928 | 0,210596 | 0,307734 | 2175,782 | 0,014238 | 0,1503 | 0,09473 | 0,924529 | 0,940084 | 2055,7121 | 0,485223399 | 0,17118 | 2,834585 | 0,004589 | 0,008556 |
| 80 | NA | RM25_0080 | Hypothetical protein | 82 | gene | 97142 | 97231 | + | 30,49216 | 0,05593249 | 0,473879 | 0,118031 | 0,906043 | 0,939526 | 18,8003581 | -0,730551335 | 0,645763 | -1,1313 | 0,257929 | 0,322857 | 42,18396 | 0,394342727 | 0,280358 | 1,406567 | 0,159556 | 0,247164 | 31,49721 | 0,548475 | 0,377848 | 1,451576 | 0,14662 | 0,203005 | 30,37568237 | 1,741243081 | 0,531687 | 3,274941 | 0,001057 | 0,002218 |
| 81 | RM25_RS00400 | RM25_0081 | Transposase IS204/IS1001/IS1096/IS1165 family protein | 83 | gene | 97375 | 98682 | - | 22,40396 | -1,744756368 | 0,8825 | -1,97706 | 0,048035 | 0,121609 | 14,93323525 | -6,149647368 | 1,335812 | -4,60368 | 4,15E-06 | 1,88E-05 | 29,87469 | -0,974691646 | 0,463333 | -2,10365 | 0,035409 | 0,078586 | 36,40972 | 0,35327 | 0,34731 | 1,017159 | 0,309078 | 0,384101 | 10,07412406 | 5,596187511 | 1,520866 | 3,679607 | 0,000234 | 0,000557 |
| 82 | RM25_RS00405 | RM25_0082 | Hypothetical protein | 84 | gene | 98817 | 99272 | + | 184,5209 | -0,764520203 | 0,344687 | -2,21802 | 0,026554 | 0,084154 | 136,2351269 | -1,312212289 | 0,298994 | -4,38875 | 1,14E-05 | 4,72E-05 | 232,8066 | -0,474540585 | 0,252164 | -1,88187 | 0,059853 | 0,115729 | 245,011 | 0,409773 | 0,262366 | 1,561834 | 0,118327 | 0,170475 | 134,1839621 | 1,313103934 | 0,32011 | 4,102035 | 4,1E-05 | 0,00011 |
| 83 | RM25_RS00410 | RM25_0083 | Transposase IS204/IS1001/IS1096/IS1165 family protein | 85 | gene | 99413 | 100705 | - | 591,8099 | -1,190329239 | 0,62173 | -1,91454 | 0,055551 | 0,132792 | 336,0900955 | -2,762773078 | 0,350649 | -7,87902 | 3,3E-15 | 5,68E-14 | 847,5298 | -0,740384862 | 0,43303 | -1,70978 | 0,087307 | 0,154957 | 863,0235 | 0,786529 | 0,390738 | 2,012932 | 0,044122 | 0,077033 | 356,0084248 | 2,88478977 | 0,419952 | 6,869329 | 6,45E-12 | 4,9E-11 |
| 84 | RM25_RS00415 | RM25_0084 | Hypothetical protein | 86 | gene | 100968 | 101336 | - | 367,64 | -0,659714693 | 0,35654 | -1,85032 | 0,064267 | 0,145232 | 252,4709397 | -0,961859265 | 0,27528 | -3,49411 | 0,000476 | 0,001389 | 482,809 | -0,511641516 | 0,274651 | -1,86288 | 0,062479 | 0,119922 | 472,5617 | 0,697366 | 0,311099 | 2,241621 | 0,024986 | 0,048751 | 279,5461106 | 1,21189232 | 0,244621 | 4,954156 | 7,26E-07 | 2,61E-06 |
| 85 | RM25_RS00420 | RM25_0085 | Hypothetical protein | 87 | gene | 101438 | 102328 | - | 798,8824 | -0,607704673 | 0,261625 | -2,32281 | 0,020189 | 0,072329 | 637,6280169 | -0,98032801 | 0,207453 | -4,72554 | 2,3E-06 | 1,1E-05 | 960,1368 | -0,374709895 | 0,196061 | -1,91119 | 0,055981 | 0,109681 | 1016,466 | 0,291113 | 0,214813 | 1,35519 | 0,175357 | 0,236117 | 619,4567354 | 0,953221722 | 0,189164 | 5,03913 | 4,68E-07 | 1,73E-06 |
| 86 | RM25_RS00425 | RM25_0086 | Transposase | 88 | gene | 102512 | 103891 | + | 29,74117 | -0,781992767 | 0,533139 | -1,46677 | 0,142438 | 0,24328 | 18,91105537 | -1,389660668 | 0,766406 | -1,81322 | 0,069798 | 0,104057 | 40,57129 | -0,547023471 | 0,35582 | -1,53736 | 0,124206 | 0,2033 | 39,58402 | 0,718307 | 0,576489 | 1,246003 | 0,212763 | 0,278447 | 21,39238677 | 1,650271861 | 0,573292 | 2,87859 | 0,003995 | 0,007502 |
| 87 | RM25_RS00430 | RM25_0087 | Putative cell wall protein | 89 | gene | 103834 | 105066 | - | 1276,047 | -1,086520763 | 0,468767 | -2,31783 | 0,020459 | 0,072518 | 650,767613 | -0,604713561 | 0,209489 | -2,88661 | 0,003894 | 0,008792 | 1901,326 | -1,265566004 | 0,363423 | -3,48235 | 0,000497 | 0,002783 | 1818,34 | 1,716823 | 0,371277 | 4,624107 | 3,76E-06 | 2,55E-05 | 798,0895638 | 1,098764845 | 0,193809 | 5,669318 | 1,43E-08 | 6,67E-08 |
| 88 | RM25_RS11645 | RM25_0088 | Transposase IS204/IS1001/IS1096/IS1165 family protein | 90 | gene | 105634 | 106983 | + | 542,0717 | -0,658227397 | 0,512734 | -1,28376 | 0,199226 | 0,302149 | 338,1505789 | -2,055709874 | 0,281012 | -7,31538 | 2,57E-13 | 3,64E-12 | 745,9927 | -0,142766074 | 0,233417 | -0,61163 | 0,54078 | 0,638109 | 698,0122 | 0,453114 | 0,325484 | 1,392124 | 0,163885 | 0,222638 | 411,5009099 | 2,427722463 | 0,170428 | 14,24486 | 4,83E-46 | 8,75E-44 |
| 89 | RM25_RS00440 | RM25_0089 | Hypothetical protein | 91 | gene | 107075 | 107647 | + | 451,9969 | -1,499415226 | 0,514129 | -2,91642 | 0,003541 | 0,023247 | 296,6455592 | -2,583544209 | 0,365931 | -7,0602 | 1,66E-12 | 2,13E-11 | 607,3482 | -1,096039579 | 0,33202 | -3,30113 | 0,000963 | 0,004699 | 701,4282 | 0,632499 | 0,3943 | 1,604105 | 0,108691 | 0,158825 | 232,3354531 | 2,191597132 | 0,347333 | 6,309786 | 2,79E-10 | 1,66E-09 |
| 90 | NA | RM25_0090 | Hypothetical protein | 92 | gene | 107861 | 107998 | + | 30,79201 | -0,706013505 | 0,389607 | -1,81212 | 0,069968 | 0,153432 | 22,27124455 | -1,215946841 | 0,526562 | -2,30922 | 0,020931 | 0,037122 | 39,31278 | -0,457058336 | 0,373524 | -1,22364 | 0,221089 | 0,31833 | 40,26142 | 0,480069 | 0,33558 | 1,430567 | 0,152554 | 0,210241 | 22,88837109 | 1,304374019 | 0,582633 | 2,238757 | 0,025172 | 0,04036 |
| 91 | RM25_RS00445 | RM25_0091 | Transposase IS204/IS1001/IS1096/IS1165 family protein | 93 | gene | 108342 | 109418 | + | 48,90257 | -1,749128637 | 0,843221 | -2,07434 | 0,038048 | 0,105876 | 26,40131214 | -4,70434379 | 0,978274 | -4,80882 | 1,52E-06 | 7,49E-06 | 71,40383 | -1,224982601 | 0,419031 | -2,92337 | 0,003463 | 0,01358 | 79,02951 | 0,899658 | 0,596085 | 1,509277 | 0,131228 | 0,185768 | 22,04654046 | 4,455983814 | 0,921544 | 4,835346 | 1,33E-06 | 4,53E-06 |
| 92 | RM25_RS00450 | RM25_0092 | Hypothetical protein | 94 | gene | 109646 | 110659 | + | 233,8481 | -0,668448582 | 0,691882 | -0,96613 | 0,333978 | 0,444487 | 107,9338489 | -2,583797222 | 0,387328 | -6,67082 | 2,54E-11 | 2,76E-10 | 359,7574 | -0,238785393 | 0,418659 | -0,57036 | 0,568435 | 0,662319 | 300,4442 | 1,006562 | 0,328729 | 3,061984 | 0,002199 | 0,006828 | 178,2043996 | 3,420914434 | 0,470629 | 7,268821 | 3,63E-13 | 3,35E-12 |
| 93 | RM25_RS00455 | RM25_0093 | Lipoprotein-releasing system ATP-binding protein LolD | 95 | gene | 110656 | 111348 | + | 207,9559 | -1,172387759 | 0,837876 | -1,39924 | 0,161742 | 0,264453 | 85,91763375 | -4,001386993 | 0,493401 | -8,1098 | 5,07E-16 | 9,19E-15 | 329,9942 | -0,756705535 | 0,484303 | -1,56246 | 0,118179 | 0,196298 | 301,3425 | 1,283739 | 0,381837 | 3,362004 | 0,000774 | 0,002828 | 126,3767752 | 4,605925683 | 0,594693 | 7,745044 | 9,55E-15 | 1,12E-13 |
| 94 | RM25_RS00460 | RM25_0094 | ABC-type antimicrobial peptide transporter, permease com | 96 | gene | 111345 | 112493 | + | 730,7988 | -1,501872619 | 0,854145 | -1,75833 | 0,078691 | 0,165628 | 387,8520381 | -5,024430934 | 0,435904 | -11,5265 | 9,71E-31 | 1,2E-28 | 1073,746 | -0,929095328 | 0,423794 | -2,19233 | 0,028356 | 0,066568 | 1132,763 | 0,8318 | 0,445422 | 1,867443 | 0,06184 | 0,100939 | 376,5986009 | 5,004228988 | 0,45457 | 11,00871 | 3,47E-28 | 1,74E-26 |
| 95 | RM25_RS00465 | RM25_0095 | Putative helicase | 97 | gene | 112651 | 116049 | - | 3111,197 | 0,023704041 | 0,301101 | 0,078724 | 0,937252 | 0,957161 | 3887,048937 | 0,32397121 | 0,1301 | 2,49017 | 0,012768 | 0,024487 | 2335,346 | -0,476717542 | 0,301357 | -1,5819 | 0,113672 | 0,190558 | 3262,649 | -0,40612 | 0,171263 | -2,37134 | 0,017724 | 0,037266 | 3084,056322 | -1,144907771 | 0,25725 | -4,45057 | 8,56E-06 | 2,53E-05 |
| 96 | RM25_RS00470 | RM25_0096 | Transposase insI for insertion sequence element IS30B/C/D | 98 | gene | 116715 | 117749 | + | 1,414518 | 0,316145685 | 1,080663 | 0,292548 | 0,769868 | 0,835441 | 1,452035908 | 1,079481593 | 2,131786 | 0,506374 | 0,612594 | 0,671262 | 1,376999 | -0,382020663 | 1,307462 | -0,29218 | 0,770145 | 0,832294 | 1,28435 | 0,586206 | 1,628826 | 0,359895 | 0,718926 | 0,766052 | 1,587901629 | -0,709314917 | 1,708278 | -0,41522 | 0,677979 | 0,729013 |
| 97 | RM25_RS11650 | RM25_0097 | Hypothetical protein | 99 | gene | 117730 | 119466 | + | 869,144 | -1,079569387 | 0,361222 | -2,98866 | 0,002802 | 0,019893 | 625,3022093 | -1,580498829 | 0,24533 | -6,44234 | 1,18E-10 | 1,14E-09 | 1112,986 | -0,826450891 | 0,282664 | -2,92379 | 0,003458 | 0,01358 | 1239,904 | 0,535132 | 0,305598 | 1,751101 | 0,079929 | 0,123293 | 548,3841635 | 1,355123268 | 0,236623 | 5,726936 | 1,02E-08 | 4,88E-08 |
| 98 | RM25_RS00480 | RM25_0098 | Hypothetical protein | 100 | gene | 119569 | 120288 | + | 3287,242 | 0,426293448 | 0,29866 | 1,427353 | 0,153478 | 0,25466 | 3784,921144 | 0,234387161 | 0,220277 | 1,064055 | 0,287304 | 0,355099 | 2789,564 | 0,693578411 | 0,424307 | 1,634616 | 0,10213 | 0,174942 | 2982,53 | -0,75401 | 0,375567 | -2,00767 | 0,044678 | 0,077832 | 3681,134619 | -0,267867022 | 0,284086 | -0,94291 | 0,345729 | 0,409694 |
| 99 | RM25_RS00485 | RM25_0099 | putative endonuclease 4 | 101 | gene | 120520 | 121323 | + | 1750,594 | 0,431091485 | 0,422983 | 1,01917 | 0,308122 | 0,417142 | 2497,323904 | 0,714159298 | 0,184978 | 3,860786 | 0,000113 | 0,000379 | 1003,864 | -0,246010372 | 0,264348 | -0,93063 | 0,352045 | 0,458437 | 1583,613 | -0,84606 | 0,292367 | -2,89382 | 0,003806 | 0,010641 | 1972,505782 | -1,768368625 | 0,117875 | -15,002 | 7,12E-51 | 1,87E-48 |
| 100 | RM25_RS00490 | RM25_0100 | AMP-binding enzyme family protein | 102 | gene | 121320 | 123326 | + | 2543,425 | 0,333093625 | 0,345199 | 0,964933 | 0,334579 | 0,445035 | 3460,719476 | 0,436344801 | 0,1625 | 2,685206 | 0,007249 | 0,015039 | 1626,131 | 0,117589309 | 0,172589 | 0,681326 | 0,495665 | 0,595758 | 2393,036 | -0,97193 | 0,198476 | -4,89697 | 9,73E-07 | 7,52E-06 | 2781,175272 | -1,244620182 | 0,119506 | -10,4147 | 2,13E-25 | 8,53E-24 |
| 101 | RM25_RS00495 | RM25_0101 | Major myo-inositol transporter IolT | 103 | gene | 123366 | 124739 | - | 437,8903 | -1,463091814 | 0,750523 | -1,94943 | 0,051244 | 0,12623 | 228,2746717 | -3,830214292 | 0,421454 | -9,0881 | 1,01E-19 | 2,97E-18 | 647,5059 | -0,978208692 | 0,466813 | -2,0955 | 0,036126 | 0,079803 | 674,5464 | 0,938505 | 0,406479 | 2,308866 | 0,020951 | 0,042534 | 230,5896887 | 3,869394155 | 0,514487 | 7,520871 | 5,44E-14 | 5,73E-13 |
| 102 | RM25_RS00500 | RM25_0102 | Hydroxypyruvate isomerase | 104 | gene | 125139 | 125936 | - | 595,8206 | -1,499720668 | 0,677256 | -2,21441 | 0,026801 | 0,084203 | 351,1238711 | -3,67438233 | 0,37998 | -9,66993 | 4,05E-22 | 1,8E-20 | 840,5173 | -0,957428995 | 0,311055 | -3,07801 | 0,002084 | 0,008941 | 923,6514 | 0,703139 | 0,39449 | 1,7824 | 0,074684 | 0,11706 | 306,1835078 | 3,486169662 | 0,338945 | 10,28537 | 8,2E-25 | 3,12E-23 |
| 103 | RM25_RS00505 | RM25_0103 | AscBF operon repressor | 105 | gene | 126428 | 127450 | + | 534,5455 | -0,167132595 | 0,205595 | -0,81292 | 0,416264 | 0,525807 | 480,6452697 | 0,013280023 | 0,137252 | 0,096756 | 0,92292 | 0,941698 | 588,4457 | -0,31462896 | 0,341074 | -0,92247 | 0,356286 | 0,463191 | 594,9229 | 0,383957 | 0,183988 | 2,086859 | 0,036901 | 0,066699 | 495,4753963 | 0,122629177 | 0,31131 | 0,393913 | 0,693645 | 0,743146 |
| 104 | RM25_RS00515 | RM25_0104 | Pyrazinamidase/nicotinamidas pnca | 106 | gene | 127716 | 128288 | + | 862,3587 | -0,371269427 | 0,231108 | -1,60648 | 0,108169 | 0,202184 | 787,4027344 | -0,668085932 | 0,260699 | -2,56267 | 0,010387 | 0,020506 | 937,3147 | -0,128604705 | 0,281717 | -0,4565 | 0,648029 | 0,728723 | 1031,886 | -0,03721 | 0,371578 | -0,10014 | 0,920231 | 0,937331 | 736,0066037 | 0,549333033 | 0,139285 | 3,943958 | 8,01E-05 | 0,000208 |
| 105 | RM25_RS00520 | RM25_0105 | ABC transporter solute-binding protein | 107 | gene | 128669 | 129529 | + | 308,1237 | -0,374587803 | 0,241691 | -1,54986 | 0,121175 | 0,217744 | 243,3409321 | -0,561874581 | 0,190842 | -2,94418 | 0,003238 | 0,007527 | 372,9065 | -0,258825451 | 0,232674 | -1,1124 | 0,265968 | 0,367891 | 366,4108 | 0,422768 | 0,170971 | 2,472755 | 0,013408 | 0,029965 | 263,4311059 | 0,786243707 | 0,251726 | 3,12341 | 0,001788 | 0,003577 |
| 106 | RM25_RS00525 | RM25_0106 | Arginine/ornithine binding protein | 108 | gene | 129823 | 130734 | + | 3034,744 | 0,172950414 | 0,278019 | 0,622082 | 0,533888 | 0,629817 | 3739,039337 | 0,022715858 | 0,19565 | 0,116104 | 0,90757 | 0,929658 | 2330,449 | 0,416972027 | 0,278604 | 1,496646 | 0,134485 | 0,216075 | 3031,173 | -0,94475 | 0,272677 | -3,46471 | 0,000531 | 0,002082 | 3148,461508 | -0,514814438 | 0,18825 | -2,73474 | 0,006243 | 0,011345 |
| 107 | RM25_RS00530 | RM25_0107 | Arginine/ornithine binding protein | 109 | gene | 130901 | 131812 | + | 1659,131 | 0,031053062 | 0,437138 | 0,071037 | 0,943368 | 0,96173 | 2404,357416 | -0,054521167 | 0,17506 | -0,31144 | 0,755464 | 0,801363 | 913,9042 | 0,259526211 | 0,28268 | 0,918092 | 0,358571 | 0,465648 | 1750,074 | -1,61017 | 0,291571 | -5,52239 | 3,34E-08 | 3,4E-07 | 1644,782038 | -1,255585385 | 0,14305 | -8,77725 | 1,68E-18 | 3,32E-17 |
| 108 | RM25_RS00535 | RM25_0108 | Putative polar amino acid ABC transporter substrate-binding | 110 | gene | 131894 | 132874 | + | 2352,88 | 0,530123126 | 0,320785 | 1,65258 | 0,098416 | 0,191077 | 2974,135259 | 0,266602358 | 0,185392 | 1,438043 | 0,150422 | 0,204348 | 1731,625 | 1,008078414 | 0,255835 | 3,94034 | 8,14E-05 | 0,000617 | 2050,265 | -1,2885 | 0,250768 | -5,13819 | 2,77E-07 | 2,34E-06 | 2723,135226 | -0,502300037 | 0,175735 | -2,85829 | 0,004259 | 0,007961 |
| 109 | RM25_RS00540 | RM25_0109 | Arginine/ornithine transport ATP-binding protein | 111 | gene | 132876 | 133703 | + | 1341,685 | 0,386971609 | 0,417422 | 0,927051 | 0,3539 | 0,466073 | 1884,126536 | 0,169732669 | 0,182432 | 0,930387 | 0,352171 | 0,420926 | 799,2443 | 0,924395202 | 0,216512 | 4,269485 | 1,96E-05 | 0,000184 | 1239,871 | -1,74171 | 0,217852 | -7,99493 | 1,3E-15 | 5,56E-14 | 1490,285743 | -0,942165697 | 0,157383 | -5,98644 | 2,14E-09 | 1,14E-08 |
| 110 | RM25_RS12160 | RM25_0110 | Hypothetical protein | 112 | gene | 134566 | 135447 | + | 393,8789 | 0,012568068 | 0,252219 | 0,04983 | 0,960258 | 0,974312 | 326,9829203 | -0,549702004 | 0,206183 | -2,66609 | 0,007674 | 0,015811 | 460,7748 | 0,405690387 | 0,204925 | 1,979707 | 0,047737 | 0,09753 | 414,8416 | -0,03729 | 0,210413 | -0,17724 | 0,85932 | 0,889121 | 387,6760241 | 0,980211228 | 0,211878 | 4,626294 | 3,72E-06 | 1,17E-05 |
| 111 | RM25_RS00555 | RM25_0111 | NAD+ synthetase | 113 | gene | 135705 | 137819 | - | 1812,703 | -0,846291259 | 0,316223 | -2,67625 | 0,007445 | 0,038683 | 1360,25829 | -1,224545825 | 0,196886 | -6,21958 | 4,98E-10 | 4,42E-09 | 2265,147 | -0,633701824 | 0,294797 | -2,14962 | 0,031585 | 0,072277 | 2448,981 | 0,465108 | 0,269008 | 1,728973 | 0,083814 | 0,12829 | 1273,522702 | 1,122286089 | 0,238195 | 4,711637 | 2,46E-06 | 7,99E-06 |
| 112 | RM25_RS00560 | RM25_0112 | Putative cystathionine beta-lyase PatB | 114 | gene | 137987 | 139192 | + | 2001,911 | 0,336636719 | 0,148695 | 2,263936 | 0,023578 | 0,078716 | 2076,681005 | 0,2490263 | 0,143519 | 1,735151 | 0,082714 | 0,120644 | 1927,141 | 0,432063139 | 0,271887 | 1,589125 | 0,112032 | 0,188345 | 1874,801 | -0,26425 | 0,2436 | -1,08476 | 0,278029 | 0,351186 | 2186,393387 | -0,039537419 | 0,174475 | -0,22661 | 0,820729 | 0,854561 |
| 113 | RM25_RS00565 | RM25_0113 | Ribose-5-phosphate isomerase B | 115 | gene | 139295 | 139741 | + | 753,9934 | 0,392644382 | 0,218753 | 1,794922 | 0,072666 | 0,157711 | 912,6792183 | 0,367883835 | 0,13776 | 2,670465 | 0,007575 | 0,015647 | 595,3075 | 0,434172878 | 0,175946 | 2,467653 | 0,0136 | 0,038621 | 692,94 | -0,71414 | 0,180259 | -3,96172 | 7,44E-05 | 0,000376 | 839,3677066 | -0,595523484 | 0,137256 | -4,33879 | 1,43E-05 | 4,13E-05 |
| 114 | RM25_RS00570 | RM25_0114 | 4-hydroxythreonine-4-phosphate dehydrogenase 2 | 116 | gene | 140034 | 141044 | + | 526,1791 | -1,289263984 | 0,416416 | -3,09609 | 0,001961 | 0,015153 | 358,2246915 | -1,702693512 | 0,332227 | -5,12509 | 2,97E-07 | 1,66E-06 | 694,1336 | -1,096358217 | 0,364289 | -3,00958 | 0,002616 | 0,010799 | 783,9685 | 0,719726 | 0,379607 | 1,895975 | 0,057963 | 0,095941 | 300,9091111 | 1,397277798 | 0,351967 | 3,969909 | 7,19E-05 | 0,000188 |
| 115 | RM25_RS00575 | RM25_0115 | High-affinity gluconate transporter | 117 | gene | 141050 | 142393 | + | 592,3887 | -1,399206161 | 0,443546 | -3,15459 | 0,001607 | 0,013108 | 395,9665723 | -1,962066072 | 0,343713 | -5,70844 | 1,14E-08 | 8,27E-08 | 788,8108 | -1,153367457 | 0,355731 | -3,24225 | 0,001186 | 0,005579 | 902,2843 | 0,718028 | 0,396066 | 1,812902 | 0,069847 | 0,111456 | 320,6912723 | 1,60008813 | 0,343238 | 4,661741 | 3,14E-06 | 1E-05 |
| 116 | RM25_RS00580 | RM25_0116 | Putative hexulose-6-phosphate isomerase | 118 | gene | 142438 | 143418 | + | 465,1368 | -1,436577377 | 0,498908 | -2,87945 | 0,003984 | 0,025577 | 300,8678112 | -2,484083152 | 0,309381 | -8,02921 | 9,81E-16 | 1,73E-14 | 629,4058 | -1,048605359 | 0,324877 | -3,2277 | 0,001248 | 0,005813 | 713,2857 | 0,664959 | 0,348949 | 1,905605 | 0,056701 | 0,094116 | 246,71111 | 2,168484477 | 0,313772 | 6,911011 | 4,81E-12 | 3,72E-11 |
| 117 | RM25_RS00585 | RM25_0117 | Transposase IS204/IS1001/IS1096/IS1165 family protein | 119 | gene | 143764 | 145071 | - | 19,03907 | -1,736922993 | 0,929876 | -1,86791 | 0,061775 | 0,142191 | 12,96946843 | -5,943890768 | 1,466568 | -4,05293 | 5,06E-05 | 0,000185 | 25,10867 | -0,959662396 | 0,54893 | -1,74824 | 0,080422 | 0,146036 | 31,02456 | 0,288265 | 0,60873 | 0,473552 | 0,63582 | 0,692526 | 8,607310521 | 5,359098104 | 1,559265 | 3,436938 | 0,000588 | 0,001306 |
| 118 | RM25_RS00590 | RM25_0118 | Putative NADH oxidase-related oxidoreductase protein | 120 | gene | 145195 | 145842 | + | 263,9095 | -0,902888912 | 0,307031 | -2,94071 | 0,003275 | 0,021921 | 205,7996566 | -1,362040243 | 0,284857 | -4,78149 | 1,74E-06 | 8,46E-06 | 322,0194 | -0,636801746 | 0,224079 | -2,84186 | 0,004485 | 0,016492 | 362,5111 | 0,33767 | 0,284442 | 1,18713 | 0,235177 | 0,3034 | 180,3247744 | 1,123197556 | 0,256912 | 4,371923 | 1,23E-05 | 3,57E-05 |
| 119 | RM25_RS00595 | RM25_0119 | MarR family protein | 121 | gene | 145845 | 146189 | + | 238,1501 | -1,06592403 | 0,325132 | -3,27844 | 0,001044 | 0,010209 | 181,8256557 | -1,311402018 | 0,339078 | -3,86755 | 0,00011 | 0,00037 | 294,4745 | -0,926630304 | 0,284797 | -3,25365 | 0,001139 | 0,005414 | 339,2893 | 0,502941 | 0,347568 | 1,44703 | 0,147889 | 0,204397 | 151,2262774 | 0,957451652 | 0,320138 | 2,990743 | 0,002783 | 0,005403 |
| 120 | RM25_RS12905 | RM25_0120 | Hypothetical protein | 122 | gene | 146233 | 147786 | + | 1871,836 | 0,171878337 | 0,092142 | 1,865368 | 0,06213 | 0,142451 | 1872,471282 | 0,080841479 | 0,120764 | 0,669416 | 0,50323 | 0,56888 | 1871,2 | 0,263278753 | 0,194372 | 1,354508 | 0,175574 | 0,266814 | 1858,225 | -0,15869 | 0,172263 | -0,9212 | 0,356945 | 0,433446 | 1943,910863 | 0,077684808 | 0,105986 | 0,73297 | 0,463576 | 0,528105 |
| 121 | RM25_RS00605 | RM25_0121 | Alanine racemase | 123 | gene | 147959 | 149107 | + | 973,1329 | 0,162087634 | 0,282254 | 0,574262 | 0,565791 | 0,659856 | 692,3187627 | -0,070014591 | 0,140363 | -0,49881 | 0,617912 | 0,675832 | 1253,947 | 0,289291634 | 0,142079 | 2,03613 | 0,041737 | 0,088547 | 966,4072 | 0,609942 | 0,154325 | 3,952317 | 7,74E-05 | 0,000388 | 1006,458367 | 1,021558713 | 0,122304 | 8,352629 | 6,68E-17 | 1,04E-15 |
| 122 | RM25_RS00610 | RM25_0122 | Peptidase, S8/S53 family | 124 | gene | 149117 | 150733 | + | 728,3259 | -0,239627723 | 0,23096 | -1,03753 | 0,29949 | 0,409217 | 591,3654619 | -0,55365727 | 0,167669 | -3,30209 | 0,00096 | 0,002591 | 865,2863 | -0,03190755 | 0,211574 | -0,15081 | 0,880125 | 0,913052 | 830,6902 | 0,251301 | 0,19145 | 1,312619 | 0,189311 | 0,251384 | 655,1076052 | 0,831264544 | 0,183783 | 4,523078 | 6,09E-06 | 1,85E-05 |
| 123 | RM25_RS11660 | RM25_0123 | Hypothetical protein | 125 | gene | 150912 | 151781 | + | 329,7907 | -0,702380769 | 0,482106 | -1,4569 | 0,145144 | 0,245765 | 179,0050679 | -1,08559926 | 0,221886 | -4,8926 | 9,95E-07 | 5,08E-06 | 480,5764 | -0,568585694 | 0,392207 | -1,44971 | 0,147139 | 0,231514 | 428,307 | 1,17259 | 0,189976 | 6,172311 | 6,73E-10 | 9,98E-09 | 247,3837189 | 1,757767146 | 0,405154 | 4,338515 | 1,43E-05 | 4,13E-05 |
| 124 | RM25_RS00620 | RM25_0124 | Hypothetical protein | 126 | gene | 151892 | 152695 | - | 1202,06 | 0,266988573 | 0,212702 | 1,255221 | 0,209398 | 0,313964 | 1242,804607 | 0,803885311 | 0,128663 | 6,24798 | 4,16E-10 | 3,71E-09 | 1161,316 | -0,285379477 | 0,244688 | -1,1663 | 0,243493 | 0,34325 | 1151,021 | 0,441419 | 0,251023 | 1,758479 | 0,078666 | 0,121744 | 1287,538079 | -0,603073274 | 0,110135 | -5,47575 | 4,36E-08 | 1,91E-07 |
| 125 | RM25_RS00625 | RM25_0125 | Citrate (Si)-synthase | 127 | gene | 153214 | 154518 | + | 1583,295 | -0,077614066 | 0,103283 | -0,75147 | 0,452371 | 0,558533 | 1500,231391 | -0,243070904 | 0,134253 | -1,81054 | 0,070212 | 0,104543 | 1666,359 | 0,069182672 | 0,14457 | 0,478542 | 0,632264 | 0,715777 | 1718,086 | -0,06151 | 0,127169 | -0,4837 | 0,628596 | 0,686244 | 1508,942571 | 0,303021054 | 0,132473 | 2,287418 | 0,022171 | 0,03614 |
| 126 | RM25_RS00630 | RM25_0126 | Putative secreted peptidoglycan-binding protein | 128 | gene | 154731 | 155339 | + | 25432,37 | 2,859488455 | 0,596757 | 4,791711 | 1,65E-06 | 6,5E-05 | 28182,42247 | 2,041320317 | 0,17012 | 11,99928 | 3,58E-33 | 6,03E-31 | 22682,31 | 5,081410939 | 0,356897 | 14,23776 | 5,34E-46 | 1,1E-42 | 6604,389 | -3,13755 | 0,276123 | -11,3629 | 6,4E-30 | 2,16E-27 | 43671,11281 | -0,059698225 | 0,249351 | -0,23941 | 0,810784 | 0,846332 |
| 127 | RM25_RS00635 | RM25_0127 | Hypothetical protein | 129 | gene | 155650 | 157707 | + | 1607,406 | 0,143734377 | 0,113321 | 1,268386 | 0,20466 | 0,308627 | 1507,328811 | 0,013211435 | 0,126231 | 0,10466 | 0,916645 | 0,936512 | 1707,483 | 0,258644322 | 0,188693 | 1,370718 | 0,170463 | 0,260897 | 1611,654 | -0,00815 | 0,153592 | -0,05304 | 0,957703 | 0,966727 | 1655,28912 | 0,293122269 | 0,149627 | 1,95902 | 0,05011 | 0,07419 |
| 128 | RM25_RS00640 | RM25_0128 | Leucyltransferase | 130 | gene | 158116 | 158841 | + | 348,7017 | -0,239865463 | 0,214799 | -1,1167 | 0,264125 | 0,371889 | 296,9428287 | -0,1691112 | 0,247019 | -0,68461 | 0,493592 | 0,559863 | 400,4606 | -0,293689765 | 0,250556 | -1,17215 | 0,241136 | 0,340538 | 398,4464 | 0,417973 | 0,244597 | 1,708825 | 0,087483 | 0,133111 | 314,0991298 | 0,361362822 | 0,280638 | 1,28765 | 0,197868 | 0,252231 |
| 129 | RM25_RS00645 | RM25_0129 | DNA-formamidopyrimidine glycosylase | 131 | gene | 158834 | 159697 | + | 433,2985 | -0,010571233 | 0,254153 | -0,04159 | 0,966822 | 0,97873 | 343,5967433 | 0,21447798 | 0,16695 | 1,284685 | 0,198902 | 0,258727 | 523,0002 | -0,159318169 | 0,300855 | -0,52955 | 0,596423 | 0,684133 | 456,1334 | 0,733135 | 0,190802 | 3,842387 | 0,000122 | 0,000581 | 424,4780012 | 0,421224003 | 0,277496 | 1,517947 | 0,129028 | 0,173444 |
| 130 | RM25_RS00650 | RM25_0130 | Gluconokinase | 132 | gene | 159845 | 161539 | + | 2393,849 | -1,510740583 | 0,533605 | -2,8312 | 0,004637 | 0,028244 | 1078,288109 | 0,057053616 | 0,133174 | 0,428414 | 0,66835 | 0,722284 | 3709,411 | -2,119218573 | 0,425086 | -4,98539 | 6,18E-07 | 9,22E-06 | 3712,653 | 2,463423 | 0,420559 | 5,857502 | 4,7E-09 | 5,54E-08 | 1217,340017 | 0,325435055 | 0,154269 | 2,109524 | 0,034899 | 0,053728 |
| 131 | RM25_RS12490 | RM25_0131 | Hypothetical protein | 133 | gene | 161570 | 162217 | - | 683,9181 | -0,754463661 | 0,369507 | -2,04181 | 0,04117 | 0,109896 | 469,2656711 | -1,289506874 | 0,241378 | -5,34227 | 9,18E-08 | 5,6E-07 | 898,5705 | -0,498576825 | 0,22841 | -2,18282 | 0,029049 | 0,067693 | 902,4931 | 0,594923 | 0,26509 | 2,244231 | 0,024818 | 0,048463 | 498,3447403 | 1,442953779 | 0,206816 | 6,976997 | 3,02E-12 | 2,4E-11 |
| 132 | RM25_RS00655 | RM25_0132 | Type I site-specific deoxyribonuclease, HsdR family | 134 | gene | 162315 | 165374 | - | 3195,3 | -0,406559107 | 0,151929 | -2,67598 | 0,007451 | 0,038683 | 2881,322985 | -0,560094567 | 0,139806 | -4,00623 | 6,17E-05 | 0,000221 | 3509,276 | -0,283080354 | 0,204641 | -1,3833 | 0,166572 | 0,25616 | 3838,875 | 0,10211 | 0,198676 | 0,513954 | 0,607284 | 0,666994 | 2693,515521 | 0,435760358 | 0,121595 | 3,5837 | 0,000339 | 0,000782 |
| 133 | RM25_RS00660 | RM25_0133 | Type I restriction enzyme, S subunit | 135 | gene | 165371 | 166555 | - | 1479,255 | -0,653634989 | 0,231941 | -2,81811 | 0,004831 | 0,028963 | 1237,440577 | -0,959639426 | 0,163967 | -5,85262 | 4,84E-09 | 3,71E-08 | 1721,07 | -0,444451673 | 0,250429 | -1,77476 | 0,075937 | 0,139286 | 1904,991 | 0,213183 | 0,248192 | 0,858944 | 0,390371 | 0,466348 | 1126,535248 | 0,787638741 | 0,150326 | 5,239527 | 1,61E-07 | 6,34E-07 |
| 134 | RM25_RS00665 | RM25_0134 | Type I restriction-modification system, M subunit | 136 | gene | 166552 | 168120 | - | 1544,561 | -0,33683248 | 0,123649 | -2,72409 | 0,006448 | 0,035179 | 1419,42108 | -0,441148658 | 0,13649 | -3,23211 | 0,001229 | 0,003218 | 1669,702 | -0,250400847 | 0,17649 | -1,41878 | 0,155963 | 0,242483 | 1818,599 | 0,087251 | 0,16513 | 0,528381 | 0,597235 | 0,658719 | 1337,526609 | 0,332608444 | 0,12953 | 2,567804 | 0,010234 | 0,017855 |
| 135 | RM25_RS00670 | RM25_0135 | Putative ABC-type transport system ATP-binding component | 137 | gene | 168175 | 169992 | - | 713,857 | -0,380696037 | 0,353429 | -1,07715 | 0,281413 | 0,390401 | 487,7398207 | -0,540555135 | 0,178929 | -3,02106 | 0,002519 | 0,006027 | 939,9741 | -0,300806942 | 0,368286 | -0,81677 | 0,414057 | 0,519113 | 847,7184 | 0,777592 | 0,215979 | 3,600308 | 0,000318 | 0,001352 | 610,8258274 | 1,086441405 | 0,341311 | 3,183143 | 0,001457 | 0,00297 |
| 136 | RM25_RS00675 | RM25_0136 | ABC-2 type transporter | 138 | gene | 169989 | 171176 | - | 854,0761 | -0,640150743 | 0,240507 | -2,66167 | 0,007775 | 0,039597 | 653,05538 | -0,558410395 | 0,146491 | -3,81192 | 0,000138 | 0,000455 | 1055,097 | -0,692820054 | 0,20796 | -3,3315 | 0,000864 | 0,004332 | 1093,896 | 0,684859 | 0,174694 | 3,92033 | 8,84E-05 | 0,000437 | 654,5780614 | 0,604093295 | 0,182096 | 3,317448 | 0,000908 | 0,001955 |

| | | | | | | | | | | | | | | | | | | | | | | | | | | | | | | | | | | | | | | |
|---|---|---|---|---|---|---|---|---|---|---|---|---|---|---|---|---|---|---|---|---|---|---|---|---|---|---|---|---|---|---|---|---|---|---|---|---|---|---|
| 137 | NA | RM25_0137 | Hypothetical protein | 139 | gene | 171264 | 171488 | + | 84,46154 | -1,297747068 | 0,489234 | -2,65261 | 0,007987 | 0,040226 | 57,39358902 | -2,207168833 | 0,454745 | -4,85364 | 1,21E-06 | 6,07E-06 | 111,5295 | -0,92532061 | 0,406952 | -2,27378 | 0,022979 | 0,056952 | 126,3064 | 0,560927 | 0,384849 | 1,457524 | 0,144972 | 0,201473 | 48,00637706 | 1,913650681 | 0,524652 | 3,647469 | 0,000265 | 0,000625 |
| 138 | RM25_RS00680 | RM25_0138 | Hypothetical protein | 140 | gene | 171377 | 172366 | - | 1192,609 | 0,446994481 | 0,121026 | 3,693371 | 0,000221 | 0,003105 | 1246,218045 | 0,441097412 | 0,14383 | 3,066791 | 0,002164 | 0,00528 | 1138,999 | 0,454024062 | 0,225905 | 2,009798 | 0,044453 | 0,092502 | 1066,969 | -0,19782 | 0,125549 | -1,57561 | 0,115116 | 0,166562 | 1346,264636 | -0,139625349 | 0,191061 | -0,73079 | 0,464909 | 0,529367 |
| 139 | RM25_RS00685 | RM25_0139 | Aminodeoxychorismate synthase, component I | 141 | gene | 172603 | 174336 | + | 811,2429 | -0,967079194 | 0,484795 | -1,99482 | 0,046062 | 0,118138 | 484,6499585 | -1,659912644 | 0,259725 | -6,39104 | 1,65E-10 | 1,57E-09 | 1137,836 | -0,706587534 | 0,40456 | -1,74656 | 0,080714 | 0,146228 | 1125,658 | 0,869594 | 0,296056 | 2,937266 | 0,003311 | 0,009603 | 541,4364231 | 1,896375483 | 0,382741 | 4,954718 | 7,24E-07 | 2,6E-06 |
| 140 | RM25_RS11665 | RM25_0140 | Branched-chain amino acid aminotransferase/4-amino-4-de | 142 | gene | 174329 | 175282 | + | 459,475 | -0,302066594 | 0,317218 | -0,95224 | 0,340977 | 0,452015 | 335,0979487 | -0,388276011 | 0,187641 | -2,06924 | 0,038523 | 0,063585 | 583,8521 | -0,252904319 | 0,36243 | -0,6978 | 0,485301 | 0,585691 | 531,7339 | 0,683035 | 0,205366 | 3,325938 | 0,000881 | 0,003157 | 405,1858095 | 0,881898097 | 0,339162 | 2,600227 | 0,009316 | 0,016375 |
| 141 | RM25_RS11670 | RM25_0141 | Hypothetical protein | 143 | gene | 175576 | 176256 | - | 661,8794 | -0,092802463 | 0,207817 | -0,44656 | 0,655193 | 0,738119 | 540,5321205 | 0,001137909 | 0,193872 | 0,005869 | 0,995317 | 0,996585 | 783,2267 | -0,158671609 | 0,174223 | -0,91074 | 0,362433 | 0,46937 | 717,7473 | 0,555638 | 0,156045 | 3,560747 | 0,00037 | 0,001537 | 628,5805269 | 0,448750671 | 0,20241 | 2,217042 | 0,02662 | 0,042481 |
| 142 | RM25_RS12175 | RM25_0142 | Prephenate dehydratase | 144 | gene | 176735 | 177721 | + | 1037,475 | -0,059392938 | 0,093336 | -0,63633 | 0,524558 | 0,621298 | 998,5504103 | -0,011258069 | 0,122296 | -0,09206 | 0,926654 | 0,944314 | 1076,4 | -0,103563423 | 0,192538 | -0,53788 | 0,590657 | 0,679774 | 1116,643 | 0,091605 | 0,148579 | 0,61654 | 0,537538 | 0,604186 | 997,2626652 | 0,057194091 | 0,161698 | 0,353709 | 0,723557 | 0,769943 |
| 143 | RM25_RS00705 | RM25_0143 | PA-phosphatase like phosphoesterase | 145 | gene | 177601 | 179226 | - | 1391,835 | 0,047394654 | 0,095546 | 0,496038 | 0,619867 | 0,707967 | 1415,776401 | 0,187756449 | 0,139404 | 1,346847 | 0,17803 | 0,235211 | 1367,894 | -0,096121344 | 0,165472 | -0,58089 | 0,561312 | 0,656582 | 1446,693 | 0,036705 | 0,176809 | 0,207597 | 0,835543 | 0,87025 | 1387,425012 | -0,195301112 | 0,134967 | -1,44703 | 0,147889 | 0,196141 |
| 144 | RM25_RS00710 | RM25_0144 | Serine--tRNA ligase | 146 | gene | 179353 | 180627 | + | 2325,219 | 0,384958879 | 0,173653 | 2,216824 | 0,026635 | 0,084154 | 2637,425637 | 0,423860325 | 0,160624 | 2,63883 | 0,008319 | 0,016962 | 2013,012 | 0,335239861 | 0,216089 | 1,551398 | 0,120806 | 0,199818 | 2137,363 | -0,39419 | 0,224885 | -1,75283 | 0,079632 | 0,122915 | 2580,116072 | -0,439348026 | 0,137595 | -3,19306 | 0,001408 | 0,002883 |
| 145 | RM25_RS11675 | RM25_0145 | Hypothetical protein | 147 | gene | 180748 | 181428 | + | 1075,545 | -0,125622987 | 0,1876 | -0,66963 | 0,503093 | 0,600097 | 1023,80327 | 0,382340954 | 0,122002 | 3,13389 | 0,001725 | 0,004312 | 1127,287 | -0,590125208 | 0,197579 | -2,98678 | 0,002819 | 0,011497 | 1182,585 | 0,550603 | 0,198886 | 2,768439 | 0,005633 | 0,01462 | 1009,435167 | -0,370662891 | 0,135091 | -2,7438 | 0,006073 | 0,011062 |
| 146 | RM25_RS00720 | RM25_0146 | PTS system mannose-specific EIIBCA component (EIIBCA-Ma | 148 | gene | 181568 | 184540 | - | 2986,468 | -0,140400942 | 0,421626 | -0,333 | 0,739135 | 0,809546 | 1736,012888 | -0,573254913 | 0,119617 | -4,79242 | 1,65E-06 | 8,06E-06 | 4236,923 | 0,034166901 | 0,307578 | 0,111084 | 0,91155 | 0,93822 | 3295,535 | 0,959522 | 0,268155 | 3,578238 | 0,000346 | 0,001456 | 2771,670463 | 1,604661876 | 0,183906 | 8,725441 | 2,65E-18 | 5,12E-17 |
| 147 | RM25_RS00725 | RM25_0147 | Putative exonuclease of the beta-lactamase fold involved in | 149 | gene | 184790 | 186184 | + | 1017,693 | -1,034611938 | 0,417922 | -2,47561 | 0,013301 | 0,056794 | 579,5616648 | -1,030623567 | 0,241108 | -4,27454 | 1,92E-05 | 7,66E-05 | 1455,825 | -1,036938256 | 0,214628 | -4,83133 | 1,36E-06 | 1,82E-05 | 1433,219 | 1,269093 | 0,27562 | 4,604498 | 4,13E-06 | 2,77E-05 | 654,9972545 | 1,319524405 | 0,203815 | 6,47412 | 9,54E-11 | 6,26E-10 |
| 148 | RM25_RS00730 | RM25_0148 | Alkaline phosphatase synthesis transcriptional regulatory pr | 150 | gene | 186181 | 186900 | + | 387,5599 | -0,60486925 | 0,342615 | -1,76545 | 0,077488 | 0,164837 | 260,4583772 | -0,685331337 | 0,240412 | -2,85066 | 0,004363 | 0,009661 | 514,6613 | -0,566654247 | 0,262361 | -2,15983 | 0,030786 | 0,070793 | 490,4389 | 0,873888 | 0,239222 | 3,653036 | 0,000259 | 0,001127 | 302,1104634 | 1,052630372 | 0,270218 | 3,89548 | 9,8E-05 | 0,000249 |
| 149 | RM25_RS00735 | RM25_0149 | Two-component system sensor kinase TcsS4 | 151 | gene | 186897 | 188186 | + | 748,4779 | -0,445064364 | 0,302259 | -1,47246 | 0,140897 | 0,241408 | 537,6173174 | -0,360463477 | 0,158075 | -2,28033 | 0,022588 | 0,039702 | 959,3385 | -0,494232148 | 0,311192 | -1,58819 | 0,112244 | 0,188566 | 905,1204 | 0,82892 | 0,18775 | 4,415009 | 1,01E-05 | 6,22E-05 | 623,4663122 | 0,759373869 | 0,285508 | 2,659727 | 0,00782 | 0,013943 |
| 150 | RM25_RS00740 | RM25_0150 | Putative periplasmic or exported protein | 152 | gene | 188248 | 188895 | - | 396,0368 | -1,195815111 | 0,68273 | -1,75152 | 0,079857 | 0,167129 | 185,5839554 | -2,795197279 | 0,39145 | -7,14063 | 9,29E-13 | 1,24E-11 | 606,4897 | -0,84165924 | 0,424101 | -1,98457 | 0,047192 | 0,096891 | 576,2621 | 1,193178 | 0,41069 | 2,9053 | 0,003669 | 0,010344 | 237,4494585 | 3,220409988 | 0,427101 | 7,540159 | 4,69E-14 | 4,98E-13 |
| 151 | RM25_RS00745 | RM25_0151 | putative thiol peroxidase | 153 | gene | 189229 | 189726 | + | 2624,603 | 2,309939847 | 0,278449 | 8,295728 | 1,08E-16 | 6,36E-14 | 3105,290669 | 2,473332303 | 0,250238 | 9,88391 | 4,89E-23 | 2,4E-21 | 2143,916 | 2,094486513 | 0,383065 | 5,467701 | 4,56E-08 | 8,6E-07 | 934,6074 | -0,27137 | 0,389124 | -0,69739 | 0,485556 | 0,555291 | 4274,835946 | -0,614762012 | 0,246126 | -2,49775 | 0,012498 | 0,021425 |
| 152 | RM25_RS00750 | RM25_0152 | Transcriptional regulator, Sir2 family | 154 | gene | 189894 | 190640 | - | 1068,444 | 0,077367298 | 0,195807 | 0,39512 | 0,692754 | 0,770199 | 1095,860507 | 0,490608029 | 0,131776 | 3,723046 | 0,000197 | 0,000629 | 1041,028 | -0,351601614 | 0,278096 | -1,26432 | 0,206116 | 0,302689 | 1098,189 | 0,301103 | 0,255545 | 1,178279 | 0,238686 | 0,307422 | 1074,53781 | -0,498259202 | 0,166234 | -2,99733 | 0,002724 | 0,005297 |
| 153 | RM25_RS00755 | RM25_0153 | Putative CONSERVED PROTEIN CPSA | 155 | gene | 190841 | 192550 | + | 1620,386 | 0,092653379 | 0,121921 | 0,759948 | 0,447286 | 0,554286 | 1696,213369 | -0,097623148 | 0,144235 | -0,67683 | 0,498511 | 0,564628 | 1544,541 | 0,30167854 | 0,17481 | 1,725756 | 0,084391 | 0,150918 | 1659,013 | -0,40044 | 0,177879 | -2,2512 | 0,024373 | 0,047872 | 1639,026713 | 0,050886727 | 0,107351 | 0,474024 | 0,635483 | 0,69152 |
| 154 | RM25_RS00760 | RM25_0154 | Argininosuccinate synthase | 156 | gene | 193204 | 194640 | - | 3587,232 | 0,558518189 | 0,191943 | 2,909808 | 0,003617 | 0,02353 | 4038,616242 | 0,553458023 | 0,172761 | 3,203613 | 0,001357 | 0,003496 | 3135,848 | 0,565534554 | 0,270635 | 2,089658 | 0,036649 | 0,080579 | 3076,615 | -0,42616 | 0,248734 | -1,7133 | 0,086657 | 0,131944 | 4182,444428 | -0,374341409 | 0,18877 | -1,98305 | 0,047361 | 0,070474 |
| 155 | RM25_RS00765 | RM25_0155 | NAD(P)H quinone oxidoreductase, PIG3 family | 157 | gene | 194802 | 195809 | - | 2801,368 | 0,217739749 | 0,305925 | 0,711742 | 0,476625 | 0,577158 | 2379,480195 | 0,864510324 | 0,163945 | 5,27317 | 1,34E-07 | 7,92E-07 | 3223,255 | -0,241549336 | 0,377733 | -0,63947 | 0,522517 | 0,622195 | 2729,236 | 0,999917 | 0,364414 | 2,743905 | 0,006071 | 0,015538 | 2946,981981 | -0,070296086 | 0,192376 | -0,36541 | 0,714805 | 0,763041 |
| 157 | RM25_RS00775 | RM25_0157 | Membrane protein without function | 161 | gene | 196023 | 196460 | + | 1669,033 | 0,092055102 | 0,217336 | 0,423561 | 0,671886 | 0,752321 | 1620,358419 | 0,280185986 | 0,18285 | 1,532329 | 0,125441 | 0,174434 | 1717,708 | -0,083531883 | 0,367825 | -0,2271 | 0,820348 | 0,870068 | 1708,626 | 0,222015 | 0,349136 | 0,635897 | 0,524843 | 0,59246 | 1683,675136 | -0,107569102 | 0,209593 | -0,51323 | 0,607791 | 0,66724 |
| 158 | RM25_RS00780 | RM25_0158 | Cell envelope-related transcriptional attenuator | 162 | gene | 196842 | 199460 | - | 1967,388 | 0,182566079 | 0,160906 | 1,134611 | 0,256538 | 0,364338 | 2227,640003 | 0,206239497 | 0,146217 | 1,410503 | 0,158391 | 0,213065 | 1707,135 | 0,152690523 | 0,188582 | 0,809678 | 0,418125 | 0,522934 | 1948,309 | -0,41371 | 0,188334 | -2,19667 | 0,028044 | 0,05347 | 2051,841557 | -0,415108453 | 0,112581 | -3,68721 | 0,000227 | 0,000541 |
| 161 | RM25_RS11680 | RM25_0161 | Hypothetical protein | 169 | gene | 200418 | 201773 | - | 1748,945 | 0,183816699 | 0,300884 | 0,610923 | 0,541251 | 0,636908 | 2271,491036 | 0,154383259 | 0,184412 | 0,837167 | 0,402499 | 0,471985 | 1226,398 | 0,238737928 | 0,186956 | 1,276971 | 0,201612 | 0,298868 | 1740,322 | -0,99422 | 0,191452 | -5,19304 | 2,07E-07 | 1,81E-06 | 1825,002898 | -0,858100681 | 0,176892 | -4,85098 | 1,23E-06 | 4,2E-06 |
| 162 | RM25_RS00800 | RM25_0162 | Oxidoreductase, short chain dehydrogenase/reductase fam | 170 | gene | 201906 | 202589 | + | 724,8444 | -0,029834489 | 0,161077 | -0,18522 | 0,853058 | 0,904036 | 633,6465891 | 0,179684276 | 0,144374 | 1,244577 | 0,213287 | 0,27456 | 816,0423 | -0,190232643 | 0,178274 | -1,06708 | 0,285936 | 0,390696 | 771,8902 | 0,49243 | 0,189904 | 2,593042 | 0,009513 | 0,02281 | 703,1646018 | 0,173364213 | 0,137144 | 1,2641 | 0,206194 | 0,260729 |
| 163 | RM25_RS00805 | RM25_0163 | Glycerophosphoryl diester phosphodiesterase family protein | 171 | gene | 202586 | 203305 | + | 522,8857 | -0,19824972 | 0,122833 | -1,61398 | 0,106532 | 0,200556 | 476,4790485 | -0,063383713 | 0,147676 | -0,42921 | 0,667772 | 0,72213 | 569,2923 | -0,311663703 | 0,168507 | -1,84955 | 0,064378 | 0,122853 | 588,9791 | 0,310131 | 0,155488 | 1,994563 | 0,046091 | 0,079645 | 477,8725924 | 0,118009718 | 0,167223 | 0,705702 | 0,480374 | 0,542521 |
| 164 | RM25_RS00810 | RM25_0164 | Shikimate dehydrogenase | 172 | gene | 203396 | 204214 | - | 1276,01 | 0,123137032 | 0,224134 | 0,549389 | 0,582738 | 0,674574 | 1090,593129 | 0,366001949 | 0,188484 | 1,941824 | 0,052158 | 0,081958 | 1461,427 | -0,055304206 | 0,28802 | -0,19202 | 0,84773 | 0,889626 | 1287,525 | 0,594456 | 0,304467 | 1,952449 | 0,050885 | 0,086099 | 1301,504034 | 0,210588257 | 0,145778 | 1,444577 | 0,148577 | 0,196628 |
| 165 | RM25_RS00815 | RM25_0165 | 3-carboxymuconate cyclase (Precursor) | 173 | gene | 204301 | 205350 | + | 885,4456 | -0,388459509 | 0,209406 | -1,85505 | 0,063589 | 0,144252 | 729,1712357 | -0,320413546 | 0,160176 | -2,00038 | 0,045459 | 0,073389 | 1041,72 | -0,437652416 | 0,247448 | -1,76866 | 0,07695 | 0,140598 | 1055,873 | 0,500345 | 0,196716 | 2,543487 | 0,010975 | 0,025486 | 753,0049276 | 0,444943186 | 0,217883 | 2,042116 | 0,04114 | 0,062158 |
| 166 | RM25_RS00820 | RM25_0166 | 2, 5-diketo-D-gluconic acid reductase A | 174 | gene | 205455 | 206297 | + | 1440,188 | 0,991955932 | 0,312434 | 3,174925 | 0,001499 | 0,012787 | 1902,296288 | 1,11093564 | 0,200119 | 5,551368 | 2,83E-08 | 1,9E-07 | 978,0788 | 0,771069527 | 0,199503 | 3,864954 | 0,000111 | 0,000806 | 1023,071 | -0,79354 | 0,186612 | -4,25237 | 2,12E-05 | 0,000121 | 1879,178529 | -1,08741316 | 0,202618 | -5,3668 | 8,01E-08 | 3,34E-07 |
| 167 | RM25_RS00825 | RM25_0167 | ABC-type antimicrobial peptide transport system | 175 | gene | 206451 | 208904 | - | 3139,915 | 0,600069653 | 0,286052 | 2,097762 | 0,035926 | 0,101532 | 4047,154783 | 0,779998704 | 0,141979 | 5,493754 | 3,93E-08 | 2,58E-07 | 2232,675 | 0,285009986 | 0,212315 | 1,342393 | 0,179469 | 0,271789 | 2644,592 | -0,62913 | 0,194779 | -3,22997 | 0,001238 | 0,004235 | 3713,829699 | -1,066977639 | 0,141427 | -7,54436 | 4,55E-14 | 4,85E-13 |
| 168 | RM25_RS00830 | RM25_0168 | Ectoine/hydroxyectoine ABC transporter, ATP-binding prote | 176 | gene | 208904 | 209842 | - | 869,4974 | 0,349213613 | 0,17311 | 2,017293 | 0,043665 | 0,114094 | 970,3696008 | 0,567476329 | 0,137691 | 4,121381 | 3,77E-05 | 0,000142 | 768,6253 | 0,080589753 | 0,234848 | 0,343157 | 0,73148 | 0,800789 | 806,2242 | -0,1267 | 0,191271 | -0,66239 | 0,507719 | 0,574781 | 956,6962791 | -0,556707608 | 0,170008 | -3,2746 | 0,001058 | 0,002219 |
| 169 | RM25_RS00835 | RM25_0169 | Two-component system response regulator | 177 | gene | 210197 | 210904 | - | 498,9117 | 0,190948152 | 0,222458 | 0,858358 | 0,390695 | 0,501836 | 400,7786777 | -0,110393544 | 0,161053 | -0,68545 | 0,49306 | 0,559798 | 597,0448 | 0,391848351 | 0,201362 | 1,945989 | 0,051656 | 0,103356 | 490,4743 | 0,249766 | 0,183661 | 1,359925 | 0,173854 | 0,234827 | 520,819358 | 0,805587064 | 0,146828 | 5,48661 | 4,1E-08 | 1,81E-07 |
| 170 | RM25_RS00840 | RM25_0170 | Two-component system sensor kinase | 178 | gene | 210873 | 212192 | - | 1929,31 | 0,735467336 | 0,234881 | 3,131229 | 0,001741 | 0,013919 | 1498,680465 | 0,448611254 | 0,133186 | 3,368308 | 0,000756 | 0,00209 | 2359,94 | 0,923810347 | 0,230693 | 4,004503 | 6,21E-05 | 0,0005 | 1525,5 | 0,300703 | 0,143796 | 2,091178 | 0,036512 | 0,066346 | 2354,490034 | 0,822141775 | 0,17985 | 4,571253 | 4,85E-06 | 1,5E-05 |
| 171 | RM25_RS00845 | RM25_0171 | Carboxylesterase type B | 179 | gene | 212356 | 213060 | - | 422,2589 | 0,001345083 | 0,323047 | 0,004164 | 0,996678 | 0,998797 | 293,9301127 | -0,239904431 | 0,260254 | -0,92181 | 0,356628 | 0,425822 | 550,5877 | 0,125570566 | 0,205486 | 0,611089 | 0,541141 | 0,638109 | 443,5743 | 0,657553 | 0,28672 | 2,293367 | 0,021827 | 0,04378 | 413,6400975 | 1,081788437 | 0,179642 | 6,021898 | 1,72E-09 | 9,26E-09 |
| 172 | RM25_RS00850 | RM25_0172 | Membrane protein containing DUF1211 | 180 | gene | 213138 | 213743 | + | 277,0954 | -0,799646929 | 0,364523 | -2,19368 | 0,028258 | 0,086721 | 191,7713204 | -1,33837739 | 0,267689 | -4,99975 | 5,74E-07 | 3E-06 | 362,4194 | -0,541999329 | 0,210346 | -2,5767 | 0,009975 | 0,03018 | 370,2758 | 0,576643 | 0,268672 | 2,146266 | 0,031852 | 0,059344 | 198,0693857 | 1,433670817 | 0,223161 | 6,424367 | 1,32E-10 | 8,44E-10 |
| 173 | RM25_RS00855 | RM25_0173 | Acetyltransferase | 181 | gene | 213907 | 214221 | + | 2556,371 | 0,534750183 | 0,447392 | 1,195262 | 0,231985 | 0,33962 | 1562,661471 | 0,499726447 | 0,185638 | 2,691937 | 0,007104 | 0,014779 | 3550,081 | 0,550813464 | 0,516412 | 1,066617 | 0,286145 | 0,390705 | 2199,495 | 1,104769 | 0,483567 | 2,284622 | 0,022335 | 0,044538 | 2946,335815 | 1,186401845 | 0,260866 | 4,547938 | 5,42E-06 | 1,66E-05 |
| 174 | RM25_RS00860 | RM25_0174 | Hypothetical protein | 182 | gene | 214310 | 214543 | + | 1908,512 | 1,228527388 | 0,504547 | 2,43491 | 0,014895 | 0,060797 | 983,8569582 | 0,433226709 | 0,227551 | 1,903866 | 0,056928 | 0,087641 | 2833,167 | 1,54593759 | 0,52057 | 2,969704 | 0,002981 | 0,011912 | 1205,523 | 0,741382 | 0,489361 | 1,514999 | 0,129773 | 0,183929 | 2602,428644 | 1,882322827 | 0,288327 | 6,528437 | 6,65E-11 | 4,46E-10 |
| 175 | RM25_RS00865 | RM25_0175 | Peptidyl-prolyl cis-trans isomerase | 183 | gene | 214677 | 215051 | - | 4233,619 | 1,587215471 | 0,480674 | 3,302066 | 0,00096 | 0,009713 | 5164,92586 | 1,021360161 | 0,25135 | 4,063498 | 4,83E-05 | 0,000178 | 3302,313 | 2,821348867 | 0,420151 | 6,715082 | 1,88E-11 | 7,38E-10 | 2261,331 | -2,10719 | 0,381745 | -5,51989 | 3,39E-08 | 3,43E-07 | 6203,587609 | -0,279078847 | 0,295453 | -0,94458 | 0,344873 | 0,408887 |
| 176 | RM25_RS00870 | RM25_0176 | Panthothenate kinase | 184 | gene | 215125 | 215748 | - | 432,5911 | -0,563620815 | 0,302961 | -1,86037 | 0,062833 | 0,142908 | 319,0212 | -0,739473906 | 0,18103 | -4,08481 | 4,41E-05 | 0,000165 | 546,1609 | -0,46618216 | 0,32011 | -1,45632 | 0,145305 | 0,229824 | 542,6997 | 0,5993 | 0,220929 | 2,712635 | 0,006675 | 0,016809 | 343,3995489 | 0,941115858 | 0,303132 | 3,104637 | 0,001905 | 0,003799 |
| 177 | RM25_RS00875 | RM25_0177 | DNA polymerase III, epsilon subunit | 185 | gene | 215842 | 216498 | + | 642,1592 | 0,122304929 | 0,151437 | 0,80763 | 0,419304 | 0,528785 | 561,6114491 | 0,206641224 | 0,142532 | 1,449785 | 0,147119 | 0,200554 | 722,7069 | 0,05619569 | 0,179699 | 0,312722 | 0,754492 | 0,819511 | 646,9267 | 0,381898 | 0,148158 | 2,577637 | 0,009948 | 0,023636 | 656,722227 | 0,287407567 | 0,159014 | 1,807438 | 0,070694 | 0,100863 |
| 178 | RM25_RS00880 | RM25_0178 | Cytochrome bd-type menaquinol oxidase subunit I | 186 | gene | 216850 | 218352 | + | 13673,72 | -0,081194176 | 0,416904 | -0,19475 | 0,845585 | 0,898172 | 9042,491095 | 0,413644934 | 0,22777 | 1,816068 | 0,06936 | 0,103647 | 18304,95 | -0,325489525 | 0,464886 | -0,70015 | 0,483834 | 0,584519 | 14795,09 | 1,342851 | 0,432887 | 3,102081 | 0,001922 | 0,006098 | 12956,58375 | 0,634062193 | 0,282316 | 2,245933 | 0,024708 | 0,039698 |
| 179 | RM25_RS00885 | RM25_0179 | Cytochrome d ubiquinol oxidase, subunit II | 187 | gene | 218369 | 219502 | + | 9062,538 | 0,334207258 | 0,349566 | 0,956062 | 0,339041 | 0,449955 | 7085,092281 | 0,609864845 | 0,22752 | 2,680486 | 0,007352 | 0,015226 | 11039,98 | 0,160940825 | 0,470083 | 0,342367 | 0,732075 | 0,801068 | 8460,898 | 0,844691 | 0,446497 | 1,891818 | 0,058515 | 0,096718 | 9862,051841 | 0,426523547 | 0,271827 | 1,569102 | 0,116624 | 0,158343 |
| 180 | RM25_RS12185 | RM25_0180 | ABC transporter | 188 | gene | 219526 | 223245 | + | 2331,764 | 0,160957023 | 0,191441 | 0,840766 | 0,400479 | 0,511615 | 2003,265333 | 0,294582732 | 0,142181 | 2,071884 | 0,038276 | 0,063222 | 2660,263 | 0,060812908 | 0,254237 | 0,239198 | 0,810952 | 0,865677 | 2310,985 | 0,471267 | 0,202977 | 2,32178 | 0,020245 | 0,041385 | 2416,67515 | 0,295902438 | 0,186202 | 1,589148 | 0,112027 | 0,152717 |
| 181 | RM25_RS00895 | RM25_0181 | Succinyl-diaminopimelate desuccinylase | 189 | gene | 223594 | 224775 | - | 1091,909 | -0,064842474 | 0,086239 | -0,75189 | 0,452117 | 0,558512 | 1061,015867 | 0,029594577 | 0,135125 | 0,219016 | 0,826638 | 0,862499 | 1122,802 | -0,153309237 | 0,164153 | -0,93394 | 0,350336 | 0,456655 | 1178,994 | 0,112442 | 0,158334 | 0,710157 | 0,477607 | 0,549131 | 1046,356231 | -0,018791866 | 0,142034 | -0,13231 | 0,894743 | 0,915325 |
| 182 | RM25_RS00900 | RM25_0182 | Multidrug resistance membrane efflux protein emrB | 190 | gene | 224913 | 226610 | + | 811,4523 | -0,759485502 | 0,462314 | -1,64279 | 0,100426 | 0,193298 | 493,9686207 | -1,387587157 | 0,28777 | -4,82186 | 1,42E-06 | 7,04E-06 | 1128,936 | -0,509440605 | 0,363613 | -1,40105 | 0,161199 | 0,249145 | 1070,201 | 0,823051 | 0,34531 | 2,383516 | 0,017148 | 0,036249 | 593,6658253 | 1,7730328 | 0,32856 | 5,396375 | 6,8E-08 | 2,89E-07 |
| 183 | RM25_RS00905 | RM25_0183 | Transcriptional regulator, TetR family | 191 | gene | 227064 | 227687 | - | 497,0036 | 0,720722543 | 0,242335 | 2,974074 | 0,002939 | 0,020514 | 586,0997446 | 0,500512705 | 0,17068 | 2,932456 | 0,003363 | 0,007787 | 407,9074 | 1,055361208 | 0,23248 | 4,539576 | 5,64E-06 | 6,15E-05 | 398,8279 | -0,93182 | 0,224385 | -4,15276 | 3,28E-05 | 0,00018 | 606,0099026 | -0,327096066 | 0,159276 | -2,05364 | 0,040011 | 0,060764 |
| 184 | RM25_RS00910 | RM25_0184 | Protein of hypothetical function DUF77 | 192 | gene | 227891 | 228208 | + | 510,9403 | -0,269035373 | 0,210269 | -1,27948 | 0,200728 | 0,303669 | 474,37996 | 0,158401495 | 0,194765 | 0,813296 | 0,416048 | 0,48426 | 547,5006 | -0,646267602 | 0,277179 | -2,33159 | 0,019722 | 0,050707 | 589,0456 | 0,517907 | 0,237617 | 2,179589 | 0,029288 | 0,055425 | 453,7007718 | -0,243000861 | 0,241351 | -1,00684 | 0,314014 | 0,377237 |
| 185 | RM25_RS00915 | RM25_0185 | TRNA uridine 5-carboxymethylaminomethyl modification en | 193 | gene | 228534 | 230165 | + | 4762,967 | -0,675345365 | 0,457468 | -1,47627 | 0,139872 | 0,240115 | 6414,160456 | -0,240002901 | 0,202349 | -1,18608 | 0,235589 | 0,299022 | 3111,774 | -1,712497216 | 0,327941 | -5,22196 | 1,77E-07 | 2,98E-06 | 6223,873 | -0,59271 | 0,333071 | -1,77953 | 0,075154 | 0,117387 | 3602,096979 | -2,024201905 | 0,207282 | -9,76543 | 1,58E-22 | 4,61E-21 |
| 186 | RM25_RS00920 | RM25_0186 | Pyruvate-flavodoxin oxidoreductase | 194 | gene | 230171 | 233938 | + | 13790,06 | -0,586571206 | 0,53809 | -1,0901 | 0,27567 | 0,384697 | 20403,72387 | -0,305936978 | 0,212184 | -1,44184 | 0,149346 | 0,203238 | 7176,4 | -1,472701027 | 0,402061 | -3,66288 | 0,000249 | 0,001568 | 17638,86 | -1,14384 | 0,411346 | -2,78073 | 0,005424 | 0,014157 | 10824,93838 | -2,274184919 | 0,19884 | -11,4372 | 2,72E-30 | 1,69E-28 |
| 187 | RM25_RS00925 | RM25_0187 | Dihydroorotate dehydrogenase 2 | 195 | gene | 233940 | 235088 | + | 4844,881 | -0,331940854 | 0,639604 | -0,51898 | 0,603775 | 0,694194 | 7349,014908 | 0,059406192 | 0,228425 | 0,260068 | 0,794811 | 0,836699 | 2340,747 | -1,736857326 | 0,584757 | -2,97022 | 0,002976 | 0,011912 | 5752,018 | -1,04277 | 0,598355 | -1,74273 | 0,08138 | 0,125122 | 4213,992422 | -2,805826677 | 0,188064 | -14,9195 | 2,46E-50 | 5,8E-48 |
| 188 | RM25_RS00930 | RM25_0188 | Gamma-aminobutyraldehyde dehydrogenase | 196 | gene | 235336 | 236760 | - | 1236,413 | 0,283442325 | 0,352506 | 0,804078 | 0,421352 | 0,530855 | 820,6331413 | -0,422382293 | 0,146342 | -2,88627 | 0,003898 | 0,008793 | 1652,193 | 0,639674674 | 0,204238 | 3,132008 | 0,001736 | 0,007649 | 1174,91 | 0,394118 | 0,189216 | 2,082905 | 0,03726 | 0,067193 | 1325,930598 | 1,507217094 | 0,136841 | 11,01435 | 3,26E-28 | 1,67E-26 |
| 189 | RM25_RS00935 | RM25_0189 | putative secreted protein | 197 | gene | 237259 | 237558 | + | 189,0396 | 0,700795278 | 0,613966 | 1,141424 | 0,253694 | 0,361959 | 65,87044223 | 0,009786495 | 0,300118 | 0,032609 | 0,973987 | 0,978971 | 312,2087 | 0,854120509 | 0,205912 | 4,147983 | 3,35E-05 | 0,000291 | 150,7536 | 1,691955 | 0,243467 | 6,949427 | 3,67E-12 | 8,73E-11 | 229,3679868 | 2,596853366 | 0,23068 | 11,25736 | 2,13E-29 | 1,26E-27 |
| 190 | RM25_RS00940 | RM25_0190 | putative secreted protein | 198 | gene | 237593 | 237802 | + | 283,9953 | 0,783390577 | 0,472138 | 1,659242 | 0,097067 | 0,189569 | 144,0657692 | 0,026092982 | 0,256885 | 0,101575 | 0,919094 | 0,938201 | 423,9249 | 1,059248497 | 0,278712 | 3,800508 | 0,000144 | 0,00101 | 219,7628 | 0,875472 | 0,248236 | 3,526777 | 0,000421 | 0,001712 | 349,9873877 | 1,958195267 | 0,263027 | 7,444835 | 9,71E-14 | 9,74E-13 |
| 191 | RM25_RS00945 | RM25_0191 | Hypothetical protein | 199 | gene | 237874 | 238245 | + | 774,0049 | 1,147903849 | 0,390161 | 2,942129 | 0,00326 | 0,021889 | 870,1788262 | 0,617291567 | 0,246678 | 2,50242 | 0,012335 | 0,023714 | 677,8309 | 1,9712728 | 0,467703 | 4,214794 | 2,5E-05 | 0,000226 | 512,3128 | -1,36395 | 0,459221 | -2,97013 | 0,002977 | 0,008748 | 1042,794816 | 0,021420028 | 0,255989 | 0,083676 | 0,933314 | 0,946025 |
| 192 | RM25_RS00950 | RM25_0192 | Hypothetical protein | 200 | gene | 238331 | 238531 | + | 477,3844 | 0,404482443 | 0,324654 | 1,245889 | 0,212805 | 0,317859 | 586,557114 | 0,143125801 | 0,273831 | 0,52268 | 0,601197 | 0,661233 | 368,2117 | 0,838310552 | 0,317374 | 2,6414 | 0,008256 | 0,026441 | 437,0984 | -1,1305 | 0,290461 | -3,8921 | 9,94E-05 | 0,000482 | 531,5220404 | -0,400469852 | 0,295791 | -1,3539 | 0,175769 | 0,228007 |
| 193 | RM25_RS00955 | RM25_0193 | Hypothetical protein | 201 | gene | 238786 | 239433 | + | 369,9499 | -0,848307987 | 0,471691 | -1,79844 | 0,072107 | 0,156642 | 217,6635209 | -1,614738921 | 0,29039 | -5,56058 | 2,69E-08 | 1,82E-07 | 522,2363 | -0,56720049 | 0,252923 | -2,24258 | 0,024924 | 0,060813 | 499,2504 | 0,858499 | 0,304712 | 2,817408 | 0,004841 | 0,012836 | 259,3818866 | 1,969861934 | 0,258624 | 7,616711 | 2,6E-14 | 2,85E-13 |
| 194 | RM25_RS00960 | RM25_0194 | Hypothetical protein | 202 | gene | 239430 | 240386 | + | 605,5167 | -0,945317505 | 0,419142 | -2,25536 | 0,02411 | 0,079348 | 399,2621234 | -1,629765964 | 0,288607 | -5,647 | 1,63E-08 | 1,14E-07 | 811,7713 | -0,648852046 | 0,236199 | -2,74706 | 0,006013 | 0,020813 | 836,7752 | 0,649895 | 0,333149 | 1,950763 | 0,051085 | 0,086376 | 405,9044344 | 1,692567256 | 0,196862 | 8,597755 | 8,13E-18 | 1,43E-16 |
| 195 | RM25_RS00965 | RM25_0195 | Multidrug resistance ABC transporter, ATP-binding/permea | 203 | gene | 240620 | 242683 | - | 1111,626 | 0,069544329 | 0,13677 | 0,508476 | 0,61112 | 0,700247 | 1036,450777 | -0,030306982 | 0,166714 | -0,18179 | 0,855748 | 0,886592 | 1186,801 | 0,155674793 | 0,213321 | 0,729769 | 0,465532 | 0,568028 | 1143,549 | 0,036768 | 0,21023 | 0,174894 | 0,861163 | 0,890637 | 1116,562425 | 0,27976156 | 0,155905 | 1,794439 | 0,072743 | 0,103473 |
| 196 | RM25_RS00970 | RM25_0196 | Multidrug resistance ABC transporter, ATP-binding/permea | 204 | gene | 242680 | 244413 | - | 807,5862 | -0,638742172 | 0,350394 | -1,82293 | 0,068314 | 0,151189 | 575,8541637 | -1,013424109 | 0,269674 | -3,75796 | 0,000171 | 0,000555 | 1039,318 | -0,442392099 | 0,291847 | -1,51584 | 0,129561 | 0,21104 | 1033,45 | 0,566126 | 0,334144 | 1,694255 | 0,090217 | 0,136396 | 620,3847027 | 1,205300882 | 0,241299 | 4,995052 | 5,88E-07 | 2,15E-06 |
| 197 | RM25_RS11685 | RM25_0197 | Transcriptional regulator, TetR family | 205 | gene | 244413 | 245105 | - | 472,4215 | -0,997738434 | 0,443242 | -2,251 | 0,024386 | 0,079665 | 307,334559 | -1,729103484 | 0,299564 | -5,77207 | 7,83E-09 | 5,82E-08 | 637,5084 | -0,691875626 | 0,312965 | -2,21072 | 0,027056 | 0,064541 | 660,6763 | 0,667513 | 0,35006 | 1,906854 | 0,05654 | 0,093914 | 309,6238252 | 1,77283706 | 0,278157 | 6,373514 | 1,85E-10 | 1,15E-09 |
| 198 | RM25_RS00980 | RM25_0198 | Dihydropteroate synthase | 206 | gene | 245183 | 246073 | - | 589,8459 | -0,204933919 | 0,297458 | -0,68895 | 0,490854 | 0,590355 | 432,9628566 | -0,337138298 | 0,170346 | -1,97914 | 0,047801 | 0,076177 | 746,7289 | -0,130817279 | 0,312788 | -0,41823 | 0,67578 | 0,754582 | 662,5084 | 0,628166 | 0,224103 | 2,803016 | 0,005063 | 0,013303 | 538,5823239 | 0,897380887 | 0,268089 | 3,34733 | 0,000816 | 0,001768 |
| 199 | RM25_RS00985 | RM25_0199 | Amino acid permease | 207 | gene | 246326 | 247828 | + | 877,6155 | -0,53629286 | 0,159454 | -3,3633 | 0,00077 | 0,008405 | 849,4514265 | -0,480269588 | 0,165565 | -2,9008 | 0,003722 | 0,008509 | 905,7797 | -0,590061627 | 0,291358 | -2,02521 | 0,042846 | 0,089927 | 1095,187 | 0,075895 | 0,185735 | 0,408621 | 0,682818 | 0,73165 | 704,4283525 | 0,027761888 | 0,269953 | 0,10284 | 0,91809 | 0,933134 |
| 200 | RM25_RS00990 | RM25_0200 | Cupin 2 barrel domain protein | 208 | gene | 247992 | 248351 | + | 197,8293 | 1,152904871 | 0,193365 | 5,962322 | 2,49E-09 | 2,55E-07 | 174,8360319 | 1,068911287 | 0,255241 | 4,187843 | 2,82E-05 | 0,00011 | 220,8226 | 1,220250558 | 0,257726 | 4,734685 | 2,19E-06 | 2,71E-05 | 129,1772 | 0,1737 | 0,254038 | 0,683755 | 0,49413 | 0,562096 | 267,4541698 | 0,380300651 | 0,238971 | 1,591409 | 0,111517 | 0,15211 |
| 201 | RM25_RS00995 | RM25_0201 | SAM-dependent methyl transferase | 209 | gene | 248351 | 249283 | + | 387,0262 | -0,030606683 | 0,426031 | -0,07184 | 0,942728 | 0,96173 | 228,8893453 | 0,174144379 | 0,268064 | 0,649637 | 0,515927 | 0,581003 | 545,1631 | -0,11911877 | 0,348958 | -0,34136 | 0,732836 | 0,801528 | 408,7194 | 1,331772 | 0,294401 | 4,523665 | 6,08E-06 | 3,95E-05 | 377,0061969 | 1,108147523 | 0,330967 | 3,348213 | 0,000813 | 0,001765 |
| 202 | RM25_RS01000 | RM25_0202 | VarM | 210 | gene | 249573 | 251267 | + | 868,366 | -0,195207941 | 0,270287 | -0,72222 | 0,470156 | 0,570743 | 689,6102732 | -0,044462427 | 0,180919 | -0,24576 | 0,805869 | 0,846633 | 1047,122 | -0,296269855 | 0,349728 | -0,84714 | 0,396915 | 0,502972 | 971,8172 | 0,655523 | 0,265152 | 2,472252 | 0,013426 | 0,029965 | 796,5382893 | 0,47044119 | 0,285595 | 1,647229 | 0,099511 | 0,137323 |
| 203 | RM25_RS01005 | RM25_0203 | Lysine--tRNA ligase | 211 | gene | 251408 | 253018 | + | 2678,899 | 0,265029844 | 0,23487 | 1,128411 | 0,259146 | 0,366852 | 3121,926216 | 0,476585742 | 0,172631 | 2,760716 | 0,005767 | 0,012437 | 2235,873 | -0,024887314 | 0,30048 | -0,08283 | 0,933991 | 0,956657 | 2578,815 | -0,26292 | 0,290474 | -0,90514 | 0,365394 | 0,440824 | 2864,140259 | -0,72658712 | 0,179642 | -4,04465 | 5,24E-05 | 0,000138 |
| 204 | RM25_RS01010 | RM25_0204 | CheR methyltransferase SAM binding domain-containing pr | 212 | gene | 253204 | 253926 | - | 967,7933 | 0,350453142 | 0,101981 | 3,436468 | 0,000589 | 0,00698 | 1010,405649 | 0,249253225 | 0,127297 | 1,958051 | 0,050224 | 0,079236 | 925,181 | 0,461656142 | 0,186288 | 2,478183 | 0,013205 | 0,037819 | 900,5351 | -0,30918 | 0,14665 | -2,10828 | 0,035007 | 0,064411 | 1062,216996 | -0,044531708 | 0,138928 | -0,32054 | 0,74856 | 0,792257 |
| 205 | RM25_RS01015 | RM25_0205 | Pyruvate dehydrogenase E1 component, alpha subunit | 213 | gene | 254071 | 255183 | + | 2207,023 | 1,744418585 | 0,412519 | 4,228694 | 2,35E-05 | 0,000538 | 3201,679672 | 1,938663276 | 0,182583 | 10,61799 | 2,46E-26 | 1,66E-24 | 1212,367 | 1,290188894 | 0,316965 | 4,070444 | 4,69E-05 | 0,000388 | 1079,285 | -0,9768 | 0,24261 | -4,02621 | 5,67E-05 | 0,000294 | 3330,146321 | -1,58037131 | 0,238792 | -6,61818 | 3,64E-11 | 2,51E-10 |
| 206 | RM25_RS01020 | RM25_0206 | Branched-chain alpha-keto acid dehydrogenase E1 beta sub | 214 | gene | 255180 | 256154 | + | 2304,4 | 1,804229629 | 0,322029 | 5,602694 | 2,11E-08 | 1,6E-06 | 3037,33103 | 2,321295274 | 0,124691 | 18,61637 | 2,37E-77 | 2,79E-74 | 1571,469 | 1,018458575 | 0,338534 | 3,008434 | 0,002626 | 0,010819 | 1080,603 | -0,02812 | 0,229883 | -0,12233 | 0,902637 | 0,922599 | 3508,595191 | -1,285244581 | 0,22887 | -5,61561 | 1,96E-08 | 8,96E-08 |
| 207 | RM25_RS01025 | RM25_0207 | Dihydrolipoamide acyltransferase | 215 | gene | 256165 | 257511 | + | 3023,231 | 2,541716124 | 0,389265 | 6,52953 | 6,6E-11 | 8,64E-09 | 4402,828164 | 3,110343996 | 0,138459 | 22,46397 | 9,3E-112 | 2,2E-108 | 1643,635 | 1,498864779 | 0,322995 | 4,640517 | 3,48E-06 | 4,06E-05 | 936,3329 | -0,15608 | 0,219534 | -0,71096 | 0,477106 | 0,548824 | 5056,789836 | -1,719612753 | 0,230691 | -7,4542 | 9,04E-14 | 9,19E-13 |
| 208 | RM25_RS11690 | RM25_0208 | UDP-N-acetylglucosamine diphosphorylase | 216 | gene | 257737 | 258771 | - | 1396,763 | -0,197383071 | 0,584683 | -0,33759 | 0,735672 | 0,806918 | 540,6419458 | 0,0049039 | 0,154395 | 0,031762 | 0,974662 | 0,979232 | 2252,885 | -0,245385656 | 0,333662 | -0,73543 | 0,462076 | 0,56548 | 1562,529 | 2,128615 | 0,330597 | 6,438701 | 1,21E-10 | 2,1E-09 | 1271,332225 | 1,918329183 | 0,14843 | 12,92414 | 3,29E-38 | 4,08E-36 |
| 209 | RM25_RS01035 | RM25_0209 | Putative membrane protein | 217 | gene | 259087 | 259374 | - | 769,2806 | -0,209775848 | 0,102487 | -2,04685 | 0,040673 | 0,109562 | 763,0766096 | -0,290864802 | 0,143104 | -2,03254 | 0,042099 | 0,068985 | 775,4846 | -0,130617091 | 0,206587 | -0,63226 | 0,527215 | 0,626018 | 872,7367 | -0,11382 | 0,169645 | -0,67096 | 0,502248 | 0,569407 | 698,1897007 | 0,097325454 | 0,16534 | 0,588637 | 0,556105 | 0,61769 |
| 210 | RM25_RS01040 | RM25_0210 | Membrane spanning protein DUF881 | 218 | gene | 259553 | 260287 | + | 839,8867 | 0,247071379 | 0,069391 | 3,560562 | 0,00037 | 0,004715 | 856,0900875 | 0,30435298 | 0,134272 | 2,266685 | 0,023409 | 0,040932 | 823,6833 | 0,190221137 | 0,14935 | 1,273662 | 0,202783 | 0,30003 | 810,55 | -0,05281 | 0,133656 | -0,39511 | 0,692758 | 0,740513 | 893,8943773 | -0,114836157 | 0,120827 | -0,95042 | 0,341899 | 0,405973 |
| 211 | RM25_RS01045 | RM25_0211 | Anthranilate synthase component II trpG | 219 | gene | 260383 | 261027 | + | 752,8841 | 0,375858645 | 0,081874 | 4,590698 | 4,42E-06 | 0,000141 | 738,6405867 | 0,374753235 | 0,151128 | 2,479708 | 0,013149 | 0,025075 | 767,1275 | 0,376313565 | 0,161881 | 2,324626 | 0,020092 | 0,051423 | 692,6677 | -0,00716 | 0,164585 | -0,04352 | 0,965289 | 0,973133 | 832,5094853 | 0,045399232 | 0,131273 | 0,345839 | 0,729464 | 0,774829 |
| 212 | RM25_RS01050 | RM25_0212 | Putative oxidoreductase YeaE, aldo/keto reductase family | 220 | gene | 261070 | 261954 | - | 1139,579 | 0,027972136 | 0,288293 | 0,097027 | 0,922705 | 0,948749 | 812,2198275 | 0,109523733 | 0,124041 | 0,882962 | 0,377257 | 0,445712 | 1466,937 | -0,017662569 | 0,232043 | -0,07612 | 0,939326 | 0,960516 | 1187,154 | 0,858896 | 0,220384 | 3,897273 | 9,73E-05 | 0,000476 | 1125,173913 | 0,777742792 | 0,133325 | 5,833427 | 5,43E-09 | 2,68E-08 |
| 213 | RM25_RS01055 | RM25_0213 | Hypothetical protein | 221 | gene | 262053 | 265448 | - | 7664,386 | -0,114399734 | 0,337383 | -0,33908 | 0,73455 | 0,806397 | 5298,297922 | -0,165583674 | 0,117821 | -1,40539 | 0,159907 | 0,214758 | 10030,47 | -0,087251773 | 0,366654 | -0,23797 | 0,811906 | 0,865899 | 8394,751 | 0,831799 | 0,30964 | 2,686337 | 0,007224 | 0,018018 | 7180,099006 | 0,944961781 | 0,221762 | 4,261144 | 2,03E-05 | 5,75E-05 |
| 214 | RM25_RS01060 | RM25_0214 | Hypothetical protein | 222 | gene | 265606 | 266346 | + | 675,8608 | 0,090875857 | 0,125855 | 0,72207 | 0,470252 | 0,570743 | 630,7506305 | -0,124404833 | 0,154199 | -0,80678 | 0,419793 | 0,487415 | 720,9709 | 0,276735391 | 0,166313 | 1,663944 | 0,096124 | 0,167329 | 691,0101 | -0,07092 | 0,148356 | -0,47801 | 0,632642 | 0,690021 | 682,3935099 | 0,380541048 | 0,135815 | 2,801914 | 0,00508 | 0,009406 |
| 215 | RM25_RS01065 | RM25_0215 | Secreted serine protease, trypsin-like serine proteases | 223 | gene | 266350 | 267792 | - | 2173,686 | -0,167983172 | 0,271239 | -0,61932 | 0,535707 | 0,631331 | 1712,375244 | -0,481179161 | 0,194406 | -2,47513 | 0,013319 | 0,025357 | 2634,997 | 0,033295884 | 0,286197 | 0,116339 | 0,907384 | 0,936285 | 2428,003 | 0,330975 | 0,313109 | 1,057061 | 0,290484 | 0,364574 | 2002,128659 | 0,887958644 | 0,139354 | 6,371966 | 1,87E-10 | 1,16E-09 |
| 216 | RM25_RS12190 | RM25_0216 | Trypsin-like serine protease | 224 | gene | 268345 | 269670 | - | 1148,63 | 0,453533712 | 0,141356 | 3,208441 | 0,001335 | 0,01196 | 1078,125075 | 0,156731184 | 0,146635 | 1,068854 | 0,285136 | 0,35279 | 1219,134 | 0,721985851 | 0,186024 | 3,881147 | 0,000104 | 0,000766 | 1024,14 | -0,20779 | 0,151877 | -1,36816 | 0,171262 | 0,231857 | 1299,688856 | 0,407685802 | 0,141065 | 2,890053 | 0,003852 | 0,007257 |
| 217 | RM25_RS01075 | RM25_0217 | Inner membrane protein ygcE | 225 | gene | 270300 | 271535 | + | 1090,849 | -0,519296529 | 0,263822 | -1,96836 | 0,049026 | 0,123062 | 1325,743194 | -0,361186791 | 0,120445 | -2,99877 | 0,002711 | 0,006425 | 855,9539 | -0,769080991 | 0,296085 | -2,5975 | 0,00939 | 0,028819 | 1359,633 | -0,52825 | 0,190833 | -2,7681 | 0,005638 | 0,01462 | 881,6520293 | -0,873076917 | 0,245938 | -3,54999 | 0,000385 | 0,000881 |
| 218 | RM25_RS12920 | RM25_0218 | Hypothetical protein | 226 | gene | 271977 | 272549 | + | 264,7893 | -1,425069697 | 0,71574 | -1,99104 | 0,046476 | 0,119006 | 137,2771703 | -3,531705974 | 0,420195 | -8,40491 | 4,28E-17 | 9,34E-16 | 392,3014 | -0,967880916 | 0,439573 | -2,20187 | 0,027675 | 0,065398 | 403,9698 | 0,967114 | 0,399064 | 2,423456 | 0,015374 | 0,033428 | 141,8167681 | 3,608144438 | 0,489461 | 7,371674 | 1,68E-13 | 1,63E-12 |
| 219 | RM25_RS01085 | RM25_0219 | Crotonobetainyl-CoA dehydrogenase | 227 | gene | 272680 | 273816 | + | 353,2128 | -1,44350315 | 0,641146 | -2,25144 | 0,024358 | 0,079665 | 203,7650084 | -3,136336568 | 0,411622 | -7,61945 | 2,55E-14 | 4,03E-13 | 502,6606 | -0,981242197 | 0,405409 | -2,42038 | 0,015504 | 0,042247 | 541,9336 | 0,798964 | 0,423324 | 1,887361 | 0,059112 | 0,097499 | 187,3090091 | 3,02642674 | 0,434829 | 6,960038 | 3,4E-12 | 2,68E-11 |
| 220 | NA | RM25_0220 | Hypothetical protein | 228 | gene | 273888 | 274367 | + | 152,8948 | -1,245803231 | 0,447926 | -2,78127 | 0,005415 | 0,031164 | 99,66690777 | -1,95189198 | 0,370895 | -5,26265 | 1,42E-07 | 8,35E-07 | 206,1227 | -0,957942845 | 0,29637 | -3,23225 | 0,001228 | 0,005737 | 226,0983 | 0,709986 | 0,332896 | 2,132754 | 0,032945 | 0,061095 | 89,03040507 | 1,771438924 | 0,374859 | 4,725616 | 2,29E-06 | 7,5E-06 |
| 221 | NA | RM25_0221 | Carnitine CoA-transferase | 229 | gene | 274371 | 275114 | + | 181,8452 | -1,10815095 | 0,407432 | -2,71984 | 0,006531 | 0,035553 | 128,8511114 | -1,778667727 | 0,332401 | -5,35097 | 8,75E-08 | 5,4E-07 | 234,8393 | -0,792836697 | 0,330317 | -2,40023 | 0,016385 | 0,044136 | 261,6753 | 0,512023 | 0,289586 | 1,768121 | 0,077041 | 0,1197 | 113,2887056 | 1,561838769 | 0,403349 | 3,872182 | 0,000108 | 0,000272 |
| 222 | RM25_RS01095 | RM25_0222 | putative crotonobetaine/carnitine-CoA ligase | 230 | gene | 275145 | 276740 | + | 393,8212 | -1,402668908 | 0,652815 | -2,14865 | 0,031662 | 0,093637 | 213,0093109 | -3,035386792 | 0,361386 | -8,39929 | 4,49E-17 | 9,71E-16 | 574,6032 | -0,984798429 | 0,458803 | -2,14645 | 0,031837 | 0,072643 | 599,7883 | 0,934637 | 0,36581 | 2,554981 | 0,010619 | 0,024856 | 213,4552829 | 3,064019378 | 0,481745 | 6,360251 | 2,01E-10 | 1,23E-09 |
| 223 | RM25_RS01100 | RM25_0223 | UfaA2 protein | 231 | gene | 276737 | 277615 | + | 319,5579 | -1,225749954 | 0,451847 | -2,71276 | 0,006673 | 0,036155 | 213,3385796 | -2,103216651 | 0,299017 | -7,03378 | 2,01E-12 | 2,55E-11 | 425,7773 | -0,864117864 | 0,298009 | -2,89964 | 0,003736 | 0,014388 | 470,4538 | 0,599581 | 0,301375 | 1,989485 | 0,046648 | 0,080489 | 188,1338261 | 1,905561348 | 0,323701 | 5,886795 | 3,94E-09 | 1,97E-08 |
| 224 | RM25_RS01105 | RM25_0224 | Putative metabolite transport protein YaaU | 232 | gene | 277826 | 278941 | + | 454,0631 | -1,464898163 | 0,511797 | -2,86226 | 0,004206 | 0,026368 | 295,4460719 | -2,511552048 | 0,340325 | -7,37986 | 1,58E-13 | 2,33E-12 | 612,6802 | -1,075068735 | 0,370278 | -2,90341 | 0,003691 | 0,014263 | 700,6692 | 0,655146 | 0,367503 | 1,782695 | 0,074636 | 0,11706 | 237,9681631 | 2,164933819 | 0,383497 | 5,645248 | 1,65E-08 | 7,64E-08 |
| 225 | RM25_RS01110 | RM25_0225 | Metallo-beta-lactamase family protein | 233 | gene | 279187 | 279831 | + | 131,1615 | -0,751406408 | 0,638682 | -1,1765 | 0,239397 | 0,347022 | 61,01159322 | -2,01585837 | 0,393995 | -5,11646 | 3,11E-07 | 1,71E-06 | 201,3115 | -0,435627013 | 0,464647 | -0,93754 | 0,348478 | 0,454797 | 172,1444 | 1,171109 | 0,302977 | 3,865336 | 0,000111 | 0,000533 | 96,54463113 | 2,828201365 | 0,536956 | 5,267097 | 1,39E-07 | 5,51E-07 |
| 226 | RM25_RS01115 | RM25_0226 | Putative electron transfer flavoprotein FixA | 234 | gene | 280015 | 280770 | + | 474,3124 | 0,2443512 | 0,260983 | 0,936273 | 0,349133 | 0,461788 | 353,6475743 | 0,004722468 | 0,178472 | 0,026461 | 0,97889 | 0,981806 | 594,9773 | 0,384738717 | 0,182312 | 2,110327 | 0,03483 | 0,077448 | 457,0669 | 0,484202 | 0,190398 | 2,543102 | 0,010987 | 0,025486 | 503,9018109 | 0,919553006 | 0,162573 | 5,656249 | 1,55E-08 | 7,18E-08 |
| 227 | RM25_RS01120 | RM25_0227 | Protein FixB | 235 | gene | 280807 | 281679 | + | 337,2465 | 0,381031174 | 0,284235 | 1,340552 | 0,180066 | 0,284652 | 244,3434568 | 0,092502512 | 0,22882 | 0,404259 | 0,686022 | 0,73901 | 430,1495 | 0,542995726 | 0,171 | 3,175412 | 0,001496 | 0,006769 | 308,3142 | 0,499417 | 0,213207 | 2,342403 | 0,01916 | 0,039649 | 373,8212131 | 1,008738175 | 0,193015 | 5,226225 | 1,73E-07 | 6,76E-07 |
| 228 | RM25_RS01125 | RM25_0228 | Putative electron transfer flavoprotein-quinone oxidoreduct | 236 | gene | 281682 | 282986 | + | 763,8344 | 0,246519663 | 0,158711 | 1,553264 | 0,12036 | 0,216888 | 672,742889 | 0,078912033 | 0,166651 | 0,473518 | 0,635844 | 0,693193 | 854,9259 | 0,377509592 | 0,187363 | 2,014857 | 0,04392 | 0,091609 | 737,6485 | 0,120942 | 0,175506 | 0,689108 | 0,490755 | 0,559608 | 811,4197519 | 0,471944885 | 0,166922 | 2,82733 | 0,004694 | 0,008739 |
| 229 | RM25_RS01130 | RM25_0229 | Ferredoxin-like protein fixX | 237 | gene | 283000 | 283293 | + | 542,4834 | 0,988588384 | 0,313845 | 3,149925 | 0,001633 | 0,013207 | 586,5701067 | 0,458185048 | 0,200407 | 2,286269 | 0,022239 | 0,039175 | 498,3967 | 1,715923613 | 0,36703 | 4,675165 | 2,94E-06 | 3,51E-05 | 388,514 | -1,14407 | 0,355853 | -3,21501 | 0,001304 | 0,004443 | 705,2035823 | 0,156311703 | 0,225909 | 0,691923 | 0,488986 | 0,550401 |
| 230 | NA | RM25_0230 | Hypothetical protein | 238 | gene | 283560 | 284369 | + | 320,1849 | -1,741988147 | 0,833408 | -2,0902 | 0,0366 | 0,102658 | 171,651724 | -5,160613713 | 0,465769 | -11,0798 | 1,57E-28 | 1,28E-26 | 468,718 | -1,19274751 | 0,335551 | -3,5546 | 0,000379 | 0,002253 | 517,1059 | 0,895109 | 0,365465 | 2,449236 | 0,014316 | 0,031506 | 145,0789961 | 4,93335651 | 0,481236 | 10,25142 | 1,17E-24 | 4,23E-23 |
| 231 | NA | RM25_0231 | Hypothetical protein | 239 | gene | 285149 | 285391 | - | 162,1509 | -1,585668967 | 0,82377 | -1,92489 | 0,054243 | 0,131128 | 89,03212368 | -4,875375552 | 0,568017 | -8,58315 | 9,23E-18 | 2,11E-16 | 235,2697 | -1,01086565 | 0,358165 | -2,82235 | 0,004767 | 0,017234 | 255,7184 | 0,794268 | 0,382164 | 2,078345 | 0,037678 | 0,067739 | 79,80062454 | 4,731741596 | 0,600637 | 7,877869 | 3,33E-15 | 4,29E-14 |
| 232 | RM25_RS01140 | RM25_0232 | Type III restriction enzyme, res subunit | 240 | gene | 285395 | 287167 | + | 1185,871 | 0,04500131 | 0,20178 | 0,223021 | 0,823519 | 0,877899 | 1081,657158 | 0,38260759 | 0,119997 | 3,188464 | 0,00143 | 0,003652 | 1290,085 | -0,236007766 | 0,306272 | -0,77058 | 0,440955 | 0,544722 | 1226,062 | 0,507779 | 0,188044 | 2,700326 | 0,006927 | 0,017388 | 1184,339162 | -0,046346089 | 0,257714 | -0,17984 | 0,857282 | 0,884062 |
| 233 | RM25_RS01145 | RM25_0233 | SanA protein | 241 | gene | 287369 | 288091 | + | 711,2695 | 0,035554839 | 0,140624 | 0,252837 | 0,800394 | 0,859841 | 665,4178761 | 0,278977944 | 0,163219 | 1,709222 | 0,08741 | 0,126086 | 757,121 | -0,17709426 | 0,185992 | -0,95216 | 0,341016 | 0,447039 | 738,7563 | 0,354604 | 0,198675 | 1,784846 | 0,074286 | 0,116806 | 706,1434176 | -0,047621614 | 0,122264 | -0,3895 | 0,696908 | 0,745964 |
| 234 | RM25_RS01150 | RM25_0234 | UDP-glucose 6-dehydrogenase udgA | 242 | gene | 288179 | 289705 | - | 402,6796 | -0,143756919 | 0,354194 | -0,40587 | 0,684838 | 0,763198 | 277,1824107 | -0,247468534 | 0,182986 | -1,35239 | 0,176251 | 0,233384 | 528,1768 | -0,089694703 | 0,394983 | -0,22709 | 0,820358 | 0,870068 | 442,4914 | 0,79412 | 0,243141 | 3,266089 | 0,00109 | 0,003791 | 376,9877192 | 1,020566275 | 0,354174 | 2,881543 | 0,003957 | 0,007438 |
| 235 | RM25_RS01155 | RM25_0235 | Arginine--tRNA ligase | 243 | gene | 289746 | 291338 | + | 1859,953 | 0,070071773 | 0,084229 | 0,831916 | 0,405456 | 0,516573 | 1896,640347 | 0,017551346 | 0,152311 | 0,115234 | 0,90826 | 0,92996 | 1823,265 | 0,123903456 | 0,141312 | 0,876806 | 0,380592 | 0,485945 | 1917,654 | -0,1697 | 0,129027 | -1,31522 | 0,188437 | 0,250506 | 1867,320631 | -0,012504219 | 0,147412 | -0,08482 | 0,932401 | 0,945641 |
| 236 | RM25_RS01160 | RM25_0236 | Transcriptional repressor RbsR | 244 | gene | 291360 | 292397 | - | 609,341 | -0,59307582 | 0,31467 | -1,88476 | 0,059463 | 0,13852 | 428,9375815 | -0,723932826 | 0,197806 | -3,65982 | 0,000252 | 0,000792 | 789,7443 | -0,525605239 | 0,261537 | -2,00968 | 0,044465 | 0,092502 | 770,2125 | 0,736247 | 0,219127 | 3,359909 | 0,00078 | 0,002845 | 477,2394311 | 0,998292055 | 0,254446 | 3,923397 | 8,73E-05 | 0,000224 |
| 237 | RM25_RS01165 | RM25_0237 | Inosose dehydratase | 245 | gene | 292635 | 293552 | + | 545,2703 | -1,737227978 | 0,7924 | -2,19236 | 0,028353 | 0,086721 | 287,3213012 | -4,555312502 | 0,407081 | -11,1902 | 4,56E-29 | 4,29E-27 | 803,2193 | -1,231620927 | 0,42313 | -2,91074 | 0,003606 | 0,014024 | 878,9332 | 0,96063 | 0,398636 | 2,409791 | 0,015962 | 0,034484 | 248,4995817 | 4,36202203 | 0,467548 | 9,329571 | 1,06E-20 | 2,58E-19 |
| 238 | RM25_RS12195 | RM25_0238 | Inositol 2-dehydrogenase | 246 | gene | 293582 | 294859 | + | 454,9737 | -1,366593881 | 0,782479 | -1,74649 | 0,080726 | 0,168108 | 229,7249619 | -4,086182788 | 0,422707 | -9,6667 | 4,18E-22 | 1,82E-20 | 680,2224 | -0,862593916 | 0,417466 | -2,06626 | 0,038804 | 0,084218 | 687,7895 | 0,944827 | 0,395828 | 2,386962 | 0,016988 | 0,035976 | 250,941433 | 4,242776338 | 0,477566 | 8,884176 | 6,44E-19 | 1,33E-17 |
| 239 | RM25_RS01175 | RM25_0239 | Putative metabolite transport protein CsbC | 247 | gene | 294865 | 296148 | + | 587,1034 | -1,510768028 | 0,53055 | -2,84755 | 0,004406 | 0,027184 | 361,1903093 | -2,531416028 | 0,353962 | -7,15165 | 8,57E-13 | 1,15E-11 | 813,0166 | -1,157718852 | 0,349534 | -3,31217 | 0,000926 | 0,004594 | 912,2998 | 0,797868 | 0,392164 | 2,034526 | 0,041899 | 0,073588 | 300,4018023 | 2,242881929 | 0,352819 | 6,357039 | 2,06E-10 | 1,25E-09 |
| 240 | RM25_RS01180 | RM25_0240 | Ferric vibriobactin ABC transporter, ATP-binding protein | 248 | gene | 296578 | 297483 | - | 509,4305 | -0,533551402 | 0,243934 | -2,18727 | 0,028722 | 0,087076 | 423,9731981 | -0,324473651 | 0,225926 | -1,4362 | 0,150946 | 0,204707 | 594,8878 | -0,687435461 | 0,31496 | -2,18261 | 0,029065 | 0,067693 | 632,9398 | 0,572004 | 0,277923 | 2,058141 | 0,039577 | 0,070084 | 409,3928046 | 0,275636542 | 0,281475 | 0,979256 | 0,327453 | 0,391582 |
| 241 | RM25_RS01185 | RM25_0241 | Ferric enterobactin transport system permease protein fepG | 249 | gene | 297480 | 298550 | - | 595,4352 | -0,316478109 | 0,294483 | -1,07469 | 0,282513 | 0,391466 | 463,0632762 | -0,096435516 | 0,217642 | -0,44309 | 0,657698 | 0,712733 | 727,8071 | -0,459144959 | 0,368033 | -1,24757 | 0,21219 | 0,309488 | 692,7423 | 0,750768 | 0,243674 | 3,081038 | 0,002063 | 0,006444 | 522,145687 | 0,457170586 | 0,345449 | 1,323408 | 0,1857 | 0,238526 |
| 242 | RM25_RS01190 | RM25_0242 | Iron-enterobactin transporter membrane protein | 250 | gene | 298547 | 299605 | - | 554,3194 | -0,25020356 | 0,307751 | -0,81301 | 0,416215 | 0,525807 | 413,7185788 | -0,037352619 | 0,228781 | -0,16327 | 0,870307 | 0,899699 | 694,9202 | -0,379400208 | 0,343604 | -1,10418 | 0,269515 | 0,371925 | 631,6726 | 0,837735 | 0,267041 | 3,137099 | 0,001706 | 0,005497 | 498,1646884 | 0,566070659 | 0,318645 | 1,776495 | 0,075651 | 0,106901 |
| 243 | RM25_RS01195 | RM25_0243 | Putative iron-siderophore transport system, periplasmic bin | 251 | gene | 299853 | 300848 | + | 1085,926 | 1,515244707 | 0,22809 | 6,643185 | 3,07E-11 | 4,52E-09 | 1081,639091 | 1,876305419 | 0,274067 | 6,846152 | 7,59E-12 | 8,85E-11 | 1090,213 | 1,195354267 | 0,250317 | 4,775358 | 1,79E-06 | 2,27E-05 | 591,3489 | 0,44867 | 0,288881 | 1,553131 | 0,120392 | 0,172816 | 1576,174506 | -0,174917894 | 0,245211 | -0,71334 | 0,475638 | 0,538462 |
| 244 | RM25_RS01200 | RM25_0244 | Polyphosphate--glucose phosphotransferase | 252 | gene | 301019 | 301804 | - | 1214,542 | 0,473118115 | 0,342191 | 1,382614 | 0,166783 | 0,269437 | 872,5392898 | 0,673643925 | 0,171735 | 3,922578 | 8,76E-05 | 0,000303 | 1556,546 | 0,364238909 | 0,398153 | 0,914822 | 0,360285 | 0,467057 | 1072,148 | 0,968048 | 0,382444 | 2,531214 | 0,011367 | 0,026189 | 1379,146755 | 0,693873567 | 0,205911 | 3,369773 | 0,000752 | 0,001645 |
| 245 | RM25_RS01205 | RM25_0245 | Transporter, DASS family | 253 | gene | 301884 | 303281 | - | 1713,076 | 0,091392678 | 0,234448 | 0,389821 | 0,696669 | 0,773821 | 1964,437 | -0,294738031 | 0,172579 | -1,70785 | 0,087665 | 0,1263 | 1461,715 | 0,617396234 | 0,195617 | 3,15615 | 0,001599 | 0,007137 | 1762,679 | -0,96754 | 0,180084 | -5,3727 | 7,76E-08 | 7,4E-07 | 1729,942143 | -0,007317808 | 0,165804 | -0,04414 | 0,964797 | 0,971391 |
| 246 | RM25_RS01210 | RM25_0246 | UDP-glucose 4-epimerase, galE | 254 | gene | 303594 | 304568 | + | 1646,01 | -0,009567809 | 0,189197 | -0,05057 | 0,959668 | 0,974138 | 1936,586974 | 0,019287767 | 0,136956 | 0,140832 | 0,888003 | 0,914782 | 1355,432 | -0,049290348 | 0,164534 | -0,29958 | 0,7645 | 0,827695 | 1749,291 | -0,54046 | 0,166644 | -3,24318 | 0,001182 | 0,004067 | 1609,385874 | -0,555411491 | 0,11495 | -4,83178 | 1,35E-06 | 4,59E-06 |
| 247 | RM25_RS01215 | RM25_0247 | Transcriptional regulator TetR | 255 | gene | 304569 | 305288 | - | 503,6921 | -0,622759115 | 0,478686 | -1,30098 | 0,193266 | 0,297342 | 294,7541292 | -1,54234916 | 0,252631 | -6,10515 | 1,03E-09 | 8,77E-09 | 712,6301 | -0,287337938 | 0,264642 | -1,08576 | 0,277586 | 0,38061 | 641,2268 | 0,765029 | 0,287203 | 2,663727 | 0,007728 | 0,019033 | 388,3854229 | 2,082786794 | 0,232607 | 8,954114 | 3,42E-19 | 7,34E-18 |
| 248 | RM25_RS01220 | RM25_0248 | Thermostable beta-glucosidase B | 256 | gene | 305658 | 308057 | + | 1416,095 | 0,211505576 | 0,199656 | 1,05935 | 0,289441 | 0,39881 | 1178,70283 | 0,186871527 | 0,144563 | 1,292664 | 0,196127 | 0,255258 | 1653,488 | 0,229086728 | 0,240005 | 0,954508 | 0,339826 | 0,445727 | 1379,071 | 0,404635 | 0,180282 | 2,244452 | 0,024803 | 0,048463 | 1491,679329 | 0,504845309 | 0,193331 | 2,611302 | 0,00902 | 0,015925 |
| 249 | RM25_RS01225 | RM25_0249 | Putative metabolite transport protein yaaU | 257 | gene | 308458 | 309846 | + | 740,902 | -1,200361112 | 0,390736 | -3,07205 | 0,002126 | 0,016009 | 519,9100264 | -1,567343627 | 0,306086 | -5,1206 | 3,05E-07 | 1,69E-06 | 961,8939 | -1,018501304 | 0,357993 | -2,84503 | 0,004441 | 0,01642 | 1084,292 | 0,6584 | 0,369845 | 1,780207 | 0,075042 | 0,117364 | 442,3054478 | 1,279658349 | 0,324583 | 3,942467 | 8,06E-05 | 0,000209 |
| 250 | RM25_RS01230 | RM25_0250 | Putative metallophosphoesterase | 258 | gene | 309839 | 311527 | + | 818,8322 | -0,143392562 | 0,206266 | -0,69518 | 0,486942 | 0,58647 | 712,4096797 | -0,113781971 | 0,188409 | -0,60391 | 0,545903 | 0,612129 | 925,2547 | -0,167632818 | 0,297121 | -0,56419 | 0,572625 | 0,664538 | 905,7748 | 0,33423 | 0,224188 | 1,49085 | 0,136001 | 0,19092 | 764,9357233 | 0,347766502 | 0,274241 | 1,268106 | 0,20476 | 0,259194 |
| 251 | RM25_RS01235 | RM25_0251 | MgtC family magnesium transporter | 259 | gene | 311534 | 312262 | + | 268,7437 | 0,150032307 | 0,220732 | 0,679702 | 0,496693 | 0,595173 | 318,1311659 | 0,36037319 | 0,160677 | 2,242837 | 0,024907 | 0,043262 | 219,3562 | -0,14463048 | 0,240939 | -0,60028 | 0,548321 | 0,645551 | 269,477 | -0,3336 | 0,156735 | -2,12843 | 0,033302 | 0,061562 | 277,8725714 | -0,779432586 | 0,241708 | -3,22469 | 0,001261 | 0,002598 |
| 252 | RM25_RS01240 | RM25_0252 | Glycerol-3-phosphate transporter | 260 | gene | 312551 | 313969 | + | 6033,48 | 1,548811953 | 0,490028 | 3,160662 | 0,001574 | 0,01296 | 8699,913105 | 1,37682618 | 0,200927 | 6,852382 | 7,26E-12 | 8,52E-11 | 3367,048 | 2,050754551 | 0,357501 | 5,736355 | 9,67E-09 | 2,11E-07 | 3284,986 | -1,93919 | 0,307414 | -6,30806 | 2,83E-10 | 4,59E-09 | 8808,88646 | -1,229458231 | 0,251638 | -4,88582 | 1,03E-06 | 3,59E-06 |
| 253 | RM25_RS01245 | RM25_0253 | Hypothetical protein | 261 | gene | 314293 | 314496 | - | 967,8937 | 0,812924107 | 0,3401 | 2,390249 | 0,016837 | 0,064844 | 665,9886387 | 0,706016565 | 0,16089 | 4,388198 | 1,14E-05 | 4,73E-05 | 1269,799 | 0,870816963 | 0,372642 | 2,336871 | 0,019446 | 0,050363 | 740,0503 | 0,770079 | 0,282739 | 2,723644 | 0,006457 | 0,016399 | 1200,929054 | 0,970829013 | 0,273113 | 3,554678 | 0,000378 | 0,000868 |
| 254 | RM25_RS01250 | RM25_0254 | NUDIX hydrolase | 262 | gene | 314589 | 315032 | - | 304,2902 | -0,509088198 | 0,145915 | -3,48894 | 0,000485 | 0,005985 | 278,9419394 | -0,296244063 | 0,184342 | -1,60704 | 0,108046 | 0,15222 | 329,6385 | -0,691471443 | 0,208658 | -3,3139 | 0,00092 | 0,004575 | 376,3526 | 0,340117 | 0,18486 | 1,839865 | 0,065788 | 0,106645 | 246,6647186 | 0,004518913 | 0,215974 | 0,020923 | 0,983307 | 0,986656 |
| 255 | RM25_RS01255 | RM25_0255 | putative sugar kinase YegV, PfkB family | 263 | gene | 315205 | 316209 | - | 793,7282 | -0,804001092 | 0,230559 | -3,48718 | 0,000488 | 0,005992 | 909,6522252 | -0,998685908 | 0,177788 | -5,61729 | 1,94E-08 | 1,34E-07 | 677,8042 | -0,552117228 | 0,338196 | -1,63253 | 0,102567 | 0,175436 | 1067,61 | -0,64468 | 0,185354 | -3,47809 | 0,000505 | 0,001987 | 568,7023583 | -0,137540318 | 0,317 | -0,43388 | 0,664375 | 0,717659 |
| 256 | RM25_RS01260 | RM25_0256 | Permease, cytosine/purine, uracil, thiamine, allantoin fam | 264 | gene | 316206 | 317726 | - | 925,0116 | -1,433313064 | 0,307042 | -4,66814 | 3,04E-06 | 0,000102 | 986,7015068 | -2,203941196 | 0,185419 | -11,8863 | 1,4E-32 | 2,19E-30 | 863,3218 | -0,736816522 | 0,369215 | -1,99563 | 0,045974 | 0,095054 | 1428,174 | -0,6473 | 0,224486 | -2,88347 | 0,003933 | 0,010881 | 491,8724165 | 0,884408204 | 0,338495 | 2,612764 | 0,008981 | 0,015869 |
| 257 | RM25_RS01265 | RM25_0257 | Inositol monophosphatase family protein | 265 | gene | 317904 | 318755 | - | 390,4471 | -0,855076456 | 0,418263 | -2,04435 | 0,040919 | 0,109812 | 257,3993936 | -1,297498073 | 0,309943 | -4,18625 | 2,84E-05 | 0,00011 | 523,4949 | -0,654515824 | 0,365899 | -1,78879 | 0,073649 | 0,13583 | 528,5701 | 0,734863 | 0,322193 | 2,28082 | 0,022559 | 0,044947 | 273,6751638 | 1,451639196 | 0,379568 | 3,82445 | 0,000131 | 0,000327 |
| 258 | RM25_RS01270 | RM25_0258 | Hexose phosphate transport protein | 266 | gene | 318748 | 320001 | - | 713,6513 | -1,162908051 | 0,415678 | -2,79762 | 0,005148 | 0,03026 | 463,4692595 | -1,66749368 | 0,225085 | -7,4083 | 1,28E-13 | 1,9E-12 | 963,8334 | -0,944695569 | 0,339123 | -2,7857 | 0,005341 | 0,018846 | 1038,477 | 0,788307 | 0,365014 | 2,159661 | 0,030799 | 0,057797 | 431,1539652 | 1,559391584 | 0,178944 | 8,714393 | 2,92E-18 | 5,56E-17 |
| 259 | RM25_RS01275 | RM25_0259 | Transposase insI for insertion sequence element IS30B/C/D | 267 | gene | 320055 | 321089 | + | 22,44666 | -1,766281556 | 0,713846 | -2,47432 | 0,013349 | 0,056794 | 11,92708672 | -3,489315895 | 0,985166 | -3,54185 | 0,000397 | 0,001187 | 32,96623 | -1,377787538 | 0,428847 | -3,21277 | 0,001315 | 0,006087 | 36,34264 | 1,050351 | 0,421084 | 2,494401 | 0,012617 | 0,028567 | 9,952248408 | 3,221497058 | 1,081 | 2,980109 | 0,002881 | 0,005562 |
| 260 | NA | RM25_0260 | Hypothetical protein | 268 | gene | 321211 | 321324 | + | 77,24769 | -1,864901929 | 0,694846 | -2,68391 | 0,007277 | 0,038456 | 42,26511063 | -3,781113046 | 0,623217 | -6,06709 | 1,3E-09 | 1,1E-08 | 112,2303 | -1,42919907 | 0,360798 | -3,96121 | 7,46E-05 | 0,00058 | 127,2522 | 0,978247 | 0,396869 | 2,464913 | 0,013705 | 0,030416 | 32,68152264 | 3,403200836 | 0,665749 | 5,11184 | 3,19E-07 | 1,2E-06 |
| 261 | RM25_RS01280 | RM25_0261 | Acetyl-CoA C-acetyltransferase vraB | 269 | gene | 321684 | 322838 | - | 382,8402 | -0,77389543 | 0,925786 | -0,83593 | 0,403192 | 0,513967 | 137,3055038 | -4,555548251 | 0,461547 | -9,87016 | 5,61E-23 | 2,7E-21 | 628,375 | -0,344113255 | 0,45814 | -0,75111 | 0,452586 | 0,555887 | 503,5549 | 1,348215 | 0,405324 | 3,326269 | 0,00088 | 0,003157 | 279,2244883 | 5,632944887 | 0,50899 | 11,06691 | 1,82E-28 | 9,73E-27 |
| 262 | RM25_RS01285 | RM25_0262 | Mycobactin-specific salicyl-AMP ligase | 270 | gene | 322835 | 323917 | - | 550,6449 | -1,138008834 | 0,696545 | -1,63379 | 0,102303 | 0,195562 | 280,2491603 | -3,273661176 | 0,291946 | -11,2132 | 3,51E-29 | 3,45E-27 | 821,0406 | -0,663867272 | 0,405154 | -1,63856 | 0,101306 | 0,174505 | 793,4167 | 0,916613 | 0,313785 | 2,921146 | 0,003487 | 0,009976 | 339,4297395 | 3,600882683 | 0,399425 | 9,01517 | 1,97E-19 | 4,37E-18 |
| 263 | RM25_RS01290 | RM25_0263 | Hypothetical protein | 271 | gene | 324088 | 324306 | + | 1697,843 | -0,146465428 | 0,430054 | -0,34057 | 0,733424 | 0,805536 | 1864,200927 | -1,163981406 | 0,158986 | -7,32127 | 2,46E-13 | 3,51E-12 | 1531,486 | 1,066492431 | 0,494316 | 2,157513 | 0,030966 | 0,071137 | 1905,07 | -1,42889 | 0,44011 | -3,24666 | 0,001168 | 0,004024 | 1573,442817 | 0,834525 | 0,285135 | 2,926768 | 0,003425 | 0,006521 |
| 264 | NA | RM25_0264 | Hypothetical protein | 272 | gene | 324361 | 324567 | + | 36,08258 | -1,351362168 | 0,782399 | -1,7272 | 0,084131 | 0,172732 | 17,89071835 | -3,457759423 | 0,827141 | -4,18038 | 2,91E-05 | 0,000113 | 54,27444 | -0,915456187 | 0,508395 | -1,80068 | 0,071753 | 0,13329 | 54,45293 | 1,024045 | 0,360922 | 2,8373 | 0,00455 | 0,012214 | 19,96971311 | 3,664200019 | 0,980268 | 3,737958 | 0,000186 | 0,00045 |
| 265 | RM25_RS01295 | RM25_0265 | CAAX amino terminal protease self-immunity | 273 | gene | 324543 | 325385 | + | 359,4744 | -1,547659063 | 0,634411 | -2,43952 | 0,014707 | 0,06039 | 208,3974402 | -3,264677267 | 0,338465 | -9,64553 | 5,13E-22 | 2,12E-20 | 510,5514 | -1,08573189 | 0,409838 | -2,64918 | 0,008069 | 0,026088 | 561,9921 | 0,807956 | 0,332048 | 2,433254 | 0,014964 | 0,032718 | 180,7159807 | 3,062362331 | 0,444797 | 6,884854 | 5,78E-12 | 4,41E-11 |
| 266 | RM25_RS01300 | RM25_0266 | Ferritin-like protein | 274 | gene | 325543 | 326070 | + | 29094,17 | -1,377199284 | 0,296088 | -4,65132 | 3,3E-06 | 0,000108 | 30552,41585 | -1,281457753 | 0,264996 | -4,83577 | 1,33E-06 | 6,6E-06 | 27635,93 | -1,486390525 | 0,458709 | -3,24038 | 0,001194 | 0,005605 | 44527,16 | -0,13118 | 0,472457 | -0,27765 | 0,781278 | 0,823188 | 15826,79948 | -0,307488056 | 0,247326 | -1,24325 | 0,213775 | 0,269448 |
| 267 | RM25_RS01305 | RM25_0267 | Transposase insI for insertion sequence element IS30B/C/D | 275 | gene | 326415 | 327449 | - | 6,142606 | -0,834564869 | 0,499581 | -1,67053 | 0,094815 | 0,186543 | 4,908452744 | -1,379937651 | 1,04376 | -1,32208 | 0,18614 | 0,244419 | 7,376759 | -0,552621428 | 0,577942 | -0,95619 | 0,338977 | 0,445108 | 8,423025 | 0,234287 | 0,534801 | 0,438083 | 0,661326 | 0,715338 | 4,26405965 | 1,12345865 | 1,159899 | 0,968583 | 0,332753 | 0,396511 |
| 268 | RM25_RS01310 | RM25_0268 | Hypothetical protein | 276 | gene | 327648 | 328445 | - | 2068,79 | -0,022635188 | 0,35129 | -0,06443 | 0,948624 | 0,966252 | 1520,665473 | 0,325814031 | 0,195946 | 1,662771 | 0,096358 | 0,137563 | 2616,915 | -0,223979996 | 0,43648 | -0,51315 | 0,607846 | 0,69582 | 2194,676 | 1,016362 | 0,40483 | 2,510587 | 0,012053 | 0,027502 | 2002,987152 | 0,49556276 | 0,246868 | 2,007401 | 0,044707 | 0,067075 |
| 269 | RM25_RS01315 | RM25_0269 | Hypothetical protein | 277 | gene | 328553 | 330088 | - | 2275,225 | 0,339894005 | 0,2227 | 1,52624 | 0,12695 | 0,224809 | 1920,312059 | -0,011047046 | 0,127961 | -0,08633 | 0,931203 | 0,94797 | 2630,138 | 0,601965275 | 0,292955 | 2,054806 | 0,039898 | 0,085862 | 2124,093 | 0,061029 | 0,221742 | 0,275224 | 0,783144 | 0,824714 | 2481,832292 | 0,713543088 | 0,210377 | 3,39173 | 0,000695 | 0,001527 |
| 270 | RM25_RS01320 | RM25_0270 | Transposase IS204/IS1001/IS1096/IS1165 family protein | 278 | gene | 330586 | 331551 | + | 4,353005 | -1,717752751 | 0,795869 | -2,15834 | 0,030902 | 0,092431 | 3,227483814 | -2,821972653 | 1,723271 | -1,63757 | 0,101512 | 0,144135 | 5,478525 | -1,297658449 | 0,773392 | -1,67788 | 0,09337 | 0,163746 | 7,05785 | 0,390098 | 0,698096 | 0,558803 | 0,576296 | 0,639817 | 1,951167209 | 1,960825104 | 2,118614 | 0,925523 | 0,354694 | 0,418635 |
| 271 | NA | RM25_0271 | Hypothetical protein | 279 | gene | 332175 | 332432 | + | 378,3037 | 0,158490649 | 0,568178 | 0,278945 | 0,780287 | 0,842867 | 163,5252168 | -0,496297219 | 0,213173 | -2,32815 | 0,019904 | 0,035541 | 593,0282 | 0,338226727 | 0,411143 | 0,822649 | 0,410708 | 0,516012 | 375,8511 | 1,399666 | 0,337372 | 4,148731 | 3,34E-05 | 0,000183 | 387,4702317 | 2,264878536 | 0,307042 | 7,376446 | 1,63E-13 | 1,58E-12 |
| 272 | RM25_RS01325 | RM25_0272 | Transposase | 280 | gene | 332375 | 333754 | - | 59,41561 | 0,134859867 | 0,264859 | 0,509175 | 0,610629 | 0,700247 | 50,80071771 | -0,452005503 | 0,34885 | -1,2957 | 0,195078 | 0,254173 | 68,03051 | 0,554938469 | 0,274237 | 2,023571 | 0,043014 | 0,0902 | 60,01941 | -0,15982 | 0,261731 | -0,61061 | 0,541457 | 0,607721 | 60,73575709 | 0,907374578 | 0,351517 | 2,581311 | 0,009843 | 0,017236 |
| 273 | NA | RM25_0273 | Hypothetical protein | 281 | gene | 333862 | 333969 | - | 33,38234 | -1,120484653 | 0,411016 | -2,72614 | 0,006408 | 0,035179 | 23,26660401 | -1,638388337 | 0,562111 | -2,91471 | 0,00356 | 0,008187 | 43,49807 | -0,899761027 | 0,336375 | -2,67488 | 0,007476 | 0,024644 | 48,01133 | 0,615319 | 0,361498 | 1,702137 | 0,08873 | 0,134579 | 20,54146211 | 1,421185015 | 0,565589 | 2,512753 | 0,011979 | 0,020655 |
| 274 | RM25_RS11695 | RM25_0274 | Hypothetical protein | 282 | gene | 334694 | 335374 | - | 1095,072 | -0,307721293 | 0,365576 | -0,84174 | 0,399932 | 0,511193 | 738,4131173 | -0,440072119 | 0,20383 | -2,15901 | 0,030849 | 0,052424 | 1451,731 | -0,241085957 | 0,379385 | -0,63547 | 0,525125 | 0,62448 | 1274,926 | 0,838602 | 0,285299 | 2,939381 | 0,003289 | 0,009558 | 953,1235041 | 1,066548946 | 0,300891 | 3,544639 | 0,000393 | 0,000895 |
| 276 | RM25_RS01350 | RM25_0275 | Hypothetical protein | 286 | gene | 336326 | 337018 | - | 796,3051 | 0,314066199 | 0,148784 | 2,110887 | 0,034782 | 0,099492 | 870,1339741 | 0,457538412 | 0,195868 | 2,335951 | 0,019494 | 0,034941 | 722,4763 | 0,14581345 | 0,153268 | 0,951365 | 0,341419 | 0,447318 | 749,8591 | -0,15192 | 0,216964 | -0,70021 | 0,483799 | 0,554326 | 865,7744686 | -0,412581891 | 0,1359 | -3,03591 | 0,002398 | 0,004722 |
| 277 | RM25_RS01355 | RM25_0276 | Hypothetical protein | 287 | gene | 337203 | 338759 | - | 598,2778 | -1,246230197 | 0,326785 | -3,81361 | 0,000137 | 0,002138 | 446,4734398 | -1,250803765 | 0,259913 | -4,8124 | 1,49E-06 | 7,37E-06 | 750,0822 | -1,245312019 | 0,381609 | -3,26332 | 0,001101 | 0,00525 | 884,4039 | 0,683859 | 0,284615 | 2,402752 | 0,016272 | 0,035018 | 349,5105571 | 0,757115519 | 0,377455 | 2,005846 | 0,044873 | 0,067238 |
| 278 | RM25_RS01360 | RM25_0277 | HAD-superfamily hydrolase, subfamily IA, variant 1 | 288 | gene | 339267 | 339920 | + | 783,4561 | 0,244211371 | 0,186403 | 1,310125 | 0,190154 | 0,294091 | 674,7926422 | 0,52477488 | 0,140256 | 3,741553 | 0,000183 | 0,00059 | 892,1195 | 0,035813129 | 0,196119 | 0,182609 | 0,855105 | 0,894577 | 754,3575 | 0,613677 | 0,181182 | 3,387073 | 0,000706 | 0,00264 | 833,3051643 | 0,175630025 | 0,152928 | 1,148451 | 0,250782 | 0,308988 |
| 279 | RM25_RS01365 | RM25_0278 | D-isomer specific 2-hydroxyacid dehydrogenase NAD-bindin | 289 | gene | 339999 | 341048 | + | 843,1089 | -0,074764731 | 0,206409 | -0,36222 | 0,71719 | 0,792507 | 696,4698715 | 0,089287007 | 0,157346 | 0,567455 | 0,570405 | 0,632085 | 989,7479 | -0,191668441 | 0,227818 | -0,84132 | 0,400167 | 0,505734 | 908,1685 | 0,582001 | 0,200506 | 2,902656 | 0,0037 | 0,01042 | 805,9541297 | 0,359067848 | 0,178931 | 2,006742 | 0,044777 | 0,067137 |
| 280 | RM25_RS01370 | RM25_0279 | Putative 2-deoxy-D-gluconate 3-dehydrogenase | 290 | gene | 341088 | 341900 | + | 512,5075 | 0,201330178 | 0,148767 | 1,353328 | 0,175951 | 0,279647 | 476,8520711 | -0,013150179 | 0,172834 | -0,07609 | 0,939351 | 0,95228 | 548,163 | 0,386249464 | 0,220047 | 1,755303 | 0,079208 | 0,144275 | 503,3915 | -0,07443 | 0,193918 | -0,38383 | 0,701103 | 0,747737 | 537,6218976 | 0,384298278 | 0,190758 | 2,014583 | 0,043948 | 0,066105 |
| 281 | RM25_RS01375 | RM25_0280 | Succinate CoA transferase | 291 | gene | 342128 | 343651 | - | 16520,99 | 0,271838916 | 0,321443 | 0,845684 | 0,397729 | 0,508779 | 19578,94374 | 0,048083505 | 0,203649 | 0,23611 | 0,813347 | 0,852026 | 13463,04 | 0,604068388 | 0,444923 | 1,357693 | 0,174561 | 0,265788 | 15927,76 | -0,89586 | 0,406419 | -2,20427 | 0,027505 | 0,052664 | 17668,25689 | -0,311499749 | 0,265714 | -1,17231 | 0,241071 | 0,298427 |
| 282 | RM25_RS01380 | RM25_0281 | Uncharacterized conserved protein UCP033563 | 292 | gene | 343775 | 344965 | - | 3049,215 | 0,479282506 | 0,181269 | 2,644045 | 0,008192 | 0,040909 | 3015,190087 | 0,444878984 | 0,175431 | 2,535919 | 0,011215 | 0,021832 | 3083,24 | 0,51342937 | 0,323471 | 1,58725 | 0,112456 | 0,188789 | 2696,377 | -0,05992 | 0,284237 | -0,21082 | 0,833031 | 0,868017 | 3471,769096 | 0,04585883 | 0,22076 | 0,207731 | 0,835439 | 0,867076 |
| 283 | RM25_RS01385 | RM25_0282 | Membrane family protein | 293 | gene | 345278 | 346618 | + | 1159,481 | 0,126202029 | 0,152436 | 0,827902 | 0,407726 | 0,518904 | 1289,41147 | 0,264959513 | 0,124934 | 2,120802 | 0,033938 | 0,056894 | 1029,55 | -0,044320763 | 0,207314 | -0,21379 | 0,830714 | 0,876845 | 1171,07 | -0,22523 | 0,168131 | -1,33962 | 0,18037 | 0,241416 | 1187,707494 | -0,478200438 | 0,151491 | -3,15662 | 0,001596 | 0,003224 |
| 284 | RM25_RS13105 | RM25_0283 | Hypothetical protein | 294 | gene | 346756 | 347001 | - | 412,884 | 0,366890645 | 0,458376 | 0,800413 | 0,423471 | 0,5329 | 217,4023832 | 0,366978987 | 0,207672 | 1,767109 | 0,07721 | 0,113527 | 608,3656 | 0,364772952 | 0,300958 | 1,21204 | 0,225497 | 0,322706 | 378,3639 | 1,430271 | 0,296149 | 4,829561 | 1,37E-06 | 1,02E-05 | 453,8222297 | 1,465434374 | 0,196654 | 7,451822 | 9,21E-14 | 9,31E-13 |

| | | | | | | | | | | |
|---|---|---|---|---|---|---|---|---|---|---|
| 285 | RM25_RS13110 | RM25_0284 | Hypothetical protein | 295 | gene | 347016 | 347246 | + | 138,9173 | -1,23877161 |
| 286 | RM25_RS01395 | RM25_0285 | Hypothetical protein | 296 | gene | 347498 | 347968 | - | 187,1516 | -1,460594568 |
| 287 | RM25_RS01400 | RM25_0286 | 2, 5-didehydrogluconate reductase | 297 | gene | 348234 | 349067 | - | 2496,322 | 0,659838597 |
| 288 | RM25_RS01405 | RM25_0287 | putative membrane spanning protein | 298 | gene | 349217 | 349990 | - | 796,6853 | -0,618502579 |
| 289 | RM25_RS01415 | RM25_0288 | Ribosomal RNA large subunit methyltransferase G | 299 | gene | 350197 | 351186 | - | 712,6931 | -0,264760829 |
| 290 | RM25_RS01415 | RM25_0289 | DNA-damage-inducible protein F DinF | 300 | gene | 351237 | 352601 | + | 480,811 | -0,352240117 |
| 291 | RM25_RS01420 | RM25_0290 | Putative insertion element ISCmi2 transposase | 301 | gene | 352728 | 353771 | - | 25,31472 | -1,332999966 |
| 292 | RM25_RS01425 | RM25_0291 | Mrr restriction system protein | 302 | gene | 353927 | 354868 | - | 503,9312 | -0,492432523 |
| 293 | RM25_RS01430 | RM25_0292 | Protein of hypothetical function DUF299 | 303 | gene | 355380 | 356273 | + | 1061,534 | -0,38824186 |
| 294 | RM25_RS01435 | RM25_0293 | Pyruvate, phosphate dikinase | 304 | gene | 356482 | 359139 | + | 6101,739 | -0,057024312 |
| 295 | RM25_RS01445 | RM25_0294 | Hypothetical protein | 305 | gene | 359601 | 362327 | + | 2000,85 | 0,247857334 |
| 296 | RM25_RS01450 | RM25_0295 | Ribokinase | 306 | gene | 362546 | 363538 | + | 487,3263 | -0,050571671 |
| 297 | RM25_RS01455 | RM25_0296 | Two component system sensor kinase phoR | 307 | gene | 363794 | 365659 | - | 1424,295 | 0,129969635 |
| 298 | RM25_RS01460 | RM25_0297 | Two component system transcriptional regulator trcR | 308 | gene | 365640 | 366368 | - | 1780,162 | -0,043354582 |
| 300 | RM25_RS01465 | RM25_0298 | NLPA lipoprotein | 309 | gene | 366919 | 367884 | + | 1214,746 | -0,371062378 |
| 301 | RM25_RS01470 | RM25_0299 | Methionine import ATP-binding protein MetN 2 | 310 | gene | 367881 | 369146 | + | 1857,642 | 0,810236269 |
| 302 | RM25_RS01475 | RM25_0300 | ABC D-methionine transporter, permease component | 311 | gene | 369143 | 369799 | + | 1074,147 | 0,837242424 |
| 303 | RM25_RS01480 | RM25_0301 | Hypothetical protein | 312 | gene | 369924 | 371543 | - | 1237,602 | -0,818040182 |
| 304 | RM25_RS01485 | RM25_0302 | Hypothetical protein | 313 | gene | 371616 | 372278 | - | 1381,239 | 0,466126487 |
| 305 | RM25_RS01490 | RM25_0303 | PadR, Transcriptional regulator PadR-like family | 314 | gene | 372573 | 373145 | + | 375,0073 | -1,534646743 |
| 306 | RM25_RS01495 | RM25_0304 | ABC-1 domain protein | 315 | gene | 373188 | 374885 | + | 783,0516 | -0,61779544 |
| 307 | RM25_RS01500 | RM25_0305 | Putative hydroxymethylpyrimidine transporter CytX | 316 | gene | 374998 | 376314 | - | 486,2552 | -1,072912551 |
| 308 | RM25_RS01505 | RM25_0306 | Phosphomethylpyrimidine kinase | 317 | gene | 376393 | 377259 | - | 374,2508 | -0,811356504 |
| 309 | RM25_RS01510 | RM25_0307 | Thiamine-phosphate synthase 2 | 318 | gene | 377256 | 378071 | - | 237,0058 | -0,245186834 |
| 310 | RM25_RS01515 | RM25_0308 | Hydroxyethylthiazole kinase | 319 | gene | 378104 | 378964 | - | 264,2983 | -0,871835969 |
| 311 | RM25_RS01520 | RM25_0309 | Aromatic amino acid transport protein AroP | 320 | gene | 379392 | 380786 | + | 891,3621 | -0,371175898 |
| 312 | RM25_RS01525 | RM25_0310 | Cobalt-zinc-cadmium efflux permease | 321 | gene | 380903 | 381853 | + | 637,3415 | -0,227636116 |
| 313 | RM25_RS01530 | RM25_0311 | Hypothetical protein | 322 | gene | 382334 | 382888 | + | 1840,136 | 0,488986744 |
| 314 | NA | RM25_0312 | IS1096, TnpA family protein | 323 | gene | 383348 | 383965 | - | 60,29989 | -1,333586441 |
| 315 | RM25_RS01540 | RM25_0313 | Hypothetical protein | 324 | gene | 385177 | 385470 | + | 255,003 | -0,213837393 |
| 316 | RM25_RS01545 | RM25_0314 | Hypothetical protein | 325 | gene | 385474 | 385779 | - | 446,4945 | 0,567700732 |
| 317 | RM25_RS01550 | RM25_0315 | Hypothetical protein | 326 | gene | 385776 | 386051 | - | 1745,798 | 0,322573524 |
| 318 | RM25_RS01555 | RM25_0316 | Endonuclease and methylase LlaGI | 327 | gene | 386128 | 389904 | + | 4336,505 | -0,11017902 |
| 319 | NA | RM25_0317 | IS3 family element | 328 | gene | 389956 | 390300 | + | 0 | NA |
| 320 | RM25_RS01565 | RM25_0318 | Transposase IS6110 | 329 | gene | 390333 | 391220 | + | 19,53992 | -1,899505544 |
| 321 | RM25_RS01570 | RM25_0319 | Endonuclease and methylase LlaGI | 330 | gene | 391264 | 392343 | + | 998,7762 | -0,61287302 |
| 322 | RM25_RS01575 | RM25_0320 | HipA domain protein | 331 | gene | 392340 | 393311 | + | 387,0407 | -0,785090551 |
| 323 | RM25_RS11700 | RM25_0321 | HIRAN domain family | 332 | gene | 393289 | 394098 | + | 4086,889 | 0,438614116 |
| 324 | RM25_RS12690 | RM25_0322 | Hypothetical protein | 333 | gene | 394108 | 394257 | + | 90,8194 | 0,56935757 |
| 325 | RM25_RS01585 | RM25_0323 | Hypothetical protein | 334 | gene | 394254 | 394760 | + | 640,1352 | -0,180606104 |
| 326 | RM25_RS12520 | RM25_0324 | Hypothetical protein | 335 | gene | 394870 | 396123 | - | 2059,297 | -0,404905745 |
| 327 | RM25_RS01590 | RM25_0325 | Transposon Tn21 resolvase | 336 | gene | 396171 | 396746 | - | 549,7377 | -1,568998602 |
| 328 | RM25_RS01595 | RM25_0326 | Ribosomal protein L6P/L9E | 337 | gene | 397043 | 397957 | - | 1621,449 | -0,413795081 |
| 329 | RM25_RS01600 | RM25_0327 | Hypothetical protein | 338 | gene | 398113 | 398526 | - | 2633,452 | -0,601417544 |
| 330 | RM25_RS01605 | RM25_0328 | Transposase IS204/IS1001/IS1096/IS1165 family protein | 339 | gene | 398849 | 400156 | - | 8,845863 | -1,900431704 |
| 331 | RM25_RS01610 | RM25_0329 | Transposase IS204/IS1001/IS1096/IS1165 family protein | 340 | gene | 400281 | 401357 | - | 15,77527 | -1,319366047 |
| 332 | RM25_RS01615 | RM25_0330 | Hypothetical protein | 341 | gene | 402033 | 402272 | + | 94,33873 | -0,544735062 |
| 333 | RM25_RS01620 | RM25_0331 | Type III restriction protein res subunit | 342 | gene | 402273 | 405437 | - | 2139,566 | -0,177675133 |
| 334 | RM25_RS01625 | RM25_0332 | Putative insertion element ISCmi2 transposase | 343 | gene | 405666 | 406709 | + | 45,439 | 0,135617153 |
| 335 | RM25_RS01630 | RM25_0333 | Transporter, anaerobic C4-dicarboxylate uptake family prot | 344 | gene | 406772 | 408088 | - | 1104,005 | -1,491418411 |
| 336 | RM25_RS01635 | RM25_0334 | Hypothetical protein | 345 | gene | 408485 | 408760 | - | 234,4078 | -0,671396522 |
| 337 | RM25_RS01640 | RM25_0335 | PF07510 family protein | 346 | gene | 409218 | 409967 | + | 775,9532 | -1,043349228 |
| 338 | RM25_RS01645 | RM25_0336 | Hypothetical protein | 347 | gene | 410057 | 410299 | + | 2023,029 | -1,449334031 |
| 339 | RM25_RS12945 | RM25_0337 | Hypothetical protein | 348 | gene | 410413 | 410550 | + | 146,4533 | -0,694596934 |
| 340 | RM25_RS01650 | RM25_0338 | Zinc-binding alcohol dehydrogenase family protein | 349 | gene | 410693 | 411427 | + | 676,8507 | -0,851210881 |
| 341 | RM25_RS01655 | RM25_0339 | Hypothetical protein | 350 | gene | 411683 | 412198 | + | 191,999 | -0,470863219 |
| 342 | RM25_RS01660 | RM25_0340 | Hypothetical protein | 351 | gene | 412233 | 412739 | - | 1811,906 | 0,49255662 |
| 344 | NA | RM25_0341 | Hypothetical protein | 352 | gene | 414215 | 414487 | + | 27,98107 | -1,372976548 |
| 345 | NA | RM25_0342 | Hypothetical protein | 353 | gene | 414895 | 415008 | + | 36,26543 | -1,271637606 |
| 346 | NA | RM25_0343 | Hypothetical protein | 354 | gene | 416057 | 416320 | + | 62,06273 | -1,901797755 |
| 347 | RM25_RS01665 | RM25_0344 | Transposase IS204/IS1001/IS1096/IS1165 family protein | 355 | gene | 416462 | 417769 | + | 12,84683 | -1,932252678 |
| 348 | NA | RM25_0345 | Hypothetical protein | 356 | gene | 418696 | 418818 | + | 23,4696 | -1,59864031 |
| 349 | RM25_RS01670 | RM25_0346 | CRISPR-associated endoribonuclease Cas2 | 357 | gene | 419700 | 420002 | - | 187,2551 | -0,206921919 |
| 350 | RM25_RS01675 | RM25_0347 | CRISPR-associated endonuclease Cas1 | 358 | gene | 419999 | 421600 | - | 1074,619 | -0,348216517 |
| 351 | RM25_RS01680 | RM25_0348 | Putative CRISPR-associated Csb3 family protein | 359 | gene | 421590 | 422591 | - | 669,8471 | -0,359316931 |
| 352 | RM25_RS01685 | RM25_0349 | CRISPR-associated helicase Cas3, Anaes-subtype | 360 | gene | 422588 | 425203 | - | 1560,9 | -0,559866634 |
| 353 | RM25_RS01690 | RM25_0350 | Putative CRISPR-associated Csb2 family protein | 361 | gene | 425200 | 426741 | - | 904,2559 | -1,122405849 |
| 354 | RM25_RS01695 | RM25_0351 | Putative CRISPR-associated Csb1 family protein | 362 | gene | 426743 | 427942 | - | 1066,205 | -0,745037894 |
| 356 | RM25_RS01705 | RM25_0352 | Threonine ammonia-lyase | 366 | gene | 428471 | 429745 | + | 534,3648 | -0,768455046 |
| 357 | RM25_RS01710 | RM25_0353 | Hemerythrin HHE cation binding region | 367 | gene | 430219 | 430506 | + | 1004,004 | 0,825914473 |
| 358 | RM25_RS11705 | RM25_0354 | Putative SufR family transcriptional regulator | 368 | gene | 430512 | 431522 | + | 712,0206 | -0,15050958 |
| 359 | RM25_RS01720 | RM25_0355 | Hypothetical protein | 369 | gene | 431570 | 432874 | + | 1059,464 | -0,497171556 |
| 360 | RM25_RS01725 | RM25_0356 | Copper-containing nitrite reductase (Major outer membrane | 370 | gene | 432871 | 435507 | + | 1154,97 | -0,583763557 |
| 361 | RM25_RS01730 | RM25_0357 | TQXA domain-containing protein | 371 | gene | 435661 | 437040 | + | 673,1867 | -1,044317132 |
| 362 | RM25_RS11710 | RM25_0358 | Peptidase C60, sortase A and B | 372 | gene | 437182 | 437916 | + | 256,343 | -0,698870354 |
| 363 | RM25_RS12205 | RM25_0359 | Hypothetical protein | 373 | gene | 437986 | 441390 | + | 3214,443 | 0,308177007 |
| 364 | RM25_RS01745 | RM25_0360 | Hypothetical protein | 374 | gene | 441641 | 441949 | + | 373,0281 | 0,853007252 |
| 365 | RM25_RS01750 | RM25_0361 | Recombination protein RecR | 375 | gene | 441951 | 442577 | + | 627,6909 | 0,543770183 |
| 366 | RM25_RS01755 | RM25_0362 | Hypothetical protein | 376 | gene | 442601 | 443368 | - | 1407,675 | 0,547448263 |
| 367 | RM25_RS01760 | RM25_0363 | Isoleucine--tRNA ligase | 377 | gene | 443689 | 446949 | + | 2166,28 | -0,312715527 |
| 368 | RM25_RS01765 | RM25_0364 | Trans-1, 2-dihydrobenzene-1, 2-diol dehydrogenase | 378 | gene | 447076 | 448062 | - | 968,6393 | -0,084357875 |
| 369 | RM25_RS01770 | RM25_0365 | Aspartate kinase | 379 | gene | 448231 | 449505 | + | 2886,067 | 0,143866496 |
| 370 | RM25_RS01775 | RM25_0366 | Integral membrane protein MviN | 380 | gene | 449592 | 451535 | + | 1319,847 | -0,038652581 |
| 371 | RM25_RS01780 | RM25_0367 | Integral membrane lysyl-tRNA synthetase | 381 | gene | 451599 | 453674 | + | 1226,975 | -0,413346799 |
| 372 | RM25_RS01785 | RM25_0368 | Hypothetical protein | 382 | gene | 453671 | 454069 | - | 293,4035 | 0,126497038 |
| 373 | RM25_RS01790 | RM25_0369 | Fructokinase | 383 | gene | 454223 | 455188 | - | 478,6738 | -0,206933433 |
| 374 | RM25_RS01795 | RM25_0370 | Agmatine/putrescine antiporter | 384 | gene | 455371 | 456909 | + | 1032,036 | -0,494194954 |
| 375 | RM25_RS01800 | RM25_0371 | MFS permease | 385 | gene | 457107 | 458858 | - | 1465,695 | -0,094267947 |
| 376 | RM25_RS01805 | RM25_0372 | Hypothetical protein | 386 | gene | 459158 | 459850 | + | 1015,784 | -0,210879407 |
| 377 | RM25_RS01810 | RM25_0373 | Hypothetical protein | 387 | gene | 459909 | 460673 | + | 460,7604 | -0,395415094 |
| 378 | RM25_RS01815 | RM25_0374 | Hypothetical protein | 388 | gene | 460735 | 461310 | + | 413,6476 | -0,424738394 |
| 379 | RM25_RS01820 | RM25_0375 | YidE/YbjL family protein | 389 | gene | 461835 | 463403 | - | 1179,404 | 0,088742488 |
| 380 | RM25_RS01825 | RM25_0376 | Phosphoribosylamine--glycine ligase | 390 | gene | 463664 | 464917 | + | 1164,445 | 0,460000476 |
| 381 | RM25_RS01830 | RM25_0377 | Adenylosuccinate lyase | 391 | gene | 464946 | 466379 | + | 2642,85 | 0,77302065 |
| 382 | RM25_RS01835 | RM25_0378 | Hypothetical protein | 392 | gene | 466389 | 467495 | - | 859,8333 | -0,178147982 |
| 383 | RM25_RS01840 | RM25_0379 | Phosphoribosylaminoimidazolesuccinocarboxamide synthase | 393 | gene | 467593 | 468471 | + | 1949,876 | 0,301752432 |
| 384 | RM25_RS01845 | RM25_0380 | Phosphoribosylformylglycinamidine synthetase PurS | 394 | gene | 468554 | 468799 | + | 1395,142 | 0,711120116 |
| 385 | RM25_RS01850 | RM25_0381 | Phosphoribosylformylglycinamidine synthase 1 | 395 | gene | 468801 | 469478 | + | 990,1452 | 0,689759932 |
| 386 | RM25_RS01855 | RM25_0382 | Hypothetical protein | 396 | gene | 469557 | 470255 | - | 320,6527 | -0,178418742 |
| 387 | RM25_RS01860 | RM25_0383 | Prophage Lp2 protein 6 | 397 | gene | 470434 | 471531 | - | 6482,416 | -0,452866984 |
| 388 | RM25_RS01865 | RM25_0384 | Polyol permease | 398 | gene | 471996 | 473279 | + | 821,5138 | -0,518070473 |
| 389 | RM25_RS01870 | RM25_0385 | Hypothetical protein | 399 | gene | 473365 | 474300 | - | 581,9719 | -0,590104318 |
| 390 | RM25_RS01875 | RM25_0386 | Hypothetical protein | 400 | gene | 474728 | 475537 | + | 434,2939 | 0,321297605 |
| 391 | RM25_RS01880 | RM25_0387 | Hypothetical protein | 401 | gene | 475982 | 476212 | + | 1014,662 | 0,266613637 |
| 392 | RM25_RS12530 | RM25_0388 | Hypothetical protein | 402 | gene | 476253 | 476771 | - | 427,9401 | -1,73690638 |
| 393 | RM25_RS01885 | RM25_0389 | Transposase insI for insertion sequence element IS30B/C/D | 403 | gene | 476866 | 477900 | + | 20,58407 | -0,822685867 |
| 395 | RM25_RS11720 | RM25_0390 | Hypothetical protein | 407 | gene | 480758 | 481456 | + | 311,2311 | -0,319763406 |
| 396 | RM25_RS11725 | RM25_0391 | Cutinase | 408 | gene | 481978 | 483216 | - | 1092,617 | -0,137837349 |
| 397 | RM25_RS01910 | RM25_0392 | Putative kinase inhibitor protein | 409 | gene | 483362 | 483826 | - | 2455,591 | 0,21431014 |
| 398 | RM25_RS01915 | RM25_0393 | DEAD/DEAH box helicase | 410 | gene | 483860 | 486412 | - | 1576,73 | -0,458932391 |
| 399 | RM25_RS11730 | RM25_0394 | 2-dehydro-3-deoxyphosphogluconate aldolase/4-hydroxy-2- | 411 | gene | 486549 | 487247 | + | 571,8841 | -0,304846308 |
| 400 | RM25_RS11735 | RM25_0395 | Hypothetical protein | 412 | gene | 487473 | 488477 | + | 4484,52 | 0,747661401 |
| 401 | RM25_RS01930 | RM25_0396 | Glucose-6-phosphate isomerase | 413 | gene | 488669 | 490351 | + | 3079,025 | -0,244167949 |
| 402 | RM25_RS01935 | RM25_0397 | OmpA family membrane associated protein | 414 | gene | 490355 | 491140 | + | 253,3143 | -0,048617058 |
| 403 | RM25_RS01940 | RM25_0398 | UPF0214 protein YfeW | 415 | gene | 491137 | 492360 | + | 536,2424 | -0,656344311 |
| 404 | RM25_RS01945 | RM25_0399 | putative D-lactate dehydrogenase Fe-S protein FAD/FMN-cc | 416 | gene | 492755 | 495541 | + | 5652,224 | -0,824132771 |
| 405 | RM25_RS01950 | RM25_0400 | Acyl-CoA dehydrogenase, short-chain specific | 417 | gene | 495700 | 496848 | - | 585,4444 | -1,034049332 |
| 406 | RM25_RS01955 | RM25_0401 | Carboxylic ester hydrolase | 418 | gene | 497234 | 498565 | + | 408,7273 | -1,243925327 |
| 407 | RM25_RS01960 | RM25_0402 | Hypothetical protein | 419 | gene | 498562 | 499062 | + | 111,4753 | -1,052840206 |
| 408 | RM25_RS01965 | RM25_0403 | ATP-dependent dsDNA exonuclease SbcC | 420 | gene | 499087 | 502251 | - | 1311,771 | -0,044462204 |
| 409 | RM25_RS01970 | RM25_0404 | Nuclease SbcCD, D subunit | 421 | gene | 502251 | 503531 | - | 757,8805 | 0,144430121 |
| 410 | RM25_RS01975 | RM25_0405 | Adenosine deaminase | 422 | gene | 503482 | 504699 | - | 491,4012 | 0,012806413 |
| 411 | RM25_RS11745 | RM25_0406 | Hypothetical protein | 423 | gene | 505303 | 506553 | + | 521,8796 | -0,428558416 |
| 412 | RM25_RS01985 | RM25_0407 | Hypothetical protein | 424 | gene | 506791 | 507162 | + | 116,2356 | -1,137875328 |
| 413 | RM25_RS01990 | RM25_0408 | Hypothetical protein | 425 | gene | 507177 | 507836 | - | 219,2945 | -1,68119088 |
| 414 | RM25_RS11750 | RM25_0409 | Hypothetical protein | 426 | gene | 508085 | 508525 | - | 239,9387 | -1,794800453 |
| 415 | RM25_RS02000 | RM25_0410 | Pyruvate oxidase | 427 | gene | 508910 | 510667 | - | 999,4914 | -0,691900204 |
| 416 | RM25_RS02005 | RM25_0411 | Putative solute/hydrogen antiporter | 428 | gene | 510807 | 512663 | - | 1136,728 | -0,482440805 |
| 417 | RM25_RS02010 | RM25_0412 | Hypothetical protein | 429 | gene | 512781 | 513155 | + | 308,0579 | -0,768656297 |
| 418 | RM25_RS02015 | RM25_0413 | Transcriptional regulator, MarR family | 430 | gene | 513339 | 513797 | - | 308,0528 | -0,04364924 |
| 419 | RM25_RS02020 | RM25_0414 | Peptidase M50B-like protein | 431 | gene | 514193 | 514939 | + | 262,0212 | 0,063837777 |
| 420 | RM25_RS02025 | RM25_0415 | Low affinity inorganic phosphate transporter membrane pro | 432 | gene | 515151 | 516422 | + | 2400,6 | 1,100068662 |
| 421 | RM25_RS02030 | RM25_0416 | Hypothetical protein | 433 | gene | 516419 | 516649 | + | 96,20084 | 0,282939805 |
| 422 | RM25_RS02035 | RM25_0417 | Azoreductase B | 434 | gene | 516905 | 517516 | + | 607,68 | 0,502340824 |
| 423 | RM25_RS02040 | RM25_0418 | Azoreductase B | 435 | gene | 517522 | 518133 | + | 342,8093 | 0,219555612 |
| 424 | RM25_RS02045 | RM25_0419 | Uncharacterized conserved protein UCP007542 (Precursor) | 436 | gene | 518187 | 520046 | - | 1246,847 | -0,078429976 |
| 425 | RM25_RS02050 | RM25_0420 | ATP-dependent helicase HrpB | 437 | gene | 520136 | 522775 | - | 960,4145 | -0,470030489 |
| 426 | RM25_RS02055 | RM25_0421 | Multifunctional siroheme synthase cysG | 438 | gene | 522814 | 523587 | - | 985,8074 | 0,388641298 |
| 427 | RM25_RS02060 | RM25_0422 | Cobalt ABC transporter, ATPase subunit | 439 | gene | 523584 | 524477 | - | 994,1502 | 0,07301443 |
| 428 | RM25_RS02065 | RM25_0423 | Cobalt ABC transporter, permease protein CbiQ | 440 | gene | 524474 | 525268 | - | 762,2411 | 0,31419015 |
| 429 | RM25_RS02070 | RM25_0424 | Cobalt transport protein CbiN | 441 | gene | 525272 | 525772 | - | 2190,699 | 0,37704165 |
| 430 | RM25_RS02075 | RM25_0425 | Cobalt transport protein CbiM | 442 | gene | 525765 | 526472 | - | 4654,544 | 0,252141555 |
| 431 | RM25_RS02080 | RM25_0426 | Putative GntR-famly transcriptional regulator | 443 | gene | 526782 | 527546 | - | 942,4064 | -1,022351292 |
| 432 | NA | RM25_0427 | Hypothetical protein | 444 | gene | 527751 | 527879 | + | 688,9172 | -3,392695887 |

| | | | | | | | | | | | | | | | | | | | | | | | | | | | | | | | | | | | | | | |
|---|---|---|---|---|---|---|---|---|---|---|---|---|---|---|---|---|---|---|---|---|---|---|---|---|---|---|---|---|---|---|---|---|---|---|---|---|---|---|
| 433 | RM25_RS02085 | RM25_0428 | Gluconate permease | 445 | gene | 527895 | 529238 | + | 2917,51 | -3,177629486 | 0,705116 | -4,50653 | 6,59E-06 | 0,000202 | 568,7672445 | -0,808827222 | 0,192498 | -4,20175 | 2,65E-05 | 0,000104 | 5266,254 | -3,710442357 | 0,411883 | -9,00849 | 2,09E-19 | 3,28E-17 | 5493,798 | 3,706386 | 0,433013 | 8,559529 | 1,13E-17 | 7,03E-16 | 568,7918165 | 0,845354595 | 0,146466 | 5,77166 | 7,85E-09 | 3,8E-08 |
| 434 | RM25_RS02090 | RM25_0429 | Phosphogluconate dehydrogenase (decarboxylating) | 446 | gene | 529365 | 530255 | + | 1838,264 | -1,592989824 | 0,508779 | -3,131 | 0,001742 | 0,013919 | 1010,417804 | 0,029236554 | 0,187203 | 0,156175 | 0,875895 | 0,904286 | 2666,111 | -2,479372015 | 0,493988 | -5,0191 | 5,19E-07 | 7,79E-06 | 2898,405 | 2,128947 | 0,507729 | 4,193079 | 2,75E-05 | 0,000153 | 896,7653975 | -0,342865662 | 0,128239 | -2,67364 | 0,007503 | 0,013438 |
| 435 | RM25_RS02095 | RM25_0430 | Chaperone protein DnaK | 447 | gene | 530663 | 532537 | + | 8277,229 | 0,332633783 | 0,496509 | 0,669945 | 0,502893 | 0,600097 | 11843,25535 | 0,229033027 | 0,24482 | 0,935517 | 0,349522 | 0,418609 | 4711,203 | 0,597953266 | 0,530225 | 1,127735 | 0,259432 | 0,360543 | 7830,402 | -1,58898 | 0,531829 | -2,98777 | 0,00281 | 0,008401 | 9040,437971 | -1,187948346 | 0,244604 | -4,85661 | 1,19E-06 | 4,1E-06 |
| 436 | RM25_RS02100 | RM25_0431 | Protein GrpE | 448 | gene | 532702 | 533352 | + | 1212,62 | 0,633964991 | 0,423235 | 1,497903 | 0,134158 | 0,233022 | 1697,964668 | 0,663535166 | 0,214248 | 3,097047 | 0,001955 | 0,004818 | 727,2763 | 0,568856309 | 0,404575 | 1,406058 | 0,159707 | 0,247164 | 1013,177 | -1,21941 | 0,418125 | -2,91638 | 0,003541 | 0,010068 | 1445,962916 | -1,273953962 | 0,179405 | -7,10099 | 1,24E-12 | 1,07E-11 |
| 437 | RM25_RS02105 | RM25_0432 | Chaperone protein DnaJ | 449 | gene | 533414 | 534586 | + | 2715,71 | 0,658836292 | 0,408865 | 1,611378 | 0,107097 | 0,200978 | 3782,651494 | 0,659777402 | 0,190977 | 3,454739 | 0,000551 | 0,001576 | 1648,769 | 0,658277148 | 0,37554 | 1,752882 | 0,079622 | 0,144807 | 2245,353 | -1,25146 | 0,37859 | -3,30559 | 0,000948 | 0,003364 | 3260,192749 | -1,211535663 | 0,176643 | -6,85867 | 6,95E-12 | 5,27E-11 |
| 438 | RM25_RS02110 | RM25_0433 | Transcriptional regulator HspR | 450 | gene | 534679 | 535113 | + | 718,4484 | 0,524979279 | 0,295437 | 1,776961 | 0,075575 | 0,162527 | 932,9141564 | 0,642704303 | 0,155942 | 4,121429 | 3,77E-05 | 0,000142 | 503,9827 | 0,314235768 | 0,218627 | 1,437315 | 0,150628 | 0,235588 | 624,8075 | -0,75626 | 0,198247 | -3,81472 | 0,000136 | 0,000643 | 831,4724622 | -1,035329611 | 0,155343 | -6,6648 | 2,65E-11 | 1,86E-10 |
| 439 | RM25_RS02115 | RM25_0434 | BadF/BadG/BcrA/BcrD ATPase family protein | 451 | gene | 535272 | 536228 | - | 936,4691 | -0,276471563 | 0,200168 | -1,3812 | 0,167219 | 0,269586 | 1010,169108 | 0,109616769 | 0,162309 | 0,675357 | 0,499449 | 0,565147 | 862,7691 | -0,73801503 | 0,216537 | -3,40826 | 0,000654 | 0,003455 | 1082,615 | 0,099094 | 0,215014 | 0,460872 | 0,64489 | 0,700464 | 831,7277693 | -0,700749646 | 0,169895 | -4,12461 | 3,71E-05 | 0,000101 |
| 440 | RM25_RS02120 | RM25_0435 | Phosphoribosylformylglycinamidine synthase 2 | 452 | gene | 536368 | 538923 | + | 3697,988 | 0,497584343 | 0,157099 | 3,167326 | 0,001538 | 0,012859 | 4077,76243 | 0,3719059 | 0,161625 | 2,301035 | 0,02139 | 0,037825 | 3318,213 | 0,654836682 | 0,181621 | 3,605507 | 0,000312 | 0,001912 | 3252,443 | -0,52064 | 0,203814 | -2,55448 | 0,010635 | 0,024867 | 4240,083976 | -0,191975087 | 0,125415 | -1,53072 | 0,125838 | 0,169582 |
| 441 | RM25_RS02125 | RM25_0436 | Peptidase dimerization domain protein | 453 | gene | 538928 | 540274 | + | 2453,255 | 0,71740867 | 0,281301 | 2,550326 | 0,010762 | 0,04913 | 2954,778646 | 0,435349796 | 0,193178 | 2,253619 | 0,02422 | 0,042193 | 1951,731 | 1,174795981 | 0,225263 | 5,215229 | 1,84E-07 | 3,07E-06 | 1972,768 | -1,12174 | 0,236014 | -4,75284 | 2,01E-06 | 1,44E-05 | 2986,22771 | -0,34227652 | 0,162719 | -2,10348 | 0,035423 | 0,054393 |
| 442 | RM25_RS02130 | RM25_0437 | Hypothetical protein | 454 | gene | 540274 | 540750 | + | 289,5817 | 0,252246159 | 0,199974 | 1,261394 | 0,207167 | 0,311808 | 283,7392621 | 0,202498746 | 0,206884 | 0,978804 | 0,327677 | 0,395056 | 295,4242 | 0,302573036 | 0,351517 | 0,860765 | 0,389368 | 0,49554 | 278,1628 | -0,05738 | 0,291945 | -0,19655 | 0,844176 | 0,877303 | 309,3623255 | 0,108183132 | 0,277648 | 0,389641 | 0,696802 | 0,745964 |
| 443 | RM25_RS02135 | RM25_0438 | Formate--tetrahydrofolate ligase | 455 | gene | 540881 | 542557 | + | 4715,07 | -0,024457511 | 0,260826 | -0,09377 | 0,925292 | 0,949875 | 3940,543433 | 0,285141663 | 0,15216 | 1,873954 | 0,060937 | 0,092484 | 5489,597 | -0,246176745 | 0,365166 | -0,67415 | 0,500216 | 0,600004 | 5014,781 | 0,693618 | 0,334078 | 2,076214 | 0,037874 | 0,068041 | 4568,22612 | 0,198577291 | 0,205714 | 0,965309 | 0,33439 | 0,398059 |
| 444 | RM25_RS11755 | RM25_0439 | Kinase | 456 | gene | 542659 | 543219 | - | 623,6994 | 0,300464609 | 0,272324 | 1,103336 | 0,269881 | 0,377962 | 787,3454359 | 0,384765491 | 0,145946 | 2,636358 | 0,00838 | 0,017066 | 460,0533 | 0,159870904 | 0,248839 | 0,642467 | 0,52057 | 0,620629 | 592,1926 | -0,71556 | 0,218621 | -3,27307 | 0,001064 | 0,003709 | 675,841218 | -0,880410558 | 0,165887 | -5,30731 | 1,11E-07 | 4,52E-07 |
| 445 | RM25_RS02145 | RM25_0440 | UPF0210 protein | 457 | gene | 543278 | 544627 | - | 1813,272 | 0,415314747 | 0,251887 | 1,648815 | 0,099186 | 0,192093 | 2255,870534 | 0,49208177 | 0,179281 | 2,744756 | 0,006056 | 0,012964 | 1370,674 | 0,291765493 | 0,173097 | 1,685561 | 0,09188 | 0,161373 | 1648,185 | -0,66132 | 0,20884 | -3,16662 | 0,001542 | 0,005063 | 2031,694173 | -0,816159694 | 0,131679 | -6,19809 | 5,72E-10 | 3,31E-09 |
| 446 | RM25_RS02150 | RM25_0441 | UPF0237 protein | 458 | gene | 544631 | 544900 | - | 655,5754 | 0,146428881 | 0,170076 | 0,860961 | 0,389259 | 0,500264 | 727,105302 | 0,36823483 | 0,154705 | 2,380233 | 0,017302 | 0,031711 | 584,0454 | -0,123654928 | 0,216044 | -0,57236 | 0,567078 | 0,661358 | 658,6518 | -0,12084 | 0,190166 | -0,63546 | 0,525131 | 0,5925 | 674,7197265 | -0,564837369 | 0,173825 | -3,24946 | 0,001156 | 0,002403 |
| 447 | RM25_RS11760 | RM25_0442 | Hypothetical protein | 459 | gene | 545170 | 546009 | + | 992,2187 | 0,475586898 | 0,211404 | 2,249657 | 0,024471 | 0,079665 | 1153,122958 | 0,347799341 | 0,196616 | 1,768931 | 0,076905 | 0,113202 | 831,3144 | 0,657337768 | 0,199039 | 3,30256 | 0,000958 | 0,004691 | 880,2911 | -0,70838 | 0,243047 | -2,91459 | 0,003562 | 0,010102 | 1131,124414 | -0,35297243 | 0,13146 | -2,68501 | 0,007253 | 0,013019 |
| 448 | RM25_RS02160 | RM25_0443 | Amidophosphoribosyltransferase | 460 | gene | 546110 | 547651 | + | 3135,898 | 0,932205259 | 0,212487 | 4,387119 | 1,15E-05 | 0,00033 | 3462,046318 | 0,785596896 | 0,154074 | 5,098838 | 3,42E-07 | 1,86E-06 | 2809,75 | 1,120250253 | 0,320331 | 3,497162 | 0,00047 | 0,002669 | 2290,06 | -0,5761 | 0,263704 | -2,18466 | 0,028913 | 0,05487 | 4022,080429 | -0,204501119 | 0,217149 | -0,94176 | 0,346317 | 0,40998 |
| 449 | RM25_RS02165 | RM25_0444 | Phosphoribosylformylglycinamidine cyclo-ligase | 461 | gene | 547648 | 548709 | + | 1580,495 | 0,934494916 | 0,215955 | 4,327265 | 1,51E-05 | 0,000412 | 1795,46153 | 0,703059666 | 0,168489 | 4,172722 | 3,01E-05 | 0,000116 | 1365,529 | 1,258545751 | 0,214375 | 5,870762 | 4,34E-09 | 1,09E-07 | 1152,705 | -0,8218 | 0,185157 | -4,43841 | 9,06E-06 | 5,62E-05 | 2030,938995 | -0,220882628 | 0,169192 | -1,30552 | 0,191717 | 0,245451 |
| 450 | RM25_RS02170 | RM25_0445 | Hypothetical protein | 462 | gene | 548943 | 549173 | + | 7555,933 | 0,797452959 | 0,522868 | 1,52515 | 0,127222 | 0,225121 | 10149,8168 | 0,283411273 | 0,230007 | 1,232183 | 0,217881 | 0,279861 | 4962,049 | 2,095640997 | 0,459323 | 4,562456 | 5,06E-06 | 5,57E-05 | 5922,931 | -2,33142 | 0,460265 | -5,06539 | 4,08E-07 | 3,31E-06 | 9385,023634 | -0,486235931 | 0,235784 | -2,06221 | 0,039188 | 0,059629 |
| 451 | RM25_RS02175 | RM25_0446 | Domain family protein, hydrolase, alpha/beta fold family | 463 | gene | 549425 | 550507 | + | 951,089 | -0,209188557 | 0,133436 | -1,5677 | 0,116951 | 0,212454 | 874,2161909 | -0,263766353 | 0,177432 | -1,48657 | 0,137127 | 0,188696 | 1027,962 | -0,163745391 | 0,175381 | -0,93365 | 0,350483 | 0,456655 | 1075,912 | 0,130418 | 0,173653 | 0,751026 | 0,452637 | 0,526847 | 864,4383209 | 0,28057038 | 0,174339 | 1,60934 | 0,107542 | 0,147199 |
| 455 | RM25_RS02195 | RM25_0447 | Cell-wall peptidases, NlpC/P60 family secreted protein | 473 | gene | 551145 | 552863 | - | 6472,018 | 0,211297258 | 0,394568 | 0,535515 | 0,592294 | 0,682669 | 8587,700728 | 0,064338789 | 0,180793 | 0,35587 | 0,721938 | 0,771704 | 4356,336 | 0,505570188 | 0,455711 | 1,109409 | 0,267254 | 0,369236 | 6385,306 | -1,27056 | 0,440532 | -2,88415 | 0,003925 | 0,01087 | 6800,293992 | -0,795706292 | 0,199486 | -3,98879 | 6,64E-05 | 0,000174 |
| 456 | RM25_RS02200 | RM25_0448 | ABC-2 type transporter | 474 | gene | 553301 | 554107 | - | 554,3713 | -1,917157373 | 0,856244 | -2,23903 | 0,025154 | 0,081105 | 319,5818056 | -5,434330025 | 0,48953 | -11,1011 | 1,24E-28 | 1,04E-26 | 789,1608 | -1,3268903 | 0,360469 | -3,68101 | 0,000232 | 0,001484 | 919,1247 | 0,783828 | 0,436769 | 1,794606 | 0,072717 | 0,114798 | 228,6389989 | 4,962580165 | 0,478786 | 10,36493 | 3,58E-25 | 1,38E-23 |
| 457 | RM25_RS02205 | RM25_0449 | Daunorubicin resistance ABC transporter ATPase subunit | 475 | gene | 554104 | 555132 | - | 364,4457 | -1,248345459 | 0,848533 | -1,47118 | 0,141242 | 0,241763 | 152,4190922 | -4,163024592 | 0,458636 | -9,07697 | 1,12E-19 | 3,21E-18 | 576,4724 | -0,827811716 | 0,45231 | -1,83019 | 0,067222 | 0,127262 | 536,0218 | 1,28492 | 0,360169 | 3,567545 | 0,00036 | 0,001503 | 213,1957252 | 4,6888362 | 0,555702 | 8,437683 | 3,24E-17 | 5,19E-16 |
| 458 | RM25_RS02210 | RM25_0450 | Transcription regulator, TetR | 476 | gene | 555301 | 556068 | + | 710,4651 | -0,055791547 | 0,155727 | -0,35826 | 0,720145 | 0,794283 | 636,7995094 | 0,216519325 | 0,134818 | 1,606011 | 0,108271 | 0,152447 | 784,1306 | -0,275659682 | 0,191184 | -1,44186 | 0,149343 | 0,234511 | 763,5758 | 0,485342 | 0,197261 | 2,460401 | 0,013878 | 0,030743 | 682,808613 | 0,042523409 | 0,131383 | 0,323659 | 0,746196 | 0,790465 |
| 460 | RM25_RS02220 | RM25_0452 | HTH-type transcriptional regulator CmtR | 480 | gene | 556622 | 557035 | + | 1479,094 | 1,998072609 | 0,251893 | 7,932241 | 2,15E-15 | 7,25E-13 | 1457,797121 | 1,688802705 | 0,14635 | 11,53944 | 8,35E-31 | 1,09E-28 | 1500,391 | 2,343868533 | 0,411884 | 5,690609 | 1,27E-08 | 2,63E-07 | 627,2734 | -0,54396 | 0,204148 | -2,66455 | 0,007709 | 0,019007 | 2308,718048 | 0,152343288 | 0,334777 | 0,455059 | 0,649067 | 0,704351 |
| 462 | RM25_RS02225 | RM25_0453 | Putative cation-transporting P-type ATPase | 481 | gene | 557032 | 558903 | + | 2641,955 | 1,725565438 | 0,172796 | 9,986149 | 1,75E-23 | 2,07E-20 | 2693,494468 | 1,785393278 | 0,152144 | 11,73493 | 8,44E-32 | 1,17E-29 | 2590,416 | 1,665477032 | 0,320366 | 5,198664 | 2,01E-07 | 3,33E-06 | 1293,561 | -0,02717 | 0,185414 | -0,14652 | 0,883507 | 0,907375 | 3969,579084 | -0,098301007 | 0,255977 | -0,38402 | 0,700961 | 0,749621 |
| 463 | RM25_RS02230 | RM25_0454 | Aspartate/tyrosine/aromatic aminotransferase | 482 | gene | 559041 | 560279 | - | 588,548 | -0,935296316 | 0,463954 | -2,01592 | 0,043808 | 0,114094 | 355,8290379 | -1,626055262 | 0,258014 | -6,30221 | 2,93E-10 | 2,66E-09 | 821,2669 | -0,67037686 | 0,336366 | -1,993 | 0,046262 | 0,095564 | 810,5613 | 0,839517 | 0,297673 | 2,820263 | 0,004798 | 0,012736 | 397,6387458 | 1,864529006 | 0,314631 | 5,926086 | 3,1E-09 | 1,6E-08 |
| 464 | RM25_RS02235 | RM25_0455 | Putative insertion element ISCmi2 transposase | 483 | gene | 560308 | 561309 | + | 140,5828 | 0,29796968 | 0,256731 | 1,160628 | 0,245793 | 0,354792 | 144,6028977 | -0,249407382 | 0,28687 | -0,86941 | 0,384623 | 0,453732 | 136,5627 | 0,89675039 | 0,265074 | 3,383025 | 0,000717 | 0,0037 | 134,7087 | -0,78411 | 0,30166 | -2,59932 | 0,009341 | 0,022466 | 151,3297783 | 0,41528977 | 0,264393 | 1,570728 | 0,116246 | 0,15792 |
| 465 | RM25_RS13125 | RM25_0456 | Hypothetical protein | 484 | gene | 561369 | 561863 | - | 206,8116 | -1,16940728 | 0,407585 | -2,86911 | 0,004116 | 0,025941 | 136,2774256 | -1,457856253 | 0,312335 | -4,6676 | 3,05E-06 | 1,42E-05 | 277,3457 | -1,040991641 | 0,381581 | -2,7281 | 0,00637 | 0,021771 | 300,3691 | 0,829305 | 0,335423 | 2,472412 | 0,01342 | 0,029965 | 125,1589684 | 1,317468749 | 0,391114 | 3,368505 | 0,000756 | 0,001651 |
| 466 | RM25_RS13130 | RM25_0457 | Hypothetical protein | 485 | gene | 561805 | 562335 | - | 160,1787 | -0,019204618 | 0,310903 | -0,06177 | 0,950746 | 0,967577 | 120,0950962 | 0,131134145 | 0,264738 | 0,495335 | 0,620364 | 0,677885 | 200,2623 | -0,107649628 | 0,355367 | -0,30293 | 0,761947 | 0,826085 | 168,602 | 0,803149 | 0,264209 | 3,039825 | 0,002367 | 0,007284 | 156,4861278 | 0,624422393 | 0,350305 | 1,782508 | 0,074666 | 0,105826 |
| 467 | RM25_RS02245 | RM25_0458 | Dehydrogenase | 486 | gene | 562332 | 563357 | - | 1263,915 | 0,039314709 | 0,212772 | 0,184774 | 0,853407 | 0,904036 | 1071,730603 | 0,399795409 | 0,124659 | 3,207121 | 0,001341 | 0,003461 | 1456,099 | -0,223240567 | 0,246997 | -0,90382 | 0,366092 | 0,472857 | 1312,524 | 0,708516 | 0,245472 | 2,886336 | 0,003898 | 0,01082 | 1254,687339 | 0,130037209 | 0,11257 | 1,15517 | 0,248021 | 0,306066 |
| 468 | RM25_RS02250 | RM25_0459 | Hypothetical protein | 487 | gene | 563674 | 564027 | - | 183,7159 | -0,36913677 | 0,224725 | -1,64262 | 0,100462 | 0,193298 | 161,6533366 | 0,046728112 | 0,201811 | 0,231544 | 0,816892 | 0,85498 | 205,7784 | -0,70036027 | 0,308374 | -2,27114 | 0,023139 | 0,057227 | 217,5871 | 0,621341 | 0,175001 | 3,550495 | 0,000385 | 0,001582 | 157,6229554 | -0,066806212 | 0,328373 | -0,20345 | 0,838786 | 0,869784 |
| 469 | RM25_RS02255 | RM25_0460 | Transaldolase | 488 | gene | 564366 | 565448 | - | 3688,765 | 0,426241934 | 0,242083 | 1,760728 | 0,078285 | 0,165486 | 3944,130949 | 0,48199008 | 0,180665 | 2,667864 | 0,007634 | 0,015741 | 3433,4 | 0,363174814 | 0,411688 | 0,88216 | 0,37769 | 0,483025 | 3338,741 | -0,18012 | 0,38525 | -0,46755 | 0,640109 | 0,696554 | 4135,703496 | -0,267568102 | 0,228528 | -1,17083 | 0,241665 | 0,298897 |
| 470 | RM25_RS02260 | RM25_0461 | Hypothetical protein | 489 | gene | 565648 | 566307 | - | 1244,412 | -0,311920448 | 0,169495 | -1,84029 | 0,065725 | 0,147269 | 1117,992205 | -0,033457597 | 0,146737 | -0,22801 | 0,819639 | 0,856923 | 1370,831 | -0,541767277 | 0,231761 | -2,33762 | 0,019407 | 0,050363 | 1453,02 | 0,468359 | 0,231245 | 2,025384 | 0,042828 | 0,074941 | 1087,278686 | 0,00362756 | 0,134392 | 0,026992 | 0,978466 | 0,983054 |
| 471 | RM25_RS11765 | RM25_0462 | Hypothetical protein | 490 | gene | 566428 | 566775 | - | 467,894 | -0,188578498 | 0,138073 | -1,36579 | 0,172004 | 0,275792 | 447,0393333 | -0,326822317 | 0,16426 | -1,98967 | 0,046627 | 0,074915 | 488,7487 | -0,065030351 | 0,241057 | -0,26977 | 0,787336 | 0,845444 | 525,4451 | -0,05756 | 0,219679 | -0,26204 | 0,793294 | 0,832129 | 428,4577634 | 0,25733069 | 0,157641 | 1,632381 | 0,102599 | 0,141419 |
| 472 | RM25_RS02270 | RM25_0463 | Phospholipase | 491 | gene | 566897 | 568486 | - | 1822,036 | 0,564431032 | 0,129852 | 4,346739 | 1,38E-05 | 0,000384 | 1680,98942 | 0,492022616 | 0,135929 | 3,619694 | 0,000295 | 0,00091 | 1963,083 | 0,627259615 | 0,216662 | 2,895111 | 0,00379 | 0,014479 | 1552,625 | 0,087724 | 0,188402 | 0,465618 | 0,641489 | 0,697091 | 2126,688379 | 0,266522245 | 0,146921 | 1,814049 | 0,06967 | 0,099644 |
| 473 | RM25_RS02275 | RM25_0464 | Cation diffusion facilitator family transporter | 492 | gene | 568677 | 569645 | + | 893,5059 | 0,329674369 | 0,169119 | 1,949359 | 0,051253 | 0,12623 | 1020,274493 | 0,321253757 | 0,150896 | 2,128968 | 0,033257 | 0,05591 | 766,7373 | 0,342824118 | 0,191943 | 1,786068 | 0,074088 | 0,13632 | 840,0332 | -0,48284 | 0,195066 | -2,47529 | 0,013313 | 0,029884 | 974,6159545 | -0,414589436 | 0,133586 | -3,10355 | 0,001912 | 0,003807 |
| 474 | RM25_RS02280 | RM25_0465 | Cytidine and deoxycytidylate deaminase zinc-binding region | 493 | gene | 569674 | 570138 | + | 564,0416 | 0,66176713 | 0,187138 | 3,536246 | 0,000406 | 0,005143 | 513,1295101 | 0,710659941 | 0,221214 | 3,212548 | 0,001316 | 0,003415 | 614,9537 | 0,622766738 | 0,266554 | 2,336366 | 0,019472 | 0,050363 | 460,513 | 0,262429 | 0,28698 | 0,914449 | 0,360481 | 0,43684 | 676,1446503 | 0,216199887 | 0,177184 | 1,220202 | 0,222388 | 0,278369 |
| 475 | RM25_RS02285 | RM25_0466 | Hypothetical protein | 494 | gene | 570164 | 570562 | - | 801,441 | 0,489878439 | 0,118707 | 4,12678 | 3,68E-05 | 0,000774 | 792,5146295 | 0,504472219 | 0,176562 | 2,857189 | 0,004274 | 0,009504 | 810,3674 | 0,475151708 | 0,211037 | 2,251513 | 0,024353 | 0,059854 | 704,5963 | -0,0063 | 0,231537 | -0,02721 | 0,978292 | 0,982042 | 916,367612 | 0,010227446 | 0,1298 | 0,078794 | 0,937196 | 0,948871 |
| 476 | RM25_RS02290 | RM25_0467 | PimH protein | 495 | gene | 570725 | 572011 | - | 361,2331 | -1,370720876 | 0,478176 | -2,86656 | 0,00415 | 0,026082 | 230,2470198 | -1,803981359 | 0,361466 | -4,99073 | 6,02E-07 | 3,14E-06 | 492,2191 | -1,189026429 | 0,467599 | -2,54283 | 0,010996 | 0,032848 | 547,7608 | 0,860631 | 0,375476 | 2,292107 | 0,021899 | 0,043855 | 198,9487348 | 1,558265945 | 0,492141 | 3,166303 | 0,001544 | 0,003134 |
| 477 | RM25_RS12225 | RM25_0468 | Putative membrane protein | 496 | gene | 572348 | 572602 | - | 397,9645 | 0,097407528 | 0,128135 | 0,760192 | 0,44714 | 0,554286 | 377,3302845 | 0,161898451 | 0,216159 | 0,748978 | 0,45387 | 0,522095 | 418,5987 | 0,039046393 | 0,1583 | 0,246661 | 0,805171 | 0,860285 | 405,7311 | 0,155046 | 0,184208 | 0,841693 | 0,39996 | 0,475394 | 403,0256415 | 0,084120251 | 0,202023 | 0,41639 | 0,677125 | 0,728759 |
| 478 | RM25_RS02300 | RM25_0469 | 1, 4-dihydroxy-2-naphthoate octaprenyltransferase | 497 | gene | 572783 | 573664 | + | 513,1424 | -0,454578085 | 0,286255 | -1,58802 | 0,112282 | 0,206881 | 403,9424925 | -0,386286944 | 0,155495 | -2,48424 | 0,012983 | 0,024798 | 622,3423 | -0,500665248 | 0,396617 | -1,26234 | 0,206827 | 0,303544 | 622,5613 | 0,610232 | 0,207763 | 2,937162 | 0,003312 | 0,009603 | 426,4509528 | 0,562672775 | 0,363704 | 1,547063 | 0,121848 | 0,164771 |
| 479 | RM25_RS02305 | RM25_0470 | Nuclease, RecB family | 498 | gene | 573681 | 575450 | - | 1382,972 | 0,03052826 | 0,135657 | 0,22504 | 0,821949 | 0,876621 | 1504,07293 | 0,052897579 | 0,146004 | 0,362303 | 0,717126 | 0,767256 | 1261,872 | 0,005137569 | 0,208669 | 0,024621 | 0,980358 | 0,988324 | 1444,378 | -0,28922 | 0,186727 | -1,54887 | 0,121412 | 0,174068 | 1371,177862 | -0,281762606 | 0,140107 | -2,01106 | 0,044319 | 0,066578 |
| 480 | RM25_RS02310 | RM25_0471 | Demethylmenaquinone methyltransferase | 499 | gene | 575577 | 576224 | - | 1524,12 | 0,410856977 | 0,185589 | 2,213805 | 0,026842 | 0,084203 | 1725,291977 | 0,473672236 | 0,175346 | 2,701352 | 0,006906 | 0,01443 | 1322,948 | 0,330704336 | 0,245915 | 1,344793 | 0,178692 | 0,270853 | 1384,07 | -0,35412 | 0,208935 | -1,69489 | 0,090096 | 0,1363 | 1702,642504 | -0,459410475 | 0,186044 | -2,46936 | 0,013535 | 0,022985 |
| 481 | RM25_RS02315 | RM25_0472 | Heme ABC exporter, ATP-binding protein CcmA | 500 | gene | 576426 | 577118 | + | 2949,581 | 0,530902165 | 0,406166 | 1,307105 | 0,191177 | 0,295091 | 4089,997595 | 0,444426309 | 0,185892 | 2,390781 | 0,016813 | 0,031007 | 1809,164 | 0,731766442 | 0,352608 | 2,075296 | 0,037959 | 0,083073 | 2573,249 | -1,39965 | 0,338056 | -4,14027 | 3,47E-05 | 0,000188 | 3416,235841 | -1,076310267 | 0,204032 | -5,2752 | 1,33E-07 | 5,28E-07 |
| 482 | RM25_RS02320 | RM25_0473 | Hypothetical protein | 501 | gene | 577115 | 578920 | + | 2579,588 | 0,64410588 | 0,376087 | 1,712651 | 0,086777 | 0,176421 | 3592,05372 | 0,655722862 | 0,184633 | 3,551486 | 0,000383 | 0,001149 | 1567,122 | 0,618980764 | 0,185752 | 3,332291 | 0,000861 | 0,004329 | 2141,114 | -1,2329 | 0,226027 | -5,45466 | 4,91E-08 | 4,8E-07 | 3087,250163 | -1,218928495 | 0,128584 | -9,47963 | 2,55E-21 | 6,76E-20 |
| 483 | RM25_RS02325 | RM25_0474 | Isochorismate synthase dhbC | 502 | gene | 578966 | 580153 | - | 617,0548 | -0,689527341 | 0,397858 | -1,7331 | 0,083078 | 0,171018 | 403,1048448 | -1,288814713 | 0,225467 | -5,71619 | 1,09E-08 | 7,96E-08 | 831,0048 | -0,422904702 | 0,260455 | -1,62372 | 0,104437 | 0,177731 | 801,26 | 0,665561 | 0,248997 | 2,672969 | 0,007518 | 0,018634 | 463,6125384 | 1,596791737 | 0,254931 | 6,263624 | 3,76E-10 | 2,23E-09 |
| 484 | RM25_RS02330 | RM25_0475 | Geranylgeranyl reductase family protein | 503 | gene | 580551 | 581882 | + | 3200,078 | 0,464461972 | 0,453067 | 1,02515 | 0,305292 | 0,41474 | 4708,612956 | 0,638710508 | 0,17287 | 3,694752 | 0,00022 | 0,000698 | 1691,543 | -0,005285849 | 0,293293 | -0,01802 | 0,985621 | 0,991934 | 2861,312 | -1,17268 | 0,296751 | -3,95174 | 7,76E-05 | 0,000388 | 3640,393103 | -1,777325088 | 0,141768 | -12,5368 | 4,69E-36 | 4,1E-34 |
| 485 | RM25_RS02335 | RM25_0476 | NADH-quinone oxidoreductase subunit A | 504 | gene | 581939 | 582304 | + | 2312,831 | 0,431849921 | 0,578249 | 0,746824 | 0,45517 | 0,560227 | 3575,845836 | 0,260747482 | 0,216523 | 1,204247 | 0,228494 | 0,291955 | 1049,817 | 1,048657797 | 0,467801 | 2,241675 | 0,024982 | 0,060862 | 2110,424 | -2,301 | 0,414472 | -5,55164 | 2,83E-08 | 2,91E-07 | 2602,235978 | -1,481247647 | 0,299943 | -4,93843 | 7,88E-07 | 2,81E-06 |
| 486 | RM25_RS02340 | RM25_0477 | NADH-quinone oxidoreductase subunit B | 505 | gene | 582323 | 582880 | + | 2484,006 | 0,453360856 | 0,512849 | 0,884004 | 0,376694 | 0,487981 | 3671,483394 | 0,290469727 | 0,202718 | 1,432879 | 0,151892 | 0,205872 | 1296,528 | 0,936454446 | 0,447046 | 2,094762 | 0,036192 | 0,079874 | 2241,223 | -1,94264 | 0,415085 | -4,68011 | 2,87E-06 | 2,01E-05 | 2813,052789 | -1,263075493 | 0,254836 | -4,95643 | 7,18E-07 | 2,58E-06 |
| 487 | RM25_RS02345 | RM25_0478 | NADH-quinone oxidoreductase subunit C | 506 | gene | 582877 | 583644 | + | 2719,589 | 0,316717969 | 0,468407 | 0,676159 | 0,49894 | 0,596955 | 3888,319165 | 0,10758148 | 0,192236 | 0,559633 | 0,57573 | 0,637387 | 1550,859 | 0,863046098 | 0,410634 | 2,10174 | 0,035576 | 0,078883 | 2587,902 | -1,81958 | 0,374529 | -4,85832 | 1,18E-06 | 8,91E-06 | 2953,582024 | -1,029034641 | 0,241593 | -4,25937 | 2,05E-05 | 5,79E-05 |
| 488 | RM25_RS02350 | RM25_0479 | NADH-quinone oxidoreductase subunit D | 507 | gene | 583641 | 585014 | + | 5708,597 | 0,451057251 | 0,564758 | 0,798673 | 0,42448 | 0,533315 | 8675,405149 | 0,273962991 | 0,208007 | 1,317086 | 0,18781 | 0,246337 | 2741,788 | 1,042933875 | 0,504586 | 2,066909 | 0,038743 | 0,084163 | 5158,109 | -2,18193 | 0,480297 | -4,54288 | 5,55E-06 | 3,63E-05 | 6460,403875 | -1,380532811 | 0,251605 | -5,4869 | 4,09E-08 | 1,81E-07 |
| 489 | RM25_RS02355 | RM25_0480 | NADH dehydrogenase subunit E | 508 | gene | 585070 | 585801 | + | 2362,705 | 0,558691815 | 0,571365 | 0,977819 | 0,328164 | 0,438482 | 3641,68313 | 0,417445169 | 0,207673 | 2,010106 | 0,04442 | 0,071908 | 1083,727 | 1,060683203 | 0,473701 | 2,23914 | 0,025147 | 0,061104 | 2043,232 | -2,20176 | 0,464023 | -4,74493 | 2,09E-06 | 1,49E-05 | 2759,495557 | -1,522524759 | 0,217754 | -6,99196 | 2,71E-12 | 2,19E-11 |
| 490 | RM25_RS02360 | RM25_0481 | NADH-quinone oxidoreductase, F subunit | 509 | gene | 585798 | 587138 | + | 4347,593 | 0,423486796 | 0,556044 | 0,761607 | 0,446295 | 0,553932 | 6672,941391 | 0,287368102 | 0,203771 | 1,410253 | 0,158465 | 0,213065 | 2022,245 | 0,891319563 | 0,450466 | 1,97866 | 0,047854 | 0,097604 | 3970,765 | -2,13803 | 0,424928 | -5,03152 | 4,87E-07 | 3,9E-06 | 4881,749739 | -1,498046733 | 0,239367 | -6,25836 | 3,89E-10 | 2,29E-09 |
| 491 | RM25_RS02365 | RM25_0482 | NADH dehydrogenase, G subunit | 510 | gene | 587135 | 589546 | + | 6721,537 | 0,445524723 | 0,540887 | 0,823693 | 0,410114 | 0,520538 | 10122,78539 | 0,365588297 | 0,228058 | 1,603047 | 0,108924 | 0,153183 | 3320,289 | 0,694787289 | 0,506872 | 1,370734 | 0,170458 | 0,260897 | 6076,458 | -1,85338 | 0,516514 | -3,58824 | 0,000333 | 0,00141 | 7602,168779 | -1,486547679 | 0,191493 | -7,76293 | 8,3E-15 | 9,78E-14 |
| 492 | RM25_RS02370 | RM25_0483 | NADH-quinone oxidoreductase subunit H | 511 | gene | 589543 | 590913 | + | 6039,817 | 0,079975248 | 0,501501 | 0,159472 | 0,873297 | 0,913204 | 8369,043348 | 0,099112926 | 0,192269 | 0,515492 | 0,606209 | 0,665813 | 3710,59 | 0,037456026 | 0,64966 | 0,057655 | 0,954024 | 0,971566 | 6261,032 | -1,19234 | 0,635567 | -1,87602 | 0,060653 | 0,099342 | 6080,428851 | -1,220520563 | 0,238242 | -5,12302 | 3,01E-07 | 1,14E-06 |
| 493 | RM25_RS02375 | RM25_0484 | NADH-quinone oxidoreductase subunit I | 512 | gene | 590914 | 591486 | + | 1991,983 | -0,05168632 | 0,517744 | -0,09983 | 0,920479 | 0,947824 | 2722,667285 | -0,017730313 | 0,207429 | -0,08548 | 0,931882 | 0,94797 | 1261,298 | -0,123577701 | 0,701876 | -0,17607 | 0,860241 | 0,898524 | 2160,726 | -1,10697 | 0,684591 | -1,61697 | 0,105884 | 0,155494 | 1916,053009 | -1,182197419 | 0,268396 | -4,40467 | 1,06E-05 | 3,1E-05 |
| 494 | RM25_RS02380 | RM25_0485 | NADH dehydrogenase subunit I J nuoJ | 513 | gene | 591483 | 592352 | + | 2196,3 | 0,042782131 | 0,475098 | 0,090049 | 0,928248 | 0,951253 | 2917,547439 | 0,159615736 | 0,183924 | 0,867835 | 0,385485 | 0,454294 | 1475,053 | -0,186668572 | 0,654397 | -0,28525 | 0,77545 | 0,836493 | 2301,793 | -0,86197 | 0,642434 | -1,34172 | 0,179686 | 0,240891 | 2183,783371 | -1,173588232 | 0,223994 | -5,23937 | 1,61E-07 | 6,34E-07 |
| 495 | RM25_RS02385 | RM25_0486 | NADH-quinone oxidoreductase subunit K | 514 | gene | 592349 | 592648 | + | 651,7687 | 0,058995611 | 0,480454 | 0,122791 | 0,902272 | 0,936028 | 872,5987948 | 0,157942012 | 0,206809 | 0,763708 | 0,445041 | 0,513442 | 430,9386 | -0,136184474 | 0,651541 | -0,20902 | 0,834433 | 0,879159 | 679,7003 | -0,9234 | 0,633973 | -1,45652 | 0,145248 | 0,201619 | 651,1123008 | -1,183519738 | 0,25991 | -4,55357 | 5,27E-06 | 1,61E-05 |
| 496 | RM25_RS02390 | RM25_0487 | NADH dehydrogenase subunit I L nuoL | 515 | gene | 592659 | 594566 | + | 5465,965 | 0,031928591 | 0,566802 | 0,056331 | 0,955078 | 0,971147 | 7913,446024 | 0,068466132 | 0,198791 | 0,344413 | 0,730535 | 0,779833 | 3018,484 | -0,063013809 | 0,716392 | -0,08796 | 0,929908 | 0,952954 | 5766,303 | -1,37236 | 0,712934 | -1,92495 | 0,054235 | 0,090662 | 5416,805079 | -1,47193177 | 0,208177 | -7,07057 | 1,54E-12 | 1,3E-11 |
| 497 | RM25_RS02395 | RM25_0488 | NADH dehydrogenase subunit M | 516 | gene | 594581 | 596122 | + | 4672,952 | 0,225982893 | 0,538372 | 0,419753 | 0,674666 | 0,754717 | 6717,996655 | 0,212384185 | 0,189131 | 1,122948 | 0,26146 | 0,326409 | 2627,908 | 0,261818222 | 0,665648 | 0,393329 | 0,694077 | 0,770579 | 4595,126 | -1,43 | 0,653665 | -2,18766 | 0,028695 | 0,054499 | 4935,988654 | -1,348417742 | 0,225347 | -5,98373 | 2,18E-09 | 1,16E-08 |
| 498 | RM25_RS02400 | RM25_0489 | NADH-quinone oxidoreductase subunit N | 517 | gene | 596126 | 597682 | + | 4359,664 | 0,235007586 | 0,515926 | 0,455506 | 0,648745 | 0,733729 | 6290,575276 | 0,272167166 | 0,217728 | 1,250036 | 0,211287 | 0,272579 | 2428,753 | 0,140087896 | 0,595276 | 0,235333 | 0,813951 | 0,866132 | 4272,932 | -1,35398 | 0,601361 | -2,25153 | 0,024352 | 0,047872 | 4620,986826 | -1,449244449 | 0,185169 | -7,82661 | 5,01E-15 | 6,19E-14 |
| 499 | RM25_RS02405 | RM25_0490 | Putative POLYPRENYL-DIPHOSPHATE SYNTHASE GRCC1 | 518 | gene | 597696 | 598673 | + | 4967,921 | 0,697963438 | 0,681232 | 1,024561 | 0,30557 | 0,414879 | 8116,447454 | 0,595992319 | 0,227555 | 2,619115 | 0,008816 | 0,017775 | 1819,394 | 1,180634094 | 0,602685 | 1,958956 | 0,050118 | 0,101224 | 4057,594 | -2,58418 | 0,603799 | -4,27987 | 1,87E-05 | 0,000108 | 6029,916952 | -1,967120699 | 0,218107 | -9,01908 | 1,9E-19 | 4,26E-18 |
| 500 | RM25_RS02410 | RM25_0491 | Putative membrane protein | 519 | gene | 598803 | 599258 | - | 552,3991 | 0,621374492 | 0,232138 | 2,676749 | 0,007434 | 0,038683 | 556,3564711 | 0,114585164 | 0,167484 | 0,684157 | 0,493876 | 0,559916 | 548,4418 | 1,177444168 | 0,29002 | 4,059877 | 4,91E-05 | 0,000405 | 462,0185 | -0,72423 | 0,206491 | -3,50731 | 0,000453 | 0,001802 | 653,6411649 | 0,379322823 | 0,230519 | 1,645518 | 0,099863 | 0,137728 |
| 501 | RM25_RS12885 | RM25_0492 | Hypothetical protein | 520 | gene | 599331 | 600536 | - | 1427,875 | 0,643108451 | 0,205923 | 3,12305 | 0,00179 | 0,014109 | 1513,594938 | 0,248060144 | 0,16289 | 1,522869 | 0,127791 | 0,177283 | 1342,155 | 1,119616793 | 0,22484 | 4,979616 | 6,37E-07 | 9,33E-06 | 1181,689 | -0,77032 | 0,203101 | -3,79276 | 0,000149 | 0,000693 | 1703,407055 | 0,15028773 | 0,145294 | 1,034372 | 0,300962 | 0,362663 |
| 502 | RM25_RS02420 | RM25_0493 | Peptidase family M13 | 521 | gene | 600624 | 602597 | - | 2505,446 | 0,503386083 | 0,132908 | 3,787473 | 0,000152 | 0,002314 | 2304,519169 | 0,289664706 | 0,106119 | 2,729634 | 0,00634 | 0,013451 | 2706,373 | 0,689320974 | 0,208609 | 3,304373 | 0,000952 | 0,004674 | 2191,179 | -0,06163 | 0,123146 | -0,50048 | 0,616738 | 0,6753 | 2872,841314 | 0,384878173 | 0,159074 | 2,419492 | 0,015542 | 0,026129 |
| 503 | RM25_RS02425 | RM25_0494 | 2-oxoglutarate ferredoxin oxidoreductase subunit beta | 522 | gene | 602727 | 603806 | - | 894,5142 | 0,148517978 | 0,211918 | 0,700826 | 0,483411 | 0,582814 | 754,5653819 | 0,052815371 | 0,205334 | 0,257217 | 0,797011 | 0,838549 | 1034,463 | 0,216538681 | 0,252837 | 0,856437 | 0,391756 | 0,497773 | 892,3217 | 0,302771 | 0,277681 | 1,090354 | 0,275557 | 0,348999 | 921,0500555 | 0,522797217 | 0,145597 | 3,590712 | 0,00033 | 0,000765 |
| 504 | RM25_RS02430 | RM25_0495 | 2-oxoacid:acceptor oxidoreductase, alpha subunit | 523 | gene | 603803 | 605692 | - | 1030,814 | -0,062104818 | 0,308765 | -0,20114 | 0,84059 | 0,894919 | 729,3322976 | -0,165600311 | 0,212882 | -0,7779 | 0,436629 | 0,504973 | 1332,295 | -0,007137487 | 0,245691 | -0,02905 | 0,976824 | 0,98645 | 1104,391 | 0,726612 | 0,273855 | 2,653274 | 0,007972 | 0,019491 | 989,4457526 | 0,945690662 | 0,176111 | 5,369869 | 7,88E-08 | 3,29E-07 |
| 505 | RM25_RS02435 | RM25_0496 | Mannosyltransferase MptA | 524 | gene | 605819 | 607453 | + | 1585,63 | -0,125725799 | 0,153261 | -0,82034 | 0,412023 | 0,522129 | 1690,573462 | 0,180957262 | 0,130041 | 1,391544 | 0,164061 | 0,219213 | 1480,686 | -0,47692271 | 0,16902 | -2,82169 | 0,004777 | 0,017243 | 1748,768 | 0,062338 | 0,180745 | 0,344894 | 0,730174 | 0,775584 | 1488,308605 | -0,542887775 | 0,132168 | -4,10754 | 4E-05 | 0,000108 |
| 506 | RM25_RS02440 | RM25_0497 | Hypothetical protein | 525 | gene | 607620 | 608006 | + | 715,6896 | -0,427308861 | 0,380355 | -1,12345 | 0,261247 | 0,36887 | 461,8596188 | 0,195222051 | 0,171768 | 1,136543 | 0,255729 | 0,320955 | 969,5196 | -0,734304168 | 0,34608 | -2,12178 | 0,033857 | 0,07564 | 862,0318 | 1,439307 | 0,298768 | 4,817477 | 1,45E-06 | 1,08E-05 | 595,8675674 | 0,544952317 | 0,23842 | 2,285682 | 0,022273 | 0,036255 |
| 507 | RM25_RS02445 | RM25_0498 | Exodeoxyribonuclease III | 526 | gene | 608065 | 609006 | - | 673,7479 | 0,212095117 | 0,152776 | 1,388277 | 0,165053 | 0,267374 | 749,073689 | 0,246162725 | 0,138657 | 1,775335 | 0,075843 | 0,111866 | 598,4221 | 0,171890136 | 0,226948 | 0,757399 | 0,448811 | 0,552689 | 660,2863 | -0,34823 | 0,19783 | -1,76025 | 0,078365 | 0,121357 | 709,9227264 | -0,361271095 | 0,166623 | -2,16819 | 0,030144 | 0,04743 |
| 508 | RM25_RS02450 | RM25_0499 | Hypothetical protein | 527 | gene | 609085 | 609579 | + | 247,7538 | -0,649195608 | 0,348012 | -1,86544 | 0,06212 | 0,142451 | 168,6278605 | -0,72674132 | 0,262348 | -2,77014 | 0,005603 | 0,012139 | 326,8797 | -0,613331275 | 0,311908 | -1,96639 | 0,049254 | 0,099906 | 317,0784 | 0,845784 | 0,2683 | 3,15238 | 0,001619 | 0,005279 | 189,4711399 | 1,02140892 | 0,311262 | 3,281512 | 0,001033 | 0,002179 |
| 510 | RM25_RS02460 | RM25_0501 | Alkyl hydroperoxide reductase subunit C | 531 | gene | 610090 | 610653 | + | 17749,93 | -0,219811372 | 0,44325 | -0,49591 | 0,619959 | 0,707967 | 23883,75157 | -0,017940629 | 0,263494 | -0,06809 | 0,945716 | 0,956675 | 11616,12 | -0,643893122 | 0,501668 | -1,28351 | 0,199315 | 0,296208 | 20334,09 | -0,80679 | 0,494363 | -1,63197 | 0,102685 | 0,152028 | 16076,16048 | -1,403619209 | 0,286391 | -4,90107 | 9,53E-07 | 3,34E-06 |
| 511 | RM25_RS02465 | RM25_0502 | Alkyl hydroperoxide reductase, F subunit | 532 | gene | 610877 | 611532 | + | 3097,538 | 1,006184802 | 0,39242 | 2,564049 | 0,010346 | 0,048288 | 4355,153911 | 1,076504022 | 0,226344 | 4,756058 | 1,97E-06 | 9,52E-06 | 1839,921 | 0,84446185 | 0,236554 | 3,569849 | 0,000357 | 0,002148 | 2190,306 | -1,15383 | 0,272407 | -4,23567 | 2,28E-05 | 0,000129 | 4056,317647 | -1,33335956 | 0,180464 | -7,38849 | 1,49E-13 | 1,45E-12 |
| 514 | RM25_RS02480 | RM25_0505 | Hypothetical protein | 539 | gene | 612931 | 613101 | + | 2577,163 | 0,736247635 | 0,567757 | 1,296766 | 0,194712 | 0,298204 | 3804,082617 | 0,392919963 | 0,213748 | 1,838243 | 0,066027 | 0,099124 | 1350,243 | 1,880449309 | 0,41737 | 4,505475 | 6,62E-06 | 7,06E-05 | 2072,85 | -2,56084 | 0,417002 | -6,14108 | 8,2E-10 | 1,19E-08 | 3155,169274 | -1,039748849 | 0,213058 | -4,88013 | 1,06E-06 | 3,68E-06 |
| 515 | RM25_RS02485 | RM25_0506 | UPF0336 protein | 540 | gene | 613271 | 613705 | + | 641,6386 | 0,17662467 | 0,116185 | 1,520206 | 0,128459 | 0,2268 | 615,5336696 | 0,290308003 | 0,149292 | 1,94456 | 0,051828 | 0,081493 | 667,7435 | 0,074443175 | 0,209818 | 0,354798 | 0,722741 | 0,794172 | 634,3702 | 0,17335 | 0,171397 | 1,011395 | 0,311827 | 0,386423 | 666,9586163 | 0,003117417 | 0,155356 | 0,020066 | 0,98399 | 0,986922 |
| 516 | RM25_RS02490 | RM25_0507 | MaoC domain protein dehydratase | 541 | gene | 613702 | 614133 | + | 570,935 | 0,425614986 | 0,162428 | 2,620337 | 0,008784 | 0,042939 | 646,1465923 | 0,509490735 | 0,161776 | 3,149365 | 0,001636 | 0,004112 | 495,7235 | 0,31983402 | 0,194124 | 1,647575 | 0,09944 | 0,171959 | 514,9194 | -0,33188 | 0,168713 | -1,96713 | 0,049168 | 0,083796 | 641,2624957 | -0,474285847 | 0,147073 | -3,22483 | 0,00126 | 0,002598 |
| 517 | RM25_RS02495 | RM25_0508 | UDP-N-acetylmuramate dehydrogenase | 542 | gene | 614203 | 615336 | + | 1081,729 | 0,699079414 | 0,288204 | 2,42564 | 0,015281 | 0,061278 | 1326,920345 | 0,412265498 | 0,148961 | 2,767613 | 0,005647 | 0,012211 | 836,5386 | 1,188351291 | 0,208463 | 5,70053 | 1,19E-08 | 2,54E-07 | 877,4606 | -1,21623 | 0,167232 | -7,27271 | 3,52E-13 | 9,55E-12 | 1310,736634 | -0,394454836 | 0,156877 | -2,51442 | 0,011923 | 0,020572 |
| 518 | RM25_RS02500 | RM25_0509 | Hypothetical protein | 543 | gene | 615358 | 616842 | + | 860,4875 | -0,120131193 | 0,155972 | -0,77021 | 0,441175 | 0,549023 | 870,800575 | -0,025367852 | 0,151283 | -0,16768 | 0,866831 | 0,896499 | 850,1745 | -0,215175369 | 0,292389 | -0,73592 | 0,461778 | 0,565409 | 945,0504 | -0,00766 | 0,206232 | -0,03716 | 0,970356 | 0,976988 | 810,1907967 | -0,133358941 | 0,245293 | -0,54367 | 0,586667 | 0,64676 |
| 520 | RM25_RS12230 | RM25_0511 | Preprotein translocase, SecE subunit | 547 | gene | 617275 | 617862 | + | 2556,114 | 0,555132713 | 0,524102 | 1,059208 | 0,289505 | 0,39881 | 3772,638182 | 0,282949958 | 0,237896 | 1,189385 | 0,234288 | 0,297531 | 1339,589 | 1,400678901 | 0,341912 | 4,096601 | 4,19E-05 | 0,00035 | 2213,571 | -2,26133 | 0,346845 | -6,51971 | 7,04E-11 | 1,33E-09 | 2979,792382 | -1,107878472 | 0,223796 | -4,95039 | 7,41E-07 | 2,65E-06 |
| 521 | RM25_RS02515 | RM25_0512 | Transcription antitermination protein nusG | 548 | gene | 617882 | 618760 | + | 6436,811 | 0,607873004 | 0,533239 | 1,139965 | 0,254301 | 0,362168 | 9292,67376 | 0,255243495 | 0,216929 | 1,17662 | 0,239347 | 0,303139 | 3580,948 | 1,653562044 | 0,419618 | 3,940633 | 8,13E-05 | 0,000617 | 5458,799 | -2,34447 | 0,375387 | -6,24549 | 4,22E-10 | 6,51E-09 | 7605,014459 | -0,915991024 | 0,272108 | -3,36628 | 0,000762 | 0,001663 |
| 522 | RM25_RS02520 | RM25_0513 | 50S ribosomal protein L11 | 549 | gene | 618870 | 619298 | + | 4120,947 | 0,717510467 | 0,384242 | 1,86734 | 0,061854 | 0,142196 | 5145,30776 | 0,396937761 | 0,187075 | 2,121811 | 0,033854 | 0,056833 | 3096,587 | 1,29745652 | 0,456809 | 2,840262 | 0,004508 | 0,016523 | 3329,302 | -1,35964 | 0,430524 | -3,15811 | 0,001588 | 0,005198 | 5012,188063 | -0,427333535 | 0,242378 | -1,76309 | 0,077886 | 0,109795 |
| 523 | RM25_RS02525 | RM25_0514 | 50S ribosomal protein L1 | 550 | gene | 619418 | 620128 | + | 4421,973 | 0,754873258 | 0,497723 | 1,516653 | 0,129354 | 0,227868 | 6054,307769 | 0,442788789 | 0,219674 | 2,015664 | 0,043835 | 0,071255 | 2789,638 | 1,512077326 | 0,535014 | 2,826241 | 0,00471 | 0,017105 | 3519,634 | -1,8714 | 0,529767 | -3,5325 | 0,000412 | 0,001681 | 5436,819314 | -0,771843579 | 0,234067 | -3,29753 | 0,000975 | 0,002075 |
| 524 | RM25_RS02530 | RM25_0515 | Octanoyltransferase | 551 | gene | 620218 | 621168 | + | 846,597 | -0,209233822 | 0,321916 | -0,64996 | 0,515715 | 0,612468 | 621,1104065 | -0,509047259 | 0,276323 | -1,84222 | 0,065443 | 0,098411 | 1072,084 | -0,040128137 | 0,269427 | -0,14894 | 0,881602 | 0,914182 | 952,89 | 0,507192 | 0,35563 | 1,426179 | 0,153817 | 0,211274 | 769,63067 | 1,037869194 | 0,152707 | 6,796489 | 1,07E-11 | 7,9E-11 |
| 525 | RM25_RS02535 | RM25_0516 | Lipoyl synthase | 552 | gene | 621165 | 622115 | + | 1826,416 | 1,580162027 | 0,302361 | 5,226071 | 1,73E-07 | 9,49E-06 | 1899,816373 | 0,986378723 | 0,161745 | 6,098359 | 1,07E-09 | 9,12E-09 | 1753,015 | 2,407720622 | 0,280117 | 8,595398 | 8,3E-18 | 8,89E-16 | 974,5493 | -1,25307 | 0,204369 | -6,1314 | 8,71E-10 | 1,24E-08 | 2674,323037 | 0,208548421 | 0,213876 | 0,975091 | 0,329515 | 0,39317 |
| 526 | RM25_RS02540 | RM25_0517 | 50S ribosomal protein L10 | 553 | gene | 622484 | 623131 | + | 13765,28 | 0,907203774 | 0,705179 | 1,286488 | 0,198273 | 0,301114 | 20544,56242 | 0,611458047 | 0,222934 | 2,742776 | 0,006092 | 0,013018 | 6986,006 | 1,936946946 | 0,844324 | 2,294081 | 0,021786 | 0,054685 | 10261,74 | -2,53385 | 0,83677 | -3,02813 | 0,002461 | 0,007513 | 17586,06167 | -1,182959279 | 0,264638 | -4,4701 | 7,82E-06 | 2,33E-05 |
| 527 | RM25_RS02545 | RM25_0518 | 50S ribosomal protein L7/L12 | 554 | gene | 623148 | 623543 | + | 4629,841 | 0,804067339 | 0,694512 | 1,157744 | 0,246969 | 0,355593 | 7014,139526 | 0,524669684 | 0,255999 | 2,049495 | 0,040414 | 0,066519 | 2245,543 | 1,820713285 | 0,787628 | 2,311642 | 0,020797 | 0,052766 | 3606,094 | -2,57874 | 0,78806 | -3,27226 | 0,001067 | 0,003715 | 5770,071097 | -1,257685297 | 0,258062 | -4,87358 | 1,1E-06 | 3,79E-06 |
| 528 | RM25_RS02550 | RM25_0519 | Hypothetical protein | 555 | gene | 623814 | 624440 | + | 496,3375 | -0,120585386 | 0,129649 | -0,93009 | 0,352325 | 0,465226 | 470,3635573 | -0,129154914 | 0,200229 | -0,64504 | 0,518904 | 0,583798 | 522,3114 | -0,11344452 | 0,191545 | -0,59226 | 0,553677 | 0,650232 | 544,4796 | 0,086362 | 0,172614 | 0,500316 | 0,616852 | 0,6753 | 466,2257536 | 0,152270606 | 0,198712 | 0,766287 | 0,443506 | 0,50893 |
| 529 | RM25_RS02555 | RM25_0520 | DNA-directed RNA polymerase, beta subunit | 556 | gene | 624766 | 628242 | + | 10755,29 | 0,694003633 | 0,479779 | 1,446507 | 0,148035 | 0,248517 | 15372,65517 | 0,397065396 | 0,195708 | 2,02887 | 0,042471 | 0,069469 | 6137,916 | 1,526377282 | 0,272947 | 5,592215 | 2,24E-08 | 4,48E-07 | 8782,505 | -2,12744 | 0,229259 | -9,27962 | 1,7E-20 | 1,38E-18 | 13014,42936 | -0,956848806 | 0,212218 | -4,50881 | 6,52E-06 | 1,96E-05 |
| 530 | RM25_RS02560 | RM25_0521 | DNA-directed RNA polymerase, beta' subunit | 557 | gene | 628344 | 632234 | + | 9529,605 | 1,004354161 | 0,510734 | 1,96649 | 0,049242 | 0,123341 | 13843,75104 | 0,723884859 | 0,197477 | 3,66566 | 0,000247 | 0,000775 | 5215,459 | 1,871814617 | 0,275083 | 6,80455 | 1,01E-11 | 4,59E-10 | 6780,918 | -2,28127 | 0,219488 | -10,3936 | 2,65E-25 | 4,98E-23 | 12459,56225 | -1,089998163 | 0,225044 | -4,84349 | 1,28E-06 | 4,36E-06 |
| 531 | RM25_RS02565 | RM25_0522 | 30S ribosomal protein S12 | 558 | gene | 632694 | 633065 | + | 3334,734 | 0,64199398 | 0,606385 | 1,058724 | 0,289726 | 0,398881 | 4635,386101 | -0,009409024 | 0,181713 | -0,05178 | 0,958704 | 0,966909 | 2034,083 | 2,652081293 | 0,405934 | 6,533284 | 6,43E-11 | 2,37E-09 | 2796,608 | -3,11311 | 0,320985 | -9,69861 | 3,06E-22 | 3,43E-20 | 3970,468763 | -0,417866003 | 0,283373 | -1,47461 | 0,140316 | 0,186557 |
| 532 | RM25_RS02570 | RM25_0523 | 30S ribosomal protein S7 | 559 | gene | 633065 | 633535 | + | 2045,19 | 0,980215327 | 0,568616 | 1,723861 | 0,084733 | 0,173597 | 2647,191604 | 0,253623663 | 0,174207 | 1,455872 | 0,145428 | 0,198365 | 1443,188 | 2,923768574 | 0,370382 | 7,893923 | 2,93E-15 | 2,47E-13 | 1476,276 | -2,90292 | 0,295356 | -9,82852 | 8,49E-23 | 1E-20 | 2651,24259 | -0,194721357 | 0,255382 | -0,76247 | 0,445779 | 0,510545 |
| 533 | RM25_RS02575 | RM25_0524 | Translation elongation factor G | 560 | gene | 633706 | 635805 | + | 11878,31 | 0,971488763 | 0,586636 | 1,656032 | 0,097715 | 0,1905 | 16961,06145 | 0,491949706 | 0,196112 | 2,508519 | 0,012124 | 0,023385 | 6795,565 | 2,578270071 | 0,359352 | 7,17477 | 7,24E-13 | 4,27E-11 | 8605,522 | -2,91026 | 0,31837 | -9,14113 | 6,18E-20 | 4,55E-18 | 15398,12068 | -0,785483432 | 0,236816 | -3,31685 | 0,00091 | 0,001958 |
| 534 | RM25_RS02580 | RM25_0525 | Translation elongation factor Tu | 561 | gene | 636040 | 637230 | + | 33692,7 | 1,546565992 | 0,547068 | 2,827009 | 0,004699 | 0,028469 | 39034,43426 | 0,7403021 | 0,226337 | 3,27079 | 0,001072 | 0,002864 | 28350,97 | 3,327863571 | 0,46782 | 7,11355 | 1,13E-12 | 6,2E-11 | 18427,04 | -2,55875 | 0,447657 | -5,71588 | 1,09E-08 | 1,2E-07 | 49022,21265 | 0,059305476 | 0,260079 | 0,228028 | 0,819624 | 0,854047 |
| 535 | RM25_RS11780 | RM25_0526 | Membrane protein, PF09852 family | 562 | gene | 637424 | 638854 | - | 986,7171 | -0,350883864 | 0,15209 | -2,30708 | 0,021051 | 0,073505 | 924,1837435 | -0,345330396 | 0,137813 | -2,50579 | 0,012218 | 0,023527 | 1049,251 | -0,356117961 | 0,276646 | -1,28727 | 0,198 | 0,294439 | 1166,084 | 0,121259 | 0,21423 | 0,566023 | 0,571378 | 0,635283 | 851,8280605 | 0,175263764 | 0,220255 | 0,795731 | 0,426189 | 0,491692 |
| 536 | RM25_RS02590 | RM25_0527 | Hypothetical protein | 563 | gene | 638896 | 639585 | + | 511,2015 | -0,015080403 | 0,336245 | -0,04485 | 0,964227 | 0,977498 | 340,2290402 | -0,259796745 | 0,228384 | -1,13754 | 0,255312 | 0,320602 | 682,1739 | 0,103770958 | 0,161794 | 0,641376 | 0,521279 | 0,62116 | 539,4677 | 0,765621 | 0,193378 | 3,959192 | 7,52E-05 | 0,00038 | 498,1656755 | 1,18334028 | 0,204115 | 5,797415 | 6,73E-09 | 3,29E-08 |
| 537 | RM25_RS02595 | RM25_0528 | Glutamate--tRNA ligase | 564 | gene | 639778 | 641172 | + | 2150,735 | 0,173420174 | 0,179843 | 0,964289 | 0,334901 | 0,445213 | 2499,530074 | 0,138947316 | 0,122362 | 1,135547 | 0,256146 | 0,321246 | 1801,94 | 0,221892302 | 0,185318 | 1,197362 | 0,231165 | 0,329617 | 2143,029 | -0,57487 | 0,128042 | -4,48971 | 7,13E-06 | 4,54E-05 | 2233,121041 | -0,445314271 | 0,14065 | -3,16613 | 0,001545 | 0,003134 |
| 538 | RM25_RS02600 | RM25_0529 | Aminotransferase, class IV | 565 | gene | 641219 | 642091 | + | 1298,792 | 0,322835658 | 0,191888 | 1,682414 | 0,092489 | 0,183704 | 1490,974352 | 0,419614154 | 0,170216 | 2,465183 | 0,013694 | 0,026009 | 1106,61 | 0,195560958 | 0,235226 | 0,831373 | 0,405763 | 0,511435 | 1222,469 | -0,35935 | 0,234486 | -1,53248 | 0,125403 | 0,179028 | 1414,278329 | -0,541437089 | 0,160512 | -3,37319 | 0,000743 | 0,001626 |
| 539 | RM25_RS02605 | RM25_0530 | 30S ribosomal protein S10 | 566 | gene | 642526 | 642837 | + | 3864,155 | 0,644345331 | 0,633747 | 1,016724 | 0,309285 | 0,417973 | 5832,50571 | 0,251298519 | 0,217052 | 1,157782 | 0,246953 | 0,311433 | 1895,805 | 2,132980641 | 0,496091 | 4,299575 | 1,71E-05 | 0,000164 | 3234,17 | -2,97043 | 0,494864 | -6,00252 | 1,94E-09 | 2,56E-08 | 4616,629785 | -1,053817297 | 0,210325 | -5,01043 | 5,43E-07 | 2E-06 |
| 540 | RM25_RS02610 | RM25_0531 | 50S ribosomal protein L3 | 567 | gene | 642849 | 643508 | + | 3325,107 | 0,685445092 | 0,645852 | 1,061304 | 0,288552 | 0,397962 | 5081,083275 | 0,305353529 | 0,223539 | 1,365997 | 0,17194 | 0,228445 | 1569,132 | 2,224173389 | 0,461489 | 4,81956 | 1,44E-06 | 1,92E-05 | 2732,741 | -3,08905 | 0,462364 | -6,68099 | 2,37E-11 | 4,91E-10 | 4017,376474 | -1,135504905 | 0,208515 | -5,44569 | 5,16E-08 | 2,23E-07 |
| 541 | RM25_RS02615 | RM25_0532 | 50S ribosomal protein L4 | 568 | gene | 643505 | 644206 | + | 4782,178 | 0,770091732 | 0,666464 | 1,155489 | 0,24789 | 0,35584 | 7103,399362 | 0,277367456 | 0,221617 | 1,25156 | 0,21073 | 0,27201 | 2460,957 | 2,723415122 | 0,470177 | 5,79232 | 6,94E-09 | 1,59E-07 | 3791,137 | -3,36038 | 0,452954 | -7,41881 | 1,18E-13 | 3,42E-12 | 5902,931824 | -0,882102491 | 0,245693 | -3,59026 | 0,00033 | 0,000766 |
| 542 | RM25_RS02620 | RM25_0533 | 50S ribosomal protein L23 | 569 | gene | 644203 | 644514 | + | 2234,539 | 0,761234152 | 0,588692 | 1,293095 | 0,195978 | 0,299559 | 3262,244415 | 0,30557689 | 0,204398 | 1,495012 | 0,134911 | 0,186174 | 1206,834 | 2,339821383 | 0,323633 | 7,229871 | 4,83E-13 | 2,92E-11 | 1777,665 | -2,93175 | 0,264445 | -11,0864 | 1,46E-28 | 4,3E-26 | 2751,419824 | -0,857134591 | 0,247989 | -3,45635 | 0,000548 | 0,001224 |
| 543 | RM25_RS02625 | RM25_0534 | 50S ribosomal protein L2 | 570 | gene | 644553 | 645389 | + | 4894,552 | 0,827975666 | 0,626332 | 1,321944 | 0,186187 | 0,291031 | 7074,313656 | 0,299506522 | 0,17417 | 1,719623 | 0,085501 | 0,12394 | 2714,791 | 2,729889297 | 0,393661 | 6,934622 | 4,07E-12 | 2E-10 | 3785,715 | -3,21183 | 0,343212 | -9,35815 | 8,11E-21 | 7,08E-19 | 6126,206329 | -0,744809206 | 0,238807 | -3,11887 | 0,001815 | 0,003629 |
| 544 | RM25_RS02630 | RM25_0535 | 30S ribosomal protein S19 | 571 | gene | 645407 | 645688 | + | 2535,015 | 0,910481632 | 0,6233 | 1,460744 | 0,144086 | 0,244676 | 3604,240315 | 0,360650397 | 0,191304 | 1,885226 | 0,059399 | 0,090735 | 1465,79 | 2,815918569 | 0,415398 | 6,77884 | 1,21E-11 | 5,02E-10 | 1890,652 | -3,16982 | 0,374898 | -8,45515 | 2,79E-17 | 1,56E-15 | 3238,592475 | -0,676123014 | 0,248242 | -2,72365 | 0,006457 | 0,011679 |
| 545 | RM25_RS02635 | RM25_0536 | 50S ribosomal protein L22 | 572 | gene | 645714 | 646199 | + | 2121,473 | 0,92898413 | 0,656061 | 1,416004 | 0,156774 | 0,258404 | 3056,528416 | 0,373204502 | 0,202201 | 1,845709 | 0,064934 | 0,097633 | 1186,418 | 3,034518093 | 0,412646 | 7,353806 | 1,93E-13 | 1,26E-11 | 1566,755 | -3,42215 | 0,361023 | -9,47902 | 2,57E-21 | 2,42E-19 | 2721,25181 | -0,725134747 | 0,259969 | -2,78931 | 0,005282 | 0,009734 |
| 546 | RM25_RS02640 | RM25_0537 | 30S ribosomal protein S3 | 573 | gene | 646202 | 647020 | + | 2401,475 | 1,016759162 | 0,637907 | 1,5939 | 0,110958 | 0,205605 | 3503,092407 | 0,533323933 | 0,2228 | 2,393733 | 0,016678 | 0,030783 | 1299,858 | 2,866238989 | 0,379494 | 7,552793 | 4,26E-14 | 2,95E-12 | 1702,341 | -3,24226 | 0,377663 | -8,58507 | 9,08E-18 | 5,78E-16 | 3148,817579 | -0,872731371 | 0,214022 | -4,07777 | 4,55E-05 | 0,000121 |
| 547 | RM25_RS02645 | RM25_0538 | 50S ribosomal protein L16 | 574 | gene | 647026 | 647445 | + | 1796,44 | 0,963903224 | 0,621388 | 1,551211 | 0,120851 | 0,217606 | 2566,348125 | 0,442761853 | 0,212619 | 2,08242 | 0,037304 | 0,061833 | 1026,532 | 2,792503447 | 0,411305 | 6,789381 | 1,13E-11 | 5,01E-10 | 1306,215 | -3,12468 | 0,383208 | -8,154 | 3,52E-16 | 1,73E-14 | 2324,433069 | -0,738388697 | 0,2459 | -3,00279 | 0,002675 | 0,005216 |
| 548 | RM25_RS02650 | RM25_0539 | 50S ribosomal protein L29 | 575 | gene | 647447 | 647695 | + | 1264,634 | 0,89385772 | 0,731577 | 1,221823 | 0,221775 | 0,327726 | 1851,555406 | 0,332166646 | 0,229465 | 1,447571 | 0,147737 | 0,201165 | 677,7119 | 3,237345152 | 0,611731 | 5,292109 | 1,21E-07 | 2,1E-06 | 950,4459 | -3,70581 | 0,602189 | -6,15389 | 7,56E-10 | 1,1E-08 | 1609,93052 | -0,768782363 | 0,254035 | -3,02629 | 0,002476 | 0,004867 |
| 549 | RM25_RS02655 | RM25_0540 | 30S ribosomal protein S17 | 576 | gene | 647692 | 647967 | + | 1857,768 | 0,858662698 | 0,69178 | 1,241237 | 0,214518 | 0,319608 | 2692,866012 | 0,286689254 | 0,223637 | 1,28194 | 0,199864 | 0,259834 | 1022,67 | 3,093794011 | 0,521244 | 5,935404 | 2,93E-09 | 7,51E-08 | 1416,443 | -3,54921 | 0,487557 | -7,27959 | 3,35E-13 | 9,18E-12 | 2342,75041 | -0,712127987 | 0,281406 | -2,53061 | 0,011387 | 0,019691 |
| 550 | RM25_RS02660 | RM25_0541 | 50S ribosomal protein L14 | 577 | gene | 648098 | 648469 | + | 5475,274 | 0,690354245 | 0,659714 | 1,046445 | 0,295356 | 0,405684 | 7968,776628 | 0,162912917 | 0,225356 | 0,722913 | 0,469733 | 0,538241 | 2981,772 | 2,570306789 | 0,562077 | 4,572873 | 4,81E-06 | 5,43E-05 | 4497,023 | -3,17692 | 0,545563 | -5,82321 | 5,77E-09 | 6,67E-08 | 6614,490064 | -0,740909405 | 0,262117 | -2,82663 | 0,004704 | 0,008751 |
| 551 | RM25_RS02665 | RM25_0542 | 50S ribosomal protein L24 | 578 | gene | 648471 | 648842 | + | 1836,779 | 0,930171925 | 0,619581 | 1,501292 | 0,13328 | 0,231838 | 2616,739187 | 0,458606235 | 0,213443 | 2,148613 | 0,031665 | 0,053646 | 1056,818 | 2,467392064 | 0,566601 | 4,354724 | 1,33E-05 | 0,000133 | 1355,24 | -2,81602 | 0,543236 | -5,18379 | 2,17E-07 | 1,88E-06 | 2357,523891 | -0,777521259 | 0,26632 | -2,9195 | 0,003506 | 0,006659 |
| 552 | RM25_RS02670 | RM25_0543 | 50S ribosomal protein L5 | 579 | gene | 648842 | 649504 | + | 4209,736 | 0,814317823 | 0,595729 | 1,366926 | 0,171649 | 0,275409 | 6139,32733 | 0,377795516 | 0,208281 | 1,813878 | 0,069696 | 0,104037 | 2280,145 | 2,313522199 | 0,418809 | 5,524048 | 3,31E-08 | 6,45E-07 | 3267,961 | -2,85842 | 0,365948 | -7,811 | 5,67E-15 | 2,23E-13 | 5251,599267 | -0,889882018 | 0,268872 | -3,30969 | 0,000934 | 0,002001 |
| 553 | RM25_RS02675 | RM25_0544 | 30S ribosomal protein S14 type Z | 580 | gene | 649513 | 649698 | + | 993,0708 | 0,941943416 | 0,616789 | 1,527172 | 0,126718 | 0,224791 | 1408,362247 | 0,421623875 | 0,211173 | 1,996578 | 0,045871 | 0,073952 | 577,7793 | 2,688159163 | 0,468925 | 5,732599 | 9,89E-09 | 2,14E-07 | 729,0542 | -3,0143 | 0,428067 | -7,04167 | 1,9E-12 | 4,81E-11 | 1277,711564 | -0,707879614 | 0,266024 | -2,66096 | 0,007792 | 0,013903 |
| 554 | RM25_RS02680 | RM25_0545 | 30S ribosomal protein S8 | 581 | gene | 649858 | 650265 | + | 5072,962 | 0,957509642 | 0,732484 | 1,307209 | 0,191142 | 0,295091 | 7736,978891 | 0,510731961 | 0,2199 | 2,322562 | 0,020203 | 0,03602 | 2408,945 | 3,062385764 | 0,579134 | 5,287873 | 1,24E-07 | 2,13E-06 | 3704,059 | -3,68011 | 0,559127 | -6,58189 | 4,65E-11 | 9,05E-10 | 6559,050893 | -1,097251412 | 0,265922 | -4,12621 | 3,69E-05 | 0,0001 |
| 555 | RM25_RS02685 | RM25_0546 | 50S ribosomal protein L6 | 582 | gene | 650296 | 650838 | + | 3973,544 | 0,850279738 | 0,683043 | 1,24484 | 0,21319 | 0,318233 | 6055,888914 | 0,439471164 | 0,221435 | 1,984648 | 0,047184 | 0,0755 | 1891,199 | 2,619263625 | 0,519338 | 5,043463 | 4,57E-07 | 6,95E-06 | 3040,683 | -3,33072 | 0,500225 | -6,65844 | 2,77E-11 | 5,59E-10 | 5006,57418 | -1,117456327 | 0,252108 | -4,43245 | 9,32E-06 | 2,73E-05 |
| 556 | RM25_RS02690 | RM25_0547 | 50S ribosomal protein L18 | 583 | gene | 650841 | 651224 | + | 2478,579 | 1,029910592 | 0,7724 | 1,33339 | 0,182404 | 0,287581 | 3786,925874 | 0,5818426 | 0,236549 | 2,459716 | 0,013905 | 0,026388 | 1170,233 | 3,233537747 | 0,683689 | 4,729542 | 2,25E-06 | 2,75E-05 | 1749,326 | -3,79956 | 0,673464 | -5,64181 | 1,68E-08 | 1,81E-07 | 3258,135594 | -1,120553865 | 0,264676 | -4,23369 | 2,3E-05 | 6,44E-05 |
| 557 | RM25_RS02695 | RM25_0548 | 30S ribosomal protein S5 | 584 | gene | 651247 | 651858 | + | 4526,132 | 1,062842043 | 0,73846 | 1,439268 | 0,150075 | 0,250514 | 6839,664308 | 0,628632168 | 0,220393 | 2,852324 | 0,00434 | 0,009627 | 2212,6 | 3,014065116 | 0,675132 | 4,464406 | 8,03E-06 | 8,41E-05 | 3145,035 | -3,50916 | 0,668849 | -5,24656 | 1,55E-07 | 1,4E-06 | 5998,41864 | -1,092421315 | 0,240465 | -4,54296 | 5,55E-06 | 1,69E-05 |
| 558 | RM25_RS02700 | RM25_0549 | Hypothetical protein | 585 | gene | 651858 | 652040 | + | 1889,598 | 1,04347926 | 0,84813 | 1,23033 | 0,218574 | 0,324215 | 2973,891978 | 0,664346673 | 0,221946 | 2,993283 | 0,00276 | 0,006512 | 805,3041 | 3,109540893 | 0,894645 | 3,475726 | 0,000509 | 0,002825 | 1328,414 | -3,82779 | 1,176914 | -3,2524 | 0,001144 | 0,003955 | 2491,135706 | -1,358949781 | 0,287172 | -4,73218 | 2,22E-06 | 7,28E-06 |
| 559 | RM25_RS02705 | RM25_0550 | 50S ribosomal protein L15 | 586 | gene | 652042 | 652485 | + | 2077,026 | 0,995324861 | 0,757909 | 1,313251 | 0,189098 | 0,293267 | 3245,456647 | 0,625692025 | 0,244159 | 2,562637 | 0,010388 | 0,020506 | 908,5955 | 2,84411352 | 0,677115 | 4,200339 | 2,67E-05 | 0,000236 | 1488,097 | -3,56775 | 0,673177 | -5,29988 | 1,16E-07 | 1,07E-06 | 2711,768928 | -1,319656474 | 0,255745 | -5,16006 | 2,47E-07 | 9,49E-07 |
| 560 | RM25_RS02710 | RM25_0551 | FMN reductase | 587 | gene | 652842 | 653489 | - | 375,9588 | -0,220406698 | 0,35568 | -0,61968 | 0,53547 | 0,631331 | 250,6597015 | -0,704808078 | 0,237923 | -2,96234 | 0,003053 | 0,007118 | 501,2579 | 0,012013139 | 0,176113 | 0,068213 | 0,945616 | 0,965692 | 425,9415 | 0,614389 | 0,229404 | 2,678194 | 0,007402 | 0,018404 | 339,8250651 | 1,389284925 | 0,198754 | 6,989973 | 2,75E-12 | 2,21E-11 |
| 561 | RM25_RS02715 | RM25_0552 | Putative luciferase-like monooxygenase, FMN-dependent, ( | 588 | gene | 653600 | 654796 | - | 2076,029 | -0,34699169 | 0,496425 | -0,69898 | 0,484564 | 0,583905 | 1008,836015 | -0,415026427 | 0,12865 | -3,22601 | 0,001255 | 0,003284 | 3143,221 | -0,325246947 | 0,343847 | -0,9459 | 0,344197 | 0,450707 | 2440,402 | 1,544077 | 0,28825 | 5,356736 | 8,47E-08 | 8,02E-07 | 1781,69789 | 1,674557673 | 0,226705 | 7,386519 | 1,51E-13 | 1,47E-12 |
| 562 | RM25_RS02720 | RM25_0553 | Putative extracellular solute-binding dependent transport lip | 589 | gene | 655047 | 656765 | - | 2325,747 | 0,254117657 | 0,207639 | 1,223841 | 0,221012 | 0,326804 | 2750,58614 | 0,335641459 | 0,132113 | 2,540564 | 0,011067 | 0,021633 | 1900,908 | 0,136941126 | 0,232542 | 0,588887 | 0,555937 | 0,652237 | 2244,649 | -0,4839 | 0,15166 | -3,19065 | 0,00142 | 0,004726 | 2477,798684 | -0,638568759 | 0,169421 | -3,76913 | 0,000164 | 0,000403 |
| 563 | RM25_RS02725 | RM25_0554 | Oligopeptide/dipeptide ABC transporter, ATPase subunit | 590 | gene | 656836 | 658935 | - | 1061,45 | -1,561108419 | 0,532001 | -2,93441 | 0,003342 | 0,022188 | 659,6194036 | -2,565957397 | 0,337434 | -7,60433 | 2,86E-14 | 4,5E-13 | 1463,28 | -1,208785264 | 0,387461 | -3,11976 | 0,00181 | 0,007944 | 1662,546 | 0,786462 | 0,413696 | 1,90106 | 0,057294 | 0,094966 | 529,5258922 | 2,217483068 | 0,342788 | 6,468965 | 9,87E-11 | 6,42E-10 |
| 564 | RM25_RS02730 | RM25_0555 | Oligopeptide transport system permease protein OppC | 591 | gene | 658932 | 659918 | - | 360,6304 | -1,501247437 | 0,471239 | -3,18574 | 0,001444 | 0,012604 | 248,3860486 | -2,37332907 | 0,33689 | -7,04482 | 1,86E-12 | 2,37E-11 | 472,8747 | -1,134940129 | 0,382126 | -2,97007 | 0,002977 | 0,011912 | 559,1554 | 0,576219 | 0,364938 | 1,578952 | 0,114347 | 0,165754 | 185,3383345 | 1,884016779 | 0,386854 | 4,8701 | 1,12E-06 | 3,85E-06 |
| 565 | RM25_RS02735 | RM25_0556 | Oligopeptide-transport membrane protein ABC transporter ( | 592 | gene | 659920 | 660903 | - | 541,6949 | -0,811548087 | 0,173049 | -4,68969 | 2,74E-06 | 9,35E-05 | 528,1953477 | -0,672772224 | 0,189375 | -3,5526 | 0,000381 | 0,001145 | 555,1945 | -0,946565943 | 0,299658 | -3,15882 | 0,001584 | 0,007098 | 726,2786 | 0,108874 | 0,248754 | 0,437679 | 0,661619 | 0,715338 | 386,6491852 | -0,103324957 | 0,25233 | -0,40948 | 0,682185 | 0,732533 |
| 566 | RM25_RS02835 | RM25_0557 | Transposase | 593 | gene | 660972 | 662351 | - | 67,08853 | -0,595749259 | 0,258435 | -2,30521 | 0,021155 | 0,07365 | 63,06844246 | -0,86098741 | 0,331788 | -2,595 | 0,009459 | 0,018943 | 71,10862 | -0,382377616 | 0,389895 | -0,98072 | 0,326732 | 0,432401 | 85,17364 | -0,08432 | 0,366386 | -0,23013 | 0,817988 | 0,853096 | 52,50581135 | 0,464103907 | 0,384366 | 1,207452 | 0,227258 | 0,283711 |
| 567 | RM25_RS02745 | RM25_0558 | Protein translocase subunit SecY | 594 | gene | 662780 | 664093 | + | 3202,44 | 0,519959669 | 0,437322 | 1,188962 | 0,234455 | 0,341539 | 4364,74085 | 0,153454164 | 0,21321 | 0,719733 | 0,471689 | 0,538954 | 2040,14 | 1,384327914 | 0,293035 | 4,724112 | 2,31E-06 | 2,81E-05 | 2808 | -1,93025 | 0,240798 | -8,01602 | 1,09E-15 | 4,96E-14 | 3692,013783 | -0,657261494 | 0,236947 | -2,77387 | 0,005539 | 0,010184 |
| 568 | RM25_RS02750 | RM25_0559 | Adenylate kinase | 595 | gene | 664105 | 664665 | + | 1070,899 | 0,792158577 | 0,409448 | 1,9347 | 0,053027 | 0,128983 | 1379,41778 | 0,3774098 | 0,231038 | 1,633543 | 0,102355 | 0,145069 | 762,3801 | 1,651843377 | 0,318528 | 5,18586 | 2,15E-07 | 3,54E-06 | 834,582 | -1,7647 | 0,294231 | -5,99768 | 2E-09 | 2,61E-08 | 1327,904952 | -0,447060529 | 0,227211 | -1,9676 | 0,049114 | 0,072806 |
| 569 | RM25_RS02755 | RM25_0560 | Methionine aminopeptidase Map | 596 | gene | 664677 | 665507 | + | 1153,263 | 0,753357052 | 0,459126 | 1,640851 | 0,100828 | 0,193702 | 1669,175031 | 0,58037985 | 0,181744 | 3,193391 | 0,001406 | 0,003609 | 637,3501 | 1,238226294 | 0,244667 | 5,060873 | 4,17E-07 | 6,47E-06 | 916,4105 | -1,87946 | 0,237229 | -7,92256 | 2,33E-15 | 9,62E-14 | 1419,055026 | -1,172854418 | 0,153909 | -7,62044 | 2,53E-14 | 2,8E-13 |
| 570 | RM25_RS02760 | RM25_0561 | Hypothetical protein | 597 | gene | 665533 | 665919 | + | 318,0341 | 0,536477852 | 0,168855 | 3,177148 | 0,001487 | 0,012778 | 348,2201822 | 0,753367547 | 0,173833 | 4,333864 | 1,47E-05 | 6,01E-05 | 287,848 | 0,288033427 | 0,249311 | 1,155319 | 0,24796 | 0,348504 | 274,6251 | -0,05913 | 0,243961 | -0,24236 | 0,808503 | 0,845824 | 368,8435023 | -0,477842991 | 0,174271 | -2,74195 | 0,006108 | 0,011108 |
| 571 | RM25_RS02765 | RM25_0562 | Hypothetical protein | 598 | gene | 665950 | 666234 | - | 1197,238 | -0,305915431 | 0,429945 | -0,71152 | 0,476761 | 0,577158 | 1334,026862 | 0,682850465 | 0,21056 | 3,243025 | 0,001183 | 0,003118 | 1060,449 | -1,701712952 | 0,366857 | -4,63863 | 3,51E-06 | 4,07E-05 | 1396,429 | 0,617458 | 0,390294 | 1,582031 | 0,113642 | 0,164936 | 1050,681026 | -1,73294327 | 0,155372 | -11,1535 | 6,88E-29 | 3,86E-27 |
| 572 | RM25_RS02770 | RM25_0563 | Translation initiation factor IF-1 | 599 | gene | 666593 | 666814 | + | 6344,121 | 1,053734558 | 0,542345 | 1,942922 | 0,052026 | 0,127418 | 7739,932867 | 0,33995969 | 0,208218 | 1,632711 | 0,10253 | 0,14523 | 4948,309 | 2,580978548 | 0,569266 | 4,533868 | 5,79E-06 | 6,26E-05 | 4419,971 | -2,31645 | 0,547008 | -4,23477 | 2,29E-05 | 0,000129 | 8363,06203 | -0,047564573 | 0,263067 | -0,18081 | 0,856518 | 0,883894 |
| 573 | RM25_RS12235 | RM25_0564 | Hypothetical protein | 600 | gene | 666907 | 667020 | + | 1617,383 | 0,983560628 | 0,525507 | 1,871641 | 0,061256 | 0,1416 | 2065,884093 | 0,374839936 | 0,223604 | 1,676359 | 0,093668 | 0,134373 | 1168,883 | 2,396359968 | 0,500588 | 4,787094 | 1,69E-06 | 2,18E-05 | 1165,761 | -2,32012 | 0,500379 | -4,63672 | 3,54E-06 | 2,43E-05 | 2101,706682 | -0,262635075 | 0,230047 | -1,14166 | 0,253596 | 0,311478 |
| 574 | RM25_RS02775 | RM25_0565 | 30S ribosomal protein S13 | 601 | gene | 667273 | 667647 | + | 2682,392 | 1,041928976 | 0,539036 | 1,93295 | 0,053242 | 0,12924 | 3340,818235 | 0,306659452 | 0,215197 | 1,425017 | 0,154152 | 0,208344 | 2023,965 | 2,762621456 | 0,430645 | 6,41508 | 1,41E-10 | 4,67E-09 | 1878,421 | -2,57543 | 0,385934 | -6,67324 | 2,5E-11 | 5,13E-10 | 3528,083197 | -0,084987245 | 0,273785 | -0,31042 | 0,756244 | 0,799313 |
| 575 | RM25_RS02780 | RM25_0566 | 30S ribosomal protein S11 | 602 | gene | 667693 | 668100 | + | 1931,656 | 0,9169688 | 0,477454 | 1,920538 | 0,05479 | 0,132045 | 2429,442494 | 0,278687079 | 0,200533 | 1,389731 | 0,164611 | 0,219574 | 1433,869 | 2,317109235 | 0,356057 | 6,507683 | 7,63E-11 | 2,65E-09 | 1430,835 | -2,2517 | 0,295373 | -7,62327 | 2,47E-14 | 8,1E-13 | 2468,669376 | -0,174864015 | 0,258059 | -0,67761 | 0,498017 | 0,559231 |
| 576 | RM25_RS02785 | RM25_0567 | 30S ribosomal protein S4 | 603 | gene | 668131 | 668736 | + | 3389,575 | 0,870435082 | 0,516245 | 1,686088 | 0,091779 | 0,182889 | 4379,009966 | 0,226581056 | 0,197108 | 1,149529 | 0,250338 | 0,315196 | 2400,14 | 2,415723254 | 0,393778 | 6,134742 | 8,53E-10 | 2,42E-08 | 2565,161 | -2,47 | 0,323292 | -7,64017 | 2,17E-14 | 7,41E-13 | 4281,084971 | -0,244002124 | 0,270761 | -0,90117 | 0,367496 | 0,432446 |
| 577 | RM25_RS02790 | RM25_0568 | DNA-directed RNA polymerase, alpha subunit | 604 | gene | 668861 | 669880 | + | 5015,817 | 0,983004793 | 0,563132 | 1,745602 | 0,08088 | 0,168108 | 6948,477584 | 0,460403926 | 0,196032 | 2,348615 | 0,018843 | 0,034034 | 3083,156 | 2,56635504 | 0,367386 | 6,985448 | 2,84E-12 | 1,45E-10 | 3608,137 | -2,77201 | 0,292425 | -9,47941 | 2,56E-21 | 2,42E-19 | 6515,947759 | -0,627762373 | 0,264117 | -2,37683 | 0,017462 | 0,029005 |
| 578 | RM25_RS02795 | RM25_0569 | 50S ribosomal protein L17 | 605 | gene | 669924 | 670511 | + | 3996,226 | 1,115376954 | 0,64298 | 1,734699 | 0,082794 | 0,170582 | 5578,102506 | 0,519451203 | 0,214221 | 2,424838 | 0,015315 | 0,028604 | 2414,35 | 3,23807742 | 0,390955 | 8,282483 | 1,21E-16 | 1,18E-14 | 2710,001 | -3,36422 | 0,365231 | -9,21123 | 3,22E-20 | 2,53E-18 | 5352,34836 | -0,605926165 | 0,242379 | -2,49991 | 0,012423 | 0,021357 |
| 579 | RM25_RS02800 | RM25_0570 | Glycosyltransferase, group 1 family | 606 | gene | 670552 | 671793 | + | 657,0224 | -0,533748462 | 0,361687 | -1,47572 | 0,140019 | 0,240193 | 413,1911804 | -0,458751348 | 0,173432 | -2,64513 | 0,008166 | 0,016678 | 900,8537 | -0,570348661 | 0,255873 | -2,22903 | 0,025812 | 0,062144 | 815,1711 | 1,107897 | 0,223399 | 4,959271 | 7,08E-07 | 5,52E-06 | 527,0094198 | 1,055011576 | 0,224292 | 4,703734 | 2,55E-06 | 8,29E-06 |
| 580 | RM25_RS02805 | RM25_0571 | Resuscitation-promoting factor RpfB | 607 | gene | 671903 | 672988 | - | 22212,95 | 1,668871462 | 0,520821 | 3,204311 | 0,001354 | 0,012087 | 29152,20172 | 1,214974127 | 0,191733 | 6,336803 | 2,35E-10 | 2,19E-09 | 15273,7 | 2,857986412 | 0,469493 | 6,087394 | 1,15E-09 | 3,15E-08 | 11374,49 | -2,29182 | 0,436965 | -5,24485 | 1,56E-07 | 1,4E-06 | 33046,93637 | -0,621580318 | 0,248755 | -2,49876 | 0,012463 | 0,021379 |
| 581 | RM25_RS02810 | RM25_0572 | Superoxide dismutase | 608 | gene | 673283 | 673993 | - | 27148,83 | -0,63558987 | 0,310028 | -2,0501 | 0,040354 | 0,109562 | 28144,19198 | -0,469541512 | 0,256008 | -1,83409 | 0,066641 | 0,099983 | 26153,46 | -0,819016312 | 0,489285 | -1,6739 | 0,09415 | 0,164256 | 35073,15 | -0,01899 | 0,487008 | -0,03899 | 0,968898 | 0,975937 | 20806,11306 | -0,33380979 | 0,274644 | -1,21543 | 0,224204 | 0,280344 |
| 582 | RM25_RS02815 | RM25_0573 | ABC transporter domain-containing ATP-binding subunit | 609 | gene | 674089 | 675129 | + | 675,7496 | -0,071742007 | 0,124459 | -0,57643 | 0,564324 | 0,658472 | 657,3727631 | 0,044242186 | 0,15168 | 0,29168 | 0,770531 | 0,81551 | 694,1265 | -0,179349081 | 0,230911 | -0,7767 | 0,437335 | 0,541385 | 729,1124 | 0,126069 | 0,176306 | 0,715059 | 0,474572 | 0,546565 | 646,3261221 | -0,043948309 | 0,186669 | -0,23543 | 0,813872 | 0,848428 |
| 583 | RM25_RS02820 | RM25_0574 | Putative integral membrane transport protein | 610 | gene | 675099 | 676691 | + | 852,228 | -0,000990548 | 0,170442 | -0,00581 | 0,995363 | 0,998328 | 783,6345324 | 0,128146569 | 0,133588 | 0,959265 | 0,337425 | 0,405771 | 920,8214 | -0,109694465 | 0,291802 | -0,37592 | 0,706975 | 0,780853 | 896,2415 | 0,287886 | 0,213467 | 1,34862 | 0,177459 | 0,238537 | 836,7331119 | 0,112279302 | 0,226232 | 0,496302 | 0,619681 | 0,676366 |
| 584 | RM25_RS02825 | RM25_0575 | Substrate-specific component CbrT of predicted cobalamin E | 611 | gene | 676706 | 677557 | - | 639,6548 | -0,489984589 | 0,208403 | -2,35113 | 0,018716 | 0,069362 | 554,8316602 | -0,588807696 | 0,183491 | -3,20892 | 0,001332 | 0,003447 | 724,478 | -0,416703805 | 0,294429 | -1,41529 | 0,156982 | 0,2438 | 786,8399 | 0,246767 | 0,232237 | 1,062565 | 0,287979 | 0,361816 | 522,5132396 | 0,484064986 | 0,256548 | 1,886838 | 0,059182 | 0,086 |
| 585 | RM25_RS02830 | RM25_0576 | Cobalt ABC transporter, ATP-binding protein | 612 | gene | 677554 | 679170 | - | 1523,112 | 0,142955854 | 0,185521 | 0,770564 | 0,440966 | 0,549023 | 1762,69405 | 0,12004153 | 0,142759 | 0,840868 | 0,400422 | 0,469783 | 1283,53 | 0,175536044 | 0,214216 | 0,819436 | 0,412538 | 0,517759 | 1532,484 | -0,54928 | 0,198178 | -2,77166 | 0,005577 | 0,014511 | 1567,887612 | -0,437770687 | 0,134503 | -3,25472 | 0,001135 | 0,002368 |

| | | | | | | | | | | | | | | | | | | | | | | | | | | | | | | | | | | | | | | |
|---|---|---|---|---|---|---|---|---|---|---|---|---|---|---|---|---|---|---|---|---|---|---|---|---|---|---|---|---|---|---|---|---|---|---|---|---|---|---|
| 586 | RM25_RS02835 | RM25_0577 | Cobalt transport protein | 613 | gene | 679167 | 680267 | - | 1003,659 | 0,306726172 | 0,205194 | 1,494814 | 0,134963 | 0,233903 | 1136,116808 | 0,037470688 | 0,164441 | 0,227867 | 0,819749 | 0,856923 | 871,2011 | 0,666686812 | 0,210805 | 3,162583 | 0,001564 | 0,007047 | 952,3527 | -0,79761 | 0,21547 | -3,70171 | 0,000214 | 0,000949 | 1087,510033 | -0,114756062 | 0,132689 | -0,86485 | 0,387123 | 0,452155 |
| 587 | RM25_RS02840 | RM25_0578 | Hypothetical protein | 614 | gene | 680313 | 680855 | - | 1898,416 | 0,807897993 | 0,387559 | 2,084578 | 0,037108 | 0,103875 | 2272,372316 | 0,225463495 | 0,221856 | 1,016259 | 0,309506 | 0,376811 | 1524,46 | 1,841468589 | 0,298497 | 6,169128 | 6,87E-10 | 2E-08 | 1471,499 | -1,71142 | 0,308076 | -5,55518 | 2,77E-08 | 2,87E-07 | 2364,455202 | -0,05275948 | 0,192865 | -0,27356 | 0,784425 | 0,824661 |
| 588 | RM25_RS02845 | RM25_0579 | YbxB protein | 615 | gene | 681147 | 681746 | - | 390,6083 | -0,14410026 | 0,177574 | -0,8115 | 0,417081 | 0,526545 | 420,1875778 | 0,072808291 | 0,166758 | 0,43661 | 0,662395 | 0,717163 | 361,029 | -0,394288088 | 0,274647 | -1,43562 | 0,151111 | 0,236185 | 432,6945 | -0,06104 | 0,191558 | -0,31866 | 0,749986 | 0,79341 | 364,9138743 | -0,46573652 | 0,256864 | -1,81316 | 0,069806 | 0,099778 |
| 589 | RM25_RS02850 | RM25_0580 | TRNA pseudouridine synthase A | 616 | gene | 681784 | 682659 | - | 322,2177 | -0,002065348 | 0,251914 | -0,0082 | 0,993459 | 0,996842 | 272,4492626 | 0,101189852 | 0,184295 | 0,549064 | 0,582962 | 0,644787 | 371,9862 | -0,078852677 | 0,377237 | -0,20903 | 0,834427 | 0,879159 | 338,3339 | 0,47836 | 0,239972 | 1,993401 | 0,046218 | 0,079806 | 317,0501741 | 0,364067058 | 0,34198 | 1,064586 | 0,287063 | 0,347692 |
| 590 | RM25_RS02855 | RM25_0581 | TRNA (guanine-N(7)-)-methyltransferase | 617 | gene | 682652 | 683506 | - | 582,0753 | 0,204015903 | 0,195725 | 1,042361 | 0,297244 | 0,407328 | 668,2054529 | 0,402133055 | 0,14149 | 2,842133 | 0,004481 | 0,00989 | 495,9451 | -0,057208596 | 0,246039 | -0,23252 | 0,816136 | 0,867673 | 571,97 | -0,25158 | 0,189781 | -1,32563 | 0,184961 | 0,24658 | 612,112147 | -0,648562222 | 0,20578 | -3,15173 | 0,001623 | 0,003272 |
| 591 | RM25_RS02860 | RM25_0582 | PF03733 domain protein | 618 | gene | 683661 | 684056 | + | 335,4247 | 0,602814998 | 0,192905 | 3,124939 | 0,001778 | 0,014109 | 339,5157852 | 0,347443648 | 0,212022 | 1,638712 | 0,101273 | 0,143882 | 331,3337 | 0,872877971 | 0,296932 | 2,939657 | 0,003286 | 0,012994 | 282,6906 | -0,4056 | 0,287017 | -1,41315 | 0,157613 | 0,215883 | 395,1874178 | 0,159055519 | 0,21456 | 0,741309 | 0,458506 | 0,523595 |
| 592 | RM25_RS02865 | RM25_0583 | Elongation factor G fusA2 | 619 | gene | 684188 | 686284 | - | 3211,241 | -1,039647104 | 0,430838 | -2,41308 | 0,015818 | 0,062873 | 2128,377231 | 0,429754689 | 0,132091 | 3,253462 | 0,00114 | 0,003019 | 4294,104 | -1,954550079 | 0,3193 | -6,12135 | 9,28E-10 | 2,59E-08 | 4530,253 | 1,864429 | 0,324754 | 5,741051 | 9,41E-09 | 1,05E-07 | 2060,585754 | -0,480945572 | 0,104407 | -4,60644 | 4,1E-06 | 1,29E-05 |
| 593 | RM25_RS02870 | RM25_0584 | 50S ribosomal protein L13 | 620 | gene | 686585 | 687028 | + | 7295,746 | 1,015032505 | 0,613686 | 1,653993 | 0,098129 | 0,190833 | 10699,91739 | 0,625003591 | 0,246805 | 2,532382 | 0,011329 | 0,021977 | 3891,575 | 2,397487303 | 0,48646 | 4,928434 | 8,29E-07 | 1,19E-05 | 5190,415 | -2,81165 | 0,493055 | -5,70251 | 1,18E-08 | 1,29E-07 | 9557,110081 | -1,006042545 | 0,23862 | -4,21609 | 2,49E-05 | 6,93E-05 |
| 594 | RM25_RS02875 | RM25_0585 | 30S ribosomal protein S9 | 621 | gene | 687025 | 687576 | + | 5665,111 | 1,068402684 | 0,669669 | 1,595418 | 0,110619 | 0,205136 | 8258,195226 | 0,57424936 | 0,226413 | 2,536292 | 0,011203 | 0,021832 | 3072,027 | 3,006992199 | 0,501664 | 5,994033 | 2,05E-09 | 5,3E-08 | 3926,297 | -3,33569 | 0,496701 | -6,71569 | 1,87E-11 | 4,08E-10 | 7513,133846 | -0,870505498 | 0,234201 | -3,71691 | 0,000202 | 0,000486 |
| 595 | RM25_RS02880 | RM25_0586 | Phosphoglucosamine mutase | 622 | gene | 687728 | 689071 | + | 2443,928 | 0,565799056 | 0,218405 | 2,590591 | 0,009581 | 0,045438 | 2728,907839 | 0,423387918 | 0,175933 | 2,406531 | 0,016105 | 0,029842 | 2158,948 | 0,750623647 | 0,316687 | 2,370239 | 0,017777 | 0,047131 | 2091,544 | -0,5855 | 0,293297 | -1,99627 | 0,045905 | 0,07944 | 2852,726099 | -0,222827472 | 0,195855 | -1,13772 | 0,255239 | 0,313007 |
| 596 | RM25_RS02885 | RM25_0587 | Pantothenate kinase | 623 | gene | 689210 | 690292 | - | 710,9518 | 0,314992261 | 0,225693 | 1,39567 | 0,162814 | 0,265538 | 865,1764499 | 0,426332345 | 0,12673 | 3,3641 | 0,000768 | 0,002117 | 556,7271 | 0,144526823 | 0,203457 | 0,710357 | 0,477483 | 0,579221 | 670,8286 | -0,54269 | 0,169503 | -3,20165 | 0,001366 | 0,004614 | 773,607653 | -0,768538197 | 0,142886 | -5,3787 | 7,5E-08 | 3,15E-07 |
| 597 | RM25_RS02890 | RM25_0588 | Glutamine--fructose-6-phosphate transaminase (isomerizing | 624 | gene | 690441 | 692291 | + | 4022,092 | 1,187868706 | 0,19968 | 5,948854 | 2,7E-09 | 2,65E-07 | 4041,200382 | 0,842683921 | 0,160889 | 5,23768 | 1,63E-07 | 9,48E-07 | 4002,984 | 1,572891332 | 0,283617 | 5,545835 | 2,93E-08 | 5,75E-07 | 2602,695 | -0,57575 | 0,241989 | -2,37923 | 0,017349 | 0,036576 | 5465,249414 | 0,193050947 | 0,194677 | 0,991645 | 0,321371 | 0,385089 |
| 598 | RM25_RS02895 | RM25_0589 | Holo-[acyl-carrier-protein] synthase | 625 | gene | 692332 | 692799 | + | 752,9186 | 1,05793546 | 0,20057 | 5,274636 | 1,33E-07 | 7,84E-06 | 825,2463577 | 1,031808003 | 0,199489 | 5,172268 | 2,31E-07 | 1,32E-06 | 680,5907 | 1,091647519 | 0,307231 | 3,553178 | 0,000381 | 0,00226 | 518,3965 | -0,36944 | 0,326246 | -1,1324 | 0,257468 | 0,328383 | 995,2921196 | -0,271972298 | 0,160291 | -1,69674 | 0,089745 | 0,125092 |
| 599 | RM25_RS02900 | RM25_0590 | YjeF protein | 626 | gene | 692801 | 694219 | + | 642,6467 | 0,815115517 | 0,166851 | 4,885301 | 1,03E-06 | 4,2E-05 | 577,2991533 | 0,928553286 | 0,158438 | 5,860682 | 4,61E-09 | 3,56E-08 | 707,9942 | 0,72746558 | 0,241664 | 3,010236 | 0,00261 | 0,010794 | 489,2982 | 0,362821 | 0,201786 | 1,798045 | 0,07217 | 0,11422 | 804,5221753 | 0,220639911 | 0,186572 | 1,182601 | 0,236967 | 0,294119 |
| 600 | RM25_RS02905 | RM25_0591 | Mannosyltransferase pimB | 627 | gene | 694248 | 695393 | - | 401,6094 | -0,818896041 | 0,49302 | -1,66098 | 0,096718 | 0,189496 | 228,5555684 | -1,440610505 | 0,283697 | -5,07799 | 3,81E-07 | 2,05E-06 | 574,6632 | -0,5962 | 0,3935 | -1,51512 | 0,129742 | 0,211189 | 537,0721 | 0,983932 | 0,270794 | 3,633504 | 0,00028 | 0,001205 | 286,4978388 | 1,897480963 | 0,407049 | 4,661557 | 3,14E-06 | 1E-05 |
| 601 | RM25_RS13145 | RM25_0592 | Hypothetical protein | 628 | gene | 695671 | 698802 | + | 2437,915 | 0,092729764 | 0,124523 | 0,744679 | 0,456466 | 0,561236 | 2386,711652 | -0,055359762 | 0,13496 | -0,41019 | 0,681664 | 0,735403 | 2489,117 | 0,234943554 | 0,228841 | 1,026669 | 0,304577 | 0,408819 | 2489,88 | -0,15276 | 0,197841 | -0,77212 | 0,440044 | 0,515429 | 2467,507522 | 0,195271896 | 0,149141 | 1,309311 | 0,190429 | 0,244068 |
| 602 | RM25_RS11790 | RM25_0593 | Hydrolase, P-loop family | 629 | gene | 699033 | 700028 | + | 702,9912 | 0,026223225 | 0,149883 | 0,174958 | 0,861113 | 0,907665 | 717,0983063 | 0,27244975 | 0,127444 | 2,137794 | 0,032533 | 0,054851 | 688,884 | -0,227310988 | 0,257425 | -0,88302 | 0,377227 | 0,482767 | 734,6738 | 0,130217 | 0,161783 | 0,804887 | 0,420885 | 0,496013 | 696,9809142 | -0,30883256 | 0,230024 | -1,34261 | 0,179397 | 0,231186 |
| 603 | RM25_RS02925 | RM25_0594 | Universal bacterial protein YeaZ | 630 | gene | 700025 | 700711 | + | 304,2207 | 0,010174142 | 0,239315 | 0,042514 | 0,966089 | 0,978544 | 255,9076956 | 0,176948047 | 0,212744 | 0,831742 | 0,405554 | 0,474749 | 352,5336 | -0,112839282 | 0,324348 | -0,3479 | 0,727918 | 0,798373 | 318,11 | 0,546166 | 0,248557 | 2,19999 | 0,027808 | 0,053088 | 300,3153307 | 0,31930372 | 0,297471 | 1,073395 | 0,283094 | 0,344299 |
| 604 | RM25_RS02930 | RM25_0595 | Ribosomal-protein-alanine acetyltransferase | 631 | gene | 700708 | 701229 | + | 170,0011 | -0,067387271 | 0,286571 | -0,23515 | 0,814092 | 0,870772 | 127,8684335 | 0,168844843 | 0,233883 | 0,72192 | 0,470344 | 0,538417 | 212,1339 | -0,210208323 | 0,296976 | -0,70783 | 0,479051 | 0,580284 | 182,3234 | 0,860604 | 0,195241 | 4,407904 | 1,04E-05 | 6,36E-05 | 163,2081735 | 0,540947897 | 0,319838 | 1,691318 | 0,090776 | 0,126155 |
| 605 | RM25_RS02935 | RM25_0596 | putative tRNA threonylcarbamoyladenosine biosynthesis prc | 632 | gene | 701226 | 702299 | + | 497,6322 | 0,092105558 | 0,160585 | 0,573563 | 0,566263 | 0,66008 | 482,295049 | 0,351551467 | 0,147231 | 2,38775 | 0,016952 | 0,031191 | 512,9693 | -0,147605071 | 0,268968 | -0,54878 | 0,583155 | 0,674433 | 506,6813 | 0,287691 | 0,162136 | 1,774383 | 0,076 | 0,118341 | 504,5620219 | -0,154253609 | 0,243373 | -0,63381 | 0,526202 | 0,586687 |
| 606 | RM25_RS02940 | RM25_0597 | Cob(II)yrinic acid a, c-diamide reductase | 633 | gene | 702373 | 704253 | + | 2423,02 | -0,278861754 | 0,20966 | -1,33007 | 0,183496 | 0,288333 | 2850,896281 | -0,175870521 | 0,178896 | -0,98309 | 0,325564 | 0,393498 | 1995,144 | -0,425770571 | 0,183159 | -2,3246 | 0,020094 | 0,051423 | 2813,168 | -0,45557 | 0,223651 | -2,03696 | 0,041654 | 0,073267 | 2148,6856 | -0,658880878 | 0,13915 | -4,73505 | 2,19E-06 | 7,19E-06 |
| 607 | RM25_RS02945 | RM25_0598 | Bleomycin hydrolase | 634 | gene | 704437 | 705885 | + | 1263,163 | 0,154990916 | 0,082693 | 1,874298 | 0,060889 | 0,140979 | 1295,634624 | 0,209057944 | 0,145559 | 1,436243 | 0,150933 | 0,204707 | 1230,692 | 0,09881848 | 0,150585 | 0,65623 | 0,511676 | 0,612193 | 1263,384 | -0,07494 | 0,140711 | -0,5326 | 0,594308 | 0,655797 | 1304,384834 | -0,135389022 | 0,145136 | -0,93284 | 0,350901 | 0,414573 |
| 608 | RM25_RS02950 | RM25_0599 | Hydrolase, NUDIX family | 635 | gene | 705940 | 706425 | + | 453,9876 | -0,227585066 | 0,133411 | -1,7059 | 0,088027 | 0,178095 | 429,1704843 | -0,245205937 | 0,201449 | -1,21721 | 0,223525 | 0,286175 | 478,8048 | -0,212652849 | 0,201283 | -1,05649 | 0,290745 | 0,394202 | 516,2544 | 0,079534 | 0,194704 | 0,408489 | 0,682915 | 0,73165 | 410,3394029 | 0,169030362 | 0,206537 | 0,818404 | 0,413127 | 0,478731 |
| 609 | RM25_RS02955 | RM25_0600 | Tetracycline resistance determinant TetV | 636 | gene | 706569 | 707795 | + | 695,9277 | -0,393869138 | 0,199885 | -1,97047 | 0,048784 | 0,122846 | 614,7590798 | -0,655193085 | 0,179968 | -3,6406 | 0,000272 | 0,000846 | 777,0964 | -0,194284777 | 0,25827 | -0,75225 | 0,451898 | 0,555621 | 833,4706 | 0,07806 | 0,189284 | 0,412397 | 0,680049 | 0,730239 | 590,3501859 | 0,598651519 | 0,243318 | 2,460363 | 0,01388 | 0,023536 |
| 610 | RM25_RS02960 | RM25_0601 | Putative SAM-dependent methyltransferase | 637 | gene | 707820 | 709184 | + | 335,7427 | -0,258740613 | 0,342154 | -0,75621 | 0,449522 | 0,55633 | 239,2717015 | -0,670818453 | 0,20463 | -3,2782 | 0,001045 | 0,002798 | 432,2137 | -0,039989749 | 0,346597 | -0,11538 | 0,908145 | 0,936285 | 384,2973 | 0,514263 | 0,214097 | 2,402015 | 0,016305 | 0,035033 | 300,9464303 | 1,209295103 | 0,337127 | 3,587059 | 0,000334 | 0,000773 |
| 611 | RM25_RS02965 | RM25_0602 | ABC transporter, permease protein OpuCB | 638 | gene | 709403 | 710044 | + | 614,5934 | -0,260615749 | 0,11093 | -2,34938 | 0,018805 | 0,069555 | 621,0307481 | -0,279727825 | 0,152916 | -1,82929 | 0,067356 | 0,100991 | 608,1561 | -0,239559235 | 0,221835 | -1,0799 | 0,280187 | 0,383731 | 706,5628 | -0,11267 | 0,208554 | -0,54025 | 0,589026 | 0,650579 | 548,5761542 | -0,016889715 | 0,140578 | -0,12015 | 0,904368 | 0,922769 |
| 612 | RM25_RS02970 | RM25_0603 | Osmoprotectant transport ATP-binding protein ABC transpo | 639 | gene | 710041 | 711375 | + | 798,5829 | 0,014209832 | 0,19261 | 0,073775 | 0,941189 | 0,960755 | 862,6215638 | 0,091368303 | 0,178916 | 0,510676 | 0,609578 | 0,668578 | 734,5443 | -0,075060438 | 0,317934 | -0,23609 | 0,813364 | 0,865899 | 838,1788 | -0,2108 | 0,222483 | -0,94747 | 0,343401 | 0,418725 | 789,0866082 | -0,313717763 | 0,275204 | -1,13995 | 0,254308 | 0,31219 |
| 613 | RM25_RS02975 | RM25_0604 | ProWL | 640 | gene | 711372 | 712109 | + | 356,7518 | -0,227705245 | 0,156845 | -1,45179 | 0,14656 | 0,247275 | 384,6656309 | -0,179445322 | 0,150237 | -1,19442 | 0,232315 | 0,295343 | 328,838 | -0,282261799 | 0,277062 | -1,01877 | 0,308313 | 0,412192 | 406,4576 | -0,24215 | 0,206374 | -1,17335 | 0,240655 | 0,309282 | 323,0541366 | -0,283024462 | 0,237503 | -1,19167 | 0,233393 | 0,289988 |
| 614 | RM25_RS02980 | RM25_0605 | ProXL | 641 | gene | 712113 | 713144 | + | 718,6233 | -0,040658157 | 0,243975 | -0,16665 | 0,867646 | 0,910121 | 888,153918 | 0,011921806 | 0,129562 | 0,092016 | 0,926685 | 0,944314 | 549,0928 | -0,124530785 | 0,21269 | -0,5855 | 0,558209 | 0,653601 | 771,8723 | -0,68752 | 0,185384 | -3,70862 | 0,000208 | 0,000929 | 695,1491264 | -0,769611128 | 0,128404 | -5,99368 | 2,05E-09 | 1,09E-08 |
| 615 | RM25_RS02985 | RM25_0606 | 10 kDa chaperonin | 642 | gene | 713381 | 713677 | + | 4334,841 | 0,86690014 | 0,747082 | 1,160381 | 0,245894 | 0,354792 | 7065,5538 | 0,593008705 | 0,205615 | 2,884079 | 0,003926 | 0,008837 | 1604,129 | 2,416557779 | 0,582745 | 4,146856 | 3,37E-05 | 0,000291 | 3296,41 | -3,52304 | 0,556789 | -6,32743 | 2,49E-10 | 4,17E-09 | 5489,946264 | -1,669686307 | 0,271801 | -6,14305 | 8,1E-10 | 4,58E-09 |
| 616 | RM25_RS02990 | RM25_0607 | 60 kDa chaperonin 1 | 643 | gene | 713699 | 715291 | + | 22561,17 | 0,88466024 | 0,775667 | 1,140515 | 0,254072 | 0,362159 | 37747,78024 | 0,669839999 | 0,209863 | 3,191794 | 0,001414 | 0,003622 | 7374,568 | 2,247328041 | 0,619678 | 3,626607 | 0,000287 | 0,001786 | 17008,36 | -3,55176 | 0,596706 | -5,95228 | 2,64E-09 | 3,35E-08 | 28704,37277 | -1,945507872 | 0,269703 | -7,21352 | 5,45E-13 | 4,89E-12 |
| 617 | RM25_RS02995 | RM25_0608 | Inosine-5'-monophosphate dehydrogenase | 644 | gene | 715438 | 716958 | + | 6964,646 | 1,510268105 | 0,321354 | 4,699695 | 2,61E-06 | 9,03E-05 | 6901,587062 | 0,870390302 | 0,202841 | 4,290992 | 1,78E-05 | 7,17E-05 | 7027,706 | 2,310902523 | 0,37125 | 6,224645 | 4,83E-10 | 1,5E-08 | 3855,411 | -1,10162 | 0,322249 | -3,41855 | 0,00063 | 0,002397 | 10056,15197 | 0,3708384 | 0,256452 | 1,446032 | 0,148168 | 0,196308 |
| 618 | RM25_RS03000 | RM25_0609 | IMP dehydrogenase family protein | 645 | gene | 717110 | 718213 | + | 1743,381 | 0,340643168 | 0,225749 | 1,508947 | 0,131312 | 0,229943 | 2027,216337 | 0,40801772 | 0,184363 | 2,213124 | 0,026889 | 0,04643 | 1459,545 | 0,249105056 | 0,2982 | 0,835362 | 0,403514 | 0,509417 | 1632,056 | -0,43846 | 0,300077 | -1,46117 | 0,143969 | 0,200434 | 1907,176115 | -0,559002828 | 0,169551 | -3,29695 | 0,000977 | 0,002077 |
| 619 | RM25_RS03005 | RM25_0610 | Chorismate mutase | 646 | gene | 718245 | 718574 | + | 585,7115 | 0,262919106 | 0,108398 | 2,425508 | 0,015287 | 0,061278 | 582,2731346 | 0,304710434 | 0,151055 | 2,017218 | 0,043673 | 0,07104 | 589,1499 | 0,223111524 | 0,219617 | 1,015913 | 0,309671 | 0,413537 | 563,3622 | -0,0006 | 0,198481 | -0,00304 | 0,997574 | 0,99817 | 625,3412972 | -0,032512389 | 0,168678 | -0,19275 | 0,847156 | 0,87692 |
| 620 | RM25_RS03010 | RM25_0611 | Transcription initiation factor | 647 | gene | 718878 | 720194 | + | 1449,306 | -0,17836475 | 0,184515 | -0,96667 | 0,333711 | 0,44441 | 1309,214035 | -0,265664305 | 0,153694 | -1,72853 | 0,083893 | 0,12206 | 1589,399 | -0,106649296 | 0,298296 | -0,35753 | 0,720696 | 0,792295 | 1625,667 | 0,151614 | 0,269217 | 0,563167 | 0,573321 | 0,637113 | 1328,7204 | 0,349381498 | 0,183004 | 1,909149 | 0,056243 | 0,082185 |
| 621 | RM25_RS03015 | RM25_0612 | Hypothetical protein | 648 | gene | 720219 | 721931 | + | 920,2003 | -0,260254025 | 0,257 | -1,01266 | 0,311221 | 0,420131 | 740,8301605 | -0,766558534 | 0,158578 | -4,83395 | 1,34E-06 | 6,64E-06 | 1099,57 | 0,067063255 | 0,197524 | 0,33952 | 0,734218 | 0,802668 | 1057,162 | 0,140477 | 0,169577 | 0,8284 | 0,407444 | 0,483558 | 819,1150964 | 1,026052764 | 0,158519 | 6,472762 | 9,62E-11 | 6,3E-10 |
| 622 | RM25_RS03020 | RM25_0613 | Hypothetical protein | 649 | gene | 721944 | 723161 | + | 361,9274 | -0,016775174 | 0,408052 | -0,04111 | 0,967208 | 0,97873 | 231,2108292 | -0,777124072 | 0,230197 | -3,37591 | 0,000736 | 0,002043 | 492,644 | 0,328796582 | 0,310387 | 1,059313 | 0,289457 | 0,393682 | 382,057 | 0,516022 | 0,273863 | 1,884231 | 0,059534 | 0,097921 | 353,4907982 | 1,684682886 | 0,265971 | 6,334075 | 2,39E-10 | 1,44E-09 |
| 623 | RM25_RS03025 | RM25_0614 | AmiC, N-acetylmuramoyl-L-alanine amidase | 650 | gene | 723229 | 724068 | + | 347,4281 | -0,604888744 | 0,320085 | -1,88978 | 0,058788 | 0,137599 | 255,6104414 | -0,816426382 | 0,230474 | -3,54238 | 0,000397 | 0,001186 | 439,2457 | -0,488953485 | 0,335261 | -1,45842 | 0,144724 | 0,229397 | 440,6107 | 0,586438 | 0,263413 | 2,226305 | 0,025994 | 0,050219 | 270,9832035 | 0,979934132 | 0,320295 | 3,059473 | 0,002217 | 0,004388 |
| 624 | RM25_RS03030 | RM25_0615 | Serine acetyltransferase | 651 | gene | 724103 | 724654 | - | 1827,884 | 0,683948081 | 0,301362 | 2,269521 | 0,023237 | 0,078018 | 2258,611284 | 0,690418439 | 0,220371 | 3,132983 | 0,00173 | 0,00432 | 1397,157 | 0,675037181 | 0,359785 | 1,876226 | 0,060624 | 0,117028 | 1490,715 | -0,73283 | 0,359384 | -2,03913 | 0,041437 | 0,072995 | 2205,553875 | -0,714555196 | 0,217177 | -3,29019 | 0,001001 | 0,002122 |
| 625 | RM25_RS03035 | RM25_0616 | Cysteine synthase | 652 | gene | 724651 | 725577 | - | 511,7319 | -0,218169784 | 0,383931 | -0,56825 | 0,569864 | 0,661986 | 310,726301 | -0,489697203 | 0,193731 | -2,52772 | 0,011481 | 0,022217 | 712,7375 | -0,10295398 | 0,190091 | -0,5416 | 0,588092 | 0,677484 | 577,5269 | 0,966633 | 0,198644 | 4,866147 | 1,14E-06 | 8,6E-06 | 463,8083646 | 1,405604457 | 0,183362 | 7,665744 | 1,78E-14 | 2,01E-13 |
| 626 | RM25_RS13160 | RM25_0617 | Ribulokinase | 653 | gene | 726024 | 726908 | + | 258,4671 | -1,554036012 | 0,62973 | -2,46778 | 0,013595 | 0,057324 | 142,1393906 | -3,030899816 | 0,417741 | -7,25546 | 4E-13 | 5,58E-12 | 374,7949 | -1,161669925 | 0,364123 | -3,19032 | 0,001421 | 0,0065 | 404,857 | 0,961191 | 0,362488 | 2,65165 | 0,00801 | 0,019564 | 129,2647855 | 2,902626654 | 0,464937 | 6,24306 | 4,29E-10 | 2,5E-09 |
| 627 | RM25_RS13165 | RM25_0618 | Ribulokinase domain protein | 654 | gene | 726905 | 727549 | + | 152,2884 | -1,775929345 | 0,795945 | -2,23122 | 0,025667 | 0,082377 | 84,69571627 | -4,637463798 | 0,551547 | -8,4081 | 4,17E-17 | 9,18E-16 | 219,8812 | -1,236259914 | 0,494502 | -2,50001 | 0,012419 | 0,036049 | 247,631 | 0,847708 | 0,397541 | 2,132377 | 0,032976 | 0,061104 | 68,06257477 | 4,332073783 | 0,683098 | 6,341808 | 2,27E-10 | 1,37E-09 |
| 628 | RM25_RS03045 | RM25_0619 | Putative L-ribulose-5-phosphate 4-epimerase | 655 | gene | 727546 | 728193 | + | 245,4388 | -1,778769688 | 0,8258 | -2,15399 | 0,031241 | 0,092882 | 135,4437202 | -4,933999683 | 0,543549 | -9,07738 | 1,11E-19 | 3,21E-18 | 355,4339 | -1,226733853 | 0,432261 | -2,83795 | 0,00454 | 0,016592 | 398,9891 | 0,850559 | 0,449745 | 1,891204 | 0,058597 | 0,096731 | 109,351284 | 4,635775062 | 0,585346 | 7,919716 | 2,38E-15 | 3,13E-14 |
| 629 | NA | RM25_0620 | Hypothetical protein | 656 | gene | 728153 | 728251 | + | 15,51821 | -2,074398604 | 0,899575 | -2,30598 | 0,021112 | 0,073611 | 8,553453122 | -6,326964691 | 1,474266 | -4,2916 | 1,77E-05 | 7,16E-05 | 22,48297 | -1,497416552 | 0,41579 | -3,60138 | 0,000317 | 0,001938 | 26,42686 | 0,861412 | 0,410683 | 2,09751 | 0,035948 | 0,065632 | 5,764834827 | 5,775199155 | 1,664552 | 3,469522 | 0,000521 | 0,001168 |
| 630 | RM25_RS03050 | RM25_0621 | L-arabinose isomerase | 657 | gene | 728321 | 729841 | + | 339,1497 | -1,576297229 | 0,745814 | -2,11353 | 0,034556 | 0,098965 | 183,6345883 | -3,95110036 | 0,441892 | -8,94133 | 3,85E-19 | 1,03E-17 | 494,6647 | -1,076041682 | 0,45952 | -2,34167 | 0,019198 | 0,049944 | 533,5939 | 0,8856 | 0,416036 | 2,128663 | 0,033282 | 0,061562 | 168,4228257 | 3,843313106 | 0,527552 | 7,285178 | 3,21E-13 | 2,99E-12 |
| 631 | RM25_RS03055 | RM25_0622 | Lactose operon repressor | 658 | gene | 729868 | 730905 | + | 406,5221 | -0,745171644 | 0,426794 | -1,74598 | 0,080815 | 0,168108 | 255,0748055 | -1,257320327 | 0,289797 | -4,33863 | 1,43E-05 | 5,89E-05 | 557,9694 | -0,529644552 | 0,3127 | -1,69378 | 0,090307 | 0,158965 | 533,5525 | 0,806316 | 0,333168 | 2,420151 | 0,015514 | 0,03364 | 298,4046694 | 1,59921489 | 0,277414 | 5,764722 | 8,18E-09 | 3,95E-08 |
| 632 | RM25_RS03060 | RM25_0623 | ABC transporter periplasmic-binding protein ytfQ | 659 | gene | 731149 | 732129 | + | 286,4903 | -1,33061103 | 0,402757 | -3,30376 | 0,000954 | 0,009713 | 198,7375382 | -1,802912767 | 0,303308 | -5,94417 | 2,78E-09 | 2,23E-08 | 374,2431 | -1,108634653 | 0,360931 | -3,0716 | 0,002129 | 0,009108 | 430,6248 | 0,661817 | 0,294475 | 2,247448 | 0,024611 | 0,048287 | 160,4382229 | 1,424461471 | 0,396143 | 3,595824 | 0,000323 | 0,000752 |
| 633 | RM25_RS13170 | RM25_0624 | ATP binding protein of ABC transporter | 660 | gene | 732267 | 732620 | + | 107,2146 | -1,636347099 | 0,584153 | -2,80123 | 0,005091 | 0,030098 | 63,14727413 | -2,744547083 | 0,535642 | -5,12384 | 2,99E-07 | 1,66E-06 | 151,282 | -1,292033806 | 0,376148 | -3,43491 | 0,000593 | 0,003203 | 170,0429 | 0,901588 | 0,438573 | 2,055732 | 0,039808 | 0,070442 | 51,3205005 | 2,41969471 | 0,540634 | 4,475658 | 7,62E-06 | 2,28E-05 |
| 634 | RM25_RS13175 | RM25_0625 | YtfR | 661 | gene | 732714 | 733799 | + | 466,8993 | -1,29059781 | 0,393298 | -3,28147 | 0,001033 | 0,010184 | 335,1751521 | -1,701983983 | 0,317941 | -5,35315 | 8,64E-08 | 5,36E-07 | 598,6234 | -1,082886126 | 0,377069 | -2,87185 | 0,004081 | 0,015389 | 695,5413 | 0,596627 | 0,368922 | 1,617216 | 0,105832 | 0,155494 | 266,6111033 | 1,287286988 | 0,355337 | 3,622717 | 0,000292 | 0,000684 |
| 635 | RM25_RS03070 | RM25_0626 | YtfT | 662 | gene | 733796 | 734854 | + | 323,1079 | -1,171776775 | 0,367239 | -3,19078 | 0,001419 | 0,012433 | 237,0867911 | -1,509207559 | 0,292743 | -5,1554 | 2,53E-07 | 1,43E-06 | 409,129 | -0,993187401 | 0,385371 | -2,57722 | 0,00996 | 0,030174 | 470,3236 | 0,566192 | 0,323506 | 1,750172 | 0,080089 | 0,123459 | 195,6248194 | 1,154094512 | 0,386001 | 2,989877 | 0,002791 | 0,00541 |
| 636 | RM25_RS03075 | RM25_0627 | ABC transporter, permease protein YjfF | 663 | gene | 734851 | 735879 | + | 318,5147 | -1,422962047 | 0,479662 | -2,9666 | 0,003011 | 0,020813 | 202,9709238 | -2,012969417 | 0,341787 | -5,88954 | 3,87E-09 | 3,05E-08 | 434,0584 | -1,185731489 | 0,438853 | -2,70189 | 0,006895 | 0,023229 | 487,1958 | 0,820408 | 0,356836 | 2,299119 | 0,021498 | 0,04342 | 170,7423901 | 1,723493872 | 0,460471 | 3,74289 | 0,000182 | 0,000442 |
| 637 | RM25_RS03080 | RM25_0628 | GMP synthase (glutamine-hydrolyzing) | 664 | gene | 736288 | 737727 | + | 3633,486 | 1,087351413 | 0,341976 | 3,179609 | 0,001475 | 0,012732 | 4661,567306 | 0,873976483 | 0,199134 | 4,388883 | 1,14E-05 | 4,72E-05 | 2605,405 | 1,502842749 | 0,260189 | 5,775957 | 7,65E-09 | 1,72E-07 | 2475,687 | -1,32988 | 0,259493 | -5,12492 | 2,98E-07 | 2,5E-06 | 4838,925648 | -0,6612572 | 0,183918 | -3,59539 | 0,000324 | 0,000753 |
| 638 | NA | RM25_0629 | Hypothetical protein | 665 | gene | 737616 | 737879 | - | 87,89781 | -0,81564404 | 0,344035 | -2,37082 | 0,017749 | 0,066827 | 67,09492473 | -0,988257936 | 0,34152 | -2,89371 | 0,003807 | 0,008637 | 108,7007 | -0,720659459 | 0,421272 | -1,71067 | 0,087141 | 0,15478 | 118,0919 | 0,531499 | 0,285109 | 1,8642 | 0,062294 | 0,101539 | 62,63918244 | 0,870103903 | 0,501591 | 1,73469 | 0,082796 | 0,116368 |
| 639 | RM25_RS03085 | RM25_0630 | YwqG | 666 | gene | 737932 | 738819 | + | 1199,7 | 0,318098539 | 0,181304 | 1,754508 | 0,079343 | 0,166529 | 1303,685835 | 0,024730711 | 0,145381 | 0,17011 | 0,864924 | 0,895312 | 1095,713 | 0,675670865 | 0,221406 | 3,051727 | 0,002275 | 0,009585 | 1131,442 | -0,67504 | 0,168957 | -3,99535 | 6,46E-05 | 0,000332 | 1302,760714 | 0,023241502 | 0,156195 | 0,148798 | 0,881713 | 0,902779 |
| 640 | RM25_RS03090 | RM25_0631 | NAD(P)H quinone oxidoreductase, PIG3 family | 667 | gene | 738870 | 739814 | - | 820,1554 | -0,857046295 | 0,379689 | -2,25723 | 0,023993 | 0,079316 | 551,5915457 | 0,172114556 | 0,156857 | 1,097272 | 0,272523 | 0,338606 | 1088,719 | -1,451966896 | 0,336363 | -4,31667 | 1,58E-05 | 0,000153 | 1109,608 | 1,565025 | 0,338508 | 4,623301 | 3,78E-06 | 2,55E-05 | 571,3924782 | -0,015103044 | 0,157472 | -0,09591 | 0,923593 | 0,937205 |
| 641 | RM25_RS03095 | RM25_0632 | putative stress-response transcriptional regulator protein Ps | 668 | gene | 740111 | 740401 | - | 1546,548 | 1,392830761 | 0,350337 | 3,975686 | 7,02E-05 | 0,001272 | 2051,247335 | 1,657386976 | 0,216285 | 7,662979 | 1,82E-14 | 2,91E-13 | 1041,848 | 0,928371952 | 0,382189 | 2,429092 | 0,015137 | 0,041485 | 905,4794 | -0,51134 | 0,362326 | -1,41126 | 0,158167 | 0,216492 | 2193,302123 | -1,207159482 | 0,238521 | -5,06101 | 4,17E-07 | 1,55E-06 |
| 642 | RM25_RS03100 | RM25_0633 | Sensor histidine kinase | 669 | gene | 740643 | 741986 | + | 988,4393 | 0,208555564 | 0,165883 | 1,257249 | 0,208664 | 0,313661 | 1087,662579 | 0,528092742 | 0,136482 | 3,869328 | 0,000109 | 0,000368 | 889,2161 | -0,171627453 | 0,156104 | -1,09944 | 0,271576 | 0,37433 | 968,53 | 0,023082 | 0,159515 | 0,144702 | 0,884946 | 0,908457 | 1040,071017 | -0,624793621 | 0,136474 | -4,57813 | 4,69E-06 | 1,46E-05 |
| 643 | RM25_RS03105 | RM25_0634 | Two component transcriptional regulator, LuxR family | 670 | gene | 741983 | 742681 | + | 839,9931 | 0,319762053 | 0,219424 | 1,457278 | 0,14504 | 0,245764 | 974,252473 | 0,540325224 | 0,164795 | 3,278774 | 0,001043 | 0,002796 | 705,7338 | 0,024508025 | 0,274544 | 0,089268 | 0,928869 | 0,952303 | 791,9564 | -0,23636 | 0,275664 | -0,85743 | 0,391207 | 0,466962 | 913,9613921 | -0,709906816 | 0,159027 | -4,46408 | 8,04E-06 | 2,38E-05 |
| 644 | RM25_RS03110 | RM25_0635 | Hypothetical protein | 671 | gene | 742918 | 743169 | - | 3024,142 | 0,734226748 | 0,281362 | 2,609544 | 0,009066 | 0,043611 | 3243,056533 | 0,678495704 | 0,221243 | 3,066743 | 0,002164 | 0,00528 | 2805,227 | 0,799977857 | 0,457533 | 1,748461 | 0,080384 | 0,146036 | 2410,775 | -0,3318 | 0,43566 | -0,76159 | 0,446303 | 0,521277 | 3689,603685 | -0,182643783 | 0,258263 | -0,7072 | 0,479442 | 0,541728 |
| 645 | RM25_RS03115 | RM25_0636 | Iron compound ABC transporter, iron compound-binding pr | 672 | gene | 743459 | 744475 | + | 512,0652 | 0,076516976 | 0,240743 | 0,317836 | 0,750609 | 0,819447 | 444,2308294 | 0,284778491 | 0,310064 | 0,918452 | 0,358382 | 0,427267 | 579,8995 | -0,082879272 | 0,212491 | -0,39004 | 0,696509 | 0,772915 | 524,2103 | 0,506714 | 0,298746 | 1,696136 | 0,08986 | 0,136031 | 515,9315095 | 0,202263492 | 0,273409 | 0,739783 | 0,459432 | 0,524397 |
| 646 | RM25_RS03120 | RM25_0637 | Ferric anguibactin transport system permease protein fatD | 673 | gene | 744472 | 745503 | + | 385,2903 | -0,339410179 | 0,183851 | -1,84612 | 0,064875 | 0,146165 | 349,8118721 | -0,311610459 | 0,245921 | -1,26712 | 0,205113 | 0,265487 | 420,7687 | -0,363356278 | 0,240241 | -1,51246 | 0,130416 | 0,21141 | 452,8099 | 0,226864 | 0,216226 | 1,049202 | 0,294085 | 0,368114 | 333,7410878 | 0,233386497 | 0,266435 | 0,875961 | 0,381051 | 0,446404 |
| 647 | RM25_RS03125 | RM25_0638 | ABC-type cobalamin/Fe3+-siderophores transport system, p | 674 | gene | 745496 | 746779 | + | 453,8875 | -0,771522807 | 0,372889 | -2,06904 | 0,038542 | 0,106699 | 323,3628447 | -0,677211443 | 0,390291 | -1,73514 | 0,082715 | 0,120644 | 584,4121 | -0,825429201 | 0,323734 | -2,54971 | 0,010781 | 0,032289 | 600,1708 | 0,841677 | 0,362223 | 2,323642 | 0,020145 | 0,041252 | 329,8021485 | 0,762118476 | 0,395761 | 1,925703 | 0,054142 | 0,079311 |
| 648 | RM25_RS03130 | RM25_0639 | Transposase InsI for insertion sequence element IS30C | 675 | gene | 746776 | 747702 | + | 0,733185 | 0,509677947 | 1,754999 | 0,290415 | 0,771499 | 0,836826 | 0,292244603 | -1,510987052 | 4,047733 | -0,37329 | 0,708931 | 0,759523 | 1,174126 | 1,513674977 | 1,702057 | 0,889321 | 0,37383 | 0,479912 | 0,663104 | -0,15106 | 1,834846 | -0,08233 | 0,934386 | 0,948062 | 0,84000742 | 2,99592508 | 3,952554 | 0,757972 | 0,448468 | 0,513375 |
| 649 | RM25_RS03135 | RM25_0640 | Putative iron-siderophore ABC transporter (ATP-binding pro | 676 | gene | 747720 | 748475 | + | 235,9487 | -1,42508571 | 0,469441 | -3,0357 | 0,0024 | 0,017566 | 157,5996389 | -2,140006762 | 0,370208 | -5,78056 | 7,45E-09 | 5,55E-08 | 314,2977 | -1,124937102 | 0,39313 | -2,86149 | 0,004217 | 0,01575 | 361,1878 | 0,67956 | 0,343102 | 1,980634 | 0,047632 | 0,081829 | 126,1293653 | 1,766426279 | 0,456668 | 3,868077 | 0,00011 | 0,000277 |
| 650 | RM25_RS03140 | RM25_0641 | Cardiolipin synthase 1 | 677 | gene | 748548 | 749801 | - | 1147,675 | 0,056681365 | 0,126057 | 0,449647 | 0,652965 | 0,737082 | 1261,189132 | 0,025470968 | 0,134552 | 0,189302 | 0,849856 | 0,882039 | 1034,161 | 0,095295475 | 0,166152 | 0,573544 | 0,566276 | 0,660749 | 1190,842 | -0,38273 | 0,150581 | -2,54168 | 0,011032 | 0,025543 | 1146,684772 | -0,261406932 | 0,120058 | -2,17733 | 0,029456 | 0,046502 |
| 651 | RM25_RS03145 | RM25_0642 | ATP-dependent DNA helicase PcrA | 678 | gene | 749938 | 752427 | + | 2067,974 | -0,071801736 | 0,178841 | -0,40148 | 0,688064 | 0,766128 | 2278,436881 | 0,174009932 | 0,119094 | 1,461115 | 0,143984 | 0,196622 | 1857,512 | -0,374853523 | 0,246059 | -1,52343 | 0,127651 | 0,208506 | 2236,786 | -0,09337 | 0,183497 | -0,50884 | 0,610861 | 0,669674 | 1981,072512 | -0,581518911 | 0,191068 | -3,04351 | 0,002338 | 0,004616 |
| 652 | RM25_RS03150 | RM25_0643 | Putative membrane protein | 679 | gene | 752521 | 753999 | + | 1858,217 | 0,281193137 | 0,30281 | 0,928614 | 0,353089 | 0,46579 | 2418,023171 | 0,324568064 | 0,185815 | 1,74673 | 0,080684 | 0,1179 | 1298,411 | 0,202387416 | 0,199964 | 1,012118 | 0,311482 | 0,41528 | 1779,13 | -0,88698 | 0,240653 | -3,68574 | 0,000228 | 0,001003 | 2000,068818 | -0,960460286 | 0,102615 | -9,35982 | 7,99E-21 | 2,01E-19 |
| 653 | RM25_RS03155 | RM25_0644 | Phosphoribosylglycinamide formyltransferase | 680 | gene | 754010 | 754621 | + | 787,3537 | 0,534281858 | 0,189748 | 2,815748 | 0,004866 | 0,029038 | 919,4262743 | 0,580445061 | 0,150816 | 3,848685 | 0,000119 | 0,000396 | 655,2812 | 0,471775893 | 0,206982 | 2,279313 | 0,022648 | 0,05643 | 680,5658 | -0,48554 | 0,191601 | -2,53409 | 0,011274 | 0,026001 | 912,9015293 | -0,542724087 | 0,127343 | -4,26192 | 2,03E-05 | 5,73E-05 |
| 654 | RM25_RS03160 | RM25_0645 | Bifunctional purine biosynthesis protein PurH | 681 | gene | 754743 | 756296 | + | 1714,219 | 0,4845045 | 0,253503 | 1,911237 | 0,055974 | 0,132999 | 2137,874846 | 0,515513037 | 0,147236 | 3,501281 | 0,000463 | 0,001357 | 1290,564 | 0,433974697 | 0,209412 | 2,072351 | 0,038233 | 0,083439 | 1516,709 | -0,74543 | 0,19904 | -3,74513 | 0,00018 | 0,000825 | 1959,869745 | -0,773166969 | 0,129039 | -5,99171 | 2,08E-09 | 1,1E-08 |
| 655 | RM25_RS03165 | RM25_0646 | Bifunctional protein FolD | 682 | gene | 756370 | 757230 | + | 1181,906 | 0,626070938 | 0,357511 | 1,751191 | 0,079913 | 0,167129 | 1630,292287 | 0,711914237 | 0,160571 | 4,43364 | 9,27E-06 | 3,91E-05 | 733,5203 | 0,441575272 | 0,179521 | 2,459736 | 0,013904 | 0,039242 | 988,0242 | -1,05135 | 0,203069 | -5,17731 | 2,25E-07 | 1,94E-06 | 1407,254151 | -1,269491157 | 0,104882 | -12,104 | 1,01E-33 | 7,19E-32 |
| 656 | RM25_RS03170 | RM25_0647 | Hypothetical protein | 683 | gene | 757236 | 757559 | + | 286,3689 | 0,691568505 | 0,300372 | 2,302374 | 0,021314 | 0,073808 | 363,6126733 | 1,00292121 | 0,24061 | 4,168238 | 3,07E-05 | 0,000118 | 209,1252 | 0,193714102 | 0,24031 | 0,806102 | 0,420184 | 0,524883 | 231,5079 | -0,36811 | 0,288612 | -1,27544 | 0,202153 | 0,266485 | 347,2683229 | -1,121592616 | 0,188286 | -5,95684 | 2,57E-09 | 1,36E-08 |
| 657 | RM25_RS03175 | RM25_0648 | Malate dehydrogenase | 684 | gene | 757741 | 758727 | - | 17788,54 | -0,275209795 | 0,375086 | -0,73372 | 0,463117 | 0,566165 | 22412,01877 | -0,127392436 | 0,207566 | -0,61374 | 0,539384 | 0,605395 | 13165,06 | -0,530493086 | 0,495003 | -1,0717 | 0,283856 | 0,388304 | 20715,99 | -0,63466 | 0,473113 | -1,34145 | 0,179774 | 0,240891 | 15765,27896 | -1,008398255 | 0,255601 | -3,94521 | 7,97E-05 | 0,000207 |
| 658 | RM25_RS03180 | RM25_0649 | Isocitrate dehydrogenase [NADP] | 685 | gene | 758905 | 760122 | + | 4241,222 | 0,765689097 | 0,273367 | 2,800959 | 0,005095 | 0,030098 | 4987,909704 | 0,507876214 | 0,166656 | 3,047451 | 0,002308 | 0,005579 | 3494,535 | 1,157456244 | 0,315448 | 3,669246 | 0,000243 | 0,001541 | 3341,809 | -0,98271 | 0,253146 | -3,88197 | 0,000104 | 0,000501 | 5221,519837 | -0,29682698 | 0,225933 | -1,31378 | 0,188919 | 0,242396 |
| 659 | RM25_RS12545 | RM25_0650 | Hypothetical protein | 686 | gene | 760314 | 760625 | - | 197,8435 | 0,020509596 | 0,189397 | 0,108289 | 0,913766 | 0,942972 | 220,5180963 | 0,242586463 | 0,176537 | 1,374142 | 0,169398 | 0,225322 | 175,1688 | -0,249665252 | 0,273004 | -0,91451 | 0,360448 | 0,467057 | 207,1537 | -0,14545 | 0,19461 | -0,7474 | 0,454821 | 0,528606 | 195,9987707 | -0,57777265 | 0,260346 | -2,21925 | 0,02647 | 0,042326 |
| 660 | RM25_RS13180 | RM25_0651 | Hypothetical protein | 687 | gene | 760703 | 760921 | + | 98,03262 | -1,119126932 | 0,401281 | -2,78888 | 0,005289 | 0,030857 | 66,71627285 | -1,528867923 | 0,354348 | -4,31459 | 1,6E-05 | 6,51E-05 | 129,349 | -0,932259822 | 0,367532 | -2,53654 | 0,011195 | 0,033234 | 141,3038 | 0,704032 | 0,27675 | 2,543931 | 0,010961 | 0,025486 | 60,74097446 | 1,372871143 | 0,472543 | 2,905161 | 0,003671 | 0,006938 |
| 661 | NA | RM25_0652 | Hypothetical protein | 688 | gene | 761096 | 761344 | + | 101,1132 | -1,066079678 | 0,547789 | -1,94615 | 0,051637 | 0,126779 | 59,88992917 | -1,976643401 | 0,450202 | -4,39057 | 1,13E-05 | 4,7E-05 | 142,3364 | -0,748903657 | 0,468954 | -1,59697 | 0,110273 | 0,185786 | 143,7118 | 0,830228 | 0,374826 | 2,21497 | 0,026762 | 0,05145 | 64,47927401 | 2,133684616 | 0,567383 | 3,76057 | 0,00017 | 0,000415 |
| 662 | RM25_RS03200 | RM25_0653 | Putative hemolysin | 689 | gene | 761583 | 763103 | + | 1252,348 | 0,317252298 | 0,173429 | 1,829289 | 0,067356 | 0,149773 | 1421,890204 | 0,537462141 | 0,145586 | 3,691709 | 0,000223 | 0,000705 | 1082,806 | 0,035203758 | 0,176189 | 0,199807 | 0,841631 | 0,885196 | 1177,201 | -0,17522 | 0,156609 | -1,11884 | 0,263209 | 0,3348 | 1363,085286 | -0,626581549 | 0,137212 | -4,56651 | 4,96E-06 | 1,53E-05 |
| 663 | RM25_RS03205 | RM25_0654 | Hypothetical protein | 690 | gene | 763184 | 764473 | + | 444,7206 | -0,316584504 | 0,292044 | -1,08403 | 0,278352 | 0,387751 | 349,9559765 | -0,078682036 | 0,191864 | -0,41009 | 0,681738 | 0,735403 | 539,4852 | -0,474991171 | 0,388962 | -1,22118 | 0,222019 | 0,318991 | 517,1574 | 0,738715 | 0,250116 | 2,95349 | 0,003142 | 0,009177 | 389,9885977 | 0,410013785 | 0,352129 | 1,164384 | 0,244268 | 0,301751 |
| 664 | RM25_RS03210 | RM25_0655 | Basic membrane protein | 691 | gene | 764723 | 765667 | + | 549,7226 | -0,222951664 | 0,116276 | -1,91743 | 0,055183 | 0,132317 | 524,6929862 | -0,156854925 | 0,154144 | -1,01759 | 0,308873 | 0,376811 | 574,7522 | -0,284169375 | 0,216003 | -1,31558 | 0,188315 | 0,282533 | 625,7024 | 0,124884 | 0,178405 | 0,699999 | 0,483928 | 0,554326 | 498,2940523 | 0,06029702 | 0,213839 | 0,281974 | 0,777964 | 0,81933 |
| 665 | RM25_RS03215 | RM25_0656 | Ribonuclease H | 692 | gene | 765739 | 766770 | + | 678,0523 | 0,192612454 | 0,295734 | 0,651302 | 0,514851 | 0,611954 | 877,8084243 | 0,266892179 | 0,191187 | 1,395976 | 0,162722 | 0,217794 | 478,2962 | 0,061139847 | 0,176405 | 0,346587 | 0,728901 | 0,799079 | 672,8593 | -0,82636 | 0,23031 | -3,58802 | 0,000333 | 0,00141 | 709,4672489 | -0,981379706 | 0,145654 | -6,73775 | 1,61E-11 | 1,16E-10 |
| 666 | RM25_RS03220 | RM25_0657 | Hypothetical protein | 693 | gene | 766847 | 767362 | + | 549,2851 | -0,721933799 | 0,421463 | -1,71292 | 0,086727 | 0,176421 | 323,4520665 | -0,055343998 | 0,172642 | -0,32057 | 0,748535 | 0,795087 | 775,1181 | -1,020116793 | 0,3863 | -2,64074 | 0,008273 | 0,026457 | 718,081 | 1,605323 | 0,371232 | 4,324309 | 1,53E-05 | 9,02E-05 | 405,4351523 | 0,677313777 | 0,207728 | 3,260582 | 0,001112 | 0,002321 |
| 667 | RM25_RS03225 | RM25_0658 | Enoyl-CoA hydratase echA9 | 694 | gene | 767438 | 768370 | - | 919,2587 | 0,046726685 | 0,099238 | 0,470854 | 0,637745 | 0,722634 | 870,0755447 | 0,091298403 | 0,144809 | 0,630477 | 0,528383 | 0,594178 | 968,4418 | 0,006473238 | 0,169629 | 0,038161 | 0,969559 | 0,982481 | 955,0125 | 0,140448 | 0,182261 | 0,77059 | 0,44095 | 0,516047 | 914,7657724 | 0,104349563 | 0,116011 | 0,899482 | 0,368396 | 0,433288 |
| 668 | RM25_RS03230 | RM25_0659 | Methionine--tRNA ligase | 695 | gene | 768411 | 770357 | + | 2696,498 | 0,390741842 | 0,218561 | 1,787791 | 0,07381 | 0,159751 | 3186,165327 | 0,254414082 | 0,139715 | 1,820951 | 0,068614 | 0,102747 | 2206,83 | 0,591413183 | 0,225474 | 2,622976 | 0,008717 | 0,02754 | 2476,631 | -0,78283 | 0,189385 | -4,13354 | 3,57E-05 | 0,000194 | 2995,854848 | -0,398504961 | 0,14963 | -2,66327 | 0,007738 | 0,013818 |
| 669 | RM25_RS03235 | RM25_0660 | Silent information regulator protein Sir2 | 696 | gene | 770585 | 771508 | + | 670,9432 | 0,052116259 | 0,137359 | 0,379416 | 0,704379 | 0,780546 | 664,8985307 | 0,156084994 | 0,134625 | 1,159409 | 0,24629 | 0,310763 | 676,9878 | -0,047890609 | 0,269602 | -0,17763 | 0,85901 | 0,897865 | 694,0291 | 0,066015 | 0,202119 | 0,326615 | 0,743959 | 0,788804 | 671,0731417 | -0,075897743 | 0,213939 | -0,35476 | 0,722767 | 0,76945 |
| 670 | RM25_RS03240 | RM25_0661 | PF04239 family protein | 697 | gene | 771527 | 772171 | - | 235,2602 | -0,714001492 | 0,395788 | -1,804 | 0,071231 | 0,155744 | 156,3887328 | -1,015416095 | 0,320757 | -3,16569 | 0,001547 | 0,003917 | 314,1317 | -0,574951032 | 0,350384 | -1,64092 | 0,100815 | 0,173953 | 306,4018 | 0,777463 | 0,332268 | 2,339868 | 0,019291 | 0,039798 | 175,1585693 | 1,283865863 | 0,354602 | 3,620581 | 0,000294 | 0,000689 |
| 671 | RM25_RS03245 | RM25_0662 | Putative membrane protein | 698 | gene | 772168 | 772623 | - | 216,2419 | -1,181978937 | 0,348552 | -3,39111 | 0,000696 | 0,007854 | 156,7511108 | -1,559247164 | 0,29446 | -5,29527 | 1,19E-07 | 7,07E-07 | 275,7328 | -0,990166188 | 0,291926 | -3,39185 | 0,000694 | 0,00362 | 315,5741 | 0,579406 | 0,293987 | 1,970855 | 0,04874 | 0,083308 | 129,869476 | 1,214554354 | 0,321083 | 3,782677 | 0,000155 | 0,000383 |
| 672 | RM25_RS03250 | RM25_0663 | Transcriptional regulator, MerR family | 699 | gene | 772796 | 773161 | - | 298,6918 | -1,086343922 | 0,422533 | -2,57103 | 0,01014 | 0,047609 | 200,8216596 | -1,645267444 | 0,333726 | -4,93 | 8,22E-07 | 4,21E-06 | 396,5619 | -0,83620938 | 0,330348 | -2,5313 | 0,011364 | 0,033496 | 426,4741 | 0,6726 | 0,346583 | 1,94066 | 0,0523 | 0,08805 | 187,9785992 | 1,548868399 | 0,347248 | 4,460415 | 8,18E-06 | 2,42E-05 |
| 673 | RM25_RS03255 | RM25_0664 | Carboxymuconolactone decarboxylase | 700 | gene | 773274 | 773681 | + | 120,4433 | -1,382964359 | 0,598859 | -2,30933 | 0,020925 | 0,073505 | 68,83399063 | -2,663375773 | 0,467 | -5,70316 | 1,18E-08 | 8,48E-08 | 172,0526 | -1,001952251 | 0,415975 | -2,40868 | 0,01601 | 0,043425 | 183,0831 | 0,870231 | 0,383297 | 2,270383 | 0,023184 | 0,046083 | 65,7867618 | 2,612940217 | 0,543439 | 4,808158 | 1,52E-06 | 5,09E-06 |
| 674 | RM25_RS12700 | RM25_0665 | Hypothetical protein | 701 | gene | 773678 | 773848 | + | 59,00971 | -1,25553919 | 0,379993 | -3,30411 | 0,000953 | 0,009713 | 41,63607649 | -1,565184177 | 0,442944 | -3,5336 | 0,00041 | 0,00122 | 76,38335 | -1,11270433 | 0,340876 | -3,26425 | 0,001098 | 0,005247 | 87,57869 | 0,669949 | 0,339444 | 1,973665 | 0,04842 | 0,08294 | 34,05930348 | 1,190895254 | 0,486971 | 2,445518 | 0,014464 | 0,024439 |
| 675 | RM25_RS03260 | RM25_0666 | Transposase IS204/IS1001/IS1096/IS1165 family protein | 702 | gene | 774099 | 775175 | - | 45,47276 | -1,790580564 | 0,698729 | -2,56263 | 0,010388 | 0,04839 | 22,14928363 | -3,22808181 | 0,753554 | -4,28381 | 1,84E-05 | 7,38E-05 | 68,79623 | -1,47967348 | 0,401061 | -3,6894 | 0,000225 | 0,001452 | 73,85818 | 1,261707 | 0,433538 | 2,910257 | 0,003611 | 0,01021 | 19,97031954 | 3,080489559 | 0,78371 | 3,930649 | 8,47E-05 | 0,000218 |
| 676 | RM25_RS03270 | RM25_0667 | Putative insertion element ISCmi2 transposase | 703 | gene | 776332 | 777375 | - | 110,9456 | -1,288923202 | 0,60806 | -2,11973 | 0,034029 | 0,097812 | 67,12685603 | -2,841763432 | 0,519218 | -5,47316 | 4,42E-08 | 2,86E-07 | 154,7643 | -0,818255244 | 0,353817 | -2,31265 | 0,020742 | 0,052682 | 165,241 | 0,668785 | 0,459087 | 1,456771 | 0,145179 | 0,201619 | 63,19720412 | 2,767365585 | 0,452373 | 6,117443 | 9,51E-10 | 5,34E-09 |
| 677 | RM25_RS11795 | RM25_0668 | Puromycin N-acetyltransferase | 704 | gene | 777524 | 778144 | - | 262,3489 | -0,102911557 | 0,183574 | -0,5606 | 0,575071 | 0,667705 | 232,2676291 | -0,014550919 | 0,209129 | -0,06958 | 0,944529 | 0,955885 | 292,4302 | -0,172032196 | 0,259273 | -0,66352 | 0,506998 | 0,606905 | 285,5569 | 0,352895 | 0,201712 | 1,749503 | 0,080204 | 0,123556 | 248,4117968 | 0,251588996 | 0,256526 | 0,980754 | 0,326714 | 0,390896 |
| 678 | NA | RM25_0669 | Hypothetical protein | 705 | gene | 778465 | 778671 | + | 193,7259 | -0,032356739 | 0,223599 | -0,14471 | 0,884941 | 0,922107 | 216,8694524 | 0,133319314 | 0,223472 | 0,596583 | 0,550786 | 0,61643 | 170,5824 | -0,238386617 | 0,324491 | -0,73465 | 0,462554 | 0,565771 | 207,2571 | -0,2209 | 0,250662 | -0,88128 | 0,378165 | 0,455227 | 187,4988464 | -0,554720028 | 0,298485 | -1,85845 | 0,063105 | 0,091251 |
| 679 | RM25_RS03280 | RM25_0670 | Transposase IS204/IS1001/IS1096/IS1165 family protein | 706 | gene | 778651 | 779727 | - | 91,0621 | -1,681664349 | 0,789923 | -2,1289 | 0,033263 | 0,096671 | 51,08307858 | -4,516502902 | 0,689936 | -6,54626 | 5,9E-11 | 6,01E-10 | 131,0411 | -1,13282318 | 0,421337 | -2,68864 | 0,007174 | 0,023884 | 145,572 | 0,802975 | 0,49606 | 1,618706 | 0,10551 | 0,15543 | 42,58401131 | 4,263041837 | 0,682552 | 6,245741 | 4,22E-10 | 2,46E-09 |
| 680 | RM25_RS03285 | RM25_0671 | Chlorophyll synthesis pathway protein BchC | 707 | gene | 780442 | 781485 | - | 1683,511 | -0,023631364 | 0,24445 | -0,09667 | 0,922987 | 0,948749 | 1956,080416 | 0,422642615 | 0,121333 | 3,483323 | 0,000495 | 0,001436 | 1410,941 | -0,646261981 | 0,170864 | -3,78232 | 0,000155 | 0,00107 | 1794,152 | -0,01428 | 0,180099 | -0,07927 | 0,936821 | 0,949714 | 1638,409067 | -1,034282507 | 0,105096 | -9,84132 | 7,47E-23 | 2,29E-21 |
| 681 | RM25_RS03290 | RM25_0672 | Putative membrane protein (DUF2207) (Precursor) | 708 | gene | 781623 | 783410 | - | 1694,306 | -0,137953005 | 0,091046 | -1,5152 | 0,129722 | 0,228346 | 1708,79041 | 0,034751528 | 0,12357 | 0,281229 | 0,778535 | 0,822504 | 1679,822 | -0,31255449 | 0,153703 | -2,03349 | 0,042003 | 0,088949 | 1872,426 | 0,081705 | 0,119189 | 0,68551 | 0,493022 | 0,561108 | 1583,232394 | -0,211875202 | 0,141823 | -1,49394 | 0,135192 | 0,180537 |
| 682 | RM25_RS03295 | RM25_0673 | Pyruvate-flavodoxin oxidoreductase | 709 | gene | 783722 | 787441 | + | 5213,444 | 0,596902061 | 0,222291 | 2,685232 | 0,007248 | 0,03839 | 6335,139926 | 0,67964759 | 0,157088 | 4,326541 | 1,51E-05 | 6,19E-05 | 4091,748 | 0,471172365 | 0,166238 | 2,834319 | 0,004592 | 0,016704 | 4398,845 | -0,56306 | 0,177303 | -3,17572 | 0,001495 | 0,004934 | 6152,028056 | -0,725627408 | 0,126901 | -5,71807 | 1,08E-08 | 5,1E-08 |
| 683 | RM25_RS03300 | RM25_0674 | Dihydroorotate oxidase | 710 | gene | 787445 | 788452 | + | 1289,597 | 0,630471495 | 0,288253 | 2,187217 | 0,028727 | 0,087076 | 1662,707524 | 0,71392412 | 0,182301 | 3,916193 | 9E-05 | 0,00031 | 916,4875 | 0,484748689 | 0,191481 | 2,531576 | 0,011355 | 0,033496 | 1073,726 | -0,77469 | 0,225383 | -3,4372 | 0,000588 | 0,002249 | 1536,196313 | -0,961632145 | 0,122869 | -7,82648 | 5,02E-15 | 6,19E-14 |
| 684 | RM25_RS03305 | RM25_0675 | Hypothetical protein | 711 | gene | 788480 | 789022 | - | 265,572 | -0,699299919 | 0,239016 | -2,92574 | 0,003436 | 0,022688 | 232,4632255 | -0,51339612 | 0,219582 | -2,33806 | 0,019384 | 0,03477 | 298,6807 | -0,849812898 | 0,367793 | -2,31057 | 0,020857 | 0,052859 | 345,2456 | 0,429839 | 0,235648 | 1,824072 | 0,068141 | 0,10963 | 199,2519525 | 0,156508499 | 0,364 | 0,429968 | 0,667219 | 0,719412 |
| 685 | RM25_RS03310 | RM25_0676 | Bifunctional acetyl-/propionyl-coenzyme A carboxylase subu | 712 | gene | 789075 | 790856 | - | 3695,841 | 0,413714643 | 0,261647 | 1,581197 | 0,113833 | 0,208635 | 4523,742907 | 0,336391749 | 0,173316 | 1,940913 | 0,052269 | 0,082022 | 2867,94 | 0,53767965 | 0,273284 | 1,967477 | 0,049128 | 0,099738 | 3366,937 | -0,8266 | 0,256197 | -3,22643 | 0,001253 | 0,004282 | 4133,121289 | -0,587022533 | 0,177033 | -3,31589 | 0,000914 | 0,001961 |
| 686 | RM25_RS03315 | RM25_0677 | DNA-formamidopyrimidine glycosylase | 713 | gene | 791012 | 791827 | - | 430,6227 | -0,087064726 | 0,156383 | -0,55674 | 0,577704 | 0,670352 | 402,5615701 | 0,143395034 | 0,183818 | 0,780092 | 0,435337 | 0,503725 | 458,6839 | -0,289317776 | 0,229348 | -1,26148 | 0,207136 | 0,303809 | 467,8783 | 0,331793 | 0,213305 | 1,555489 | 0,11983 | 0,172219 | 410,5656372 | -0,036149735 | 0,227108 | -0,15917 | 0,873532 | 0,897521 |
| 687 | NA | RM25_0678 | Hypothetical protein | 714 | gene | 791978 | 792193 | - | 1217,065 | 0,43806588 | 0,428275 | 1,022862 | 0,306373 | 0,415729 | 800,8975673 | 0,145577825 | 0,245475 | 0,593044 | 0,553152 | 0,618197 | 1633,233 | 0,58458991 | 0,521066 | 1,121911 | 0,2619 | 0,363331 | 1091,417 | 0,731734 | 0,439207 | 1,666034 | 0,095707 | 0,143317 | 1360,709114 | 1,196034619 | 0,364999 | 3,276818 | 0,00105 | 0,002211 |
| 688 | RM25_RS12555 | RM25_0679 | Hypothetical protein | 715 | gene | 792474 | 792953 | + | 381,7971 | -0,59538748 | 0,246171 | -2,41859 | 0,015581 | 0,062243 | 293,8782981 | -0,482448219 | 0,219775 | -2,19519 | 0,02815 | 0,04829 | 469,7159 | -0,668786202 | 0,194064 | -3,44622 | 0,000568 | 0,003108 | 483,1551 | 0,684129 | 0,206506 | 3,312878 | 0,000923 | 0,003293 | 298,2089042 | 0,55802191 | 0,224389 | 2,48685 | 0,012888 | 0,021996 |
| 689 | RM25_RS03325 | RM25_0680 | MutT/NUDIX family protein | 716 | gene | 793044 | 793544 | - | 328,8931 | 0,090109047 | 0,14258 | 0,631991 | 0,527393 | 0,623403 | 324,2355217 | 0,355917044 | 0,202219 | 1,76006 | 0,078398 | 0,114915 | 333,5508 | -0,161584142 | 0,186326 | -0,86721 | 0,385826 | 0,491298 | 336,2791 | 0,244409 | 0,176404 | 1,385509 | 0,165897 | 0,224853 | 332,944653 | -0,215515254 | 0,227091 | -0,94903 | 0,342607 | 0,406608 |
| 690 | RM25_RS03330 | RM25_0681 | Septum formation protein Maf | 717 | gene | 793548 | 794276 | - | 662,6553 | 0,101234105 | 0,1695 | 0,597252 | 0,550339 | 0,645027 | 726,3619752 | 0,415061007 | 0,139415 | 2,977165 | 0,002909 | 0,006809 | 598,9487 | -0,272207354 | 0,186333 | -1,46087 | 0,144052 | 0,228486 | 674,3929 | 0,017107 | 0,176724 | 0,096802 | 0,922884 | 0,938816 | 673,1093876 | -0,619995566 | 0,130455 | -4,75258 | 2,01E-06 | 6,61E-06 |
| 691 | RM25_RS03335 | RM25_0682 | Hypothetical protein | 718 | gene | 794327 | 794581 | - | 287,7838 | 0,533874438 | 0,317398 | 1,682036 | 0,092562 | 0,183704 | 367,0600264 | 0,3663979 | 0,243877 | 1,502386 | 0,132997 | 0,183964 | 208,5075 | 0,837582121 | 0,237665 | 3,524209 | 0,000425 | 0,002458 | 249,4881 | -1,15795 | 0,281522 | -4,11318 | 3,9E-05 | 0,000208 | 333,7698472 | -0,631847023 | 0,178688 | -3,53603 | 0,000406 | 0,000921 |
| 692 | RM25_RS03340 | RM25_0683 | Propionyl-CoA carboxylase subunit beta 5 accD5 | 719 | gene | 794587 | 796161 | - | 3379,083 | 0,325523801 | 0,214115 | 1,520326 | 0,128429 | 0,2268 | 3988,74319 | 0,27066998 | 0,157015 | 1,723846 | 0,084736 | 0,123057 | 2769,422 | 0,405644369 | 0,233947 | 1,733912 | 0,082934 | 0,149103 | 3181,454 | -0,65771 | 0,204826 | -3,21106 | 0,001322 | 0,004498 | 3679,861308 | -0,480508951 | 0,164476 | -2,92146 | 0,003484 | 0,006622 |
| 693 | RM25_RS03345 | RM25_0684 | hypothetical protein | 720 | gene | 796293 | 797639 | + | 919,758 | -0,432308144 | 0,206859 | -2,08987 | 0,036629 | 0,102658 | 803,9619037 | -0,205656395 | 0,158469 | -1,29777 | 0,194366 | 0,253386 | 1035,554 | -0,612660232 | 0,308649 | -1,98498 | 0,047147 | 0,096888 | 1111,003 | 0,474634 | 0,20912 | 2,269675 | 0,023227 | 0,046083 | 769,6832422 | 0,132384815 | 0,271492 | 0,487619 | 0,62582 | 0,681949 |
| 694 | RM25_RS03350 | RM25_0685 | Putative exporter of polyketide antibiotics | 721 | gene | 797747 | 799219 | - | 453,2307 | -1,402212156 | 0,629639 | -2,22701 | 0,025947 | 0,082825 | 238,2792287 | -2,818269699 | 0,322102 | -8,74962 | 2,14E-18 | 5,31E-17 | 668,1822 | -1,035386582 | 0,437908 | -2,36439 | 0,01806 | 0,047614 | 689,6781 | 1,033924 | 0,333868 | 3,096807 | 0,001956 | 0,006164 | 245,736712 | 2,894643447 | 0,450117 | 6,430864 | 1,27E-10 | 8,13E-10 |
| 695 | RM25_RS03355 | RM25_0686 | Transposase IS204/IS1001/IS1096/IS1165 family protein | 722 | gene | 799583 | 800659 | + | 11,54689 | 1,011364839 | 1,304907 | 0,775047 | 0,438312 | 0,546903 | 1,383634083 | -3,679953192 | 2,400392 | -1,53306 | 0,12526 | 0,174285 | 21,71015 | 1,283362821 | 0,779707 | 1,645956 | 0,099773 | 0,172408 | 7,865177 | 2,191353 | 0,820857 | 2,669591 | 0,007594 | 0,018802 | 15,36109757 | 7,189516973 | 1,606475 | 4,475338 | 7,63E-06 | 2,28E-05 |
| 696 | RM25_RS12560 | RM25_0687 | Hypothetical protein | 723 | gene | 800625 | 800834 | - | 36,2658 | -0,811241395 | 0,414445 | -1,95742 | 0,050299 | 0,125197 | 23,82582976 | -1,028730777 | 0,584677 | -1,75949 | 0,078495 | 0,114986 | 48,70577 | -0,722628444 | 0,272736 | -2,64955 | 0,00806 | 0,026088 | 48,51825 | 0,846338 | 0,355006 | 2,384012 | 0,017125 | 0,036233 | 25,76786065 | 1,219818548 | 0,533899 | 2,284736 | 0,022328 | 0,036298 |
| 697 | RM25_RS03360 | RM25_0688 | Tetronasin-transport ATP-binding protein ABC transporter | 724 | gene | 800831 | 801775 | - | 451,2345 | -0,994148016 | 0,46167 | -2,15337 | 0,031289 | 0,092882 | 248,7019485 | -1,295444261 | 0,256202 | -5,05635 | 4,27E-07 | 2,28E-06 | 653,767 | -0,886718888 | 0,293983 | -3,01623 | 0,002559 | 0,010621 | 631,0176 | 1,198296 | 0,293899 | 4,077235 | 4,56E-05 | 0,00024 | 296,330685 | 1,669410215 | 0,286927 | 5,818241 | 5,95E-09 | 2,92E-08 |
| 698 | RM25_RS03365 | RM25_0689 | Putative membrane protein | 725 | gene | 801880 | 802398 | + | 177,594 | -0,651535619 | 0,284065 | -2,29362 | 0,021813 | 0,075137 | 138,8045676 | -0,742607976 | 0,329862 | -2,25127 | 0,024368 | 0,04242 | 216,3835 | -0,597562135 | 0,252705 | -2,36466 | 0,018046 | 0,047614 | 228,5792 | 0,520467 | 0,318521 | 1,634009 | 0,102257 | 0,151489 | 135,5907698 | 0,723355174 | 0,301231 | 2,401334 | 0,016335 | 0,027385 |
| 699 | RM25_RS03370 | RM25_0690 | Transcription regulator | 726 | gene | 802513 | 803091 | + | 420,2911 | -0,347983821 | 0,127735 | -2,72425 | 0,006445 | 0,035179 | 457,6857927 | -0,35740329 | 0,177557 | -2,01289 | 0,044126 | 0,07158 | 382,8965 | -0,334621627 | 0,168579 | -1,98496 | 0,047149 | 0,096888 | 498,6022 | -0,32501 | 0,182715 | -1,77877 | 0,075277 | 0,117503 | 362,8002929 | -0,248572521 | 0,180668 | -1,37585 | 0,168867 | 0,220817 |
| 700 | RM25_RS11800 | RM25_0691 | Transcriptional regulator, TetR family | 727 | gene | 803105 | 803710 | - | 621,0649 | 0,299356779 | 0,337625 | 0,886655 | 0,375265 | 0,486523 | 404,9874935 | 0,180460692 | 0,156574 | 1,152561 | 0,24909 | 0,313793 | 837,1422 | 0,356116722 | 0,229738 | 1,5501 | 0,121118 | 0,200052 | 585,3758 | 0,894934 | 0,196959 | 4,543761 | 5,53E-06 | 3,63E-05 | 669,3603525 | 1,112796703 | 0,173138 | 6,427223 | 1,3E-10 | 8,3E-10 |
| 701 | RM25_RS03380 | RM25_0692 | Biotin-[acetyl-CoA-carboxylase] ligase | 728 | gene | 803860 | 804651 | + | 737,2196 | 0,198338226 | 0,194386 | 1,020331 | 0,307572 | 0,416636 | 842,6443347 | 0,290289161 | 0,223459 | 1,299072 | 0,193919 | 0,252943 | 631,7948 | 0,077666075 | 0,181768 | 0,427281 | 0,669174 | 0,749285 | 727,5634 | -0,36078 | 0,192701 | -1,87221 | 0,061178 | 0,099997 | 772,3258362 | -0,524532884 | 0,227566 | -2,30497 | 0,021168 | 0,034673 |
| 702 | RM25_RS03385 | RM25_0693 | Putative membrane protein | 729 | gene | 804685 | 805188 | + | 584,3826 | -0,01853931 | 0,21012 | -0,08823 | 0,929692 | 0,952318 | 624,570823 | 0,415966515 | 0,213333 | 1,94985 | 0,051194 | 0,080604 | 544,1944 | -0,513713671 | 0,191357 | -2,68458 | 0,007262 | 0,02404 | 619,8968 | 0,206214 | 0,235928 | 0,874055 | 0,382088 | 0,458779 | 569,3265425 | -0,680392298 | 0,167827 | -4,05413 | 5,03E-05 | 0,000133 |
| 703 | RM25_RS03390 | RM25_0694 | Transcriptional regulator, TetR family | 730 | gene | 805270 | 805854 | + | 1057,62 | 0,646934672 | 0,475798 | 1,359684 | 0,17393 | 0,277558 | 1556,177178 | 0,742555724 | 0,223095 | 3,328435 | 0,000873 | 0,002377 | 559,062 | 0,392549601 | 0,406222 | 0,966342 | 0,333873 | 0,43914 | 878,8918 | -1,31749 | 0,424945 | -3,10038 | 0,001933 | 0,006115 | 1266,027581 | -1,631607427 | 0,177425 | -9,19603 | 3,71E-20 | 8,75E-19 |
| 704 | RM25_RS03395 | RM25_0695 | Putative insertion element ISCmi2 transposase | 731 | gene | 805963 | 807006 | + | 71,49441 | 0,088780293 | 0,285477 | 0,31099 | 0,755808 | 0,823216 | 62,69390141 | -0,50194971 | 0,403629 | -1,24359 | 0,21365 | 0,274726 | 80,29491 | 0,536438434 | 0,240452 | 2,230961 | 0,025684 | 0,061904 | 73,62829 | -0,22459 | 0,276725 | -0,81161 | 0,417013 | 0,492681 | 71,83571161 | 0,865610703 | 0,391435 | 2,211379 | 0,02701 | 0,043044 |
| 705 | RM25_RS03400 | RM25_0696 | Peptidase family S51 | 732 | gene | 807003 | 807719 | - | 922,1743 | 0,083556273 | 0,187349 | 0,445992 | 0,655603 | 0,738119 | 927,6548899 | 0,321041622 | 0,170478 | 1,883187 | 0,059675 | 0,091038 | 916,6937 | -0,153266295 | 0,313689 | -0,48859 | 0,62513 | 0,711116 | 947,4764 | 0,174267 | 0,321333 | 0,542326 | 0,587594 | 0,649605 | 928,9366724 | -0,259822135 | 0,152074 | -1,70853 | 0,087538 | 0,122377 |
| 706 | RM25_RS03405 | RM25_0697 | Phosphoribosylaminoimidazole carboxylase, ATPase subunit | 733 | gene | 807797 | 808969 | + | 1991,49 | 0,522034777 | 0,289728 | 1,801807 | 0,071576 | 0,156063 | 2525,10962 | 0,71087055 | 0,179714 | 3,955562 | 7,64E-05 | 0,000267 | 1457,871 | 0,206234017 | 0,274465 | 0,751404 | 0,45241 | 0,555887 | 1735,83 | -0,5538 | 0,278564 | -1,98804 | 0,046807 | 0,080647 | 2300,622062 | -1,019145735 | 0,164951 | -6,17848 | 6,47E-10 | 3,72E-09 |
| 707 | RM25_RS03410 | RM25_0698 | N5-carboxyaminoimidazole ribonucleotide mutase | 734 | gene | 808962 | 809522 | + | 566,3009 | 0,320580603 | 0,164061 | 1,954036 | 0,050697 | 0,12544 | 636,0006463 | 0,515578023 | 0,172236 | 2,993435 | 0,002759 | 0,006512 | 496,6012 | 0,080037111 | 0,166018 | 0,482098 | 0,629736 | 0,7136 | 532,628 | -0,17447 | 0,186714 | -0,93443 | 0,350081 | 0,425769 | 617,1331781 | -0,557603248 | 0,159795 | -3,4895 | 0,000484 | 0,001089 |
| 708 | RM25_RS03415 | RM25_0699 | Cell envelope-related transcriptional regulator | 735 | gene | 809560 | 810918 | - | 1204,772 | 0,103732197 | 0,163637 | 0,633916 | 0,526136 | 0,622541 | 1357,915466 | 0,20023866 | 0,1664 | 1,203359 | 0,228837 | 0,292183 | 1051,629 | -0,018509183 | 0,179912 | -0,10288 | 0,918059 | 0,943272 | 1230,24 | -0,31211 | 0,214347 | -1,45612 | 0,14536 | 0,201656 | 1223,235817 | -0,484359881 | 0,118891 | -4,07399 | 4,62E-05 | 0,000123 |
| 709 | RM25_RS03420 | RM25_0700 | Mannose-1-phosphate guanylyltransferase | 736 | gene | 811044 | 812123 | + | 1707,374 | 0,341784781 | 0,38308 | 0,892202 | 0,372285 | 0,484258 | 2365,688343 | 0,483554016 | 0,207135 | 2,334483 | 0,01957 | 0,035038 | 1049,059 | 0,029956116 | 0,260934 | 0,114803 | 0,908601 | 0,936285 | 1600,938 | -0,97778 | 0,305536 | -3,20021 | 0,001373 | 0,004624 | 1872,169862 | -1,391174669 | 0,125168 | -11,1145 | 1,07E-28 | 5,85E-27 |
| 710 | RM25_RS03425 | RM25_0701 | Putative membrane protein | 737 | gene | 812120 | 813187 | + | 1009,312 | 0,323553393 | 0,24639 | 1,313175 | 0,189124 | 0,293267 | 1214,645702 | 0,350386181 | 0,21552 | 1,625773 | 0,103998 | 0,146868 | 803,9791 | 0,285734615 | 0,253903 | 1,125368 | 0,260433 | 0,361721 | 949,5783 | -0,61128 | 0,303553 | -2,01375 | 0,044035 | 0,076939 | 1100,843105 | -0,629239965 | 0,131661 | -4,77924 | 1,76E-06 | 5,86E-06 |
| 711 | RM25_RS03430 | RM25_0702 | Transcription factor WhiB | 738 | gene | 813520 | 813780 | + | 1236,193 | 0,708325444 | 0,263701 | 2,686095 | 0,007229 | 0,038377 | 1426,189934 | 0,387667012 | 0,216057 | 1,794279 | 0,072769 | 0,107736 | 1046,197 | 1,178274513 | 0,24633 | 4,783322 | 1,72E-06 | 2,21E-05 | 999,5292 | -1,00235 | 0,291189 | -3,44228 | 0,000577 | 0,002213 | 1501,468344 | -0,167491536 | 0,160551 | -1,04323 | 0,296841 | 0,358246 |
| 712 | RM25_RS03435 | RM25_0703 | Hypothetical protein | 739 | gene | 813977 | 817216 | + | 1796,835 | 0,158695197 | 0,1938 | 0,81886 | 0,412866 | 0,522624 | 1928,542787 | 0,432343302 | 0,165847 | 2,606879 | 0,009137 | 0,018391 | 1665,127 | -0,153059691 | 0,28853 | -0,53048 | 0,595778 | 0,684133 | 1787,837 | 0,036514 | 0,22445 | 0,162681 | 0,870769 | 0,8986 | 1862,855018 | -0,490658909 | 0,229231 | -2,14046 | 0,032318 | 0,050279 |
| 713 | RM25_RS03440 | RM25_0704 | Hypothetical protein | 740 | gene | 817213 | 818565 | + | 489,3974 | 0,072249364 | 0,176281 | 0,409853 | 0,681914 | 0,760658 | 479,8679909 | 0,192785335 | 0,171841 | 1,121883 | 0,261912 | 0,326629 | 498,9267 | -0,040674621 | 0,317796 | -0,12799 | 0,898157 | 0,928897 | 502,3174 | 0,117533 | 0,203085 | 0,578739 | 0,562765 | 0,627751 | 493,2206194 | -0,054473871 | 0,286413 | -0,19019 | 0,849158 | 0,878606 |
| 714 | RM25_RS03445 | RM25_0705 | Hypothetical protein | 741 | gene | 818670 | 819110 | - | 147,1671 | 0,136905742 | 0,248635 | 0,55063 | 0,581888 | 0,67396 | 115,6182462 | 0,363756179 | 0,248814 | 1,461962 | 0,143752 | 0,196419 | 178,716 | -0,005978214 | 0,228243 | -0,02619 | 0,979104 | 0,987483 | 147,6444 | 0,760619 | 0,231129 | 3,290885 | 0,000999 | 0,003524 | 151,2106134 | 0,448573462 | 0,258856 | 1,732906 | 0,083112 | 0,116744 |
| 715 | RM25_RS13190 | RM25_0706 | Hypothetical protein | 742 | gene | 819177 | 819677 | + | 770,8616 | 0,707035031 | 0,115803 | 6,105486 | 1,02E-09 | 1,21E-07 | 796,1947477 | 0,735881763 | 0,163401 | 4,503526 | 6,68E-06 | 2,9E-05 | 745,5285 | 0,676956701 | 0,211309 | 3,203635 | 0,001357 | 0,006247 | 618,9986 | -0,1148 | 0,186691 | -0,61492 | 0,538611 | 0,605103 | 935,3850648 | -0,129865454 | 0,158576 | -0,81895 | 0,412815 | 0,478606 |
| 716 | RM25_RS03455 | RM25_0707 | Eukaryotic phosphomannomutase | 743 | gene | 819793 | 820608 | + | 1999,74 | 0,652111806 | 0,230151 | 2,833406 | 0,004605 | 0,028122 | 2308,445172 | 0,602357111 | 0,177914 | 3,38567 | 0,00071 | 0,001976 | 1691,034 | 0,721669147 | 0,316592 | 2,279492 | 0,022638 | 0,05643 | 1650,367 | -0,57768 | 0,235219 | -2,45593 | 0,014052 | 0,031041 | 2388,482068 | -0,42218161 | 0,244927 | -1,72371 | 0,084761 | 0,118565 |
| 717 | RM25_RS03460 | RM25_0708 | Hypothetical protein | 744 | gene | 820669 | 820869 | + | 1358,586 | 0,899045198 | 0,279363 | 3,218195 | 0,00129 | 0,011785 | 1606,632769 | 0,688380376 | 0,195404 | 3,522865 | 0,000427 | 0,001264 | 1110,54 | 1,223461632 | 0,325541 | 3,75824 | 0,000171 | 0,001166 | 1009,692 | -0,93995 | 0,317462 | -2,96083 | 0,003068 | 0,008983 | 1731,023705 | -0,364809562 | 0,203425 | -1,79333 | 0,07292 | 0,103599 |
| 718 | RM25_RS03465 | RM25_0709 | Phospho-glucose isomerase C-terminal domain protein | 745 | gene | 820877 | 821893 | + | 1301,586 | 1,085351604 | 0,219673 | 4,940769 | 7,78E-07 | 3,33E-05 | 1502,354443 | 0,975735997 | 0,22162 | 4,402745 | 1,07E-05 | 4,48E-05 | 1100,818 | 1,240860484 | 0,201131 | 6,169418 | 6,85E-10 | 2E-08 | 883,1913 | -0,68235 | 0,26462 | -2,5786 | 0,00992 | 0,023594 | 1733,052037 | -0,375427306 | 0,13385 | -2,80483 | 0,005034 | 0,009329 |
| 719 | RM25_RS03470 | RM25_0710 | Cation diffusion facilitator family transporter | 746 | gene | 822005 | 823045 | + | 1323,879 | 0,549076462 | 0,222394 | 2,468936 | 0,013552 | 0,057324 | 1583,400521 | 0,491327844 | 0,135808 | 3,617817 | 0,000297 | 0,000915 | 1064,358 | 0,636955974 | 0,233209 | 2,73127 | 0,006309 | 0,021646 | 1139,635 | -0,72147 | 0,174233 | -4,14083 | 3,46E-05 | 0,000188 | 1539,912773 | -0,527826477 | 0,158619 | -3,32764 | 0,000876 | 0,00189 |
| 720 | RM25_RS03475 | RM25_0711 | Hypothetical protein | 747 | gene | 823201 | 823464 | - | 273,9194 | 0,75332303 | 0,201823 | 3,732598 | 0,00019 | 0,002792 | 272,8616231 | 0,786351124 | 0,240955 | 3,263472 | 0,001101 | 0,002928 | 274,9772 | 0,721075466 | 0,335623 | 2,148471 | 0,031676 | 0,072349 | 216,1452 | -0,01456 | 0,246547 | -0,05905 | 0,952915 | 0,963544 | 335,6056108 | -0,035660437 | 0,315706 | -0,11295 | 0,910067 | 0,926578 |
| 721 | RM25_RS03480 | RM25_0712 | Multidrug resistance efflux protein | 748 | gene | 823523 | 825034 | + | 1214,827 | 0,639178489 | 0,309913 | 2,062447 | 0,039165 | 0,10757 | 1579,474499 | 0,962049809 | 0,151212 | 6,362257 | 1,99E-10 | 1,87E-09 | 850,1789 | 0,076774456 | 0,237864 | 0,322767 | 0,746872 | 0,813859 | 1004,438 | -0,43477 | 0,213521 | -2,0362 | 0,04173 | 0,073346 | 1453,16697 | -1,261296268 | 0,165479 | -7,62209 | 2,5E-14 | 2,79E-13 |
| 722 | RM25_RS03485 | RM25_0713 | Hypothetical protein | 749 | gene | 825208 | 826260 | + | 1137,704 | 0,533075896 | 0,221341 | 2,408395 | 0,016023 | 0,063048 | 1330,721322 | 0,514730601 | 0,179549 | 2,866793 | 0,004147 | 0,009281 | 944,687 | 0,560933695 | 0,27451 | 2,043399 | 0,041013 | 0,087561 | 985,832 | -0,57311 | 0,255029 | -2,24723 | 0,024625 | 0,048287 | 1317,190272 | -0,488944536 | 0,187063 | -2,61379 | 0,008954 | 0,015833 |
| 723 | RM25_RS03490 | RM25_0714 | 4-alpha-glucanotransferase malQ | 750 | gene | 826391 | 828580 | + | 1883,174 | 0,356303986 | 0,250386 | 1,423019 | 0,154731 | 0,25575 | 2336,876595 | 0,541762091 | 0,136548 | 3,967565 | 7,26E-05 | 0,000256 | 1429,471 | 0,060297488 | 0,197219 | 0,305739 | 0,759803 | 0,824519 | 1748,709 | -0,50499 | 0,174213 | -2,89873 | 0,003747 | 0,010526 | 2075,314989 | -0,930722881 | 0,143258 | -6,49682 | 8,2E-11 | 5,43E-10 |
| 724 | RM25_RS03495 | RM25_0715 | O-succinylbenzoate synthase | 751 | gene | 828577 | 829575 | + | 563,4935 | 0,365526286 | 0,191078 | 1,912971 | 0,055752 | 0,1328 | 591,522225 | 0,691517647 | 0,149631 | 4,621486 | 3,81E-06 | 1,74E-05 | 535,4648 | 0,019209662 | 0,298487 | 0,064357 | 0,948686 | 0,967988 | 517,8159 | 0,172933 | 0,204872 | 0,844102 | 0,398612 | 0,474031 | 623,8881081 | -0,438482806 | 0,249509 | -1,75738 | 0,078853 | 0,111025 |
| 725 | RM25_RS03500 | RM25_0716 | 2-succinyl-5-enolpyruvyl-6-hydroxy-3-cyclohexene-1-carbox | 752 | gene | 829568 | 831355 | + | [illegible] | [illegible] | [illegible] | [illegible] | [illegible] | [illegible] | [illegible] | [illegible] | [illegible] | [illegible] | [illegible] | [illegible] | [illegible] | [illegible] | [illegible] | [illegible] | [illegible] | [illegible] | [illegible] | [illegible] | [illegible] | [illegible] | [illegible] | [illegible] | [illegible] | [illegible] | [illegible] | [illegible] | [illegible] | [illegible] | [illegible] |
| 726 | RM25_RS03505 | RM25_0717 | Excinuclease ABC, A subunit | 753 | gene | 831463 | 834413 | + | [illegible] | [illegible] | [illegible] | [illegible] | [illegible] | [illegible] | [illegible] | [illegible] | [illegible] | [illegible] | [illegible] | [illegible] | [illegible] | [illegible] | [illegible] | [illegible] | [illegible] | [illegible] | [illegible] | [illegible] | [illegible] | [illegible] | [illegible] | [illegible] | [illegible] | [illegible] | [illegible] | [illegible] | [illegible] | [illegible] | [illegible] |
| 727 | RM25_RS03510 | RM25_0718 | Hypothetical protein | 754 | gene | 834406 | 834669 | - | [illegible] | [illegible] | [illegible] | [illegible] | [illegible] | [illegible] | [illegible] | [illegible] | [illegible] | [illegible] | [illegible] | [illegible] | [illegible] | [illegible] | [illegible] | [illegible] | [illegible] | [illegible] | [illegible] | [illegible] | [illegible] | [illegible] | [illegible] | [illegible] | [illegible] | [illegible] | [illegible] | [illegible] | [illegible] | [illegible] | [illegible] |
| 728 | RM25_RS03515 | RM25_0719 | Isochorismate synthase | 755 | gene | 834665 | 836116 | - | [illegible] | [illegible] | [illegible] | [illegible] | [illegible] | [illegible] | [illegible] | [illegible] | [illegible] | [illegible] | [illegible] | [illegible] | [illegible] | [illegible] | [illegible] | [illegible] | [illegible] | [illegible] | [illegible] | [illegible] | [illegible] | [illegible] | [illegible] | [illegible] | [illegible] | [illegible] | [illegible] | [illegible] | [illegible] | [illegible] | [illegible] |
| 729 | RM25_RS03520 | RM25_0720 | O-succinylbenzoic acid--CoA ligase | 756 | gene | 836130 | 837362 | - | [illegible] | [illegible] | [illegible] | [illegible] | [illegible] | [illegible] | [illegible] | [illegible] | [illegible] | [illegible] | [illegible] | [illegible] | [illegible] | [illegible] | [illegible] | [illegible] | [illegible] | [illegible] | [illegible] | [illegible] | [illegible] | [illegible] | [illegible] | [illegible] | [illegible] | [illegible] | [illegible] | [illegible] | [illegible] | [illegible] | [illegible] |

| | | | | | | | | | | | | | | | | | | | | | | | | | | | | | | | | | | | | | | |
|---|---|---|---|---|---|---|---|---|---|---|---|---|---|---|---|---|---|---|---|---|---|---|---|---|---|---|---|---|---|---|---|---|---|---|---|---|---|---|
| 730 | RM25_RS03525 | RM25_0721 | Naphthoate synthase | 757 | gene | 837435 | 838520 | - | 1308,27 | 0,128849987 | 0,086999 | 1,481048 | 0,138594 | 0,238346 | 1254,637976 | 0,169752929 | 0,138982 | 1,2214 | 0,221934 | 0,28491 | 1361,902 | 0,091608 | 0,156061 | 0,587002 | 0,557202 | 0,653071 | 1318,126 | 0,101211 | 0,13734 | 0,736936 | 0,461162 | 0,533869 | 1338,767527 | 0,072111562 | 0,126352 | 0,57072 | 0,568189 | 0,629038 |
| 731 | RM25_RS03530 | RM25_0722 | Nicotinate phosphoribosyltransferase | 758 | gene | 838602 | 839957 | - | 801,8501 | -0,326651134 | 0,205701 | -1,58799 | 0,112288 | 0,206881 | 671,9342341 | 0,057549608 | 0,135886 | 0,423514 | 0,67192 | 0,725809 | 931,7659 | -0,60890209 | 0,212364 | -2,86726 | 0,00414 | 0,01554 | 938,1926 | 0,71739 | 0,184192 | 3,894798 | 9,83E-05 | 0,00048 | 698,1880195 | 0,102661667 | 0,175216 | 0,585914 | 0,557933 | 0,619429 |
| 732 | RM25_RS03535 | RM25_0723 | ATP-dependent Clp protease adapter protein ClpS | 759 | gene | 840084 | 840377 | + | 735,3791 | 0,836352068 | 0,229627 | 3,642213 | 0,00027 | 0,003641 | 646,9311575 | 1,011277017 | 0,171306 | 5,903326 | 3,56E-09 | 2,83E-08 | 823,827 | 0,70419962 | 0,345012 | 2,041087 | 0,041242 | 0,087971 | 558,0297 | 0,48907 | 0,301462 | 1,622328 | 0,104733 | 0,154575 | 920,9629696 | 0,222868718 | 0,237541 | 0,938231 | 0,348126 | 0,411914 |
| 733 | RM25_RS11805 | RM25_0724 | Hypothetical protein | 760 | gene | 840374 | 840982 | + | 489,5763 | -0,046459808 | 0,199322 | -0,23309 | 0,815692 | 0,871526 | 420,0637386 | 0,087631922 | 0,231854 | 0,377961 | 0,705459 | 0,756147 | 559,0889 | -0,148988895 | 0,20647 | -0,7216 | 0,470539 | 0,572862 | 523,4273 | 0,46156 | 0,252036 | 1,83133 | 0,067051 | 0,108321 | 471,9153941 | 0,281449751 | 0,189975 | 1,481511 | 0,13847 | 0,184497 |
| 734 | RM25_RS03545 | RM25_0725 | Glutamate racemase | 761 | gene | 841070 | 841888 | + | 758,9532 | 0,140468088 | 0,139945 | 1,003737 | 0,315506 | 0,424941 | 838,9088736 | 0,294938485 | 0,12605 | 2,339857 | 0,019291 | 0,034683 | 678,9975 | -0,046503618 | 0,178639 | -0,26032 | 0,794616 | 0,85212 | 762,7514 | -0,18835 | 0,139682 | -1,34846 | 0,177511 | 0,238537 | 780,5352284 | -0,477292656 | 0,137375 | -3,47439 | 0,000512 | 0,001148 |
| 735 | RM25_RS03550 | RM25_0726 | TRNA nucleotidyltransferase | 762 | gene | 842042 | 842803 | + | 1166,035 | 0,929937047 | 0,175799 | 5,289786 | 1,22E-07 | 7,6E-06 | 1215,934205 | 0,725082009 | 0,164898 | 4,39716 | 1,1E-05 | 4,58E-05 | 1116,137 | 1,164176718 | 0,274929 | 4,234466 | 2,29E-05 | 0,000211 | 851,4947 | -0,4689 | 0,213987 | -2,19127 | 0,028432 | 0,054076 | 1494,565203 | 0,010710174 | 0,21186 | 0,050553 | 0,959682 | 0,967481 |
| 736 | RM25_RS03545 | RM25_0727 | Non-canonical purine NTP pyrophosphatase | 763 | gene | 842800 | 843606 | + | 1065,652 | 0,785836111 | 0,178436 | 4,404019 | 1,06E-05 | 0,000309 | 1059,139709 | 0,426374725 | 0,132769 | 3,211405 | 0,001321 | 0,003425 | 1072,164 | 1,164872378 | 0,260902 | 4,464781 | 8,02E-06 | 8,41E-05 | 829,9057 | -0,5141 | 0,167815 | -3,0635 | 0,002188 | 0,006802 | 1317,718606 | 0,271734563 | 0,201352 | 1,349547 | 0,177161 | 0,22893 |
| 737 | RM25_RS03560 | RM25_0728 | Thymidylate synthase | 764 | gene | 843608 | 844420 | + | 1265,941 | 0,608539928 | 0,166234 | 3,660748 | 0,000251 | 0,003426 | 1358,220952 | 0,386788532 | 0,13666 | 2,830296 | 0,004651 | 0,010187 | 1173,661 | 0,874466068 | 0,237352 | 3,684262 | 0,000229 | 0,001469 | 1061,864 | -0,56837 | 0,171741 | -3,30944 | 0,000935 | 0,003328 | 1495,727259 | -0,032727973 | 0,163649 | -0,19999 | 0,84149 | 0,872204 |
| 738 | RM25_RS03565 | RM25_0729 | Dihydrofolate reductase | 765 | gene | 844535 | 845137 | + | 1990,067 | 1,478526386 | 0,376274 | 3,92939 | 8,52E-05 | 0,001487 | 1937,634435 | 0,590663618 | 0,172947 | 3,415289 | 0,000637 | 0,001801 | 2042,499 | 2,667538006 | 0,322991 | 8,258868 | 1,47E-16 | 1,39E-14 | 1118,611 | -1,53585 | 0,219448 | -6,99871 | 2,58E-12 | 6,34E-11 | 2857,330731 | 0,582106328 | 0,24345 | 2,391068 | 0,016799 | 0,028043 |
| 739 | RM25_RS03570 | RM25_0730 | DNA polymerase IV | 766 | gene | 845141 | 846592 | + | 985,8438 | 0,382858074 | 0,135029 | 2,835379 | 0,004577 | 0,028022 | 1002,561527 | 0,560226502 | 0,121978 | 4,592856 | 4,37E-06 | 1,97E-05 | 969,1261 | 0,205199326 | 0,253175 | 0,810503 | 0,417651 | 0,522785 | 901,0737 | 0,088292 | 0,198561 | 0,44466 | 0,656565 | 0,710852 | 1096,035659 | -0,205370992 | 0,183628 | -1,11841 | 0,263392 | 0,322669 |
| 740 | RM25_RS03575 | RM25_0731 | LAO/AO transport system ATPase | 767 | gene | 846796 | 847809 | - | 4392,184 | 0,628510761 | 0,521314 | 1,205628 | 0,227961 | 0,335396 | 6038,891193 | 0,294312082 | 0,230381 | 1,277501 | 0,201425 | 0,261557 | 2745,477 | 1,445356275 | 0,585042 | 2,470517 | 0,013492 | 0,038423 | 3686,934 | -1,92574 | 0,580475 | -3,31752 | 0,000908 | 0,003243 | 5219,586144 | -0,745480772 | 0,236871 | -3,14721 | 0,001648 | 0,003321 |
| 741 | RM25_RS03580 | RM25_0732 | Methylmalonyl-CoA mutase | 768 | gene | 847802 | 849988 | - | 10450,06 | 0,423296023 | 0,366339 | 1,155477 | 0,247895 | 0,35584 | 12484,19744 | 0,021650979 | 0,181893 | 0,119032 | 0,90525 | 0,928088 | 8415,918 | 1,056701613 | 0,475413 | 2,222701 | 0,026236 | 0,062972 | 9517,285 | -1,22963 | 0,450782 | -2,72778 | 0,006376 | 0,016212 | 11703,48827 | -0,162800467 | 0,231861 | -0,70215 | 0,482588 | 0,544761 |
| 742 | RM25_RS03585 | RM25_0733 | Methylmalonyl-CoA mutase, small subunit | 769 | gene | 849985 | 851901 | - | 7074,024 | 0,342063536 | 0,309379 | 1,105647 | 0,268879 | 0,376783 | 7931,947201 | 0,137627916 | 0,183304 | 0,750818 | 0,452762 | 0,521075 | 6216,101 | 0,608610954 | 0,484318 | 1,256634 | 0,208886 | 0,305425 | 6633,414 | -0,66444 | 0,4578 | -1,45137 | 0,146677 | 0,203005 | 7729,274957 | -0,163930679 | 0,239249 | -0,68519 | 0,493224 | 0,554907 |
| 743 | NA | RM25_0734 | Hypothetical protein | 770 | gene | 851898 | 852161 | - | 905,0342 | 0,365100276 | 0,277958 | 1,31351 | 0,189011 | 0,293267 | 875,4820947 | 0,531179586 | 0,251517 | 2,111905 | 0,034695 | 0,058079 | 934,5863 | 0,213292233 | 0,442295 | 0,48224 | 0,629635 | 0,7136 | 836,615 | 0,22668 | 0,444256 | 0,510246 | 0,609879 | 0,668909 | 995,8729373 | -0,060311112 | 0,24802 | -0,24317 | 0,807874 | 0,843668 |
| 744 | RM25_RS03590 | RM25_0735 | Precorrin-2 C20-methyltransferase CbiL | 771 | gene | 852736 | 853509 | + | 1279,807 | 0,468495073 | 0,278483 | 1,682311 | 0,092509 | 0,183704 | 1606,303893 | 0,402210972 | 0,202817 | 1,983119 | 0,047354 | 0,07567 | 953,3105 | 0,583241006 | 0,213969 | 2,725821 | 0,006414 | 0,021865 | 1139,874 | -0,9154 | 0,258996 | -3,53443 | 0,000409 | 0,001672 | 1457,40228 | -0,684705119 | 0,141657 | -4,83354 | 1,34E-06 | 4,57E-06 |
| 745 | RM25_RS03595 | RM25_0736 | Tetrapyrrole methylase, precorrin-3 methylase, CbiF | 772 | gene | 853539 | 854498 | + | 1288,955 | 0,164178098 | 0,280013 | 0,586322 | 0,557659 | 0,651985 | 1639,060057 | 0,076535228 | 0,161501 | 0,473898 | 0,635572 | 0,693193 | 938,8499 | 0,319055822 | 0,203101 | 1,570922 | 0,116201 | 0,193832 | 1291,255 | -0,99526 | 0,216816 | -4,59034 | 4,43E-06 | 2,95E-05 | 1335,989385 | -0,698199981 | 0,110586 | -6,31366 | 2,73E-10 | 1,63E-09 |
| 746 | RM25_RS03600 | RM25_0737 | Bifunctional cbiH protein and precorrin-3B C17-methyltrans | 773 | gene | 854495 | 857140 | + | 2829,737 | 0,201331565 | 0,24543 | 0,820322 | 0,412032 | 0,522129 | 3446,416181 | 0,136795885 | 0,186956 | 0,731701 | 0,464351 | 0,532332 | 2213,057 | 0,302681093 | 0,219363 | 1,37982 | 0,167642 | 0,257158 | 2787,554 | -0,78728 | 0,228677 | -3,44275 | 0,000576 | 0,002213 | 2969,893153 | -0,568655522 | 0,155466 | -3,65775 | 0,000254 | 0,000603 |
| 747 | RM25_RS03605 | RM25_0738 | Sirohydrochlorin cobaltochelatase | 774 | gene | 857133 | 858374 | + | 1116,966 | 0,266855038 | 0,235224 | 1,13447 | 0,256598 | 0,364338 | 1330,65381 | 0,185464232 | 0,208928 | 0,887695 | 0,374705 | 0,443586 | 903,2782 | 0,388798552 | 0,229212 | 1,696236 | 0,089841 | 0,158571 | 1073,736 | -0,72768 | 0,27062 | -2,68896 | 0,007168 | 0,017953 | 1196,577891 | -0,473154557 | 0,137752 | -3,43483 | 0,000593 | 0,001315 |
| 748 | RM25_RS03610 | RM25_0739 | Putative cobalt-precorrin-6A synthase | 775 | gene | 858406 | 859698 | - | 751,6092 | -0,047347305 | 0,137667 | -0,34393 | 0,730902 | 0,804265 | 709,7119877 | 0,086516127 | 0,147398 | 0,586955 | 0,557234 | 0,621875 | 793,5065 | -0,166656779 | 0,239414 | -0,6961 | 0,486364 | 0,586673 | 803,6205 | 0,22386 | 0,19379 | 1,155167 | 0,248022 | 0,317193 | 725,8807549 | 0,027908158 | 0,187168 | 0,149107 | 0,881469 | 0,902779 |
| 749 | RM25_RS03615 | RM25_0740 | Precorrin-8X methylmutase | 776 | gene | 859695 | 860348 | - | 574,9714 | 0,614308658 | 0,120698 | 5,089622 | 3,59E-07 | 1,8E-05 | 547,736142 | 0,487567974 | 0,162548 | 2,999524 | 0,002704 | 0,006418 | 602,2067 | 0,730312319 | 0,208694 | 3,499443 | 0,000466 | 0,002661 | 480,1445 | -0,06983 | 0,173745 | -0,40191 | 0,687749 | 0,735826 | 680,2302733 | 0,221105068 | 0,166338 | 1,329255 | 0,183764 | 0,236296 |
| 750 | RM25_RS03620 | RM25_0741 | Precorrin-6Y C5, 15-methyltransferase, CbiT subunit | 777 | gene | 860445 | 861035 | - | 632,3601 | 0,002683164 | 0,16576 | 0,016187 | 0,987085 | 0,99256 | 552,6828307 | 0,2169198 | 0,160571 | 1,350928 | 0,176719 | 0,233741 | 712,0374 | -0,164607607 | 0,17821 | -0,92367 | 0,355656 | 0,462628 | 664,8979 | 0,495155 | 0,180387 | 2,744954 | 0,006052 | 0,015505 | 619,8139812 | 0,163102079 | 0,140531 | 1,160613 | 0,245799 | 0,303483 |
| 751 | RM25_RS03625 | RM25_0742 | Precorrin-6x reductase CbiJ/CobK | 778 | gene | 861108 | 861875 | - | 263,0354 | -0,537139121 | 0,33195 | -1,61813 | 0,105634 | 0,199663 | 193,3259602 | -0,61799998 | 0,242419 | -2,54931 | 0,010794 | 0,021201 | 332,7449 | -0,494780211 | 0,390216 | -1,26797 | 0,20481 | 0,3019 | 326,5568 | 0,667351 | 0,287783 | 2,318935 | 0,020399 | 0,041519 | 211,2271889 | 0,858190379 | 0,367001 | 2,338389 | 0,019367 | 0,031899 |
| 752 | RM25_RS03630 | RM25_0743 | Putative L-ribulose-5-phosphate 4-epimerase | 779 | gene | 861856 | 862548 | - | 572,3665 | -0,333451376 | 0,127538 | -2,61453 | 0,008935 | 0,043245 | 594,2844098 | -0,259041211 | 0,146792 | -1,76469 | 0,077617 | 0,113917 | 550,4486 | -0,412355104 | 0,241821 | -1,7052 | 0,088156 | 0,155994 | 673,0989 | -0,10565 | 0,20316 | -0,52003 | 0,603041 | 0,663261 | 497,6384877 | -0,20021519 | 0,187196 | -1,06955 | 0,284823 | 0,346045 |
| 754 | RM25_RS03640 | RM25_0744 | Bacterioferritin comigratory protein bcp | 783 | gene | 862781 | 863269 | - | 641,0093 | 0,214923188 | 0,129864 | 1,654988 | 0,097927 | 0,190598 | 592,642047 | 0,40045848 | 0,141155 | 2,837016 | 0,004554 | 0,010022 | 689,3765 | 0,058971815 | 0,185609 | 0,31772 | 0,750697 | 0,815765 | 624,9385 | 0,342315 | 0,161247 | 2,122922 | 0,03376 | 0,062362 | 674,6394718 | 0,050436314 | 0,144358 | 0,349384 | 0,726801 | 0,773046 |
| 755 | RM25_RS12705 | RM25_0745 | Hypothetical protein | 784 | gene | 863443 | 863742 | + | 2070,106 | 0,120743073 | 0,278596 | 0,433398 | 0,664726 | 0,746787 | 2244,301183 | 0,216836843 | 0,22071 | 0,98245 | 0,325878 | 0,393498 | 1895,911 | 0,008231996 | 0,449389 | 0,018318 | 0,985385 | 0,991934 | 2103,017 | -0,18014 | 0,424234 | -0,42462 | 0,671114 | 0,723281 | 2108,576921 | -0,360799575 | 0,264113 | -1,36608 | 0,171914 | 0,224006 |
| 756 | RM25_RS03650 | RM25_0746 | 10 kDa chaperonin | 785 | gene | 863769 | 864218 | + | 1231,034 | 0,483125774 | 0,402257 | 1,201039 | 0,229736 | 0,33729 | 1666,920549 | 0,365497871 | 0,227322 | 1,607841 | 0,10787 | 0,152063 | 795,1483 | 0,738626481 | 0,395435 | 1,867885 | 0,061778 | 0,118769 | 1093,836 | -1,33687 | 0,410914 | -3,25341 | 0,00114 | 0,003947 | 1406,616709 | -0,928957277 | 0,194718 | -4,77078 | 1,84E-06 | 6,07E-06 |
| 758 | RM25_RS03660 | RM25_0748 | Cobyric acid synthase cobQ2 | 789 | gene | 864480 | 865253 | - | 731,5247 | 0,407016527 | 0,161653 | 2,517836 | 0,011808 | 0,051924 | 822,4960529 | 0,300556584 | 0,150786 | 1,993265 | 0,046232 | 0,074433 | 640,5533 | 0,54570724 | 0,182994 | 2,98211 | 0,002863 | 0,011593 | 666,4298 | -0,56029 | 0,147158 | -3,80739 | 0,00014 | 0,000658 | 816,7399414 | -0,266449815 | 0,14955 | -1,78168 | 0,074802 | 0,105955 |
| 759 | RM25_RS03665 | RM25_0749 | Mur ligase middle domain protein | 790 | gene | 865247 | 866569 | - | 1227,607 | 0,323389806 | 0,099509 | 3,249858 | 0,001155 | 0,011018 | 1154,050388 | 0,347468207 | 0,15227 | 2,281918 | 0,022494 | 0,039596 | 1301,163 | 0,301956586 | 0,141357 | 2,136133 | 0,032669 | 0,073896 | 1148,951 | 0,142383 | 0,144171 | 0,987598 | 0,323349 | 0,398606 | 1337,233624 | 0,14545353 | 0,119638 | 1,215784 | 0,224067 | 0,280322 |
| 760 | RM25_RS03670 | RM25_0750 | Protein tyrosine/serine phosphatase | 791 | gene | 866673 | 867488 | - | 419,4411 | -1,056592704 | 0,548029 | -1,92799 | 0,053857 | 0,130463 | 233,1981307 | -2,073006958 | 0,334698 | -6,19366 | 5,88E-10 | 5,19E-09 | 605,6841 | -0,734287431 | 0,361296 | -2,03237 | 0,042116 | 0,08911 | 593,3792 | 0,938671 | 0,357323 | 2,626954 | 0,008615 | 0,020913 | 268,089439 | 2,345106317 | 0,359524 | 6,522803 | 6,9E-11 | 4,61E-10 |
| 761 | RM25_RS03675 | RM25_0751 | Daunorubicin resistance protein | 792 | gene | 867559 | 869892 | - | 1332,512 | -0,461475648 | 0,349723 | -1,31955 | 0,186986 | 0,291293 | 910,7193738 | -0,931351578 | 0,205436 | -4,53353 | 5,8E-06 | 2,56E-05 | 1754,304 | -0,229802995 | 0,209966 | -1,09448 | 0,273745 | 0,376659 | 1621,782 | 0,598358 | 0,260555 | 2,296472 | 0,021649 | 0,0436 | 1097,988466 | 1,35789867 | 0,132573 | 10,24264 | 1,28E-24 | 4,51E-23 |
| 762 | RM25_RS03680 | RM25_0752 | putative GTP pyrophosphokinase | 793 | gene | 869967 | 870473 | + | 200,5928 | -0,838978271 | 0,424953 | -1,97429 | 0,048349 | 0,122012 | 119,1186623 | -0,884613062 | 0,317047 | -2,79017 | 0,005268 | 0,011487 | 282,0669 | -0,823421926 | 0,330745 | -2,4896 | 0,012789 | 0,036895 | 269,3784 | 1,15755 | 0,276829 | 4,181471 | 2,9E-05 | 0,000161 | 141,4997406 | 1,282129355 | 0,376583 | 3,404635 | 0,000663 | 0,001464 |
| 764 | RM25_RS03690 | RM25_0753 | Oligoribonuclease | 797 | gene | 871199 | 871879 | - | 4006,255 | 0,674880049 | 0,397798 | 1,69654 | 0,089784 | 0,180563 | 5553,444382 | 0,792543157 | 0,19664 | 4,030422 | 5,57E-05 | 0,000203 | 2459,065 | 0,417392303 | 0,369788 | 1,128734 | 0,25901 | 0,360382 | 3283,606 | -0,999 | 0,316069 | -3,1607 | 0,001574 | 0,00516 | 4825,346842 | -1,343317873 | 0,257316 | -5,22049 | 1,78E-07 | 6,96E-07 |
| 765 | RM25_RS03695 | RM25_0754 | High-affinity glucose transporter | 798 | gene | 871891 | 873342 | - | 758,5214 | -0,312121336 | 0,170418 | -1,83151 | 0,067025 | 0,149458 | 790,8883613 | -0,132449745 | 0,149567 | -0,88556 | 0,375856 | 0,444503 | 726,1544 | -0,510217445 | 0,293652 | -1,73749 | 0,0823 | 0,148255 | 887,23 | -0,01954 | 0,178912 | -0,10921 | 0,91304 | 0,93109 | 665,4714019 | -0,334077531 | 0,270671 | -1,23425 | 0,217108 | 0,273211 |
| 767 | RM25_RS03705 | RM25_0756 | Membrane protein-like protein | 802 | gene | 873396 | 874898 | + | 1146,528 | -0,207746727 | 0,075521 | -2,75086 | 0,005944 | 0,033356 | 1110,104003 | -0,160319708 | 0,126739 | -1,26496 | 0,205887 | 0,266057 | 1182,952 | -0,251684492 | 0,157305 | -1,59997 | 0,109604 | 0,184923 | 1296,954 | 0,07154 | 0,13589 | 0,526451 | 0,598575 | 0,659579 | 1043,632563 | 0,036887663 | 0,140202 | 0,263104 | 0,792471 | 0,831636 |
| 768 | RM25_RS03710 | RM25_0757 | ATP/GTP-binding family protein | 803 | gene | 874968 | 876815 | + | 847,3569 | 0,035794313 | 0,203707 | 0,175715 | 0,860518 | 0,907665 | 760,7941382 | 0,206602571 | 0,169955 | 1,215629 | 0,224126 | 0,286789 | 933,9196 | -0,10169729 | 0,318003 | -0,3198 | 0,74912 | 0,814802 | 878,1617 | 0,394462 | 0,220577 | 1,788316 | 0,073725 | 0,116079 | 843,7307121 | 0,147126237 | 0,267911 | 0,54916 | 0,582896 | 0,643205 |
| 769 | RM25_RS03715 | RM25_0758 | ATP/GTP-binding family protein | 804 | gene | 876812 | 878455 | + | 908,3996 | 0,285282473 | 0,175316 | 1,62725 | 0,103684 | 0,197083 | 1029,411002 | 0,412423059 | 0,161309 | 2,556731 | 0,010566 | 0,02084 | 787,3882 | 0,123428059 | 0,217654 | 0,567083 | 0,570658 | 0,663235 | 864,3968 | -0,28703 | 0,2008 | -1,42943 | 0,152882 | 0,210241 | 979,538906 | -0,521294155 | 0,159655 | -3,26514 | 0,001094 | 0,002288 |
| 770 | RM25_RS11810 | RM25_0759 | Hypothetical protein | 805 | gene | 878452 | 879114 | + | 970,2488 | 0,59530957 | 0,364103 | 1,635003 | 0,102048 | 0,195234 | 1278,189769 | 0,383232733 | 0,18455 | 2,076575 | 0,037841 | 0,06259 | 662,3079 | 1,029272971 | 0,311067 | 3,308846 | 0,000937 | 0,004619 | 822,2212 | -1,4039 | 0,311234 | -4,51076 | 6,46E-06 | 4,16E-05 | 1144,120068 | -0,714223828 | 0,152055 | -4,69713 | 2,64E-06 | 8,53E-06 |
| 771 | RM25_RS03725 | RM25_0760 | Putative insertion element ISCmi2 transposase | 806 | gene | 879132 | 880175 | - | 56,64451 | -0,061440753 | 0,240956 | -0,25499 | 0,798733 | 0,859249 | 51,14709481 | -0,616992976 | 0,33311 | -1,85222 | 0,063994 | 0,096442 | 62,14193 | 0,365973254 | 0,281604 | 1,299601 | 0,193738 | 0,289563 | 61,36114 | -0,2542 | 0,234265 | -1,0851 | 0,277877 | 0,351186 | 53,96603964 | 0,780217027 | 0,365995 | 2,131769 | 0,033026 | 0,051144 |
| 772 | RM25_RS03730 | RM25_0761 | Hypothetical protein | 807 | gene | 880429 | 880929 | + | 225,3221 | -1,167560604 | 0,639444 | -1,8259 | 0,067865 | 0,150478 | 108,9208929 | -2,462039405 | 0,379771 | -6,48295 | 8,99E-11 | 8,8E-10 | 341,7234 | -0,848871981 | 0,428608 | -1,98053 | 0,047644 | 0,09748 | 326,2097 | 1,185108 | 0,32464 | 3,650535 | 0,000262 | 0,001136 | 136,9471468 | 2,870170271 | 0,487948 | 5,882118 | 4,05E-09 | 2,02E-08 |
| 773 | RM25_RS03735 | RM25_0762 | ATP-binding cassette protein, ChvD family | 808 | gene | 881125 | 882813 | + | 3312,865 | 0,808084349 | 0,441689 | 1,829531 | 0,06732 | 0,149773 | 4373,30774 | 0,359881132 | 0,203718 | 1,766564 | 0,077301 | 0,113591 | 2252,422 | 1,835345289 | 0,297292 | 6,173542 | 6,68E-10 | 2E-08 | 2574,525 | -2,016 | 0,254555 | -7,91969 | 2,38E-15 | 9,68E-14 | 4126,09316 | -0,498878135 | 0,230938 | -2,16023 | 0,030755 | 0,048326 |
| 774 | RM25_RS03740 | RM25_0763 | Dihydroxyacetone kinase, DhaK subunit | 809 | gene | 882985 | 883980 | + | 1499,787 | -0,656867188 | 0,430263 | -1,52666 | 0,126845 | 0,224791 | 934,9068463 | 0,554660202 | 0,138806 | 3,995947 | 6,44E-05 | 0,00023 | 2064,667 | -1,260184588 | 0,338967 | -3,71772 | 0,000201 | 0,001321 | 1924,016 | 1,894664 | 0,324279 | 5,8427 | 5,14E-09 | 6E-08 | 1138,817351 | 0,114960924 | 0,1625 | 0,70745 | 0,479287 | 0,541728 |
| 775 | RM25_RS03745 | RM25_0764 | Dihydroxyacetone kinase, L subunit | 810 | gene | 883980 | 884609 | + | 1019,33 | -0,431410556 | 0,337098 | -1,27978 | 0,200623 | 0,303669 | 747,6860933 | 0,461618028 | 0,144016 | 3,205325 | 0,001349 | 0,003479 | 1290,973 | -0,976198487 | 0,275845 | -3,53894 | 0,000402 | 0,002361 | 1229,036 | 1,391573 | 0,274437 | 5,070641 | 3,96E-07 | 3,26E-06 | 850,308461 | -0,004332268 | 0,145967 | -0,02968 | 0,976322 | 0,981737 |
| 776 | RM25_RS03750 | RM25_0765 | Dihydroxyacetone kinase subunit M | 811 | gene | 884622 | 886142 | + | 1064,342 | -0,388962412 | 0,241723 | -1,60913 | 0,107589 | 0,201419 | 830,7612769 | 0,008658487 | 0,140283 | 0,061722 | 0,950785 | 0,960977 | 1297,922 | -0,650013006 | 0,20377 | -3,18994 | 0,001423 | 0,0065 | 1268,397 | 0,877048 | 0,158951 | 5,517728 | 3,43E-08 | 3,46E-07 | 904,4948484 | 0,273364372 | 0,193055 | 1,415995 | 0,156777 | 0,206322 |
| 777 | RM25_RS03755 | RM25_0766 | UTP--hexose-1-phosphate uridylyltransferase | 812 | gene | 886129 | 887499 | - | 1270,069 | 0,108506331 | 0,061453 | 1,765679 | 0,07745 | 0,164837 | 1252,259029 | 0,096907716 | 0,12336 | 0,785568 | 0,432121 | 0,500495 | 1287,88 | 0,120535569 | 0,148897 | 0,809524 | 0,418214 | 0,522934 | 1290,183 | -0,031 | 0,117005 | -0,26491 | 0,791078 | 0,830174 | 1291,27035 | 0,042383914 | 0,117428 | 0,360935 | 0,718148 | 0,76557 |
| 778 | RM25_RS03760 | RM25_0767 | Peptidyl-dipeptidase Dcp | 813 | gene | 887556 | 889592 | + | 2046,379 | 0,313546499 | 0,120124 | 2,610195 | 0,009049 | 0,043611 | 1856,506602 | 0,321548875 | 0,124924 | 2,573958 | 0,010054 | 0,019968 | 2236,251 | 0,307019757 | 0,16799 | 1,82761 | 0,067608 | 0,127789 | 1922,71 | 0,219244 | 0,12048 | 1,819753 | 0,068797 | 0,110357 | 2219,801201 | 0,250750919 | 0,121811 | 2,058522 | 0,03954 | 0,060126 |
| 779 | RM25_RS03765 | RM25_0768 | FAD linked oxidase domain protein | 814 | gene | 889713 | 892541 | - | 2510,251 | 0,231150003 | 0,227423 | 1,016386 | 0,309446 | 0,417973 | 3037,07097 | 0,406747173 | 0,13635 | 2,983112 | 0,002853 | 0,006699 | 1983,43 | -0,033595465 | 0,194782 | -0,17248 | 0,863062 | 0,90018 | 2443,775 | -0,44277 | 0,186896 | -2,36907 | 0,017833 | 0,037395 | 2660,308395 | -0,826757677 | 0,120922 | -6,83713 | 8,08E-12 | 6,08E-11 |
| 780 | RM25_RS03770 | RM25_0769 | Transcriptional regulator, MarR family | 815 | gene | 892547 | 893005 | - | 838,9744 | 0,021456543 | 0,256226 | 0,083741 | 0,933263 | 0,954451 | 638,4018406 | 0,036394229 | 0,14827 | 0,24546 | 0,8061 | 0,846633 | 1039,547 | 0,011086319 | 0,254238 | 0,043606 | 0,965218 | 0,979767 | 877,2266 | 0,657418 | 0,23189 | 2,835039 | 0,004582 | 0,012286 | 826,2598695 | 0,676943963 | 0,172848 | 3,91641 | 8,99E-05 | 0,000231 |
| 781 | RM25_RS03775 | RM25_0770 | Long-chain-fatty-acid--CoA ligase | 816 | gene | 893171 | 895060 | + | 4643,729 | 0,176989715 | 0,238998 | 0,740548 | 0,458967 | 0,56343 | 5286,8017 | 0,256702265 | 0,188995 | 1,358251 | 0,174384 | 0,231172 | 4000,656 | 0,072541462 | 0,353433 | 0,205248 | 0,837378 | 0,881116 | 4622,273 | -0,35394 | 0,330251 | -1,07174 | 0,283839 | 0,357376 | 4821,000797 | -0,505821555 | 0,214976 | -2,35293 | 0,018626 | 0,030809 |
| 782 | RM25_RS03780 | RM25_0771 | Acyl-CoA thioester hydrolase, YbgC/YbaW family | 817 | gene | 895066 | 895947 | + | 899,9967 | 0,237543612 | 0,199246 | 1,192212 | 0,233178 | 0,340521 | 1057,752161 | 0,291427184 | 0,16383 | 1,778834 | 0,075267 | 0,111225 | 742,2413 | 0,163566803 | 0,200719 | 0,814903 | 0,415128 | 0,520179 | 875,5248 | -0,50236 | 0,210374 | -2,38792 | 0,016944 | 0,035915 | 954,6840711 | -0,579401102 | 0,138381 | -4,18699 | 2,83E-05 | 7,8E-05 |
| 783 | RM25_RS03785 | RM25_0772 | Phosphomethylpyrimidine kinase | 818 | gene | 896094 | 896915 | - | 1321,152 | 1,33298318 | 0,158081 | 8,43229 | 3,39E-17 | 2,66E-14 | 1399,262607 | 1,306297837 | 0,159434 | 8,193349 | 2,54E-16 | 4,87E-15 | 1243,041 | 1,363697403 | 0,274045 | 4,976176 | 6,49E-07 | 9,41E-06 | 795,9228 | -0,27314 | 0,205247 | -1,3308 | 0,183255 | 0,244582 | 1848,826114 | -0,171944071 | 0,214535 | -0,80147 | 0,422857 | 0,488806 |
| 784 | RM25_RS03790 | RM25_0773 | Phosphogluconate dehydrogenase (decarboxylating) | 819 | gene | 897045 | 898502 | - | 2683,47 | 0,48950907 | 0,177188 | 2,762647 | 0,005733 | 0,032407 | 2676,767728 | 0,593159628 | 0,149909 | 3,956791 | 7,6E-05 | 0,000266 | 2690,171 | 0,388379007 | 0,325181 | 1,194347 | 0,232342 | 0,330695 | 2361,441 | 0,073423 | 0,267576 | 0,274402 | 0,783775 | 0,824714 | 3063,08999 | -0,095035971 | 0,219897 | -0,43218 | 0,665608 | 0,718332 |
| 785 | RM25_RS03795 | RM25_0774 | Aminopeptidase N | 820 | gene | 898715 | 901318 | - | 5149,717 | 0,311968392 | 0,242707 | 1,28537 | 0,198663 | 0,301512 | 5964,327784 | 0,071169196 | 0,15251 | 0,466652 | 0,640749 | 0,696929 | 4335,106 | 0,651660894 | 0,304091 | 2,142983 | 0,032114 | 0,072993 | 4880,045 | -0,83975 | 0,265129 | -3,16733 | 0,001538 | 0,005057 | 5580,192489 | -0,221060445 | 0,188251 | -1,17428 | 0,240281 | 0,297918 |
| 786 | RM25_RS03800 | RM25_0775 | DSBA oxidoreductase | 821 | gene | 901661 | 902275 | + | 2081,017 | 0,877179898 | 0,393568 | 2,228786 | 0,025828 | 0,082601 | 1360,654791 | 0,905643824 | 0,163138 | 5,551399 | 2,83E-08 | 1,9E-07 | 2801,379 | 0,864491943 | 0,445128 | 1,942121 | 0,052123 | 0,103848 | 1545,999 | 1,017313 | 0,395745 | 2,570625 | 0,010152 | 0,023999 | 2626,051236 | 1,00927913 | 0,259698 | 3,886353 | 0,000102 | 0,000258 |
| 787 | RM25_RS03805 | RM25_0776 | Hypothetical protein | 822 | gene | 902500 | 903294 | + | 2197,513 | 0,78934353 | 0,239438 | 3,296648 | 0,000978 | 0,009772 | 2461,853125 | 0,650747365 | 0,186253 | 3,493893 | 0,000476 | 0,001389 | 1933,174 | 0,972039724 | 0,35115 | 2,76816 | 0,005637 | 0,019714 | 1709,894 | -0,60354 | 0,306758 | -1,96747 | 0,049129 | 0,08379 | 2721,43094 | -0,249586989 | 0,235358 | -1,06045 | 0,288938 | 0,349603 |
| 788 | RM25_RS03810 | RM25_0777 | DNA ligase | 823 | gene | 903371 | 905665 | + | 1969,004 | -0,086299578 | 0,128841 | -0,66981 | 0,502976 | 0,600097 | 1997,020862 | 0,215212147 | 0,10993 | 1,957728 | 0,050262 | 0,079242 | 1940,986 | -0,397056926 | 0,172788 | -2,29794 | 0,021565 | 0,054304 | 2136,387 | 0,194335 | 0,143261 | 1,356505 | 0,174938 | 0,23576 | 1874,600494 | -0,362749083 | 0,126224 | -2,87385 | 0,004055 | 0,007592 |
| 789 | RM25_RS03815 | RM25_0778 | Hypothetical protein | 824 | gene | 905662 | 907185 | - | 644,1531 | -0,537391676 | 0,3051 | -1,76136 | 0,078177 | 0,165486 | 491,5309055 | -0,599659754 | 0,171178 | -3,50314 | 0,00046 | 0,00135 | 796,7752 | -0,501486378 | 0,393802 | -1,27345 | 0,202859 | 0,30003 | 800,728 | 0,592238 | 0,231475 | 2,558541 | 0,010511 | 0,024659 | 517,7578963 | 0,759598811 | 0,355725 | 2,135355 | 0,032732 | 0,050756 |
| 790 | NA | RM25_0779 | Hypothetical protein | 825 | gene | 907420 | 907611 | - | 96,14389 | -1,61934366 | 0,769319 | -2,10491 | 0,035299 | 0,100242 | 46,14057191 | -3,793791162 | 0,626888 | -6,05178 | 1,43E-09 | 1,2E-08 | 146,1472 | -1,213560274 | 0,477679 | -2,54054 | 0,011068 | 0,032981 | 152,1222 | 1,168669 | 0,423784 | 2,7577 | 0,005821 | 0,014995 | 46,54547086 | 3,826493403 | 0,713543 | 5,362668 | 8,2E-08 | 3,41E-07 |
| 791 | RM25_RS03825 | RM25_0780 | Multidrug-transport integral membrane protein Mmr-like pr | 826 | gene | 907739 | 908053 | + | 375,21 | 0,442093107 | 0,41062 | 1,076648 | 0,281638 | 0,390482 | 530,1013821 | 0,401890913 | 0,232072 | 1,731753 | 0,083317 | 0,121447 | 220,3186 | 0,548381873 | 0,250924 | 2,185451 | 0,028856 | 0,067443 | 339,16 | -1,40858 | 0,282454 | -4,98691 | 6,14E-07 | 4,84E-06 | 423,5945172 | -1,219772513 | 0,195171 | -6,24976 | 4,11E-10 | 2,4E-09 |
| 794 | RM25_RS03840 | RM25_0782 | Trigger factor | 833 | gene | 908788 | 910371 | + | 10671,28 | 0,770760483 | 0,538829 | 1,430436 | 0,152592 | 0,25346 | 15405,59467 | 0,42291085 | 0,204599 | 2,06702 | 0,038732 | 0,063841 | 5936,96 | 1,828486293 | 0,418401 | 4,370177 | 1,24E-05 | 0,000124 | 8448,239 | -2,38499 | 0,400875 | -5,94946 | 2,69E-09 | 3,39E-08 | 13173,39761 | -0,947001028 | 0,229767 | -4,12158 | 3,76E-05 | 0,000102 |
| 795 | RM25_RS03845 | RM25_0783 | Hypothetical protein | 834 | gene | 910596 | 911789 | + | 1661,581 | 0,697350061 | 0,215427 | 3,237059 | 0,001208 | 0,011314 | 1892,308144 | 0,490506723 | 0,185407 | 2,645564 | 0,008155 | 0,016674 | 1430,854 | 0,98258319 | 0,226611 | 4,335994 | 1,45E-05 | 0,000144 | 1343,031 | -0,7673 | 0,206808 | -3,71023 | 0,000207 | 0,000924 | 2011,453869 | -0,232033973 | 0,169807 | -1,36646 | 0,171794 | 0,224006 |
| 796 | RM25_RS03850 | RM25_0784 | Pheromone autoinducer 2 transporter | 835 | gene | 912022 | 913254 | + | 987,2275 | -0,189402393 | 0,071315 | -2,65587 | 0,00791 | 0,040096 | 991,9547989 | -0,159648591 | 0,132715 | -1,20295 | 0,228997 | 0,292228 | 982,5001 | -0,217736052 | 0,160681 | -1,35508 | 0,175392 | 0,266709 | 1109,366 | -0,04527 | 0,14041 | -0,32241 | 0,747145 | 0,79147 | 904,4945341 | -0,051576989 | 0,114072 | -0,45214 | 0,651166 | 0,70621 |
| 797 | RM25_RS03855 | RM25_0785 | ATP-dependent Clp protease proteolytic subunit 2 | 836 | gene | 913341 | 913976 | + | 2447,977 | 1,10973284 | 0,336208 | 3,300733 | 0,000964 | 0,009713 | 3269,519769 | 1,339490544 | 0,139903 | 9,574457 | 1,02E-21 | 4,08E-20 | 1626,435 | 0,68501304 | 0,332343 | 2,06116 | 0,039288 | 0,0848 | 1646,42 | -0,62406 | 0,26099 | -2,39114 | 0,016796 | 0,035633 | 3276,548253 | -1,242815186 | 0,226665 | -5,48305 | 4,18E-08 | 1,83E-07 |
| 798 | RM25_RS03860 | RM25_0786 | ATP-dependent Clp protease proteolytic subunit 1 | 837 | gene | 914000 | 914689 | + | 2338,946 | 1,277633514 | 0,378435 | 3,3761 | 0,000735 | 0,008252 | 3217,371978 | 1,294554185 | 0,169594 | 7,633272 | 2,29E-14 | 3,64E-13 | 1460,52 | 1,242831977 | 0,323264 | 3,844632 | 0,000121 | 0,000865 | 1454,238 | -1,15387 | 0,278473 | -4,14356 | 3,42E-05 | 0,000187 | 3244,375813 | -1,171074814 | 0,217254 | -5,39036 | 7,03E-08 | 2,98E-07 |
| 799 | RM25_RS03865 | RM25_0787 | ATP-dependent Clp protease ATP-binding subunit ClpX | 838 | gene | 914811 | 916088 | + | 2352,189 | 1,071229927 | 0,190994 | 5,608697 | 2,04E-08 | 1,6E-06 | 2615,256615 | 0,955207018 | 0,137846 | 6,929525 | 4,22E-12 | 5,16E-11 | 2089,122 | 1,222216151 | 0,276902 | 4,413897 | 1,02E-05 | 0,000103 | 1609,95 | -0,56169 | 0,204498 | -2,74668 | 0,00602 | 0,01544 | 3116,462384 | -0,25511412 | 0,202066 | -1,26253 | 0,206758 | 0,261302 |
| 801 | RM25_RS03870 | RM25_0788 | Hypothetical protein | 839 | gene | 916200 | 916544 | - | 149,1072 | -0,396517733 | 0,25863 | -1,53314 | 0,12524 | 0,222786 | 123,0157107 | -0,526391032 | 0,277453 | -1,89723 | 0,057798 | 0,088691 | 175,1987 | -0,310903635 | 0,318583 | -0,9759 | 0,329116 | 0,435068 | 178,3372 | 0,362139 | 0,205998 | 1,757972 | 0,078752 | 0,121797 | 126,4464969 | 0,635980173 | 0,377967 | 1,682634 | 0,092446 | 0,128174 |
| 802 | RM25_RS03875 | RM25_0789 | Threonine synthase | 840 | gene | 916633 | 918075 | + | 1733,703 | 0,276362698 | 0,136402 | 2,026096 | 0,042755 | 0,113102 | 1864,080606 | 0,419952923 | 0,131748 | 3,187545 | 0,001435 | 0,003659 | 1603,325 | 0,112766975 | 0,218869 | 0,515226 | 0,606395 | 0,694834 | 1657,55 | -0,10587 | 0,170418 | -0,62123 | 0,534451 | 0,601003 | 1859,345589 | -0,370333115 | 0,157385 | -2,35303 | 0,018621 | 0,030809 |
| 803 | RM25_RS03880 | RM25_0790 | 6-phosphofructokinase 2 | 841 | gene | 918154 | 919275 | + | 1725,118 | 0,456910257 | 0,175728 | 2,600101 | 0,00932 | 0,044377 | 1960,803836 | 0,385860523 | 0,152477 | 2,530613 | 0,011386 | 0,022052 | 1489,433 | 0,551775236 | 0,206867 | 2,667289 | 0,007647 | 0,024997 | 1538,664 | -0,55277 | 0,189344 | -2,91938 | 0,003507 | 0,009996 | 1955,048069 | -0,338189033 | 0,122685 | -2,75655 | 0,005841 | 0,01069 |
| 804 | RM25_RS03885 | RM25_0791 | Hydrolase | 842 | gene | 919402 | 920109 | + | 606,1855 | 0,457213 | 0,22123 | 2,066682 | 0,038764 | 0,106987 | 721,7987442 | 0,331000459 | 0,177278 | 1,867131 | 0,061883 | 0,093619 | 490,5723 | 0,647189304 | 0,17648 | 3,667206 | 0,000245 | 0,001545 | 542,9483 | -0,80425 | 0,209454 | -3,83975 | 0,000123 | 0,000586 | 687,2407869 | -0,434890801 | 0,130159 | -3,34123 | 0,000834 | 0,001805 |
| 805 | RM25_RS03890 | RM25_0792 | Valine--tRNA ligase | 843 | gene | 920216 | 922834 | - | 4868,598 | 0,367576495 | 0,277582 | 1,32421 | 0,185433 | 0,290796 | 6158,073755 | 0,290429271 | 0,149867 | 1,937917 | 0,052633 | 0,082485 | 3579,123 | 0,502179416 | 0,223392 | 2,247974 | 0,024578 | 0,060277 | 4519,764 | -0,96019 | 0,185232 | -5,18372 | 2,18E-07 | 1,88E-06 | 5371,851445 | -0,705658362 | 0,163981 | -4,30328 | 1,68E-05 | 4,8E-05 |
| 806 | RM25_RS03895 | RM25_0793 | DSBA-like thioredoxin domain protein | 844 | gene | 922933 | 923727 | - | 946,3257 | 0,182516977 | 0,310501 | 0,587814 | 0,556657 | 0,65146 | 664,5932302 | -0,277366282 | 0,143863 | -1,92799 | 0,053856 | 0,084051 | 1228,058 | 0,432911373 | 0,223685 | 1,935361 | 0,052946 | 0,105045 | 933,949 | 0,462853 | 0,167375 | 2,765366 | 0,005686 | 0,014687 | 982,3183711 | 1,216945502 | 0,175697 | 6,92637 | 4,32E-12 | 3,35E-11 |
| 807 | RM25_RS03900 | RM25_0794 | DSBA-like thioredoxin domain protein | 845 | gene | 923991 | 924785 | - | 1503,904 | 0,751910744 | 0,352519 | 2,132963 | 0,032928 | 0,095934 | 1876,695958 | 0,340731631 | 0,176748 | 1,92778 | 0,053882 | 0,084051 | 1131,112 | 1,519413288 | 0,246504 | 6,16385 | 7,1E-10 | 2,04E-08 | 1194,276 | -1,56343 | 0,233211 | -6,70392 | 2,03E-11 | 4,35E-10 | 1847,668063 | -0,334692808 | 0,158741 | -2,10842 | 0,034995 | 0,05384 |
| 808 | RM25_RS03905 | RM25_0795 | Putative type IV conjugative transfer system protein TraL | 846 | gene | 924821 | 925432 | - | 857,9229 | 0,382186483 | 0,263431 | 1,450805 | 0,146834 | 0,247383 | 1054,234135 | 0,169630586 | 0,149452 | 1,135014 | 0,25637 | 0,321246 | 661,6116 | 0,730843481 | 0,202474 | 3,609575 | 0,000307 | 0,001887 | 792,0284 | -1,05904 | 0,199566 | -5,30672 | 1,12E-07 | 1,03E-06 | 951,4848413 | -0,44350023 | 0,116892 | -3,7941 | 0,000148 | 0,000367 |
| 809 | RM25_RS03910 | RM25_0796 | Folylpolyglutamate synthase folC | 847 | gene | 925547 | 927052 | + | 1760,553 | 0,623019632 | 0,421099 | 1,479507 | 0,139005 | 0,2388 | 2557,212667 | 0,79375178 | 0,178658 | 4,442857 | 8,88E-06 | 3,77E-05 | 963,8928 | 0,188855968 | 0,16692 | 1,131414 | 0,257881 | 0,359235 | 1472,359 | -1,11244 | 0,203099 | -5,47732 | 4,32E-08 | 4,29E-07 | 2095,183329 | -1,668579578 | 0,116219 | -14,3572 | 9,61E-47 | 1,89E-44 |
| 810 | RM25_RS03915 | RM25_0797 | Nucleoside diphosphate kinase | 848 | gene | 927163 | 927639 | + | 1152,664 | 0,760135843 | 0,145248 | 5,233353 | 1,66E-07 | 9,34E-06 | 1204,414079 | 0,608007186 | 0,128697 | 4,724344 | 2,31E-06 | 1,1E-05 | 1100,915 | 0,931721736 | 0,251851 | 3,699499 | 0,000216 | 0,001407 | 906,3481 | -0,39638 | 0,155546 | -2,5483 | 0,010825 | 0,025262 | 1417,243501 | -0,025073127 | 0,19317 | -0,1298 | 0,896726 | 0,916558 |
| 811 | RM25_RS03920 | RM25_0798 | Hypothetical protein | 849 | gene | 927846 | 928976 | + | 1480,28 | 0,576597715 | 0,195289 | 2,952543 | 0,003152 | 0,021532 | 1600,223467 | 0,261446141 | 0,136794 | 1,911233 | 0,055975 | 0,086456 | 1360,337 | 0,966208108 | 0,246802 | 3,914915 | 9,04E-05 | 0,000677 | 1259,901 | -0,7208 | 0,172512 | -4,17825 | 2,94E-05 | 0,000163 | 1732,891292 | 0,030327367 | 0,175096 | 0,173204 | 0,862491 | 0,888502 |
| 812 | RM25_RS03925 | RM25_0799 | Radical SAM domain protein | 850 | gene | 929053 | 930996 | + | 2002,71 | 0,86468551 | 0,293035 | 2,950797 | 0,00317 | 0,021591 | 2496,471048 | 0,646780359 | 0,163572 | 3,954104 | 7,68E-05 | 0,000269 | 1508,95 | 1,24947503 | 0,196271 | 6,366082 | 1,94E-10 | 6,35E-09 | 1508,55 | -1,18258 | 0,198036 | -5,97153 | 2,35E-09 | 3,01E-08 | 2533,512747 | -0,530767084 | 0,116021 | -4,57476 | 4,77E-06 | 1,48E-05 |
| 813 | RM25_RS03930 | RM25_0800 | Calcium/proton antiporter | 851 | gene | 931060 | 932160 | - | 964,4692 | -0,220500387 | 0,154538 | -1,42684 | 0,153626 | 0,25466 | 1062,019854 | -0,14728502 | 0,144505 | -1,01924 | 0,308089 | 0,376252 | 866,9186 | -0,308848578 | 0,231142 | -1,33618 | 0,181489 | 0,274211 | 1095,962 | -0,28077 | 0,192353 | -1,45964 | 0,144388 | 0,200899 | 874,5864385 | -0,384307681 | 0,171502 | -2,24084 | 0,025037 | 0,040171 |
| 814 | RM25_RS03935 | RM25_0801 | Integral membrane protein TerC | 852 | gene | 932157 | 933413 | - | 2022,919 | 0,117527895 | 0,205681 | 0,571408 | 0,567723 | 0,660802 | 2394,630126 | 0,011321348 | 0,140816 | 0,080398 | 0,935921 | 0,950028 | 1651,208 | 0,27280855 | 0,189279 | 1,441305 | 0,149499 | 0,234599 | 2058,969 | -0,733 | 0,154016 | -4,75926 | 1,94E-06 | 1,4E-05 | 2061,47096 | -0,424548262 | 0,139519 | -3,04294 | 0,002343 | 0,004621 |
| 815 | RM25_RS03940 | RM25_0802 | Transcriptional regulator, MarR family | 853 | gene | 933628 | 934083 | + | 293,1828 | -0,327788352 | 0,234961 | -1,39508 | 0,162993 | 0,265538 | 240,8425266 | -0,487309788 | 0,213584 | -2,28159 | 0,022514 | 0,0396 | 345,5231 | -0,220522979 | 0,276433 | -0,79774 | 0,425019 | 0,529555 | 343,3825 | 0,342244 | 0,195154 | 1,753715 | 0,079479 | 0,122761 | 255,4150924 | 0,670569735 | 0,287504 | 2,332381 | 0,019681 | 0,032371 |
| 816 | RM25_RS03945 | RM25_0803 | Radical SAM domain protein | 854 | gene | 934147 | 934899 | + | 579,8393 | -0,031812145 | 0,132521 | -0,24005 | 0,810289 | 0,868465 | 573,4774876 | -0,135747268 | 0,165306 | -0,82119 | 0,411538 | 0,480196 | 586,201 | 0,069347077 | 0,24458 | 0,283535 | 0,776766 | 0,837146 | 619,0101 | -0,13524 | 0,223328 | -0,60555 | 0,544814 | 0,610327 | 562,1928525 | 0,129093345 | 0,177196 | 0,728535 | 0,466286 | 0,529912 |
| 817 | RM25_RS03950 | RM25_0804 | Hypothetical protein | 855 | gene | 935153 | 937939 | + | 4320,975 | 0,837274188 | 0,398085 | 2,103256 | 0,035443 | 0,10053 | 5721,722488 | 0,508069605 | 0,189847 | 2,676209 | 0,007446 | 0,015405 | 2920,227 | 1,56202676 | 0,249202 | 6,268127 | 3,65E-10 | 1,15E-08 | 3302,152 | -1,73619 | 0,241871 | -7,17818 | 7,06E-13 | 1,87E-11 | 5426,399546 | -0,636468502 | 0,157104 | -4,05126 | 5,09E-05 | 0,000134 |
| 818 | RM25_RS03955 | RM25_0805 | 50S ribosomal protein L21 | 856 | gene | 938136 | 938483 | + | 9354,101 | 0,894593678 | 0,486124 | 1,840258 | 0,06573 | 0,147269 | 12617,95804 | 0,579574963 | 0,248989 | 2,327718 | 0,019927 | 0,035555 | 6090,243 | 1,633390876 | 0,51349 | 3,180962 | 0,001468 | 0,006666 | 6996,306 | -1,81458 | 0,524705 | -3,45829 | 0,000544 | 0,002111 | 11905,21635 | -0,732041953 | 0,224357 | -3,26284 | 0,001103 | 0,002305 |
| 819 | RM25_RS03960 | RM25_0806 | 50S ribosomal protein L27 | 857 | gene | 938497 | 938760 | + | 3834,452 | 0,919937644 | 0,518202 | 1,775249 | 0,075857 | 0,162837 | 4979,726674 | 0,411496204 | 0,222485 | 1,849542 | 0,06438 | 0,09696 | 2689,177 | 2,080561234 | 0,556352 | 3,739651 | 0,000184 | 0,001234 | 2837,666 | -2,10128 | 0,529808 | -3,96612 | 7,31E-05 | 0,00037 | 4903,93081 | -0,405452378 | 0,28089 | -1,44346 | 0,148892 | 0,196935 |
| 820 | RM25_RS03965 | RM25_0807 | GTPase obg | 858 | gene | 938909 | 940510 | + | 2747,432 | 0,818441265 | 0,223204 | 3,666781 | 0,000246 | 0,003386 | 3184,462463 | 0,696345864 | 0,179241 | 3,884962 | 0,000102 | 0,000348 | 2310,401 | 0,992433052 | 0,258596 | 3,837778 | 0,000124 | 0,000884 | 2109,669 | -0,70511 | 0,247807 | -2,8454 | 0,004436 | 0,011975 | 3431,751451 | -0,371117398 | 0,172564 | -2,15061 | 0,031507 | 0,049311 |
| 821 | RM25_RS03970 | RM25_0808 | Glutamate 5-kinase | 859 | gene | 940503 | 941672 | + | 1323,396 | 0,727065993 | 0,253136 | 2,872234 | 0,004076 | 0,025812 | 1624,749859 | 0,623146719 | 0,174132 | 3,578592 | 0,000345 | 0,001052 | 1022,042 | 0,896877252 | 0,186643 | 4,805302 | 1,55E-06 | 2,03E-05 | 1057,791 | -0,90011 | 0,202505 | -4,44486 | 8,79E-06 | 5,47E-05 | 1616,84517 | -0,577448403 | 0,125011 | -4,61917 | 3,85E-06 | 1,21E-05 |
| 822 | RM25_RS03975 | RM25_0809 | Hypothetical protein | 860 | gene | 941705 | 942331 | + | 416,8945 | 0,092429492 | 0,168353 | 0,549023 | 0,58299 | 0,674574 | 471,5788848 | 0,235688856 | 0,155786 | 1,512901 | 0,130305 | 0,180452 | 362,21 | -0,090350699 | 0,202936 | -0,44522 | 0,656162 | 0,736464 | 426,3845 | -0,27626 | 0,177673 | -1,5549 | 0,11997 | 0,172315 | 422,1316621 | -0,548114587 | 0,163003 | -3,36261 | 0,000772 | 0,001683 |
| 823 | RM25_RS03980 | RM25_0810 | Gamma-glutamyl phosphate reductase | 861 | gene | 942384 | 943625 | + | 2350,779 | 0,759303864 | 0,188619 | 4,025598 | 5,68E-05 | 0,001089 | 2678,412854 | 0,849605561 | 0,166065 | 5,116103 | 3,12E-07 | 1,71E-06 | 2023,145 | 0,64302554 | 0,253308 | 2,538509 | 0,011133 | 0,033089 | 1848,84 | -0,32798 | 0,244636 | -1,34067 | 0,180029 | 0,241095 | 2893,729424 | -0,495494993 | 0,157097 | -3,15406 | 0,00161 | 0,003249 |
| 824 | RM25_RS03985 | RM25_0811 | putative nicotinate-nucleotide adenylyltransferase | 862 | gene | 943699 | 944469 | + | 1311,118 | 0,621565972 | 0,259831 | 2,392197 | 0,016748 | 0,064713 | 1571,600311 | 0,900748098 | 0,155368 | 5,7975 | 6,73E-09 | 5,05E-08 | 1050,636 | 0,226003856 | 0,326814 | 0,691536 | 0,489229 | 0,589224 | 1094,387 | -0,22969 | 0,304557 | -0,75418 | 0,450741 | 0,525159 | 1556,511137 | -0,868542015 | 0,183916 | -4,7225 | 2,33E-06 | 7,61E-06 |
| 825 | RM25_RS03990 | RM25_0812 | Ribosome-associated protein, iojap family | 863 | gene | 944466 | 944849 | + | 590,1511 | 0,834927801 | 0,380268 | 2,195632 | 0,028118 | 0,086721 | 801,551377 | 0,919204237 | 0,201997 | 4,550579 | 5,35E-06 | 2,37E-05 | 378,7507 | 0,668515303 | 0,380301 | 1,757858 | 0,078772 | 0,143592 | 451,0409 | -0,97411 | 0,359808 | -2,70731 | 0,006783 | 0,017044 | 741,0108787 | -1,188102029 | 0,230071 | -5,16406 | 2,42E-07 | 9,34E-07 |
| 826 | RM25_RS03995 | RM25_0813 | Phosphoglycerate mutase | 864 | gene | 944849 | 945457 | + | 670,5778 | 0,432021628 | 0,229838 | 1,879677 | 0,060152 | 0,139408 | 812,5210977 | 0,649591883 | 0,178509 | 3,638984 | 0,000274 | 0,00085 | 528,6346 | 0,108925517 | 0,154654 | 0,704319 | 0,481234 | 0,581676 | 603,8451 | -0,3721 | 0,155199 | -2,39755 | 0,016505 | 0,035206 | 755,3876363 | -0,863858978 | 0,165782 | -5,2108 | 1,88E-07 | 7,31E-07 |
| 827 | RM25_RS04000 | RM25_0814 | Hypothetical protein | 865 | gene | 945517 | 945954 | + | 245,5319 | -0,5063877 | 0,179021 | -2,82864 | 0,004675 | 0,028397 | 217,4662561 | -0,414958643 | 0,209166 | -1,98388 | 0,04727 | 0,075586 | 273,5975 | -0,58202178 | 0,258435 | -2,2521 | 0,024316 | 0,059825 | 303,9134 | 0,33228 | 0,221732 | 1,498564 | 0,133987 | 0,189106 | 199,2971801 | 0,229785907 | 0,268749 | 0,855021 | 0,392539 | 0,457801 |
| 828 | RM25_RS04005 | RM25_0815 | NADPH-dependent glutamate synthase beta chain and oxidc | 866 | gene | 946069 | 946989 | + | 408,7463 | -0,920093763 | 0,417771 | -2,20239 | 0,027638 | 0,086167 | 257,7545642 | -1,394883541 | 0,291507 | -4,78507 | 1,71E-06 | 8,32E-06 | 559,7381 | -0,719770112 | 0,262706 | -2,73984 | 0,006147 | 0,021151 | 561,6563 | 0,83259 | 0,302377 | 2,753485 | 0,005896 | 0,015156 | 277,6585304 | 1,571861023 | 0,280874 | 5,596324 | 2,19E-08 | 9,96E-08 |
| 829 | NA | RM25_0816 | Hypothetical protein | 867 | gene | 946989 | 947099 | + | 22,40486 | -1,429319798 | 0,346043 | -4,13047 | 3,62E-05 | 0,000769 | 19,00277502 | -1,639804084 | 0,618827 | -2,64986 | 0,008053 | 0,016504 | 25,80694 | -1,303830996 | 0,391148 | -3,33334 | 0,000858 | 0,004322 | 34,53499 | 0,273618 | 0,366777 | 0,746006 | 0,455664 | 0,529324 | 11,84377921 | 0,677283989 | 0,7404 | 0,914754 | 0,360321 | 0,424638 |
| 831 | RM25_RS04015 | RM25_0817 | Hypothetical protein | 871 | gene | 947292 | 948032 | - | 313,8775 | -0,80805951 | 0,258175 | -3,12989 | 0,001749 | 0,013925 | 248,8961821 | -0,86662057 | 0,208231 | -4,16183 | 3,16E-05 | 0,000121 | 378,8587 | -0,773488617 | 0,305084 | -2,53533 | 0,011234 | 0,033303 | 419,9597 | 0,507418 | 0,229502 | 2,210956 | 0,027039 | 0,051856 | 224,100384 | 0,66361314 | 0,297532 | 2,230396 | 0,025721 | 0,041157 |
| 832 | RM25_RS04020 | RM25_0818 | Hypothetical protein | 872 | gene | 948201 | 949259 | + | 262,3777 | -1,294521081 | 0,787663 | -1,6435 | 0,10028 | 0,193263 | 129,6706029 | -3,980903022 | 0,468666 | -8,49411 | 1,99E-17 | 4,44E-16 | 395,0848 | -0,799125259 | 0,450864 | -1,77243 | 0,076323 | 0,139777 | 391,1041 | 0,967519 | 0,406617 | 2,379435 | 0,017339 | 0,036576 | 150,0383913 | 4,226765414 | 0,540408 | 7,821438 | 5,22E-15 | 6,41E-14 |
| 833 | RM25_RS04025 | RM25_0819 | Leucine--tRNA ligase | 873 | gene | 949576 | 952104 | + | 4604,681 | 0,473258103 | 0,382125 | 1,238491 | 0,215534 | 0,320706 | 6365,949081 | 0,360843737 | 0,182321 | 1,979167 | 0,047797 | 0,076177 | 2843,413 | 0,731359807 | 0,210478 | 3,474764 | 0,000511 | 0,002829 | 4106,674 | -1,44029 | 0,221555 | -6,50081 | 7,99E-11 | 1,46E-09 | 5246,54924 | -1,025307744 | 0,144434 | -7,09878 | 1,26E-12 | 1,08E-11 |
| 834 | RM25_RS04030 | RM25_0820 | Dihydroorotase | 874 | gene | 952161 | 953453 | + | 1746,558 | 0,432736795 | 0,286536 | 1,510237 | 0,130983 | 0,229602 | 2242,538029 | 0,408343256 | 0,171134 | 2,386104 | 0,017028 | 0,031282 | 1250,577 | 0,477412963 | 0,182496 | 2,616017 | 0,008896 | 0,027884 | 1579,552 | -0,94507 | 0,207717 | -4,5498 | 5,37E-06 | 3,55E-05 | 1968,043334 | -0,821414646 | 0,129166 | -6,3594 | 2,03E-10 | 1,24E-09 |
| 835 | RM25_RS04035 | RM25_0821 | Malic enzyme, NAD binding domain protein | 875 | gene | 953471 | 955225 | + | 2621,258 | 0,324101201 | 0,290654 | 1,115077 | 0,264818 | 0,372419 | 3372,377909 | 0,492876645 | 0,144525 | 3,41031 | 0,000649 | 0,001827 | 1870,139 | 0,025823064 | 0,219889 | 0,117437 | 0,906514 | 0,936285 | 2469,858 | -0,65126 | 0,192283 | -3,38697 | 0,000707 | 0,00264 | 2856,720428 | -1,076268874 | 0,154184 | -6,98041 | 2,94E-12 | 2,35E-11 |
| 836 | RM25_RS11815 | RM25_0822 | Putative competence protein ComEA | 876 | gene | 955410 | 956321 | + | 171,5654 | -0,933980114 | 0,596581 | -1,56556 | 0,117453 | 0,213115 | 88,22700997 | -1,721892155 | 0,443583 | -3,88178 | 0,000104 | 0,000352 | 254,9039 | -0,698040898 | 0,504661 | -1,38319 | 0,166607 | 0,25616 | 236,0951 | 1,143383 | 0,390833 | 2,925501 | 0,003439 | 0,009872 | 116,5612502 | 2,250499926 | 0,572551 | 3,930653 | 8,47E-05 | 0,000218 |
| 837 | RM25_RS04045 | RM25_0823 | DNA internalization competence protein ComEC/Rec2-like p | 877 | gene | 956305 | 958761 | + | 748,5412 | -1,444181549 | 0,764646 | -1,88869 | 0,058933 | 0,137666 | 376,0171827 | -3,820478073 | 0,414013 | -9,22792 | 2,76E-20 | 9,29E-19 | 1121,065 | -0,97888925 | 0,483297 | -2,02544 | 0,042822 | 0,089927 | 1148,707 | 1,008893 | 0,434663 | 2,32109 | 0,020282 | 0,041425 | 397,9217884 | 3,931829881 | 0,497061 | 7,910157 | 2,57E-15 | 3,35E-14 |
| 838 | RM25_RS04050 | RM25_0824 | DNA polymerase III, delta subunit | 878 | gene | 958844 | 959842 | + | 738,9664 | 0,100301917 | 0,167107 | 0,600226 | 0,548356 | 0,643342 | 792,1404038 | 0,44390662 | 0,153398 | 2,893823 | 0,003806 | 0,008637 | 685,7924 | -0,290496227 | 0,180863 | -1,60617 | 0,108237 | 0,183009 | 753,048 | 0,104194 | 0,162101 | 0,642774 | 0,520371 | 0,587974 | 750,5300729 | -0,572268408 | 0,177059 | -3,23208 | 0,001229 | 0,002539 |
| 839 | RM25_RS04055 | RM25_0825 | Hypothetical protein | 879 | gene | 959856 | 960518 | - | 380,304 | -0,695876228 | 0,316359 | -2,19964 | 0,027832 | 0,086316 | 275,1077611 | -0,734716239 | 0,238169 | -3,08485 | 0,002037 | 0,004995 | 485,5002 | -0,677678237 | 0,315844 | -2,14561 | 0,031904 | 0,072725 | 493,7915 | 0,73105 | 0,236677 | 3,088809 | 0,00201 | 0,006307 | 285,3708134 | 0,853122558 | 0,324823 | 2,626422 | 0,008629 | 0,01528 |
| 840 | RM25_RS04060 | RM25_0826 | 30S ribosomal protein S20 | 880 | gene | 960652 | 960912 | - | 7711,331 | 0,725836448 | 0,596505 | 1,216815 | 0,223675 | 0,329913 | 11754,45812 | 0,408244129 | 0,217196 | 1,879611 | 0,060161 | 0,091484 | 3668,204 | 1,936497286 | 0,351832 | 5,504044 | 3,71E-08 | 7,17E-07 | 6245,293 | -2,78687 | 0,36124 | -7,71472 | 1,21E-14 | 4,33E-13 | 9418,04622 | -1,220109125 | 0,205904 | -5,92562 | 3,11E-09 | 1,6E-08 |
| 841 | RM25_RS04065 | RM25_0827 | Transposase IS204/IS1001/IS1096/IS1165 family protein | 881 | gene | 961457 | 962533 | + | 32,70729 | -2,068119964 | 1,05451 | -1,96121 | 0,049854 | 0,12474 | 16,63864525 | -7,286326843 | 1,44868 | -5,02963 | 4,91E-07 | 2,6E-06 | 48,77593 | -1,523824166 | 0,370133 | -4,11696 | 3,84E-05 | 0,000323 | 55,34624 | 1,040271 | 0,519541 | 2,002286 | 0,045254 | 0,078497 | 12,33849917 | 6,872910912 | 1,46948 | 4,677105 | 2,91E-06 | 9,33E-06 |
| 842 | RM25_RS04070 | RM25_0828 | Putative membrane protein | 882 | gene | 962614 | 963234 | + | 16173,61 | 0,909611161 | 0,275325 | 3,303772 | 0,000954 | 0,009713 | 15810,30899 | 0,780605201 | 0,193139 | 4,041684 | 5,31E-05 | 0,000194 | 16536,92 | 1,036443672 | 0,469352 | 2,208242 | 0,027227 | 0,064823 | 11916,44 | -0,14889 | 0,429216 | -0,34688 | 0,728682 | 0,774529 | 20604,54218 | 0,134308253 | 0,265472 | 0,505923 | 0,612911 | 0,671787 |
| 843 | RM25_RS04075 | RM25_0829 | Elongation factor 4 | 883 | gene | 963535 | 965364 | + | 1739,698 | -0,028020455 | 0,268073 | -0,10453 | 0,916752 | 0,944812 | 2201,443532 | 0,040418812 | 0,146973 | 0,275008 | 0,78331 | 0,826808 | 1277,953 | -0,144837149 | 0,18594 | -0,77894 | 0,436012 | 0,540885 | 1861,344 | -0,75333 | 0,183509 | -4,10513 | 4,04E-05 | 0,000214 | 1689,849711 | -0,887166372 | 0,104487 | -8,49065 | 2,05E-17 | 3,39E-16 |
| 844 | RM25_RS04080 | RM25_0830 | Coproporphyrinogen dehydrogenase | 884 | gene | 965367 | 966566 | + | 571,6995 | -0,367551907 | 0,19331 | -1,90136 | 0,057255 | 0,135086 | 552,0000049 | -0,111725993 | 0,139883 | -0,79871 | 0,424457 | 0,492345 | 591,399 | -0,611198083 | 0,335853 | -1,81984 | 0,068784 | 0,129039 | 678,2789 | 0,253771 | 0,192646 | 1,317287 | 0,187742 | 0,249864 | 491,1650741 | -0,180592683 | 0,304282 | -0,59351 | 0,552843 | 0,614357 |
| 845 | RM25_RS04085 | RM25_0831 | PF11296 family protein | 885 | gene | 966661 | 967542 | + | 912,3357 | 0,516079678 | 0,296324 | 1,741609 | 0,081577 | 0,168664 | 1181,245297 | 0,564314061 | 0,17501 | 3,224464 | 0,001262 | 0,003298 | 643,4261 | 0,430044611 | 0,219115 | 1,962645 | 0,049687 | 0,100699 | 797,1384 | -0,85722 | 0,192209 | -4,45981 | 8,2E-06 | 5,14E-05 | 1052,145685 | -0,945390583 | 0,175824 | -5,37692 | 7,58E-08 | 3,17E-07 |
| 846 | RM25_RS04090 | RM25_0832 | Inositol monophosphatase | 886 | gene | 967620 | 968396 | + | 436,304 | 0,261363078 | 0,138699 | 1,884392 | 0,059512 | 0,13852 | 459,8166506 | 0,370857672 | 0,182913 | 2,027505 | 0,042611 | 0,069601 | 412,7914 | 0,143249113 | 0,227412 | 0,62991 | 0,528753 | 0,626897 | 419,2163 | -0,09337 | 0,168025 | -0,55569 | 0,578423 | 0,641272 | 467,3447633 | -0,26153805 | 0,235707 | -1,10959 | 0,267175 | 0,326794 |
| 847 | RM25_RS04095 | RM25_0833 | Heat-inducible transcription repressor hrcA | 887 | gene | 968654 | 969664 | + | 1106,335 | 0,530916907 | 0,211667 | 2,508265 | 0,012133 | 0,052858 | 1262,149497 | 0,305669887 | 0,145404 | 2,102217 | 0,035534 | 0,05919 | 950,5208 | 0,841882257 | 0,243732 | 3,45413 | 0,000552 | 0,003033 | 961,7179 | -0,78932 | 0,203983 | -3,86951 | 0,000109 | 0,000525 | 1279,111032 | -0,208065797 | 0,169392 | -1,22831 | 0,219332 | 0,275567 |
| 848 | RM25_RS04100 | RM25_0834 | Chaperone protein DnaJ | 888 | gene | 969677 | 970885 | + | 1260,286 | 0,585732993 | 0,231546 | 2,529657 | 0,011417 | 0,051153 | 1497,349152 | 0,420483411 | 0,175613 | 2,394372 | 0,016649 | 0,030753 | 1023,222 | 0,835027081 | 0,194383 | 4,295779 | 1,74E-05 | 0,000166 | 1068,833 | -0,85985 | 0,202851 | -4,23883 | 2,25E-05 | 0,000128 | 1481,839751 | -0,396897523 | 0,130385 | -3,04404 | 0,002334 | 0,004612 |
| 849 | RM25_RS04105 | RM25_0835 | Ribosomal RNA small subunit methyltransferase E | 889 | gene | 970882 | 971631 | + | 381,9322 | -0,127403714 | 0,222346 | -0,573 | 0,566646 | 0,6602 | 392,8574039 | 0,219434084 | 0,208706 | 1,051401 | 0,293075 | 0,361474 | 371,0069 | -0,493547341 | 0,330497 | -1,49335 | 0,135346 | 0,21731 | 419,7926 | 0,199369 | 0,170519 | 1,16919 | 0,242327 | 0,310753 | 359,1081896 | -0,452702243 | 0,33782 | -1,34007 | 0,180223 | 0,232123 |
| 850 | RM25_RS04110 | RM25_0836 | Putative Tat (Twin-arginine translocation) pathway signal se | 890 | gene | 971620 | 972948 | - | 1218,268 | -0,644766885 | 0,255007 | -2,52842 | 0,011458 | 0,051153 | 992,9199211 | -0,461097946 | 0,253927 | -1,81587 | 0,069391 | 0,103647 | 1443,616 | -0,774889225 | 0,285486 | -2,71428 | 0,006642 | 0,022493 | 1559,217 | 0,598811 | 0,321877 | 1,860369 | 0,062833 | 0,102349 | 933,9685667 | 0,348316725 | 0,2248 | 1,549453 | 0,121273 | 0,164182 |
| 851 | RM25_RS04115 | RM25_0837 | TRNA-guanine transglycosylase | 891 | gene | 973068 | 974336 | + | 844,0901 | 0,159141916 | 0,126734 | 1,255715 | 0,209219 | 0,313895 | 880,0413283 | 0,267063682 | 0,129171 | 2,067527 | 0,038685 | 0,063806 | 808,1388 | 0,04424648 | 0,240733 | 0,183799 | 0,854171 | 0,894061 | 840,4338 | -0,06927 | 0,213497 | -0,32445 | 0,745597 | 0,790186 | 874,3991599 | -0,233017036 | 0,143827 | -1,62012 | 0,105207 | 0,144507 |
| 852 | RM25_RS04120 | RM25_0838 | Hypothetical protein | 892 | gene | 974449 | 975000 | + | 4555,435 | -0,099977326 | 0,328374 | -0,30446 | 0,760776 | 0,826336 | 4513,627548 | 0,581732664 | 0,183076 | 3,177539 | 0,001485 | 0,003768 | 4597,243 | -0,779353915 | 0,451881 | -1,72469 | 0,084584 | 0,151147 | 4973,658 | 0,635483 | 0,392455 | 1,619248 | 0,105394 | 0,15543 | 4298,992145 | -0,697787148 | 0,279247 | -2,49882 | 0,012461 | 0,021379 |
| 853 | RM25_RS04125 | RM25_0839 | Prolyl aminopeptidase | 893 | gene | 975082 | 976140 | + | 1747,011 | 0,275022236 | 0,206329 | 1,332932 | 0,182554 | 0,287581 | 2035,205963 | 0,492295389 | 0,139946 | 3,517747 | 0,000435 | 0,001282 | 1458,815 | -0,021523548 | 0,231207 | -0,09309 | 0,925831 | 0,949601 | 1673,131 | -0,25711 | 0,199754 | -1,28711 | 0,198055 | 0,261521 | 1874,325777 | -0,729211547 | 0,158112 | -4,612 | 3,99E-06 | 1,25E-05 |
| 854 | RM25_RS04130 | RM25_0840 | Hypothetical protein | 894 | gene | 976196 | 976687 | + | 416,4797 | 0,390179711 | 0,102856 | 3,793449 | 0,000149 | 0,002274 | 433,8791335 | 0,488528383 | 0,151744 | 3,219425 | 0,001284 | 0,003353 | 399,0802 | 0,290349723 | 0,193594 | 1,499788 | 0,133669 | 0,21491 | 380,1761 | -0,06807 | 0,16133 | -0,42192 | 0,673086 | 0,724082 | 463,3594696 | -0,214834439 | 0,157268 | -1,36604 | 0,171925 | 0,224006 |
| 855 | RM25_RS04135 | RM25_0841 | Inner membrane metabolite transport protein yhjE | 895 | gene | 976856 | 978238 | + | 759,7435 | -0,932048293 | 0,365775 | -2,54815 | 0,01083 | 0,049182 | 550,0112735 | -0,650332204 | 0,327535 | -1,98554 | 0,047085 | 0,075393 | 969,4757 | -1,100605642 | 0,409324 | -2,68884 | 0,00717 | 0,023884 | 1045,348 | 0,905456 | 0,401786 | 2,253576 | 0,024223 | 0,047657 | 514,9450503 | 0,531970106 | 0,368767 | 1,442565 | 0,149143 | 0,197157 |
| 856 | RM25_RS04140 | RM25_0842 | Phosphate starvation-inducible protein phoH1 | 896 | gene | 978370 | 979377 | + | 2081,796 | 0,220496347 | 0,237708 | 0,927592 | 0,353619 | 0,466073 | 1989,663238 | 0,637789906 | 0,150812 | 4,229048 | 2,35E-05 | 9,27E-05 | 2173,928 | -0,151774525 | 0,363683 | -0,41733 | 0,67644 | 0,754582 | 2028,799 | 0,506127 | 0,321509 | 1,574224 | 0,115436 | 0,166819 | 2190,539808 | -0,251337092 | 0,214743 | -1,17041 | 0,241837 | 0,298903 |
| 857 | RM25_RS04145 | RM25_0843 | putative rRNA maturation factor | 897 | gene | 979417 | 979875 | + | 1528,681 | 0,384478474 | 0,307192 | 1,251591 | 0,210719 | 0,315543 | 1886,753308 | 0,432259528 | 0,212826 | 2,031049 | 0,04225 | 0,069155 | 1170,608 | 0,309519176 | 0,385754 | 0,802375 | 0,422336 | 0,526691 | 1408,654 | -0,6667 | 0,371693 | -1,79369 | 0,072863 | 0,114953 | 1694,173171 | -0,75959045 | 0,228809 | -3,31976 | 0,000901 | 0,001941 |
| 858 | RM25_RS04150 | RM25_0844 | CBS domain protein | 898 | gene | 980029 | 981333 | + | 1643,763 | 0,168837942 | 0,240706 | 0,701428 | 0,483036 | 0,582659 | 2026,564568 | 0,308947898 | 0,143585 | 2,151672 | 0,031423 | 0,053322 | 1260,961 | -0,053328455 | 0,165724 | -0,32179 | 0,747612 | 0,814289 | 1639,405 | -0,55344 | 0,154806 | -3,57505 | 0,00035 | 0,001471 | 1705,691113 | -0,868284289 | 0,121282 | -7,15919 | 8,12E-13 | 7,14E-12 |
| 859 | RM25_RS04155 | RM25_0845 | GTPase Era | 899 | gene | 981326 | 982270 | + | 1140,462 | 0,403055679 | 0,26008 | 1,549736 | 0,121205 | 0,217744 | 1426,097988 | 0,410207005 | 0,159372 | 2,573891 | 0,010056 | 0,019968 | 854,8251 | 0,3931583 | 0,215771 | 1,82211 | 0,068438 | 0,128738 | 1041,459 | -0,78543 | 0,17483 | -4,4925 | 7,04E-06 | 4,5E-05 | 1272,104409 | -0,759012949 | 0,163426 | -4,6444 | 3,41E-06 | 1,08E-05 |
| 860 | RM25_RS04160 | RM25_0846 | Hypothetical protein | 900 | gene | 982320 | 982928 | + | 16025,9 | -0,317058266 | 0,430477 | -0,73653 | 0,46141 | 0,564957 | 18600,58472 | 0,359098594 | 0,224359 | 1,600554 | 0,109476 | 0,153867 | 13451,22 | -1,333861733 | 0,529296 | -2,52007 | 0,011733 | 0,03444 | 18832,51 | 0,194828 | 0,475909 | 0,40938 | 0,682261 | 0,731615 | 13969,82914 | -1,474814181 | 0,318126 | -4,63594 | 3,55E-06 | 1,13E-05 |
| 861 | RM25_RS13195 | RM25_0847 | Hypothetical protein | 901 | gene | 983039 | 983173 | + | 1006,766 | -0,212261356 | 0,41612 | -0,5101 | 0,609984 | 0,699967 | 1091,151581 | 0,545763792 | 0,225723 | 2,417847 | 0,015613 | 0,02909 | 922,3806 | -1,155565104 | 0,547277 | -2,11148 | 0,034731 | 0,0773 | 1142,495 | 0,476361 | 0,548857 | 0,867915 | 0,385441 | 0,462098 | 914,4052052 | -1,193382355 | 0,226167 | -5,27655 | 1,32E-07 | 5,25E-07 |
| 862 | RM25_RS04165 | RM25_0848 | Amidohydrolase family protein | 902 | gene | 983303 | 984694 | - | 726,0016 | -0,559720556 | 0,268578 | -2,08402 | 0,037159 | 0,103894 | 575,8234432 | -0,538101382 | 0,204444 | -2,63203 | 0,008488 | 0,017234 | 876,1798 | -0,575955247 | 0,33627 | -1,71278 | 0,086754 | 0,154323 | 909,6302 | 0,551317 | 0,269219 | 2,047841 | 0,040576 | 0,071692 | 577,1349756 | 0,58295388 | 0,294911 | 1,976708 | 0,048075 | 0,0714 |
| 863 | RM25_RS04170 | RM25_0849 | 2-isopropylmalate synthase | 903 | gene | 985040 | 986806 | + | 1151,527 | 0,135041861 | 0,126285 | 1,069342 | 0,284916 | 0,3941 | 1079,563264 | 0,170777106 | 0,159592 | 1,070082 | 0,284582 | 0,35229 | 1223,49 | 0,103511986 | 0,206955 | 0,500166 | 0,616958 | 0,703518 | 1157,593 | 0,147574 | 0,225261 | 0,655123 | 0,512388 | 0,579789 | 1181,18274 | 0,137907221 | 0,114331 | 1,206207 | 0,227738 | 0,284039 |
| 864 | RM25_RS04175 | RM25_0850 | Protein of hypothetical function DUF488 | 904 | gene | 986921 | 987283 | - | 2061,554 | 1,637086383 | 0,281074 | 5,824401 | 5,73E-09 | 5,2E-07 | 2343,107235 | 1,601426903 | 0,241645 | 6,627184 | 3,42E-11 | 3,6E-10 | 1780,002 | 1,685832823 | 0,406718 | 4,144972 | 3,4E-05 | 0,000292 | 1065,394 | -0,50958 | 0,388426 | -1,31191 | 0,189549 | 0,251558 | 3047,974167 | -0,396144766 | 0,263379 | -1,50409 | 0,132559 | 0,177378 |
| 865 | RM25_RS04180 | RM25_0851 | Isoprenyl transferase 2 | 905 | gene | 987344 | 988033 | - | 408,4259 | 0,008089836 | 0,128515 | 0,062949 | 0,949807 | 0,967039 | 445,4536817 | 0,162920984 | 0,152475 | 1,068508 | 0,285291 | 0,352797 | 371,3982 | -0,170316035 | 0,160373 | -1,062 | 0,288237 | 0,392476 | 430,3336 | -0,1537 | 0,152332 | -1,00897 | 0,31299 | 0,387661 | 402,0119788 | -0,433255574 | 0,158467 | -2,73405 | 0,006256 | 0,01136 |
| 866 | RM25_RS04185 | RM25_0852 | DNA repair protein RecO | 906 | gene | 988217 | 988960 | + | 440,4358 | -0,192960114 | 0,243876 | -0,79122 | 0,428815 | 0,537329 | 343,0735792 | -0,075339926 | 0,211157 | -0,3568 | 0,721244 | 0,771312 | 537,798 | -0,270812576 | 0,209424 | -1,29313 | 0,195966 | 0,291599 | 494,1338 | 0,672385 | 0,232243 | 2,895181 | 0,003789 | 0,010608 | 403,1457874 | 0,5378267 | 0,20288 | 2,650964 | 0,008026 | 0,014278 |
| 867 | RM25_RS11820 | RM25_0853 | Transcriptional regulator | 907 | gene | 989124 | 990512 | + | 2734,544 | 0,011423344 | 0,31168 | 0,036651 | 0,970763 | 0,98075 | 2961,557832 | 0,734624176 | 0,142487 | 5,155735 | 2,53E-07 | 1,43E-06 | 2507,529 | -0,846543252 | 0,345344 | -2,4513 | 0,014234 | 0,039863 | 2875,588 | 0,485911 | 0,324117 | 1,499184 | 0,133826 | 0,189013 | 2690,549481 | -1,058930089 | 0,182918 | -5,7891 | 7,08E-09 | 3,43E-08 |
| 868 | RM25_RS04195 | RM25_0854 | Propanol dehydrogenase | 908 | gene | 990469 | 991599 | + | 555,9551 | -0,342923643 | 0,16519 | -2,07594 | 0,0379 | 0,105591 | 496,4574765 | -0,326852638 | 0,184402 | -1,7725 | 0,076312 | 0,112417 | 615,4526 | -0,357543125 | 0,234808 | -1,5227 | 0,127833 | 0,208513 | 654,2825 | 0,262075 | 0,20835 | 1,257859 | 0,208443 | 0,273248 | 481,1868878 | 0,288613758 | 0,207936 | 1,387994 | 0,165139 | 0,21636 |
| 869 | RM25_RS04200 | RM25_0855 | Ethanolamine utilization protein | 909 | gene | 991652 | 992077 | - | 83,6859 | -1,653685194 | 0,861024 | -1,9206 | 0,054782 | 0,132045 | 46,95100366 | -4,31246981 | 0,750483 | -5,74626 | 9,12E-09 | 6,74E-08 | 120,4208 | -1,110169404 | 0,497175 | -2,23295 | 0,025552 | 0,06177 | 133,4106 | 0,799811 | 0,45592 | 1,754281 | 0,079382 | 0,122691 | 39,8559981 | 4,096012855 | 0,896316 | 4,569831 | 4,88E-06 | 1,51E-05 |
| 870 | RM25_RS04205 | RM25_0856 | Propanediol utilization protein PduU | 910 | gene | 992074 | 992280 | - | 54,92095 | -0,793067278 | 0,387702 | -2,04556 | 0,0408 | 0,109652 | 39,33655821 | -1,462496346 | 0,402346 | -3,63493 | 0,000278 | 0,000861 | 70,50534 | -0,471685655 | 0,340309 | -1,38605 | 0,165731 | 0,255479 | 73,51324 | 0,434831 | 0,284044 | 1,53086 | 0,125804 | 0,179383 | 39,36128069 | 1,492271126 | 0,476854 | 3,129407 | 0,001752 | 0,003511 |
| 871 | RM25_RS04210 | RM25_0857 | Transposase IS204/IS1001/IS1096/IS1165 family protein | 911 | gene | 992460 | 993767 | + | 16,63101 | -1,477714665 | 0,756101 | -1,95439 | 0,050655 | 0,12544 | 9,014222436 | -3,870152129 | 1,152675 | -3,35754 | 0,000786 | 0,002163 | 24,24779 | -0,985719276 | 0,483983 | -2,03668 | 0,041682 | 0,088547 | 25,7265 | 0,843508 | 0,469527 | 1,796506 | 0,072414 | 0,114473 | 8,575633701 | 3,810403843 | 1,211892 | 3,144178 | 0,001666 | 0,00335 |
| 872 | NA | RM25_0858 | Hypothetical protein | 912 | gene | 993739 | 993867 | - | 15,77559 | -1,378731482 | 0,49252 | -2,79934 | 0,005121 | 0,030174 | 12,02692702 | -1,952965189 | 0,805296 | -2,42515 | 0,015302 | 0,028602 | 19,52425 | -1,101194233 | 0,501175 | -2,19722 | 0,028004 | 0,065941 | 24,11108 | 0,389561 | 0,433251 | 0,899157 | 0,368569 | 0,444357 | 8,533796431 | 1,324595341 | 0,965721 | 1,371613 | 0,170184 | 0,222106 |
| 873 | RM25_RS04215 | RM25_0859 | 3-isopropylmalate dehydrogenase | 913 | gene | 993901 | 994965 | - | 1000,165 | 0,30174995 | 0,288013 | 1,047697 | 0,294778 | 0,405127 | 1271,327275 | 0,629398429 | 0,120134 | 5,239128 | 1,61E-07 | 9,44E-07 | 729,002 | -0,253139732 | 0,168739 | -1,50018 | 0,133567 | 0,21491 | 948,3015 | -0,39374 | 0,140445 | -2,80351 | 0,005055 | 0,013298 | 1084,328244 | -1,219639953 | 0,137296 | -8,88327 | 6,49E-19 | 1,33E-17 |
| 874 | RM25_RS04220 | RM25_0860 | Branched-chain-amino-acid aminotransferase | 914 | gene | 995131 | 996216 | + | 2171,822 | 0,674614579 | 0,291229 | 2,316441 | 0,020534 | 0,072671 | 2765,160663 | 0,751072627 | 0,170284 | 4,410695 | 1,03E-05 | 4,33E-05 | 1578,483 | 0,544739306 | 0,287551 | 1,894408 | 0,058171 | 0,11322 | 1777,739 | -0,73472 | 0,285807 | -2,57069 | 0,010149 | 0,023999 | 2616,426419 | -0,902881982 | 0,161316 | -5,59698 | 2,18E-08 | 9,94E-08 |
| 875 | RM25_RS04225 | RM25_0861 | 2-isopropylmalate synthase/homocitrate synthase family prc | 915 | gene | 996248 | 997846 | + | 1736,351 | 0,825723672 | 0,33459 | 2,467866 | 0,013592 | 0,057324 | 2332,065422 | 0,827232696 | 0,146629 | 5,641673 | 1,68E-08 | 1,17E-07 | 1140,637 | 0,823919221 | 0,221504 | 3,719655 | 0,000199 | 0,001317 | 1331,031 | -1,09082 | 0,188897 | -5,7747 | 7,71E-09 | 8,78E-08 | 2175,922087 | -1,045856419 | 0,140289 | -7,45501 | 8,99E-14 | 9,17E-13 |
| 876 | RM25_RS04230 | RM25_0862 | FAH family protein | 916 | gene | 997906 | 998751 | + | 1560,194 | 0,556256696 | 0,172806 | 3,21896 | 0,001287 | 0,011785 | 1682,532182 | 0,612671874 | 0,16414 | 3,732613 | 0,00019 | 0,000609 | 1437,855 | 0,492030484 | 0,289374 | 1,700328 | 0,089069 | 0,157373 | 1336,742 | -0,21031 | 0,228637 | -0,91982 | 0,357665 | 0,434007 | 1816,115305 | -0,292154693 | 0,214122 | -1,36443 | 0,172431 | 0,224542 |
| 877 | RM25_RS04235 | RM25_0863 | CAAX amino terminal protease family protein | 917 | gene | 998961 | 999855 | + | 1189,332 | [illegible] | [illegible] | [illegible] | [illegible] | [illegible] | [illegible] | [illegible] | [illegible] | [illegible] | [illegible] | [illegible] | [illegible] | [illegible] | [illegible] | [illegible] | [illegible] | [illegible] | [illegible] | [illegible] | [illegible] | [illegible] | [illegible] | [illegible] | [illegible] | [illegible] | [illegible] | [illegible] | [illegible] | [illegible] |
| 878 | RM25_RS04240 | RM25_0864 | IclR family transcriptional regulator | 920 | gene | 1000255 | 1000983 | + | 725,9475 | [illegible] | [illegible] | [illegible] | [illegible] | [illegible] | [illegible] | [illegible] | [illegible] | [illegible] | [illegible] | [illegible] | [illegible] | [illegible] | [illegible] | [illegible] | [illegible] | [illegible] | [illegible] | [illegible] | [illegible] | [illegible] | [illegible] | [illegible] | [illegible] | [illegible] | [illegible] | [illegible] | [illegible] | [illegible] |
| 880 | RM25_RS04250 | RM25_0866 | 3-isopropylmalate dehydratase large subunit | 924 | gene | 1001027 | 1003426 | + | 1391,055 | [illegible] | [illegible] | [illegible] | [illegible] | [illegible] | [illegible] | [illegible] | [illegible] | [illegible] | [illegible] | [illegible] | [illegible] | [illegible] | [illegible] | [illegible] | [illegible] | [illegible] | [illegible] | [illegible] | [illegible] | [illegible] | [illegible] | [illegible] | [illegible] | [illegible] | [illegible] | [illegible] | [illegible] | [illegible] |
| 881 | RM25_RS04255 | RM25_0867 | 3-isopropylmalate dehydratase small subunit | 925 | gene | 1003446 | 1004047 | + | 364,9644 | [illegible] | [illegible] | [illegible] | [illegible] | [illegible] | [illegible] | [illegible] | [illegible] | [illegible] | [illegible] | [illegible] | [illegible] | [illegible] | [illegible] | [illegible] | [illegible] | [illegible] | [illegible] | [illegible] | [illegible] | [illegible] | [illegible] | [illegible] | [illegible] | [illegible] | [illegible] | [illegible] | [illegible] | [illegible] |
| 883 | RM25_RS04265 | RM25_0869 | Acyltransferase | 927 | gene | 1003177 | 1003980 | + | 755,2141 | 0,028987799 | 0,15338 | 0,188994 | 0,850098 | 0,901746 | 798,8897262 | 0,232780993 | 0,141154 | 1,649122 | 0,099123 | 0,141082 | 711,5385 | -0,19604578 | 0,251105 | -0,78073 | 0,43496 | 0,539864 | 791,2263 | -0,00772 | 0,258278 | -0,02989 | 0,976155 | 0,980732 | 742,6223909 | -0,389806959 | 0,125591 | -3,10378 | 0,001911 | 0,003807 |

| | | | | | | | | | |
|---|---|---|---|---|---|---|---|---|---|
| 884 | RM25_RS04270 | RM25_0870 | D-alanine--D-alanine ligase | 928 | gene | 1004054 | 1005181 | + | [illegible] |
| 885 | RM25_RS04275 | RM25_0871 | Putative insertion element ISCmi2 transposase | 929 | gene | 1005237 | 1006280 | + | [illegible] |
| 886 | RM25_RS04280 | RM25_0872 | Putative membrane protein | 930 | gene | 1006318 | 1007319 | - | [illegible] |
| 887 | RM25_RS04285 | RM25_0873 | putative transcriptional regulator | 931 | gene | 1007423 | 1008157 | + | [illegible] |
| 888 | RM25_RS04290 | RM25_0874 | Thiamine-phosphate kinase | 932 | gene | 1008168 | 1009151 | + | [illegible] |
| 889 | RM25_RS04295 | RM25_0875 | FtsK/SpoIIIE family protein | 933 | gene | 1009385 | 1011706 | + | [illegible] |
| 890 | RM25_RS04300 | RM25_0876 | Ribosomal protein S12 methylthiotransferase RimO | 934 | gene | 1011786 | 1013219 | + | [illegible] |
| 891 | RM25_RS04305 | RM25_0877 | Phosphatidylglycerophosphate synthase pgsA3 | 935 | gene | 1013224 | 1013877 | + | [illegible] |
| 892 | RM25_RS04310 | RM25_0878 | Competence/damage-inducible protein CinA | 936 | gene | 1013870 | 1014373 | + | [illegible] |
| 893 | RM25_RS04315 | RM25_0879 | Helix-turn-helix XRE-family like proteins | 937 | gene | 1014541 | 1014846 | + | [illegible] |
| 894 | RM25_RS04320 | RM25_0880 | Putative ATP-DEPENDENT HELICASE LHR | 938 | gene | 1015068 | 1019840 | - | [illegible] |
| 895 | RM25_RS04325 | RM25_0881 | Hypothetical protein | 939 | gene | 1019852 | 1020019 | - | [illegible] |
| 896 | RM25_RS04330 | RM25_0882 | RecA protein | 940 | gene | 1020328 | 1021446 | + | [illegible] |
| 897 | RM25_RS04335 | RM25_0883 | Regulatory protein RecX | 941 | gene | 1021449 | 1022063 | + | [illegible] |
| 898 | RM25_RS04340 | RM25_0884 | Ribonuclease Y | 942 | gene | 1022225 | 1023760 | + | [illegible] |
| 899 | RM25_RS04345 | RM25_0885 | Putative glutamate ABC transporter ATP-binding protein | 943 | gene | 1023770 | 1024585 | - | [illegible] |
| 900 | RM25_RS04350 | RM25_0886 | (Dimethylallyl)adenosine tRNA methylthiotransferase MiaB | 944 | gene | 1024689 | 1026203 | + | [illegible] |
| 901 | RM25_RS04355 | RM25_0887 | TRNA dimethylallyltransferase | 945 | gene | 1026200 | 1027186 | + | [illegible] |
| 902 | RM25_RS04360 | RM25_0888 | Diaminopimelate epimerase | 946 | gene | 1027201 | 1028043 | + | [illegible] |
| 903 | RM25_RS04365 | RM25_0889 | GTPase HflX | 947 | gene | 1028087 | 1029523 | + | [illegible] |
| 904 | RM25_RS04370 | RM25_0890 | ATP-dependent helicase DinG | 948 | gene | 1029690 | 1031753 | + | [illegible] |
| 905 | RM25_RS04375 | RM25_0891 | LexA repressor | 949 | gene | 1031757 | 1032599 | - | [illegible] |
| 906 | RM25_RS04380 | RM25_0892 | Hypothetical protein | 950 | gene | 1032869 | 1033672 | + | [illegible] |
| 907 | RM25_RS04385 | RM25_0893 | Transcriptional repressor NrdR | 951 | gene | 1034007 | 1034504 | + | [illegible] |
| 908 | RM25_RS04390 | RM25_0894 | Vitamin B12-dependent ribonucleotide reductase | 952 | gene | 1034690 | 1037581 | + | [illegible] |
| 909 | RM25_RS04395 | RM25_0895 | Phospholipase, patatin family | 953 | gene | 1037690 | 1038631 | - | [illegible] |
| 910 | RM25_RS04400 | RM25_0896 | Glycerol-3-phosphate dehydrogenase, anaerobic, A subunit | 954 | gene | 1038823 | 1040451 | + | [illegible] |
| 911 | RM25_RS04405 | RM25_0897 | Anaerobic glycerol-3-phosphate dehydrogenase subunit B | 955 | gene | 1040448 | 1041758 | + | [illegible] |
| 912 | RM25_RS04410 | RM25_0898 | Glycerol-3-phosphate dehydrogenase, anaerobic, C subunit | 956 | gene | 1041755 | 1043146 | + | [illegible] |
| 913 | RM25_RS04415 | RM25_0899 | Deoxyribonucleoside regulator | 957 | gene | 1043228 | 1044196 | - | [illegible] |
| 914 | RM25_RS04420 | RM25_0900 | Auxin Efflux Carrier | 958 | gene | 1044384 | 1045319 | + | [illegible] |
| 915 | RM25_RS04425 | RM25_0901 | putative phosphoketolase | 959 | gene | 1045379 | 1048009 | + | [illegible] |
| 916 | RM25_RS04430 | RM25_0902 | ATP-dependent helicase HrpA | 960 | gene | 1048147 | 1052115 | + | [illegible] |
| 917 | RM25_RS04435 | RM25_0903 | Acyl-CoA thioesterase II | 961 | gene | 1052223 | 1053098 | + | [illegible] |
| 918 | RM25_RS04440 | RM25_0904 | Solute-binding lipoprotein | 962 | gene | 1053113 | 1054453 | - | [illegible] |
| 919 | RM25_RS12955 | RM25_0905 | Hypothetical protein | 963 | gene | 1054750 | 1055100 | + | [illegible] |
| 920 | RM25_RS12960 | RM25_0906 | LacI-type response repressor | 964 | gene | 1055097 | 1055738 | + | [illegible] |
| 921 | RM25_RS04450 | RM25_0907 | RNA polymerase sigma factor SigB | 965 | gene | 1055841 | 1056770 | - | [illegible] |
| 922 | RM25_RS04455 | RM25_0908 | RNA polymerase sigma factor | 966 | gene | 1057129 | 1058658 | + | [illegible] |
| 923 | RM25_RS04460 | RM25_0909 | L-lactate dehydrogenase | 967 | gene | 1058746 | 1059711 | - | [illegible] |
| 924 | RM25_RS04465 | RM25_0910 | Hypothetical protein | 968 | gene | 1059857 | 1060048 | - | [illegible] |
| 925 | RM25_RS04470 | RM25_0911 | DNA gyrase, B subunit | 969 | gene | 1060211 | 1062316 | + | [illegible] |
| 926 | RM25_RS04475 | RM25_0912 | 2-dehydropantoate 2-reductase (Precursor) | 970 | gene | 1062341 | 1063258 | - | [illegible] |
| 927 | RM25_RS04480 | RM25_0913 | DNA gyrase/topoisomerase IV, A subunit | 971 | gene | 1063363 | 1065825 | - | [illegible] |
| 928 | RM25_RS04485 | RM25_0914 | PF09949 family protein | 972 | gene | 1065951 | 1067036 | - | [illegible] |
| 929 | RM25_RS04490 | RM25_0915 | Hypothetical protein | 973 | gene | 1067092 | 1067685 | + | [illegible] |
| 930 | RM25_RS04495 | RM25_0916 | Type I phosphodiesterase/nucleotide pyrophosphatase | 974 | gene | 1067689 | 1068840 | + | [illegible] |
| 931 | RM25_RS04500 | RM25_0917 | Hypothetical protein | 975 | gene | 1068943 | 1070253 | - | [illegible] |
| 932 | RM25_RS04505 | RM25_0918 | PF13834 domain protein | 976 | gene | 1070469 | 1070843 | + | [illegible] |
| 933 | RM25_RS12255 | RM25_0919 | Hypothetical protein | 977 | gene | 1070955 | 1071275 | - | [illegible] |
| 934 | RM25_RS04515 | RM25_0920 | Hypothetical protein | 978 | gene | 1071565 | 1072386 | + | [illegible] |
| 935 | RM25_RS04520 | RM25_0921 | Nucleic acid binding, OB-fold, tRNA/helicase-type | 979 | gene | 1072383 | 1072763 | + | [illegible] |
| 936 | RM25_RS04525 | RM25_0922 | Trk system potassium uptake protein ceoC | 980 | gene | 1072770 | 1073438 | - | [illegible] |
| 937 | RM25_RS04530 | RM25_0923 | Trk family potassium (K+) transporter, NAD+ binding proteir | 981 | gene | 1073452 | 1074114 | - | [illegible] |
| 938 | RM25_RS04535 | RM25_0924 | Conserved alanine, valine and leucine rich membrane protei | 982 | gene | 1074287 | 1076281 | + | [illegible] |
| 939 | RM25_RS04540 | RM25_0925 | Conserved alanine, valine and glycine rich protein | 983 | gene | 1076278 | 1077456 | + | [illegible] |
| 940 | RM25_RS04545 | RM25_0926 | Hypothetical protein | 984 | gene | 1077453 | 1077713 | + | [illegible] |
| 941 | RM25_RS04550 | RM25_0927 | Putative glutathione S-transferase with thioredoxin-like dom | 985 | gene | 1077748 | 1078821 | - | [illegible] |
| 942 | RM25_RS11825 | RM25_0928 | Acetyltransferase | 986 | gene | 1078818 | 1079396 | - | [illegible] |
| 943 | RM25_RS04560 | RM25_0929 | Cysteine synthase | 987 | gene | 1079454 | 1080287 | - | [illegible] |
| 944 | NA | RM25_0930 | Hypothetical protein | 988 | gene | 1080309 | 1080428 | - | [illegible] |
| 945 | RM25_RS04565 | RM25_0931 | Putative glutathione S-transferase | 989 | gene | 1080920 | 1082035 | + | [illegible] |
| 946 | RM25_RS04570 | RM25_0932 | Alpha-ketoglutarate transporter, MFS super | 990 | gene | 1082056 | 1083486 | + | [illegible] |
| 947 | RM25_RS04575 | RM25_0933 | Aconitate hydratase 1 | 991 | gene | 1083740 | 1086412 | + | [illegible] |
| 948 | RM25_RS04580 | RM25_0934 | Hypothetical protein | 992 | gene | 1086524 | 1086826 | + | [illegible] |
| 949 | RM25_RS04585 | RM25_0935 | Hypothetical protein | 993 | gene | 1086976 | 1087506 | + | [illegible] |
| 950 | RM25_RS04590 | RM25_0936 | 1-deoxy-D-xylulose-5-phosphate synthase | 994 | gene | 1087557 | 1089572 | + | [illegible] |
| 951 | RM25_RS04595 | RM25_0937 | 3'-5' exonuclease | 995 | gene | 1089624 | 1090889 | - | [illegible] |
| 952 | RM25_RS04600 | RM25_0938 | PF11452 family protein | 996 | gene | 1090958 | 1091557 | - | [illegible] |
| 953 | RM25_RS04605 | RM25_0939 | YbaK/prolyl-tRNA synthetase-associated domain protein | 997 | gene | 1091643 | 1092167 | - | [illegible] |
| 954 | RM25_RS04610 | RM25_0940 | Methionine-R-sulfoxide reductase | 998 | gene | 1092191 | 1092643 | - | [illegible] |
| 955 | RM25_RS04615 | RM25_0941 | Polyphosphate:nucleotide phosphotransferase, PPK2 family | 999 | gene | 1092791 | 1093666 | + | [illegible] |
| 956 | RM25_RS04620 | RM25_0942 | Putative insertion element ISCmi2 transposase | 1000 | gene | 1093663 | 1094706 | - | [illegible] |
| 957 | RM25_RS04625 | RM25_0943 | Cobalamin biosynthesis protein CbiX | 1001 | gene | 1094891 | 1095733 | - | [illegible] |
| 962 | RM25_RS04650 | RM25_0947 | Threonine--tRNA ligase | 1014 | gene | 1097461 | 1099503 | + | [illegible] |
| 963 | RM25_RS04655 | RM25_0948 | Histidine triad domain protein | 1015 | gene | 1099496 | 1100101 | + | [illegible] |
| 964 | RM25_RS04660 | RM25_0949 | Phosphatidylinositol synthase pgsA1 | 1016 | gene | 1100094 | 1100723 | + | [illegible] |
| 965 | RM25_RS04665 | RM25_0950 | Putative acyltransferase | 1017 | gene | 1100756 | 1101598 | + | [illegible] |
| 966 | RM25_RS04670 | RM25_0951 | Phosphatidylinositol alpha-mannosyltransferase | 1018 | gene | 1101595 | 1102740 | + | [illegible] |
| 967 | RM25_RS04675 | RM25_0952 | Putative CDP-diacylglycerol--glycerol-3-phosphate 3-phosph | 1019 | gene | 1102780 | 1103457 | + | [illegible] |
| 968 | RM25_RS13200 | RM25_0953 | Hypothetical protein | 1020 | gene | 1103508 | 1104305 | + | [illegible] |
| 969 | RM25_RS04685 | RM25_0954 | Small basic protein (Membrane) | 1021 | gene | 1104333 | 1104665 | + | [illegible] |
| 970 | RM25_RS04690 | RM25_0955 | PF05949 family protein | 1022 | gene | 1104658 | 1105392 | + | [illegible] |
| 971 | RM25_RS04695 | RM25_0956 | Glycine cleavage system H protein | 1023 | gene | 1105504 | 1105887 | + | [illegible] |
| 972 | RM25_RS04700 | RM25_0957 | Oxoglutarate dehydrogenase inhibitor | 1024 | gene | 1106053 | 1106541 | + | [illegible] |
| 973 | RM25_RS04705 | RM25_0958 | Transcriptional regulator, MerR family | 1025 | gene | 1106541 | 1107275 | + | [illegible] |
| 974 | RM25_RS04710 | RM25_0959 | Protein of hypothetical function | 1026 | gene | 1107277 | 1107738 | + | [illegible] |
| 975 | RM25_RS04715 | RM25_0960 | Transcriptional regulator, MerR family | 1027 | gene | 1107919 | 1108494 | + | [illegible] |
| 976 | RM25_RS04720 | RM25_0961 | Glycine dehydrogenase | 1028 | gene | 1108646 | 1111561 | + | [illegible] |
| 977 | RM25_RS04725 | RM25_0962 | Inner membrane protein yhjD | 1029 | gene | 1111579 | 1113027 | + | [illegible] |
| 978 | RM25_RS04730 | RM25_0963 | YhgE/Pip C-terminal domain protein | 1030 | gene | 1113000 | 1115207 | - | [illegible] |
| 979 | RM25_RS04735 | RM25_0964 | Hypothetical protein | 1031 | gene | 1115212 | 1116024 | - | [illegible] |
| 980 | RM25_RS04740 | RM25_0965 | Transcriptional regulator, TetR family | 1032 | gene | 1116021 | 1116635 | - | [illegible] |
| 981 | RM25_RS04745 | RM25_0966 | Adenosylcobinamide-GDP ribazoletransferase | 1033 | gene | 1116729 | 1117550 | - | [illegible] |
| 982 | RM25_RS04750 | RM25_0967 | Cobinamide kinase/cobinamide phosphate guanylyltransfera | 1034 | gene | 1117510 | 1118181 | - | [illegible] |
| 983 | RM25_RS04755 | RM25_0968 | Cobyrinic acid a, c-diamide synthase | 1035 | gene | 1118178 | 1120763 | - | [illegible] |
| 984 | RM25_RS04760 | RM25_0969 | Cob(I)yrinic acid a, c-diamide adenosyltransferase | 1036 | gene | 1120757 | 1121359 | - | [illegible] |
| 985 | RM25_RS04765 | RM25_0970 | Adenosylcobyric acid synthase (glutamine-hydrolyzing) | 1037 | gene | 1121356 | 1122801 | - | [illegible] |
| 986 | RM25_RS04770 | RM25_0971 | Cobalamin biosynthesis protein CobD | 1038 | gene | 1122798 | 1123829 | - | [illegible] |
| 987 | RM25_RS04775 | RM25_0972 | Phosphoglycerate mutase family protein | 1039 | gene | 1123833 | 1124411 | + | [illegible] |
| 988 | RM25_RS04780 | RM25_0973 | Alpha, alpha-trehalose-phosphate synthase | 1040 | gene | 1124670 | 1125857 | - | [illegible] |
| 989 | RM25_RS04785 | RM25_0974 | Transposase IS204/IS1001/IS1096/IS1165 family protein | 1041 | gene | 1126228 | 1127304 | + | [illegible] |
| 990 | RM25_RS04790 | RM25_0975 | Hypothetical protein | 1042 | gene | 1127284 | 1127598 | - | [illegible] |
| 991 | RM25_RS04795 | RM25_0976 | Trehalose-6-phosphate phosphatase | 1043 | gene | 1127760 | 1128617 | + | [illegible] |
| 993 | RM25_RS04810 | RM25_0978 | ADP-ribosyl-(Dinitrogen reductase) hydrolase | 1047 | gene | 1129670 | 1130626 | - | [illegible] |
| 994 | NA | RM25_0979 | Hypothetical protein | 1048 | gene | 1130731 | 1132731 | - | [illegible] |
| 995 | RM25_RS04825 | RM25_0980 | Hypothetical protein | 1049 | gene | 1133047 | 1133628 | - | [illegible] |
| 996 | RM25_RS04830 | RM25_0981 | Hypothetical protein | 1050 | gene | 1133898 | 1134230 | - | [illegible] |
| 997 | RM25_RS04835 | RM25_0982 | Transcriptional activator, TenA family | 1051 | gene | 1134268 | 1134963 | - | [illegible] |
| 998 | RM25_RS04840 | RM25_0983 | DNA primase | 1052 | gene | 1135133 | 1137001 | + | [illegible] |
| 999 | RM25_RS11830 | RM25_0984 | RNA polymerase sigma factor | 1053 | gene | 1137074 | 1137928 | + | [illegible] |
| 1000 | RM25_RS11835 | RM25_0985 | Hypothetical protein | 1054 | gene | 1138008 | 1138574 | + | [illegible] |
| 1002 | RM25_RS04860 | RM25_0987 | Phosphofructokinase (Precursor) | 1058 | gene | 1138905 | 1140119 | - | [illegible] |
| 1003 | RM25_RS11840 | RM25_0988 | Hypothetical protein | 1059 | gene | 1140343 | 1140765 | + | [illegible] |
| 1004 | RM25_RS04870 | RM25_0989 | Sua5/YciO/YrdC/YwlC family protein | 1060 | gene | 1140834 | 1141451 | + | [illegible] |
| 1005 | RM25_RS04875 | RM25_0990 | Putative membrane protein | 1061 | gene | 1141501 | 1142118 | - | [illegible] |
| 1006 | RM25_RS04880 | RM25_0991 | 1D-myo-inositol 2-acetamido-2-deoxy-alpha-D-glucopyranos | 1062 | gene | 1142566 | 1143366 | + | [illegible] |
| 1007 | RM25_RS04885 | RM25_0992 | Proteasome assembly chaperones 2 | 1063 | gene | 1143505 | 1144542 | - | [illegible] |
| 1008 | RM25_RS04890 | RM25_0993 | ABC-type uncharacterized transport system, ATPase compo | 1064 | gene | 1144708 | 1145499 | - | [illegible] |
| 1009 | RM25_RS04895 | RM25_0994 | Inner-membrane translocator | 1065 | gene | 1145501 | 1146454 | - | [illegible] |
| 1010 | RM25_RS04900 | RM25_0995 | ABC transporter substrate binding protein | 1066 | gene | 1146461 | 1147441 | - | [illegible] |
| 1011 | RM25_RS04905 | RM25_0996 | Phosphoglycerate mutase | 1067 | gene | 1147630 | 1148253 | + | [illegible] |
| 1012 | RM25_RS04910 | RM25_0997 | Glutamine--tRNA ligase | 1068 | gene | 1148275 | 1149987 | + | [illegible] |
| 1013 | RM25_RS04915 | RM25_0998 | TIGR03085 family protein | 1069 | gene | 1150032 | 1150778 | - | [illegible] |
| 1014 | RM25_RS04920 | RM25_0999 | Phosphoribosyl-AMP cyclohydrolase | 1070 | gene | 1150794 | 1151246 | + | [illegible] |
| 1015 | RM25_RS04925 | RM25_1000 | Anthranilate synthase component I | 1071 | gene | 1151243 | 1152775 | + | [illegible] |
| 1016 | RM25_RS04930 | RM25_1001 | Indole-3-glycerol phosphate synthase | 1072 | gene | 1152772 | 1153632 | + | [illegible] |
| 1017 | RM25_RS04935 | RM25_1002 | Tryptophan synthase beta chain | 1073 | gene | 1153625 | 1154878 | + | [illegible] |
| 1018 | RM25_RS04940 | RM25_1003 | Tryptophan synthase alpha chain | 1074 | gene | 1154875 | 1155771 | + | [illegible] |
| 1019 | RM25_RS04945 | RM25_1004 | Prolipoprotein diacylglyceryl transferase | 1075 | gene | 1155768 | 1156739 | + | [illegible] |
| 1020 | RM25_RS04950 | RM25_1005 | Ferredoxin-dependent glutamate synthase [NADPH] large su | 1076 | gene | 1156883 | 1161391 | + | [illegible] |
| 1021 | RM25_RS04955 | RM25_1006 | Glutamate synthase small subunit | 1077 | gene | 1161384 | 1162853 | + | [illegible] |
| 1022 | RM25_RS04960 | RM25_1007 | Hypothetical protein | 1078 | gene | 1162873 | 1163544 | - | [illegible] |
| 1023 | RM25_RS04965 | RM25_1008 | DNA polymerase III, alpha subunit | 1079 | gene | 1163770 | 1167342 | + | [illegible] |
| 1024 | RM25_RS11845 | RM25_1009 | Putative membrane protein | 1080 | gene | 1167415 | 1168140 | + | [illegible] |
| 1025 | RM25_RS04975 | RM25_1010 | Cys-tRNA(Pro)/Cys-tRNA(Cys) deacylase ybaK | 1081 | gene | 1168177 | 1168695 | - | [illegible] |
| 1026 | NA | RM25_1011 | Hypothetical protein | 1082 | gene | 1168741 | 1169094 | - | [illegible] |
| 1027 | NA | RM25_1012 | Hypothetical protein | 1083 | gene | 1169151 | 1169531 | - | [illegible] |
| 1028 | RM25_RS04985 | RM25_1013 | Histidinol dehydrogenase | 1084 | gene | 1169609 | 1170922 | + | [illegible] |
| 1029 | RM25_RS04990 | RM25_1014 | Histidinol-phosphate transaminase | 1085 | gene | 1170922 | 1172142 | + | [illegible] |
| 1030 | RM25_RS04995 | RM25_1015 | Imidazoleglycerol-phosphate dehydratase | 1086 | gene | 1172139 | 1172747 | + | [illegible] |
| 1031 | RM25_RS05000 | RM25_1016 | Imidazole glycerol phosphate synthase, glutamine amidotra | 1087 | gene | 1172744 | 1173421 | + | [illegible] |
| 1032 | RM25_RS05005 | RM25_1017 | Phosphoribosyl isomerase A | 1088 | gene | 1173490 | 1174215 | + | [illegible] |
| 1033 | RM25_RS05010 | RM25_1018 | Mycothiol acetyltransferase | 1089 | gene | 1174212 | 1175249 | + | [illegible] |

| | | | | | | | | | | | | | | | | | | | | | | | | | | | | | | | | | | | | | | |
|---|---|---|---|---|---|---|---|---|---|---|---|---|---|---|---|---|---|---|---|---|---|---|---|---|---|---|---|---|---|---|---|---|---|---|---|---|---|---|
| 1034 | RM25_RS05015 | RM25_1019 | putative transcriptional regulatory protein | 1090 | gene | 1175413 | 1176213 | + | 3971,745 | 0,53573642 | 0,515976 | 1,038297 | 0,299132 | 0,409217 | 5767,526003 | 0,296147048 | 0,232311 | 1,274786 | 0,202385 | 0,262388 | 2175,963 | 1,220198571 | 0,457299 | 2,668273 | 0,007624 | 0,024994 | 3464,724 | -2,03169 | 0,440263 | -4,61472 | 3,94E-06 | 2,65E-05 | 4602,021025 | -1,080988462 | 0,256596 | -4,2128 | 2,52E-05 | 7,02E-05 |
| 1035 | RM25_RS05020 | RM25_1020 | Crossover junction endodeoxyribonuclease RuvC | 1091 | gene | 1176482 | 1177093 | + | 457,3797 | 0,652491292 | 0,239424 | 2,725252 | 0,006425 | 0,035179 | 529,0584232 | 0,412887093 | 0,198887 | 2,075992 | 0,037895 | 0,062635 | 385,701 | 0,999175194 | 0,250359 | 3,990964 | 6,58E-05 | 0,000518 | 378,4047 | -0,879 | 0,267673 | -3,28384 | 0,001024 | 0,003592 | 547,0157175 | -0,245952505 | 0,162048 | -1,51777 | 0,129072 | 0,173444 |
| 1036 | RM25_RS05025 | RM25_1021 | Holliday junction ATP-dependent DNA helicase RuvA | 1092 | gene | 1177090 | 1177701 | + | 414,8152 | 0,204642918 | 0,184531 | 1,108987 | 0,267436 | 0,375206 | 471,7497172 | -0,000789354 | 0,164624 | -0,00479 | 0,996174 | 0,99702 | 357,8807 | 0,477802128 | 0,183914 | 2,597968 | 0,009378 | 0,028818 | 408,8975 | -0,71734 | 0,163816 | -4,37896 | 1,19E-05 | 7,17E-05 | 435,0453979 | -0,187134553 | 0,159844 | -1,17074 | 0,241705 | 0,298897 |
| 1037 | RM25_RS05030 | RM25_1022 | Holliday junction ATP-dependent DNA helicase RuvB | 1093 | gene | 1177765 | 1178790 | + | 794,0621 | 0,437835464 | 0,241463 | 1,813262 | 0,069791 | 0,153432 | 983,7880769 | 0,53624444 | 0,155244 | 3,454196 | 0,000552 | 0,001577 | 604,3361 | 0,282507451 | 0,170774 | 1,654274 | 0,098072 | 0,170217 | 714,7259 | -0,6185 | 0,178579 | -3,46347 | 0,000533 | 0,002084 | 896,1935449 | -0,818732782 | 0,125736 | -6,51151 | 7,44E-11 | 4,95E-10 |
| 1038 | RM25_RS11860 | RM25_1023 | Preprotein translocase, YajC subunit | 1094 | gene | 1178901 | 1179404 | + | 2364,462 | 0,561427136 | 0,384578 | 1,459853 | 0,144331 | 0,244915 | 3117,591359 | 0,294178732 | 0,180747 | 1,62757 | 0,103616 | 0,146417 | 1611,333 | 1,113713694 | 0,352871 | 3,156147 | 0,001599 | 0,007137 | 2037,731 | -1,51138 | 0,316558 | -4,77441 | 1,8E-06 | 1,32E-05 | 2757,915608 | -0,656893683 | 0,22246 | -2,95287 | 0,003148 | 0,006041 |
| 1039 | RM25_RS05040 | RM25_1024 | Protein translocase subunit SecD | 1095 | gene | 1179432 | 1181150 | + | 3082,653 | 0,456513282 | 0,410713 | 1,111514 | 0,266347 | 0,373901 | 4295,438688 | 0,234415875 | 0,155685 | 1,505707 | 0,132142 | 0,182889 | 1869,867 | 0,993687265 | 0,240801 | 4,126588 | 3,68E-05 | 0,000313 | 2770,237 | -1,71908 | 0,202792 | -8,47703 | 2,31E-17 | 1,33E-15 | 3494,81427 | -0,91190121 | 0,158848 | -5,74073 | 9,43E-09 | 4,52E-08 |
| 1040 | RM25_RS05045 | RM25_1025 | Protein translocase subunit SecF | 1096 | gene | 1181150 | 1182397 | + | 3037,824 | 0,740752894 | 0,486637 | 1,522187 | 0,127962 | 0,226262 | 4393,171122 | 0,474315281 | 0,191977 | 2,470682 | 0,013486 | 0,025654 | 1682,476 | 1,516770003 | 0,264338 | 5,737987 | 9,58E-09 | 2,11E-07 | 2426,688 | -2,13487 | 0,230035 | -9,28062 | 1,68E-20 | 1,38E-18 | 3723,404406 | -1,050418166 | 0,191836 | -5,47561 | 4,36E-08 | 1,91E-07 |
| 1041 | RM25_RS05050 | RM25_1026 | Adenine phosphoribosyltransferase | 1097 | gene | 1182400 | 1182939 | + | 925,7493 | 0,860412803 | 0,40569 | 2,120862 | 0,033933 | 0,097657 | 1244,812641 | 0,572126851 | 0,190004 | 3,011134 | 0,002603 | 0,006219 | 606,6859 | 1,522217876 | 0,259947 | 5,855881 | 4,74E-09 | 1,15E-07 | 701,8619 | -1,73387 | 0,23304 | -7,44022 | 1,01E-13 | 3E-12 | 1168,609651 | -0,741403879 | 0,196701 | -3,76919 | 0,000164 | 0,000403 |
| 1042 | RM25_RS05055 | RM25_1027 | GTP diphosphokinase | 1098 | gene | 1183002 | 1185353 | + | 4192,765 | 0,704537121 | 0,31774 | 2,217338 | 0,0266 | 0,084154 | 5274,149213 | 0,44262421 | 0,175377 | 2,52385 | 0,011608 | 0,022444 | 3111,381 | 1,179538137 | 0,246912 | 4,777161 | 1,78E-06 | 2,27E-05 | 3392,39 | -1,28585 | 0,205752 | -6,24951 | 4,12E-10 | 6,47E-09 | 5085,538892 | -0,508913628 | 0,186983 | -2,72171 | 0,006494 | 0,011721 |
| 1043 | RM25_RS05060 | RM25_1028 | Conserved alanine and arginine rich protein | 1099 | gene | 1185434 | 1186573 | - | 3642,907 | 0,205959505 | 0,268109 | 0,768194 | 0,442372 | 0,550222 | 4119,782732 | 0,535003777 | 0,180292 | 2,967427 | 0,003003 | 0,007015 | 3166,031 | -0,214577133 | 0,38182 | -0,56198 | 0,574126 | 0,665953 | 3582,524 | -0,03185 | 0,366774 | -0,08683 | 0,930807 | 0,944838 | 3821,227005 | -0,748953415 | 0,203252 | -3,68486 | 0,000229 | 0,000546 |
| 1044 | RM25_RS05065 | RM25_1029 | Metallo-beta-lactamase family protein | 1100 | gene | 1186677 | 1187396 | + | 960,5823 | -0,002180805 | 0,151845 | -0,01436 | 0,988541 | 0,993176 | 1052,098591 | 0,151625342 | 0,148367 | 1,021962 | 0,306799 | 0,375259 | 869,0661 | -0,185623976 | 0,211158 | -0,87908 | 0,379359 | 0,484633 | 1017,279 | -0,1649 | 0,211648 | -0,77914 | 0,435896 | 0,511148 | 940,4561664 | -0,456744693 | 0,134183 | -3,4039 | 0,000664 | 0,001466 |
| 1045 | RM25_RS05070 | RM25_1030 | Histidine--tRNA ligase | 1101 | gene | 1187396 | 1188727 | + | 1459,822 | 0,14365433 | 0,188516 | 0,762029 | 0,446043 | 0,553911 | 1704,732783 | 0,08605819 | 0,143436 | 0,599976 | 0,548522 | 0,614188 | 1214,912 | 0,225070827 | 0,182735 | 1,231679 | 0,218069 | 0,314978 | 1471,598 | -0,62153 | 0,131939 | -4,71075 | 2,47E-06 | 1,74E-05 | 1500,893719 | -0,435440935 | 0,159918 | -2,7229 | 0,006471 | 0,011697 |
| 1046 | RM25_RS05075 | RM25_1031 | Alpha-acetolactate decarboxylase | 1102 | gene | 1188841 | 1189656 | + | 893,3455 | 0,418269061 | 0,097295 | 4,298974 | 1,72E-05 | 0,00043 | 929,6673149 | 0,37681355 | 0,125411 | 3,004621 | 0,002659 | 0,006337 | 857,0237 | 0,463975373 | 0,195694 | 2,370927 | 0,017744 | 0,047096 | 808,9492 | -0,22658 | 0,13722 | -1,65126 | 0,098687 | 0,147125 | 1000,181594 | -0,092693586 | 0,148698 | -0,62337 | 0,533041 | 0,593191 |
| 1047 | RM25_RS05080 | RM25_1032 | Hypothetical protein | 1103 | gene | 1189865 | 1190938 | + | 3271,55 | 0,039099334 | 0,344429 | 0,113519 | 0,909619 | 0,941556 | 3927,364234 | 0,553364981 | 0,201248 | 2,749666 | 0,005966 | 0,012798 | 2615,737 | -0,737542243 | 0,38584 | -1,91152 | 0,055938 | 0,109681 | 3416,775 | -0,00904 | 0,37006 | -0,02443 | 0,98051 | 0,983366 | 3248,214603 | -1,268862121 | 0,221881 | -5,71865 | 1,07E-08 | 5,1E-08 |
| 1048 | RM25_RS05085 | RM25_1033 | Aspartate--tRNA ligase | 1104 | gene | 1191138 | 1192985 | + | 3377,783 | 0,39894884 | 0,39067 | 1,021192 | 0,307163 | 0,416439 | 4599,71133 | 0,156536427 | 0,190861 | 0,82016 | 0,412125 | 0,480642 | 2155,855 | 0,941288047 | 0,245498 | 3,834193 | 0,000126 | 0,000894 | 3107,657 | -1,61709 | 0,245381 | -6,59012 | 4,39E-11 | 8,63E-10 | 3762,567719 | -0,78885554 | 0,169123 | -4,66438 | 3,1E-06 | 9,9E-06 |
| 1049 | RM25_RS05090 | RM25_1034 | Recombination factor protein RarA | 1105 | gene | 1193110 | 1194534 | + | 970,0975 | -0,228543627 | 0,13313 | -1,7167 | 0,086034 | 0,175568 | 974,9310691 | 0,058752139 | 0,127604 | 0,460425 | 0,645211 | 0,700819 | 965,2639 | -0,519531678 | 0,196528 | -2,64355 | 0,008204 | 0,026341 | 1102,867 | 0,19288 | 0,134857 | 1,430257 | 0,152643 | 0,210241 | 877,4493121 | -0,329475717 | 0,178547 | -1,84532 | 0,064991 | 0,093682 |
| 1050 | RM25_RS05095 | RM25_1035 | Lactate/malate dehydrogenase, NAD binding domain protei | 1106 | gene | 1194589 | 1195569 | + | 950,9584 | 0,049210167 | 0,192433 | 0,255726 | 0,798163 | 0,859027 | 1119,07686 | 0,128327555 | 0,142025 | 0,903556 | 0,366231 | 0,434427 | 782,8399 | -0,061336325 | 0,173604 | -0,35331 | 0,723854 | 0,795025 | 989,626 | -0,47961 | 0,165515 | -2,89772 | 0,003759 | 0,010547 | 948,4768814 | -0,616765204 | 0,125065 | -4,93155 | 8,16E-07 | 2,9E-06 |
| 1051 | RM25_RS05100 | RM25_1036 | Alanine--tRNA ligase | 1107 | gene | 1195734 | 1198421 | + | 3937,999 | 0,843852804 | 0,316191 | 2,668805 | 0,007612 | 0,038919 | 4733,756284 | 0,474018936 | 0,183436 | 2,584105 | 0,009763 | 0,019502 | 3142,241 | 1,463278027 | 0,283024 | 5,170148 | 2,34E-07 | 3,83E-06 | 3000,155 | -1,30103 | 0,268076 | -4,8532 | 1,21E-06 | 9,12E-06 | 4948,765798 | -0,270117523 | 0,179646 | -1,50361 | 0,132682 | 0,177386 |
| 1052 | RM25_RS05105 | RM25_1037 | Putative crossover junction endodeoxyribonuclease | 1108 | gene | 1198405 | 1198950 | + | 336,3485 | 0,85345758 | 0,327797 | 2,603618 | 0,009225 | 0,044102 | 419,8598807 | 0,60928576 | 0,225804 | 2,698293 | 0,00697 | 0,014537 | 252,8372 | 1,295219055 | 0,300068 | 4,316415 | 1,59E-05 | 0,000153 | 255,6683 | -1,23954 | 0,286215 | -4,33079 | 1,49E-05 | 8,8E-05 | 423,9403977 | -0,510414076 | 0,244709 | -2,0858 | 0,036997 | 0,056588 |
| 1053 | RM25_RS05110 | RM25_1038 | Amino deoxychorismate lyase | 1109 | gene | 1198925 | 1200157 | + | 1190,857 | 0,202550205 | 0,172503 | 1,174187 | 0,24032 | 0,347933 | 1354,286333 | 0,261961876 | 0,129471 | 2,023325 | 0,04304 | 0,070155 | 1027,427 | 0,125503712 | 0,230579 | 0,544299 | 0,586236 | 0,676375 | 1171,086 | -0,38421 | 0,130898 | -2,93516 | 0,003334 | 0,009641 | 1247,630746 | -0,476515219 | 0,178028 | -2,67663 | 0,007437 | 0,013329 |
| 1054 | RM25_RS05115 | RM25_1039 | Chorismate synthase | 1110 | gene | 1200295 | 1201497 | + | 1569,198 | 0,440012556 | 0,245863 | 1,789666 | 0,073508 | 0,15939 | 1927,417907 | 0,520476209 | 0,183874 | 2,830617 | 0,004646 | 0,010186 | 1210,978 | 0,315420235 | 0,210291 | 1,499925 | 0,133634 | 0,21491 | 1412,501 | -0,60714 | 0,248154 | -2,44661 | 0,014421 | 0,031589 | 1770,790371 | -0,769634864 | 0,117782 | -6,53441 | 6,39E-11 | 4,3E-10 |
| 1055 | RM25_RS05120 | RM25_1040 | 3-dehydroquinate synthase | 1111 | gene | 1201494 | 1203143 | + | 1551,548 | 0,333434727 | 0,361357 | 0,92273 | 0,356148 | 0,467134 | 2134,716666 | 0,360989324 | 0,183246 | 1,969967 | 0,048842 | 0,07747 | 968,3798 | 0,274578647 | 0,186238 | 1,474345 | 0,140389 | 0,224033 | 1458,746 | -1,15131 | 0,223331 | -5,15516 | 2,53E-07 | 2,15E-06 | 1697,223901 | -1,187702839 | 0,111288 | -10,6723 | 1,37E-26 | 6,31E-25 |
| 1056 | RM25_RS05125 | RM25_1041 | Translation elongation factor P | 1112 | gene | 1203302 | 1203883 | + | 2603,467 | 0,594742249 | 0,474261 | 1,25404 | 0,209827 | 0,314408 | 3750,505209 | 0,379186526 | 0,221193 | 1,714278 | 0,086478 | 0,124818 | 1456,428 | 1,1903354 | 0,326803 | 3,642358 | 0,00027 | 0,001689 | 2214,637 | -1,93187 | 0,305256 | -6,32867 | 2,47E-10 | 4,16E-09 | 3067,377562 | -1,08389758 | 0,230753 | -4,69722 | 2,64E-06 | 8,53E-06 |
| 1057 | RM25_RS05130 | RM25_1042 | N utilization substance protein B-like protein | 1113 | gene | 1203883 | 1204509 | + | 1003,277 | 0,484435619 | 0,544781 | 0,88923 | 0,37388 | 0,484994 | 1573,957252 | 0,416885746 | 0,211706 | 1,969176 | 0,048933 | 0,077562 | 432,5966 | 0,737423258 | 0,198871 | 3,708044 | 0,000209 | 0,001364 | 893,6252 | -2,11516 | 0,245418 | -8,6186 | 6,78E-18 | 4,56E-16 | 1148,172716 | -1,745970268 | 0,152417 | -11,4552 | 2,21E-30 | 1,41E-28 |
| 1058 | RM25_RS05135 | RM25_1043 | Carbamoyl-phosphate synthase small chain | 1114 | gene | 1204688 | 1205872 | + | 2256,257 | 0,460610981 | 0,369439 | 1,246783 | 0,212477 | 0,317658 | 3093,782746 | 0,38445265 | 0,175245 | 2,193799 | 0,02825 | 0,04839 | 1418,732 | 0,630007637 | 0,233231 | 2,701214 | 0,006909 | 0,023229 | 2020,994 | -1,32744 | 0,203875 | -6,51108 | 7,46E-11 | 1,39E-09 | 2560,681221 | -1,037480439 | 0,175066 | -5,92624 | 3,1E-09 | 1,6E-08 |
| 1059 | RM25_RS05140 | RM25_1044 | Carbamoyl-phosphate synthase large chain | 1115 | gene | 1205872 | 1209213 | + | 6215,315 | 0,623004554 | 0,404847 | 1,538865 | 0,123837 | 0,221531 | 8620,871164 | 0,436842685 | 0,179262 | 2,436902 | 0,014814 | 0,027777 | 3809,759 | 1,067269061 | 0,240746 | 4,433178 | 9,29E-06 | 9,56E-05 | 5215,866 | -1,63206 | 0,211135 | -7,72994 | 1,08E-14 | 3,9E-13 | 7383,913564 | -0,957218664 | 0,179849 | -5,32236 | 1,02E-07 | 4,18E-07 |
| 1060 | RM25_RS05145 | RM25_1045 | Dihydroorotate dehydrogenase (quinone) | 1116 | gene | 1209295 | 1210395 | + | 1684,051 | 0,62037466 | 0,452722 | 1,370321 | 0,170587 | 0,274265 | 2464,119475 | 0,535542669 | 0,173554 | 3,085737 | 0,00203 | 0,004985 | 903,9831 | 0,859594246 | 0,25158 | 3,416777 | 0,000634 | 0,003364 | 1415,854 | -1,70692 | 0,226748 | -7,52784 | 5,16E-14 | 1,62E-12 | 2000,182515 | -1,338008431 | 0,170218 | -7,86056 | 3,82E-15 | 4,85E-14 |
| 1061 | RM25_RS05150 | RM25_1046 | Orotidine 5'-phosphate decarboxylase | 1117 | gene | 1210392 | 1211201 | + | 807,6758 | 0,404014367 | 0,334993 | 1,206037 | 0,227803 | 0,335373 | 1083,010421 | 0,382679708 | 0,165615 | 2,310665 | 0,020851 | 0,037064 | 532,3411 | 0,449051661 | 0,212021 | 2,117962 | 0,034178 | 0,076269 | 739,1242 | -1,12318 | 0,190198 | -5,90535 | 3,52E-09 | 4,32E-08 | 901,9075914 | -1,007423846 | 0,158001 | -6,37608 | 1,82E-10 | 1,14E-09 |
| 1062 | RM25_RS05155 | RM25_1047 | Integration host factor family protein | 1118 | gene | 1211364 | 1211675 | + | 590,7964 | 0,598971866 | 0,321871 | 1,860906 | 0,062757 | 0,142908 | 464,3465121 | 0,355855231 | 0,216741 | 1,641848 | 0,100621 | 0,143129 | 717,2463 | 0,762448151 | 0,450121 | 1,693873 | 0,090289 | 0,158965 | 497,4885 | 0,330382 | 0,418035 | 0,790322 | 0,42934 | 0,504212 | 693,6950829 | 0,773863324 | 0,27708 | 2,792926 | 0,005223 | 0,009664 |
| 1063 | RM25_RS05160 | RM25_1048 | Guanylate kinase | 1119 | gene | 1211679 | 1212299 | + | 911,4452 | 0,406573014 | 0,216146 | 1,88101 | 0,059971 | 0,139399 | 1082,90017 | 0,496028191 | 0,160841 | 3,083972 | 0,002043 | 0,005005 | 739,9902 | 0,279988518 | 0,236995 | 1,181411 | 0,23744 | 0,336325 | 830,5852 | -0,48536 | 0,235422 | -2,06165 | 0,039241 | 0,0697 | 1018,297319 | -0,653508072 | 0,141788 | -4,60905 | 4,05E-06 | 1,27E-05 |
| 1064 | RM25_RS05165 | RM25_1049 | DNA-directed RNA polymerase, omega subunit | 1120 | gene | 1212359 | 1212727 | + | 2571,814 | 0,942821726 | 0,509881 | 1,849103 | 0,064443 | 0,145351 | 3652,306726 | 0,626261107 | 0,230591 | 2,715896 | 0,00661 | 0,013947 | 1491,322 | 1,850256787 | 0,378326 | 4,890638 | 1,01E-06 | 1,43E-05 | 1883,619 | -2,19917 | 0,377239 | -5,82966 | 5,55E-09 | 6,45E-08 | 3311,505625 | -0,943213006 | 0,225367 | -4,18523 | 2,85E-05 | 7,85E-05 |
| 1065 | RM25_RS05170 | RM25_1050 | S-adenosylmethionine synthase | 1121 | gene | 1212793 | 1213983 | + | 2256,61 | 0,885916548 | 0,486131 | 1,822381 | 0,068397 | 0,151231 | 3314,290199 | 0,715882042 | 0,214051 | 3,344448 | 0,000824 | 0,00226 | 1198,93 | 1,395154448 | 0,259422 | 5,377943 | 7,53E-08 | 1,36E-06 | 1692,932 | -1,98488 | 0,262754 | -7,55413 | 4,22E-14 | 1,36E-12 | 2869,577148 | -1,261702828 | 0,190712 | -6,61576 | 3,7E-11 | 2,54E-10 |
| 1066 | RM25_RS05175 | RM25_1051 | Primosomal protein N' | 1122 | gene | 1214094 | 1216115 | + | 1148,042 | -0,203671771 | 0,162955 | -1,24986 | 0,21135 | 0,316287 | 1062,657085 | 0,106976979 | 0,141869 | 0,754056 | 0,450816 | 0,519595 | 1233,427 | -0,474347845 | 0,229362 | -2,06812 | 0,038629 | 0,083992 | 1292,323 | 0,426156 | 0,164463 | 2,591197 | 0,009564 | 0,022886 | 1048,297294 | -0,096887454 | 0,202247 | -0,47906 | 0,631899 | 0,688256 |
| 1067 | RM25_RS05180 | RM25_1052 | Methionyl-tRNA formyltransferase | 1123 | gene | 1216193 | 1217140 | + | 954,0655 | 0,518557264 | 0,195482 | 2,652712 | 0,007985 | 0,040226 | 1097,530221 | 0,42793439 | 0,174952 | 2,446009 | 0,014445 | 0,027193 | 810,6008 | 0,643631676 | 0,21571 | 2,983776 | 0,002847 | 0,01155 | 831,0407 | -0,62033 | 0,185323 | -3,34732 | 0,000816 | 0,00295 | 1099,633017 | -0,36320199 | 0,171057 | -2,12328 | 0,033731 | 0,052099 |
| 1068 | RM25_RS05185 | RM25_1053 | Sun protein | 1124 | gene | 1217133 | 1218602 | + | 857,546 | -0,20154962 | 0,154398 | -1,30539 | 0,19176 | 0,295604 | 833,1828561 | -0,030534352 | 0,19412 | -0,1573 | 0,875012 | 0,90377 | 881,9092 | -0,363452584 | 0,241549 | -1,50467 | 0,132408 | 0,213757 | 965,9335 | 0,173292 | 0,211734 | 0,818443 | 0,413104 | 0,488554 | 783,633292 | -0,098806759 | 0,223487 | -0,44211 | 0,658407 | 0,712518 |
| 1069 | RM25_RS05190 | RM25_1054 | Ribulose-phosphate 3-epimerase (Precursor) | 1125 | gene | 1218637 | 1219299 | + | 529,3485 | 0,163716106 | 0,141198 | 1,159477 | 0,246262 | 0,354792 | 582,8674053 | 0,22410125 | 0,179953 | 1,245334 | 0,213009 | 0,274351 | 475,8295 | 0,091824459 | 0,167522 | 0,548132 | 0,583601 | 0,674619 | 527,7069 | -0,27883 | 0,168927 | -1,65057 | 0,098827 | 0,147241 | 548,1101839 | -0,364163728 | 0,159252 | -2,28671 | 0,022213 | 0,036182 |
| 1070 | RM25_RS05195 | RM25_1055 | Aminopeptidase C | 1126 | gene | 1219359 | 1220690 | + | 4290,038 | 1,087892127 | 0,438573 | 2,480528 | 0,013119 | 0,056425 | 6283,6392 | 1,186796704 | 0,181919 | 6,523761 | 6,86E-11 | 6,88E-10 | 2296,436 | 0,830013829 | 0,297786 | 2,787287 | 0,005315 | 0,018833 | 2921,513 | -1,26712 | 0,241557 | -5,24563 | 1,56E-07 | 1,4E-06 | 5719,002274 | -1,587690657 | 0,222467 | -7,13675 | 9,56E-13 | 8,34E-12 |
| 1071 | RM25_RS05200 | RM25_1056 | Proteasome assembly chaperones 2 | 1127 | gene | 1220791 | 1221600 | - | 1372,222 | 0,690904364 | 0,217952 | 3,169981 | 0,001524 | 0,012787 | 1595,360254 | 0,805686664 | 0,187386 | 4,299619 | 1,71E-05 | 6,93E-05 | 1149,083 | 0,53697359 | 0,279598 | 1,920523 | 0,054792 | 0,107926 | 1112,466 | -0,3607 | 0,275536 | -1,30908 | 0,190506 | 0,252686 | 1658,891006 | -0,591527215 | 0,179344 | -3,29828 | 0,000973 | 0,002071 |
| 1072 | RM25_RS05205 | RM25_1057 | Methionine synthase | 1128 | gene | 1221773 | 1225240 | + | 3214,751 | 0,465108093 | 0,167633 | 2,77456 | 0,005528 | 0,0317 | 3682,322692 | 0,470120226 | 0,117896 | 3,987595 | 6,67E-05 | 0,000237 | 2747,18 | 0,458828535 | 0,19822 | 2,314742 | 0,020627 | 0,05256 | 2858,095 | -0,4787 | 0,17057 | -2,80644 | 0,005009 | 0,013192 | 3655,0531 | -0,436434852 | 0,106434 | -4,10053 | 4,12E-05 | 0,00011 |
| 1073 | RM25_RS05210 | RM25_1058 | Signal peptide peptidase SppA | 1129 | gene | 1225603 | 1226859 | - | 1810,548 | 0,549190235 | 0,229225 | 2,395857 | 0,016582 | 0,064439 | 1952,301954 | 0,305285118 | 0,188914 | 1,616 | 0,106094 | 0,149739 | 1668,793 | 0,845675565 | 0,340517 | 2,483505 | 0,01301 | 0,037349 | 1560,236 | -0,59837 | 0,327207 | -1,82871 | 0,067443 | 0,108729 | 2103,510305 | -0,02370647 | 0,199508 | -0,11882 | 0,905414 | 0,923436 |
| 1074 | RM25_RS05215 | RM25_1059 | Hydrolase | 1130 | gene | 1227043 | 1227768 | + | 435,6457 | -0,07927689 | 0,235397 | -0,33678 | 0,736283 | 0,807172 | 512,8157622 | 0,156957638 | 0,161149 | 0,973989 | 0,330062 | 0,397728 | 358,4757 | -0,413284644 | 0,269542 | -1,53328 | 0,125206 | 0,204653 | 473,6674 | -0,3077 | 0,194644 | -1,58085 | 0,113913 | 0,165226 | 416,5045403 | -0,812888689 | 0,249924 | -3,25254 | 0,001144 | 0,002384 |
| 1075 | RM25_RS05220 | RM25_1060 | TRNA (Adenine-N(1)-)-methyltransferase | 1131 | gene | 1227765 | 1228835 | + | 488,9449 | -0,225711956 | 0,212242 | -1,06346 | 0,287571 | 0,397074 | 539,0303308 | 0,056805792 | 0,148918 | 0,381456 | 0,702865 | 0,754739 | 438,8596 | -0,576501979 | 0,304029 | -1,89621 | 0,057932 | 0,112849 | 555,555 | -0,06874 | 0,198304 | -0,34665 | 0,728852 | 0,774529 | 443,4439501 | -0,640787403 | 0,267394 | -2,39642 | 0,016556 | 0,027676 |
| 1076 | RM25_RS05225 | RM25_1061 | Proteasome subunit beta | 1132 | gene | 1228828 | 1229694 | + | 739,5853 | 0,460052509 | 0,320311 | 1,436268 | 0,150926 | 0,251295 | 975,0633833 | 0,756153587 | 0,16286 | 4,642962 | 3,43E-06 | 1,59E-05 | 504,1073 | -0,086303915 | 0,165705 | -0,52083 | 0,602487 | 0,690691 | 659,559 | -0,53806 | 0,180807 | -2,9759 | 0,002921 | 0,008607 | 840,4300448 | -1,33408628 | 0,134382 | -9,92759 | 3,16E-23 | 9,79E-22 |
| 1077 | RM25_RS05230 | RM25_1062 | Proteasome subunit alpha | 1133 | gene | 1229691 | 1230416 | + | 417,4775 | 0,100412913 | 0,26939 | 0,372742 | 0,709341 | 0,784954 | 516,6890876 | 0,36928957 | 0,158511 | 2,329735 | 0,01982 | 0,035445 | 318,2659 | -0,329893449 | 0,224708 | -1,4681 | 0,142078 | 0,226115 | 426,2445 | -0,40652 | 0,162154 | -2,50702 | 0,012175 | 0,027646 | 424,5423273 | -1,047537896 | 0,214802 | -4,87677 | 1,08E-06 | 3,73E-06 |
| 1078 | RM25_RS05235 | RM25_1063 | Prephenate dehydrogenase | 1134 | gene | 1230418 | 1231512 | + | 566,4552 | -0,033806299 | 0,299975 | -0,1127 | 0,910271 | 0,941556 | 696,1268692 | 0,311910659 | 0,131683 | 2,368641 | 0,017854 | 0,03252 | 436,7835 | -0,588454604 | 0,318246 | -1,84905 | 0,06445 | 0,122853 | 605,3361 | -0,30604 | 0,206828 | -1,47967 | 0,138961 | 0,194496 | 550,4611303 | -1,143236343 | 0,267063 | -4,28078 | 1,86E-05 | 5,3E-05 |
| 1079 | RM25_RS05240 | RM25_1064 | Cytidylate kinase | 1135 | gene | 1231552 | 1232223 | + | 625,1886 | 0,504270908 | 0,381717 | 1,321059 | 0,186482 | 0,291031 | 873,5248293 | 0,734496738 | 0,143496 | 5,118593 | 3,08E-07 | 1,7E-06 | 376,8524 | -0,009001732 | 0,227817 | -0,03951 | 0,968481 | 0,981811 | 549,3433 | -0,85667 | 0,166173 | -5,15527 | 2,53E-07 | 2,15E-06 | 720,3012808 | -1,542642119 | 0,200743 | -7,68468 | 1,53E-14 | 1,75E-13 |
| 1080 | RM25_RS05245 | RM25_1065 | GTPase Der | 1136 | gene | 1232338 | 1233687 | + | 2288,249 | 0,390149555 | 0,26661 | 1,463369 | 0,143366 | 0,243982 | 2723,268326 | 0,664843856 | 0,192965 | 3,44541 | 0,00057 | 0,001625 | 1853,23 | -0,0007794 | 0,324602 | -0,0024 | 0,998084 | 0,998932 | 2097,858 | -0,23273 | 0,335247 | -0,69421 | 0,487551 | 0,556863 | 2543,047846 | -0,8635628 | 0,162148 | -5,32576 | 1,01E-07 | 4,12E-07 |
| 1081 | RM25_RS05250 | RM25_1066 | Protein of hypothetical function DUF1706 | 1137 | gene | 1233722 | 1234231 | - | 425,3974 | -0,3553088 | 0,135157 | -2,62886 | 0,008567 | 0,042422 | 445,2885654 | -0,148626956 | 0,156832 | -0,94769 | 0,34329 | 0,412193 | 405,5062 | -0,579890202 | 0,208113 | -2,78641 | 0,005329 | 0,018833 | 504,3478 | -0,00529 | 0,172014 | -0,03076 | 0,975458 | 0,98045 | 366,1413773 | -0,386582541 | 0,190685 | -2,02734 | 0,042628 | 0,064241 |
| 1082 | RM25_RS05255 | RM25_1067 | Antiseptic resistance protein | 1138 | gene | 1234524 | 1235858 | + | 1339,607 | -0,145015295 | 0,509703 | -0,28451 | 0,77602 | 0,840226 | 2052,460423 | -0,014975873 | 0,168379 | -0,08894 | 0,929128 | 0,946394 | 626,7531 | -0,573512482 | 0,212825 | -2,69477 | 0,007044 | 0,023583 | 1498,527 | -1,51793 | 0,187846 | -8,08072 | 6,44E-16 | 3,1E-14 | 1250,316912 | -2,020703446 | 0,192258 | -10,5104 | 7,74E-26 | 3,32E-24 |
| 1083 | RM25_RS05260 | RM25_1068 | Regulatory protein, TetR | 1139 | gene | 1235884 | 1236462 | + | 295,3818 | -0,53904103 | 0,232357 | -2,31988 | 0,020347 | 0,072518 | 298,43935 | -0,281596529 | 0,172644 | -1,63108 | 0,102874 | 0,14563 | 292,3242 | -0,810113063 | 0,39787 | -2,03613 | 0,041738 | 0,088547 | 368,7677 | 0,123474 | 0,194782 | 0,633906 | 0,526142 | 0,593074 | 237,2842375 | -0,337744211 | 0,39112 | -0,86353 | 0,387845 | 0,452774 |
| 1084 | RM25_RS05265 | RM25_1069 | Oxidoreductase, zinc-binding dehydrogenase family | 1140 | gene | 1236508 | 1237458 | - | 278,1596 | -0,079473601 | 0,185972 | -0,42734 | 0,669131 | 0,750156 | 243,6338213 | -0,01112591 | 0,196965 | -0,05649 | 0,954954 | 0,964365 | 312,6854 | -0,133080249 | 0,264474 | -0,50319 | 0,614832 | 0,702183 | 300,8373 | 0,358516 | 0,167065 | 2,145975 | 0,031875 | 0,059344 | 265,7489039 | 0,297760349 | 0,279234 | 1,066348 | 0,286266 | 0,347083 |
| 1085 | RM25_RS05270 | RM25_1070 | Inner membrane transporter rhtA | 1141 | gene | 1237567 | 1238442 | - | 493,0543 | -0,401802019 | 0,147147 | -2,73062 | 0,006322 | 0,034894 | 498,4872114 | -0,168469299 | 0,142792 | -1,17982 | 0,238071 | 0,301846 | 487,6215 | -0,642556839 | 0,253759 | -2,53216 | 0,011336 | 0,033496 | 592,0563 | 0,107792 | 0,178221 | 0,60482 | 0,545298 | 0,610579 | 417,7494955 | -0,303238038 | 0,235973 | -1,28505 | 0,198773 | 0,252975 |
| 1086 | RM25_RS05275 | RM25_1071 | Hypothetical protein | 1142 | gene | 1238613 | 1239134 | + | 236,8351 | -0,230777832 | 0,234783 | -0,98294 | 0,325636 | 0,435713 | 187,196106 | -0,071970462 | 0,222012 | -0,32417 | 0,745806 | 0,79326 | 286,4741 | -0,338471995 | 0,214734 | -1,57624 | 0,11497 | 0,192188 | 268,8993 | 0,673884 | 0,202561 | 3,326822 | 0,000878 | 0,003157 | 213,9223125 | 0,465579991 | 0,242067 | 1,923352 | 0,054436 | 0,079693 |
| 1087 | RM25_RS05280 | RM25_1072 | Transcriptional regulator | 1143 | gene | 1239127 | 1239432 | + | 98,06834 | -0,624773908 | 0,350765 | -1,78117 | 0,074884 | 0,161337 | 70,92649363 | -1,03738419 | 0,376495 | -2,75537 | 0,005863 | 0,012619 | 125,2102 | -0,409332486 | 0,271293 | -1,50882 | 0,131345 | 0,212478 | 125,7981 | 0,503003 | 0,329854 | 1,524924 | 0,127278 | 0,181265 | 75,69455957 | 1,204870972 | 0,36484 | 3,302461 | 0,000958 | 0,002044 |
| 1089 | RM25_RS05290 | RM25_1073 | S-ribosylhomocysteine lyase | 1147 | gene | 1239829 | 1240293 | + | 1000,598 | 1,070321498 | 0,251537 | 4,255134 | 2,09E-05 | 0,000492 | 1112,462333 | 0,794489325 | 0,187931 | 4,22755 | 2,36E-05 | 9,31E-05 | 888,734 | 1,446129716 | 0,326705 | 4,426405 | 9,58E-06 | 9,82E-05 | 685,122 | -0,82406 | 0,234707 | -3,51104 | 0,000446 | 0,001789 | 1324,078783 | -0,136524388 | 0,258114 | -0,52893 | 0,596854 | 0,657069 |
| 1090 | RM25_RS05295 | RM25_1074 | Antiseptic resistance protein | 1148 | gene | 1240485 | 1242065 | + | 2822,571 | 0,874559541 | 0,42795 | 2,043602 | 0,040993 | 0,109812 | 4130,365123 | 0,994560451 | 0,169196 | 5,878152 | 4,15E-09 | 3,24E-08 | 1514,776 | 0,561595245 | 0,214632 | 2,616555 | 0,008882 | 0,027877 | 2119,357 | -1,23696 | 0,229192 | -5,39705 | 6,77E-08 | 6,49E-07 | 3584,073396 | -1,617122206 | 0,107706 | -15,0142 | 5,92E-51 | 1,75E-48 |
| 1091 | RM25_RS05300 | RM25_1075 | ABC-type nitrate/sulfonate/bicarbonate transport system, p | 1149 | gene | 1242372 | 1243262 | + | 968,0802 | -0,301009581 | 0,220711 | -1,36382 | 0,172626 | 0,276064 | 807,8368934 | -0,44864619 | 0,222696 | -2,01461 | 0,043946 | 0,071385 | 1128,323 | -0,198094319 | 0,21888 | -0,90504 | 0,365446 | 0,472667 | 1124,772 | 0,304142 | 0,271233 | 1,121332 | 0,262147 | 0,333628 | 849,8732044 | 0,612488254 | 0,156657 | 3,909739 | 9,24E-05 | 0,000235 |
| 1092 | RM25_RS05305 | RM25_1076 | Putative sulfonate/nitrate/taurine transport system substrat | 1150 | gene | 1243259 | 1244266 | + | 978,8154 | 1,384685245 | 0,250158 | 5,53525 | 3,11E-08 | 2,22E-06 | 812,286339 | 0,701991214 | 0,129449 | 5,422917 | 5,86E-08 | 3,71E-07 | 1145,345 | 1,969477219 | 0,268031 | 7,347953 | 2,01E-13 | 1,28E-11 | 574,09 | -0,472 | 0,181737 | -2,59718 | 0,009399 | 0,02256 | 1382,287862 | 0,844877584 | 0,19205 | 4,399252 | 1,09E-05 | 3,17E-05 |
| 1093 | RM25_RS05310 | RM25_1077 | Homoserine O-acetyltransferase | 1151 | gene | 1244448 | 1245701 | + | 1089,633 | 0,050613455 | 0,112786 | 0,448757 | 0,653607 | 0,737459 | 1132,235585 | 0,086822347 | 0,132197 | 0,656765 | 0,511332 | 0,57638 | 1047,031 | 0,013273639 | 0,222153 | 0,05975 | 0,952355 | 0,970768 | 1128,892 | -0,13606 | 0,184264 | -0,73839 | 0,46028 | 0,533372 | 1086,38976 | -0,160609071 | 0,130689 | -1,22894 | 0,219093 | 0,275414 |
| 1094 | RM25_RS05315 | RM25_1078 | Methionine biosynthesis protein MetW | 1152 | gene | 1245698 | 1246384 | + | 694,8906 | 0,004018883 | 0,130412 | 0,030817 | 0,975416 | 0,984606 | 714,3132637 | -0,04365551 | 0,155173 | -0,28134 | 0,778453 | 0,822504 | 675,468 | 0,054994667 | 0,247827 | 0,221908 | 0,824386 | 0,871938 | 731,6994 | -0,19348 | 0,224442 | -0,86207 | 0,388651 | 0,464687 | 682,5190818 | -0,039104868 | 0,15297 | -0,25564 | 0,798231 | 0,836563 |
| 1095 | RM25_RS05320 | RM25_1079 | AsnC-family transcriptional regulatory protein | 1153 | gene | 1246520 | 1246933 | + | 776,2701 | 0,432933512 | 0,1403 | 3,085764 | 0,00203 | 0,015537 | 839,9182761 | 0,583021934 | 0,123603 | 4,716881 | 2,39E-06 | 1,14E-05 | 712,6219 | 0,261147465 | 0,231528 | 1,127928 | 0,25935 | 0,360543 | 698,1717 | -0,11584 | 0,144722 | -0,80043 | 0,423461 | 0,4988 | 873,4322571 | -0,38920308 | 0,184354 | -2,11117 | 0,034758 | 0,053545 |
| 1096 | RM25_RS05325 | RM25_1080 | Apolipoprotein N-acyltransferase | 1154 | gene | 1247027 | 1248661 | + | 1110,023 | 0,332417187 | 0,119151 | 2,789874 | 0,005273 | 0,030839 | 1180,97328 | 0,517941323 | 0,124237 | 4,168992 | 3,06E-05 | 0,000118 | 1039,072 | 0,127911696 | 0,183133 | 0,698465 | 0,484886 | 0,58549 | 1036,734 | -0,02822 | 0,149939 | -0,18822 | 0,850701 | 0,882916 | 1214,528534 | -0,362240109 | 0,14496 | -2,4989 | 0,012458 | 0,021379 |
| 1097 | RM25_RS05330 | RM25_1081 | Polyprenol-phosphate-mannose synthase domain 1 | 1155 | gene | 1248748 | 1249554 | + | 2114 | 0,734240867 | 0,501443 | 1,464255 | 0,143124 | 0,243746 | 3061,50217 | 0,480189453 | 0,20331 | 2,361855 | 0,018184 | 0,033096 | 1166,498 | 1,473235236 | 0,372515 | 3,954833 | 7,66E-05 | 0,00059 | 1697,757 | -2,1084 | 0,273664 | -7,70433 | 1,32E-14 | 4,63E-13 | 2584,666199 | -1,075878644 | 0,291734 | -3,68787 | 0,000226 | 0,000541 |
| 1098 | RM25_RS05335 | RM25_1082 | Putative membrane-associated GTPase | 1156 | gene | 1249650 | 1249988 | - | 19820,05 | -1,359663618 | 0,517585 | -2,62694 | 0,008616 | 0,042478 | 10210,10137 | -1,120397638 | 0,275248 | -4,07051 | 4,69E-05 | 0,000174 | 29429,99 | -1,446761293 | 0,520375 | -2,78023 | 0,005432 | 0,019081 | 29981,94 | 1,573754 | 0,500506 | 3,144325 | 0,001665 | 0,00539 | 10806,04812 | 1,273475623 | 0,310118 | 4,106428 | 4,02E-05 | 0,000108 |
| 1099 | RM25_RS05340 | RM25_1083 | Glycerophosphoryl diester phosphodiesterase glpQ2 | 1157 | gene | 1250095 | 1250892 | - | 713,2116 | 0,390642734 | 0,222061 | 1,759169 | 0,078549 | 0,165599 | 841,8770674 | 0,659451064 | 0,194262 | 3,394646 | 0,000687 | 0,001921 | 584,5462 | 0,017873097 | 0,179989 | 0,099301 | 0,920899 | 0,944954 | 652,1581 | -0,22309 | 0,207334 | -1,07601 | 0,281924 | 0,355726 | 793,0911425 | -0,820612866 | 0,15243 | -5,38354 | 7,3E-08 | 3,07E-07 |
| 1100 | RM25_RS11865 | RM25_1084 | Hypothetical protein | 1158 | gene | 1250937 | 1251461 | - | 317,6973 | 0,389117008 | 0,136726 | 2,845963 | 0,004428 | 0,027249 | 300,9126733 | 0,618421091 | 0,191171 | 3,234913 | 0,001217 | 0,00319 | 334,4819 | 0,190371338 | 0,180461 | 1,054915 | 0,291464 | 0,39459 | 289,9725 | 0,328423 | 0,205309 | 1,59965 | 0,109676 | 0,160066 | 353,3926607 | -0,043454731 | 0,174913 | -0,24844 | 0,803796 | 0,841511 |
| 1101 | RM25_RS05350 | RM25_1085 | Pyridoxal kinase | 1159 | gene | 1251599 | 1252303 | + | 289,7368 | -0,739291227 | 0,29387 | -2,51571 | 0,011879 | 0,052141 | 213,947406 | -0,790863602 | 0,25015 | -3,16156 | 0,001569 | 0,003964 | 365,5261 | -0,713853303 | 0,264246 | -2,70147 | 0,006903 | 0,023229 | 381,1735 | 0,675753 | 0,235807 | 2,865698 | 0,004161 | 0,011351 | 213,1277816 | 0,817181758 | 0,29707 | 2,750806 | 0,005945 | 0,010862 |
| 1102 | RM25_RS05355 | RM25_1086 | UDP-N-acetylglucosamine transferase | 1160 | gene | 1252263 | 1253732 | - | 2090,859 | 0,122732643 | 0,155502 | 0,789265 | 0,429957 | 0,538189 | 2117,171292 | -0,229913699 | 0,140289 | -1,63885 | 0,101244 | 0,143882 | 2064,546 | 0,487536861 | 0,193858 | 2,514919 | 0,011906 | 0,03486 | 2119,877 | -0,47172 | 0,146938 | -3,21033 | 0,001326 | 0,004503 | 2132,458494 | 0,294602973 | 0,147056 | 2,003344 | 0,04514 | 0,06751 |
| 1103 | RM25_RS05360 | RM25_1087 | Dihydrolipoyl dehydrogenase | 1161 | gene | 1253945 | 1255345 | + | 5964,187 | 1,39066118 | 0,206523 | 6,733699 | 1,65E-11 | 2,6E-09 | 6104,302325 | 1,307339177 | 0,156142 | 8,372783 | 5,63E-17 | 1,18E-15 | 5824,071 | 1,480759695 | 0,368139 | 4,02228 | 5,76E-05 | 0,000467 | 3488,26 | -0,24346 | 0,282265 | -0,86251 | 0,388408 | 0,464687 | 8432,624306 | -0,039154553 | 0,258057 | -0,15173 | 0,879401 | 0,90198 |
| 1104 | RM25_RS05365 | RM25_1088 | Receptor ligand binding region family protein | 1162 | gene | 1255541 | 1256704 | + | 2185,754 | 0,689207436 | 0,311558 | 2,212133 | 0,026957 | 0,084269 | 2827,5521 | 0,572848111 | 0,172553 | 3,319838 | 0,000901 | 0,002443 | 1543,956 | 0,909177836 | 0,215966 | 4,209828 | 2,56E-05 | 0,000229 | 1779,021 | -1,14204 | 0,191873 | -5,95209 | 2,65E-09 | 3,35E-08 | 2642,936575 | -0,76060322 | 0,166062 | -4,58023 | 4,64E-06 | 1,45E-05 |
| 1105 | RM25_RS05370 | RM25_1089 | ABC transporter membrane-spanning permease-branched cl | 1163 | gene | 1256801 | 1257685 | + | 1380,503 | 0,417420138 | 0,21159 | 1,972774 | 0,048521 | 0,122315 | 1633,16581 | 0,28302678 | 0,132369 | 2,138169 | 0,032503 | 0,054839 | 1127,84 | 0,615707326 | 0,194898 | 3,159118 | 0,001582 | 0,007098 | 1255,181 | -0,78725 | 0,178482 | -4,4108 | 1,03E-05 | 6,32E-05 | 1547,072161 | -0,403493032 | 0,12326 | -3,27352 | 0,001062 | 0,002225 |
| 1106 | RM25_RS05375 | RM25_1090 | Branched-chain amino acid transport system / permease co | 1164 | gene | 1257682 | 1258701 | + | 1208,392 | 0,547706297 | 0,240791 | 2,274609 | 0,022929 | 0,077762 | 1402,78283 | 0,340556944 | 0,139186 | 2,446781 | 0,014414 | 0,027157 | 1014,001 | 0,844328158 | 0,306469 | 2,755017 | 0,005869 | 0,020381 | 1040,4 | -0,83597 | 0,254667 | -3,28259 | 0,001029 | 0,003602 | 1404,337237 | -0,281052487 | 0,17362 | -1,61878 | 0,105494 | 0,144732 |
| 1107 | RM25_RS05380 | RM25_1091 | High-affinity branched-chain amino acid transport ATP-bindi | 1165 | gene | 1258698 | 1259489 | + | 959,6833 | 0,903394248 | 0,30437 | 2,968076 | 0,002997 | 0,020774 | 1188,46157 | 0,656103234 | 0,181135 | 3,622169 | 0,000292 | 0,000902 | 730,9051 | 1,33769227 | 0,246661 | 5,423199 | 5,85E-08 | 1,07E-06 | 711,217 | -1,21463 | 0,20719 | -5,86242 | 4,56E-09 | 5,43E-08 | 1224,00507 | -0,487892731 | 0,189404 | -2,57594 | 0,009997 | 0,017455 |
| 1108 | RM25_RS05385 | RM25_1092 | High-affinity branched-chain amino acid transport ATP-bindi | 1166 | gene | 1259482 | 1260186 | + | 1035,926 | 0,625532317 | 0,180466 | 3,4662 | 0,000528 | 0,006342 | 1161,501691 | 0,548680966 | 0,157275 | 3,488673 | 0,000485 | 0,001411 | 910,3502 | 0,725319193 | 0,249431 | 2,907901 | 0,003639 | 0,014106 | 862,867 | -0,51853 | 0,180531 | -2,87227 | 0,004075 | 0,011156 | 1229,864242 | -0,297068003 | 0,191907 | -1,54798 | 0,121628 | 0,164568 |
| 1109 | RM25_RS05390 | RM25_1093 | AcuB family protein | 1167 | gene | 1260188 | 1260838 | + | 1029,098 | 0,432457253 | 0,231077 | 1,871485 | 0,061278 | 0,1416 | 1083,08572 | 0,762765555 | 0,185521 | 4,111489 | 3,93E-05 | 0,000148 | 975,11 | 0,080857665 | 0,358603 | 0,22548 | 0,821606 | 0,870349 | 925,3451 | 0,189089 | 0,333168 | 0,567548 | 0,570342 | 0,634701 | 1156,469898 | -0,45950434 | 0,214651 | -2,1407 | 0,032298 | 0,050279 |
| 1110 | RM25_RS12735 | RM25_1094 | PTS system, mannose/fructose/sorbose family, IIA compon | 1168 | gene | 1260870 | 1261292 | - | 680,9513 | -0,502741752 | 0,424752 | -1,18361 | 0,236566 | 0,343977 | 881,3572444 | -0,028042602 | 0,225631 | -0,12429 | 0,90109 | 0,925836 | 480,5453 | -1,471636105 | 0,3664 | -4,01647 | 5,91E-05 | 0,000477 | 846,8863 | -0,38503 | 0,372659 | -1,0332 | 0,301512 | 0,376212 | 553,3390134 | -1,785495474 | 0,231816 | -7,70221 | 1,34E-14 | 1,53E-13 |
| 1111 | RM25_RS05400 | RM25_1095 | Dipeptidase pepE | 1169 | gene | 1261765 | 1262898 | + | 1003,02 | -0,20246275 | 0,137512 | -1,47233 | 0,140932 | 0,241408 | 936,9343697 | 0,074727859 | 0,128026 | 0,583692 | 0,559428 | 0,624028 | 1069,106 | -0,446469812 | 0,1919 | -2,32657 | 0,019988 | 0,051264 | 1130,589 | 0,374273 | 0,177884 | 2,104032 | 0,035376 | 0,06481 | 914,3267787 | -0,09610118 | 0,131755 | -0,72939 | 0,465762 | 0,529912 |
| 1112 | RM25_RS05405 | RM25_1096 | Putative esterase | 1170 | gene | 1262928 | 1264244 | - | 1017,183 | 0,435457903 | 0,202856 | 2,146635 | 0,031822 | 0,093992 | 1156,88871 | 0,255027553 | 0,138657 | 1,839275 | 0,065875 | 0,098959 | 877,4772 | 0,679099546 | 0,260172 | 2,610195 | 0,009049 | 0,028138 | 916,2568 | -0,70585 | 0,180234 | -3,91631 | 8,99E-05 | 0,000442 | 1144,055225 | -0,235360651 | 0,183645 | -1,28161 | 0,199981 | 0,2541 |
| 1113 | RM25_RS05410 | RM25_1097 | Hypothetical protein | 1171 | gene | 1264241 | 1266724 | - | 2694,569 | 0,521796802 | 0,296322 | 1,760909 | 0,078254 | 0,165486 | 3502,996215 | 0,502306506 | 0,136779 | 3,672394 | 0,00024 | 0,000756 | 1886,142 | 0,558952041 | 0,195204 | 2,863428 | 0,004191 | 0,015704 | 2350,728 | -0,98821 | 0,171993 | -5,74562 | 9,16E-09 | 1,03E-07 | 3113,990852 | -0,880782079 | 0,12453 | -7,07284 | 1,52E-12 | 1,28E-11 |
| 1114 | RM25_RS05415 | RM25_1098 | Aspartyl/glutamyl-tRNA(Asn/Gln) amidotransferase subunit | 1172 | gene | 1266933 | 1268444 | - | 4214,046 | 0,720878538 | 0,515029 | 1,399685 | 0,161608 | 0,264453 | 6135,454267 | 0,422751658 | 0,19819 | 2,133063 | 0,03292 | 0,055462 | 2292,638 | 1,627120537 | 0,334231 | 4,868252 | 1,13E-06 | 1,56E-05 | 3401,213 | -2,27661 | 0,318631 | -7,14496 | 9E-13 | 2,33E-11 | 5137,338817 | -1,037277434 | 0,20123 | -5,15468 | 2,54E-07 | 9,72E-07 |
| 1115 | RM25_RS05420 | RM25_1099 | Glutamyl-tRNA(Gln) amidotransferase subunit A | 1173 | gene | 1268441 | 1269973 | - | 2211,665 | 0,437309864 | 0,363491 | 1,203082 | 0,228945 | 0,336423 | 2999,653506 | 0,305088209 | 0,193186 | 1,579247 | 0,114279 | 0,160113 | 1423,676 | 0,723869045 | 0,204668 | 3,536804 | 0,000405 | 0,002375 | 1999,142 | -1,3784 | 0,233694 | -5,89833 | 3,67E-09 | 4,46E-08 | 2493,630813 | -0,914847753 | 0,134246 | -6,81469 | 9,45E-12 | 7,02E-11 |
| 1116 | RM25_RS05425 | RM25_1100 | Aspartyl/glutamyl-tRNA(Asn/Gln) amidotransferase subunit | 1174 | gene | 1269981 | 1270283 | - | 968,5597 | 0,439544458 | 0,368406 | 1,193097 | 0,232831 | 0,340281 | 1314,895676 | 0,372239315 | 0,190007 | 1,959083 | 0,050103 | 0,079151 | 622,2237 | 0,586523777 | 0,29203 | 2,008435 | 0,044597 | 0,092694 | 875,2675 | -1,26116 | 0,224364 | -5,62105 | 1,9E-08 | 2,01E-07 | 1091,478732 | -1,007182382 | 0,237194 | -4,24623 | 2,17E-05 | 6,13E-05 |
| 1117 | RM25_RS05430 | RM25_1101 | Amino acid-binding ACT domain protein | 1175 | gene | 1270504 | 1271130 | + | 1514,894 | -0,12231054 | 0,440105 | -0,27791 | 0,78108 | 0,842951 | 879,0057422 | 0,137813586 | 0,183103 | 0,752655 | 0,451657 | 0,520057 | 2150,782 | -0,227991331 | 0,417721 | -0,5458 | 0,585205 | 0,67581 | 1660,248 | 1,423522 | 0,400358 | 3,555624 | 0,000377 | 0,001557 | 1414,204786 | 1,088370884 | 0,211002 | 5,158108 | 2,49E-07 | 9,58E-07 |
| 1118 | RM25_RS05435 | RM25_1102 | putative membrane anchored protein | 1176 | gene | 1271151 | 1271354 | - | 321,2129 | 0,038712546 | 0,15815 | 0,244783 | 0,806624 | 0,864974 | 294,8635154 | -0,043441643 | 0,194121 | -0,22379 | 0,822923 | 0,859068 | 347,5622 | 0,10695496 | 0,247819 | 0,431585 | 0,666043 | 0,746843 | 334,1484 | 0,09435 | 0,229837 | 0,410507 | 0,681434 | 0,731393 | 318,8017804 | 0,30076706 | 0,199114 | 1,510527 | 0,130909 | 0,175413 |
| 1119 | RM25_RS05440 | RM25_1103 | Methionine synthase, vitamin-B12 independent | 1177 | gene | 1271423 | 1272409 | - | 436,5107 | -0,368608921 | 0,210858 | -1,74814 | 0,08044 | 0,167893 | 371,6343156 | -0,250269616 | 0,17915 | -1,39699 | 0,162417 | 0,21751 | 501,387 | -0,456977094 | 0,295994 | -1,54387 | 0,122619 | 0,201403 | 517,1954 | 0,466317 | 0,160852 | 2,899042 | 0,003743 | 0,010526 | 374,6881127 | 0,318983699 | 0,294479 | 1,083215 | 0,278713 | 0,340024 |
| 1120 | RM25_RS05445 | RM25_1104 | TRNA (5-methylaminomethyl-2-thiouridylate)-methyltransfe | 1178 | gene | 1272455 | 1273528 | - | 894,9727 | 0,008972299 | 0,083309 | 0,1077 | 0,914234 | 0,943041 | 907,5114435 | -0,1341157 | 0,121613 | -1,1028 | 0,270112 | 0,336143 | 882,4339 | 0,153829584 | 0,162693 | 0,945519 | 0,344394 | 0,450714 | 943,5889 | -0,2452 | 0,124741 | -1,9657 | 0,049333 | 0,084016 | 879,4136427 | 0,094437169 | 0,128211 | 0,736578 | 0,461379 | 0,526111 |
| 1121 | RM25_RS05450 | RM25_1105 | Glycogen debranching enzyme GlgX | 1179 | gene | 1273785 | 1275977 | + | 3003,814 | 0,206885095 | 0,191429 | 1,080741 | 0,279812 | 0,389096 | 3455,396777 | 0,179204161 | 0,135181 | 1,325661 | 0,184952 | 0,243402 | 2552,232 | 0,244955449 | 0,251642 | 0,97343 | 0,33034 | 0,436025 | 2954,668 | -0,53037 | 0,147061 | -3,60646 | 0,00031 | 0,001325 | 3149,027593 | -0,422885503 | 0,196233 | -2,15502 | 0,03116 | 0,048898 |
| 1122 | RM25_RS05455 | RM25_1106 | Hypothetical protein | 1180 | gene | 1276195 | 1276512 | + | 665,6902 | -0,745104982 | 0,350447 | -2,12616 | 0,03349 | 0,097092 | 702,7207924 | -0,111762103 | 0,216131 | -0,5171 | 0,605083 | 0,664887 | 628,6597 | -1,548468309 | 0,443514 | -3,49136 | 0,000481 | 0,002703 | 880,5672 | 0,311845 | 0,445814 | 0,699497 | 0,484241 | 0,554326 | 487,674685 | -1,090373897 | 0,204847 | -5,32286 | 1,02E-07 | 4,18E-07 |
| 1123 | NA | RM25_1107 | Transposase IS3/IS911 family protein | 1181 | gene | 1276564 | 1276893 | + | 0 | NA | NA | NA | NA | NA | 0 | NA | NA | NA | NA | NA | 0 | NA | NA | NA | NA | NA | 0 | NA | NA | NA | NA | NA | 0 | NA | NA | NA | NA | NA |
| 1124 | RM25_RS05465 | RM25_1108 | IS3 family transposase OrfB | 1182 | gene | 1276890 | 1277810 | + | 7,61001 | -1,661069063 | 0,699935 | -2,37318 | 0,017636 | 0,066722 | 4,599713148 | -1,889337532 | 1,15558 | -1,63497 | 0,102055 | 0,144732 | 10,62031 | -1,601334096 | 0,747877 | -2,14117 | 0,03226 | 0,073112 | 12,2204 | 1,055219 | 0,508332 | 2,075844 | 0,037908 | 0,06805 | 3,560851562 | 1,418230853 | 1,505888 | 0,94179 | 0,3463 | 0,40998 |
| 1125 | RM25_RS05470 | RM25_1109 | putative secreted protein | 1183 | gene | 1278257 | 1278949 | + | 652,706 | -2,063527788 | 0,861397 | -2,39556 | 0,016595 | 0,064439 | 352,4032409 | -5,622258656 | 0,446675 | -12,5869 | 2,49E-36 | 5,34E-34 | 953,0087 | -1,531924329 | 0,383357 | -3,99608 | 6,44E-05 | 0,000513 | 1101,485 | 0,965706 | 0,461861 | 2,090903 | 0,036537 | 0,066346 | 247,8615402 | 5,128023262 | 0,400063 | 12,81803 | 1,3E-37 | 1,39E-35 |
| 1126 | RM25_RS05475 | RM25_1110 | Alpha-glucan phosphorylase | 1184 | gene | 1279091 | 1281682 | - | 4324,718 | 0,865557398 | 0,274117 | 3,157625 | 0,001591 | 0,013018 | 5389,912307 | 0,826440399 | 0,161275 | 5,124403 | 2,98E-07 | 1,66E-06 | 3259,524 | 0,931666434 | 0,272002 | 3,425225 | 0,000614 | 0,003283 | 3255,716 | -0,84995 | 0,191867 | -4,42993 | 9,43E-06 | 5,83E-05 | 5465,247023 | -0,704884173 | 0,214348 | -3,2885 | 0,001007 | 0,002133 |
| 1127 | RM25_RS05480 | RM25_1111 | Alpha-1, 4-glucan:maltose-1-phosphate maltosyltransferase | 1185 | gene | 1281808 | 1283922 | + | 2604,413 | -0,66490007 | 0,26019 | -2,55544 | 0,010605 | 0,048918 | 1941,389254 | -0,497884913 | 0,153381 | -3,24607 | 0,00117 | 0,003088 | 3267,437 | -0,766791474 | 0,225965 | -3,3934 | 0,00069 | 0,003608 | 3356,614 | 0,796689 | 0,227149 | 3,507335 | 0,000453 | 0,001802 | 1970,601953 | 0,575029663 | 0,130101 | 4,419872 | 9,88E-06 | 2,89E-05 |
| 1128 | RM25_RS05485 | RM25_1112 | Maltose alpha-D-glucosyltransferase | 1186 | gene | 1284008 | 1285813 | + | 2340,036 | 0,025653361 | 0,201709 | 0,12718 | 0,898798 | 0,933245 | 2719,527614 | 0,074977584 | 0,13233 | 0,566595 | 0,57099 | 0,632435 | 1960,543 | -0,042081803 | 0,260573 | -0,1615 | 0,871702 | 0,906552 | 2451,529 | -0,47567 | 0,217098 | -2,19104 | 0,028449 | 0,054076 | 2312,742904 | -0,542409684 | 0,14458 | -3,75163 | 0,000176 | 0,000428 |
| 1129 | RM25_RS05490 | RM25_1113 | Maltokinase | 1187 | gene | 1285829 | 1287166 | + | 1439,58 | -0,153676906 | 0,231126 | -0,66491 | 0,50611 | 0,603085 | 1310,050567 | 0,375919278 | 0,129624 | 2,900067 | 0,003731 | 0,008513 | 1569,109 | -0,600679347 | 0,302175 | -1,98785 | 0,046828 | 0,09648 | 1596,35 | 0,675278 | 0,29448 | 2,29312 | 0,021841 | 0,04378 | 1335,226638 | -0,257920788 | 0,132007 | -1,95385 | 0,050719 | 0,074809 |
| 1130 | RM25_RS05495 | RM25_1114 | 1, 4-alpha-glucan branching enzyme GlgB | 1188 | gene | 1287159 | 1289063 | + | 3248,837 | -0,110438059 | 0,266221 | -0,41484 | 0,678261 | 0,75766 | 2839,240844 | 0,321813372 | 0,135096 | 2,382111 | 0,017214 | 0,031574 | 3658,434 | -0,44753373 | 0,392386 | -1,14055 | 0,254059 | 0,35496 | 3556,371 | 0,691866 | 0,350077 | 1,976327 | 0,048118 | 0,082483 | 3053,558795 | -0,04409694 | 0,216444 | -0,20373 | 0,838562 | 0,869784 |
| 1131 | RM25_RS05500 | RM25_1115 | Hydrolase, NUDIX family | 1189 | gene | 1289099 | 1289884 | + | 991,8038 | 0,30211229 | 0,189817 | 1,591594 | 0,111476 | 0,20622 | 1112,202276 | 0,449161374 | 0,155034 | 2,89718 | 0,003765 | 0,008575 | 871,4054 | 0,119411675 | 0,278037 | 0,429482 | 0,667573 | 0,747846 | 940,4008 | -0,22634 | 0,248373 | -0,91128 | 0,362149 | 0,437512 | 1072,097384 | -0,515206793 | 0,179196 | -2,87509 | 0,004039 | 0,007574 |
| 1132 | RM25_RS05505 | RM25_1116 | Phosphoglucomutase | 1190 | gene | 1290000 | 1291631 | - | 5329,207 | -0,111065352 | 0,472677 | -0,23497 | 0,814231 | 0,870772 | 3059,399824 | 0,630349299 | 0,155093 | 4,064336 | 4,82E-05 | 0,000178 | 7599,014 | -0,407844392 | 0,453727 | -0,89888 | 0,368718 | 0,474647 | 5809,584 | 1,805136 | 0,386676 | 4,668342 | 3,04E-06 | 2,1E-05 | 4994,067632 | 0,794070545 | 0,276395 | 2,872959 | 0,004066 | 0,007607 |
| 1133 | RM25_RS05510 | RM25_1117 | Thioredoxin TrxA | 1191 | gene | 1291763 | 1292755 | + | 876,5423 | 0,451201187 | 0,116219 | 3,882337 | 0,000103 | 0,001767 | 865,4909354 | 0,536454299 | 0,176133 | 3,04574 | 0,002321 | 0,005594 | 887,5938 | 0,370319635 | 0,194553 | 1,903436 | 0,056984 | 0,111092 | 780,7319 | 0,071903 | 0,164859 | 0,436149 | 0,662729 | 0,716023 | 991,584364 | -0,044506662 | 0,176188 | -0,25261 | 0,800571 | 0,838642 |
| 1134 | RM25_RS05515 | RM25_1118 | Methylmalonyl-CoA epimerase | 1192 | gene | 1292987 | 1293433 | + | 27140,59 | 0,11733277 | 0,464893 | 0,252386 | 0,800742 | 0,859841 | 37621,07021 | 0,370341633 | 0,23583 | 1,570375 | 0,116328 | 0,162721 | 16660,1 | -0,453233668 | 0,510105 | -0,88851 | 0,374266 | 0,48021 | 27687,63 | -0,81404 | 0,44593 | -1,82548 | 0,067929 | 0,109363 | 27658,23239 | -1,61543816 | 0,330809 | -4,8833 | 1,04E-06 | 3,63E-06 |
| 1135 | RM25_RS05520 | RM25_1119 | Phospholipid/glycerol acyltransferase | 1193 | gene | 1293589 | 1294164 | + | 379,6942 | -0,536868883 | 0,134998 | -3,97688 | 6,98E-05 | 0,001272 | 342,8442806 | -0,344966008 | 0,170826 | -2,01939 | 0,043446 | 0,07072 | 416,5442 | -0,697150497 | 0,167045 | -4,17344 | 3E-05 | 0,000264 | 473,005 | 0,365454 | 0,155752 | 2,346382 | 0,018957 | 0,039297 | 303,9444584 | 0,066050301 | 0,171104 | 0,386024 | 0,699479 | 0,748376 |
| 1136 | RM25_RS12740 | RM25_1120 | Hypothetical protein | 1194 | gene | 1294323 | 1296116 | + | 6500,143 | 0,716143513 | 0,223131 | 3,209516 | 0,00133 | 0,01196 | 6981,83013 | 0,536630508 | 0,185682 | 2,890051 | 0,003852 | 0,008713 | 6088,456 | 0,927029743 | 0,354074 | 2,618178 | 0,00884 | 0,027849 | 5216,495 | -0,47558 | 0,317779 | -1,49658 | 0,134503 | 0,189494 | 7898,008558 | -0,053217514 | 0,225731 | -0,23576 | 0,813622 | 0,848428 |
| 1137 | RM25_RS05530 | RM25_1121 | Endonuclease NucS | 1195 | gene | 1296157 | 1296804 | - | 893,0031 | 0,939448684 | 0,237275 | 3,959319 | 7,52E-05 | 0,001352 | 1096,094626 | 1,106627237 | 0,155934 | 7,096748 | 1,28E-12 | 1,67E-11 | 689,9116 | 0,68654831 | 0,231733 | 2,962665 | 0,00305 | 0,012163 | 647,5912 | -0,45591 | 0,175532 | -2,5973 | 0,009396 | 0,02256 | 1150,269177 | -0,827084572 | 0,176176 | -4,69465 | 2,67E-06 | 8,61E-06 |
| 1138 | RM25_RS05535 | RM25_1122 | Hypothetical protein | 1196 | gene | 1296961 | 1297239 | + | 874,9884 | 0,736857287 | 0,186847 | 3,943636 | 8,03E-05 | 0,001422 | 803,6925898 | 0,801556133 | 0,142177 | 5,637719 | 1,72E-08 | 1,19E-07 | 946,2842 | 0,684312519 | 0,32152 | 2,128367 | 0,033307 | 0,074766 | 692,7778 | 0,253177 | 0,230071 | 1,10043 | 0,271145 | 0,343966 | 1067,794756 | 0,173645609 | 0,239334 | 0,725536 | 0,468124 | 0,531231 |
| 1139 | RM25_RS05540 | RM25_1123 | Esterase YbfF | 1197 | gene | 1297236 | 1298066 | + | 635,4466 | -0,4164696 | 0,281231 | -1,48088 | 0,138639 | 0,238346 | 477,7023086 | -0,409844062 | 0,240137 | -1,70671 | 0,087876 | 0,126526 | 793,1909 | -0,422259056 | 0,251805 | -1,67693 | 0,093557 | 0,163895 | 761,9754 | 0,670394 | 0,287247 | 2,333855 | 0,019603 | 0,04025 | 533,6785374 | 0,717339299 | 0,195001 | 3,678637 | 0,000234 | 0,000558 |
| 1140 | RM25_RS05545 | RM25_1124 | ATP:cob(I)alamin adenosyltransferase | 1198 | gene | 1298145 | 1298753 | + | 2075,201 | 0,199012994 | 0,267655 | 0,743543 | 0,457153 | 0,561788 | 2045,604907 | 0,750132204 | 0,167082 | 4,489617 | 7,14E-06 | 3,06E-05 | 2104,796 | -0,321156857 | 0,378079 | -0,84944 | 0,395634 | 0,501619 | 2037,482 | 0,565092 | 0,329651 | 1,714212 | 0,08649 | 0,131775 | 2169,453878 | -0,472263538 | 0,236736 | -1,99489 | 0,046054 | 0,068616 |
| 1141 | RM25_RS05550 | RM25_1125 | Lipoate--protein ligase | 1199 | gene | 1298928 | 1299992 | - | 1617,757 | 0,640474571 | 0,358874 | 1,784679 | 0,074313 | 0,160537 | 2227,056968 | 0,710704603 | 0,157471 | 4,513237 | 6,38E-06 | 2,78E-05 | 1008,456 | 0,488547359 | 0,217958 | 2,241474 | 0,024995 | 0,060862 | 1341,85 | -1,06656 | 0,169739 | -6,28353 | 3,31E-10 | 5,31E-09 | 1932,459782 | -1,244422917 | 0,159285 | -7,81253 | 5,6E-15 | 6,77E-14 |
| 1142 | RM25_RS05555 | RM25_1126 | putative secreted protein | 1200 | gene | 1300016 | 1300474 | - | 856,2273 | 0,953653234 | 0,592465 | 1,609638 | 0,107477 | 0,201419 | 1365,810791 | 0,865046489 | 0,20252 | 4,271404 | 1,94E-05 | 7,76E-05 | 346,6439 | 1,330978461 | 0,333097 | 3,995765 | 6,45E-05 | 0,000513 | 623,0575 | -2,35173 | 0,293384 | -8,01588 | 1,09E-15 | 4,96E-14 | 1107,27378 | -1,847825997 | 0,226461 | -8,15958 | 3,36E-16 | 4,86E-15 |
| 1143 | RM25_RS05560 | RM25_1127 | ATP synthase F1, epsilon subunit | 1201 | gene | 1300516 | 1300983 | - | 1558,494 | 1,110223968 | 0,579568 | 1,915605 | 0,055415 | 0,132603 | 2345,756818 | 0,88818839 | 0,215306 | 4,12523 | 3,7E-05 | 0,00014 | 771,2305 | 1,894965465 | 0,47883 | 3,957493 | 7,57E-05 | 0,000585 | 1055,206 | -2,3796 | 0,469235 | -5,07124 | 3,95E-07 | 3,26E-06 | 2087,745682 | -1,340720719 | 0,227439 | -5,89486 | 3,75E-09 | 1,89E-08 |
| 1144 | RM25_RS05565 | RM25_1128 | ATP synthase subunit beta | 1202 | gene | 1300983 | 1302434 | - | 6361,122 | 1,041078106 | 0,590466 | 1,763146 | 0,077876 | 0,165215 | 9391,557263 | 0,748693109 | 0,201427 | 3,71694 | 0,000202 | 0,000644 | 3330,686 | 2,027000652 | 0,542043 | 3,73956 | 0,000184 | 0,001234 | 4448,319 | -2,46545 | 0,513351 | -4,80267 | 1,57E-06 | 1,16E-05 | 8383,504661 | -1,157088291 | 0,255727 | -4,52471 | 6,05E-06 | 1,83E-05 |
| 1145 | RM25_RS05570 | RM25_1129 | ATP synthase gamma chain | 1203 | gene | 1302435 | 1303370 | - | 2486,105 | 0,980980599 | 0,506412 | 1,937118 | 0,052731 | 0,128661 | 3597,40059 | 0,694896617 | 0,194319 | 3,576064 | 0,000349 | 0,001059 | 1374,809 | 1,85301564 | 0,278232 | 6,659961 | 2,74E-11 | 1,02E-09 | 1785,715 | -2,26177 | 0,22689 | -9,96856 | 2,09E-23 | 2,65E-21 | 3233,278234 | -1,061485914 | 0,217073 | -4,88999 | 1,01E-06 | 3,52E-06 |
| 1146 | RM25_RS05575 | RM25_1130 | ATP synthase subunit alpha | 1204 | gene | 1303373 | 1305010 | - | 5676,706 | 0,911287855 | 0,541586 | 1,682627 | 0,092447 | 0,183704 | 8513,628307 | 0,666128068 | 0,195166 | 3,413131 | 0,000642 | 0,001813 | 2839,784 | 1,753078084 | 0,292288 | 5,997772 | 2E-09 | 5,24E-08 | 4209,323 | -2,39867 | 0,247914 | -9,6754 | 3,84E-22 | 4,11E-20 | 7263,250968 | -1,269662865 | 0,210407 | -6,03431 | 1,6E-09 | 8,67E-09 |
| 1147 | RM25_RS05580 | RM25_1131 | ATP synthase F1, delta subunit | 1205 | gene | 1305085 | 1305864 | - | 3513,764 | 1,02508969 | 0,576822 | 1,777133 | 0,075546 | 0,162527 | 5323,631428 | 0,749181407 | 0,192895 | 3,88388 | 0,000103 | 0,000349 | 1703,897 | 2,061793668 | 0,308368 | 6,68614 | 2,29E-11 | 8,85E-10 | 2478,181 | -2,64961 | 0,257422 | -10,2929 | 7,59E-25 | 1,19E-22 | 4616,87589 | -1,296651511 | 0,225112 | -5,76004 | 8,41E-09 | 4,05E-08 |
| 1148 | RM25_RS05585 | RM25_1132 | ATP synthase subunit b | 1206 | gene | 1305861 | 1306415 | - | 2451,17 | 1,157006889 | 0,611267 | 1,892802 | 0,058384 | 0,136791 | 3673,918927 | 0,807770246 | 0,204751 | 3,945131 | 7,98E-05 | 0,000278 | 1228,421 | 2,527344507 | 0,359061 | 7,038762 | 1,94E-12 | 1,02E-10 | 1626,727 | -2,93287 | 0,320414 | -9,15339 | 5,52E-20 | 4,2E-18 | 3315,808204 | -1,175686965 | 0,243395 | -4,83037 | 1,36E-06 | 4,61E-06 |
| 1149 | RM25_RS05590 | RM25_1133 | ATP synthase C chain (F1F0-ATPase subunit c) | 1207 | gene | 1306426 | 1306644 | - | 1432,231 | 1,001877525 | 0,706077 | 1,418934 | 0,155918 | 0,257352 | 2207,492465 | 0,61997183 | 0,232287 | 2,668993 | 0,007608 | 0,015702 | 656,9691 | 2,782066224 | 0,540637 | 5,145909 | 2,66E-07 | 4,33E-06 | 1023,86 | -3,431 | 0,50344 | -6,81511 | 9,42E-12 | 2,09E-10 | 1870,293797 | -1,241594729 | 0,301002 | -4,12487 | 3,71E-05 | 0,000101 |
| 1150 | RM25_RS05595 | RM25_1134 | ATP synthase subunit a | 1208 | gene | 1306690 | 1307571 | - | 3497,598 | 0,635452852 | 0,437854 | 1,451288 | 0,1467 | 0,247333 | 4987,900521 | 0,512717216 | 0,217748 | 2,35464 | 0,018541 | 0,03359 | 2007,295 | 0,951984137 | 0,290869 | 3,272896 | 0,001065 | 0,00511 | 2922,955 | -1,64834 | 0,236625 | -6,96605 | 3,26E-12 | 7,84E-11 | 4167,740038 | -1,165423213 | 0,247171 | -4,71505 | 2,42E-06 | 7,88E-06 |
| 1151 | RM25_RS05600 | RM25_1135 | Hypothetical protein | 1209 | gene | 1307564 | 1307788 | - | 241,7946 | 0,706225953 | 0,379676 | 1,860077 | 0,062875 | 0,142908 | 328,8678374 | 0,643866433 | 0,230093 | 2,798288 | 0,005137 | 0,011212 | 154,7213 | 0,855052267 | 0,3016 | 2,835049 | 0,004582 | 0,016692 | 195,6381 | -1,27381 | 0,309042 | -4,12182 | 3,76E-05 | 0,000201 | 293,591037 | -1,0257825 | 0,209925 | -4,88643 | 1,03E-06 | 3,58E-06 |
| 1152 | RM25_RS05605 | RM25_1136 | Hypothetical protein | 1210 | gene | 1307785 | 1308222 | - | 1187,372 | 0,605245902 | 0,527309 | 1,147801 | 0,251051 | 0,359057 | 1824,173047 | 0,636085285 | 0,251112 | 2,533072 | 0,011307 | 0,021952 | 550,5706 | 0,508087316 | 0,360667 | 1,408743 | 0,158911 | 0,246417 | 1005,527 | -1,70365 | 0,351583 | -4,84567 | 1,26E-06 | 9,44E-06 | 1404,190247 | -1,798696758 | 0,25794 | -6,97332 | 3,1E-12 | 2,45E-11 |
| 1153 | RM25_RS05610 | RM25_1137 | Undecaprenyl-phosphate alpha-N-acetylglucosaminyltransfe | 1211 | gene | 1308300 | 1309436 | - | 822,7766 | -0,126156882 | 0,109528 | -1,15183 | 0,249393 | 0,35762 | 842,6105457 | -0,011594617 | 0,152152 | -0,0762 | 0,939257 | 0,95228 | 802,9426 | -0,243772131 | 0,196555 | -1,24022 | 0,214893 | 0,312446 | 905,0543 | -0,02135 | 0,191233 | -0,11165 | 0,911102 | 0,929713 | 772,2785722 | -0,198888294 | 0,133798 | -1,48648 | 0,137151 | 0,182946 |
| 1154 | RM25_RS05615 | RM25_1138 | Sua5/YciO/YrdC/YwlC family protein | 1212 | gene | 1309445 | 1310323 | - | 935,5747 | 0,397365439 | 0,207228 | 1,917531 | 0,05517 | 0,132317 | 1111,60735 | 0,507798394 | 0,155765 | 3,260034 | 0,001114 | 0,00296 | 759,542 | 0,240500682 | 0,199565 | 1,205123 | 0,228156 | 0,32572 | 856,1747 | -0,45572 | 0,202972 | -2,24523 | 0,024753 | 0,048418 | 1042,235769 | -0,677837391 | 0,138245 | -4,90315 | 9,43E-07 | 3,31E-06 |
| 1155 | RM25_RS05620 | RM25_1139 | Release factor glutamine methyltransferase | 1213 | gene | 1310320 | 1311198 | - | 349,2999 | -0,117407339 | 0,149289 | -0,78644 | 0,431607 | 0,539681 | 376,004875 | -0,18079463 | 0,176182 | -1,02618 | 0,304806 | 0,373598 | 322,5949 | -0,042630708 | 0,247299 | -0,17239 | 0,863134 | 0,90018 | 384,1289 | -0,34789 | 0,200391 | -1,73606 | 0,082554 | 0,126515 | 328,6573445 | -0,15441609 | 0,211871 | -0,72882 | 0,466111 | 0,529912 |
| 1156 | RM25_RS05625 | RM25_1140 | Peptide chain release factor 1 | 1214 | gene | 1311195 | 1312274 | - | 770,9804 | -0,508312307 | 0,236922 | -2,14548 | 0,031914 | 0,094028 | 935,4378303 | -0,650232032 | 0,194566 | -3,34197 | 0,000832 | 0,002275 | 606,523 | -0,291628173 | 0,210801 | -1,38343 | 0,166534 | 0,25616 | 961,7407 | -0,83628 | 0,233106 | -3,58755 | 0,000334 | 0,00141 | 624,8306373 | -0,419202147 | 0,184361 | -2,27382 | 0,022977 | 0,03717 |
| 1157 | RM25_RS05630 | RM25_1141 | 50S ribosomal protein L31 | 1215 | gene | 1312382 | 1312606 | - | 5913,285 | 0,298547148 | 0,406021 | 0,7353 | 0,462157 | 0,565419 | 7357,047048 | -0,086186659 | 0,215714 | -0,39954 | 0,689494 | 0,741733 | 4469,522 | 0,963596967 | 0,509569 | 1,891004 | 0,058624 | 0,113913 | 5665,144 | -1,36777 | 0,513038 | -2,66602 | 0,007675 | 0,018963 | 6383,341087 | -0,287068792 | 0,212706 | -1,3496 | 0,177143 | 0,22893 |
| 1158 | RM25_RS05635 | RM25_1142 | Transcription termination factor Rho | 1216 | gene | 1312743 | 1314713 | - | 9470,307 | -0,112486135 | 0,361465 | -0,31119 | 0,755653 | 0,823216 | 10693,722 | -0,737887884 | 0,278511 | -2,6494 | 0,008063 | 0,016512 | 8246,892 | 0,691212211 | 0,423144 | 1,633516 | 0,102361 | 0,17521 | 10498,71 | -1,13477 | 0,423591 | -2,67892 | 0,007386 | 0,018383 | 8876,181607 | 0,324711231 | 0,283693 | 1,144586 | 0,252381 | 0,31047 |
| 1159 | RM25_RS05640 | RM25_1143 | Phosphoserine phosphatase/homoserine phosphotransferas | 1217 | gene | 1315107 | 1315730 | - | 2403,596 | 0,487861857 | 0,359396 | 1,357449 | 0,174639 | 0,278312 | 3205,623043 | 0,378788716 | 0,184274 | 2,055577 | 0,039823 | 0,065593 | 1601,57 | 0,712266056 | 0,316901 | 2,247598 | 0,024602 | 0,060277 | 2133,098 | -1,2518 | 0,294175 | -4,25529 | 2,09E-05 | 0,000119 | 2747,442028 | -0,881894423 | 0,206271 | -4,27542 | 1,91E-05 | 5,4E-05 |
| 1160 | RM25_RS05645 | RM25_1144 | Homoserine dehydrogenase | 1218 | gene | 1315739 | 1317064 | - | 2391,691 | 0,653073202 | 0,399048 | 1,636577 | 0,101719 | 0,195078 | 3331,428747 | 0,557444784 | 0,196923 | 2,830777 | 0,004644 | 0,010186 | 1451,954 | 0,88004375 | 0,262247 | 3,355782 | 0,000791 | 0,004029 | 1979,981 | -1,45153 | 0,249365 | -5,8209 | 5,85E-09 | 6,73E-08 | 2864,443309 | -1,091693017 | 0,19154 | -5,69955 | 1,2E-08 | 5,63E-08 |
| 1161 | RM25_RS05650 | RM25_1145 | Diaminopimelate decarboxylase | 1219 | gene | 1317075 | 1318469 | - | 3639,553 | 0,812166493 | 0,52136 | 1,557784 | 0,119285 | 0,215443 | 5479,881518 | 0,634122372 | 0,20532 | 3,088461 | 0,002012 | 0,004945 | 1799,224 | 1,403150609 | 0,277923 | 5,0487 | 4,45E-07 | 6,85E-06 | 2823,766 | -2,17698 | 0,261921 | -8,31161 | 9,44E-17 | 5,06E-15 | 4544,244606 | -1,368735797 | 0,203777 | -6,71684 | 1,86E-11 | 1,33E-10 |
| 1163 | RM25_RS05660 | RM25_1147 | Brp/Blh family beta-carotene 15, 15'-monooxygenase | 1223 | gene | 1319734 | 1321152 | + | 2040,793 | 0,242336049 | 0,279762 | 0,866221 | 0,386369 | 0,497619 | 2576,494906 | 0,150542178 | 0,168185 | 0,895099 | 0,370734 | 0,439547 | 1505,091 | 0,401235607 | 0,23094 | 1,737402 | 0,082316 | 0,148255 | 1985,471 | -0,9739 | 0,199242 | -4,888 | 1,02E-06 | 7,81E-06 | 2166,050971 | -0,675613312 | 0,166553 | -4,05646 | 4,98E-05 | 0,000132 |
| 1164 | RM25_RS05665 | RM25_1148 | Brp/Blh family beta-carotene 15, 15'-monooxygenase | 1224 | gene | 1321293 | 1322588 | + | 672,0329 | -0,323044193 | 0,1686 | -1,91604 | 0,05536 | 0,132603 | 674,1298134 | -0,244313676 | 0,14243 | -1,71533 | 0,086285 | 0,124693 | 669,9361 | -0,401716362 | 0,321155 | -1,25085 | 0,210989 | 0,308117 | 787,114 | -0,00248 | 0,212042 | -0,0117 | 0,990664 | 0,991926 | 587,1422012 | -0,095113901 | 0,272652 | -0,34885 | 0,727204 | 0,773126 |
| 1165 | RM25_RS05670 | RM25_1149 | Hypothetical protein | 1225 | gene | 1322581 | 1323987 | + | 738,9426 | -0,602980706 | 0,18588 | -3,24392 | 0,001179 | 0,01116 | 755,5256005 | -0,495936648 | 0,134339 | -3,69169 | 0,000223 | 0,000705 | 722,3596 | -0,716773578 | 0,345562 | -2,07423 | 0,038058 | 0,083213 | 939,3893 | -0,04223 | 0,2362 | -0,1788 | 0,858094 | 0,888243 | 577,1394989 | -0,196967317 | 0,284253 | -0,69293 | 0,488354 | 0,549953 |
| 1166 | RM25_RS05675 | RM25_1150 | 2-oxoglutarate decarboxylase | 1226 | gene | 1324057 | 1327842 | + | 4113,677 | 0,415586509 | 0,250383 | 1,659805 | 0,096954 | 0,189569 | 4503,863265 | 1,022611751 | 0,115573 | 8,8482 | 8,89E-19 | 2,28E-17 | 3723,491 | -0,275721926 | 0,267035 | -1,03253 | 0,301823 | 0,406194 | 3716,878 | 0,403323 | 0,209084 | 1,929002 | 0,053731 | 0,090009 | 4605,539219 | -0,856159771 | 0,177799 | -4,81533 | 1,47E-06 | 4,95E-06 |
| 1167 | RM25_RS05680 | RM25_1151 | Transcription factor WhiB | 1227 | gene | 1328087 | 1328335 | + | 922,1007 | 0,794676598 | 0,34662 | 2,292643 | 0,021869 | 0,075137 | 1065,603606 | 0,729471742 | 0,266514 | 2,737082 | 0,006199 | 0,01321 | 778,5978 | 0,887177883 | 0,512414 | 1,731369 | 0,083386 | 0,149802 | 717,4473 | -0,60174 | 0,512817 | -1,1734 | 0,240637 | 0,309282 | 1143,601159 | -0,412337811 | 0,271865 | -1,5167 | 0,129342 | 0,17361 |
| 1168 | RM25_RS05685 | RM25_1152 | Putative sensor histidine kinase pdtaS | 1228 | gene | 1328440 | 1329948 | - | 1348,962 | 0,069667702 | 0,141619 | 0,491938 | 0,622763 | 0,709451 | 1459,961795 | 0,282850377 | 0,145849 | 1,939337 | 0,05246 | 0,082268 | 1237,962 | -0,178748444 | 0,170276 | -1,04976 | 0,29383 | 0,397121 | 1389,145 | -0,05896 | 0,145231 | -0,40598 | 0,684759 | 0,733293 | 1353,700011 | -0,475859543 | 0,131273 | -3,62495 | 0,000289 | 0,000679 |
| 1169 | RM25_RS05690 | RM25_1153 | Anti-sigma factor | 1229 | gene | 1330434 | 1330745 | - | 3085,039 | -0,915358723 | 0,418544 | -2,18701 | 0,028742 | 0,087076 | 4035,746072 | -1,025378074 | 0,283853 | -3,61235 | 0,000303 | 0,000933 | 2134,333 | -0,713346782 | 0,530146 | -1,34557 | 0,178442 | 0,270649 | 4298,134 | -1,07329 | 0,554523 | -1,93551 | 0,052928 | 0,089025 | 2094,206929 | -0,731218148 | 0,246782 | -2,96302 | 0,003046 | 0,005866 |
| 1170 | RM25_RS05695 | RM25_1154 | RNA polymerase sigma factor | 1230 | gene | 1330738 | 1331325 | - | 10140,47 | -1,347755915 | 0,308065 | -4,37491 | 1,21E-05 | 0,000345 | 10456,89299 | -1,207140008 | 0,268145 | -4,50182 | 6,74E-06 | 2,91E-05 | 9824,044 | -1,504180088 | 0,479671 | -3,13586 | 0,001714 | 0,007563 | 15457,85 | -0,05405 | 0,496512 | -0,10886 | 0,913313 | 0,93109 | 5601,94755 | -0,316129475 | 0,256213 | -1,23385 | 0,217257 | 0,273253 |
| 1171 | RM25_RS05700 | RM25_1155 | UPF0361 protein yqgW | 1231 | gene | 1331466 | 1332218 | - | 404,5371 | -0,338450141 | 0,164248 | -2,06061 | 0,03934 | 0,10757 | 370,3203979 | -0,252884721 | 0,184921 | -1,36753 | 0,17146 | 0,227937 | 438,7538 | -0,41214766 | 0,262016 | -1,57299 | 0,115722 | 0,19317 | 475,7724 | 0,248106 | 0,235265 | 1,054579 | 0,291618 | 0,365733 | 350,8537242 | 0,150264495 | 0,221572 | 0,678174 | 0,497662 | 0,559231 |
| 1172 | RM25_RS05705 | RM25_1156 | 3-phosphoshikimate 1-carboxyvinyltransferase 2 | 1232 | gene | 1332366 | 1333718 | + | 1033,294 | -0,225557441 | 0,197505 | -1,14203 | 0,25344 | 0,361816 | 931,2038132 | 0,233123219 | 0,172658 | 1,350204 | 0,176951 | 0,233916 | 1135,385 | -0,606669782 | 0,203097 | -2,9871 | 0,002816 | 0,011497 | 1170,354 | 0,626254 | 0,196129 | 3,193077 | 0,001408 | 0,004693 | 935,1988357 | -0,16416295 | 0,176504 | -0,93008 | 0,352328 | 0,416051 |
| 1173 | RM25_RS05710 | RM25_1157 | Putative ribosome biogenesis GTPase RsgA | 1233 | gene | 1333715 | 1334785 | + | 931,4417 | -0,35172412 | 0,147338 | -2,3872 | 0,016977 | 0,065132 | 917,5941913 | -0,047983518 | 0,161678 | -0,29679 | 0,76663 | 0,811766 | 945,2892 | -0,652086319 | 0,200531 | -3,25179 | 0,001147 | 0,005428 | 1098,639 | 0,24983 | 0,188675 | 1,324127 | 0,185461 | 0,247106 | 803,3872975 | -0,30195837 | 0,155762 | -1,93859 | 0,052551 | 0,077126 |
| 1174 | RM25_RS05715 | RM25_1158 | Histidinol-phosphate phosphatase HisN | 1234 | gene | 1334839 | 1335654 | + | 678,3391 | -0,005487206 | 0,181333 | -0,03026 | 0,975859 | 0,984632 | 778,1438561 | 0,139548661 | 0,169187 | 0,824819 | 0,409474 | 0,478244 | 578,5343 | -0,196660614 | 0,174983 | -1,12388 | 0,261062 | 0,362381 | 719,9579 | -0,32334 | 0,200412 | -1,6134 | 0,106658 | 0,156533 | 664,5833529 | -0,601238081 | 0,159876 | -3,76065 | 0,000169 | 0,000415 |
| 1175 | RM25_RS05720 | RM25_1159 | PF14262 domain protein | 1235 | gene | 1335708 | 1337441 | - | 473,1565 | -0,955691596 | 0,482083 | -1,98242 | 0,047432 | 0,120602 | 286,5425686 | -1,635117172 | 0,323412 | -5,05583 | 4,29E-07 | 2,28E-06 | 659,7705 | -0,694788032 | 0,376338 | -1,84618 | 0,064866 | 0,123397 | 654,9202 | 0,843158 | 0,350949 | 2,402507 | 0,016283 | 0,035018 | 317,1037719 | 1,854728865 | 0,373202 | 4,969769 | 6,7E-07 | 2,42E-06 |
| 1176 | RM25_RS05725 | RM25_1160 | VTC domain protein | 1236 | gene | 1337438 | 1338067 | - | 172,2072 | -1,10966243 | 0,67389 | -1,64665 | 0,09963 | 0,192532 | 92,11925469 | -2,903303864 | 0,501174 | -5,793 | 6,91E-09 | 5,17E-08 | 252,2951 | -0,651588301 | 0,404055 | -1,61262 | 0,106827 | 0,181144 | 246,8154 | 0,849661 | 0,450647 | 1,885426 | 0,059372 | 0,097724 | 107,4671649 | 3,177173971 | 0,493156 | 6,442531 | 1,17E-10 | 7,57E-10 |
| 1177 | RM25_RS05730 | RM25_1161 | Hypothetical protein | 1237 | gene | 1338246 | 1338881 | - | 197,6391 | -0,984893201 | 0,560986 | -1,75565 | 0,079149 | 0,166417 | 109,4659796 | -2,019676641 | 0,440012 | -4,59005 | 4,43E-06 | 1,99E-05 | 285,8121 | -0,658519433 | 0,332201 | -1,98229 | 0,047446 | 0,097244 | 275,5595 | 0,918281 | 0,442908 | 2,073298 | 0,038145 | 0,06837 | 130,2930386 | 2,352393789 | 0,361643 | 6,504746 | 7,78E-11 | 5,17E-10 |
| 1178 | RM25_RS05735 | RM25_1162 | Exopolysaccharide biosynthesis protein related to N-acetylgl | 1238 | gene | 1339038 | 1339985 | + | 243,0899 | -1,200284766 | 0,577649 | -2,07788 | 0,037721 | 0,105216 | 135,798238 | -2,464552744 | 0,383656 | -6,42386 | 1,33E-10 | 1,28E-09 | 350,3815 | -0,825716837 | 0,334047 | -2,47186 | 0,013441 | 0,038355 | 355,1004 | 0,891375 | 0,376402 | 2,368146 | 0,017877 | 0,037422 | 144,7763555 | 2,598259599 | 0,368302 | 7,054695 | 1,73E-12 | 1,44E-11 |
| 1179 | RM25_RS11875 | RM25_1163 | Glycosyltransferase, family 2 | 1239 | gene | 1339995 | 1340543 | + | 64,89825 | -1,148990924 | 0,826819 | -1,38965 | 0,164635 | 0,267091 | 26,69004367 | -3,855571878 | 0,689842 | -5,58906 | 2,28E-08 | 1,56E-07 | 103,1064 | -0,742882568 | 0,453361 | -1,63861 | 0,101294 | 0,174505 | 93,70742 | 1,284999 | 0,338286 | 3,798558 | 0,000146 | 0,000681 | 39,72609343 | 4,482479646 | 0,770339 | 5,818842 | 5,93E-09 | 2,92E-08 |

| | | | | | | | | | | | | | | | | | | | | | | | | | | | | | | | | | | | | | | |
|---|---|---|---|---|---|---|---|---|---|---|---|---|---|---|---|---|---|---|---|---|---|---|---|---|---|---|---|---|---|---|---|---|---|---|---|---|---|---|
| 1180 | RM25_RS11880 | RM25_1164 | Dolichyl-phosphate mannose synthase | 1240 | gene | 1340497 | 1341153 | + | 207,2085 | -1,547250248 | 0,515457 | -3,00171 | 0,002685 | 0,019175 | 127,7031294 | -2,438905351 | 0,415127 | -5,87508 | 4,23E-09 | 3,29E-08 | 286,7139 | -1,23188186 | 0,319537 | -3,85521 | 0,000116 | 0,000833 | 324,3063 | 0,824658 | 0,411645 | 2,003324 | 0,045143 | 0,078409 | 103,7824277 | 2,104596913 | 0,376861 | 5,584546 | 2,34E-08 | 1,06E-07 |
| 1181 | RM25_RS05745 | RM25_1165 | Phosphoglycerate dehydrogenase | 1241 | gene | 1341143 | 1342363 | - | 2538,134 | 0,585461138 | 0,307406 | 1,904522 | 0,056842 | 0,134515 | 3235,702099 | 0,421802435 | 0,16404 | 2,571332 | 0,010131 | 0,020083 | 1840,565 | 0,8839011 | 0,253751 | 3,483341 | 0,000495 | 0,002779 | 2161,653 | -1,15679 | 0,210434 | -5,49716 | 3,86E-08 | 3,87E-07 | 2981,345188 | -0,650351497 | 0,185956 | -3,49733 | 0,00047 | 0,00106 |
| 1182 | RM25_RS05750 | RM25_1166 | Oligopeptide/dipeptide ABC transporter, ATP-binding prote | 1242 | gene | 1342397 | 1343224 | - | 627,4552 | -0,091599202 | 0,181423 | -0,50489 | 0,613634 | 0,701763 | 703,8081267 | 0,088186832 | 0,143422 | 0,614874 | 0,538638 | 0,604845 | 551,1022 | -0,319056792 | 0,245825 | -1,2979 | 0,194321 | 0,289171 | 682,6465 | -0,21909 | 0,215424 | -1,01703 | 0,309138 | 0,384101 | 596,4104333 | -0,569367738 | 0,154388 | -3,6879 | 0,000226 | 0,000541 |
| 1183 | RM25_RS05755 | RM25_1167 | Oligopeptide transport ATP-binding protein OppD | 1243 | gene | 1343217 | 1344074 | - | 498,8995 | 0,541902659 | 0,285536 | 1,897843 | 0,057717 | 0,135555 | 635,4389514 | 0,619463415 | 0,168121 | 3,684619 | 0,000229 | 0,000724 | 362,3601 | 0,411773263 | 0,266471 | 1,545286 | 0,122277 | 0,201262 | 430,0783 | -0,7488 | 0,233618 | -3,20525 | 0,001349 | 0,00457 | 580,9127192 | -0,896223696 | 0,191236 | -4,68648 | 2,78E-06 | 8,95E-06 |
| 1184 | RM25_RS05760 | RM25_1168 | Putative oligopeptide ABC transport system membrane prot | 1244 | gene | 1344076 | 1344891 | - | 727,1412 | 0,0356035 | 0,316839 | 0,112371 | 0,910529 | 0,941556 | 943,691203 | 0,241531552 | 0,146358 | 1,650282 | 0,098885 | 0,140829 | 510,5912 | -0,341346951 | 0,266556 | -1,28058 | 0,20034 | 0,297356 | 759,8905 | -0,66077 | 0,217746 | -3,0346 | 0,002409 | 0,007382 | 723,130812 | -1,184072797 | 0,192099 | -6,16387 | 7,1E-10 | 4,04E-09 |
| 1185 | RM25_RS05765 | RM25_1169 | Putative glutathione ABC transporter, permease protein Gsi | 1245 | gene | 1344888 | 1345841 | - | 1163,082 | 0,466401841 | 0,47458 | 0,982768 | 0,325722 | 0,435713 | 1747,092254 | 0,452438638 | 0,18178 | 2,48893 | 0,012813 | 0,024553 | 579,0713 | 0,510995921 | 0,21284 | 2,400844 | 0,016357 | 0,044112 | 1040,552 | -1,6878 | 0,22958 | -7,35168 | 1,96E-13 | 5,43E-12 | 1324,021875 | -1,578498013 | 0,122876 | -12,8462 | 9,03E-38 | 1,01E-35 |
| 1186 | RM25_RS05770 | RM25_1170 | ABC-type dipeptide transport system, periplasmic compone | 1246 | gene | 1345863 | 1347386 | - | 4795,083 | 0,687369513 | 0,353361 | 1,945233 | 0,051747 | 0,126917 | 6087,695455 | 0,355656455 | 0,183969 | 1,933243 | 0,053206 | 0,083217 | 3502,47 | 1,317021842 | 0,291782 | 4,513724 | 6,37E-06 | 6,82E-05 | 3914,63 | -1,46827 | 0,215243 | -6,82143 | 9,01E-12 | 2,02E-10 | 5786,251707 | -0,468708568 | 0,233908 | -2,00381 | 0,04509 | 0,067489 |
| 1188 | RM25_RS05780 | RM25_1172 | Biotin synthase | 1250 | gene | 1347771 | 1348916 | - | 586,9777 | -0,666749084 | 0,278534 | -2,39378 | 0,016676 | 0,064646 | 488,2494888 | -1,323997698 | 0,158574 | -8,34939 | 6,86E-17 | 1,43E-15 | 685,706 | -0,243573683 | 0,208104 | -1,17044 | 0,241824 | 0,341305 | 759,6616 | 0,029893 | 0,174622 | 0,171184 | 0,864079 | 0,892869 | 444,5518531 | 1,16763199 | 0,182574 | 6,395397 | 1,6E-10 | 1,01E-09 |
| 1189 | RM25_RS05785 | RM25_1173 | Acetyltransferase | 1251 | gene | 1349079 | 1349369 | - | 1083,96 | -1,029743903 | 0,409644 | -2,51375 | 0,011945 | 0,052327 | 691,9332713 | -0,814344452 | 0,272818 | -2,98493 | 0,002836 | 0,006665 | 1475,986 | -1,134066724 | 0,438961 | -2,58352 | 0,00978 | 0,029743 | 1535,601 | 1,149776 | 0,48504 | 2,370476 | 0,017765 | 0,03732 | 696,1262889 | 0,867371826 | 0,190321 | 4,557425 | 5,18E-06 | 1,59E-05 |
| 1190 | RM25_RS05790 | RM25_1174 | Prolyl aminopeptidase | 1252 | gene | 1349536 | 1350750 | - | 1658,554 | -0,00030171 | 0,400061 | -0,00075 | 0,999398 | 0,999649 | 1007,716535 | 0,301024066 | 0,127052 | 2,369292 | 0,017822 | 0,032488 | 2309,392 | -0,131008876 | 0,345671 | -0,379 | 0,704689 | 0,779424 | 1742,031 | 1,366893 | 0,30592 | 4,468139 | 7,89E-06 | 4,98E-05 | 1618,620189 | 0,971463296 | 0,198505 | 4,893897 | 9,89E-07 | 3,46E-06 |
| 1191 | RM25_RS05795 | RM25_1175 | Peptidase M16 inactive domain protein | 1253 | gene | 1350895 | 1352169 | + | 1444,277 | 0,489578344 | 0,102367 | 4,782594 | 1,73E-06 | 6,62E-05 | 1512,747708 | 0,586312271 | 0,129502 | 4,527452 | 5,97E-06 | 2,61E-05 | 1375,807 | 0,386240717 | 0,188059 | 2,053824 | 0,039993 | 0,08587 | 1268,823 | -0,07812 | 0,140984 | -0,55411 | 0,579501 | 0,641561 | 1651,915245 | -0,232371216 | 0,136928 | -1,69703 | 0,08969 | 0,125089 |
| 1192 | RM25_RS05800 | RM25_1176 | Peptidase M16 inactive domain protein | 1254 | gene | 1352166 | 1353530 | + | 723,0063 | 0,552727289 | 0,113836 | 4,855469 | 1,2E-06 | 4,8E-05 | 774,4265573 | 0,729273742 | 0,132744 | 5,493849 | 3,93E-08 | 2,58E-07 | 671,586 | 0,357358616 | 0,165025 | 2,165476 | 0,030351 | 0,070067 | 618,8248 | -0,04752 | 0,140163 | -0,33907 | 0,73456 | 0,77954 | 843,9545238 | -0,363274241 | 0,149475 | -2,43034 | 0,015085 | 0,025433 |
| 1193 | RM25_RS05805 | RM25_1177 | UPF0182 protein | 1255 | gene | 1353614 | 1356553 | - | 5696,478 | 0,585276691 | 0,417209 | 1,402839 | 0,160665 | 0,263894 | 7956,314503 | 0,396646344 | 0,178989 | 2,216035 | 0,026689 | 0,046153 | 3436,641 | 1,045575037 | 0,276247 | 3,784929 | 0,000154 | 0,001063 | 4856,627 | -1,67128 | 0,225302 | -7,41794 | 1,19E-13 | 3,42E-12 | 6695,190156 | -0,981768711 | 0,204111 | -4,80998 | 1,51E-06 | 5,05E-06 |
| 1194 | RM25_RS05810 | RM25_1178 | Hypothetical protein | 1256 | gene | 1356550 | 1357149 | - | 1123,244 | 0,657703241 | 0,487335 | 1,349591 | 0,177147 | 0,28098 | 1693,879863 | 0,639932968 | 0,191602 | 3,339913 | 0,000838 | 0,002289 | 552,608 | 0,715065831 | 0,273537 | 2,614147 | 0,008945 | 0,027999 | 929,8885 | -1,72198 | 0,22512 | -7,64918 | 2,02E-14 | 7,01E-13 | 1347,623126 | -1,604054168 | 0,214225 | -7,48769 | 7,01E-14 | 7,25E-13 |
| 1195 | RM25_RS05815 | RM25_1179 | putative secreted protein containing a PDZ domain | 1257 | gene | 1357266 | 1358309 | - | 1802,023 | 0,315817173 | 0,308007 | 1,025358 | 0,305194 | 0,41474 | 2338,143906 | 0,618091421 | 0,132376 | 4,669203 | 3,02E-06 | 1,41E-05 | 1265,902 | -0,228028073 | 0,184187 | -1,23803 | 0,215706 | 0,312874 | 1700,493 | -0,49418 | 0,146007 | -3,38467 | 0,000713 | 0,002649 | 1959,968235 | -1,291270792 | 0,134643 | -9,5903 | 8,78E-22 | 2,35E-20 |
| 1196 | RM25_RS05820 | RM25_1180 | Putative hydrolase | 1258 | gene | 1358550 | 1359953 | + | 1693,979 | 0,663666123 | 0,120819 | 5,493046 | 3,95E-08 | 2,74E-06 | 1619,108502 | 0,743298591 | 0,152853 | 4,862818 | 1,16E-06 | 5,82E-06 | 1768,849 | 0,592295563 | 0,205876 | 2,876955 | 0,004015 | 0,015191 | 1383,715 | 0,166416 | 0,204859 | 0,812342 | 0,416595 | 0,492435 | 2033,209326 | 0,058267192 | 0,133037 | 0,437978 | 0,661402 | 0,715431 |
| 1197 | RM25_RS05825 | RM25_1181 | Putative insertion element ISCmi2 transposase | 1259 | gene | 1359950 | 1360993 | - | 76,79069 | 0,369302939 | 0,413706 | 0,892671 | 0,372034 | 0,484198 | 91,74688696 | -0,302849934 | 0,358426 | -0,84494 | 0,398142 | 0,467342 | 61,8345 | 1,47857072 | 0,353927 | 4,177615 | 2,95E-05 | 0,00026 | 71,7149 | -1,69127 | 0,36517 | -4,63144 | 3,63E-06 | 2,47E-05 | 84,2965336 | 0,128308217 | 0,355089 | 0,361341 | 0,717845 | 0,76557 |
| 1198 | RM25_RS05830 | RM25_1182 | Metal-dependent hydrolase | 1260 | gene | 1361084 | 1361731 | - | 772,4832 | 0,740912574 | 0,293702 | 2,522666 | 0,011647 | 0,051408 | 1000,992134 | 0,818885851 | 0,183959 | 4,451463 | 8,53E-06 | 3,64E-05 | 543,9742 | 0,604254887 | 0,201218 | 3,00298 | 0,002674 | 0,010978 | 613,9551 | -0,8057 | 0,226745 | -3,55332 | 0,00038 | 0,001568 | 947,6166392 | -0,971694276 | 0,137802 | -7,05137 | 1,77E-12 | 1,47E-11 |
| 1199 | RM25_RS05835 | RM25_1183 | Hypothetical protein | 1261 | gene | 1362168 | 1362329 | + | 681,3656 | 0,748319347 | 0,400808 | 1,867028 | 0,061898 | 0,142196 | 848,09973 | 0,760995826 | 0,262499 | 2,89904 | 0,003743 | 0,008532 | 514,6315 | 0,730372141 | 0,562528 | 1,298375 | 0,194158 | 0,289771 | 541,3626 | -0,74246 | 0,563919 | -1,3166 | 0,187972 | 0,250028 | 836,3760512 | -0,746824854 | 0,271613 | -2,74959 | 0,005967 | 0,010894 |
| 1200 | RM25_RS05840 | RM25_1184 | Hypothetical protein | 1262 | gene | 1362486 | 1363613 | + | 381,8814 | -0,823358714 | 0,535928 | -1,53632 | 0,124459 | 0,221821 | 193,2826586 | -1,304813107 | 0,262813 | -4,9648 | 6,88E-07 | 3,55E-06 | 570,4802 | -0,673072627 | 0,448397 | -1,50106 | 0,133339 | 0,214819 | 509,6461 | 1,282339 | 0,287318 | 4,463143 | 8,08E-06 | 5,08E-05 | 272,1630069 | 1,986448871 | 0,433268 | 4,584799 | 4,54E-06 | 1,42E-05 |
| 1201 | RM25_RS05845 | RM25_1185 | ATP-dependent DNA helicase II uvrD2 | 1263 | gene | 1363670 | 1365826 | - | 1101,153 | 0,074490594 | 0,156337 | 0,476473 | 0,633737 | 0,718825 | 1105,027396 | 0,391240215 | 0,136424 | 2,867826 | 0,004133 | 0,00926 | 1097,279 | -0,241689163 | 0,235817 | -1,0249 | 0,30541 | 0,409705 | 1129,067 | 0,25324 | 0,164959 | 1,535172 | 0,124742 | 0,178192 | 1109,793859 | -0,321735681 | 0,202716 | -1,58713 | 0,112484 | 0,153163 |
| 1202 | RM25_RS05850 | RM25_1186 | Exodeoxyribonuclease V, gamma subunit | 1264 | gene | 1365929 | 1369303 | + | 2010,883 | -0,405315147 | 0,148409 | -2,73107 | 0,006313 | 0,034894 | 1823,817695 | -0,311473842 | 0,14116 | -2,20653 | 0,027347 | 0,047138 | 2197,947 | -0,483932177 | 0,233372 | -2,07365 | 0,038112 | 0,083252 | 2411,588 | 0,281563 | 0,175256 | 1,606583 | 0,108146 | 0,158225 | 1698,610658 | 0,167886195 | 0,193206 | 0,86895 | 0,384874 | 0,450198 |
| 1203 | RM25_RS05855 | RM25_1187 | Exodeoxyribonuclease V, beta subunit | 1265 | gene | 1369300 | 1372662 | + | 1922,316 | -0,441154814 | 0,142701 | -3,09147 | 0,001992 | 0,015341 | 1814,807005 | -0,316143021 | 0,16116 | -1,96167 | 0,049801 | 0,078779 | 2029,826 | -0,554329134 | 0,238907 | -2,32027 | 0,020326 | 0,051962 | 2330,762 | 0,19939 | 0,218797 | 0,911304 | 0,362135 | 0,437512 | 1601,199728 | 0,020931466 | 0,179386 | 0,116684 | 0,90711 | 0,924366 |
| 1204 | RM25_RS05860 | RM25_1188 | Exodeoxyribonuclease V, alpha subunit | 1266 | gene | 1372649 | 1374502 | + | 956,9635 | -0,211092279 | 0,15413 | -1,36957 | 0,170821 | 0,274454 | 956,1718606 | 0,041056327 | 0,137377 | 0,298859 | 0,765048 | 0,811164 | 957,7551 | -0,463750978 | 0,260518 | -1,78011 | 0,075058 | 0,137889 | 1082,292 | 0,175023 | 0,145677 | 1,201441 | 0,22958 | 0,297482 | 871,8113458 | -0,26878016 | 0,241612 | -1,11244 | 0,265947 | 0,325461 |
| 1205 | RM25_RS05865 | RM25_1189 | NAD(+) diphosphatase | 1267 | gene | 1374554 | 1375444 | - | 862,3734 | 0,068916956 | 0,17736 | 0,38857 | 0,697594 | 0,774484 | 939,8639933 | 0,400682336 | 0,147775 | 2,711443 | 0,006699 | 0,014085 | 784,8829 | -0,32232941 | 0,200884 | -1,60455 | 0,108592 | 0,183478 | 889,202 | 0,050702 | 0,213055 | 0,237977 | 0,811899 | 0,848623 | 865,9672127 | -0,624951788 | 0,122117 | -5,11767 | 3,09E-07 | 1,17E-06 |
| 1206 | RM25_RS05870 | RM25_1190 | UvrD/REP helicase | 1268 | gene | 1375454 | 1378810 | - | 1944,423 | 0,039052389 | 0,102658 | 0,380412 | 0,70364 | 0,780094 | 1827,632855 | 0,192695486 | 0,130551 | 1,476012 | 0,139941 | 0,192152 | 2061,212 | -0,096308877 | 0,136077 | -0,70776 | 0,479097 | 0,580284 | 2021,465 | 0,25961 | 0,125781 | 2,063978 | 0,03902 | 0,069411 | 1932,987847 | 0,024899155 | 0,134868 | 0,184619 | 0,853528 | 0,881194 |
| 1207 | RM25_RS05875 | RM25_1191 | ATP-dependent DNA helicase | 1269 | gene | 1378807 | 1382178 | - | 1951,401 | 0,215391626 | 0,129717 | 1,660479 | 0,096818 | 0,189535 | 1868,53019 | 0,502914703 | 0,112167 | 4,48364 | 7,34E-06 | 3,14E-05 | 2034,273 | -0,044096317 | 0,175818 | -0,25081 | 0,801964 | 0,857247 | 1902,462 | 0,355044 | 0,137528 | 2,581607 | 0,009834 | 0,023437 | 2057,956045 | -0,134571436 | 0,153098 | -0,87899 | 0,379406 | 0,444905 |
| 1208 | RM25_RS05880 | RM25_1192 | Cold shock-like protein CspA | 1270 | gene | 1382518 | 1382724 | + | 11164,15 | 0,804436384 | 0,455587 | 1,765716 | 0,077444 | 0,164837 | 14286,00035 | 0,371543155 | 0,218933 | 1,697066 | 0,089684 | 0,128894 | 8042,308 | 1,691624457 | 0,500058 | 3,382855 | 0,000717 | 0,0037 | 8689,944 | -1,75585 | 0,499477 | -3,51539 | 0,000439 | 0,001766 | 13893,57011 | -0,406590039 | 0,222743 | -1,82538 | 0,067944 | 0,097471 |
| 1209 | RM25_RS05885 | RM25_1193 | Serine/threonine phosphatase stp | 1271 | gene | 1382890 | 1383762 | + | 606,0646 | 0,303120832 | 0,178384 | 1,699261 | 0,08927 | 0,179837 | 679,8687755 | 0,55212709 | 0,172907 | 3,193203 | 0,001407 | 0,003609 | 532,2603 | -0,004120623 | 0,195412 | -0,02109 | 0,983176 | 0,990743 | 572,4115 | -0,10458 | 0,152052 | -0,68778 | 0,49159 | 0,559747 | 657,4265593 | -0,607725966 | 0,197398 | -3,07868 | 0,002079 | 0,004125 |
| 1210 | RM25_RS05890 | RM25_1194 | Hypothetical protein | 1272 | gene | 1383819 | 1384103 | + | 2443,325 | 1,329210741 | 0,541123 | 2,456393 | 0,014034 | 0,058236 | 3742,457105 | 1,31984526 | 0,250436 | 5,270182 | 1,36E-07 | 8,03E-07 | 1144,193 | 1,361843051 | 0,434185 | 3,136547 | 0,00171 | 0,00756 | 1486,928 | -1,79153 | 0,347713 | -5,15232 | 2,57E-07 | 2,17E-06 | 3422,732085 | -1,721937591 | 0,339334 | -5,07446 | 3,89E-07 | 1,45E-06 |
| 1211 | RM25_RS05895 | RM25_1195 | Hypothetical protein | 1273 | gene | 1384290 | 1385045 | - | 1557,394 | 0,714446314 | 0,209971 | 3,402598 | 0,000667 | 0,007637 | 1616,182834 | 0,619814166 | 0,184095 | 3,366812 | 0,00076 | 0,002099 | 1498,606 | 0,819042192 | 0,359422 | 2,278775 | 0,02268 | 0,05645 | 1250,461 | -0,28383 | 0,324529 | -0,8746 | 0,38179 | 0,458654 | 1890,853967 | -0,050978805 | 0,227613 | -0,22397 | 0,82278 | 0,856199 |
| 1212 | RM25_RS05900 | RM25_1196 | DEAD-box ATP-dependent RNA helicase CshA | 1274 | gene | 1385321 | 1386826 | + | 2000,552 | 0,32997425 | 0,389639 | 0,846872 | 0,397066 | 0,508357 | 2771,85264 | 0,166496758 | 0,164656 | 1,01118 | 0,311931 | 0,378979 | 1229,251 | 0,709334443 | 0,232359 | 3,05275 | 0,002268 | 0,009585 | 1888,712 | -1,54448 | 0,197959 | -7,802 | 6,09E-15 | 2,35E-13 | 2183,17062 | -0,956514534 | 0,166179 | -5,75594 | 8,62E-09 | 4,14E-08 |
| 1213 | RM25_RS05905 | RM25_1197 | Tryptophan--tRNA ligase | 1275 | gene | 1386964 | 1388136 | - | 3090,037 | 0,558233709 | 0,401742 | 1,389532 | 0,164671 | 0,267091 | 4335,966395 | 0,693419739 | 0,195867 | 3,540255 | 0,0004 | 0,001193 | 1844,108 | 0,250214607 | 0,32198 | 0,777112 | 0,437093 | 0,54137 | 2659,573 | -1,02744 | 0,297459 | -3,45405 | 0,000552 | 0,002137 | 3606,840572 | -1,435082803 | 0,218641 | -6,56366 | 5,25E-11 | 3,56E-10 |
| 1214 | RM25_RS05910 | RM25_1198 | PHP domain protein | 1276 | gene | 1388105 | 1388986 | - | 1457,981 | 0,155521102 | 0,2795 | 0,556427 | 0,577919 | 0,670352 | 1349,403061 | 0,693527039 | 0,208043 | 3,333582 | 0,000857 | 0,002336 | 1566,559 | -0,296317622 | 0,38085 | -0,77804 | 0,436543 | 0,540974 | 1455,79 | 0,692947 | 0,389314 | 1,779919 | 0,075089 | 0,117364 | 1503,470663 | -0,260171968 | 0,193016 | -1,34793 | 0,177681 | 0,229224 |
| 1215 | RM25_RS05915 | RM25_1199 | Hypothetical protein | 1277 | gene | 1389118 | 1389594 | + | 1327,201 | -0,51855583 | 0,384407 | -1,34898 | 0,177345 | 0,281104 | 1067,866709 | 0,452539504 | 0,204371 | 2,214301 | 0,026808 | 0,046325 | 1586,535 | -1,228450659 | 0,458043 | -2,68195 | 0,007319 | 0,024196 | 1644,877 | 1,254496 | 0,41819 | 2,999819 | 0,002701 | 0,008174 | 1066,776316 | -0,397360242 | 0,276518 | -1,43701 | 0,150714 | 0,19901 |
| 1216 | RM25_RS05920 | RM25_1200 | Hypothetical protein | 1278 | gene | 1389653 | 1389856 | - | 59,14535 | 0,089615628 | 0,217276 | 0,41245 | 0,68001 | 0,759253 | 61,60969727 | 0,008333099 | 0,322399 | 0,025847 | 0,979379 | 0,981879 | 56,68101 | 0,169802928 | 0,357706 | 0,4747 | 0,635001 | 0,718185 | 61,06403 | -0,2609 | 0,311517 | -0,83752 | 0,402298 | 0,477932 | 59,66887044 | -0,038476043 | 0,401066 | -0,09593 | 0,923573 | 0,937205 |
| 1217 | RM25_RS05925 | RM25_1201 | Hypothetical protein | 1279 | gene | 1389858 | 1390367 | - | 503,8402 | 0,163030379 | 0,211117 | 0,772229 | 0,439979 | 0,548113 | 592,9272364 | 0,345200497 | 0,183571 | 1,880471 | 0,060044 | 0,091423 | 414,7531 | -0,090262854 | 0,193259 | -0,46706 | 0,640459 | 0,722279 | 503,5546 | -0,34503 | 0,218712 | -1,57754 | 0,114671 | 0,166122 | 521,8487039 | -0,733951533 | 0,154434 | -4,75252 | 2,01E-06 | 6,61E-06 |
| 1218 | RM25_RS05935 | RM25_1202 | Hypothetical protein | 1280 | gene | 1390448 | 1390990 | + | 344,2625 | -0,337525322 | 0,140142 | -2,40845 | 0,01602 | 0,063048 | 316,5887948 | -0,159748914 | 0,178686 | -0,89402 | 0,371311 | 0,439789 | 371,9362 | -0,48931983 | 0,203552 | -2,40391 | 0,016221 | 0,043804 | 404,8264 | 0,313826 | 0,185491 | 1,691866 | 0,090671 | 0,13682 | 298,7806854 | 0,044894366 | 0,206785 | 0,217107 | 0,828125 | 0,861002 |
| 1219 | RM25_RS05940 | RM25_1203 | Protein translocase subunit SecA | 1281 | gene | 1391030 | 1393879 | - | 6753,443 | 0,746220603 | 0,381674 | 1,955127 | 0,050568 | 0,12544 | 9009,094717 | 0,514806776 | 0,195285 | 2,636179 | 0,008385 | 0,017066 | 4497,791 | 1,244038767 | 0,280004 | 4,442938 | 8,87E-06 | 9,25E-05 | 5376,321 | -1,52854 | 0,241648 | -6,32551 | 2,52E-10 | 4,17E-09 | 8281,893771 | -0,762022674 | 0,213808 | -3,56406 | 0,000365 | 0,00084 |
| 1220 | RM25_RS05945 | RM25_1204 | S30AE family protein | 1282 | gene | 1394124 | 1394720 | - | 9083,578 | 0,056275779 | 0,309073 | 0,182079 | 0,855521 | 0,90513 | 9200,885654 | 0,573918345 | 0,191722 | 2,993496 | 0,002758 | 0,006512 | 8966,271 | -0,471075343 | 0,460867 | -1,02215 | 0,30671 | 0,410806 | 9408,02 | 0,450133 | 0,418531 | 1,075508 | 0,282147 | 0,355816 | 9058,198861 | -0,56824096 | 0,265994 | -2,13629 | 0,032656 | 0,050671 |
| 1221 | RM25_RS05950 | RM25_1205 | Phosphoribosyltransferase | 1283 | gene | 1394876 | 1395685 | - | 190,1092 | -0,948832272 | 0,617244 | -1,53721 | 0,124242 | 0,221821 | 88,17726371 | -1,946288011 | 0,353575 | -5,5046 | 3,7E-08 | 2,46E-07 | 292,0411 | -0,695951017 | 0,445068 | -1,56369 | 0,117889 | 0,196232 | 261,4648 | 1,302992 | 0,300091 | 4,341993 | 1,41E-05 | 8,43E-05 | 128,0762908 | 2,622570115 | 0,489436 | 5,358355 | 8,4E-08 | 3,47E-07 |
| 1222 | RM25_RS05955 | RM25_1206 | DNA-binding response regulator MtrA | 1284 | gene | 1395767 | 1396507 | - | 553,8062 | 0,411050698 | 0,228041 | 1,802527 | 0,071463 | 0,15596 | 662,067695 | 0,621128513 | 0,161128 | 3,854866 | 0,000116 | 0,000388 | 445,5448 | 0,108820722 | 0,239335 | 0,454679 | 0,64934 | 0,72985 | 502,3609 | -0,3408 | 0,151198 | -2,25396 | 0,024199 | 0,047649 | 619,6316547 | -0,807127013 | 0,206313 | -3,91214 | 9,15E-05 | 0,000234 |
| 1223 | RM25_RS05960 | RM25_1207 | UPF0301 protein | 1285 | gene | 1396643 | 1397209 | - | 512,6041 | 0,363144038 | 0,178587 | 2,033426 | 0,04201 | 0,111691 | 576,5931004 | 0,569839832 | 0,187124 | 3,045253 | 0,002325 | 0,005597 | 448,615 | 0,106833535 | 0,200941 | 0,531666 | 0,594957 | 0,683722 | 473,8756 | -0,16191 | 0,208765 | -0,77556 | 0,438009 | 0,51337 | 565,135375 | -0,577235176 | 0,162527 | -3,55163 | 0,000383 | 0,000877 |
| 1225 | RM25_RS05965 | RM25_1208 | Putative membrane protein | 1286 | gene | 1397915 | 1398556 | - | 1320,825 | 1,164129278 | 0,209521 | 5,556138 | 2,76E-08 | 2,03E-06 | 1504,23654 | 1,099486535 | 0,161402 | 6,812095 | 9,62E-12 | 1,11E-10 | 1137,412 | 1,252874826 | 0,289336 | 4,330171 | 1,49E-05 | 0,000147 | 865,2303 | -0,56699 | 0,2197 | -2,58077 | 0,009858 | 0,02347 | 1786,046386 | -0,372734678 | 0,220298 | -1,69196 | 0,090654 | 0,126126 |
| 1226 | RM25_RS05970 | RM25_1209 | SsrA-binding protein 2 | 1287 | gene | 1398646 | 1399128 | - | 981,9817 | 1,030715937 | 0,403726 | 2,553008 | 0,01068 | 0,048973 | 1399,189394 | 1,061228475 | 0,191163 | 5,551434 | 2,83E-08 | 1,9E-07 | 564,774 | 0,960088393 | 0,216308 | 4,438524 | 9,06E-06 | 9,4E-05 | 687,247 | -1,29894 | 0,219539 | -5,91665 | 3,29E-09 | 4,06E-08 | 1293,434153 | -1,35334363 | 0,18011 | -7,51398 | 5,74E-14 | 6,01E-13 |
| 1227 | RM25_RS05975 | RM25_1210 | MutG family lantibiotic protection ABC superfamily ATP binc | 1288 | gene | 1399279 | 1401231 | - | 2282,875 | 0,591333394 | 0,260808 | 2,26731 | 0,023371 | 0,078136 | 2854,664613 | 0,667760379 | 0,174553 | 3,825552 | 0,00013 | 0,000432 | 1711,086 | 0,466947666 | 0,21691 | 2,152722 | 0,031341 | 0,071858 | 1930,101 | -0,67172 | 0,215655 | -3,11479 | 0,001841 | 0,005871 | 2688,697998 | -0,832733573 | 0,14624 | -5,69428 | 1,24E-08 | 5,78E-08 |
| 1228 | RM25_RS05980 | RM25_1211 | Hypothetical protein | 1289 | gene | 1401415 | 1402734 | - | 2117,567 | 0,790368831 | 0,376153 | 2,101189 | 0,035624 | 0,1008 | 2776,21517 | 0,495105536 | 0,188824 | 2,622044 | 0,00874 | 0,017653 | 1458,919 | 1,4081325 | 0,259097 | 5,434769 | 5,49E-08 | 1,01E-06 | 1652,508 | -1,58945 | 0,237634 | -6,68863 | 2,25E-11 | 4,7E-10 | 2626,934177 | -0,631013896 | 0,176429 | -3,57659 | 0,000348 | 0,000802 |
| 1229 | RM25_RS05985 | RM25_1212 | Cell division protein FtsX | 1290 | gene | 1402801 | 1403718 | - | 1136,556 | 0,613338388 | 0,375102 | 1,635125 | 0,102023 | 0,195234 | 1503,932694 | 0,34390981 | 0,168177 | 2,044925 | 0,040862 | 0,06721 | 769,1797 | 1,17888151 | 0,287943 | 4,09415 | 4,24E-05 | 0,000353 | 957,5386 | -1,55347 | 0,22127 | -7,02071 | 2,21E-12 | 5,48E-11 | 1345,159668 | -0,673177042 | 0,208214 | -3,23311 | 0,001225 | 0,002534 |
| 1230 | RM25_RS05990 | RM25_1213 | Cell division ATP-binding protein FtsE | 1291 | gene | 1403757 | 1404443 | - | 1353,969 | 0,529921078 | 0,474965 | 1,115705 | 0,264548 | 0,372263 | 1857,888368 | 0,141289229 | 0,243516 | 0,580205 | 0,561777 | 0,62576 | 850,0489 | 1,479845484 | 0,391298 | 3,781885 | 0,000156 | 0,00107 | 1185,111 | -2,03231 | 0,344636 | -5,89696 | 3,7E-09 | 4,48E-08 | 1563,834523 | -0,65985695 | 0,291253 | -2,26558 | 0,023477 | 0,037849 |
| 1231 | RM25_RS05995 | RM25_1214 | Peptide chain release factor 2 | 1292 | gene | 1404595 | 1405704 | - | 2184,566 | 0,841059265 | 0,180621 | 4,656484 | 3,22E-06 | 0,000107 | 2352,75171 | 0,738644018 | 0,151216 | 4,884705 | 1,04E-06 | 5,26E-06 | 2016,379 | 0,963883469 | 0,295401 | 3,262967 | 0,001103 | 0,00525 | 1660,892 | -0,42254 | 0,249471 | -1,69376 | 0,090311 | 0,13645 | 2741,192403 | -0,158550717 | 0,200394 | -0,79119 | 0,428831 | 0,493773 |
| 1232 | RM25_RS06000 | RM25_1215 | GatB/YqeY domain protein | 1293 | gene | 1405799 | 1406260 | + | 1066,782 | 0,279329181 | 0,242533 | 1,151714 | 0,249439 | 0,35762 | 1236,100044 | 0,335955051 | 0,195734 | 1,716383 | 0,086092 | 0,124566 | 897,4643 | 0,203672094 | 0,339941 | 0,599139 | 0,54908 | 0,645799 | 1022,678 | -0,44238 | 0,31635 | -1,39838 | 0,162 | 0,220875 | 1143,988135 | -0,539686978 | 0,219826 | -2,45506 | 0,014086 | 0,023849 |
| 1234 | RM25_RS06010 | RM25_1217 | Fructosamine kinase | 1297 | gene | 1406449 | 1407480 | - | 370,2959 | -0,775260344 | 0,318687 | -2,43267 | 0,014988 | 0,060854 | 261,5127082 | -0,752035483 | 0,234071 | -3,21285 | 0,001314 | 0,003415 | 479,0792 | -0,790855212 | 0,29406 | -2,68943 | 0,007157 | 0,023884 | 490,6736 | 0,82232 | 0,25746 | 3,193969 | 0,001403 | 0,00469 | 268,4119072 | 0,847213081 | 0,287921 | 2,94252 | 0,003256 | 0,006233 |
| 1235 | RM25_RS06015 | RM25_1218 | 4-aminobutyrate transaminase | 1298 | gene | 1407559 | 1408896 | + | 2319,364 | 0,424958682 | 0,354817 | 1,197683 | 0,23104 | 0,338451 | 2981,035233 | 0,077325827 | 0,211288 | 0,365974 | 0,714384 | 0,76467 | 1657,693 | 1,090854901 | 0,239026 | 4,563756 | 5,02E-06 | 5,57E-05 | 2111,771 | -1,51477 | 0,251627 | -6,01988 | 1,75E-09 | 2,36E-08 | 2601,701422 | -0,455299469 | 0,181763 | -2,50491 | 0,012248 | 0,021088 |
| 1236 | RM25_RS06020 | RM25_1219 | Aliphatic amidase | 1299 | gene | 1409075 | 1409935 | - | 535,3503 | -0,115770985 | 0,197235 | -0,58697 | 0,557224 | 0,6518 | 622,3063415 | -0,22210373 | 0,190704 | -1,16465 | 0,244159 | 0,308902 | 448,3942 | 0,031868699 | 0,19605 | 0,162554 | 0,87087 | 0,906552 | 590,3467 | -0,65659 | 0,219727 | -2,9882 | 0,002806 | 0,008401 | 503,8220907 | -0,346744855 | 0,158361 | -2,18958 | 0,028555 | 0,045231 |
| 1237 | RM25_RS06025 | RM25_1220 | Putrescine importer | 1300 | gene | 1409932 | 1411323 | - | 1186,072 | -0,733395966 | 0,198271 | -3,69895 | 0,000216 | 0,003074 | 1385,406876 | -0,8744925 | 0,145496 | -6,01043 | 1,85E-09 | 1,52E-08 | 986,7368 | -0,540022166 | 0,241737 | -2,23393 | 0,025488 | 0,06177 | 1571,417 | -0,67996 | 0,198884 | -3,41888 | 0,000629 | 0,002397 | 874,6703596 | -0,28502187 | 0,197004 | -1,44678 | 0,147959 | 0,196141 |
| 1238 | RM25_RS06030 | RM25_1221 | Putative heat shock protein Hsp18 | 1301 | gene | 1411596 | 1412048 | - | 14513,38 | -0,553202851 | 0,316039 | -1,75042 | 0,080045 | 0,167258 | 14901,04019 | -0,154847178 | 0,206869 | -0,74853 | 0,454143 | 0,522154 | 14125,72 | -0,997659458 | 0,487063 | -2,04832 | 0,040529 | 0,086685 | 18294,19 | 0,213016 | 0,448391 | 0,475067 | 0,634739 | 0,691669 | 11498,18862 | -0,601797464 | 0,280832 | -2,14291 | 0,03212 | 0,050101 |
| 1239 | RM25_RS06035 | RM25_1222 | Transport protein MsbA | 1302 | gene | 1412298 | 1414091 | + | 862,4065 | -0,845277314 | 0,273706 | -3,08827 | 0,002013 | 0,015457 | 678,7252416 | -0,835707189 | 0,238975 | -3,49705 | 0,00047 | 0,001377 | 1046,088 | -0,853103593 | 0,310619 | -2,74646 | 0,006024 | 0,02082 | 1162,932 | 0,565434 | 0,313669 | 1,802647 | 0,071444 | 0,113396 | 606,0991536 | 0,611928115 | 0,246466 | 2,482814 | 0,013035 | 0,022224 |
| 1240 | RM25_RS06040 | RM25_1223 | Hypothetical protein | 1303 | gene | 1414126 | 1414551 | - | 2928,591 | 0,932669061 | 0,352687 | 2,644466 | 0,008182 | 0,040909 | 3820,456392 | 0,897499584 | 0,200705 | 4,47174 | 7,76E-06 | 3,31E-05 | 2036,725 | 1,000739434 | 0,381282 | 2,624673 | 0,008673 | 0,02744 | 2145,143 | -1,02544 | 0,363201 | -2,82334 | 0,004753 | 0,012658 | 3762,920193 | -0,888758197 | 0,226972 | -3,91572 | 9,01E-05 | 0,000231 |
| 1241 | NA | RM25_1224 | Hypothetical protein | 1304 | gene | 1414556 | 1414690 | - | 10,56784 | 1,241949479 | 0,565293 | 2,197003 | 0,02802 | 0,086672 | 13,84449356 | 0,750262671 | 0,637115 | 1,177594 | 0,238958 | 0,302809 | 7,291188 | 2,422018098 | 0,74624 | 3,24563 | 0,001172 | 0,005529 | 6,867842 | -2,22793 | 0,709341 | -3,14084 | 0,001685 | 0,005447 | 14,42284381 | -0,504711839 | 0,693325 | -0,72796 | 0,466639 | 0,530057 |
| 1242 | RM25_RS06045 | RM25_1225 | Pyruvate dehydrogenase E1 component | 1305 | gene | 1414653 | 1417400 | - | 12737,99 | 0,86239505 | 0,279708 | 3,083193 | 0,002048 | 0,0156 | 15170,84435 | 0,96557341 | 0,188681 | 5,117501 | 3,1E-07 | 1,71E-06 | 10305,13 | 0,714498249 | 0,395539 | 1,806389 | 0,070858 | 0,132025 | 9586,774 | -0,44346 | 0,348013 | -1,27425 | 0,202574 | 0,266592 | 16077,79592 | -0,666145236 | 0,250593 | -2,65827 | 0,007854 | 0,013993 |
| 1243 | RM25_RS06050 | RM25_1226 | Aromatic-amino-acid transaminase | 1306 | gene | 1417655 | 1418851 | + | 1698,052 | 0,260524793 | 0,370577 | 0,703025 | 0,48204 | 0,582054 | 2354,617999 | 0,257448456 | 0,172492 | 1,492527 | 0,135561 | 0,186852 | 1041,485 | 0,268603502 | 0,18643 | 1,440774 | 0,149648 | 0,234678 | 1643,862 | -1,24264 | 0,200044 | -6,21185 | 5,24E-10 | 7,91E-09 | 1814,927699 | -1,18318842 | 0,131092 | -9,02565 | 1,79E-19 | 4,09E-18 |
| 1244 | RM25_RS06055 | RM25_1227 | Transcriptional regulator, PucR family | 1307 | gene | 1418937 | 1420151 | + | 610,0437 | -0,421495247 | 0,25286 | -1,66691 | 0,095532 | 0,187641 | 642,1214018 | 0,003741723 | 0,132852 | 0,028165 | 0,977531 | 0,98086 | 577,9661 | -0,914705709 | 0,37388 | -2,44652 | 0,014424 | 0,040139 | 735,0732 | 0,172154 | 0,215223 | 0,799886 | 0,423777 | 0,498922 | 513,4132243 | -0,6780084 | 0,328144 | -2,06619 | 0,038811 | 0,059093 |
| 1245 | RM25_RS06060 | RM25_1228 | Malonyl-CoA--acyl carrier protein transacylase | 1308 | gene | 1420308 | 1421264 | + | 3173,941 | 0,120929874 | 0,700983 | 0,172515 | 0,863033 | 0,908517 | 5397,009056 | -0,013388456 | 0,179658 | -0,07452 | 0,940595 | 0,953131 | 950,8733 | 0,913890359 | 0,384337 | 2,377837 | 0,017415 | 0,046327 | 3257,846 | -3,09941 | 0,234601 | -13,2114 | 7,54E-40 | 4,44E-37 | 3241,25905 | -2,134651563 | 0,300933 | -7,09344 | 1,31E-12 | 1,12E-11 |
| 1246 | RM25_RS06065 | RM25_1229 | 3-oxoacyl-[acyl-carrier-protein] synthase 3 | 1309 | gene | 1421268 | 1422272 | + | 4849,833 | -0,114121568 | 0,757566 | -0,15064 | 0,880258 | 0,918533 | 8410,816029 | -0,331960541 | 0,191882 | -1,73003 | 0,083626 | 0,121745 | 1288,851 | 1,399578956 | 0,332757 | 4,20601 | 2,6E-05 | 0,000232 | 5411,183 | -3,78542 | 0,233999 | -16,177 | 7,32E-59 | 1,44E-55 | 4566,020419 | -2,015443363 | 0,257749 | -7,8194 | 5,31E-15 | 6,48E-14 |
| 1247 | RM25_RS06070 | RM25_1230 | Acyl carrier protein | 1310 | gene | 1422334 | 1422585 | + | 7431,847 | -0,369517894 | 0,828756 | -0,44587 | 0,655691 | 0,738119 | 13113,22995 | -0,650226271 | 0,200761 | -3,2388 | 0,0012 | 0,003152 | 1750,464 | 1,906698648 | 0,417199 | 4,570236 | 4,87E-06 | 5,47E-05 | 9012,768 | -4,49508 | 0,29173 | -15,4084 | 1,44E-53 | 1,13E-50 | 6351,28618 | -1,911112119 | 0,323202 | -5,91306 | 3,36E-09 | 1,72E-08 |
| 1248 | RM25_RS06075 | RM25_1231 | 3-oxoacyl-[acyl-carrier-protein] synthase II | 1311 | gene | 1422637 | 1423896 | + | 14027,42 | -0,210199391 | 0,906374 | -0,23191 | 0,816606 | 0,872039 | 25578,41869 | -0,407301065 | 0,213875 | -1,90439 | 0,05686 | 0,087594 | 2476,412 | 2,114695011 | 0,363464 | 5,818162 | 5,95E-09 | 1,4E-07 | 16173,03 | -5,02653 | 0,311325 | -16,1456 | 1,22E-58 | 1,44E-55 | 12757,75475 | -2,47211063 | 0,255134 | -9,68945 | 3,34E-22 | 9,38E-21 |
| 1249 | RM25_RS06080 | RM25_1232 | PF11343 family protein | 1312 | gene | 1424080 | 1424574 | - | 911,8363 | 0,826891857 | 0,205155 | 4,030573 | 5,56E-05 | 0,001082 | 989,1201654 | 0,667213213 | 0,183365 | 3,638712 | 0,000274 | 0,00085 | 834,5525 | 1,023796766 | 0,309332 | 3,3097 | 0,000934 | 0,004615 | 698,5031 | -0,53186 | 0,30565 | -1,7401 | 0,081842 | 0,125669 | 1140,505741 | -0,132244882 | 0,17965 | -0,73612 | 0,461655 | 0,526171 |
| 1250 | RM25_RS06085 | RM25_1233 | Acyl-peptide hydrolase | 1313 | gene | 1424725 | 1426704 | + | 1342,125 | -0,302618192 | 0,216304 | -1,39904 | 0,1618 | 0,264453 | 1160,802174 | 0,22886364 | 0,152678 | 1,498992 | 0,133876 | 0,184962 | 1523,449 | -0,718637635 | 0,195545 | -3,67505 | 0,000238 | 0,001515 | 1558,489 | 0,773546 | 0,207556 | 3,726916 | 0,000194 | 0,000876 | 1178,611502 | -0,128911281 | 0,132056 | -0,97619 | 0,328972 | 0,392999 |
| 1251 | RM25_RS06090 | RM25_1234 | Pyridoxal phosphate enzyme, YggS family | 1314 | gene | 1426753 | 1427505 | - | 464,9815 | 0,334001213 | 0,299982 | 1,113405 | 0,265535 | 0,372983 | 605,2936669 | 0,464307314 | 0,17991 | 2,580781 | 0,009858 | 0,019655 | 324,6693 | 0,099483478 | 0,187857 | 0,52957 | 0,59641 | 0,684133 | 435,7149 | -0,75371 | 0,19755 | -3,81529 | 0,000136 | 0,000642 | 509,012349 | -1,066354896 | 0,154407 | -6,90612 | 4,98E-12 | 3,84E-11 |
| 1252 | RM25_RS06095 | RM25_1235 | UPF0353 protein | 1315 | gene | 1427502 | 1428467 | - | 853,8653 | 0,214530178 | 0,352605 | 0,608415 | 0,542912 | 0,638531 | 1164,31383 | 0,33286843 | 0,158319 | 2,102524 | 0,035507 | 0,05919 | 543,4167 | -0,034117628 | 0,211648 | -0,1612 | 0,871936 | 0,906552 | 838,2528 | -0,96626 | 0,208511 | -4,63409 | 3,59E-06 | 2,45E-05 | 900,1430888 | -1,279504223 | 0,117253 | -10,9123 | 1,01E-27 | 4,94E-26 |
| 1253 | RM25_RS06100 | RM25_1236 | UPF0353 protein | 1316 | gene | 1428467 | 1429441 | - | 798,5281 | 0,050012537 | 0,327161 | 0,152868 | 0,878502 | 0,917832 | 1047,616828 | 0,286820878 | 0,137804 | 2,081368 | 0,0374 | 0,061948 | 549,4394 | -0,399807945 | 0,244165 | -1,63745 | 0,101536 | 0,174559 | 831,0604 | -0,65425 | 0,173329 | -3,7746 | 0,00016 | 0,000741 | 798,1420486 | -1,283319432 | 0,202486 | -6,33783 | 2,33E-10 | 1,4E-09 |
| 1254 | RM25_RS06105 | RM25_1237 | Hypothetical protein | 1317 | gene | 1429441 | 1430505 | - | 851,2078 | 0,206475503 | 0,301047 | 0,685859 | 0,492802 | 0,591714 | 1086,819176 | 0,471175073 | 0,158228 | 2,977823 | 0,002903 | 0,006802 | 615,5964 | -0,251587174 | 0,246521 | -1,02055 | 0,307467 | 0,411294 | 836,2461 | -0,50871 | 0,225009 | -2,26082 | 0,02377 | 0,047042 | 895,5852886 | -1,17140211 | 0,169821 | -6,89787 | 5,28E-12 | 4,04E-11 |
| 1255 | RM25_RS06110 | RM25_1238 | Transcriptional regulator moxR1 | 1318 | gene | 1430502 | 1431554 | - | 1853,381 | 0,508308405 | 0,305289 | 1,665008 | 0,095911 | 0,188229 | 2403,991274 | 0,48703926 | 0,176359 | 2,761639 | 0,005751 | 0,012414 | 1302,77 | 0,54955732 | 0,235752 | 2,331081 | 0,019749 | 0,050707 | 1625,506 | -0,97604 | 0,231668 | -4,21308 | 2,52E-05 | 0,000141 | 2132,20532 | -0,87200709 | 0,155029 | -5,62479 | 1,86E-08 | 8,53E-08 |
| 1256 | RM25_RS06115 | RM25_1239 | Hypothetical protein | 1319 | gene | 1431799 | 1432725 | + | 1013,848 | 0,108054032 | 0,332177 | 0,325291 | 0,744961 | 0,814412 | 1317,743621 | 0,424503666 | 0,184631 | 2,299201 | 0,021494 | 0,037976 | 709,9515 | -0,47481628 | 0,233312 | -2,03511 | 0,04184 | 0,088684 | 1032,146 | -0,50492 | 0,204222 | -2,47239 | 0,013421 | 0,029965 | 1033,076388 | -1,349424021 | 0,204438 | -6,60064 | 4,09E-11 | 2,79E-10 |
| 1257 | RM25_RS06120 | RM25_1240 | Putative hydrolase | 1320 | gene | 1432733 | 1433575 | + | 628,0376 | 0,008030071 | 0,289002 | 0,027785 | 0,977833 | 0,986202 | 719,2591299 | 0,601663275 | 0,160867 | 3,74014 | 0,000184 | 0,000592 | 536,8161 | -0,787370406 | 0,26144 | -3,01166 | 0,002598 | 0,010763 | 660,7999 | 0,192563 | 0,202352 | 0,951627 | 0,341286 | 0,416793 | 619,1223298 | -1,137854917 | 0,23841 | -4,77267 | 1,82E-06 | 6,04E-06 |
| 1258 | RM25_RS06125 | RM25_1241 | Dihydrolipoyl dehydrogenase | 1321 | gene | 1433596 | 1435026 | - | 2213,654 | 0,495432851 | 0,216626 | 2,287042 | 0,022193 | 0,076142 | 2516,987844 | 0,599524529 | 0,20622 | 2,907212 | 0,003647 | 0,008369 | 1910,321 | 0,361156154 | 0,292606 | 1,234274 | 0,217101 | 0,314316 | 1945,153 | -0,30796 | 0,305831 | -1,00695 | 0,313961 | 0,388454 | 2535,739905 | -0,511353915 | 0,172806 | -2,95912 | 0,003085 | 0,005931 |
| 1259 | RM25_RS06130 | RM25_1242 | Hypothetical protein | 1322 | gene | 1435435 | 1436076 | + | 459,0348 | -0,272237681 | 0,260581 | -1,04473 | 0,296147 | 0,406297 | 341,7401935 | -0,122609797 | 0,187375 | -0,65436 | 0,512883 | 0,577851 | 576,3293 | -0,365121935 | 0,182596 | -1,99962 | 0,045542 | 0,094325 | 527,6622 | 0,802145 | 0,208377 | 3,849479 | 0,000118 | 0,000566 | 407,0921812 | 0,610518292 | 0,150102 | 4,067351 | 4,76E-05 | 0,000126 |
| 1260 | RM25_RS06135 | RM25_1243 | TIGR00245 family protein | 1323 | gene | 1436084 | 1436842 | + | 472,0646 | -0,082705512 | 0,1626 | -0,50864 | 0,611003 | 0,700247 | 448,9522655 | 0,301856842 | 0,17572 | 1,717828 | 0,085828 | 0,124161 | 495,1769 | -0,430077219 | 0,176797 | -2,43261 | 0,014991 | 0,041133 | 510,7796 | 0,439293 | 0,161863 | 2,713975 | 0,006648 | 0,016777 | 450,2150389 | -0,238963613 | 0,18782 | -1,2723 | 0,203266 | 0,25758 |
| 1261 | RM25_RS06140 | RM25_1244 | Deoxyguanosinetriphosphate triphosphohydrolase-like prote | 1324 | gene | 1436859 | 1438004 | - | 1032,725 | -0,00164178 | 0,16413 | -0,01 | 0,992019 | 0,995821 | 932,643465 | 0,36001683 | 0,140324 | 2,565604 | 0,0103 | 0,020383 | 1132,806 | -0,298765652 | 0,153298 | -1,94892 | 0,051305 | 0,102828 | 1087,589 | 0,553279 | 0,166322 | 3,326557 | 0,000879 | 0,003157 | 1011,964 | -0,054639899 | 0,126134 | -0,43319 | 0,664877 | 0,717873 |
| 1262 | RM25_RS06145 | RM25_1245 | TIM-barrel protein, nifR3 family | 1325 | gene | 1438055 | 1439242 | - | 872,8006 | 0,600547313 | 0,078115 | 7,687948 | 1,5E-14 | 3,92E-12 | 880,0456294 | 0,638801673 | 0,127763 | 4,999892 | 5,74E-07 | 3E-06 | 865,5557 | 0,565751815 | 0,179354 | 3,154389 | 0,001608 | 0,007166 | 732,1347 | -0,04431 | 0,156615 | -0,28292 | 0,777235 | 0,819662 | 1031,101739 | -0,064066686 | 0,128119 | -0,50005 | 0,617037 | 0,675816 |
| 1263 | RM25_RS06150 | RM25_1246 | Succinate dehydrogenase/fumarate reductase iron-sulfur su | 1326 | gene | 1439319 | 1440077 | - | 1834,072 | -1,543083354 | 0,639826 | -2,41173 | 0,015877 | 0,062895 | 3008,555779 | -1,493716571 | 0,237241 | -6,2962 | 3,05E-10 | 2,75E-09 | 659,5873 | -1,775680763 | 0,379051 | -4,68454 | 2,81E-06 | 3,37E-05 | 2924,156 | -2,17145 | 0,405752 | -5,35167 | 8,71E-08 | 8,22E-07 | 921,130781 | -2,404549197 | 0,217899 | -11,0351 | 2,59E-28 | 1,35E-26 |
| 1264 | RM25_RS06155 | RM25_1247 | Succinate dehydrogenase flavoprotein subunit | 1327 | gene | 1440074 | 1442158 | - | 4008,763 | -1,539695839 | 0,603027 | -2,55328 | 0,010671 | 0,048973 | 6500,678064 | -1,500262301 | 0,214504 | -6,99411 | 2,67E-12 | 3,31E-11 | 1516,848 | -1,712956661 | 0,262278 | -6,53107 | 6,53E-11 | 2,37E-09 | 6378,816 | -2,09853 | 0,297091 | -7,06357 | 1,62E-12 | 4,16E-11 | 2016,533332 | -2,262963141 | 0,195104 | -11,5987 | 4,18E-31 | 2,9E-29 |
| 1265 | RM25_RS06160 | RM25_1248 | Succinate dehydrogenase cytochrome B subunit, b558 fami | 1328 | gene | 1442155 | 1442910 | - | 3053,495 | -1,596078812 | 0,53365 | -2,99087 | 0,002782 | 0,019809 | 4770,451765 | -1,568916978 | 0,177452 | -8,84135 | 9,46E-19 | 2,4E-17 | 1336,538 | -1,69287108 | 0,191219 | -8,85306 | 8,52E-19 | 1,12E-16 | 4908,269 | -1,86255 | 0,225112 | -8,27388 | 1,3E-16 | 6,64E-15 | 1491,368666 | -1,933992754 | 0,168467 | -11,48 | 1,66E-30 | 1,09E-28 |
| 1266 | RM25_RS06165 | RM25_1249 | NUDIX hydrolase | 1329 | gene | 1443158 | 1443910 | - | 456,4639 | -0,298335298 | 0,143237 | -2,08281 | 0,037269 | 0,104078 | 409,1344176 | -0,173496919 | 0,152344 | -1,13885 | 0,254766 | 0,320088 | 503,7933 | -0,401386189 | 0,206335 | -1,94531 | 0,051737 | 0,103431 | 530,8767 | 0,341655 | 0,158666 | 2,153301 | 0,031295 | 0,058635 | 401,9357149 | 0,170932479 | 0,208486 | 0,819875 | 0,412287 | 0,478464 |
| 1267 | RM25_RS06170 | RM25_1250 | Quinolinate synthase A | 1330 | gene | 1443977 | 1444957 | + | 458,85 | -1,791567103 | 0,670668 | -2,67132 | 0,007555 | 0,038785 | 258,6757591 | -3,637094885 | 0,388778 | -9,35519 | 8,34E-21 | 2,89E-19 | 659,0242 | -1,343404983 | 0,400262 | -3,35632 | 0,00079 | 0,004029 | 746,4304 | 0,911411 | 0,397461 | 2,293079 | 0,021843 | 0,04378 | 202,6676765 | 3,277519119 | 0,433326 | 7,563628 | 3,92E-14 | 4,24E-13 |
| 1268 | RM25_RS06175 | RM25_1251 | L-aspartate oxidase | 1331 | gene | 1444954 | 1446636 | + | 438,0843 | -1,230233649 | 0,645164 | -1,90685 | 0,05654 | 0,134068 | 215,1419524 | -2,548114832 | 0,336851 | -7,56452 | 3,89E-14 | 6,08E-13 | 661,0267 | -0,902945193 | 0,473687 | -1,90621 | 0,056623 | 0,110573 | 642,5157 | 1,16188 | 0,326445 | 3,559188 | 0,000372 | 0,001538 | 258,8061453 | 2,881991067 | 0,489599 | 5,886426 | 3,95E-09 | 1,97E-08 |
| 1269 | RM25_RS06180 | RM25_1252 | Nicotinate-nucleotide pyrophosphatase nadC | 1332 | gene | 1446633 | 1447553 | + | 166,0145 | -0,943250904 | 0,615224 | -1,53318 | 0,125231 | 0,222786 | 85,11301718 | -2,227060251 | 0,381698 | -5,83461 | 5,39E-09 | 4,11E-08 | 246,916 | -0,591320481 | 0,45517 | -1,29912 | 0,193903 | 0,289626 | 229,4014 | 1,000669 | 0,338076 | 2,959895 | 0,003077 | 0,008999 | 111,961967 | 2,719508385 | 0,512833 | 5,30291 | 1,14E-07 | 4,61E-07 |
| 1270 | RM25_RS06185 | RM25_1253 | Glycine--tRNA ligase | 1333 | gene | 1447678 | 1449066 | - | 3812,499 | 0,422153532 | 0,338637 | 1,246624 | 0,212535 | 0,317658 | 4990,608343 | 0,312223948 | 0,172353 | 1,81154 | 0,070057 | 0,104378 | 2634,389 | 0,634877129 | 0,318716 | 1,991981 | 0,046373 | 0,095711 | 3468,593 | -1,16243 | 0,271601 | -4,2799 | 1,87E-05 | 0,000108 | 4274,1143 | -0,80365423 | 0,217035 | -3,70288 | 0,000213 | 0,000511 |
| 1271 | RM25_RS06190 | RM25_1254 | Metal transport system ABC transporter substrate-binding p | 1334 | gene | 1449299 | 1450246 | + | 787,9455 | -0,716447424 | 0,18655 | -3,84052 | 0,000123 | 0,001982 | 704,5981964 | -0,478502957 | 0,196511 | -2,43499 | 0,014892 | 0,027902 | 871,2929 | -0,916387559 | 0,259409 | -3,5326 | 0,000411 | 0,002401 | 1029,629 | 0,407576 | 0,261201 | 1,560394 | 0,118667 | 0,17086 | 585,7615603 | 0,032707774 | 0,198124 | 0,165087 | 0,868875 | 0,893906 |
| 1272 | RM25_RS06195 | RM25_1255 | Zinc uptake system ATP-binding protein zurA | 1335 | gene | 1450341 | 1451129 | + | 300,2397 | -0,743006157 | 0,191034 | -3,88939 | 0,0001 | 0,001729 | 275,8647335 | -0,70885075 | 0,204034 | -3,47417 | 0,000512 | 0,00148 | 324,6146 | -0,775750986 | 0,315484 | -2,45892 | 0,013935 | 0,039242 | 396,179 | 0,19396 | 0,236342 | 0,820672 | 0,411833 | 0,487294 | 220,9255943 | 0,19301586 | 0,309193 | 0,624258 | 0,532458 | 0,592822 |
| 1273 | RM25_RS06200 | RM25_1256 | Metal transport ABC transporter | 1336 | gene | 1451131 | 1452138 | + | 315,5946 | -0,986801548 | 0,396349 | -2,48973 | 0,012784 | 0,05522 | 225,8990132 | -1,347227749 | 0,330533 | -4,07592 | 4,58E-05 | 0,000171 | 405,2901 | -0,80149549 | 0,413333 | -1,9391 | 0,052489 | 0,104314 | 440,8032 | 0,596742 | 0,364867 | 1,635504 | 0,101943 | 0,151215 | 208,5713995 | 1,217498646 | 0,411565 | 2,958215 | 0,003094 | 0,005944 |
| 1274 | RM25_RS06205 | RM25_1257 | Zinc uptake regulator, Fur family | 1337 | gene | 1452104 | 1452616 | + | 129,5863 | -1,051477764 | 0,377921 | -2,78227 | 0,005398 | 0,031164 | 92,53040127 | -1,323034059 | 0,383149 | -3,45305 | 0,000554 | 0,001582 | 166,6422 | -0,915586506 | 0,350625 | -2,6113 | 0,00902 | 0,028122 | 183,3773 | 0,654838 | 0,330787 | 1,979633 | 0,047745 | 0,081925 | 82,85479347 | 1,126068184 | 0,431587 | 2,609132 | 0,009077 | 0,016002 |
| 1275 | RM25_RS06210 | RM25_1258 | Phage integrase | 1338 | gene | 1452869 | 1453543 | + | 958,0946 | 1,125830328 | 0,26885 | 4,187578 | 2,82E-05 | 0,000627 | 917,51713 | 0,507386716 | 0,269636 | 1,88175 | 0,05987 | 0,091218 | 998,6721 | 1,794524422 | 0,203618 | 8,81318 | 1,22E-18 | 1,37E-16 | 637,1872 | -0,81444 | 0,300706 | -2,70843 | 0,00676 | 0,017005 | 1285,999498 | 0,516483834 | 0,165646 | 3,117989 | 0,001821 | 0,003637 |
| 1276 | RM25_RS06215 | RM25_1259 | Ethanolamine utilization protein EutE | 1339 | gene | 1453757 | 1455166 | - | 516,6124 | -0,735471843 | 0,311909 | -2,35797 | 0,018375 | 0,068517 | 389,7867517 | -0,709309049 | 0,285478 | -2,48464 | 0,012968 | 0,02479 | 643,4379 | -0,75343189 | 0,331133 | -2,27531 | 0,022887 | 0,056784 | 677,5254 | 0,674499 | 0,314233 | 2,146491 | 0,031834 | 0,059344 | 381,5224372 | 0,696854416 | 0,323726 | 2,152605 | 0,03135 | 0,049163 |
| 1277 | RM25_RS06220 | RM25_1260 | PduO | 1340 | gene | 1455163 | 1456248 | - | 369,5503 | -0,960915031 | 0,475593 | -2,02046 | 0,043336 | 0,113818 | 217,670634 | -1,456952081 | 0,338678 | -4,30188 | 1,69E-05 | 6,87E-05 | 521,43 | -0,772325744 | 0,361417 | -2,13694 | 0,032603 | 0,073819 | 511,7422 | 0,97296 | 0,360974 | 2,695373 | 0,007031 | 0,01763 | 246,9288286 | 1,729834521 | 0,365851 | 4,728247 | 2,26E-06 | 7,41E-06 |
| 1278 | RM25_RS06225 | RM25_1261 | Propanediol utilization protein PduN | 1341 | gene | 1456251 | 1456523 | - | 91,47549 | -0,817340984 | 0,335721 | -2,43458 | 0,014909 | 0,060797 | 63,78820488 | -0,672107674 | 0,38087 | -1,76466 | 0,07762 | 0,113917 | 119,1628 | -0,903337617 | 0,259816 | -3,47684 | 0,000507 | 0,00282 | 122,3851 | 0,917624 | 0,297349 | 3,086021 | 0,002029 | 0,00635 | 64,94099148 | 0,745657529 | 0,36354 | 2,051099 | 0,040257 | 0,061099 |
| 1279 | RM25_RS11885 | RM25_1262 | Ethanolamine utilization flavoprotein | 1342 | gene | 1456525 | 1457409 | - | 149,1537 | -1,162138559 | 0,547085 | -2,12424 | 0,03365 | 0,097153 | 82,21017785 | -1,900979353 | 0,419826 | -4,52802 | 5,95E-06 | 2,61E-05 | 216,0972 | -0,921482206 | 0,41746 | -2,20735 | 0,027289 | 0,06484 | 216,2623 | 1,047656 | 0,367565 | 2,850263 | 0,004368 | 0,011821 | 90,76273001 | 2,106880878 | 0,498948 | 4,224763 | 2,39E-05 | 6,68E-05 |
| 1280 | RM25_RS06235 | RM25_1263 | Ethanolamine utilization protein EutJ | 1343 | gene | 1457406 | 1458263 | - | 450,5276 | -0,999961703 | 0,395494 | -2,52839 | 0,011459 | 0,051153 | 302,4443063 | -1,122882782 | 0,331308 | -3,38924 | 0,000701 | 0,001955 | 598,6109 | -0,940918084 | 0,368004 | -2,55682 | 0,010564 | 0,031677 | 629,3017 | 0,860825 | 0,340137 | 2,530815 | 0,01138 | 0,026193 | 295,7397783 | 1,109906469 | 0,381219 | 2,911466 | 0,003597 | 0,00681 |
| 1281 | RM25_RS06240 | RM25_1264 | Propanediol utilization protein PduJ | 1344 | gene | 1458410 | 1458688 | - | 151,9178 | 0,034269171 | 0,369282 | 0,092799 | 0,926063 | 0,950245 | 96,69370295 | 0,439807914 | 0,269999 | 1,628927 | 0,103329 | 0,146098 | 207,142 | -0,156401708 | 0,192602 | -0,81205 | 0,416764 | 0,521952 | 156,9615 | 1,346889 | 0,212175 | 6,34801 | 2,18E-10 | 3,74E-09 | 150,7545598 | 0,808492937 | 0,232561 | 3,476477 | 0,000508 | 0,001142 |
| 1282 | RM25_RS06245 | RM25_1265 | Propanediol utilization protein PduK | 1345 | gene | 1458733 | 1459350 | - | 104,1774 | -0,115416947 | 0,23402 | -0,49319 | 0,621876 | 0,709284 | 85,59264543 | 0,22183589 | 0,283561 | 0,782322 | 0,434026 | 0,502455 | 122,7621 | -0,353671447 | 0,228955 | -1,54472 | 0,122414 | 0,201329 | 113,6246 | 0,732543 | 0,230375 | 3,179786 | 0,001474 | 0,004872 | 98,13605491 | 0,214506409 | 0,273812 | 0,783406 | 0,433389 | 0,498534 |
| 1283 | RM25_RS06250 | RM25_1266 | Glycerol dehydratase reactivation factor DhaG | 1346 | gene | 1459367 | 1459711 | - | 64,47537 | -0,821704445 | 0,428413 | -1,91802 | 0,055109 | 0,132317 | 43,02897257 | -1,287678754 | 0,44875 | -2,86948 | 0,004112 | 0,009221 | 85,92177 | -0,613914209 | 0,3815 | -1,60921 | 0,10757 | 0,182179 | 86,59756 | 0,693836 | 0,336846 | 2,0598 | 0,039418 | 0,069908 | 45,7799158 | 1,4387337 | 0,510943 | 2,815842 | 0,004865 | 0,009036 |
| 1284 | RM25_RS06255 | RM25_1267 | Glycerol dehydratase reactivation factor large subunit | 1347 | gene | 1459708 | 1461522 | - | 575,1865 | -1,260609938 | 0,459252 | -2,74492 | 0,006053 | 0,033886 | 369,2045733 | -1,789802255 | 0,304834 | -5,8714 | 4,32E-09 | 3,35E-08 | 781,1685 | -1,039050863 | 0,424554 | -2,44739 | 0,014389 | 0,04009 | 851,3317 | 0,803462 | 0,36869 | 2,179233 | 0,029314 | 0,055425 | 333,9762588 | 1,631479337 | 0,393614 | 4,144869 | 3,4E-05 | 9,29E-05 |
| 1285 | RM25_RS06260 | RM25_1268 | Propanediol dehydratase small subunit | 1348 | gene | 1461558 | 1462091 | - | 127,9102 | -0,232201366 | 0,254699 | -0,91167 | 0,361942 | 0,473552 | 123,9446757 | 0,330829982 | 0,296179 | 1,116995 | 0,263996 | 0,329053 | 131,8758 | -0,769611861 | 0,275658 | -2,79191 | 0,00524 | 0,018656 | 145,1032 | 0,529162 | 0,290758 | 1,819941 | 0,068768 | 0,110357 | 115,9128216 | -0,504581272 | 0,310173 | -1,62677 | 0,103785 | 0,142771 |
| 1286 | RM25_RS06265 | RM25_1269 | Propanediol dehydratase medium subunit | 1349 | gene | 1462100 | 1462783 | - | 182,138 | -0,926765942 | 0,369824 | -2,50596 | 0,012212 | 0,053106 | 129,0263124 | -0,896566564 | 0,398611 | -2,24922 | 0,024498 | 0,042614 | 235,2498 | -0,946365756 | 0,33267 | -2,84476 | 0,004444 | 0,01642 | 250,4927 | 0,81571 | 0,390773 | 2,087426 | 0,03685 | 0,066699 | 123,5149352 | 0,833536801 | 0,384252 | 2,169246 | 0,030064 | 0,047367 |
| 1287 | RM25_RS06270 | RM25_1270 | Glycerol dehydratase large subunit | 1350 | gene | 1462794 | 1464455 | - | 466,4662 | -0,862157985 | 0,317247 | -2,71762 | 0,006575 | 0,03571 | 357,2342598 | -0,835718315 | 0,340099 | -2,45728 | 0,013999 | 0,026525 | 575,6981 | -0,880014106 | 0,299759 | -2,93574 | 0,003327 | 0,013137 | 632,6099 | 0,635934 | 0,378138 | 1,68175 | 0,092617 | 0,139133 | 325,5920007 | 0,660089225 | 0,303492 | 2,174981 | 0,029632 | 0,046748 |
| 1288 | RM25_RS06275 | RM25_1271 | Propanediol utilization protein PduB | 1351 | gene | 1464470 | 1465279 | - | 289,9399 | -1,215794075 | 0,458124 | -2,65385 | 0,007958 | 0,040226 | 184,6116096 | -1,718622092 | 0,362067 | -4,7467 | 2,07E-06 | 9,95E-06 | 395,2683 | -1,006528781 | 0,376404 | -2,67406 | 0,007494 | 0,024669 | 425,1883 | 0,824145 | 0,369508 | 2,230383 | 0,025722 | 0,049817 | 171,7187114 | 1,611918487 | 0,40473 | 3,982701 | 6,81E-05 | 0,000178 |
| 1289 | RM25_RS06280 | RM25_1272 | Propanediol utilization protein PduA | 1352 | gene | 1465294 | 1465578 | - | 110,8169 | -0,565006625 | 0,398544 | -1,41768 | 0,156285 | 0,257778 | 79,53817643 | -0,203803281 | 0,404923 | -0,50331 | 0,614743 | 0,673304 | 142,0956 | -0,777877599 | 0,450394 | -1,72711 | 0,084149 | 0,150599 | 138,8221 | 0,997416 | 0,385241 | 2,589072 | 0,009623 | 0,022981 | 88,14492201 | 0,505313712 | 0,503886 | 1,002834 | 0,315941 | 0,379354 |
| 1290 | RM25_RS12285 | RM25_1273 | Hypothetical protein | 1353 | gene | 1465905 | 1465997 | + | 30,99355 | -0,359558354 | 0,388733 | -0,92495 | 0,354992 | 0,466539 | 20,38852312 | -0,531034525 | 0,547507 | -0,96991 | 0,332089 | 0,399966 | 41,59857 | -0,283435566 | 0,300667 | -0,94269 | 0,34584 | 0,452105 | 36,72589 | 0,836126 | 0,348357 | 2,4002 | 0,016386 | 0,035069 | 26,55031848 | 1,152409069 | 0,49609 | 2,322986 | 0,02018 | 0,033169 |
| 1291 | RM25_RS06285 | RM25_1274 | Transposase IS204/IS1001/IS1096/IS1165 family protein | 1354 | gene | 1466341 | 1467417 | + | 0 | NA | NA | NA | NA | NA | 0 | NA | NA | NA | NA | NA | 0 | NA | NA | NA | NA | NA | 0 | NA | NA | NA | NA | NA | 0 | NA | NA | NA | NA | NA |
| 1292 | RM25_RS06290 | RM25_1275 | putative secreted protein | 1355 | gene | 1467414 | 1467980 | + | 219,3471 | -1,905427802 | 0,817866 | -2,32975 | 0,019819 | 0,07121 | 124,7048944 | -4,996709215 | 0,55125 | -9,06433 | 1,25E-19 | 3,56E-18 | 313,9892 | -1,344941593 | 0,360557 | -3,73018 | 0,000191 | 0,001274 | 363,3059 | 0,826623 | 0,405236 | 2,039853 | 0,041365 | 0,072922 | 91,03629993 | 4,553101302 | 0,581508 | 7,829812 | 4,89E-15 | 6,09E-14 |
| 1293 | RM25_RS06295 | RM25_1276 | putative secreted protein | 1356 | gene | 1467962 | 1468822 | + | 719,2193 | -1,995978102 | 0,820591 | -2,43237 | 0,015 | 0,060854 | 417,4936388 | -5,298843443 | 0,425808 | -12,4442 | 1,5E-35 | 2,95E-33 | 1020,945 | -1,419352951 | 0,370999 | -3,82576 | 0,00013 | 0,00092 | 1203,961 | 0,79809 | 0,458726 | 1,739799 | 0,081894 | 0,125669 | 283,2914932 | 4,74825986 | 0,368501 | 12,88533 | 5,44E-38 | 6,42E-36 |
| 1294 | RM25_RS06300 | RM25_1277 | Ketol-acid reductoisomerase | 1357 | gene | 1468981 | 1470012 | - | 3307,177 | 0,837427714 | 0,555794 | 1,506722 | 0,131882 | 0,230139 | 5003,995968 | 0,5934063 | 0,20493 | 2,895648 | 0,003784 | 0,0086 | 1610,358 | 1,702965396 | 0,32084 | 5,307838 | 1,11E-07 | 1,97E-06 | 2541,624 | -2,45328 | 0,306432 | -8,00595 | 1,19E-15 | 5,26E-14 | 4155,882151 | -1,304569824 | 0,21113 | -6,17899 | 6,45E-10 | 3,72E-09 |
| 1295 | RM25_RS06305 | RM25_1278 | Acetolactate synthase, small subunit | 1358 | gene | 1470057 | 1470584 | - | 741,6564 | 0,292577407 | 0,284809 | 1,027277 | 0,30429 | 0,413855 | 950,852224 | 0,328018735 | 0,15497 | 2,11666 | 0,034289 | 0,05744 | 532,4607 | 0,2310272 | 0,223262 | 1,034781 | 0,300771 | 0,405096 | 708,2863 | -0,84564 | 0,17815 | -4,74682 | 2,07E-06 | 1,48E-05 | 800,0615831 | -0,893678149 | 0,175265 | -5,09902 | 3,41E-07 | 1,28E-06 |
| 1296 | RM25_RS06310 | RM25_1279 | Acetolactate synthase | 1359 | gene | 1470581 | 1472341 | - | 1043,569 | -0,322414455 | 0,147722 | -2,18257 | 0,029068 | 0,087724 | 1080,77666 | -0,085349766 | 0,156414 | -0,54566 | 0,585297 | 0,646459 | 1006,361 | -0,58040588 | 0,219555 | -2,64355 | 0,008204 | 0,026341 | 1223,206 | 0,05189 | 0,19334 | 0,26839 | 0,788399 | 0,8281 | 911,1680653 | -0,382612865 | 0,188353 | -2,03136 | 0,042219 | 0,063707 |
| 1297 | RM25_RS06315 | RM25_1280 | Putative membrane protein | 1360 | gene | 1472640 | 1473665 | - | 343,069 | -0,275901949 | 0,28322 | -0,97416 | 0,329977 | 0,440298 | 308,5701581 | 0,128283804 | 0,186357 | 0,688378 | 0,491215 | 0,557973 | 377,5679 | -0,612075728 | 0,440032 | -1,39098 | 0,164231 | 0,253499 | 393,9176 | 0,57171 | 0,231709 | 2,467359 | 0,013611 | 0,030295 | 305,8996796 | -0,102160068 | 0,411697 | -0,24814 | 0,804023 | 0,841511 |
| 1298 | RM25_RS06320 | RM25_1281 | Imidazole glycerol phosphate synthase subunit HisF | 1361 | gene | 1473850 | 1474614 | - | 1121,958 | 0,691609486 | 0,422415 | 1,637274 | 0,101573 | 0,194958 | 1603,301333 | 0,828248961 | 0,219054 | 3,781022 | 0,000156 | 0,000509 | 640,6142 | 0,365522014 | 0,316918 | 1,153366 | 0,24876 | 0,349213 | 912,9564 | -1,09729 | 0,311544 | -3,52211 | 0,000428 | 0,001737 | 1358,292976 | -1,523904612 | 0,216952 | -7,02415 | 2,15E-12 | 1,77E-11 |
| 1299 | RM25_RS06325 | RM25_1282 | Voltage-dependent potassium channel beta subunit | 1362 | gene | 1474699 | 1475715 | + | 1639,798 | 0,387765899 | 0,195767 | 1,980757 | 0,047619 | 0,120867 | 1473,598681 | 0,362286676 | 0,166442 | 2,176652 | 0,029507 | 0,05047 | 1805,998 | 0,409472788 | 0,315522 | 1,297761 | 0,194369 | 0,289771 | 1501,111 | 0,212782 | 0,259213 | 0,820875 | 0,411717 | 0,487294 | 1814,80467 | 0,296918303 | 0,224734 | 1,3212 | 0,186435 | 0,239339 |
| 1300 | RM25_RS06330 | RM25_1283 | Antibiotic biosynthesis monooxygenase | 1363 | gene | 1475785 | 1476162 | - | 1733,229 | 0,367617426 | 0,292578 | 1,256478 | 0,208943 | 0,31388 | 1593,958126 | 0,707440018 | 0,220425 | 3,209437 | 0,00133 | 0,003445 | 1872,499 | 0,087619793 | 0,446259 | 0,196343 | 0,844342 | 0,887255 | 1595,896 | 0,536082 | 0,375037 | 1,429411 | 0,152886 | 0,210241 | 1904,579166 | -0,058261207 | 0,308869 | -0,18863 | 0,850385 | 0,878854 |
| 1301 | RM25_RS06335 | RM25_1284 | ATP-dependent DNA helicase helY | 1364 | gene | 1476174 | 1478975 | - | 1600,979 | 0,038529811 | 0,143717 | 0,268095 | 0,788626 | 0,850705 | 1686,399842 | 0,350537547 | 0,107931 | 3,247789 | 0,001163 | 0,003077 | 1515,559 | -0,306270788 | 0,178403 | -1,71674 | 0,086027 | 0,153262 | 1666,664 | 0,115884 | 0,133173 | 0,870177 | 0,384204 | 0,461084 | 1592,52313 | -0,485647797 | 0,152371 | -3,18727 | 0,001436 | 0,002936 |
| 1302 | RM25_RS06340 | RM25_1285 | Phosphatase YidA | 1365 | gene | 1479111 | 1479968 | + | 293,7516 | -0,80425909 | 0,325489 | -2,47093 | 0,013476 | 0,057129 | 213,5432779 | -0,755488025 | 0,309558 | -2,44053 | 0,014666 | 0,027521 | 373,9599 | -0,83448327 | 0,305903 | -2,72793 | 0,006373 | 0,021771 | 393,2419 | 0,765545 | 0,297612 | 2,572292 | 0,010103 | 0,023932 | 210,3583041 | 0,757455945 | 0,354552 | 2,136374 | 0,032649 | 0,050671 |
| 1303 | RM25_RS06345 | RM25_1286 | Diacylglycerol kinase catalytic region | 1366 | gene | 1480009 | 1481022 | - | 507,1321 | -0,96672815 | 0,348855 | -2,77115 | 0,005586 | 0,031802 | 359,7259302 | -0,043599337 | 0,18453 | -0,23627 | 0,813221 | 0,852026 | 654,5383 | -1,55620175 | 0,324143 | -4,80097 | 1,58E-06 | 2,07E-05 | 703,4593 | 1,364048 | 0,246498 | 5,533718 | 3,14E-08 | 3,21E-07 | 337,4608177 | -0,095201415 | 0,294731 | -0,32301 | 0,746687 | 0,790629 |
| 1304 | RM25_RS06350 | RM25_1287 | Sec-independent protein translocase, TatC subunit | 1367 | gene | 1481038 | 1481910 | - | 1171,47 | 0,126672908 | 0,122523 | 1,033872 | 0,301196 | 0,411071 | 1205,882738 | 0,052339711 | 0,111399 | 0,469841 | 0,638468 | 0,695733 | 1137,058 | 0,206448696 | 0,249215 | 0,828396 | 0,407446 | 0,51246 | 1182,867 | -0,22909 | 0,184645 | -1,24068 | 0,214723 | 0,280544 | 1196,708483 | -0,025266551 | 0,155633 | -0,16235 | 0,871033 | 0,895344 |
| 1305 | RM25_RS12290 | RM25_1288 | Hypothetical protein | 1368 | gene | 1481929 | 1482237 | - | 1015,807 | 0,869000329 | 0,362724 | 2,395763 | 0,016586 | 0,064439 | 1191,428304 | 0,368251792 | 0,220208 | 1,672291 | 0,094467 | 0,135355 | 840,185 | 1,693220695 | 0,369959 | 4,576783 | 4,72E-06 | 5,38E-05 | 766,5247 | -1,44184 | 0,343595 | -4,19635 | 2,71E-05 | 0,000152 | 1282,381797 | -0,083126526 | 0,246776 | -0,33685 | 0,736229 | 0,781155 |
| 1306 | RM25_RS06360 | RM25_1289 | Protein pafC | 1369 | gene | 1482339 | 1483298 | - | 1032,577 | 0,364401248 | 0,162404 | 2,243801 | 0,024845 | 0,080661 | 1134,191819 | 0,312246269 | 0,157941 | 1,976977 | 0,048044 | 0,076462 | 930,9625 | 0,429442571 | 0,245775 | 1,747297 | 0,080586 | 0,146108 | 957,4292 | -0,4072 | 0,221424 | -1,83899 | 0,065917 | 0,106714 | 1137,678316 | -0,245879172 | 0,177348 | -1,38642 | 0,165618 | 0,216868 |
| 1307 | RM25_RS06365 | RM25_1290 | Protein pafB | 1370 | gene | 1483285 | 1484253 | - | 1870,225 | 0,346239328 | 0,185179 | 1,869751 | 0,061518 | 0,142017 | 1920,42645 | 0,571079779 | 0,165946 | 3,441355 | 0,000579 | 0,001646 | 1820,024 | 0,114710585 | 0,309881 | 0,370176 | 0,711252 | 0,784841 | 1742,15 | 0,124095 | 0,292775 | 0,423859 | 0,671669 | 0,723548 | 2047,95601 | -0,294203069 | 0,185193 | -1,58863 | 0,112143 | 0,152787 |
| 1308 | RM25_RS06370 | RM25_1291 | Glutamyl-Q tRNA(Asp) synthetase | 1371 | gene | 1484322 | 1485281 | + | 307,3472 | -0,69476691 | 0,321111 | -2,16363 | 0,030493 | 0,091672 | 220,2441655 | -0,705010463 | 0,268118 | -2,62947 | 0,008552 | 0,017316 | 394,4503 | -0,692067692 | 0,303337 | -2,28151 | 0,022518 | 0,056322 | 398,7853 | 0,770228 | 0,275805 | 2,792653 | 0,005228 | 0,013691 | 230,6745417 | 0,846839506 | 0,310151 | 2,730412 | 0,006326 | 0,01146 |
| 1309 | RM25_RS06375 | RM25_1292 | Peptidyl-prolyl cis-trans isomerase | 1372 | gene | 1485424 | 1486482 | - | 2720,878 | 0,548681328 | 0,294214 | 1,864904 | 0,062195 | 0,142462 | 3467,43774 | 0,672860944 | 0,178055 | 3,778943 | 0,000157 | 0,000513 | 1974,318 | 0,33650183 | 0,286134 | 1,17603 | 0,239583 | 0,338954 | 2344,128 | -0,66668 | 0,262183 | -2,54281 | 0,010997 | 0,025486 | 3165,556096 | -0,966963717 | 0,189742 | -5,09619 | 3,47E-07 | 1,3E-06 |
| 1310 | NA | RM25_1293 | Peptidyl-prolyl cis-trans isomerase | 1373 | gene | 1486568 | 1486978 | - | 536,5713 | 0,420478379 | 0,153727 | 2,735222 | 0,006234 | 0,034572 | 591,1542331 | 0,491862983 | 0,185114 | 2,657085 | 0,007882 | 0,016199 | 481,9884 | 0,336007128 | 0,205365 | 1,636145 | 0,101809 | 0,174885 | 486,0066 | -0,26363 | 0,216534 | -1,21751 | 0,22341 | 0,290606 | 601,6558916 | -0,371262709 | 0,168399 | -2,20466 | 0,027478 | 0,04376 |
| 1311 | RM25_RS06385 | RM25_1294 | Pseudouridylate synthase | 1374 | gene | 1487275 | 1488057 | - | 1096,508 | 0,381555014 | 0,260413 | 1,465194 | 0,142868 | 0,243485 | 1369,757707 | 0,354204682 | 0,145988 | 2,426267 | 0,015255 | 0,028537 | 823,2591 | 0,42897321 | 0,225201 | 1,904848 | 0,0568 | 0,110825 | 1011,011 | -0,83599 | 0,179556 | -4,65588 | 3,23E-06 | 2,22E-05 | 1215,332165 | -0,715096879 | 0,164657 | -4,34294 | 1,41E-05 | 4,06E-05 |
| 1312 | RM25_RS06390 | RM25_1295 | Transcriptional regulator | 1375 | gene | 1488050 | 1488805 | - | 566,1515 | 0,341511757 | 0,245829 | 1,389223 | 0,164765 | 0,267091 | 700,0943532 | 0,460494917 | 0,173289 | 2,657389 | 0,007875 | 0,016199 | 432,2086 | 0,15549608 | 0,184095 | 0,844649 | 0,398307 | 0,504465 | 529,8744 | -0,58716 | 0,200696 | -2,92561 | 0,003438 | 0,009872 | 620,3613304 | -0,842394231 | 0,14753 | -5,71 | 1,13E-08 | 5,32E-08 |
| 1313 | RM25_RS06395 | RM25_1296 | Segregation and condensation protein A | 1376 | gene | 1488802 | 1489716 | - | 533,3162 | -0,201762205 | 0,174519 | -1,1561 | 0,247639 | 0,35584 | 579,7719375 | -0,011432778 | 0,138684 | -0,08244 | 0,934299 | 0,949275 | 486,8605 | -0,427511438 | 0,274417 | -1,55789 | 0,11926 | 0,197537 | 602,2246 | -0,12421 | 0,207848 | -0,59759 | 0,550113 | 0,615093 | 487,5212066 | -0,47813781 | 0,223542 | -2,13892 | 0,032442 | 0,050439 |
| 1314 | RM25_RS06400 | RM25_1297 | Hypothetical protein | 1377 | gene | 1489697 | 1489972 | - | 192,0289 | 0,379979167 | 0,415125 | 0,915338 | 0,360014 | 0,471681 | 274,2499244 | 0,362566725 | 0,217027 | 1,670607 | 0,094799 | 0,135749 | 109,8078 | 0,441690192 | 0,228296 | 1,934726 | 0,053024 | 0,105054 | 177,7227 | -1,42064 | 0,246292 | -5,7681 | 8,02E-09 | 9,04E-08 | 213,1808205 | -1,290602261 | 0,206354 | -6,25433 | 3,99E-10 | 2,34E-09 |
| 1315 | RM25_RS06405 | RM25_1298 | Initiation inhibitor protein | 1378 | gene | 1489957 | 1490853 | - | 1575,703 | 0,104255319 | 0,30107 | 0,346282 | 0,729131 | 0,803066 | 2030,853459 | 0,27583053 | 0,152498 | 1,808749 | 0,07049 | 0,104776 | 1120,552 | -0,203471074 | 0,238829 | -0,85195 | 0,394239 | 0,500389 | 1612,14 | -0,67064 | 0,213083 | -3,14729 | 0,001648 | 0,00535 | 1600,417304 | -1,107286915 | 0,168963 | -6,55341 | 5,62E-11 | 3,8E-10 |
| 1316 | RM25_RS06410 | RM25_1299 | Tyrosine recombinase XerD | 1379 | gene | 1490974 | 1491969 | - | 365,2563 | -0,092821822 | 0,304239 | -0,30509 | 0,760294 | 0,826193 | 391,3778543 | 0,41142193 | 0,208412 | 1,97408 | 0,048373 | 0,076917 | 339,1347 | -0,682799301 | 0,427487 | -1,59724 | 0,110212 | 0,185786 | 396,8437 | 0,248781 | 0,239754 | 1,037652 | 0,299432 | 0,374013 | 348,2125302 | -0,773885616 | 0,406812 | -1,90232 | 0,05713 | 0,083326 |
| 1317 | RM25_RS06415 | RM25_1300 | ADP-ribose pyrophosphatase (ADPRase) | 1380 | gene | 1491962 | 1492624 | - | 1267,757 | 1,270986758 | 0,516979 | 2,458488 | 0,013952 | 0,057999 | 1986,400697 | 1,392152532 | 0,20022 | 6,953107 | 3,57E-12 | 4,39E-11 | 549,1128 | 0,865910157 | 0,223024 | 3,882586 | 0,000103 | 0,000764 | 792,8494 | -1,55612 | 0,230385 | -6,75441 | 1,43E-11 | 3,16E-10 | 1758,827163 | -2,034168128 | 0,181529 | -11,2058 | 3,82E-29 | 2,2E-27 |
| 1318 | RM25_RS06420 | RM25_1301 | CTP synthase | 1381 | gene | 1492617 | 1494371 | - | 5378,979 | 1,16033759 | 0,394684 | 2,939918 | 0,003283 | 0,021921 | 7395,13051 | 1,05280017 | 0,181383 | 5,804297 | 6,46E-09 | 4,9E-08 | 3362,828 | 1,409159379 | 0,288669 | 4,882814 | 1,05E-06 | 1,48E-05 | 3546,357 | -1,44261 | 0,24932 | -5,78616 | 7,2E-09 | 8,24E-08 | 7278,49973 | -1,046917393 | 0,209122 | -5,00626 | 5,55E-07 | 2,03E-06 |
| 1319 | RM25_RS06425 | RM25_1302 | DNA repair protein RecN | 1382 | gene | 1494458 | 1496149 | - | 936,015 | -0,077047139 | 0,15196 | -0,50702 | 0,61214 | 0,700735 | 1013,351394 | 0,102959023 | 0,120423 | 0,854978 | 0,392563 | 0,461713 | 858,6786 | -0,288507111 | 0,23681 | -1,21831 | 0,223107 | 0,319869 | 1015,034 | -0,11344 | 0,167455 | -0,67745 | 0,498123 | 0,565312 | 894,6037514 | -0,445720084 | 0,196079 | -2,27317 | 0,023016 | 0,037185 |
| 1320 | RM25_RS06430 | RM25_1303 | putative inorganic polyphosphate/ATP-NAD kinase | 1383 | gene | 1496143 | 1497066 | - | 1523,78 | 0,365139952 | 0,268888 | 1,357961 | 0,174476 | 0,278241 | 1846,065469 | 0,605877855 | 0,185914 | 3,258911 | 0,001118 | 0,002969 | 1201,495 | 0,006173786 | 0,307944 | 0,020048 | 0,984005 | 0,991154 | 1411,306 | -0,33801 | 0,305646 | -1,10588 | 0,268778 | 0,341331 | 1680,443155 | -0,902511463 | 0,175952 | -5,1293 | 2,91E-07 | 1,1E-06 |
| 1321 | RM25_RS06435 | RM25_1304 | Hypothetical protein | 1384 | gene | 1497131 | 1497370 | - | 159,7578 | -0,064367259 | 0,160353 | -0,40141 | 0,688118 | 0,766128 | 169,7747959 | 0,190931342 | 0,205146 | 0,930712 | 0,352003 | 0,420926 | 149,7409 | -0,341828333 | 0,233322 | -1,46505 | 0,142907 | 0,227128 | 171,9329 | 0,01296 | 0,21588 | 0,060034 | 0,952128 | 0,963162 | 153,6040111 | -0,460163121 | 0,219104 | -2,10021 | 0,035711 | 0,054798 |
| 1322 | RM25_RS06440 | RM25_1305 | Cobalt transport protein | 1385 | gene | 1497443 | 1498051 | - | 291,7384 | -0,720835145 | 0,221618 | -3,2526 | 0,001144 | 0,010957 | 259,8894593 | -1,040043485 | 0,199344 | -5,23741 | 1,63E-07 | 9,48E-07 | 323,5874 | -0,480903758 | 0,297966 | -1,61396 | 0,106537 | 0,180783 | 382,8024 | 0,044744 | 0,24481 | 0,18277 | 0,854978 | 0,885795 | 216,2758175 | 0,668816291 | 0,267989 | 2,495681 | 0,012572 | 0,021534 |
| 1323 | RM25_RS06445 | RM25_1306 | Biotin ABC transport system ATP-binding protein | 1386 | gene | 1498048 | 1498776 | - | 418,1989 | -0,364521913 | 0,147056 | -2,4788 | 0,013183 | 0,056597 | 441,337462 | -0,665132289 | 0,165418 | -4,02093 | 5,8E-05 | 0,00021 | 395,0604 | -0,042414561 | 0,186576 | -0,22733 | 0,820166 | 0,870068 | 498,4979 | -0,49166 | 0,170614 | -2,88169 | 0,003956 | 0,010917 | 358,0629105 | 0,18541665 | 0,162336 | 1,142176 | 0,253381 | 0,311376 |
| 1324 | RM25_RS06450 | RM25_1307 | BioY family protein | 1387 | gene | 1498790 | 1499419 | - | 479,7508 | -0,41751128 | 0,163766 | -2,54943 | 0,01079 | 0,04913 | 482,2974314 | -0,77996382 | 0,153458 | -5,08259 | 3,72E-07 | 2,01E-06 | 477,2042 | -0,067840868 | 0,230949 | -0,29375 | 0,76895 | 0,831383 | 580,4332 | -0,38241 | 0,210062 | -1,82048 | 0,068687 | 0,110357 | 402,087427 | 0,385751167 | 0,14905 | 2,588069 | 0,009652 | 0,016914 |
| 1325 | RM25_RS06455 | RM25_1308 | HAD-superfamily hydrolase, subfamily IIA | 1388 | gene | 1499608 | 1500639 | - | 844,0268 | -0,105371495 | 0,218499 | -0,48225 | 0,629627 | 0,716231 | 968,0544304 | -0,37033255 | 0,149233 | -2,48158 | 0,01308 | 0,024963 | 719,9992 | 0,247846243 | 0,274357 | 0,903371 | 0,366329 | 0,472857 | 927,063 | -0,79133 | 0,213204 | -3,71162 | 0,000206 | 0,000921 | 795,928759 | -0,123925633 | 0,188193 | -0,6585 | 0,510215 | 0,571295 |
| 1326 | RM25_RS12295 | RM25_1309 | Hypothetical protein | 1389 | gene | 1500636 | 1501748 | - | 1549,849 | 0,48439017 | 0,389868 | 1,242448 | 0,214071 | 0,319346 | 1974,030316 | 0,023762276 | 0,183529 | 0,129474 | 0,896982 | 0,922823 | 1125,668 | 1,376740794 | 0,327458 | 4,204333 | 2,62E-05 | 0,000233 | 1378,224 | -1,70045 | 0,288031 | -5,90372 | 3,55E-09 | 4,34E-08 | 1768,071323 | -0,308298008 | 0,216846 | -1,42174 | 0,155102 | 0,204346 |

| | | | | | | | | | | | | | | | | | | | | | | | | | | | | | | | | | | | | | | |
|---|---|---|---|---|---|---|---|---|---|---|---|---|---|---|---|---|---|---|---|---|---|---|---|---|---|---|---|---|---|---|---|---|---|---|---|---|---|---|
| 1327 | RM25_RS12300 | RM25_1310 | Hypothetical protein | 1390 | gene | 1501823 | 1502881 | - | 944,103 | 0,379247196 | 0,250668 | 1,512944 | 0,130294 | 0,228857 | 1095,906077 | 0,174150813 | 0,201351 | 0,864912 | 0,387087 | 0,455954 | 792,2999 | 0,671285557 | 0,307069 | 2,186109 | 0,028808 | 0,067428 | 871,3282 | -0,80728 | 0,314896 | -2,56363 | 0,010358 | 0,024366 | 1044,411993 | -0,26971993 | 0,165672 | -1,62804 | 0,103517 | 0,142518 |
| 1328 | RM25_RS06465 | RM25_1311 | 5S ribosomal RNA | 1392 | rRNA | 1502835 | 1502950 | - | 0 | NA | NA | NA | NA | NA | 0 | NA | NA | NA | NA | NA | 0 | NA | NA | NA | NA | NA | 0 | NA | NA | NA | NA | NA | 0 | NA | NA | NA | NA | NA |
| 1329 | RM25_RS06465 | RM25_1312 | 23S ribosomal RNA | 1395 | rRNA | 1503095 | 1506220 | - | 0 | NA | NA | NA | NA | NA | 0 | NA | NA | NA | NA | NA | 0 | NA | NA | NA | NA | NA | 0 | NA | NA | NA | NA | NA | 0 | NA | NA | NA | NA | NA |
| 1330 | NA | RM25_1313 | Hypothetical protein | 1397 | gene | 1505812 | 1506807 | + | 0 | NA | NA | NA | NA | NA | 0 | NA | NA | NA | NA | NA | 0 | NA | NA | NA | NA | NA | 0 | NA | NA | NA | NA | NA | 0 | NA | NA | NA | NA | NA |
| 1331 | RM25_RS06475 | RM25_1314 | 16S ribosomal RNA | 1399 | rRNA | 1506567 | 1508073 | - | 0 | NA | NA | NA | NA | NA | 0 | NA | NA | NA | NA | NA | 0 | NA | NA | NA | NA | NA | 0 | NA | NA | NA | NA | NA | 0 | NA | NA | NA | NA | NA |
| 1332 | RM25_RS06480 | RM25_1315 | Tyrosine--tRNA ligase | 1401 | gene | 1508760 | 1510028 | - | 3421,072 | 0,388149076 | 0,356498 | 1,088782 | 0,27625 | 0,385279 | 4608,053159 | 0,457764206 | 0,19005 | 2,408655 | 0,016011 | 0,029716 | 2234,09 | 0,24702054 | 0,307352 | 0,803706 | 0,421567 | 0,52601 | 3150,919 | -0,97718 | 0,295878 | -3,30266 | 0,000958 | 0,003389 | 3801,202478 | -1,15145974 | 0,195078 | -5,90257 | 3,58E-09 | 1,82E-08 |
| 1333 | RM25_RS06485 | RM25_1316 | Argininosuccinate lyase | 1402 | gene | 1510156 | 1511577 | - | 1343,089 | 0,195097023 | 0,196344 | 0,993647 | 0,320395 | 0,429807 | 1576,027754 | 0,165426502 | 0,162958 | 1,01515 | 0,310035 | 0,37687 | 1110,15 | 0,238434293 | 0,184208 | 1,294374 | 0,195536 | 0,291143 | 1326,921 | -0,60058 | 0,179842 | -3,33948 | 0,000839 | 0,00303 | 1404,724341 | -0,482527091 | 0,141883 | -3,40087 | 0,000672 | 0,00148 |
| 1334 | RM25_RS06490 | RM25_1317 | Arginine repressor | 1403 | gene | 1511581 | 1512135 | - | 282,8457 | 0,132841244 | 0,14621 | 0,908564 | 0,363581 | 0,474507 | 313,024816 | 0,212817213 | 0,174285 | 1,221085 | 0,222054 | 0,28491 | 252,6665 | 0,039548168 | 0,205553 | 0,192399 | 0,84743 | 0,889626 | 284,9735 | -0,27718 | 0,194543 | -1,42477 | 0,154225 | 0,211711 | 290,4193929 | -0,392734955 | 0,171961 | -2,28386 | 0,022379 | 0,036353 |
| 1335 | RM25_RS06495 | RM25_1318 | Acetylornithine transaminase | 1404 | gene | 1512132 | 1513355 | - | 785,3337 | 0,217251087 | 0,16973 | 1,279981 | 0,200552 | 0,303669 | 884,8742713 | 0,206223837 | 0,16267 | 1,267741 | 0,20489 | 0,265344 | 685,7931 | 0,231935108 | 0,224119 | 1,034874 | 0,300728 | 0,405096 | 768,7458 | -0,4463 | 0,204885 | -2,17831 | 0,029383 | 0,055493 | 827,5795635 | -0,365342556 | 0,161138 | -2,26727 | 0,023374 | 0,037709 |
| 1336 | RM25_RS06500 | RM25_1319 | Acetylglutamate kinase | 1405 | gene | 1513348 | 1514259 | - | 497,052 | 0,437176175 | 0,251417 | 1,73885 | 0,082061 | 0,169516 | 606,899663 | 0,337312033 | 0,201574 | 1,673387 | 0,094251 | 0,135128 | 387,2042 | 0,595085049 | 0,207863 | 2,862873 | 0,004198 | 0,015707 | 448,3752 | -0,85672 | 0,168928 | -5,0715 | 3,95E-07 | 3,26E-06 | 560,0620428 | -0,552469442 | 0,215186 | -2,56741 | 0,010246 | 0,017863 |
| 1337 | RM25_RS06505 | RM25_1320 | Arginine biosynthesis bifunctional protein ArgJ | 1406 | gene | 1514256 | 1515407 | - | 696,2263 | 0,151164422 | 0,122903 | 1,229951 | 0,218716 | 0,324222 | 753,4572714 | 0,212656856 | 0,158323 | 1,343187 | 0,179211 | 0,236639 | 638,9954 | 0,080383953 | 0,172541 | 0,465884 | 0,641298 | 0,722879 | 697,6212 | -0,23109 | 0,142854 | -1,61766 | 0,105737 | 0,155494 | 718,341898 | -0,310835893 | 0,168467 | -1,84509 | 0,065025 | 0,093682 |
| 1338 | RM25_RS06510 | RM25_1321 | N-acetyl-gamma-glutamyl-phosphate reductase | 1407 | gene | 1515404 | 1516435 | - | 933,7357 | -0,056960038 | 0,106679 | -0,53394 | 0,593383 | 0,683579 | 955,6775532 | -0,083844711 | 0,181898 | -0,46094 | 0,644839 | 0,700819 | 911,7939 | -0,028789308 | 0,163704 | -0,17586 | 0,860402 | 0,898524 | 1004,828 | -0,1523 | 0,138881 | -1,09663 | 0,272805 | 0,345885 | 897,4267755 | -0,047461689 | 0,182804 | -0,25963 | 0,795148 | 0,833702 |
| 1339 | RM25_RS06515 | RM25_1322 | Aminotransferase AlaT | 1408 | gene | 1516600 | 1517814 | + | 1521,551 | 1,267384209 | 0,160783 | 7,882594 | 3,21E-15 | 9,45E-13 | 1583,001966 | 1,173646475 | 0,141683 | 8,283583 | 1,2E-16 | 2,39E-15 | 1460,101 | 1,371551758 | 0,288896 | 4,74757 | 2,06E-06 | 2,57E-05 | 944,308 | -0,31719 | 0,149535 | -2,12114 | 0,03391 | 0,062588 | 2101,264745 | -0,076389553 | 0,230903 | -0,33083 | 0,740773 | 0,785426 |
| 1340 | RM25_RS06520 | RM25_1323 | Phenylalanine--tRNA ligase beta subunit | 1409 | gene | 1517942 | 1520470 | - | 3254,572 | 0,282971531 | 0,277149 | 1,02101 | 0,30725 | 0,416439 | 4142,973852 | 0,243204648 | 0,154366 | 1,575502 | 0,115141 | 0,161156 | 2366,17 | 0,353562414 | 0,196312 | 1,80102 | 0,0717 | 0,13329 | 3116,637 | -0,92825 | 0,186477 | -4,9778 | 6,43E-07 | 5,05E-06 | 3501,490216 | -0,76992512 | 0,126271 | -6,09738 | 1,08E-09 | 5,98E-09 |
| 1341 | RM25_RS06525 | RM25_1324 | Phenylalanine--tRNA ligase alpha subunit | 1410 | gene | 1520467 | 1521561 | - | 1579,434 | 0,326213153 | 0,272124 | 1,198768 | 0,230618 | 0,33825 | 1998,490464 | 0,394847791 | 0,163439 | 2,415876 | 0,015697 | 0,029225 | 1160,378 | 0,210710438 | 0,216486 | 0,973319 | 0,330395 | 0,436025 | 1487,163 | -0,74052 | 0,207571 | -3,56758 | 0,00036 | 0,001503 | 1722,502172 | -0,878130362 | 0,15302 | -5,73868 | 9,54E-09 | 4,56E-08 |
| 1342 | RM25_RS12750 | RM25_1325 | Hypothetical protein | 1411 | gene | 1521717 | 1521893 | - | 123,3881 | 0,678711953 | 0,380382 | 1,784289 | 0,074377 | 0,160537 | 169,5895382 | 0,855786775 | 0,222305 | 3,849601 | 0,000118 | 0,000396 | 77,18656 | 0,318545665 | 0,333542 | 0,955039 | 0,339558 | 0,445623 | 100,5758 | -0,8663 | 0,231436 | -3,74315 | 0,000182 | 0,00083 | 149,1502553 | -1,35056097 | 0,318541 | -4,23983 | 2,24E-05 | 6,28E-05 |
| 1343 | RM25_RS06530 | RM25_1326 | Putative 23S rRNA methyltransferase tsnR | 1412 | gene | 1521893 | 1522753 | - | 1317,753 | 0,779849559 | 0,478545 | 1,629627 | 0,10318 | 0,196601 | 1949,478265 | 0,648559517 | 0,201709 | 3,215327 | 0,001303 | 0,003396 | 686,0286 | 1,175526578 | 0,21627 | 5,435466 | 5,47E-08 | 1,01E-06 | 1034,185 | -1,90809 | 0,239153 | -7,97854 | 1,48E-15 | 6,23E-14 | 1633,389896 | -1,335127703 | 0,152328 | -8,76479 | 1,87E-18 | 3,64E-17 |
| 1344 | RM25_RS06535 | RM25_1327 | 50S ribosomal protein L20 | 1413 | gene | 1522764 | 1523135 | - | 3441,92 | 1,081233278 | 0,746851 | 1,447723 | 0,147694 | 0,248123 | 4805,809036 | 0,405248455 | 0,226131 | 1,792094 | 0,073118 | 0,108185 | 2078,032 | 3,777092264 | 0,612188 | 6,169823 | 6,84E-10 | 2E-08 | 2367,808 | -3,91552 | 0,601462 | -6,51 | 7,51E-11 | 1,39E-09 | 4574,347151 | -0,515999872 | 0,24896 | -2,07262 | 0,038208 | 0,058288 |
| 1345 | RM25_RS06540 | RM25_1328 | 50S ribosomal protein L35 | 1414 | gene | 1523181 | 1523375 | - | 2688,412 | 1,105054012 | 0,699166 | 1,580532 | 0,113985 | 0,208718 | 3765,788097 | 0,475255763 | 0,239012 | 1,98842 | 0,046765 | 0,074984 | 1611,037 | 3,504498637 | 0,516997 | 6,778563 | 1,21E-11 | 5,02E-10 | 1827,804 | -3,64145 | 0,488491 | -7,4545 | 9,02E-14 | 2,73E-12 | 3589,898345 | -0,583889636 | 0,282208 | -2,069 | 0,038546 | 0,058766 |
| 1346 | RM25_RS06545 | RM25_1329 | Translation initiation factor IF-3 | 1415 | gene | 1523443 | 1524171 | - | 5759,655 | 0,733391802 | 0,56368 | 1,301078 | 0,193232 | 0,297342 | 8198,802679 | 0,358770172 | 0,244234 | 1,468959 | 0,141844 | 0,194263 | 3320,507 | 1,81763192 | 0,544223 | 3,339864 | 0,000838 | 0,004249 | 4623,018 | -2,3351 | 0,538722 | -4,33451 | 1,46E-05 | 8,67E-05 | 7041,998519 | -0,849956002 | 0,251597 | -3,37824 | 0,00073 | 0,001601 |
| 1347 | RM25_RS06550 | RM25_1330 | Hypothetical protein | 1416 | gene | 1524477 | 1524959 | + | 434,5559 | 0,225802554 | 0,172967 | 1,305467 | 0,191734 | 0,295604 | 436,9050027 | 0,610674113 | 0,207058 | 2,949295 | 0,003185 | 0,007411 | 432,2067 | -0,148940197 | 0,178904 | -0,83252 | 0,405117 | 0,510894 | 420,9772 | 0,342338 | 0,207906 | 1,646597 | 0,099641 | 0,148266 | 459,8154071 | -0,368775425 | 0,169252 | -2,17886 | 0,029342 | 0,04636 |
| 1348 | RM25_RS06555 | RM25_1331 | ATP phosphoribosyltransferase | 1417 | gene | 1525023 | 1525886 | - | 1509,705 | 0,553119164 | 0,211133 | 2,619762 | 0,008799 | 0,042939 | 1781,222779 | 0,5353374 | 0,167627 | 3,193622 | 0,001405 | 0,003609 | 1238,186 | 0,579568612 | 0,223976 | 2,587634 | 0,009664 | 0,029504 | 1294,637 | -0,61004 | 0,215538 | -2,8303 | 0,00465 | 0,012441 | 1759,197413 | -0,517838631 | 0,128904 | -4,01724 | 5,89E-05 | 0,000155 |
| 1349 | RM25_RS06560 | RM25_1332 | Phosphoribosyl-ATP diphosphatase | 1418 | gene | 1525953 | 1526219 | - | 1577,348 | 0,769644247 | 0,33415 | 2,303292 | 0,021262 | 0,073808 | 1974,679623 | 0,768834815 | 0,212886 | 3,611492 | 0,000304 | 0,000933 | 1180,016 | 0,772849117 | 0,425796 | 1,81507 | 0,069513 | 0,129787 | 1240,787 | -0,79446 | 0,398211 | -1,99508 | 0,046034 | 0,079606 | 1944,57141 | -0,76086663 | 0,253657 | -2,99959 | 0,002703 | 0,005262 |
| 1350 | RM25_RS06565 | RM25_1333 | ATP-dependent zinc metalloprotease FtsH | 1419 | gene | 1526392 | 1528542 | + | 3913,623 | 0,53675693 | 0,14299 | 3,753806 | 0,000174 | 0,002598 | 4258,53118 | 0,494080564 | 0,135685 | 3,64138 | 0,000271 | 0,000844 | 3568,716 | 0,588338866 | 0,223277 | 2,635022 | 0,008413 | 0,026833 | 3381,917 | -0,36561 | 0,18305 | -1,9973 | 0,045792 | 0,079304 | 4535,305304 | -0,229636767 | 0,15567 | -1,47515 | 0,140171 | 0,186552 |
| 1351 | RM25_RS06570 | RM25_1334 | L-cysteine:1D-myo-inositol 2-amino-2-deoxy-alpha-D-glucop | 1420 | gene | 1528652 | 1529908 | - | 2478,531 | 0,64736339 | 0,314733 | 2,056863 | 0,039699 | 0,108203 | 3197,38185 | 0,604912681 | 0,199132 | 3,037752 | 0,002383 | 0,005727 | 1759,679 | 0,726657384 | 0,278713 | 2,607184 | 0,009129 | 0,02828 | 2055,721 | -0,98943 | 0,291716 | -3,39176 | 0,000694 | 0,002606 | 2963,047844 | -0,828936654 | 0,173696 | -4,77234 | 1,82E-06 | 6,04E-06 |
| 1352 | RM25_RS06575 | RM25_1335 | Aldo/keto reductase | 1421 | gene | 1529975 | 1530964 | + | 1756,939 | 0,135513897 | 0,258813 | 0,523599 | 0,600558 | 0,691169 | 2104,264495 | 0,298321573 | 0,178767 | 1,66877 | 0,095163 | 0,136186 | 1409,614 | -0,104675029 | 0,317273 | -0,32992 | 0,741459 | 0,809083 | 1776,817 | -0,42221 | 0,295339 | -1,42959 | 0,152836 | 0,210241 | 1800,963342 | -0,788326892 | 0,20388 | -3,86663 | 0,00011 | 0,000278 |
| 1353 | RM25_RS06580 | RM25_1336 | Hypothetical protein | 1422 | gene | 1530982 | 1531233 | - | 923,3568 | 1,385858564 | 0,309384 | 4,479416 | 7,48E-06 | 0,000223 | 880,2601153 | 0,698254234 | 0,192733 | 3,622904 | 0,000291 | 0,000901 | 966,4536 | 2,174436568 | 0,361991 | 6,006881 | 1,89E-09 | 5,07E-08 | 545,2297 | -0,99267 | 0,330458 | -3,00393 | 0,002665 | 0,008085 | 1303,030824 | 0,521305936 | 0,236518 | 2,204087 | 0,027518 | 0,043765 |
| 1355 | RM25_RS06590 | RM25_1338 | Hypothetical protein | 1426 | gene | 1532138 | 1532458 | + | 527,7234 | -1,308935953 | 0,470445 | -2,78234 | 0,005397 | 0,031164 | 311,5658235 | -1,776200799 | 0,364681 | -4,87055 | 1,11E-06 | 5,62E-06 | 743,8809 | -1,134535546 | 0,248302 | -4,56918 | 4,9E-06 | 5,47E-05 | 788,3503 | 1,018102 | 0,406094 | 2,507062 | 0,012174 | 0,027646 | 296,8374606 | 1,715807046 | 0,220533 | 7,780285 | 7,24E-15 | 8,61E-14 |
| 1356 | RM25_RS12600 | RM25_1339 | Hypothetical protein | 1427 | gene | 1532505 | 1533002 | + | 445,6263 | -1,180438732 | 0,435928 | -2,70788 | 0,006772 | 0,036523 | 289,5947887 | -1,900036482 | 0,281388 | -6,75237 | 1,45E-11 | 1,66E-10 | 601,6579 | -0,884206492 | 0,267763 | -3,3022 | 0,000959 | 0,004691 | 649,455 | 0,707001 | 0,313429 | 2,255697 | 0,02409 | 0,047554 | 267,7819657 | 1,785173196 | 0,25533 | 6,991626 | 2,72E-12 | 2,19E-11 |
| 1357 | RM25_RS06605 | RM25_1340 | Undecaprenyl-diphosphatase UppP | 1428 | gene | 1533517 | 1534353 | + | 1137,162 | 1,239272885 | 0,49099 | 2,524027 | 0,011602 | 0,051402 | 1726,679031 | 1,242081812 | 0,174646 | 7,111995 | 1,14E-12 | 1,51E-11 | 547,6447 | 1,234452345 | 0,258709 | 4,771579 | 1,83E-06 | 2,3E-05 | 722,5139 | -1,71255 | 0,207312 | -8,26075 | 1,45E-16 | 7,26E-15 | 1566,146409 | -1,674853672 | 0,199872 | -8,37963 | 5,31E-17 | 8,4E-16 |
| 1358 | RM25_RS06610 | RM25_1341 | Hypothetical protein | 1429 | gene | 1534424 | 1534810 | - | 3465,217 | 0,848645682 | 0,41955 | 2,022752 | 0,043099 | 0,113593 | 4572,849783 | 0,632029402 | 0,219846 | 2,874869 | 0,004042 | 0,009082 | 2357,585 | 1,300905351 | 0,46075 | 2,823453 | 0,004751 | 0,017228 | 2639,294 | -1,44266 | 0,423509 | -3,40646 | 0,000658 | 0,002486 | 4356,5593 | -0,749855592 | 0,276686 | -2,71014 | 0,006726 | 0,012119 |
| 1359 | RM25_RS06615 | RM25_1342 | Pyridoxal-phosphate dependent TrpB-like enzyme | 1430 | gene | 1534813 | 1536135 | - | 2500,359 | 0,534204607 | 0,258276 | 2,068348 | 0,038607 | 0,106699 | 3071,632925 | 0,566931619 | 0,155284 | 3,650938 | 0,000261 | 0,000816 | 1929,086 | 0,483782049 | 0,286604 | 1,68798 | 0,091415 | 0,160795 | 2168,094 | -0,67381 | 0,25363 | -2,65669 | 0,007891 | 0,019375 | 2896,5834 | -0,719777518 | 0,184464 | -3,90199 | 9,54E-05 | 0,000243 |
| 1360 | RM25_RS06620 | RM25_1343 | DNA protecting protein DprA | 1431 | gene | 1536298 | 1537458 | - | 316,6179 | -1,347671081 | 0,538549 | -2,50241 | 0,012335 | 0,053543 | 197,2719368 | -2,27670969 | 0,382948 | -5,94521 | 2,76E-09 | 2,22E-08 | 435,9638 | -1,009294599 | 0,469491 | -2,14976 | 0,031574 | 0,072277 | 477,2421 | 0,758947 | 0,41895 | 1,811544 | 0,070057 | 0,111645 | 176,3521792 | 2,106686953 | 0,475398 | 4,43142 | 9,36E-06 | 2,74E-05 |
| 1361 | RM25_RS06625 | RM25_1344 | Mg chelatase-like protein | 1432 | gene | 1537455 | 1538993 | - | 361,329 | -1,252232438 | 0,587807 | -2,13035 | 0,033143 | 0,096442 | 202,6289407 | -2,329419515 | 0,388331 | -5,99854 | 1,99E-09 | 1,63E-08 | 520,029 | -0,919451279 | 0,46988 | -1,95678 | 0,050374 | 0,101566 | 533,7549 | 0,937138 | 0,404107 | 2,319035 | 0,020393 | 0,041519 | 211,1022706 | 2,425784256 | 0,486549 | 4,98569 | 6,17E-07 | 2,24E-06 |
| 1362 | RM25_RS06630 | RM25_1345 | TIGR00252 family protein | 1433 | gene | 1538990 | 1539352 | - | 144,8619 | -0,569916504 | 0,284561 | -2,00279 | 0,0452 | 0,116687 | 139,6761055 | -0,305469364 | 0,263231 | -1,16046 | 0,245862 | 0,310717 | 150,0478 | -0,82684847 | 0,461713 | -1,79083 | 0,073321 | 0,135331 | 182,5809 | 0,245709 | 0,248428 | 0,989056 | 0,322636 | 0,398143 | 115,0695215 | -0,201973519 | 0,497759 | -0,40577 | 0,684915 | 0,735129 |
| 1363 | RM25_RS06635 | RM25_1346 | Protein often found in Actinomycetes clustered with signal p | 1434 | gene | 1539506 | 1539820 | - | 1099,034 | 0,705602908 | 0,431996 | 1,633357 | 0,102394 | 0,195577 | 1562,02179 | 0,697683894 | 0,228476 | 3,053649 | 0,002261 | 0,005488 | 636,0458 | 0,72690246 | 0,353632 | 2,055536 | 0,039827 | 0,085807 | 892,8049 | -1,37375 | 0,298008 | -4,60976 | 4,03E-06 | 2,71E-05 | 1334,140054 | -1,304674287 | 0,285765 | -4,56554 | 4,98E-06 | 1,54E-05 |
| 1364 | RM25_RS06640 | RM25_1347 | Ribonuclease HII | 1435 | gene | 1539817 | 1540452 | - | 261,7086 | 0,340860331 | 0,326108 | 1,045239 | 0,295912 | 0,406212 | 334,5441565 | 0,788358961 | 0,161854 | 4,870815 | 1,11E-06 | 5,62E-06 | 188,873 | -0,422388601 | 0,247626 | -1,70575 | 0,088054 | 0,155931 | 243,9907 | -0,24149 | 0,188284 | -1,2826 | 0,199632 | 0,263455 | 287,5370585 | -1,39296615 | 0,224994 | -6,19112 | 5,97E-10 | 3,45E-09 |
| 1365 | RM25_RS06645 | RM25_1348 | Signal peptidase I | 1436 | gene | 1540458 | 1541282 | - | 1316,199 | 0,442045451 | 0,428421 | 1,031801 | 0,302165 | 0,411917 | 1908,316117 | 0,521152625 | 0,192543 | 2,706687 | 0,006796 | 0,014251 | 724,0826 | 0,237925738 | 0,239196 | 0,99469 | 0,319887 | 0,425013 | 1187,176 | -1,29609 | 0,206102 | -6,28856 | 3,2E-10 | 5,17E-09 | 1486,338707 | -1,536753376 | 0,195467 | -7,86197 | 3,78E-15 | 4,82E-14 |
| 1366 | RM25_RS06650 | RM25_1349 | 50S ribosomal protein L19 | 1437 | gene | 1541294 | 1541650 | - | 4082,117 | 0,701185362 | 0,453458 | 1,546308 | 0,12203 | 0,21906 | 5440,01303 | 0,468254171 | 0,244817 | 1,912672 | 0,05579 | 0,086228 | 2724,221 | 1,19937834 | 0,521503 | 2,29985 | 0,021457 | 0,054089 | 3320,722 | -1,51142 | 0,515143 | -2,93399 | 0,003346 | 0,009666 | 4944,281169 | -0,753515295 | 0,258627 | -2,91352 | 0,003574 | 0,006771 |
| 1367 | RM25_RS06655 | RM25_1350 | Succinate dehydrogenase/fumarate reductase iron-sulfur su | 1438 | gene | 1541924 | 1542688 | - | 9043,568 | 0,553505747 | 0,478449 | 1,156876 | 0,247323 | 0,355806 | 12254,6618 | 0,14957943 | 0,215619 | 0,693722 | 0,487857 | 0,55496 | 5832,473 | 1,505219047 | 0,456916 | 3,294301 | 0,000987 | 0,004795 | 7836,425 | -1,98045 | 0,420925 | -4,705 | 2,54E-06 | 1,78E-05 | 10524,24939 | -0,597149235 | 0,275341 | -2,16876 | 0,030101 | 0,047393 |
| 1368 | RM25_RS06660 | RM25_1351 | Succinate dehydrogenase or fumarate reductase, flavoprote | 1439 | gene | 1542685 | 1544706 | - | 16694,96 | 0,493153098 | 0,417202 | 1,182048 | 0,237187 | 0,344667 | 21747,87864 | 0,154324475 | 0,204878 | 0,753252 | 0,451298 | 0,519897 | 11642,04 | 1,173620804 | 0,450254 | 2,606575 | 0,009145 | 0,02828 | 14806,54 | -1,57192 | 0,388852 | -4,04246 | 5,29E-05 | 0,000276 | 19088,78642 | -0,524911125 | 0,295838 | -1,77432 | 0,07601 | 0,107344 |
| 1369 | RM25_RS06665 | RM25_1352 | Succinate dehydrogenase cytochrome B subunit, b558 fami | 1440 | gene | 1544703 | 1545464 | - | 10964,06 | 0,249709629 | 0,302434 | 0,825666 | 0,408994 | 0,520237 | 12790,86871 | 0,10649729 | 0,195774 | 0,54398 | 0,586455 | 0,647435 | 9137,244 | 0,45261948 | 0,443404 | 1,020784 | 0,307357 | 0,411294 | 10653,52 | -0,72266 | 0,383794 | -1,88294 | 0,059708 | 0,098003 | 11635,94598 | -0,349293941 | 0,284774 | -1,22657 | 0,219985 | 0,276166 |
| 1370 | RM25_RS06670 | RM25_1353 | Pectin acetylesterase | 1441 | gene | 1545562 | 1546260 | + | 347,7472 | -0,266257951 | 0,186665 | -1,42639 | 0,153755 | 0,254674 | 305,3698959 | -0,188604712 | 0,16614 | -1,13522 | 0,256284 | 0,321246 | 390,1245 | -0,328958618 | 0,285592 | -1,15185 | 0,249383 | 0,349286 | 400,0078 | 0,354585 | 0,190007 | 1,866166 | 0,062018 | 0,10116 | 310,2896877 | 0,276309916 | 0,274436 | 1,006828 | 0,314017 | 0,377237 |
| 1371 | RM25_RS06675 | RM25_1354 | TRNA (guanine-N(1)-)-methyltransferase | 1442 | gene | 1546268 | 1546993 | - | 656,9503 | 0,529969736 | 0,299496 | 1,769541 | 0,076804 | 0,164271 | 845,4522028 | 0,717707934 | 0,170624 | 4,206383 | 2,59E-05 | 0,000102 | 468,4483 | 0,205433012 | 0,268882 | 0,764027 | 0,444851 | 0,548673 | 568,8892 | -0,61846 | 0,254456 | -2,43051 | 0,015078 | 0,032875 | 762,2482751 | -1,069788643 | 0,170898 | -6,25982 | 3,85E-10 | 2,28E-09 |
| 1372 | RM25_RS06680 | RM25_1355 | Ribosome maturation factor RimM | 1443 | gene | 1547059 | 1547610 | - | 720,9521 | 0,51511305 | 0,335275 | 1,536388 | 0,124443 | 0,221821 | 965,7832108 | 0,633048952 | 0,175275 | 3,611745 | 0,000304 | 0,000933 | 476,121 | 0,282182237 | 0,246772 | 1,143494 | 0,252834 | 0,353457 | 629,5574 | -0,88125 | 0,233698 | -3,77092 | 0,000163 | 0,000749 | 832,4523072 | -1,175079606 | 0,166016 | -7,0781 | 1,46E-12 | 1,24E-11 |
| 1373 | RM25_RS06685 | RM25_1356 | UPF0109 protein | 1444 | gene | 1547619 | 1547873 | - | 1084,255 | 0,960747275 | 0,660818 | 1,453876 | 0,145981 | 0,24665 | 1617,143463 | 0,533348856 | 0,204649 | 2,606169 | 0,009156 | 0,018414 | 551,3665 | 2,673398384 | 0,511916 | 5,22234 | 1,77E-07 | 2,98E-06 | 789,6994 | -3,19162 | 0,504708 | -6,32369 | 2,55E-10 | 4,18E-09 | 1403,027758 | -1,020927377 | 0,218085 | -4,68133 | 2,85E-06 | 9,16E-06 |
| 1374 | RM25_RS06690 | RM25_1357 | 30S ribosomal protein S16 | 1445 | gene | 1547876 | 1548367 | - | 4521,768 | 0,976469361 | 0,560402 | 1,742444 | 0,081431 | 0,16851 | 6369,190781 | 0,537016466 | 0,226573 | 2,370171 | 0,01778 | 0,032461 | 2674,345 | 2,312007595 | 0,430597 | 5,369308 | 7,9E-08 | 1,41E-06 | 3264,005 | -2,5832 | 0,419369 | -6,15973 | 7,29E-10 | 1,07E-08 | 5867,678901 | -0,77821142 | 0,240604 | -3,23441 | 0,001219 | 0,002527 |
| 1375 | RM25_RS06695 | RM25_1358 | M38 family peptidase | 1446 | gene | 1548545 | 1549603 | - | 1393,325 | 0,665463059 | 0,292279 | 2,27681 | 0,022798 | 0,077571 | 1788,02052 | 0,64986662 | 0,184295 | 3,52623 | 0,000422 | 0,001253 | 998,6296 | 0,694735739 | 0,214955 | 3,231998 | 0,001229 | 0,005737 | 1142,756 | -0,92876 | 0,236984 | -3,91908 | 8,89E-05 | 0,000438 | 1676,143078 | -0,830941469 | 0,121207 | -6,85555 | 7,1E-12 | 5,37E-11 |
| 1376 | RM25_RS06700 | RM25_1359 | Signal recognition particle protein | 1447 | gene | 1549668 | 1551239 | - | 4032,961 | 0,850399024 | 0,493564 | 1,722977 | 0,084893 | 0,1736 | 5662,486915 | 0,481282535 | 0,205714 | 2,339566 | 0,019306 | 0,034683 | 2403,435 | 1,878294291 | 0,297742 | 6,308471 | 2,82E-10 | 9,1E-09 | 3075,576 | -2,25705 | 0,261997 | -8,61477 | 7,01E-18 | 4,59E-16 | 5079,014221 | -0,820663878 | 0,223166 | -3,67736 | 0,000236 | 0,00056 |
| 1377 | RM25_RS06705 | RM25_1360 | Signal recognition particle receptor FtsY | 1448 | gene | 1551366 | 1552520 | - | 4540,669 | 0,477624816 | 0,280358 | 1,703622 | 0,088452 | 0,1788 | 5431,374251 | 0,571741099 | 0,202997 | 2,816503 | 0,004855 | 0,010605 | 3649,965 | 0,339982917 | 0,377305 | 0,901083 | 0,367544 | 0,473907 | 4029,486 | -0,48741 | 0,370685 | -1,31489 | 0,188546 | 0,250509 | 5173,893456 | -0,687014855 | 0,208957 | -3,28783 | 0,00101 | 0,002136 |
| 1378 | RM25_RS06710 | RM25_1361 | Putative insertion element ISCmi2 transposase | 1449 | gene | 1552530 | 1553573 | - | 4,017333 | 0,299887788 | 0,626337 | 0,478796 | 0,632084 | 0,718146 | 5,233760794 | 0,203001696 | 1,010443 | 0,200904 | 0,840774 | 0,874539 | 2,800906 | 0,446262352 | 0,899978 | 0,495859 | 0,619994 | 0,706637 | 3,812846 | -1,05331 | 0,833639 | -1,26351 | 0,206407 | 0,271183 | 4,35480063 | -0,76507833 | 1,1026 | -0,69389 | 0,487754 | 0,549539 |
| 1379 | RM25_RS06715 | RM25_1362 | Hypothetical protein | 1450 | gene | 1553711 | 1553908 | - | 1111,689 | 0,127392293 | 0,502102 | 0,253718 | 0,799714 | 0,859841 | 1586,094496 | 0,172494395 | 0,257908 | 0,668822 | 0,503609 | 0,569035 | 637,2827 | 0,018485559 | 0,568665 | 0,032507 | 0,974068 | 0,985032 | 1133,534 | -1,2812 | 0,560778 | -2,28468 | 0,022332 | 0,044538 | 1136,82495 | -1,408491018 | 0,279138 | -5,04586 | 4,51E-07 | 1,67E-06 |
| 1380 | RM25_RS06720 | RM25_1363 | Chromosome segregation protein SMC | 1451 | gene | 1553990 | 1557535 | - | 1569,499 | -0,242704864 | 0,207592 | -1,16914 | 0,242346 | 0,35065 | 1556,012242 | 0,117146905 | 0,141011 | 0,830762 | 0,406108 | 0,474875 | 1582,986 | -0,603479079 | 0,32857 | -1,83669 | 0,066256 | 0,125839 | 1789,113 | 0,292234 | 0,196788 | 1,485024 | 0,137537 | 0,192847 | 1414,371123 | -0,364214036 | 0,281572 | -1,2935 | 0,195837 | 0,250318 |
| 1381 | RM25_RS06725 | RM25_1364 | Hypothetical protein | 1452 | gene | 1557573 | 1558187 | - | 676,2985 | 0,072765778 | 0,151825 | 0,479274 | 0,631743 | 0,718146 | 713,1275729 | 0,266009171 | 0,156035 | 1,704809 | 0,08823 | 0,126959 | 639,4695 | -0,139922083 | 0,245699 | -0,56949 | 0,569027 | 0,662319 | 696,3186 | -0,00977 | 0,180679 | -0,05407 | 0,956875 | 0,966305 | 678,8737533 | -0,371496885 | 0,202371 | -1,83573 | 0,066398 | 0,095485 |
| 1382 | RM25_RS06730 | RM25_1365 | DNA-formamidopyrimidine glycosylase | 1453 | gene | 1558199 | 1559080 | - | 464,729 | -0,801143382 | 0,247361 | -3,23877 | 0,0012 | 0,011314 | 402,2883022 | -0,694811004 | 0,240869 | -2,8846 | 0,003919 | 0,008831 | 527,1696 | -0,886051981 | 0,358741 | -2,46989 | 0,013515 | 0,038427 | 621,2341 | 0,39269 | 0,289332 | 1,357229 | 0,174708 | 0,23576 | 333,5308891 | 0,269906672 | 0,334871 | 0,806002 | 0,420241 | 0,48602 |
| 1383 | RM25_RS06735 | RM25_1366 | Ribonuclease III | 1454 | gene | 1559104 | 1559826 | - | 1206,441 | 0,822316598 | 0,521828 | 1,575837 | 0,115063 | 0,20991 | 1819,323136 | 0,64599926 | 0,214549 | 3,010957 | 0,002604 | 0,006219 | 593,5594 | 1,413021836 | 0,247739 | 5,703672 | 1,17E-08 | 2,51E-07 | 930,8712 | -2,18845 | 0,25589 | -8,55233 | 1,21E-17 | 7,29E-16 | 1510,773346 | -1,377415231 | 0,182862 | -7,53254 | 4,98E-14 | 5,26E-13 |
| 1384 | RM25_RS12305 | RM25_1367 | Hypothetical protein | 1455 | gene | 1559813 | 1560025 | - | 5848,733 | 0,637131308 | 0,593531 | 1,07346 | 0,283065 | 0,392 | 8648,129524 | 0,271335447 | 0,275807 | 0,983789 | 0,325219 | 0,393497 | 3049,336 | 1,858766391 | 0,479989 | 3,872521 | 0,000108 | 0,000788 | 4909,723 | -2,6187 | 0,438034 | -5,9783 | 2,25E-09 | 2,9E-08 | 6970,138306 | -1,003654982 | 0,33897 | -2,9609 | 0,003067 | 0,005902 |
| 1385 | RM25_RS06740 | RM25_1368 | Putative metal-binding protein, possibly nucleic-acid bindin | 1456 | gene | 1560176 | 1560664 | - | 1087,753 | 0,997710948 | 0,197558 | 5,05022 | 4,41E-07 | 2,08E-05 | 947,8451779 | 0,711283765 | 0,156679 | 4,539753 | 5,63E-06 | 2,49E-05 | 1227,661 | 1,233176734 | 0,28846 | 4,275037 | 1,91E-05 | 0,000181 | 766,0923 | -0,03014 | 0,198095 | -0,15214 | 0,879074 | 0,903998 | 1415,066908 | 0,533827557 | 0,218336 | 2,444984 | 0,014486 | 0,024458 |
| 1386 | RM25_RS06745 | RM25_1369 | Pantetheine-phosphate adenylyltransferase | 1457 | gene | 1560824 | 1561309 | - | 656,6084 | 0,089470538 | 0,218167 | 0,410102 | 0,681731 | 0,760658 | 791,0955398 | 0,176917512 | 0,141599 | 1,249423 | 0,21151 | 0,272719 | 522,1213 | -0,041787588 | 0,197629 | -0,21144 | 0,83254 | 0,877985 | 672,9135 | -0,54663 | 0,170929 | -3,19798 | 0,001384 | 0,004653 | 664,1459144 | -0,712902306 | 0,131912 | -5,40438 | 6,5E-08 | 2,77E-07 |
| 1387 | RM25_RS06750 | RM25_1370 | RNA methyltransferase, RsmD family | 1458 | gene | 1561302 | 1561901 | - | 640,6008 | -0,020676634 | 0,186233 | -0,11103 | 0,911596 | 0,941556 | 731,3700738 | 0,203401011 | 0,157161 | 1,294223 | 0,195588 | 0,254697 | 549,8315 | -0,316807438 | 0,177225 | -1,7876 | 0,07384 | 0,13597 | 681,4004 | -0,21307 | 0,152355 | -1,39848 | 0,16197 | 0,220875 | 623,7238794 | -0,686505005 | 0,153645 | -4,46812 | 7,89E-06 | 2,35E-05 |
| 1388 | RM25_RS06755 | RM25_1371 | Pyridoxamine 5'-phosphate oxidase family protein | 1459 | gene | 1561977 | 1562399 | + | 4164,443 | 0,536661244 | 0,289931 | 1,850994 | 0,06417 | 0,145232 | 4610,48714 | 0,720541238 | 0,212097 | 3,397225 | 0,000681 | 0,001906 | 3718,399 | 0,314940972 | 0,459676 | 0,685137 | 0,493258 | 0,59347 | 3605,509 | -0,11834 | 0,440359 | -0,26873 | 0,788136 | 0,8281 | 4820,357837 | -0,495560917 | 0,248954 | -1,99057 | 0,046528 | 0,069278 |
| 1389 | RM25_RS06760 | RM25_1372 | ATP-dependent DNA helicase RecG | 1460 | gene | 1562506 | 1564785 | - | 1398,348 | -0,070027005 | 0,148843 | -0,47048 | 0,638015 | 0,722634 | 1311,623448 | 0,20450491 | 0,136533 | 1,497841 | 0,134175 | 0,185266 | 1485,073 | -0,313084677 | 0,216852 | -1,44377 | 0,148803 | 0,233819 | 1507,27 | 0,368524 | 0,185829 | 1,983131 | 0,047353 | 0,081468 | 1339,550857 | -0,089288468 | 0,171213 | -0,52151 | 0,602014 | 0,662131 |
| 1390 | RM25_RS06765 | RM25_1373 | 50S ribosomal protein L28 | 1461 | gene | 1564915 | 1565100 | - | 3290,122 | 0,375151929 | 0,677215 | 0,553963 | 0,579604 | 0,671976 | 5344,275069 | 0,174938501 | 0,249515 | 0,701113 | 0,483232 | 0,550763 | 1235,969 | 1,316754733 | 0,540239 | 2,437357 | 0,014795 | 0,040738 | 3074,863 | -2,87107 | 0,543802 | -5,27962 | 1,29E-07 | 1,18E-06 | 3642,908532 | -1,699810273 | 0,248994 | -6,82671 | 8,69E-12 | 6,48E-11 |
| 1391 | RM25_RS06770 | RM25_1374 | Metallo-beta-lactamase family protein | 1462 | gene | 1565235 | 1566986 | - | 4236,065 | 0,417309352 | 0,38238 | 1,091346 | 0,275121 | 0,384158 | 5853,295809 | 0,518086093 | 0,189618 | 2,732261 | 0,00629 | 0,013369 | 2618,835 | 0,196322636 | 0,304289 | 0,645184 | 0,518808 | 0,619155 | 3857,005 | -1,031 | 0,276874 | -3,7237 | 0,000196 | 0,000883 | 4746,348983 | -1,317915815 | 0,207174 | -6,36138 | 2E-10 | 1,23E-09 |
| 1392 | RM25_RS06775 | RM25_1375 | Putative GCN5-related N-acetyltransferase | 1463 | gene | 1567074 | 1567562 | + | 421,1873 | 0,623775617 | 0,247459 | 2,520721 | 0,011711 | 0,051596 | 515,3992108 | 0,608710143 | 0,161276 | 3,77433 | 0,00016 | 0,000521 | 326,9753 | 0,652742244 | 0,254395 | 2,565862 | 0,010292 | 0,031021 | 352,4476 | -0,74352 | 0,219656 | -3,38494 | 0,000712 | 0,002649 | 500,0508699 | -0,654825206 | 0,1898 | -3,45007 | 0,00056 | 0,001251 |
| 1393 | RM25_RS06780 | RM25_1376 | Hypothetical protein | 1464 | gene | 1567575 | 1568300 | - | 739,7675 | 0,274657479 | 0,189439 | 1,449846 | 0,147101 | 0,247656 | 865,7385537 | 0,343468327 | 0,143715 | 2,389926 | 0,016852 | 0,031055 | 613,7965 | 0,180745788 | 0,196584 | 0,919433 | 0,357869 | 0,464993 | 708,8898 | -0,46523 | 0,16619 | -2,79937 | 0,00512 | 0,013439 | 793,5576441 | -0,582128336 | 0,142873 | -4,07444 | 4,61E-05 | 0,000123 |
| 1394 | RM25_RS06785 | RM25_1377 | Dihydrodipicolinate reductase | 1465 | gene | 1568306 | 1569046 | - | 1904,291 | 0,367183689 | 0,277411 | 1,323607 | 0,185634 | 0,290917 | 2264,057125 | 0,524743192 | 0,199936 | 2,624562 | 0,008676 | 0,017553 | 1544,525 | 0,141086051 | 0,375017 | 0,376212 | 0,706759 | 0,780853 | 1763,557 | -0,38631 | 0,334193 | -1,15595 | 0,247701 | 0,317041 | 2098,665841 | -0,740613213 | 0,248387 | -2,9817 | 0,002867 | 0,005538 |
| 1395 | RM25_RS06790 | RM25_1378 | Polyribonucleotide nucleotidyltransferase | 1466 | gene | 1569251 | 1571461 | - | 8581,5 | 0,74946911 | 0,605014 | 1,238763 | 0,215433 | 0,320706 | 13457,05501 | 0,513554378 | 0,197313 | 2,602735 | 0,009248 | 0,018552 | 3705,944 | 1,73211908 | 0,318566 | 5,437242 | 5,41E-08 | 1,01E-06 | 6855,817 | -2,74714 | 0,301316 | -9,11714 | 7,71E-20 | 5,51E-18 | 10551,41866 | -1,48912255 | 0,204635 | -7,27697 | 3,41E-13 | 3,17E-12 |
| 1396 | RM25_RS06795 | RM25_1379 | 30S ribosomal protein S15 | 1467 | gene | 1571786 | 1572049 | - | 7031,863 | 0,551093726 | 0,591635 | 0,931475 | 0,351608 | 0,46454 | 10354,03827 | 0,164309969 | 0,235803 | 0,696809 | 0,485922 | 0,553294 | 3709,688 | 1,812706226 | 0,510591 | 3,550215 | 0,000385 | 0,00228 | 6113,461 | -2,61569 | 0,5069 | -5,16018 | 2,47E-07 | 2,11E-06 | 8182,416272 | -0,939758537 | 0,243125 | -3,86533 | 0,000111 | 0,000279 |
| 1397 | RM25_RS06800 | RM25_1380 | Riboflavin biosynthesis protein RibF | 1468 | gene | 1572153 | 1573130 | - | 931,9105 | 0,5415902 | 0,307085 | 1,763647 | 0,077791 | 0,165184 | 1219,041786 | 0,525358479 | 0,16865 | 3,115079 | 0,001839 | 0,004558 | 644,7793 | 0,574324708 | 0,199856 | 2,87369 | 0,004057 | 0,015325 | 806,8423 | -1,00796 | 0,191744 | -5,25679 | 1,47E-07 | 1,33E-06 | 1082,49424 | -0,91185915 | 0,150749 | -6,04887 | 1,46E-09 | 7,96E-09 |
| 1398 | RM25_RS06805 | RM25_1381 | Glycerol kinase | 1469 | gene | 1573456 | 1574982 | - | 2284,283 | 0,163499056 | 0,231692 | 0,705675 | 0,48039 | 0,580656 | 2206,600138 | -0,156873566 | 0,158576 | -0,98927 | 0,322533 | 0,390452 | 2361,966 | 0,466485368 | 0,379636 | 1,22877 | 0,219158 | 0,31613 | 2286,141 | -0,2815 | 0,34738 | -0,81036 | 0,417736 | 0,493288 | 2359,240521 | 0,37924162 | 0,219131 | 1,730661 | 0,083512 | 0,117166 |
| 1399 | RM25_RS06810 | RM25_1382 | Glycerol uptake facilitator protein | 1470 | gene | 1575040 | 1575831 | - | 476,1487 | -1,843405965 | 0,453639 | -4,0636 | 4,83E-05 | 0,000973 | 317,9323934 | -2,627670702 | 0,264759 | -9,92477 | 3,25E-23 | 1,7E-21 | 634,3651 | -1,530922272 | 0,322223 | -4,75113 | 2,02E-06 | 2,54E-05 | 783,0046 | 0,719601 | 0,263064 | 2,735457 | 0,006229 | 0,015907 | 204,1378853 | 1,882010736 | 0,354519 | 5,308637 | 1,1E-07 | 4,5E-07 |
| 1400 | RM25_RS06815 | RM25_1383 | TRNA pseudouridine synthase B | 1471 | gene | 1576076 | 1576972 | - | 674,5788 | 0,394713985 | 0,283026 | 1,394619 | 0,163131 | 0,265538 | 853,2094951 | 0,420434539 | 0,212075 | 1,982484 | 0,047425 | 0,075732 | 495,948 | 0,352998014 | 0,22955 | 1,537782 | 0,124102 | 0,203272 | 617,0373 | -0,80134 | 0,237434 | -3,37499 | 0,000738 | 0,002731 | 752,4758872 | -0,813717811 | 0,197409 | -4,122 | 3,76E-05 | 0,000102 |
| 1401 | RM25_RS06820 | RM25_1384 | Ribosome-binding factor A | 1472 | gene | 1576969 | 1577409 | - | 1633,934 | 1,185939939 | 0,472128 | 2,511906 | 0,012008 | 0,052413 | 2079,07396 | 0,627192873 | 0,206875 | 3,031751 | 0,002431 | 0,00583 | 1188,794 | 2,483376689 | 0,314863 | 7,887172 | 3,09E-15 | 2,51E-13 | 1067,446 | -2,23486 | 0,302026 | -7,39957 | 1,37E-13 | 3,88E-12 | 2221,141028 | -0,338573853 | 0,210958 | -1,60494 | 0,108508 | 0,148349 |
| 1402 | RM25_RS06825 | RM25_1385 | Translation initiation factor IF-2 | 1473 | gene | 1577409 | 1580330 | - | 7742,389 | 1,071733686 | 0,448213 | 2,391124 | 0,016797 | 0,064796 | 9777,370838 | 0,533135338 | 0,208976 | 2,551185 | 0,010736 | 0,02114 | 5707,407 | 2,241923885 | 0,328063 | 6,833812 | 8,27E-12 | 3,82E-10 | 5330,088 | -2,05563 | 0,324314 | -6,33838 | 2,32E-10 | 3,94E-09 | 10265,83131 | -0,310863753 | 0,197312 | -1,57549 | 0,115143 | 0,156512 |
| 1403 | RM25_RS06830 | RM25_1386 | Hypothetical protein | 1474 | gene | 1580526 | 1580723 | - | 140,7546 | 0,19309413 | 0,150451 | 1,283437 | 0,199339 | 0,302149 | 157,8126251 | 0,24509704 | 0,208513 | 1,175453 | 0,239813 | 0,303566 | 123,6966 | 0,133102227 | 0,195156 | 0,682029 | 0,495221 | 0,595528 | 139,0203 | -0,3531 | 0,192783 | -1,83158 | 0,067014 | 0,108321 | 147,4050458 | -0,407833794 | 0,215371 | -1,89364 | 0,058273 | 0,084784 |
| 1404 | RM25_RS06835 | RM25_1387 | Transcription termination factor NusA | 1475 | gene | 1580870 | 1581928 | - | 1900,717 | 0,070710423 | 0,229588 | 0,307988 | 0,758092 | 0,824795 | 2075,88756 | 0,281234691 | 0,194094 | 1,448961 | 0,147348 | 0,200752 | 1725,546 | -0,180462037 | 0,350757 | -0,51449 | 0,606907 | 0,695082 | 1963,206 | -0,08108 | 0,342628 | -0,23663 | 0,812941 | 0,849336 | 1905,782397 | -0,508851352 | 0,2007 | -2,53538 | 0,011233 | 0,019439 |
| 1405 | RM25_RS06840 | RM25_1388 | Ribosome maturation factor RimP | 1476 | gene | 1581930 | 1582403 | - | 1488,189 | 0,217432693 | 0,270438 | 0,804002 | 0,421396 | 0,530855 | 1702,39803 | 0,503392246 | 0,194963 | 2,581994 | 0,009823 | 0,019605 | 1273,98 | -0,157710249 | 0,381463 | -0,41343 | 0,679288 | 0,757371 | 1458,389 | -0,12035 | 0,313342 | -0,38408 | 0,70092 | 0,747737 | 1565,326501 | -0,748283467 | 0,279315 | -2,67899 | 0,007384 | 0,013246 |
| 1406 | RM25_RS06845 | RM25_1389 | Hypothetical protein | 1477 | gene | 1582673 | 1583659 | + | 404,346 | -0,37087377 | 0,250236 | -1,48209 | 0,138315 | 0,238346 | 326,5706879 | -0,384963432 | 0,17525 | -2,19666 | 0,028045 | 0,048245 | 482,1213 | -0,3631694 | 0,33007 | -1,10028 | 0,27121 | 0,374045 | 479,1093 | 0,489349 | 0,209675 | 2,333389 | 0,019604 | 0,04025 | 346,8218914 | 0,576119769 | 0,307166 | 1,875599 | 0,06071 | 0,08795 |
| 1407 | RM25_RS06850 | RM25_1390 | Proline--tRNA ligase | 1478 | gene | 1583718 | 1585538 | - | 2463,775 | 0,337368319 | 0,274425 | 1,229364 | 0,218935 | 0,324344 | 3108,27547 | 0,232902795 | 0,158253 | 1,471712 | 0,141099 | 0,19358 | 1819,275 | 0,518922212 | 0,195498 | 2,654367 | 0,007946 | 0,025796 | 2313,096 | -0,99316 | 0,195454 | -5,08131 | 3,75E-07 | 3,11E-06 | 2694,755701 | -0,660483112 | 0,124847 | -5,29032 | 1,22E-07 | 4,91E-07 |
| 1408 | RM25_RS06855 | RM25_1391 | Acetyltransferase, GNAT family | 1479 | gene | 1585623 | 1586465 | - | 717,8649 | 0,527950907 | 0,111793 | 4,722576 | 2,33E-06 | 8,32E-05 | 772,8014052 | 0,549110513 | 0,131495 | 4,175894 | 2,97E-05 | 0,000115 | 662,9285 | 0,505072775 | 0,18727 | 2,697023 | 0,006996 | 0,023457 | 620,9543 | -0,25528 | 0,14673 | -1,73979 | 0,081896 | 0,125669 | 830,761459 | -0,249634071 | 0,127991 | -1,9504 | 0,051128 | 0,075318 |
| 1409 | RM25_RS06860 | RM25_1392 | 4-hydroxy-3-methylbut-2-en-1-yl diphosphate synthase | 1480 | gene | 1586605 | 1587771 | - | 2641,484 | 0,933344538 | 0,215971 | 4,321614 | 1,55E-05 | 0,000412 | 3015,016899 | 0,830005748 | 0,147834 | 5,614461 | 1,97E-08 | 1,35E-07 | 2267,951 | 1,075440357 | 0,297289 | 3,617487 | 0,000297 | 0,001835 | 1928,105 | -0,62665 | 0,24467 | -2,56122 | 0,010431 | 0,024512 | 3391,058939 | -0,343395166 | 0,20252 | -1,69561 | 0,08996 | 0,125316 |
| 1410 | RM25_RS06865 | RM25_1393 | 1-deoxy-D-xylulose 5-phosphate reductoisomerase | 1481 | gene | 1587844 | 1589037 | - | 1368,539 | 0,266801124 | 0,167667 | 1,591252 | 0,111553 | 0,20622 | 1549,552835 | 0,269673986 | 0,152469 | 1,768717 | 0,076941 | 0,113202 | 1187,526 | 0,264336989 | 0,209029 | 1,264593 | 0,206017 | 0,302689 | 1316,606 | -0,44005 | 0,181898 | -2,41923 | 0,015553 | 0,03364 | 1463,219083 | -0,399713308 | 0,157378 | -2,53984 | 0,01109 | 0,019207 |
| 1411 | RM25_RS06870 | RM25_1394 | Ribosomal RNA large subunit methyltransferase N | 1482 | gene | 1589108 | 1590244 | - | 1515,699 | 0,783328898 | 0,26931 | 2,908655 | 0,00363 | 0,02353 | 1890,505994 | 0,694549019 | 0,169257 | 4,103525 | 4,07E-05 | 0,000152 | 1140,893 | 0,934891268 | 0,208219 | 4,48995 | 7,12E-06 | 7,56E-05 | 1183,474 | -0,94063 | 0,190998 | -4,92482 | 8,44E-07 | 6,55E-06 | 1877,792742 | -0,654720725 | 0,158783 | -4,12338 | 3,73E-05 | 0,000101 |
| 1412 | RM25_RS06875 | RM25_1395 | Phosphatidate cytidylyltransferase | 1483 | gene | 1590241 | 1591086 | - | 895,2499 | 0,310260651 | 0,363611 | 0,853276 | 0,393506 | 0,504694 | 1232,372782 | 0,341280894 | 0,174537 | 1,955348 | 0,050542 | 0,079631 | 558,127 | 0,245165657 | 0,212034 | 1,156254 | 0,247577 | 0,348174 | 849,3172 | -1,14826 | 0,21838 | -5,25807 | 1,46E-07 | 1,32E-06 | 972,2150724 | -1,1955382 | 0,139638 | -8,56172 | 1,11E-17 | 1,9E-16 |
| 1413 | RM25_RS06880 | RM25_1396 | Ribosome-recycling factor | 1484 | gene | 1591095 | 1591652 | - | 2115,952 | 0,490993124 | 0,339013 | 1,4483 | 0,147533 | 0,248028 | 2712,651279 | 0,347810627 | 0,181133 | 1,920191 | 0,054834 | 0,085309 | 1519,253 | 0,754583429 | 0,368065 | 2,050138 | 0,040351 | 0,086382 | 1873,035 | -1,12786 | 0,343069 | -3,28757 | 0,001011 | 0,003555 | 2421,486932 | -0,685686361 | 0,216924 | -3,16096 | 0,001573 | 0,003179 |
| 1414 | RM25_RS06885 | RM25_1397 | UMP kinase | 1485 | gene | 1591725 | 1592474 | - | 4253,223 | 0,366186594 | 0,29797 | 1,228937 | 0,219095 | 0,324377 | 5123,275481 | 0,33658062 | 0,178023 | 1,890655 | 0,05867 | 0,089913 | 3383,17 | 0,411756498 | 0,41545 | 0,991111 | 0,321632 | 0,42661 | 3950,118 | -0,68949 | 0,349647 | -1,97197 | 0,048614 | 0,083151 | 4682,154189 | -0,586181786 | 0,271721 | -2,1573 | 0,030983 | 0,048652 |
| 1415 | RM25_RS06890 | RM25_1398 | Translation elongation factor Ts | 1486 | gene | 1592611 | 1593423 | - | 3529,16 | 0,959040275 | 0,438495 | 2,187117 | 0,028734 | 0,087076 | 4556,294579 | 0,518169497 | 0,211263 | 2,452727 | 0,014178 | 0,026798 | 2502,025 | 1,917042433 | 0,376652 | 5,089687 | 3,59E-07 | 5,64E-06 | 2561,432 | -1,89118 | 0,363128 | -5,20803 | 1,91E-07 | 1,68E-06 | 4560,78334 | -0,455672438 | 0,22257 | -2,04732 | 0,040627 | 0,06158 |
| 1416 | RM25_RS06895 | RM25_1399 | 30S ribosomal protein S2 | 1487 | gene | 1593462 | 1594439 | - | 3486,771 | 0,676071162 | 0,431306 | 1,567496 | 0,116999 | 0,212454 | 4660,162894 | 0,429140152 | 0,223805 | 1,917469 | 0,055178 | 0,085575 | 2313,378 | 1,210592673 | 0,452959 | 2,672631 | 0,007526 | 0,02474 | 2866,124 | -1,55817 | 0,458011 | -3,40202 | 0,000669 | 0,002523 | 4200,692119 | -0,741577681 | 0,210931 | -3,51574 | 0,000439 | 0,000992 |
| 1417 | RM25_RS12755 | RM25_1400 | Peptidase, M23 family | 1488 | gene | 1594733 | 1595917 | + | 281,5352 | -1,321915492 | 0,671318 | -1,96913 | 0,048938 | 0,12297 | 151,884943 | -2,963263304 | 0,429521 | -6,89899 | 5,24E-12 | 6,3E-11 | 411,1855 | -0,898817236 | 0,495123 | -1,81534 | 0,069472 | 0,129787 | 422,531 | 0,914752 | 0,3906 | 2,341916 | 0,019185 | 0,039666 | 159,0635371 | 3,065126733 | 0,562721 | 5,446973 | 5,12E-08 | 2,22E-07 |
| 1418 | RM25_RS06910 | RM25_1401 | Tyrosine recombinase XerC | 1489 | gene | 1595961 | 1596887 | - | 824,7729 | 0,501763027 | 0,101943 | 4,921992 | 8,57E-07 | 3,61E-05 | 797,199499 | 0,495744507 | 0,153071 | 3,238652 | 0,001201 | 0,003152 | 852,3464 | 0,506089258 | 0,192607 | 2,627572 | 0,0086 | 0,027258 | 722,3276 | 0,033396 | 0,206427 | 0,161783 | 0,871477 | 0,898937 | 946,3857172 | 0,09142354 | 0,127454 | 0,717304 | 0,473187 | 0,535945 |
| 1419 | RM25_RS06915 | RM25_1402 | Peptide deformylase | 1490 | gene | 1597047 | 1597667 | + | 1428,149 | 0,512623429 | 0,331868 | 1,54466 | 0,122428 | 0,21944 | 1865,178204 | 0,525697706 | 0,195593 | 2,687715 | 0,007194 | 0,01494 | 991,1193 | 0,490705085 | 0,324014 | 1,514456 | 0,12991 | 0,211317 | 1251,432 | -0,94369 | 0,331228 | -2,84906 | 0,004385 | 0,011852 | 1645,324975 | -0,940326981 | 0,169634 | -5,54328 | 2,97E-08 | 1,33E-07 |
| 1420 | RM25_RS06920 | RM25_1403 | RNA methyltransferase, TrmH family | 1491 | gene | 1597701 | 1598510 | - | 649,5545 | 0,094944502 | 0,128739 | 0,737495 | 0,460822 | 0,564889 | 706,1811185 | 0,100526663 | 0,160661 | 0,625705 | 0,531508 | 0,597124 | 592,928 | 0,089758322 | 0,184288 | 0,487054 | 0,62622 | 0,712012 | 664,7914 | -0,30697 | 0,189778 | -1,61753 | 0,105764 | 0,155494 | 657,299193 | -0,266380938 | 0,135575 | -1,96482 | 0,049435 | 0,073236 |
| 1421 | RM25_RS06925 | RM25_1404 | Sodium Bile acid symporter family protein | 1492 | gene | 1598651 | 1599694 | + | 1294,893 | -0,204063195 | 0,120778 | -1,68957 | 0,09111 | 0,182183 | 1238,502587 | -0,304408636 | 0,15544 | -1,95837 | 0,050187 | 0,07923 | 1351,283 | -0,113222012 | 0,205789 | -0,55019 | 0,582192 | 0,673651 | 1464,337 | -0,01922 | 0,161588 | -0,11893 | 0,90533 | 0,924951 | 1176,810506 | 0,215062026 | 0,169427 | 1,269349 | 0,204316 | 0,258772 |
| 1422 | RM25_RS06930 | RM25_1405 | ABC-type molybdenum transport system, ATPase componer | 1493 | gene | 1599776 | 1600570 | - | 2661,2 | 0,422455364 | 0,259336 | 1,628989 | 0,103315 | 0,1967 | 2857,748373 | 0,827643905 | 0,173142 | 4,780138 | 1,75E-06 | 8,5E-06 | 2464,652 | -0,027137962 | 0,389541 | -0,06967 | 0,944459 | 0,964929 | 2406,419 | 0,221574 | 0,35069 | 0,631822 | 0,527503 | 0,594324 | 2981,339535 | -0,600116123 | 0,235993 | -2,54294 | 0,010992 | 0,019051 |
| 1423 | RM25_RS06935 | RM25_1406 | Hypothetical protein | 1494 | gene | 1600723 | 1601472 | + | 243,0645 | -0,009248412 | 0,461254 | -0,02005 | 0,984003 | 0,990305 | 134,589066 | -0,172026629 | 0,314424 | -0,54712 | 0,584299 | 0,64566 | 351,5399 | 0,048933584 | 0,35503 | 0,137829 | 0,890375 | 0,921253 | 254,4382 | 1,212199 | 0,320617 | 3,780829 | 0,000156 | 0,000724 | 238,3513329 | 1,498161946 | 0,342273 | 4,377099 | 1,2E-05 | 3,5E-05 |
| 1424 | RM25_RS06940 | RM25_1407 | Enoyl-[acyl-carrier-protein] reductase (NADH) | 1495 | gene | 1601547 | 1602320 | - | 3070,897 | 0,231185713 | 0,322376 | 0,71713 | 0,473294 | 0,573844 | 3936,081874 | 0,287298058 | 0,199975 | 1,436669 | 0,150812 | 0,204707 | 2205,712 | 0,132556914 | 0,346957 | 0,382056 | 0,70242 | 0,777644 | 3003,734 | -0,80241 | 0,339485 | -2,3636 | 0,018098 | 0,037783 | 3247,803978 | -0,922858598 | 0,204505 | -4,51265 | 6,4E-06 | 1,93E-05 |
| 1425 | RM25_RS06945 | RM25_1408 | 3-oxoacyl-[acyl-carrier protein] reductase fabG1 | 1496 | gene | 1602410 | 1603105 | - | 1120,112 | 0,496401046 | 0,221623 | 2,239841 | 0,025101 | 0,081105 | 1324,199707 | 0,437755514 | 0,210712 | 2,07751 | 0,037755 | 0,062491 | 916,0241 | 0,583002889 | 0,190898 | 3,053998 | 0,002258 | 0,009573 | 985,0734 | -0,67087 | 0,242051 | -2,77163 | 0,005578 | 0,014511 | 1284,033468 | -0,483410226 | 0,147602 | -3,27509 | 0,001056 | 0,002218 |
| 1426 | RM25_RS06950 | RM25_1409 | Short-chain dehydrogenase/reductase SDR | 1497 | gene | 1603214 | 1603960 | + | 609,3413 | 0,200443949 | 0,220254 | 0,910058 | 0,362792 | 0,47374 | 599,167723 | 0,537979998 | 0,214659 | 2,506212 | 0,012203 | 0,023518 | 619,5149 | -0,117561358 | 0,327279 | -0,35921 | 0,719439 | 0,791651 | 599,2434 | 0,349832 | 0,351134 | 0,996291 | 0,319109 | 0,39441 | 638,2622256 | -0,266069937 | 0,177038 | -1,50289 | 0,132866 | 0,177532 |
| 1427 | RM25_RS06955 | RM25_1410 | Ribosomal RNA large subunit methyltransferase A | 1498 | gene | 1603968 | 1604801 | + | 427,6215 | 0,404302022 | 0,306192 | 1,320421 | 0,186695 | 0,291031 | 556,4409185 | 0,49127608 | 0,230411 | 2,132175 | 0,032992 | 0,055545 | 298,802 | 0,250998583 | 0,168975 | 1,485415 | 0,137434 | 0,220393 | 390,1602 | -0,81425 | 0,232901 | -3,49615 | 0,000472 | 0,001873 | 478,1238265 | -1,005913815 | 0,188493 | -5,33662 | 9,47E-08 | 3,9E-07 |
| 1428 | RM25_RS06960 | RM25_1411 | Macrolide ABC transporter ATP-binding protein | 1499 | gene | 1604952 | 1606550 | - | 1002,15 | 0,378971689 | 0,092102 | 4,114677 | 3,88E-05 | 0,000809 | 1016,041835 | 0,376055043 | 0,129416 | 2,905793 | 0,003663 | 0,008399 | 988,2587 | 0,383713964 | 0,201076 | 1,908304 | 0,056352 | 0,110225 | 919,0916 | -0,10602 | 0,168362 | -0,62969 | 0,5289 | 0,595612 | 1111,22069 | -0,043513843 | 0,137945 | -0,31544 | 0,752425 | 0,79599 |
| 1429 | RM25_RS06965 | RM25_1412 | Putative aromatic ring hydroxylating enzyme | 1500 | gene | 1606824 | 1607273 | - | 1741,688 | 1,638649899 | 0,482809 | 3,393995 | 0,000689 | 0,007843 | 2137,305764 | 1,050170729 | 0,213762 | 4,912793 | 8,98E-07 | 4,59E-06 | 1346,069 | 2,991516395 | 0,355387 | 8,417624 | 3,84E-17 | 3,94E-15 | 906,7931 | -2,26754 | 0,327541 | -6,92292 | 4,42E-12 | 1,02E-10 | 2577,402975 | -0,286512995 | 0,238971 | -1,19894 | 0,230549 | 0,287364 |
| 1430 | RM25_RS06970 | RM25_1413 | SUF system FeS assembly protein, NifU family | 1501 | gene | 1607270 | 1607722 | - | 2757,428 | 1,564568785 | 0,374797 | 4,174441 | 2,99E-05 | 0,000652 | 3114,803042 | 0,986316462 | 0,171422 | 5,753729 | 8,73E-09 | 6,47E-08 | 2400,053 | 2,558842412 | 0,330439 | 7,743759 | 9,65E-15 | 7,34E-13 | 1489,885 | -1,6443 | 0,251189 | -6,54604 | 5,91E-11 | 1,13E-09 | 4024,101518 | -0,031811707 | 0,247606 | -0,12848 | 0,897772 | 0,916832 |
| 1431 | RM25_RS06975 | RM25_1414 | Cysteine desulfurase, SufS family protein | 1502 | gene | 1607719 | 1608975 | - | 3718,956 | 1,704623628 | 0,43246 | 3,941692 | 8,09E-05 | 0,001423 | 4160,441549 | 0,987448023 | 0,181514 | 5,440072 | 5,33E-08 | 3,4E-07 | 3277,47 | 3,058952778 | 0,289671 | 10,5601 | 4,56E-26 | 1,5E-23 | 1865,5 | -2,04954 | 0,193811 | -10,5749 | 3,89E-26 | 9,18E-24 | 5559,563135 | 0,064500629 | 0,23474 | 0,274775 | 0,783489 | 0,824045 |
| 1432 | RM25_RS06980 | RM25_1415 | FeS assembly ATPase SufC | 1503 | gene | 1608965 | 1609714 | - | 1933,288 | 1,63703599 | 0,405925 | 4,032855 | 5,51E-05 | 0,001082 | 2049,911528 | 0,850253448 | 0,179756 | 4,730047 | 2,24E-06 | 1,07E-05 | 1816,664 | 2,941787131 | 0,284544 | 10,3386 | 4,71E-25 | 1,23E-22 | 1005,152 | -1,8676 | 0,20505 | -9,10801 | 8,39E-20 | 5,82E-18 | 2857,835481 | 0,270030041 | 0,226594 | 1,19169 | 0,233383 | 0,289988 |
| 1433 | RM25_RS06985 | RM25_1416 | Dioxygenase | 1504 | gene | 1609719 | 1610066 | - | 867,8873 | 1,54690611 | 0,384876 | 4,019233 | 5,84E-05 | 0,001101 | 848,3826204 | 0,685366419 | 0,219765 | 3,11863 | 0,001817 | 0,004517 | 887,3919 | 2,713728654 | 0,349052 | 7,774558 | 7,57E-15 | 5,95E-13 | 471,1383 | -1,53185 | 0,244718 | -6,25964 | 3,86E-10 | 6,1E-09 | 1261,045921 | 0,539948041 | 0,284825 | 1,89572 | 0,057997 | 0,084539 |
| 1434 | RM25_RS06990 | RM25_1417 | FeS assembly protein SufD | 1505 | gene | 1610063 | 1611280 | - | 3060,567 | 1,608803691 | 0,37377 | 4,304258 | 1,68E-05 | 0,000429 | 3301,00663 | 0,920541634 | 0,173543 | 5,304395 | 1,13E-07 | 6,76E-07 | 2820,128 | 2,726477158 | 0,283559 | 9,615205 | 6,9E-22 | 1,16E-19 | 1612,035 | -1,68411 | 0,174537 | -9,64902 | 4,96E-22 | 5,09E-20 | 4501,289017 | 0,164537371 | 0,231552 | 0,710585 | 0,477342 | 0,540132 |
| 1435 | RM25_RS06995 | RM25_1418 | FeS assembly protein SufB | 1506 | gene | 1611280 | 1612737 | - | 4439,591 | 1,653049553 | 0,410285 | 4,029032 | 5,6E-05 | 0,001082 | 4918,597502 | 0,950413678 | 0,179113 | 5,306224 | 1,12E-07 | 6,71E-07 | 3960,585 | 2,911440058 | 0,285866 | 10,18464 | 2,32E-24 | 4,98E-22 | 2287,141 | -1,9114 | 0,188237 | -10,1542 | 3,17E-24 | 4,67E-22 | 6579,361116 | 0,091647452 | 0,232036 | 0,394971 | 0,692864 | 0,742985 |
| 1436 | RM25_RS07000 | RM25_1419 | Iron-sulfur cluster biosynthesis transcriptional regulator SufI | 1507 | gene | 1612734 | 1613498 | - | 1532,989 | 1,465722735 | 0,38332 | 3,823756 | 0,000131 | 0,002093 | 1838,933026 | 1,042784723 | 0,262933 | 3,965975 | 7,31E-05 | 0,000258 | 1227,045 | 2,243663045 | 0,336779 | 6,662129 | 2,7E-11 | 1,02E-09 | 869,7853 | -1,54964 | 0,324966 | -4,76863 | 1,85E-06 | 1,36E-05 | 2199,875686 | -0,306549209 | 0,268243 | -1,14281 | 0,25312 | 0,311217 |
| 1437 | RM25_RS07005 | RM25_1420 | Transcriptional regulator | 1508 | gene | 1613767 | 1614141 | + | 185,6452 | -1,003774034 | 0,304442 | -3,2971 | 0,000977 | 0,009772 | 139,1806898 | -0,939720646 | 0,251935 | -3,73001 | 0,000191 | 0,000615 | 232,1097 | -1,048226805 | 0,359169 | -2,91848 | 0,003517 | 0,013726 | 260,6105 | 0,707688 | 0,246417 | 2,871907 | 0,00408 | 0,011156 | 121,333435 | 0,662053543 | 0,389958 | 1,697757 | 0,089554 | 0,124972 |
| 1438 | RM25_RS07010 | RM25_1421 | ABC transporter, ATP-binding protein | 1509 | gene | 1614138 | 1614836 | + | 486,0749 | -1,524642462 | 0,424778 | -3,58927 | 0,000332 | 0,004271 | 318,5040805 | -1,909976475 | 0,311553 | -6,13049 | 8,76E-10 | 7,51E-09 | 653,6457 | -1,356797184 | 0,35789 | -3,7911 | 0,00015 | 0,00104 | 756,4603 | 0,834697 | 0,343321 | 2,431244 | 0,015047 | 0,032869 | 246,8211125 | 1,45739525 | 0,359576 | 4,053098 | 5,05E-05 | 0,000133 |
| 1439 | RM25_RS07015 | RM25_1422 | Hypothetical protein | 1510 | gene | 1614833 | 1615648 | + | 323,6006 | -0,995519079 | 0,26098 | -3,81454 | 0,000136 | 0,002138 | 257,3426514 | -1,024472678 | 0,226216 | -4,52873 | 5,93E-06 | 2,6E-05 | 389,8586 | -0,981063342 | 0,308547 | -3,17963 | 0,001475 | 0,006684 | 453,8731 | 0,516952 | 0,238975 | 2,163205 | 0,030525 | 0,057421 | 212,5532983 | 0,62791069 | 0,322301 | 1,948211 | 0,05139 | 0,075609 |
| 1440 | RM25_RS07020 | RM25_1423 | Glutamine amidotransferase subunit PdxT | 1511 | gene | 1615800 | 1616420 | - | 777,8919 | -0,527338485 | 0,387487 | -1,36092 | 0,173539 | 0,277121 | 637,9966481 | -1,534754343 | 0,210255 | -7,2995 | 2,89E-13 | 4,08E-12 | 917,7872 | 0,09245272 | 0,391092 | 0,236396 | 0,813125 | 0,865899 | 975,7247 | -0,14923 | 0,350746 | -0,42546 | 0,670503 | 0,722953 | 627,0523605 | 1,53646397 | 0,307616 | 4,994749 | 5,89E-07 | 2,15E-06 |
| 1441 | RM25_RS07025 | RM25_1424 | Pyridoxal biosynthesis lyase PdxS | 1512 | gene | 1616425 | 1617315 | - | 2485,045 | -1,558857319 | 0,578876 | -2,69291 | 0,007083 | 0,037944 | 1063,894154 | -1,827549838 | 0,170373 | -10,7267 | 7,62E-27 | 5,45E-25 | 3906,196 | -1,489720167 | 0,410208 | -3,63162 | 0,000282 | 0,001756 | 3898,893 | 1,740037 | 0,393065 | 4,426841 | 9,56E-06 | 5,9E-05 | 1238,491718 | 2,134619667 | 0,248945 | 8,574666 | 9,94E-18 | 1,71E-16 |
| 1442 | RM25_RS07030 | RM25_1425 | HTH-type pyridoxine biosynthesis transcriptional regulator p | 1513 | gene | 1617411 | 1618841 | + | 393,6047 | -1,077961205 | 0,642157 | -1,67866 | 0,093219 | 0,184258 | 186,7211085 | -2,350473018 | 0,360624 | -6,5178 | 7,13E-11 | 7,13E-10 | 600,4882 | -0,766180855 | 0,434772 | -1,76226 | 0,078026 | 0,142452 | 557,9372 | 1,20692 | 0,377533 | 3,196857 | 0,001389 | 0,004665 | 249,6018847 | 2,865468937 | 0,432914 | 6,619032 | 3,62E-11 | 2,5E-10 |
| 1443 | RM25_RS07035 | RM25_1426 | Glucose-6-phosphate 1-dehydrogenase Zwf | 1514 | gene | 1618994 | 1620589 | + | 2028,884 | 0,246465196 | 0,205247 | 1,20082 | 0,229821 | 0,33729 | 2336,351426 | 0,360661574 | 0,162345 | 2,221582 | 0,026312 | 0,045533 | 1721,416 | 0,093922793 | 0,269227 | 0,348861 | 0,727194 | 0,798264 | 1965,419 | -0,35082 | 0,239149 | -1,46697 | 0,142384 | 0,198579 | 2155,450392 | -0,578330991 | 0,180187 | -3,20961 | 0,001329 | 0,002734 |
| 1444 | RM25_RS07040 | RM25_1427 | Putative OpcA protein | 1515 | gene | 1620600 | 1621502 | + | 853,9439 | 0,235026886 | 0,120072 | 1,957386 | 0,050302 | 0,125197 | 876,1973018 | 0,385636499 | 0,180314 | 2,138696 | 0,03246 | 0,054806 | 831,6905 | 0,079548049 | 0,173481 | 0,45854 | 0,646564 | 0,727771 | 828,0748 | 0,035926 | 0,197631 | 0,181783 | 0,855753 | 0,886208 | 904,5495034 | -0,225925387 | 0,143174 | -1,57798 | 0,114571 | 0,155824 |
| 1445 | RM25_RS07045 | RM25_1428 | 6-phosphogluconolactonase devB | 1516 | gene | 1621499 | 1622257 | + | 643,4816 | -0,579897319 | 0,146909 | -3,94733 | 7,9E-05 | 0,001411 | 605,346865 | -0,520652577 | 0,157006 | -3,31614 | 0,000913 | 0,002473 | 681,6163 | -0,633070781 | 0,262639 | -2,41042 | 0,015934 | 0,043318 | 811,6353 | 0,152341 | 0,221318 | 0,688334 | 0,491242 | 0,559622 | 506,5341053 | 0,100188188 | 0,201958 | 0,496084 | 0,619835 | 0,676366 |
| 1446 | RM25_RS07050 | RM25_1429 | Electron transport protein | 1517 | gene | 1622574 | 1622930 | - | 2339,979 | 0,885230722 | 0,597725 | 1,480999 | 0,138607 | 0,238346 | 3643,746566 | 0,652893295 | 0,197367 | 3,308014 | 0,00094 | 0,002543 | 1036,211 | 1,835182773 | 0,315919 | 5,809027 | 6,28E-09 | 1,47E-07 | 1760,626 | -2,69681 | 0,260647 | -10,3466 | 4,34E-25 | 7,3E-23 | 2974,677418 | -1,478696911 | 0,239881 | -6,16428 | 7,08E-10 | 4,04E-09 |
| 1447 | RM25_RS07055 | RM25_1430 | Hypothetical protein | 1518 | gene | 1623003 | 1623263 | - | 3248,201 | 0,766426395 | 0,525169 | 1,45939 | 0,144458 | 0,244955 | 4753,712957 | 0,51626757 | 0,259426 | 1,990041 | 0,046586 | 0,074901 | 1742,688 | 1,527813074 | 0,384796 | 3,970445 | 7,17E-05 | 0,00056 | 2573,838 | -2,17686 | 0,356246 | -6,11053 | 9,93E-10 | 1,41E-08 | 4002,230566 | -1,134245608 | 0,285891 | -3,96741 | 7,27E-05 | 0,00019 |
| 1448 | RM25_RS07060 | RM25_1431 | Triosephosphate isomerase 2 | 1519 | gene | 1623386 | 1624174 | - | 3653,283 | 0,979675322 | 0,506222 | 1,935269 | 0,052957 | 0,128947 | 4950,159477 | 0,536638241 | 0,235861 | 2,27523 | 0,022892 | 0,040146 | 2356,407 | 2,130759809 | 0,404143 | 5,272292 | 1,35E-07 | 2,3E-06 | 2628,082 | -2,25321 | 0,403288 | -5,58711 | 2,31E-08 | 2,43E-07 | 4744,178762 | -0,627744061 | 0,230081 | -2,72837 | 0,006365 | 0,011522 |
| 1449 | RM25_RS07065 | RM25_1432 | Phosphoglycerate kinase | 1520 | gene | 1624177 | 1625388 | - | 4524,96 | 0,844650494 | 0,47777 | 1,767901 | 0,077078 | 0,164707 | 6023,902238 | 0,385490598 | 0,207048 | 1,861844 | 0,062625 | 0,094681 | 3026,017 | 1,942944191 | 0,400481 | 4,851524 | 1,23E-06 | 1,66E-05 | 3460,932 | -2,11407 | 0,358961 | -5,88941 | 3,88E-09 | 4,66E-08 | 5683,025629 | -0,525793647 | 0,257239 | -2,04399 | 0,040955 | 0,061958 |
| 1450 | RM25_RS07070 | RM25_1433 | Glyceraldehyde-3-phosphate dehydrogenase, type I | 1521 | gene | 1625552 | 1626556 | - | 35153,95 | 0,494324684 | 0,342544 | 1,443097 | 0,148993 | 0,249238 | 38166,67348 | 0,107993553 | 0,231049 | 0,467405 | 0,64021 | 0,696873 | 32141,23 | 0,978593225 | 0,504163 | 1,941026 | 0,052255 | 0,103937 | 31082,52 | -0,80859 | 0,475457 | -1,70066 | 0,089008 | 0,134827 | 40119,50743 | 0,086213545 | 0,283924 | 0,30365 | 0,761394 | 0,803675 |
| 1451 | RM25_RS07075 | RM25_1434 | Phosphotransferase system, mannitol-specific IIBC compon | 1522 | gene | 1626670 | 1628049 | - | 967,1823 | 1,339786102 | 0,294949 | 4,542436 | 5,56E-06 | 0,000172 | 1242,991339 | 1,872544526 | 0,153803 | 12,17498 | 4,23E-34 | 7,66E-32 | 691,3733 | 0,546684814 | 0,20887 | 2,617349 | 0,008862 | 0,027849 | 577,376 | 0,013854 | 0,177315 | 0,078134 | 0,937722 | 0,950219 | 1361,548035 | -1,254486223 | 0,169153 | -7,4163 | 1,2E-13 | 1,2E-12 |
| 1452 | RM25_RS07080 | RM25_1435 | ABC transporter, ATP-binding protein | 1523 | gene | 1628046 | 1628771 | - | 2403,651 | 2,171706337 | 0,499659 | 4,346374 | 1,38E-05 | 0,000384 | 3766,611837 | 2,326345566 | 0,182332 | 12,75886 | 2,78E-37 | 7,29E-35 | 1040,69 | 1,68810587 | 0,262407 | 6,433146 | 1,25E-10 | 4,21E-09 | 931,2763 | -1,40476 | 0,232658 | -6,03787 | 1,56E-09 | 2,14E-08 | 3858,760547 | -1,999256522 | 0,196857 | -10,1559 | 3,12E-24 | 1,07E-22 |
| 1453 | RM25_RS07085 | RM25_1436 | Putative sporulation transcription regulator WhiA | 1524 | gene | 1628834 | 1629817 | - | 737,4884 | 0,158483259 | 0,180918 | 0,875997 | 0,381032 | 0,49181 | 853,7259144 | 0,179887129 | 0,164639 | 1,092614 | 0,274563 | 0,340782 | 621,251 | 0,131709833 | 0,179512 | 0,733709 | 0,463126 | 0,565883 | 738,4943 | -0,49158 | 0,17019 | -2,88844 | 0,003872 | 0,010761 | 762,8068768 | -0,488786682 | 0,166393 | -2,93755 | 0,003308 | 0,006314 |
| 1454 | RM25_RS07090 | RM25_1437 | UPF0052 protein | 1525 | gene | 1629861 | 1630829 | - | 830,6705 | 0,230770552 | 0,14447 | 1,597355 | 0,110187 | 0,204657 | 875,9522558 | 0,270027358 | 0,191196 | 1,412306 | 0,15786 | 0,212493 | 785,3888 | 0,187835561 | 0,22672 | 0,828493 | 0,407392 | 0,51246 | 809,1851 | -0,16867 | 0,23609 | -0,71443 | 0,474961 | 0,546623 | 877,9002923 | -0,20735308 | 0,174456 | -1,18857 | 0,234609 | 0,291346 |
| 1455 | RM25_RS07095 | RM25_1438 | UPF0042 nucleotide-binding protein | 1526 | gene | 1630826 | 1631710 | - | 947,3848 | 0,04942941 | 0,171863 | 0,287609 | 0,773646 | 0,838769 | 928,9107124 | 0,378100306 | 0,16863 | 2,242194 | 0,024949 | 0,043302 | 965,8588 | -0,262767083 | 0,248714 | -1,0565 | 0,290739 | 0,394202 | 983,0097 | 0,329599 | 0,259127 | 1,271961 | 0,203387 | 0,267364 | 944,0336732 | -0,268519293 | 0,149779 | -1,79277 | 0,07301 | 0,10364 |
| 1456 | RM25_RS07100 | RM25_1439 | UvrABC system protein C | 1527 | gene | 1631906 | 1633900 | - | 1260,638 | 0,212322982 | 0,13579 | 1,563614 | 0,117908 | 0,213777 | 1250,973626 | 0,498148415 | 0,127204 | 3,916132 | 9E-05 | 0,00031 | 1270,302 | -0,063788679 | 0,206054 | -0,30957 | 0,756886 | 0,821732 | 1229,942 | 0,262006 | 0,156628 | 1,672795 | 0,094368 | 0,141581 | 1328,854039 | -0,24214021 | 0,173493 | -1,39567 | 0,162813 | 0,213788 |
| 1457 | RM25_RS07105 | RM25_1440 | Histidine kinase | 1528 | gene | 1634042 | 1635361 | + | 651,8564 | -0,108225892 | 0,16478 | -0,65679 | 0,511316 | 0,608673 | 699,2870629 | 0,029757361 | 0,127583 | 0,23324 | 0,815575 | 0,853981 | 604,4257 | -0,266178882 | 0,286194 | -0,93007 | 0,352337 | 0,458564 | 713,9308 | -0,13105 | 0,193464 | -0,67739 | 0,498156 | 0,565312 | 616,6798489 | -0,364691644 | 0,240653 | -1,51543 | 0,129665 | 0,173944 |
| 1458 | RM25_RS07110 | RM25_1441 | Two-component system response regulator | 1529 | gene | 1635364 | 1636017 | + | 341,0003 | -0,254157285 | 0,176975 | -1,43612 | 0,150969 | 0,251295 | 344,4203153 | 0,04635211 | 0,175041 | 0,264807 | 0,791158 | 0,834153 | 337,5802 | -0,562199595 | 0,278258 | -2,02042 | 0,043339 | 0,09064 | 390,5596 | 0,185252 | 0,195436 | 0,947891 | 0,343185 | 0,418678 | 306,0237159 | -0,360792093 | 0,266734 | -1,35263 | 0,176174 | 0,22828 |
| 1459 | RM25_RS07115 | RM25_1442 | Na+-efflux ABC transporter (ATP-binding protein) | 1530 | gene | 1636120 | 1637073 | + | 267,8519 | -1,216520247 | 0,407667 | -2,9841 | 0,002844 | 0,020131 | 187,0312023 | -1,757221242 | 0,274072 | -6,41154 | 1,44E-10 | 1,38E-09 | 348,6726 | -0,961745642 | 0,394881 | -2,43553 | 0,01487 | 0,040897 | 394,5284 | 0,60256 | 0,267316 | 2,254113 | 0,024189 | 0,047649 | 158,714912 | 1,473975065 | 0,428047 | 3,443492 | 0,000574 | 0,001276 |
| 1460 | RM25_RS07120 | RM25_1443 | ABC transporter (Permease) (Na+ exclusion) | 1531 | gene | 1637070 | 1638236 | + | 394,4349 | -1,213751198 | 0,350282 | -3,46507 | 0,00053 | 0,006342 | 288,3934818 | -1,494045246 | 0,27709 | -5,39191 | 6,97E-08 | 4,37E-07 | 500,4762 | -1,064947229 | 0,348425 | -3,05646 | 0,00224 | 0,009512 | 579,3354 | 0,602725 | 0,306943 | 1,96364 | 0,049572 | 0,084362 | 233,8250197 | 1,100701853 | 0,348091 | 3,16211 | 0,001566 | 0,003172 |
| 1461 | RM25_RS07125 | RM25_1444 | Acetyltransferase | 1532 | gene | 1638378 | 1638914 | + | 617,5907 | -0,427295758 | 0,208635 | -2,04806 | 0,040554 | 0,109562 | 584,6220226 | 0,128198528 | 0,151624 | 0,8455 | 0,397831 | 0,467209 | 650,5594 | -0,9493049 | 0,223334 | -4,25061 | 2,13E-05 | 0,000197 | 746,1217 | 0,562377 | 0,213237 | 2,637332 | 0,008356 | 0,020346 | 517,314618 | -0,463672766 | 0,18157 | -2,55369 | 0,010659 | 0,018541 |
| 1462 | RM25_RS07130 | RM25_1445 | UvrABC system protein A | 1533 | gene | 1638929 | 1641994 | - | 3016,253 | 0,676825072 | 0,131539 | 5,145414 | 2,67E-07 | 1,37E-05 | 3270,355261 | 0,605174919 | 0,117476 | 5,151458 | 2,58E-07 | 1,46E-06 | 2762,15 | 0,762869253 | 0,203587 | 3,747135 | 0,000179 | 0,001211 | 2454,724 | -0,40151 | 0,150538 | -2,66716 | 0,007649 | 0,018919 | 3634,395487 | -0,194578596 | 0,126107 | -1,54297 | 0,122838 | 0,165729 |
| 1463 | RM25_RS07135 | RM25_1446 | TIGR03083 family protein | 1534 | gene | 1642206 | 1642952 | + | 1599,791 | 0,417752681 | 0,291469 | 1,433268 | 0,151781 | 0,252291 | 2053,583869 | 0,641902748 | 0,167238 | 3,838254 | 0,000124 | 0,000413 | 1145,998 | 0,029177772 | 0,204257 | 0,142848 | 0,88641 | 0,91836 | 1451,844 | -0,55398 | 0,216701 | -2,55641 | 0,010576 | 0,024779 | 1793,983531 | -1,124967276 | 0,131011 | -8,58682 | 8,94E-18 | 1,56E-16 |
| 1464 | RM25_RS07140 | RM25_1447 | Metallo-beta-lactamase domain protein | 1535 | gene | 1642954 | 1643547 | + | 723,0689 | 0,568964071 | 0,199027 | 2,858735 | 0,004253 | 0,026522 | 846,7991366 | 0,659677259 | 0,205642 | 3,207894 | 0,001337 | 0,003456 | 599,3387 | 0,444945747 | 0,156528 | 2,842594 | 0,004475 | 0,01648 | 615,9716 | -0,42519 | 0,218538 | -1,94562 | 0,051701 | 0,087354 | 846,9865536 | -0,593033527 | 0,146021 | -4,06128 | 4,88E-05 | 0,000129 |
| 1465 | RM25_RS07145 | RM25_1448 | Integral membrane protein, TerC family | 1536 | gene | 1643582 | 1644616 | - | 938,1139 | 0,400032328 | 0,107988 | 3,704409 | 0,000212 | 0,003045 | 893,6022467 | 0,409115397 | 0,128208 | 3,191033 | 0,001418 | 0,003628 | 982,6255 | 0,393317145 | 0,211668 | 1,858179 | 0,063144 | 0,121036 | 853,1191 | 0,082632 | 0,156534 | 0,527886 | 0,597577 | 0,658788 | 1044,848953 | 0,118503435 | 0,155264 | 0,763239 | 0,445321 | 0,510496 |
| 1466 | RM25_RS07150 | RM25_1449 | UvrABC system protein B | 1537 | gene | 1644807 | 1646891 | - | 1802,166 | 0,630217663 | 0,240411 | 2,621419 | 0,008756 | 0,042939 | 1414,920474 | 0,32118341 | 0,172599 | 1,860864 | 0,062763 | 0,094829 | 2189,411 | 0,835991197 | 0,237244 | 3,523763 | 0,000425 | 0,002458 | 1488,548 | 0,258244 | 0,228752 | 1,128927 | 0,258929 | 0,330068 | 2140,148806 | 0,822705331 | 0,142021 | 5,792855 | 6,92E-09 | 3,36E-08 |
| 1467 | RM25_RS07155 | RM25_1450 | Dephospho-CoA kinase | 1538 | gene | 1646966 | 1647634 | - | 452,7179 | 0,108329744 | 0,233094 | 0,464746 | 0,642113 | 0,726926 | 538,133389 | 0,273009324 | 0,178836 | 1,526587 | 0,126864 | 0,176308 | 367,3023 | -0,127230729 | 0,263128 | -0,48353 | 0,628718 | 0,713475 | 460,4705 | -0,40744 | 0,216041 | -1,88592 | 0,059306 | 0,097682 | 461,2231381 | -0,748557083 | 0,220107 | -3,40088 | 0,000672 | 0,00148 |
| 1468 | RM25_RS07160 | RM25_1451 | Phosphorylase, Pnp/Udp family | 1539 | gene | 1647638 | 1648414 | - | 720,0491 | 0,36342272 | 0,324494 | 1,119969 | 0,262727 | 0,370284 | 952,900959 | 0,331035066 | 0,171986 | 1,92478 | 0,054257 | 0,084523 | 487,1972 | 0,427463211 | 0,231214 | 1,848775 | 0,06449 | 0,122853 | 668,6011 | -1,08236 | 0,207333 | -5,22041 | 1,79E-07 | 1,59E-06 | 794,2935438 | -0,936443926 | 0,161379 | -5,80275 | 6,52E-09 | 3,19E-08 |
| 1469 | RM25_RS07165 | RM25_1452 | 30S ribosomal protein S1 | 1540 | gene | 1648531 | 1649985 | - | 11474,5 | 0,939578068 | 0,580819 | 1,617677 | 0,105732 | 0,199677 | 16727,94903 | 0,546105054 | 0,187078 | 2,919135 | 0,00351 | 0,008085 | 6221,057 | 2,276025685 | 0,415978 | 5,471508 | 4,46E-08 | 8,48E-07 | 8437,34 | -2,72634 | 0,393126 | -6,93503 | 4,06E-12 | 9,57E-11 | 14768,85477 | -0,962334886 | 0,223561 | -4,30457 | 1,67E-05 | 4,78E-05 |
| 1470 | RM25_RS07170 | RM25_1453 | Succinate-semialdehyde dehydrogenase | 1541 | gene | 1650348 | 1651814 | - | 6136,392 | 0,03233116 | 0,231783 | 0,139489 | 0,889064 | 0,925584 | 6174,273206 | 0,16041638 | 0,226833 | 0,707201 | 0,479442 | 0,546707 | 6098,511 | -0,096993743 | 0,374344 | -0,2591 | 0,795555 | 0,852329 | 6424,781 | 0,065608 | 0,415766 | 0,1578 | 0,874614 | 0,90099 | 6074,977939 | -0,157533261 | 0,128661 | -1,2244 | 0,2208 | 0,276823 |
| 1471 | RM25_RS07175 | RM25_1454 | Sortase family protein | 1542 | gene | 1652034 | 1652747 | + | 221,5727 | -0,308232189 | 0,225965 | -1,36407 | 0,172545 | 0,276064 | 188,1872923 | -0,483355579 | 0,225539 | -2,14312 | 0,032104 | 0,054281 | 254,9581 | -0,186454286 | 0,294383 | -0,63337 | 0,52649 | 0,625788 | 258,5991 | 0,24019 | 0,229465 | 1,046741 | 0,295219 | 0,369282 | 194,6187622 | 0,603534882 | 0,306548 | 1,968808 | 0,048975 | 0,072646 |

| | | | | | | | | | | | | | | | | | | | | | | | | | | | | | | | | | | | | | | |
|---|---|---|---|---|---|---|---|---|---|---|---|---|---|---|---|---|---|---|---|---|---|---|---|---|---|---|---|---|---|---|---|---|---|---|---|---|---|---|
| 1472 | RM25_RS07180 | RM25_1455 | DNA-directed DNA polymerase | 1543 | gene | 1652767 | 1655445 | - | 2554,252 | -0,070531929 | 0,139403 | -0,50596 | 0,612887 | 0,70125 | 2841,573648 | 0,062424087 | 0,117454 | 0,531475 | 0,59509 | 0,655739 | 2266,93 | -0,236528474 | 0,144405 | -1,63796 | 0,101431 | 0,174505 | 2765,986 | -0,24052 | 0,115975 | -2,07389 | 0,038089 | 0,068323 | 2444,937428 | -0,486111381 | 0,123461 | -3,93737 | 8,24E-05 | 0,000213 |
| 1473 | RM25_RS07185 | RM25_1456 | Putative transcriptional regulatory protein pdtaR | 1544 | gene | 1655552 | 1656175 | - | 1412,869 | 0,423827139 | 0,356236 | 1,189738 | 0,234149 | 0,341516 | 1828,532818 | 0,720583171 | 0,209535 | 3,438971 | 0,000584 | 0,001658 | 997,2043 | -0,099173324 | 0,391454 | -0,25335 | 0,800001 | 0,856313 | 1279,989 | -0,46831 | 0,402418 | -1,16374 | 0,24453 | 0,313237 | 1587,297588 | -1,253318481 | 0,180162 | -6,9566 | 3,49E-12 | 2,73E-11 |
| 1475 | RM25_RS07195 | RM25_1458 | Pyruvate kinase | 1548 | gene | 1656711 | 1658144 | - | 3073,141 | 0,814684343 | 0,349542 | 2,330717 | 0,019768 | 0,07121 | 3899,767646 | 0,558640815 | 0,177784 | 3,142244 | 0,001677 | 0,004204 | 2246,514 | 1,295180034 | 0,344368 | 3,761032 | 0,000169 | 0,001159 | 2374,794 | -1,32999 | 0,284499 | -4,67486 | 2,94E-06 | 2,05E-05 | 3832,524671 | -0,559072624 | 0,242663 | -2,30391 | 0,021228 | 0,034746 |
| 1476 | RM25_RS07200 | RM25_1459 | Pseudouridine synthase (Precursor) | 1549 | gene | 1658460 | 1659380 | - | 681,9498 | 0,444427828 | 0,253535 | 1,752926 | 0,079615 | 0,166802 | 845,6284958 | 0,536014836 | 0,152301 | 3,519439 | 0,000432 | 0,001277 | 518,2712 | 0,29923416 | 0,242906 | 1,231892 | 0,217989 | 0,314978 | 611,1284 | -0,6297 | 0,2108 | -2,98716 | 0,002816 | 0,008401 | 771,8639045 | -0,810533735 | 0,168511 | -4,80996 | 1,51E-06 | 5,05E-06 |
| 1477 | RM25_RS07205 | RM25_1460 | Signal peptidase II | 1550 | gene | 1659377 | 1659922 | - | 726,6699 | 0,685918108 | 0,337578 | 2,031877 | 0,042166 | 0,111921 | 957,8574531 | 0,546771585 | 0,194105 | 2,81689 | 0,004849 | 0,010602 | 495,4823 | 0,965839806 | 0,210005 | 4,599119 | 4,24E-06 | 4,85E-05 | 592,8283 | -1,27504 | 0,230748 | -5,52566 | 3,28E-08 | 3,35E-07 | 878,2840287 | -0,805845677 | 0,148332 | -5,43273 | 5,55E-08 | 2,4E-07 |
| 1478 | RM25_RS07210 | RM25_1461 | C4-type zinc finger protein, DksA/TraR family | 1551 | gene | 1659979 | 1660407 | - | 5291,441 | 1,001995869 | 0,412986 | 2,426225 | 0,015257 | 0,061278 | 6543,876937 | 0,559894989 | 0,217468 | 2,574614 | 0,010035 | 0,01996 | 4039,004 | 1,845719442 | 0,427575 | 4,316715 | 1,58E-05 | 0,000153 | 3761,539 | -1,63525 | 0,412059 | -3,96849 | 7,23E-05 | 0,000367 | 6899,56243 | -0,321413139 | 0,237829 | -1,35145 | 0,176553 | 0,228645 |
| 1479 | RM25_RS07215 | RM25_1462 | DivIVA domain protein | 1552 | gene | 1660482 | 1661480 | - | 6692,765 | 0,692196132 | 0,409973 | 1,688395 | 0,091335 | 0,182284 | 9128,32589 | 0,761496229 | 0,214122 | 3,556362 | 0,000376 | 0,001133 | 4257,205 | 0,546727883 | 0,451841 | 1,210001 | 0,226278 | 0,323628 | 5451,59 | -1,01665 | 0,438606 | -2,31791 | 0,020454 | 0,041597 | 8097,329285 | -1,201679443 | 0,23687 | -5,07316 | 3,91E-07 | 1,46E-06 |
| 1480 | RM25_RS07220 | RM25_1463 | Hypothetical protein | 1553 | gene | 1661617 | 1661910 | - | 228,1432 | 0,469223401 | 0,273768 | 1,713944 | 0,086539 | 0,176421 | 286,456596 | 0,541339778 | 0,212947 | 2,542134 | 0,011018 | 0,021569 | 169,8298 | 0,351164405 | 0,244159 | 1,438261 | 0,15036 | 0,235324 | 202,9589 | -0,70374 | 0,176116 | -3,9959 | 6,45E-05 | 0,000332 | 259,5137189 | -0,846976739 | 0,252088 | -3,35985 | 0,00078 | 0,001699 |
| 1481 | RM25_RS07225 | RM25_1464 | Cell division protein sepF | 1554 | gene | 1661914 | 1662435 | - | 2032,404 | 0,521921421 | 0,171045 | 3,051371 | 0,002278 | 0,016884 | 2093,361418 | 0,517089014 | 0,175648 | 2,943886 | 0,003241 | 0,007527 | 1971,447 | 0,527571934 | 0,302689 | 1,74295 | 0,081342 | 0,14714 | 1766,336 | -0,14641 | 0,249445 | -0,58695 | 0,557239 | 0,622765 | 2342,33322 | -0,098096488 | 0,224424 | -0,4371 | 0,662037 | 0,71579 |
| 1482 | RM25_RS07230 | RM25_1465 | YfiH family protein | 1555 | gene | 1662587 | 1663375 | - | 600,3138 | 0,579494479 | 0,247155 | 2,344663 | 0,019044 | 0,069848 | 728,118737 | 0,838107685 | 0,220927 | 3,793591 | 0,000148 | 0,000486 | 472,5088 | 0,200277888 | 0,187599 | 1,067587 | 0,285707 | 0,39061 | 507,6869 | -0,3041 | 0,248756 | -1,22246 | 0,221533 | 0,288323 | 705,7224094 | -0,891385815 | 0,13576 | -6,56588 | 5,17E-11 | 3,51E-10 |
| 1483 | RM25_RS07235 | RM25_1466 | Cell division protein FtsZ | 1556 | gene | 1663385 | 1664626 | - | 4364,611 | 0,526316431 | 0,426113 | 1,235156 | 0,216772 | 0,321949 | 6095,623274 | 0,785732933 | 0,189578 | 4,144634 | 3,4E-05 | 0,00013 | 2633,599 | -0,048331854 | 0,420393 | -0,11497 | 0,90847 | 0,936285 | 3805,374 | -0,78958 | 0,419313 | -1,88304 | 0,059695 | 0,098003 | 5051,444406 | -1,589883164 | 0,185415 | -8,57475 | 9,93E-18 | 1,71E-16 |
| 1484 | RM25_RS07240 | RM25_1467 | POTRA domain protein, FtsQ-type | 1557 | gene | 1664844 | 1665575 | - | 658,889 | 0,361607033 | 0,228228 | 1,584414 | 0,1131 | 0,207775 | 772,6492347 | 0,234560536 | 0,164595 | 1,425073 | 0,154136 | 0,208344 | 545,1288 | 0,545267366 | 0,278789 | 1,955841 | 0,050484 | 0,101595 | 611,9681 | -0,74134 | 0,198914 | -3,72695 | 0,000194 | 0,000876 | 724,6684872 | -0,385175958 | 0,218132 | -1,76579 | 0,07743 | 0,109218 |
| 1485 | RM25_RS07245 | RM25_1468 | UDP-N-acetylmuramate--L-alanine ligase | 1558 | gene | 1665572 | 1667014 | - | 1587,83 | 0,464983734 | 0,251921 | 1,845754 | 0,064928 | 0,146165 | 1947,249869 | 0,394268698 | 0,176299 | 2,23636 | 0,025328 | 0,043928 | 1228,41 | 0,579472131 | 0,222424 | 2,605261 | 0,00918 | 0,028322 | 1415,553 | -0,83297 | 0,243891 | -3,41536 | 0,000637 | 0,002422 | 1804,977257 | -0,597382295 | 0,110685 | -5,39716 | 6,77E-08 | 2,88E-07 |
| 1486 | RM25_RS07250 | RM25_1469 | UDP-N-acetylglucosamine--N-acetylmuramyl-(pentapeptide | 1559 | gene | 1667011 | 1668111 | - | 805,1187 | 0,374201272 | 0,24319 | 1,538718 | 0,123873 | 0,221531 | 984,3659631 | 0,31174534 | 0,162587 | 1,917406 | 0,055186 | 0,085575 | 625,8715 | 0,475559061 | 0,218658 | 2,1749 | 0,029638 | 0,068756 | 744,4174 | -0,80802 | 0,230174 | -3,51048 | 0,000447 | 0,00179 | 891,2022249 | -0,590279327 | 0,120241 | -4,90913 | 9,15E-07 | 3,23E-06 |
| 1487 | RM25_RS07255 | RM25_1470 | Cell division protein FtsW | 1560 | gene | 1668111 | 1669376 | - | 2320,492 | 0,426085723 | 0,358367 | 1,188965 | 0,234454 | 0,341539 | 3142,95507 | 0,34913072 | 0,182518 | 1,912856 | 0,055767 | 0,086228 | 1498,03 | 0,591121832 | 0,246847 | 2,394685 | 0,016635 | 0,044707 | 2106,902 | -1,2662 | 0,242799 | -5,21502 | 1,84E-07 | 1,62E-06 | 2606,674361 | -0,981757913 | 0,159987 | -6,1365 | 8,44E-10 | 4,76E-09 |
| 1488 | RM25_RS07260 | RM25_1471 | UDP-N-acetylmuramoyl-L-alanine--D-glutamate ligase | 1561 | gene | 1669569 | 1670990 | - | 3456,73 | 0,647731785 | 0,398882 | 1,623867 | 0,104404 | 0,198132 | 4727,511522 | 0,449762558 | 0,193699 | 2,32197 | 0,020235 | 0,036049 | 2185,949 | 1,101594629 | 0,279854 | 3,936315 | 8,27E-05 | 0,000623 | 2873,141 | -1,57915 | 0,262379 | -6,01861 | 1,76E-09 | 2,37E-08 | 4131,529209 | -0,888314136 | 0,196409 | -4,52277 | 6,1E-06 | 1,85E-05 |
| 1489 | RM25_RS07265 | RM25_1472 | Phospho-N-acetylmuramoyl-pentappeptidetransferase Mur | 1562 | gene | 1671050 | 1672126 | - | 2065,423 | 0,621287267 | 0,383275 | 1,620998 | 0,105018 | 0,198977 | 2799,719895 | 0,445220538 | 0,174837 | 2,546488 | 0,010881 | 0,021337 | 1331,127 | 1,010639201 | 0,291989 | 3,461219 | 0,000538 | 0,002961 | 1735,078 | -1,48193 | 0,21236 | -6,9784 | 2,99E-12 | 7,25E-11 | 2449,621931 | -0,877922002 | 0,229935 | -3,81813 | 0,000134 | 0,000335 |
| 1490 | RM25_RS07270 | RM25_1473 | UDP-N-acetylmuramoyl-tripeptide--D-alanyl-D-alanine ligase | 1563 | gene | 1672123 | 1673607 | - | 1994,572 | 0,65678262 | 0,290715 | 2,259196 | 0,023871 | 0,079134 | 2535,682166 | 0,541406857 | 0,170026 | 3,184255 | 0,001451 | 0,003694 | 1453,462 | 0,86369807 | 0,203136 | 4,251817 | 2,12E-05 | 0,000197 | 1644,548 | -1,06289 | 0,190382 | -5,58294 | 2,36E-08 | 2,48E-07 | 2392,060172 | -0,691408272 | 0,151486 | -4,56418 | 5,01E-06 | 1,54E-05 |
| 1491 | RM25_RS07275 | RM25_1474 | UDP-N-acetylmuramoyl-L-alanyl-D-glutamate--2, 6-diamino | 1564 | gene | 1673609 | 1675198 | - | 1674,44 | 0,794481405 | 0,240411 | 3,304675 | 0,000951 | 0,009713 | 2018,941541 | 0,707540797 | 0,172521 | 4,101177 | 4,11E-05 | 0,000154 | 1329,938 | 0,930263472 | 0,223604 | 4,160313 | 3,18E-05 | 0,000277 | 1297,877 | -0,80289 | 0,228104 | -3,51985 | 0,000432 | 0,001746 | 2081,195246 | -0,532711375 | 0,122328 | -4,35479 | 1,33E-05 | 3,85E-05 |
| 1492 | RM25_RS07280 | RM25_1475 | Penicillin binding protein transpeptidase domain protein | 1565 | gene | 1675198 | 1677261 | - | 3141,758 | 0,395751839 | 0,223127 | 1,773664 | 0,076119 | 0,163102 | 3677,348348 | 0,195239643 | 0,128428 | 1,520222 | 0,128455 | 0,178099 | 2606,168 | 0,686096103 | 0,243067 | 2,822657 | 0,004763 | 0,017234 | 2879,208 | -0,83873 | 0,175989 | -4,76582 | 1,88E-06 | 1,37E-05 | 3493,268479 | -0,303065358 | 0,163572 | -1,8528 | 0,063911 | 0,092247 |
| 1493 | RM25_RS12605 | RM25_1476 | Hypothetical protein | 1566 | gene | 1677258 | 1677776 | - | 680,6559 | 0,635667009 | 0,222812 | 2,852935 | 0,004332 | 0,026868 | 807,680908 | 0,567298456 | 0,18314 | 3,097629 | 0,001951 | 0,004815 | 553,6308 | 0,739156613 | 0,220726 | 3,348752 | 0,000812 | 0,004124 | 565,1891 | -0,7074 | 0,224646 | -3,14896 | 0,001639 | 0,005327 | 811,1062856 | -0,489556441 | 0,15062 | -3,25028 | 0,001153 | 0,002401 |
| 1494 | RM25_RS07290 | RM25_1477 | Ribosomal RNA small subunit methyltransferase H | 1567 | gene | 1677773 | 1678801 | - | 1884,7 | 0,698983793 | 0,338457 | 2,065209 | 0,038903 | 0,107196 | 2468,315402 | 0,670072048 | 0,219847 | 3,047897 | 0,002304 | 0,005577 | 1301,084 | 0,7561639 | 0,306094 | 2,470362 | 0,013498 | 0,038423 | 1527,825 | -1,02866 | 0,329537 | -3,12154 | 0,001799 | 0,005769 | 2284,289219 | -0,90661326 | 0,170931 | -5,30399 | 1,13E-07 | 4,59E-07 |
| 1495 | RM25_RS07295 | RM25_1478 | Protein MraZ | 1568 | gene | 1678980 | 1679417 | - | 5172,097 | 0,445127728 | 0,350647 | 1,269448 | 0,204281 | 0,308253 | 6354,477 | 0,71142876 | 0,229032 | 3,106237 | 0,001895 | 0,004686 | 3989,717 | 0,034957458 | 0,470082 | 0,074365 | 0,94072 | 0,961525 | 4646,955 | -0,3322 | 0,455559 | -0,72922 | 0,465869 | 0,537471 | 5834,544997 | -0,986062614 | 0,250583 | -3,93508 | 8,32E-05 | 0,000215 |
| 1496 | RM25_RS07300 | RM25_1479 | Putative membrane protein | 1569 | gene | 1679640 | 1680023 | - | 2765,893 | 0,013165468 | 0,370757 | 0,03551 | 0,971673 | 0,981249 | 2284,115469 | 0,869743477 | 0,153884 | 5,651924 | 1,59E-08 | 1,11E-07 | 3247,67 | -0,578203138 | 0,466488 | -1,23948 | 0,215167 | 0,312446 | 2899,731 | 1,220831 | 0,408133 | 2,991257 | 0,002778 | 0,008331 | 2712,565729 | -0,20164378 | 0,27089 | -0,74437 | 0,45665 | 0,521981 |
| 1497 | RM25_RS07305 | RM25_1480 | DNA polymerase IV | 1570 | gene | 1680063 | 1681304 | - | 387,4332 | -1,025462137 | 0,461294 | -2,22301 | 0,026215 | 0,083385 | 245,6902532 | -1,74755838 | 0,305736 | -5,71591 | 1,09E-08 | 7,96E-08 | 529,1762 | -0,734605499 | 0,357556 | -2,05452 | 0,039926 | 0,085862 | 546,0815 | 0,736069 | 0,31972 | 2,302231 | 0,021322 | 0,043213 | 251,2235626 | 1,817643619 | 0,368852 | 4,927834 | 8,31E-07 | 2,95E-06 |
| 1498 | RM25_RS07310 | RM25_1481 | DNA polymerase III, alpha subunit | 1571 | gene | 1681331 | 1684951 | - | 1455,267 | -0,573204942 | 0,357713 | -1,60242 | 0,109064 | 0,203212 | 1006,641738 | -1,013094959 | 0,236476 | -4,28413 | 1,83E-05 | 7,38E-05 | 1903,892 | -0,35302025 | 0,26396 | -1,3374 | 0,181092 | 0,273963 | 1829,71 | 0,597881 | 0,294216 | 2,032119 | 0,042142 | 0,073905 | 1149,482306 | 1,32445292 | 0,224444 | 5,901043 | 3,61E-09 | 1,83E-08 |
| 1499 | RM25_RS07315 | RM25_1482 | Hypothetical protein | 1572 | gene | 1684963 | 1685451 | - | 421,3636 | 0,711417268 | 0,119831 | 5,936844 | 2,91E-09 | 2,74E-07 | 443,7206096 | 0,676958957 | 0,185358 | 3,652178 | 0,00026 | 0,000813 | 399,0067 | 0,748265863 | 0,189366 | 3,951424 | 7,77E-05 | 0,000596 | 338,0327 | -0,2543 | 0,172314 | -1,47579 | 0,14 | 0,195718 | 513,185683 | -0,128679167 | 0,194843 | -0,66042 | 0,508981 | 0,570185 |
| 1500 | RM25_RS07320 | RM25_1483 | DNA or RNA helicase | 1573 | gene | 1685578 | 1687788 | + | 2930,236 | 0,343032563 | 0,20094 | 1,707139 | 0,087796 | 0,17778 | 3373,013292 | 0,475546327 | 0,135739 | 3,50339 | 0,000459 | 0,00135 | 2487,46 | 0,166602183 | 0,272663 | 0,611018 | 0,541188 | 0,638109 | 2737,894 | -0,32171 | 0,260933 | -1,23292 | 0,217606 | 0,283828 | 3210,07808 | -0,589363776 | 0,141595 | -4,16231 | 3,15E-05 | 8,63E-05 |
| 1501 | RM25_RS07325 | RM25_1484 | Methylenetetrahydrofolate reductase (NAD(P)H) | 1574 | gene | 1687742 | 1688683 | - | 770,0772 | 0,318258327 | 0,230421 | 1,381201 | 0,167217 | 0,269586 | 941,6763232 | 0,373169633 | 0,129095 | 2,89065 | 0,003844 | 0,008705 | 598,4782 | 0,233960359 | 0,207111 | 1,129636 | 0,25863 | 0,360065 | 726,7529 | -0,64089 | 0,185003 | -3,46424 | 0,000532 | 0,002082 | 838,040818 | -0,725177277 | 0,124879 | -5,80706 | 6,36E-09 | 3,12E-08 |
| 1502 | RM25_RS07330 | RM25_1485 | Putative GERANYLGERANYL PYROPHOSPHATE SYNTHETASE | 1575 | gene | 1688729 | 1689799 | + | 695,9933 | -0,307796484 | 0,158693 | -1,93957 | 0,052431 | 0,128063 | 638,8030639 | -0,152279339 | 0,135496 | -1,12387 | 0,26107 | 0,326095 | 753,1836 | -0,440596941 | 0,267269 | -1,64851 | 0,099247 | 0,171752 | 809,8635 | 0,302304 | 0,207376 | 1,457758 | 0,144907 | 0,201473 | 610,9401081 | 0,076079371 | 0,209573 | 0,363021 | 0,716589 | 0,7646 |
| 1503 | NA | RM25_1486 | Hypothetical protein | 1576 | gene | 1689923 | 1692193 | + | 2731,043 | 0,804951524 | 0,222149 | 3,623481 | 0,000291 | 0,003849 | 3185,099848 | 0,632044506 | 0,155154 | 4,073661 | 4,63E-05 | 0,000172 | 2276,986 | 1,05762365 | 0,222874 | 4,745393 | 2,08E-06 | 2,58E-05 | 2110,437 | -0,81641 | 0,218948 | -3,72879 | 0,000192 | 0,000875 | 3401,693457 | -0,344136043 | 0,124192 | -2,77099 | 0,005589 | 0,010258 |
| 1504 | RM25_RS07340 | RM25_1487 | Membrane spanning protein DUF6 | 1577 | gene | 1692238 | 1693161 | + | 479,1592 | -0,292201753 | 0,219149 | -1,33335 | 0,182417 | 0,287581 | 411,9730428 | -0,068317455 | 0,174492 | -0,39152 | 0,695412 | 0,747758 | 546,3454 | -0,464371494 | 0,314918 | -1,47458 | 0,140326 | 0,224033 | 553,6255 | 0,524735 | 0,228065 | 2,30081 | 0,021402 | 0,043301 | 423,5144481 | 0,189933223 | 0,272931 | 0,695903 | 0,48649 | 0,548377 |
| 1505 | RM25_RS07345 | RM25_1488 | Zn-dependent hydrolase of the beta-lactamase fold-like prot | 1578 | gene | 1693174 | 1693830 | - | 763,9676 | -0,230536389 | 0,27031 | -0,85286 | 0,393737 | 0,504694 | 577,9417995 | 0,173016631 | 0,15378 | 1,125091 | 0,26055 | 0,325619 | 949,9934 | -0,480862448 | 0,238433 | -2,01676 | 0,04372 | 0,091274 | 866,5956 | 0,967726 | 0,232193 | 4,167771 | 3,08E-05 | 0,000169 | 688,0142266 | 0,359588311 | 0,151056 | 2,380504 | 0,017289 | 0,028741 |
| 1506 | RM25_RS07350 | RM25_1489 | 3-deoxy-7-phosphoheptulonate synthase | 1579 | gene | 1693949 | 1695295 | - | 2025,457 | 1,038252957 | 0,184243 | 5,635242 | 1,75E-08 | 1,42E-06 | 2038,610576 | 0,665628098 | 0,143073 | 4,65237 | 3,28E-06 | 1,52E-05 | 2012,303 | 1,453269803 | 0,239293 | 6,073184 | 1,25E-09 | 3,4E-08 | 1408,028 | -0,61049 | 0,180788 | -3,37682 | 0,000733 | 0,002722 | 2662,428117 | 0,222257912 | 0,178083 | 1,248057 | 0,21201 | 0,26751 |
| 1507 | RM25_RS11895 | RM25_1490 | Acyltransferase | 1580 | gene | 1695444 | 1696385 | - | 1002,397 | 0,540855831 | 0,185969 | 2,908319 | 0,003634 | 0,02353 | 1126,079057 | 0,454492179 | 0,226065 | 2,010451 | 0,044383 | 0,071898 | 878,7147 | 0,652757039 | 0,159234 | 4,099362 | 4,14E-05 | 0,000348 | 863,8332 | -0,52865 | 0,252124 | -2,09678 | 0,036013 | 0,065699 | 1164,072686 | -0,286146996 | 0,123972 | -2,30816 | 0,02099 | 0,034428 |
| 1508 | RM25_RS07360 | RM25_1491 | 1-acyl-sn-glycerol-3-phosphate acyltransferase | 1581 | gene | 1696378 | 1697223 | - | 1703,646 | 0,402011869 | 0,305451 | 1,316124 | 0,188132 | 0,292692 | 2129,795692 | 0,230807355 | 0,208315 | 1,107972 | 0,267874 | 0,33371 | 1277,496 | 0,695834174 | 0,301512 | 2,307815 | 0,021009 | 0,053132 | 1560,917 | -1,05869 | 0,291209 | -3,63549 | 0,000277 | 0,001198 | 1898,108258 | -0,557276308 | 0,20912 | -2,66486 | 0,007702 | 0,013769 |
| 1509 | RM25_RS11900 | RM25_1492 | Peptidase M23B family / metalloendopeptidase | 1582 | gene | 1697340 | 1698461 | - | 15100,82 | 0,783597169 | 0,460668 | 1,701003 | 0,088942 | 0,179363 | 19932,47279 | 0,462979163 | 0,235555 | 1,965482 | 0,049358 | 0,078184 | 10269,17 | 1,475056541 | 0,511972 | 2,88113 | 0,003963 | 0,015016 | 11841,67 | -1,66931 | 0,479874 | -3,47864 | 0,000504 | 0,001986 | 18670,49576 | -0,634375376 | 0,288089 | -2,20201 | 0,027665 | 0,043939 |
| 1510 | RM25_RS07370 | RM25_1493 | Transcriptional regulator, AsnC family | 1583 | gene | 1698721 | 1698990 | + | 1050,744 | 0,072750696 | 0,368498 | 0,197425 | 0,843495 | 0,89676 | 1261,711364 | -0,139345064 | 0,259831 | -0,53629 | 0,591758 | 0,652678 | 839,7769 | 0,395185498 | 0,513674 | 0,769332 | 0,441696 | 0,545353 | 1091,045 | -0,91155 | 0,503711 | -1,80967 | 0,070347 | 0,112032 | 1053,220843 | -0,348366437 | 0,286964 | -1,21397 | 0,224759 | 0,28074 |
| 1511 | RM25_RS07375 | RM25_1494 | Peptidyl-prolyl cis-trans isomerase | 1584 | gene | 1699164 | 1699676 | - | 6760,935 | 0,880787223 | 0,33349 | 2,641118 | 0,008263 | 0,041177 | 8306,252643 | 0,725589085 | 0,212334 | 3,417205 | 0,000633 | 0,00179 | 5215,617 | 1,139443837 | 0,413608 | 2,754885 | 0,005871 | 0,020381 | 5063,737 | -0,98967 | 0,379827 | -2,60558 | 0,009172 | 0,022082 | 8570,6363 | -0,546837694 | 0,258627 | -2,11439 | 0,034482 | 0,05319 |
| 1512 | RM25_RS11905 | RM25_1495 | Putative regulatory protein | 1585 | gene | 1700003 | 1701019 | + | 7457,192 | 1,946068035 | 0,536761 | 3,625574 | 0,000288 | 0,003849 | 11469,80906 | 1,934371189 | 0,194552 | 9,942684 | 2,71E-23 | 1,45E-21 | 3444,575 | 1,987171593 | 0,413206 | 4,809152 | 1,52E-06 | 2,01E-05 | 3281,612 | -1,82578 | 0,383334 | -4,76291 | 1,91E-06 | 1,38E-05 | 11611,45323 | -1,739318317 | 0,240295 | -7,23826 | 4,54E-13 | 4,15E-12 |
| 1513 | RM25_RS07385 | RM25_1496 | Phosphoribosyl-ATP diphosphatase | 1586 | gene | 1701209 | 1702306 | + | 1294,016 | 0,005354603 | 0,112983 | 0,047393 | 0,9622 | 0,975863 | 1335,39708 | 0,220235044 | 0,121437 | 1,81358 | 0,069742 | 0,10404 | 1252,635 | -0,221291642 | 0,176986 | -1,25034 | 0,211177 | 0,3082 | 1362,18 | 0,065495 | 0,147812 | 0,443098 | 0,657695 | 0,711748 | 1272,694745 | -0,318203985 | 0,144479 | -2,20242 | 0,027636 | 0,043922 |
| 1514 | RM25_RS12760 | RM25_1497 | Hypothetical protein | 1587 | gene | 1702398 | 1703048 | + | 260,5923 | 0,012845692 | 0,319547 | 0,0402 | 0,967934 | 0,97873 | 192,9847156 | 0,173849311 | 0,243561 | 0,713781 | 0,475363 | 0,542581 | 328,1999 | -0,080702402 | 0,364624 | -0,22133 | 0,824835 | 0,871938 | 270,8944 | 0,839671 | 0,289969 | 2,895723 | 0,003783 | 0,010602 | 257,4889773 | 0,645863444 | 0,318527 | 2,027656 | 0,042595 | 0,064234 |
| 1515 | RM25_RS11910 | RM25_1498 | Hypothetical protein | 1588 | gene | 1703074 | 1703757 | + | 265,0239 | -0,681353703 | 0,266026 | -2,56123 | 0,01043 | 0,048394 | 210,5891666 | -0,786370283 | 0,220158 | -3,57184 | 0,000354 | 0,001075 | 319,4586 | -0,616748667 | 0,318658 | -1,93545 | 0,052935 | 0,105045 | 343,8749 | 0,469974 | 0,195718 | 2,401278 | 0,016338 | 0,035069 | 200,1231346 | 0,707007066 | 0,345757 | 2,044807 | 0,040874 | 0,061875 |
| 1516 | RM25_RS07400 | RM25_1499 | Hypothetical protein | 1589 | gene | 1703897 | 1704322 | + | 679,1999 | -0,913614349 | 0,630561 | -1,44889 | 0,147368 | 0,247927 | 233,573057 | -0,469520384 | 0,212803 | -2,20636 | 0,027359 | 0,047138 | 1124,827 | -1,01187673 | 0,411452 | -2,45928 | 0,013921 | 0,039242 | 929,6621 | 2,418047 | 0,40146 | 6,023134 | 1,71E-09 | 2,33E-08 | 459,0223523 | 1,910380481 | 0,230659 | 8,282285 | 1,21E-16 | 1,81E-15 |
| 1517 | RM25_RS07405 | RM25_1500 | Anthranilate phosphoribosyltransferase | 1590 | gene | 1704363 | 1705412 | - | 2177,751 | 0,704694032 | 0,380163 | 1,853661 | 0,063788 | 0,144565 | 2943,138358 | 0,545108203 | 0,198676 | 2,743711 | 0,006075 | 0,012993 | 1412,364 | 1,054629441 | 0,28061 | 3,75835 | 0,000171 | 0,001166 | 1764,913 | -1,43787 | 0,295451 | -4,86669 | 1,13E-06 | 8,6E-06 | 2644,470091 | -0,887008481 | 0,158628 | -5,59175 | 2,25E-08 | 1,02E-07 |
| 1518 | RM25_RS07410 | RM25_1501 | Response regulator | 1591 | gene | 1705412 | 1705936 | - | 2369,688 | 0,549373952 | 0,344294 | 1,595652 | 0,110567 | 0,205136 | 3055,899802 | 0,457072796 | 0,200851 | 2,275677 | 0,022865 | 0,040129 | 1683,477 | 0,72121438 | 0,374536 | 1,92562 | 0,054152 | 0,106898 | 2050,392 | -1,07074 | 0,344576 | -3,1074 | 0,001887 | 0,006004 | 2754,105328 | -0,773813257 | 0,238943 | -3,23849 | 0,001202 | 0,002493 |
| 1519 | RM25_RS07415 | RM25_1502 | IMP dehydrogenase family protein | 1592 | gene | 1706149 | 1707612 | + | 2441,559 | 1,195018821 | 0,291304 | 4,102307 | 4,09E-05 | 0,000846 | 2879,256589 | 0,845829796 | 0,173682 | 4,869993 | 1,12E-06 | 5,62E-06 | 2003,861 | 1,767370943 | 0,219211 | 8,062408 | 7,48E-16 | 6,53E-14 | 1578,986 | -1,23589 | 0,202576 | -6,1009 | 1,05E-09 | 1,49E-08 | 3326,208007 | -0,269684202 | 0,159216 | -1,69382 | 0,090299 | 0,125714 |
| 1520 | RM25_RS07420 | RM25_1503 | Iron-sulfur cluster assembly accessory protein | 1593 | gene | 1707751 | 1708101 | - | 6182,659 | 2,654467955 | 0,523613 | 5,069522 | 3,99E-07 | 1,92E-05 | 6187,391455 | 1,738357281 | 0,209826 | 8,284746 | 1,18E-16 | 2,38E-15 | 6177,927 | 4,465985064 | 0,414841 | 10,76554 | 5,01E-27 | 2,36E-24 | 1815,616 | -2,46574 | 0,39917 | -6,17715 | 6,53E-10 | 9,74E-09 | 10415,89149 | 0,293884595 | 0,228418 | 1,286606 | 0,198232 | 0,252442 |
| 1521 | RM25_RS07425 | RM25_1504 | Hypothetical protein | 1594 | gene | 1708259 | 1709191 | - | 429,2837 | 0,53900207 | 0,148932 | 3,619112 | 0,000296 | 0,00389 | 462,1257323 | 0,699504907 | 0,192004 | 3,643178 | 0,000269 | 0,00084 | 396,4417 | 0,360234807 | 0,2124 | 1,696023 | 0,089882 | 0,158571 | 369,1363 | -0,07919 | 0,212939 | -0,3719 | 0,709969 | 0,75685 | 499,279148 | -0,361351497 | 0,184826 | -1,95509 | 0,050572 | 0,074733 |
| 1522 | RM25_RS07430 | RM25_1505 | Hypothetical protein | 1595 | gene | 1709195 | 1709485 | - | 199,0975 | 0,036413921 | 0,322019 | 0,11308 | 0,909967 | 0,941556 | 156,478947 | 0,017643712 | 0,294723 | 0,059865 | 0,952263 | 0,962059 | 241,716 | 0,051420991 | 0,417494 | 0,123166 | 0,901976 | 0,932437 | 207,7309 | 0,5632 | 0,453156 | 1,242838 | 0,213927 | 0,27966 | 197,122318 | 0,637661658 | 0,232447 | 2,743256 | 0,006083 | 0,011072 |
| 1523 | RM25_RS07435 | RM25_1506 | PspA/IM30 family protein | 1596 | gene | 1709482 | 1710276 | - | 2575,799 | 0,109154461 | 0,26865 | 0,406307 | 0,684517 | 0,763198 | 2365,68785 | 0,222817743 | 0,213009 | 1,04605 | 0,295538 | 0,36418 | 2785,911 | 0,013439824 | 0,435105 | 0,030889 | 0,975358 | 0,985815 | 2621,725 | 0,297346 | 0,406279 | 0,731876 | 0,464245 | 0,535859 | 2610,479604 | 0,117450598 | 0,261299 | 0,449488 | 0,65308 | 0,707728 |
| 1524 | RM25_RS07440 | RM25_1507 | Hypothetical protein | 1597 | gene | 1710576 | 1711130 | + | 1542,368 | 0,596124306 | 0,242141 | 2,461886 | 0,013821 | 0,057861 | 1635,342607 | 0,717583592 | 0,218185 | 3,288883 | 0,001006 | 0,002709 | 1449,394 | 0,462719894 | 0,394219 | 1,173765 | 0,240489 | 0,339828 | 1300,79 | -0,06938 | 0,373576 | -0,18571 | 0,852671 | 0,884571 | 1814,752326 | -0,293460691 | 0,245642 | -1,19467 | 0,232216 | 0,288983 |
| 1525 | RM25_RS07445 | RM25_1508 | Hypothetical protein | 1598 | gene | 1711130 | 1711345 | + | 253,0046 | 0,694626193 | 0,290969 | 2,387286 | 0,016973 | 0,065132 | 313,2457945 | 0,655406682 | 0,195294 | 3,356006 | 0,000791 | 0,002172 | 192,7634 | 0,766649403 | 0,343647 | 2,230922 | 0,025686 | 0,061904 | 205,6802 | -0,82721 | 0,291502 | -2,83775 | 0,004543 | 0,012214 | 306,1091263 | -0,672721753 | 0,263597 | -2,55209 | 0,010708 | 0,018613 |
| 1526 | RM25_RS07450 | RM25_1509 | Aminomethyltransferase | 1599 | gene | 1711499 | 1712620 | - | 2912,276 | -0,248996123 | 0,364509 | -0,6831 | 0,494544 | 0,593248 | 2804,603876 | 0,763032923 | 0,198939 | 3,835511 | 0,000125 | 0,000417 | 3019,948 | -1,24191764 | 0,366387 | -3,38964 | 0,0007 | 0,003641 | 3328,41 | 0,98333 | 0,359939 | 2,731932 | 0,006296 | 0,016061 | 2608,018562 | -0,989639944 | 0,203227 | -4,86964 | 1,12E-06 | 3,85E-06 |
| 1527 | RM25_RS07455 | RM25_1510 | Hypothetical protein | 1600 | gene | 1712921 | 1714690 | + | 5564,189 | 0,89344959 | 0,24648 | 3,624831 | 0,000289 | 0,003849 | 5707,822181 | 0,770811039 | 0,197824 | 3,896452 | 9,76E-05 | 0,000333 | 5420,556 | 1,026384231 | 0,414739 | 2,47477 | 0,013332 | 0,038136 | 4132,469 | -0,28927 | 0,394577 | -0,73312 | 0,463485 | 0,535476 | 7067,419067 | -0,002680523 | 0,230163 | -0,01165 | 0,990708 | 0,991651 |
| 1528 | RM25_RS07460 | RM25_1511 | Hypothetical protein | 1601 | gene | 1714974 | 1715744 | + | 1647,663 | 0,291318943 | 0,225845 | 1,289908 | 0,197083 | 0,299693 | 1840,949442 | 0,463907396 | 0,18329 | 2,531005 | 0,011374 | 0,022046 | 1454,376 | 0,07724924 | 0,341062 | 0,226496 | 0,820816 | 0,870068 | 1569,78 | -0,17778 | 0,340502 | -0,52211 | 0,601596 | 0,662196 | 1775,521168 | -0,529191255 | 0,175242 | -3,01978 | 0,00253 | 0,004948 |
| 1529 | RM25_RS07465 | RM25_1512 | Macrolide-transport ATP-binding protein ABC transporter | 1602 | gene | 1715777 | 1717588 | + | 1317,19 | 0,340595545 | 0,255846 | 1,331252 | 0,183106 | 0,287913 | 1641,803909 | 0,513421147 | 0,129334 | 3,969731 | 7,2E-05 | 0,000254 | 992,5767 | 0,061629091 | 0,215777 | 0,285615 | 0,775173 | 0,836493 | 1231,626 | -0,5422 | 0,194797 | -2,78342 | 0,005379 | 0,014056 | 1445,255357 | -0,933198436 | 0,148356 | -6,29028 | 3,17E-10 | 1,88E-09 |
| 1530 | RM25_RS07470 | RM25_1513 | Glutamate--ammonia ligase | 1603 | gene | 1717782 | 1719203 | + | 5337,849 | -0,554168587 | 0,258379 | -2,14479 | 0,031969 | 0,094072 | 5689,784215 | -1,005355061 | 0,237649 | -4,23042 | 2,33E-05 | 9,22E-05 | 4985,915 | -0,069267441 | 0,333496 | -0,2077 | 0,835462 | 0,879493 | 6751,985 | -0,62442 | 0,38957 | -1,60284 | 0,108969 | 0,159133 | 4235,050115 | 0,352996835 | 0,143796 | 2,454841 | 0,014095 | 0,023849 |
| 1531 | RM25_RS07475 | RM25_1514 | Hypothetical protein | 1604 | gene | 1719448 | 1720038 | + | 407,5676 | -0,032085619 | 0,194184 | -0,16523 | 0,868761 | 0,910885 | 335,4414226 | 0,086281229 | 0,16855 | 0,511903 | 0,608719 | 0,667947 | 479,6937 | -0,112846124 | 0,175443 | -0,64321 | 0,52009 | 0,62037 | 433,7437 | 0,558937 | 0,180887 | 3,089982 | 0,002002 | 0,006291 | 394,8522576 | 0,410110391 | 0,154692 | 2,651138 | 0,008022 | 0,014278 |
| 1532 | RM25_RS07480 | RM25_1515 | Tetracycline repressor protein | 1605 | gene | 1720129 | 1720713 | + | 632,7244 | 0,070660601 | 0,20029 | 0,352791 | 0,724245 | 0,798431 | 738,7548382 | 0,273172484 | 0,164698 | 1,658628 | 0,097191 | 0,138667 | 526,6941 | -0,207541131 | 0,168304 | -1,23313 | 0,217527 | 0,314739 | 652,9395 | -0,30013 | 0,189597 | -1,58299 | 0,113425 | 0,164822 | 635,7766851 | -0,733993491 | 0,132894 | -5,52315 | 3,33E-08 | 1,48E-07 |
| 1533 | RM25_RS07485 | RM25_1516 | Putative oxidoreductase in MprA 5'region | 1606 | gene | 1720733 | 1721548 | - | 203,7563 | 0,026770192 | 0,376561 | 0,071091 | 0,943325 | 0,96173 | 133,6951301 | 0,280155251 | 0,277876 | 1,008202 | 0,313358 | 0,380125 | 273,8175 | -0,095493099 | 0,336087 | -0,28413 | 0,776309 | 0,837036 | 210,4807 | 1,175189 | 0,277951 | 4,228047 | 2,36E-05 | 0,000133 | 202,2654607 | 0,85700928 | 0,321614 | 2,664713 | 0,007705 | 0,013769 |
| 1534 | RM25_RS07490 | RM25_1517 | [glutamate--ammonia-ligase] adenylyltransferase | 1607 | gene | 1721545 | 1724511 | - | 1747,699 | 0,026709403 | 0,20294 | 0,131612 | 0,895291 | 0,930014 | 1423,332816 | 0,117749257 | 0,132748 | 0,887016 | 0,37507 | 0,443795 | 2072,065 | -0,036666847 | 0,198601 | -0,18463 | 0,853523 | 0,894061 | 1819,431 | 0,557869 | 0,165426 | 3,372317 | 0,000745 | 0,002745 | 1729,913349 | 0,459579506 | 0,155647 | 2,9527 | 0,00315 | 0,006041 |
| 1535 | RM25_RS11915 | RM25_1518 | Hypothetical protein | 1608 | gene | 1724753 | 1725490 | + | 426,0218 | -0,287646451 | 0,325992 | -0,88237 | 0,377575 | 0,488443 | 292,3441569 | -0,47604085 | 0,225523 | -2,11083 | 0,034787 | 0,058192 | 559,6994 | -0,193876281 | 0,222412 | -0,8717 | 0,383374 | 0,488703 | 492,0392 | 0,745185 | 0,220888 | 3,373586 | 0,000742 | 0,002737 | 376,1924294 | 1,087666586 | 0,233203 | 4,66404 | 3,1E-06 | 9,9E-06 |
| 1536 | RM25_RS07500 | RM25_1519 | Glutamine synthetase, beta-Grasp domain protein | 1609 | gene | 1725546 | 1726880 | - | 1828,413 | 0,200320281 | 0,174116 | 1,150501 | 0,249938 | 0,358117 | 1665,446812 | 0,468011755 | 0,118985 | 3,933379 | 8,38E-05 | 0,00029 | 1991,378 | -0,020029358 | 0,25873 | -0,07741 | 0,938294 | 0,960295 | 1794,178 | 0,464001 | 0,21561 | 2,152038 | 0,031394 | 0,058727 | 1912,573926 | 0,018516385 | 0,169266 | 0,109392 | 0,912892 | 0,929052 |
| 1537 | RM25_RS11920 | RM25_1520 | Hypothetical protein | 1610 | gene | 1727087 | 1727968 | + | 419,0727 | 0,063431492 | 0,133428 | 0,4754 | 0,634502 | 0,719346 | 460,7886553 | 0,052346121 | 0,157365 | 0,332641 | 0,739405 | 0,787875 | 377,3568 | 0,077352811 | 0,187327 | 0,41293 | 0,679658 | 0,757425 | 433,9452 | -0,36475 | 0,178253 | -2,04625 | 0,040732 | 0,071914 | 420,6708886 | -0,28026636 | 0,177079 | -1,58272 | 0,113486 | 0,154438 |
| 1538 | RM25_RS11925 | RM25_1521 | Hypothetical protein | 1611 | gene | 1728053 | 1728997 | + | 440,4493 | -0,649269262 | 0,227365 | -2,85563 | 0,004295 | 0,026712 | 363,7575132 | -0,727319141 | 0,206616 | -3,52015 | 0,000431 | 0,001275 | 517,1411 | -0,59766216 | 0,268314 | -2,22747 | 0,025916 | 0,06233 | 564,9644 | 0,398927 | 0,222822 | 1,790339 | 0,073399 | 0,115721 | 336,9365616 | 0,586638354 | 0,25035 | 2,343271 | 0,019116 | 0,031573 |
| 1539 | RM25_RS07515 | RM25_1522 | Methionyl aminopeptidase | 1612 | gene | 1729076 | 1729984 | + | 1888,828 | 0,887675018 | 0,187133 | 4,74355 | 2,1E-06 | 7,62E-05 | 2052,477756 | 0,863260089 | 0,143341 | 6,022432 | 1,72E-09 | 1,43E-08 | 1725,178 | 0,917468913 | 0,317308 | 2,891418 | 0,003835 | 0,014603 | 1404,183 | -0,34127 | 0,231867 | -1,47183 | 0,141068 | 0,196976 | 2396,90909 | -0,249572667 | 0,23076 | -1,08153 | 0,279463 | 0,340763 |
| 1540 | RM25_RS11930 | RM25_1523 | S-layer domain protein | 1613 | gene | 1730123 | 1730782 | - | 532,3014 | 0,148916078 | 0,21802 | 0,683038 | 0,494583 | 0,593248 | 621,7664145 | 0,037177855 | 0,204459 | 0,181836 | 0,855712 | 0,886592 | 442,8365 | 0,30873692 | 0,233716 | 1,320992 | 0,186504 | 0,280351 | 536,7913 | -0,69239 | 0,260957 | -2,65327 | 0,007972 | 0,019491 | 548,0628057 | -0,373656469 | 0,175731 | -2,12629 | 0,033479 | 0,051744 |
| 1541 | NA | RM25_1524 | Hypothetical protein | 1614 | gene | 1730840 | 1731550 | + | 515,5849 | -0,531606774 | 0,329785 | -1,61198 | 0,106966 | 0,200978 | 344,0793238 | -0,453325846 | 0,196216 | -2,31035 | 0,020869 | 0,037067 | 687,0904 | -0,571883392 | 0,236152 | -2,42167 | 0,015449 | 0,042146 | 641,6332 | 0,986782 | 0,258914 | 3,811233 | 0,000138 | 0,000651 | 413,1481707 | 0,919258771 | 0,191175 | 4,80846 | 1,52E-06 | 5,09E-06 |
| 1542 | RM25_RS07525 | RM25_1525 | Zn-ribbon protein, possibly nucleic acid-binding | 1615 | gene | 1731361 | 1732104 | - | 1284,123 | 0,425072294 | 0,286847 | 1,481878 | 0,138373 | 0,238346 | 1567,786358 | 0,835307285 | 0,1484 | 5,628761 | 1,82E-08 | 1,25E-07 | 1000,459 | -0,187177881 | 0,29868 | -0,62668 | 0,530867 | 0,629087 | 1161,392 | -0,1331 | 0,289485 | -0,45978 | 0,645674 | 0,700992 | 1442,822086 | -1,115940084 | 0,159574 | -6,99326 | 2,69E-12 | 2,18E-11 |
| 1543 | RM25_RS07530 | RM25_1526 | Xaa-Pro aminopeptidase | 1616 | gene | 1732283 | 1733737 | + | 1807,75 | 0,066804093 | 0,129661 | 0,515222 | 0,606398 | 0,69619 | 1952,928704 | 0,277075058 | 0,130432 | 2,124282 | 0,033647 | 0,056525 | 1662,57 | -0,17734322 | 0,139613 | -1,27025 | 0,203997 | 0,301266 | 1865,782 | -0,05934 | 0,147222 | -0,40305 | 0,686913 | 0,735265 | 1813,593761 | -0,464171605 | 0,110459 | -4,2022 | 2,64E-05 | 7,34E-05 |
| 1544 | RM25_RS12870 | RM25_1527 | Hypothetical protein | 1617 | gene | 1733864 | 1734739 | - | 2157,963 | 0,904103851 | 0,181297 | 4,986869 | 6,14E-07 | 2,68E-05 | 2211,89218 | 0,668615234 | 0,191464 | 3,492122 | 0,000479 | 0,001394 | 2104,035 | 1,165022196 | 0,264061 | 4,411942 | 1,02E-05 | 0,000103 | 1591,675 | -0,45008 | 0,233946 | -1,92359 | 0,054369 | 0,090756 | 2749,099679 | 0,084579351 | 0,199676 | 0,423582 | 0,671871 | 0,723435 |
| 1545 | RM25_RS07540 | RM25_1528 | Citrate lyase beta subunit | 1618 | gene | 1734884 | 1735747 | - | 1079,119 | 0,454283905 | 0,177871 | 2,554009 | 0,010649 | 0,048973 | 1165,628525 | 0,764626559 | 0,180245 | 4,242154 | 2,21E-05 | 8,77E-05 | 992,6096 | 0,105188311 | 0,221115 | 0,475717 | 0,634276 | 0,717709 | 961,6987 | 0,092177 | 0,239269 | 0,385244 | 0,700057 | 0,747298 | 1222,485623 | -0,523433869 | 0,147578 | -3,54683 | 0,00039 | 0,00089 |
| 1546 | RM25_RS07545 | RM25_1529 | MgtE intracellular region | 1619 | gene | 1735928 | 1737205 | + | 1128,401 | 0,130077796 | 0,100834 | 1,290024 | 0,197042 | 0,299693 | 1153,501242 | 0,052156722 | 0,15264 | 0,341698 | 0,732578 | 0,78166 | 1103,302 | 0,211097115 | 0,183277 | 1,151795 | 0,249405 | 0,349286 | 1139,246 | -0,20615 | 0,141826 | -1,45352 | 0,146079 | 0,202534 | 1153,896614 | -0,000160884 | 0,155423 | -0,00104 | 0,999174 | 0,999174 |
| 1547 | RM25_RS07550 | RM25_1530 | PF06210 family protein | 1620 | gene | 1737202 | 1737765 | + | 382,2008 | -0,055430936 | 0,196795 | -0,28167 | 0,778197 | 0,841381 | 331,39721 | -0,385066026 | 0,166933 | -2,3067 | 0,021071 | 0,037314 | 433,0044 | 0,191929266 | 0,231799 | 0,827997 | 0,407672 | 0,512471 | 411,0138 | 0,04162 | 0,210135 | 0,198061 | 0,842997 | 0,876958 | 367,335604 | 0,679335313 | 0,1801 | 3,771986 | 0,000162 | 0,000399 |
| 1548 | RM25_RS07555 | RM25_1531 | Putative ATP-binding protein Mrp | 1621 | gene | 1737909 | 1739066 | + | 3109,568 | 0,988866528 | 0,265217 | 3,728525 | 0,000193 | 0,00282 | 3704,606017 | 0,81447262 | 0,179753 | 4,531057 | 5,87E-06 | 2,58E-05 | 2514,529 | 1,260430269 | 0,288287 | 4,37214 | 1,23E-05 | 0,000123 | 2215,989 | -0,91413 | 0,262111 | -3,48758 | 0,000487 | 0,001927 | 4047,144375 | -0,427577068 | 0,199419 | -2,14411 | 0,032024 | 0,050053 |
| 1549 | RM25_RS07560 | RM25_1532 | Sec-independent protein translocase protein TatB | 1622 | gene | 1739248 | 1739697 | - | 359,8096 | 0,575179203 | 0,358891 | 1,602656 | 0,109011 | 0,203212 | 479,5851599 | 0,455518158 | 0,2262 | 2,013785 | 0,044032 | 0,071476 | 240,034 | 0,827250631 | 0,27251 | 3,035672 | 0,0024 | 0,01003 | 307,6725 | -1,28282 | 0,276538 | -4,63887 | 3,5E-06 | 2,41E-05 | 421,730184 | -0,866739821 | 0,2116 | -4,09613 | 4,2E-05 | 0,000112 |
| 1550 | RM25_RS07565 | RM25_1533 | Dihydrodipicolinate synthase | 1623 | gene | 1739789 | 1740670 | - | 1434,759 | 0,515537248 | 0,16509 | 3,122768 | 0,001792 | 0,014109 | 1394,91163 | 0,277913044 | 0,147393 | 1,885529 | 0,059358 | 0,090731 | 1474,606 | 0,746834502 | 0,280283 | 2,664577 | 0,007709 | 0,025165 | 1249,694 | -0,2517 | 0,249827 | -1,00748 | 0,313705 | 0,388342 | 1650,707461 | 0,259903572 | 0,16658 | 1,560232 | 0,118705 | 0,16089 |
| 1551 | RM25_RS07570 | RM25_1534 | Putative O-methyltransferase | 1624 | gene | 1740814 | 1741479 | + | 2007,314 | 0,079982865 | 0,229895 | 0,347911 | 0,727907 | 0,802093 | 2015,614101 | 0,324537126 | 0,174579 | 1,858973 | 0,063031 | 0,095111 | 1999,014 | -0,164225994 | 0,386725 | -0,42466 | 0,671086 | 0,751068 | 2063,538 | 0,188485 | 0,32592 | 0,578318 | 0,563049 | 0,627771 | 2015,532417 | -0,268599865 | 0,257725 | -1,0422 | 0,297321 | 0,35862 |
| 1552 | RM25_RS07575 | RM25_1535 | Glucose-1-phosphate adenylyltransferase | 1625 | gene | 1741617 | 1742807 | - | 5471,147 | 0,077398908 | 0,421853 | 0,183474 | 0,854426 | 0,904709 | 3561,92746 | 0,62837061 | 0,192061 | 3,271731 | 0,001069 | 0,00286 | 7380,367 | -0,183737571 | 0,45705 | -0,40201 | 0,687678 | 0,765278 | 5596,73 | 1,444317 | 0,411791 | 3,507398 | 0,000453 | 0,001802 | 5476,434619 | 0,65664846 | 0,265963 | 2,468948 | 0,013551 | 0,022995 |
| 1553 | RM25_RS07580 | RM25_1536 | Starch synthase | 1626 | gene | 1742962 | 1744179 | + | 1734,539 | 0,093143187 | 0,248421 | 0,374941 | 0,707705 | 0,783863 | 1418,801514 | 0,318805111 | 0,135146 | 2,358961 | 0,018326 | 0,033329 | 2050,277 | -0,061162688 | 0,330372 | -0,18513 | 0,853125 | 0,894061 | 1769,8 | 0,677759 | 0,29454 | 2,301074 | 0,021387 | 0,043301 | 1749,179379 | 0,333960215 | 0,193017 | 1,730215 | 0,083592 | 0,117177 |
| 1554 | RM25_RS12310 | RM25_1537 | Hypothetical protein | 1627 | gene | 1744358 | 1744525 | - | 2782,51 | 0,317912387 | 0,404677 | 0,785596 | 0,432104 | 0,540016 | 3585,787668 | 0,410985032 | 0,246754 | 1,665567 | 0,0958 | 0,136931 | 1979,232 | 0,151607587 | 0,530542 | 0,28576 | 0,775062 | 0,836493 | 2637,987 | -0,75905 | 0,500424 | -1,51682 | 0,129313 | 0,183609 | 3023,02579 | -0,992584089 | 0,309867 | -3,20326 | 0,001359 | 0,00279 |
| 1555 | RM25_RS11935 | RM25_1538 | Putative membrane protein | 1628 | gene | 1744585 | 1744977 | - | 1872,426 | 0,326984812 | 0,297981 | 1,097334 | 0,272495 | 0,38117 | 2114,171655 | 0,586004159 | 0,215365 | 2,720975 | 0,006509 | 0,013759 | 1630,68 | -0,000302378 | 0,449058 | -0,00067 | 0,999463 | 0,999775 | 1761,536 | -0,09875 | 0,431664 | -0,22877 | 0,819044 | 0,85382 | 2038,235818 | -0,656094568 | 0,248064 | -2,64486 | 0,008173 | 0,014527 |
| 1556 | RM25_RS07590 | RM25_1539 | Chloride transporter, chloride channel family protein | 1629 | gene | 1745000 | 1746448 | - | 489,6012 | -0,164207394 | 0,192368 | -0,85361 | 0,393321 | 0,504694 | 464,6323333 | -0,119398894 | 0,143585 | -0,83155 | 0,405661 | 0,474749 | 514,57 | -0,206148314 | 0,353821 | -0,58263 | 0,560139 | 0,655535 | 545,4575 | 0,122864 | 0,21708 | 0,565985 | 0,571404 | 0,635283 | 454,3671367 | 0,103808078 | 0,314411 | 0,330167 | 0,741274 | 0,785603 |
| 1557 | RM25_RS07595 | RM25_1540 | TIGR00730 family protein | 1630 | gene | 1746540 | 1747376 | - | 2106,593 | 0,532051114 | 0,363783 | 1,462552 | 0,14359 | 0,244186 | 2824,928704 | 0,4404339 | 0,203437 | 2,16496 | 0,030391 | 0,051784 | 1388,258 | 0,72368336 | 0,303833 | 2,381845 | 0,017226 | 0,045982 | 1833,134 | -1,24719 | 0,289133 | -4,31354 | 1,61E-05 | 9,4E-05 | 2438,946134 | -0,927525614 | 0,206098 | -4,50041 | 6,78E-06 | 2,04E-05 |
| 1558 | RM25_RS07600 | RM25_1541 | Succinyl-diaminopimelate desuccinylase | 1631 | gene | 1747538 | 1748626 | - | 776,1834 | -0,334747617 | 0,137121 | -2,44126 | 0,014636 | 0,060205 | 748,9005944 | -0,045073406 | 0,136934 | -0,32916 | 0,742034 | 0,790319 | 803,4663 | -0,608716343 | 0,203896 | -2,98542 | 0,002832 | 0,01151 | 912,7765 | 0,288901 | 0,145258 | 1,988878 | 0,046715 | 0,080546 | 674,3997422 | -0,216204886 | 0,200188 | -1,08001 | 0,280139 | 0,341411 |
| 1559 | RM25_RS07605 | RM25_1542 | 2, 3, 4, 5-tetrahydropyridine-2, 6-dicarboxylate N-succinyltr | 1632 | gene | 1748726 | 1749664 | + | 1636,754 | 0,464736493 | 0,219115 | 2,120971 | 0,033924 | 0,097657 | 1932,631476 | 0,608623973 | 0,17155 | 3,547788 | 0,000388 | 0,001163 | 1340,876 | 0,263035388 | 0,249023 | 1,056271 | 0,290845 | 0,394202 | 1458,119 | -0,38413 | 0,241785 | -1,58873 | 0,112122 | 0,163231 | 1858,643429 | -0,688561077 | 0,167448 | -4,11209 | 3,92E-05 | 0,000106 |
| 1560 | RM25_RS07610 | RM25_1543 | Conserved protein, putative lipoprotein | 1633 | gene | 1749671 | 1750516 | + | 616,6139 | 0,326528501 | 0,335994 | 0,971828 | 0,331136 | 0,441452 | 825,9316845 | 0,549064682 | 0,136897 | 4,010781 | 6,05E-05 | 0,000218 | 407,2961 | -0,110867426 | 0,217659 | -0,50936 | 0,610498 | 0,697839 | 580,5569 | -0,72666 | 0,162647 | -4,46768 | 7,91E-06 | 4,98E-05 | 673,9602593 | -1,326772277 | 0,1909 | -6,95011 | 3,65E-12 | 2,85E-11 |
| 1561 | RM25_RS07615 | RM25_1544 | Succinyldiaminopimelate transaminase | 1634 | gene | 1750544 | 1751635 | - | 1960,738 | 0,657616219 | 0,418875 | 1,569958 | 0,116425 | 0,211739 | 2721,140776 | 0,439423028 | 0,220935 | 1,988921 | 0,04671 | 0,074984 | 1200,335 | 1,187914692 | 0,231956 | 5,121283 | 3,03E-07 | 4,87E-06 | 1620,862 | -1,71207 | 0,280886 | -6,09527 | 1,09E-09 | 1,52E-08 | 2353,327198 | -0,919072486 | 0,150887 | -6,09113 | 1,12E-09 | 6,19E-09 |
| 1562 | RM25_RS07620 | RM25_1545 | Ferredoxin fdxC | 1635 | gene | 1751632 | 1751952 | - | 6126,116 | 1,044849879 | 0,424973 | 2,458625 | 0,013947 | 0,057999 | 7325,802655 | 0,527075266 | 0,241028 | 2,18678 | 0,028759 | 0,049226 | 4926,429 | 1,976013478 | 0,46322 | 4,265825 | 1,99E-05 | 0,000186 | 4274,638 | -1,63651 | 0,434558 | -3,76593 | 0,000166 | 0,000762 | 8058,673164 | -0,158576064 | 0,285565 | -0,55531 | 0,578685 | 0,639457 |
| 1563 | RM25_RS07625 | RM25_1546 | Glyoxylate reductase | 1636 | gene | 1752188 | 1753156 | + | 1645,727 | -0,817194613 | 0,362198 | -2,25621 | 0,024058 | 0,079347 | 1224,025584 | 0,211635885 | 0,160994 | 1,314555 | 0,18866 | 0,247039 | 2067,429 | -1,516863625 | 0,359376 | -4,22083 | 2,43E-05 | 0,000222 | 2206,92 | 1,384371 | 0,3725 | 3,716427 | 0,000202 | 0,000907 | 1167,974901 | -0,304129168 | 0,111378 | -2,73061 | 0,006322 | 0,01146 |
| 1564 | RM25_RS07630 | RM25_1547 | Fumarate hydratase class II | 1637 | gene | 1753320 | 1754744 | + | 4773,528 | -0,028097595 | 0,32801 | -0,08566 | 0,931736 | 0,953583 | 5858,34749 | 0,366111769 | 0,174388 | 2,099414 | 0,03578 | 0,059537 | 3688,708 | -0,661661185 | 0,364334 | -1,81608 | 0,069357 | 0,129743 | 5108,993 | -0,22973 | 0,311935 | -0,73646 | 0,461451 | 0,533942 | 4631,543651 | -1,223038721 | 0,244129 | -5,00981 | 5,45E-07 | 2E-06 |
| 1565 | RM25_RS07635 | RM25_1548 | Hypothetical protein | 1638 | gene | 1754908 | 1755354 | + | 265,8941 | 0,546811574 | 0,192243 | 2,844383 | 0,00445 | 0,027313 | 223,7711284 | 0,27337996 | 0,193 | 1,416475 | 0,156637 | 0,21124 | 308,0171 | 0,750484911 | 0,207393 | 3,618657 | 0,000296 | 0,001832 | 228,5867 | 0,120298 | 0,206435 | 0,582741 | 0,560068 | 0,625038 | 308,4360747 | 0,646100146 | 0,177217 | 3,645818 | 0,000267 | 0,000629 |
| 1566 | RM25_RS07640 | RM25_1549 | Aspartate ammonia-lyase | 1639 | gene | 1755482 | 1756951 | + | 1455,347 | -0,007774702 | 0,196501 | -0,03957 | 0,968439 | 0,978821 | 1207,743008 | -0,087633345 | 0,147403 | -0,59451 | 0,552168 | 0,617391 | 1702,952 | 0,047531058 | 0,214902 | 0,221175 | 0,824956 | 0,871938 | 1536,321 | 0,36699 | 0,180566 | 2,032416 | 0,04211 | 0,073903 | 1419,432845 | 0,550381868 | 0,155 | 3,550859 | 0,000384 | 0,000879 |
| 1567 | RM25_RS07645 | RM25_1550 | Aspartate ammonia-lyase | 1640 | gene | 1757281 | 1758753 | + | 13431,91 | -1,807939039 | 0,558788 | -3,23546 | 0,001214 | 0,011314 | 9677,459871 | -0,065839287 | 0,204821 | -0,32145 | 0,747871 | 0,794739 | 17186,35 | -3,684836527 | 0,454437 | -8,10857 | 5,12E-16 | 4,64E-14 | 21953,41 | 1,645934 | 0,425985 | 3,863832 | 0,000112 | 0,000535 | 5851,08696 | -1,948760552 | 0,251616 | -7,74499 | 9,56E-15 | 1,12E-13 |
| 1568 | RM25_RS07650 | RM25_1551 | Anaerobic C4-dicarboxylate transporter | 1641 | gene | 1758758 | 1760053 | + | 7941,429 | -1,411317891 | 0,46821 | -3,01429 | 0,002576 | 0,018623 | 7306,335508 | -0,232305484 | 0,193856 | -1,19834 | 0,230784 | 0,29419 | 8576,522 | -2,953448786 | 0,430312 | -6,8635 | 6,72E-12 | 3,23E-10 | 12164,17 | 0,901652 | 0,42163 | 2,138494 | 0,032477 | 0,060369 | 4256,048282 | -1,791774842 | 0,205417 | -8,7226 | 2,72E-18 | 5,21E-17 |
| 1569 | RM25_RS07655 | RM25_1552 | Aspartate racemase | 1642 | gene | 1760066 | 1760788 | + | 3852,976 | -0,952970114 | 0,420255 | -2,2676 | 0,023354 | 0,078136 | 4323,332991 | -0,216958877 | 0,213124 | -1,01804 | 0,308657 | 0,376751 | 3382,618 | -2,139296537 | 0,419198 | -5,10331 | 3,34E-07 | 5,28E-06 | 5375,646 | 0,199904 | 0,401166 | 0,498307 | 0,618267 | 0,676535 | 2572,682438 | -1,693647677 | 0,241539 | -7,0119 | 2,35E-12 | 1,92E-11 |
| 1570 | RM25_RS07660 | RM25_1553 | Integral membrane efflux protein | 1643 | gene | 1760892 | 1762229 | - | 411,6506 | -1,005327881 | 0,360586 | -2,78804 | 0,005303 | 0,030861 | 304,8521754 | -1,239415691 | 0,26591 | -4,66103 | 3,15E-06 | 1,46E-05 | 518,4491 | -0,876825773 | 0,424771 | -2,06423 | 0,038996 | 0,084401 | 577,4301 | 0,579988 | 0,30642 | 1,892788 | 0,058386 | 0,096573 | 269,8469693 | 1,016165662 | 0,411016 | 2,472328 | 0,013424 | 0,022811 |
| 1571 | RM25_RS07665 | RM25_1554 | Regulatory protein, MarR | 1644 | gene | 1762226 | 1762771 | - | 296,5834 | -0,785429597 | 0,379016 | -2,07229 | 0,038239 | 0,106159 | 206,254638 | -1,18823863 | 0,240254 | -4,94575 | 7,59E-07 | 3,89E-06 | 386,9121 | -0,58695694 | 0,38596 | -1,52077 | 0,128318 | 0,209159 | 394,0856 | 0,628354 | 0,274 | 2,293261 | 0,021833 | 0,04378 | 214,2852452 | 1,298747644 | 0,373755 | 3,474863 | 0,000511 | 0,001147 |
| 1572 | RM25_RS07670 | RM25_1555 | Phosphinothricin acetyltransferase | 1645 | gene | 1762912 | 1763424 | + | 207,645 | -0,729205436 | 0,397506 | -1,83445 | 0,066587 | 0,148905 | 137,3022092 | -1,08585784 | 0,31332 | -3,46565 | 0,000529 | 0,00152 | 277,9878 | -0,566512471 | 0,340313 | -1,66468 | 0,095977 | 0,167197 | 272,0009 | 0,759236 | 0,315308 | 2,407921 | 0,016044 | 0,034629 | 153,5710888 | 1,347113493 | 0,360859 | 3,733073 | 0,000189 | 0,000457 |
| 1573 | RM25_RS07675 | RM25_1556 | Purine phosphoribosyltransferase | 1646 | gene | 1763484 | 1763990 | + | 1351,729 | 0,289883634 | 0,398787 | 0,726913 | 0,467279 | 0,569305 | 1856,598343 | 0,55825655 | 0,175546 | 3,180112 | 0,001472 | 0,003739 | 846,8604 | -0,284019442 | 0,356076 | -0,79764 | 0,425082 | 0,529555 | 1294,044 | -0,74335 | 0,338438 | -2,19642 | 0,028062 | 0,05347 | 1457,828992 | -1,549417433 | 0,203356 | -7,61923 | 2,55E-14 | 2,81E-13 |
| 1574 | RM25_RS07680 | RM25_1557 | Hippurate hydrolase | 1647 | gene | 1764066 | 1765235 | - | 661,2885 | 0,101806527 | 0,139902 | 0,727696 | 0,4668 | 0,569191 | 631,4823997 | 0,352269572 | 0,143412 | 2,456348 | 0,014036 | 0,026572 | 691,0946 | -0,123737201 | 0,218066 | -0,56743 | 0,570422 | 0,663235 | 672,6019 | 0,314432 | 0,150448 | 2,089974 | 0,03662 | 0,066395 | 672,6483507 | -0,10093583 | 0,215613 | -0,46813 | 0,639689 | 0,695135 |
| 1575 | RM25_RS07685 | RM25_1558 | Hypothetical protein | 1648 | gene | 1765396 | 1766547 | + | 593,9355 | -0,943487552 | 0,471482 | -2,00111 | 0,04538 | 0,117018 | 341,4014883 | -1,334969994 | 0,286062 | -4,66664 | 3,06E-06 | 1,43E-05 | 846,4695 | -0,796847937 | 0,379986 | -2,09705 | 0,035989 | 0,07965 | 818,7073 | 1,065425 | 0,332816 | 3,201242 | 0,001368 | 0,004614 | 400,2515735 | 1,676506282 | 0,356475 | 4,703012 | 2,56E-06 | 8,31E-06 |
| 1576 | NA | RM25_1559 | Hypothetical protein | 1649 | gene | 1766433 | 1766675 | - | 69,47364 | -1,159321761 | 0,734002 | -1,57945 | 0,114232 | 0,208718 | 27,18605165 | -2,039686605 | 0,741949 | -2,74909 | 0,005976 | 0,012805 | 111,7612 | -0,97984604 | 0,419617 | -2,3351 | 0,019538 | 0,050409 | 100,2594 | 1,672508 | 0,503961 | 3,318726 | 0,000904 | 0,003234 | 42,27099484 | 2,815825359 | 0,674507 | 4,174644 | 2,98E-05 | 8,22E-05 |
| 1577 | RM25_RS11940 | RM25_1560 | Diacylglycerol kinase catalytic region | 1650 | gene | 1766638 | 1767693 | - | 314,71 | -0,246985455 | 0,277489 | -0,89007 | 0,373427 | 0,484674 | 242,6288313 | -0,265360721 | 0,200559 | -1,3231 | 0,185801 | 0,24411 | 386,7912 | -0,238740214 | 0,33215 | -0,71877 | 0,472281 | 0,574686 | 358,9893 | 0,598755 | 0,189645 | 3,157245 | 0,001593 | 0,005207 | 283,0833188 | 0,687448586 | 0,33259 | 2,066955 | 0,038738 | 0,059022 |
| 1578 | RM25_RS07695 | RM25_1561 | Cysteine synthase | 1651 | gene | 1768138 | 1769106 | + | 2634,225 | 0,917657349 | 0,453365 | 2,024103 | 0,04296 | 0,113388 | 3892,192025 | 1,054266724 | 0,265972 | 3,963819 | 7,38E-05 | 0,000259 | 1376,257 | 0,549926514 | 0,210902 | 2,6075 | 0,009121 | 0,02828 | 1926,089 | -1,22742 | 0,307852 | -3,98706 | 6,69E-05 | 0,000342 | 3379,911308 | -1,695398277 | 0,123342 | -13,7455 | 5,42E-43 | 7,98E-41 |
| 1579 | RM25_RS07700 | RM25_1562 | SPFH/Band 7/PHB domain protein | 1652 | gene | 1769326 | 1770294 | + | 559,2686 | -0,74810524 | 0,239908 | -3,11831 | 0,001819 | 0,014243 | 445,9488074 | -0,277535895 | 0,153448 | -1,80867 | 0,070502 | 0,104776 | 672,5885 | -1,079537499 | 0,251563 | -4,29133 | 1,78E-05 | 0,000169 | 737,159 | 0,847144 | 0,235215 | 3,601566 | 0,000316 | 0,001348 | 409,4810332 | 0,09511005 | 0,187264 | 0,507892 | 0,611529 | 0,671031 |
| 1580 | RM25_RS07705 | RM25_1563 | Hypothetical protein | 1653 | gene | 1770307 | 1770531 | + | 68,72811 | 0,155645713 | 0,454646 | 0,342345 | 0,732091 | 0,805067 | 91,25500606 | 0,30626903 | 0,333803 | 0,917515 | 0,358873 | 0,427636 | 46,20121 | -0,114289425 | 0,575559 | -0,19857 | 0,842598 | 0,885818 | 69,21477 | -0,81605 | 0,570492 | -1,43044 | 0,152592 | 0,210241 | 71,07291916 | -1,185543559 | 0,365244 | -3,24589 | 0,001171 | 0,002431 |
| 1581 | RM25_RS07710 | RM25_1564 | Cyanate MFS transporter | 1654 | gene | 1770634 | 1771911 | - | 765,351 | 0,464048262 | 0,108736 | 4,267673 | 1,98E-05 | 0,000483 | 783,5327663 | 0,411004879 | 0,149194 | 2,754834 | 0,005872 | 0,012628 | 747,1693 | 0,520565587 | 0,21207 | 2,45469 | 0,014101 | 0,039566 | 679,6856 | -0,19689 | 0,203507 | -0,9675 | 0,333292 | 0,408142 | 869,9094242 | -0,029085188 | 0,135302 | -0,21496 | 0,829795 | 0,861977 |
| 1582 | RM25_RS07715 | RM25_1565 | Putative OSMOPROTECTANT (GLYCINE BETAINE/CARNITINE | 1655 | gene | 1772059 | 1772958 | - | 496,4375 | 0,347861623 | 0,120984 | 2,87527 | 0,004037 | 0,025716 | 532,9295118 | 0,439716259 | 0,149743 | 2,936474 | 0,00332 | 0,007701 | 459,9456 | 0,245504635 | 0,198119 | 1,239179 | 0,215279 | 0,312446 | 460,9709 | -0,16474 | 0,159025 | -1,03594 | 0,300229 | 0,374809 | 545,7549619 | -0,304674941 | 0,169188 | -1,8008 | 0,071734 | 0,102285 |
| 1583 | RM25_RS07720 | RM25_1566 | Peptidyl-tRNA hydrolase PTH2 | 1656 | gene | 1772996 | 1773709 | - | 451,3935 | 0,092281539 | 0,14107 | 0,654154 | 0,513012 | 0,610076 | 435,1795947 | 0,341953389 | 0,144188 | 2,371581 | 0,017712 | 0,032362 | 467,6073 | -0,13399987 | 0,232357 | -0,5767 | 0,564144 | 0,65924 | 459,5812 | 0,289815 | 0,164221 | 1,764782 | 0,0776 | 0,120332 | 457,4836406 | -0,128689284 | 0,202944 | -0,63508 | 0,525377 | 0,586045 |
| 1584 | RM25_RS07725 | RM25_1567 | 3-demethylubiquinone-9 3-methyltransferase domain prote | 1657 | gene | 1773810 | 1774703 | + | 721,3337 | -0,067703534 | 0,1502 | -0,45075 | 0,652166 | 0,736892 | 647,7089347 | -0,240515977 | 0,157606 | -1,52605 | 0,126996 | 0,176388 | 794,9584 | 0,070703542 | 0,194021 | 0,364412 | 0,71555 | 0,788109 | 779,484 | 0,079826 | 0,155085 | 0,514725 | 0,606745 | 0,666713 | 688,9999722 | 0,443059671 | 0,172 | 2,575921 | 0,009997 | 0,017455 |
| 1585 | RM25_RS07730 | RM25_1568 | Major facilitator family transmembrane efflux protein | 1658 | gene | 1774905 | 1776674 | + | 862,2338 | -0,115878835 | 0,375517 | -0,30858 | 0,757638 | 0,824795 | 553,375912 | -0,144468309 | 0,263927 | -0,54738 | 0,584118 | 0,64566 | 1171,092 | -0,103444786 | 0,285047 | -0,3629 | 0,716676 | 0,78898 | 941,0771 | 0,991549 | 0,310307 | 3,195385 | 0,001396 | 0,004677 | 813,4004146 | 1,101749053 | 0,262911 | 4,190581 | 2,78E-05 | 7,69E-05 |
| 1586 | RM25_RS07735 | RM25_1569 | Glycerate kinase | 1659 | gene | 1776752 | 1777870 | - | 445,3733 | -0,759772942 | 0,443798 | -1,71198 | 0,086901 | 0,176421 | 242,417229 | -0,594048767 | 0,19323 | -3,07431 | 0,00211 | 0,005159 | 648,3294 | -0,82288734 | 0,341493 | -2,40968 | 0,015967 | 0,043156 | 585,7414 | 1,450244 | 0,181191 | 8,003968 | 1,2E-15 | 5,26E-14 | 325,4544698 | 1,280699832 | 0,339374 | 3,773712 | 0,000161 | 0,000397 |
| 1587 | RM25_RS07740 | RM25_1570 | Carbon starvation protein A | 1660 | gene | 1778241 | 1780637 | + | 1976,695 | -0,764891631 | 0,336676 | -2,2719 | 0,023093 | 0,07779 | 1281,528106 | -0,335336722 | 0,186883 | -1,79437 | 0,072755 | 0,107736 | 2671,863 | -0,982985863 | 0,185454 | -5,30042 | 1,16E-07 | 2,03E-06 | 2611,134 | 1,252561 | 0,231401 | 5,412955 | 6,2E-08 | 5,99E-07 | 1433,241057 | 0,652350842 | 0,141809 | 4,600208 | 4,22E-06 | 1,32E-05 |
| 1588 | RM25_RS07745 | RM25_1571 | Hypothetical protein | 1661 | gene | 1780627 | 1780842 | + | 86,7484 | -0,535854343 | 0,273756 | -1,95742 | 0,050299 | 0,125197 | 66,72380449 | -0,112419022 | 0,297309 | -0,37812 | 0,70534 | 0,756147 | 106,773 | -0,806110729 | 0,267992 | -3,00796 | 0,00263 | 0,010819 | 107,9277 | 0,910847 | 0,267126 | 3,409798 | 0,00065 | 0,002462 | 69,38513999 | 0,268773028 | 0,30684 | 0,87594 | 0,381063 | 0,446404 |
| 1589 | RM25_RS07750 | RM25_1572 | Transcriptional regulator, LysR family | 1662 | gene | 1780846 | 1781799 | - | 715,8467 | -0,159453257 | 0,094381 | -1,68947 | 0,09113 | 0,182183 | 693,6644168 | -0,040481823 | 0,153002 | -0,26458 | 0,791331 | 0,834153 | 738,029 | -0,269635689 | 0,15914 | -1,69433 | 0,090202 | 0,158965 | 797,158 | 0,13674 | 0,15812 | 0,864785 | 0,387157 | 0,463683 | 663,0987867 | -0,038074583 | 0,153756 | -0,24763 | 0,804421 | 0,841553 |
| 1590 | RM25_RS07755 | RM25_1573 | UPF0324 membrane protein | 1663 | gene | 1781836 | 1782918 | + | 762,6669 | -0,121769959 | 0,135359 | -0,89961 | 0,368328 | 0,47964 | 741,6778019 | -0,007256315 | 0,191244 | -0,03794 | 0,969733 | 0,975944 | 783,656 | -0,229144391 | 0,213756 | -1,07199 | 0,283725 | 0,388304 | 836,6025 | 0,129637 | 0,176179 | 0,735824 | 0,461838 | 0,534127 | 716,8205603 | -0,039412168 | 0,209128 | -0,18846 | 0,850516 | 0,878854 |
| 1591 | RM25_RS07760 | RM25_1574 | Putative adenylyltransferase/sulfurtransferase MoeZ | 1664 | gene | 1782986 | 1784197 | - | 585,9877 | -0,375309714 | 0,13029 | -2,88058 | 0,003969 | 0,025563 | 593,6501982 | -0,265688725 | 0,14098 | -1,88458 | 0,059486 | 0,090809 | 578,3252 | -0,486737222 | 0,254665 | -1,91129 | 0,055968 | 0,109681 | 698,0593 | -0,00041 | 0,144134 | -0,00282 | 0,997746 | 0,99817 | 501,365909 | -0,163282894 | 0,240891 | -0,67783 | 0,49788 | 0,559231 |
| 1592 | RM25_RS07765 | RM25_1575 | Thiazole synthase | 1665 | gene | 1784206 | 1785036 | - | 463,6552 | -0,399059774 | 0,170941 | -2,33449 | 0,01957 | 0,071001 | 411,5942485 | -0,473985607 | 0,17116 | -2,76926 | 0,005618 | 0,01216 | 515,7161 | -0,342944604 | 0,247839 | -1,38374 | 0,166439 | 0,25616 | 555,4206 | 0,20588 | 0,199633 | 1,031294 | 0,302403 | 0,377053 | 392,48493 | 0,395461469 | 0,219887 | 1,798477 | 0,072101 | 0,102747 |
| 1593 | RM25_RS07770 | RM25_1576 | Thiamine biosynthesis protein | 1666 | gene | 1785039 | 1785272 | - | 63,4603 | -0,532244733 | 0,37129 | -1,4335 | 0,151714 | 0,252291 | 45,11497062 | -0,863140209 | 0,454545 | -1,89891 | 0,057576 | 0,088409 | 81,80563 | -0,368478243 | 0,301961 | -1,22028 | 0,222357 | 0,319181 | 78,76762 | 0,59611 | 0,344965 | 1,72803 | 0,083983 | 0,128454 | 50,90195363 | 1,153574036 | 0,42712 | 2,700817 | 0,006917 | 0,012455 |
| 1594 | RM25_RS07775 | RM25_1577 | Glycine oxidase ThiO | 1667 | gene | 1785269 | 1786522 | - | 481,6304 | -0,462945737 | 0,311649 | -1,48547 | 0,137418 | 0,237286 | 351,5699787 | -0,562008882 | 0,221013 | -2,54288 | 0,010994 | 0,021541 | 611,6908 | -0,408990653 | 0,321708 | -1,27131 | 0,203619 | 0,300896 | 586,2618 | 0,671579 | 0,224134 | 2,996328 | 0,002733 | 0,008236 | 398,2696666 | 0,887955996 | 0,318218 | 2,790402 | 0,005264 | 0,009709 |
| 1595 | RM25_RS07780 | RM25_1578 | Thiamine-phosphate diphosphorylase | 1668 | gene | 1786698 | 1787345 | + | 508,9653 | -0,14916243 | 0,198671 | -0,7508 | 0,452772 | 0,558602 | 424,1863564 | -0,165268177 | 0,199156 | -0,82984 | 0,406628 | 0,475173 | 593,7443 | -0,139905905 | 0,198534 | -0,70469 | 0,481001 | 0,581676 | 562,8239 | 0,411575 | 0,212983 | 1,932435 | 0,053306 | 0,089552 | 473,5039525 | 0,493009984 | 0,180422 | 2,732536 | 0,006285 | 0,011404 |
| 1596 | RM25_RS12765 | RM25_1579 | Hypothetical protein | 1669 | gene | 1787858 | 1788031 | + | 82,08179 | -1,312815606 | 0,563418 | -2,33009 | 0,019801 | 0,07121 | 48,98644959 | -2,537867495 | 0,459872 | -5,51864 | 3,42E-08 | 2,28E-07 | 115,1771 | -0,924460925 | 0,400754 | -2,30681 | 0,021066 | 0,053217 | 122,7961 | 0,781079 | 0,331019 | 2,359619 | 0,018294 | 0,038124 | 46,28018041 | 2,462492652 | 0,552671 | 4,455618 | 8,37E-06 | 2,47E-05 |
| 1597 | RM25_RS07785 | RM25_1580 | UPF0324 membrane protein | 1670 | gene | 1788028 | 1789191 | + | 319,6015 | -1,363594315 | 0,508981 | -2,67907 | 0,007383 | 0,038683 | 199,5973828 | -2,09711605 | 0,384764 | -5,4504 | 5,03E-08 | 3,23E-07 | 439,6056 | -1,083154509 | 0,426135 | -2,54181 | 0,011028 | 0,032903 | 482,7101 | 0,814128 | 0,390467 | 2,085014 | 0,037068 | 0,066899 | 176,3984628 | 1,904139678 | 0,457494 | 4,162107 | 3,15E-05 | 8,63E-05 |
| 1598 | RM25_RS07790 | RM25_1581 | Putative arsenite efflux pump | 1671 | gene | 1789257 | 1789982 | - | 191,5108 | -1,191847771 | 0,514163 | -2,31803 | 0,020447 | 0,072518 | 123,0724962 | -2,346193995 | 0,36135 | -6,49285 | 8,42E-11 | 8,27E-10 | 259,9492 | -0,767869705 | 0,345904 | -2,21989 | 0,026426 | 0,063235 | 279,996 | 0,602136 | 0,319194 | 1,886423 | 0,059238 | 0,097639 | 114,5723023 | 2,247302222 | 0,415345 | 5,410688 | 6,28E-08 | 2,69E-07 |
| 1599 | RM25_RS07795 | RM25_1582 | Glycerol-3-phosphate dehydrogenase glpD2 | 1672 | gene | 1790122 | 1791807 | + | 648,048 | -0,399160948 | 0,302216 | -1,32078 | 0,186574 | 0,291031 | 495,1611945 | -0,844096132 | 0,18568 | -4,54597 | 5,47E-06 | 2,42E-05 | 800,9348 | -0,136130884 | 0,297326 | -0,45785 | 0,647059 | 0,72798 | 776,0705 | 0,332815 | 0,20923 | 1,590664 | 0,111685 | 0,162696 | 548,7553991 | 1,104721939 | 0,275053 | 4,016391 | 5,91E-05 | 0,000155 |
| 1600 | RM25_RS07800 | RM25_1583 | Alkylglycerone-phosphate synthase | 1673 | gene | 1791804 | 1793336 | + | 695,7308 | -0,536898138 | 0,209823 | -2,55882 | 0,010503 | 0,048578 | 666,4944751 | -0,91945728 | 0,196108 | -4,68852 | 2,75E-06 | 1,29E-05 | 724,9672 | -0,203385852 | 0,287858 | -0,70655 | 0,479847 | 0,580894 | 871,18 | -0,23151 | 0,170648 | -1,35665 | 0,174891 | 0,23576 | 556,7881878 | 0,543498038 | 0,289226 | 1,879144 | 0,060225 | 0,0873 |
| 1601 | RM25_RS07805 | RM25_1584 | Transcriptional regulator GlpP family | 1674 | gene | 1793425 | 1794084 | + | 290,4456 | -0,606434702 | 0,361608 | -1,67705 | 0,093533 | 0,184637 | 195,2235274 | -0,889834915 | 0,21556 | -4,12801 | 3,66E-05 | 0,000139 | 385,6678 | -0,470185234 | 0,321329 | -1,46325 | 0,143399 | 0,227757 | 368,103 | 0,754468 | 0,224947 | 3,353977 | 0,000797 | 0,002897 | 226,2789528 | 1,240686234 | 0,322364 | 3,848707 | 0,000119 | 0,000297 |
| 1602 | RM25_RS07810 | RM25_1585 | Hypothetical protein | 1675 | gene | 1794063 | 1794329 | - | 202,3009 | -0,33032675 | 0,356755 | -0,92592 | 0,354487 | 0,466388 | 128,5670778 | -0,278901797 | 0,240402 | -1,16015 | 0,245989 | 0,310717 | 276,0347 | -0,356507868 | 0,212965 | -1,67402 | 0,094126 | 0,164256 | 236,3153 | 1,07922 | 0,172407 | 6,259733 | 3,86E-10 | 6,1E-09 | 175,3367146 | 1,052149416 | 0,255322 | 4,120869 | 3,77E-05 | 0,000102 |
| 1603 | RM25_RS12315 | RM25_1586 | Hypothetical protein | 1676 | gene | 1794408 | 1795280 | - | 1472,864 | -0,705267004 | 0,471274 | -1,49651 | 0,13452 | 0,233479 | 766,2499187 | -0,142994531 | 0,149208 | -0,95836 | 0,337881 | 0,406112 | 2179,477 | -0,914081205 | 0,394589 | -2,31654 | 0,020529 | 0,052367 | 1915,707 | 1,772172 | 0,309409 | 5,727609 | 1,02E-08 | 1,13E-07 | 1090,25222 | 1,032139687 | 0,277777 | 3,715712 | 0,000203 | 0,000488 |
| 1604 | RM25_RS07815 | RM25_1587 | FrnE protein | 1677 | gene | 1795853 | 1796548 | - | 608,0281 | 1,380420213 | 0,268248 | 5,146058 | 2,66E-07 | 1,37E-05 | 443,6736951 | 0,963060583 | 0,19711 | 4,885893 | 1,03E-06 | 5,24E-06 | 772,3826 | 1,642674769 | 0,232049 | 7,079004 | 1,45E-12 | 7,78E-11 | 355,7173 | 0,256324 | 0,214947 | 1,192502 | 0,233064 | 0,301004 | 857,9046223 | 0,979610348 | 0,186407 | 5,255222 | 1,48E-07 | 5,87E-07 |
| 1605 | RM25_RS07820 | RM25_1588 | Inner membrane transport protein ynfM | 1678 | gene | 1796955 | 1798106 | - | 458,0521 | -1,313113277 | 0,656551 | -2,00002 | 0,045498 | 0,117074 | 236,7344544 | -2,900924294 | 0,331711 | -8,74533 | 2,22E-18 | 5,46E-17 | 679,3697 | -0,921604857 | 0,466079 | -1,97736 | 0,048001 | 0,097702 | 684,5345 | 1,018196 | 0,340004 | 2,994655 | 0,002748 | 0,008271 | 259,7417601 | 3,074582067 | 0,477617 | 6,437335 | 1,22E-10 | 7,81E-10 |
| 1606 | RM25_RS07825 | RM25_1589 | DNA-binding transcriptional regulator OxyR | 1679 | gene | 1798264 | 1799229 | + | 340,0784 | -1,079870121 | 0,653711 | -1,65191 | 0,098554 | 0,191186 | 166,5795972 | -2,546412792 | 0,320901 | -7,93521 | 2,1E-15 | 3,64E-14 | 513,5772 | -0,720840801 | 0,489827 | -1,47162 | 0,141123 | 0,225052 | 483,0331 | 1,097147 | 0,334664 | 3,278357 | 0,001044 | 0,003651 | 215,9010131 | 3,000364095 | 0,490694 | 6,114533 | 9,68E-10 | 5,42E-09 |
| 1607 | RM25_RS07830 | RM25_1590 | Hypothetical protein | 1680 | gene | 1799193 | 1799963 | - | 314,3183 | -1,40603854 | 0,576534 | -2,43878 | 0,014737 | 0,060409 | 188,744808 | -2,830545703 | 0,345334 | -8,19654 | 2,47E-16 | 4,78E-15 | 439,8918 | -0,968381534 | 0,354688 | -2,73023 | 0,006329 | 0,021682 | 478,9561 | 0,743963 | 0,359401 | 2,070007 | 0,038452 | 0,068816 | 169,4233606 | 2,67717866 | 0,371935 | 7,197982 | 6,11E-13 | 5,46E-12 |
| 1608 | RM25_RS07835 | RM25_1591 | Hypothetical protein | 1681 | gene | 1800057 | 1801166 | - | 382,11 | -1,502544227 | 0,742238 | -2,02434 | 0,042935 | 0,113388 | 201,8711716 | -3,887677783 | 0,415077 | -9,36661 | 7,49E-21 | 2,63E-19 | 562,3488 | -1,011127572 | 0,438804 | -2,30428 | 0,021207 | 0,053517 | 592,2475 | 0,915427 | 0,425541 | 2,151208 | 0,03146 | 0,058803 | 196,7883771 | 3,872700779 | 0,465564 | 8,318296 | 8,92E-17 | 1,35E-15 |
| 1609 | RM25_RS07840 | RM25_1592 | Hypothetical protein | 1682 | gene | 1801166 | 1801390 | - | 57,81223 | -1,168562823 | 0,772397 | -1,5129 | 0,130304 | 0,228857 | 28,16757352 | -3,594829581 | 0,702816 | -5,1149 | 3,14E-07 | 1,72E-06 | 87,45688 | -0,698281069 | 0,43218 | -1,61572 | 0,106155 | 0,180369 | 83,82611 | 0,981141 | 0,397973 | 2,465349 | 0,013688 | 0,030408 | 35,06871038 | 3,95181362 | 0,747418 | 5,28729 | 1,24E-07 | 4,98E-07 |
| 1610 | NA | RM25_1593 | Hypothetical protein | 1683 | gene | 1801592 | 1801732 | - | 87,54764 | -1,689411898 | 0,721531 | -2,34143 | 0,01921 | 0,070199 | 47,01117742 | -3,7611618 | 0,635018 | -5,92292 | 3,16E-09 | 2,53E-08 | 128,0841 | -1,236428902 | 0,360013 | -3,4344 | 0,000594 | 0,003203 | 139,8609 | 0,966747 | 0,443642 | 2,179115 | 0,029323 | 0,055425 | 40,68930871 | 3,5587122 | 0,61763 | 5,761885 | 8,32E-09 | 4,01E-08 |
| 1611 | NA | RM25_1594 | Hypothetical protein | 1684 | gene | 1802178 | 1802735 | - | 222,9274 | -1,175165628 | 0,33334 | -3,52542 | 0,000423 | 0,005329 | 165,4895966 | -1,616837114 | 0,285268 | -5,66777 | 1,45E-08 | 1,03E-07 | 280,3651 | -0,946332128 | 0,245877 | -3,84881 | 0,000119 | 0,000853 | 325,116 | 0,496606 | 0,294199 | 1,687992 | 0,091413 | 0,137762 | 133,9064523 | 1,226525205 | 0,247738 | 4,950893 | 7,39E-07 | 2,65E-06 |
| 1612 | NA | RM25_1595 | Hypothetical protein | 1685 | gene | 1802748 | 1802876 | - | 61,08245 | -0,73907736 | 0,212078 | -3,48494 | 0,000492 | 0,006012 | 55,04224407 | -0,895211156 | 0,355054 | -2,52134 | 0,011691 | 0,022587 | 67,12265 | -0,627298475 | 0,284624 | -2,20396 | 0,027527 | 0,065142 | 80,41398 | 0,121711 | 0,255939 | 0,475548 | 0,634396 | 0,691615 | 44,76001224 | 0,444644044 | 0,390493 | 1,138673 | 0,25484 | 0,312679 |
| 1613 | RM25_RS07845 | RM25_1596 | gbkey=CDS | 1686 | gene | 1803059 | 1805005 | - | 3656,5 | -0,720410479 | 0,386842 | -1,86229 | 0,062563 | 0,14275 | 2340,293383 | -0,189984102 | 0,168822 | -1,12535 | 0,260439 | 0,325619 | 4972,706 | -0,98516949 | 0,402093 | -2,45011 | 0,014281 | 0,039883 | 4783,668 | 1,357937 | 0,348749 | 3,89374 | 9,87E-05 | 0,00048 | 2694,92221 | 0,592497684 | 0,252774 | 2,343984 | 0,019079 | 0,031535 |
| 1614 | NA | RM25_1597 | Transposase IS3/IS911 | 1687 | gene | 1805398 | 1805742 | + | 165,7 | -0,245596311 | 0,247628 | -0,9918 | 0,321297 | 0,430528 | 128,6575268 | -0,166449027 | 0,231268 | -0,71972 | 0,471694 | 0,538954 | 202,7425 | -0,297466468 | 0,24922 | -1,19359 | 0,232639 | 0,330718 | 189,5595 | 0,654427 | 0,252462 | 2,592181 | 0,009537 | 0,022844 | 148,3146232 | 0,572246653 | 0,231011 | 2,477146 | 0,013244 | 0,022522 |
| 1615 | RM25_RS07855 | RM25_1598 | IS3 family transposase OrfB | 1688 | gene | 1805739 | 1806365 | + | 72,01756 | -0,20894802 | 0,2931 | -0,71289 | 0,475914 | 0,576724 | 55,05072089 | -0,161984062 | 0,395465 | -0,4096 | 0,682096 | 0,735453 | 88,98439 | -0,241357041 | 0,243392 | -0,99164 | 0,321373 | 0,426507 | 81,24588 | 0,668039 | 0,238828 | 2,797158 | 0,005155 | 0,013517 | 65,50167379 | 0,645791955 | 0,41339 | 1,562185 | 0,118244 | 0,160358 |
| 1616 | RM25_RS07860 | RM25_1599 | Transposase insf for insertion sequence element IS30B/C/D | 1689 | gene | 1806411 | 1807850 | + | 2,689189 | -1,607440007 | 1,106741 | -1,55047 | 0,121028 | 0,217744 | 1,227014258 | -2,277840986 | 2,241176 | -1,01636 | 0,309458 | 0,376811 | 4,151363 | -1,479145334 | 0,922638 | -1,60317 | 0,108897 | 0,183861 | 4,243051 | 1,608437 | 0,842649 | 1,908787 | 0,05629 | 0,093697 | 1,26945403 | 2,384893471 | 2,889941 | 0,82524 | 0,409236 | 0,475391 |

| | | | | | | | | | | | | | | | | | | | | | | | | | | | | | | | | | | | | | | |
|---|---|---|---|---|---|---|---|---|---|---|---|---|---|---|---|---|---|---|---|---|---|---|---|---|---|---|---|---|---|---|---|---|---|---|---|---|---|---|
| 1617 | RM25_RS07865 | RM25_1600 | Hypothetical protein | 1690 | gene | 1807876 | 1808148 | + | 25,10379 | -0,66264282 | 0,732827 | -0,90423 | 0,365875 | 0,477237 | 14,45287477 | -3,252444846 | 0,834701 | -3,89654 | 9,76E-05 | 0,000333 | 35,7547 | 0,004199153 | 0,444359 | 0,00945 | 0,99246 | 0,995678 | 32,38858 | 0,37739 | 0,473444 | 0,797117 | 0,425383 | 0,500563 | 19,01256348 | 3,705707869 | 0,805416 | 4,600984 | 4,21E-06 | 1,32E-05 |
| 1618 | RM25_RS07870 | RM25_1601 | IS1096, TnpA family protein | 1691 | gene | 1808198 | 1808737 | + | 282,519 | -1,575743365 | 0,668411 | -2,35745 | 0,018401 | 0,068517 | 168,0428524 | -3,613018211 | 0,434098 | -8,32305 | 8,57E-17 | 1,77E-15 | 396,9951 | -1,052883946 | 0,353163 | -2,9813 | 0,00287 | 0,011604 | 443,6063 | 0,714856 | 0,451093 | 1,584718 | 0,11303 | 0,164452 | 139,4499787 | 3,344108679 | 0,36756 | 9,098125 | 9,19E-20 | 2,14E-18 |
| 1619 | RM25_RS07875 | RM25_1602 | Type III restriction-modification system StyLTI enzyme res | 1692 | gene | 1808835 | 1811810 | - | 933,5629 | -1,353789141 | 0,465177 | -2,91026 | 0,003611 | 0,02353 | 588,0321776 | -2,038137937 | 0,304864 | -6,6854 | 2,3E-11 | 2,54E-10 | 1279,094 | -1,085449329 | 0,319752 | -3,39466 | 0,000687 | 0,003607 | 1408,094 | 0,808355 | 0,37704 | 2,143949 | 0,032037 | 0,059598 | 516,214392 | 1,831321943 | 0,271416 | 6,747277 | 1,51E-11 | 1,09E-10 |
| 1620 | RM25_RS07880 | RM25_1603 | Type 3 restriction-modification system methylation subunit | 1693 | gene | 1811814 | 1812746 | - | 189,8541 | -1,29853629 | 0,405816 | -3,19982 | 0,001375 | 0,012231 | 133,7827821 | -1,88491931 | 0,342525 | -5,50301 | 3,73E-08 | 2,47E-07 | 245,9254 | -1,024129302 | 0,314879 | -3,25246 | 0,001144 | 0,005426 | 283,5128 | 0,587293 | 0,27289 | 2,152123 | 0,031388 | 0,058727 | 107,7870399 | 1,507968312 | 0,410312 | 3,675172 | 0,000238 | 0,000564 |
| 1621 | RM25_RS07885 | RM25_1604 | Transposase IS204/IS1001/IS1096/IS1165 family protein | 1694 | gene | 1812756 | 1813721 | - | 0 | NA | NA | NA | NA | NA | 0 | NA | NA | NA | NA | NA | 0 | NA | NA | NA | NA | NA | 0 | NA | NA | NA | NA | NA | 0 | NA | NA | NA | NA | NA |
| 1622 | RM25_RS12980 | RM25_1605 | Type III restriction-modification system methyltransferase | 1695 | gene | 1814122 | 1815213 | - | 1111,837 | 0,211967949 | 0,422473 | 0,501732 | 0,615856 | 0,703964 | 1601,758884 | 0,209925997 | 0,157848 | 1,329926 | 0,183543 | 0,241817 | 621,915 | 0,218654398 | 0,261364 | 0,836591 | 0,402823 | 0,508817 | 1095,88 | -1,43035 | 0,235574 | -6,07177 | 1,27E-09 | 1,74E-08 | 1170,322347 | -1,372743233 | 0,145166 | -9,4564 | 3,19E-21 | 8,35E-20 |
| 1623 | RM25_RS07895 | RM25_1606 | Hypothetical protein | 1696 | gene | 1815266 | 1816192 | - | 996,4549 | 0,283926938 | 0,442035 | 0,642318 | 0,520667 | 0,61762 | 1450,492995 | 0,237950881 | 0,192884 | 1,23365 | 0,217333 | 0,27931 | 542,4169 | 0,407732583 | 0,265047 | 1,538342 | 0,123965 | 0,203189 | 956,4936 | -1,57364 | 0,241173 | -6,52495 | 6,8E-11 | 1,29E-09 | 1072,920454 | -1,35582535 | 0,18784 | -7,21798 | 5,28E-13 | 4,75E-12 |
| 1624 | NA | RM25_1607 | Hypothetical protein | 1697 | gene | 1816247 | 1816702 | - | 374,7929 | 0,273284184 | 0,397256 | 0,68793 | 0,491497 | 0,590448 | 529,0060169 | 0,326970196 | 0,205593 | 1,590376 | 0,11175 | 0,156783 | 220,5798 | 0,149262021 | 0,221633 | 0,673464 | 0,500652 | 0,600222 | 361,2842 | -1,22685 | 0,236303 | -5,19183 | 2,08E-07 | 1,82E-06 | 402,232514 | -1,353800049 | 0,180234 | -7,51136 | 5,85E-14 | 6,1E-13 |
| 1625 | RM25_RS07905 | RM25_1608 | GTP-binding protein YchF | 1698 | gene | 1817122 | 1818213 | - | 1189,802 | 0,427729272 | 0,366897 | 1,165802 | 0,243694 | 0,352385 | 1649,718218 | 0,512020809 | 0,179152 | 2,858018 | 0,004263 | 0,009497 | 729,8861 | 0,241327759 | 0,16778 | 1,438356 | 0,150333 | 0,235324 | 1078,348 | -1,08062 | 0,187301 | -5,76943 | 7,95E-09 | 9,01E-08 | 1338,991756 | -1,304341526 | 0,150332 | -8,67642 | 4,08E-18 | 7,57E-17 |
| 1626 | RM25_RS07910 | RM25_1609 | M18 family aminopeptidase | 1699 | gene | 1818376 | 1819647 | - | 1127,193 | 0,31549161 | 0,092065 | 3,426833 | 0,000611 | 0,007161 | 1138,970134 | 0,489779603 | 0,126803 | 3,862524 | 0,000112 | 0,000377 | 1115,415 | 0,143249524 | 0,159706 | 0,896956 | 0,369742 | 0,475441 | 1057,872 | 0,102315 | 0,134556 | 0,760394 | 0,447019 | 0,521596 | 1226,422284 | -0,191581294 | 0,119596 | -1,60191 | 0,109176 | 0,149175 |
| 1627 | RM25_RS07915 | RM25_1610 | Putative flagellar protein FliS | 1700 | gene | 1819797 | 1821146 | + | 534,0065 | -0,383534616 | 0,257557 | -1,48913 | 0,136454 | 0,23614 | 409,8093081 | -0,263643594 | 0,195074 | -1,35151 | 0,176533 | 0,233626 | 658,2036 | -0,46030831 | 0,254658 | -1,80756 | 0,070676 | 0,13179 | 634,8805 | 0,706518 | 0,201086 | 3,513512 | 0,000442 | 0,001776 | 455,2762275 | 0,570902614 | 0,250485 | 2,279193 | 0,022656 | 0,036701 |
| 1628 | RM25_RS07920 | RM25_1611 | 4-hydroxy-3-methylbut-2-enyl diphosphate reductase | 1701 | gene | 1821078 | 1822316 | - | 997,7883 | 0,698630396 | 0,092359 | 7,56429 | 3,9E-14 | 8,36E-12 | 989,3286865 | 0,789210435 | 0,164653 | 4,793169 | 1,64E-06 | 8,04E-06 | 1006,248 | 0,611088264 | 0,144925 | 4,216586 | 2,48E-05 | 0,000225 | 802,7031 | 0,078165 | 0,174505 | 0,447925 | 0,654207 | 0,70895 | 1209,402979 | -0,052097991 | 0,112399 | -0,46351 | 0,642999 | 0,69841 |
| 1629 | RM25_RS07925 | RM25_1612 | Exodeoxyribonuclease 7 large subunit | 1702 | gene | 1822333 | 1823484 | + | 641,8706 | -0,47077785 | 0,27844 | -1,69077 | 0,090881 | 0,181994 | 475,4445038 | -0,337282054 | 0,19794 | -1,70396 | 0,088389 | 0,127109 | 808,2967 | -0,551698695 | 0,258624 | -2,1332 | 0,032908 | 0,074224 | 782,6296 | 0,791934 | 0,216111 | 3,664469 | 0,000248 | 0,001086 | 528,5097007 | 0,639461534 | 0,246345 | 2,595794 | 0,009437 | 0,016563 |
| 1630 | RM25_RS07930 | RM25_1613 | Exodeoxyribonuclease 7 small subunit | 1703 | gene | 1823532 | 1823774 | + | 230,1074 | 0,279660432 | 0,184832 | 1,513053 | 0,130266 | 0,228857 | 194,9315092 | 0,036333036 | 0,199037 | 0,182545 | 0,855155 | 0,886592 | 265,2833 | 0,453982082 | 0,189258 | 2,398751 | 0,016451 | 0,044264 | 219,2502 | 0,155795 | 0,164751 | 0,94564 | 0,344332 | 0,41926 | 246,8379977 | 0,624765299 | 0,194283 | 3,215756 | 0,001301 | 0,002678 |
| 1631 | RM25_RS07935 | RM25_1614 | Channel protein, hemolysin III family protein | 1704 | gene | 1823819 | 1824556 | - | 814,5515 | 0,626508095 | 0,290652 | 2,155527 | 0,031121 | 0,09285 | 961,8954397 | 1,265284803 | 0,187657 | 6,742539 | 1,56E-11 | 1,76E-10 | 667,2075 | -0,204943274 | 0,226511 | -0,90478 | 0,36558 | 0,472667 | 675,6143 | 0,280478 | 0,228925 | 1,225196 | 0,220501 | 0,287139 | 969,5415413 | -1,143171339 | 0,182641 | -6,25911 | 3,87E-10 | 2,28E-09 |
| 1632 | RM25_RS07940 | RM25_1615 | Isoprenyl transferase 1 | 1705 | gene | 1824755 | 1825564 | + | 1357,432 | 0,666704415 | 0,191093 | 3,488909 | 0,000485 | 0,005985 | 1569,645205 | 0,792566856 | 0,131776 | 6,014494 | 1,8E-09 | 1,49E-08 | 1145,219 | 0,499703058 | 0,250161 | 1,997529 | 0,045768 | 0,09471 | 1110,557 | -0,33667 | 0,149226 | -2,25612 | 0,024063 | 0,047541 | 1630,727431 | -0,583097052 | 0,198701 | -2,93455 | 0,00334 | 0,00637 |
| 1633 | RM25_RS07945 | RM25_1616 | Hypothetical protein | 1706 | gene | 1825577 | 1826095 | - | 396,3997 | 0,073899192 | 0,10033 | 0,736561 | 0,461389 | 0,564957 | 388,1879765 | 0,172860349 | 0,167577 | 1,03153 | 0,302293 | 0,371483 | 404,6115 | -0,019788438 | 0,18428 | -0,10738 | 0,914486 | 0,94042 | 408,4541 | 0,09863 | 0,198492 | 0,496897 | 0,619261 | 0,676994 | 398,039282 | -0,043710856 | 0,152413 | -0,28679 | 0,774271 | 0,815806 |
| 1634 | RM25_RS07950 | RM25_1617 | YyaL | 1707 | gene | 1826133 | 1828817 | - | 1703,23 | -0,151882644 | 0,174599 | -0,86989 | 0,384359 | 0,495316 | 1476,040224 | 0,116475685 | 0,128743 | 0,904718 | 0,365615 | 0,433914 | 1930,421 | -0,357747589 | 0,19686 | -1,81727 | 0,069176 | 0,129609 | 1884,469 | 0,55761 | 0,126478 | 4,408736 | 1,04E-05 | 6,36E-05 | 1584,144942 | 0,137080523 | 0,179189 | 0,765003 | 0,44427 | 0,509559 |
| 1635 | RM25_RS07955 | RM25_1618 | Mycothiol conjugate amidase Mca | 1708 | gene | 1828926 | 1829852 | - | 579,1946 | -0,520235402 | 0,253831 | -2,04953 | 0,04041 | 0,109562 | 455,2766416 | -0,700987699 | 0,207951 | -3,37093 | 0,000749 | 0,002075 | 703,1125 | -0,408142319 | 0,235132 | -1,7358 | 0,082599 | 0,148615 | 718,2863 | 0,442832 | 0,241971 | 1,830101 | 0,067235 | 0,108543 | 467,1343144 | 0,797567093 | 0,213388 | 3,737634 | 0,000186 | 0,00045 |
| 1636 | RM25_RS07960 | RM25_1619 | Transcription elongation factor | 1709 | gene | 1829947 | 1830450 | + | 1959,435 | 0,377062153 | 0,279345 | 1,349808 | 0,177078 | 0,28098 | 2303,303213 | 0,397128135 | 0,200472 | 1,98097 | 0,047595 | 0,075952 | 1615,566 | 0,349681245 | 0,400403 | 0,873323 | 0,382487 | 0,487837 | 1811,09 | -0,53359 | 0,377607 | -1,41309 | 0,15763 | 0,215883 | 2165,519054 | -0,550060602 | 0,234972 | -2,34096 | 0,019234 | 0,031725 |
| 1637 | RM25_RS07965 | RM25_1620 | Putative membrane protein | 1710 | gene | 1830537 | 1830971 | + | 1888,871 | 0,853891063 | 0,397886 | 2,146068 | 0,031868 | 0,094007 | 2423,581768 | 0,451769959 | 0,210283 | 2,148395 | 0,031682 | 0,053646 | 1354,16 | 1,682057376 | 0,300883 | 5,590407 | 2,27E-08 | 4,49E-07 | 1436,063 | -1,7223 | 0,289996 | -5,93904 | 2,87E-09 | 3,59E-08 | 2379,887092 | -0,455035918 | 0,212418 | -2,14217 | 0,032179 | 0,05013 |
| 1638 | RM25_RS07970 | RM25_1621 | Hypothetical protein | 1711 | gene | 1831087 | 1832049 | - | 904,9056 | 0,339207742 | 0,188077 | 1,803557 | 0,071301 | 0,155751 | 926,0162218 | 0,747895626 | 0,177988 | 4,20195 | 2,65E-05 | 0,000104 | 883,7949 | -0,071152954 | 0,235909 | -0,30161 | 0,762948 | 0,82641 | 842,7968 | 0,338559 | 0,269367 | 1,256867 | 0,208802 | 0,273414 | 991,3465022 | -0,435838136 | 0,117126 | -3,7211 | 0,000198 | 0,000479 |
| 1639 | RM25_RS07975 | RM25_1622 | Hypothetical protein | 1712 | gene | 1832135 | 1833565 | - | 1306,366 | 0,287050213 | 0,175507 | 1,635547 | 0,101935 | 0,195234 | 1116,816936 | 0,074479143 | 0,130458 | 0,570907 | 0,568063 | 0,630379 | 1495,916 | 0,447749461 | 0,201619 | 2,220767 | 0,026367 | 0,063169 | 1242,423 | 0,15813 | 0,163456 | 0,967416 | 0,333336 | 0,408142 | 1403,543863 | 0,578481671 | 0,145956 | 3,963395 | 7,39E-05 | 0,000193 |
| 1640 | RM25_RS07980 | RM25_1623 | Uracil-DNA glycosylase | 1713 | gene | 1833746 | 1834435 | + | 397,7317 | -0,078315007 | 0,15832 | -0,49466 | 0,620838 | 0,708627 | 352,4241442 | -0,149748564 | 0,163335 | -0,91682 | 0,359237 | 0,427778 | 443,0393 | -0,024466274 | 0,224964 | -0,10876 | 0,913396 | 0,939709 | 430,4726 | 0,207568 | 0,171754 | 1,208514 | 0,22685 | 0,294592 | 379,5476109 | 0,390093997 | 0,208622 | 1,869863 | 0,061503 | 0,088988 |
| 1641 | RM25_RS07985 | RM25_1624 | Peptide methionine sulfoxide reductase MsrA | 1714 | gene | 1834547 | 1835182 | + | 797,8263 | 0,789610883 | 0,412687 | 1,913343 | 0,055704 | 0,1328 | 456,7108454 | 0,04479996 | 0,212903 | 0,210424 | 0,833337 | 0,867568 | 1138,942 | 1,111143977 | 0,238595 | 4,657038 | 3,21E-06 | 3,8E-05 | 614,9838 | 0,615482 | 0,242723 | 2,535736 | 0,011221 | 0,025904 | 985,8705204 | 1,730183915 | 0,185497 | 9,327282 | 1,09E-20 | 2,61E-19 |
| 1642 | RM25_RS07990 | RM25_1625 | Hypothetical protein | 1715 | gene | 1835309 | 1835839 | + | 290,1046 | -0,038868702 | 0,231082 | -0,1682 | 0,866424 | 0,909648 | 231,086887 | -0,237333874 | 0,204698 | -1,15943 | 0,246279 | 0,310763 | 349,1224 | 0,087927155 | 0,210556 | 0,417596 | 0,676243 | 0,754582 | 309,3637 | 0,372849 | 0,201494 | 1,850428 | 0,064252 | 0,104396 | 280,471676 | 0,755151168 | 0,200795 | 3,760816 | 0,000169 | 0,000415 |
| 1643 | RM25_RS07995 | RM25_1626 | Pyridoxamine 5'-phosphate oxidase-related, FMN-binding p | 1716 | gene | 1835929 | 1836576 | - | 470,5391 | 0,522413903 | 0,185849 | 2,810961 | 0,004939 | 0,029325 | 535,2246833 | 0,513863447 | 0,21224 | 2,421141 | 0,015472 | 0,028851 | 405,8536 | 0,535327241 | 0,20231 | 2,64608 | 0,008143 | 0,026256 | 409,2451 | -0,46862 | 0,222741 | -2,10389 | 0,035388 | 0,06481 | 544,3324244 | -0,393878427 | 0,202332 | -1,94669 | 0,051572 | 0,075829 |
| 1644 | RM25_RS08000 | RM25_1627 | Putative hydrolase | 1717 | gene | 1836754 | 1837368 | + | 443,1561 | -0,531247051 | 0,188284 | -2,82152 | 0,00478 | 0,028886 | 370,8456102 | -0,480561567 | 0,180923 | -2,65617 | 0,007903 | 0,016227 | 515,4665 | -0,570801659 | 0,200228 | -2,85076 | 0,004362 | 0,016215 | 551,633 | 0,448035 | 0,21435 | 2,090198 | 0,0366 | 0,066395 | 355,5586672 | 0,416068868 | 0,17535 | 2,372788 | 0,017654 | 0,029304 |
| 1645 | RM25_RS08005 | RM25_1628 | Hypothetical protein | 1718 | gene | 1837365 | 1837871 | + | 264,5995 | 0,219045954 | 0,120418 | 1,819052 | 0,068904 | 0,152089 | 246,8600743 | 0,31535348 | 0,177366 | 1,77798 | 0,075407 | 0,111363 | 282,3389 | 0,137496078 | 0,192446 | 0,714468 | 0,474938 | 0,577025 | 257,8113 | 0,230586 | 0,169988 | 1,356486 | 0,174945 | 0,23576 | 279,2897403 | 0,110798827 | 0,195927 | 0,565511 | 0,571726 | 0,632657 |
| 1646 | RM25_RS08010 | RM25_1629 | Hypothetical protein | 1719 | gene | 1837897 | 1838352 | - | 699,4528 | 0,506443248 | 0,208687 | 2,426809 | 0,015232 | 0,061278 | 628,0806761 | 0,500127552 | 0,21286 | 2,349563 | 0,018795 | 0,033973 | 770,825 | 0,512986789 | 0,315612 | 1,625372 | 0,104083 | 0,177258 | 610,8761 | 0,234467 | 0,32655 | 0,718011 | 0,472751 | 0,545131 | 802,1997263 | 0,28651604 | 0,188245 | 1,52204 | 0,127999 | 0,172298 |
| 1647 | RM25_RS08015 | RM25_1630 | Hypothetical protein | 1720 | gene | 1838486 | 1838890 | - | 3252,463 | -0,425851958 | 0,452152 | -0,94183 | 0,346277 | 0,458784 | 2078,020233 | 0,685005732 | 0,195894 | 3,496812 | 0,000471 | 0,001377 | 4426,906 | -0,971326908 | 0,441331 | -2,2009 | 0,027743 | 0,065402 | 3913,37 | 1,833022 | 0,404613 | 4,530308 | 5,89E-06 | 3,84E-05 | 2709,873236 | 0,204784879 | 0,256608 | 0,798045 | 0,424844 | 0,490862 |
| 1648 | NA | RM25_1631 | Hypothetical protein | 1721 | gene | 1838903 | 1838992 | - | 631,3201 | -0,065797611 | 0,489033 | -0,13455 | 0,892971 | 0,928421 | 428,7633889 | 0,881559721 | 0,309081 | 2,852198 | 0,004342 | 0,009627 | 833,8768 | -0,543628907 | 0,56134 | -0,96845 | 0,33282 | 0,438244 | 678,6904 | 1,66966 | 0,482323 | 3,461706 | 0,000537 | 0,002095 | 599,9781498 | 0,261145437 | 0,41694 | 0,626338 | 0,531093 | 0,591861 |
| 1649 | RM25_RS08020 | RM25_1632 | Putative 3-methyladenine DNA glycosylase | 1722 | gene | 1839020 | 1839670 | - | 548,4671 | 0,515310635 | 0,15011 | 3,432883 | 0,000597 | 0,007038 | 583,5007293 | 0,782926778 | 0,183404 | 4,268875 | 1,96E-05 | 7,84E-05 | 513,4335 | 0,226394254 | 0,182147 | 1,24292 | 0,213897 | 0,311564 | 476,7439 | 0,08499 | 0,205174 | 0,414236 | 0,678701 | 0,729457 | 632,5628554 | -0,423273168 | 0,155698 | -2,71856 | 0,006557 | 0,011824 |
| 1650 | RM25_RS11945 | RM25_1633 | Hypothetical protein | 1723 | gene | 1839879 | 1840829 | + | 898,1827 | 0,094012283 | 0,148074 | 0,634899 | 0,525494 | 0,622094 | 985,103821 | 0,300375585 | 0,138747 | 2,16492 | 0,030394 | 0,051784 | 811,2616 | -0,153167585 | 0,181667 | -0,84312 | 0,39916 | 0,504899 | 918,1 | -0,10868 | 0,138346 | -0,78556 | 0,432126 | 0,507232 | 908,968649 | -0,512816377 | 0,153196 | -3,34744 | 0,000816 | 0,001768 |
| 1651 | RM25_RS12320 | RM25_1634 | Hydrolase, NUDIX family | 1724 | gene | 1840959 | 1841582 | + | 678,0824 | 0,577286872 | 0,182011 | 3,171709 | 0,001515 | 0,012787 | 560,7735669 | 0,535166176 | 0,140958 | 3,796624 | 0,000147 | 0,000482 | 795,3913 | 0,607810065 | 0,165609 | 3,670152 | 0,000242 | 0,00154 | 573,413 | 0,39984 | 0,144598 | 2,765184 | 0,005689 | 0,014687 | 794,6224718 | 0,523088622 | 0,1378 | 3,795991 | 0,000147 | 0,000365 |
| 1652 | RM25_RS08035 | RM25_1635 | Putative CONSERVED TRANSMEMBRANE PROTEIN | 1725 | gene | 1841757 | 1842569 | - | 2231,785 | 1,25284037 | 0,302988 | 4,134948 | 3,55E-05 | 0,000761 | 2825,149952 | 1,132276962 | 0,176607 | 6,411268 | 1,44E-10 | 1,38E-09 | 1638,419 | 1,473331254 | 0,245522 | 6,000807 | 1,96E-09 | 5,2E-08 | 1404,449 | -1,08324 | 0,252034 | -4,298 | 1,72E-05 | 0,0001 | 3080,547862 | -0,699506143 | 0,159057 | -4,39783 | 1,09E-05 | 3,19E-05 |
| 1653 | RM25_RS08040 | RM25_1636 | Methyltransferase type 11 | 1726 | gene | 1842706 | 1843455 | - | 325,8859 | -0,532086041 | 0,342388 | -1,55404 | 0,120174 | 0,216718 | 225,7759035 | -0,83238581 | 0,278256 | -2,99143 | 0,002777 | 0,006538 | 425,9959 | -0,382030689 | 0,229059 | -1,66782 | 0,095351 | 0,166229 | 405,0453 | 0,667809 | 0,30185 | 2,212387 | 0,02694 | 0,051708 | 261,2434129 | 1,180832625 | 0,220377 | 5,358231 | 8,4E-08 | 3,47E-07 |
| 1654 | RM25_RS08045 | RM25_1637 | putative nucleoside-diphosphate-sugar epimerases | 1727 | gene | 1843758 | 1844531 | + | 896,761 | -0,15540453 | 0,36767 | -0,42267 | 0,672533 | 0,752688 | 552,2580135 | -0,138449493 | 0,148762 | -0,93068 | 0,352021 | 0,420926 | 1241,264 | -0,163263704 | 0,2217 | -0,73642 | 0,461477 | 0,565334 | 993,3537 | 1,122297 | 0,223736 | 5,016156 | 5,27E-07 | 4,2E-06 | 829,7781144 | 1,143786247 | 0,147981 | 7,729279 | 1,08E-14 | 1,26E-13 |
| 1655 | RM25_RS08050 | RM25_1638 | Putative CATIONIC AMINO ACID TRANSPORT INTEGRAL MEI | 1728 | gene | 1844748 | 1846211 | + | 2186,327 | 0,743821395 | 0,27495 | 2,705295 | 0,006824 | 0,036724 | 1860,822716 | -0,035491752 | 0,177239 | -0,20025 | 0,841287 | 0,874686 | 2511,831 | 1,391936339 | 0,244445 | 5,694269 | 1,24E-08 | 2,61E-07 | 1733,718 | -0,50015 | 0,211452 | -2,36529 | 0,018016 | 0,037678 | 2671,73506 | 0,969386374 | 0,190131 | 5,09851 | 3,42E-07 | 1,28E-06 |
| 1656 | RM25_RS08055 | RM25_1639 | Phosphopyruvate hydratase | 1729 | gene | 1846269 | 1847540 | - | 499,9723 | -1,241728804 | 0,489226 | -2,53815 | 0,011144 | 0,050223 | 300,567673 | -2,05664796 | 0,316153 | -6,50522 | 7,76E-11 | 7,68E-10 | 699,3769 | -0,948154105 | 0,281337 | -3,37017 | 0,000751 | 0,003858 | 736,8599 | 0,859147 | 0,351016 | 2,447599 | 0,014381 | 0,031589 | 291,8848509 | 2,032980091 | 0,27101 | 7,501495 | 6,31E-14 | 6,55E-13 |
| 1657 | RM25_RS08060 | RM25_1640 | Transcriptional regulator | 1730 | gene | 1847831 | 1848469 | - | 522,9708 | 0,382261563 | 0,209212 | 1,827146 | 0,067678 | 0,150204 | 414,6893414 | 0,250307032 | 0,145989 | 1,71456 | 0,086426 | 0,124818 | 631,2524 | 0,467874455 | 0,154417 | 3,029933 | 0,002446 | 0,010204 | 477,9895 | 0,421859 | 0,140957 | 2,992824 | 0,002764 | 0,008302 | 579,5966429 | 0,693254433 | 0,132806 | 5,220046 | 1,79E-07 | 6,97E-07 |
| 1658 | RM25_RS08065 | RM25_1641 | Actinorhodin transporter | 1731 | gene | 1848534 | 1849961 | + | 545,7795 | -1,734480354 | 0,767988 | -2,25847 | 0,023916 | 0,079172 | 292,1612441 | -4,363077866 | 0,391014 | -11,1584 | 6,52E-29 | 5,69E-27 | 799,3977 | -1,232682687 | 0,431079 | -2,85953 | 0,004243 | 0,015823 | 879,5423 | 0,937537 | 0,401101 | 2,337411 | 0,019418 | 0,040007 | 249,1149286 | 4,145604258 | 0,456474 | 9,081798 | 1,07E-19 | 2,47E-18 |
| 1659 | RM25_RS08070 | RM25_1642 | Multidrug ABC transporter, ATP-binding protein | 1732 | gene | 1850109 | 1850909 | + | 326,3077 | -1,163729157 | 0,288056 | -4,03994 | 5,35E-05 | 0,001059 | 253,6507689 | -1,125372905 | 0,255711 | -4,40095 | 1,08E-05 | 4,5E-05 | 398,9646 | -1,192332101 | 0,342613 | -3,48011 | 0,000501 | 0,002793 | 473,9853 | 0,609167 | 0,283456 | 2,149071 | 0,031629 | 0,059072 | 198,137317 | 0,608542074 | 0,343034 | 1,774002 | 0,076063 | 0,107353 |
| 1660 | RM25_RS08075 | RM25_1643 | Putative ABC-2 | 1733 | gene | 1851027 | 1851800 | + | 455,2169 | -0,956593422 | 0,391063 | -2,44614 | 0,01444 | 0,0595 | 298,8223875 | -1,276909163 | 0,304739 | -4,19017 | 2,79E-05 | 0,000109 | 611,6114 | -0,811238042 | 0,265853 | -3,05145 | 0,002277 | 0,009585 | 630,6528 | 0,81219 | 0,321107 | 2,529345 | 0,011428 | 0,026278 | 303,8007207 | 1,342982209 | 0,278212 | 4,827188 | 1,38E-06 | 4,68E-06 |
| 1661 | RM25_RS08080 | RM25_1644 | Tetracycline repressor protein | 1734 | gene | 1851797 | 1852435 | + | 552,1689 | -1,341077056 | 0,468732 | -2,86107 | 0,004222 | 0,026396 | 336,5351123 | -1,932794531 | 0,322021 | -6,00207 | 1,95E-09 | 1,6E-08 | 767,8027 | -1,115610613 | 0,313055 | -3,56362 | 0,000366 | 0,002188 | 830,2547 | 0,909537 | 0,338502 | 2,686942 | 0,007211 | 0,018006 | 307,3649445 | 1,795294657 | 0,331227 | 5,420135 | 5,96E-08 | 2,56E-07 |
| 1663 | RM25_RS08090 | RM25_1645 | Ppx/GppA phosphatase family protein | 1738 | gene | 1852732 | 1853688 | - | 993,1298 | 0,668426712 | 0,097778 | 6,836193 | 8,13E-12 | 1,37E-09 | 992,5570684 | 0,568510487 | 0,130819 | 4,345778 | 1,39E-05 | 5,71E-05 | 993,7024 | 0,769612527 | 0,200794 | 3,832852 | 0,000127 | 0,000897 | 810,9191 | -0,1841 | 0,164619 | -1,11834 | 0,26342 | 0,334887 | 1193,22557 | 0,067842018 | 0,133898 | 0,50667 | 0,612386 | 0,671659 |
| 1664 | RM25_RS08095 | RM25_1646 | PF04417 family protein | 1739 | gene | 1853685 | 1854200 | - | 773,075 | 0,934157499 | 0,163481 | 5,714183 | 1,1E-08 | 9,62E-07 | 668,7807962 | 0,916689131 | 0,155089 | 5,910735 | 3,41E-09 | 2,71E-08 | 877,3693 | 0,948899059 | 0,197814 | 4,796924 | 1,61E-06 | 2,1E-05 | 560,1754 | 0,31083 | 0,17462 | 1,780039 | 0,07507 | 0,117364 | 992,5612195 | 0,390418132 | 0,154814 | 2,521846 | 0,011674 | 0,020158 |
| 1666 | RM25_RS08100 | RM25_1647 | Hypothetical protein | 1740 | gene | 1854280 | 1854921 | - | 740,1427 | 0,035112073 | 0,185052 | 0,189742 | 0,849512 | 0,90153 | 657,9199288 | 0,354137919 | 0,178076 | 1,988694 | 0,046735 | 0,074984 | 822,3655 | -0,216217278 | 0,209452 | -1,0323 | 0,301931 | 0,406194 | 769,1733 | 0,55764 | 0,224988 | 2,478536 | 0,013192 | 0,029739 | 734,6500326 | 0,038137675 | 0,156113 | 0,244295 | 0,807003 | 0,843506 |
| 1667 | RM25_RS08105 | RM25_1648 | Phosphopyruvate hydratase | 1741 | gene | 1855199 | 1856488 | - | 15513,03 | 0,533111755 | 0,390626 | 1,364761 | 0,172328 | 0,276064 | 20539,32036 | 0,468747988 | 0,213562 | 2,194899 | 0,028171 | 0,04829 | 10486,74 | 0,661002379 | 0,447156 | 1,478235 | 0,139345 | 0,222819 | 13509,9 | -1,13041 | 0,415786 | -2,71872 | 0,006554 | 0,016574 | 17954,30305 | -0,911127073 | 0,263678 | -3,45546 | 0,000549 | 0,001227 |
| 1668 | RM25_RS08110 | RM25_1649 | Nucleoside triphosphate pyrophosphohydrolase | 1742 | gene | 1856603 | 1857274 | - | 397,5425 | 0,065681692 | 0,151993 | 0,432137 | 0,665642 | 0,747264 | 433,714694 | 0,065473352 | 0,163647 | 0,400088 | 0,689091 | 0,741639 | 361,3702 | 0,067858324 | 0,246839 | 0,274909 | 0,783386 | 0,843124 | 411,1481 | -0,33127 | 0,220974 | -1,49913 | 0,133841 | 0,189013 | 399,0984606 | -0,266160081 | 0,195815 | -1,35924 | 0,17407 | 0,226148 |
| 1669 | RM25_RS08115 | RM25_1650 | Hypothetical protein | 1743 | gene | 1857274 | 1857807 | - | 555,0494 | 0,190899617 | 0,235863 | 0,809366 | 0,418305 | 0,527808 | 662,2176131 | 0,037858543 | 0,200571 | 0,188754 | 0,850286 | 0,882097 | 447,8813 | 0,42244924 | 0,217325 | 1,943859 | 0,051912 | 0,103605 | 551,3594 | -0,83059 | 0,251404 | -3,30382 | 0,000954 | 0,003381 | 579,6728662 | -0,39701734 | 0,165498 | -2,39892 | 0,016444 | 0,027527 |
| 1670 | RM25_RS08120 | RM25_1651 | Putative insertion element ISCmi2 transposase | 1744 | gene | 1857929 | 1858972 | - | 60,52485 | -0,388346186 | 0,298638 | -1,30039 | 0,193466 | 0,297456 | 53,75278277 | -0,735771171 | 0,428207 | -1,71826 | 0,085749 | 0,124222 | 67,29691 | -0,130357151 | 0,353065 | -0,36922 | 0,711967 | 0,785262 | 72,75141 | -0,00863 | 0,359676 | -0,02399 | 0,980862 | 0,983366 | 51,30783025 | 0,669734639 | 0,461411 | 1,451492 | 0,146643 | 0,194616 |
| 1671 | RM25_RS08125 | RM25_1652 | Transcription-repair coupling factor | 1745 | gene | 1859028 | 1862723 | - | 2731,341 | 0,193263106 | 0,110124 | 1,75496 | 0,079266 | 0,166515 | 2881,426526 | 0,287320671 | 0,127871 | 2,246959 | 0,024643 | 0,042834 | 2581,255 | 0,089649946 | 0,19284 | 0,464893 | 0,642008 | 0,723334 | 2689,586 | -0,11575 | 0,161285 | -0,71765 | 0,472971 | 0,545131 | 2857,631564 | -0,258668437 | 0,132323 | -1,95482 | 0,050604 | 0,074733 |
| 1672 | RM25_RS08130 | RM25_1653 | Hypothetical protein | 1746 | gene | 1862802 | 1863167 | - | 202,8792 | -0,184161817 | 0,32185 | -0,5722 | 0,567188 | 0,660504 | 140,8112866 | 0,009447393 | 0,256693 | 0,036804 | 0,970641 | 0,976024 | 264,947 | -0,284913449 | 0,251457 | -1,13305 | 0,257193 | 0,3587 | 225,944 | 1,001757 | 0,232292 | 4,312492 | 1,61E-05 | 9,42E-05 | 186,4149389 | 0,758987744 | 0,262044 | 2,89641 | 0,003775 | 0,007129 |
| 1673 | RM25_RS08135 | RM25_1654 | Aminoacyl-tRNA hydrolase | 1747 | gene | 1863235 | 1863825 | - | 322,9109 | -0,129952918 | 0,17974 | -0,72301 | 0,469676 | 0,570743 | 354,2258598 | -0,06477773 | 0,191148 | -0,33889 | 0,734694 | 0,783563 | 291,596 | -0,207413356 | 0,279972 | -0,74084 | 0,458793 | 0,56263 | 357,1271 | -0,27537 | 0,196235 | -1,40325 | 0,160541 | 0,21936 | 303,0548764 | -0,358152886 | 0,280933 | -1,27487 | 0,202356 | 0,256564 |
| 1674 | RM25_RS08140 | RM25_1655 | 50S ribosomal protein L25 | 1748 | gene | 1864033 | 1864692 | - | 2806,813 | 0,962871296 | 0,468151 | 2,056755 | 0,03971 | 0,108203 | 3874,215753 | 0,682390523 | 0,221642 | 3,078789 | 0,002078 | 0,005087 | 1739,41 | 1,670945351 | 0,402743 | 4,148916 | 3,34E-05 | 0,000291 | 2034,928 | -1,89174 | 0,420831 | -4,49525 | 6,95E-06 | 4,45E-05 | 3634,991744 | -0,86441514 | 0,175789 | -4,91736 | 8,77E-07 | 3,1E-06 |
| 1675 | RM25_RS08145 | RM25_1656 | Ribose-phosphate diphosphokinase | 1749 | gene | 1865070 | 1866065 | - | 2707,868 | 0,547931193 | 0,373729 | 1,466118 | 0,142616 | 0,243286 | 3504,317459 | 0,188324962 | 0,189097 | 0,995915 | 0,319291 | 0,386925 | 1911,419 | 1,26544517 | 0,282931 | 4,472635 | 7,73E-06 | 8,17E-05 | 2346,309 | -1,60095 | 0,270017 | -5,92908 | 3,05E-09 | 3,8E-08 | 3148,436531 | -0,481611328 | 0,188245 | -2,55843 | 0,010515 | 0,018317 |
| 1676 | RM25_RS08150 | RM25_1657 | Bifunctional protein GlmU | 1750 | gene | 1866099 | 1867646 | - | 3704,671 | 0,521439638 | 0,271245 | 1,922394 | 0,054556 | 0,131751 | 4370,104247 | 0,267903609 | 0,188362 | 1,422279 | 0,154945 | 0,209167 | 3039,237 | 0,902251309 | 0,307428 | 2,934842 | 0,003337 | 0,013153 | 3231,365 | -0,95672 | 0,30473 | -3,13958 | 0,001692 | 0,005463 | 4274,653546 | -0,283322602 | 0,17167 | -1,65039 | 0,098863 | 0,136509 |
| 1678 | RM25_RS08155 | RM25_1658 | Aspartate carbamoyltransferase | 1751 | gene | 1868073 | 1869119 | + | 1163,619 | 0,303696065 | 0,087614 | 3,466294 | 0,000528 | 0,006342 | 1212,164868 | 0,322936174 | 0,11684 | 2,763919 | 0,005711 | 0,012338 | 1115,074 | 0,284968021 | 0,18419 | 1,547144 | 0,121828 | 0,200939 | 1099,842 | -0,16172 | 0,150794 | -1,07245 | 0,283517 | 0,357161 | 1260,295388 | -0,146832389 | 0,118647 | -1,23756 | 0,215879 | 0,271809 |
| 1679 | RM25_RS08160 | RM25_1659 | Transcriptional regulator, TetR family | 1752 | gene | 1869262 | 1870023 | + | 505,2193 | -0,134993542 | 0,127605 | -1,0579 | 0,2901 | 0,399163 | 483,1006054 | 0,049232601 | 0,1659 | 0,296761 | 0,766649 | 0,811766 | 527,3381 | -0,301312479 | 0,20352 | -1,48051 | 0,138738 | 0,22215 | 556,7184 | 0,233165 | 0,158187 | 1,473981 | 0,140487 | 0,196282 | 472,6830289 | -0,061467157 | 0,196166 | -0,31334 | 0,754021 | 0,79732 |
| 1680 | RM25_RS08165 | RM25_1660 | 4-(cytidine 5'-diphospho)-2-C-methyl-D-erythritol kinase | 1753 | gene | 1870058 | 1871035 | - | 677,952 | -0,245961015 | 0,217057 | -1,13316 | 0,257146 | 0,364457 | 581,753179 | 0,124711959 | 0,17329 | 0,719671 | 0,471727 | 0,538954 | 774,1508 | -0,527328482 | 0,266385 | -1,97957 | 0,047751 | 0,09753 | 773,5456 | 0,660752 | 0,174176 | 3,793584 | 0,000148 | 0,000692 | 609,6262848 | 0,063643988 | 0,261225 | 0,243636 | 0,807513 | 0,843665 |
| 1681 | RM25_RS08170 | RM25_1661 | Ribosomal RNA small subunit methyltransferase A | 1754 | gene | 1871112 | 1871981 | - | 420,7064 | -0,153530267 | 0,205345 | -0,74767 | 0,454659 | 0,560183 | 385,7906107 | -0,052746797 | 0,149653 | -0,35246 | 0,724492 | 0,774084 | 455,6221 | -0,237828854 | 0,357562 | -0,66514 | 0,505961 | 0,605971 | 466,3576 | 0,26612 | 0,196725 | 1,352752 | 0,176135 | 0,236958 | 392,1350015 | 0,149124512 | 0,330034 | 0,451847 | 0,65138 | 0,70621 |
| 1682 | RM25_RS08175 | RM25_1662 | Hydrolase, TatD family | 1755 | gene | 1871978 | 1872934 | - | 615,4379 | -0,276828877 | 0,14433 | -1,91802 | 0,055108 | 0,132317 | 620,4068123 | -0,221840173 | 0,139193 | -1,59376 | 0,11099 | 0,15581 | 610,4689 | -0,332736102 | 0,285452 | -1,16565 | 0,243758 | 0,343471 | 711,1956 | -0,03505 | 0,189409 | -0,18503 | 0,853203 | 0,884734 | 547,0492366 | -0,084604884 | 0,247943 | -0,34123 | 0,732932 | 0,778163 |
| 1683 | RM25_RS08180 | RM25_1663 | Ribosomal RNA small subunit methyltransferase I | 1756 | gene | 1873139 | 1874089 | - | 677,8704 | -0,154563388 | 0,145218 | -1,06435 | 0,287169 | 0,396751 | 682,4750848 | 0,041914556 | 0,147578 | 0,284016 | 0,776398 | 0,820983 | 673,2657 | -0,35266144 | 0,257853 | -1,36769 | 0,17141 | 0,262006 | 752,2745 | 0,107913 | 0,156114 | 0,691246 | 0,489411 | 0,558345 | 630,8531417 | -0,227081514 | 0,239954 | -0,94635 | 0,343968 | 0,408018 |
| 1684 | RM25_RS08185 | RM25_1664 | Alpha-L-fucosidase (Precursor) | 1757 | gene | 1874103 | 1876490 | - | 1030,002 | 0,53470472 | 0,259815 | 2,058019 | 0,039588 | 0,108122 | 983,3219349 | 0,888900496 | 0,243119 | 3,656236 | 0,000256 | 0,000802 | 1076,682 | 0,227181341 | 0,373474 | 0,608291 | 0,542994 | 0,639919 | 880,3859 | 0,468829 | 0,320579 | 1,462442 | 0,14362 | 0,200067 | 1199,289049 | -0,132760035 | 0,297312 | -0,44653 | 0,655211 | 0,709712 |
| 1685 | RM25_RS08190 | RM25_1665 | Dolichyl-phosphate-mannose-protein mannosyltransferase | 1758 | gene | 1876778 | 1878337 | + | 1545,031 | -0,142992648 | 0,072598 | -1,96966 | 0,048877 | 0,122949 | 1499,386509 | -0,184295558 | 0,127952 | -1,44035 | 0,149768 | 0,203577 | 1590,676 | -0,105085798 | 0,148971 | -0,70541 | 0,480542 | 0,581452 | 1713,506 | -0,01386 | 0,128922 | -0,10754 | 0,914364 | 0,931758 | 1438,293444 | 0,117495963 | 0,131068 | 0,896453 | 0,370011 | 0,434753 |
| 1686 | RM25_RS08195 | RM25_1666 | Integral membrane protein with DUF218 domain protein | 1759 | gene | 1878351 | 1879451 | + | 853,7404 | -0,277324209 | 0,160708 | -1,72564 | 0,084412 | 0,173158 | 743,9104855 | -0,291763161 | 0,138629 | -2,10463 | 0,035323 | 0,058964 | 963,5704 | -0,267288702 | 0,212083 | -1,2603 | 0,20756 | 0,30424 | 986,2672 | 0,296876 | 0,192845 | 1,53945 | 0,123694 | 0,17691 | 757,0554856 | 0,381187606 | 0,167013 | 2,282377 | 0,022467 | 0,036445 |
| 1687 | RM25_RS08200 | RM25_1667 | Pyrimidine utilization transport protein G | 1760 | gene | 1879573 | 1881126 | + | 1061,254 | -0,38874817 | 0,140441 | -2,76806 | 0,005639 | 0,03195 | 1139,043926 | -0,355129878 | 0,13938 | -2,54792 | 0,010837 | 0,021268 | 983,4649 | -0,42679034 | 0,242509 | -1,75989 | 0,078426 | 0,143072 | 1271,149 | -0,24351 | 0,183362 | -1,32801 | 0,184175 | 0,245671 | 902,7410615 | -0,255383073 | 0,199095 | -1,28272 | 0,199591 | 0,253742 |
| 1688 | RM25_RS08205 | RM25_1668 | Hypothetical protein | 1761 | gene | 1881157 | 1881678 | - | 755,4049 | 0,218337813 | 0,235545 | 0,926946 | 0,353955 | 0,466073 | 614,5356974 | 0,426691984 | 0,143808 | 2,967085 | 0,003007 | 0,007016 | 896,2742 | 0,078542121 | 0,286188 | 0,274443 | 0,783744 | 0,843124 | 736,0882 | 0,677811 | 0,269779 | 2,512472 | 0,011989 | 0,027382 | 794,4004123 | 0,372347393 | 0,166 | 2,243052 | 0,024893 | 0,039969 |
| 1689 | RM25_RS08210 | RM25_1669 | Galactokinase | 1762 | gene | 1881942 | 1883138 | + | 2294,369 | -0,083206715 | 0,355967 | -0,23375 | 0,81518 | 0,871374 | 2258,578666 | 0,953756887 | 0,174039 | 5,480123 | 4,25E-08 | 2,77E-07 | 2330,159 | -1,105334302 | 0,326212 | -3,38839 | 0,000703 | 0,003642 | 2483,684 | 0,999006 | 0,34937 | 2,859451 | 0,004244 | 0,011537 | 2186,559348 | -1,01759891 | 0,108997 | -9,33606 | 1E-20 | 2,48E-19 |
| 1690 | RM25_RS08215 | RM25_1670 | UspA domain protein | 1763 | gene | 1883295 | 1883828 | + | 429,2168 | -0,40384467 | 0,125709 | -3,21252 | 0,001316 | 0,011882 | 420,6530444 | -0,391833196 | 0,169829 | -2,30722 | 0,021043 | 0,037292 | 437,7806 | -0,41579173 | 0,240428 | -1,72938 | 0,083741 | 0,150177 | 515,9467 | 0,008381 | 0,153754 | 0,054511 | 0,956528 | 0,966305 | 362,6198734 | 0,041504866 | 0,241925 | 0,171561 | 0,863783 | 0,889292 |
| 1691 | RM25_RS08220 | RM25_1671 | CrcB-like protein | 1764 | gene | 1883825 | 1884241 | + | 176,4608 | -0,559236896 | 0,277637 | -2,01428 | 0,043981 | 0,114418 | 140,6635619 | -0,436752278 | 0,319841 | -1,36553 | 0,172087 | 0,228512 | 212,258 | -0,643330948 | 0,295037 | -2,18051 | 0,02922 | 0,067987 | 221,2731 | 0,605237 | 0,279237 | 2,167469 | 0,030199 | 0,056898 | 140,2188548 | 0,468974524 | 0,362884 | 1,292353 | 0,196235 | 0,250692 |
| 1692 | RM25_RS08225 | RM25_1672 | CrcB-like protein | 1765 | gene | 1884238 | 1884624 | + | 173,9554 | -1,155789273 | 0,449076 | -2,5737 | 0,010062 | 0,047336 | 116,0212182 | -1,744937489 | 0,366875 | -4,75622 | 1,97E-06 | 9,52E-06 | 231,8896 | -0,89860684 | 0,394191 | -2,27962 | 0,02263 | 0,05643 | 252,4913 | 0,684752 | 0,300811 | 2,276251 | 0,022831 | 0,04545 | 106,1087016 | 1,604924713 | 0,484963 | 3,309376 | 0,000935 | 0,002002 |
| 1693 | RM25_RS08230 | RM25_1673 | Acetyltransferase | 1766 | gene | 1884684 | 1885157 | + | 228,8212 | -0,633529808 | 0,189933 | -3,33554 | 0,000851 | 0,009121 | 192,6693409 | -0,540521297 | 0,220219 | -2,45447 | 0,014109 | 0,02669 | 264,9731 | -0,706472869 | 0,205326 | -3,44073 | 0,00058 | 0,003158 | 292,6407 | 0,459139 | 0,220169 | 2,085388 | 0,037034 | 0,066888 | 175,9739765 | 0,352481116 | 0,208749 | 1,688538 | 0,091308 | 0,126745 |
| 1694 | RM25_RS08235 | RM25_1674 | Transposase IS204/IS1001/IS1096/IS1165 family protein | 1767 | gene | 1885232 | 1886308 | - | 11,7604 | -1,845085077 | 0,686073 | -2,68934 | 0,007159 | 0,038248 | 6,038056899 | -2,485563249 | 1,2665 | -1,96254 | 0,049699 | 0,078671 | 17,48275 | -1,673008656 | 0,424662 | -3,93963 | 8,16E-05 | 0,000617 | 19,35827 | 1,253959 | 0,468282 | 2,677784 | 0,007411 | 0,018407 | 4,933125403 | 2,161985694 | 1,343185 | 1,609597 | 0,107486 | 0,147199 |
| 1695 | RM25_RS08240 | RM25_1675 | Glutamate dehydrogenase | 1768 | gene | 1886724 | 1888088 | - | 2692,016 | 0,802309453 | 0,460369 | 1,742751 | 0,081377 | 0,16851 | 4010,623342 | 0,857511556 | 0,179118 | 4,787412 | 1,69E-06 | 8,24E-06 | 1373,408 | 0,646354855 | 0,25032 | 2,582116 | 0,00982 | 0,029788 | 2091,648 | -1,46869 | 0,251375 | -5,84264 | 5,14E-09 | 6E-08 | 3355,661674 | -1,639601085 | 0,157438 | -10,4143 | 2,13E-25 | 8,53E-24 |
| 1696 | RM25_RS08245 | RM25_1676 | Multidrug resistance transporter, MFS superfamily protein | 1769 | gene | 1888243 | 1889730 | - | 489,7534 | -0,867953756 | 0,385799 | -2,24976 | 0,024464 | 0,079665 | 334,3162862 | -1,10152433 | 0,275828 | -3,99353 | 6,51E-05 | 0,000232 | 645,1905 | -0,753617006 | 0,389785 | -1,93342 | 0,053185 | 0,105253 | 664,9171 | 0,757313 | 0,308937 | 2,451354 | 0,014232 | 0,03138 | 341,6116888 | 1,180097635 | 0,381625 | 3,092295 | 0,001986 | 0,003951 |
| 1697 | RM25_RS08250 | RM25_1677 | Regulatory protein, TetR | 1770 | gene | 1889843 | 1890436 | + | 310,7455 | -0,766507421 | 0,298625 | -2,56679 | 0,010265 | 0,048003 | 244,7288954 | -0,795553621 | 0,266521 | -2,98496 | 0,002836 | 0,006665 | 376,762 | -0,750054073 | 0,369783 | -2,02836 | 0,042523 | 0,089729 | 411,8633 | 0,538085 | 0,259588 | 2,072846 | 0,038187 | 0,068393 | 226,3740149 | 0,653799684 | 0,388162 | 1,684348 | 0,092114 | 0,127789 |
| 1698 | RM25_RS08255 | RM25_1678 | Proline/betaine transporter | 1771 | gene | 1890512 | 1891951 | - | 1119,646 | -0,054105206 | 0,123914 | -0,43664 | 0,662375 | 0,7445 | 1167,595107 | 0,029622754 | 0,122201 | 0,242411 | 0,808462 | 0,848417 | 1071,697 | -0,143468472 | 0,239379 | -0,59934 | 0,548949 | 0,645799 | 1202,386 | -0,1014 | 0,195285 | -0,51925 | 0,60359 | 0,663555 | 1077,625558 | -0,219719434 | 0,149565 | -1,46905 | 0,141818 | 0,188425 |
| 1699 | RM25_RS08260 | RM25_1679 | Drug resistance transporter, EmrB/QacA subfamily | 1772 | gene | 1892090 | 1893556 | - | 418,4267 | -1,341563867 | 0,588819 | -2,2784 | 0,022703 | 0,077551 | 233,0924466 | -2,592173395 | 0,364892 | -7,10395 | 1,21E-12 | 1,6E-11 | 603,7609 | -0,97553024 | 0,374308 | -2,60623 | 0,009155 | 0,02828 | 629,1222 | 0,932049 | 0,380464 | 2,449767 | 0,014295 | 0,031489 | 233,2547704 | 2,620779828 | 0,391857 | 6,688105 | 2,26E-11 | 1,61E-10 |
| 1700 | RM25_RS08265 | RM25_1680 | Transcriptional regulator | 1773 | gene | 1893596 | 1893868 | - | 257,3643 | 0,183699447 | 0,245102 | 0,749481 | 0,453567 | 0,559131 | 213,4312877 | 0,052248871 | 0,254209 | 0,205535 | 0,837154 | 0,871158 | 301,2973 | 0,273863901 | 0,303579 | 0,902117 | 0,366995 | 0,473457 | 254,4963 | 0,314822 | 0,234506 | 1,342489 | 0,179437 | 0,240891 | 267,0919002 | 0,581645963 | 0,307551 | 1,891216 | 0,058596 | 0,0852 |
| 1701 | RM25_RS08270 | RM25_1681 | Transcriptional regulator | 1774 | gene | 1893984 | 1894403 | - | 249,5172 | -0,47871825 | 0,142952 | -3,34881 | 0,000812 | 0,008775 | 266,3457389 | -0,295018686 | 0,200758 | -1,46953 | 0,14169 | 0,194263 | 232,6888 | -0,684482696 | 0,182707 | -3,74634 | 0,000179 | 0,001212 | 306,8602 | -0,097 | 0,176383 | -0,54994 | 0,58236 | 0,644424 | 204,9547052 | -0,430997862 | 0,222468 | -1,93734 | 0,052703 | 0,0773 |
| 1702 | RM25_RS08275 | RM25_1682 | Lactoylglutathione lyase and lyase | 1775 | gene | 1894482 | 1894850 | + | 199,0451 | -1,041069518 | 0,573528 | -1,8152 | 0,069493 | 0,153079 | 107,7969179 | -2,30673426 | 0,35092 | -6,57339 | 4,92E-11 | 5,11E-10 | 290,2933 | -0,673299451 | 0,32519 | -2,07048 | 0,038407 | 0,083743 | 280,6683 | 0,9217 | 0,339326 | 2,716267 | 0,006602 | 0,016679 | 127,7933559 | 2,621671275 | 0,347321 | 7,548264 | 4,41E-14 | 4,73E-13 |
| 1703 | NA | RM25_1683 | Hypothetical protein | 1776 | gene | 1894900 | 1895163 | - | 45,77231 | -1,678864715 | 0,720218 | -2,33105 | 0,019751 | 0,07121 | 25,34958452 | -3,937859715 | 0,7172 | -5,4906 | 4,01E-08 | 2,62E-07 | 66,19504 | -1,190871972 | 0,40753 | -2,92217 | 0,003476 | 0,013609 | 73,38423 | 0,871864 | 0,308319 | 2,827798 | 0,004687 | 0,012511 | 21,38846912 | 3,691991819 | 0,84586 | 4,364778 | 1,27E-05 | 3,68E-05 |
| 1704 | RM25_RS12330 | RM25_1684 | SEFIR domain protein | 1777 | gene | 1895543 | 1896496 | + | 1572,323 | -0,891719937 | 0,341452 | -2,61155 | 0,009013 | 0,043534 | 1046,176543 | -0,982156117 | 0,171911 | -5,71315 | 1,11E-08 | 8,07E-08 | 2098,469 | -0,848621889 | 0,259687 | -3,26786 | 0,001084 | 0,005191 | 2147,195 | 0,897053 | 0,254814 | 3,520427 | 0,000431 | 0,001745 | 1076,109469 | 1,078023673 | 0,169386 | 6,364293 | 1,96E-10 | 1,21E-09 |
| 1705 | NA | RM25_1685 | Hypothetical protein | 1778 | gene | 1897593 | 1897802 | - | 91,89275 | -1,640440281 | 0,614019 | -2,67164 | 0,007548 | 0,038785 | 44,52686043 | -2,578318687 | 0,512963 | -5,02632 | 5E-07 | 2,64E-06 | 139,2586 | -1,406672532 | 0,341184 | -4,12291 | 3,74E-05 | 0,000316 | 145,7966 | 1,330675 | 0,357111 | 3,726228 | 0,000194 | 0,000876 | 43,78895773 | 2,572017093 | 0,538412 | 4,777046 | 1,78E-06 | 5,91E-06 |
| 1706 | RM25_RS08295 | RM25_1686 | Hypothetical protein | 1779 | gene | 1897850 | 1898629 | - | 225,838 | -1,414676917 | 0,480237 | -2,94579 | 0,003221 | 0,021755 | 151,951294 | -2,330317034 | 0,346316 | -6,72888 | 1,71E-11 | 1,92E-10 | 299,7247 | -1,045385799 | 0,38355 | -2,72555 | 0,006419 | 0,021865 | 344,9773 | 0,603124 | 0,381189 | 1,582219 | 0,1136 | 0,164936 | 121,1414544 | 1,957510717 | 0,384209 | 5,09491 | 3,49E-07 | 1,31E-06 |
| 1707 | RM25_RS08300 | RM25_1687 | Putative response regulator | 1780 | gene | 1898792 | 1899451 | - | 308,9737 | -0,218420392 | 0,15782 | -1,38398 | 0,166364 | 0,268944 | 282,083633 | -0,317103527 | 0,197709 | -1,60389 | 0,108738 | 0,153012 | 335,8637 | -0,1393833 | 0,234383 | -0,59468 | 0,552056 | 0,648652 | 349,7111 | 0,110623 | 0,227525 | 0,486202 | 0,626824 | 0,684626 | 280,0830141 | 0,342321911 | 0,181821 | 1,882739 | 0,059736 | 0,086698 |
| 1708 | RM25_RS08305 | RM25_1688 | Sensor kinase-Membrane protein | 1781 | gene | 1899448 | 1900635 | - | 548,8998 | -0,565740071 | 0,211071 | -2,68033 | 0,007355 | 0,038683 | 457,3125111 | -0,542093982 | 0,173225 | -3,12943 | 0,001751 | 0,004368 | 640,487 | -0,584935438 | 0,269788 | -2,16813 | 0,030149 | 0,069798 | 689,2405 | 0,43971 | 0,23108 | 1,902852 | 0,05706 | 0,094645 | 434,4368615 | 0,456063941 | 0,223988 | 2,036104 | 0,04174 | 0,063024 |
| 1709 | RM25_RS08310 | RM25_1689 | Putative membrane protein | 1782 | gene | 1900786 | 1901349 | + | 194,186 | -0,852064057 | 0,421623 | -2,02092 | 0,043289 | 0,113818 | 127,2308623 | -1,406549063 | 0,356101 | -3,94986 | 7,82E-05 | 0,000273 | 261,1411 | -0,609487764 | 0,296772 | -2,05373 | 0,040002 | 0,08587 | 262,1783 | 0,703095 | 0,354972 | 1,980703 | 0,047625 | 0,081829 | 135,9141727 | 1,565292793 | 0,315743 | 4,957483 | 7,14E-07 | 2,58E-06 |
| 1710 | RM25_RS08315 | RM25_1690 | Hypothetical protein | 1783 | gene | 1901396 | 1901947 | + | 136,5817 | -1,606882349 | 0,739566 | -2,17274 | 0,0298 | 0,089819 | 73,05336234 | -3,81378833 | 0,545392 | -6,99274 | 2,7E-12 | 3,33E-11 | 200,11 | -1,136283041 | 0,479993 | -2,36729 | 0,017919 | 0,047383 | 215,5078 | 0,941678 | 0,432847 | 2,175544 | 0,029589 | 0,055834 | 66,66579078 | 3,693095028 | 0,632805 | 5,836074 | 5,34E-09 | 2,64E-08 |
| 1711 | RM25_RS08325 | RM25_1691 | Heat shock protein transcriptional repressor HspR1 (Hspr1 p | 1784 | gene | 1902848 | 1903267 | - | 849,3658 | 0,131114612 | 0,194837 | 0,672945 | 0,500982 | 0,598791 | 866,4733133 | -0,248953511 | 0,176744 | -1,40856 | 0,158966 | 0,213617 | 832,2584 | 0,530823613 | 0,265462 | 1,999621 | 0,045541 | 0,094325 | 859,988 | -0,52348 | 0,249929 | -2,09452 | 0,036214 | 0,066014 | 868,6856163 | 0,301411745 | 0,18616 | 1,619103 | 0,105425 | 0,144722 |
| 1712 | RM25_RS08330 | RM25_1692 | Chaperone protein DnaJ | 1785 | gene | 1903269 | 1904366 | - | 2231,294 | 0,16971235 | 0,26526 | 0,639797 | 0,522305 | 0,619252 | 2405,508705 | -0,191644586 | 0,169297 | -1,132 | 0,257633 | 0,322658 | 2057,08 | 0,60048062 | 0,398607 | 1,506448 | 0,131952 | 0,213167 | 2227,79 | -0,702 | 0,358583 | -1,9577 | 0,050265 | 0,085356 | 2307,551284 | 0,125143361 | 0,222176 | 0,563263 | 0,573256 | 0,634052 |
| 1713 | RM25_RS08335 | RM25_1693 | Protein GrpE | 1786 | gene | 1904363 | 1904932 | - | 1643,183 | -0,00239853 | 0,207018 | -0,01159 | 0,990756 | 0,994977 | 1666,194788 | -0,271737782 | 0,149188 | -1,82144 | 0,06854 | 0,102701 | 1620,171 | 0,274700617 | 0,349082 | 0,786923 | 0,431327 | 0,536484 | 1741,587 | -0,36721 | 0,306233 | -1,19913 | 0,230477 | 0,298152 | 1604,101351 | 0,216571741 | 0,203766 | 1,062847 | 0,287851 | 0,348467 |
| 1714 | RM25_RS08340 | RM25_1694 | Chaperone protein DnaK | 1787 | gene | 1905093 | 1906922 | - | 10098,83 | 0,3602558 | 0,342297 | 1,052466 | 0,292586 | 0,402348 | 9096,884611 | 0,190416666 | 0,208589 | 0,912878 | 0,361307 | 0,429667 | 11100,78 | 0,501408472 | 0,557821 | 0,898869 | 0,368722 | 0,474647 | 9363,525 | 0,070323 | 0,533862 | 0,131726 | 0,895201 | 0,916191 | 11077,92437 | 0,405427459 | 0,262741 | 1,54307 | 0,122814 | 0,165729 |
| 1715 | RM25_RS08345 | RM25_1695 | Putative SERINE PROTEASE PEPD | 1788 | gene | 1907302 | 1908393 | - | 966,6613 | 0,181854929 | 0,199436 | 0,911847 | 0,361849 | 0,473552 | 988,4663163 | -0,286800799 | 0,145298 | -1,97388 | 0,048396 | 0,076917 | 944,8564 | 0,681545439 | 0,23556 | 2,893296 | 0,003812 | 0,014539 | 959,5031 | -0,64277 | 0,194287 | -3,30836 | 0,000938 | 0,003336 | 1004,778025 | 0,374337013 | 0,150135 | 2,493336 | 0,012655 | 0,02163 |
| 1716 | RM25_RS08355 | RM25_1696 | Hypothetical protein | 1789 | gene | 1908456 | 1908977 | - | 660,7756 | -0,331042088 | 0,209883 | -1,57727 | 0,114734 | 0,209472 | 577,6483203 | -0,684616269 | 0,166789 | -4,10469 | 4,05E-05 | 0,000152 | 743,903 | -0,067209467 | 0,251372 | -0,26737 | 0,789184 | 0,847043 | 776,2504 | 0,029909 | 0,242174 | 0,123503 | 0,901709 | 0,922051 | 572,9396015 | 0,703681052 | 0,148129 | 4,750465 | 2,03E-06 | 6,67E-06 |
| 1717 | RM25_RS08360 | RM25_1697 | Oxygen regulatory protein NreC | 1790 | gene | 1909200 | 1909859 | - | 638,1367 | -0,439379715 | 0,263333 | -1,66853 | 0,09521 | 0,187165 | 516,2621419 | -0,83466107 | 0,210798 | -3,95952 | 7,51E-05 | 0,000264 | 760,0112 | -0,184106689 | 0,248897 | -0,73969 | 0,459488 | 0,563189 | 773,0568 | 0,223269 | 0,271633 | 0,821948 | 0,411106 | 0,487294 | 530,2129205 | 0,929946716 | 0,1543 | 6,026862 | 1,67E-09 | 9,02E-09 |
| 1718 | RM25_RS08365 | RM25_1698 | Sensor kinase | 1791 | gene | 1909862 | 1910812 | - | 789,1416 | -0,791743605 | 0,331714 | -2,38683 | 0,016995 | 0,065132 | 576,0471009 | -1,168557765 | 0,234193 | -4,98972 | 6,05E-07 | 3,15E-06 | 1002,236 | -0,589416686 | 0,283093 | -2,08206 | 0,037337 | 0,082016 | 1050,579 | 0,527581 | 0,31563 | 1,671518 | 0,094619 | 0,141869 | 566,4238822 | 1,167757863 | 0,184932 | 6,314533 | 2,71E-10 | 1,62E-09 |
| 1719 | RM25_RS08370 | RM25_1699 | Putative insertion element ISCmi2 transposase | 1792 | gene | 1910885 | 1911928 | + | 28,15601 | -0,229845678 | 0,307603 | -0,74722 | 0,454933 | 0,560227 | 28,5589112 | -0,856848178 | 0,457573 | -1,87259 | 0,061125 | 0,09271 | 27,75311 | 0,364730381 | 0,375819 | 0,970494 | 0,331801 | 0,437294 | 32,43679 | -0,6803 | 0,333947 | -2,03715 | 0,041635 | 0,073267 | 25,20629063 | 0,600027648 | 0,523339 | 1,146538 | 0,251573 | 0,3098 |
| 1720 | RM25_RS08375 | RM25_1700 | Hypothetical protein | 1793 | gene | 1912008 | 1912775 | - | 800,0626 | 0,013475364 | 0,192206 | 0,070109 | 0,944107 | 0,962067 | 911,4561406 | -0,083761546 | 0,172086 | -0,48674 | 0,626441 | 0,683892 | 688,6691 | 0,14283595 | 0,247563 | 0,576968 | 0,563961 | 0,65924 | 843,5298 | -0,57604 | 0,190492 | -3,02394 | 0,002495 | 0,007608 | 786,7890687 | -0,305381167 | 0,203056 | -1,50393 | 0,132601 | 0,177378 |
| 1721 | RM25_RS08380 | RM25_1701 | ATP-dependent chaperone protein ClpB | 1794 | gene | 1912998 | 1915724 | + | 7839,472 | 0,302328247 | 0,359864 | 0,840118 | 0,400842 | 0,511802 | 8455,012545 | -0,265384545 | 0,180629 | -1,46922 | 0,141772 | 0,194263 | 7223,932 | 1,002131654 | 0,512453 | 1,955557 | 0,050517 | 0,101595 | 7471,409 | -0,98852 | 0,493111 | -2,00465 | 0,045 | 0,07822 | 8458,492293 | 0,311355156 | 0,223546 | 1,392802 | 0,16368 | 0,214568 |
| 1722 | RM25_RS08385 | RM25_1702 | Hypothetical protein | 1795 | gene | 1915793 | 1916527 | - | 2936,724 | 0,357040319 | 0,253502 | 1,40843 | 0,159004 | 0,261712 | 3284,399826 | 0,337310957 | 0,175467 | 1,922363 | 0,05456 | 0,084939 | 2589,048 | 0,382932386 | 0,407413 | 0,939912 | 0,347263 | 0,453462 | 2733,803 | -0,41767 | 0,357128 | -1,16951 | 0,242197 | 0,310753 | 3223,481586 | -0,341738825 | 0,253483 | -1,34817 | 0,177602 | 0,229224 |
| 1723 | RM25_RS08390 | RM25_1703 | Thioredoxin | 1796 | gene | 1916632 | 1917057 | - | 689,9671 | 0,433069476 | 0,186492 | 2,322189 | 0,020223 | 0,072329 | 683,9113993 | 0,271564244 | 0,179175 | 1,515641 | 0,12961 | 0,179595 | 696,0228 | 0,594548932 | 0,324557 | 1,831878 | 0,06697 | 0,126887 | 622,4373 | -0,21465 | 0,283272 | -0,75776 | 0,448596 | 0,522918 | 774,2937858 | 0,146639391 | 0,226977 | 0,646053 | 0,518245 | 0,579461 |
| 1724 | RM25_RS08395 | RM25_1704 | Transposase IS204/IS1001/IS1096/IS1165 family protein | 1797 | gene | 1917738 | 1918991 | + | 53,47378 | -1,356651961 | 0,841778 | -1,61165 | 0,107038 | 0,200978 | 27,79409783 | -4,801817757 | 0,827702 | -5,80139 | 6,58E-09 | 4,95E-08 | 79,15347 | -0,788434357 | 0,419906 | -1,87764 | 0,06043 | 0,116749 | 80,75152 | 0,825304 | 0,374891 | 2,201449 | 0,027704 | 0,052959 | 29,55772748 | 4,914360428 | 0,887382 | 5,538041 | 3,06E-08 | 1,37E-07 |
| 1725 | RM25_RS08400 | RM25_1705 | Hypothetical protein | 1798 | gene | 1919160 | 1919531 | - | 260,5069 | -0,758622582 | 0,372031 | -2,03914 | 0,041436 | 0,110481 | 178,3726656 | -1,253349525 | 0,25012 | -5,01099 | 5,42E-07 | 2,85E-06 | 342,6411 | -0,523102246 | 0,287386 | -1,82021 | 0,068728 | 0,129039 | 343,9458 | 0,61847 | 0,24718 | 2,502101 | 0,012346 | 0,028007 | 190,045936 | 1,413606978 | 0,300025 | 4,711625 | 2,46E-06 | 7,99E-06 |
| 1726 | RM25_RS12995 | RM25_1706 | Hypothetical protein | 1799 | gene | 1919728 | 1921893 | + | 558,1935 | -1,691353471 | 0,884958 | -1,91122 | 0,055976 | 0,132999 | 286,41582 | -5,407236197 | 0,469854 | -11,5083 | 1,2E-30 | 1,41E-28 | 829,9711 | -1,154226022 | 0,429509 | -2,68732 | 0,007203 | 0,023911 | 893,2026 | 0,96102 | 0,437725 | 2,195489 | 0,028129 | 0,053553 | 260,6820846 | 5,292008117 | 0,503062 | 10,5196 | 7,02E-26 | 3,06E-24 |
| 1727 | RM25_RS08410 | RM25_1707 | Hypothetical protein | 1800 | gene | 1921897 | 1923366 | + | 658,6279 | -1,590584844 | 0,706389 | -2,25171 | 0,02434 | 0,079665 | 356,1767469 | -3,697307147 | 0,370079 | -9,9906 | 1,68E-23 | 9,19E-22 | 961,079 | -1,120785603 | 0,419814 | -2,66972 | 0,007591 | 0,02492 | 1036,002 | 0,926663 | 0,38712 | 2,393737 | 0,016678 | 0,035478 | 324,1228018 | 3,575272774 | 0,432858 | 8,259692 | 1,46E-16 | 2,18E-15 |
| 1728 | RM25_RS08415 | RM25_1708 | Hypothetical protein | 1801 | gene | 1923370 | 1925064 | - | 6458,837 | 0,250402969 | 0,306316 | 0,817467 | 0,413661 | 0,523349 | 8153,857648 | 0,407426518 | 0,189573 | 2,149177 | 0,03162 | 0,053618 | 4763,817 | -0,014929065 | 0,332347 | -0,04492 | 0,964171 | 0,979146 | 6260,945 | -0,59837 | 0,319096 | -1,87519 | 0,060766 | 0,099394 | 6873,671271 | -0,987745591 | 0,197875 | -4,99176 | 5,98E-07 | 2,18E-06 |
| 1729 | RM25_RS08420 | RM25_1709 | Hypothetical protein | 1802 | gene | 1925201 | 1926028 | + | 160,3705 | -1,434602111 | 0,654373 | -2,19233 | 0,028356 | 0,086721 | 88,34210535 | -3,008797209 | 0,437913 | -6,87076 | 6,39E-12 | 7,56E-11 | 232,3988 | -1,019010428 | 0,471626 | -2,16063 | 0,030724 | 0,070719 | 245,6063 | 0,913556 | 0,330198 | 2,766693 | 0,005663 | 0,014667 | 85,49534824 | 2,983290503 | 0,597264 | 4,994924 | 5,89E-07 | 2,15E-06 |
| 1730 | RM25_RS08425 | RM25_1710 | Type IV secretory pathway VirB4 protein-like protein | 1803 | gene | 1926025 | 1927524 | + | 381,6698 | -1,303378454 | 0,769203 | -1,69445 | 0,090179 | 0,180895 | 185,2007578 | -3,673908098 | 0,442667 | -8,29948 | 1,05E-16 | 2,12E-15 | 578,1389 | -0,847418222 | 0,493801 | -1,71611 | 0,086141 | 0,15335 | 568,5395 | 1,044356 | 0,429674 | 2,430578 | 0,015075 | 0,032875 | 217,7011096 | 3,94646436 | 0,531493 | 7,425239 | 1,13E-13 | 1,12E-12 |
| 1731 | RM25_RS08430 | RM25_1711 | Putative MmyB family DNA-binding protein | 1804 | gene | 1927512 | 1928213 | - | 404,5789 | -0,962117577 | 0,306381 | -3,14026 | 0,001688 | 0,013579 | 293,0238767 | -1,043536526 | 0,238834 | -4,3693 | 1,25E-05 | 5,15E-05 | 516,134 | -0,919833919 | 0,267752 | -3,43539 | 0,000592 | 0,003203 | 561,6023 | 0,709235 | 0,260573 | 2,721834 | 0,006492 | 0,016454 | 269,569039 | 0,896698923 | 0,265615 | 3,375934 | 0,000736 | 0,001613 |
| 1732 | RM25_RS08435 | RM25_1712 | 2, 5-didehydrogluconate reductase | 1805 | gene | 1928540 | 1929385 | + | 472,1761 | -1,22748078 | 0,479131 | -2,56189 | 0,01041 | 0,048394 | 296,0738439 | -2,041834255 | 0,330381 | -6,18025 | 6,4E-10 | 5,63E-09 | 648,2784 | -0,915949485 | 0,307651 | -2,97723 | 0,002909 | 0,011739 | 694,1597 | 0,762705 | 0,373156 | 2,04393 | 0,04096 | 0,072263 | 277,7387112 | 1,955898979 | 0,289861 | 6,747725 | 1,5E-11 | 1,09E-10 |
| 1733 | RM25_RS08440 | RM25_1713 | Hypothetical protein | 1806 | gene | 1929610 | 1930161 | + | 162,2639 | -1,517843879 | 0,646514 | -2,34773 | 0,018888 | 0,069561 | 86,71678027 | -2,818568303 | 0,517038 | -5,45137 | 5E-08 | 3,22E-07 | 237,8111 | -1,172239704 | 0,449067 | -2,61039 | 0,009044 | 0,028138 | 252,0669 | 1,04366 | 0,467184 | 2,23394 | 0,025487 | 0,049483 | 82,83453197 | 2,766295426 | 0,550796 | 5,022357 | 5,1E-07 | 1,88E-06 |
| 1734 | RM25_RS08445 | RM25_1714 | Type IV secretory pathway, VirD4 component | 1807 | gene | 1930158 | 1931936 | + | 970,2694 | -1,642022577 | 0,92079 | -1,78328 | 0,074541 | 0,160745 | 459,4035958 | -5,634568282 | 0,428921 | -13,1366 | 2,03E-39 | 7,98E-37 | 1481,135 | -1,142260706 | 0,439169 | -2,60096 | 0,009296 | 0,028642 | 1538,03 | 1,105877 | 0,437864 | 2,525615 | 0,01155 | 0,026455 | 465,2400315 | 5,67793494 | 0,462916 | 12,26557 | 1,39E-34 | 1,09E-32 |
| 1735 | RM25_RS08450 | RM25_1715 | Single-strand binding protein | 1808 | gene | 1932021 | 1932461 | + | 156,0222 | -1,525241206 | 0,979971 | -1,55641 | 0,119609 | 0,215865 | 69,50460699 | -5,164467113 | 0,692874 | -7,45369 | 9,08E-14 | 1,38E-12 | 242,5399 | -1,060004328 | 0,464659 | -2,28125 | 0,022533 | 0,056322 | 242,3927 | 1,197134 | 0,413899 | 2,892334 | 0,003824 | 0,010679 | 79,51686959 | 5,395801209 | 0,792955 | 6,804672 | 1,01E-11 | 7,51E-11 |
| 1736 | RM25_RS08455 | RM25_1716 | Hypothetical protein | 1809 | gene | 1932481 | 1933038 | + | 135,2773 | -1,517805676 | 0,937044 | -1,61978 | 0,10528 | 0,199312 | 67,90134914 | -5,40209257 | 0,732507 | -7,3748 | 1,65E-13 | 2,41E-12 | 202,6532 | -0,973287311 | 0,530207 | -1,83567 | 0,066406 | 0,12592 | 210,4849 | 0,940308 | 0,450003 | 2,089559 | 0,036657 | 0,066412 | 69,33718824 | 5,446448312 | 0,84835 | 6,420048 | 1,36E-10 | 8,63E-10 |
| 1737 | RM25_RS08460 | RM25_1717 | Hypothetical protein | 1810 | gene | 1933043 | 1933270 | + | 75,82553 | -1,775388696 | 0,859786 | -2,06492 | 0,038931 | 0,107196 | 39,51675091 | -5,048870254 | 0,802112 | -6,29447 | 3,08E-10 | 2,77E-09 | 112,1343 | -1,257810164 | 0,528938 | -2,37799 | 0,017407 | 0,046327 | 123,1263 | 0,963033 | 0,459979 | 2,093644 | 0,036292 | 0,066092 | 33,85690553 | 4,837648435 | 0,909379 | 5,319725 | 1,04E-07 | 4,24E-07 |
| 1738 | NA | RM25_1718 | | 1811 | gene | 1933453 | 1935207 | + | 1047,647 | -1,784406729 | 0,727288 | -2,45351 | 0,014147 | 0,058499 | 554,8833552 | -3,993990698 | 0,379803 | -10,516 | 7,29E-26 | 4,78E-24 | 1540,411 | -1,329130766 | 0,387484 | -3,43016 | 0,000603 | 0,003239 | 1700,029 | 1,008765 | 0,434459 | 2,321887 | 0,020239 | 0,041385 | 464,8657496 | 3,74677908 | 0,370558 | 10,11119 | 4,93E-24 | 1,66E-22 |
| 1740 | RM25_RS08480 | RM25_1719 | Hypothetical protein | 1815 | gene | 1935255 | 1936058 | - | 4168,636 | 0,498808861 | 0,362271 | 1,376894 | 0,168545 | 0,271168 | 5393,769788 | 0,392530459 | 0,217775 | 1,802458 | 0,071473 | 0,106085 | 2943,501 | 0,698116089 | 0,407822 | 1,711817 | 0,08693 | 0,154521 | 3678,471 | -1,10269 | 0,383014 | -2,87899 | 0,00399 | 0,010985 | 4776,125375 | -0,767848807 | 0,249275 | -3,08032 | 0,002068 | 0,004106 |
| 1741 | RM25_RS08485 | RM25_1720 | Ribosomal-protein-alanine acetyltransferase rimJ | 1816 | gene | 1936133 | 1936843 | - | 393,1483 | -0,032868926 | 0,21649 | -0,15183 | 0,879324 | 0,918284 | 328,7523135 | 0,048773202 | 0,20292 | 0,240357 | 0,810054 | 0,849709 | 457,5443 | -0,091893133 | 0,267559 | -0,34345 | 0,73126 | 0,800789 | 417,2189 | 0,485832 | 0,216843 | 2,240483 | 0,02506 | 0,048814 | 381,6161951 | 0,403761385 | 0,242501 | 1,664991 | 0,095915 | 0,132749 |
| 1742 | RM25_RS08490 | RM25_1721 | Molybdopterin biosynthesis protein moeA1 | 1817 | gene | 1936847 | 1938133 | - | 1132,044 | 0,31448647 | 0,123667 | 2,543001 | 0,010991 | 0,049721 | 1071,827786 | 0,493231638 | 0,142503 | 3,46119 | 0,000538 | 0,001544 | 1192,261 | 0,157513112 | 0,189927 | 0,829336 | 0,406914 | 0,512338 | 1064,146 | 0,281496 | 0,179701 | 1,56647 | 0,117239 | 0,169114 | 1228,984139 | -0,006211999 | 0,13613 | -0,04563 | 0,963603 | 0,97063 |
| 1743 | RM25_RS08495 | RM25_1722 | 5-formyltetrahydrofolate cyclo-ligase family protein | 1818 | gene | 1938148 | 1938861 | + | 724,3478 | 0,066758572 | 0,197779 | 0,33754 | 0,73571 | 0,806918 | 598,043438 | 0,035219895 | 0,183511 | 0,191923 | 0,847803 | 0,880684 | 850,6521 | 0,090393365 | 0,176304 | 0,512714 | 0,608151 | 0,695832 | 744,7993 | 0,427438 | 0,193738 | 2,206268 | 0,027365 | 0,052439 | 726,448255 | 0,530598977 | 0,161621 | 3,282981 | 0,001027 | 0,002171 |
| 1744 | RM25_RS08500 | RM25_1723 | Type I antifreeze protein | 1819 | gene | 1939017 | 1939424 | + | 853,6501 | 0,431241505 | 0,315143 | 1,3684 | 0,171187 | 0,274855 | 1065,953658 | 0,140900127 | 0,202592 | 0,695487 | 0,48675 | 0,553969 | 641,3466 | 0,93659937 | 0,26259 | 3,566779 | 0,000361 | 0,002167 | 774,6795 | -1,26797 | 0,250299 | -5,06584 | 4,07E-07 | 3,31E-06 | 959,096846 | -0,423309528 | 0,19671 | -2,15195 | 0,031401 | 0,049211 |
| 1745 | RM25_RS11960 | RM25_1724 | Hypothetical protein | 1820 | gene | 1939617 | 1940165 | + | 137,4689 | -1,268012948 | 0,547075 | -2,3178 | 0,02046 | 0,072518 | 79,29440762 | -2,250757723 | 0,413546 | -5,44259 | 5,25E-08 | 3,36E-07 | 195,6435 | -0,950304725 | 0,388873 | -2,44374 | 0,014536 | 0,040307 | 203,6749 | 0,902714 | 0,368934 | 2,446819 | 0,014412 | 0,031589 | 79,28523062 | 2,274724932 | 0,463202 | 4,910866 | 9,07E-07 | 3,2E-06 |
| 1746 | RM25_RS08510 | RM25_1725 | Large conductance mechanosensitive channel | 1821 | gene | 1940258 | 1940701 | + | 474,7211 | -0,33501791 | 0,306952 | -1,09143 | 0,275082 | 0,384158 | 351,1964046 | -0,781512287 | 0,216756 | -3,6055 | 0,000312 | 0,000954 | 598,2457 | -0,084652317 | 0,230622 | -0,36706 | 0,713573 | 0,786666 | 557,4115 | 0,411209 | 0,211696 | 1,942447 | 0,052083 | 0,087874 | 409,9245682 | 1,153378898 | 0,213338 | 5,406357 | 6,43E-08 | 2,75E-07 |
| 1747 | RM25_RS08515 | RM25_1726 | Hypothetical protein | 1822 | gene | 1940755 | 1941165 | - | 235,0407 | -0,213191979 | 0,141379 | -1,50795 | 0,131567 | 0,230047 | 237,8298347 | 0,030398036 | 0,222698 | 0,136499 | 0,891427 | 0,917909 | 232,2515 | -0,45773628 | 0,174656 | -2,62079 | 0,008773 | 0,02768 | 265,6142 | 0,132155 | 0,171218 | 0,77185 | 0,440203 | 0,515429 | 213,7729828 | -0,304004023 | 0,223696 | -1,35901 | 0,174145 | 0,226148 |
| 1748 | RM25_RS08520 | RM25_1727 | UTP--glucose-1-phosphate uridylyltransferase | 1823 | gene | 1941158 | 1942531 | - | 2481,173 | -0,182313697 | 0,250316 | -0,72834 | 0,466408 | 0,569008 | 2361,478381 | 0,316146374 | 0,170065 | 1,858979 | 0,06303 | 0,095111 | 2600,869 | -0,642709272 | 0,35714 | -1,7996 | 0,071924 | 0,133484 | 2781,533 | 0,543529 | 0,329159 | 1,651263 | 0,098685 | 0,147125 | 2274,659155 | -0,382513441 | 0,208026 | -1,83877 | 0,065948 | 0,094896 |
| 1750 | RM25_RS08530 | RM25_1729 | Hypothetical protein | 1827 | gene | 1943106 | 1943438 | + | 601,2757 | 0,159068123 | 0,172341 | 0,922987 | 0,356014 | 0,467134 | 591,7170481 | 0,497668756 | 0,166185 | 2,994672 | 0,002747 | 0,006502 | 610,8344 | -0,160553269 | 0,252078 | -0,63692 | 0,524178 | 0,623669 | 598,7967 | 0,34116 | 0,228726 | 1,491567 | 0,135813 | 0,190769 | 621,4547264 | -0,27471468 | 0,181741 | -1,51158 | 0,130642 | 0,175155 |
| 1751 | RM25_RS08535 | RM25_1730 | Isochorismatase hydrolase | 1828 | gene | 1943583 | 1944203 | - | 1905,439 | 1,979409106 | 0,642117 | 3,082628 | 0,002052 | 0,0156 | 570,2763312 | 0,189463711 | 0,168315 | 1,125652 | 0,260313 | 0,325619 | 3240,602 | 2,439544579 | 0,433875 | 5,622691 | 1,88E-08 | 3,85E-07 | 813,0028 | 0,864472 | 0,388595 | 2,224607 | 0,026108 | 0,050398 | 2952,525865 | 3,151532607 | 0,251904 | 12,51083 | 6,51E-36 | 5,48E-34 |
| 1752 | RM25_RS08540 | RM25_1731 | Esterase/lipase/thioesterase family protein | 1829 | gene | 1944200 | 1945009 | - | 968,6125 | 0,791803756 | 0,583802 | 1,356288 | 0,175007 | 0,278711 | 406,6330511 | -0,595460363 | 0,18682 | -3,18735 | 0,001436 | 0,003659 | 1530,592 | 1,196847091 | 0,411118 | 2,911204 | 0,0036 | 0,014024 | 745,3181 | 0,867388 | 0,279858 | 3,099383 | 0,001939 | 0,006127 | 1190,052324 | 2,692841808 | 0,320278 | 8,407839 | 4,18E-17 | 6,65E-16 |
| 1753 | RM25_RS08545 | RM25_1732 | Hypothetical protein | 1830 | gene | 1945162 | 1945521 | + | 3343,472 | 2,813489007 | 0,591872 | 4,753545 | 2E-06 | 7,36E-05 | 1633,661829 | 0,602344217 | 0,205622 | 2,929381 | 0,003396 | 0,007856 | 5053,282 | 4,728353689 | 0,33302 | 14,1984 | 9,37E-46 | 1,1E-42 | 889,7733 | -1,87547 | 0,251204 | -7,46593 | 8,27E-14 | 2,57E-12 | 5687,821794 | 2,286005492 | 0,265114 | 8,622729 | 6,54E-18 | 1,17E-16 |
| 1754 | RM25_RS08550 | RM25_1733 | Thioredoxin | 1831 | gene | 1945659 | 1945994 | - | 6629,66 | -1,406993846 | 0,312408 | -4,50371 | 6,68E-06 | 0,000202 | 6176,798329 | -1,708505397 | 0,270235 | -6,32229 | 2,58E-10 | 2,35E-09 | 7082,522 | -1,165701888 | 0,451155 | -2,58382 | 0,009771 | 0,029743 | 10221,12 | 0,000856 | 0,483842 | 0,001768 | 0,998589 | 0,998589 | 3548,187318 | 0,579593188 | 0,227837 | 2,543896 | 0,010962 | 0,019027 |
| 1755 | RM25_RS08555 | RM25_1734 | Iron-dependent repressor and activator IdeR | 1832 | gene | 1946110 | 1946787 | - | 5273,122 | -0,274237247 | 0,307761 | -0,89107 | 0,37289 | 0,484511 | 5117,574777 | -0,763038463 | 0,277456 | -2,75012 | 0,005957 | 0,012798 | 5428,668 | 0,171998143 | 0,417017 | 0,412449 | 0,68001 | 0,75746 | 6141,844 | -0,38772 | 0,449883 | -0,86183 | 0,388783 | 0,464687 | 4659,398387 | 0,582946758 | 0,240564 | 2,423247 | 0,015382 | 0,025897 |
| 1756 | RM25_RS08560 | RM25_1735 | Phosphoserine transaminase | 1833 | gene | 1946964 | 1948085 | + | 706,0064 | -0,526934086 | 0,182045 | -2,89453 | 0,003797 | 0,024521 | 627,9788996 | -0,707062226 | 0,139141 | -5,08161 | 3,74E-07 | 2,01E-06 | 784,0339 | -0,388525976 | 0,266903 | -1,45568 | 0,145481 | 0,229824 | 878,3809 | 0,125431 | 0,214125 | 0,585785 | 0,55802 | 0,623343 | 567,7461463 | 0,506190982 | 0,207532 | 2,439093 | 0,014724 | 0,024842 |
| 1757 | RM25_RS08565 | RM25_1736 | Hypothetical protein | 1834 | gene | 1948128 | 1948514 | - | 213,4096 | 0,661495641 | 0,203681 | 3,247702 | 0,001163 | 0,011057 | 229,1049768 | 0,587840611 | 0,31165 | 1,886221 | 0,059265 | 0,090658 | 197,7142 | 0,743040035 | 0,217206 | 3,420906 | 0,000624 | 0,003321 | 174,5893 | -0,35701 | 0,303687 | -1,17558 | 0,239761 | 0,30847 | 256,1401365 | -0,154823513 | 0,240324 | -0,64423 | 0,519427 | 0,580325 |
| 1758 | RM25_RS08570 | RM25_1737 | Putative hypoxanthine/guanine permease | 1835 | gene | 1948676 | 1950238 | - | 1766,24 | 0,283522299 | 0,303233 | 0,934998 | 0,349789 | 0,462397 | 2146,362359 | -0,132365175 | 0,179675 | -0,73669 | 0,46131 | 0,529619 | 1386,117 | 0,959163435 | 0,224509 | 4,272266 | 1,93E-05 | 0,000182 | 1697,538 | -1,31448 | 0,232761 | -5,64733 | 1,63E-08 | 1,76E-07 | 1897,876789 | -0,174935955 | 0,146096 | -1,19741 | 0,231148 | 0,287957 |
| 1759 | RM25_RS08575 | RM25_1738 | Hypothetical protein | 1836 | gene | 1950284 | 1950625 | + | 451,926 | 0,045327263 | 0,231995 | 0,19538 | 0,845095 | 0,898057 | 428,1988394 | 0,067472407 | 0,233695 | 0,28872 | 0,772795 | 0,81754 | 475,6531 | 0,025522329 | 0,379665 | 0,067223 | 0,946404 | 0,966078 | 471,6232 | 0,118768 | 0,361401 | 0,328633 | 0,742433 | 0,78754 | 448,6896463 | 0,116501072 | 0,273386 | 0,426141 | 0,670005 | 0,721756 |
| 1760 | RM25_RS08580 | RM25_1739 | Hypothetical protein | 1837 | gene | 1950703 | 1951482 | - | 2397,001 | 0,805583122 | 0,399267 | 2,017653 | 0,043627 | 0,114094 | 3176,241749 | 0,58791538 | 0,215165 | 2,732387 | 0,006288 | 0,013369 | 1617,76 | 1,265322795 | 0,385299 | 3,284003 | 0,001023 | 0,004943 | 1860,755 | -1,46636 | 0,396269 | -3,70042 | 0,000215 | 0,00095 | 2986,990378 | -0,752938354 | 0,188417 | -3,99612 | 6,44E-05 | 0,000169 |
| 1761 | RM25_RS12620 | RM25_1740 | Cold shock protein B cspB | 1838 | gene | 1951486 | 1951872 | - | 1517,217 | 0,722683716 | 0,353025 | 2,047121 | 0,040646 | 0,109562 | 1813,637486 | 0,423933349 | 0,2391 | 1,77304 | 0,076222 | 0,112355 | 1220,797 | 1,200492751 | 0,443337 | 2,707855 | 0,006772 | 0,0229 | 1218,497 | -1,1092 | 0,447862 | -2,47666 | 0,013262 | 0,029798 | 1847,800099 | -0,304729298 | 0,225948 | -1,34867 | 0,177442 | 0,229168 |
| 1762 | RM25_RS08590 | RM25_1741 | Hypothetical protein | 1839 | gene | 1952099 | 1953127 | + | 454,6188 | -0,46210047 | 0,393119 | -1,17547 | 0,239806 | 0,347402 | 291,4840971 | -0,93950809 | 0,248731 | -3,77721 | 0,000159 | 0,000516 | 617,7536 | -0,250386388 | 0,267473 | -0,93612 | 0,349213 | 0,455503 | 552,5697 | 0,741029 | 0,28893 | 2,564734 | 0,010326 | 0,024313 | 375,084304 | 1,493190679 | 0,225983 | 6,607547 | 3,91E-11 | 2,67E-10 |
| 1763 | RM25_RS08595 | RM25_1742 | DeaD/DeaH box family protein | 1840 | gene | 1953124 | 1955445 | + | 1009,183 | 0,154126459 | 0,223874 | 0,68845 | 0,491169 | 0,590355 | 826,3543336 | 0,103872594 | 0,155663 | 0,667291 | 0,504586 | 0,569866 | 1192,012 | 0,188545874 | 0,281519 | 0,669745 | 0,503021 | 0,602755 | 1002,814 | 0,424313 | 0,205378 | 2,06601 | 0,038828 | 0,069215 | 1044,689892 | 0,568666194 | 0,229078 | 2,482422 | 0,013049 | 0,022224 |
| 1764 | RM25_RS08600 | RM25_1743 | Putative DNA HELICASE ERCC3 | 1841 | gene | 1955518 | 1957197 | + | 1146,902 | 0,234615161 | 0,121501 | 1,93098 | 0,053485 | 0,129697 | 1076,655928 | 0,291546967 | 0,151974 | 1,9184 | 0,05506 | 0,085509 | 1217,147 | 0,184640779 | 0,20149 | 0,916376 | 0,359469 | 0,466301 | 1110,167 | 0,169203 | 0,209821 | 0,806418 | 0,420002 | 0,495468 | 1215,621681 | 0,118658942 | 0,113886 | 1,041906 | 0,297455 | 0,35862 |
| 1765 | RM25_RS13000 | RM25_1744 | S-layer domain protein | 1842 | gene | 1957360 | 1958259 | - | 1772,341 | 0,576053425 | 0,411258 | 1,400711 | 0,1613 | 0,264385 | 2368,182312 | 0,506009802 | 0,189092 | 2,675996 | 0,007451 | 0,015405 | 1176,5 | 0,721303212 | 0,491072 | 1,468834 | 0,141878 | 0,22595 | 1515,449 | -1,19077 | 0,452451 | -2,63183 | 0,008493 | 0,020658 | 2076,418029 | -0,942437367 | 0,260947 | -3,61161 | 0,000304 | 0,000711 |

| | | | | | | | | | |
|---|---|---|---|---|---|---|---|---|---|
| 1766 | RM25_RS11970 | RM25_1745 | Hypothetical protein | 1843 | gene | 1958284 | 1959237 | - | [illegible] |
| 1767 | RM25_RS11975 | RM25_1746 | Surface layer protein A (S-layer protein A) | 1844 | gene | 1959635 | 1961017 | - | [illegible] |
| 1768 | RM25_RS11980 | RM25_1747 | Surface layer protein A (S-layer protein A) | 1845 | gene | 1961666 | 1963201 | - | [illegible] |
| 1769 | RM25_RS08625 | RM25_1748 | IS407A, transposase OrfB | 1846 | gene | 1963198 | 1964013 | - | [illegible] |
| 1770 | NA | RM25_1749 | Transposase IS3/IS911 | 1847 | gene | 1964064 | 1964348 | - | [illegible] |
| 1771 | NA | RM25_1750 | Hypothetical protein | 1848 | gene | 1964411 | 1964596 | - | [illegible] |
| 1772 | RM25_RS08635 | RM25_1751 | MutT/NUDIX family protein | 1849 | gene | 1965146 | 1965706 | - | [illegible] |
| 1773 | RM25_RS08640 | RM25_1752 | 5S ribosomal RNA | 1851 | rRNA | 1965947 | 1966062 | - | [illegible] |
| 1774 | RM25_RS08645 | RM25_1753 | 23S ribosomal RNA | 1854 | rRNA | 1966207 | 1969332 | - | [illegible] |
| 1775 | NA | RM25_1754 | Hypothetical protein | 1856 | gene | 1968924 | 1969919 | + | [illegible] |
| 1776 | RM25_RS08650 | RM25_1755 | 16S ribosomal RNA | 1858 | rRNA | 1969679 | 1971185 | - | [illegible] |
| 1777 | RM25_RS08655 | RM25_1756 | 60 kDa chaperonin 1 | 1860 | gene | 1972490 | 1974112 | - | [illegible] |
| 1778 | RM25_RS08660 | RM25_1757 | Hypothetical protein | 1861 | gene | 1974456 | 1974737 | - | [illegible] |
| 1779 | RM25_RS08665 | RM25_1758 | Mannose-6-phosphate isomerase, class I | 1862 | gene | 1974956 | 1976155 | + | [illegible] |
| 1781 | RM25_RS08675 | RM25_1760 | Conserved transmembrane transport protein MmpL13 | 1866 | gene | 1976644 | 1978974 | + | [illegible] |
| 1782 | RM25_RS08680 | RM25_1761 | Ktn nad-binding domain involved with k+ transport | 1867 | gene | 1979095 | 1981236 | - | [illegible] |
| 1783 | RM25_RS08685 | RM25_1762 | Hypothetical protein | 1868 | gene | 1981243 | 1981626 | - | [illegible] |
| 1784 | RM25_RS08690 | RM25_1763 | Putative RNase III regulator YmdB | 1869 | gene | 1981711 | 1982220 | + | [illegible] |
| 1785 | RM25_RS08695 | RM25_1764 | SNARE-like domain protein | 1870 | gene | 1982245 | 1983258 | - | [illegible] |
| 1786 | RM25_RS08700 | RM25_1765 | Amidohydrolase | 1871 | gene | 1983328 | 1984602 | - | [illegible] |
| 1787 | RM25_RS08705 | RM25_1766 | L-serine dehydratase 1 | 1872 | gene | 1984778 | 1986169 | + | [illegible] |
| 1788 | RM25_RS08710 | RM25_1767 | ATP binding protein of ABC transporter for sugars | 1873 | gene | 1986385 | 1987494 | + | [illegible] |
| 1789 | RM25_RS08715 | RM25_1768 | Cystathionine gamma-synthase | 1874 | gene | 1987780 | 1988967 | + | [illegible] |
| 1790 | RM25_RS08720 | RM25_1769 | Lipopolysaccharide kinase, Kdo/WaaP family | 1875 | gene | 1989081 | 1990463 | + | [illegible] |
| 1791 | RM25_RS08725 | RM25_1770 | Hypothetical protein | 1876 | gene | 1990520 | 1990921 | - | [illegible] |
| 1792 | RM25_RS08730 | RM25_1771 | Protein of hypothetical function DUF162 | 1877 | gene | 1991163 | 1991819 | - | [illegible] |
| 1793 | RM25_RS08735 | RM25_1772 | Iron-sulfur cluster-binding protein | 1878 | gene | 1991825 | 1993363 | - | [illegible] |
| 1794 | RM25_RS08740 | RM25_1773 | Lactate utilization protein A | 1879 | gene | 1993360 | 1994169 | - | [illegible] |
| 1795 | RM25_RS08745 | RM25_1774 | L-lactate permease | 1880 | gene | 1994210 | 1995913 | - | [illegible] |
| 1796 | RM25_RS08750 | RM25_1775 | FCD domain protein | 1881 | gene | 1996418 | 1997419 | + | [illegible] |
| 1797 | RM25_RS08755 | RM25_1776 | Cysteine--tRNA ligase | 1882 | gene | 1997546 | 1998949 | + | [illegible] |
| 1798 | RM25_RS08760 | RM25_1777 | RNA methyltransferase, TrmH family, group 3 | 1883 | gene | 1998949 | 1999872 | + | [illegible] |
| 1799 | RM25_RS08765 | RM25_1778 | L-asparagine permease | 1884 | gene | 2000085 | 2001587 | + | [illegible] |
| 1800 | RM25_RS08770 | RM25_1779 | L-asparagine permease | 1885 | gene | 2001906 | 2003396 | + | [illegible] |
| 1801 | RM25_RS08775 | RM25_1780 | Cytoplasmic asparaginase I | 1886 | gene | 2003407 | 2004408 | + | [illegible] |
| 1802 | RM25_RS08780 | RM25_1781 | Glycine hydroxymethyltransferase | 1887 | gene | 2004628 | 2006076 | + | [illegible] |
| 1803 | RM25_RS08785 | RM25_1782 | 2-C-methyl-D-erythritol 2, 4-cyclodiphosphate synthase | 1888 | gene | 2006281 | 2006781 | - | [illegible] |
| 1804 | RM25_RS08790 | RM25_1783 | 2-C-methyl-D-erythritol 4-phosphate cytidylyltransferase | 1889 | gene | 2006778 | 2007542 | - | [illegible] |
| 1805 | RM25_RS08795 | RM25_1784 | Transcriptional regulator, CarD family | 1890 | gene | 2007606 | 2008091 | - | [illegible] |
| 1806 | RM25_RS08800 | RM25_1785 | Transposase IS204/IS1001/IS1096/IS1165 family protein | 1891 | gene | 2008280 | 2009356 | - | [illegible] |
| 1807 | RM25_RS08805 | RM25_1786 | Sensory transduction protein RegX3 | 1892 | gene | 2009700 | 2010356 | - | [illegible] |
| 1808 | RM25_RS08810 | RM25_1787 | Two-component histidine kinase SenX3 | 1893 | gene | 2010362 | 2011549 | - | [illegible] |
| 1809 | RM25_RS08815 | RM25_1788 | Phosphate-transport system transcriptional regulator phoY2 | 1894 | gene | 2011844 | 2012521 | + | [illegible] |
| 1810 | RM25_RS08820 | RM25_1789 | 2, 3-bisphosphoglycerate-dependent phosphoglycerate mut | 1895 | gene | 2012721 | 2013470 | - | [illegible] |
| 1811 | RM25_RS11985 | RM25_1790 | Putative transcription regulator TetR/AcrR family | 1896 | gene | 2013801 | 2014532 | + | [illegible] |
| 1812 | RM25_RS08830 | RM25_1791 | Short-chain dehydrogenase/reductase yusR | 1897 | gene | 2014647 | 2015366 | + | [illegible] |
| 1813 | RM25_RS08835 | RM25_1792 | Biotin-requiring enzyme | 1898 | gene | 2015497 | 2015868 | - | [illegible] |
| 1814 | RM25_RS08840 | RM25_1793 | Hypothetical protein | 1899 | gene | 2015885 | 2016133 | - | [illegible] |
| 1815 | RM25_RS08845 | RM25_1794 | Carboxyl transferase | 1900 | gene | 2016149 | 2017723 | - | [illegible] |
| 1816 | RM25_RS08850 | RM25_1795 | Oxaloacetate decarboxylase alpha subunit | 1901 | gene | 2017739 | 2019256 | - | [illegible] |
| 1817 | RM25_RS08855 | RM25_1796 | Glutamate-1-semialdehyde 2, 1-aminomutase 2 | 1902 | gene | 2019525 | 2020850 | - | [illegible] |
| 1818 | RM25_RS08860 | RM25_1797 | Transcriptional regulator, TetR family | 1903 | gene | 2020908 | 2021687 | - | [illegible] |
| 1819 | RM25_RS08865 | RM25_1798 | Membrane-bound transporter | 1904 | gene | 2021739 | 2023199 | + | [illegible] |
| 1820 | RM25_RS08870 | RM25_1799 | Porphobilinogen synthase | 1905 | gene | 2023228 | 2024226 | - | [illegible] |
| 1821 | RM25_RS08875 | RM25_1800 | Ferrochelatase | 1906 | gene | 2024349 | 2025407 | - | [illegible] |
| 1822 | RM25_RS08880 | RM25_1801 | Protoporphyrinogen oxidase | 1907 | gene | 2025404 | 2026987 | - | [illegible] |
| 1823 | RM25_RS08885 | RM25_1802 | Coenzyme PQQ synthesis protein | 1908 | gene | 2026991 | 2028112 | - | [illegible] |
| 1824 | RM25_RS12355 | RM25_1803 | Uroporphyrinogen decarboxylase | 1909 | gene | 2028114 | 2030012 | - | [illegible] |
| 1825 | RM25_RS08895 | RM25_1804 | Hydroxymethylbilane synthase | 1910 | gene | 2030009 | 2030941 | - | [illegible] |
| 1826 | RM25_RS08900 | RM25_1805 | Glutamyl-tRNA reductase | 1911 | gene | 2030938 | 2032191 | - | [illegible] |
| 1827 | RM25_RS08905 | RM25_1806 | Aspartate--ammonia ligase | 1912 | gene | 2032400 | 2034310 | + | [illegible] |
| 1828 | RM25_RS08910 | RM25_1807 | Dihydroxy-acid dehydratase | 1913 | gene | 2034443 | 2036284 | + | [illegible] |
| 1829 | RM25_RS08915 | RM25_1808 | Lysine exporter protein | 1914 | gene | 2036612 | 2037220 | - | [illegible] |
| 1830 | RM25_RS08920 | RM25_1809 | Inosose dehydratase | 1915 | gene | 2037467 | 2038390 | - | [illegible] |
| 1831 | RM25_RS08925 | RM25_1810 | Hypothetical protein | 1916 | gene | 2038565 | 2038912 | - | [illegible] |
| 1832 | RM25_RS08930 | RM25_1811 | Inositol 2-dehydrogenase | 1917 | gene | 2039056 | 2040045 | - | [illegible] |
| 1833 | RM25_RS08935 | RM25_1812 | Transaldolase | 1918 | gene | 2040359 | 2041435 | - | [illegible] |
| 1834 | RM25_RS08940 | RM25_1813 | UbiC transcription regulator-associated | 1919 | gene | 2041629 | 2042381 | - | [illegible] |
| 1835 | RM25_RS08945 | RM25_1814 | Putative 5-dehydro-2-deoxygluconokinase | 1920 | gene | 2042655 | 2043644 | + | [illegible] |
| 1836 | RM25_RS08950 | RM25_1815 | Deoxyribose-phosphate aldolase | 1921 | gene | 2043647 | 2044516 | + | [illegible] |
| 1837 | RM25_RS08955 | RM25_1816 | Myo-inositol catabolism protein | 1922 | gene | 2044513 | 2045385 | + | [illegible] |
| 1838 | RM25_RS08960 | RM25_1817 | 3D-(3, 5/4)-trihydroxycyclohexane-1, 2-dione hydrolase | 1923 | gene | 2045485 | 2047392 | + | [illegible] |
| 1839 | RM25_RS08965 | RM25_1818 | Methylmalonate-semialdehyde dehydrogenase (acylating) | 1924 | gene | 2047658 | 2049163 | + | [illegible] |
| 1840 | RM25_RS08970 | RM25_1819 | MFS transporter, sugar porter family protein | 1925 | gene | 2049501 | 2051099 | + | [illegible] |
| 1841 | RM25_RS08975 | RM25_1820 | Hypothetical protein | 1926 | gene | 2051282 | 2052574 | + | [illegible] |
| 1842 | RM25_RS08980 | RM25_1821 | Dehydrogenase | 1927 | gene | 2052700 | 2053848 | + | [illegible] |
| 1843 | NA | RM25_1822 | Hypothetical protein | 1928 | gene | 2054020 | 2054187 | + | [illegible] |
| 1844 | RM25_RS08985 | RM25_1823 | Hypothetical protein | 1929 | gene | 2054148 | 2054423 | - | [illegible] |
| 1845 | RM25_RS08990 | RM25_1824 | Fructose-bisphosphate aldolase, class II | 1930 | gene | 2054677 | 2055699 | - | [illegible] |
| 1846 | RM25_RS08995 | RM25_1825 | Hypothetical protein | 1931 | gene | 2055928 | 2056554 | - | [illegible] |
| 1847 | RM25_RS09000 | RM25_1826 | RNA methyltransferase, TrmH family | 1932 | gene | 2056560 | 2057252 | - | [illegible] |
| 1848 | RM25_RS09005 | RM25_1827 | SNARE-like domain protein | 1933 | gene | 2057328 | 2058071 | + | [illegible] |
| 1849 | RM25_RS09010 | RM25_1828 | LemA family protein | 1934 | gene | 2058279 | 2059148 | - | [illegible] |
| 1850 | RM25_RS09015 | RM25_1829 | Orotate phosphoribosyltransferase | 1935 | gene | 2059418 | 2059975 | - | [illegible] |
| 1851 | RM25_RS09020 | RM25_1830 | Peptidase C60 family protein | 1936 | gene | 2060055 | 2060663 | - | [illegible] |
| 1852 | RM25_RS09025 | RM25_1831 | Spermidine synthase | 1937 | gene | 2060867 | 2061784 | - | [illegible] |
| 1853 | RM25_RS09030 | RM25_1832 | Oligopeptide transporter, OPT family | 1938 | gene | 2061999 | 2063990 | + | [illegible] |
| 1854 | RM25_RS09035 | RM25_1833 | ATP-dependent chaperone protein ClpB | 1939 | gene | 2064111 | 2066711 | - | [illegible] |
| 1855 | RM25_RS09040 | RM25_1834 | Glutamate-1-semialdehyde 2, 1-aminomutase 2 | 1940 | gene | 2067069 | 2068436 | - | [illegible] |
| 1856 | RM25_RS09045 | RM25_1835 | Amino acid permease-associated region membrane protein | 1941 | gene | 2068578 | 2070233 | - | [illegible] |
| 1857 | RM25_RS09050 | RM25_1836 | Adenylosuccinate synthase | 1942 | gene | 2070768 | 2072051 | - | [illegible] |
| 1858 | RM25_RS09055 | RM25_1837 | Hypothetical protein | 1943 | gene | 2072248 | 2072463 | - | [illegible] |
| 1859 | RM25_RS09060 | RM25_1838 | Putative aldehyde dehydrogenase AldA | 1944 | gene | 2072658 | 2074154 | - | [illegible] |
| 1860 | RM25_RS09065 | RM25_1839 | Ribonucleoside hydrolase 1 | 1945 | gene | 2074573 | 2075586 | + | [illegible] |
| 1861 | RM25_RS09070 | RM25_1840 | putative ACR | 1946 | gene | 2075710 | 2076390 | - | [illegible] |
| 1862 | RM25_RS09075 | RM25_1841 | Methylated-DNA--[protein]-cysteine S-methyltransferase | 1947 | gene | 2076668 | 2077222 | + | [illegible] |
| 1863 | RM25_RS09080 | RM25_1842 | Methylated-DNA--[protein]-cysteine S-methyltransferase | 1948 | gene | 2077219 | 2077854 | + | [illegible] |
| 1864 | RM25_RS09085 | RM25_1843 | Multicopper oxidase | 1949 | gene | 2078061 | 2079581 | + | [illegible] |
| 1865 | NA | RM25_1844 | IS3 family element | 1950 | gene | 2080007 | 2080351 | + | [illegible] |
| 1866 | RM25_RS09095 | RM25_1845 | Transposase IS6110 | 1951 | gene | 2080384 | 2081271 | + | [illegible] |
| 1867 | RM25_RS09100 | RM25_1846 | Hypothetical protein | 1952 | gene | 2081321 | 2081500 | + | [illegible] |
| 1868 | RM25_RS09105 | RM25_1847 | Putative polysaccharide syntase | 1953 | gene | 2081497 | 2083032 | + | [illegible] |
| 1870 | RM25_RS09115 | RM25_1848 | Ser/Thr protein phosphatase family protein | 1957 | gene | 2083374 | 2084345 | - | [illegible] |
| 1871 | RM25_RS09120 | RM25_1849 | Glycosyl transferase family 51 | 1958 | gene | 2084342 | 2086549 | - | [illegible] |
| 1872 | RM25_RS12360 | RM25_1850 | Transcription factor WhiB | 1959 | gene | 2086743 | 2087126 | + | [illegible] |
| 1873 | RM25_RS09130 | RM25_1851 | Hypothetical protein | 1960 | gene | 2087116 | 2087307 | + | [illegible] |
| 1874 | RM25_RS09135 | RM25_1852 | Endoribonuclease L-PSP | 1961 | gene | 2087304 | 2087759 | + | [illegible] |
| 1875 | RM25_RS09140 | RM25_1853 | Endonuclease III | 1962 | gene | 2087785 | 2088519 | + | [illegible] |
| 1876 | RM25_RS09145 | RM25_1854 | NTP pyrophosphohydrolase | 1963 | gene | 2088605 | 2089303 | + | [illegible] |
| 1877 | RM25_RS09150 | RM25_1855 | S4 domain protein | 1964 | gene | 2089342 | 2089695 | + | [illegible] |
| 1878 | RM25_RS09155 | RM25_1856 | Phosphoenolpyruvate-protein phosphotransferase | 1965 | gene | 2089799 | 2091544 | - | [illegible] |
| 1879 | RM25_RS09160 | RM25_1857 | Phosphocarrier, HPr family | 1966 | gene | 2091783 | 2092049 | + | [illegible] |
| 1880 | RM25_RS09165 | RM25_1858 | Putative cation-transporting P-type ATPase | 1967 | gene | 2092163 | 2094808 | + | [illegible] |
| 1881 | RM25_RS09170 | RM25_1859 | Mannosyltransferase PIG-V domain protein | 1968 | gene | 2094940 | 2096157 | + | [illegible] |
| 1882 | RM25_RS09175 | RM25_1860 | Hypothetical protein | 1969 | gene | 2096196 | 2097332 | - | [illegible] |
| 1883 | RM25_RS11990 | RM25_1861 | Hypothetical protein | 1970 | gene | 2097329 | 2097808 | - | [illegible] |
| 1884 | RM25_RS09180 | RM25_1862 | Transposase IS204/IS1001/IS1096/IS1165 family protein | 1971 | gene | 2098286 | 2099251 | + | [illegible] |
| 1885 | NA | RM25_1863 | Hypothetical protein | 1972 | gene | 2099456 | 2099806 | - | [illegible] |
| 1886 | RM25_RS09190 | RM25_1864 | Glycoside hydrolase family 2 sugar binding protein | 1973 | gene | 2099855 | 2101762 | + | [illegible] |
| 1887 | RM25_RS09195 | RM25_1865 | Thioredoxin | 1974 | gene | 2101900 | 2102328 | + | [illegible] |
| 1888 | RM25_RS09200 | RM25_1866 | FHA domain protein | 1975 | gene | 2102499 | 2103215 | + | [illegible] |
| 1889 | RM25_RS09205 | RM25_1867 | FHA domain protein | 1976 | gene | 2103218 | 2103691 | + | [illegible] |
| 1890 | NA | RM25_1868 | Putative SERINE/THREONINE PHOSPHATASE PPP | 1977 | gene | 2103694 | 2105229 | + | [illegible] |
| 1891 | RM25_RS09215 | RM25_1869 | Cell division protein RodA | 1978 | gene | 2105226 | 2106614 | + | [illegible] |
| 1892 | RM25_RS09220 | RM25_1870 | Penicillin-binding protein pbpA | 1979 | gene | 2106611 | 2108035 | + | [illegible] |
| 1893 | RM25_RS09225 | RM25_1871 | Non-specific serine/threonine protein kinase | 1980 | gene | 2108127 | 2109911 | + | [illegible] |
| 1894 | RM25_RS11995 | RM25_1872 | Hypothetical protein | 1981 | gene | 2110036 | 2111808 | + | [illegible] |
| 1895 | RM25_RS09235 | RM25_1873 | Galactokinase | 1982 | gene | 2112025 | 2113173 | - | [illegible] |
| 1896 | RM25_RS09240 | RM25_1874 | NifU-like protein | 1983 | gene | 2113363 | 2114010 | - | [illegible] |
| 1897 | RM25_RS09245 | RM25_1875 | Ferrous iron transport protein B | 1984 | gene | 2114137 | 2116101 | - | [illegible] |
| 1898 | RM25_RS09250 | RM25_1876 | Ferrous iron transport protein A | 1985 | gene | 2116280 | 2116639 | - | [illegible] |
| 1899 | RM25_RS09255 | RM25_1877 | Hypothetical protein | 1986 | gene | 2117003 | 2117602 | + | [illegible] |
| 1900 | RM25_RS09260 | RM25_1878 | Potassium/proton antiporter | 1987 | gene | 2117684 | 2119264 | - | [illegible] |
| 1901 | RM25_RS12000 | RM25_1879 | Hypothetical protein | 1988 | gene | 2119280 | 2121184 | - | [illegible] |
| 1903 | RM25_RS09275 | RM25_1880 | Deaminase | 1992 | gene | 2121660 | 2122091 | - | [illegible] |
| 1904 | RM25_RS09280 | RM25_1881 | Putative tRNA adenosine deaminase-associated protein | 1993 | gene | 2122163 | 2122891 | - | [illegible] |
| 1905 | RM25_RS09285 | RM25_1882 | Uracil phosphoribosyltransferase | 1994 | gene | 2123080 | 2123721 | + | [illegible] |
| 1906 | RM25_RS09290 | RM25_1883 | Putative fimbrial associated sortase-like protein | 1995 | gene | 2123664 | 2124590 | - | [illegible] |
| 1907 | RM25_RS09295 | RM25_1884 | Hypothetical protein | 1996 | gene | 2124787 | 2125089 | - | [illegible] |
| 1908 | NA | RM25_1885 | Hypothetical protein | 1997 | gene | 2125996 | 2126268 | - | [illegible] |
| 1909 | RM25_RS09305 | RM25_1886 | Putative surface-anchored fimbrial subunit | 1998 | gene | 2126361 | 2127500 | - | [illegible] |
| 1910 | RM25_RS09310 | RM25_1887 | Surface-anchored fimbrial subunit | 1999 | gene | 2127497 | 2130385 | - | [illegible] |
| 1911 | RM25_RS09315 | RM25_1888 | Transposase insI for insertion sequence element IS30B/C/D | 2000 | gene | 2131210 | 2132244 | + | [illegible] |
| 1912 | RM25_RS09320 | RM25_1889 | GTP-binding translation elongation factor typA | 2001 | gene | 2132537 | 2134420 | + | [illegible] |

| | | | | | | | | | | | | | | | | | | | | | | | | | | | | | | | | | | | | | | |
|---|---|---|---|---|---|---|---|---|---|---|---|---|---|---|---|---|---|---|---|---|---|---|---|---|---|---|---|---|---|---|---|---|---|---|---|---|---|---|
| 1913 | RM25_RS09325 | RM25_1890 | Folate-binding protein YgfZ | 2002 | gene | 2134510 | 2135559 | - | 1124,441 | 0,458690407 | 0,212484 | 2,158703 | 0,030873 | 0,092431 | 1349,75791 | 0,476420707 | 0,168703 | 2,824024 | 0,004742 | 0,010379 | 899,1231 | 0,434346569 | 0,15304 | 2,838133 | 0,004538 | 0,016592 | 1003,395 | -0,61839 | 0,178994 | -3,45481 | 0,000551 | 0,002135 | 1276,84504 | -0,610086717 | 0,135846 | -4,49102 | 7,09E-06 | 2,13E-05 |
| 1914 | RM25_RS09330 | RM25_1891 | UPF0678 fatty acid-binding protein-like protein | 2003 | gene | 2135735 | 2136238 | - | 791,0558 | 0,777416529 | 0,236747 | 3,283747 | 0,001024 | 0,010145 | 895,5167678 | 0,727190342 | 0,21903 | 3,320053 | 0,0009 | 0,002443 | 686,5949 | 0,845634483 | 0,334092 | 2,531143 | 0,011369 | 0,033496 | 618,2773 | -0,5077 | 0,336496 | -1,50879 | 0,131352 | 0,185833 | 977,432143 | -0,354588833 | 0,20563 | -1,7244 | 0,084635 | 0,118459 |
| 1915 | RM25_RS09335 | RM25_1892 | D-tyrosyl-tRNA(Tyr) deacylase | 2004 | gene | 2136551 | 2136982 | + | 650,3418 | 0,04551564 | 0,227537 | 0,200036 | 0,841452 | 0,89531 | 516,3904586 | 0,117260172 | 0,135014 | 0,868501 | 0,38512 | 0,454091 | 784,2932 | -0,001573336 | 0,247589 | -0,00635 | 0,99493 | 0,996621 | 674,4334 | 0,606408 | 0,225689 | 2,686912 | 0,007212 | 0,018006 | 646,0272343 | 0,53179127 | 0,157582 | 3,3747 | 0,000739 | 0,001619 |
| 1916 | RM25_RS09340 | RM25_1893 | Transcriptional regulatory protein GlnR | 2005 | gene | 2137147 | 2137806 | + | 1151,593 | -0,249311654 | 0,0938 | -2,65791 | 0,007863 | 0,03994 | 1076,846518 | -0,243457334 | 0,127726 | -1,90609 | 0,056639 | 0,087311 | 1226,34 | -0,255020832 | 0,155919 | -1,6356 | 0,101924 | 0,174885 | 1319,504 | 0,131084 | 0,153806 | 0,852265 | 0,394067 | 0,469811 | 1031,335794 | 0,172182435 | 0,111583 | 1,543089 | 0,122809 | 0,165729 |
| 1917 | RM25_RS09345 | RM25_1894 | Hypothetical protein | 2006 | gene | 2137982 | 2138503 | - | 416,7607 | 0,042176782 | 0,535841 | 0,078711 | 0,937262 | 0,957161 | 220,1915009 | -1,300574967 | 0,239519 | -5,42995 | 5,64E-08 | 3,58E-07 | 613,3298 | 0,499053193 | 0,341148 | 1,462864 | 0,143505 | 0,227771 | 432,2743 | 0,642787 | 0,28826 | 2,229886 | 0,025755 | 0,04984 | 411,1077499 | 2,477165695 | 0,277286 | 8,933608 | 4,12E-19 | 8,76E-18 |
| 1918 | RM25_RS09350 | RM25_1895 | Inositol-3-phosphate synthase | 2007 | gene | 2139115 | 2140194 | + | 929,3134 | -0,897349848 | 0,749242 | -1,19768 | 0,231043 | 0,338451 | 327,8877113 | -2,298710377 | 0,181681 | -12,6524 | 1,08E-36 | 2,56E-34 | 1530,739 | -0,657911065 | 0,589781 | -1,11552 | 0,264629 | 0,366684 | 1270,173 | 1,731149 | 0,567479 | 3,050598 | 0,002284 | 0,007064 | 633,4216803 | 3,409854048 | 0,26758 | 12,74333 | 3,4E-37 | 3,34E-35 |
| 1919 | RM25_RS09355 | RM25_1896 | 3-hydroxyisobutyrate dehydrogenase related beta-hydroxya | 2008 | gene | 2140551 | 2141411 | + | 1003,544 | -0,476460433 | 0,470799 | -1,01203 | 0,311526 | 0,420237 | 522,3929052 | -0,278272066 | 0,147824 | -1,88245 | 0,059775 | 0,091131 | 1484,695 | -0,547999499 | 0,383056 | -1,4306 | 0,152545 | 0,238112 | 1226,293 | 1,570228 | 0,353212 | 4,445572 | 8,77E-06 | 5,47E-05 | 819,479902 | 1,338283236 | 0,217211 | 6,161226 | 7,22E-10 | 4,1E-09 |
| 1920 | RM25_RS09360 | RM25_1897 | putative gamma-carboxymuconolactone decarboxylase subu | 2009 | gene | 2141570 | 2142295 | + | 501,9323 | -0,804176205 | 0,343963 | -2,33797 | 0,019389 | 0,070633 | 339,2486742 | -0,864069813 | 0,260183 | -3,321 | 0,000897 | 0,002438 | 664,6158 | -0,776474943 | 0,255516 | -3,03885 | 0,002375 | 0,009942 | 669,4225 | 0,875184 | 0,279185 | 3,134778 | 0,00172 | 0,00553 | 358,5846943 | 1,02060239 | 0,248965 | 4,099389 | 4,14E-05 | 0,000111 |
| 1921 | RM25_RS09365 | RM25_1898 | UDP-glucose 4-epimerase (UDP-galactose 4-epimerase) (Gal | 2010 | gene | 2142327 | 2143148 | - | 282,2537 | -1,066021202 | 0,474894 | -2,24476 | 0,024784 | 0,080573 | 177,7327835 | -1,85949628 | 0,332206 | -5,59743 | 2,18E-08 | 1,49E-07 | 386,7747 | -0,755764397 | 0,349432 | -2,16284 | 0,030554 | 0,070396 | 401,398 | 0,731186 | 0,328926 | 2,222953 | 0,026219 | 0,050571 | 179,4285883 | 1,904999594 | 0,380361 | 5,008401 | 5,49E-07 | 2,01E-06 |
| 1922 | NA | RM25_1899 | Hypothetical protein | 2011 | gene | 2143239 | 2143355 | - | 186,2404 | -1,410905331 | 0,644061 | -2,19064 | 0,028478 | 0,086721 | 101,7872826 | -3,019113088 | 0,455392 | -6,6297 | 3,36E-11 | 3,57E-10 | 270,6935 | -0,993802707 | 0,354833 | -2,80076 | 0,005098 | 0,018207 | 283,6447 | 0,916855 | 0,408257 | 2,245777 | 0,024718 | 0,04839 | 100,0228257 | 3,015662879 | 0,441439 | 6,831437 | 8,41E-12 | 6,31E-11 |
| 1923 | RM25_RS09370 | RM25_1900 | Mycothiol acetyltransferase | 2012 | gene | 2143567 | 2144505 | + | 1524,105 | -0,020416201 | 0,209071 | -0,09765 | 0,922209 | 0,948749 | 1284,403781 | 0,058583274 | 0,145287 | 0,403225 | 0,686783 | 0,739492 | 1763,806 | -0,077047389 | 0,284588 | -0,27073 | 0,786596 | 0,845382 | 1617,201 | 0,471725 | 0,26178 | 1,801991 | 0,071547 | 0,113407 | 1479,594646 | 0,376358496 | 0,160878 | 2,339409 | 0,019314 | 0,031835 |
| 1924 | RM25_RS09375 | RM25_1901 | Polyphosphate kinase | 2013 | gene | 2144502 | 2146691 | + | 3525,277 | -0,166952455 | 0,177382 | -0,9412 | 0,346602 | 0,458955 | 3066,169755 | -0,44670148 | 0,117587 | -3,7989 | 0,000145 | 0,000478 | 3984,384 | 0,045530682 | 0,192088 | 0,237031 | 0,812633 | 0,865899 | 3933,938 | 0,089423 | 0,120087 | 0,744655 | 0,45648 | 0,530012 | 3247,30483 | 0,62833105 | 0,141235 | 4,448844 | 8,63E-06 | 2,54E-05 |
| 1925 | RM25_RS09380 | RM25_1902 | Putative HYDROLASE MUTT1 | 2014 | gene | 2146688 | 2147578 | + | 1105,648 | 0,180459831 | 0,167235 | 1,079081 | 0,280552 | 0,389664 | 1195,721575 | -0,094262261 | 0,159912 | -0,58946 | 0,555552 | 0,620291 | 1015,574 | 0,507544879 | 0,191918 | 2,644595 | 0,008179 | 0,026336 | 1097,04 | -0,61484 | 0,177782 | -3,45841 | 0,000543 | 0,002111 | 1150,067227 | 0,034103016 | 0,128025 | 0,266377 | 0,789949 | 0,829728 |
| 1926 | RM25_RS09385 | RM25_1903 | Phosphate ABC transporter, phosphate-binding protein Pst! | 2015 | gene | 2147766 | 2148860 | + | 406,3426 | -1,073666286 | 0,4161 | -2,58031 | 0,009871 | 0,046626 | 264,6555318 | -1,542909982 | 0,255902 | -6,02931 | 1,65E-09 | 1,37E-08 | 548,0297 | -0,868872961 | 0,358116 | -2,42623 | 0,015256 | 0,041765 | 578,0577 | 0,778453 | 0,282248 | 2,758044 | 0,005815 | 0,014995 | 257,733031 | 1,522303092 | 0,352829 | 4,314558 | 1,6E-05 | 4,59E-05 |
| 1927 | RM25_RS09390 | RM25_1904 | Phosphate ABC transporter, permease protein PstC | 2016 | gene | 2148857 | 2149837 | + | 312,304 | -1,244302183 | 0,526841 | -2,36182 | 0,018186 | 0,068145 | 186,1993924 | -2,194543218 | 0,344054 | -6,37849 | 1,79E-10 | 1,69E-09 | 438,4085 | -0,916617566 | 0,389593 | -2,35275 | 0,018635 | 0,048853 | 460,6062 | 0,837184 | 0,359616 | 2,327991 | 0,019913 | 0,040812 | 182,6844751 | 2,187862916 | 0,404254 | 5,412101 | 6,23E-08 | 2,67E-07 |
| 1928 | RM25_RS09395 | RM25_1905 | Phosphate-transport membrane ABC transporter pstA1 | 2017 | gene | 2149834 | 2150994 | + | 309,8865 | -1,359122117 | 0,545328 | -2,4923 | 0,012692 | 0,05499 | 179,4836086 | -2,317575413 | 0,37719 | -6,14432 | 8,03E-10 | 6,93E-09 | 440,2894 | -1,045238008 | 0,376557 | -2,77578 | 0,005507 | 0,019315 | 467,0842 | 0,916909 | 0,403937 | 2,269931 | 0,023212 | 0,046083 | 171,1605813 | 2,262761096 | 0,382732 | 5,912128 | 3,38E-09 | 1,73E-08 |
| 1929 | RM25_RS09400 | RM25_1906 | Phosphate-transport ATP-binding protein ABC transporter p | 2018 | gene | 2150999 | 2151778 | + | 374,3707 | -1,147953216 | 0,494263 | -2,32255 | 0,020203 | 0,072329 | 226,3155217 | -2,041119022 | 0,323722 | -6,30517 | 2,88E-10 | 2,62E-09 | 522,4258 | -0,826530308 | 0,290504 | -2,84516 | 0,004439 | 0,01642 | 541,4154 | 0,806848 | 0,342574 | 2,355249 | 0,01851 | 0,038516 | 228,5649956 | 2,087005468 | 0,297469 | 7,015879 | 2,29E-12 | 1,87E-11 |
| 1930 | RM25_RS09405 | RM25_1907 | Putative heavy metal reductase | 2019 | gene | 2151779 | 2152210 | - | 179,0759 | -0,523479455 | 0,222013 | -2,35788 | 0,01838 | 0,068517 | 151,175448 | -0,426823283 | 0,276972 | -1,54103 | 0,123309 | 0,171671 | 206,9763 | -0,597256421 | 0,255149 | -2,34081 | 0,019242 | 0,050003 | 222,7462 | 0,458271 | 0,275857 | 1,66126 | 0,096661 | 0,144654 | 144,3824625 | 0,349198527 | 0,292083 | 1,195546 | 0,231874 | 0,288709 |
| 1931 | RM25_RS09410 | RM25_1908 | Transcriptional regulator ArsR | 2020 | gene | 2152210 | 2152584 | - | 264,9324 | -0,350252128 | 0,159392 | -2,19743 | 0,02799 | 0,086672 | 237,9847939 | -0,108129899 | 0,204415 | -0,52897 | 0,596825 | 0,657344 | 291,88 | -0,548920221 | 0,189561 | -2,89574 | 0,003783 | 0,014479 | 312,6519 | 0,431694 | 0,179318 | 2,407415 | 0,016066 | 0,034645 | 228,5687238 | 0,041472729 | 0,221048 | 0,187619 | 0,851176 | 0,87915 |
| 1932 | RM25_RS09415 | RM25_1909 | Arsenical resistance operon trans-acting repressor ArsD | 2021 | gene | 2152681 | 2153148 | + | 189,7999 | -0,796962282 | 0,270418 | -2,94715 | 0,003207 | 0,021722 | 154,3519296 | -0,854931716 | 0,225199 | -3,79634 | 0,000147 | 0,000482 | 225,2478 | -0,76408157 | 0,369985 | -2,06517 | 0,038907 | 0,084287 | 253,8221 | 0,44767 | 0,189002 | 2,368606 | 0,017855 | 0,037409 | 136,4208763 | 0,604848812 | 0,406837 | 1,48671 | 0,137092 | 0,182946 |
| 1933 | RM25_RS09420 | RM25_1910 | Arsenical pump-driving ATPase | 2022 | gene | 2153145 | 2154920 | + | 834,7594 | -1,180074722 | 0,380419 | -3,10204 | 0,001922 | 0,014951 | 585,1500664 | -1,549812177 | 0,263541 | -5,88074 | 4,08E-09 | 3,2E-08 | 1084,369 | -0,996836487 | 0,357172 | -2,79092 | 0,005256 | 0,018685 | 1216,361 | 0,659725 | 0,320223 | 2,060202 | 0,039379 | 0,069892 | 503,3317379 | 1,284112379 | 0,32562 | 3,943587 | 8,03E-05 | 0,000208 |
| 1934 | RM25_RS09425 | RM25_1911 | putative B-hexosaminidase | 2023 | gene | 2155003 | 2156232 | - | 404,6239 | 0,054625998 | 0,302473 | 0,180598 | 0,856683 | 0,905472 | 294,573272 | 0,065508021 | 0,220139 | 0,297576 | 0,766027 | 0,811766 | 514,6746 | 0,047984016 | 0,299458 | 0,160236 | 0,872695 | 0,906941 | 416,265 | 0,760215 | 0,176716 | 4,301894 | 1,69E-05 | 9,86E-05 | 405,5230344 | 0,799165782 | 0,308761 | 2,588297 | 0,009645 | 0,016914 |
| 1935 | RM25_RS09430 | RM25_1912 | Mycothione reductase | 2024 | gene | 2156286 | 2157806 | - | 1731,793 | -0,332862271 | 0,238918 | -1,39321 | 0,163558 | 0,266049 | 1403,718625 | 0,100098033 | 0,13495 | 0,741743 | 0,458243 | 0,526611 | 2059,868 | -0,634575636 | 0,256611 | -2,47291 | 0,013402 | 0,038289 | 2031,322 | 0,831971 | 0,258748 | 3,21537 | 0,001303 | 0,004443 | 1501,744066 | 0,143063798 | 0,124057 | 1,153213 | 0,248823 | 0,306895 |
| 1936 | RM25_RS09435 | RM25_1913 | Phosphoglycerate mutase | 2025 | gene | 2157922 | 2158581 | - | 612,1428 | 0,344331166 | 0,209441 | 1,64405 | 0,100166 | 0,193263 | 698,8113489 | 0,16111431 | 0,157847 | 1,020702 | 0,307396 | 0,375794 | 525,4742 | 0,592274311 | 0,267956 | 2,210342 | 0,027081 | 0,064541 | 571,5349 | -0,71876 | 0,246525 | -2,91555 | 0,003551 | 0,010083 | 671,6517536 | -0,228879229 | 0,162667 | -1,40705 | 0,159414 | 0,209559 |
| 1937 | RM25_RS09440 | RM25_1914 | Uroporphyrin-III C-methyltransferase/uroporphyrinogen-III s | 2026 | gene | 2158636 | 2160252 | - | 3026,565 | 0,247731171 | 0,359783 | 0,688557 | 0,491102 | 0,590355 | 4122,843787 | 0,190455971 | 0,186012 | 1,023892 | 0,305886 | 0,374337 | 1930,287 | 0,371564718 | 0,230913 | 1,609114 | 0,107591 | 0,182179 | 2941,812 | -1,2529 | 0,249551 | -5,02062 | 5,15E-07 | 4,12E-06 | 3221,032409 | -1,025219174 | 0,123064 | -8,3308 | 8,03E-17 | 1,22E-15 |
| 1938 | RM25_RS09445 | RM25_1915 | Pyrroline-5-carboxylate reductase | 2027 | gene | 2160741 | 2161601 | - | 661,9307 | 0,658439536 | 0,154622 | 4,25839 | 2,06E-05 | 0,00049 | 602,4387793 | 0,730922094 | 0,211224 | 3,46042 | 0,000539 | 0,001546 | 721,4226 | 0,599321478 | 0,169617 | 3,533382 | 0,00041 | 0,0024 | 539,7983 | 0,28856 | 0,199552 | 1,446039 | 0,148166 | 0,204586 | 794,4727334 | 0,203747145 | 0,170902 | 1,192189 | 0,233187 | 0,289988 |
| 1939 | RM25_RS09450 | RM25_1916 | Aspartate-semialdehyde dehydrogenase | 2028 | gene | 2161605 | 2162639 | - | 2010,69 | 0,785077759 | 0,164224 | 4,780543 | 1,75E-06 | 6,62E-05 | 1888,31683 | 0,794353801 | 0,170247 | 4,665887 | 3,07E-06 | 1,43E-05 | 2133,064 | 0,77746139 | 0,279017 | 2,78643 | 0,005329 | 0,018833 | 1560,876 | 0,133393 | 0,261378 | 0,510347 | 0,609808 | 0,668909 | 2487,916359 | 0,155283321 | 0,1822 | 0,852267 | 0,394066 | 0,459354 |
| 1940 | RM25_RS09455 | RM25_1917 | Hypothetical membrane protein, DUF1211 family | 2029 | gene | 2162731 | 2163426 | - | 524,6496 | -0,499397722 | 0,2136 | -2,338 | 0,019387 | 0,070633 | 426,0445189 | -0,405350722 | 0,1794 | -2,25949 | 0,023853 | 0,041615 | 623,2548 | -0,566062527 | 0,227294 | -2,49044 | 0,012759 | 0,036853 | 645,8324 | 0,553743 | 0,189672 | 2,919475 | 0,003506 | 0,009996 | 426,7279251 | 0,451139044 | 0,215448 | 2,093962 | 0,036263 | 0,055574 |
| 1941 | RM25_RS09460 | RM25_1918 | CAAX amino terminal protease family protein | 2030 | gene | 2163585 | 2164463 | - | 381,7236 | 0,340752924 | 0,18127 | 1,879804 | 0,060135 | 0,139408 | 374,1818222 | 0,713746753 | 0,201749 | 3,537787 | 0,000403 | 0,001202 | 389,2655 | -0,003706839 | 0,231326 | -0,01602 | 0,987215 | 0,992267 | 353,161 | 0,401925 | 0,222701 | 1,804778 | 0,071109 | 0,113094 | 419,0831561 | -0,262127747 | 0,190542 | -1,3757 | 0,168915 | 0,220817 |
| 1942 | RM25_RS09465 | RM25_1919 | AP endonuclease, family 2 | 2031 | gene | 2164465 | 2165334 | - | 1433,302 | 0,63886129 | 0,130558 | 4,893296 | 9,92E-07 | 4,1E-05 | 1503,560016 | 0,818085573 | 0,142398 | 5,745044 | 9,19E-09 | 6,77E-08 | 1363,044 | 0,448501942 | 0,218678 | 2,050973 | 0,04027 | 0,086287 | 1183,228 | 0,028539 | 0,188175 | 0,151664 | 0,879452 | 0,903998 | 1709,085453 | -0,299091465 | 0,146382 | -2,04323 | 0,04103 | 0,062031 |
| 1943 | RM25_RS09470 | RM25_1920 | Hypothetical protein | 2032 | gene | 2165553 | 2166191 | + | 1649,771 | 0,524745123 | 0,284487 | 1,844533 | 0,065105 | 0,146425 | 2000,279914 | 0,438594509 | 0,220684 | 1,98743 | 0,046875 | 0,075108 | 1299,262 | 0,660570107 | 0,333162 | 1,982727 | 0,047398 | 0,09723 | 1437,644 | -0,80435 | 0,324726 | -2,47702 | 0,013248 | 0,029796 | 1904,333685 | -0,548697766 | 0,224123 | -2,4482 | 0,014357 | 0,024276 |
| 1944 | RM25_RS09475 | RM25_1921 | DNA repair protein radA | 2033 | gene | 2166307 | 2167701 | + | 831,1504 | 0,296023822 | 0,162968 | 1,816456 | 0,0693 | 0,152798 | 748,2892127 | 0,40248871 | 0,16867 | 2,386254 | 0,017021 | 0,031282 | 914,0117 | 0,210699266 | 0,233198 | 0,90352 | 0,36625 | 0,472857 | 785,2064 | 0,337051 | 0,167911 | 2,007318 | 0,044716 | 0,07784 | 899,5720177 | 0,20134394 | 0,219105 | 0,91894 | 0,358127 | 0,422264 |
| 1945 | RM25_RS09480 | RM25_1922 | DNA integrity scanning protein DisA | 2034 | gene | 2167739 | 2168809 | + | 1175,641 | -0,048805355 | 0,117474 | -0,41546 | 0,677809 | 0,757513 | 1146,615151 | 0,228145572 | 0,114446 | 1,993473 | 0,04621 | 0,074433 | 1204,666 | -0,310772698 | 0,166449 | -1,86707 | 0,061891 | 0,11889 | 1261,072 | 0,277897 | 0,161394 | 1,721851 | 0,085096 | 0,129989 | 1133,066729 | -0,211538007 | 0,115288 | -1,83487 | 0,066525 | 0,095552 |
| 1946 | RM25_RS09485 | RM25_1923 | Adenine glycosylase mutY | 2035 | gene | 2168875 | 2169753 | + | 226,039 | -0,427273821 | 0,48346 | -0,88378 | 0,376813 | 0,487981 | 131,132495 | -0,895440724 | 0,315543 | -2,83777 | 0,004543 | 0,010007 | 320,9456 | -0,247434727 | 0,456225 | -0,54235 | 0,587576 | 0,677221 | 270,7883 | 0,968447 | 0,333189 | 2,906599 | 0,003654 | 0,010314 | 190,2502802 | 1,683807141 | 0,445109 | 3,782912 | 0,000155 | 0,000383 |
| 1947 | RM25_RS09490 | RM25_1924 | HMP/thiamine permease protein ykoE | 2036 | gene | 2169985 | 2170620 | + | 359,5654 | -0,863703356 | 0,273127 | -3,16227 | 0,001565 | 0,012946 | 284,8718497 | -1,166186598 | 0,200203 | -5,82501 | 5,71E-09 | 4,34E-08 | 434,2589 | -0,679215427 | 0,282556 | -2,40383 | 0,016224 | 0,043804 | 488,7898 | 0,372848 | 0,22026 | 1,692764 | 0,0905 | 0,136649 | 250,4118735 | 0,925216175 | 0,280396 | 3,299676 | 0,000968 | 0,002063 |
| 1949 | RM25_RS09495 | RM25_1925 | HMP/thiamine import ATP-binding protein | 2037 | gene | 2170646 | 2172316 | + | 450,3359 | -0,643710596 | 0,498927 | -1,29019 | 0,196984 | 0,299693 | 271,8732498 | -1,374713031 | 0,268878 | -5,11278 | 3,17E-07 | 1,73E-06 | 628,7986 | -0,358022519 | 0,492812 | -0,72649 | 0,467539 | 0,570026 | 575,9887 | 0,77958 | 0,291639 | 2,673098 | 0,007515 | 0,018634 | 347,3496351 | 1,872372676 | 0,477198 | 3,92368 | 8,72E-05 | 0,000224 |
| 1950 | RM25_RS09500 | RM25_1926 | Cobalt transport protein CbiQ | 2038 | gene | 2172313 | 2173140 | + | 252,1154 | -0,809541279 | 0,489067 | -1,65528 | 0,097868 | 0,190598 | 147,5008602 | -1,270539843 | 0,238224 | -5,33337 | 9,64E-08 | 5,87E-07 | 356,7299 | -0,634136179 | 0,499629 | -1,26921 | 0,204365 | 0,301431 | 336,2787 | 0,987039 | 0,264763 | 3,728011 | 0,000193 | 0,000876 | 180,8730284 | 1,700919147 | 0,498177 | 3,414289 | 0,000639 | 0,001414 |
| 1951 | RM25_RS09505 | RM25_1927 | Para-aminobenzoate synthase | 2039 | gene | 2173151 | 2173732 | + | 215,9224 | -0,900143482 | 0,483109 | -1,86323 | 0,06243 | 0,142584 | 120,5999049 | -1,222182548 | 0,304716 | -4,01088 | 6,05E-05 | 0,000218 | 311,2449 | -0,783576514 | 0,410926 | -1,90685 | 0,056539 | 0,1105 | 294,2097 | 1,154723 | 0,266665 | 4,330236 | 1,49E-05 | 8,8E-05 | 148,4274099 | 1,663874749 | 0,444583 | 3,742551 | 0,000182 | 0,000442 |
| 1952 | RM25_RS09510 | RM25_1928 | 3-dehydroquinate dehydratase | 2040 | gene | 2173886 | 2174641 | + | 629,4418 | -0,745994995 | 0,444366 | -1,67878 | 0,093194 | 0,184258 | 366,43306 | -1,152218368 | 0,267608 | -4,30561 | 1,67E-05 | 6,77E-05 | 892,4505 | -0,58955169 | 0,28954 | -2,03616 | 0,041734 | 0,088547 | 826,014 | 1,019267 | 0,298778 | 3,411448 | 0,000646 | 0,002453 | 461,867522 | 1,645045231 | 0,269738 | 6,098681 | 1,07E-09 | 5,95E-09 |
| 1953 | RM25_RS09515 | RM25_1929 | Putative membrane protein | 2041 | gene | 2174638 | 2175324 | + | 161,2784 | -0,279215978 | 0,29115 | -0,95901 | 0,337553 | 0,448232 | 120,9976605 | -0,410686884 | 0,295277 | -1,39085 | 0,16427 | 0,219365 | 201,5591 | -0,205961724 | 0,255645 | -0,80566 | 0,420441 | 0,524883 | 186,2028 | 0,572851 | 0,271016 | 2,113719 | 0,034539 | 0,063601 | 142,9545393 | 0,842518349 | 0,297321 | 2,833697 | 0,004601 | 0,008573 |
| 1954 | RM25_RS09520 | RM25_1930 | ATP-dependent protease ATP-binding subunit clpC1 | 2042 | gene | 2175475 | 2178051 | - | 16155,56 | 1,05107293 | 0,347261 | 3,026752 | 0,002472 | 0,017983 | 18271,76758 | 0,578494151 | 0,21181 | 2,731198 | 0,00631 | 0,0134 | 14039,34 | 1,766552148 | 0,408987 | 4,319337 | 1,56E-05 | 0,000153 | 11212,5 | -1,24934 | 0,396635 | -3,14984 | 0,001634 | 0,005318 | 21298,81978 | -0,029319183 | 0,228033 | -0,12857 | 0,897695 | 0,916832 |
| 1955 | RM25_RS09525 | RM25_1931 | Hypothetical protein | 2043 | gene | 2178915 | 2179271 | - | 3495,814 | 1,354949242 | 0,356082 | 3,805166 | 0,000142 | 0,002183 | 3691,036652 | 1,292202618 | 0,638508 | 2,023786 | 0,042992 | 0,070126 | 3300,592 | 1,426333914 | 0,400375 | 3,562494 | 0,000367 | 0,002192 | 2071,729 | -0,29169 | 0,520904 | -0,55996 | 0,575506 | 0,63924 | 4908,851978 | -0,143829275 | 0,335037 | -0,42929 | 0,66771 | 0,719612 |
| 1956 | RM25_RS09530 | RM25_1932 | 2-amino-4-hydroxy-6-hydroxymethyldihydropteridine dipho: | 2044 | gene | 2179845 | 2180390 | - | 669,6278 | 0,000792406 | 0,263418 | 0,003008 | 0,9976 | 0,998871 | 528,4022032 | 0,309713723 | 0,16327 | 1,896942 | 0,057836 | 0,088691 | 810,8534 | -0,198006219 | 0,302376 | -0,65483 | 0,512574 | 0,612956 | 704,9437 | 0,819241 | 0,28872 | 2,837496 | 0,004547 | 0,012214 | 654,9612375 | 0,356106142 | 0,181859 | 1,958149 | 0,050213 | 0,074248 |
| 1957 | RM25_RS09535 | RM25_1933 | Dihydroneopterin aldolase | 2045 | gene | 2180387 | 2180761 | - | 654,6844 | -0,189599449 | 0,262428 | -0,72248 | 0,469999 | 0,570743 | 520,9461417 | 0,286675084 | 0,163297 | 1,75554 | 0,079167 | 0,115898 | 788,4227 | -0,507608479 | 0,273329 | -1,85714 | 0,063292 | 0,121185 | 733,2951 | 0,925528 | 0,249201 | 3,713987 | 0,000204 | 0,000914 | 598,4874992 | 0,171827903 | 0,183916 | 0,934272 | 0,350164 | 0,414117 |
| 1958 | RM25_RS09540 | RM25_1934 | Dihydropteroate synthase | 2046 | gene | 2180908 | 2181741 | - | 562,4694 | -0,08100114 | 0,193622 | -0,41835 | 0,675694 | 0,755508 | 494,4293755 | 0,257572912 | 0,164228 | 1,568387 | 0,116791 | 0,163272 | 630,5095 | -0,344545187 | 0,237412 | -1,45125 | 0,14671 | 0,230992 | 608,1795 | 0,592345 | 0,175454 | 3,376073 | 0,000735 | 0,002725 | 537,1685469 | 0,045452791 | 0,22917 | 0,198337 | 0,842782 | 0,87307 |
| 1959 | RM25_RS09535 | RM25_1935 | Regulatory protein GntR, HTH | 2047 | gene | 2181878 | 2182345 | - | 168,2449 | 0,880715885 | 0,23374 | 3,767935 | 0,000165 | 0,002471 | 159,4190603 | 0,988380702 | 0,330329 | 2,992106 | 0,002771 | 0,00653 | 177,0707 | 0,788910852 | 0,305256 | 2,584426 | 0,009754 | 0,029742 | 124,583 | 0,231511 | 0,32692 | 0,708159 | 0,478846 | 0,55002 | 214,2604767 | 0,088209378 | 0,315813 | 0,279309 | 0,780008 | 0,821116 |
| 1960 | RM25_RS09550 | RM25_1936 | GTP cyclohydrolase 1 | 2048 | gene | 2182356 | 2183057 | - | 1477,913 | 0,603964944 | 0,206375 | 2,926539 | 0,003428 | 0,022688 | 1692,259755 | 0,678240839 | 0,19422 | 3,492132 | 0,000479 | 0,001394 | 1263,566 | 0,506823873 | 0,268907 | 1,884757 | 0,059463 | 0,115163 | 1243,834 | -0,37435 | 0,254459 | -1,47116 | 0,141247 | 0,197111 | 1744,826727 | -0,505123085 | 0,20252 | -2,49418 | 0,012625 | 0,02161 |
| 1961 | RM25_RS09555 | RM25_1937 | PIF1 helicase | 2049 | gene | 2183084 | 2185561 | - | 1373,145 | -0,505600663 | 0,267844 | -1,88767 | 0,059071 | 0,137852 | 1021,994735 | -0,414070462 | 0,17194 | -2,40823 | 0,01603 | 0,029727 | 1724,296 | -0,56133472 | 0,237148 | -2,36702 | 0,017932 | 0,047383 | 1693,096 | 0,753665 | 0,197381 | 3,818324 | 0,000134 | 0,000636 | 1114,931984 | 0,66647782 | 0,221843 | 3,00428 | 0,002662 | 0,005203 |
| 1962 | RM25_RS09560 | RM25_1938 | Hypoxanthine phosphoribosyltransferase | 2050 | gene | 2185633 | 2186184 | - | 1104,08 | 0,354948019 | 0,150047 | 2,365571 | 0,018002 | 0,067566 | 1191,491452 | 0,338218512 | 0,161433 | 2,095107 | 0,036161 | 0,060065 | 1016,669 | 0,375165046 | 0,24215 | 1,549308 | 0,121308 | 0,200226 | 1026,347 | -0,30932 | 0,169086 | -1,82938 | 0,067343 | 0,108643 | 1212,354028 | -0,227432461 | 0,207735 | -1,09482 | 0,273595 | 0,333953 |
| 1963 | RM25_RS09565 | RM25_1939 | TRNA(Ile)-lysidine synthase | 2051 | gene | 2186351 | 2187340 | - | 175,5884 | -0,075158401 | 0,668193 | -0,11248 | 0,910443 | 0,941556 | 68,39998356 | -1,197543379 | 0,362172 | -3,30656 | 0,000944 | 0,002553 | 282,7768 | 0,179681264 | 0,521263 | 0,344704 | 0,730317 | 0,800259 | 187,2337 | 1,42187 | 0,355163 | 4,003429 | 6,24E-05 | 0,000322 | 169,1539256 | 2,866845197 | 0,509871 | 5,622684 | 1,88E-08 | 8,62E-08 |
| 1964 | RM25_RS09570 | RM25_1940 | Coenzyme F420 biosynthesis-associated protein | 2052 | gene | 2187340 | 2188248 | - | 427,8852 | -0,501320305 | 0,266061 | -1,88423 | 0,059534 | 0,13852 | 383,1066028 | -0,129143039 | 0,16165 | -0,7989 | 0,424346 | 0,492345 | 472,6638 | -0,813722281 | 0,422118 | -1,92771 | 0,053891 | 0,106472 | 527,183 | 0,524781 | 0,201305 | 2,606903 | 0,009137 | 0,022019 | 349,1395341 | -0,0866831 | 0,401926 | -0,21567 | 0,829246 | 0,861787 |
| 1965 | RM25_RS13290 | RM25_1941 | Hypothetical protein | 2053 | gene | 2188200 | 2188430 | - | 281,8989 | 1,57200334 | 0,451394 | 3,482555 | 0,000497 | 0,006034 | 277,7195663 | 2,004748116 | 0,92172 | 2,175008 | 0,02963 | 0,050643 | 286,0783 | 1,206950621 | 0,236385 | 5,105877 | 3,29E-07 | 5,24E-06 | 148,2819 | 0,612491 | 0,58299 | 1,050604 | 0,293441 | 0,367698 | 413,7118671 | -0,163093884 | 0,484524 | -0,33661 | 0,736414 | 0,781155 |
| 1966 | RM25_RS09575 | RM25_1942 | D-alanyl-D-alanine carboxypeptidase, serine-type, PBP4 fan | 2054 | gene | 2188592 | 2189962 | - | 1186,513 | 0,15148384 | 0,208552 | 0,72636 | 0,467618 | 0,569305 | 1390,99325 | 0,253797195 | 0,170731 | 1,486531 | 0,137139 | 0,188696 | 982,0321 | 0,009626045 | 0,229796 | 0,041889 | 0,966587 | 0,980312 | 1189,334 | -0,433 | 0,21874 | -1,97952 | 0,047757 | 0,081925 | 1225,518865 | -0,623446674 | 0,174113 | -3,58071 | 0,000343 | 0,00079 |
| 1967 | RM25_RS09580 | RM25_1943 | Inorganic diphosphatase | 2055 | gene | 2190024 | 2190581 | + | 3434,227 | 0,440722367 | 0,346353 | 1,272464 | 0,203208 | 0,30683 | 4312,650608 | 0,713494615 | 0,209145 | 3,411484 | 0,000646 | 0,001822 | 2555,804 | -0,003961323 | 0,434222 | -0,00912 | 0,992721 | 0,995678 | 3099,213 | -0,4054 | 0,393411 | -1,03047 | 0,302788 | 0,377205 | 3869,96722 | -1,090598847 | 0,276114 | -3,94982 | 7,82E-05 | 0,000204 |
| 1968 | RM25_RS09585 | RM25_1944 | Hypothetical protein | 2056 | gene | 2190731 | 2191987 | + | 575,4801 | -0,369528852 | 0,363436 | -1,01676 | 0,309266 | 0,417973 | 394,4013707 | -0,752253308 | 0,196378 | -3,83065 | 0,000128 | 0,000424 | 756,5587 | -0,178440173 | 0,356553 | -0,50046 | 0,616752 | 0,703518 | 681,0053 | 0,631518 | 0,260998 | 2,419631 | 0,015536 | 0,03364 | 494,2261979 | 1,273640578 | 0,312987 | 4,069313 | 4,72E-05 | 0,000125 |
| 1969 | RM25_RS09590 | RM25_1945 | Phosphoglycerate mutase family protein | 2057 | gene | 2191989 | 2192531 | + | 463,7459 | -0,02706165 | 0,192672 | -0,14045 | 0,888301 | 0,925199 | 379,5727935 | 0,089031071 | 0,153249 | 0,580958 | 0,561269 | 0,625489 | 547,9191 | -0,10852567 | 0,160505 | -0,67615 | 0,498944 | 0,598783 | 492,1812 | 0,566488 | 0,144878 | 3,910103 | 9,23E-05 | 0,000452 | 450,0521548 | 0,422163529 | 0,14687 | 2,874402 | 0,004048 | 0,007584 |
| 1970 | RM25_RS09595 | RM25_1946 | CorA-like Mg2+ transporter family protein | 2058 | gene | 2192535 | 2193437 | + | 782,1617 | 0,085259666 | 0,120042 | 0,710248 | 0,47755 | 0,577816 | 751,1579352 | 0,309673325 | 0,139118 | 2,225969 | 0,026016 | 0,045055 | 813,1655 | -0,119118559 | 0,183277 | -0,64994 | 0,515733 | 0,61635 | 798,9089 | 0,27278 | 0,175316 | 1,555933 | 0,119724 | 0,172172 | 789,7628401 | -0,102958932 | 0,121178 | -0,84965 | 0,395519 | 0,46082 |
| 1971 | RM25_RS09600 | RM25_1947 | Hypothetical protein | 2059 | gene | 2193594 | 2194658 | + | 523,6584 | -0,009824188 | 0,124198 | -0,0791 | 0,936952 | 0,957161 | 516,3555642 | -0,057338336 | 0,175082 | -0,32749 | 0,743294 | 0,790945 | 530,9612 | 0,035219113 | 0,228636 | 0,15404 | 0,877578 | 0,911212 | 554,3056 | -0,06301 | 0,197243 | -0,31947 | 0,749373 | 0,79318 | 511,9615114 | 0,084769575 | 0,196692 | 0,430975 | 0,666486 | 0,718951 |
| 1972 | RM25_RS09605 | RM25_1948 | Ammonium-transport integral membrane protein Amt | 2060 | gene | 2194820 | 2196181 | + | 1649,724 | -1,386942756 | 0,633034 | -2,19095 | 0,028456 | 0,086721 | 718,3010755 | -2,386873571 | 0,156126 | -15,2881 | 9,18E-53 | 5,41E-50 | 2581,147 | -1,16226138 | 0,417722 | -2,78238 | 0,005396 | 0,019012 | 2504,488 | 1,506746 | 0,238217 | 6,325089 | 2,53E-10 | 4,17E-09 | 898,879857 | 2,790818049 | 0,377502 | 7,392856 | 1,44E-13 | 1,42E-12 |
| 1973 | RM25_RS09610 | RM25_1949 | Nitrogen regulatory protein P-II | 2061 | gene | 2196178 | 2196516 | + | 220,7942 | -0,227946676 | 0,611574 | -0,37272 | 0,709356 | 0,784954 | 125,4448265 | -2,112936936 | 0,316432 | -6,67737 | 2,43E-11 | 2,66E-10 | 316,1437 | 0,399027423 | 0,459313 | 0,868748 | 0,384985 | 0,490492 | 252,0618 | 0,358778 | 0,357761 | 1,002843 | 0,315937 | 0,390694 | 200,478689 | 2,93805598 | 0,455476 | 6,450518 | 1,11E-10 | 7,22E-10 |
| 1974 | RM25_RS09615 | RM25_1950 | Putative flavin-containing monoamine oxidase AofH | 2062 | gene | 2197326 | 2198681 | - | 1678,285 | 0,076222264 | 0,21173 | 0,359997 | 0,718849 | 0,793596 | 1951,88006 | 0,317459087 | 0,169798 | 1,869627 | 0,061536 | 0,093213 | 1404,69 | -0,255383641 | 0,199085 | -1,28279 | 0,199567 | 0,296395 | 1728,437 | -0,23745 | 0,23362 | -1,0164 | 0,309439 | 0,384173 | 1689,880842 | -0,765512881 | 0,114865 | -6,66444 | 2,66E-11 | 1,86E-10 |
| 1975 | RM25_RS09620 | RM25_1951 | D-serine/D-alanine/glycine transporter | 2063 | gene | 2198857 | 2200308 | - | 4553,018 | 1,297487033 | 0,385782 | 3,363263 | 0,00077 | 0,008405 | 3196,519288 | 0,155956517 | 0,186935 | 0,834282 | 0,404122 | 0,473653 | 5909,516 | 2,094703374 | 0,372947 | 5,61663 | 1,95E-08 | 3,96E-07 | 2796,023 | -0,48351 | 0,334096 | -1,44722 | 0,147837 | 0,204397 | 6301,306068 | 1,488652378 | 0,235051 | 6,333326 | 2,4E-10 | 1,44E-09 |
| 1976 | RM25_RS09625 | RM25_1952 | Antimicrobial peptide ABC transporter, permease protein | 2064 | gene | 2200572 | 2202086 | + | 620,0737 | -0,120955494 | 0,494122 | -0,24479 | 0,80662 | 0,864974 | 342,6249822 | -1,220599628 | 0,240089 | -5,08394 | 3,7E-07 | 2E-06 | 897,5224 | 0,271820986 | 0,239194 | 1,136403 | 0,255788 | 0,357163 | 678,8424 | 0,693448 | 0,268356 | 2,584059 | 0,009765 | 0,023294 | 581,9224819 | 2,246518338 | 0,212177 | 10,58797 | 3,39E-26 | 1,51E-24 |
| 1977 | RM25_RS09630 | RM25_1953 | ABC transporter, ATP-binding protein | 2065 | gene | 2202086 | 2202928 | + | 199,5163 | -0,155016829 | 0,547939 | -0,28291 | 0,777247 | 0,840739 | 102,6067282 | -1,410666585 | 0,298171 | -4,73106 | 2,23E-06 | 1,07E-05 | 296,4259 | 0,241975204 | 0,323196 | 0,748696 | 0,45404 | 0,557092 | 220,1013 | 0,800265 | 0,277763 | 2,881103 | 0,003963 | 0,010925 | 185,5511721 | 2,516485876 | 0,326323 | 7,711642 | 1,24E-14 | 1,43E-13 |
| 1979 | RM25_RS09640 | RM25_1954 | PSP1 domain protein | 2069 | gene | 2203236 | 2204285 | - | 523,8993 | 0,315084415 | 0,216474 | 1,455528 | 0,145523 | 0,246053 | 419,1685389 | 0,370992896 | 0,148919 | 2,491247 | 0,01273 | 0,024433 | 628,6301 | 0,278719949 | 0,210691 | 1,322884 | 0,185874 | 0,279582 | 490,7686 | 0,575695 | 0,15726 | 3,660793 | 0,000251 | 0,001097 | 570,303801 | 0,542866301 | 0,195926 | 2,770771 | 0,005592 | 0,010258 |
| 1980 | RM25_RS12010 | RM25_1955 | Hypothetical protein | 2070 | gene | 2204415 | 2206127 | - | 1618,461 | 0,475474617 | 0,262331 | 1,812497 | 0,069909 | 0,153432 | 1826,35064 | 0,433218488 | 0,224256 | 1,931803 | 0,053384 | 0,083439 | 1410,572 | 0,531623011 | 0,390294 | 1,362108 | 0,173164 | 0,264173 | 1436,186 | -0,47826 | 0,419711 | -1,1395 | 0,254497 | 0,324945 | 1844,278244 | -0,342353145 | 0,14745 | -2,32182 | 0,020242 | 0,033225 |
| 1981 | RM25_RS09650 | RM25_1956 | DNA polymerase III, delta' subunit | 2071 | gene | 2206275 | 2207657 | - | 701,2326 | 0,144716171 | 0,186951 | 0,774087 | 0,438879 | 0,547032 | 620,2831977 | 0,117684119 | 0,170752 | 0,689211 | 0,49069 | 0,557646 | 782,182 | 0,16625744 | 0,282797 | 0,587904 | 0,556597 | 0,652686 | 701,831 | 0,246699 | 0,200142 | 1,23262 | 0,217718 | 0,283828 | 723,6201959 | 0,358372533 | 0,256972 | 1,394595 | 0,163138 | 0,214096 |
| 1982 | RM25_RS09655 | RM25_1957 | DTMP kinase | 2072 | gene | 2207900 | 2208538 | - | 437,0991 | 0,334055176 | 0,20686 | 1,614885 | 0,106336 | 0,200346 | 475,4896076 | 0,289233105 | 0,257811 | 1,121882 | 0,261913 | 0,326629 | 398,7085 | 0,388041548 | 0,28078 | 1,382013 | 0,166968 | 0,256547 | 408,4366 | -0,36862 | 0,315109 | -1,16981 | 0,242077 | 0,310753 | 478,4960754 | -0,210101989 | 0,219552 | -0,95696 | 0,338588 | 0,402447 |
| 1983 | RM25_RS09660 | RM25_1958 | Hypothetical protein | 2073 | gene | 2208535 | 2209980 | - | 875,9704 | 0,338182947 | 0,187364 | 1,804947 | 0,071083 | 0,155564 | 826,3355609 | 0,347724137 | 0,225571 | 1,541528 | 0,123188 | 0,171605 | 925,6051 | 0,330474934 | 0,286731 | 1,152562 | 0,24909 | 0,34926 | 813,5911 | 0,117071 | 0,259633 | 0,450908 | 0,652056 | 0,706944 | 961,1445687 | 0,157443886 | 0,247308 | 0,636631 | 0,524365 | 0,585194 |
| 1984 | RM25_RS09665 | RM25_1959 | DNA topoisomerase | 2074 | gene | 2210024 | 2212819 | - | 3009,684 | 0,117944173 | 0,176026 | 0,670038 | 0,502833 | 0,600097 | 3414,204504 | -0,05580634 | 0,132669 | -0,42064 | 0,674015 | 0,727738 | 2605,163 | 0,3472536 | 0,193637 | 1,793324 | 0,072921 | 0,134698 | 3057,787 | -0,66266 | 0,18101 | -3,66087 | 0,000251 | 0,001097 | 3072,174172 | -0,205383348 | 0,112823 | -1,82041 | 0,068697 | 0,098431 |
| 1985 | RM25_RS09670 | RM25_1960 | RRNA or tRNA methyltransferase | 2075 | gene | 2213227 | 2214744 | - | 1229,7 | 0,085474688 | 0,143283 | 0,596544 | 0,550812 | 0,64526 | 1099,639116 | 0,071674138 | 0,159478 | 0,449429 | 0,653122 | 0,708752 | 1359,761 | 0,096749463 | 0,181623 | 0,532692 | 0,594246 | 0,683239 | 1257,245 | 0,234492 | 0,171504 | 1,36727 | 0,171541 | 0,231975 | 1242,138794 | 0,315255949 | 0,165708 | 1,90248 | 0,057108 | 0,083326 |
| 1986 | RM25_RS09680 | RM25_1961 | DEAD/DEAH box helicase | 2076 | gene | 2214994 | 2217456 | + | 1338,473 | 0,210151759 | 0,237885 | 0,883418 | 0,37701 | 0,487981 | 1062,840644 | 0,22605656 | 0,151351 | 1,493592 | 0,135282 | 0,186577 | 1614,105 | 0,198683283 | 0,271218 | 0,73256 | 0,463827 | 0,566446 | 1304,981 | 0,552604 | 0,195397 | 2,828108 | 0,004682 | 0,012511 | 1410,985415 | 0,588912296 | 0,236344 | 2,49176 | 0,012711 | 0,02171 |
| 1987 | RM25_RS09685 | RM25_1962 | Anaerobic dimethyl sulfoxide reductase chain C | 2077 | gene | 2217453 | 2218457 | - | 402,2477 | -0,500230239 | 0,214056 | -2,33691 | 0,019444 | 0,070724 | 357,4457473 | -0,771761702 | 0,199396 | -3,87049 | 0,000109 | 0,000367 | 447,0496 | -0,290937078 | 0,289834 | -1,00381 | 0,315472 | 0,420332 | 496,7855 | 0,071326 | 0,204441 | 0,348883 | 0,727177 | 0,773796 | 327,2259695 | 0,610647715 | 0,282601 | 2,160814 | 0,03071 | 0,048257 |
| 1988 | RM25_RS09690 | RM25_1963 | Dimethylsulfoxide reductase, chain B | 2078 | gene | 2218457 | 2219101 | - | 292,0855 | 0,067497624 | 0,212079 | 0,318266 | 0,750283 | 0,819447 | 279,7941398 | -0,303235197 | 0,214223 | -1,41551 | 0,156918 | 0,211467 | 304,3768 | 0,405456743 | 0,307389 | 1,319036 | 0,187157 | 0,281153 | 300,9766 | -0,29457 | 0,246153 | -1,19669 | 0,231427 | 0,299218 | 293,5720675 | 0,473326273 | 0,27359 | 1,730057 | 0,08362 | 0,117177 |
| 1989 | RM25_RS09695 | RM25_1964 | Anaerobic dimethyl sulfoxide reductase chain A | 2079 | gene | 2219115 | 2221718 | - | 2109,885 | -0,524795944 | 0,13548 | -3,8736 | 0,000107 | 0,001805 | 1963,184092 | -0,743152466 | 0,137146 | -5,41871 | 6E-08 | 3,78E-07 | 2256,586 | -0,340990669 | 0,172627 | -1,9753 | 0,048234 | 0,098092 | 2625,324 | -0,02185 | 0,159397 | -0,13705 | 0,890992 | 0,913073 | 1696,409001 | 0,433534595 | 0,12576 | 3,447315 | 0,000566 | 0,001263 |
| 1990 | RM25_RS09700 | RM25_1965 | 4Fe-4S ferredoxin iron-sulfur binding domain protein | 2080 | gene | 2221852 | 2223063 | - | 545,4558 | -0,878995584 | 0,632666 | -1,38935 | 0,164726 | 0,267091 | 265,1252955 | -2,237138211 | 0,336639 | -6,64552 | 3,02E-11 | 3,24E-10 | 825,7863 | -0,532030748 | 0,428155 | -1,24261 | 0,21401 | 0,311564 | 737,456 | 1,093476 | 0,358873 | 3,046972 | 0,002312 | 0,007131 | 379,0203997 | 2,866881676 | 0,413391 | 6,935038 | 4,06E-12 | 3,16E-11 |
| 1991 | NA | RM25_1966 | Cytoplasmic chaperone TorD | 2081 | gene | 2223279 | 2224175 | - | 702,5765 | 0,102046971 | 0,258144 | 0,39531 | 0,692614 | 0,770199 | 522,8916239 | -0,15800062 | 0,149387 | -1,05766 | 0,29021 | 0,358316 | 882,2614 | 0,255882838 | 0,140249 | 1,824485 | 0,068079 | 0,128164 | 713,6964 | 0,482418 | 0,141908 | 3,399514 | 0,000675 | 0,002542 | 712,2479707 | 0,949385723 | 0,139546 | 6,80338 | 1,02E-11 | 7,55E-11 |
| 1992 | RM25_RS09710 | RM25_1967 | Hypothetical protein | 2082 | gene | 2224257 | 2224523 | + | 239,8727 | -0,403194054 | 0,199363 | -2,02241 | 0,043134 | 0,113593 | 213,0523332 | -0,61344139 | 0,250904 | -2,44493 | 0,014488 | 0,027253 | 266,6932 | -0,24514386 | 0,233835 | -1,04836 | 0,294471 | 0,397747 | 288,1784 | 0,09951 | 0,228252 | 0,435968 | 0,66286 | 0,716023 | 202,4747628 | 0,52875295 | 0,264904 | 1,996021 | 0,045932 | 0,068563 |
| 1993 | RM25_RS12790 | RM25_1968 | Hypothetical protein | 2083 | gene | 2224685 | 2225017 | - | 120,8649 | -1,1142998 | 0,555786 | -2,00491 | 0,044973 | 0,116357 | 68,7528103 | -2,011134224 | 0,455387 | -4,41632 | 1E-05 | 4,23E-05 | 172,977 | -0,819763001 | 0,423738 | -1,9346 | 0,05304 | 0,105054 | 173,4448 | 0,932365 | 0,370616 | 2,515718 | 0,011879 | 0,027183 | 75,28863775 | 2,196293186 | 0,53364 | 4,115682 | 3,86E-05 | 0,000104 |
| 1994 | RM25_RS09720 | RM25_1969 | Putative membrane protein | 2084 | gene | 2225179 | 2225637 | - | 145,2872 | -0,54158895 | 0,618499 | -0,87565 | 0,38122 | 0,49181 | 69,33927081 | -1,665889463 | 0,379064 | -4,39475 | 1,11E-05 | 4,62E-05 | 221,235 | -0,236276898 | 0,49972 | -0,47282 | 0,636343 | 0,719357 | 180,5123 | 1,107544 | 0,321364 | 3,446391 | 0,000568 | 0,002195 | 117,0159862 | 2,617459889 | 0,547211 | 4,78327 | 1,72E-06 | 5,75E-06 |
| 1995 | RM25_RS09725 | RM25_1970 | Hypothetical protein | 2085 | gene | 2225768 | 2226079 | - | 91,70733 | -1,120941731 | 0,64267 | -1,74419 | 0,081125 | 0,168321 | 46,47656031 | -2,523300916 | 0,520416 | -4,84862 | 1,24E-06 | 6,21E-06 | 136,9381 | -0,761047509 | 0,370785 | -2,05253 | 0,040118 | 0,08604 | 131,8822 | 1,0393 | 0,356639 | 2,914153 | 0,003567 | 0,010104 | 56,82092382 | 2,882073862 | 0,558265 | 5,162551 | 2,44E-07 | 9,38E-07 |
| 1996 | RM25_RS09730 | RM25_1971 | Mcbg protein, putative | 2086 | gene | 2226238 | 2226843 | + | 246,2663 | -0,619477617 | 0,392052 | -1,58009 | 0,114086 | 0,208718 | 160,0440257 | -0,908082773 | 0,242797 | -3,7401 | 0,000184 | 0,000592 | 332,4885 | -0,488295013 | 0,36619 | -1,33345 | 0,182385 | 0,275036 | 312,5068 | 0,831468 | 0,248139 | 3,350812 | 0,000806 | 0,002926 | 191,1794975 | 1,317935547 | 0,367203 | 3,589119 | 0,000332 | 0,000768 |
| 1997 | RM25_RS09735 | RM25_1972 | Type II secretion system protein F domain protein | 2087 | gene | 2226976 | 2227719 | - | 173,6826 | -0,620944246 | 0,837115 | -0,74177 | 0,458228 | 0,562816 | 55,72076329 | -3,092206039 | 0,447534 | -6,90944 | 4,87E-12 | 5,88E-11 | 291,6444 | -0,295977012 | 0,463024 | -0,63923 | 0,522676 | 0,622195 | 219,4266 | 1,618096 | 0,2911 | 5,558551 | 2,72E-08 | 2,82E-07 | 135,2591469 | 4,485275974 | 0,564926 | 7,939586 | 2,03E-15 | 2,69E-14 |
| 1998 | RM25_RS09740 | RM25_1973 | Hypothetical protein | 2088 | gene | 2227716 | 2228513 | - | 203,9272 | -1,091169168 | 0,776074 | -1,40601 | 0,15972 | 0,262708 | 90,36175242 | -3,445491596 | 0,421031 | -8,18346 | 2,76E-16 | 5,24E-15 | 317,4927 | -0,669268084 | 0,512557 | -1,30574 | 0,19164 | 0,287338 | 290,6675 | 1,164211 | 0,315062 | 3,695178 | 0,00022 | 0,000968 | 128,9674082 | 4,01518317 | 0,60218 | 6,667748 | 2,6E-11 | 1,83E-10 |
| 1999 | RM25_RS09745 | RM25_1974 | Helicase/secretion neighborhood ATPase | 2089 | gene | 2228510 | 2229661 | - | 230,444 | -0,76981271 | 0,652252 | -1,18024 | 0,237905 | 0,345498 | 113,5356124 | -2,330983828 | 0,408801 | -5,702 | 1,18E-08 | 8,51E-08 | 347,3524 | -0,370111295 | 0,43533 | -0,85019 | 0,395222 | 0,501366 | 305,0724 | 0,970905 | 0,365551 | 2,656007 | 0,007907 | 0,019394 | 168,214194 | 3,014773323 | 0,492224 | 6,124798 | 9,08E-10 | 5,11E-09 |
| 2000 | RM25_RS12375 | RM25_1975 | Putative septum site-determining protein | 2090 | gene | 2229648 | 2231141 | - | 395,9206 | -0,740655381 | 0,801201 | -0,92443 | 0,355262 | 0,466539 | 143,3938541 | -3,008258931 | 0,373981 | -8,04389 | 8,7E-16 | 1,55E-14 | 648,4473 | -0,391916009 | 0,49897 | -0,78545 | 0,43219 | 0,537274 | 516,3871 | 1,457927 | 0,359033 | 4,060702 | 4,89E-05 | 0,000256 | 293,3289662 | 4,149398767 | 0,509254 | 8,147989 | 3,7E-16 | 5,32E-15 |
| 2001 | RM25_RS09755 | RM25_1976 | Putative methyltransferase | 2091 | gene | 2231266 | 2232015 | + | 648,1838 | -0,648803873 | 0,245297 | -2,64497 | 0,00817 | 0,040909 | 510,7459385 | -0,75416723 | 0,221312 | -3,4077 | 0,000655 | 0,001842 | 785,6217 | -0,582980275 | 0,195106 | -2,98801 | 0,002808 | 0,011497 | 833,0087 | 0,489723 | 0,221176 | 2,214177 | 0,026817 | 0,051513 | 495,1129706 | 0,720661807 | 0,215224 | 3,348426 | 0,000813 | 0,001765 |
| 2002 | RM25_RS09760 | RM25_1977 | Na(+)/H(+) antiporter NhaA | 2092 | gene | 2232319 | 2233542 | + | 1061,456 | 0,408432564 | 0,137942 | 2,960895 | 0,003067 | 0,021017 | 1135,519266 | 0,561664025 | 0,135122 | 4,156712 | 3,23E-05 | 0,000123 | 987,3934 | 0,23701259 | 0,230099 | 1,030047 | 0,302988 | 0,407383 | 961,8637 | -0,07686 | 0,172127 | -0,44652 | 0,655223 | 0,709724 | 1189,188184 | -0,343944533 | 0,185354 | -1,85561 | 0,06351 | 0,091723 |
| 2003 | RM25_RS09765 | RM25_1978 | Hypothetical protein | 2093 | gene | 2233877 | 2234350 | + | 2519,09 | 0,17715176 | 0,26084 | 0,679158 | 0,497038 | 0,595283 | 2875,30799 | 0,21383367 | 0,195205 | 1,09543 | 0,273328 | 0,339428 | 2162,871 | 0,129312609 | 0,395885 | 0,326642 | 0,743939 | 0,811413 | 2511,626 | -0,4151 | 0,382449 | -1,08536 | 0,277761 | 0,351186 | 2615,233028 | -0,467169756 | 0,217871 | -2,14425 | 0,032013 | 0,050053 |
| 2004 | RM25_RS09770 | RM25_1979 | Hypothetical protein | 2094 | gene | 2234466 | 2234693 | + | 282,0891 | 0,078183301 | 0,160069 | 0,488434 | 0,625243 | 0,711587 | 289,0137562 | 0,035567476 | 0,231781 | 0,153453 | 0,878041 | 0,905312 | 275,1644 | 0,123276402 | 0,256219 | 0,481137 | 0,630419 | 0,714031 | 289,1345 | -0,17107 | 0,2284 | -0,74897 | 0,453874 | 0,527765 | 283,8373338 | -0,03702323 | 0,235954 | -0,15691 | 0,875317 | 0,898572 |
| 2005 | RM25_RS12015 | RM25_1980 | Hypothetical protein | 2095 | gene | 2234809 | 2235588 | - | 1253,986 | 0,318239157 | 0,274453 | 1,159541 | 0,246236 | 0,354792 | 1554,572842 | 0,286361712 | 0,219594 | 1,304052 | 0,192216 | 0,250999 | 953,3989 | 0,372493752 | 0,25078 | 1,485342 | 0,137453 | 0,220393 | 1184,157 | -0,80502 | 0,307483 | -2,6181 | 0,008842 | 0,021419 | 1364,920806 | -0,674854218 | 0,125902 | -5,36015 | 8,32E-08 | 3,44E-07 |
| 2006 | RM25_RS09780 | RM25_1981 | Copper-exporting ATPase | 2096 | gene | 2235737 | 2238292 | - | 1283,586 | -0,449759158 | 0,352509 | -1,27588 | 0,201998 | 0,305395 | 941,5591431 | -1,154321812 | 0,152722 | -7,55833 | 4,08E-14 | 6,33E-13 | 1625,613 | -0,070466645 | 0,311393 | -0,22629 | 0,820972 | 0,870068 | 1558,633 | 0,295136 | 0,210313 | 1,403319 | 0,160522 | 0,21936 | 1066,827229 | 1,442337914 | 0,263042 | 5,483289 | 4,17E-08 | 1,83E-07 |
| 2007 | RM25_RS09785 | RM25_1982 | Heavy metal transport/detoxification protein | 2097 | gene | 2238402 | 2238614 | - | 396,8218 | 0,185514373 | 0,387688 | 0,478514 | 0,632284 | 0,718146 | 304,250194 | -1,068228443 | 0,210471 | -5,07543 | 3,87E-07 | 2,07E-06 | 489,3933 | 0,966346309 | 0,224607 | 4,302384 | 1,69E-05 | 0,000163 | 392,279 | -0,37055 | 0,20385 | -1,81776 | 0,069101 | 0,110721 | 412,3328149 | 1,711310909 | 0,19003 | 9,00548 | 2,15E-19 | 4,73E-18 |
| 2008 | RM25_RS12380 | RM25_1983 | Hypothetical protein | 2098 | gene | 2238749 | 2239471 | - | 575,8369 | -0,254507022 | 0,309367 | -0,82267 | 0,410695 | 0,520995 | 477,7172558 | -1,168851013 | 0,147268 | -7,93691 | 2,07E-15 | 3,62E-14 | 673,9566 | 0,351383147 | 0,176442 | 1,991492 | 0,046427 | 0,095738 | 662,6186 | -0,22032 | 0,155596 | -1,416 | 0,156776 | 0,214962 | 514,1458401 | 1,355206838 | 0,153212 | 8,845284 | 9,13E-19 | 1,86E-17 |
| 2009 | RM25_RS09795 | RM25_1984 | Replicative DNA helicase | 2099 | gene | 2240067 | 2241422 | + | 2163,12 | 0,912854344 | 0,273782 | 3,334236 | 0,000855 | 0,009121 | 2238,523222 | 0,296972307 | 0,164083 | 1,809893 | 0,070312 | 0,104625 | 2087,717 | 1,680954171 | 0,24776 | 6,78461 | 1,16E-11 | 5,02E-10 | 1595,359 | -1,07813 | 0,199589 | -5,40175 | 6,6E-08 | 6,35E-07 | 2761,045141 | 0,352144244 | 0,179797 | 1,958567 | 0,050163 | 0,074222 |
| 2010 | RM25_RS12020 | RM25_1985 | Hypothetical protein | 2100 | gene | 2241551 | 2242081 | - | 278,88 | -1,115678231 | 0,508602 | -2,19362 | 0,028263 | 0,086721 | 159,7380403 | -1,712149991 | 0,334751 | -5,1147 | 3,14E-07 | 1,72E-06 | 398,0219 | -0,90463364 | 0,425646 | -2,12532 | 0,03356 | 0,075119 | 400,7889 | 1,0094 | 0,343287 | 2,940393 | 0,003278 | 0,00955 | 173,7068674 | 1,895663766 | 0,448007 | 4,231326 | 2,32E-05 | 6,5E-05 |
| 2011 | RM25_RS09805 | RM25_1986 | Lincomycin resistance protein | 2101 | gene | 2242228 | 2243631 | + | 655,1754 | -1,500431683 | 0,592332 | -2,53309 | 0,011306 | 0,050856 | 373,5079251 | -2,788102079 | 0,335911 | -8,30012 | 1,04E-16 | 2,12E-15 | 936,8429 | -1,122880938 | 0,409308 | -2,74337 | 0,006081 | 0,020956 | 1014,615 | 0,908567 | 0,378503 | 2,400423 | 0,016376 | 0,035069 | 337,512423 | 2,646624469 | 0,396825 | 6,669507 | 2,57E-11 | 1,82E-10 |
| 2012 | RM25_RS09810 | RM25_1987 | NADH:flavin oxidoreductase/NADH oxidase | 2102 | gene | 2243663 | 2244775 | - | 741,3672 | -0,000916917 | 0,210732 | -0,00435 | 0,996528 | 0,998797 | 611,7988472 | 0,072556301 | 0,159824 | 0,453976 | 0,649846 | 0,705522 | 870,9356 | -0,052263414 | 0,250562 | -0,20858 | 0,834772 | 0,879159 | 779,2606 | 0,511987 | 0,157124 | 3,258499 | 0,00112 | 0,003882 | 727,9477001 | 0,446316975 | 0,235523 | 1,895004 | 0,058092 | 0,084625 |
| 2013 | RM25_RS09815 | RM25_1988 | UbiE/COQ5 methyltransferase, putative | 2103 | gene | 2244930 | 2245748 | + | 219,3745 | -0,851218018 | 0,550751 | -1,54556 | 0,122211 | 0,219217 | 122,3581995 | -1,773611455 | 0,341964 | -5,18654 | 2,14E-07 | 1,23E-06 | 316,3908 | -0,545289986 | 0,462083 | -1,18007 | 0,237973 | 0,336878 | 295,8913 | 0,916553 | 0,307906 | 2,976731 | 0,002913 | 0,008594 | 154,4679783 | 2,222762993 | 0,501128 | 4,435524 | 9,18E-06 | 2,7E-05 |
| 2014 | RM25_RS09820 | RM25_1989 | Hypothetical protein | 2104 | gene | 2245831 | 2246400 | - | 248,0853 | -0,962306163 | 0,438983 | -2,19213 | 0,02837 | 0,086721 | 152,6552364 | -1,323733742 | 0,293072 | -4,51675 | 6,28E-06 | 2,74E-05 | 343,5155 | -0,813910817 | 0,38192 | -2,1311 | 0,033081 | 0,0744 | 344,1011 | 0,934395 | 0,277215 | 3,370652 | 0,00075 | 0,002757 | 165,7057359 | 1,517473568 | 0,412478 | 3,678918 | 0,000234 | 0,000558 |
| 2015 | RM25_RS09830 | RM25_1990 | Hypothetical protein | 2105 | gene | 2246919 | 2247251 | + | 478,5983 | -0,947597909 | 0,399124 | -2,37419 | 0,017587 | 0,066722 | 300,5987714 | -1,384016297 | 0,252412 | -5,48316 | 4,18E-08 | 2,73E-07 | 656,5978 | -0,765064026 | 0,144451 | -5,29636 | 1,18E-07 | 2,06E-06 | 662,1975 | 0,864036 | 0,24562 | 3,51777 | 0,000435 | 0,001756 | 319,9880523 | 1,539088256 | 0,193158 | 7,968029 | 1,61E-15 | 2,18E-14 |
| 2016 | RM25_RS09835 | RM25_1991 | PF06224 family protein | 2106 | gene | 2247512 | 2248672 | + | 354,6995 | -0,350257376 | 0,482712 | -0,7256 | 0,468082 | 0,569576 | 186,3631967 | -0,500907699 | 0,227842 | -2,19849 | 0,027914 | 0,04806 | 523,0358 | -0,299290535 | 0,398963 | -0,75017 | 0,453152 | 0,556291 | 415,0335 | 1,33938 | 0,245999 | 5,444648 | 5,19E-08 | 5,06E-07 | 307,4373499 | 1,607458579 | 0,380082 | 4,229236 | 2,34E-05 | 6,56E-05 |
| 2017 | RM25_RS09840 | RM25_1992 | Phosphomethylpyrimidine synthase | 2107 | gene | 2248799 | 2250691 | - | 921,5308 | -0,575490945 | 0,303024 | -1,89916 | 0,057544 | 0,13536 | 694,4328145 | -0,968742966 | 0,17129 | -5,65558 | 1,55E-08 | 1,09E-07 | 1148,629 | -0,350783181 | 0,287285 | -1,22103 | 0,222075 | 0,318991 | 1159,743 | 0,419866 | 0,233811 | 1,79575 | 0,072534 | 0,114587 | 726,8857095 | 1,101710216 | 0,237609 | 4,636649 | 3,54E-06 | 1,12E-05 |
| 2018 | RM25_RS09845 | RM25_1993 | Na+/galactose cotransporter / Sodium/glucose cotransporte | 2108 | gene | 2251029 | 2252708 | + | 4070,417 | -0,303965706 | 0,234143 | -1,2982 | 0,194217 | 0,298027 | 4026,082815 | -0,734549562 | 0,158586 | -4,63186 | 3,62E-06 | 1,67E-05 | 4114,752 | 0,104533246 | 0,352077 | 0,296904 | 0,76654 | 0,829157 | 4771,261 | -0,39585 | 0,306542 | -1,29135 | 0,196582 | 0,259868 | 3554,662036 | 0,47809561 | 0,218314 | 2,189945 | 0,028528 | 0,045219 |
| 2019 | RM25_RS09850 | RM25_1994 | Hypothetical protein | 2109 | gene | 2252730 | 2253155 | + | 1173,317 | -0,006863982 | 0,273821 | -0,02507 | 0,980001 | 0,987709 | 1183,305789 | -0,549235327 | 0,20935 | -2,62353 | 0,008702 | 0,017592 | 1163,328 | 0,544470544 | 0,369296 | 1,474347 | 0,140388 | 0,224033 | 1249,437 | -0,62234 | 0,350068 | -1,77776 | 0,075443 | 0,117683 | 1142,249787 | 0,505072411 | 0,231811 | 2,178807 | 0,029346 | 0,04636 |
| 2020 | RM25_RS09855 | RM25_1995 | Putative TRANSCRIPTIONAL REPRESSOR SIRR | 2110 | gene | 2253260 | 2253937 | - | 447,3349 | -0,441405284 | 0,209322 | -2,10874 | 0,034967 | 0,099658 | 379,5969589 | -0,59512937 | 0,168842 | -3,52477 | 0,000424 | 0,001258 | 515,0729 | -0,333155833 | 0,273297 | -1,21903 | 0,222834 | 0,319671 | 543,5093 | 0,265446 | 0,197835 | 1,341755 | 0,179675 | 0,240891 | 372,3225649 | 0,58833002 | 0,257653 | 2,283418 | 0,022406 | 0,036371 |
| 2021 | RM25_RS09860 | RM25_1996 | Divalent metal cation transporter MntH | 2111 | gene | 2254159 | 2255574 | + | 612,1543 | -1,162593912 | 0,361473 | -3,21626 | 0,001299 | 0,011819 | 448,2173776 | -1,464182324 | 0,273604 | -5,35146 | 8,72E-08 | 5,4E-07 | 776,0913 | -1,00100866 | 0,378219 | -2,64664 | 0,00813 | 0,026249 | 889,46 | 0,584421 | 0,321778 | 1,816224 | 0,069336 | 0,111022 | 372,352561 | 1,120912099 | 0,358377 | 3,127742 | 0,001762 | 0,003528 |
| 2022 | RM25_RS12630 | RM25_1997 | Hypothetical protein | 2112 | gene | 2255783 | 2256028 | - | 216,977 | -0,51234302 | 0,247716 | -2,06827 | 0,038615 | 0,106699 | 176,3331065 | -0,774440989 | 0,222491 | -3,48077 | 0,0005 | 0,001448 | 257,6209 | -0,343880382 | 0,271919 | -1,26464 | 0,206 | 0,302689 | 268,7649 | 0,305822 | 0,237405 | 1,288188 | 0,197681 | 0,261173 | 175,4397866 | 0,799447999 | 0,267529 | 2,988265 | 0,002806 | 0,005432 |
| 2023 | RM25_RS09865 | RM25_1998 | Hypothetical protein | 2113 | gene | 2256588 | 2257424 | + | 511,0347 | -0,765088067 | 0,484335 | -1,57967 | 0,114183 | 0,208718 | 301,089033 | -1,49846029 | 0,265584 | -5,64214 | 1,68E-08 | 1,17E-07 | 720,9804 | -0,491415548 | 0,387682 | -1,26757 | 0,20495 | 0,301918 | 673,7133 | 0,857187 | 0,298086 | 2,875638 | 0,004032 | 0,011077 | 373,1206105 | 1,93219402 | 0,365126 | 5,291858 | 1,21E-07 | 4,88E-07 |
| 2024 | RM25_RS09870 | RM25_1999 | Hypothetical protein | 2114 | gene | 2257446 | 2258309 | - | 297,596 | -1,176694432 | 0,851518 | -1,38188 | 0,167009 | 0,269586 | 119,8527387 | -4,091167074 | 0,475949 | -8,59581 | 8,27E-18 | 1,91E-16 | 475,3392 | -0,768794522 | 0,462236 | -1,66321 | 0,096271 | 0,167338 | 431,5265 | 1,32825 | 0,404247 | 3,28574 | 0,001017 | 0,003573 | 180,40999 | 4,729194496 | 0,547474 | 8,638208 | 5,71E-18 | 1,03E-16 |
| 2025 | RM25_RS09875 | RM25_2000 | Von Willebrand factor type A domain protein | 2115 | gene | 2258306 | 2259313 | - | 297,4288 | -0,973550427 | 0,861591 | -1,12994 | 0,2585 | 0,366156 | 109,5076794 | -3,954133127 | 0,442541 | -8,93506 | 4,07E-19 | 1,08E-17 | 485,3499 | -0,587004803 | 0,439836 | -1,3346 | 0,182007 | 0,274642 | 411,1857 | 1,434744 | 0,347999 | 4,122843 | 3,74E-05 | 0,000201 | 198,2461668 | 4,870671292 | 0,521755 | 9,335163 | 1,01E-20 | 2,48E-19 |
| 2026 | RM25_RS09880 | RM25_2001 | Von Willebrand factor type A domain protein | 2116 | gene | 2259310 | 2260320 | - | 309,0155 | -1,314088871 | 0,810417 | -1,6215 | 0,104911 | 0,198935 | 135,5312365 | -3,867579973 | 0,496641 | -7,78747 | 6,84E-15 | 1,13E-13 | 482,4998 | -0,897767085 | 0,464815 | -1,93145 | 0,053427 | 0,105645 | 461,1104 | 1,23275 | 0,475993 | 2,58985 | 0,009602 | 0,022953 | 175,1890099 | 4,283348427 | 0,518917 | 8,254393 | 1,53E-16 | 2,26E-15 |
| 2027 | RM25_RS09885 | RM25_2002 | Hypothetical protein | 2117 | gene | 2260311 | 2260931 | - | 149,9913 | -1,05209558 | 0,853552 | -1,23261 | 0,217722 | 0,323155 | 58,20858513 | -3,887818913 | 0,538842 | -7,21513 | 5,39E-13 | 7,34E-12 | 241,774 | -0,656623502 | 0,517209 | -1,26955 | 0,204245 | 0,301431 | 211,1175 | 1,369076 | 0,380786 | 3,595393 | 0,000324 | 0,001376 | 96,6128242 | 4,67577123 | 0,654191 | 7,14741 | 8,84E-13 | 7,75E-12 |
| 2028 | RM25_RS09890 | RM25_2003 | PF01882 family protein | 2118 | gene | 2260915 | 2261976 | - | 367,3342 | -1,184332151 | 0,834434 | -1,41932 | 0,155805 | 0,257345 | 157,5179609 | -4,090920459 | 0,404942 | -10,1025 | 5,39E-24 | 3,1E-22 | 577,1504 | -0,745106418 | 0,503126 | -1,48095 | 0,138619 | 0,222111 | 533,3833 | 1,208631 | 0,367593 | 3,287961 | 0,001009 | 0,003555 | 222,1664523 | 4,632435188 | 0,54316 | 8,528675 | 1,48E-17 | 2,49E-16 |
| 2029 | RM25_RS09900 | RM25_2004 | Transposase IS204/IS1001/IS1096/IS1165 family protein | 2119 | gene | 2262225 | 2263190 | - | 47,02524 | -1,655865445 | 0,725187 | -2,28336 | 0,022409 | 0,07677 | 24,37060997 | -3,548623604 | 0,770832 | -4,60363 | 4,15E-06 | 1,88E-05 | 69,67986 | -1,244679984 | 0,436273 | -2,85298 | 0,004331 | 0,016127 | 74,97661 | 1,040431 | 0,477276 | 2,179935 | 0,029262 | 0,055425 | 22,26446679 | 3,426701734 | 0,795366 | 4,308334 | 1,64E-05 | 4,71E-05 |
| 2030 | RM25_RS09905 | RM25_2005 | Modification methylase Cfr9I | 2120 | gene | 2263802 | 2264974 | + | 2337,344 | 0,600935342 | 0,24931 | 2,410396 | 0,015935 | 0,063009 | 2559,38602 | 0,185142013 | 0,177006 | 1,045965 | 0,295577 | 0,36418 | 2115,301 | 1,141793722 | 0,285425 | 4,000322 | 6,33E-05 | 0,000507 | 1975,432 | -0,91621 | 0,239612 | -3,82372 | 0,000131 | 0,000623 | 2753,759459 | 0,080807073 | 0,213973 | 0,377652 | 0,705689 | 0,754336 |
| 2031 | RM25_RS09910 | RM25_2006 | 5-methylcytosine restriction system component-like protein | 2121 | gene | 2264945 | 2266345 | - | 1942,339 | -0,200682314 | 0,151813 | -1,3219 | 0,1862 | 0,291031 | 1924,204125 | -0,436102325 | 0,122455 | -3,56134 | 0,000369 | 0,001115 | 1960,473 | 0,026719279 | 0,265507 | 0,100635 | 0,91984 | 0,944278 | 2194,364 | -0,25319 | 0,228515 | -1,10797 | 0,267876 | 0,340368 | 1769,077448 | 0,262778585 | 0,142884 | 1,839104 | 0,0659 | 0,094885 |
| 2032 | RM25_RS09915 | RM25_2007 | 5-methylcytosine-specific restriction enzyme B | 2122 | gene | 2266342 | 2267325 | - | 2043,3 | 0,639146048 | 0,327269 | 1,952966 | 0,050824 | 0,125567 | 2254,74699 | 0,037790865 | 0,198531 | 0,190353 | 0,849033 | 0,881573 | 1831,852 | 1,472337436 | 0,348221 | 4,228174 | 2,36E-05 | 0,000216 | 1698,113 | -1,25158 | 0,208873 | -5,99206 | 2,07E-09 | 2,68E-08 | 2427,416196 | 0,217211362 | 0,292292 | 0,743131 | 0,457402 | 0,522587 |
| 2033 | RM25_RS13015 | RM25_2008 | Hypothetical protein | 2123 | gene | 2267306 | 2268031 | - | 1981,332 | 0,489748278 | 0,270273 | 1,812051 | 0,069978 | 0,153432 | 2195,551143 | 0,207679963 | 0,199307 | 1,042009 | 0,297408 | 0,366172 | 1767,113 | 0,854731575 | 0,387646 | 2,204927 | 0,027459 | 0,065047 | 1750,158 | -0,74567 | 0,280332 | -2,65994 | 0,007816 | 0,019229 | 2259,084788 | -0,068907666 | 0,30428 | -0,22646 | 0,820843 | 0,854561 |
| 2034 | RM25_RS09925 | RM25_2009 | Fic family toxin-antitoxin system | 2124 | gene | 2268235 | 2268498 | + | 80,31722 | -1,202921771 | 0,282364 | -4,26018 | 2,04E-05 | 0,00049 | 66,10921042 | -1,327735996 | 0,373102 | -3,55864 | 0,000373 | 0,001125 | 94,52522 | -1,127119763 | 0,32191 | -3,50135 | 0,000463 | 0,002648 | 118,2949 | 0,380468 | 0,272422 | 1,396613 | 0,16253 | 0,221307 | 47,65106765 | 0,653519624 | 0,472807 | 1,382211 | 0,166907 | 0,218434 |
| 2035 | NA | RM25_2010 | Transposase IS3/IS911 family protein | 2125 | gene | 2268557 | 2268886 | + | 0 | NA | NA | NA | NA | NA | 0 | NA | NA | NA | NA | NA | 0 | NA | NA | NA | NA | NA | 0 | NA | NA | NA | NA | NA | 0 | NA | NA | NA | NA | NA |
| 2036 | RM25_RS09935 | RM25_2011 | IS3 family transposase OrfB | 2126 | gene | 2268883 | 2269803 | + | 18,38324 | -1,112858934 | 0,374137 | -2,97447 | 0,002935 | 0,020514 | 14,54283607 | -1,084735563 | 0,618525 | -1,75374 | 0,079474 | 0,116204 | 22,22365 | -1,147111654 | 0,469734 | -2,44205 | 0,014604 | 0,040401 | 26,2158 | 0,565738 | 0,398564 | 1,419441 | 0,15577 | 0,213708 | 11,43262201 | 0,562884853 | 0,727102 | 0,774148 | 0,438843 | 0,504317 |
| 2037 | RM25_RS09940 | RM25_2012 | Hypothetical protein | 2127 | gene | 2269957 | 2270379 | + | 136,8558 | -1,470942949 | 0,661112 | -2,22495 | 0,026084 | 0,083082 | 84,6114318 | -3,494828153 | 0,476179 | -7,33931 | 2,15E-13 | 3,09E-12 | 189,1003 | -0,912607421 | 0,417787 | -2,18439 | 0,028934 | 0,067522 | 211,6519 | 0,594858 | 0,390451 | 1,523517 | 0,127629 | 0,181656 | 71,52787256 | 3,254582822 | 0,550373 | 5,913418 | 3,35E-09 | 1,72E-08 |
| 2038 | RM25_RS09945 | RM25_2013 | Hypothetical protein | 2128 | gene | 2270485 | 2271585 | + | 453,7348 | -0,580725753 | 0,327948 | -1,77079 | 0,076596 | 0,163976 | 318,0942276 | -0,569951629 | 0,218741 | -2,6056 | 0,009171 | 0,018429 | 589,3754 | -0,58923689 | 0,324394 | -1,81643 | 0,069305 | 0,129743 | 571,2201 | 0,83083 | 0,238052 | 3,490122 | 0,000483 | 0,001913 | 357,9348332 | 0,879795874 | 0,32059 | 2,744305 | 0,006064 | 0,011054 |
| 2039 | RM25_RS09950 | RM25_2014 | 2-oxoglutarate/malate translocator | 2129 | gene | 2272359 | 2273570 | + | 521,0832 | -1,351454423 | 0,518715 | -2,60539 | 0,009177 | 0,043964 | 289,4931843 | -2,176251014 | 0,315778 | -6,89172 | 5,51E-12 | 6,56E-11 | 752,6732 | -1,087203805 | 0,230983 | -4,70686 | 2,52E-06 | 3,04E-05 | 785,7023 | 1,04483 | 0,329746 | 3,168586 | 0,001532 | 0,005043 | 287,9815059 | 2,195142605 | 0,262741 | 8,354778 | 6,56E-17 | 1,02E-15 |
| 2040 | RM25_RS09955 | RM25_2015 | 6, 7-dimethyl-8-ribityllumazine synthase | 2130 | gene | 2273727 | 2274194 | - | 476,9192 | 2,890436548 | 0,549027 | 5,264651 | 1,4E-07 | 8,07E-06 | 166,5923665 | 1,613142852 | 0,301687 | 5,347077 | 8,94E-08 | 5,49E-07 | 787,246 | 3,302063872 | 0,322198 | 10,24857 | 1,2E-24 | 2,83E-22 | 118,7265 | 0,755779 | 0,257024 | 2,940494 | 0,003277 | 0,00955 | 821,0287544 | 2,498169652 | 0,311149 | 8,02885 | 9,84E-16 | 1,36E-14 |
| 2041 | RM25_RS12025 | RM25_2016 | Riboflavin biosynthesis protein ribBA | 2131 | gene | 2274191 | 2275660 | - | 1898,652 | 2,487735678 | 0,536503 | 4,636945 | 3,54E-06 | 0,000114 | 700,9015139 | 1,351759841 | 0,262741 | 5,144847 | 2,68E-07 | 1,51E-06 | 3096,403 | 2,847962693 | 0,268158 | 10,62048 | 2,39E-26 | 9,4E-24 | 602,6588 | 0,867454 | 0,248755 | 3,487177 | 0,000488 | 0,001927 | 3148,407899 | 2,420968002 | 0,269347 | 8,988302 | 2,51E-19 | 5,48E-18 |
| 2042 | RM25_RS09965 | RM25_2017 | Riboflavin synthase, alpha subunit | 2132 | gene | 2275657 | 2276319 | - | 888,36 | 1,777521822 | 0,561741 | 3,164311 | 0,001555 | 0,012946 | 332,7131889 | 0,656223099 | 0,287947 | 2,278975 | 0,022669 | 0,039814 | 1444,007 | 2,104363255 | 0,284406 | 7,399155 | 1,37E-13 | 9,23E-12 | 420,132 | 1,005824 | 0,31018 | 3,242709 | 0,001184 | 0,004068 | 1341,778277 | 2,509917381 | 0,245055 | 10,24224 | 1,28E-24 | 4,51E-23 |
| 2043 | RM25_RS09970 | RM25_2018 | Riboflavin biosynthesis protein RibD | 2133 | gene | 2276409 | 2277653 | - | 522,8029 | 0,411868931 | 0,597579 | 0,689229 | 0,490679 | 0,590355 | 217,2610973 | -0,89846392 | 0,333764 | -2,69192 | 0,007104 | 0,014779 | 828,3448 | 0,759442166 | 0,231171 | 3,285193 | 0,001019 | 0,004932 | 469,6666 | 1,050413 | 0,350483 | 2,997045 | 0,002726 | 0,008227 | 584,3630167 | 2,76943911 | 0,205392 | 13,48367 | 1,95E-41 | 2,56E-39 |
| 2044 | RM25_RS09975 | RM25_2019 | Hypothetical protein | 2134 | gene | 2278054 | 2278413 | + | 230,9855 | -0,948135527 | 0,370576 | -2,55855 | 0,010511 | 0,048578 | 154,7282629 | -1,268317495 | 0,281824 | -4,50038 | 6,78E-06 | 2,93E-05 | 307,2428 | -0,800969336 | 0,269516 | -2,97188 | 0,00296 | 0,011912 | 319,4519 | 0,768624 | 0,266403 | 2,885193 | 0,003912 | 0,010847 | 154,8017723 | 1,299942432 | 0,30389 | 4,277676 | 1,89E-05 | 5,36E-05 |
| 2045 | RM25_RS12030 | RM25_2020 | Hypothetical protein | 2135 | gene | 2278471 | 2278953 | + | 456,1757 | -0,396134506 | 0,379732 | -1,0432 | 0,296857 | 0,407035 | 291,8177152 | -0,681477909 | 0,278016 | -2,45122 | 0,014237 | 0,026889 | 620,5336 | -0,268345893 | 0,225906 | -1,18787 | 0,234886 | 0,333589 | 543,7612 | 0,847471 | 0,31564 | 2,684934 | 0,007254 | 0,018075 | 385,9221666 | 1,320374703 | 0,184179 | 7,168985 | 7,56E-13 | 6,67E-12 |
| 2046 | RM25_RS09985 | RM25_2021 | 4-alpha-glucanotransferase malQ | 2136 | gene | 2279024 | 2280979 | + | 1599,449 | 0,000685032 | 0,284264 | 0,00241 | 0,998077 | 0,998925 | 1152,467384 | -0,082114215 | 0,186879 | -0,4394 | 0,660374 | 0,715304 | 2046,431 | 0,045996118 | 0,18246 | 0,252089 | 0,800972 | 0,856575 | 1680,311 | 0,698198 | 0,234479 | 2,977656 | 0,002905 | 0,00859 | 1568,407496 | 0,886386351 | 0,137414 | 6,450471 | 1,12E-10 | 7,22E-10 |
| 2047 | RM25_RS12385 | RM25_2022 | Hypothetical protein | 2137 | gene | 2281654 | 2282031 | + | 173,9894 | -1,335455611 | 0,578573 | -2,30819 | 0,020989 | 0,073505 | 106,2223707 | -2,778544058 | 0,393284 | -7,06499 | 1,61E-12 | 2,07E-11 | 241,7565 | -0,878777745 | 0,359472 | -2,44463 | 0,0145 | 0,040302 | 261,6739 | 0,681087 | 0,348305 | 1,955435 | 0,050532 | 0,085686 | 97,06841772 | 2,656745849 | 0,440755 | 6,027711 | 1,66E-09 | 8,99E-09 |
| 2048 | RM25_RS10000 | RM25_2023 | NAD(P) transhydrogenase subunit beta | 2138 | gene | 2282239 | 2283651 | - | 8279,587 | 0,805592187 | 0,457461 | 1,761009 | 0,078237 | 0,165486 | 11027,68795 | 0,402863629 | 0,211311 | 1,906501 | 0,056585 | 0,087285 | 5531,485 | 1,736463044 | 0,401589 | 4,323982 | 1,53E-05 | 0,00015 | 6445,976 | -1,94454 | 0,382692 | -5,08122 | 3,75E-07 | 3,11E-06 | 10307,12846 | -0,577790883 | 0,239703 | -2,41045 | 0,015933 | 0,026748 |
| 2049 | RM25_RS10005 | RM25_2024 | NAD(P)(+) transhydrogenase (AB-specific), alpha subunit | 2139 | gene | 2283658 | 2285211 | - | 7967,894 | 0,832828252 | 0,342618 | 2,430775 | 0,015067 | 0,060889 | 9696,645491 | 0,484192635 | 0,213526 | 2,267603 | 0,023353 | 0,040864 | 6239,143 | 1,431739244 | 0,347276 | 4,12277 | 3,74E-05 | 0,000316 | 6106,446 | -1,31122 | 0,322893 | -4,06085 | 4,89E-05 | 0,000256 | 9979,702324 | -0,330067136 | 0,23676 | -1,3941 | 0,163288 | 0,214174 |
| 2050 | NA | RM25_2025 | Transcription regulator | 2140 | gene | 2285570 | 2286604 | + | 915,7055 | -0,404641155 | 0,163349 | -2,47715 | 0,013243 | 0,056754 | 813,7057997 | -0,451544115 | 0,132519 | -3,4074 | 0,000656 | 0,001842 | 1017,705 | -0,369398241 | 0,249704 | -1,47935 | 0,139048 | 0,222496 | 1100,255 | 0,225106 | 0,157382 | 1,430321 | 0,152625 | 0,210241 | 774,0384553 | 0,367277581 | 0,230031 | 1,596643 | 0,110345 | 0,150599 |
| 2051 | RM25_RS10015 | RM25_2026 | Daunorubicin resistance ABC transporter ATPase subunit | 2141 | gene | 2286822 | 2287877 | + | 660,2788 | -1,434057375 | 0,747392 | -1,91875 | 0,055016 | 0,132317 | 348,6830265 | -3,971585209 | 0,369862 | -10,738 | 6,75E-27 | 4,97E-25 | 971,8745 | -0,922043864 | 0,389472 | -2,36742 | 0,017913 | 0,047383 | 1009,646 | 0,888111 | 0,388465 | 2,286206 | 0,022242 | 0,044428 | 351,7754268 | 4,010010609 | 0,399636 | 10,03415 | 1,08E-23 | 3,48E-22 |
| 2052 | RM25_RS10020 | RM25_2027 | ABC-2 type transporter | 2142 | gene | 2287874 | 2288704 | + | 239,618 | -1,323545114 | 0,57172 | -2,31502 | 0,020612 | 0,072836 | 139,8465254 | -2,570077292 | 0,380872 | -6,74788 | 1,5E-11 | 1,7E-10 | 339,3895 | -0,93495544 | 0,379602 | -2,46299 | 0,013778 | 0,038986 | 359,5604 | 0,824303 | 0,367779 | 2,241302 | 0,025006 | 0,048751 | 134,7410593 | 2,533806066 | 0,428735 | 5,909962 | 3,42E-09 | 1,75E-08 |
| 2053 | RM25_RS10025 | RM25_2028 | Sulfate/thiosulfate ABC transporter, ATP-binding protein Cy | 2143 | gene | 2289018 | 2290070 | - | 454,3338 | -0,758721544 | 0,357368 | -2,12308 | 0,033747 | 0,097477 | 330,5957994 | -0,614577483 | 0,226979 | -2,70764 | 0,006776 | 0,014223 | 578,0717 | -0,844543283 | 0,463871 | -1,82064 | 0,068661 | 0,129039 | 598,3979 | 0,826915 | 0,267476 | 3,09155 | 0,001991 | 0,006266 | 332,8834097 | 0,670099562 | 0,439211 | 1,52569 | 0,127087 | 0,171168 |
| 2054 | RM25_RS10030 | RM25_2029 | Sulfate ABC transporter, permease protein CysW | 2144 | gene | 2290067 | 2290876 | - | 322,1007 | -1,303031355 | 0,403607 | -3,22846 | 0,001245 | 0,011549 | 222,1964356 | -1,657168287 | 0,298764 | -5,54674 | 2,91E-08 | 1,95E-07 | 422,0051 | -1,133314073 | 0,403002 | -2,81218 | 0,004921 | 0,017626 | 481,4655 | 0,714549 | 0,28731 | 2,487028 | 0,012882 | 0,029082 | 183,0395414 | 1,310287609 | 0,43334 | 3,023696 | 0,002497 | 0,004901 |
| 2055 | RM25_RS10035 | RM25_2030 | Sulfate ABC transporter, permease protein CysT | 2145 | gene | 2290866 | 2291900 | - | 320,9631 | -1,028946408 | 0,433997 | -2,37086 | 0,017747 | 0,066827 | 209,4552974 | -1,286356941 | 0,329639 | -3,90233 | 9,53E-05 | 0,000326 | 432,471 | -0,912417963 | 0,442378 | -2,06253 | 0,039157 | 0,084665 | 452,4788 | 0,856178 | 0,346112 | 2,473699 | 0,013372 | 0,029965 | 208,2433228 | 1,306810866 | 0,454107 | 2,877757 | 0,004005 | 0,007516 |
| 2056 | RM25_RS10040 | RM25_2031 | SubI | 2146 | gene | 2291897 | 2293003 | - | 461,954 | -1,082205059 | 0,491812 | -2,20044 | 0,027775 | 0,086254 | 281,2495485 | -1,745216809 | 0,30621 | -5,69941 | 1,2E-08 | 8,61E-08 | 642,6584 | -0,827973475 | 0,438336 | -1,8889 | 0,058905 | 0,114331 | 658,2545 | 0,851309 | 0,351265 | 2,423551 | 0,01537 | 0,033428 | 292,4250483 | 1,847546691 | 0,422143 | 4,376587 | 1,21E-05 | 3,5E-05 |
| 2057 | RM25_RS10045 | RM25_2032 | Ferredoxin--nitrite reductase | 2147 | gene | 2293791 | 2295596 | + | 1147,692 | -0,444770048 | 0,221083 | -2,01178 | 0,044244 | 0,114848 | 1162,001454 | -0,215440976 | 0,188897 | -1,14052 | 0,25407 | 0,319383 | 1133,382 | -0,685173978 | 0,36536 | -1,87534 | 0,060746 | 0,117072 | 1390,758 | 0,101823 | 0,258414 | 0,394029 | 0,69356 | 0,740698 | 956,5397428 | -0,304168691 | 0,304701 | -0,99825 | 0,318157 | 0,38182 |
| 2058 | RM25_RS10050 | RM25_2033 | Phosphoadenosine phosphosulfate reductase | 2148 | gene | 2295593 | 2296342 | + | 499,6401 | -0,324758759 | 0,252307 | -1,28716 | 0,19804 | 0,300954 | 565,2425806 | -0,041983665 | 0,24829 | -0,16909 | 0,865725 | 0,895748 | 434,0376 | -0,700161513 | 0,290383 | -2,41117 | 0,015902 | 0,043279 | 584,2734 | -0,1458 | 0,269679 | -0,54063 | 0,588764 | 0,650579 | 436,3428701 | -0,750424536 | 0,269633 | -2,78313 | 0,005384 | 0,009914 |

| | | | | | | | | | | | | | | | | | | | | | | | | | | | | | | | | | | | | | | |
|---|---|---|---|---|---|---|---|---|---|---|---|---|---|---|---|---|---|---|---|---|---|---|---|---|---|---|---|---|---|---|---|---|---|---|---|---|---|---|
| 2059 | RM25_RS10055 | RM25_2034 | Sulfate adenylyltransferase subunit 2 | 2149 | gene | 2296369 | 2297436 | + | 1074,644 | 0,067998893 | 0,384275 | 0,176954 | 0,859545 | 0,907276 | 1442,355615 | 0,333608217 | 0,251117 | 1,328498 | 0,184014 | 0,242302 | 706,9321 | -0,472928876 | 0,307076 | -1,5401 | 0,123535 | 0,202625 | 1105,479 | -0,68544 | 0,298189 | -2,29867 | 0,021524 | 0,043435 | 1081,508242 | -1,440210047 | 0,252116 | -5,71249 | 1,11E-08 | 5,25E-08 |
| 2060 | RM25_RS10060 | RM25_2035 | Sulfate adenylyltransferase subunit 1 | 2150 | gene | 2297439 | 2298788 | + | 1693,822 | 0,422204112 | 0,488464 | 0,86435 | 0,387395 | 0,49814 | 2533,388859 | 0,620832767 | 0,281053 | 2,208953 | 0,027178 | 0,046895 | 854,2561 | -0,14954676 | 0,255142 | -0,58613 | 0,557787 | 0,653431 | 1528,502 | -1,19941 | 0,31996 | -3,74863 | 0,000178 | 0,000815 | 1906,402821 | -1,924540134 | 0,214389 | -8,97686 | 2,79E-19 | 6,02E-18 |
| 2061 | RM25_RS10065 | RM25_2036 | Ribose-5-phosphate isomerase B | 2151 | gene | 2299121 | 2299603 | - | 759,0497 | 0,879447966 | 0,174788 | 5,031524 | 4,87E-07 | 2,25E-05 | 826,9370162 | 1,284797936 | 0,153067 | 8,393692 | 4,71E-17 | 1,01E-15 | 691,1623 | 0,437530983 | 0,180543 | 2,423418 | 0,015375 | 0,042041 | 562,3098 | 0,227792 | 0,154282 | 1,476468 | 0,139818 | 0,19558 | 965,4321212 | -0,565328122 | 0,154711 | -3,6541 | 0,000258 | 0,000611 |
| 2062 | RM25_RS10070 | RM25_2037 | Methyltransferase | 2152 | gene | 2299938 | 2300618 | - | 598,6495 | -0,759987655 | 0,249029 | -3,05181 | 0,002275 | 0,016884 | 705,4869396 | -0,993477053 | 0,268639 | -3,69819 | 0,000217 | 0,00069 | 491,812 | -0,439819361 | 0,244681 | -1,79752 | 0,072253 | 0,133884 | 799,7032 | -0,78929 | 0,320667 | -2,4614 | 0,01384 | 0,030686 | 436,3237561 | -0,177970632 | 0,220653 | -0,80656 | 0,419919 | 0,485886 |
| 2063 | RM25_RS10075 | RM25_2038 | Nodulin 21-like protein | 2153 | gene | 2300629 | 2301426 | - | 1406,438 | -1,360885308 | 0,177565 | -7,66417 | 1,8E-14 | 4,24E-12 | 1469,214556 | -1,53629521 | 0,236138 | -6,50591 | 7,72E-11 | 7,68E-10 | 1343,661 | -1,179791249 | 0,237119 | -4,97553 | 6,51E-07 | 9,41E-06 | 2145,915 | -0,28205 | 0,318288 | -0,88615 | 0,375535 | 0,452293 | 773,1027022 | 0,123872278 | 0,158578 | 0,781143 | 0,434719 | 0,49982 |
| 2064 | RM25_RS10080 | RM25_2039 | Putative insertion element ISCmi2 transposase | 2154 | gene | 2301582 | 2302625 | - | 1,660961 | 0,353319939 | 1,082449 | 0,326408 | 0,744116 | 0,813866 | 1,083941443 | -0,91864966 | 2,795045 | -0,32867 | 0,742405 | 0,790356 | 2,23798 | 1,031628913 | 1,030928 | 1,00068 | 0,316981 | 0,421866 | 1,533933 | -0,07761 | 1,433558 | -0,05414 | 0,956827 | 0,966305 | 1,818267675 | 1,938773234 | 1,855689 | 1,044773 | 0,296128 | 0,357752 |
| 2065 | RM25_RS10085 | RM25_2040 | L-aspartate oxidase | 2155 | gene | 2303136 | 2304725 | + | 973,9457 | 0,124103338 | 0,251733 | 0,492996 | 0,622015 | 0,709284 | 745,8727426 | -0,007834266 | 0,194874 | -0,0402 | 0,967932 | 0,974548 | 1202,019 | 0,206422799 | 0,194712 | 1,060144 | 0,289079 | 0,393395 | 978,9556 | 0,523738 | 0,169692 | 3,086407 | 0,002026 | 0,00635 | 996,993652 | 0,79003585 | 0,205097 | 3,852019 | 0,000117 | 0,000294 |
| 2066 | RM25_RS10090 | RM25_2041 | Putative CoA-substrate-specific enzyme activase | 2156 | gene | 2304776 | 2309152 | - | 13170,63 | 2,299250077 | 0,15179 | 15,14758 | 7,86E-52 | 1,85E-48 | 12738,91095 | 2,119022669 | 0,120727 | 17,55217 | 5,73E-69 | 4,5E-66 | 13602,36 | 2,482894957 | 0,278258 | 8,922982 | 4,54E-19 | 6,69E-17 | 4701,48 | -0,26627 | 0,124046 | -2,14653 | 0,031831 | 0,059344 | 21390,4661 | 0,139451146 | 0,216505 | 0,6441 | 0,51951 | 0,580325 |
| 2067 | RM25_RS10095 | RM25_2042 | Molybdate ABC transporter, permease protein | 2157 | gene | 2309345 | 2311465 | - | 371,7057 | -0,152779621 | 0,539969 | -0,28294 | 0,777222 | 0,840739 | 185,6592362 | -0,800143775 | 0,253255 | -3,15944 | 0,001581 | 0,003989 | 557,7521 | 0,055287149 | 0,469951 | 0,117644 | 0,906349 | 0,936285 | 407,8665 | 1,156992 | 0,295037 | 3,921518 | 8,8E-05 | 0,000437 | 347,6712162 | 2,078643542 | 0,430062 | 4,833352 | 1,34E-06 | 4,57E-06 |
| 2068 | RM25_RS10100 | RM25_2043 | Molybdate ABC transporter, periplasmic molybdate-binding | 2158 | gene | 2311462 | 2312238 | - | 543,4338 | -0,419816552 | 0,206481 | -2,0332 | 0,042032 | 0,111691 | 473,4925921 | -0,402373289 | 0,209726 | -1,91856 | 0,05504 | 0,085509 | 613,375 | -0,434311044 | 0,287085 | -1,51283 | 0,130322 | 0,211404 | 654,2759 | 0,323079 | 0,206777 | 1,562452 | 0,118182 | 0,170369 | 456,7855554 | 0,354159555 | 0,284509 | 1,244809 | 0,213202 | 0,268869 |
| 2069 | RM25_RS10105 | RM25_2044 | Molybdenum cofactor synthesis domain protein | 2159 | gene | 2312458 | 2312964 | + | 747,4588 | 0,129727241 | 0,140136 | 0,925724 | 0,354589 | 0,466388 | 670,0575533 | 0,259300131 | 0,15037 | 1,724409 | 0,084634 | 0,122985 | 824,86 | 0,026568227 | 0,17339 | 0,153228 | 0,878218 | 0,911475 | 752,5016 | 0,366052 | 0,177063 | 2,067359 | 0,0387 | 0,069103 | 764,9579695 | 0,18055845 | 0,133416 | 1,353349 | 0,175944 | 0,228108 |
| 2070 | RM25_RS10110 | RM25_2045 | MobA-like NTP transferase domain protein | 2160 | gene | 2313337 | 2314011 | - | 403,8317 | -0,908488845 | 0,542266 | -1,67536 | 0,093864 | 0,184982 | 186,7257808 | -1,213012878 | 0,264584 | -4,5846 | 4,55E-06 | 2,04E-05 | 620,9376 | -0,821583214 | 0,348575 | -2,35698 | 0,018424 | 0,048467 | 550,6049 | 1,541247 | 0,222337 | 6,932023 | 4,15E-12 | 9,68E-11 | 276,4045028 | 1,9976454 | 0,370868 | 5,386404 | 7,19E-08 | 3,03E-07 |
| 2071 | RM25_RS13310 | RM25_2046 | Molybdenum cofactor biosynthesis protein A | 2161 | gene | 2314110 | 2315285 | + | 960,1256 | -1,26574954 | 0,627866 | -2,01596 | 0,043805 | 0,114094 | 480,758575 | -2,621789319 | 0,306058 | -8,56631 | 1,07E-17 | 2,42E-16 | 1439,493 | -0,924795854 | 0,394428 | -2,34465 | 0,019045 | 0,049656 | 1416,3 | 1,120616 | 0,378084 | 2,96393 | 0,003037 | 0,008904 | 555,6220117 | 2,888393847 | 0,339133 | 8,516997 | 1,64E-17 | 2,74E-16 |
| 2072 | RM25_RS10120 | RM25_2047 | Nitrite extrusion protein | 2162 | gene | 2315429 | 2316787 | + | 2758,51 | -0,427650382 | 0,386208 | -1,1073 | 0,268162 | 0,376001 | 1674,172103 | -0,343787403 | 0,131503 | -2,61429 | 0,008941 | 0,018013 | 3842,847 | -0,464005541 | 0,308515 | -1,50399 | 0,132583 | 0,213893 | 3326,447 | 1,198446 | 0,280657 | 4,270141 | 1,95E-05 | 0,000112 | 2296,475958 | 1,115988873 | 0,175076 | 6,374325 | 1,84E-10 | 1,15E-09 |
| 2073 | RM25_RS10125 | RM25_2048 | Nitrate reductase, alpha subunit | 2163 | gene | 2316821 | 2320288 | + | 4185,78 | -0,104414043 | 0,286051 | -0,36502 | 0,715098 | 0,790565 | 3074,033945 | -0,493485755 | 0,107917 | -4,57284 | 4,81E-06 | 2,14E-05 | 5297,526 | 0,118751964 | 0,217062 | 0,547088 | 0,584318 | 0,675117 | 4570,058 | 0,443019 | 0,187493 | 2,362857 | 0,018135 | 0,037826 | 3942,50962 | 1,09869635 | 0,126392 | 8,692765 | 3,54E-18 | 6,67E-17 |
| 2074 | RM25_RS10130 | RM25_2049 | Transposase IS204/IS1001/IS1096/IS1165 family protein | 2164 | gene | 2320370 | 2321677 | + | 0 | NA | NA | NA | NA | NA | 0 | NA | NA | NA | NA | NA | 0 | NA | NA | NA | NA | NA | 0 | NA | NA | NA | NA | NA | 0 | NA | NA | NA | NA | NA |
| 2075 | RM25_RS10135 | RM25_2050 | Nitrate reductase subunit alpha | 2165 | gene | 2321688 | 2322071 | + | 151,602 | -1,158919101 | 0,486168 | -2,38378 | 0,017136 | 0,065567 | 103,1204279 | -2,256658396 | 0,388827 | -5,80376 | 6,48E-09 | 4,9E-08 | 200,0836 | -0,715158586 | 0,327268 | -2,18524 | 0,028871 | 0,067443 | 220,7285 | 0,471045 | 0,3574 | 1,317977 | 0,187511 | 0,249697 | 92,11169695 | 2,083112444 | 0,403423 | 5,163592 | 2,42E-07 | 9,35E-07 |
| 2076 | RM25_RS10140 | RM25_2051 | Nitrate reductase, beta subunit | 2166 | gene | 2322071 | 2323582 | + | 585,0974 | -1,498583879 | 0,789704 | -1,89765 | 0,057742 | 0,135555 | 293,3711493 | -4,109379769 | 0,42867 | -9,58635 | 9,13E-22 | 3,71E-20 | 876,8236 | -1,018345503 | 0,460367 | -2,21203 | 0,026965 | 0,064458 | 905,8561 | 1,008537 | 0,408843 | 2,466806 | 0,013632 | 0,030313 | 302,3156655 | 4,181488418 | 0,513466 | 8,143655 | 3,84E-16 | 5,48E-15 |
| 2077 | RM25_RS10145 | RM25_2052 | Respiratory nitrate reductase subunit delta narJ | 2167 | gene | 2323582 | 2324406 | + | 254,9174 | -1,315616971 | 0,808737 | -1,62675 | 0,103789 | 0,197124 | 115,1959938 | -4,035829065 | 0,468421 | -8,61581 | 6,94E-18 | 1,64E-16 | 394,6388 | -0,87249501 | 0,444978 | -1,96076 | 0,049907 | 0,100959 | 380,6916 | 1,161195 | 0,387237 | 2,998667 | 0,002712 | 0,008194 | 144,2752946 | 4,398716914 | 0,54292 | 8,101964 | 5,41E-16 | 7,59E-15 |
| 2078 | RM25_RS10150 | RM25_2053 | Respiratory nitrate reductase subunit gamma narI | 2168 | gene | 2324412 | 2325131 | + | 367,5324 | -1,605091902 | 0,719023 | -2,23232 | 0,025594 | 0,082298 | 190,0623249 | -3,67503797 | 0,391644 | -9,38362 | 6,37E-21 | 2,28E-19 | 545,0025 | -1,165980416 | 0,449349 | -2,59482 | 0,009464 | 0,029008 | 579,8108 | 1,023829 | 0,370192 | 2,765672 | 0,005681 | 0,014687 | 179,6038258 | 3,613659109 | 0,499964 | 7,227832 | 4,91E-13 | 4,45E-12 |
| 2079 | RM25_RS10155 | RM25_2054 | ThiS family protein | 2169 | gene | 2325260 | 2326375 | + | 760,1447 | -0,458419364 | 0,271896 | -1,68601 | 0,091794 | 0,182889 | 602,370258 | -0,605369978 | 0,194813 | -3,10745 | 0,001887 | 0,004672 | 917,9191 | -0,365145543 | 0,329776 | -1,10725 | 0,268185 | 0,370305 | 924,5392 | 0,442518 | 0,230156 | 1,922685 | 0,05452 | 0,090943 | 629,7137352 | 0,748696344 | 0,29914 | 2,502828 | 0,012321 | 0,021197 |
| 2080 | RM25_RS10160 | RM25_2055 | Putative molybdenum cofactor biosynthesis protein e2 moa | 2170 | gene | 2326477 | 2326896 | + | 138,7191 | -0,814821664 | 0,463056 | -1,75966 | 0,078466 | 0,165599 | 81,94816819 | -1,24580906 | 0,36564 | -3,4072 | 0,000656 | 0,001842 | 195,4901 | -0,650204471 | 0,356234 | -1,82522 | 0,067968 | 0,12816 | 185,8249 | 0,97279 | 0,326246 | 2,981764 | 0,002866 | 0,008518 | 98,87644671 | 1,644224995 | 0,419005 | 3,924122 | 8,7E-05 | 0,000224 |
| 2081 | RM25_RS10165 | RM25_2056 | Molybdenum cofactor biosynthesis protein moeB1 | 2171 | gene | 2326896 | 2327771 | + | 245,9142 | -0,694755353 | 0,598896 | -1,16006 | 0,246024 | 0,354792 | 120,9240322 | -1,667938614 | 0,309917 | -5,38189 | 7,37E-08 | 4,61E-07 | 370,9043 | -0,419332885 | 0,512436 | -0,81831 | 0,413178 | 0,518287 | 317,7765 | 1,140714 | 0,2896 | 3,938929 | 8,18E-05 | 0,000408 | 185,7968749 | 2,462221033 | 0,52438 | 4,695491 | 2,66E-06 | 8,59E-06 |
| 2082 | RM25_RS10170 | RM25_2057 | Molybdopterin molybdenumtransferase | 2172 | gene | 2327768 | 2329294 | + | 478,5709 | -0,39814332 | 0,499459 | -0,79715 | 0,425364 | 0,533996 | 257,8805615 | -0,922937679 | 0,216544 | -4,26212 | 2,02E-05 | 8,06E-05 | 699,2612 | -0,213662408 | 0,440735 | -0,48479 | 0,627828 | 0,713152 | 568,4843 | 1,092812 | 0,242159 | 4,512793 | 6,4E-06 | 4,14E-05 | 407,5603767 | 1,870105931 | 0,411428 | 4,545401 | 5,48E-06 | 1,67E-05 |
| 2083 | RM25_RS10175 | RM25_2058 | Molybdenum cofactor biosynthesis protein C | 2173 | gene | 2329287 | 2329835 | + | 295,0107 | -0,431815321 | 0,21192 | -2,03763 | 0,041587 | 0,110757 | 253,5254128 | -0,513794787 | 0,176486 | -2,91125 | 0,0036 | 0,00827 | 336,496 | -0,371482702 | 0,310416 | -1,19673 | 0,231414 | 0,329772 | 357,2 | 0,285833 | 0,191592 | 1,491888 | 0,135729 | 0,190765 | 246,5874866 | 0,491641431 | 0,304087 | 1,616776 | 0,105927 | 0,145156 |
| 2084 | RM25_RS10180 | RM25_2059 | Molybdopterin biosynthesis Mog protein | 2174 | gene | 2329890 | 2330474 | - | 260,5764 | -0,1670039 | 0,307757 | -0,54265 | 0,587372 | 0,678978 | 184,9738129 | -0,275097228 | 0,20508 | -1,34142 | 0,179785 | 0,237181 | 336,1791 | -0,110733364 | 0,279474 | -0,39622 | 0,691942 | 0,768933 | 289,3241 | 0,72428 | 0,186856 | 3,876142 | 0,000106 | 0,000512 | 241,2792698 | 0,949842916 | 0,288533 | 3,291977 | 0,000995 | 0,002111 |
| 2085 | RM25_RS10185 | RM25_2060 | Histidine triad protein | 2175 | gene | 2330478 | 2330927 | - | 428,8595 | -0,284656047 | 0,287345 | -0,99064 | 0,321861 | 0,431037 | 321,3968076 | 0,137192282 | 0,157211 | 0,872664 | 0,382846 | 0,451862 | 536,3221 | -0,541303153 | 0,296407 | -1,82621 | 0,067818 | 0,128083 | 495,1213 | 0,999115 | 0,296768 | 3,366654 | 0,000761 | 0,002793 | 378,9142088 | 0,369131159 | 0,153067 | 2,411564 | 0,015884 | 0,026685 |
| 2086 | NA | RM25_2061 | IS3 family element | 2176 | gene | 2331113 | 2331457 | + | 5,620817 | -1,66750462 | 1,159355 | -1,4383 | 0,150348 | 0,250615 | 2,609050345 | -4,617386516 | 2,407 | -1,91832 | 0,055071 | 0,085509 | 8,632584 | -1,155293266 | 0,739881 | -1,56146 | 0,118416 | 0,196443 | 9,052568 | 1,029259 | 0,636126 | 1,618011 | 0,10566 | 0,155494 | 2,616171795 | 4,636702332 | 2,324081 | 1,995069 | 0,046035 | 0,068616 |
| 2087 | RM25_RS10195 | RM25_2062 | Transposase IS6110 | 2177 | gene | 2331490 | 2332377 | + | 64,98415 | 0,433467092 | 0,428432 | 1,011753 | 0,311656 | 0,420237 | 43,0335556 | -0,479513619 | 0,39345 | -1,21874 | 0,222943 | 0,285895 | 86,93474 | 0,895260625 | 0,398559 | 2,246244 | 0,024688 | 0,060426 | 58,29165 | 0,21197 | 0,2785 | 0,761114 | 0,446589 | 0,521352 | 72,51337863 | 1,629643035 | 0,459594 | 3,545834 | 0,000391 | 0,000892 |
| 2088 | NA | RM25_2063 | Hypothetical protein | 2178 | gene | 2332469 | 2332675 | - | 48,89769 | -1,371161081 | 0,512091 | -2,67757 | 0,007416 | 0,038683 | 34,46065394 | -2,648569 | 0,566486 | -4,67544 | 2,93E-06 | 1,38E-05 | 63,33473 | -0,864942994 | 0,299749 | -2,88556 | 0,003907 | 0,01483 | 74,41089 | 0,387678 | 0,338567 | 1,145056 | 0,252186 | 0,32217 | 26,61190628 | 2,232727006 | 0,602028 | 3,708679 | 0,000208 | 0,000501 |
| 2089 | RM25_RS10200 | RM25_2064 | Asp/Glu/Hydantoin racemase | 2179 | gene | 2332659 | 2333435 | - | 178,3397 | -1,235357782 | 0,650239 | -1,89985 | 0,057453 | 0,13528 | 104,934552 | -3,065948586 | 0,457517 | -6,70127 | 2,07E-11 | 2,29E-10 | 251,7448 | -0,719886761 | 0,411975 | -1,7474 | 0,080567 | 0,146108 | 263,583 | 0,663455 | 0,386022 | 1,718698 | 0,085669 | 0,130694 | 104,7763894 | 3,090577216 | 0,524782 | 5,889256 | 3,88E-09 | 1,95E-08 |
| 2090 | RM25_RS10205 | RM25_2065 | D-amino acid dehydrogenase | 2180 | gene | 2333432 | 2334685 | - | 398,5377 | -1,478234756 | 0,696531 | -2,12228 | 0,033814 | 0,097552 | 221,897331 | -3,595580856 | 0,403495 | -8,91109 | 5,05E-19 | 1,32E-17 | 575,178 | -0,979883437 | 0,401924 | -2,43798 | 0,014769 | 0,040715 | 616,1185 | 0,824798 | 0,398602 | 2,069224 | 0,038525 | 0,068895 | 207,741615 | 3,517683839 | 0,448922 | 7,835852 | 4,66E-15 | 5,84E-14 |
| 2091 | RM25_RS10210 | RM25_2066 | Translation initiation inhibitor involved in amino acid or puri | 2181 | gene | 2334682 | 2335065 | - | 75,19199 | -1,286321146 | 0,551541 | -2,33223 | 0,019689 | 0,071175 | 44,01540565 | -2,301935664 | 0,501084 | -4,59391 | 4,35E-06 | 1,96E-05 | 106,3686 | -0,957693139 | 0,369266 | -2,5935 | 0,0095 | 0,029081 | 112,1112 | 0,861305 | 0,360127 | 2,391673 | 0,016772 | 0,035614 | 42,96352684 | 2,281886154 | 0,552609 | 4,129296 | 3,64E-05 | 9,92E-05 |
| 2092 | RM25_RS10220 | RM25_2067 | Glutamine ABC transporter, ATP-binding protein GlnQ | 2182 | gene | 2335073 | 2335876 | - | 243,277 | -1,352806096 | 0,659868 | -2,05011 | 0,040353 | 0,109562 | 138,094085 | -3,192823252 | 0,434879 | -7,34187 | 2,11E-13 | 3,05E-12 | 348,46 | -0,872109333 | 0,388559 | -2,24447 | 0,024802 | 0,060619 | 367,6205 | 0,783454 | 0,367097 | 2,134189 | 0,032827 | 0,060924 | 134,9626578 | 3,178137298 | 0,499079 | 6,368006 | 1,92E-10 | 1,19E-09 |
| 2093 | RM25_RS10215 | RM25_2068 | Lysine-arginine-ornithine-binding periplasmic protein | 2183 | gene | 2335946 | 2336806 | - | 283,1759 | -1,534034034 | 0,7189 | -2,13386 | 0,032854 | 0,095934 | 159,8514904 | -3,87869118 | 0,447154 | -8,67418 | 4,17E-18 | 1E-16 | 406,5003 | -1,005582787 | 0,352834 | -2,85001 | 0,004372 | 0,016227 | 441,855 | 0,786376 | 0,398654 | 1,97258 | 0,048543 | 0,083091 | 143,1345651 | 3,733037432 | 0,452686 | 8,246409 | 1,63E-16 | 2,4E-15 |
| 2094 | RM25_RS10225 | RM25_2069 | Putative ABC transporter membrane protein | 2184 | gene | 2336873 | 2337529 | - | 304,1433 | -1,64279263 | 0,81863 | -2,00676 | 0,044775 | 0,115973 | 152,3541796 | -4,659054188 | 0,474595 | -9,8169 | 9,52E-23 | 4,32E-21 | 455,9325 | -1,149093089 | 0,41433 | -2,77338 | 0,005548 | 0,019429 | 482,8318 | 1,02591 | 0,417451 | 2,457558 | 0,013989 | 0,03093 | 145,6094546 | 4,614421665 | 0,511476 | 9,021778 | 1,85E-19 | 4,19E-18 |
| 2095 | RM25_RS10230 | RM25_2070 | Amino acid ABC transporter, permease protein | 2185 | gene | 2337526 | 2338164 | - | 262,0032 | -1,638583708 | 0,626423 | -2,61578 | 0,008902 | 0,043175 | 153,886799 | -3,375423096 | 0,343589 | -9,82401 | 8,87E-23 | 4,1E-21 | 370,1196 | -1,172152002 | 0,373072 | -3,14189 | 0,001679 | 0,007465 | 416,6722 | 0,80199 | 0,284955 | 2,814444 | 0,004886 | 0,012926 | 125,4673016 | 3,073843686 | 0,452709 | 6,789885 | 1,12E-11 | 8,24E-11 |
| 2096 | RM25_RS10235 | RM25_2071 | Dehydrogenase | 2186 | gene | 2338281 | 2339327 | - | 272,662 | -1,101137364 | 0,840485 | -1,31012 | 0,190155 | 0,294091 | 126,5607703 | -4,127472637 | 0,505527 | -8,16469 | 3,22E-16 | 5,98E-15 | 418,7632 | -0,600430324 | 0,507323 | -1,18353 | 0,236601 | 0,33554 | 390,3752 | 0,996409 | 0,433154 | 2,300357 | 0,021428 | 0,043315 | 171,6295283 | 4,608303209 | 0,603245 | 7,639196 | 2,19E-14 | 2,47E-13 |
| 2097 | RM25_RS10240 | RM25_2072 | D-isomer specific 2-hydroxyacid dehydrogenase NAD-bindin | 2187 | gene | 2339331 | 2340410 | - | 317,2006 | -1,552017754 | 0,730204 | -2,12546 | 0,033548 | 0,097142 | 179,9162209 | -3,969346934 | 0,431772 | -9,19316 | 3,81E-20 | 1,23E-18 | 454,4849 | -1,014683394 | 0,431512 | -2,35146 | 0,0187 | 0,048919 | 497,053 | 0,771143 | 0,417081 | 1,848906 | 0,064471 | 0,104655 | 159,2942359 | 3,806511576 | 0,490216 | 7,764966 | 8,17E-15 | 9,67E-14 |
| 2098 | RM25_RS10245 | RM25_2073 | 4-aminobutyrate transaminase | 2188 | gene | 2340407 | 2341741 | - | 393,8456 | -1,479522058 | 0,73241 | -2,02007 | 0,043376 | 0,113818 | 213,6947455 | -3,860631613 | 0,386249 | -9,99519 | 1,6E-23 | 8,98E-22 | 573,9965 | -0,969274026 | 0,451149 | -2,14846 | 0,031678 | 0,072349 | 608,3455 | 0,856501 | 0,377343 | 2,269819 | 0,023219 | 0,046083 | 205,497859 | 3,823974103 | 0,487267 | 7,847801 | 4,23E-15 | 5,34E-14 |
| 2099 | RM25_RS10250 | RM25_2074 | Transcriptional regulator, PucR family | 2189 | gene | 2341921 | 2343417 | + | 673,4641 | -0,761384862 | 0,457731 | -1,66339 | 0,096234 | 0,188706 | 390,6037914 | -0,944338981 | 0,221778 | -4,25805 | 2,06E-05 | 8,2E-05 | 956,3244 | -0,690135055 | 0,443952 | -1,55453 | 0,120059 | 0,198721 | 886,9622 | 1,133068 | 0,257042 | 4,408109 | 1,04E-05 | 6,36E-05 | 493,2382537 | 1,460287906 | 0,417254 | 3,499761 | 0,000466 | 0,001051 |
| 2100 | RM25_RS13315 | RM25_2075 | Hypothetical protein | 2190 | gene | 2343920 | 2344048 | + | 123,8896 | 0,245161585 | 0,287483 | 0,852787 | 0,393777 | 0,504694 | 97,01027077 | 0,198413037 | 0,349299 | 0,568032 | 0,570013 | 0,631948 | 150,769 | 0,274321941 | 0,273746 | 1,002103 | 0,316294 | 0,421189 | 118,6792 | 0,549577 | 0,31432 | 1,748466 | 0,080383 | 0,123751 | 131,9087808 | 0,674801407 | 0,294721 | 2,289628 | 0,022043 | 0,035955 |
| 2101 | RM25_RS10255 | RM25_2076 | Hypothetical protein | 2191 | gene | 2344135 | 2346018 | - | 719,5701 | -1,286218006 | 0,719303 | -1,78814 | 0,073753 | 0,159751 | 355,4075991 | -3,384326066 | 0,368947 | -9,17293 | 4,6E-20 | 1,43E-18 | 1083,733 | -0,845989806 | 0,414964 | -2,03871 | 0,041479 | 0,088397 | 1069,692 | 1,031055 | 0,391969 | 2,630447 | 0,008527 | 0,02072 | 413,1267488 | 3,646654419 | 0,420856 | 8,66485 | 4,52E-18 | 8,26E-17 |
| 2102 | RM25_RS10260 | RM25_2077 | Histidine kinase | 2192 | gene | 2346392 | 2347801 | + | 507,7522 | 0,934336185 | 0,280297 | 3,333378 | 0,000858 | 0,009121 | 578,6203688 | 1,510326877 | 0,164642 | 9,173414 | 4,58E-20 | 1,43E-18 | 436,8839 | 0,271526303 | 0,357926 | 0,758611 | 0,448085 | 0,552084 | 366,8128 | 0,335705 | 0,201178 | 1,668699 | 0,095177 | 0,142614 | 656,1621892 | -0,835676322 | 0,324378 | -2,57624 | 0,009988 | 0,017455 |
| 2103 | RM25_RS10265 | RM25_2078 | Two-component system response regulator | 2193 | gene | 2347835 | 2348497 | + | 1086,169 | 0,197599361 | 0,239637 | 0,824577 | 0,409612 | 0,520538 | 1138,767942 | 0,78382883 | 0,16464 | 4,760869 | 1,93E-06 | 9,33E-06 | 1033,569 | -0,42781244 | 0,260073 | -1,64497 | 0,099976 | 0,172632 | 1067,165 | 0,449057 | 0,266946 | 1,682203 | 0,09253 | 0,139089 | 1137,066252 | -0,721503567 | 0,140428 | -5,13788 | 2,78E-07 | 1,06E-06 |
| 2104 | RM25_RS10270 | RM25_2079 | hypothetical protein | 2194 | gene | 2348851 | 2349816 | + | 357,5797 | -1,141722797 | 0,814606 | -1,40156 | 0,161046 | 0,26415 | 160,3258188 | -3,914737096 | 0,443427 | -8,82837 | 1,06E-18 | 2,66E-17 | 554,8335 | -0,682856131 | 0,465344 | -1,46742 | 0,142262 | 0,226255 | 515,1995 | 1,114363 | 0,382925 | 2,910135 | 0,003613 | 0,01021 | 220,547402 | 4,421453777 | 0,538523 | 8,210342 | 2,21E-16 | 3,21E-15 |
| 2105 | RM25_RS10275 | RM25_2080 | Transporter, major facilitator family protein | 2195 | gene | 2350150 | 2351577 | + | 1093,961 | 0,023650892 | 0,11761 | 0,201096 | 0,840624 | 0,894919 | 1184,34423 | -0,088995388 | 0,127151 | -0,69992 | 0,483977 | 0,551346 | 1003,578 | 0,156130434 | 0,162866 | 0,958645 | 0,337738 | 0,443975 | 1150,182 | -0,42578 | 0,163061 | -2,61116 | 0,009023 | 0,021787 | 1081,417717 | -0,124293922 | 0,126485 | -0,98268 | 0,325765 | 0,389959 |
| 2106 | RM25_RS10280 | RM25_2081 | Putative antibiotic transporter | 2196 | gene | 2351577 | 2352167 | + | 376,0791 | -0,558854959 | 0,185119 | -3,01889 | 0,002537 | 0,018399 | 332,7263183 | -0,541897183 | 0,21666 | -2,50114 | 0,012379 | 0,02378 | 419,4318 | -0,575007918 | 0,251995 | -2,28182 | 0,0225 | 0,056322 | 471,8875 | 0,278969 | 0,253228 | 1,10165 | 0,270614 | 0,343477 | 298,5839145 | 0,309767273 | 0,227541 | 1,361369 | 0,173397 | 0,22555 |
| 2108 | RM25_RS12040 | RM25_2082 | Hypothetical protein | 2197 | gene | 2352178 | 2355888 | - | 1326,725 | -1,072449563 | 0,63998 | -1,67576 | 0,093786 | 0,184982 | 657,3957895 | -2,479667892 | 0,301693 | -8,21917 | 2,05E-16 | 3,99E-15 | 1996,054 | -0,717020202 | 0,45549 | -1,57417 | 0,115448 | 0,192849 | 1882,718 | 1,085046 | 0,361189 | 3,004097 | 0,002664 | 0,008085 | 845,2153279 | 2,925076954 | 0,416073 | 7,030194 | 2,06E-12 | 1,71E-11 |
| 2109 | RM25_RS10290 | RM25_2083 | Putative membrane protein | 2198 | gene | 2355885 | 2356283 | - | 134,1651 | -1,295641547 | 0,938392 | -1,3807 | 0,16737 | 0,269645 | 52,80621109 | -4,895725677 | 0,677728 | -7,22373 | 5,06E-13 | 6,93E-12 | 215,524 | -0,877694498 | 0,490958 | -1,78772 | 0,073822 | 0,13597 | 199,2039 | 1,374741 | 0,380696 | 3,611129 | 0,000305 | 0,001306 | 76,79514709 | 5,477338563 | 0,774902 | 7,06843 | 1,57E-12 | 1,31E-11 |
| 2110 | RM25_RS12045 | RM25_2084 | Hypothetical protein | 2199 | gene | 2356508 | 2356930 | - | 83,32207 | -1,16562442 | 0,776332 | -1,50145 | 0,133239 | 0,231838 | 34,88808632 | -3,189042117 | 0,561064 | -5,68392 | 1,32E-08 | 9,4E-08 | 131,7561 | -0,8086323 | 0,53847 | -1,50172 | 0,133169 | 0,214691 | 120,4375 | 1,338691 | 0,337187 | 3,970175 | 7,18E-05 | 0,000366 | 50,83536778 | 3,805072027 | 0,731839 | 5,199333 | 2E-07 | 7,77E-07 |
| 2111 | RM25_RS10300 | RM25_2085 | Hypothetical protein | 2200 | gene | 2356905 | 2357129 | - | 65,50434 | -1,264943945 | 0,933241 | -1,35543 | 0,17528 | 0,278957 | 27,2953088 | -4,461934023 | 0,840194 | -5,3106 | 1,09E-07 | 6,57E-07 | 103,7134 | -0,833363787 | 0,4443 | -1,87568 | 0,060699 | 0,117072 | 97,06167 | 1,271971 | 0,351088 | 3,622943 | 0,000291 | 0,00125 | 37,98438048 | 4,968750385 | 0,92665 | 5,362056 | 8,23E-08 | 3,41E-07 |
| 2112 | RM25_RS10305 | RM25_2086 | Type II secretion system F domain protein | 2201 | gene | 2357214 | 2358119 | - | 366,5738 | -1,418497921 | 0,87101 | -1,62857 | 0,103405 | 0,196711 | 164,7182626 | -4,628247547 | 0,483468 | -9,57302 | 1,04E-21 | 4,08E-20 | 568,4293 | -0,95825643 | 0,457897 | -2,09273 | 0,036373 | 0,080123 | 558,6028 | 1,17193 | 0,382878 | 3,060841 | 0,002207 | 0,006845 | 197,3080579 | 4,920483018 | 0,583013 | 8,439744 | 3,18E-17 | 5,13E-16 |
| 2113 | RM25_RS10310 | RM25_2087 | Bacterial type II secretion system domain protein F | 2202 | gene | 2358112 | 2359002 | - | 250,2222 | -1,125441814 | 0,918357 | -1,2255 | 0,220389 | 0,326087 | 106,0989493 | -4,879406323 | 0,541611 | -9,00906 | 2,08E-19 | 5,7E-18 | 394,3454 | -0,64598773 | 0,47357 | -1,36408 | 0,172542 | 0,263566 | 359,0437 | 1,15723 | 0,385442 | 3,002349 | 0,002679 | 0,008116 | 155,4725422 | 5,466982837 | 0,632505 | 8,643379 | 5,46E-18 | 9,89E-17 |
| 2114 | RM25_RS10315 | RM25_2088 | Secretion system protein | 2203 | gene | 2358999 | 2360774 | - | 622,182 | -1,362978739 | 0,901374 | -1,51211 | 0,130505 | 0,22904 | 275,3418599 | -5,00460671 | 0,444588 | -11,2567 | 2,15E-29 | 2,2E-27 | 969,0222 | -0,887626859 | 0,486225 | -1,82555 | 0,067919 | 0,12816 | 938,3509 | 1,162129 | 0,432511 | 2,686936 | 0,007211 | 0,018006 | 344,6207093 | 5,361042383 | 0,52162 | 10,27767 | 8,88E-25 | 3,32E-23 |
| 2115 | RM25_RS12050 | RM25_2089 | Hypothetical protein | 2204 | gene | 2360771 | 2361808 | - | 321,1086 | -0,791930611 | 0,953092 | -0,83091 | 0,406026 | 0,51702 | 105,6866149 | -4,632366761 | 0,467785 | -9,90277 | 4,05E-23 | 2,03E-21 | 536,5306 | -0,404141645 | 0,501461 | -0,80593 | 0,420284 | 0,524883 | 424,2513 | 1,52157 | 0,326684 | 4,657615 | 3,2E-06 | 2,21E-05 | 232,8309119 | 5,818475386 | 0,603885 | 9,635079 | 5,68E-22 | 1,56E-20 |
| 2116 | RM25_RS10325 | RM25_2090 | SAF domain protein | 2205 | gene | 2361808 | 2362323 | - | 179,7328 | -1,378907241 | 0,762156 | -1,80922 | 0,070417 | 0,15425 | 84,71919484 | -3,629580325 | 0,4881 | -7,43614 | 1,04E-13 | 1,56E-12 | 274,7463 | -0,956173372 | 0,471796 | -2,02667 | 0,042696 | 0,089925 | 271,8425 | 1,136318 | 0,393232 | 2,889691 | 0,003856 | 0,010744 | 98,5841694 | 3,888286936 | 0,583926 | 6,658875 | 2,76E-11 | 1,93E-10 |
| 2117 | RM25_RS10330 | RM25_2091 | Transposase IS204/IS1001/IS1096/IS1165 family protein | 2206 | gene | 2362730 | 2363806 | + | 35,65382 | -1,951046297 | 1,060831 | -1,83917 | 0,065891 | 0,147487 | 18,09047871 | -7,406909358 | 1,37672 | -5,38011 | 7,44E-08 | 4,64E-07 | 53,21716 | -1,396047202 | 0,41573 | -3,35807 | 0,000785 | 0,004013 | 59,43759 | 1,003704 | 0,430775 | 2,329997 | 0,019806 | 0,04063 | 14,39750404 | 7,096379779 | 1,462659 | 4,851698 | 1,22E-06 | 4,19E-06 |
| 2118 | RM25_RS10335 | RM25_2092 | YghA protein | 2207 | gene | 2364509 | 2365198 | - | 565,9589 | -0,01263493 | 0,144559 | -0,0874 | 0,930351 | 0,952579 | 600,8559312 | -0,215658188 | 0,177815 | -1,21282 | 0,225198 | 0,288004 | 531,0618 | 0,214405558 | 0,196754 | 1,089716 | 0,275838 | 0,378876 | 603,0718 | -0,45804 | 0,202443 | -2,26257 | 0,023663 | 0,046868 | 551,74177 | 0,027665861 | 0,170409 | 0,16235 | 0,87103 | 0,895344 |
| 2119 | RM25_RS12055 | RM25_2093 | Transcriptional Regulator, TetR family | 2208 | gene | 2365378 | 2365986 | + | 498,9562 | -0,610895521 | 0,247817 | -2,46511 | 0,013697 | 0,057651 | 386,1780291 | -0,593601986 | 0,205341 | -2,89081 | 0,003843 | 0,008705 | 611,7344 | -0,624492717 | 0,222036 | -2,81258 | 0,004915 | 0,017626 | 634,7158 | 0,60928 | 0,238146 | 2,558433 | 0,010515 | 0,024659 | 387,5868438 | 0,640679638 | 0,211945 | 3,022865 | 0,002504 | 0,004906 |
| 2120 | RM25_RS10345 | RM25_2094 | Hypothetical protein | 2209 | gene | 2366132 | 2366356 | + | 395,3135 | -1,247144359 | 0,51708 | -2,4119 | 0,01587 | 0,062895 | 178,4346822 | 0,000499266 | 0,255506 | 0,001954 | 0,998441 | 0,998865 | 612,1922 | -1,689830508 | 0,288625 | -5,85477 | 4,78E-09 | 1,15E-07 | 582,1271 | 2,333202 | 0,33148 | 7,038745 | 1,94E-12 | 4,86E-11 | 229,221682 | 0,683291482 | 0,198287 | 3,445979 | 0,000569 | 0,001265 |
| 2121 | RM25_RS10350 | RM25_2095 | ABC-type Fe3+-siderophores transport systems | 2210 | gene | 2366416 | 2367225 | - | 750,5303 | 0,304078206 | 0,09516 | 3,195443 | 0,001396 | 0,012325 | 768,447017 | 0,263514955 | 0,139651 | 1,886959 | 0,059166 | 0,090613 | 732,6135 | 0,346823456 | 0,195627 | 1,772881 | 0,076249 | 0,139749 | 710,7295 | -0,17444 | 0,164435 | -1,06082 | 0,288773 | 0,36262 | 811,5191085 | -0,043873263 | 0,147058 | -0,29834 | 0,765444 | 0,806866 |
| 2122 | RM25_RS10355 | RM25_2096 | Hypothetical protein | 2211 | gene | 2367281 | 2367886 | - | 958,8188 | 0,578036559 | 0,235384 | 2,455718 | 0,01406 | 0,058243 | 1154,239608 | 0,52849056 | 0,195778 | 2,699436 | 0,006946 | 0,0145 | 763,3981 | 0,656148128 | 0,209662 | 3,12955 | 0,001751 | 0,007699 | 815,6557 | -0,72408 | 0,247986 | -2,91986 | 0,003502 | 0,009996 | 1125,112897 | -0,556543058 | 0,129163 | -4,30884 | 1,64E-05 | 4,7E-05 |
| 2123 | RM25_RS10360 | RM25_2097 | Hypothetical protein | 2212 | gene | 2368005 | 2368496 | - | 445,7277 | 0,062471325 | 0,114453 | 0,545824 | 0,585187 | 0,676784 | 473,2773833 | -0,048899009 | 0,154338 | -0,31683 | 0,751372 | 0,797381 | 418,178 | 0,187142947 | 0,184625 | 1,013639 | 0,310755 | 0,41475 | 460,9757 | -0,35906 | 0,162069 | -2,21547 | 0,026728 | 0,051427 | 446,2504499 | -0,071474236 | 0,143911 | -0,49665 | 0,619433 | 0,676366 |
| 2124 | RM25_RS10365 | RM25_2098 | Metal cation transporting P-type ATPase ctpG | 2213 | gene | 2368548 | 2370662 | - | 828,8418 | 0,058287216 | 0,3399 | 0,171483 | 0,863844 | 0,908558 | 560,396524 | 0,02521853 | 0,164153 | 0,153628 | 0,877903 | 0,905312 | 1097,287 | 0,074572975 | 0,325547 | 0,22907 | 0,818815 | 0,869737 | 849,9648 | 0,880252 | 0,237387 | 3,708092 | 0,000209 | 0,000929 | 831,215736 | 0,993870572 | 0,264789 | 3,753442 | 0,000174 | 0,000425 |
| 2125 | RM25_RS10370 | RM25_2099 | Transcriptional regulator ArsR | 2214 | gene | 2370659 | 2371006 | - | 263,4147 | 0,165889638 | 0,127836 | 1,297672 | 0,1944 | 0,298114 | 272,820921 | 0,162629054 | 0,200145 | 0,812556 | 0,416473 | 0,484514 | 254,0086 | 0,171840383 | 0,221659 | 0,775248 | 0,438193 | 0,542164 | 262,576 | -0,17484 | 0,193189 | -0,90502 | 0,365452 | 0,440824 | 272,990437 | -0,109704199 | 0,222661 | -0,4927 | 0,622227 | 0,678349 |
| 2126 | RM25_RS10375 | RM25_2100 | N-acetyltransferase yafP | 2215 | gene | 2371138 | 2371683 | + | 626,2736 | 0,071097863 | 0,132769 | 0,535502 | 0,592303 | 0,682669 | 683,310238 | 0,13173918 | 0,144717 | 0,910325 | 0,362651 | 0,430831 | 569,237 | 0,001415159 | 0,203397 | 0,006958 | 0,994449 | 0,996621 | 645,0225 | -0,25782 | 0,184694 | -1,39594 | 0,162732 | 0,221455 | 629,6760012 | -0,330996262 | 0,141305 | -2,34243 | 0,019159 | 0,031623 |
| 2127 | RM25_RS10380 | RM25_2101 | Hypothetical protein | 2216 | gene | 2371754 | 2372308 | - | 477,2705 | 0,257270714 | 0,194757 | 1,320983 | 0,186507 | 0,291031 | 402,6973896 | 0,352247457 | 0,18547 | 1,899219 | 0,057536 | 0,088404 | 551,8437 | 0,18674065 | 0,218683 | 0,853935 | 0,393141 | 0,499264 | 458,2092 | 0,478477 | 0,197603 | 2,42141 | 0,01546 | 0,033585 | 509,0100022 | 0,367799416 | 0,206301 | 1,782825 | 0,074615 | 0,105817 |
| 2128 | RM25_RS10385 | RM25_2102 | Transcriptional regulator, MerR | 2217 | gene | 2372391 | 2372735 | - | 99,61215 | -1,500097657 | 0,810665 | -1,85045 | 0,064248 | 0,145232 | 49,87782883 | -4,249256637 | 0,583779 | -7,27888 | 3,37E-13 | 4,72E-12 | 149,3465 | -1,011023333 | 0,515712 | -1,96044 | 0,049944 | 0,100959 | 154,8693 | 0,993425 | 0,320319 | 3,101364 | 0,001926 | 0,006103 | 51,46828493 | 4,324099318 | 0,776874 | 5,566023 | 2,61E-08 | 1,17E-07 |
| 2129 | RM25_RS10390 | RM25_2103 | Protein AraJ | 2218 | gene | 2372773 | 2374020 | + | 367,866 | -1,318235089 | 0,783813 | -1,68182 | 0,092603 | 0,183704 | 167,347031 | -3,823302647 | 0,415728 | -9,19664 | 3,69E-20 | 1,23E-18 | 568,3849 | -0,887008617 | 0,43684 | -2,03051 | 0,042304 | 0,089347 | 549,8811 | 1,167563 | 0,380294 | 3,070163 | 0,002139 | 0,00667 | 208,0435509 | 4,18004152 | 0,492323 | 8,490439 | 2,06E-17 | 3,39E-16 |
| 2130 | RM25_RS10395 | RM25_2104 | Glutamine amidotransferase class-I | 2219 | gene | 2374046 | 2374735 | + | 335,9138 | -0,264779262 | 0,511405 | -0,51775 | 0,604634 | 0,694842 | 166,5376223 | -0,649542214 | 0,275655 | -2,35636 | 0,018455 | 0,033486 | 505,2899 | -0,143071584 | 0,357689 | -0,39999 | 0,689165 | 0,76657 | 382,472 | 1,320284 | 0,291443 | 4,530168 | 5,89E-06 | 3,84E-05 | 300,3731134 | 1,889666795 | 0,33512 | 5,638769 | 1,71E-08 | 7,91E-08 |
| 2131 | RM25_RS10400 | RM25_2105 | Ribose-phosphate diphosphokinase | 2220 | gene | 2374819 | 2375799 | + | 1884,801 | 0,395913416 | 0,205527 | 1,926333 | 0,054063 | 0,130827 | 2127,92653 | 0,540176423 | 0,166672 | 3,240947 | 0,001191 | 0,003134 | 1641,675 | 0,212784713 | 0,299482 | 0,710509 | 0,477388 | 0,579221 | 1726,612 | -0,24639 | 0,287409 | -0,8573 | 0,391282 | 0,466962 | 2096,480984 | -0,531993866 | 0,180832 | -2,94192 | 0,003262 | 0,00624 |
| 2132 | RM25_RS10405 | RM25_2106 | Brp/Blh family beta-carotene 15, 15'-monooxygenase | 2221 | gene | 2375950 | 2376561 | - | 1476,934 | 0,194508064 | 0,300301 | 0,647709 | 0,517173 | 0,613785 | 1815,366598 | 0,439803426 | 0,220972 | 1,99031 | 0,046557 | 0,074901 | 1138,502 | -0,190059098 | 0,333466 | -0,56995 | 0,568711 | 0,662319 | 1461,281 | -0,39532 | 0,363065 | -1,08884 | 0,276225 | 0,349658 | 1544,605747 | -0,988213875 | 0,164377 | -6,01186 | 1,83E-09 | 9,82E-09 |
| 2133 | RM25_RS10410 | RM25_2107 | 1D-myo-inositol 2-acetamido-2-deoxy-alpha-D-glucopyranos | 2222 | gene | 2376732 | 2377550 | - | 1722,801 | 0,477280619 | 0,209606 | 2,27704 | 0,022784 | 0,077571 | 1910,41038 | 0,313796005 | 0,1951 | 1,60839 | 0,10775 | 0,151985 | 1535,191 | 0,685760576 | 0,286929 | 2,390002 | 0,016848 | 0,045229 | 1527,119 | -0,58591 | 0,281655 | -2,08022 | 0,037505 | 0,067481 | 1961,367635 | -0,1761067 | 0,183095 | -0,96183 | 0,336135 | 0,399934 |
| 2134 | RM25_RS10415 | RM25_2108 | D-inositol-3-phosphate glycosyltransferase | 2223 | gene | 2377547 | 2378776 | - | 1453,892 | 0,450812056 | 0,171568 | 2,627599 | 0,008599 | 0,042478 | 1504,100586 | 0,645994901 | 0,237157 | 2,723911 | 0,006451 | 0,013662 | 1403,683 | 0,247148533 | 0,2039 | 1,212107 | 0,225471 | 0,322706 | 1296,008 | 0,081671 | 0,293596 | 0,278174 | 0,780878 | 0,823135 | 1645,241577 | -0,278040316 | 0,12316 | -2,25756 | 0,023973 | 0,038596 |
| 2135 | RM25_RS10420 | RM25_2109 | Hypothetical protein | 2224 | gene | 2378949 | 2379806 | + | 1015,124 | -0,096984534 | 0,112091 | -0,86523 | 0,386912 | 0,49779 | 960,0926886 | 0,013625177 | 0,158724 | 0,085842 | 0,931592 | 0,94797 | 1070,156 | -0,195825249 | 0,170684 | -1,1473 | 0,25126 | 0,351674 | 1107,624 | 0,201794 | 0,192828 | 1,046496 | 0,295332 | 0,369282 | 961,3132027 | 0,041178093 | 0,136632 | 0,30138 | 0,763125 | 0,804781 |
| 2136 | RM25_RS10425 | RM25_2110 | Hypothetical protein | 2225 | gene | 2379807 | 2380547 | - | 181,3687 | -0,858343235 | 0,753484 | -1,13917 | 0,254634 | 0,362423 | 80,34775279 | -3,041142926 | 0,480945 | -6,32326 | 2,56E-10 | 2,35E-09 | 282,3897 | -0,434329598 | 0,43427 | -1,00014 | 0,317244 | 0,421978 | 244,6798 | 1,109765 | 0,337149 | 3,291615 | 0,000996 | 0,00352 | 127,3444064 | 3,787978661 | 0,570194 | 6,64331 | 3,07E-11 | 2,13E-10 |
| 2137 | RM25_RS10430 | RM25_2111 | Transposase | 2226 | gene | 2380661 | 2382040 | + | 47,1489 | -0,079932915 | 0,439541 | -0,18186 | 0,855696 | 0,90513 | 28,08668454 | -0,477040739 | 0,457901 | -1,0418 | 0,297505 | 0,366172 | 66,21112 | 0,076804104 | 0,330092 | 0,232675 | 0,816014 | 0,867673 | 50,78648 | 0,912717 | 0,30498 | 2,992709 | 0,002765 | 0,008302 | 45,0171521 | 1,533613576 | 0,457879 | 3,349385 | 0,00081 | 0,001761 |
| 2138 | RM25_RS10435 | RM25_2112 | Hypothetical protein | 2227 | gene | 2382045 | 2382377 | - | 126,5676 | -0,82112162 | 0,38123 | -2,15388 | 0,03125 | 0,092882 | 87,04809612 | -0,82561703 | 0,351853 | -2,34648 | 0,018952 | 0,034177 | 166,0871 | -0,824334212 | 0,399736 | -2,06219 | 0,039189 | 0,084665 | 169,2428 | 0,861865 | 0,338579 | 2,545537 | 0,010911 | 0,025437 | 89,9754957 | 0,933054674 | 0,435282 | 2,143563 | 0,032068 | 0,050089 |
| 2139 | RM25_RS10440 | RM25_2113 | Aminoglycoside adenyltransferase | 2228 | gene | 2382370 | 2383203 | - | 1071,458 | -0,310812583 | 0,245328 | -1,26693 | 0,205181 | 0,309214 | 939,6938961 | 0,269410688 | 0,154369 | 1,745238 | 0,080943 | 0,118205 | 1203,222 | -0,777345319 | 0,292337 | -2,65907 | 0,007836 | 0,025544 | 1249,68 | 0,782531 | 0,303656 | 2,577035 | 0,009965 | 0,023654 | 937,6205728 | -0,220164104 | 0,127212 | -1,73069 | 0,083507 | 0,117166 |
| 2140 | RM25_RS10445 | RM25_2114 | Aldose 1-epimerase | 2229 | gene | 2383318 | 2384214 | + | 1571,683 | 0,273230212 | 0,248422 | 1,099862 | 0,271392 | 0,379852 | 1745,528063 | 0,668841451 | 0,182004 | 3,674877 | 0,000238 | 0,00075 | 1397,838 | -0,206026311 | 0,333828 | -0,61716 | 0,537127 | 0,63459 | 1505,15 | 0,102768 | 0,333973 | 0,307713 | 0,7583 | 0,801486 | 1685,325866 | -0,735433069 | 0,176417 | -4,16872 | 3,06E-05 | 8,42E-05 |
| 2141 | RM25_RS10450 | RM25_2115 | Putative excinuclease ABC, A subunit | 2230 | gene | 2384290 | 2386572 | - | 2086,376 | 0,152556754 | 0,093439 | 1,632682 | 0,102536 | 0,19569 | 2140,342277 | 0,287232012 | 0,124065 | 2,315177 | 0,020603 | 0,036678 | 2032,41 | 0,013160651 | 0,17168 | 0,076658 | 0,938896 | 0,960494 | 2082,945 | 0,010032 | 0,144162 | 0,069587 | 0,944522 | 0,956699 | 2153,571569 | -0,213070712 | 0,106791 | -1,99522 | 0,046019 | 0,068616 |
| 2142 | RM25_RS10455 | RM25_2116 | Zinc finger protein | 2231 | gene | 2386748 | 2387059 | + | 170,7582 | -0,885306933 | 0,416004 | -2,12812 | 0,033327 | 0,096739 | 115,2114004 | -1,427383411 | 0,384418 | -3,7131 | 0,000205 | 0,000652 | 226,305 | -0,637002825 | 0,297105 | -2,14403 | 0,03203 | 0,072872 | 232,9172 | 0,641988 | 0,37269 | 1,722578 | 0,084965 | 0,129872 | 117,8082279 | 1,500707545 | 0,345226 | 4,347024 | 1,38E-05 | 3,99E-05 |
| 2143 | RM25_RS10460 | RM25_2117 | Transketolase | 2232 | gene | 2387340 | 2389406 | - | 8046,647 | 0,867859607 | 0,226564 | 3,830521 | 0,000128 | 0,00205 | 9013,113834 | 0,755859033 | 0,185936 | 4,065146 | 4,8E-05 | 0,000177 | 7080,18 | 1,014480243 | 0,326789 | 3,104386 | 0,001907 | 0,008307 | 6041,833 | -0,56436 | 0,303772 | -1,85785 | 0,063191 | 0,102859 | 10169,74482 | -0,27316765 | 0,204541 | -1,33552 | 0,181708 | 0,233908 |
| 2144 | RM25_RS10465 | RM25_2118 | Ribulokinase | 2233 | gene | 2389526 | 2391310 | - | 1231,106 | -0,209034345 | 0,119596 | -1,74784 | 0,080492 | 0,167893 | 1142,13506 | -0,232119327 | 0,114212 | -2,03236 | 0,042117 | 0,068985 | 1320,077 | -0,189815387 | 0,213757 | -0,888 | 0,374543 | 0,480304 | 1391,978 | 0,128067 | 0,155013 | 0,826167 | 0,408709 | 0,484815 | 1121,016594 | 0,228884334 | 0,176714 | 1,295227 | 0,195242 | 0,249694 |
| 2145 | RM25_RS10470 | RM25_2119 | Sn-glycerol-1-phosphate dehydrogenase | 2234 | gene | 2391548 | 2392924 | - | 1519,233 | -0,159093029 | 0,240083 | -0,66266 | 0,507549 | 0,604493 | 1823,432788 | -0,426930842 | 0,157928 | -2,70333 | 0,006865 | 0,014362 | 1215,033 | 0,240085541 | 0,211642 | 1,134395 | 0,256629 | 0,358126 | 1702,93 | -0,96761 | 0,223202 | -4,33515 | 1,46E-05 | 8,67E-05 | 1407,856269 | -0,247297047 | 0,126542 | -1,95427 | 0,050669 | 0,074782 |
| 2146 | RM25_RS10475 | RM25_2120 | FGGY-family pentulose kinase | 2235 | gene | 2392937 | 2394511 | - | 1180,606 | -0,499757387 | 0,135218 | -3,69593 | 0,000219 | 0,003092 | 1180,231111 | -0,754291371 | 0,141753 | -5,32116 | 1,03E-07 | 6,25E-07 | 1180,982 | -0,255085801 | 0,212646 | -1,19958 | 0,230303 | 0,328586 | 1461,901 | -0,26642 | 0,16118 | -1,65294 | 0,098343 | 0,146891 | 959,8292848 | 0,29064252 | 0,185615 | 1,565836 | 0,117387 | 0,159287 |
| 2147 | RM25_RS10480 | RM25_2121 | HAD-superfamily hydrolase, subfamily IIA | 2236 | gene | 2394543 | 2395415 | - | 864,125 | -0,758080325 | 0,189617 | -3,99797 | 6,39E-05 | 0,001195 | 779,9127054 | -1,021400188 | 0,196293 | -5,20344 | 1,96E-07 | 1,12E-06 | 948,3373 | -0,553192786 | 0,222463 | -2,48668 | 0,012894 | 0,037154 | 1144,372 | 0,048042 | 0,242723 | 0,197929 | 0,8431 | 0,876958 | 629,8336018 | 0,573736239 | 0,164224 | 3,493611 | 0,000477 | 0,001074 |
| 2148 | RM25_RS10485 | RM25_2122 | Hypothetical protein | 2237 | gene | 2395475 | 2396554 | - | 1122,465 | -0,992418857 | 0,223412 | -4,4421 | 8,91E-06 | 0,000262 | 964,0361159 | -1,229587467 | 0,209895 | -5,85812 | 4,68E-09 | 3,61E-08 | 1280,894 | -0,823998064 | 0,241493 | -3,4121 | 0,000645 | 0,003414 | 1572,988 | 0,21428 | 0,253063 | 0,846746 | 0,397136 | 0,472514 | 737,276818 | 0,679499706 | 0,205577 | 3,305337 | 0,000949 | 0,002029 |
| 2149 | RM25_RS10490 | RM25_2123 | Ribose 5-phosphate isomerase | 2238 | gene | 2397100 | 2397588 | - | 686,3848 | 0,245773204 | 0,119985 | 2,048365 | 0,040524 | 0,109562 | 713,4907667 | 0,214528398 | 0,153297 | 1,39943 | 0,161684 | 0,216651 | 659,2789 | 0,279623655 | 0,224662 | 1,24464 | 0,213264 | 0,310862 | 664,2776 | -0,20573 | 0,19349 | -1,06325 | 0,287669 | 0,361619 | 729,1535318 | -0,093435421 | 0,16901 | -0,55284 | 0,580373 | 0,641021 |
| 2150 | RM25_RS10495 | RM25_2124 | Serine 3-dehydrogenase | 2239 | gene | 2398429 | 2399112 | + | 1696,342 | 0,676515889 | 0,174915 | 3,867684 | 0,00011 | 0,001837 | 1869,664917 | 0,820032682 | 0,170557 | 4,807963 | 1,52E-06 | 7,5E-06 | 1523,019 | 0,506078709 | 0,260665 | 1,941492 | 0,052199 | 0,103912 | 1382,696 | -0,15798 | 0,253008 | -0,6244 | 0,532366 | 0,598943 | 2043,329226 | -0,430302825 | 0,172569 | -2,49351 | 0,012649 | 0,02163 |
| 2151 | RM25_RS10500 | RM25_2125 | Putative aga operon transcriptional repressor | 2240 | gene | 2399257 | 2400087 | - | 639,0626 | -0,345120808 | 0,116449 | -2,96372 | 0,003039 | 0,020947 | 634,5630639 | -0,305709564 | 0,163485 | -1,86996 | 0,06149 | 0,093203 | 643,5622 | -0,384626396 | 0,223554 | -1,72051 | 0,08534 | 0,152268 | 755,5353 | -0,00582 | 0,186534 | -0,0312 | 0,975113 | 0,98045 | 552,7880761 | -0,026753112 | 0,211477 | -0,12651 | 0,899332 | 0,918027 |
| 2152 | RM25_RS10505 | RM25_2126 | Triose-phosphate isomerase | 2241 | gene | 2400249 | 2401001 | - | 701,4359 | -0,349002806 | 0,170823 | -2,04307 | 0,041046 | 0,109812 | 692,8357721 | -0,033590007 | 0,157583 | -0,21316 | 0,831204 | 0,866113 | 710,036 | -0,661195632 | 0,263477 | -2,5095 | 0,01209 | 0,035224 | 829,4678 | 0,257955 | 0,265964 | 0,969885 | 0,332104 | 0,407479 | 605,5431695 | -0,319411627 | 0,163849 | -1,94943 | 0,051244 | 0,075442 |
| 2153 | RM25_RS10510 | RM25_2127 | DAK1 domain protein | 2242 | gene | 2401076 | 2402794 | - | 823,3437 | -0,651583728 | 0,307589 | -2,11836 | 0,034145 | 0,098025 | 574,6359589 | -0,352171602 | 0,149648 | -2,35334 | 0,018606 | 0,033682 | 1072,052 | -0,817604819 | 0,282383 | -2,89538 | 0,003787 | 0,014479 | 1056,102 | 1,028699 | 0,159008 | 6,46947 | 9,83E-11 | 1,74E-09 | 629,8143212 | 0,621373826 | 0,272504 | 2,280236 | 0,022594 | 0,036625 |
| 2154 | RM25_RS10515 | RM25_2128 | Transcriptional regulator | 2243 | gene | 2402942 | 2403772 | + | 1068,2 | -0,814420825 | 0,22641 | -3,5971 | 0,000322 | 0,00419 | 865,2317118 | -0,887818104 | 0,187181 | -4,7431 | 2,1E-06 | 1,01E-05 | 1271,168 | -0,766366097 | 0,227849 | -3,36348 | 0,00077 | 0,003944 | 1432,329 | 0,445391 | 0,230745 | 1,930233 | 0,053578 | 0,089817 | 759,8715357 | 0,628506529 | 0,188155 | 3,340361 | 0,000837 | 0,001809 |
| 2155 | RM25_RS10520 | RM25_2129 | Putative MFS-family membrane transport protein | 2244 | gene | 2403811 | 2405103 | - | 535,0634 | -1,084106544 | 0,45536 | -2,38077 | 0,017277 | 0,065999 | 336,8311078 | -1,820682494 | 0,285498 | -6,37721 | 1,8E-10 | 1,7E-09 | 733,2956 | -0,792086962 | 0,3003 | -2,63765 | 0,008348 | 0,026662 | 763,8805 | 0,755614 | 0,307817 | 2,454752 | 0,014098 | 0,031114 | 337,1555224 | 1,852367952 | 0,305372 | 6,065945 | 1,31E-09 | 7,21E-09 |
| 2156 | RM25_RS10525 | RM25_2130 | Cyclase family protein | 2245 | gene | 2405093 | 2405818 | - | 396,5574 | -1,766955866 | 0,777234 | -2,27339 | 0,023003 | 0,07779 | 231,4536223 | -4,589122051 | 0,502116 | -9,13957 | 6,27E-20 | 1,89E-18 | 561,6612 | -1,194769149 | 0,348221 | -3,43107 | 0,000601 | 0,003235 | 644,1434 | 0,740443 | 0,480753 | 1,540171 | 0,123519 | 0,176766 | 177,2670934 | 4,212411901 | 0,446536 | 9,433534 | 3,96E-21 | 1,03E-19 |
| 2157 | RM25_RS12060 | RM25_2131 | Hypothetical protein | 2246 | gene | 2405958 | 2406272 | - | 169,822 | -1,79345848 | 0,565161 | -3,17336 | 0,001507 | 0,012787 | 107,9723407 | -3,171933911 | 0,427867 | -7,41337 | 1,23E-13 | 1,84E-12 | 231,6717 | -1,354409154 | 0,291088 | -4,65292 | 3,27E-06 | 3,84E-05 | 277,1683 | 0,703915 | 0,364331 | 1,932075 | 0,05335 | 0,089563 | 74,57029145 | 2,590041606 | 0,4244 | 6,102834 | 1,04E-09 | 5,81E-09 |
| 2158 | RM25_RS12065 | RM25_2132 | Hypothetical protein | 2247 | gene | 2406315 | 2406599 | - | 134,3974 | -0,44989704 | 0,23266 | -1,93371 | 0,053149 | 0,129147 | 141,4231317 | -0,214073561 | 0,331668 | -0,64545 | 0,518638 | 0,583778 | 127,3717 | -0,709448359 | 0,27987 | -2,53492 | 0,011247 | 0,033303 | 163,2995 | 0,010143 | 0,215162 | 0,047141 | 0,962401 | 0,970637 | 111,6468063 | -0,436614574 | 0,397638 | -1,09802 | 0,272196 | 0,332417 |
| 2159 | RM25_RS10535 | RM25_2133 | Uma4 protein | 2248 | gene | 2406599 | 2407945 | - | 0 | NA | NA | NA | NA | NA | 0 | NA | NA | NA | NA | NA | 0 | NA | NA | NA | NA | NA | 0 | NA | NA | NA | NA | NA | 0 | NA | NA | NA | NA | NA |
| 2160 | RM25_RS10540 | RM25_2134 | Putative integrase/recombinase | 2249 | gene | 2407965 | 2408372 | - | 92,16458 | -1,318395932 | 0,615525 | -2,1419 | 0,032201 | 0,094518 | 53,70518554 | -2,730214253 | 0,524655 | -5,20383 | 1,95E-07 | 1,12E-06 | 130,624 | -0,896081849 | 0,424915 | -2,10885 | 0,034958 | 0,077658 | 138,3185 | 0,786198 | 0,400966 | 1,960761 | 0,049907 | 0,084809 | 51,94647479 | 2,700494195 | 0,591961 | 4,561943 | 5,07E-06 | 1,56E-05 |
| 2161 | RM25_RS12070 | RM25_2135 | Phage integrase/recombinase | 2250 | gene | 2408365 | 2408898 | - | 189,8639 | -1,33774038 | 0,715775 | -1,86894 | 0,061631 | 0,142061 | 97,97812028 | -3,412797887 | 0,497836 | -6,85526 | 7,12E-12 | 8,39E-11 | 281,7498 | -0,880344545 | 0,400006 | -2,20083 | 0,027748 | 0,065402 | 285,2845 | 0,952489 | 0,453086 | 2,102228 | 0,035533 | 0,065025 | 106,060817 | 3,560973693 | 0,487493 | 7,304663 | 2,78E-13 | 2,63E-12 |
| 2162 | RM25_RS12075 | RM25_2136 | Hypothetical protein | 2251 | gene | 2408949 | 2409296 | - | 115,7272 | -0,996947596 | 0,512262 | -1,94617 | 0,051635 | 0,126779 | 75,45072849 | -2,233927223 | 0,421907 | -5,29483 | 1,19E-07 | 7,07E-07 | 156,0037 | -0,528643714 | 0,305587 | -1,72993 | 0,083643 | 0,15015 | 162,1267 | 0,4952 | 0,358469 | 1,381431 | 0,167146 | 0,226416 | 75,75573848 | 2,265597462 | 0,39774 | 5,696172 | 1,23E-08 | 5,73E-08 |
| 2163 | RM25_RS10550 | RM25_2137 | Tyrosine recombinase XerD | 2252 | gene | 2409293 | 2410234 | - | 899,5039 | 0,329030988 | 0,189625 | 1,735171 | 0,082711 | 0,170559 | 899,213346 | -0,103907531 | 0,17229 | -0,6031 | 0,546445 | 0,612444 | 899,7945 | 0,775849341 | 0,221892 | 3,496525 | 0,000471 | 0,002669 | 844,4454 | -0,54858 | 0,183625 | -2,98751 | 0,002813 | 0,008401 | 979,6518815 | 0,378795151 | 0,178575 | 2,12121 | 0,033904 | 0,052333 |
| 2164 | RM25_RS10555 | RM25_2138 | Hypothetical protein | 2253 | gene | 2410818 | 2411120 | + | 1208,282 | 0,128505169 | 0,221311 | 0,580655 | 0,561473 | 0,655794 | 1229,852762 | 0,469556349 | 0,163842 | 2,865906 | 0,004158 | 0,009299 | 1186,712 | -0,218876261 | 0,352193 | -0,62147 | 0,534293 | 0,631876 | 1218,257 | 0,255859 | 0,291163 | 0,87875 | 0,379537 | 0,45618 | 1234,192444 | -0,397991187 | 0,237627 | -1,67486 | 0,093962 | 0,130199 |
| 2165 | RM25_RS10560 | RM25_2139 | Hypothetical protein | 2254 | gene | 2411124 | 2411387 | + | 212,5175 | 0,087880457 | 0,199893 | 0,439637 | 0,6602 | 0,74241 | 183,6145878 | 0,304484446 | 0,220512 | 1,380807 | 0,167338 | 0,222834 | 241,4204 | -0,073386088 | 0,243256 | -0,30168 | 0,762894 | 0,82641 | 216,8254 | 0,530256 | 0,183477 | 2,89004 | 0,003852 | 0,010744 | 215,1913174 | 0,21419474 | 0,278755 | 0,768398 | 0,442251 | 0,507738 |
| 2166 | NA | RM25_2140 | Hypothetical protein | 2255 | gene | 2411384 | 2411911 | + | 222,9177 | 0,228683774 | 0,092521 | 2,471685 | 0,013448 | 0,057111 | 226,4264249 | 0,304291381 | 0,184066 | 1,653168 | 0,098297 | 0,140076 | 219,409 | 0,157934795 | 0,177967 | 0,887437 | 0,374844 | 0,480428 | 216,2951 | -0,025 | 0,164155 | -0,15231 | 0,878944 | 0,903998 | 236,2233651 | -0,116992252 | 0,183523 | -0,63748 | 0,523812 | 0,584854 |
| 2167 | RM25_RS10570 | RM25_2141 | Glycogen debranching enzyme GlgX | 2256 | gene | 2411865 | 2413997 | + | 11413,99 | 0,351885292 | 0,480427 | 0,732442 | 0,463899 | 0,566827 | 16854,26318 | 0,199321337 | 0,212116 | 0,939683 | 0,34738 | 0,416468 | 5973,708 | 0,79663038 | 0,297121 | 2,681169 | 0,007337 | 0,024219 | 10701,06 | -1,89943 | 0,294832 | -6,44243 | 1,18E-10 | 2,07E-09 | 12543,68985 | -1,263239734 | 0,198541 | -6,36262 | 1,98E-10 | 1,22E-09 |
| 2168 | RM25_RS10575 | RM25_2142 | Hypothetical protein | 2257 | gene | 2414067 | 2414495 | + | 394,7924 | -0,321203832 | 0,327533 | -0,98068 | 0,326753 | 0,436844 | 503,7322565 | -0,020833591 | 0,201986 | -0,10314 | 0,917849 | 0,937335 | 285,8526 | -0,864231063 | 0,235651 | -3,66742 | 0,000245 | 0,001545 | 463,6437 | -0,51579 | 0,197553 | -2,61089 | 0,009031 | 0,021787 | 344,9570738 | -1,302810752 | 0,236769 | -5,50245 | 3,75E-08 | 1,66E-07 |
| 2169 | RM25_RS10585 | RM25_2143 | Uma4 protein | 2258 | gene | 2415142 | 2416488 | + | 0 | NA | NA | NA | NA | NA | 0 | NA | NA | NA | NA | NA | 0 | NA | NA | NA | NA | NA | 0 | NA | NA | NA | NA | NA | 0 | NA | NA | NA | NA | NA |
| 2170 | RM25_RS10590 | RM25_2144 | Hypothetical protein | 2259 | gene | 2416427 | 2416906 | + | 266,1061 | -0,548866894 | 0,144063 | -3,80991 | 0,000139 | 0,002156 | 265,7401827 | -0,581194748 | 0,212586 | -2,73392 | 0,006258 | 0,013325 | 266,4719 | -0,51633291 | 0,249605 | -2,0686 | 0,038584 | 0,083992 | 333,3458 | -0,08621 | 0,218566 | -0,39443 | 0,693265 | 0,740698 | 212,0091523 | 0,037538935 | 0,241727 | 0,155295 | 0,876589 | 0,899486 |
| 2171 | NA | RM25_2145 | Hypothetical protein | 2260 | gene | 2416866 | 2417498 | + | 47,58696 | -1,612289035 | 0,615144 | -2,62099 | 0,008767 | 0,042939 | 24,93853252 | -2,712888563 | 0,597958 | -4,53692 | 5,71E-06 | 2,52E-05 | 70,23539 | -1,327162289 | 0,438877 | -3,024 | 0,002495 | 0,01037 | 74,96909 | 1,140264 | 0,384314 | 2,967014 | 0,003007 | 0,008826 | 22,99506476 | 2,597664257 | 0,687852 | 3,77649 | 0,000159 | 0,000393 |
| 2172 | RM25_RS10595 | RM25_2146 | DNA-invertase hin | 2261 | gene | 2417530 | 2418111 | - | 738,3935 | 0,432427639 | 0,232048 | 1,863528 | 0,062388 | 0,142584 | 899,8181067 | 0,525384464 | 0,152617 | 3,442499 | 0,000576 | 0,001641 | 576,969 | 0,293129128 | 0,215726 | 1,358802 | 0,174209 | 0,265424 | 666,3508 | -0,56588 | 0,205677 | -2,75133 | 0,005935 | 0,015239 | 831,2722571 | -0,752881995 | 0,142688 | -5,27641 | 1,32E-07 | 5,25E-07 |
| 2173 | RM25_RS10600 | RM25_2147 | Nucleotidyl transferase, PF08843 family | 2262 | gene | 2418108 | 2418887 | - | 1230,388 | 0,520621968 | 0,2301 | 2,26259 | 0,023661 | 0,07877 | 1372,959693 | 0,79798547 | 0,1985 | 4,02008 | 5,82E-05 | 0,00021 | 1087,816 | 0,185352594 | 0,325774 | 0,56896 | 0,569383 | 0,662407 | 1069,482 | -0,0277 | 0,343823 | -0,08057 | 0,935782 | 0,94907 | 1420,919841 | -0,601684336 | 0,160029 | -3,75985 | 0,00017 | 0,000415 |
| 2174 | RM25_RS10605 | RM25_2148 | Transcriptional regulator | 2263 | gene | 2418884 | 2419216 | - | 2129,654 | 0,013574955 | 0,3338 | 0,040668 | 0,967561 | 0,97873 | 1972,159167 | 0,627303137 | 0,205795 | 3,048198 | 0,002302 | 0,005577 | 2287,149 | -0,511499522 | 0,479682 | -1,06633 | 0,286274 | 0,390705 | 2239,748 | 0,74699 | 0,456702 | 1,635618 | 0,10192 | 0,151215 | 2091,494287 | -0,362420589 | 0,257637 | -1,40671 | 0,159514 | 0,209573 |
| 2175 | RM25_RS10610 | RM25_2149 | Hsp20/alpha crystallin family protein | 2264 | gene | 2419576 | 2420058 | + | 35409,78 | 0,351316331 | 0,306468 | 1,146339 | 0,251655 | 0,359703 | 31979,58656 | 0,312907265 | 0,247179 | 1,265915 | 0,205544 | 0,265898 | 38839,98 | 0,383071154 | 0,48129 | 0,795926 | 0,426075 | 0,530512 | 32935,04 | 0,196852 | 0,455269 | 0,432386 | 0,665461 | 0,718173 | 38712,44315 | 0,290504161 | 0,287969 | 1,008802 | 0,31307 | 0,376482 |
| 2176 | RM25_RS10615 | RM25_2150 | Putative insertion element ISCmi2 transposase | 2265 | gene | 2420257 | 2421300 | + | 301,1019 | 0,618832842 | 0,335982 | 1,841863 | 0,065495 | 0,147161 | 292,3587581 | -0,003840075 | 0,325229 | -0,01181 | 0,990579 | 0,992263 | 309,845 | 1,263248274 | 0,409478 | 3,085023 | 0,002035 | 0,008762 | 252,4131 | -0,72909 | 0,432805 | -1,68456 | 0,092073 | 0,138491 | 355,4177313 | 0,56578563 | 0,29703 | 1,904808 | 0,056805 | 0,082955 |
| 2177 | RM25_RS10620 | RM25_2151 | Heat shock protein Hsp20 | 2266 | gene | 2421410 | 2421874 | + | 3556,092 | -0,360923172 | 0,370026 | -0,9754 | 0,329362 | 0,439834 | 4169,363719 | -0,93325565 | 0,30136 | -3,09682 | 0,001956 | 0,004818 | 2942,821 | 0,413045573 | 0,427225 | 0,966811 | 0,333638 | 0,439076 | 4270,386 | -1,16758 | 0,461846 | -2,52807 | 0,011469 | 0,026322 | 3041,628949 | 0,213089353 | 0,268777 | 0,79281 | 0,427888 | 0,493002 |
| 2178 | RM25_RS10625 | RM25_2152 | putative potassium transport system protein kup | 2267 | gene | 2422031 | 2423908 | - | 4437,813 | 0,221031992 | 0,305152 | 0,724335 | 0,46886 | 0,570229 | 5032,269707 | -0,392814061 | 0,167709 | -2,34224 | 0,019169 | 0,034515 | 3843,357 | 1,066518643 | 0,275903 | 3,865555 | 0,000111 | 0,000806 | 4356,902 | -1,26324 | 0,223915 | -5,6416 | 1,68E-08 | 1,81E-07 | 4668,951773 | 0,242659476 | 0,187906 | 1,291386 | 0,19657 | 0,250848 |
| 2179 | RM25_RS10630 | RM25_2153 | Uma4 protein | 2268 | gene | 2424231 | 2425577 | - | 28,19474 | -2,140119519 | 0,86059 | -2,48681 | 0,01289 | 0,055541 | 13,78899903 | -5,156335646 | 1,257555 | -4,10029 | 4,13E-05 | 0,000154 | 42,60049 | -1,70799633 | 0,374331 | -4,5628 | 5,05E-06 | 5,57E-05 | 48,09801 | 1,19992 | 0,409443 | 2,930613 | 0,003383 | 0,00976 | 10,12648732 | 4,726264877 | 1,332237 | 3,547614 | 0,000389 | 0,000888 |
| 2180 | RM25_RS10635 | RM25_2154 | Hypothetical protein | 2269 | gene | 2425626 | 2426348 | - | 392,6961 | -0,202070059 | 0,152994 | -1,32077 | 0,186578 | 0,291031 | 369,7823219 | -0,390276666 | 0,191409 | -2,03897 | 0,041454 | 0,068088 | 415,6099 | -0,039953258 | 0,231496 | -0,17259 | 0,862976 | 0,90018 | 443,0805 | -0,05615 | 0,221312 | -0,25373 | 0,799703 | 0,837733 | 358,2686869 | 0,352169607 | 0,191887 | 1,835299 | 0,066461 | 0,095518 |
| 2181 | RM25_RS10640 | RM25_2155 | Hypothetical protein | 2270 | gene | 2427275 | 2427754 | + | 3321,555 | 1,201611362 | 0,354355 | 3,390985 | 0,000696 | 0,007854 | 3545,353709 | 0,640874994 | 0,237737 | 2,695732 | 0,007023 | 0,014637 | 3097,757 | 1,976889956 | 0,428674 | 4,611644 | 3,99E-06 | 4,62E-05 | 2144,132 | -1,18976 | 0,416508 | -2,85652 | 0,004283 | 0,011617 | 4519,404948 | 0,174821316 | 0,251136 | 0,696123 | 0,486352 | 0,548377 |
| 2182 | RM25_RS10645 | RM25_2156 | Transposase, IS30 family | 2271 | gene | 2428150 | 2429307 | + | 2418,601 | 0,328262552 | 0,339597 | 0,966625 | 0,333732 | 0,44441 | 3263,43816 | 0,37422215 | 0,155981 | 2,399152 | 0,016433 | 0,030426 | 1573,764 | 0,234514949 | 0,221912 | 1,056793 | 0,290606 | 0,394202 | 2276,613 | -1,03724 | 0,219851 | -4,71791 | 2,38E-06 | 1,69E-05 | 2643,453719 | -1,119708781 | 0,130917 | -8,55278 | 1,2E-17 | 2,04E-16 |
| 2183 | RM25_RS12415 | RM25_2157 | Hypothetical protein | 2272 | gene | 2429269 | 2429577 | - | 838,7682 | 0,36046416 | 0,295502 | 1,219838 | 0,222526 | 0,328456 | 688,6994838 | 0,501230056 | 0,184704 | 2,71369 | 0,006654 | 0,014015 | 988,8369 | 0,265437088 | 0,430952 | 0,615933 | 0,537939 | 0,635232 | 774,8091 | 0,606539 | 0,385061 | 1,575177 | 0,115216 | 0,166603 | 920,3210927 | 0,402539384 | 0,262047 | 1,536133 | 0,124506 | 0,167883 |
| 2184 | RM25_RS12805 | RM25_2158 | Hypothetical protein | 2273 | gene | 2429583 | 2429759 | - | 79,35054 | 0,164291904 | 0,217518 | 0,755303 | 0,450067 | 0,556523 | 76,11594186 | 0,406829424 | 0,310874 | 1,308662 | 0,190649 | 0,24909 | 82,58514 | -0,051156151 | 0,329399 | -0,1553 | 0,876584 | 0,910581 | 78,92274 | 0,295915 | 0,341943 | 0,865393 | 0,386823 | 0,463519 | 82,34775875 | -0,110228245 | 0,310077 | -0,35549 | 0,722225 | 0,769221 |
| 2185 | RM25_RS13325 | RM25_2159 | Hypothetical protein | 2274 | gene | 2430048 | 2430332 | + | 89,14276 | -0,889014414 | 0,501155 | -1,77393 | 0,076075 | 0,163102 | 54,9705161 | -1,519426742 | 0,443283 | -3,42767 | 0,000609 | 0,001725 | 123,315 | -0,643352479 | 0,455788 | -1,41152 | 0,158093 | 0,245309 | 121,6987 | 0,807811 | 0,323346 | 2,498284 | 0,01248 | 0,028283 | 61,56039843 | 1,759597209 | 0,582073 | 3,022983 | 0,002503 | 0,004906 |
| 2186 | NA | RM25_2160 | Hypothetical protein | 2275 | gene | 2430406 | 2430573 | + | 60,07199 | -1,310157844 | 0,340089 | -3,85239 | 0,000117 | 0,001919 | 45,17208736 | -1,404824219 | 0,429565 | -3,27034 | 0,001074 | 0,002864 | 74,97189 | -1,26443173 | 0,351611 | -3,59611 | 0,000323 | 0,001972 | 90,03541 | 0,620159 | 0,317408 | 1,953822 | 0,050722 | 0,085885 | 33,81813539 | 0,829115104 | 0,516353 | 1,605713 | 0,108337 | 0,148201 |
| 2187 | RM25_RS12420 | RM25_2161 | Putative integrase/recombinase y4rC | 2276 | gene | 2430700 | 2431365 | - | 418,0941 | -1,312346858 | 0,548574 | -2,39229 | 0,016744 | 0,064713 | 257,7834647 | -2,535283404 | 0,344652 | -7,35607 | 1,89E-13 | 2,76E-12 | 578,4046 | -0,898421445 | 0,37841 | -2,3742 | 0,017587 | 0,046733 | 624,4712 | 0,707239 | 0,395622 | 1,787662 | 0,073831 | 0,116167 | 236,205727 | 2,414062973 | 0,349737 | 6,902504 | 5,11E-12 | 3,92E-11 |
| 2188 | RM25_RS10670 | RM25_2162 | Transposase IS204/IS1001/IS1096/IS1165 family protein | 2277 | gene | 2431469 | 2432776 | + | 17,07418 | -1,65299234 | 0,738156 | -2,23935 | 0,025133 | 0,081105 | 11,83184188 | -3,040852284 | 1,012315 | -3,00386 | 0,002666 | 0,006344 | 22,31651 | -1,156513985 | 0,672543 | -1,71961 | 0,085503 | 0,152443 | 27,35088 | 0,459055 | 0,445221 | 1,031074 | 0,302506 | 0,377053 | 8,052014414 | 2,4563196 | 1,40713 | 1,745623 | 0,080876 | 0,113806 |
| 2189 | RM25_RS12430 | RM25_2163 | Hypothetical protein | 2278 | gene | 2433036 | 2433371 | + | 135,6033 | -1,884081381 | 0,68758 | -2,74016 | 0,006141 | 0,034218 | 74,27154476 | -3,826568248 | 0,532554 | -7,18531 | 6,71E-13 | 9,08E-12 | 196,935 | -1,450603532 | 0,298798 | -4,85479 | 1,21E-06 | 1,66E-05 | 223,6952 | 0,983452 | 0,410656 | 2,394832 | 0,016628 | 0,035404 | 56,63372605 | 3,427606243 | 0,491286 | 6,97681 | 3,02E-12 | 2,4E-11 |
| 2190 | RM25_RS12435 | RM25_2164 | Restriction endonuclease PvuRts1 I | 2279 | gene | 2433390 | 2434328 | - | 3782,401 | -0,587811334 | 0,544531 | -1,07948 | 0,280373 | 0,389646 | 1712,743807 | 0,025184106 | 0,171668 | 0,146703 | 0,883367 | 0,910405 | 5852,059 | -0,774987595 | 0,453736 | -1,70801 | 0,087634 | 0,15542 | 4765,422 | 2,072265 | 0,368298 | 5,626599 | 1,84E-08 | 1,95E-07 | 2939,966361 | 1,299769373 | 0,306485 | 4,240893 | 2,23E-05 | 6,26E-05 |
| 2191 | RM25_RS10685 | RM25_2165 | Transposase IS204/IS1001/IS1096/IS1165 family protein | 2280 | gene | 2435402 | 2436184 | + | 75,13505 | -1,428630095 | 0,606856 | -2,35415 | 0,018565 | 0,069019 | 41,82160347 | -2,918028109 | 0,51493 | -5,66684 | 1,45E-08 | 1,03E-07 | 108,4485 | -1,036200592 | 0,269794 | -3,84071 | 0,000123 | 0,000876 | 115,1139 | 0,903064 | 0,315101 | 2,865951 | 0,004158 | 0,011351 | 39,66711949 | 2,85192135 | 0,505792 | 5,63853 | 1,72E-08 | 7,91E-08 |
| 2192 | RM25_RS10690 | RM25_2166 | Putative IS1236 transposase | 2281 | gene | 2436403 | 2437377 | - | 226,6373 | -0,17801338 | 0,113936 | -1,5624 | 0,118193 | 0,214129 | 220,5753359 | -0,157410285 | 0,192255 | -0,81876 | 0,412926 | 0,481338 | 232,6992 | -0,193763297 | 0,215527 | -0,89902 | 0,368641 | 0,474647 | 253,6242 | 0,030801 | 0,215965 | 0,142621 | 0,88659 | 0,909352 | 208,8156046 | 0,048111497 | 0,185293 | 0,259652 | 0,795133 | 0,833702 |
| 2193 | RM25_RS10700 | RM25_2167 | Transposase IS3/IS911 | 2282 | gene | 2437374 | 2437718 | - | 345,1415 | 0,377980361 | 0,202264 | 1,868745 | 0,061658 | 0,142061 | 368,9877116 | 0,388554588 | 0,207117 | 1,876011 | 0,060654 | 0,092114 | 321,2953 | 0,368767345 | 0,337195 | 1,093633 | 0,274116 | 0,37695 | 318,4155 | -0,24208 | 0,325519 | -0,74368 | 0,45707 | 0,530435 | 381,5946053 | -0,220170833 | 0,225722 | -0,97541 | 0,329358 | 0,39317 |
| 2194 | NA | RM25_2168 | Hypothetical protein | 2283 | gene | 2437767 | 2437964 | + | 87,08669 | -1,175831993 | 0,842387 | -1,39583 | 0,162764 | 0,265538 | 43,51182369 | -4,034617008 | 0,6763 | -5,96572 | 2,44E-09 | 1,97E-08 | 130,6616 | -0,649281171 | 0,630998 | -1,02898 | 0,303491 | 0,407827 | 126,8704 | 0,874459 | 0,49097 | 1,781086 | 0,074898 | 0,117299 | 52,97540484 | 4,354744362 | 0,832017 | 5,233958 | 1,66E-07 | 6,51E-07 |
| 2195 | RM25_RS10705 | RM25_2169 | Nitric oxide reductase | 2284 | gene | 2438694 | 2441108 | + | 4463,641 | -0,046994419 | 0,167504 | -0,28056 | 0,77905 | 0,841917 | 4502,007475 | 0,104112558 | 0,132054 | 0,78841 | 0,430457 | 0,498814 | 4425,275 | -0,200421668 | 0,308554 | -0,64955 | 0,515982 | 0,61635 | 4794,357 | 0,072525 | 0,255513 | 0,28384 | 0,776533 | 0,819288 | 4294,649188 | -0,195079216 | 0,197719 | -0,98665 | 0,323814 | 0,38782 |
| 2196 | RM25_RS10710 | RM25_2170 | Putative transposase | 2285 | gene | 2441113 | 2441982 | + | 1153,02 | 0,009587348 | 0,114715 | 0,083575 | 0,933394 | 0,954451 | 1155,541438 | 0,068985761 | 0,137529 | 0,501611 | 0,615942 | 0,674303 | 1150,498 | -0,048952554 | 0,231453 | -0,2115 | 0,832496 | 0,877985 | 1214,537 | -0,00365 | 0,183638 | -0,01988 | 0,984138 | 0,986025 | 1132,211339 | -0,077409381 | 0,174458 | -0,44371 | 0,657249 | 0,711592 |
| 2197 | RM25_RS10715 | RM25_2171 | SPFH/Band 7/PHB domain protein | 2286 | gene | 2442117 | 2442881 | + | 658,4042 | -0,157213721 | 0,121745 | -1,29134 | 0,196587 | 0,299693 | 660,8994476 | -0,370240982 | 0,140753 | -2,63044 | 0,008528 | 0,017285 | 655,909 | 0,053476782 | 0,209218 | 0,255604 | 0,798257 | 0,854835 | 734,0517 | -0,2739 | 0,172973 | -1,58346 | 0,113316 | 0,164766 | 609,4300499 | 0,20341368 | 0,149874 | 1,357228 | 0,174709 | 0,226756 |
| 2198 | RM25_RS10720 | RM25_2172 | Hypothetical protein | 2287 | gene | 2442995 | 2443417 | + | 732,1221 | -0,08690433 | 0,305508 | -0,28446 | 0,776059 | 0,840226 | 594,0076299 | 0,480580791 | 0,153026 | 3,140511 | 0,001687 | 0,00422 | 870,2366 | -0,472914259 | 0,391185 | -1,20893 | 0,22669 | 0,32402 | 793,9656 | 0,974977 | 0,347188 | 2,80821 | 0,004982 | 0,013134 | 693,7826098 | 0,058177374 | 0,235746 | 0,24678 | 0,805079 | 0,841868 |
| 2199 | RM25_RS10725 | RM25_2173 | Hypothetical protein | 2288 | gene | 2443374 | 2443667 | + | 231,4062 | -0,544082306 | 0,136336 | -3,99073 | 6,59E-05 | 0,001222 | 218,7977561 | -0,385172797 | 0,198929 | -1,93623 | 0,052839 | 0,082697 | 244,0146 | -0,685306623 | 0,221238 | -3,09759 | 0,001951 | 0,008484 | 288,9138 | 0,221671 | 0,204303 | 1,085013 | 0,277916 | 0,351186 | 185,0119373 | -0,025419719 | 0,217058 | -0,11711 | 0,906773 | 0,924366 |
| 2200 | RM25_RS10730 | RM25_2174 | Hypothetical protein | 2289 | gene | 2443685 | 2444545 | - | 550,4226 | -1,319919811 | 0,438232 | -3,01192 | 0,002596 | 0,018712 | 381,2114886 | -2,110561676 | 0,310114 | -6,80576 | 1,01E-11 | 1,16E-10 | 719,6337 | -0,97265127 | 0,315309 | -3,08475 | 0,002037 | 0,008762 | 826,3107 | 0,553971 | 0,365587 | 1,515293 | 0,129698 | 0,183929 | 309,310614 | 1,760084926 | 0,286236 | 6,149059 | 7,79E-10 | 4,42E-09 |
| 2201 | RM25_RS10735 | RM25_2175 | Putative transposase for insertion sequence element IS986/ | 2290 | gene | 2444600 | 2444944 | - | 18,07857 | 0,040875048 | 0,435506 | 0,093856 | 0,925223 | 0,949875 | 11,73417903 | -0,147283551 | 0,646685 | -0,22775 | 0,819839 | 0,856923 | 24,42295 | 0,122828616 | 0,452549 | 0,271415 | 0,786072 | 0,845242 | 18,87669 | 0,847983 | 0,497554 | 1,704303 | 0,088325 | 0,134051 | 17,84407402 | 1,166239472 | 0,575724 | 2,025691 | 0,042796 | 0,064454 |
| 2202 | RM25_RS10740 | RM25_2176 | Transposase, IS30 family | 2291 | gene | 2444925 | 2446274 | - | 0 | NA | NA | NA | NA | NA | 0 | NA | NA | NA | NA | NA | 0 | NA | NA | NA | NA | NA | 0 | NA | NA | NA | NA | NA | 0 | NA | NA | NA | NA | NA |
| 2203 | NA | RM25_2177 | OrfB | 2292 | gene | 2446214 | 2446987 | - | 50,83662 | -0,814962953 | 0,555909 | -1,466 | 0,142648 | 0,243286 | 25,52260932 | -1,212210327 | 0,535308 | -2,26451 | 0,023543 | 0,041134 | 76,15062 | -0,698284074 | 0,428029 | -1,63139 | 0,102807 | 0,175719 | 67,85498 | 1,3166 | 0,373454 | 3,525467 | 0,000423 | 0,001718 | 36,22687025 | 1,912178363 | 0,578738 | 3,304049 | 0,000953 | 0,002035 |

| | | | | | | | | | | | | | | | | | | | | | | | | | | | | | | | | | | | | | | |
|---|---|---|---|---|---|---|---|---|---|---|---|---|---|---|---|---|---|---|---|---|---|---|---|---|---|---|---|---|---|---|---|---|---|---|---|---|---|---|
| 2204 | NA | RM25_2178 | IS3 family element | 2293 | gene | 2447020 | 2447364 | - | 40,4381 | 0,399143716 | 0,457072 | 0,873262 | 0,382521 | 0,493217 | 55,95122303 | 0,364847602 | 0,480859 | 0,758742 | 0,448007 | 0,516611 | 24,92498 | 0,470039042 | 0,38285 | 1,227738 | 0,219545 | 0,316301 | 37,14845 | -1,27408 | 0,349956 | -3,64069 | 0,000272 | 0,001176 | 45,08336882 | -1,124966552 | 0,548953 | -2,04929 | 0,040433 | 0,061326 |
| 2205 | RM25_RS12450 | RM25_2179 | Hypothetical protein | 2294 | gene | 2447676 | 2447834 | + | 86,52647 | -0,710354158 | 0,41763 | -1,70092 | 0,088959 | 0,179363 | 61,63439721 | -1,724013203 | 0,376255 | -4,58203 | 4,6E-06 | 2,06E-05 | 111,4185 | -0,239678766 | 0,225467 | -1,06303 | 0,287767 | 0,392062 | 113,3275 | 0,275199 | 0,264365 | 1,040983 | 0,297883 | 0,372275 | 64,1253814 | 1,823442124 | 0,353749 | 5,154619 | 2,54E-07 | 9,72E-07 |
| 2206 | RM25_RS10760 | RM25_2180 | Hypothetical protein | 2295 | gene | 2447831 | 2448394 | + | 122,7662 | 0,161741728 | 0,283599 | 0,570318 | 0,568462 | 0,661335 | 107,8132406 | -0,610514241 | 0,249878 | -2,44325 | 0,014555 | 0,027337 | 137,7192 | 0,758256835 | 0,280785 | 2,700486 | 0,006924 | 0,023247 | 122,8573 | -0,41722 | 0,279164 | -1,49454 | 0,135034 | 0,190048 | 126,9119922 | 1,014538398 | 0,241614 | 4,198999 | 2,68E-05 | 7,43E-05 |
| 2207 | RM25_RS10765 | RM25_2181 | Ribonuclease BN | 2296 | gene | 2448457 | 2449551 | + | 1612,85 | -0,054576946 | 0,172253 | -0,31684 | 0,751364 | 0,819512 | 1747,700374 | -0,376717632 | 0,120718 | -3,12064 | 0,001805 | 0,004491 | 1477,999 | 0,324546284 | 0,214456 | 1,51335 | 0,130191 | 0,211336 | 1740,881 | -0,65106 | 0,174706 | -3,72663 | 0,000194 | 0,000876 | 1549,252053 | 0,101732155 | 0,122809 | 0,828379 | 0,407456 | 0,474025 |
| 2208 | RM25_RS12645 | RM25_2182 | Hypothetical protein | 2297 | gene | 2449598 | 2449852 | + | 310,3427 | 0,431688962 | 0,375615 | 1,149284 | 0,250439 | 0,358617 | 410,9528525 | 0,120762556 | 0,230689 | 0,523486 | 0,600636 | 0,660924 | 209,7326 | 1,086373543 | 0,197954 | 5,488006 | 4,06E-08 | 7,79E-07 | 281,5225 | -1,6069 | 0,257636 | -6,23707 | 4,46E-10 | 6,82E-09 | 349,2344837 | -0,595164382 | 0,164438 | -3,61939 | 0,000295 | 0,000691 |
| 2209 | RM25_RS10775 | RM25_2183 | Transposase IS204/IS1001/IS1096/IS1165 family protein | 2298 | gene | 2449776 | 2451131 | - | 472,2406 | -1,361001167 | 0,563465 | -2,41542 | 0,015717 | 0,062577 | 291,2866775 | -2,636234982 | 0,340915 | -7,73281 | 1,05E-14 | 1,71E-13 | 653,1945 | -0,936711079 | 0,421811 | -2,22069 | 0,026372 | 0,063169 | 714,0874 | 0,701821 | 0,381779 | 1,838289 | 0,06602 | 0,106801 | 261,0532288 | 2,478466613 | 0,41583 | 5,960286 | 2,52E-09 | 1,33E-08 |
| 2210 | RM25_RS10780 | RM25_2184 | Oxidoreductase, FAD/FMN dependent domain protein | 2299 | gene | 2451098 | 2451886 | - | 398,4022 | -0,580115157 | 0,235634 | -2,46193 | 0,013819 | 0,057861 | 324,3864957 | -0,83920595 | 0,207358 | -4,04714 | 5,18E-05 | 0,00019 | 472,4179 | -0,412863698 | 0,215632 | -1,91467 | 0,055535 | 0,109079 | 503,0054 | 0,307727 | 0,241646 | 1,27346 | 0,202855 | 0,266813 | 312,8571367 | 0,793551951 | 0,175947 | 4,510185 | 6,48E-06 | 1,95E-05 |
| 2211 | RM25_RS10785 | RM25_2185 | Putative insertion element ISCmi2 transposase | 2300 | gene | 2452010 | 2453053 | + | 48,133 | -0,030286054 | 0,207574 | -0,1459 | 0,883997 | 0,92153 | 42,79379064 | -0,058522425 | 0,379404 | -0,15425 | 0,877414 | 0,905312 | 53,47222 | -0,009135491 | 0,28079 | -0,03253 | 0,974045 | 0,985032 | 51,08102 | 0,218592 | 0,284728 | 0,767724 | 0,442651 | 0,517524 | 46,78358873 | 0,330873339 | 0,36901 | 0,896651 | 0,369905 | 0,434753 |
| 2212 | RM25_RS10790 | RM25_2186 | G/U mismatch-specific DNA glycosylase | 2301 | gene | 2453269 | 2453868 | - | 422,1756 | 0,039799838 | 0,143044 | 0,278234 | 0,780833 | 0,842951 | 460,0352407 | 0,191230844 | 0,173677 | 1,101073 | 0,270865 | 0,336902 | 384,316 | -0,135880897 | 0,187116 | -0,72618 | 0,467726 | 0,570026 | 439,9822 | -0,15311 | 0,145907 | -1,04939 | 0,293997 | 0,368114 | 420,3852325 | -0,423718078 | 0,210803 | -2,01002 | 0,044429 | 0,0667 |
| 2213 | RM25_RS10795 | RM25_2187 | Ornithine carbamoyltransferase | 2302 | gene | 2453872 | 2454876 | - | 2829,884 | 0,233091067 | 0,25404 | 0,917538 | 0,358861 | 0,470431 | 3318,126471 | 0,359100006 | 0,177404 | 2,024191 | 0,04295 | 0,070107 | 2341,641 | 0,05684323 | 0,352365 | 0,161319 | 0,871842 | 0,906552 | 2759,999 | -0,39004 | 0,329207 | -1,1848 | 0,236098 | 0,304422 | 2992,966353 | -0,659447767 | 0,207291 | -3,18127 | 0,001466 | 0,002987 |
| 2214 | RM25_RS10800 | RM25_2188 | Putative membrane protein | 2303 | gene | 2455015 | 2455395 | + | 1488,129 | 0,886032543 | 0,262777 | 3,371804 | 0,000747 | 0,008342 | 1802,342059 | 0,987956645 | 0,186294 | 5,303218 | 1,14E-07 | 6,79E-07 | 1173,916 | 0,735424454 | 0,329472 | 2,232128 | 0,025607 | 0,061839 | 1109,805 | -0,51034 | 0,289156 | -1,76492 | 0,077578 | 0,120332 | 1889,24183 | -0,726195957 | 0,229729 | -3,1611 | 0,001572 | 0,003179 |
| 2215 | RM25_RS10805 | RM25_2189 | Low molecular weight phosphotyrosine protein phosphatase | 2304 | gene | 2455392 | 2455856 | + | 360,9413 | 0,519648455 | 0,164787 | 3,153448 | 0,001614 | 0,013114 | 402,2198792 | 0,758711426 | 0,168224 | 4,510113 | 6,48E-06 | 2,81E-05 | 319,6627 | 0,234124587 | 0,200373 | 1,168446 | 0,242627 | 0,342233 | 313,3724 | -0,0878 | 0,171103 | -0,51312 | 0,60787 | 0,667327 | 417,6943764 | -0,552083976 | 0,200352 | -2,75557 | 0,005859 | 0,010713 |
| 2216 | RM25_RS10810 | RM25_2190 | Cyanophycinase | 2305 | gene | 2455912 | 2456808 | + | 1906,871 | 0,169894004 | 0,167092 | 1,016768 | 0,309264 | 0,417973 | 1911,46752 | 0,407316498 | 0,16669 | 2,443551 | 0,014544 | 0,027336 | 1902,275 | -0,065178305 | 0,271606 | -0,23997 | 0,810351 | 0,865427 | 1895,462 | 0,193795 | 0,285523 | 0,678738 | 0,497304 | 0,564889 | 1978,005781 | -0,239425903 | 0,12723 | -1,88183 | 0,059858 | 0,086822 |
| 2217 | RM25_RS10815 | RM25_2191 | Two component response transcriptional regulatory protein | 2306 | gene | 2456815 | 2457507 | + | 1304,597 | 0,288401581 | 0,206751 | 1,394926 | 0,163038 | 0,265538 | 1462,817069 | 0,30599431 | 0,17031 | 1,796695 | 0,072384 | 0,107369 | 1146,376 | 0,267442931 | 0,316822 | 0,844143 | 0,398589 | 0,504552 | 1246,766 | -0,38296 | 0,315317 | -1,21451 | 0,224552 | 0,291769 | 1404,284248 | -0,382392011 | 0,168241 | -2,27288 | 0,023033 | 0,037185 |
| 2218 | RM25_RS10820 | RM25_2192 | Signal transduction histidine-protein kinase/phosphatase mı | 2307 | gene | 2457504 | 2458931 | + | 1550,681 | 0,232366296 | 0,276753 | 0,839616 | 0,401124 | 0,511884 | 1975,298895 | 0,243377748 | 0,173493 | 1,402811 | 0,160673 | 0,215542 | 1126,062 | 0,215002577 | 0,181332 | 1,185688 | 0,235746 | 0,334529 | 1513,327 | -0,85433 | 0,213321 | -4,00493 | 6,2E-05 | 0,000321 | 1643,694038 | -0,830282213 | 0,124753 | -6,65539 | 2,83E-11 | 1,97E-10 |
| 2219 | RM25_RS10825 | RM25_2193 | Membrane protein mosC | 2308 | gene | 2458960 | 2460219 | - | 396,2315 | -0,748978709 | 0,487338 | -1,53688 | 0,124323 | 0,221821 | 228,6966788 | -1,180965311 | 0,309681 | -3,81349 | 0,000137 | 0,000452 | 563,7664 | -0,585803635 | 0,44234 | -1,32433 | 0,185394 | 0,279038 | 520,5271 | 1,022301 | 0,334929 | 3,052292 | 0,002271 | 0,007034 | 291,7588516 | 1,692951053 | 0,434159 | 3,899382 | 9,64E-05 | 0,000245 |
| 2220 | RM25_RS10830 | RM25_2194 | Hypothetical protein | 2309 | gene | 2460219 | 2461142 | - | 251,4985 | -0,267777508 | 0,683523 | -0,39176 | 0,695235 | 0,772592 | 99,92434587 | -1,733048657 | 0,387557 | -4,47172 | 7,76E-06 | 3,31E-05 | 403,0727 | 0,048759562 | 0,437834 | 0,111365 | 0,911327 | 0,93822 | 286,558 | 1,307657 | 0,297464 | 4,396018 | 1,1E-05 | 6,7E-05 | 225,4245297 | 3,15385689 | 0,486943 | 6,476853 | 9,37E-11 | 6,17E-10 |
| 2221 | RM25_RS10835 | RM25_2195 | Transcriptional regulator, DeoR family | 2310 | gene | 2461932 | 2462708 | - | 516,3319 | -0,282656646 | 0,166585 | -1,69677 | 0,089739 | 0,180563 | 538,5841551 | -0,21789113 | 0,15707 | -1,38722 | 0,165375 | 0,220344 | 494,0797 | -0,352951052 | 0,30612 | -1,15298 | 0,248917 | 0,349225 | 598,535 | -0,12567 | 0,207244 | -0,60638 | 0,544261 | 0,609997 | 458,0649001 | -0,198130537 | 0,273765 | -0,72373 | 0,469234 | 0,531979 |
| 2222 | RM25_RS10840 | RM25_2196 | Glycerol-3-phosphate dehydrogenase [NAD(P)+] | 2311 | gene | 2462799 | 2463851 | - | 432,8685 | -0,918881685 | 0,407304 | -2,25601 | 0,02407 | 0,079347 | 270,5313941 | -1,002822273 | 0,187657 | -5,34391 | 9,1E-08 | 5,57E-07 | 595,2055 | -0,883255549 | 0,412384 | -2,14183 | 0,032207 | 0,073063 | 593,705 | 1,034263 | 0,189107 | 5,469195 | 4,52E-08 | 4,46E-07 | 295,2398453 | 1,221474734 | 0,408533 | 2,989904 | 0,002791 | 0,00541 |
| 2223 | RM25_RS10845 | RM25_2197 | Gluconokinase | 2312 | gene | 2463905 | 2465461 | - | 818,2306 | -1,299359562 | 0,709432 | -1,83155 | 0,067019 | 0,149458 | 401,982798 | -3,31897503 | 0,336775 | -9,85516 | 6,51E-23 | 3,07E-21 | 1234,478 | -0,872280918 | 0,421744 | -2,06827 | 0,038614 | 0,083992 | 1218,463 | 1,060355 | 0,367032 | 2,889001 | 0,003865 | 0,010754 | 466,9625912 | 3,581016689 | 0,412252 | 8,686484 | 3,74E-18 | 6,99E-17 |
| 2224 | RM25_RS10850 | RM25_2198 | Anaerobic sulfatase-maturating enzyme | 2313 | gene | 2465890 | 2467329 | - | 548,6816 | -1,097129376 | 0,524827 | -2,09046 | 0,036577 | 0,102658 | 316,128122 | -2,026030871 | 0,285807 | -7,0888 | 1,35E-12 | 1,75E-11 | 781,235 | -0,783959157 | 0,386885 | -2,02634 | 0,04273 | 0,089925 | 784,6093 | 0,892765 | 0,312301 | 2,858672 | 0,004254 | 0,011552 | 344,537432 | 2,206575323 | 0,380691 | 5,796231 | 6,78E-09 | 3,3E-08 |
| 2225 | RM25_RS10855 | RM25_2199 | Aliphatic compound ABC transporter, periplasmic substrate | 2314 | gene | 2467944 | 2468945 | + | 443,4714 | -1,425773804 | 0,644168 | -2,21336 | 0,026873 | 0,084203 | 243,650382 | -3,063793945 | 0,372634 | -8,22199 | 2E-16 | 3,93E-15 | 643,2924 | -0,999666397 | 0,410586 | -2,43473 | 0,014903 | 0,04094 | 677,7283 | 0,909153 | 0,389323 | 2,335217 | 0,019532 | 0,040205 | 237,3935313 | 3,047787643 | 0,427086 | 7,136232 | 9,59E-13 | 8,34E-12 |
| 2226 | RM25_RS12090 | RM25_2200 | ABC transporter, permease protein | 2315 | gene | 2468942 | 2469949 | + | 322,3685 | -1,160508373 | 0,584097 | -1,98684 | 0,04694 | 0,119608 | 184,9425259 | -2,32278918 | 0,377658 | -6,15052 | 7,72E-10 | 6,72E-09 | 459,7945 | -0,791419382 | 0,47145 | -1,67869 | 0,093212 | 0,16359 | 468,0552 | 0,846177 | 0,384266 | 2,202059 | 0,027661 | 0,05292 | 196,836322 | 2,458048296 | 0,494379 | 4,971988 | 6,63E-07 | 2,4E-06 |
| 2227 | RM25_RS10865 | RM25_2201 | ABC-type nitrate/sulfonate/bicarbonate transport system, ı | 2316 | gene | 2469946 | 2470767 | + | 252,857 | -1,349131567 | 0,680095 | -1,98374 | 0,047285 | 0,120357 | 123,7207877 | -2,875736022 | 0,433023 | -6,64106 | 3,11E-11 | 3,32E-10 | 381,9933 | -0,994861881 | 0,465952 | -2,13512 | 0,032751 | 0,073941 | 381,147 | 1,146636 | 0,400219 | 2,865025 | 0,00417 | 0,011362 | 140,8334553 | 3,110922539 | 0,531235 | 5,856015 | 4,74E-09 | 2,35E-08 |
| 2228 | RM25_RS10870 | RM25_2202 | Nitrate transport ATP-binding subunits C and D | 2317 | gene | 2470764 | 2471558 | + | 306,7692 | -0,921711911 | 0,67586 | -1,36376 | 0,172642 | 0,276064 | 136,4693774 | -2,392650084 | 0,381215 | -6,27637 | 3,47E-10 | 3,11E-09 | 477,0689 | -0,594975076 | 0,440846 | -1,34962 | 0,177138 | 0,268843 | 420,0283 | 1,254911 | 0,370621 | 3,385973 | 0,000709 | 0,002645 | 209,3390016 | 3,126422816 | 0,461105 | 6,780289 | 1,2E-11 | 8,78E-11 |
| 2229 | RM25_RS10875 | RM25_2203 | N-acetylgalactosamine-6-sulfatase (Precursor) | 2318 | gene | 2471594 | 2473216 | + | 561,6847 | -1,18511837 | 0,511061 | -2,31894 | 0,020399 | 0,072518 | 329,4853408 | -1,946467357 | 0,307198 | -6,3362 | 2,36E-10 | 2,19E-09 | 793,8841 | -0,915295846 | 0,405281 | -2,25842 | 0,023919 | 0,059034 | 818,8905 | 0,918947 | 0,335505 | 2,738995 | 0,006163 | 0,015754 | 338,4641408 | 2,023936095 | 0,402468 | 5,028811 | 4,94E-07 | 1,82E-06 |
| 2230 | RM25_RS10880 | RM25_2204 | SDS hydrolase SdsA1 | 2319 | gene | 2473293 | 2475311 | + | 659,8841 | -1,00586861 | 0,345779 | -2,909 | 0,003626 | 0,02353 | 469,1880793 | -1,103536692 | 0,300137 | -3,67677 | 0,000236 | 0,000745 | 850,5801 | -0,954554039 | 0,306901 | -3,1103 | 0,001869 | 0,008158 | 925,1463 | 0,745271 | 0,332949 | 2,238394 | 0,025195 | 0,049038 | 431,6027682 | 0,959153382 | 0,305511 | 3,139503 | 0,001692 | 0,003401 |
| 2231 | NA | RM25_2205 | Hypothetical protein | 2320 | gene | 2475260 | 2476069 | - | 227,2466 | -1,162312649 | 0,594883 | -1,95385 | 0,050719 | 0,12544 | 132,6347354 | -2,610121428 | 0,395912 | -6,59269 | 4,32E-11 | 4,52E-10 | 321,8585 | -0,72019351 | 0,401415 | -1,79414 | 0,072791 | 0,13467 | 330,0842 | 0,741131 | 0,378873 | 1,956146 | 0,050448 | 0,085605 | 138,3239098 | 2,706136541 | 0,451096 | 5,999024 | 1,99E-09 | 1,06E-08 |
| 2232 | RM25_RS12815 | RM25_2206 | Hypothetical protein | 2321 | gene | 2476111 | 2476422 | + | 178,4743 | -1,332500326 | 0,809281 | -1,64652 | 0,099656 | 0,192532 | 89,4999886 | -4,256580891 | 0,521798 | -8,15752 | 3,42E-16 | 6,3E-15 | 267,4485 | -0,815021513 | 0,468393 | -1,74004 | 0,081852 | 0,147836 | 267,6771 | 0,931227 | 0,416716 | 2,234677 | 0,025439 | 0,04943 | 100,2129297 | 4,449246163 | 0,596741 | 7,455904 | 8,93E-14 | 9,17E-13 |
| 2233 | RM25_RS10890 | RM25_2207 | H+ antiporter-2 family protein | 2322 | gene | 2476486 | 2477838 | - | 972,2982 | -0,371624769 | 0,163846 | -2,26814 | 0,023321 | 0,078136 | 1046,995334 | -0,301324536 | 0,118608 | -2,54052 | 0,011069 | 0,021633 | 897,6011 | -0,453686883 | 0,290566 | -1,56139 | 0,118432 | 0,196443 | 1159,639 | -0,21889 | 0,161543 | -1,35502 | 0,17541 | 0,236117 | 833,7039162 | -0,307166444 | 0,260972 | -1,17701 | 0,239191 | 0,296723 |
| 2234 | RM25_RS12095 | RM25_2208 | Hypothetical protein | 2323 | gene | 2478201 | 2479451 | - | 496,7966 | -0,251606303 | 0,22064 | -1,14035 | 0,254141 | 0,362159 | 478,0450394 | -0,165773799 | 0,168912 | -0,98142 | 0,326384 | 0,3939 | 515,5481 | -0,331944598 | 0,393819 | -0,84289 | 0,399292 | 0,504899 | 569,0385 | 0,120611 | 0,209463 | 0,575809 | 0,564744 | 0,629363 | 446,6396875 | 0,023810483 | 0,365621 | 0,065123 | 0,948076 | 0,957008 |
| 2235 | RM25_RS10905 | RM25_2209 | Inosose dehydratase | 2324 | gene | 2479535 | 2480488 | + | 886,9821 | -0,886612993 | 0,259693 | -3,41408 | 0,00064 | 0,007431 | 856,4715042 | -0,557161988 | 0,231023 | -2,41172 | 0,015878 | 0,029514 | 917,4926 | -1,21526526 | 0,39844 | -3,05006 | 0,002288 | 0,009613 | 1211,304 | 0,266323 | 0,276091 | 0,96462 | 0,334735 | 0,409642 | 613,4115012 | -0,320644391 | 0,36733 | -0,87291 | 0,382714 | 0,447893 |
| 2236 | RM25_RS10910 | RM25_2210 | Inositol 2-dehydrogenase | 2325 | gene | 2480542 | 2481552 | + | 720,3323 | -0,584182858 | 0,291992 | -2,00068 | 0,045427 | 0,117018 | 830,6686957 | -0,270113269 | 0,21161 | -1,27647 | 0,20179 | 0,26176 | 609,996 | -1,034886751 | 0,367241 | -2,818 | 0,004832 | 0,017389 | 911,471 | -0,20858 | 0,24181 | -0,86259 | 0,388361 | 0,464687 | 567,4216862 | -0,907178795 | 0,340086 | -2,6675 | 0,007642 | 0,013676 |
| 2237 | RM25_RS10915 | RM25_2211 | Protein IolH | 2326 | gene | 2481771 | 2482637 | + | 563,5222 | -0,933807706 | 0,232247 | -4,02075 | 5,8E-05 | 0,001101 | 584,1193708 | -0,920254448 | 0,189454 | -4,8574 | 1,19E-06 | 5,96E-06 | 542,925 | -0,948203118 | 0,406736 | -2,33125 | 0,01974 | 0,050707 | 780,7428 | -0,15923 | 0,226361 | -0,70345 | 0,481774 | 0,552844 | 381,4230243 | -0,116082563 | 0,384244 | -0,30211 | 0,762571 | 0,804557 |
| 2238 | RM25_RS10920 | RM25_2212 | Transcriptional regulator, LacI family | 2327 | gene | 2482773 | 2483807 | + | 830,6508 | -0,438460902 | 0,192591 | -2,27665 | 0,022807 | 0,077571 | 735,1292001 | -0,607054103 | 0,14976 | -4,0535 | 5,05E-05 | 0,000185 | 926,1724 | -0,308829842 | 0,284559 | -1,08529 | 0,277792 | 0,380673 | 1007,815 | 0,142047 | 0,204659 | 0,694068 | 0,487639 | 0,556863 | 692,9782071 | 0,504973034 | 0,245542 | 2,056565 | 0,039728 | 0,060373 |
| 2239 | RM25_RS10925 | RM25_2213 | Hypothetical protein | 2328 | gene | 2484124 | 2484450 | + | 374,8117 | -0,470420775 | 0,508914 | -0,92436 | 0,355298 | 0,466539 | 204,6401468 | -1,480528518 | 0,283895 | -5,21507 | 1,84E-07 | 1,06E-06 | 544,9832 | -0,133615844 | 0,235843 | -0,56654 | 0,571023 | 0,663333 | 456,3771 | 0,852941 | 0,299931 | 2,843791 | 0,004458 | 0,012009 | 307,7136079 | 2,260097091 | 0,209955 | 10,7647 | 5,05E-27 | 2,38E-25 |
| 2240 | RM25_RS10930 | RM25_2214 | Hydrolase, alpha/beta domain protein | 2329 | gene | 2484695 | 2485633 | + | 1580,192 | 0,570763045 | 0,255906 | 2,230366 | 0,025723 | 0,082377 | 1207,456442 | 0,498042007 | 0,132952 | 3,746037 | 0,00018 | 0,000581 | 1952,927 | 0,616364027 | 0,275233 | 2,239423 | 0,025128 | 0,061104 | 1339,046 | 0,568687 | 0,214179 | 2,655197 | 0,007926 | 0,01942 | 1844,295356 | 0,726237745 | 0,190476 | 3,812745 | 0,000137 | 0,000342 |
| 2241 | RM25_RS10935 | RM25_2215 | Glycerophosphodiester phosphodiesterase | 2330 | gene | 2485848 | 2486657 | + | 885,7446 | -0,451357066 | 0,266109 | -1,69613 | 0,08986 | 0,180563 | 695,2370439 | -0,717106304 | 0,238742 | -3,00369 | 0,002667 | 0,006344 | 1076,252 | -0,286301731 | 0,200637 | -1,42697 | 0,153589 | 0,239583 | 1077,029 | 0,385978 | 0,257207 | 1,500648 | 0,133447 | 0,188683 | 733,7484974 | 0,877306919 | 0,203228 | 4,316862 | 1,58E-05 | 4,54E-05 |
| 2242 | RM25_RS10940 | RM25_2216 | Inner membrane protein yihN | 2331 | gene | 2486852 | 2488216 | + | 1161,305 | -0,638584785 | 0,195128 | -3,27265 | 0,001065 | 0,010377 | 1371,499834 | -0,742786137 | 0,136791 | -5,43008 | 5,63E-08 | 3,58E-07 | 951,1094 | -0,489894075 | 0,198614 | -2,46656 | 0,013642 | 0,038693 | 1500,472 | -0,68909 | 0,174892 | -3,94009 | 8,15E-05 | 0,000407 | 891,800283 | -0,378020121 | 0,165917 | -2,27837 | 0,022704 | 0,036754 |
| 2243 | RM25_RS10945 | RM25_2217 | FAD-binding 9 siderophore-interacting domain protein | 2332 | gene | 2488584 | 2489300 | + | 373,1164 | 0,261407097 | 0,297018 | 0,880106 | 0,378802 | 0,48926 | 283,6280128 | 0,566208546 | 0,278474 | 2,033252 | 0,042027 | 0,068934 | 462,6047 | 0,076497488 | 0,238605 | 0,320602 | 0,748512 | 0,814802 | 355,7831 | 0,911474 | 0,255356 | 3,569426 | 0,000358 | 0,001498 | 399,5588959 | 0,484596766 | 0,275165 | 1,761116 | 0,078219 | 0,110198 |
| 2244 | RM25_RS10950 | RM25_2218 | AP endonuclease, family 2 | 2333 | gene | 2489448 | 2490476 | - | 380,6083 | -1,413801043 | 0,710693 | -1,98933 | 0,046665 | 0,119295 | 207,8643091 | -3,56273104 | 0,424921 | -8,38446 | 5,1E-17 | 1,08E-15 | 553,3522 | -0,918091258 | 0,431495 | -2,1277 | 0,033362 | 0,07478 | 581,1957 | 0,8441 | 0,418333 | 2,017769 | 0,043615 | 0,076262 | 205,2193927 | 3,569292079 | 0,47877 | 7,455128 | 8,98E-14 | 9,17E-13 |
| 2245 | RM25_RS10955 | RM25_2219 | Oxidoreductase, NAD-binding Rossmann fold family protein | 2334 | gene | 2490600 | 2491787 | - | 534,777 | -1,113664669 | 0,728958 | -1,52775 | 0,126575 | 0,224791 | 264,8039359 | -3,337830399 | 0,355153 | -9,39828 | 5,55E-21 | 2,01E-19 | 804,7502 | -0,643487874 | 0,474154 | -1,35713 | 0,17474 | 0,26589 | 766,1256 | 0,955947 | 0,3782 | 2,527625 | 0,011484 | 0,02633 | 334,3304295 | 3,727686713 | 0,468685 | 7,95351 | 1,81E-15 | 2,44E-14 |
| 2246 | RM25_RS10960 | RM25_2220 | Transcriptional regulator, LacI family | 2335 | gene | 2491919 | 2492962 | - | 360,6988 | -0,413627529 | 0,318575 | -1,29837 | 0,194161 | 0,298027 | 267,913869 | -0,42101784 | 0,182972 | -2,301 | 0,021392 | 0,037825 | 453,4838 | -0,411498844 | 0,400063 | -1,02859 | 0,303674 | 0,407841 | 432,7898 | 0,690243 | 0,199499 | 3,459889 | 0,00054 | 0,002105 | 304,7429877 | 0,7695852 | 0,389183 | 1,977436 | 0,047992 | 0,071368 |
| 2247 | RM25_RS10965 | RM25_2221 | Sulfurtransferase | 2336 | gene | 2492989 | 2493807 | - | 1436,763 | 0,131650146 | 0,17343 | 0,759095 | 0,447796 | 0,554627 | 1533,638489 | 0,384067744 | 0,162925 | 2,357327 | 0,018407 | 0,033451 | 1339,887 | -0,152930641 | 0,254363 | -0,60123 | 0,547686 | 0,645126 | 1450,088 | 0,02976 | 0,266765 | 0,111558 | 0,911174 | 0,929713 | 1471,788625 | -0,464114721 | 0,130543 | -3,55527 | 0,000378 | 0,000867 |
| 2248 | RM25_RS10970 | RM25_2222 | Glutamate decarboxylase | 2337 | gene | 2494220 | 2495632 | + | 1269,499 | -0,617224843 | 0,385469 | -1,60123 | 0,109325 | 0,203432 | 852,669085 | -1,213893111 | 0,258101 | -4,70318 | 2,56E-06 | 1,21E-05 | 1686,328 | -0,338283295 | 0,209103 | -1,61778 | 0,105709 | 0,179767 | 1616,313 | 0,592674 | 0,313394 | 1,891145 | 0,058605 | 0,096731 | 981,5220162 | 1,52850567 | 0,158916 | 9,618316 | 6,69E-22 | 1,81E-20 |
| 2249 | RM25_RS10975 | RM25_2223 | Glutamate/gamma-aminobutyrate antiporter | 2338 | gene | 2495822 | 2497219 | + | 709,9543 | -0,290988476 | 0,253274 | -1,14891 | 0,250595 | 0,358623 | 569,3243183 | -0,689852626 | 0,195802 | -3,52322 | 0,000426 | 0,001264 | 850,5844 | -0,033644286 | 0,204979 | -0,16414 | 0,869625 | 0,906124 | 824,2404 | 0,222922 | 0,233439 | 0,954949 | 0,339603 | 0,414953 | 625,7948612 | 0,940806278 | 0,173264 | 5,429887 | 5,64E-08 | 2,43E-07 |
| 2250 | RM25_RS10980 | RM25_2224 | Putative UDP-glucose 4-epimerase | 2339 | gene | 2497268 | 2498167 | - | 545,9652 | -0,354396661 | 0,220762 | -1,60533 | 0,108421 | 0,202494 | 443,2971643 | -0,434146916 | 0,200739 | -2,16274 | 0,030561 | 0,052009 | 648,6333 | -0,302357192 | 0,212753 | -1,42116 | 0,155269 | 0,241724 | 644,7845 | 0,426116 | 0,21625 | 1,970483 | 0,048783 | 0,08332 | 470,2083088 | 0,618708384 | 0,204272 | 3,028841 | 0,002455 | 0,00483 |
| 2251 | RM25_RS10985 | RM25_2225 | Major facilitator superfamily (Precursor) | 2340 | gene | 2498480 | 2499748 | + | 633,3337 | -1,516594661 | 0,603466 | -2,51314 | 0,011966 | 0,052327 | 373,2130095 | -3,051544277 | 0,357434 | -8,53737 | 1,37E-17 | 3,08E-16 | 893,4544 | -1,073997008 | 0,333244 | -3,22285 | 0,001269 | 0,005889 | 984,9302 | 0,794499 | 0,391269 | 2,030568 | 0,042299 | 0,07407 | 322,9633005 | 2,842485435 | 0,339939 | 8,361742 | 6,18E-17 | 9,71E-16 |
| 2252 | RM25_RS12100 | RM25_2226 | Conserved transmembrane protein | 2341 | gene | 2499815 | 2501344 | - | 1343,088 | -0,216644577 | 0,200888 | -1,07844 | 0,280839 | 0,389833 | 1227,739857 | -0,730522388 | 0,133519 | -5,47129 | 4,47E-08 | 2,89E-07 | 1458,437 | 0,202274317 | 0,19016 | 1,063704 | 0,287463 | 0,391945 | 1527,193 | -0,23573 | 0,143741 | -1,63996 | 0,101014 | 0,150055 | 1215,252455 | 0,746148618 | 0,152091 | 4,905932 | 9,3E-07 | 3,27E-06 |
| 2253 | RM25_RS11000 | RM25_2227 | Putative oxydoreductase | 2342 | gene | 2501560 | 2502387 | - | 1196,843 | 0,181426075 | 0,190284 | 0,953451 | 0,340362 | 0,451454 | 1082,954268 | 0,268288123 | 0,158594 | 1,691663 | 0,09071 | 0,130289 | 1310,731 | 0,111042041 | 0,309775 | 0,35846 | 0,719999 | 0,791898 | 1185,249 | 0,304599 | 0,277986 | 1,095733 | 0,273196 | 0,346195 | 1242,885748 | 0,18629437 | 0,199474 | 0,933929 | 0,350341 | 0,414119 |
| 2254 | RM25_RS11005 | RM25_2228 | Hypothetical protein | 2343 | gene | 2502777 | 2503178 | + | 229,3792 | -0,396080135 | 0,192659 | -2,05586 | 0,039796 | 0,108314 | 210,5966182 | -0,662600176 | 0,189995 | -3,48745 | 0,000488 | 0,001415 | 248,1618 | -0,182740386 | 0,293876 | -0,62183 | 0,534055 | 0,631876 | 275,3716 | -0,03333 | 0,241431 | -0,13803 | 0,890215 | 0,912674 | 194,4715269 | 0,511654299 | 0,265532 | 1,926903 | 0,053992 | 0,079141 |
| 2255 | RM25_RS11010 | RM25_2229 | Osmoprotectant proV | 2344 | gene | 2503506 | 2504396 | + | 631,2106 | -0,491518108 | 0,202247 | -2,43029 | 0,015087 | 0,060889 | 542,8885407 | -0,500774899 | 0,194365 | -2,57647 | 0,009981 | 0,01987 | 719,5327 | -0,486605973 | 0,271014 | -1,7955 | 0,072574 | 0,134373 | 776,66 | 0,333855 | 0,250045 | 1,33518 | 0,181817 | 0,243076 | 515,4325724 | 0,412379784 | 0,229511 | 1,796773 | 0,072372 | 0,10307 |
| 2256 | RM25_RS11015 | RM25_2230 | ABC transporter, permease protein | 2345 | gene | 2504393 | 2505040 | + | 440,6668 | -0,07674885 | 0,106526 | -0,72047 | 0,471236 | 0,571644 | 447,688863 | -0,072781253 | 0,142844 | -0,50951 | 0,610392 | 0,66916 | 433,6448 | -0,079100005 | 0,226986 | -0,34848 | 0,72748 | 0,798264 | 477,4989 | -0,10454 | 0,173175 | -0,60366 | 0,546071 | 0,611153 | 421,1264046 | -0,051287057 | 0,194764 | -0,26333 | 0,792297 | 0,831636 |
| 2257 | RM25_RS11020 | RM25_2231 | Osmoprotectant transport integral membrane protein ABC t | 2346 | gene | 2505037 | 2505717 | + | 429,3079 | -0,543297847 | 0,236944 | -2,29294 | 0,021851 | 0,075137 | 364,2864106 | -0,478772081 | 0,247095 | -1,9376 | 0,052672 | 0,08249 | 494,3294 | -0,593187861 | 0,30828 | -1,92418 | 0,054332 | 0,107163 | 536,0739 | 0,417101 | 0,299527 | 1,392529 | 0,163762 | 0,2226 | 343,558659 | 0,37226972 | 0,280802 | 1,325737 | 0,184927 | 0,237662 |
| 2258 | RM25_RS11025 | RM25_2232 | Putative alcohol dehydrogenase AdhA | 2347 | gene | 2505792 | 2506811 | - | 1095,603 | 0,250395456 | 0,12452 | 2,010881 | 0,044338 | 0,114967 | 1142,969415 | 0,396343735 | 0,127855 | 3,099949 | 0,001936 | 0,004782 | 1048,236 | 0,095038871 | 0,22666 | 0,419302 | 0,674996 | 0,754582 | 1056,773 | -0,0182 | 0,184234 | -0,0988 | 0,921296 | 0,937615 | 1165,69264 | -0,275578024 | 0,152276 | -1,80972 | 0,070339 | 0,100417 |
| 2259 | RM25_RS11030 | RM25_2233 | Methionine aminotransferase | 2348 | gene | 2506881 | 2508071 | - | 1200,644 | -0,169925518 | 0,164757 | -1,03137 | 0,302366 | 0,411951 | 1045,070787 | -0,269229159 | 0,144169 | -1,86746 | 0,061838 | 0,093611 | 1356,217 | -0,095392834 | 0,204144 | -0,46728 | 0,640299 | 0,722279 | 1339,889 | 0,231686 | 0,202162 | 1,146038 | 0,251779 | 0,321824 | 1106,24185 | 0,457202195 | 0,116794 | 3,91459 | 9,06E-05 | 0,000232 |
| 2260 | NA | RM25_2234 | Hypothetical protein | 2349 | gene | 2508164 | 2508328 | - | 140,1416 | -0,471141905 | 0,235709 | -1,99883 | 0,045627 | 0,117277 | 112,5628339 | -0,56808954 | 0,278022 | -2,04332 | 0,04102 | 0,067423 | 167,7203 | -0,414364557 | 0,216364 | -1,91513 | 0,055476 | 0,109055 | 171,1572 | 0,450228 | 0,23385 | 1,925286 | 0,054194 | 0,090662 | 114,9515248 | 0,655313918 | 0,252164 | 2,598759 | 0,009356 | 0,016433 |
| 2261 | RM25_RS11035 | RM25_2235 | Hypothetical protein | 2350 | gene | 2508550 | 2508753 | - | 263,7818 | 0,44814843 | 0,219459 | 2,042063 | 0,041145 | 0,109896 | 222,6592773 | 0,041252523 | 0,183548 | 0,224751 | 0,822173 | 0,858982 | 304,9043 | 0,749747994 | 0,265902 | 2,819641 | 0,004808 | 0,017327 | 235,773 | -0,01217 | 0,193615 | -0,06283 | 0,949899 | 0,961319 | 297,1604235 | 0,741503378 | 0,228165 | 3,249849 | 0,001155 | 0,002402 |
| 2262 | RM25_RS11040 | RM25_2236 | Hypothetical protein | 2351 | gene | 2508891 | 2509673 | + | 876,7487 | -0,4245843 | 0,279851 | -1,51718 | 0,129221 | 0,227804 | 660,0198032 | -0,744112895 | 0,197394 | -3,76969 | 0,000163 | 0,00053 | 1093,478 | -0,239879693 | 0,18964 | -1,26492 | 0,2059 | 0,302689 | 1057,215 | 0,453982 | 0,237938 | 1,907985 | 0,056393 | 0,093803 | 732,7521271 | 1,013323506 | 0,125705 | 8,061144 | 7,56E-16 | 1,05E-14 |
| 2263 | RM25_RS11045 | RM25_2237 | Protein of hypothetical function (DUF1697) | 2352 | gene | 2509688 | 2510293 | + | 205,7682 | -0,050939164 | 0,319022 | -0,15967 | 0,873139 | 0,913204 | 149,9668097 | -0,383416989 | 0,22437 | -1,70886 | 0,087477 | 0,126105 | 261,5697 | 0,133763965 | 0,33508 | 0,3992 | 0,689746 | 0,766854 | 219,7258 | 0,493538 | 0,258066 | 1,912454 | 0,055818 | 0,092977 | 198,6229014 | 1,072489123 | 0,30559 | 3,509574 | 0,000449 | 0,001014 |
| 2264 | RM25_RS12105 | RM25_2238 | MFS permease | 2353 | gene | 2510450 | 2511349 | - | 434,2643 | 0,17605506 | 0,149264 | 1,179486 | 0,238205 | 0,34572 | 449,5087525 | 0,494798732 | 0,171696 | 2,881833 | 0,003954 | 0,008892 | 419,0198 | -0,157213403 | 0,183456 | -0,85695 | 0,391471 | 0,497679 | 429,7191 | 0,181209 | 0,184657 | 0,98133 | 0,32643 | 0,401773 | 452,9117528 | -0,412988326 | 0,183778 | -2,24721 | 0,024626 | 0,039594 |
| 2265 | RM25_RS12110 | RM25_2239 | Major Facilitator Superfamily protein | 2354 | gene | 2511304 | 2512110 | - | 1346,463 | 0,348006685 | 0,226169 | 1,538703 | 0,123877 | 0,221531 | 1147,24412 | 0,486733426 | 0,161924 | 3,005933 | 0,002648 | 0,006316 | 1545,682 | 0,24716149 | 0,32059 | 0,770959 | 0,440731 | 0,544722 | 1250,338 | 0,50982 | 0,309495 | 1,647264 | 0,099504 | 0,148156 | 1474,250113 | 0,311439285 | 0,175111 | 1,778528 | 0,075317 | 0,106556 |
| 2266 | RM25_RS11055 | RM25_2240 | Sn-glycerol-3-phosphate transport membrane protein ABC t | 2355 | gene | 2512387 | 2513394 | + | 564,7966 | -1,703748289 | 0,846534 | -2,01262 | 0,044155 | 0,114744 | 304,6074504 | -5,095812704 | 0,447667 | -11,383 | 5,08E-30 | 5,44E-28 | 824,9858 | -1,147799566 | 0,427324 | -2,68602 | 0,007231 | 0,023971 | 906,1986 | 0,871292 | 0,413382 | 2,10772 | 0,035055 | 0,06445 | 262,0544088 | 4,893341689 | 0,50325 | 9,723485 | 2,39E-22 | 6,88E-21 |
| 2267 | RM25_RS11060 | RM25_2241 | Sn-glycerol-3-phosphate transport membrane protein ABC t | 2356 | gene | 2513391 | 2514371 | + | 439,9874 | -1,635629972 | 0,843009 | -1,94023 | 0,052352 | 0,128002 | 224,6818561 | -4,935393697 | 0,441421 | -11,1807 | 5,07E-29 | 4,6E-27 | 655,293 | -1,11354841 | 0,422814 | -2,63366 | 0,008447 | 0,026868 | 697,0114 | 0,971901 | 0,400107 | 2,429103 | 0,015136 | 0,032972 | 211,5276027 | 4,870699085 | 0,496212 | 9,815766 | 9,63E-23 | 2,87E-21 |
| 2268 | RM25_RS11065 | RM25_2242 | Sn-glycerol-3-phosphate-binding periplasmic protein UgpB | 2357 | gene | 2514404 | 2515837 | + | 442,4036 | -1,041796056 | 0,786729 | -1,32421 | 0,185433 | 0,290796 | 201,6297616 | -3,741408616 | 0,361381 | -10,3531 | 4,05E-25 | 2,51E-23 | 683,1774 | -0,568926469 | 0,446806 | -1,27332 | 0,202905 | 0,30003 | 622,403 | 1,052202 | 0,360103 | 2,921944 | 0,003479 | 0,009962 | 285,6618977 | 4,296900037 | 0,457644 | 9,389179 | 6,05E-21 | 1,55E-19 |
| 2269 | RM25_RS11070 | RM25_2243 | Drug resistance transporter, EmrB/QacA subfamily | 2358 | gene | 2516082 | 2517623 | + | 464,4367 | -1,527458137 | 0,781183 | -1,95531 | 0,050546 | 0,12544 | 251,6277634 | -4,298636448 | 0,416349 | -10,3246 | 5,46E-25 | 3,3E-23 | 677,2457 | -0,987966532 | 0,463455 | -2,13174 | 0,033028 | 0,074353 | 723,9435 | 0,838292 | 0,406095 | 2,064277 | 0,038991 | 0,069411 | 236,6088762 | 4,229771415 | 0,5075 | 8,33453 | 7,78E-17 | 1,2E-15 |
| 2270 | RM25_RS11075 | RM25_2244 | 50S ribosomal protein L9 | 2359 | gene | 2517752 | 2518204 | - | 2803,017 | 0,935226176 | 0,542589 | 1,723638 | 0,084773 | 0,173597 | 3766,058532 | 0,518675094 | 0,211777 | 2,449158 | 0,014319 | 0,027022 | 1839,976 | 1,958648588 | 0,614594 | 3,186898 | 0,001438 | 0,006556 | 2060,708 | -2,08681 | 0,612048 | -3,40955 | 0,000651 | 0,002462 | 3604,188176 | -0,614486353 | 0,221561 | -2,77344 | 0,005547 | 0,01019 |
| 2271 | RM25_RS11080 | RM25_2245 | Hypothetical protein | 2360 | gene | 2518216 | 2518455 | - | 2799,718 | 0,808195322 | 0,488764 | 1,65355 | 0,098219 | 0,190851 | 3601,794787 | 0,295773676 | 0,198789 | 1,487875 | 0,136784 | 0,188428 | 1997,641 | 1,917306213 | 0,520372 | 3,684488 | 0,000229 | 0,001469 | 2176,227 | -2,00152 | 0,485529 | -4,12235 | 3,75E-05 | 0,000201 | 3483,021825 | -0,349214611 | 0,266901 | -1,3084 | 0,190736 | 0,244329 |
| 2272 | RM25_RS11085 | RM25_2246 | Single-stranded DNA-binding protein 2 | 2361 | gene | 2518557 | 2519123 | - | 2906,964 | 0,913658313 | 0,404371 | 2,259454 | 0,023855 | 0,079134 | 3567,257318 | 0,497692734 | 0,213205 | 2,334339 | 0,019578 | 0,035038 | 2246,67 | 1,671512367 | 0,458159 | 3,648324 | 0,000264 | 0,001655 | 2149,591 | -1,5121 | 0,441557 | -3,42448 | 0,000616 | 0,002353 | 3714,813648 | -0,30480718 | 0,236917 | -1,28656 | 0,198248 | 0,252442 |
| 2273 | RM25_RS11090 | RM25_2247 | 30S ribosomal protein S6 | 2362 | gene | 2519304 | 2519591 | - | 4119,414 | 0,764863366 | 0,537371 | 1,423344 | 0,154637 | 0,25575 | 5605,34348 | 0,357859223 | 0,202919 | 1,763557 | 0,077807 | 0,11412 | 2633,485 | 1,776074544 | 0,597946 | 2,970294 | 0,002975 | 0,011912 | 3265,299 | -2,09199 | 0,587705 | -3,5596 | 0,000371 | 0,001538 | 5077,572372 | -0,643120828 | 0,23269 | -2,76385 | 0,005712 | 0,01047 |
| 2274 | RM25_RS11095 | RM25_2248 | Hypothetical protein | 2363 | gene | 2520262 | 2520558 | + | 158,7308 | -0,921847523 | 0,312786 | -2,94721 | 0,003207 | 0,021722 | 120,6698244 | -1,328957947 | 0,306793 | -4,33177 | 1,48E-05 | 6,05E-05 | 196,7917 | -0,699185285 | 0,230894 | -3,02817 | 0,00246 | 0,010246 | 218,9152 | 0,428388 | 0,261073 | 1,640876 | 0,100823 | 0,149931 | 107,4840194 | 1,120882734 | 0,308698 | 3,630998 | 0,000282 | 0,000664 |
| 2275 | RM25_RS11100 | RM25_2249 | Inorganic diphosphatase | 2364 | gene | 2520630 | 2520989 | - | 681,8164 | -0,826238509 | 0,380708 | -2,17027 | 0,029987 | 0,090266 | 435,8005425 | -0,528140771 | 0,204258 | -2,58566 | 0,009719 | 0,01943 | 927,8322 | -0,972249585 | 0,385546 | -2,52175 | 0,011677 | 0,034319 | 917,332 | 1,203043 | 0,359076 | 3,350386 | 0,000807 | 0,002926 | 480,1467655 | 0,79710651 | 0,25456 | 3,131307 | 0,00174 | 0,003491 |
| 2276 | RM25_RS11105 | RM25_2250 | Chlorite O(2)-lyase | 2365 | gene | 2521580 | 2522260 | - | 1072,763 | 1,280332827 | 0,211603 | 6,050627 | 1,44E-09 | 1,62E-07 | 901,0373824 | 1,169832791 | 0,169746 | 6,89168 | 5,51E-12 | 6,56E-11 | 1244,488 | 1,361132207 | 0,292535 | 4,652888 | 3,27E-06 | 3,84E-05 | 659,859 | 0,264809 | 0,197368 | 1,341704 | 0,179692 | 0,240891 | 1483,081635 | 0,502855462 | 0,238859 | 2,10524 | 0,03527 | 0,054193 |
| 2277 | RM25_RS11110 | RM25_2251 | Hypothetical protein | 2366 | gene | 2522446 | 2522835 | - | 373,2211 | 0,006061936 | 0,116759 | 0,051918 | 0,958594 | 0,973882 | 362,3152027 | 0,226055867 | 0,158277 | 1,428227 | 0,153227 | 0,20756 | 384,1269 | -0,195856748 | 0,186577 | -1,04974 | 0,293839 | 0,397121 | 392,5551 | 0,234716 | 0,145994 | 1,60771 | 0,107899 | 0,157961 | 367,1737893 | -0,130174323 | 0,194296 | -0,66998 | 0,502871 | 0,564144 |
| 2278 | RM25_RS12115 | RM25_2252 | Isochorismatase hydrolase | 2367 | gene | 2522979 | 2523710 | + | 714,8466 | -0,48011336 | 0,229308 | -2,09375 | 0,036282 | 0,102256 | 556,7040074 | -0,329418055 | 0,153424 | -2,14711 | 0,031785 | 0,053781 | 872,9892 | -0,578144388 | 0,198959 | -2,90585 | 0,003663 | 0,014175 | 875,6898 | 0,69322 | 0,194103 | 3,571396 | 0,000355 | 0,001489 | 585,4207489 | 0,499521181 | 0,16535 | 3,020986 | 0,00252 | 0,004932 |
| 2279 | RM25_RS12825 | RM25_2253 | Hypothetical protein | 2368 | gene | 2523956 | 2527324 | + | 6211,725 | 0,222856161 | 0,52054 | 0,428125 | 0,66856 | 0,750022 | 3807,90857 | -1,588623411 | 0,155704 | -10,2028 | 1,93E-24 | 1,13E-22 | 8615,541 | 0,997192117 | 0,28916 | 3,448584 | 0,000564 | 0,003089 | 6051,913 | -0,05626 | 0,210542 | -0,2672 | 0,789315 | 0,828693 | 6511,471606 | 2,577643597 | 0,210731 | 12,23191 | 2,1E-34 | 1,6E-32 |
| 2280 | RM25_RS11125 | RM25_2254 | Serine protease, subtilase family | 2369 | gene | 2527495 | 2528331 | + | 616,9776 | -1,021426762 | 0,683126 | -1,49522 | 0,134856 | 0,233889 | 338,5211403 | -3,293877382 | 0,344975 | -9,54816 | 1,32E-21 | 5,02E-20 | 895,434 | -0,467637227 | 0,297936 | -1,56959 | 0,116511 | 0,194075 | 865,8701 | 0,691967 | 0,368906 | 1,875729 | 0,060693 | 0,099342 | 399,5886375 | 3,580035237 | 0,279724 | 12,79848 | 1,67E-37 | 1,71E-35 |
| 2281 | RM25_RS11130 | RM25_2255 | High-affinity glucose transporter | 2370 | gene | 2528551 | 2529804 | + | 820,638 | -0,669343728 | 0,356027 | -1,88004 | 0,060103 | 0,139408 | 581,4416778 | -0,875591097 | 0,225696 | -3,87951 | 0,000105 | 0,000354 | 1059,834 | -0,560168192 | 0,388997 | -1,44003 | 0,149858 | 0,234851 | 1057,706 | 0,680985 | 0,258057 | 2,638895 | 0,008318 | 0,020274 | 624,1240574 | 1,066836505 | 0,362422 | 2,943628 | 0,003244 | 0,006216 |
| 2282 | RM25_RS11135 | RM25_2256 | DNA-3-methyladenine glycosylase I | 2371 | gene | 2529896 | 2530522 | + | 409,5913 | -1,257172959 | 0,515978 | -2,43649 | 0,014831 | 0,060688 | 259,8959642 | -2,198415423 | 0,361577 | -6,08008 | 1,2E-09 | 1,02E-08 | 559,2866 | -0,901480971 | 0,413629 | -2,17944 | 0,029299 | 0,068037 | 606,7392 | 0,69975 | 0,391226 | 1,788608 | 0,073678 | 0,116079 | 238,2759764 | 2,073921455 | 0,420822 | 4,928267 | 8,3E-07 | 2,94E-06 |
| 2283 | RM25_RS11140 | RM25_2257 | Protein RtcB | 2372 | gene | 2530904 | 2532112 | + | 940,0675 | -0,618941343 | 0,42181 | -1,46734 | 0,142282 | 0,24319 | 593,8148608 | -1,044951041 | 0,294019 | -3,55403 | 0,000379 | 0,001141 | 1286,32 | -0,433070413 | 0,326455 | -1,32659 | 0,184646 | 0,278089 | 1194,376 | 0,816114 | 0,342912 | 2,379952 | 0,017315 | 0,036569 | 729,4643842 | 1,498294078 | 0,300326 | 4,988886 | 6,07E-07 | 2,21E-06 |
| 2284 | RM25_RS11145 | RM25_2258 | Alpha amylase, catalytic domain protein | 2373 | gene | 2532142 | 2533410 | - | 996,2082 | -0,558792564 | 0,150372 | -3,71607 | 0,000202 | 0,002944 | 912,575123 | -0,509692315 | 0,126758 | -4,021 | 5,8E-05 | 0,00021 | 1079,841 | -0,601796772 | 0,262159 | -2,29555 | 0,021702 | 0,054591 | 1251,267 | 0,215841 | 0,166219 | 1,298534 | 0,194104 | 0,257024 | 791,530616 | 0,186305724 | 0,235675 | 0,79052 | 0,429224 | 0,493985 |
| 2285 | RM25_RS11150 | RM25_2259 | Mannosyltransferase PIG-V domain protein | 2374 | gene | 2533520 | 2534674 | - | 990,8239 | 0,043073919 | 0,170372 | 0,252823 | 0,800405 | 0,859841 | 1119,485287 | 0,076314357 | 0,13307 | 0,573491 | 0,566312 | 0,629028 | 862,1626 | 0,001615473 | 0,239084 | 0,006757 | 0,994609 | 0,996621 | 1033,763 | -0,40354 | 0,190649 | -2,11667 | 0,034288 | 0,063187 | 987,1321314 | -0,417434677 | 0,191061 | -2,18482 | 0,028902 | 0,04572 |
| 2286 | RM25_RS11155 | RM25_2260 | Hypothetical protein | 2375 | gene | 2534671 | 2536329 | - | 1392,294 | 0,206399043 | 0,183613 | 1,1241 | 0,260971 | 0,36877 | 1599,756232 | 0,265066728 | 0,136547 | 1,941213 | 0,052232 | 0,08202 | 1184,832 | 0,129289025 | 0,234838 | 0,550544 | 0,581946 | 0,673651 | 1367,973 | -0,42296 | 0,182387 | -2,31905 | 0,020392 | 0,041519 | 1463,80837 | -0,501379431 | 0,181623 | -2,76055 | 0,00577 | 0,010568 |
| 2287 | RM25_RS11160 | RM25_2261 | Bifunctional penicillin-binding protein ponA1 | 2376 | gene | 2536326 | 2538461 | - | 2442,118 | 0,119313889 | 0,144819 | 0,823883 | 0,410006 | 0,520538 | 2464,450491 | -0,190720574 | 0,133793 | -1,42549 | 0,154014 | 0,208344 | 2419,786 | 0,437154896 | 0,201131 | 2,17348 | 0,029744 | 0,068935 | 2477,236 | -0,41528 | 0,173037 | -2,39997 | 0,016396 | 0,035069 | 2490,485615 | 0,264668473 | 0,132277 | 2,00087 | 0,045406 | 0,067865 |
| 2288 | RM25_RS11165 | RM25_2262 | Hypothetical protein | 2377 | gene | 2538454 | 2538792 | - | 708,7712 | 0,441328408 | 0,226515 | 1,948339 | 0,051374 | 0,126398 | 721,2701327 | -0,02598036 | 0,201767 | -0,12876 | 0,897544 | 0,922998 | 696,2722 | 0,950752146 | 0,273192 | 3,480158 | 0,000501 | 0,002793 | 638,1781 | -0,67399 | 0,206626 | -3,26189 | 0,001107 | 0,003842 | 796,6662317 | 0,343750304 | 0,239667 | 1,434283 | 0,151491 | 0,199925 |
| 2290 | NA | RM25_2264 | Cassette chromosome recombinase B | 2381 | gene | 2539195 | 2540529 | - | 1317,581 | -1,535628976 | 0,631657 | -2,43111 | 0,015052 | 0,060889 | 799,5749183 | -3,395826281 | 0,323837 | -10,4862 | 9,99E-26 | 6,37E-24 | 1835,587 | -1,0198319 | 0,341366 | -2,98751 | 0,002813 | 0,011497 | 2056,608 | 0,680349 | 0,383747 | 1,77291 | 0,076244 | 0,118618 | 665,1201315 | 3,129254751 | 0,311528 | 10,04484 | 9,68E-24 | 3,17E-22 |
| 2291 | RM25_RS11180 | RM25_2265 | Transposase | 2382 | gene | 2540630 | 2542009 | - | 93,63926 | -1,729622702 | 0,618969 | -2,79436 | 0,0052 | 0,03049 | 56,17259783 | -3,359259025 | 0,499844 | -6,72061 | 1,81E-11 | 2,01E-10 | 131,1059 | -1,276741568 | 0,363639 | -3,51102 | 0,000446 | 0,00256 | 151,2316 | 0,77983 | 0,360604 | 2,162567 | 0,030575 | 0,057467 | 42,58551974 | 2,938813971 | 0,561452 | 5,234307 | 1,66E-07 | 6,51E-07 |
| 2292 | RM25_RS11185 | RM25_2266 | Ferritin-like protein | 2383 | gene | 2542566 | 2543099 | - | 15598,8 | -0,890647405 | 0,302567 | -2,94364 | 0,003244 | 0,021845 | 17360,14091 | -1,00695804 | 0,258893 | -3,88947 | 0,0001 | 0,000342 | 13837,45 | -0,748517838 | 0,460365 | -1,62592 | 0,103966 | 0,177185 | 21574,12 | -0,47024 | 0,441459 | -1,0652 | 0,286787 | 0,360702 | 10680,14545 | -0,179029568 | 0,297231 | -0,60232 | 0,546958 | 0,608104 |
| 2293 | RM25_RS11190 | RM25_2267 | Transcriptional regulator type FNR | 2384 | gene | 2543223 | 2543912 | - | 734,9986 | 0,514201364 | 0,172913 | 2,973756 | 0,002942 | 0,020514 | 692,427054 | 0,482858181 | 0,226799 | 2,129016 | 0,033253 | 0,05591 | 777,5701 | 0,542204627 | 0,25393 | 2,135253 | 0,03274 | 0,073941 | 639,6123 | 0,082182 | 0,299482 | 0,274412 | 0,783768 | 0,824714 | 847,3633968 | 0,186591942 | 0,173311 | 1,076632 | 0,281645 | 0,342891 |
| 2294 | RM25_RS11195 | RM25_2268 | Hypothetical protein | 2385 | gene | 2544101 | 2544340 | + | 120,1386 | -1,028047828 | 0,586374 | -1,75323 | 0,079563 | 0,166802 | 68,34834926 | -2,238101436 | 0,505355 | -4,42877 | 9,48E-06 | 4E-05 | 171,9288 | -0,648827962 | 0,373404 | -1,7376 | 0,082281 | 0,148255 | 169,36 | 0,822641 | 0,431228 | 1,907669 | 0,056434 | 0,093805 | 77,81572758 | 2,486278275 | 0,491535 | 5,058194 | 4,23E-07 | 1,57E-06 |
| 2295 | RM25_RS11200 | RM25_2269 | Cadmium-transporting ATPase | 2386 | gene | 2544375 | 2546381 | + | 489,6144 | -0,533398715 | 0,50022 | -1,06633 | 0,286276 | 0,395749 | 279,4817425 | -1,315138126 | 0,287526 | -4,57397 | 4,79E-06 | 2,13E-05 | 699,7471 | -0,248289892 | 0,397701 | -0,62431 | 0,532422 | 0,630612 | 608,0578 | 0,859464 | 0,282301 | 3,044497 | 0,002331 | 0,007181 | 394,494773 | 2,001857401 | 0,405436 | 4,937542 | 7,91E-07 | 2,82E-06 |
| 2296 | RM25_RS11205 | RM25_2270 | Hypothetical protein | 2387 | gene | 2546425 | 2546814 | + | 339,3514 | 0,171640061 | 0,274095 | 0,626206 | 0,53118 | 0,62725 | 246,947095 | 0,236220225 | 0,186023 | 1,269847 | 0,204139 | 0,264517 | 431,7558 | 0,134031106 | 0,186696 | 0,717912 | 0,472812 | 0,574738 | 335,0536 | 0,797977 | 0,159773 | 4,994436 | 5,9E-07 | 4,67E-06 | 352,7257287 | 0,753763178 | 0,200044 | 3,767982 | 0,000165 | 0,000404 |
| 2297 | RM25_RS11210 | RM25_2271 | Putative uroporphyrin-III c-methyltransferase | 2388 | gene | 2546837 | 2547211 | + | 286,4327 | 0,50368177 | 0,188169 | 2,676754 | 0,007434 | 0,038683 | 237,7481447 | 0,410007355 | 0,19545 | 2,097757 | 0,035927 | 0,059717 | 335,1172 | 0,570688494 | 0,174203 | 3,276006 | 0,001053 | 0,005064 | 249,5605 | 0,346618 | 0,174539 | 1,985911 | 0,047043 | 0,080994 | 328,7764253 | 0,554965778 | 0,169247 | 3,279033 | 0,001042 | 0,002196 |
| 2298 | RM25_RS11215 | RM25_2272 | Hypothetical protein | 2389 | gene | 2547347 | 2548417 | + | 630,0791 | -1,637867891 | 0,32685 | -5,01108 | 5,41E-07 | 2,45E-05 | 593,4734721 | -2,392180734 | 0,167934 | -14,2448 | 4,83E-46 | 2,28E-43 | 666,6847 | -1,107593784 | 0,39588 | -2,7978 | 0,005145 | 0,018347 | 1007,454 | -0,19096 | 0,159937 | -1,19397 | 0,23249 | 0,300427 | 301,8434474 | 1,162828074 | 0,395586 | 2,939505 | 0,003287 | 0,006279 |
| 2299 | RM25_RS11220 | RM25_2273 | Putative nitrite reductase, copper-dependent | 2390 | gene | 2548410 | 2551163 | + | 1529,726 | -0,898054378 | 0,246163 | -3,64822 | 0,000264 | 0,003577 | 1353,329374 | -1,357441461 | 0,147635 | -9,19457 | 3,76E-20 | 1,23E-18 | 1706,123 | -0,564131904 | 0,298693 | -1,88867 | 0,058936 | 0,114331 | 2098,585 | 0,0057 | 0,157591 | 0,036169 | 0,971148 | 0,977368 | 1050,46871 | 0,859832802 | 0,271831 | 3,163111 | 0,001561 | 0,003163 |
| 2300 | RM25_RS11225 | RM25_2274 | Transcriptional regulator, ArsR family | 2391 | gene | 2551160 | 2551789 | + | 279,7589 | -0,690957525 | 0,224906 | -3,07221 | 0,002125 | 0,016009 | 251,9823588 | -1,076240475 | 0,189857 | -5,66869 | 1,44E-08 | 1,02E-07 | 307,5355 | -0,399825444 | 0,299326 | -1,33575 | 0,18163 | 0,274249 | 365,0432 | -0,02798 | 0,18395 | -0,15208 | 0,879124 | 0,903998 | 210,1488836 | 0,710687941 | 0,312657 | 2,273056 | 0,023023 | 0,037185 |
| 2301 | RM25_RS11230 | RM25_2275 | Oligopeptide-transport membrane protein ABC transporter ( | 2392 | gene | 2552141 | 2553124 | + | 777,8869 | -1,025264748 | 0,290963 | -3,52369 | 0,000426 | 0,005336 | 635,9556321 | -1,093325954 | 0,255671 | -4,27631 | 1,9E-05 | 7,62E-05 | 919,8182 | -0,9803839 | 0,37954 | -2,58308 | 0,009792 | 0,029743 | 1095,4 | 0,430905 | 0,308912 | 1,39491 | 0,163043 | 0,22175 | 504,7091463 | 0,612859251 | 0,344115 | 1,780972 | 0,074917 | 0,106054 |
| 2302 | RM25_RS11235 | RM25_2276 | IS1096, TnpA family protein | 2393 | gene | 2553133 | 2553450 | - | 29,16387 | -1,093028074 | 0,413282 | -2,64475 | 0,008175 | 0,040909 | 22,17434176 | -1,497440588 | 0,583952 | -2,56432 | 0,010338 | 0,020442 | 36,1534 | -0,886279525 | 0,438615 | -2,02063 | 0,043318 | 0,09064 | 41,93615 | 0,436962 | 0,407006 | 1,073602 | 0,283001 | 0,356702 | 18,24989066 | 1,127866934 | 0,675416 | 1,669885 | 0,094942 | 0,13148 |
| 2303 | RM25_RS11240 | RM25_2277 | Putative insertion element ISCmi2 transposase | 2394 | gene | 2553630 | 2554640 | + | 1559,061 | -0,399086976 | 0,209884 | -1,90147 | 0,057241 | 0,135086 | 1365,049098 | -0,764995336 | 0,194032 | -3,94263 | 8,06E-05 | 0,00028 | 1753,073 | -0,124932453 | 0,197528 | -0,63248 | 0,527074 | 0,626018 | 1872,119 | 0,027303 | 0,165776 | 0,164696 | 0,869183 | 0,897356 | 1316,252791 | 0,720096719 | 0,210128 | 3,426946 | 0,00061 | 0,001351 |
| 2304 | RM25_RS11245 | RM25_2278 | Transmembrane protein | 2395 | gene | 2554637 | 2555029 | - | 1317,565 | 0,706162292 | 0,223474 | 3,159925 | 0,001578 | 0,01296 | 1271,874935 | 0,570898154 | 0,211089 | 2,704533 | 0,00684 | 0,014331 | 1363,255 | 0,8358124 | 0,371057 | 2,252518 | 0,02429 | 0,059823 | 1061,123 | -0,11595 | 0,340724 | -0,3403 | 0,733627 | 0,7789 | 1594,660826 | 0,182682982 | 0,247366 | 0,738514 | 0,460202 | 0,525023 |
| 2305 | NA | RM25_2279 | Hypothetical protein | 2396 | gene | 2555079 | 2555195 | - | 991,4259 | 0,790029163 | 0,245797 | 3,214148 | 0,001308 | 0,01186 | 944,2785133 | 0,529030162 | 0,21344 | 2,478589 | 0,01319 | 0,025133 | 1038,573 | 1,039299309 | 0,394573 | 2,633982 | 0,008439 | 0,026868 | 770,7237 | -0,23765 | 0,33487 | -0,70968 | 0,4779 | 0,5492 | 1225,435057 | 0,305865707 | 0,286676 | 1,066937 | 0,286 | 0,347083 |
| 2306 | RM25_RS11250 | RM25_2280 | SOS mutagenesis and repair | 2397 | gene | 2555268 | 2555978 | - | 286,0172 | -1,328320512 | 0,789973 | -1,68148 | 0,092671 | 0,183704 | 147,14879 | -4,181769017 | 0,439383 | -9,51736 | 1,78E-21 | 6,54E-20 | 424,8855 | -0,797336367 | 0,409423 | -1,94746 | 0,051479 | 0,10309 | 428,7662 | 0,88003 | 0,371356 | 2,369773 | 0,017799 | 0,037357 | 160,6177714 | 4,339492792 | 0,504288 | 8,605191 | 7,62E-18 | 1,35E-16 |
| 2307 | RM25_RS11255 | RM25_2281 | Hypothetical protein | 2398 | gene | 2556357 | 2557337 | + | 1307,763 | -0,442412339 | 0,126565 | -3,49554 | 0,000473 | 0,0059 | 1195,050424 | -0,555517435 | 0,141037 | -3,9388 | 8,19E-05 | 0,000284 | 1420,475 | -0,350207717 | 0,165412 | -2,11719 | 0,034244 | 0,076288 | 1589,556 | 0,101953 | 0,151279 | 0,673941 | 0,500349 | 0,567528 | 1085,987419 | 0,358827605 | 0,135707 | 2,644143 | 0,00819 | 0,014547 |
| 2308 | RM25_RS11260 | RM25_2282 | Putative transporter, DMT superfamily | 2399 | gene | 2558198 | 2559058 | - | 426,9952 | -1,608825707 | 0,65331 | -2,46258 | 0,013794 | 0,057861 | 246,4922399 | -3,393580879 | 0,376101 | -9,02305 | 1,83E-19 | 5,07E-18 | 607,4981 | -1,143013968 | 0,420693 | -2,71698 | 0,006588 | 0,022343 | 674,3513 | 0,823063 | 0,39559 | 2,080597 | 0,037471 | 0,06747 | 208,1126797 | 3,149944273 | 0,438749 | 7,179375 | 7E-13 | 6,21E-12 |
| 2309 | RM25_RS11265 | RM25_2283 | Transcriptional regulator, RpiR family | 2400 | gene | 2559058 | 2559852 | - | 517,1704 | -1,08107951 | 0,355207 | -3,04352 | 0,002338 | 0,017223 | 387,6337899 | -1,434957533 | 0,219636 | -6,53336 | 6,43E-11 | 6,48E-10 | 646,707 | -0,886422465 | 0,415657 | -2,13258 | 0,032959 | 0,074269 | 738,0678 | 0,500078 | 0,281847 | 1,774287 | 0,076016 | 0,118341 | 327,0666325 | 1,12152681 | 0,384832 | 2,91433 | 0,003565 | 0,006765 |
| 2310 | RM25_RS11270 | RM25_2284 | 2-dehydropantoate 2-reductase (Precursor) | 2401 | gene | 2560047 | 2560886 | + | 736,2374 | -0,939387242 | 0,171507 | -5,47724 | 4,32E-08 | 2,91E-06 | 699,7069673 | -0,604232609 | 0,150768 | -4,00769 | 6,13E-05 | 0,00022 | 772,7679 | -1,262952272 | 0,260332 | -4,85132 | 1,23E-06 | 1,66E-05 | 1020,302 | 0,311459 | 0,155 | 2,009414 | 0,044493 | 0,077567 | 496,2568387 | -0,288708974 | 0,260628 | -1,10775 | 0,267972 | 0,327428 |
| 2311 | RM25_RS11275 | RM25_2285 | Putative cation-transporting P-type ATPase | 2402 | gene | 2561093 | 2563429 | - | 1316,259 | -1,447618919 | 0,61812 | -2,34197 | 0,019182 | 0,070199 | 802,5092716 | -3,184530782 | 0,320939 | -9,92255 | 3,32E-23 | 1,7E-21 | 1830,009 | -0,941444869 | 0,384085 | -2,45114 | 0,01424 | 0,039863 | 2022,438 | 0,665375 | 0,381446 | 1,744348 | 0,081098 | 0,124771 | 696,2634703 | 2,982978847 | 0,352633 | 8,459156 | 2,69E-17 | 4,38E-16 |
| 2312 | RM25_RS11280 | RM25_2286 | Hypothetical protein | 2403 | gene | 2563498 | 2563803 | - | 426,1053 | -1,818015603 | 0,919882 | -1,97636 | 0,048114 | 0,12168 | 252,1461475 | -5,965339589 | 0,547873 | -10,8882 | 1,31E-27 | 1,03E-25 | 600,0644 | -1,172874475 | 0,398942 | -2,93996 | 0,003283 | 0,012994 | 696,8126 | 0,67192 | 0,442282 | 1,519212 | 0,128709 | 0,182862 | 185,7115383 | 5,54296836 | 0,571902 | 9,692164 | 3,26E-22 | 9,25E-21 |
| 2313 | RM25_RS11285 | RM25_2287 | Multicopper oxidase | 2404 | gene | 2563979 | 2565457 | - | 978,6038 | -1,554954428 | 0,597571 | -2,60213 | 0,009265 | 0,044205 | 623,0355455 | -3,161647519 | 0,365737 | -8,64459 | 5,4E-18 | 1,29E-16 | 1334,172 | -1,051791123 | 0,358598 | -2,93307 | 0,003356 | 0,013207 | 1533,812 | 0,614098 | 0,414689 | 1,480864 | 0,138643 | 0,194166 | 489,6320803 | 2,796759904 | 0,351929 | 7,946948 | 1,91E-15 | 2,55E-14 |
| 2314 | RM25_RS11290 | RM25_2288 | Two-component system sensor kinase TcsS4 | 2405 | gene | 2565493 | 2566632 | - | 588,7119 | -0,651269885 | 0,205322 | -3,17194 | 0,001514 | 0,012787 | 513,1137901 | -0,611498367 | 0,231456 | -2,64197 | 0,008243 | 0,016821 | 664,31 | -0,683752536 | 0,263805 | -2,59188 | 0,009545 | 0,029181 | 758,0465 | 0,331944 | 0,276453 | 1,200723 | 0,229859 | 0,29768 | 450,1564271 | 0,327193852 | 0,243544 | 1,343469 | 0,17912 | 0,230955 |
| 2315 | RM25_RS11295 | RM25_2289 | DNA-binding response regulator PhoP | 2406 | gene | 2566632 | 2567327 | - | 577,9153 | 0,099741051 | 0,153489 | 0,649827 | 0,515804 | 0,612468 | 629,1080946 | 0,319214045 | 0,136393 | 2,340399 | 0,019263 | 0,034659 | 526,7226 | -0,157468932 | 0,217218 | -0,72493 | 0,468492 | 0,570665 | 589,9879 | -0,06737 | 0,212439 | -0,3171 | 0,751164 | 0,794299 | 586,3153673 | -0,497727563 | 0,138294 | -3,59906 | 0,000319 | 0,000744 |
| 2316 | RM25_RS11300 | RM25_2290 | Permease | 2407 | gene | 2567359 | 2568066 | - | 247,9223 | -1,627300839 | 0,7076 | -2,29975 | 0,021463 | 0,074175 | 148,3663514 | -3,743380927 | 0,518104 | -7,22516 | 5,01E-13 | 6,9E-12 | 347,4782 | -1,094858221 | 0,414227 | -2,64313 | 0,008214 | 0,026341 | 393,8633 | 0,705896 | 0,42574 | 1,658043 | 0,097309 | 0,145531 | 119,6166656 | 3,429111943 | 0,57572 | 5,956214 | 2,58E-09 | 1,36E-08 |
| 2317 | RM25_RS11305 | RM25_2291 | IstB domain protein ATP-binding protein | 2408 | gene | 2568298 | 2569113 | - | 637,8294 | -1,72792113 | 0,780821 | -2,21296 | 0,026901 | 0,084203 | 369,3349638 | -4,669000889 | 0,407483 | -11,4582 | 2,14E-30 | 2,4E-28 | 906,3239 | -1,148443548 | 0,38469 | -2,98537 | 0,002832 | 0,01151 | 1027,556 | 0,743212 | 0,410818 | 1,809101 | 0,070435 | 0,112097 | 291,6069343 | 4,337521004 | 0,424031 | 10,22926 | 1,47E-24 | 5,08E-23 |
| 2318 | RM25_RS11310 | RM25_2292 | IstA | 2409 | gene | 2569110 | 2570771 | - | 894,0321 | -1,382630106 | 0,847251 | -1,6319 | 0,1027 | 0,195845 | 451,4290109 | -4,830953978 | 0,378354 | -12,7683 | 2,46E-37 | 7,26E-35 | 1336,635 | -0,834355026 | 0,429076 | -1,94454 | 0,051831 | 0,10353 | 1354,291 | 0,904111 | 0,400073 | 2,259867 | 0,02383 | 0,047119 | 489,7816019 | 4,976661643 | 0,432506 | 11,50657 | 1,22E-30 | 8,23E-29 |
| 2319 | NA | RM25_2293 | ABC peptide transporter, permease component | 2410 | gene | 2571007 | 2571909 | - | 355,494 | -1,188926387 | 0,907181 | -1,31057 | 0,190002 | 0,294091 | 166,4372527 | -5,173637161 | 0,485165 | -10,6637 | 1,51E-26 | 1,04E-24 | 544,5507 | -0,647286201 | 0,444185 | -1,45725 | 0,145049 | 0,229603 | 518,2331 | 0,96254 | 0,393327 | 2,447173 | 0,014398 | 0,031589 | 214,1736655 | 5,566116668 | 0,553681 | 10,05293 | 8,92E-24 | 2,96E-22 |
| 2320 | NA | RM25_2294 | ABC-type dipeptide transport system, periplasmic compone | 2411 | gene | 2571964 | 2572659 | - | 426,7724 | -1,544098894 | 0,885176 | -1,7444 | 0,08109 | 0,168321 | 226,2650358 | -5,450704968 | 0,462048 | -11,7968 | 4,05E-32 | 5,97E-30 | 627,2798 | -0,961401429 | 0,402692 | -2,38744 | 0,016966 | 0,045391 | 666,3584 | 0,837926 | 0,372966 | 2,246652 | 0,024662 | 0,04832 | 215,1752291 | 5,398704494 | 0,520583 | 10,3705 | 3,38E-25 | 1,33E-23 |
| 2321 | RM25_RS11320 | RM25_2295 | Pyridoxamine 5'-phosphate oxidase family protein | 2412 | gene | 2572971 | 2573456 | - | 114,9462 | -0,938441046 | 0,44488 | -2,10942 | 0,034908 | 0,09961 | 76,13351574 | -1,659174257 | 0,415314 | -3,99499 | 6,47E-05 | 0,000231 | 153,759 | -0,629119696 | 0,303465 | -2,07573 | 0,037919 | 0,083073 | 158,9097 | 0,625473 | 0,345065 | 1,812621 | 0,06989 | 0,111456 | 77,41880048 | 1,717933444 | 0,412575 | 4,163931 | 3,13E-05 | 8,58E-05 |
| 2322 | RM25_RS11325 | RM25_2296 | 2, 3-diaminopropionate biosynthesis protein SbnA | 2413 | gene | 2573755 | 2574942 | + | 1607,471 | 0,266198195 | 0,203376 | 1,308896 | 0,19057 | 0,294539 | 1847,619491 | 0,161333346 | 0,208505 | 0,773764 | 0,439071 | 0,50705 | 1367,322 | 0,409648703 | 0,2002 | 2,046195 | 0,040737 | 0,087051 | 1545,954 | -0,62577 | 0,261208 | -2,3957 | 0,016589 | 0,035353 | 1720,752918 | -0,330353352 | 0,121364 | -2,72201 | 0,006489 | 0,011719 |
| 2323 | RM25_RS11330 | RM25_2297 | Pyridoxal-phosphate dependent enzyme | 2414 | gene | 2574935 | 2576989 | + | 988,2586 | -0,039348817 | 0,219974 | -0,17888 | 0,858032 | 0,906492 | 872,3778908 | -0,183517369 | 0,181903 | -1,00888 | 0,313034 | 0,379929 | 1104,139 | 0,074200291 | 0,335065 | 0,221451 | 0,824742 | 0,871938 | 1052,835 | 0,154621 | 0,242079 | 0,638721 | 0,523004 | 0,590667 | 958,8056601 | 0,474009002 | 0,278322 | 1,703094 | 0,08855 | 0,123645 |
| 2324 | RM25_RS11335 | RM25_2298 | Decarboxylase | 2415 | gene | 2577001 | 2578710 | + | 975,14 | -0,039468636 | 0,152404 | -0,25897 | 0,795655 | 0,857111 | 996,3022192 | -0,198460471 | 0,150518 | -1,31852 | 0,187331 | 0,245846 | 953,9778 | 0,126036594 | 0,273333 | 0,461109 | 0,64472 | 0,726042 | 1044,654 | -0,28289 | 0,190624 | -1,48402 | 0,137803 | 0,193105 | 944,7123458 | 0,102041612 | 0,234175 | 0,43575 | 0,663018 | 0,716522 |
| 2325 | RM25_RS11340 | RM25_2299 | Putative insertion element ISCmi2 transposase | 2416 | gene | 2579027 | 2580070 | + | 84,1772 | -0,070075927 | 0,30573 | -0,22921 | 0,818707 | 0,873559 | 76,2488794 | -0,804181303 | 0,342529 | -2,34778 | 0,018886 | 0,034084 | 92,10552 | 0,5112137 | 0,314008 | 1,628027 | 0,103519 | 0,176552 | 91,29693 | -0,4014 | 0,230908 | -1,73835 | 0,08215 | 0,125977 | 80,01745382 | 0,953539388 | 0,399111 | 2,38916 | 0,016887 | 0,028169 |
| 2326 | RM25_RS11345 | RM25_2300 | Hypothetical protein | 2417 | gene | 2580074 | 2581099 | - | 933,0839 | -0,631647994 | 0,219422 | -2,87869 | 0,003993 | 0,025577 | 752,1179044 | -0,564913194 | 0,202851 | -2,78487 | 0,005355 | 0,011654 | 1114,05 | -0,678719733 | 0,192474 | -3,52629 | 0,000421 | 0,002449 | 1193,685 | 0,546658 | 0,245007 | 2,231191 | 0,025668 | 0,049754 | 717,9919325 | 0,491953547 | 0,164636 | 2,988132 | 0,002807 | 0,005432 |
| 2327 | RM25_RS11350 | RM25_2301 | Hypothetical protein | 2418 | gene | 2581342 | 2581983 | + | 471,1399 | -0,254056836 | 0,214505 | -1,18438 | 0,236261 | 0,343745 | 385,127927 | -0,375318352 | 0,208973 | -1,79601 | 0,072492 | 0,107462 | 557,1518 | -0,174941497 | 0,185338 | -0,94391 | 0,345218 | 0,451542 | 539,3643 | 0,378444 | 0,221501 | 1,708543 | 0,087536 | 0,133111 | 421,2274102 | 0,635778148 | 0,171572 | 3,705602 | 0,000211 | 0,000506 |
| 2328 | RM25_RS11355 | RM25_2302 | Hypothetical protein | 2419 | gene | 2581980 | 2583440 | + | 554,24 | -0,054122692 | 0,172989 | -0,31287 | 0,754381 | 0,822422 | 481,3492976 | -0,233711902 | 0,174243 | -1,3413 | 0,179823 | 0,237181 | 627,1307 | 0,080403073 | 0,192679 | 0,41729 | 0,676466 | 0,754582 | 594,7683 | 0,168823 | 0,175205 | 0,963573 | 0,33526 | 0,410072 | 533,4249162 | 0,538581228 | 0,179506 | 3,000354 | 0,002697 | 0,005253 |
| 2329 | RM25_RS11360 | RM25_2303 | DEAD-like helicase | 2420 | gene | 2583719 | 2586292 | + | 4012,816 | 0,52891101 | 0,232407 | 2,275795 | 0,022858 | 0,077633 | 4419,842565 | 0,156238712 | 0,166042 | 0,940957 | 0,346727 | 0,416108 | 3605,789 | 1,012741856 | 0,266279 | 3,803311 | 0,000143 | 0,001002 | 3489,166 | -0,86148 | 0,215096 | -4,00509 | 6,2E-05 | 0,000321 | 4634,648934 | 0,034540562 | 0,198541 | 0,173972 | 0,861888 | 0,888268 |
| 2330 | RM25_RS11370 | RM25_2304 | Hypothetical protein | 2421 | gene | [illegible] | [illegible] | - | [illegible] | [illegible] | [illegible] | [illegible] | [illegible] | [illegible] | [illegible] | [illegible] | [illegible] | [illegible] | [illegible] | [illegible] | [illegible] | [illegible] | [illegible] | [illegible] | [illegible] | [illegible] | [illegible] | [illegible] | [illegible] | [illegible] | [illegible] | [illegible] | [illegible] | [illegible] | [illegible] | [illegible] | [illegible] | [illegible] |
| 2331 | RM25_RS11375 | RM25_2305 | Low temperature requirement A | 2422 | gene | 2587221 | 2588343 | + | 658,9592 | [illegible] | [illegible] | [illegible] | [illegible] | [illegible] | [illegible] | [illegible] | [illegible] | [illegible] | [illegible] | [illegible] | [illegible] | [illegible] | [illegible] | [illegible] | [illegible] | [illegible] | [illegible] | [illegible] | [illegible] | [illegible] | [illegible] | [illegible] | [illegible] | [illegible] | [illegible] | [illegible] | [illegible] | [illegible] |
| 2332 | NA | RM25_2306 | Putative membrane protein | 2423 | gene | 2588334 | 2588705 | + | 273,7221 | 0,547506582 | 0,197556 | 2,771396 | 0,005582 | 0,031802 | 249,975069 | 0,146519756 | 0,233771 | 0,626767 | 0,530812 | 0,596625 | 297,4691 | 0,893203142 | 0,225665 | 3,958102 | 7,55E-05 | 0,000585 | 235,1711 | -0,25395 | 0,201167 | -1,26241 | 0,206801 | 0,271549 | 317,9365934 | 0,546519285 | 0,236824 | 2,307699 | 0,021016 | 0,034447 |
| 2333 | RM25_RS11385 | RM25_2307 | Cytochrome P450 | 2424 | gene | 2588806 | 2590020 | - | 1621,148 | 0,308978253 | 0,239424 | 1,290508 | 0,196874 | 0,299693 | 1239,744493 | 0,161156404 | 0,129049 | 1,248802 | 0,211738 | 0,272862 | 2002,551 | 0,401748524 | 0,193576 | 2,075408 | 0,037949 | 0,083073 | 1525,361 | 0,502628 | 0,163896 | 3,066757 | 0,002164 | 0,006738 | 1754,41968 | 0,789008756 | 0,140801 | 5,603724 | 2,1E-08 | 9,58E-08 |
| 2334 | RM25_RS11390 | RM25_2308 | Putative glyoxal reductase | 2425 | gene | 2590353 | 2591207 | - | 2350,825 | 0,686972311 | 0,297012 | 2,312944 | 0,020726 | 0,07302 | 2740,139542 | 0,274595143 | 0,182826 | 1,50195 | 0,13311 | 0,184012 | 1961,511 | 1,319072451 | 0,289325 | 4,559139 | 5,14E-06 | 5,63E-05 | 1920,328 | -1,20081 | 0,261254 | -4,59635 | 4,3E-06 | 2,87E-05 | 2836,612572 | -0,114278313 | 0,207568 | -0,55056 | 0,581937 | 0,642448 |
| 2335 | RM25_RS11395 | RM25_2309 | Hypothetical protein | 2426 | gene | 2591415 | 2592458 | - | 1172,082 | 0,611782846 | 0,143393 | 4,26647 | 1,99E-05 | 0,000483 | 1294,84759 | 0,786944361 | 0,119937 | 6,561317 | 5,33E-11 | 5,51E-10 | 1049,317 | 0,40347821 | 0,203181 | 1,985809 | 0,047055 | 0,096863 | 978,6226 | -0,13458 | 0,169245 | -0,79518 | 0,426506 | 0,501384 | 1391,027506 | -0,460251936 | 0,144496 | -3,18523 | 0,001446 | 0,002954 |
| 2336 | RM25_RS11400 | RM25_2310 | Alpha-glucosidase aglA | 2427 | gene | 2592768 | 2594555 | - | 1391,945 | 0,160659181 | 0,097462 | 1,648424 | 0,099266 | 0,192093 | 1350,665498 | 0,230739269 | 0,134022 | 1,721658 | 0,085132 | 0,12348 | 1433,224 | 0,095661693 | 0,190145 | 0,5031 | 0,614894 | 0,702183 | 1385,602 | 0,096727 | 0,13825 | 0,699652 | 0,484145 | 0,554326 | 1442,006421 | 0,017709546 | 0,16709 | 0,105988 | 0,915592 | 0,931398 |
| 2337 | RM25_RS11405 | RM25_2311 | Hypothetical protein | 2428 | gene | 2594614 | 2595234 | - | 818,4192 | 0,03713908 | 0,211743 | 0,175397 | 0,860768 | 0,907665 | 690,7266313 | 0,470891418 | 0,148935 | 3,161724 | 0,001568 | 0,003964 | 946,1117 | -0,274993513 | 0,168752 | -1,62957 | 0,103192 | 0,176122 | 848,7512 | 0,785575 | 0,179261 | 4,382303 | 1,17E-05 | 7,1E-05 | 812,7799534 | 0,086949 | 0,124926 | 0,696003 | 0,486427 | 0,548377 |
| 2338 | RM25_RS11410 | RM25_2312 | Xanthine permease | 2429 | gene | 2595732 | 2597678 | + | 1602,419 | 0,089649769 | 0,176495 | 0,507946 | 0,611491 | 0,700333 | 1814,722949 | -0,121633325 | 0,134007 | -0,90766 | 0,364056 | 0,432282 | 1390,115 | 0,366988646 | 0,177684 | 2,065397 | 0,038885 | 0,084287 | 1647,227 | -0,70107 | 0,17005 | -4,1227 | 3,74E-05 | 0,000201 | 1618,893681 | -0,159530315 | 0,112153 | -1,42243 | 0,154901 | 0,204196 |
| 2339 | RM25_RS11415 | RM25_2313 | Putative xylan esterase | 2430 | gene | 2597777 | 2598778 | - | 760,4287 | -0,423961884 | 0,240367 | -1,76381 | 0,077764 | 0,165184 | 625,9225891 | -0,803950916 | 0,14381 | -5,59035 | 2,27E-08 | 1,55E-07 | 894,9349 | -0,168330533 | 0,248911 | -0,67627 | 0,498871 | 0,598783 | 918,4556 | 0,188532 | 0,1602 | 1,176849 | 0,239256 | 0,307988 | 637,5387632 | 0,885239455 | 0,229098 | 3,864019 | 0,000112 | 0,00028 |
| 2340 | RM25_RS11420 | RM25_2314 | AMP-binding enzyme | 2431 | gene | 2599256 | 2600923 | + | 1671,434 | 0,250171174 | 0,168144 | 1,487838 | 0,136794 | 0,236554 | 1550,882941 | 0,548545734 | 0,15791 | 3,473789 | 0,000513 | 0,00148 | 1791,984 | -0,003702991 | 0,221881 | -0,01669 | 0,986685 | 0,992157 | 1605,485 | 0,444797 | 0,215318 | 2,065766 | 0,038851 | 0,069215 | 1781,751186 | -0,04851614 | 0,158893 | -0,30534 | 0,760108 | 0,803037 |
| 2341 | RM25_RS11425 | RM25_2315 | Catalase | 2432 | gene | 2601064 | 2602512 | - | 6035,746 | 1,336762778 | 0,401293 | 3,331136 | 0,000865 | 0,009142 | 3673,225261 | 1,081938221 | 0,138135 | 7,832483 | 4,78E-15 | 8E-14 | 8398,267 | 1,454934308 | 0,422246 | 3,445703 | 0,00057 | 0,003108 | 3606,35 | 0,880481 | 0,379623 | 2,319355 | 0,020376 | 0,041519 | 8429,417888 | 1,285720533 | 0,225224 | 5,708622 | 1,14E-08 | 5,35E-08 |
| 2342 | RM25_RS11430 | RM25_2316 | Dyp-type peroxidase family protein | 2433 | gene | 2602617 | 2603534 | - | 1472,787 | -0,023607082 | 0,095076 | -0,2483 | 0,803906 | 0,862844 | 1454,944278 | 0,067924981 | 0,118278 | 0,574284 | 0,565775 | 0,62887 | 1490,629 | -0,111883185 | 0,201341 | -0,55569 | 0,578423 | 0,670607 | 1569,166 | 0,066558 | 0,186364 | 0,35714 | 0,720987 | 0,767901 | 1430,603444 | -0,066227947 | 0,127249 | -0,52046 | 0,602744 | 0,662625 |
| 2343 | RM25_RS11435 | RM25_2317 | ABC-1 domain protein | 2434 | gene | 2604066 | 2606018 | + | 1414,459 | -0,043023019 | 0,232448 | -0,18509 | 0,853161 | 0,904036 | 1116,75142 | -0,303708586 | 0,161569 | -1,87975 | 0,060142 | 0,091484 | 1712,167 | 0,124879681 | 0,17115 | 0,72965 | 0,465604 | 0,568028 | 1512,215 | 0,344259 | 0,193819 | 1,776183 | 0,075703 | 0,11801 | 1365,81767 | 0,830761616 | 0,139618 | 5,950264 | 2,68E-09 | 1,4E-08 |
| 2344 | RM25_RS12125 | RM25_2318 | Hypothetical protein | 2435 | gene | 2606485 | 2607240 | - | 846,908 | 0,263058288 | 0,189801 | 1,385968 | 0,165757 | 0,26833 | 761,6406093 | 0,427998876 | 0,165857 | 2,580524 | 0,009865 | 0,019655 | 932,1753 | 0,131250358 | 0,289362 | 0,453585 | 0,650128 | 0,730386 | 812,2825 | 0,400475 | 0,257106 | 1,557624 | 0,119322 | 0,171699 | 903,2433695 | 0,143038763 | 0,198192 | 0,721719 | 0,470467 | 0,533121 |
| 2345 | RM25_RS11445 | RM25_2319 | Chloride transporter, chloride channel family protein | 2436 | gene | 2607324 | 2608580 | - | 554,745 | -0,73810854 | 0,37328 | -1,97736 | 0,048001 | 0,121609 | 389,1366632 | -1,149512873 | 0,229448 | -5,00991 | 5,45E-07 | 2,86E-06 | 720,3533 | -0,530908016 | 0,377681 | -1,4057 | 0,159812 | 0,247164 | 728,6252 | 0,600475 | 0,269532 | 2,227845 | 0,025891 | 0,050061 | 409,6585116 | 1,288385991 | 0,354009 | 3,639419 | 0,000273 | 0,000644 |
| 2346 | RM25_RS11450 | RM25_2320 | Hypothetical protein | 2437 | gene | 2609488 | 2609787 | + | 411,6146 | 0,046937782 | 0,341416 | 0,13748 | 0,890652 | 0,926419 | 281,2957816 | -0,380063028 | 0,219601 | -1,7307 | 0,083506 | 0,121646 | 541,9333 | 0,265320793 | 0,266423 | 0,995862 | 0,319317 | 0,424496 | 426,6527 | 0,566982 | 0,271067 | 2,091665 | 0,036468 | 0,066324 | 408,8940308 | 1,262042857 | 0,209503 | 6,023982 | 1,7E-09 | 9,16E-09 |
| 2347 | RM25_RS11455 | RM25_2321 | Hypothetical protein | 2438 | gene | 2610283 | 2610516 | - | 487,0021 | 0,4597374 | 0,348135 | 1,320572 | 0,186644 | 0,291031 | 336,1516806 | 0,229700509 | 0,223173 | 1,02925 | 0,303362 | 0,372519 | 637,8526 | 0,584299798 | 0,369713 | 1,580415 | 0,114012 | 0,190992 | 433,2111 | 0,666574 | 0,350878 | 1,899731 | 0,057468 | 0,095188 | 550,7700379 | 1,064344351 | 0,259284 | 4,10493 | 4,04E-05 | 0,000109 |
| 2348 | RM25_RS11460 | RM25_2322 | Hypothetical protein | 2439 | gene | 2610743 | 2611156 | - | 3732,559 | -0,85266664 | 0,468761 | -1,81898 | 0,068914 | 0,152089 | 2094,578741 | -0,135644204 | 0,236275 | -0,5741 | 0,565903 | 0,62887 | 5370,539 | -1,15710791 | 0,465598 | -2,48521 | 0,012948 | 0,037241 | 5046,274 | 1,712399 | 0,412828 | 4,147976 | 3,35E-05 | 0,000183 | 2591,753228 | 0,715343133 | 0,31171 | 2,294903 | 0,021739 | 0,035508 |

| | | | | | | | | | | | | | | | | | | | | | | | | | | | | | | | | | | | | | | |
|---|---|---|---|---|---|---|---|---|---|---|---|---|---|---|---|---|---|---|---|---|---|---|---|---|---|---|---|---|---|---|---|---|---|---|---|---|---|---|
| 2349 | RM25_RS11465 | RM25_2323 | Fructose-bisphosphate aldolase class 1 | 2440 | gene | 2611319 | 2612206 | - | 5920,822 | -1,146872043 | 0,590367 | -1,94264 | 0,052059 | 0,127418 | 2289,528858 | 0,097068239 | 0,169705 | 0,571981 | 0,567335 | 0,629868 | 9552,115 | -1,495269649 | 0,435329 | -3,4348 | 0,000593 | 0,003203 | 8546,69 | 2,624444 | 0,413462 | 6,347487 | 2,19E-10 | 3,74E-09 | 3595,764375 | 1,064991639 | 0,221325 | 4,811895 | 1,5E-06 | 5,02E-06 |
| 2350 | RM25_RS11470 | RM25_2324 | YceI family protein | 2441 | gene | 2612589 | 2613140 | - | 3975,67 | 2,157845539 | 0,509549 | 4,234816 | 2,29E-05 | 0,000531 | 1780,798702 | 0,536077524 | 0,183039 | 2,928764 | 0,003403 | 0,007864 | 6170,541 | 2,901672386 | 0,406895 | 7,131257 | 9,95E-13 | 5,58E-11 | 1542,233 | -0,04764 | 0,36054 | -0,13212 | 0,894887 | 0,916191 | 6317,481969 | 2,350666362 | 0,252152 | 9,322432 | 1,14E-20 | 2,71E-19 |
| 2351 | RM25_RS11475 | RM25_2325 | Citrate (Si)-synthase | 2442 | gene | 2613520 | 2614803 | + | 2375,768 | 0,609305807 | 0,224681 | 2,711871 | 0,00669 | 0,036168 | 2264,189043 | 1,124554987 | 0,126519 | 8,888445 | 6,2E-19 | 1,61E-17 | 2487,348 | 0,172436298 | 0,292003 | 0,59053 | 0,554836 | 0,651269 | 1979,604 | 0,660484 | 0,196261 | 3,365335 | 0,000765 | 0,002802 | 2807,514335 | -0,25207572 | 0,219333 | -1,14928 | 0,25044 | 0,308727 |
| 2352 | RM25_RS11480 | RM25_2326 | L-glyceraldehyde 3-phosphate reductase | 2443 | gene | 2615043 | 2616101 | + | 1023,991 | -0,150006772 | 0,221801 | -0,67631 | 0,498842 | 0,596955 | 813,2265037 | -0,031464172 | 0,145099 | -0,21685 | 0,828329 | 0,863881 | 1234,755 | -0,229537925 | 0,211226 | -1,08669 | 0,277173 | 0,380266 | 1132,911 | 0,638682 | 0,205894 | 3,101992 | 0,001922 | 0,006098 | 950,6346635 | 0,489180483 | 0,135178 | 3,6188 | 0,000296 | 0,000692 |
| 2353 | RM25_RS11485 | RM25_2327 | Hypothetical protein | 2444 | gene | 2616244 | 2616885 | + | 406,1173 | -0,806117876 | 0,243776 | -3,3068 | 0,000944 | 0,009713 | 339,1818939 | -0,8326061 | 0,272568 | -3,05467 | 0,002253 | 0,005475 | 473,0528 | -0,789650007 | 0,273581 | -2,88635 | 0,003897 | 0,014816 | 543,4983 | 0,398879 | 0,313009 | 1,274334 | 0,202545 | 0,266592 | 290,3607749 | 0,505922248 | 0,253255 | 1,997682 | 0,045751 | 0,068337 |
| 2354 | RM25_RS11490 | RM25_2328 | Hypothetical protein | 2445 | gene | 2616913 | 2617116 | - | 331,128 | 0,023873434 | 0,215001 | 0,111039 | 0,911586 | 0,941556 | 339,1056681 | -0,479637587 | 0,185031 | -2,59221 | 0,009536 | 0,019081 | 323,1504 | 0,552787897 | 0,249979 | 2,211341 | 0,027012 | 0,064506 | 348,8394 | -0,65132 | 0,201169 | -3,23768 | 0,001205 | 0,004128 | 325,9750186 | 0,426526537 | 0,219507 | 1,943112 | 0,052003 | 0,076415 |
| 2355 | RM25_RS11495 | RM25_2329 | Membrane protein YdfJ | 2446 | gene | 2617509 | 2620013 | - | 34209,89 | 1,950306932 | 0,610338 | 3,195456 | 0,001396 | 0,012325 | 28305,33232 | 0,395308482 | 0,202782 | 1,94943 | 0,051244 | 0,080629 | 40114,45 | 4,379643909 | 0,430814 | 10,16597 | 2,81E-24 | 5,53E-22 | 15087,79 | -2,78455 | 0,405509 | -6,86681 | 6,57E-12 | 1,49E-10 | 52951,16804 | 1,232901515 | 0,239328 | 5,151519 | 2,58E-07 | 9,85E-07 |
| 2356 | RM25_RS11500 | RM25_2330 | Transcriptional regulator, TetR family | 2447 | gene | 2620227 | 2620913 | + | 4334,655 | 0,725154373 | 0,286957 | 2,527053 | 0,011502 | 0,051201 | 3612,17423 | 0,148964869 | 0,206295 | 0,722095 | 0,470236 | 0,538417 | 5057,135 | 1,171980506 | 0,362455 | 3,233455 | 0,001223 | 0,005731 | 3460,437 | -0,18809 | 0,34429 | -0,54632 | 0,584845 | 0,646869 | 5267,165423 | 0,865385755 | 0,221313 | 3,910231 | 9,22E-05 | 0,000235 |
| 2357 | RM25_RS11505 | RM25_2331 | Putative NAD(P) binding enzyme | 2448 | gene | 2620971 | 2621627 | + | 1764,296 | 0,500095649 | 0,256152 | 1,952343 | 0,050898 | 0,125618 | 1388,996201 | 0,011860104 | 0,142991 | 0,082943 | 0,933897 | 0,949275 | 2139,595 | 0,831519186 | 0,260359 | 3,193739 | 0,001404 | 0,00644 | 1544,623 | 0,096597 | 0,194316 | 0,497113 | 0,619109 | 0,676994 | 2016,790483 | 0,958235948 | 0,196356 | 4,880106 | 1,06E-06 | 3,68E-06 |
| 2358 | RM25_RS11510 | RM25_2332 | Hypothetical protein | 2449 | gene | 2621834 | 2622151 | + | 397,6073 | 0,729498409 | 0,34489 | 2,115161 | 0,034416 | 0,098685 | 261,4709975 | 0,147576861 | 0,256279 | 0,575845 | 0,56472 | 0,628147 | 533,7436 | 1,032572831 | 0,166343 | 6,20749 | 5,38E-10 | 1,65E-08 | 314,4203 | 0,443546 | 0,172614 | 2,569577 | 0,010182 | 0,024048 | 485,1873117 | 1,374965129 | 0,230977 | 5,952834 | 2,64E-09 | 1,38E-08 |
| 2359 | RM25_RS12130 | RM25_2333 | Respiratory-chain NADH dehydrogenase 24 Kd subunit doma | 2450 | gene | 2622191 | 2622982 | - | 250,3038 | -0,409679473 | 0,529123 | -0,77426 | 0,438777 | 0,547032 | 128,7416693 | -1,142611051 | 0,298569 | -3,82696 | 0,00013 | 0,00043 | 371,8658 | -0,174955031 | 0,394162 | -0,44387 | 0,65714 | 0,73721 | 298,5422 | 1,092657 | 0,267299 | 4,087761 | 4,36E-05 | 0,00023 | 211,895991 | 2,124405175 | 0,408893 | 5,195499 | 2,04E-07 | 7,91E-07 |
| 2360 | RM25_RS11520 | RM25_2334 | Iron(III) dicitrate transport ATP-binding protein | 2451 | gene | 2622972 | 2623703 | - | 286,2834 | -0,795332522 | 0,285394 | -2,78679 | 0,005323 | 0,030904 | 248,8038747 | -1,024983128 | 0,19966 | -5,13365 | 2,84E-07 | 1,6E-06 | 323,7629 | -0,630720802 | 0,435289 | -1,44897 | 0,147346 | 0,231685 | 383,363 | 0,171694 | 0,234301 | 0,732792 | 0,463685 | 0,535476 | 206,2497958 | 0,638384173 | 0,432934 | 1,474552 | 0,140333 | 0,186557 |
| 2361 | RM25_RS11525 | RM25_2335 | Iron chelate uptake ABC transporter, FeCT family, permease | 2452 | gene | 2623700 | 2624806 | - | 518,991 | -0,372898758 | 0,237183 | -1,5722 | 0,115905 | 0,211118 | 460,70745 | -0,560287678 | 0,188288 | -2,97569 | 0,002923 | 0,006835 | 577,2745 | -0,226828643 | 0,364897 | -0,62162 | 0,534189 | 0,631876 | 617,8404 | 0,118186 | 0,194018 | 0,609148 | 0,542426 | 0,60823 | 445,2372141 | 0,520491466 | 0,354558 | 1,468002 | 0,142104 | 0,188698 |
| 2362 | RM25_RS11530 | RM25_2336 | ABC transporter domain-containing permease component | 2453 | gene | 2624814 | 2625683 | - | 1515,748 | 0,267241515 | 0,362407 | 0,737408 | 0,460874 | 0,564889 | 1920,469086 | -0,202940983 | 0,19976 | -1,01592 | 0,309666 | 0,376811 | 1111,026 | 1,132141069 | 0,24275 | 4,663821 | 3,1E-06 | 3,69E-05 | 1466,237 | -1,62261 | 0,250722 | -6,47176 | 9,69E-11 | 1,73E-09 | 1622,344378 | -0,236192518 | 0,173722 | -1,3596 | 0,173957 | 0,226148 |
| 2363 | RM25_RS12135 | RM25_2337 | MscS transporter, small conductance mechanosensitive ion | 2454 | gene | 2626314 | 2628089 | + | 2481,19 | 1,069146818 | 0,248702 | 4,298912 | 1,72E-05 | 0,00043 | 3053,78506 | 1,418851936 | 0,181061 | 7,836329 | 4,64E-15 | 7,81E-14 | 1908,596 | 0,564751711 | 0,196819 | 2,869391 | 0,004113 | 0,01546 | 1694,359 | -0,16553 | 0,231552 | -0,71489 | 0,474679 | 0,546565 | 3295,061407 | -0,975646258 | 0,133698 | -7,29741 | 2,93E-13 | 2,77E-12 |
| 2364 | RM25_RS11540 | RM25_2338 | CCA tRNA nucleotidyltransferase | 2455 | gene | 2628144 | 2629634 | + | 1309,881 | -0,243710162 | 0,130983 | -1,86062 | 0,062797 | 0,142908 | 1230,262594 | -0,301810975 | 0,132748 | -2,27357 | 0,022992 | 0,040291 | 1389,5 | -0,193592812 | 0,235821 | -0,82093 | 0,411686 | 0,516965 | 1497,628 | 0,060472 | 0,205793 | 0,293849 | 0,768873 | 0,811933 | 1177,189032 | 0,229715356 | 0,167037 | 1,375234 | 0,169059 | 0,220882 |
| 2365 | RM25_RS11545 | RM25_2339 | Major facilitator family transmembrane efflux protein | 2456 | gene | 2630037 | 2631527 | + | 1188,591 | 0,051269345 | 0,213952 | 0,23963 | 0,810617 | 0,868465 | 1224,701315 | -0,494073233 | 0,15564 | -3,17447 | 0,001501 | 0,003804 | 1152,48 | 0,634355289 | 0,200512 | 3,163682 | 0,001558 | 0,007034 | 1240,335 | -0,72993 | 0,206659 | -3,53207 | 0,000412 | 0,001681 | 1185,12162 | 0,455849495 | 0,143006 | 3,187624 | 0,001434 | 0,002935 |
| 2366 | RM25_RS11550 | RM25_2340 | Hypothetical protein | 2457 | gene | 2631701 | 2633797 | + | 1992,262 | -0,08497372 | 0,0905 | -0,93894 | 0,347763 | 0,460234 | 1976,675612 | 0,014016459 | 0,126182 | 0,111082 | 0,911552 | 0,932925 | 2007,847 | -0,181854148 | 0,184176 | -0,98739 | 0,323451 | 0,428782 | 2164,174 | 0,057831 | 0,1368 | 0,422741 | 0,672484 | 0,723765 | 1897,884581 | -0,082283242 | 0,165506 | -0,49716 | 0,619074 | 0,676366 |
| 2367 | RM25_RS12140 | RM25_2341 | Hypothetical protein | 2458 | gene | 2634313 | 2636388 | + | 1415,344 | -0,402239268 | 0,26344 | -1,52687 | 0,126793 | 0,224791 | 1161,347145 | -0,761186039 | 0,123869 | -6,14509 | 7,99E-10 | 6,93E-09 | 1669,341 | -0,161474288 | 0,333028 | -0,48487 | 0,62777 | 0,713152 | 1695,231 | 0,20597 | 0,237123 | 0,868619 | 0,385056 | 0,461871 | 1198,858089 | 0,871435248 | 0,257795 | 3,380339 | 0,000724 | 0,00159 |
| 2368 | RM25_RS11560 | RM25_2342 | Phosphoglycerate mutase family protein | 2459 | gene | 2636464 | 2637183 | + | 1387,613 | 0,134713556 | 0,221559 | 0,608025 | 0,543171 | 0,638531 | 1679,716088 | 0,167866005 | 0,15917 | 1,054636 | 0,291592 | 0,359833 | 1095,509 | 0,085580623 | 0,170814 | 0,501017 | 0,616359 | 0,703515 | 1402,211 | -0,63088 | 0,170162 | -3,70756 | 0,000209 | 0,000929 | 1423,830343 | -0,66508451 | 0,142193 | -4,67734 | 2,91E-06 | 9,33E-06 |
| 2369 | RM25_RS11565 | RM25_2343 | Putative exported solute-binding protein | 2460 | gene | 2637312 | 2638631 | - | 1303,795 | -0,013062645 | 0,131939 | -0,09901 | 0,921134 | 0,948084 | 1429,073685 | -0,032068818 | 0,143352 | -0,22371 | 0,822985 | 0,859068 | 1178,516 | 0,010949211 | 0,184377 | 0,059385 | 0,952646 | 0,970768 | 1385,466 | -0,36039 | 0,184402 | -1,95438 | 0,050656 | 0,085835 | 1272,587429 | -0,263249947 | 0,118662 | -2,21849 | 0,026522 | 0,042352 |
| 2370 | RM25_RS11570 | RM25_2344 | Thioredoxin-disulfide reductase | 2461 | gene | 2639050 | 2640027 | + | 11630,29 | -0,216253666 | 0,356188 | -0,60713 | 0,543762 | 0,638907 | 13011,39275 | -0,699278907 | 0,246557 | -2,83617 | 0,004566 | 0,010039 | 10249,19 | 0,383169348 | 0,501457 | 0,764113 | 0,4448 | 0,548673 | 13316,88 | -0,90204 | 0,499996 | -1,80409 | 0,071218 | 0,113189 | 10516,62 | 0,210516026 | 0,256677 | 0,82016 | 0,412125 | 0,478464 |
| 2371 | RM25_RS11575 | RM25_2345 | Thioredoxin | 2462 | gene | 2640201 | 2640524 | + | 2127,243 | 0,745923792 | 0,567493 | 1,31442 | 0,188705 | 0,293196 | 2421,740054 | -0,362959353 | 0,274247 | -1,32348 | 0,185677 | 0,24411 | 1832,746 | 2,825077945 | 0,433162 | 6,521993 | 6,94E-11 | 2,48E-09 | 1701,709 | -2,63766 | 0,445827 | -5,91633 | 3,29E-09 | 4,06E-08 | 2601,623823 | 0,583628243 | 0,253423 | 2,30298 | 0,02128 | 0,034807 |
| 2372 | RM25_RS11580 | RM25_2346 | OsmC family protein | 2463 | gene | 2640524 | 2640970 | + | 2072,775 | 1,141729973 | 0,542111 | 2,10608 | 0,035197 | 0,100073 | 2364,369265 | 0,167280883 | 0,249508 | 0,670444 | 0,502575 | 0,568411 | 1781,181 | 3,163816819 | 0,299901 | 10,54953 | 5,11E-26 | 1,5E-23 | 1385,183 | -2,69161 | 0,347017 | -7,75641 | 8,74E-15 | 3,32E-13 | 2790,628563 | 0,345230907 | 0,190053 | 1,816497 | 0,069294 | 0,099166 |
| 2373 | RM25_RS11585 | RM25_2347 | Chromosome partitioning protein ParB | 2464 | gene | 2641321 | 2642304 | - | 804,0528 | -0,196624223 | 0,075099 | -2,61821 | 0,008839 | 0,043045 | 817,7765949 | -0,232476734 | 0,126206 | -1,84205 | 0,065468 | 0,098411 | 790,3289 | -0,157961448 | 0,173882 | -0,90844 | 0,363645 | 0,470682 | 906,9767 | -0,14646 | 0,150007 | -0,97634 | 0,328894 | 0,404173 | 734,9401476 | -0,017068031 | 0,130378 | -0,13091 | 0,895845 | 0,916055 |
| 2374 | NA | RM25_2348 | Chromosome partitioning protein | 2465 | gene | 2642301 | 2643983 | - | 2430,615 | -0,029882701 | 0,198509 | -0,15054 | 0,880342 | 0,918533 | 2795,295961 | 0,132516763 | 0,150497 | 0,880528 | 0,378573 | 0,447043 | 2065,935 | -0,248939779 | 0,235759 | -1,05591 | 0,291011 | 0,394202 | 2601,417 | -0,30363 | 0,222075 | -1,36726 | 0,171545 | 0,231975 | 2356,873055 | -0,643072052 | 0,156829 | -4,10048 | 4,12E-05 | 0,00011 |
| 2376 | RM25_RS11600 | RM25_2349 | 16S rRNA (guanine(527)-N(7))-methyltransferase | 2466 | gene | 2644996 | 2645766 | - | 862,6504 | 0,655993286 | 0,472093 | 1,389543 | 0,164668 | 0,267091 | 1289,51353 | 0,615548538 | 0,195332 | 3,151297 | 0,001625 | 0,004093 | 435,7873 | 0,780149506 | 0,193529 | 4,031171 | 5,55E-05 | 0,000453 | 714,3523 | -1,7274 | 0,232603 | -7,42639 | 1,12E-13 | 3,29E-12 | 1035,493025 | -1,511892975 | 0,132391 | -11,4199 | 3,32E-30 | 2,01E-28 |
| 2377 | NA | RM25_2350 | Hypothetical protein | 2467 | gene | 2645789 | 2646736 | - | 2025,578 | 0,786323602 | 0,487534 | 1,612858 | 0,106775 | 0,200853 | 2930,464461 | 0,511360641 | 0,177149 | 2,886613 | 0,003894 | 0,008792 | 1120,691 | 1,597169509 | 0,233217 | 6,848421 | 7,47E-12 | 3,52E-10 | 1587,187 | -2,17572 | 0,209399 | -10,3903 | 2,75E-25 | 4,98E-23 | 2512,672997 | -1,045846837 | 0,162894 | -6,42042 | 1,36E-10 | 8,63E-10 |
| 2378 | RM25_RS11610 | RM25_2351 | Membrane protein OxaA | 2468 | gene | 2646788 | 2647879 | - | 1580,78 | 0,480539524 | 0,360603 | 1,3326 | 0,182663 | 0,287581 | 2154,837227 | 0,393266562 | 0,13819 | 2,845837 | 0,004429 | 0,009785 | 1006,722 | 0,671267772 | 0,236009 | 2,844246 | 0,004452 | 0,01642 | 1404,62 | -1,32586 | 0,212187 | -6,24852 | 4,14E-10 | 6,47E-09 | 1805,598541 | -0,995227803 | 0,122251 | -8,14085 | 3,93E-16 | 5,57E-15 |
| 2379 | RM25_RS11615 | RM25_2352 | Putative membrane protein insertion efficiency factor | 2469 | gene | 2647890 | 2648240 | - | 475,8213 | 0,566802596 | 0,390098 | 1,452976 | 0,14623 | 0,246895 | 665,8931816 | 0,546146428 | 0,196535 | 2,778879 | 0,005455 | 0,011849 | 285,7494 | 0,616092478 | 0,235841 | 2,612321 | 0,008993 | 0,028075 | 407,4994 | -1,32147 | 0,231521 | -5,70777 | 1,14E-08 | 1,25E-07 | 557,1829586 | -1,201271506 | 0,170929 | -7,02789 | 2,1E-12 | 1,73E-11 |
| 2380 | RM25_RS11620 | RM25_2353 | Ribonuclease P protein component | 2470 | gene | 2648237 | 2648665 | - | 159,3965 | 1,202353501 | 0,328044 | 3,665222 | 0,000247 | 0,003386 | 177,4351031 | 0,645826473 | 0,254689 | 2,535744 | 0,011221 | 0,021832 | 141,358 | 2,047264432 | 0,267449 | 7,654788 | 1,94E-14 | 1,43E-12 | 102,9308 | -1,39123 | 0,304889 | -4,56306 | 5,04E-06 | 3,35E-05 | 217,3433661 | 0,066594508 | 0,184946 | 0,360075 | 0,718791 | 0,765909 |
| 22 | RM25_RS00110 | RM25_2358 | tRNA-Ile | 23 | tRNA | 30167 | 30243 | + | 97,26144 | -0,510564591 | 0,285741 | -1,78681 | 0,073968 | 0,159948 | 122,1523314 | -0,621836555 | 0,290514 | -2,14047 | 0,032317 | 0,054603 | 72,37054 | -0,312617848 | 0,24815 | -1,25979 | 0,207744 | 0,304321 | 121,2275 | -0,92994 | 0,25516 | -3,64456 | 0,000268 | 0,001161 | 78,72743887 | -0,574694987 | 0,317061 | -1,81257 | 0,069898 | 0,099849 |
| 156 | RM25_RS00770 | RM25_2363 | tRNA-Ser | 159 | tRNA | 195833 | 195918 | + | 73,89842 | -0,790698823 | 0,214231 | -3,69086 | 0,000223 | 0,003115 | 76,57241177 | -0,899055447 | 0,334128 | -2,69075 | 0,007129 | 0,014818 | 71,22443 | -0,68378554 | 0,326719 | -2,09289 | 0,036359 | 0,080123 | 99,01042 | -0,22378 | 0,225158 | -0,9939 | 0,320271 | 0,395639 | 53,11296218 | 0,038330011 | 0,459931 | 0,083339 | 0,933582 | 0,946025 |
| 159 | RM25_RS00785 | RM25_2368 | tRNA-Ser | 164 | tRNA | 199700 | 199792 | + | 69,42121 | -0,830726998 | 0,31103 | -2,67089 | 0,007565 | 0,038785 | 56,36759222 | -1,25783727 | 0,351636 | -3,5771 | 0,000347 | 0,001057 | 82,47483 | -0,574975095 | 0,357495 | -1,60834 | 0,10776 | 0,182334 | 93,88632 | 0,240175 | 0,297266 | 0,807946 | 0,419122 | 0,494677 | 48,98314241 | 0,992364952 | 0,444888 | 2,230595 | 0,025708 | 0,041157 |
| 160 | RM25_RS00790 | RM25_2373 | tRNA-Arg | 167 | tRNA | 199977 | 200052 | + | 8,970433 | -1,754053836 | 0,831457 | -2,10961 | 0,034892 | 0,09961 | 4,702222872 | -4,443942223 | 1,672515 | -2,65704 | 0,007883 | 0,016199 | 13,23864 | -1,271552645 | 0,479814 | -2,65009 | 0,008047 | 0,026088 | 14,63676 | 0,931034 | 0,51349 | 1,813147 | 0,069809 | 0,111456 | 3,955832986 | 4,19619914 | 1,748756 | 2,399534 | 0,016416 | 0,0275 |
| 275 | RM25_RS01345 | RM25_2378 | tRNA-Gly | 284 | tRNA | 336166 | 336239 | - | 30,4721 | -0,666098155 | 0,295635 | -2,25311 | 0,024252 | 0,079613 | 28,74577035 | -0,550270432 | 0,480978 | -1,14406 | 0,252597 | 0,3177 | 32,19844 | -0,765797842 | 0,420856 | -1,81962 | 0,068817 | 0,129039 | 39,09226 | 0,197771 | 0,3943 | 0,501574 | 0,615967 | 0,674958 | 23,21629485 | 0,038426828 | 0,534306 | 0,071919 | 0,942666 | 0,953527 |
| 355 | RM25_RS01700 | RM25_2383 | tRNA-Ser | 364 | tRNA | 428286 | 428373 | - | 78,12612 | -1,990013232 | 0,875357 | -2,27337 | 0,023004 | 0,07779 | 43,01660477 | -5,189524352 | 0,832238 | -6,23562 | 4,5E-10 | 4E-09 | 113,2356 | -1,452528916 | 0,364035 | -3,99008 | 6,61E-05 | 0,000518 | 131,2683 | 0,909707 | 0,453558 | 2,005712 | 0,044887 | 0,07808 | 30,89813013 | 4,719834501 | 0,865475 | 5,453464 | 4,94E-08 | 2,14E-07 |
| 394 | RM25_RS01895 | RM25_2388 | tRNA-Leu | 405 | tRNA | 480541 | 480627 | - | 87,45931 | -1,633346134 | 0,640025 | -2,552 | 0,010711 | 0,049019 | 49,91682743 | -3,033519004 | 0,644127 | -4,70951 | 2,48E-06 | 1,17E-05 | 125,0018 | -1,245948019 | 0,320772 | -3,88422 | 0,000103 | 0,000761 | 138,7288 | 0,909572 | 0,529313 | 1,718402 | 0,085723 | 0,130694 | 41,77926335 | 2,765995115 | 0,527452 | 5,244074 | 1,57E-07 | 6,2E-07 |
| 452 | RM25_RS02180 | RM25_2393 | tRNA-Phe | 465 | tRNA | 550596 | 550669 | - | 52,04317 | -0,786814672 | 0,269265 | -2,92208 | 0,003477 | 0,022892 | 43,41800033 | -0,528016519 | 0,357443 | -1,47721 | 0,139621 | 0,191887 | 60,66833 | -0,979086933 | 0,364315 | -2,68748 | 0,007199 | 0,023911 | 69,18795 | 0,585838 | 0,266306 | 2,199865 | 0,027817 | 0,053088 | 37,59313549 | 0,198626522 | 0,480121 | 0,413701 | 0,679093 | 0,729878 |
| 453 | RM25_RS02185 | RM25_2398 | tRNA-Asp | 468 | tRNA | 550683 | 550756 | - | 333,8197 | -0,367055952 | 0,438943 | -0,83623 | 0,403027 | 0,513967 | 478,6991742 | -0,237179742 | 0,274668 | -0,86351 | 0,387855 | 0,456631 | 188,9403 | -0,70313281 | 0,187905 | -3,74197 | 0,000183 | 0,00123 | 400,3325 | -1,20502 | 0,226909 | -5,31057 | 1,09E-07 | 1,01E-06 | 286,276719 | -1,618944744 | 0,282948 | -5,7217 | 1,05E-08 | 5,02E-08 |
| 454 | RM25_RS02190 | RM25_2403 | tRNA-Glu | 471 | tRNA | 550958 | 551030 | - | 509,1972 | 0,297674482 | 0,39958 | 0,744968 | 0,456291 | 0,561236 | 696,0367053 | 0,228768606 | 0,245507 | 0,931821 | 0,351429 | 0,42068 | 322,3576 | 0,451907626 | 0,364645 | 1,239307 | 0,215232 | 0,312446 | 489,442 | -1,29674 | 0,382576 | -3,3895 | 0,0007 | 0,002624 | 549,9683969 | -1,0284712 | 0,237223 | -4,33545 | 1,45E-05 | 4,18E-05 |
| 459 | RM25_RS02215 | RM25_2408 | tRNA-Lys | 479 | tRNA | 556207 | 556279 | + | 65,85711 | -1,050946753 | 0,278895 | -3,76825 | 0,000164 | 0,002471 | 65,39428596 | -0,888541987 | 0,425283 | -2,08929 | 0,036681 | 0,060843 | 66,31993 | -1,207039801 | 0,365101 | -3,30604 | 0,000946 | 0,004656 | 93,87725 | 0,054024 | 0,295121 | 0,183056 | 0,854754 | 0,885795 | 42,32541102 | -0,195746047 | 0,562835 | -0,34779 | 0,728001 | 0,773624 |
| 509 | RM25_RS02455 | RM25_2413 | tRNA-Tyr | 529 | tRNA | 609677 | 609759 | + | 189,9109 | -1,049968126 | 0,166886 | -6,29153 | 3,14E-10 | 3,9E-08 | 203,7594395 | -1,181220073 | 0,232116 | -5,08893 | 3,6E-07 | 1,95E-06 | 176,0623 | -0,906147701 | 0,256988 | -3,52603 | 0,000422 | 0,002449 | 271,4328 | -0,36087 | 0,204266 | -1,76669 | 0,077281 | 0,119994 | 121,2857053 | -0,023381952 | 0,31413 | -0,07443 | 0,940665 | 0,951974 |
| 512 | RM25_RS02470 | RM25_2418 | tRNA-Thr | 534 | tRNA | 612577 | 612649 | + | 1151,919 | -0,199452924 | 0,848917 | -0,23495 | 0,814248 | 0,870772 | 2071,372232 | -0,292408819 | 0,291205 | -1,00413 | 0,315314 | 0,382302 | 232,4654 | 0,639542673 | 0,607114 | 1,053415 | 0,292151 | 0,395293 | 1325,338 | -3,69856 | 0,558645 | -6,6206 | 3,58E-11 | 7,15E-10 | 1051,865743 | -2,741996666 | 0,377577 | -7,26209 | 3,81E-13 | 3,51E-12 |
| 513 | RM25_RS02475 | RM25_2423 | tRNA-Met | 537 | tRNA | 612728 | 612803 | + | 3056,48 | 0,524277668 | 0,694442 | 0,754963 | 0,450271 | 0,556523 | 4844,469893 | 0,203343736 | 0,297538 | 0,68342 | 0,494341 | 0,560174 | 1268,491 | 1,990669801 | 0,538419 | 3,69725 | 0,000218 | 0,001414 | 2695,301 | -3,19252 | 0,523465 | -6,09881 | 1,07E-09 | 1,5E-08 | 3528,904917 | -1,377502806 | 0,32776 | -4,20277 | 2,64E-05 | 7,33E-05 |
| 519 | RM25_RS02505 | RM25_2428 | tRNA-Trp | 545 | tRNA | 617033 | 617108 | + | 599,8507 | 0,136945023 | 0,528026 | 0,259353 | 0,795363 | 0,857111 | 903,0953055 | 8,30816E-05 | 0,262495 | 0,000317 | 0,999747 | 0,999747 | 296,6061 | 0,567852491 | 0,424868 | 1,33654 | 0,181373 | 0,274211 | 610,4296 | -1,96511 | 0,439 | -4,47634 | 7,59E-06 | 4,82E-05 | 615,6986754 | -1,365900812 | 0,240378 | -5,68229 | 1,33E-08 | 6,19E-08 |
| 753 | RM25_RS03635 | RM25_2433 | tRNA-Leu | 781 | tRNA | 862613 | 862697 | - | 213,8442 | -0,000701225 | 0,21222 | -0,0033 | 0,997364 | 0,998871 | 250,5568859 | 0,127164428 | 0,191827 | 0,66291 | 0,507388 | 0,572481 | 177,1314 | -0,175433508 | 0,24787 | -0,70776 | 0,479091 | 0,580284 | 225,8602 | -0,40851 | 0,217265 | -1,88024 | 0,060075 | 0,098467 | 209,7844053 | -0,6581141 | 0,203531 | -3,23348 | 0,001223 | 0,002533 |
| 757 | RM25_RS03655 | RM25_2438 | tRNA-Lys | 787 | tRNA | 864276 | 864348 | + | 19,86133 | -0,431588384 | 0,358582 | -1,2036 | 0,228745 | 0,336339 | 23,48728729 | -0,298826777 | 0,524149 | -0,57012 | 0,568598 | 0,630676 | 16,23537 | -0,588319297 | 0,475519 | -1,23722 | 0,216007 | 0,313117 | 24,19998 | -0,48856 | 0,377836 | -1,29306 | 0,195992 | 0,259232 | 16,70196056 | -0,71186088 | 0,695757 | -1,02315 | 0,306239 | 0,368456 |
| 763 | RM25_RS03685 | RM25_2443 | tRNA-His | 795 | tRNA | 871048 | 871123 | - | 45,99699 | -0,781061106 | 0,271584 | -2,87595 | 0,004028 | 0,025716 | 43,14410167 | -0,664971183 | 0,477781 | -1,39179 | 0,163986 | 0,219213 | 48,84988 | -0,877392583 | 0,31918 | -2,7489 | 0,00598 | 0,020726 | 61,27354 | 0,183774 | 0,40149 | 0,457729 | 0,647147 | 0,701944 | 33,33805841 | 0,036603799 | 0,458371 | 0,079856 | 0,936352 | 0,948423 |
| 766 | RM25_RS03700 | RM25_2448 | tRNA-Arg | 800 | tRNA | 873250 | 873325 | + | 69,11192 | -1,12049829 | 0,290786 | -3,85335 | 0,000117 | 0,001919 | 61,19858816 | -1,159171889 | 0,368971 | -3,14164 | 0,00168 | 0,004208 | 77,02525 | -1,097985296 | 0,417899 | -2,6274 | 0,008604 | 0,027258 | 99,96174 | 0,246134 | 0,269521 | 0,913226 | 0,361124 | 0,437169 | 42,7832039 | 0,379119817 | 0,565172 | 0,670804 | 0,502345 | 0,563823 |
| 792 | RM25_RS03830 | RM25_2453 | tRNA-Pro | 828 | tRNA | 908249 | 908324 | - | 30,17245 | -0,504986852 | 0,375412 | -1,34515 | 0,178576 | 0,282675 | 37,78125567 | -0,081535974 | 0,377283 | -0,21611 | 0,828899 | 0,864094 | 22,56364 | -1,240181499 | 0,406358 | -3,05194 | 0,002274 | 0,009585 | 37,27629 | -0,35329 | 0,310766 | -1,13684 | 0,255605 | 0,326183 | 24,69266936 | -1,450328661 | 0,516972 | -2,80543 | 0,005025 | 0,009318 |
| 793 | RM25_RS03835 | RM25_2458 | tRNA-Gly | 831 | tRNA | 908584 | 908657 | + | 60,18091 | -1,269627462 | 0,240344 | -5,28255 | 1,27E-07 | 7,7E-06 | 62,6957757 | -1,283887903 | 0,356166 | -3,60475 | 0,000312 | 0,000955 | 57,66604 | -1,253339695 | 0,376719 | -3,32698 | 0,000878 | 0,004384 | 90,07152 | -0,18524 | 0,250126 | -0,74058 | 0,458948 | 0,532352 | 34,61552989 | -0,093447497 | 0,545623 | -0,17127 | 0,864013 | 0,889292 |
| 830 | RM25_RS04010 | RM25_2463 | tRNA-Ala | 869 | tRNA | 947114 | 947189 | - | 147,85 | -0,680978227 | 0,390778 | -1,74262 | 0,0814 | 0,16851 | 200,1206844 | -0,625467732 | 0,328612 | -1,90336 | 0,056993 | 0,087685 | 95,57934 | -0,786959314 | 0,288889 | -2,72409 | 0,006448 | 0,021931 | 193,2201 | -1,05691 | 0,277441 | -3,80949 | 0,000139 | 0,000654 | 111,6531994 | -1,161972625 | 0,384052 | -3,02556 | 0,002482 | 0,004875 |
| 878 | RM25_RS04240 | RM25_2468 | tRNA-Gln | 919 | tRNA | 999918 | 999989 | + | 1447,449 | 0,66145171 | 0,725811 | 0,911328 | 0,362123 | 0,473552 | 2506,25602 | 0,710250238 | 0,297375 | 2,388402 | 0,016922 | 0,03116 | 388,6422 | 0,359162418 | 0,406233 | 0,884129 | 0,376627 | 0,482451 | 1205,48 | -2,53492 | 0,462973 | -5,47531 | 4,37E-08 | 4,33E-07 | 1741,142412 | -2,849235507 | 0,227188 | -12,5413 | 4,44E-36 | 4,02E-34 |
| 879 | RM25_RS04245 | RM25_2473 | tRNA-Glu | 922 | tRNA | 1000083 | 1000158 | + | 63,94581 | 0,28617693 | 0,314334 | 0,910424 | 0,362599 | 0,47374 | 81,17459819 | 0,393227123 | 0,326562 | 1,204142 | 0,228535 | 0,291955 | 46,71701 | 0,12563747 | 0,305072 | 0,411828 | 0,680465 | 0,757608 | 61,08817 | -0,70007 | 0,292662 | -2,39207 | 0,016753 | 0,035607 | 69,00659579 | -0,921253901 | 0,361444 | -2,54881 | 0,010809 | 0,018774 |
| 958 | RM25_RS04630 | RM25_2478 | tRNA-Val | 1003 | tRNA | 1096391 | 1096465 | - | 94,00786 | -1,323594302 | 0,485341 | -2,72714 | 0,006389 | 0,035179 | 65,14682177 | -2,217568461 | 0,419579 | -5,28523 | 1,26E-07 | 7,44E-07 | 122,8689 | -0,947753301 | 0,429323 | -2,20755 | 0,027275 | 0,06484 | 141,2554 | 0,527885 | 0,29086 | 1,814913 | 0,069537 | 0,111194 | 52,81549889 | 1,8657143 | 0,586101 | 3,183264 | 0,001456 | 0,00297 |
| 959 | RM25_RS04635 | RM25_2483 | tRNA-Gly | 1006 | tRNA | 1096869 | 1096941 | + | 161,9604 | -0,141031763 | 0,428294 | -0,32929 | 0,741938 | 0,812238 | 230,1825123 | 0,030065694 | 0,255722 | 0,117572 | 0,906407 | 0,92887 | 93,73827 | -0,568649204 | 0,23803 | -2,38898 | 0,016895 | 0,045304 | 181,1924 | -1,09173 | 0,230466 | -4,73706 | 2,17E-06 | 1,54E-05 | 151,2136491 | -1,632708936 | 0,294791 | -5,53852 | 3,05E-08 | 1,36E-07 |
| 960 | RM25_RS04640 | RM25_2488 | tRNA-Cys | 1009 | tRNA | 1097003 | 1097073 | + | 72,02026 | 0,103175637 | 0,399479 | 0,258275 | 0,796194 | 0,8573 | 99,04700266 | 0,319973049 | 0,325355 | 0,983457 | 0,325382 | 0,393497 | 44,99352 | -0,353004577 | 0,271354 | -1,3009 | 0,193292 | 0,289264 | 73,75137 | -0,85763 | 0,282285 | -3,03816 | 0,00238 | 0,007315 | 73,25820662 | -1,478168945 | 0,348158 | -4,24569 | 2,18E-05 | 6,14E-05 |
| 961 | RM25_RS04645 | RM25_2493 | tRNA-Val | 1012 | tRNA | 1097075 | 1097149 | + | 12,09235 | -0,789085084 | 0,369352 | -2,1364 | 0,032647 | 0,095588 | 11,93875894 | -0,68569091 | 0,697948 | -0,98244 | 0,325884 | 0,393498 | 12,24593 | -0,891151429 | 0,502068 | -1,77496 | 0,075904 | 0,139286 | 16,30182 | 0,062601 | 0,419539 | 0,149214 | 0,881385 | 0,905591 | 8,657108489 | -0,090339763 | 0,874311 | -0,10333 | 0,917704 | 0,933134 |
| 992 | RM25_RS04800 | RM25_2498 | tRNA-Met | 1045 | tRNA | 1128693 | 1128766 | + | 31,60947 | -0,632190384 | 0,280359 | -2,25493 | 0,024138 | 0,079348 | 28,62752893 | -0,249369336 | 0,462876 | -0,53874 | 0,590067 | 0,651118 | 34,59141 | -0,94573687 | 0,36748 | -2,57358 | 0,010065 | 0,030415 | 40,38385 | 0,474772 | 0,316927 | 1,49805 | 0,13412 | 0,189181 | 24,48615036 | -0,153610104 | 0,558429 | -0,27508 | 0,783258 | 0,824045 |
| 1001 | RM25_RS04855 | RM25_2503 | tRNA-Asn | 1056 | tRNA | 1138658 | 1138733 | + | 28,24577 | -0,172657327 | 0,501295 | -0,34442 | 0,730528 | 0,80423 | 16,92458223 | -0,031380757 | 0,662555 | -0,04736 | 0,962224 | 0,969629 | 39,56695 | -0,239904184 | 0,424139 | -0,56563 | 0,571648 | 0,663731 | 31,09522 | 1,300792 | 0,455704 | 2,854468 | 0,004311 | 0,011679 | 26,08078503 | 1,136094338 | 0,599626 | 1,894671 | 0,058136 | 0,084637 |
| 1088 | RM25_RS05285 | RM25_2508 | tRNA-Pro | 1145 | tRNA | 1239593 | 1239669 | - | 114,629 | -0,833053855 | 0,295682 | -2,81739 | 0,004841 | 0,028963 | 134,5596174 | -0,57823312 | 0,303257 | -1,90674 | 0,056554 | 0,087285 | 94,69832 | -1,198314481 | 0,324158 | -3,6967 | 0,000218 | 0,001414 | 155,3199 | -0,34199 | 0,20999 | -1,62859 | 0,103401 | 0,1528 | 81,17929984 | -0,9041775 | 0,441872 | -2,04624 | 0,040733 | 0,061701 |
| 1162 | RM25_RS05655 | RM25_2513 | tRNA-Arg | 1221 | tRNA | 1318608 | 1318682 | + | 109,0597 | -0,594748415 | 0,253683 | -2,34446 | 0,019055 | 0,069848 | 123,9906684 | -0,47257528 | 0,337251 | -1,40126 | 0,161137 | 0,216041 | 94,12866 | -0,745862662 | 0,283644 | -2,62958 | 0,008549 | 0,027157 | 138,9426 | -0,34167 | 0,216762 | -1,57623 | 0,114974 | 0,166458 | 85,48979944 | -0,560608001 | 0,434098 | -1,29143 | 0,196554 | 0,250848 |
| 1187 | RM25_RS05775 | RM25_2518 | tRNA-Met | 1248 | tRNA | 1347595 | 1347671 | + | 52,75787 | -1,44743262 | 0,423729 | -3,41594 | 0,000636 | 0,007417 | 38,30370433 | -2,055647658 | 0,541648 | -3,79517 | 0,000148 | 0,000484 | 67,21203 | -1,166328283 | 0,312814 | -3,72851 | 0,000193 | 0,001279 | 81,51561 | 0,50543 | 0,40807 | 1,238585 | 0,215499 | 0,281247 | 27,64677537 | 1,470424798 | 0,529271 | 2,778206 | 0,005466 | 0,010057 |
| 1233 | RM25_RS06005 | RM25_2523 | tRNA-Val | 1295 | tRNA | 1406301 | 1406377 | + | 13,65167 | -2,02657748 | 0,649111 | -3,12208 | 0,001796 | 0,014109 | 8,961800385 | -3,849631413 | 1,139551 | -3,3782 | 0,00073 | 0,002028 | 18,34155 | -1,540827288 | 0,484103 | -3,18285 | 0,001458 | 0,006636 | 23,1008 | 0,645984 | 0,382768 | 1,687663 | 0,091476 | 0,137769 | 5,168786436 | 3,005673184 | 1,406875 | 2,136417 | 0,032645 | 0,050671 |
| 1354 | RM25_RS06585 | RM25_2528 | tRNA-Leu | 1425 | tRNA | 1531448 | 1531536 | + | 232,5164 | -0,341047949 | 0,35559 | -0,9591 | 0,337507 | 0,448232 | 309,247236 | -0,132913875 | 0,224276 | -0,59264 | 0,553425 | 0,61821 | 155,7855 | -0,759419477 | 0,227564 | -3,33718 | 0,000846 | 0,004281 | 276,0598 | -0,78436 | 0,21197 | -3,70032 | 0,000215 | 0,00095 | 201,5876646 | -1,35200935 | 0,263496 | -5,13104 | 2,88E-07 | 1,1E-06 |
| 1474 | RM25_RS07190 | RM25_2533 | tRNA-Leu | 1546 | tRNA | 1656395 | 1656480 | + | 85,53011 | -1,442540174 | 0,389686 | -3,7018 | 0,000214 | 0,003058 | 65,62446201 | -2,120644645 | 0,391731 | -5,41352 | 6,18E-08 | 3,88E-07 | 105,4358 | -1,100156658 | 0,359832 | -3,05742 | 0,002233 | 0,009498 | 131,9337 | 0,353367 | 0,347345 | 1,017338 | 0,308993 | 0,384101 | 45,06672416 | 1,449368529 | 0,462067 | 3,136705 | 0,001709 | 0,00343 |
| 1662 | RM25_RS08085 | RM25_2538 | tRNA-Leu | 1736 | tRNA | 1852547 | 1852622 | - | 122,2186 | 0,226562618 | 0,200975 | 1,127319 | 0,259608 | 0,367284 | 138,1371462 | 0,075243928 | 0,225899 | 0,333087 | 0,739069 | 0,787872 | 106,3001 | 0,422644593 | 0,273237 | 1,546805 | 0,12191 | 0,200939 | 119,3861 | -0,62702 | 0,210457 | -2,97935 | 0,002889 | 0,008564 | 128,8585139 | -0,233684777 | 0,265138 | -0,88137 | 0,378118 | 0,443616 |
| 1739 | RM25_RS08475 | RM25_2543 | tRNA-Ala | 1813 | tRNA | 1935104 | 1935179 | - | 69,64945 | -0,458576529 | 0,194891 | -2,35299 | 0,018623 | 0,069126 | 75,80987068 | -0,164032851 | 0,272421 | -0,60213 | 0,547087 | 0,612873 | 63,48904 | -0,794376219 | 0,269836 | -2,94393 | 0,003241 | 0,012881 | 84,75989 | -0,05585 | 0,240568 | -0,23215 | 0,816424 | 0,852219 | 57,94924952 | -0,625190731 | 0,31626 | -1,97683 | 0,048061 | 0,0714 |
| 1749 | RM25_RS08525 | RM25_2548 | tRNA-Arg | 1825 | tRNA | 1942934 | 1943009 | + | 102,935 | -1,481863128 | 0,387856 | -3,82066 | 0,000133 | 0,002105 | 69,32843147 | -1,743801553 | 0,39203 | -4,44813 | 8,66E-06 | 3,69E-05 | 136,5416 | -1,368042419 | 0,29918 | -4,57265 | 4,82E-06 | 5,43E-05 | 158,9734 | 0,810046 | 0,343945 | 2,355163 | 0,018515 | 0,038516 | 53,10792691 | 1,250959168 | 0,37718 | 3,31661 | 0,000911 | 0,001958 |
| 1780 | RM25_RS08670 | RM25_2553 | tRNA-Thr | 1864 | tRNA | 1976289 | 1976364 | + | 47,82984 | -1,061778718 | 0,319644 | -3,32175 | 0,000895 | 0,009413 | 39,96082088 | -1,278653496 | 0,419687 | -3,04668 | 0,002314 | 0,005582 | 55,69886 | -0,928150452 | 0,413574 | -2,24422 | 0,024818 | 0,060619 | 68,17114 | 0,300887 | 0,31731 | 0,948241 | 0,343007 | 0,418678 | 30,36690713 | 0,721493965 | 0,565504 | 1,275843 | 0,202011 | 0,256403 |
| 1869 | RM25_RS09110 | RM25_2558 | tRNA-Pro | 1955 | tRNA | 2083150 | 2083223 | - | 26,67144 | -0,033761317 | 0,353414 | -0,09553 | 0,923895 | 0,949267 | 31,29561918 | -0,052039209 | 0,483848 | -0,10755 | 0,914351 | 0,934978 | 22,04726 | 0,005032666 | 0,475683 | 0,01058 | 0,991559 | 0,995359 | 28,66452 | -0,57496 | 0,443859 | -1,29537 | 0,195193 | 0,258321 | 25,83532552 | -0,477429727 | 0,562695 | -0,84847 | 0,396176 | 0,461358 |
| 1902 | RM25_RS09270 | RM25_2563 | tRNA-Ser | 1990 | tRNA | 2121515 | 2121602 | - | 103,2257 | -0,542123232 | 0,283228 | -1,91409 | 0,055609 | 0,132797 | 123,1513057 | -0,233649572 | 0,273942 | -0,85292 | 0,393705 | 0,462824 | 83,30003 | -0,999908834 | 0,263575 | -3,79364 | 0,000148 | 0,001035 | 129,5057 | -0,31936 | 0,203197 | -1,57169 | 0,116023 | 0,167564 | 82,74216952 | -1,027641376 | 0,365874 | -2,80873 | 0,004974 | 0,009231 |
| 1978 | RM25_RS09635 | RM25_2568 | tRNA-Thr | 2067 | tRNA | 2203088 | 2203163 | - | 14,18284 | -0,011190428 | 0,729038 | -0,01535 | 0,987753 | 0,992808 | 4,915069298 | -0,439771195 | 1,025035 | -0,42903 | 0,667901 | 0,72213 | 23,45061 | 0,069506549 | 0,486446 | 0,142886 | 0,88638 | 0,91836 | 14,7901 | 1,978 | 0,524435 | 3,77168 | 0,000162 | 0,000748 | 13,89853248 | 2,532178883 | 0,852149 | 2,971522 | 0,002963 | 0,005716 |
| 2289 | RM25_RS11170 | RM25_2573 | tRNA-Ala | 2379 | tRNA | 2539184 | 2539259 | + | 51,5088 | -0,364571554 | 0,186544 | -1,95434 | 0,050661 | 0,12544 | 53,37559081 | -0,475886589 | 0,342228 | -1,39055 | 0,164361 | 0,219365 | 49,642 | -0,260802892 | 0,290438 | -0,89796 | 0,369205 | 0,475009 | 61,64553 | -0,25248 | 0,259288 | -0,97376 | 0,330176 | 0,405536 | 43,92627799 | 0,012682429 | 0,402914 | 0,031477 | 0,974889 | 0,980715 |
