## Supplemental Table 2 for "Aerobic Adaptation and Metabolic Dynamics of *Propionibacterium freudenreichii* DSM 20271: Insights from Comparative Transcriptomics and Surfaceome Analysis"

**Table S2**. List of 74 genes encoding transport proteins detected with more than two-fold higher expression under anaerobic conditions.

| Locus tag | Product/Feature name | Fold change |
|---|---|---|
| | ***Sampling point I*** | |
| RM25_RS02200 | ABC-2 type transporter | 43,24 |
| RM25_RS11055 | Sn-glycerol-3-phosphate transport membrane protein ABC transporter ugpA | 34,20 |
| RM25_RS00460 | ABC-type antimicrobial peptide transporter, permease component | 32,55 |
| RM25_RS11060 | Sn-glycerol-3-phosphate transport membrane protein ABC transporter ugpE | 30,60 |
| RM25_RS10225 | Putative ABC transporter membrane protein | 25,26 |
| RM25_RS08065 | Actinorhodin transporter | 20,58 |
| RM25_RS11070 | Drug resistance transporter, EmrB/QacA subfamily | 19,68 |
| RM25_RS02205 | Daunorubicin resistance ABC transporter ATPase subunit | 17,91 |
| RM25_RS00455 | Lipoprotein-releasing system ATP-binding protein LolD | 16,02 |
| RM25_RS10015 | Daunorubicin resistance ABC transporter ATPase subunit | 15,69 |
| RM25_RS00495 | Major myo-inositol transporter IolT | 14,22 |
| RM25_RS10390 | Protein AraJ | 14,16 |
| RM25_RS11260 | Putative transporter, DMT superfamily | 10,51 |
| RM25_RS08865 | Membrane-bound transporter | 10,42 |
| RM25_RS10230 | Amino acid ABC transporter, permease protein | 10,38 |
| RM25_RS01630 | Transporter, anaerobic C4-dicarboxylate uptake family protein | 9,62 |
| RM25_RS10220 | Glutamine ABC transporter, ATP-binding protein GlnQ | 9,14 |
| RM25_RS11275 | Putative cation-transporting P-type ATPase | 9,09 |
| RM25_RS10855 | Aliphatic compound ABC transporter, periplasmic substrate-binding protein | 8,36 |
| RM25_RS07820 | Inner membrane transport protein ynfM | 7,47 |
| RM25_RS10865 | ABC-type nitrate/sulfonate/bicarbonate transport system | 7,34 |
| RM25_RS03350 | Putative exporter of polyketide antibiotics | 7,05 |
| RM25_RS09805 | Lincomycin resistance protein | 6,91 |
| RM25_RS13170 | ATP binding protein of ABC transporter | 6,70 |
| RM25_RS08260 | Drug resistance transporter, EmrB/QacA subfamily | 6,03 |
| RM25_RS10020 | ABC-2 type transporter | 5,94 |
| RM25_RS02725 | Oligopeptide/dipeptide ABC transporter, ATPase subunit | 5,92 |
| RM25_RS01175 | Putative metabolite transport protein CsbC | 5,78 |
| RM25_RS01105 | Putative metabolite transport protein YaaU | 5,70 |
| RM25_RS10870 | Nitrate transport ATP-binding subunits C and D | 5,25 |
| RM25_RS09605 | Ammonium-transport integral membrane protein Amt | 5,23 |
| RM25_RS02730 | Oligopeptide transport system permease protein OppC | 5,18 |
| RM25_RS12090 | ABC transporter, permease protein | 5,00 |
| RM25_RS09395 | Phosphate-transport membrane ABC transporter pstA1 | 4,98 |
| RM25_RS09390 | Phosphate ABC transporter, permease protein PstC | 4,58 |
| RM25_RS09950 | 2-oxoglutarate/malate translocator | 4,52 |
| RM25_RS03135 | Putative iron-siderophore ABC transporter (ATP-binding protein) | 4,41 |
| RM25_RS01500 | Putative hydroxymethylpyrimidine transporter CytX | 4,14 |
| RM25_RS09400 | Phosphate-transport ATP-binding protein ABC transporter phoT | 4,12 |
| RM25_RS03075 | ABC transporter, permease protein YjfF | 4,04 |
| RM25_RS00575 | High-affinity gluconate transporter | 3,90 |
| RM25_RS07010 | ABC transporter, ATP-binding protein | 3,76 |
| RM25_RS00220 | D-serine/D-alanine/glycine transporter | 3,73 |
| RM25_RS10520 | Putative MFS-family membrane transport protein | 3,53 |
| RM25_RS03060 | ABC transporter periplasmic-binding protein ytfQ | 3,49 |
| RM25_RS07115 | Na+-efflux ABC transporter (ATP-binding protein) | 3,38 |
| RM25_RS08225 | CrcB-like protein | 3,35 |
| RM25_RS13175 | YtfR | 3,25 |

| | | |
|---|---|---|
| RM25_RS01270 | Hexose phosphate transport protein | 3,18 |
| RM25_RS10030 | Sulfate ABC transporter, permease protein CysW | 3,15 |
| RM25_RS01225 | Putative metabolite transport protein yaaU | 2,96 |
| RM25_RS09385 | Phosphate ABC transporter, phosphate-binding protein PstS | 2,91 |
| RM25_RS03070 | YtfT | 2,85 |
| RM25_RS07120 | ABC transporter (Permease) (Na+ exclusion) | 2,82 |
| RM25_RS09860 | Divalent metal cation transporter MntH | 2,76 |
| RM25_RS04420 | Auxin Efflux Carrier | 2,68 |
| RM25_RS09630 | ABC transporter, ATP-binding protein | 2,66 |
| RM25_RS00900 | Multidrug resistance membrane efflux protein emrB | 2,62 |
| RM25_RS09495 | HMP/thiamine import ATP-binding protein | 2,59 |
| RM25_RS06200 | Metal transport ABC transporter | 2,54 |
| RM25_RS11200 | Cadmium-transporting ATPase | 2,49 |
| RM25_RS03360 | Tetronasin-transport ATP-binding protein ABC transporter | 2,45 |
| RM25_RS10035 | Sulfate ABC transporter, permease protein CysT | 2,44 |
| RM25_RS08075 | Putative ABC-2 | 2,42 |
| RM25_RS09500 | Cobalt transport protein CbiQ | 2,41 |
| RM25_RS09625 | Antimicrobial peptide ABC transporter, permease protein | 2,33 |
| RM25_RS11445 | Chloride transporter, chloride channel family protein | 2,22 |
| RM25_RS08070 | Multidrug ABC transporter, ATP-binding protein | 2,18 |
| RM25_RS08245 | Multidrug resistance transporter, MFS superfamily protein | 2,15 |
| RM25_RS11230 | Oligopeptide-transport membrane protein ABC transporter oppB | 2,13 |
| RM25_RS09785 | Heavy metal transport/detoxification protein | 2,10 |
| RM25_RS06440 | Cobalt transport protein | 2,06 |
| RM25_RS11520 | Iron(III) dicitrate transport ATP-binding protein | 2,03 |
| RM25_RS00970 | Multidrug resistance ABC transporter, ATP-binding/permease protein | 2,02 |
| | ***sampling point III*** | |
| RM25_RS02085 | Gluconate permease | 13,09 |
| RM25_RS00220 | D-serine/D-alanine/glycine transporter | 11,37 |
| RM25_RS07650 | Anaerobic C4-dicarboxylate transporter | 7,75 |
| RM25_RS07010 | ABC transporter, ATP-binding protein | 2,56 |
| RM25_RS02200 | ABC-2 type transporter | 2,51 |
| RM25_RS13170 | ATP binding protein of ABC transporter | 2,45 |
| RM25_RS08065 | Actinorhodin transporter | 2,35 |
| RM25_RS02725 | Oligopeptide/dipeptide ABC transporter, ATPase subunit | 2,31 |
| RM25_RS08070 | Multidrug ABC transporter, ATP-binding protein | 2,29 |
| RM25_RS03075 | ABC transporter, permease protein YjfF | 2,27 |
| RM25_RS10230 | Amino acid ABC transporter, permease protein | 2,25 |
| RM25_RS09605 | Ammonium-transport integral membrane protein Amt | 2,24 |
| RM25_RS01175 | Putative metabolite transport protein CsbC | 2,23 |
| RM25_RS00575 | High-affinity gluconate transporter | 2,22 |
| RM25_RS10225 | Putative ABC transporter membrane protein | 2,22 |
| RM25_RS11055 | Sn-glycerol-3-phosphate transport membrane protein ABC transporter ugpA | 2,22 |
| RM25_RS11260 | Putative transporter, DMT superfamily | 2,21 |
| RM25_RS02730 | Oligopeptide transport system permease protein OppC | 2,20 |
| RM25_RS10030 | Sulfate ABC transporter, permease protein CysW | 2,19 |
| RM25_RS01630 | Transporter, anaerobic C4-dicarboxylate uptake family protein | 2,19 |
| RM25_RS03135 | Putative iron-siderophore ABC transporter (ATP-binding protein) | 2,18 |
| RM25_RS09805 | Lincomycin resistance protein | 2,18 |
| RM25_RS11060 | Sn-glycerol-3-phosphate transport membrane protein ABC transporter ugpE | 2,16 |
| RM25_RS03060 | ABC transporter periplasmic-binding protein ytfQ | 2,16 |
| RM25_RS04135 | Inner membrane metabolite transport protein yhjE | 2,14 |

| | | |
|---|---|---|
| RM25_RS09950 | 2-oxoglutarate/malate translocator | 2,12 |
| RM25_RS13175 | YtfR | 2,12 |
| RM25_RS01105 | Putative metabolite transport protein YaaU | 2,11 |
| RM25_RS04420 | Auxin Efflux Carrier | 2,10 |
| RM25_RS07120 | ABC transporter (Permease) (Na+ exclusion) | 2,09 |
| RM25_RS09395 | Phosphate-transport membrane ABC transporter pstA1 | 2,06 |
| RM25_RS03350 | Putative exporter of polyketide antibiotics | 2,05 |
| RM25_RS01225 | Putative metabolite transport protein yaaU | 2,03 |
| RM25_RS09860 | Divalent metal cation transporter MntH | 2,00 |
