## Supplemental Table 3 for "Aerobic Adaptation and Metabolic Dynamics of *Propionibacterium freudenreichii* DSM 20271: Insights from Comparative Transcriptomics and Surfaceome Analysis"

**Table S3.** List of all identified proteins with the detected raw intensity values for each. Reverse hits, potential contaminants, and proteins only identified by site were filtered out. Proteins were filtered to contain minimally 2/3 valid values in at least one of the growth environments. The number of transmembrane spanning domains (TMD) were predicted using TMHMM 2.0, subcellular localization of proteins were done with prediction tool PSORTb 3.0.3, and the presence of possible classical and non-classical signal peptide sequences were analysed with LipoP 1.0, SignalP 5.0, and SecretomeP 2.0. The COG categorization was accomplished with EggNOG 5.0.0.

| Protein IDs | NCBI-LocusTag | Protein name | Anaerobic 1 | Anaerobic 2 | Anaerobic 3 | Aerobic 1 | Aerobic 3 | Aerobic 4 | Razor + unique peptides | Unique peptides | Sequence coverage [%] | Unique + razor sequence coverage [%] | Unique sequence coverage [%] | Mol. weight [kDa] | Q-value | Score | Intensity | MS/MS count | Protein ID | LocusTag | PsortB | No. of TMHMM | LipoP | SecP Score | SignalP | COG |
|---|---|---|---|---|---|---|---|---|---|---|---|---|---|---|---|---|---|---|---|---|---|---|---|---|---|---|
| lcl\|CP010341.1_prot_AJQ89737.1_1 | RM25_RS00005 | Chromosomal replication initiator protein DnaA | 8572900 | 9192800 | 19858000 | 4552400 | 4755500 | 14248000 | 2 | 2 | 6,5 | 6,5 | 6,5 | 55,17 | 0 | 24,765 | 67110000 | 3 | AJQ89737.1_1 | RM25_0001 | Cytoplasmic | 0 | CYT | 0.103861 | OTHER | L |
| lcl\|CP010341.1_prot_AJQ89743.1_7 | RM25_RS00035 | ABC-type antimicrobial peptide transport system | 30511000 | 49771000 | 56692000 | 19896000 | 30831000 | 47489000 | 7 | 7 | 10,9 | 10,9 | 10,9 | 88,215 | 0 | 97,168 | 246920000 | 17 | AJQ89743.1_7 | RM25_0007 | CytoplasmicMembrane | 10 | SPI | 0.780506 | SP(Sec/SPI) | Q |
| lcl\|CP010341.1_prot_AJQ89745.1_9 | RM25_RS00040 | DNA polymerase III, beta subunit | 46471000 | 29559000 | 138150000 | 20027000 | 18762000 | 51546000 | 7 | 7 | 18,1 | 18,1 | 18,1 | 41,525 | 0 | 50,476 | 304520000 | 14 | AJQ89745.1_9 | RM25_0009 | Cytoplasmic | 0 | CYT | 0.045147 | OTHER | L |
| lcl\|CP010341.1_prot_AJQ89750.1_14 | RM25_RS00070 | DNA gyrase, B subunit | 31801000 | 62609000 | 189080000 | 28174000 | 36093000 | 81446000 | 12 | 12 | 19,6 | 19,6 | 19,6 | 75,053 | 0 | 111,6 | 443410000 | 18 | AJQ89750.1_14 | RM25_0014 | Cytoplasmic | 0 | CYT | 0.099985 | OTHER | L |
| lcl\|CP010341.1_prot_AJQ89751.1_15 | RM25_RS00075 | DNA gyrase, A subunit | 35355000 | 43781000 | 79931000 | 51184000 | 60296000 | 90733000 | 7 | 7 | 10,5 | 10,5 | 10,5 | 100,64 | 0 | 50,533 | 392330000 | 19 | AJQ89751.1_15 | RM25_0015 | Cytoplasmic | 0 | CYT | 0.111352 | OTHER | L |
| lcl\|CP010341.1_prot_AJQ89768.1_32 | RM25_RS00165 | Hypothetical protein | 1727900 | 2439400 | 5960700 | 3032000 | 0 | 4428500 | 1 | 1 | 8,8 | 8,8 | 8,8 | 14,354 | 0,0042872 | 7,5314 | 20062000 | 4 | AJQ89768.1_32 | RM25_0033 | Unknown | 0 | CYT | 0.451756 | OTHER | |
| lcl\|CP010341.1_prot_AJQ89777.1_41 | RM25_RS00200 | Putative hippurate hydrolase | 11044000 | 16032000 | 66359000 | 0 | 0 | 18207000 | 4 | 4 | 16,9 | 16,9 | 16,9 | 42,723 | 0 | 37,08 | 111640000 | 6 | AJQ89777.1_41 | RM25_0042 | Cytoplasmic | 0 | CYT | 0.054438 | OTHER | S |
| lcl\|CP010341.1_prot_AJQ89780.1_44 | RM25_RS00215 | Alanine dehydrogenase | 46584000 | 10872000 | 119010000 | 30866000 | 63283000 | 114370000 | 6 | 6 | 18,6 | 18,6 | 18,6 | 39,469 | 0 | 42,317 | 411080000 | 14 | AJQ89780.1_44 | RM25_0045 | Extracellular | 0 | CYT | 0.058250 | OTHER | E |
| lcl\|CP010341.1_prot_AJQ89782.1_46 | RM25_RS00225 | Putative conserved lipoprotein lppS | 135310000 | 194900000 | 379900000 | 136600000 | 196920000 | 369910016 | 4 | 4 | 20,6 | 20,6 | 20,6 | 42,251 | 0 | 84,683 | 1551200000 | 19 | AJQ89782.1_46 | RM25_0047 | Unknown | 0 | SPI | 0.868909 | TAT(Tat/SPI) | S |
| lcl\|CP010341.1_prot_AJQ89786.1_50 | RM25_RS00240 | Nitroreductase | 8008300 | 9162600 | 16377000 | 8935300 | 7791100 | 8631600 | 1 | 1 | 4,1 | 4,1 | 4,1 | 18,592 | 0 | 13,069 | 64155000 | 4 | AJQ89786.1_50 | RM25_0051 | Cytoplasmic | 0 | CYT | 0.157171 | OTHER | C |
| lcl\|CP010341.1_prot_AJQ89787.1_51 | RM25_RS00245 | Conserved transmembrane protein | 9441500 | 15501000 | 26231000 | 11659000 | 11185000 | 22920000 | 2 | 2 | 4,8 | 4,8 | 4,8 | 61,446 | 0 | 12,438 | 107410000 | 5 | AJQ89787.1_51 | RM25_0052 | CytoplasmicMembrane | 10 | CYT | 0.934179 | OTHER | M |
| lcl\|CP010341.1_prot_AJQ89791.1_55 | RM25_RS00265 | Glucose-1-phosphate thymidylyltransferase | 10271000 | 14001000 | 37167000 | 0 | 11889000 | 19434000 | 3 | 3 | 15,2 | 15,2 | 15,2 | 31,219 | 0 | 21,519 | 92762000 | 3 | AJQ89791.1_55 | RM25_0056 | Cytoplasmic | 0 | CYT | 0.072368 | OTHER | M |
| lcl\|CP010341.1_prot_AJQ89794.1_58 | RM25_RS00280 | Alpha-1,6-rhamnosyltransferase MigA | 6300800 | 2463600 | 12044000 | 2672600 | 5076000 | 9406400 | 2 | 2 | 7,8 | 7,8 | 7,8 | 32,589 | 0 | 12,063 | 37963000 | 1 | AJQ89794.1_58 | RM25_0059 | Cytoplasmic | 0 | CYT | 0.055567 | OTHER | |
| lcl\|CP010341.1_prot_AJQ89796.1_60 | RM25_RS11630 | Family 2 glycosyl transferase | 2502800 | 2400900 | 6955200 | 5565100 | 3630800 | 4538500 | 1 | 1 | 2,8 | 2,8 | 2,8 | 69,959 | 0,008982 | 6,111 | 25593000 | 1 | AJQ89796.1_60 | RM25_0061 | CytoplasmicMembrane | 0 | CYT | 0.059348 | OTHER | |
| lcl\|CP010341.1_prot_AJQ89799.1_63 | RM25_RS00310 | DTDP-4-keto-L-rhamnose reductase and dTDP-4-keto-6 | 26786000 | 18038000 | 42833000 | 29981000 | 68641000 | 66482000 | 8 | 8 | 23,3 | 23,3 | 23,3 | 52,496 | 0 | 61,687 | 266590000 | 15 | AJQ89799.1_63 | RM25_0064 | Cytoplasmic | 0 | CYT | 0.271305 | OTHER | M |
| lcl\|CP010341.1_prot_AJQ89800.1_64 | RM25_RS00315 | DTDP-glucose 4,6-dehydratase (Precursor) | 70985000 | 79100000 | 172760000 | 100440000 | 79094000 | 122080000 | 8 | 8 | 35,2 | 35,2 | 35,2 | 36,934 | 0 | 60,01 | 704150000 | 17 | AJQ89800.1_64 | RM25_0065 | Cytoplasmic | 0 | CYT | 0.258722 | OTHER | M |
| lcl\|CP010341.1_prot_AJQ89801.1_65 | RM25_RS11635 | Large surface protein A | 43384000512 | 52458999808 | 81229996032 | 22104000512 | 36900999168 | 63281999872 | 26 | 26 | 33,3 | 33,3 | 33,3 | 137,32 | 0 | 323,31 | 3,1594E+11 | 534 | AJQ89801.1_65 | RM25_0066 | Extracellular | 0 | SPI | 0.895517 | SP(Sec/SPI) | |
| lcl\|CP010341.1_prot_AJQ89802.1_66 | RM25_RS00335 | UDP-galactopyranose mutase | 59294000 | 59640000 | 165410000 | 55762000 | 66060000 | 101060000 | 9 | 9 | 28,1 | 28,1 | 28,1 | 45,347 | 0 | 223,77 | 536070000 | 31 | AJQ89802.1_66 | RM25_0067 | Cytoplasmic | 0 | CYT | 0.245122 | OTHER | M |
| lcl\|CP010341.1_prot_AJQ89803.1_67 | RM25_RS00340 | Bifunctional UDP-galactofuranosyl transferase glfT | 75067000 | 67553000 | 163340000 | 130600000 | 175520000 | 256530000 | 9 | 9 | 21,3 | 21,3 | 21,3 | 75,349 | 0 | 110,49 | 981220000 | 34 | AJQ89803.1_67 | RM25_0068 | Cytoplasmic | 0 | CYT | 0.349368 | OTHER | S |
| lcl\|CP010341.1_prot_AJQ89806.1_70 | RM25_RS11640 | Glycosyltransferase | 2167600 | 1690400 | 0 | 0 | 0 | 2812100 | 1 | 1 | 2,4 | 2,4 | 2,4 | 87,916 | 0,0042239 | 7,2629 | 6670200 | 1 | AJQ89806.1_70 | RM25_0071 | Cytoplasmic | 0 | CYT | 0.079184 | OTHER | |
| lcl\|CP010341.1_prot_AJQ89810.1_74 | RM25_RS00375 | Integral membrane protein | 4291100 | 1859700 | 0 | 6843600 | 4038700 | 8337200 | 1 | 1 | 7,1 | 7,1 | 7,1 | 17,209 | 0,004215 | 7,2149 | 25370000 | 2 | AJQ89810.1_74 | RM25_0075 | CytoplasmicMembrane | 3 | TMH | 0.930809 | OTHER | S |
| lcl\|CP010341.1_prot_AJQ89812.1_76 | RM25_RS00385 | Oxidoreductase, short chain dehydrogenase/reductase | 0 | 4647000 | 0 | 7797400 | 8248600 | 13070000 | 1 | 1 | 4 | 4 | 4 | 32,671 | 0,0098814 | 5,964 | 40004000 | 3 | AJQ89812.1_76 | RM25_0077 | Cytoplasmic | 0 | CYT | 0.053924 | OTHER | S |
| lcl\|CP010341.1_prot_AJQ89813.1_77 | RM25_RS00390 | Acetate kinase | 235230000 | 249210000 | 595320000 | 167690000 | 140750000 | 159600000 | 8 | 8 | 26,8 | 26,8 | 26,8 | 42,431 | 0 | 69,26 | 1622700000 | 38 | AJQ89813.1_77 | RM25_0078 | Cytoplasmic | 0 | CYT | 0.061288 | OTHER | C |
| lcl\|CP010341.1_prot_AJQ89814.1_78 | RM25_RS00395 | Phospho acetyltransferase | 232110000 | 169790000 | 503569984 | 395889984 | 407120000 | 663409984 | 16 | 16 | 36,7 | 36,7 | 36,7 | 52,851 | 0 | 128,62 | 2652700000 | 56 | AJQ89814.1_78 | RM25_0079 | Cytoplasmic | 0 | CYT | 0.066405 | OTHER | C |
| lcl\|CP010341.1_prot_AJQ89827.1_91 | RM25_RS00450 | Hypothetical protein | 0 | 6146000 | 8361200 | 3554700 | 0 | 7071200 | 1 | 1 | 6,8 | 6,8 | 6,8 | 33,706 | 0,0044101 | 8,2878 | 25133000 | 2 | AJQ89827.1_91 | RM25_0092 | Unknown | 1 | TMH | 0.234930 | OTHER | |
| lcl\|CP010341.1_prot_AJQ89830.1_94 | RM25_RS00465 | Putative helicase | 16506000 | 14405000 | 24340000 | 37817000 | 45083000 | 79708000 | 5 | 5 | 6,4 | 6,4 | 6,4 | 124,74 | 0 | 35,792 | 254380000 | 8 | AJQ89830.1_94 | RM25_0095 | Cytoplasmic | 0 | CYT | 0.082579 | OTHER | |
| lcl\|CP010341.1_prot_AJQ89833.1_97 | RM25_RS00480 | Hypothetical protein | 270700000 | 547049984 | 409420000 | 134130000 | 281900000 | 403900000 | 5 | 5 | 36 | 36 | 36 | 24,561 | 0 | 323,31 | 2235800000 | 33 | AJQ89833.1_97 | RM25_0098 | Unknown | 1 | TMH | 0.959591 | OTHER | |
| lcl\|CP010341.1_prot_AJQ89840.1_104 | RM25_RS00520 | ABC transporter solute-binding protein | 5813400 | 7566600 | 8830400 | 0 | 5441800 | 16018000 | 1 | 1 | 5,2 | 5,2 | 5,2 | 29,394 | 0,0044004 | 8,0813 | 43671000 | 2 | AJQ89840.1_104 | RM25_0105 | Unknown | 1 | SPII | 0.630163 | LIPO(Sec/SPII) | ET |
| lcl\|CP010341.1_prot_AJQ89841.1_105 | RM25_RS00525 | Arginine/ornithine binding protein | 265680000 | 324020000 | 573560000 | 158850000 | 157180000 | 312600000 | 5 | 5 | 18,2 | 18,2 | 18,2 | 31,534 | 0 | 57,177 | 1895100000 | 22 | AJQ89841.1_105 | RM25_0106 | Unknown | 0 | SPII | 0.802714 | LIPO(Sec/SPII) | ET |
| lcl\|CP010341.1_prot_AJQ89842.1_106 | RM25_RS00530 | Arginine/ornithine binding protein | 245460000 | 224270000 | 336769984 | 91733000 | 106580000 | 151520000 | 5 | 5 | 20,8 | 20,8 | 20,8 | 31,488 | 0 | 83,192 | 1207300000 | 29 | AJQ89842.1_106 | RM25_0107 | Unknown | 0 | SPII | 0.831717 | LIPO(Sec/SPII) | ET |
| lcl\|CP010341.1_prot_AJQ89844.1_108 | RM25_RS00540 | Arginine/ornithine transport ATP-binding protein | 4317200 | 3741800 | 8949800 | 6021500 | 7367400 | 9007100 | 1 | 1 | 6,5 | 6,5 | 6,5 | 30,101 | 0,004603 | 10,302 | 45588000 | 1 | AJQ89844.1_108 | RM25_0109 | CytoplasmicMembrane | 0 | CYT | 0.061319 | OTHER | E |
| lcl\|CP010341.1_prot_AJQ89845.1_109 | RM25_RS12160 | Hypothetical protein | 6344700 | 3302000 | 13771000 | 7992000 | 11087000 | 22158000 | 3 | 3 | 13,3 | 13,3 | 13,3 | 30,898 | 0 | 25,503 | 68297000 | 5 | AJQ89845.1_109 | RM25_0110 | Cytoplasmic | 0 | CYT | 0.444577 | OTHER | K |
| lcl\|CP010341.1_prot_AJQ89847.1_111 | RM25_RS00560 | Putative cystathionine beta-lyase PatB | 25696000 | 38681000 | 91683000 | 41821000 | 30545000 | 51098000 | 3 | 3 | 7 | 7 | 7 | 45,389 | 0 | 21,014 | 306640000 | 6 | AJQ89847.1_111 | RM25_0112 | Cytoplasmic | 0 | CYT | 0.084048 | OTHER | E |
| lcl\|CP010341.1_prot_AJQ89848.1_112 | RM25_RS00565 | Ribose-5-phosphate isomerase B | 0 | 2095300 | 18620000 | 7419900 | 5090100 | 0 | 2 | 2 | 26,4 | 26,4 | 26,4 | 15,486 | 0 | 14,453 | 38777000 | 1 | AJQ89848.1_112 | RM25_0113 | Cytoplasmic | 0 | CYT | 0.059166 | OTHER | |
| lcl\|CP010341.1_prot_AJQ89855.1_119 | RM25_RS12905 | Hypothetical protein | 11480000 | 13105000 | 29121000 | 17548000 | 20240000 | 29939000 | 2 | 2 | 3,7 | 3,7 | 3,7 | 53,443 | 0 | 13,324 | 134040000 | 7 | AJQ89855.1_119 | RM25_0120 | Cytoplasmic | 0 | SPI | 0.064769 | OTHER | H |
| lcl\|CP010341.1_prot_AJQ89857.1_121 | RM25_RS00610 | Peptidase, S8/S53 family | 709779968 | 1294700032 | 1814400000 | 441600000 | 699230016 | 1271200000 | 7 | 7 | 21 | 21 | 21 | 55,272 | 0 | 246,7 | 6746000000 | 64 | AJQ89857.1_121 | RM25_0122 | Unknown | 1 | SPI | 0.488491 | SP(Sec/SPI) | |
| lcl\|CP010341.1_prot_AJQ89859.1_123 | RM25_RS00620 | Hypothetical protein | 2523700 | 2851700 | 5791600 | 0 | 0 | 10669000 | 1 | 1 | 6 | 6 | 6 | 27,861 | 0,004386 | 7,9641 | 21836000 | 1 | AJQ89859.1_123 | RM25_0124 | CytoplasmicMembrane | 2 | SPI | 0.811203 | TAT(Tat/SPI) | S |
| lcl\|CP010341.1_prot_AJQ89860.1_124 | RM25_RS00625 | Citrate (Si)-synthase | 10824000 | 7521600 | 31516000 | 10615000 | 13965000 | 23452000 | 4 | 4 | 12,4 | 12,4 | 12,4 | 48,495 | 0 | 24,867 | 97893000 | 2 | AJQ89860.1_124 | RM25_0125 | Cytoplasmic | 0 | CYT | 0.112851 | OTHER | C |
| lcl\|CP010341.1_prot_AJQ89861.1_125 | RM25_RS00630 | Putative secreted peptidoglycan-binding protein | 3771200000 | 2973700096 | 4445100032 | 10289000448 | 14273999872 | 11687000064 | 4 | 4 | 34,7 | 34,7 | 34,7 | 20,027 | 0 | 322,7 | 5,4687E+10 | 129 | AJQ89861.1_125 | RM25_0126 | Extracellular | 0 | SPI | 0.875484 | SP(Sec/SPI) | |
| lcl\|CP010341.1_prot_AJQ89862.1_126 | RM25_RS00635 | Hypothetical protein | 18441000 | 12730000 | 46885000 | 21055000 | 11058000 | 27159000 | 5 | 5 | 10,8 | 10,8 | 10,8 | 68,896 | 0 | 37,415 | 152820000 | 8 | AJQ89862.1_126 | RM25_0127 | CytoplasmicMembrane | 1 | CYT | 0.925145 | OTHER | |
| lcl\|CP010341.1_prot_AJQ89864.1_128 | RM25_RS00645 | DNA-formamidopyrimidine glycosylase | 139130000 | 169220000 | 247250000 | 64179000 | 82224000 | 138780000 | 2 | 2 | 7,3 | 7,3 | 7,3 | 30,818 | 0 | 11,415 | 880270000 | 4 | AJQ89864.1_128 | RM25_0129 | Cytoplasmic | 0 | CYT | 0.059935 | OTHER | L |
| lcl\|CP010341.1_prot_AJQ89865.1_129 | RM25_RS00650 | Gluconokinase | 1504800 | 0 | 18201000 | 4106900 | 0 | 0 | 2 | 2 | 4,6 | 4,6 | 4,6 | 60,977 | 0 | 14,469 | 28292000 | 2 | AJQ89865.1_129 | RM25_0130 | Cytoplasmic | 1 | CYT | 0.138870 | OTHER | G |
| lcl\|CP010341.1_prot_AJQ89873.1_137 | RM25_RS00680 | Hypothetical protein | 0 | 0 | 0 | 0 | 3220900 | 3567000 | 1 | 1 | 3,3 | 3,3 | 3,3 | 36,978 | 0,0043621 | 7,8513 | 6787900 | 2 | AJQ89873.1_137 | RM25_0138 | Cytoplasmic | 0 | CYT | 0.086450 | OTHER | |
| lcl\|CP010341.1_prot_AJQ89876.1_140 | RM25_RS11670 | Hypothetical protein | 2359100 | 2999900 | 6725500 | 5374200 | 8957300 | 10368000 | 1 | 1 | 7,5 | 7,5 | 7,5 | 24,46 | 0,0043812 | 7,9474 | 41807000 | 4 | AJQ89876.1_140 | RM25_0141 | CytoplasmicMembrane | 2 | CYT | 0.975124 | OTHER | |
| lcl\|CP010341.1_prot_AJQ89879.1_143 | RM25_RS00710 | Serine--tRNA ligase | 65789000 | 55406000 | 191830000 | 104540000 | 94724000 | 145170000 | 7 | 7 | 24,8 | 24,8 | 24,8 | 47,319 | 0 | 79,991 | 718790000 | 28 | AJQ89879.1_143 | RM25_0144 | Cytoplasmic | 0 | CYT | 0.082324 | OTHER | J |
| lcl\|CP010341.1_prot_AJQ89880.1_144 | RM25_RS11675 | Hypothetical protein | 10548000 | 5894900 | 18010000 | 4427300 | 9902100 | 5502200 | 1 | 1 | 7,5 | 7,5 | 7,5 | 24,398 | 0,0045455 | 9,9306 | 57271000 | 7 | AJQ89880.1_144 | RM25_0145 | Cytoplasmic | 0 | CYT | 0.055365 | OTHER | |
| lcl\|CP010341.1_prot_AJQ89881.1_145 | RM25_RS00720 | PTS system mannose-specific EIIBCA component (EIIBC | 7647300 | 6553300 | 15995000 | 3530600 | 4492500 | 9687900 | 3 | 3 | 3,5 | 3,5 | 3,5 | 101,99 | 0 | 23,255 | 49508000 | 8 | AJQ89881.1_145 | RM25_0146 | CytoplasmicMembrane | 8 | CYT | 0.828886 | OTHER | |
| lcl\|CP010341.1_prot_AJQ89885.1_149 | RM25_RS00740 | Putative periplasmic or exported protein | 65252000 | 104170000 | 199490000 | 12742000 | 81597000 | 97425000 | 3 | 3 | 14,9 | 14,9 | 14,9 | 22,112 | 0 | 59,766 | 595480000 | 10 | AJQ89885.1_149 | RM25_0150 | Unknown | 0 | SPII | 0.833668 | LIPO(Sec/SPII) | |
| lcl\|CP010341.1_prot_AJQ89886.1_150 | RM25_RS00745 | putative thiol peroxidase | 31309000 | 24435000 | 209360000 | 358200000 | 301800000 | 364289984 | 4 | 4 | 45,5 | 45,5 | 45,5 | 17,439 | 0 | 96,766 | 1572600000 | 25 | AJQ89886.1_150 | RM25_0151 | Unknown | 0 | CYT | 0.060125 | OTHER | O |
| lcl\|CP010341.1_prot_AJQ89888.1_152 | RM25_RS00755 | Putative CONSERVED PROTEIN CPSA | 253220000 | 260920000 | 363030016 | 75863000 | 160400000 | 227520000 | 11 | 11 | 27,6 | 27,6 | 27,6 | 58,847 | 0 | 164,5 | 1407300000 | 36 | AJQ89888.1_152 | RM25_0153 | Unknown | 1 | CYT | 0.894593 | OTHER | |
| lcl\|CP010341.1_prot_AJQ89889.1_153 | RM25_RS00760 | Argininosuccinate synthase | 205190000 | 194050000 | 715939968 | 346609984 | 391670016 | 690969984 | 20 | 20 | 50,4 | 50,4 | 50,4 | 52,617 | 0 | 243,74 | 2880000000 | 60 | AJQ89889.1_153 | RM25_0154 | Cytoplasmic | 0 | CYT | 0.139415 | OTHER | E |
| lcl\|CP010341.1_prot_AJQ89890.1_154 | RM25_RS00765 | NAD(P)H quinone oxidoreductase, PIG3 family | 17766000 | 16607000 | 50566000 | 26594000 | 32874000 | 38482000 | 2 | 2 | 7,5 | 7,5 | 7,5 | 35,013 | 0 | 25,366 | 199020000 | 16 | AJQ89890.1_154 | RM25_0155 | Cytoplasmic | 0 | CYT | 0.043173 | OTHER | C |
| lcl\|CP010341.1_prot_AJQ89892.1_156 | RM25_RS00780 | Cell envelope-related transcriptional attenuator | 1902599936 | 2761400064 | 3472199936 | 909049984 | 1612099968 | 2238099968 | 13 | 13 | 20,8 | 20,8 | 20,8 | 90,81 | 0 | 305,68 | 1,3436E+10 | 100 | AJQ89892.1_156 | RM25_0158 | CytoplasmicMembrane | 4 | CYT | 0.865761 | OTHER | K |
| lcl\|CP010341.1_prot_AJQ89894.1_158 | RM25_RS00800 | Oxidoreductase, short chain dehydrogenase/reductase | 1443200 | 0 | 4668400 | 0 | 3521300 | 2211700 | 1 | 1 | 4,8 | 4,8 | 4,8 | 24,361 | 0,0042644 | 7,4838 | 11845000 | 2 | AJQ89894.1_158 | RM25_0162 | Cytoplasmic | 0 | CYT | 0.050720 | OTHER | S |
| lcl\|CP010341.1_prot_AJQ89896.1_160 | RM25_RS00810 | Shikimate dehydrogenase | 20565000 | 21350000 | 51666000 | 0 | 0 | 20682000 | 3 | 3 | 12,5 | 12,5 | 12,5 | 29,183 | 0 | 33,145 | 124600000 | 4 | AJQ89896.1_160 | RM25_0164 | Cytoplasmic | 0 | CYT | 0.040835 | OTHER | E |
| lcl\|CP010341.1_prot_AJQ89897.1_161 | RM25_RS00815 | 3-carboxymuconate cyclase (Precursor) | 6256800 | 8024400 | 8885200 | 0 | 0 | 0 | 3 | 3 | 15,2 | 15,2 | 15,2 | 38,268 | 0 | 18,55 | 23166000 | 3 | AJQ89897.1_161 | RM25_0165 | Cytoplasmic | 0 | CYT | 0.162919 | OTHER | |
| lcl\|CP010341.1_prot_AJQ89898.1_162 | RM25_RS00820 | 2,5-diketo-D-gluconic acid reductase A | 504889984 | 615969984 | 1113299968 | 559840000 | 937020032 | 937880000 | 12 | 12 | 66,1 | 66,1 | 66,1 | 30,462 | 0 | 232,72 | 5095000000 | 70 | AJQ89898.1_162 | RM25_0166 | Cytoplasmic | 0 | CYT | 0.104037 | OTHER | S |
| lcl\|CP010341.1_prot_AJQ89899.1_163 | RM25_RS00825 | ABC-type antimicrobial peptide transport system | 2018700032 | 5691400192 | 5526099968 | 1053100032 | 2406899968 | 2081900032 | 18 | 18 | 32,6 | 32,6 | 32,6 | 83,579 | 0 | 323,31 | 1,9846E+10 | 121 | AJQ89899.1_163 | RM25_0167 | CytoplasmicMembrane | 10 | SPI | 0.796157 | SP(Sec/SPI) | Q |
| lcl\|CP010341.1_prot_AJQ89900.1_164 | RM25_RS00830 | Ectoine/hydroxyectoine ABC transporter, ATP-binding p | 2654600 | 0 | 6974400 | 0 | 3239200 | 2251100 | 1 | 1 | 3,5 | 3,5 | 3,5 | 33,059 | 0,005123 | 6,6175 | 15119000 | 1 | AJQ89900.1_164 | RM25_0168 | CytoplasmicMembrane | 0 | CYT | 0.367215 | OTHER | V |
| lcl\|CP010341.1_prot_AJQ89901.1_165 | RM25_RS00835 | Two-component system response regulator | 8707100 | 17793000 | 28686000 | 9089200 | 18090000 | 16870000 | 3 | 3 | 23 | 23 | 23 | 25,067 | 0 | 31,752 | 109280000 | 8 | AJQ89901.1_165 | RM25_0169 | Cytoplasmic | 0 | CYT | 0.040393 | OTHER | T |
| lcl\|CP010341.1_prot_AJQ89903.1_167 | RM25_RS00845 | Carboxylesterase type B | 72583000 | 162550000 | 268590016 | 85302000 | 166680000 | 165620000 | 4 | 4 | 31,6 | 31,6 | 31,6 | 24,711 | 0 | 137,99 | 986760000 | 17 | AJQ89903.1_167 | RM25_0171 | Unknown | 1 | SPI | 0.535156 | OTHER | |
| lcl\|CP010341.1_prot_AJQ89905.1_169 | RM25_RS00855 | Acetyltransferase | 19137000 | 26156000 | 87305000 | 975720 | 10263000 | 47404000 | 3 | 3 | 36,5 | 36,5 | 36,5 | 11,668 | 0 | 28,23 | 201450000 | 7 | AJQ89905.1_169 | RM25_0173 | Cytoplasmic | 0 | CYT | 0.253136 | OTHER | |
| lcl\|CP010341.1_prot_AJQ89906.1_170 | RM25_RS00860 | Hypothetical protein | 72397000 | 89597000 | 115920000 | 73992000 | 89489000 | 94898000 | 2 | 2 | 23,4 | 23,4 | 23,4 | 8,5867 | 0 | 22,601 | 579910000 | 14 | AJQ89906.1_170 | RM25_0174 | Unknown | 0 | CYT | 0.402231 | OTHER | S |
| lcl\|CP010341.1_prot_AJQ89907.1_171 | RM25_RS00865 | Peptidyl-prolyl cis-trans isomerase | 202000000 | 309910016 | 354590016 | 266610000 | 316560000 | 411740000 | 3 | 3 | 56,5 | 56,5 | 56,5 | 13,027 | 0 | 103,48 | 2114300000 | 26 | AJQ89907.1_171 | RM25_0175 | Cytoplasmic | 0 | CYT | 0.167592 | OTHER | O |
| lcl\|CP010341.1_prot_AJQ89910.1_174 | RM25_RS00880 | Cytochrome bd-type menaquinol oxidase subunit I | 86129000 | 114270000 | 203070000 | 840190016 | 1648600064 | 2564300032 | 9 | 9 | 26,8 | 26,8 | 26,8 | 55,025 | 0 | 134,84 | 5967700000 | 41 | AJQ89910.1_174 | RM25_0178 | CytoplasmicMembrane | 9 | TMH | 0.949184 | OTHER | C |
| lcl\|CP010341.1_prot_AJQ89911.1_175 | RM25_RS00885 | Cytochrome d ubiquinol oxidase, subunit II | 7561600 | 5900400 | 11653000 | 9843700 | 18152000 | 29935000 | 2 | 2 | 11,4 | 11,4 | 11,4 | 41,282 | 0 | 12,25 | 83046000 | 2 | AJQ89911.1_175 | RM25_0179 | CytoplasmicMembrane | 8 | TMH | 0.957940 | OTHER | C |
| lcl\|CP010341.1_prot_AJQ89916.1_180 | RM25_RS00910 | Protein of hypothetical function DUF77 | 3418400 | 5022300 | 8301500 | 12703000 | 9572700 | 9311100 | 1 | 1 | 13,3 | 13,3 | 13,3 | 11,22 | 0,0044297 | 8,4086 | 59827000 | 4 | AJQ89916.1_180 | RM25_0184 | Cytoplasmic | 0 | CYT | 0.048256 | OTHER | S |
| lcl\|CP010341.1_prot_AJQ89917.1_181 | RM25_RS00915 | TRNA uridine 5-carboxymethylaminomethyl modificatic | 475809984 | 491870016 | 1002200000 | 467260000 | 515790016 | 961350016 | 22 | 22 | 45,3 | 45,3 | 45,3 | 59,855 | 0 | 213,31 | 4368100000 | 114 | AJQ89917.1_181 | RM25_0185 | Cytoplasmic | 0 | CYT | 0.184882 | OTHER | C |
| lcl\|CP010341.1_prot_AJQ89918.1_182 | RM25_RS00920 | Pyruvate-flavodoxin oxidoreductase | 463760000 | 395769984 | 1048800000 | 569910016 | 623729984 | 1353100032 | 30 | 30 | 32,6 | 32,6 | 32,6 | 136,55 | 0 | 323,31 | 4929100000 | 144 | AJQ89918.1_182 | RM25_0186 | Cytoplasmic | 0 | CYT | 0.215578 | OTHER | C |
| lcl\|CP010341.1_prot_AJQ89919.1_183 | RM25_RS00925 | Dihydroorotate dehydrogenase 2 | 541939968 | 409289984 | 894419968 | 686240000 | 392870016 | 1000899968 | 10 | 10 | 34,8 | 34,8 | 34,8 | 40,394 | 0 | 215,09 | 4305200000 | 68 | AJQ89919.1_183 | RM25_0187 | Unknown | 0 | SPI | 0.046635 | OTHER | |
| lcl\|CP010341.1_prot_AJQ89920.1_184 | RM25_RS00930 | Gamma-aminobutyraldehyde dehydrogenase | 11612000 | 7480000 | 29858000 | 19549000 | 13197000 | 22134000 | 1 | 1 | 4 | 4 | 4 | 50,327 | 0,0043573 | 7,8423 | 124350000 | 1 | AJQ89920.1_184 | RM25_0188 | Cytoplasmic | 0 | CYT | 0.090445 | OTHER | C |
| lcl\|CP010341.1_prot_AJQ89927.1_191 | RM25_RS00965 | Multidrug resistance ABC transporter, ATP-binding/pern | 1241600 | 0 | 2442700 | 0 | 0 | 1897700 | 1 | 1 | 1,9 | 1,9 | 1,9 | 73,882 | 0,008 | 6,1491 | 5581900 | 1 | AJQ89927.1_191 | RM25_0195 | CytoplasmicMembrane | 5 | CYT | 0.669969 | OTHER | V |
| lcl\|CP010341.1_prot_AJQ89934.1_198 | RM25_RS01000 | VarM | 2018800 | 2738600 | 5338300 | 6474500 | 6499500 | 9378800 | 2 | 2 | 4,6 | 4,6 | 4,6 | 60,009 | 0 | 11,243 | 38069000 | 3 | AJQ89934.1_198 | RM25_0202 | Cytoplasmic | 0 | CYT | 0.066027 | OTHER | S |
| lcl\|CP010341.1_prot_AJQ89935.1_199 | RM25_RS01005 | Lysine--tRNA ligase | 9416400 | 39091000 | 94957000 | 118260000 | 131640000 | 144490000 | 3 | 3 | 5,8 | 5,8 | 5,8 | 60,08 | 0 | 21,874 | 614850000 | 7 | AJQ89935.1_199 | RM25_0203 | Cytoplasmic | 0 | CYT | 0.077528 | OTHER | J |
| lcl\|CP010341.1_prot_AJQ89936.1_200 | RM25_RS01010 | CheR methyltransferase SAM binding domain-containin | 0 | 8780800 | 26639000 | 0 | 0 | 0 | 1 | 1 | 11,2 | 11,2 | 11,2 | 26,381 | 0 | 16,549 | 35419000 | 1 | AJQ89936.1_200 | RM25_0204 | Unknown | 0 | CYT | 0.487471 | OTHER | |
| lcl\|CP010341.1_prot_AJQ89937.1_201 | RM25_RS01015 | Pyruvate dehydrogenase E1 component, alpha subunit | 18956000 | 23715000 | 85212000 | 7566700 | 24520000 | 48112000 | 5 | 5 | 19,2 | 19,2 | 19,2 | 40,517 | 0 | 39,781 | 222980000 | 11 | AJQ89937.1_201 | RM25_0205 | Cytoplasmic | 0 | CYT | 0.419936 | OTHER | C |
| lcl\|CP010341.1_prot_AJQ89938.1_202 | RM25_RS01020 | Branched-chain alpha-keto acid dehydrogenase E1 beta | 13096000 | 3410100 | 68813000 | 25315000 | 14730000 | 38224000 | 3 | 3 | 12,3 | 12,3 | 12,3 | 35,589 | 0 | 20,236 | 163590000 | 5 | AJQ89938.1_202 | RM25_0206 | Cytoplasmic | 0 | CYT | 0.082878 | OTHER | C |
| lcl\|CP010341.1_prot_AJQ89939.1_203 | RM25_RS01025 | Dihydrolipoamide acyltransferase | 45706000 | 40681000 | 139460000 | 88901000 | 106810000 | 173190000 | 7 | 7 | 19,2 | 17,4 | 17,4 | 47,743 | 0 | 48,362 | 644440000 | 25 | AJQ89939.1_203 | RM25_0207 | Cytoplasmic | 0 | CYT | 0.141189 | OTHER | C |
| lcl\|CP010341.1_prot_AJQ89940.1_204 | RM25_RS11690 | UDP-N-acetylglucosamine diphosphorylase | 5778400 | 4146500 | 17583000 | 2416700 | 1547600 | 9330400 | 2 | 2 | 7,3 | 7,3 | 7,3 | 36,694 | 0 | 13,174 | 40803000 | 2 | AJQ89940.1_204 | RM25_0208 | Cytoplasmic | 0 | CYT | 0.120721 | OTHER | |
| lcl\|CP010341.1_prot_AJQ89942.1_206 | RM25_RS01040 | Membrane spanning protein DUF881 | 124650000 | 184740000 | 273230016 | 47847000 | 6427500 | 6027100 | 2 | 2 | 10,2 | 10,2 | 10,2 | 25,487 | 0 | 12,731 | 673700000 | 5 | AJQ89942.1_206 | RM25_0210 | CytoplasmicMembrane | 1 | SPI | 0.337388 | SP(Sec/SPI) | S |
| lcl\|CP010341.1_prot_AJQ89944.1_208 | RM25_RS01050 | Putative oxidoreductase YeaE, aldo/keto reductase fam | 3216600 | 4381900 | 8941700 | 8219900 | 5358300 | 7271800 | 1 | 1 | 6,5 | 6,5 | 6,5 | 32,094 | 0,0042105 | 7,1921 | 41736000 | 3 | AJQ89944.1_208 | RM25_0212 | Cytoplasmic | 0 | CYT | 0.151826 | OTHER | |
| lcl\|CP010341.1_prot_AJQ89945.1_209 | RM25_RS01055 | Hypothetical protein | 64423000 | 26120000 | 117910000 | 86344000 | 74848000 | 140940000 | 12 | 12 | 14,5 | 14,5 | 14,5 | 127,04 | 0 | 98,944 | 564890000 | 24 | AJQ89945.1_209 | RM25_0213 | Cytoplasmic | 0 | CYT | 0.250741 | OTHER | |
| lcl\|CP010341.1_prot_AJQ89947.1_211 | RM25_RS01065 | Secreted serine protease, trypsin-like serine proteases | 679240000 | 866390016 | 1410899968 | 1139100032 | 1159900032 | 1909600000 | 7 | 7 | 23,5 | 23,5 | 23,5 | 47,825 | 0 | 323,31 | 8369900000 | 68 | AJQ89947.1_211 | RM25_0215 | Unknown | 1 | CYT | 0.944305 | OTHER | O |
| lcl\|CP010341.1_prot_AJQ89948.1_212 | RM25_RS12190 | Trypsin-like serine protease | 740419968 | 1072800000 | 1445400064 | 756780032 | 1175399936 | 1539900032 | 5 | 5 | 18,4 | 18,4 | 18,4 | 45,09 | 0 | 240,18 | 7232600000 | 58 | AJQ89948.1_212 | RM25_0216 | CytoplasmicMembrane | 1 | CYT | 0.959651 | OTHER | O |
| lcl\|CP010341.1_prot_AJQ89951.1_215 | RM25_RS01085 | Crotonobetainyl-CoA dehydrogenase | 1080900 | 3720800 | 4747600 | 0 | 4285500 | 4971700 | 1 | 1 | 2,6 | 2,6 | 2,6 | 42,811 | 0,0070779 | 6,3204 | 21125000 | 0 | AJQ89951.1_215 | RM25_0219 | Cytoplasmic | 0 | CYT | 0.127107 | OTHER | I |
| lcl\|CP010341.1_prot_AJQ89958.1_222 | RM25_RS01115 | Putative electron transfer flavoprotein FixA | 25951000 | 39794000 | 77920000 | 39512000 | 58683000 | 62372000 | 5 | 5 | 24,7 | 24,7 | 24,7 | 25,759 | 0 | 51,789 | 343240000 | 17 | AJQ89958.1_222 | RM25_0226 | Unknown | 0 | CYT | 0.048054 | OTHER | |
| lcl\|CP010341.1_prot_AJQ89959.1_223 | RM25_RS01120 | Protein FixB | 28875000 | 50351000 | 93102000 | 14530000 | 65865000 | 82117000 | 4 | 4 | 14,1 | 14,1 | 14,1 | 29,581 | 0 | 26,305 | 403580000 | 5 | AJQ89959.1_223 | RM25_0227 | Cytoplasmic | 0 | CYT | 0.043769 | OTHER | |
| lcl\|CP010341.1_prot_AJQ89965.1_229 | RM25_RS01145 | SanA protein | 3109200 | 6503600 | 5330200 | 9611500 | 0 | 16271000 | 1 | 1 | 5,4 | 5,4 | 5,4 | 26,475 | 0,0099206 | 6,0098 | 40825000 | 1 | AJQ89965.1_229 | RM25_0233 | CytoplasmicMembrane | 1 | SPI | 0.242407 | OTHER | |
| lcl\|CP010341.1_prot_AJQ89967.1_231 | RM25_RS01155 | Arginine--tRNA ligase | 44580000 | 44268000 | 127200000 | 40100000 | 44548000 | 83909000 | 7 | 7 | 21,3 | 21,3 | 21,3 | 58,38 | 0 | 69,123 | 405990000 | 16 | AJQ89967.1_231 | RM25_0235 | Cytoplasmic | 0 | CYT | 0.088614 | OTHER | J |
| lcl\|CP010341.1_prot_AJQ89975.1_239 | RM25_RS01195 | Putative iron-siderophore transport system, periplasmi | 16621000 | 10185000 | 65565000 | 87188000 | 87018000 | 192120000 | 2 | 2 | 13 | 13 | 13 | 34,454 | 0 | 44,363 | 498810000 | 14 | AJQ89975.1_239 | RM25_0243 | Unknown | 0 | SPII | 0.816934 | LIPO(Sec/SPII) | P |
| lcl\|CP010341.1_prot_AJQ89978.1_242 | RM25_RS01210 | UDP-glucose 4-epimerase, galE | 89296000 | 129020000 | 220620000 | 122230000 | 133890000 | 162370000 | 5 | 5 | 19,8 | 19,8 | 19,8 | 35,243 | 0 | 49,496 | 938290000 | 23 | AJQ89978.1_242 | RM25_0246 | Cytoplasmic | 0 | CYT | 0.144302 | OTHER | M |

| | | | | | | | | | | | | | | | | | | | | | | | | | |
|---|---|---|---|---|---|---|---|---|---|---|---|---|---|---|---|---|---|---|---|---|---|---|---|---|---|---|
| lcl\|CP010341.1_prot_AJQ89980.1_244 | RM25_RS01220 | Thermostable beta-glucosidase B | 2454800 | 3048900 | 6807600 | 0 | 0 | 0 | 1 | 1 | 1,8 | 1,8 | 1,8 | 85,968 | 0,0043103 | 7,5665 | 18319000 | 2 | AJQ89980.1_244 | RM25_0248 | Cytoplasmic | 0 | CYT | 0.063195 | OTHER | |
| lcl\|CP010341.1_prot_AJQ89982.1_246 | RM25_RS01230 | Putative metallophosphoesterase | 74810000 | 121960000 | 158430000 | 71615000 | 132530000 | 18794000 | 6 | 6 | 20,5 | 20,5 | 20,5 | 60,434 | 0 | 50,564 | 624970000 | 8 | AJQ89982.1_246 | RM25_0250 | Cytoplasmic | 0 | CYT | 0.555971 | OTHER | M |
| lcl\|CP010341.1_prot_AJQ89984.1_248 | RM25_RS01240 | Glycerol-3-phosphate transporter | 16421000 | 10610000 | 35121000 | 35940000 | 42636000 | 80853000 | 1 | 1 | 3,6 | 3,6 | 3,6 | 51,39 | 0 | 12,05 | 265350000 | 8 | AJQ89984.1_248 | RM25_0252 | CytoplasmicMembrane | 10 | CYT | 0.956760 | OTHER | G |
| lcl\|CP010341.1_prot_AJQ89995.1_259 | RM25_RS01290 | Hypothetical protein | 129620000 | 79935000 | 119280000 | 16534000 | 13257000 | 31998000 | 5 | 5 | 65,3 | 65,3 | 65,3 | 7,4602 | 0 | 156,69 | 398480000 | 25 | AJQ89995.1_259 | RM25_0263 | Unknown | 0 | CYT | 0.754004 | OTHER | |
| lcl\|CP010341.1_prot_AJQ89998.1_262 | RM25_RS06075 | Ferritin-like protein | 101090000 | 0 | 435900000 | 193140000 | 156370000 | 203220000 | 4 | 4 | 40,6 | 40,6 | 40,6 | 19,151 | 0 | 82,623 | 1166100000 | 27 | AJQ89998.1_262 | RM25_1231 | CytoplasmicMembrane | 0 | CYT | 0.082400 | OTHER | I |
| lcl\|CP010341.1_prot_AJQ90001.1_265 | RM25_RS06075 | Hypothetical protein | 49725000 | 47957000 | 119810000 | 78492000 | 50394000 | 108200000 | 5 | 5 | 11,4 | 11,4 | 11,4 | 56,088 | 0 | 36,144 | 486670000 | 14 | AJQ90001.1_265 | RM25_1231 | CytoplasmicMembrane | 0 | CYT | 0.082400 | OTHER | I |
| lcl\|CP010341.1_prot_AJQ90006.1_270 | RM25_RS11695 | Hypothetical protein | 82715000 | 142390000 | 173640000 | 48778000 | 80957000 | 157620000 | 1 | 1 | 4 | 4 | 4 | 22,803 | 0 | 14,212 | 708320000 | 7 | AJQ90006.1_270 | RM25_0274 | CytoplasmicMembrane | 3 | TMH | 0.959732 | OTHER | |
| lcl\|CP010341.1_prot_AJQ90009.1_273 | RM25_RS01360 | HAD-superfamily hydrolase, subfamily IA, variant 1 | 49813000 | 31646000 | 128240000 | 118120000 | 84480000 | 101280000 | 3 | 3 | 24 | 24 | 24 | 23,184 | 0 | 37,872 | 591200000 | 15 | AJQ90009.1_273 | RM25_0277 | Cytoplasmic | 0 | CYT | 0.053344 | OTHER | |
| lcl\|CP010341.1_prot_AJQ90012.1_276 | RM25_RS01375 | Succinate CoA transferase | 227620000 | 153340000 | 813430016 | 280560000 | 222250000 | 383310016 | 12 | 12 | 27 | 27 | 27 | 55,685 | 0 | 124,14 | 2218300000 | 52 | AJQ90012.1_276 | RM25_0280 | Cytoplasmic | 0 | CYT | 0.154785 | OTHER | C |
| lcl\|CP010341.1_prot_AJQ90013.1_277 | RM25_RS01380 | Uncharacterized conserved protein UCP033563 | 652979968 | 915459968 | 1697200000 | 1053299968 | 1057900032 | 1160600064 | 17 | 17 | 65,4 | 65,4 | 65,4 | 43,402 | 0 | 323,31 | 7574400000 | 128 | AJQ90013.1_277 | RM25_0281 | Cytoplasmic | 0 | CYT | 0.088315 | OTHER | S |
| lcl\|CP010341.1_prot_AJQ90018.1_282 | RM25_RS01400 | 2,5-didehydrogluconate reductase | 189130000 | 197720000 | 573200000 | 240340000 | 251500000 | 307440000 | 10 | 10 | 45,8 | 45,8 | 45,8 | 30,542 | 0 | 153,12 | 1964100000 | 46 | AJQ90018.1_282 | RM25_0286 | Cytoplasmic | 0 | CYT | 0.108224 | OTHER | S |
| lcl\|CP010341.1_prot_AJQ90025.1_289 | RM25_RS01435 | Pyruvate, phosphate dikinase | 198720000 | 126620000 | 744529984 | 367680000 | 228230000 | 373209984 | 21 | 21 | 33,2 | 33,2 | 33,2 | 95,879 | 0 | 270,61 | 2224500000 | 68 | AJQ90025.1_289 | RM25_0293 | Cytoplasmic | 0 | CYT | 0.085358 | OTHER | G |
| lcl\|CP010341.1_prot_AJQ90026.1_290 | RM25_RS01445 | Hypothetical protein | 17521000 | 31652000 | 65873000 | 20231000 | 27542000 | 49721000 | 6 | 6 | 9,5 | 9,5 | 9,5 | 92,256 | 0 | 74,761 | 235060000 | 19 | AJQ90026.1_290 | RM25_0294 | Cytoplasmic | 0 | CYT | 0.736797 | OTHER | |
| lcl\|CP010341.1_prot_AJQ90028.1_292 | RM25_RS01455 | Two component system sensor kinase phoR | 49254000 | 127500000 | 123240000 | 31400000 | 67121000 | 87222000 | 3 | 3 | 5,6 | 5,6 | 5,6 | 64,615 | 0 | 24,457 | 515960000 | 13 | AJQ90028.1_292 | RM25_0296 | CytoplasmicMembrane | 2 | CYT | 0.536262 | OTHER | T |
| lcl\|CP010341.1_prot_AJQ90029.1_293 | RM25_RS01460 | Two component system transcriptional regulator trcR | 14124000 | 21497000 | 64460000 | 21285000 | 8468900 | 19824000 | 3 | 3 | 21,1 | 21,1 | 21,1 | 26,904 | 0 | 32,335 | 168500000 | 10 | AJQ90029.1_293 | RM25_0297 | Cytoplasmic | 0 | CYT | 0.076490 | OTHER | T |
| lcl\|CP010341.1_prot_AJQ90030.1_294 | RM25_RS01465 | NLPA lipoprotein | 66368000 | 53790000 | 123190000 | 59013000 | 93594000 | 168990000 | 7 | 7 | 41,1 | 41,1 | 41,1 | 34,067 | 0 | 88,3 | 630880000 | 24 | AJQ90030.1_294 | RM25_0298 | CytoplasmicMembrane | 1 | CYT | 0.957119 | OTHER | P |
| lcl\|CP010341.1_prot_AJQ90031.1_295 | RM25_RS01470 | Methionine import ATP-binding protein MetN 2 | 513490 | 1830800 | 2852800 | 0 | 861930 | 7681500 | 2 | 2 | 6,4 | 6,4 | 6,4 | 44,98 | 0 | 13,669 | 13741000 | 2 | AJQ90031.1_295 | RM25_0299 | CytoplasmicMembrane | 0 | CYT | 0.174628 | OTHER | P |
| lcl\|CP010341.1_prot_AJQ90034.1_298 | RM25_RS01485 | Hypothetical protein | 321569984 | 476609984 | 564659968 | 292169984 | 365310016 | 586209984 | 6 | 6 | 28,2 | 28,2 | 28,2 | 23,081 | 0 | 79,932 | 2831000000 | 39 | AJQ90034.1_298 | RM25_0302 | Unknown | 1 | CYT | 0.951850 | SP(Sec/SPI) | |
| lcl\|CP010341.1_prot_AJQ90043.1_307 | RM25_RS01530 | Hypothetical protein | 69232000 | 84383000 | 112590000 | 108880000 | 108470000 | 139720000 | 5 | 5 | 51,1 | 51,1 | 51,1 | 19,518 | 0 | 59,362 | 738730000 | 18 | AJQ90043.1_307 | RM25_0311 | CytoplasmicMembrane | 0 | CYT | 0.187375 | OTHER | |
| lcl\|CP010341.1_prot_AJQ90059.1_323 | RM25_RS01600 | Hypothetical protein | 20355000 | 17081000 | 53289000 | 15734000 | 25744000 | 38985000 | 3 | 3 | 21,9 | 21,9 | 21,9 | 14,975 | 0 | 36,841 | 191960000 | 12 | AJQ90059.1_323 | RM25_0327 | Unknown | 0 | CYT | 0.292168 | OTHER | |
| lcl\|CP010341.1_prot_AJQ90067.1_331 | RM25_RS01640 | PF07510 family protein | 294710016 | 502369984 | 707480000 | 220270000 | 510360000 | 586590016 | 7 | 7 | 37,3 | 37,3 | 37,3 | 26,27 | 0 | 323,31 | 2969800000 | 37 | AJQ90067.1_331 | RM25_0335 | Unknown | 0 | SPII | 0.878305 | LIPO(Sec/SPII) | L |
| lcl\|CP010341.1_prot_AJQ90071.1_335 | RM25_RS01655 | Hypothetical protein | 239720000 | 423569984 | 552220032 | 173090000 | 205870000 | 345870016 | 3 | 3 | 22,2 | 22,2 | 22,2 | 18,03 | 0 | 75,298 | 2090300000 | 37 | AJQ90071.1_335 | RM25_0339 | Unknown | 1 | TMH | 0.932836 | OTHER | |
| lcl\|CP010341.1_prot_AJQ90072.1_336 | RM25_RS01660 | Hypothetical protein | 50985000 | 75655000 | 128830000 | 58477000 | 61599000 | 43990000 | 3 | 3 | 31 | 31 | 31 | 18,363 | 0 | 71,528 | 468780000 | 19 | AJQ90072.1_336 | RM25_0340 | Unknown | 0 | CYT | 0.437868 | OTHER | |
| lcl\|CP010341.1_prot_AJQ90081.1_345 | RM25_RS01685 | CRISPR-associated helicase Cas3, Anaes-subtype | 19929000 | 5786600 | 44014000 | 10878000 | 15536000 | 37934000 | 6 | 6 | 10,1 | 10,1 | 10,1 | 93,073 | 0 | 37,589 | 146870000 | 10 | AJQ90081.1_345 | RM25_0349 | Unknown | 0 | CYT | 0.103749 | OTHER | |
| lcl\|CP010341.1_prot_AJQ90083.1_347 | RM25_RS01695 | Putative CRISPR-associated Csb1 family protein | 125770000 | 84583000 | 272870016 | 107020000 | 96258000 | 183370000 | 7 | 7 | 29,3 | 29,3 | 29,3 | 42,279 | 0 | 145,65 | 997690000 | 32 | AJQ90083.1_347 | RM25_0351 | Cytoplasmic | 0 | CYT | 0.086134 | OTHER | |
| lcl\|CP010341.1_prot_AJQ90088.1_352 | RM25_RS01725 | Copper-containing nitrite reductase (Major outer meml | 26170000 | 24384000 | 37649000 | 11952000 | 15581000 | 15750000 | 4 | 4 | 6 | 6 | 6 | 90,528 | 0 | 28,286 | 138440000 | 8 | AJQ90088.1_352 | RM25_0356 | CytoplasmicMembrane | 13 | TMH | 0.906582 | OTHER | |
| lcl\|CP010341.1_prot_AJQ90089.1_353 | RM25_RS01730 | TQXA domain-containing protein | 55550000 | 57764000 | 90513000 | 9136400 | 15216000 | 45483000 | 5 | 5 | 12,4 | 12,4 | 12,4 | 47,513 | 0 | 32,787 | 303320000 | 13 | AJQ90089.1_353 | RM25_0357 | Unknown | 2 | SPI | 0.910848 | SP(Sec/SPI) | |
| lcl\|CP010341.1_prot_AJQ90091.1_355 | RM25_RS12205 | Hypothetical protein | 27569000 | 30153000 | 60477000 | 29140000 | 46699000 | 78462000 | 9 | 9 | 12,3 | 12,3 | 12,3 | 116,86 | 0 | 98,352 | 291220000 | 25 | AJQ90091.1_355 | RM25_0359 | Cytoplasmic | 0 | CYT | 0.857751 | OTHER | L |
| lcl\|CP010341.1_prot_AJQ90092.1_356 | RM25_RS01745 | Hypothetical protein | 9592400 | 14399000 | 27309000 | 0 | 12811000 | 12254000 | 1 | 1 | 9,8 | 9,8 | 9,8 | 10,473 | 0,0042017 | 7,1589 | 76366000 | 6 | AJQ90092.1_356 | RM25_0360 | Unknown | 0 | CYT | 0.035859 | OTHER | L |
| lcl\|CP010341.1_prot_AJQ90095.1_359 | RM25_RS01760 | Isoleucine--tRNA ligase | 8875500 | 8853900 | 20725000 | 0 | 2986800 | 8593000 | 2 | 2 | 2,1 | 2,1 | 2,1 | 122,02 | 0 | 15,605 | 50034000 | 4 | AJQ90095.1_359 | RM25_0363 | Cytoplasmic | 0 | CYT | 0.203080 | OTHER | J |
| lcl\|CP010341.1_prot_AJQ90097.1_361 | RM25_RS01770 | Aspartate kinase | 126560000 | 169900000 | 394980000 | 230360000 | 190140000 | 292449984 | 10 | 10 | 30,7 | 30,7 | 30,7 | 44,924 | 0 | 80,486 | 1523600000 | 30 | AJQ90097.1_361 | RM25_0365 | Unknown | 0 | CYT | 0.047905 | OTHER | E |
| lcl\|CP010341.1_prot_AJQ90098.1_362 | RM25_RS01775 | Integral membrane protein MviN | 0 | 0 | 0 | 3260800 | 3493200 | 4677800 | 1 | 1 | 2,8 | 2,8 | 2,8 | 68,944 | 0,0045198 | 9,5313 | 11432000 | 2 | AJQ90098.1_362 | RM25_0366 | CytoplasmicMembrane | 14 | CYT | 0.962179 | OTHER | KLT |
| lcl\|CP010341.1_prot_AJQ90105.1_369 | RM25_RS01810 | Hypothetical protein | 44246000 | 56150000 | 95239000 | 24422000 | 30400000 | 44606000 | 3 | 3 | 13,4 | 13,4 | 13,4 | 26,104 | 0 | 21,07 | 310860000 | 6 | AJQ90105.1_369 | RM25_0373 | Unknown | 0 | SPII | 0.760425 | LIPO(Sec/SPII) | |
| lcl\|CP010341.1_prot_AJQ90108.1_372 | RM25_RS01825 | Phosphoribosylamine--glycine ligase | 31744000 | 33279000 | 51113000 | 17622000 | 30685000 | 40318000 | 7 | 7 | 17,5 | 17,5 | 17,5 | 42,045 | 0 | 46,303 | 217810000 | 15 | AJQ90108.1_372 | RM25_0376 | Cytoplasmic | 0 | CYT | 0.056303 | OTHER | F |
| lcl\|CP010341.1_prot_AJQ90109.1_373 | RM25_RS01830 | Adenylosuccinate lyase | 19914000 | 3012600 | 112970000 | 20364000 | 13899000 | 16836000 | 5 | 5 | 12,4 | 12,4 | 12,4 | 51,911 | 0 | 85,461 | 195590000 | 6 | AJQ90109.1_373 | RM25_0377 | Cytoplasmic | 0 | CYT | 0.040084 | OTHER | F |
| lcl\|CP010341.1_prot_AJQ90111.1_375 | RM25_RS01840 | Phosphoribosylaminoimidazolesuccinocarboxamide syr | 66197000 | 133140000 | 243250000 | 176590000 | 159290000 | 221430000 | 8 | 8 | 33,2 | 33,2 | 33,2 | 32,298 | 0 | 54,882 | 1161700000 | 27 | AJQ90111.1_375 | RM25_0379 | Cytoplasmic | 0 | CYT | 0.186959 | OTHER | F |
| lcl\|CP010341.1_prot_AJQ90113.1_377 | RM25_RS01850 | Phosphoribosylformylglycinamidine synthase 1 | 5449400 | 5461700 | 19036000 | 5883500 | 9372900 | 10054000 | 1 | 1 | 7,6 | 7,6 | 7,6 | 24,616 | 0,0044893 | 9,1213 | 55258000 | 2 | AJQ90113.1_377 | RM25_0381 | Cytoplasmic | 0 | CYT | 0.073373 | OTHER | F |
| lcl\|CP010341.1_prot_AJQ90114.1_378 | RM25_RS01855 | Hypothetical protein | 97689000 | 177970000 | 212530000 | 48447000 | 135300000 | 216940000 | 1 | 1 | 6,5 | 6,5 | 6,5 | 23,814 | 0 | 43,073 | 899470000 | 10 | AJQ90114.1_378 | RM25_0382 | Unknown | 0 | CYT | 0.918507 | OTHER | |
| lcl\|CP010341.1_prot_AJQ90115.1_379 | RM25_RS01860 | Prophage Lp2 protein 6 | 85297000 | 55860000 | 174030000 | 97893000 | 39219000 | 211440000 | 12 | 12 | 43,6 | 43,6 | 43,6 | 41,431 | 0 | 235,93 | 734570000 | 41 | AJQ90115.1_379 | RM25_0383 | Cytoplasmic | 0 | CYT | 0.138916 | OTHER | |
| lcl\|CP010341.1_prot_AJQ90123.1_387 | RM25_RS11725 | Cutinase | 9818700 | 16676000 | 1608200 | 0 | 8323800 | 16358000 | 3 | 3 | 11,2 | 11,2 | 11,2 | 42,979 | 0 | 19,119 | 52785000 | 2 | AJQ90123.1_387 | RM25_0391 | CytoplasmicMembrane | 2 | SPI | 0.540049 | SP(Sec/SPI) | |
| lcl\|CP010341.1_prot_AJQ90124.1_388 | RM25_RS01910 | Putative kinase inhibitor protein | 129700000 | 199310000 | 323100000 | 170630000 | 142750000 | 177500000 | 5 | 5 | 40,9 | 40,9 | 40,9 | 16,532 | 0 | 104,38 | 1337900000 | 21 | AJQ90124.1_388 | RM25_0392 | Cytoplasmic | 0 | CYT | 0.261538 | OTHER | |
| lcl\|CP010341.1_prot_AJQ90125.1_389 | RM25_RS01915 | DEAD/DEAH box helicase | 3637800 | 5760600 | 11469000 | 5068200 | 5812300 | 9760200 | 2 | 2 | 3,6 | 3,6 | 3,6 | 93,806 | 0 | 12,757 | 41509000 | 3 | AJQ90125.1_389 | RM25_0393 | Cytoplasmic | 0 | CYT | 0.073018 | OTHER | L |
| lcl\|CP010341.1_prot_AJQ90127.1_391 | RM25_RS11735 | Hypothetical protein | 415209984 | 629120000 | 1020400000 | 844870016 | 815600000 | 1353900032 | 4 | 4 | 21,6 | 21,6 | 21,6 | 33,804 | 0 | 191,5 | 5916600000 | 32 | AJQ90127.1_391 | RM25_0395 | Extracellular | 0 | CYT | 0.239684 | OTHER | M |
| lcl\|CP010341.1_prot_AJQ90128.1_392 | RM25_RS01930 | Glucose-6-phosphate isomerase | 54020000 | 19450000 | 113050000 | 12355000 | 22989000 | 41274000 | 6 | 6 | 13,2 | 13,2 | 13,2 | 61,894 | 0 | 54,362 | 273000000 | 6 | AJQ90128.1_392 | RM25_0396 | Cytoplasmic | 0 | CYT | 0.086973 | OTHER | G |
| lcl\|CP010341.1_prot_AJQ90131.1_395 | RM25_RS01945 | putative D-lactate dehydrogenase Fe-S protein FAD/FM | 181330000 | 145610000 | 291230016 | 120860000 | 231400000 | 390990016 | 17 | 17 | 21,1 | 21,1 | 21,1 | 100,57 | 0 | 204,51 | 1457500000 | 62 | AJQ90131.1_395 | RM25_0399 | Cytoplasmic | 0 | CYT | 0.090568 | OTHER | C |
| lcl\|CP010341.1_prot_AJQ90133.1_397 | RM25_RS01955 | Carboxylic ester hydrolase | 41748000 | 53440000 | 59105000 | 22512000 | 30787000 | 35883000 | 4 | 4 | 13,3 | 13,3 | 13,3 | 46,469 | 0 | 30,176 | 267650000 | 11 | AJQ90133.1_397 | RM25_0401 | Unknown | 0 | SPI | 0.779888 | SP(Sec/SPI) | |
| lcl\|CP010341.1_prot_AJQ90144.1_408 | RM25_RS02010 | Hypothetical protein | 0 | 3936700 | 7409000 | 7980600 | 3324100 | 7639700 | 1 | 1 | 10,5 | 10,5 | 10,5 | 13,733 | 0,0042418 | 7,3781 | 37800000 | 2 | AJQ90144.1_408 | RM25_0412 | Unknown | 0 | CYT | 0.124423 | OTHER | |
| lcl\|CP010341.1_prot_AJQ90149.1_413 | RM25_RS02035 | Azoreductase B | 1793900 | 2553300 | 9759000 | 10924000 | 3822400 | 0 | 2 | 2 | 16,3 | 16,3 | 16,3 | 21,651 | 0 | 14,583 | 28852000 | 1 | AJQ90149.1_413 | RM25_0417 | Cytoplasmic | 0 | CYT | 0.106123 | OTHER | S |
| lcl\|CP010341.1_prot_AJQ90153.1_417 | RM25_RS02055 | Multifunctional siroheme synthase cysG | 59155000 | 65290000 | 166700000 | 63432000 | 99375000 | 130000000 | 6 | 6 | 42,8 | 42,8 | 42,8 | 27,095 | 0 | 58,271 | 633970000 | 26 | AJQ90153.1_417 | RM25_0421 | Unknown | 0 | CYT | 0.055335 | OTHER | H |
| lcl\|CP010341.1_prot_AJQ90154.1_418 | RM25_RS02060 | Cobalt ABC transporter, ATPase subunit | 1818700 | 1999000 | 10490000 | 0 | 0 | 0 | 1 | 1 | 6,1 | 6,1 | 6,1 | 31,244 | 0,005071 | 6,3615 | 14308000 | 1 | AJQ90154.1_418 | RM25_0422 | CytoplasmicMembrane | 0 | CYT | 0.067773 | OTHER | P |
| lcl\|CP010341.1_prot_AJQ90156.1_420 | RM25_RS02070 | Cobalt transport protein CbiN | 6096900 | 6822500 | 19382000 | 15134000 | 13598000 | 28224000 | 1 | 1 | 22,9 | 22,9 | 22,9 | 16,469 | 0 | 49,439 | 100610000 | 1 | AJQ90156.1_420 | RM25_0424 | CytoplasmicMembrane | 2 | CYT | 0.933248 | OTHER | |
| lcl\|CP010341.1_prot_AJQ90157.1_421 | RM25_RS02075 | Cobalt transport protein CbiM | 51085000 | 49914000 | 154810000 | 121940000 | 96660000 | 185360000 | 1 | 1 | 5,1 | 5,1 | 5,1 | 24,534 | 0 | 24,886 | 731120000 | 3 | AJQ90157.1_421 | RM25_0425 | CytoplasmicMembrane | 6 | CYT | 0.974267 | OTHER | |
| lcl\|CP010341.1_prot_AJQ90162.1_426 | RM25_RS02095 | Chaperone protein DnaK | 337640000 | 254010000 | 940790016 | 457580000 | 558009984 | 735320000 | 15 | 14 | 33,8 | 33,8 | 31,2 | 67,228 | 0 | 188,17 | 3613400000 | 79 | AJQ90162.1_426 | RM25_0430 | Cytoplasmic | 0 | CYT | 0.567972 | OTHER | O |
| lcl\|CP010341.1_prot_AJQ90163.1_427 | RM25_RS02100 | Protein GrpE | 25382000 | 19348000 | 60809000 | 24568000 | 35903000 | 69690000 | 2 | 2 | 16,2 | 16,2 | 16,2 | 22,908 | 0 | 33,492 | 250200000 | 9 | AJQ90163.1_427 | RM25_0431 | Cytoplasmic | 0 | CYT | 0.130924 | OTHER | O |
| lcl\|CP010341.1_prot_AJQ90164.1_428 | RM25_RS02105 | Chaperone protein DnaJ | 17812000 | 12842000 | 44765000 | 45594000 | 38503000 | 59939000 | 3 | 3 | 13,6 | 13,6 | 13,6 | 40,755 | 0 | 24,216 | 252770000 | 10 | AJQ90164.1_428 | RM25_0432 | Cytoplasmic | 0 | CYT | 0.757186 | OTHER | O |
| lcl\|CP010341.1_prot_AJQ90167.1_431 | RM25_RS02120 | Phosphoribosylformylglycinamidine synthase 2 | 26808000 | 30559000 | 86634000 | 39976000 | 32922000 | 87428000 | 4 | 4 | 7,4 | 7,4 | 7,4 | 90,473 | 0 | 54,526 | 337990000 | 17 | AJQ90167.1_431 | RM25_0435 | Cytoplasmic | 0 | CYT | 0.103520 | OTHER | F |
| lcl\|CP010341.1_prot_AJQ90168.1_432 | RM25_RS02125 | Peptidase dimerization domain protein | 97158000 | 154460000 | 284340000 | 159890000 | 145780000 | 173910000 | 11 | 11 | 39,1 | 39,1 | 39,1 | 47,588 | 0 | 203,01 | 1109400000 | 36 | AJQ90168.1_432 | RM25_0436 | Cytoplasmic | 0 | CYT | 0.072527 | OTHER | E |
| lcl\|CP010341.1_prot_AJQ90170.1_434 | RM25_RS02135 | Formate--tetrahydrofolate ligase | 18394000 | 14422000 | 51239000 | 7956200 | 18102000 | 55840000 | 5 | 5 | 10 | 10 | 10 | 59,184 | 0 | 30,71 | 169560000 | 10 | AJQ90170.1_434 | RM25_0438 | Cytoplasmic | 0 | CYT | 0.054872 | OTHER | F |
| lcl\|CP010341.1_prot_AJQ90173.1_437 | RM25_RS02150 | UPF0237 protein | 18233000 | 15630000 | 48754000 | 40750000 | 28795000 | 32773000 | 3 | 3 | 38,2 | 38,2 | 38,2 | 9,8172 | 0 | 23,188 | 201890000 | 12 | AJQ90173.1_437 | RM25_0441 | Cytoplasmic | 0 | CYT | 0.023025 | OTHER | |
| lcl\|CP010341.1_prot_AJQ90174.1_438 | RM25_RS11760 | Hypothetical protein | 2303000 | 0 | 6847100 | 0 | 0 | 6539300 | 1 | 1 | 5,7 | 5,7 | 5,7 | 28,208 | 0,0041451 | 6,847 | 15689000 | 2 | AJQ90174.1_438 | RM25_0442 | CytoplasmicMembrane | 2 | CYT | 0.966684 | OTHER | |
| lcl\|CP010341.1_prot_AJQ90175.1_439 | RM25_RS02160 | Amidophosphoribosyltransferase | 5348100 | 3906200 | 17413000 | 2934900 | 2627300 | 8894600 | 2 | 2 | 3,9 | 3,9 | 3,9 | 55,334 | 0 | 12,119 | 44888000 | 4 | AJQ90175.1_439 | RM25_0443 | Cytoplasmic | 0 | CYT | 0.090594 | OTHER | F |
| lcl\|CP010341.1_prot_AJQ90176.1_440 | RM25_RS02165 | Phosphoribosylformylglycinamidine cyclo-ligase | 16167000 | 21294000 | 38307000 | 4983800 | 24056000 | 28185000 | 4 | 4 | 19,5 | 19,5 | 19,5 | 36,474 | 0 | 32,997 | 154360000 | 8 | AJQ90176.1_440 | RM25_0444 | Cytoplasmic | 0 | CYT | 0.053108 | OTHER | F |
| lcl\|CP010341.1_prot_AJQ90177.1_441 | RM25_RS02170 | Hypothetical protein | 40031000 | 55838000 | 68899000 | 60977000 | 49314000 | 85698000 | 3 | 3 | 69,7 | 69,7 | 69,7 | 8,5521 | 0 | 31,954 | 410580000 | 9 | AJQ90177.1_441 | RM25_0445 | Cytoplasmic | 0 | CYT | 0.945452 | OTHER | |
| lcl\|CP010341.1_prot_AJQ90179.1_443 | RM25_RS02195 | Cell-wall peptidases, NlpC/P60 family secreted protein | 11868999680 | 12636000256 | 22329999360 | 8949099520 | 10940999680 | 15692999680 | 22 | 22 | 61,2 | 61,2 | 61,2 | 58,794 | 0 | 323,31 | 9,055E+10 | 250 | AJQ90179.1_443 | RM25_0447 | Extracellular | 0 | SPI | 0.884174 | SP(Sec/SPI) | M |
| lcl\|CP010341.1_prot_AJQ90189.1_453 | RM25_RS02245 | Dehydrogenase | 22978000 | 28376000 | 70915000 | 41107000 | 5701900 | 19539000 | 4 | 4 | 12,3 | 12,3 | 12,3 | 36,254 | 0 | 30,72 | 204030000 | 10 | AJQ90189.1_453 | RM25_0458 | Cytoplasmic | 0 | CYT | 0.045928 | OTHER | C |
| lcl\|CP010341.1_prot_AJQ90190.1_454 | RM25_RS02250 | Hypothetical protein | 16310000 | 27128000 | 21074000 | 0 | 38763000 | 43545000 | 2 | 2 | 23,1 | 23,1 | 23,1 | 12,14 | 0 | 14,466 | 156930000 | 4 | AJQ90190.1_454 | RM25_0459 | Unknown | 0 | SPI | 0.127638 | OTHER | |
| lcl\|CP010341.1_prot_AJQ90191.1_455 | RM25_RS02255 | Transaldolase | 397710016 | 510809984 | 1009000000 | 561729984 | 618920000 | 623539968 | 12 | 12 | 43,6 | 43,6 | 43,6 | 38,022 | 0 | 195,66 | 3975900000 | 65 | AJQ90191.1_455 | RM25_0460 | Cytoplasmic | 0 | CYT | 0.053721 | OTHER | G |
| lcl\|CP010341.1_prot_AJQ90192.1_456 | RM25_RS02260 | Hypothetical protein | 24845000 | 19969000 | 46472000 | 23955000 | 26089000 | 70279000 | 4 | 4 | 27,4 | 27,4 | 27,4 | 23,801 | 0 | 31,45 | 234140000 | 14 | AJQ90192.1_456 | RM25_0461 | Unknown | 1 | SPI | 0.645929 | SP(Sec/SPI) | |
| lcl\|CP010341.1_prot_AJQ90197.1_461 | RM25_RS02285 | Hypothetical protein | 74990000 | 89250000 | 103420000 | 137090000 | 130450000 | 164620000 | 5 | 5 | 53,8 | 53,8 | 53,8 | 14,192 | 0 | 225,88 | 828930000 | 21 | AJQ90197.1_461 | RM25_0466 | Unknown | 0 | CYT | 0.920135 | OTHER | |
| lcl\|CP010341.1_prot_AJQ90203.1_467 | RM25_RS02315 | Heme ABC exporter, ATP-binding protein CcmA | 8644500 | 4554600 | 23516000 | 10116000 | 6888800 | 15933000 | 1 | 1 | 4,3 | 4,3 | 4,3 | 25,332 | 0,004329 | 7,6044 | 69653000 | 6 | AJQ90203.1_467 | RM25_0472 | CytoplasmicMembrane | 0 | CYT | 0.053484 | OTHER | V |
| lcl\|CP010341.1_prot_AJQ90204.1_468 | RM25_RS02320 | Hypothetical protein | 1531300 | 1954100 | 4322400 | 1664200 | 2795000 | 3193300 | 1 | 1 | 2 | 2 | 2 | 64,06 | 0,0070565 | 6,2678 | 15460000 | 5 | AJQ90204.1_468 | RM25_0473 | CytoplasmicMembrane | 12 | CYT | 0.953131 | OTHER | - |
| lcl\|CP010341.1_prot_AJQ90208.1_472 | RM25_RS02340 | NADH-quinone oxidoreductase subunit B | 51514000 | 41858000 | 105340000 | 72163000 | 47437000 | 171670000 | 2 | 2 | 11,4 | 11,4 | 11,4 | 20,311 | 0 | 30,603 | 536840000 | 13 | AJQ90208.1_472 | RM25_0477 | CytoplasmicMembrane | 0 | CYT | 0.071149 | OTHER | C |
| lcl\|CP010341.1_prot_AJQ90209.1_473 | RM25_RS02345 | NADH-quinone oxidoreductase subunit C | 160490000 | 158080000 | 441910016 | 229760000 | 374520000 | 456289984 | 10 | 10 | 52,2 | 52,2 | 52,2 | 28,677 | 0 | 146,21 | 1986100000 | 52 | AJQ90209.1_473 | RM25_0478 | Cytoplasmic | 0 | CYT | 0.144167 | OTHER | C |
| lcl\|CP010341.1_prot_AJQ90210.1_474 | RM25_RS02350 | NADH-quinone oxidoreductase subunit D | 58038000 | 49079000 | 135830000 | 90769000 | 91541000 | 245600000 | 7 | 7 | 21 | 21 | 21 | 50,313 | 0 | 79,511 | 727010000 | 26 | AJQ90210.1_474 | RM25_0479 | Cytoplasmic | 0 | CYT | 0.138191 | OTHER | C |
| lcl\|CP010341.1_prot_AJQ90211.1_475 | RM25_RS02355 | NADH dehydrogenase subunit E | 21700000 | 28265000 | 65271000 | 45323000 | 49543000 | 79300000 | 3 | 3 | 11,9 | 11,9 | 11,9 | 25,936 | 0 | 18,142 | 308670000 | 4 | AJQ90211.1_475 | RM25_0480 | Cytoplasmic | 0 | CYT | 0.098946 | OTHER | C |
| lcl\|CP010341.1_prot_AJQ90212.1_476 | RM25_RS02360 | NADH-quinone oxidoreductase, F subunit | 17902000 | 6686200 | 55845000 | 37191000 | 38574000 | 77509000 | 3 | 3 | 7,4 | 7,4 | 7,4 | 48,644 | 0 | 24,366 | 245000000 | 8 | AJQ90212.1_476 | RM25_0481 | Cytoplasmic | 0 | CYT | 0.174933 | OTHER | C |
| lcl\|CP010341.1_prot_AJQ90213.1_477 | RM25_RS02365 | NADH dehydrogenase, G subunit | 178670000 | 138530000 | 349140000 | 204310000 | 277000000 | 544460032 | 15 | 15 | 27,5 | 27,5 | 27,5 | 84,953 | 0 | 174,59 | 1824900000 | 68 | AJQ90213.1_477 | RM25_0482 | Cytoplasmic | 0 | CYT | 0.083286 | OTHER | C |
| lcl\|CP010341.1_prot_AJQ90214.1_478 | RM25_RS02370 | NADH-quinone oxidoreductase subunit H | 8867700 | 7161300 | 14756000 | 0 | 13814000 | 24194000 | 1 | 1 | 3,7 | 3,7 | 3,7 | 50,143 | 0,0023175 | 10,586 | 79410000 | 5 | AJQ90214.1_478 | RM25_0483 | CytoplasmicMembrane | 9 | TMH | 0.960024 | OTHER | C |
| lcl\|CP010341.1_prot_AJQ90215.1_479 | RM25_RS02375 | NADH-quinone oxidoreductase subunit I | 7679600 | 3320400 | 8433500 | 7908800 | 7024500 | 23601000 | 2 | 2 | 16,3 | 16,3 | 16,3 | 21,512 | 0 | 14,297 | 57968000 | 6 | AJQ90215.1_479 | RM25_0484 | Cytoplasmic | 0 | CYT | 0.263075 | OTHER | C |
| lcl\|CP010341.1_prot_AJQ90216.1_480 | RM25_RS02380 | NADH dehydrogenase subunit I J nuoJ | 25462000 | 17455000 | 59159000 | 43457000 | 53861000 | 90350000 | 3 | 3 | 21,5 | 21,5 | 21,5 | 30,687 | 0 | 51,353 | 324710000 | 11 | AJQ90216.1_480 | RM25_0485 | CytoplasmicMembrane | 5 | TMH | 0.940824 | OTHER | C |
| lcl\|CP010341.1_prot_AJQ90217.1_481 | RM25_RS02385 | NADH-quinone oxidoreductase subunit K | 4143100 | 0 | 6867500 | 5964400 | 6380300 | 15464000 | 1 | 1 | 11,1 | 11,1 | 11,1 | 10,665 | 0,0041754 | 6,957 | 38820000 | 2 | AJQ90217.1_481 | RM25_0486 | CytoplasmicMembrane | 3 | TMH | 0.968441 | OTHER | C |
| lcl\|CP010341.1_prot_AJQ90218.1_482 | RM25_RS02390 | NADH dehydrogenase subunit I L nuoL | 8863200 | 5099700 | 19470000 | 13548000 | 15674000 | 28059000 | 1 | 1 | 2,2 | 2,2 | 2,2 | 67,544 | 0,0043243 | 7,5898 | 101820000 | 5 | AJQ90218.1_482 | RM25_0487 | CytoplasmicMembrane | 14 | TMH | 0.953839 | OTHER | CP |
| lcl\|CP010341.1_prot_AJQ90219.1_483 | RM25_RS02395 | NADH dehydrogenase subunit M | 4754400 | 4677900 | 9954000 | 14237000 | 15784000 | 14564000 | 2 | 2 | 4,5 | 4,5 | 4,5 | 55,157 | 0 | 18,648 | 68189000 | 7 | AJQ90219.1_483 | RM25_0488 | CytoplasmicMembrane | 13 | TMH | 0.965649 | OTHER | C |
| lcl\|CP010341.1_prot_AJQ90221.1_485 | RM25_RS02405 | Putative POLYPRENYL-DIPHOSPHATE SYNTHASE GRCC1 | 15585000 | 62258000 | 102880000 | 11123000 | 45607000 | 86520000 | 5 | 5 | 24 | 24 | 24 | 34,523 | 0 | 35,253 | 356320000 | 8 | AJQ90221.1_485 | RM25_0490 | Cytoplasmic | 0 | CYT | 0.035045 | OTHER | H |
| lcl\|CP010341.1_prot_AJQ90224.1_488 | RM25_RS02420 | Peptidase family M13 | 90938000 | 101400000 | 268990016 | 121620000 | 144010000 | 194200000 | 12 | 12 | 25,3 | 25,3 | 25,3 | 73,523 | 0 | 145,27 | 1054000000 | 31 | AJQ90224.1_488 | RM25_0493 | Cytoplasmic | 0 | CYT | 0.216999 | OTHER | O |
| lcl\|CP010341.1_prot_AJQ90226.1_490 | RM25_RS02430 | 2-oxoacid:acceptor oxidoreductase, alpha subunit | 2524100 | 2875900 | 5202100 | 3122000 | 5352900 | 7740500 | 2 | 2 | 5,1 | 5,1 | 5,1 | 67,49 | 0 | 11,427 | 26817000 | 2 | AJQ90226.1_490 | RM25_0495 | Cytoplasmic | 0 | CYT | 0.103751 | OTHER | C |
| lcl\|CP010341.1_prot_AJQ90231.1_495 | RM25_RS02460 | Alkyl hydroperoxide reductase subunit C | 91090000 | 89229000 | 283110016 | 365150016 | 403660000 | 623320000 | 5 | 5 | 43,3 | 43,3 | 43,3 | 20,761 | 0 | 143,75 | 2080400000 | 27 | AJQ90231.1_495 | RM25_0501 | Cytoplasmic | 0 | CYT | 0.183246 | OTHER | |
| lcl\|CP010341.1_prot_AJQ90232.1_496 | RM25_RS02465 | Alkyl hydroperoxide reductase, F subunit | 7878800 | 6335500 | 16700000 | 22693000 | 33521000 | 48054000 | 5 | 5 | 14,7 | 14,7 | 14,7 | 58,196 | 0 | 47,793 | 147230000 | 8 | AJQ90232.1_496 | RM25_0502 | CytoplasmicMembrane | 0 | CYT | 0.064183 | OTHER | O |
| lcl\|CP010341.1_prot_AJQ90238.1_502 | RM25_RS12230 | Preprotein translocase, SecE subunit | 20334000 | 25693000 | 56263000 | 41344000 | 38803000 | 64122000 | 1 | 1 | 6,2 | 6,2 | 6,2 | 21,241 | 0 | 24,683 | 274090000 | 7 | AJQ90238.1_502 | RM25_0511 | Cytoplasmic | 1 | CYT | 0.903705 | OTHER | U |
| lcl\|CP010341.1_prot_AJQ90239.1_503 | RM25_RS02515 | Transcription antitermination protein nusG | 136330000 | 189120000 | 391470016 | 176660000 | 156280000 | 296100000 | 6 | 6 | 33,9 | 33,9 | 33,9 | 32,042 | 0 | 173,16 | 1502400000 | 37 | AJQ90239.1_503 | RM25_0512 | Cytoplasmic | 0 | CYT | 0.852783 | OTHER | K |
| lcl\|CP010341.1_prot_AJQ90240.1_504 | RM25_RS02520 | 50S ribosomal protein L11 | 333670016 | 268710016 | 1007900032 | 923840000 | 921430016 | 1337500032 | 5 | 5 | 38 | 38 | 38 | 14,871 | 0 | 86,225 | 5530400000 | 40 | AJQ90240.1_504 | RM25_0513 | Unknown | 0 | SPI | 0.140270 | SP(Sec/SPI) | J |
| lcl\|CP010341.1_prot_AJQ90241.1_505 | RM25_RS02525 | 50S ribosomal protein L1 | 674460032 | 710419968 | 1496899968 | 1089100032 | 1193500032 | 1820899968 | 11 | 11 | 64,4 | 64,4 | 64,4 | 25,411 | 0 | 323,31 | 7884600000 | 109 | AJQ90241.1_505 | RM25_0514 | Cytoplasmic | 0 | CYT | 0.154590 | OTHER | J |
| lcl\|CP010341.1_prot_AJQ90243.1_507 | RM25_RS02535 | Lipoyl synthase | 3032100 | 6423100 | 12862000 | 22117000 | 19107000 | 22433000 | 3 | 3 | 11,7 | 11,7 | 11,7 | 35,329 | 0 | 16,137 | 89378000 | 3 | AJQ90243.1_507 | RM25_0516 | Cytoplasmic | 0 | CYT | 0.085746 | OTHER | H |
| lcl\|CP010341.1_prot_AJQ90244.1_508 | RM25_RS02540 | 50S ribosomal protein L10 | 390910016 | 449830016 | 1074499968 | 1125900032 | 1115299968 | 1629400064 | 9 | 9 | 58,6 | 58,6 | 58,6 | 21,881 | 0 | 232,1 | 6765300000 | 57 | AJQ90244.1_508 | RM25_0517 | Cytoplasmic | 0 | CYT | 0.076266 | OTHER | J |
| lcl\|CP010341.1_prot_AJQ90245.1_509 | RM25_RS02545 | 50S ribosomal protein L7/L12 | 1662800000 | 1434200064 | 3893499904 | 2379399936 | 2976399872 | 5304699904 | 8 | 8 | 77,1 | 77,1 | 77,1 | 13,665 | 0 | 114,14 | 1,9751E+10 | 89 | AJQ90245.1_509 | RM25_0518 | Cytoplasmic | 0 | CYT | 0.041670 | OTHER | J |
| lcl\|CP010341.1_prot_AJQ90246.1_510 | RM25_RS02550 | Hypothetical protein | 102300000 | 162130000 | 187860000 | 52852000 | 89482000 | 166790000 | 2 | 2 | 8,7 | 8,7 | 8,7 | 22,183 | 0 | 18,02 | 811460000 | 11 | AJQ90246.1_510 | RM25_0519 | Unknown | 1 | TMH | 0.932468 | OTHER | |
| lcl\|CP010341.1_prot_AJQ90247.1_511 | RM25_RS02555 | DNA-directed RNA polymerase, beta subunit | 343420000 | 294870016 | 785980032 | 628700032 | 795550016 | 1562099968 | 34 | 34 | 38,9 | 38,9 | 38,9 | 128,72 | 0 | 323,31 | 4789200000 | 142 | AJQ90247.1_511 | RM25_0520 | Cytoplasmic | 0 | CYT | 0.136443 | OTHER | K |
| lcl\|CP010341.1_prot_AJQ90248.1_512 | RM25_RS02560 | DNA-directed RNA polymerase, beta' subunit | 350249984 | 277409984 | 903100032 | 726649984 | 883870016 | 1664099968 | 39 | 39 | 32,3 | 32,3 | 32,3 | 143,61 | 0 | 323,31 | 5253800000 | 165 | AJQ90248.1_512 | RM25_0521 | Cytoplasmic | 0 | CYT | 0.078299 | OTHER | K |
| lcl\|CP010341.1_prot_AJQ90249.1_513 | RM25_RS02565 | 30S ribosomal protein S12 | 103950000 | 90666000 | 278980000 | 68388000 | 91131000 | 96080000 | 2 | 2 | 13,8 | 13,8 | 13,8 | 13,611 | 0 | 67,025 | 771680000 | 11 | AJQ90249.1_513 | RM25_0522 | Cytoplasmic | 0 | CYT | 0.710030 | OTHER | J |

| | | | | | | | | | | | | | | | | | | | | | | | | | | |
|---|---|---|---|---|---|---|---|---|---|---|---|---|---|---|---|---|---|---|---|---|---|---|---|---|---|---|
| lcl\|CP010341.1_prot_AJQ90250.1_514 | RM25_RS02570 | 30S ribosomal protein S7 | 283900000 | 261280000 | 874070016 | 748849984 | 866499968 | 1386599936 | 10 | 10 | 63,5 | 63,5 | 63,5 | 17,455 | 0 | 203,19 | 5139300000 | 59 | AJQ90250.1_514 | RM25_0523 | Unknown | 0 | CYT | 0.091659 | OTHER | J |
| lcl\|CP010341.1_prot_AJQ90251.1_515 | RM25_RS02575 | Translation elongation factor G | 660369984 | 677969984 | 1566599936 | 399920000 | 699960000 | 1276899968 | 17 | 17 | 39,1 | 39,1 | 39,1 | 76,687 | 0 | 258,85 | 5726700000 | 75 | AJQ90251.1_515 | RM25_0524 | Cytoplasmic | 0 | CYT | 0.090831 | OTHER | J |
| lcl\|CP010341.1_prot_AJQ90252.1_516 | RM25_RS02580 | Translation elongation factor Tu | 3153600000 | 3075699968 | 8353800192 | 2716000000 | 3878799872 | 6039199744 | 15 | 15 | 55,8 | 55,8 | 55,8 | 43,679 | 0 | 323,31 | 2,9737E+10 | 227 | AJQ90252.1_516 | RM25_0525 | Cytoplasmic | 0 | CYT | 0.101860 | OTHER | J |
| lcl\|CP010341.1_prot_AJQ90255.1_519 | RM25_RS02595 | Glutamate--tRNA ligase | 31403000 | 22357000 | 70015000 | 39542000 | 40085000 | 75160000 | 5 | 5 | 11,6 | 11,6 | 11,6 | 52,44 | 0 | 35,59 | 304430000 | 15 | AJQ90255.1_519 | RM25_0528 | Cytoplasmic | 0 | CYT | 0.142332 | OTHER | J |
| lcl\|CP010341.1_prot_AJQ90257.1_521 | RM25_RS02605 | 30S ribosomal protein S10 | 589299968 | 445740000 | 1414200064 | 1078899968 | 1119500032 | 1777299968 | 6 | 6 | 57,3 | 57,3 | 57,3 | 11,744 | 0 | 49,602 | 7288700000 | 43 | AJQ90257.1_521 | RM25_0530 | Cytoplasmic | 0 | CYT | 0.068205 | OTHER | J |
| lcl\|CP010341.1_prot_AJQ90258.1_522 | RM25_RS02610 | 50S ribosomal protein L3 | 158180000 | 242240000 | 785910016 | 734470016 | 692129984 | 1142899968 | 6 | 6 | 38,8 | 38,8 | 38,8 | 22,967 | 0 | 323,31 | 4151100000 | 45 | AJQ90258.1_522 | RM25_0531 | Cytoplasmic | 0 | CYT | 0.345011 | OTHER | J |
| lcl\|CP010341.1_prot_AJQ90259.1_523 | RM25_RS02615 | 50S ribosomal protein L4 | 281240000 | 389640000 | 1136199936 | 843580032 | 814409984 | 1234300032 | 9 | 9 | 44,6 | 44,6 | 44,6 | 24,839 | 0 | 255,39 | 5405400000 | 78 | AJQ90259.1_523 | RM25_0532 | Cytoplasmic | 0 | CYT | 0.311203 | OTHER | J |
| lcl\|CP010341.1_prot_AJQ90260.1_524 | RM25_RS02620 | 50S ribosomal protein L23 | 274750016 | 306129984 | 795609984 | 483529984 | 403969984 | 766550016 | 6 | 6 | 60,2 | 60,2 | 60,2 | 11,526 | 0 | 80,517 | 3356600000 | 48 | AJQ90260.1_524 | RM25_0533 | Cytoplasmic | 0 | CYT | 0.118148 | OTHER | J |
| lcl\|CP010341.1_prot_AJQ90261.1_525 | RM25_RS02625 | 50S ribosomal protein L2 | 541120000 | 565670016 | 1623600000 | 1142200064 | 1231800064 | 1824899968 | 12 | 12 | 53,2 | 53,2 | 53,2 | 30,369 | 0 | 186,99 | 7555400000 | 103 | AJQ90261.1_525 | RM25_0534 | Cytoplasmic | 0 | CYT | 0.823561 | OTHER | J |
| lcl\|CP010341.1_prot_AJQ90262.1_526 | RM25_RS02630 | 30S ribosomal protein S19 | 50952000 | 153240000 | 370020000 | 332780000 | 439070016 | 606480000 | 6 | 6 | 47,3 | 47,3 | 47,3 | 10,612 | 0 | 70,916 | 2220100000 | 29 | AJQ90262.1_526 | RM25_0535 | Cytoplasmic | 0 | CYT | 0.203174 | OTHER | J |
| lcl\|CP010341.1_prot_AJQ90263.1_527 | RM25_RS02635 | 50S ribosomal protein L22 | 313790016 | 305009984 | 576630016 | 641939968 | 828760000 | 1159200000 | 5 | 5 | 34,8 | 34,8 | 34,8 | 17,268 | 0 | 84,978 | 4350000000 | 37 | AJQ90263.1_527 | RM25_0536 | Cytoplasmic | 0 | CYT | 0.090746 | OTHER | J |
| lcl\|CP010341.1_prot_AJQ90264.1_528 | RM25_RS02640 | 30S ribosomal protein S3 | 331920000 | 309320000 | 769529984 | 483180000 | 555760000 | 867560000 | 8 | 8 | 34,6 | 34,6 | 34,6 | 30,019 | 0 | 153,56 | 3648800000 | 47 | AJQ90264.1_528 | RM25_0537 | Cytoplasmic | 0 | CYT | 0.073050 | OTHER | J |
| lcl\|CP010341.1_prot_AJQ90265.1_529 | RM25_RS02645 | 50S ribosomal protein L16 | 222460000 | 167010000 | 508120000 | 295790016 | 332550016 | 508200000 | 4 | 4 | 46,8 | 46,8 | 46,8 | 15,732 | 0 | 61,346 | 2308000000 | 29 | AJQ90265.1_529 | RM25_0538 | Cytoplasmic | 0 | CYT | 0.079580 | OTHER | J |
| lcl\|CP010341.1_prot_AJQ90266.1_530 | RM25_RS02650 | 50S ribosomal protein L29 | 171890000 | 268330000 | 561779968 | 473480000 | 604320000 | 730110016 | 4 | 4 | 48,8 | 48,8 | 48,8 | 9,3074 | 0 | 96,847 | 3063100000 | 28 | AJQ90266.1_530 | RM25_0539 | Cytoplasmic | 0 | CYT | 0.051263 | OTHER | J |
| lcl\|CP010341.1_prot_AJQ90267.1_531 | RM25_RS02655 | 30S ribosomal protein S17 | 26614000 | 113720000 | 302729984 | 240890000 | 293520000 | 379000000 | 2 | 2 | 18,7 | 18,7 | 18,7 | 10,235 | 0 | 11,202 | 1568400000 | 6 | AJQ90267.1_531 | RM25_0540 | Cytoplasmic | 0 | CYT | 0.443741 | OTHER | J |
| lcl\|CP010341.1_prot_AJQ90268.1_532 | RM25_RS02660 | 50S ribosomal protein L14 | 124110000 | 131590000 | 283209984 | 224200000 | 254970000 | 372160000 | 3 | 3 | 26,8 | 26,8 | 26,8 | 13,344 | 0 | 36,727 | 1549600000 | 25 | AJQ90268.1_532 | RM25_0541 | Cytoplasmic | 0 | CYT | 0.039985 | OTHER | J |
| lcl\|CP010341.1_prot_AJQ90269.1_533 | RM25_RS02665 | 50S ribosomal protein L24 | 161920000 | 184150000 | 724200000 | 534849984 | 611240000 | 741609984 | 6 | 6 | 55,3 | 55,3 | 55,3 | 13,196 | 0 | 101,03 | 3113800000 | 40 | AJQ90269.1_533 | RM25_0542 | Cytoplasmic | 0 | CYT | 0.213444 | OTHER | J |
| lcl\|CP010341.1_prot_AJQ90270.1_534 | RM25_RS02670 | 50S ribosomal protein L5 | 308260000 | 300150016 | 800129984 | 633230016 | 655000000 | 852969984 | 11 | 11 | 52,7 | 52,7 | 52,7 | 24,709 | 0 | 207,14 | 4039800000 | 77 | AJQ90270.1_534 | RM25_0543 | Cytoplasmic | 0 | CYT | 0.100875 | OTHER | J |
| lcl\|CP010341.1_prot_AJQ90271.1_535 | RM25_RS02675 | 30S ribosomal protein S14 type Z | 64210000 | 40330000 | 182050000 | 212060000 | 220150000 | 344820000 | 1 | 1 | 21,3 | 21,3 | 21,3 | 6,9383 | 0,0045351 | 9,6632 | 1196700000 | 10 | AJQ90271.1_535 | RM25_0544 | Cytoplasmic | 0 | CYT | 0.317729 | OTHER | |
| lcl\|CP010341.1_prot_AJQ90272.1_536 | RM25_RS02680 | 30S ribosomal protein S8 | 226190000 | 189400000 | 630579968 | 536660000 | 430769984 | 691529984 | 8 | 8 | 65,2 | 65,2 | 65,2 | 14,765 | 0 | 199,73 | 3170800000 | 70 | AJQ90272.1_536 | RM25_0545 | Cytoplasmic | 0 | CYT | 0.068172 | OTHER | J |
| lcl\|CP010341.1_prot_AJQ90273.1_537 | RM25_RS02685 | 50S ribosomal protein L6 | 268400000 | 334750016 | 802659968 | 492520000 | 877409984 | 1095200000 | 7 | 7 | 45,6 | 45,6 | 45,6 | 19,743 | 0 | 74,195 | 4570800000 | 56 | AJQ90273.1_537 | RM25_0546 | Cytoplasmic | 0 | CYT | 0.127448 | OTHER | J |
| lcl\|CP010341.1_prot_AJQ90274.1_538 | RM25_RS02690 | 50S ribosomal protein L18 | 345070016 | 293369984 | 848300032 | 944230016 | 897240000 | 1252999936 | 8 | 8 | 55,9 | 55,9 | 55,9 | 14,023 | 0 | 95,318 | 5347400000 | 52 | AJQ90274.1_538 | RM25_0547 | Cytoplasmic | 0 | CYT | 0.302649 | OTHER | J |
| lcl\|CP010341.1_prot_AJQ90275.1_539 | RM25_RS02695 | 30S ribosomal protein S5 | 143370000 | 137930000 | 450310016 | 464729984 | 513720000 | 681139968 | 5 | 5 | 23,6 | 23,6 | 23,6 | 21,586 | 0 | 48,232 | 2763500000 | 17 | AJQ90275.1_539 | RM25_0548 | Cytoplasmic | 0 | CYT | 0.046736 | OTHER | J |
| lcl\|CP010341.1_prot_AJQ90276.1_540 | RM25_RS02700 | Hypothetical protein | 224480000 | 288470016 | 592780032 | 924899968 | 857139968 | 979340032 | 3 | 3 | 55 | 55 | 55 | 6,6828 | 0 | 39,117 | 4813300000 | 50 | AJQ90276.1_540 | RM25_0549 | Cytoplasmic | 0 | CYT | 0.215523 | OTHER | |
| lcl\|CP010341.1_prot_AJQ90277.1_541 | RM25_RS02705 | 50S ribosomal protein L15 | 98677000 | 335520000 | 314200000 | 338089984 | 871929984 | 1273400064 | 5 | 5 | 41,5 | 41,5 | 41,5 | 15,464 | 0 | 45,04 | 3344000000 | 21 | AJQ90277.1_541 | RM25_0550 | Cytoplasmic | 0 | CYT | 0.065481 | OTHER | J |
| lcl\|CP010341.1_prot_AJQ90279.1_543 | RM25_RS02715 | Putative luciferase-like monooxygenase, FMN-depende | 8829800 | 7033100 | 19132000 | 5828100 | 6064400 | 8553900 | 2 | 2 | 9,8 | 9,8 | 9,8 | 44,644 | 0 | 151,09 | 63113000 | 4 | AJQ90279.1_543 | RM25_0552 | CytoplasmicMembrane | 0 | CYT | 0.316542 | OTHER | C |
| lcl\|CP010341.1_prot_AJQ90280.1_544 | RM25_RS02720 | Putative extracellular solute-binding dependent transpo | 184100000 | 262820000 | 331740000 | 68055000 | 123470000 | 180180000 | 5 | 5 | 9,3 | 9,3 | 9,3 | 61,767 | 0 | 30,569 | 1196200000 | 11 | AJQ90280.1_544 | RM25_0553 | Cellwall | 0 | SPII | 0.896355 | LIPO(Sec/SPII) | E |
| lcl\|CP010341.1_prot_AJQ90285.1_549 | RM25_RS02745 | Protein translocase subunit SecY | 9736600 | 7520400 | 12457000 | 5833300 | 9886600 | 20118000 | 2 | 2 | 3,7 | 3,7 | 3,7 | 47,759 | 0 | 12,786 | 70575000 | 4 | AJQ90285.1_549 | RM25_0558 | CytoplasmicMembrane | 9 | TMH | 0.953798 | SP(Sec/SPI) | U |
| lcl\|CP010341.1_prot_AJQ90286.1_550 | RM25_RS02750 | Adenylate kinase | 198750000 | 368750016 | 423300000 | 264250000 | 249870000 | 309280000 | 8 | 8 | 61,3 | 61,3 | 61,3 | 19,836 | 0 | 107,79 | 2043500000 | 55 | AJQ90286.1_550 | RM25_0559 | Cytoplasmic | 0 | CYT | 0.090638 | OTHER | F |
| lcl\|CP010341.1_prot_AJQ90289.1_553 | RM25_RS02765 | Hypothetical protein | 53726000 | 51677000 | 81412000 | 47054000 | 33595000 | 43966000 | 3 | 3 | 52,1 | 52,1 | 52,1 | 10,637 | 0 | 48,286 | 352810000 | 12 | AJQ90289.1_553 | RM25_0562 | Cytoplasmic | 0 | CYT | 0.042787 | OTHER | S |
| lcl\|CP010341.1_prot_AJQ90290.1_554 | RM25_RS02770 | Translation initiation factor IF-1 | 56754000 | 75770000 | 132440000 | 121730000 | 177020000 | 202200000 | 4 | 4 | 65,8 | 65,8 | 65,8 | 8,3747 | 0 | 58,93 | 835920000 | 12 | AJQ90290.1_554 | RM25_0563 | Cytoplasmic | 0 | CYT | 0.045792 | OTHER | J |
| lcl\|CP010341.1_prot_AJQ90292.1_556 | RM25_RS02775 | 30S ribosomal protein S13 | 251040000 | 266220000 | 859900032 | 910489984 | 876600000 | 1273299968 | 6 | 6 | 46,8 | 46,8 | 46,8 | 14,026 | 0 | 124,71 | 5208400000 | 45 | AJQ90292.1_556 | RM25_0565 | Cytoplasmic | 0 | CYT | 0.090890 | OTHER | J |
| lcl\|CP010341.1_prot_AJQ90293.1_557 | RM25_RS02780 | 30S ribosomal protein S11 | 39686000 | 38545000 | 98731000 | 113670000 | 192430000 | 246230000 | 4 | 4 | 45,2 | 45,2 | 45,2 | 14,237 | 0 | 32,709 | 810590000 | 18 | AJQ90293.1_557 | RM25_0566 | Cytoplasmic | 0 | CYT | 0.272629 | OTHER | J |
| lcl\|CP010341.1_prot_AJQ90294.1_558 | RM25_RS02785 | 30S ribosomal protein S4 | 512400000 | 466960000 | 1375900032 | 1034099968 | 1087699968 | 1712700032 | 12 | 12 | 49,3 | 49,3 | 49,3 | 23,303 | 0 | 138,37 | 6877400000 | 58 | AJQ90294.1_558 | RM25_0567 | Cytoplasmic | 0 | CYT | 0.104076 | OTHER | J |
| lcl\|CP010341.1_prot_AJQ90295.1_559 | RM25_RS02790 | DNA-directed RNA polymerase, alpha subunit | 361200000 | 284900000 | 801129984 | 734569984 | 526740000 | 1216000000 | 13 | 13 | 47,2 | 47,2 | 47,2 | 36,975 | 0 | 323,31 | 4527400000 | 63 | AJQ90295.1_559 | RM25_0568 | Cytoplasmic | 0 | CYT | 0.075576 | OTHER | K |
| lcl\|CP010341.1_prot_AJQ90296.1_560 | RM25_RS02795 | 50S ribosomal protein L17 | 241780000 | 298160000 | 661820032 | 377160000 | 453929984 | 718480000 | 8 | 8 | 42,1 | 42,1 | 42,1 | 21,401 | 0 | 77,612 | 3054900000 | 49 | AJQ90296.1_560 | RM25_0569 | Cytoplasmic | 0 | CYT | 0.371243 | OTHER | J |
| lcl\|CP010341.1_prot_AJQ90298.1_562 | RM25_RS02805 | Resuscitation-promoting factor RpfB | 3925499904 | 5485599744 | 9288399872 | 5843200000 | 7154699776 | 11438999552 | 11 | 11 | 46,3 | 46,3 | 46,3 | 37,701 | 0 | 323,31 | 4,833E+10 | 156 | AJQ90298.1_562 | RM25_0571 | Extracellular | 1 | SPI | 0.919864 | SP(Sec/SPI) | M |
| lcl\|CP010341.1_prot_AJQ90299.1_563 | RM25_RS02810 | Superoxide dismutase | 516640000 | 244050000 | 1849200000 | 567740032 | 1173200000 | 1448600064 | 5 | 5 | 36 | 36 | 36 | 26,612 | 0 | 107,28 | 5976400000 | 50 | AJQ90299.1_563 | RM25_0572 | Extracellular | 0 | CYT | 0.573730 | OTHER | P |
| lcl\|CP010341.1_prot_AJQ90305.1_569 | RM25_RS02840 | Hypothetical protein | 28181000 | 28029000 | 126200000 | 16573000 | 41610000 | 46951000 | 1 | 1 | 6,1 | 6,1 | 6,1 | 17,944 | 0 | 10,856 | 308950000 | 1 | AJQ90305.1_569 | RM25_0578 | Extracellular | 0 | SPII | 0.896515 | LIPO(Sec/SPII) | |
| lcl\|CP010341.1_prot_AJQ90310.1_574 | RM25_RS02865 | Elongation factor G fusA2 | 24319000 | 22845000 | 68006000 | 49417000 | 48752000 | 160100000 | 4 | 4 | 5,4 | 5,4 | 5,4 | 75,085 | 0 | 24,77 | 406270000 | 10 | AJQ90310.1_574 | RM25_0583 | Cytoplasmic | 0 | CYT | 0.081190 | OTHER | J |
| lcl\|CP010341.1_prot_AJQ90311.1_575 | RM25_RS02870 | 50S ribosomal protein L13 | 110630000 | 131200000 | 339660000 | 253750000 | 341769984 | 529960000 | 4 | 4 | 34 | 34 | 34 | 16,121 | 0 | 43,589 | 1775700000 | 19 | AJQ90311.1_575 | RM25_0584 | Unknown | 0 | CYT | 0.741864 | OTHER | J |
| lcl\|CP010341.1_prot_AJQ90312.1_576 | RM25_RS02875 | 30S ribosomal protein S9 | 495580000 | 503620000 | 1218400000 | 1236600064 | 959030016 | 1443299968 | 8 | 8 | 65 | 65 | 65 | 19,467 | 0 | 166,76 | 7181900000 | 50 | AJQ90312.1_576 | RM25_0585 | Cytoplasmic | 0 | CYT | 0.819396 | OTHER | J |
| lcl\|CP010341.1_prot_AJQ90313.1_577 | RM25_RS02880 | Phosphoglucosamine mutase | 23673000 | 13223000 | 71621000 | 65117000 | 41103000 | 75771000 | 5 | 5 | 15,2 | 15,2 | 15,2 | 46,255 | 0 | 42,183 | 313870000 | 10 | AJQ90313.1_577 | RM25_0586 | Cytoplasmic | 0 | CYT | 0.040137 | OTHER | G |
| lcl\|CP010341.1_prot_AJQ90314.1_578 | RM25_RS02885 | Pantothenate kinase | 12680000 | 16097000 | 36413000 | 18573000 | 19170000 | 33994000 | 4 | 4 | 13,3 | 13,3 | 13,3 | 39,41 | 0 | 29,206 | 148590000 | 21 | AJQ90314.1_578 | RM25_0587 | Cytoplasmic | 0 | SPI | 0.111255 | OTHER | H |
| lcl\|CP010341.1_prot_AJQ90315.1_579 | RM25_RS02890 | Glutamine--fructose-6-phosphate transaminase (isomer | 25486000 | 23260000 | 85534000 | 27599000 | 24026000 | 53410000 | 5 | 5 | 9,1 | 9,1 | 9,1 | 66,893 | 0 | 37,101 | 251860000 | 12 | AJQ90315.1_579 | RM25_0588 | Cytoplasmic | 0 | CYT | 0.066484 | OTHER | M |
| lcl\|CP010341.1_prot_AJQ90319.1_583 | RM25_RS13145 | Hypothetical protein | 17077000 | 29853000 | 59118000 | 39547000 | 46239000 | 76632000 | 4 | 4 | 5 | 5 | 5 | 116,91 | 0 | 39,17 | 302670000 | 13 | AJQ90319.1_583 | RM25_0592 | Cytoplasmic | 0 | CYT | 0.790250 | OTHER | L |
| lcl\|CP010341.1_prot_AJQ90320.1_584 | RM25_RS11790 | Hydrolase, P-loop family | 12524000 | 14186000 | 28158000 | 14051000 | 17083000 | 28160000 | 4 | 4 | 13 | 13 | 13 | 35,03 | 0 | 27,372 | 118030000 | 12 | AJQ90320.1_584 | RM25_0593 | Cytoplasmic | 0 | CYT | 0.070271 | OTHER | K |
| lcl\|CP010341.1_prot_AJQ90323.1_587 | RM25_RS02935 | putative tRNA threonylcarbamoyladenosine biosynthes | 4186600 | 4514100 | 8080200 | 5819600 | 9554300 | 13079000 | 2 | 2 | 11,5 | 11,5 | 11,5 | 36,685 | 0 | 11,57 | 45234000 | 3 | AJQ90323.1_587 | RM25_0596 | Cytoplasmic | 0 | CYT | 0.044482 | OTHER | O |
| lcl\|CP010341.1_prot_AJQ90324.1_588 | RM25_RS02940 | Cob(II)yrinic acid a,c-diamide reductase | 94186000 | 116140000 | 262410000 | 143890000 | 141590000 | 193260000 | 9 | 9 | 16,5 | 16,5 | 16,5 | 66,427 | 0 | 116,78 | 1052600000 | 35 | AJQ90324.1_588 | RM25_0597 | Cytoplasmic | 0 | CYT | 0.062020 | OTHER | CH |
| lcl\|CP010341.1_prot_AJQ90325.1_589 | RM25_RS02945 | Bleomycin hydrolase | 4720600 | 0 | 13906000 | 0 | 2058300 | 3778600 | 2 | 2 | 5,6 | 5,6 | 5,6 | 53,387 | 0 | 14,798 | 24464000 | 2 | AJQ90325.1_589 | RM25_0598 | Cytoplasmic | 0 | CYT | 0.140150 | OTHER | E |
| lcl\|CP010341.1_prot_AJQ90330.1_594 | RM25_RS02970 | Osmoprotectant transport ATP-binding protein ABC trai | 13915000 | 13202000 | 22773000 | 0 | 15211000 | 19959000 | 4 | 4 | 14,9 | 14,9 | 14,9 | 47,948 | 0 | 47,672 | 90922000 | 10 | AJQ90330.1_594 | RM25_0603 | CytoplasmicMembrane | 0 | CYT | 0.072981 | OTHER | |
| lcl\|CP010341.1_prot_AJQ90332.1_596 | RM25_RS02980 | ProXL | 48066000 | 82755000 | 129130000 | 12527000 | 26721000 | 53815000 | 6 | 6 | 20,1 | 20,1 | 20,1 | 37,28 | 0 | 101,13 | 357440000 | 19 | AJQ90332.1_596 | RM25_0605 | CytoplasmicMembrane | 1 | TMH | 0.706404 | LIPO(Sec/SPII) | M |
| lcl\|CP010341.1_prot_AJQ90333.1_597 | RM25_RS02985 | 10 kDa chaperonin | 1538899968 | 1433400064 | 3767399936 | 1428400000 | 1711800064 | 2421799936 | 5 | 5 | 63,3 | 63,3 | 63,3 | 10,597 | 0 | 211,24 | 1,341E+10 | 38 | AJQ90333.1_597 | RM25_0606 | Cytoplasmic | 0 | CYT | 0.308932 | OTHER | O |
| lcl\|CP010341.1_prot_AJQ90334.1_598 | RM25_RS02990 | 60 kDa chaperonin 1 | 2744600064 | 2423600128 | 6126700032 | 5488200192 | 7000399872 | 9862500352 | 26 | 26 | 74 | 74 | 74 | 56,146 | 0 | 323,31 | 3,7612E+10 | 234 | AJQ90334.1_598 | RM25_0607 | Cytoplasmic | 0 | CYT | 0.044350 | OTHER | O |
| lcl\|CP010341.1_prot_AJQ90335.1_599 | RM25_RS02995 | Inosine-5'-monophosphate dehydrogenase | 102380000 | 37799000 | 193890000 | 161600000 | 182200000 | 294470016 | 12 | 12 | 29,2 | 29,2 | 29,2 | 53,857 | 0 | 83,39 | 1032800000 | 40 | AJQ90335.1_599 | RM25_0608 | Cytoplasmic | 0 | CYT | 0.072799 | OTHER | F |
| lcl\|CP010341.1_prot_AJQ90336.1_600 | RM25_RS03000 | IMP dehydrogenase family protein | 29092000 | 24716000 | 81795000 | 21927000 | 51506000 | 47555000 | 6 | 6 | 22,6 | 22,6 | 22,6 | 39,503 | 0 | 55,679 | 269040000 | 11 | AJQ90336.1_600 | RM25_0609 | Cytoplasmic | 0 | CYT | 0.049161 | OTHER | F |
| lcl\|CP010341.1_prot_AJQ90337.1_601 | RM25_RS03005 | Chorismate mutase | 10816000 | 8646200 | 26993000 | 16408000 | 18388000 | 16465000 | 2 | 2 | 22,9 | 22,9 | 22,9 | 12,192 | 0,0023095 | 10,425 | 109570000 | 4 | AJQ90337.1_601 | RM25_0610 | Cytoplasmic | 0 | CYT | 0.065427 | OTHER | |
| lcl\|CP010341.1_prot_AJQ90339.1_603 | RM25_RS03015 | Hypothetical protein | 88884000 | 169040000 | 306449984 | 63523000 | 142410000 | 159200000 | 4 | 4 | 6,5 | 6,5 | 6,5 | 58,461 | 0 | 31,782 | 983260000 | 20 | AJQ90339.1_603 | RM25_0612 | CytoplasmicMembrane | 1 | CYT | 0.946382 | OTHER | |
| lcl\|CP010341.1_prot_AJQ90341.1_605 | RM25_RS03025 | AmiC, N-acetylmuramoyl-L-alanine amidase | 6168300 | 10018000 | 7283800 | 0 | 8302800 | 2032200 | 2 | 2 | 10 | 10 | 10 | 28,536 | 0 | 14,665 | 33805000 | 4 | AJQ90341.1_605 | RM25_0614 | CytoplasmicMembrane | 1 | SPI | 0.627699 | SP(Sec/SPI) | |
| lcl\|CP010341.1_prot_AJQ90342.1_606 | RM25_RS03030 | Serine acetyltransferase | 2747500 | 0 | 7108900 | 5454300 | 0 | 5565600 | 1 | 1 | 5,5 | 5,5 | 5,5 | 19,84 | 0,009901 | 6,003 | 24534000 | 3 | AJQ90342.1_606 | RM25_0615 | Cytoplasmic | 0 | CYT | 0.041835 | OTHER | E |
| lcl\|CP010341.1_prot_AJQ90355.1_619 | RM25_RS03080 | GMP synthase (glutamine-hydrolyzing) | 109950000 | 123700000 | 343280000 | 139580000 | 114020000 | 151260000 | 15 | 15 | 43,4 | 43,4 | 43,4 | 51,373 | 0 | 131,09 | 1027400000 | 32 | AJQ90355.1_619 | RM25_0628 | Cytoplasmic | 0 | CYT | 0.062385 | OTHER | F |
| lcl\|CP010341.1_prot_AJQ90361.1_625 | RM25_RS03105 | Two component transcriptional regulator, LuxR family | 8261300 | 15056000 | 28126000 | 12103000 | 19249000 | 0 | 3 | 3 | 20,7 | 20,7 | 20,7 | 25,153 | 0 | 20,636 | 100540000 | 2 | AJQ90361.1_625 | RM25_0634 | Cytoplasmic | 0 | CYT | 0.044412 | OTHER | T |
| lcl\|CP010341.1_prot_AJQ90363.1_627 | RM25_RS03115 | Iron compound ABC transporter, iron compound-bindir | 196540000 | 301830016 | 488209984 | 788819968 | 1161600000 | 3219399936 | 16 | 16 | 56,2 | 56,2 | 56,2 | 35,557 | 0 | 255,25 | 7123700000 | 97 | AJQ90363.1_627 | RM25_0636 | Unknown | 1 | SPII | 0.915942 | LIPO(Sec/SPII) | |
| lcl\|CP010341.1_prot_AJQ90369.1_633 | RM25_RS03145 | ATP-dependent DNA helicase PcrA | 11699000 | 13638000 | 22070000 | 47458000 | 52483000 | 48044000 | 2 | 2 | 3 | 3 | 3 | 90,861 | 0 | 10,865 | 222270000 | 4 | AJQ90369.1_633 | RM25_0642 | Cytoplasmic | 0 | CYT | 0.085536 | OTHER | L |
| lcl\|CP010341.1_prot_AJQ90372.1_636 | RM25_RS03160 | Bifunctional purine biosynthesis protein PurH | 66264000 | 47906000 | 206880000 | 60509000 | 58762000 | 69760000 | 7 | 7 | 20,7 | 20,7 | 20,7 | 54,491 | 0 | 86,274 | 532120000 | 21 | AJQ90372.1_636 | RM25_0645 | Cytoplasmic | 0 | CYT | 0.069266 | OTHER | F |
| lcl\|CP010341.1_prot_AJQ90373.1_637 | RM25_RS03165 | Bifunctional protein FolD | 9420800 | 5844700 | 35654000 | 4559400 | 5871700 | 8080700 | 3 | 3 | 8,4 | 8,4 | 8,4 | 30,547 | 0 | 19,377 | 69431000 | 5 | AJQ90373.1_637 | RM25_0646 | Cytoplasmic | 0 | CYT | 0.079023 | OTHER | H |
| lcl\|CP010341.1_prot_AJQ90375.1_639 | RM25_RS03175 | Malate dehydrogenase | 1211800064 | 1007900032 | 3099500032 | 1038499968 | 995830016 | 1533299968 | 8 | 8 | 34,5 | 34,5 | 34,5 | 34,909 | 0 | 166,35 | 9365000000 | 61 | AJQ90375.1_639 | RM25_0648 | Cytoplasmic | 0 | CYT | 0.114624 | OTHER | C |
| lcl\|CP010341.1_prot_AJQ90376.1_640 | RM25_RS03180 | Isocitrate dehydrogenase (NADP) | 189360000 | 211100000 | 626470016 | 206300000 | 207820000 | 306420000 | 13 | 13 | 35,8 | 35,8 | 35,8 | 44,79 | 0 | 174,73 | 1862700000 | 78 | AJQ90376.1_640 | RM25_0649 | Cytoplasmic | 0 | CYT | 0.096150 | OTHER | |
| lcl\|CP010341.1_prot_AJQ90382.1_646 | RM25_RS03210 | Basic membrane protein | 74912000 | 109160000 | 169950000 | 32204000 | 66025000 | 92425000 | 2 | 2 | 9,9 | 9,9 | 9,9 | 31,261 | 0 | 28,311 | 611590000 | 16 | AJQ90382.1_646 | RM25_0655 | Unknown | 0 | CYT | 0.686081 | OTHER | |
| lcl\|CP010341.1_prot_AJQ90383.1_647 | RM25_RS03215 | Ribonuclease H | 4172900 | 4237700 | 11246000 | 0 | 0 | 10578000 | 1 | 1 | 3,2 | 3,2 | 3,2 | 36,849 | 0,005102 | 6,5247 | 37512000 | 1 | AJQ90383.1_647 | RM25_0656 | Unknown | 0 | CYT | 0.261903 | OTHER | L |
| lcl\|CP010341.1_prot_AJQ90385.1_649 | RM25_RS03225 | Enoyl-CoA hydratase echA9 | 36330000 | 59860000 | 111550000 | 122700000 | 130960000 | 73570000 | 7 | 7 | 26,5 | 26,5 | 26,5 | 33,239 | 0 | 78,379 | 574200000 | 27 | AJQ90385.1_649 | RM25_0658 | CytoplasmicMembrane | 0 | CYT | 0.039217 | OTHER | I |
| lcl\|CP010341.1_prot_AJQ90386.1_650 | RM25_RS03230 | Methionine--tRNA ligase | 56094000 | 53103000 | 187350000 | 72936000 | 98201000 | 138680000 | 9 | 9 | 19,3 | 19,3 | 19,3 | 71,839 | 0 | 112,86 | 663520000 | 22 | AJQ90386.1_650 | RM25_0659 | Cytoplasmic | 0 | CYT | 0.276409 | OTHER | J |
| lcl\|CP010341.1_prot_AJQ90398.1_662 | RM25_RS03285 | Chlorophyll synthesis pathway protein BchC | 9879500 | 8471400 | 25408000 | 2033200 | 3889500 | 2601900 | 3 | 3 | 12,1 | 12,1 | 12,1 | 37,127 | 0 | 20,804 | 52284000 | 5 | AJQ90398.1_662 | RM25_0671 | Cytoplasmic | 0 | CYT | 0.050289 | OTHER | |
| lcl\|CP010341.1_prot_AJQ90399.1_663 | RM25_RS03290 | Putative membrane protein (DUF2207) (Precursor) | 13533000 | 22440000 | 11319000 | 9824300 | 10852000 | 9123700 | 2 | 2 | 4,7 | 4,7 | 4,7 | 64,274 | 0 | 15,602 | 89844000 | 4 | AJQ90399.1_663 | RM25_0672 | CytoplasmicMembrane | 4 | SPI | 0.778701 | SP(Sec/SPI) | |
| lcl\|CP010341.1_prot_AJQ90400.1_664 | RM25_RS03295 | Pyruvate-flavodoxin oxidoreductase | 53109000 | 42605000 | 112520000 | 80573000 | 74608000 | 203910000 | 15 | 15 | 15,4 | 15,4 | 15,4 | 132,82 | 0 | 135,8 | 633410000 | 30 | AJQ90400.1_664 | RM25_0673 | Cytoplasmic | 0 | CYT | 0.121606 | OTHER | C |
| lcl\|CP010341.1_prot_AJQ90401.1_665 | RM25_RS03300 | Dihydroorotate oxidase | 9206500 | 12261000 | 17782000 | 16863000 | 14489000 | 27480000 | 3 | 3 | 10,4 | 10,4 | 10,4 | 35,394 | 0 | 18,711 | 107390000 | 7 | AJQ90401.1_665 | RM25_0674 | Cytoplasmic | 0 | CYT | 0.069491 | OTHER | |
| lcl\|CP010341.1_prot_AJQ90403.1_667 | RM25_RS03310 | Bifunctional acetyl-/propionyl-coenzyme A carboxylase | 72007000 | 59981000 | 172830000 | 70846000 | 87146000 | 123510000 | 9 | 9 | 20,6 | 20,6 | 20,6 | 63,48 | 0 | 80,538 | 653700000 | 31 | AJQ90403.1_667 | RM25_0676 | Cytoplasmic | 0 | CYT | 0.069318 | OTHER | I |
| lcl\|CP010341.1_prot_AJQ90406.1_670 | RM25_RS12555 | Hypothetical protein | 31775000 | 37455000 | 42178000 | 13517000 | 36566000 | 42768000 | 2 | 2 | 20,1 | 20,1 | 20,1 | 16,152 | 0 | 25,501 | 215360000 | 5 | AJQ90406.1_670 | RM25_0679 | Unknown | 1 | SPII | 0.928766 | LIPO(Sec/SPII) | |
| lcl\|CP010341.1_prot_AJQ90408.1_672 | RM25_RS03330 | Septum formation protein Maf | 1972700 | 1865900 | 5931700 | 0 | 0 | 4012200 | 1 | 1 | 7,9 | 7,9 | 7,9 | 25,941 | 0 | 13,674 | 13783000 | 3 | AJQ90408.1_672 | RM25_0681 | Cytoplasmic | 0 | CYT | 0.084671 | OTHER | D |
| lcl\|CP010341.1_prot_AJQ90410.1_674 | RM25_RS03340 | Propionyl-CoA carboxylase subunit beta 5 accD5 | 39671000 | 23984000 | 95280000 | 55982000 | 79366000 | 122220000 | 8 | 8 | 22,5 | 22,5 | 22,5 | 57,075 | 0 | 106,57 | 440760000 | 25 | AJQ90410.1_674 | RM25_0683 | Cytoplasmic | 0 | CYT | 0.079680 | OTHER | I |
| lcl\|CP010341.1_prot_AJQ90419.1_683 | RM25_RS03380 | Biotin-acetyl-CoA-carboxylase ligase | 1778900 | 0 | 5005300 | 0 | 4220600 | 6377300 | 1 | 1 | 6,5 | 6,5 | 6,5 | 28,499 | 0,0043337 | 7,6076 | 21132000 | 1 | AJQ90419.1_683 | RM25_0692 | Cytoplasmic | 0 | CYT | 0.072190 | OTHER | H |
| lcl\|CP010341.1_prot_AJQ90423.1_687 | RM25_RS03400 | Peptidase family S51 | 15653000 | 13540000 | 38797000 | 16058000 | 22477000 | 29734000 | 2 | 2 | 7,1 | 7,1 | 7,1 | 24,878 | 0 | 16,394 | 143930000 | 5 | AJQ90423.1_687 | RM25_0696 | Unknown | 0 | CYT | 0.067805 | OTHER | E |
| lcl\|CP010341.1_prot_AJQ90424.1_688 | RM25_RS03405 | Phosphoribosylaminoimidazole carboxylase, ATPase su | 54877000 | 86317000 | 171970000 | 123960000 | 99854000 | 120150000 | 8 | 8 | 31,8 | 31,8 | 31,8 | 41,77 | 0 | 113,56 | 728210000 | 30 | AJQ90424.1_688 | RM25_0697 | CytoplasmicMembrane | 0 | CYT | 0.069465 | OTHER | F |
| lcl\|CP010341.1_prot_AJQ90426.1_690 | RM25_RS03415 | Cell envelope-related transcriptional regulator | 5012900 | 14107000 | 13861000 | 9933000 | 0 | 14757000 | 4 | 4 | 10,4 | 10,4 | 10,4 | 48,391 | 0 | 28,826 | 63370000 | 7 | AJQ90426.1_690 | RM25_0699 | CytoplasmicMembrane | 1 | CYT | 0.679155 | OTHER | K |
| lcl\|CP010341.1_prot_AJQ90427.1_691 | RM25_RS03420 | Mannose-1-phosphate guanylyltransferase | 20814000 | 21424000 | 59918000 | 20134000 | 31525000 | 51830000 | 4 | 4 | 13,4 | 13,4 | 13,4 | 38,941 | 0 | 24,882 | 219060000 | 9 | AJQ90427.1_691 | RM25_0700 | Cytoplasmic | 0 | CYT | 0.050290 | OTHER | M |
| lcl\|CP010341.1_prot_AJQ90428.1_692 | RM25_RS03425 | Putative membrane protein | 2784600 | 3416300 | 7393500 | 6094300 | 9363900 | 10461000 | 1 | 1 | 6,8 | 6,8 | 6,8 | 36,582 | 0 | 11,163 | 39513000 | 1 | AJQ90428.1_692 | RM25_0701 | CytoplasmicMembrane | 4 | CYT | 0.937566 | OTHER | |
| lcl\|CP010341.1_prot_AJQ90430.1_694 | RM25_RS03435 | Hypothetical protein | 2593000 | 0 | 6182700 | 0 | 0 | 0 | 1 | 1 | 1 | 1 | 1 | 112,66 | 0,0041537 | 6,8551 | 8775700 | 1 | AJQ90430.1_694 | RM25_0703 | CytoplasmicMembrane | 9 | CYT | 0.822891 | OTHER | S |
| lcl\|CP010341.1_prot_AJQ90431.1_695 | RM25_RS03440 | Hypothetical protein | 18291000 | 18642000 | 12029000 | 6557500 | 14333000 | 23719000 | 5 | 5 | 14,4 | 14,4 | 14,4 | 45,249 | 0 | 40,996 | 100110000 | 4 | AJQ90431.1_695 | RM25_0704 | Unknown | 1 | CYT | 0.203353 | OTHER | |
| lcl\|CP010341.1_prot_AJQ90434.1_698 | RM25_RS03455 | Eukaryotic phosphomannomutase | 29435000 | 52954000 | 54676000 | 21271000 | 43538000 | 12385000 | 4 | 4 | 21 | 21 | 21 | 29,26 | 0 | 32,855 | 227300000 | 10 | AJQ90434.1_698 | RM25_0707 | Cytoplasmic | 0 | CYT | 0.114583 | OTHER | |
| lcl\|CP010341.1_prot_AJQ90436.1_700 | RM25_RS03465 | Phospho-glucose isomerase C-terminal domain protein | 16285000 | 23101000 | 51590000 | 16249000 | 34954000 | 53418000 | 6 | 6 | 22,2 | 22,2 | 22,2 | 34,901 | 0 | 45,822 | 210720000 | 16 | AJQ90436.1_700 | RM25_0709 | Unknown | 0 | CYT | 0.037488 | OTHER | K |
| lcl\|CP010341.1_prot_AJQ90438.1_702 | RM25_RS03475 | Hypothetical protein | 14389000 | 18037000 | 29366000 | 30981000 | 34817000 | 36485000 | 2 | 2 | 27,6 | 27,6 | 27,6 | 9,7291 | 0 | 12,266 | 181690000 | 4 | AJQ90438.1_702 | RM25_0711 | Unknown | 0 | CYT | 0.359708 | OTHER | |
| lcl\|CP010341.1_prot_AJQ90440.1_704 | RM25_RS03485 | Hypothetical protein | 7266900 | 7841300 | 17682000 | 0 | 8399600 | 11052000 | 2 | 2 | 9,7 | 9,7 | 9,7 | 37,432 | 0 | 12,764 | 55617000 | 0 | AJQ90440.1_704 | RM25_0713 | CytoplasmicMembrane | 3 | CYT | 0.970202 | OTHER | |
| lcl\|CP010341.1_prot_AJQ90442.1_706 | RM25_RS03495 | O-succinylbenzoate synthase | 2075500 | 3823600 | 9707200 | 2312400 | 3666600 | 10740000 | 2 | 2 | 7,8 | 7,8 | 7,8 | 35,17 | 0 | 11,461 | 32325000 | 1 | AJQ90442.1_706 | RM25_0715 | Cytoplasmic | 0 | CYT | 0.034377 | OTHER | M |
| lcl\|CP010341.1_prot_AJQ90448.1_712 | RM25_RS03525 | Naphthoate synthase | 9617700 | 12678000 | 29671000 | 17324000 | 27241000 | 35012000 | 3 | 3 | 11,1 | 11,1 | 11,1 | 39,348 | 0 | 27,53 | 148900000 | 11 | AJQ90448.1_712 | RM25_0721 | Unknown | 0 | CYT | 0.215309 | OTHER | H |
| lcl\|CP010341.1_prot_AJQ90449.1_713 | RM25_RS03530 | Nicotinate phosphoribosyltransferase | 4638200 | 4556500 | 14435000 | 7406100 | 4959400 | 7854200 | 3 | 3 | 7,5 | 7,5 | 7,5 | 48,289 | 0 | 20,838 | 45574000 | 6 | AJQ90449.1_713 | RM25_0722 | Cytoplasmic | 0 | CYT | 0.069907 | OTHER | H |
| lcl\|CP010341.1_prot_AJQ90452.1_716 | RM25_RS03545 | Glutamate racemase | 3193900 | 4164400 | 9306100 | 6943700 | 6232800 | 8334200 | 1 | 1 | 5,1 | 5,1 | 5,1 | 29,444 | 0,004128 | 6,7056 | 38175000 | 0 | AJQ90452.1_716 | RM25_0725 | Cytoplasmic | 0 | CYT | 0.042110 | OTHER | M |
| lcl\|CP010341.1_prot_AJQ90453.1_717 | RM25_RS03550 | TRNA nucleotidyltransferase | 3115400 | 2520400 | 10759000 | 4024100 | 2465900 | 8549200 | 1 | 1 | 5,1 | 5,1 | 5,1 | 27,272 | 0,0042553 | 7,3849 | 31434000 | 3 | AJQ90453.1_717 | RM25_0726 | Cytoplasmic | 0 | CYT | 0.059843 | OTHER | J |
| lcl\|CP010341.1_prot_AJQ90454.1_718 | RM25_RS03545 | Non-canonical purine NTP pyrophosphatase | 3999100 | 0 | 8908300 | 0 | 11455000 | 11358000 | 1 | 1 | 7,5 | 7,5 | 7,5 | 28,871 | 0,0051335 | 6,6448 | 35720000 | 1 | AJQ90454.1_718 | RM25_0727 | Cytoplasmic | 0 | CYT | 0.253390 | OTHER | F |
| lcl\|CP010341.1_prot_AJQ90456.1_720 | RM25_RS03565 | Dihydrofolate reductase | 30419000 | 37865000 | 76636000 | 32358000 | 16215000 | 48346000 | 4 | 4 | 32 | 32 | 32 | 21,478 | 0 | 51,903 | 260210000 | 18 | AJQ90456.1_720 | RM25_0729 | Unknown | 0 | CYT | 0.201251 | OTHER | |

| | | | | | | | | | | | | | | | | | | | | | | | | |
|---|---|---|---|---|---|---|---|---|---|---|---|---|---|---|---|---|---|---|---|---|---|---|---|---|
| lcl\|CP010341.1_prot_AJQ90458.1_722 | RM25_RS03575 | LAO/AO transport system ATPase | 38407000 | 30050000 | 104580000 | 44517000 | 48290000 | 84444000 | 6 | 6 | 20,2 | 20,2 | 20,2 | 36,042 | 0 | 40,458 | 375270000 | 17 | AJQ90458.1_722 | RM25_0731 | CytoplasmicMembrane | 0 | CYT | 0.045317 | OTHER | E |
| lcl\|CP010341.1_prot_AJQ90459.1_723 | RM25_RS03580 | Methylmalonyl-CoA mutase | 558729984 | 588080000 | 1453400064 | 504080000 | 613020032 | 853900032 | 23 | 23 | 41,6 | 41,6 | 41,6 | 80,177 | 0 | 323,31 | 5003700000 | 116 | AJQ90459.1_723 | RM25_0732 | Cytoplasmic | 0 | CYT | 0.085945 | OTHER | I |
| lcl\|CP010341.1_prot_AJQ90460.1_724 | RM25_RS03585 | Methylmalonyl-CoA mutase, small subunit | 412369984 | 223370000 | 929619968 | 328740000 | 443790016 | 606129984 | 19 | 19 | 34,8 | 34,8 | 34,8 | 69,542 | 0 | 315,62 | 3267500000 | 89 | AJQ90460.1_724 | RM25_0733 | Cytoplasmic | 0 | CYT | 0.076138 | OTHER | I |
| lcl\|CP010341.1_prot_AJQ90462.1_726 | RM25_RS03590 | Precorrin-2 C20-methyltransferase CbiL | 79584000 | 70713000 | 220780000 | 165790000 | 158080000 | 249580000 | 5 | 5 | 25,7 | 25,7 | 25,7 | 26,883 | 0 | 42,864 | 1057900000 | 28 | AJQ90462.1_726 | RM25_0735 | Cytoplasmic | 0 | CYT | 0.036687 | OTHER | |
| lcl\|CP010341.1_prot_AJQ90463.1_727 | RM25_RS03595 | Tetrapyrrole methylase, precorrin-3 methylase, CbiF | 22646000 | 45421000 | 143420000 | 38604000 | 57024000 | 85350000 | 6 | 6 | 29,2 | 29,2 | 29,2 | 33,891 | 0 | 51,162 | 410550000 | 14 | AJQ90463.1_727 | RM25_0736 | Cytoplasmic | 0 | CYT | 0.081466 | OTHER | H |
| lcl\|CP010341.1_prot_AJQ90464.1_728 | RM25_RS03600 | Bifunctional cbiH protein and precorrin-3B C17-methylt | 128070000 | 113990000 | 355809984 | 265180000 | 250100000 | 375710016 | 16 | 16 | 24,4 | 24,4 | 24,4 | 91,429 | 0 | 180,19 | 1671200000 | 55 | AJQ90464.1_728 | RM25_0737 | Cytoplasmic | 0 | CYT | 0.065358 | OTHER | H |
| lcl\|CP010341.1_prot_AJQ90465.1_729 | RM25_RS03605 | Sirohydrochlorin cobaltochelatase | 38482000 | 24998000 | 97266000 | 66027000 | 68735000 | 84479000 | 7 | 7 | 20,6 | 20,6 | 20,6 | 44,471 | 0 | 52,688 | 424200000 | 16 | AJQ90465.1_729 | RM25_0738 | Cytoplasmic | 0 | CYT | 0.060965 | OTHER | H |
| lcl\|CP010341.1_prot_AJQ90467.1_731 | RM25_RS03615 | Precorrin-8X methylmutase | 15632000 | 25258000 | 40161000 | 32291000 | 21724000 | 31009000 | 3 | 3 | 21,2 | 21,2 | 21,2 | 22,952 | 0 | 23,052 | 192920000 | 10 | AJQ90467.1_731 | RM25_0740 | Cytoplasmic | 0 | CYT | 0.047737 | OTHER | H |
| lcl\|CP010341.1_prot_AJQ90472.1_736 | RM25_RS12705 | Hypothetical protein | 114330000 | 85301000 | 189330000 | 64672000 | 92267000 | 197920000 | 2 | 2 | 29,3 | 29,3 | 29,3 | 10,745 | 0 | 36,042 | 803720000 | 16 | AJQ90472.1_736 | RM25_0745 | Unknown | 1 | CYT | 0.726869 | OTHER | |
| lcl\|CP010341.1_prot_AJQ90474.1_738 | RM25_RS03660 | Cobyric acid synthase cobQ2 | 955450 | 690890 | 1803600 | 0 | 1144500 | 0 | 2 | 2 | 8,9 | 8,9 | 8,9 | 27,636 | 0 | 12,72 | 4594400 | 2 | AJQ90474.1_738 | RM25_0748 | CytoplasmicMembrane | 0 | CYT | 0.070727 | OTHER | S |
| lcl\|CP010341.1_prot_AJQ90475.1_739 | RM25_RS03665 | Mur ligase middle domain protein | 2932100 | 3174400 | 6043900 | 12539000 | 12088000 | 10406000 | 2 | 2 | 6,8 | 6,8 | 6,8 | 46,781 | 0 | 12,047 | 50058000 | 4 | AJQ90475.1_739 | RM25_0749 | Unknown | 0 | CYT | 0.075225 | OTHER | M |
| lcl\|CP010341.1_prot_AJQ90479.1_743 | RM25_RS03690 | Oligoribonuclease | 71426000 | 64753000 | 191140000 | 114200000 | 144190000 | 207470000 | 7 | 7 | 43,8 | 43,8 | 43,8 | 25,179 | 0 | 50,246 | 861840000 | 27 | AJQ90479.1_743 | RM25_0753 | Cytoplasmic | 0 | CYT | 0.198649 | OTHER | L |
| lcl\|CP010341.1_prot_AJQ90483.1_747 | RM25_RS03715 | ATP/GTP-binding family protein | 3972600 | 4117600 | 11132000 | 2489800 | 5056400 | 13065000 | 3 | 3 | 8,6 | 8,6 | 8,6 | 58,088 | 0 | 18,498 | 39834000 | 4 | AJQ90483.1_747 | RM25_0758 | CytoplasmicMembrane | 2 | CYT | 0.575280 | OTHER | S |
| lcl\|CP010341.1_prot_AJQ90484.1_748 | RM25_RS11810 | Hypothetical protein | 14927000 | 25256000 | 93872000 | 41204000 | 38995000 | 45970000 | 4 | 4 | 26,4 | 26,4 | 26,4 | 24,204 | 0 | 76,195 | 287290000 | 12 | AJQ90484.1_748 | RM25_0759 | Cytoplasmic | 0 | CYT | 0.097572 | OTHER | Q |
| lcl\|CP010341.1_prot_AJQ90487.1_751 | RM25_RS03735 | ATP-binding cassette protein, ChvD family | 348969984 | 436260000 | 808840000 | 470200000 | 507060000 | 593190016 | 19 | 19 | 44,3 | 44,3 | 44,3 | 62,562 | 0 | 310,55 | 3493400000 | 89 | AJQ90487.1_751 | RM25_0762 | Cytoplasmic | 0 | CYT | 0.130922 | OTHER | S |
| lcl\|CP010341.1_prot_AJQ90488.1_752 | RM25_RS03740 | Dihydroxyacetone kinase, DhaK subunit | 5902200 | 0 | 16523000 | 5577600 | 2240800 | 2341100 | 3 | 3 | 9,4 | 9,4 | 9,4 | 34,696 | 0 | 19,679 | 33851000 | 2 | AJQ90488.1_752 | RM25_0763 | Cytoplasmic | 0 | CYT | 0.052094 | OTHER | G |
| lcl\|CP010341.1_prot_AJQ90489.1_753 | RM25_RS03745 | Dihydroxyacetone kinase, L subunit | 12094000 | 16995000 | 36936000 | 70967000 | 108830000 | 47012000 | 5 | 5 | 29,7 | 29,7 | 29,7 | 21,792 | 0 | 30,932 | 331660000 | 9 | AJQ90489.1_753 | RM25_0764 | Cytoplasmic | 0 | CYT | 0.042658 | OTHER | S |
| lcl\|CP010341.1_prot_AJQ90490.1_754 | RM25_RS03750 | Dihydroxyacetone kinase subunit M | 11237000 | 8489900 | 19932000 | 62871000 | 42069000 | 46145000 | 2 | 2 | 8,5 | 8,5 | 8,5 | 51,795 | 0 | 11,776 | 244850000 | 2 | AJQ90490.1_754 | RM25_0765 | Cytoplasmic | 0 | CYT | 0.053831 | OTHER | G |
| lcl\|CP010341.1_prot_AJQ90492.1_756 | RM25_RS03760 | Peptidyl-dipeptidase Dcp | 7232800 | 5610100 | 30775000 | 7170600 | 0 | 16765000 | 3 | 3 | 5,6 | 5,6 | 5,6 | 75,292 | 0 | 24,369 | 70263000 | 3 | AJQ90492.1_756 | RM25_0767 | Cytoplasmic | 0 | CYT | 0.086212 | OTHER | |
| lcl\|CP010341.1_prot_AJQ90493.1_757 | RM25_RS03765 | FAD linked oxidase domain protein | 3257400 | 5002300 | 16513000 | 10686000 | 10477000 | 25831000 | 3 | 3 | 4 | 4 | 4 | 101,03 | 0 | 19,687 | 71766000 | 4 | AJQ90493.1_757 | RM25_0768 | CytoplasmicMembrane | 0 | CYT | 0.099061 | OTHER | C |
| lcl\|CP010341.1_prot_AJQ90495.1_759 | RM25_RS03775 | Long-chain-fatty-acid--CoA ligase | 0 | 0 | 9154900 | 0 | 4645800 | 11495000 | 1 | 1 | 3 | 3 | 3 | 69,995 | 0,0050659 | 6,3398 | 25296000 | 1 | AJQ90495.1_759 | RM25_0770 | CytoplasmicMembrane | 0 | CYT | 0.089484 | OTHER | I |
| lcl\|CP010341.1_prot_AJQ90497.1_761 | RM25_RS03785 | Phosphomethylpyrimidine kinase | 30662000 | 49349000 | 75126000 | 79093000 | 70493000 | 110590000 | 5 | 5 | 34,1 | 34,1 | 34,1 | 28,724 | 0 | 120,5 | 488100000 | 19 | AJQ90497.1_761 | RM25_0772 | Cytoplasmic | 0 | SPI | 0.068637 | OTHER | |
| lcl\|CP010341.1_prot_AJQ90498.1_762 | RM25_RS03790 | Phosphogluconate dehydrogenase (decarboxylating) | 37604000 | 27755000 | 110780000 | 39541000 | 26983000 | 54365000 | 6 | 6 | 14,4 | 14,4 | 14,4 | 52,611 | 0 | 48,937 | 300560000 | 20 | AJQ90498.1_762 | RM25_0773 | Cytoplasmic | 0 | CYT | 0.082683 | OTHER | G |
| lcl\|CP010341.1_prot_AJQ90499.1_763 | RM25_RS03795 | Aminopeptidase N | 134670000 | 144830000 | 425120000 | 141690000 | 113310000 | 176810000 | 17 | 17 | 25 | 25 | 25 | 95,657 | 0 | 128,16 | 1170800000 | 56 | AJQ90499.1_763 | RM25_0774 | Cytoplasmic | 0 | CYT | 0.104969 | OTHER | E |
| lcl\|CP010341.1_prot_AJQ90500.1_764 | RM25_RS03800 | DSBA oxidoreductase | 30017000 | 20452000 | 40288000 | 35487000 | 37269000 | 23775000 | 3 | 3 | 11,3 | 11,3 | 11,3 | 23,032 | 0 | 22,443 | 206280000 | 10 | AJQ90500.1_764 | RM25_0775 | Cytoplasmic | 0 | CYT | 0.113585 | OTHER | Q |
| lcl\|CP010341.1_prot_AJQ90501.1_765 | RM25_RS03805 | Hypothetical protein | 5666100 | 4169400 | 10896000 | 2154000 | 0 | 0 | 1 | 1 | 6,1 | 6,1 | 6,1 | 29,053 | 0,0044543 | 8,5039 | 22886000 | 1 | AJQ90501.1_765 | RM25_0776 | Unknown | 0 | CYT | 0.084832 | OTHER | S |
| lcl\|CP010341.1_prot_AJQ90502.1_766 | RM25_RS03810 | DNA ligase | 12578000 | 13033000 | 22003000 | 13367000 | 27419000 | 15377000 | 4 | 4 | 9,4 | 9,4 | 9,4 | 83,261 | 0 | 47,263 | 118750000 | 7 | AJQ90502.1_766 | RM25_0777 | Cytoplasmic | 0 | CYT | 0.138128 | OTHER | L |
| lcl\|CP010341.1_prot_AJQ90506.1_770 | RM25_RS03840 | Trigger factor | 954569984 | 1190800000 | 2369100032 | 1443000064 | 2133500032 | 2525299968 | 18 | 18 | 56,2 | 56,2 | 56,2 | 57,471 | 0 | 323,31 | 1,1977E+10 | 157 | AJQ90506.1_770 | RM25_0782 | Cytoplasmic | 0 | CYT | 0.299075 | OTHER | D |
| lcl\|CP010341.1_prot_AJQ90509.1_773 | RM25_RS03855 | ATP-dependent Clp protease proteolytic subunit 2 | 37437000 | 19003000 | 115980000 | 88946000 | 86121000 | 180750000 | 6 | 6 | 40,3 | 40,3 | 40,3 | 22,645 | 0 | 63,213 | 590380000 | 19 | AJQ90509.1_773 | RM25_0785 | Cytoplasmic | 0 | CYT | 0.169716 | OTHER | OU |
| lcl\|CP010341.1_prot_AJQ90510.1_774 | RM25_RS03860 | ATP-dependent Clp protease proteolytic subunit 1 | 61022000 | 45558000 | 131610000 | 54242000 | 70224000 | 126220000 | 4 | 4 | 27,5 | 27,5 | 27,5 | 24,806 | 0 | 58,87 | 530780000 | 19 | AJQ90510.1_774 | RM25_0786 | Cytoplasmic | 0 | CYT | 0.196118 | OTHER | OU |
| lcl\|CP010341.1_prot_AJQ90513.1_777 | RM25_RS03875 | Threonine synthase | 63784000 | 88560000 | 192290000 | 109610000 | 90148000 | 124430000 | 9 | 9 | 22,7 | 22,7 | 22,7 | 52,506 | 0 | 80,475 | 730710000 | 26 | AJQ90513.1_777 | RM25_0789 | Cytoplasmic | 0 | CYT | 0.067874 | OTHER | |
| lcl\|CP010341.1_prot_AJQ90514.1_778 | RM25_RS03880 | 6-phosphofructokinase 2 | 5511600 | 2927400 | 13262000 | 14532000 | 10127000 | 12424000 | 1 | 1 | 3,5 | 3,5 | 3,5 | 38,965 | 0,0042373 | 7,3681 | 58784000 | 3 | AJQ90514.1_778 | RM25_0790 | Cytoplasmic | 0 | CYT | 0.044652 | OTHER | G |
| lcl\|CP010341.1_prot_AJQ90515.1_779 | RM25_RS03885 | Hydrolase | 0 | 2676000 | 4748800 | 3286700 | 3547000 | 4053100 | 1 | 1 | 6 | 6 | 6 | 25,923 | 0,0043431 | 7,6459 | 18312000 | 1 | AJQ90515.1_779 | RM25_0791 | Cytoplasmic | 0 | CYT | 0.111279 | OTHER | |
| lcl\|CP010341.1_prot_AJQ90516.1_780 | RM25_RS03890 | Valine--tRNA ligase | 42674000 | 19550000 | 113930000 | 25598000 | 53594000 | 69719000 | 8 | 8 | 10 | 10 | 10 | 97,657 | 0 | 49,518 | 351600000 | 20 | AJQ90516.1_780 | RM25_0792 | Cytoplasmic | 0 | CYT | 0.182139 | OTHER | J |
| lcl\|CP010341.1_prot_AJQ90517.1_781 | RM25_RS03895 | DSBA-like thioredoxin domain protein | 3095699968 | 4548800000 | 5527000064 | 1196800000 | 2407899904 | 3781100032 | 10 | 7 | 40,9 | 40,9 | 31,4 | 28,404 | 0 | 258,63 | 2,1343E+10 | 97 | AJQ90517.1_781 | RM25_0793 | Unknown | 1 | SPI | 0.424100 | OTHER | |
| lcl\|CP010341.1_prot_AJQ90518.1_782 | RM25_RS03900 | DSBA-like thioredoxin domain protein | 5598799872 | 9577900032 | 11233000448 | 3429499904 | 4555800064 | 8630400000 | 6 | 6 | 49,6 | 40,2 | 40,2 | 28,408 | 0 | 323,31 | 4,614E+10 | 113 | AJQ90518.1_782 | RM25_0794 | Unknown | 1 | SPI | 0.921181 | OTHER | |
| lcl\|CP010341.1_prot_AJQ90520.1_784 | RM25_RS03910 | Folylpolyglutamate synthase folC | 14108000 | 18580000 | 52004000 | 33788000 | 36783000 | 50907000 | 6 | 6 | 15 | 15 | 15 | 52,814 | 0 | 41,69 | 225050000 | 14 | AJQ90520.1_784 | RM25_0796 | Cytoplasmic | 0 | CYT | 0.044938 | OTHER | H |
| lcl\|CP010341.1_prot_AJQ90525.1_789 | RM25_RS03935 | Integral membrane protein TerC | 3696700 | 3817700 | 7844000 | 10106000 | 9996000 | 12073000 | 3 | 3 | 8,6 | 8,6 | 8,6 | 46,008 | 0 | 26,792 | 55233000 | 6 | AJQ90525.1_789 | RM25_0801 | CytoplasmicMembrane | 9 | TMH | 0.964168 | OTHER | P |
| lcl\|CP010341.1_prot_AJQ90528.1_792 | RM25_RS03950 | Hypothetical protein | 215210000 | 195100000 | 603820032 | 412360000 | 503209984 | 773950016 | 20 | 20 | 34,3 | 34,3 | 34,3 | 101,18 | 0 | 323,31 | 3035200000 | 83 | AJQ90528.1_792 | RM25_0804 | Cytoplasmic | 0 | CYT | 0.457438 | OTHER | J |
| lcl\|CP010341.1_prot_AJQ90529.1_793 | RM25_RS03955 | 50S ribosomal protein L21 | 309289984 | 354180000 | 891009984 | 1079000064 | 729160000 | 1246599936 | 4 | 4 | 37,4 | 37,4 | 37,4 | 12,756 | 0 | 70,141 | 5552600000 | 30 | AJQ90529.1_793 | RM25_0805 | Unknown | 0 | CYT | 0.819801 | OTHER | J |
| lcl\|CP010341.1_prot_AJQ90530.1_794 | RM25_RS03960 | 50S ribosomal protein L27 | 134720000 | 185880000 | 429260000 | 335590016 | 360740000 | 438060000 | 5 | 5 | 64,4 | 64,4 | 64,4 | 9,2763 | 0 | 33,495 | 2194100000 | 20 | AJQ90530.1_794 | RM25_0806 | Cytoplasmic | 0 | CYT | 0.094003 | OTHER | J |
| lcl\|CP010341.1_prot_AJQ90531.1_795 | RM25_RS03965 | GTPase obg | 19340000 | 26685000 | 68249000 | 40915000 | 13227000 | 50554000 | 5 | 5 | 12,6 | 12,6 | 12,6 | 57,313 | 0 | 43,628 | 313790000 | 12 | AJQ90531.1_795 | RM25_0807 | Cytoplasmic | 0 | CYT | 0.094936 | OTHER | S |
| lcl\|CP010341.1_prot_AJQ90534.1_798 | RM25_RS03980 | Gamma-glutamyl phosphate reductase | 13498000 | 2108800 | 38294000 | 16858000 | 13889000 | 16090000 | 2 | 2 | 5,6 | 5,6 | 5,6 | 43,614 | 0 | 18,043 | 108600000 | 5 | AJQ90534.1_798 | RM25_0810 | Cytoplasmic | 0 | CYT | 0.055376 | OTHER | E |
| lcl\|CP010341.1_prot_AJQ90541.1_805 | RM25_RS04015 | Hypothetical protein | 17279000 | 27966000 | 34258000 | 13538000 | 23960000 | 30407000 | 1 | 1 | 7,7 | 7,7 | 7,7 | 25,33 | 0 | 57,88 | 160250000 | 7 | AJQ90541.1_805 | RM25_0817 | Unknown | 1 | CYT | 0.959914 | OTHER | |
| lcl\|CP010341.1_prot_AJQ90543.1_807 | RM25_RS04025 | Leucine--tRNA ligase | 30262000 | 33652000 | 106210000 | 30825000 | 36331000 | 56432000 | 7 | 7 | 8,4 | 8,4 | 8,4 | 93,771 | 0 | 97,701 | 306170000 | 22 | AJQ90543.1_807 | RM25_0819 | Cytoplasmic | 0 | CYT | 0.321874 | OTHER | J |
| lcl\|CP010341.1_prot_AJQ90544.1_808 | RM25_RS04030 | Dihydroorotase | 8372800 | 5300000 | 25831000 | 0 | 0 | 6759100 | 3 | 3 | 10,7 | 10,7 | 10,7 | 45,5 | 0 | 26,577 | 46263000 | 2 | AJQ90544.1_808 | RM25_0820 | Cytoplasmic | 0 | CYT | 0.094165 | OTHER | F |
| lcl\|CP010341.1_prot_AJQ90545.1_809 | RM25_RS04035 | Malic enzyme, NAD binding domain protein | 7513800 | 618990 | 15175000 | 13958000 | 13094000 | 13960000 | 2 | 2 | 5,3 | 5,3 | 5,3 | 64,157 | 0 | 12,478 | 72112000 | 4 | AJQ90545.1_809 | RM25_0821 | Cytoplasmic | 0 | CYT | 0.061447 | OTHER | C |
| lcl\|CP010341.1_prot_AJQ90550.1_814 | RM25_RS04060 | 30S ribosomal protein S20 | 0 | 0 | 0 | 0 | 1193400 | 1519200 | 1 | 1 | 12,8 | 12,8 | 12,8 | 9,3957 | 0,0050813 | 6,4309 | 2712600 | 1 | AJQ90550.1_814 | RM25_0826 | Cytoplasmic | 0 | CYT | 0.860127 | OTHER | J |
| lcl\|CP010341.1_prot_AJQ90552.1_816 | RM25_RS04070 | Putative membrane protein | 330340000 | 443910016 | 842769984 | 256010000 | 279609984 | 573710016 | 4 | 4 | 30,1 | 30,1 | 30,1 | 21,558 | 0 | 76,599 | 2908200000 | 32 | AJQ90552.1_816 | RM25_0828 | CytoplasmicMembrane | 2 | SPI | 0.823898 | SP(Sec/SPI) | |
| lcl\|CP010341.1_prot_AJQ90553.1_817 | RM25_RS04075 | Elongation factor 4 | 15801000 | 18521000 | 21151000 | 69481000 | 12233000 | 27050000 | 5 | 5 | 12 | 12 | 12 | 67,058 | 0 | 33,676 | 194050000 | 2 | AJQ90553.1_817 | RM25_0829 | CytoplasmicMembrane | 0 | CYT | 0.061255 | OTHER | M |
| lcl\|CP010341.1_prot_AJQ90556.1_820 | RM25_RS04090 | Inositol monophosphatase | 3601100 | 0 | 6894700 | 4376400 | 11777000 | 14290000 | 1 | 1 | 4,3 | 4,3 | 4,3 | 27,876 | 0,0050916 | 6,448 | 40939000 | 2 | AJQ90556.1_820 | RM25_0832 | Cytoplasmic | 0 | CYT | 0.077614 | OTHER | G |
| lcl\|CP010341.1_prot_AJQ90557.1_821 | RM25_RS04095 | Heat-inducible transcription repressor hrcA | 14625000 | 15623000 | 24848000 | 974000 | 7358000 | 19550000 | 3 | 3 | 14 | 14 | 14 | 36,115 | 0 | 15,979 | 94476000 | 3 | AJQ90557.1_821 | RM25_0833 | Cytoplasmic | 0 | CYT | 0.042421 | OTHER | K |
| lcl\|CP010341.1_prot_AJQ90558.1_822 | RM25_RS04100 | Chaperone protein DnaJ | 5434300 | 6427900 | 43885000 | 6973300 | 33238000 | 46576000 | 4 | 4 | 13,4 | 13,4 | 13,4 | 42,403 | 0 | 34,088 | 199220000 | 6 | AJQ90558.1_822 | RM25_0834 | Cytoplasmic | 0 | CYT | 0.457346 | OTHER | O |
| lcl\|CP010341.1_prot_AJQ90563.1_827 | RM25_RS04125 | Prolyl aminopeptidase | 2014200 | 2166600 | 15677000 | 4572000 | 5457300 | 9060000 | 2 | 2 | 9,4 | 9,4 | 9,4 | 39,037 | 0 | 12,36 | 38948000 | 2 | AJQ90563.1_827 | RM25_0839 | Cytoplasmic | 0 | CYT | 0.154621 | OTHER | S |
| lcl\|CP010341.1_prot_AJQ90568.1_832 | RM25_RS04150 | CBS domain protein | 2563800 | 4849400 | 6079300 | 0 | 0 | 0 | 1 | 1 | 2,8 | 2,8 | 2,8 | 47,868 | 0,0041623 | 6,8941 | 13493000 | 1 | AJQ90568.1_832 | RM25_0844 | CytoplasmicMembrane | 3 | CYT | 0.360026 | SP(Sec/SPI) | S |
| lcl\|CP010341.1_prot_AJQ90574.1_838 | RM25_RS04175 | Protein of hypothetical function DUF488 | 47871000 | 67057000 | 122150000 | 125220000 | 111630000 | 121470000 | 5 | 5 | 48,3 | 48,3 | 48,3 | 13,763 | 0 | 31,473 | 665520000 | 17 | AJQ90574.1_838 | RM25_0850 | CytoplasmicMembrane | 0 | CYT | 0.542384 | OTHER | |
| lcl\|CP010341.1_prot_AJQ90583.1_847 | RM25_RS04215 | 3-isopropylmalate dehydrogenase | 11173000 | 6232800 | 32259000 | 21736000 | 14454000 | 18731000 | 3 | 3 | 13,8 | 13,8 | 13,8 | 37,444 | 0 | 22,215 | 118790000 | 5 | AJQ90583.1_847 | RM25_0859 | Cytoplasmic | 0 | CYT | 0.066064 | OTHER | CE |
| lcl\|CP010341.1_prot_AJQ90584.1_848 | RM25_RS04220 | Branched-chain-amino-acid aminotransferase | 32936000 | 36841000 | 50554000 | 49915000 | 39777000 | 45071000 | 4 | 4 | 15 | 15 | 15 | 39,833 | 0 | 29,687 | 286680000 | 13 | AJQ90584.1_848 | RM25_0860 | Cytoplasmic | 0 | CYT | 0.192482 | OTHER | E |
| lcl\|CP010341.1_prot_AJQ90585.1_849 | RM25_RS04225 | 2-isopropylmalate synthase/homocitrate synthase fami | 3344200 | 4175500 | 8579200 | 12485000 | 13721000 | 17475000 | 2 | 2 | 4,1 | 4,1 | 4,1 | 57,583 | 0 | 13,267 | 69762000 | 5 | AJQ90585.1_849 | RM25_0861 | Cytoplasmic | 0 | CYT | 0.097704 | OTHER | E |
| lcl\|CP010341.1_prot_AJQ90586.1_850 | RM25_RS04230 | FAH family protein | 10320000 | 6234500 | 30700000 | 3711300 | 6206500 | 13789000 | 2 | 2 | 5,3 | 5,3 | 5,3 | 29,599 | 0,0011682 | 10,813 | 71991000 | 3 | AJQ90586.1_850 | RM25_0862 | Cytoplasmic | 0 | CYT | 0.194208 | OTHER | Q |
| lcl\|CP010341.1_prot_AJQ90589.1_853 | RM25_RS04255 | 3-isopropylmalate dehydratase large subunit | 6311700 | 5128100 | 19456000 | 23343000 | 35396000 | 41672000 | 3 | 3 | 7 | 7 | 7 | 49,77 | 0 | 19,129 | 131310000 | 4 | AJQ90589.1_853 | RM25_0867 | Cytoplasmic | 0 | CYT | 0.094731 | OTHER | E |
| lcl\|CP010341.1_prot_AJQ90590.1_854 | RM25_RS04260 | 3-isopropylmalate dehydratase small subunit | 14779000 | 21055000 | 52443000 | 45967000 | 44359000 | 71648000 | 4 | 4 | 22,7 | 22,7 | 22,7 | 22,153 | 0 | 25,865 | 307930000 | 17 | AJQ90590.1_854 | RM25_0868 | Cytoplasmic | 0 | CYT | 0.248403 | OTHER | E |
| lcl\|CP010341.1_prot_AJQ90592.1_856 | RM25_RS04270 | D-alanine--D-alanine ligase | 44245000 | 45283000 | 56597000 | 27062000 | 27439000 | 55306000 | 9 | 9 | 28,8 | 28,8 | 28,8 | 40,437 | 0 | 77,901 | 267400000 | 9 | AJQ90592.1_856 | RM25_0870 | Cytoplasmic | 0 | CYT | 0.065339 | OTHER | |
| lcl\|CP010341.1_prot_AJQ90596.1_860 | RM25_RS04290 | Thiamine-phosphate kinase | 11858000 | 9481700 | 16882000 | 6365800 | 2270600 | 12152000 | 2 | 2 | 9,2 | 9,2 | 9,2 | 33,213 | 0 | 19,376 | 63693000 | 4 | AJQ90596.1_860 | RM25_0874 | Cytoplasmic | 0 | CYT | 0.049849 | OTHER | H |
| lcl\|CP010341.1_prot_AJQ90598.1_862 | RM25_RS04300 | Ribosomal protein S12 methylthiotransferase RimO | 8086900 | 32934000 | 34426000 | 23286000 | 18716000 | 13249000 | 5 | 5 | 15,9 | 15,9 | 15,9 | 51,192 | 0 | 32,904 | 146970000 | 8 | AJQ90598.1_862 | RM25_0876 | Cytoplasmic | 0 | CYT | 0.070140 | OTHER | J |
| lcl\|CP010341.1_prot_AJQ90604.1_868 | RM25_RS04330 | RecA protein | 45045000 | 48638000 | 91393000 | 72020000 | 69375000 | 116390000 | 6 | 6 | 23,7 | 23,7 | 23,7 | 39,289 | 0 | 49,319 | 502410000 | 13 | AJQ90604.1_868 | RM25_0882 | Cytoplasmic | 0 | CYT | 0.068881 | OTHER | L |
| lcl\|CP010341.1_prot_AJQ90608.1_872 | RM25_RS04350 | (Dimethylallyl)adenosine tRNA methylthiotransferase M | 7544100 | 11405000 | 22536000 | 3936500 | 6473000 | 14589000 | 2 | 2 | 6 | 6 | 6 | 55,662 | 0 | 12,606 | 72447000 | 2 | AJQ90608.1_872 | RM25_0886 | Cytoplasmic | 0 | CYT | 0.121978 | OTHER | J |
| lcl\|CP010341.1_prot_AJQ90610.1_874 | RM25_RS04360 | Diaminopimelate epimerase | 4438700 | 4804300 | 15772000 | 10927000 | 12348000 | 11449000 | 1 | 1 | 4,3 | 4,3 | 4,3 | 30,39 | 0,0099108 | 6,0071 | 59740000 | 5 | AJQ90610.1_874 | RM25_0888 | Cytoplasmic | 0 | CYT | 0.132520 | OTHER | E |
| lcl\|CP010341.1_prot_AJQ90613.1_877 | RM25_RS04375 | LexA repressor | 2868900 | 3984500 | 9130900 | 3051600 | 3988300 | 3778900 | 2 | 2 | 10,7 | 10,7 | 10,7 | 30,231 | 0 | 17,667 | 26803000 | 4 | AJQ90613.1_877 | RM25_0891 | Cytoplasmic | 0 | CYT | 0.104176 | OTHER | K |
| lcl\|CP010341.1_prot_AJQ90614.1_878 | RM25_RS04380 | Hypothetical protein | 1565000 | 1558400 | 3305600 | 0 | 7294600 | 10350000 | 1 | 1 | 4,5 | 4,5 | 4,5 | 25,771 | 0,0042965 | 7,532 | 24073000 | 4 | AJQ90614.1_878 | RM25_0892 | Unknown | 1 | CYT | 0.223865 | OTHER | |
| lcl\|CP010341.1_prot_AJQ90615.1_879 | RM25_RS04385 | Transcriptional repressor NrdR | 1055600 | 1510400 | 0 | 2039300 | 1835700 | 2642600 | 1 | 1 | 5,5 | 5,5 | 5,5 | 18,452 | 0,0099305 | 6,0245 | 11182000 | 1 | AJQ90615.1_879 | RM25_0893 | Cytoplasmic | 0 | CYT | 0.047578 | OTHER | K |
| lcl\|CP010341.1_prot_AJQ90616.1_880 | RM25_RS04390 | Vitamin B12-dependent ribonucleotide reductase | 412620000 | 554000000 | 1216400000 | 455529984 | 764400000 | 1176099968 | 27 | 27 | 38,4 | 38,4 | 38,4 | 104,84 | 0 | 323,31 | 5023700000 | 126 | AJQ90616.1_880 | RM25_0894 | Cytoplasmic | 0 | CYT | 0.201347 | OTHER | F |
| lcl\|CP010341.1_prot_AJQ90618.1_882 | RM25_RS04400 | Glycerol-3-phosphate dehydrogenase, anaerobic, A sub | 103910000 | 76233000 | 302729984 | 189160000 | 294660000 | 483620000 | 12 | 12 | 31,9 | 31,9 | 31,9 | 58,83 | 0 | 258,58 | 1661400000 | 38 | AJQ90618.1_882 | RM25_0896 | Cytoplasmic | 0 | CYT | 0.100970 | OTHER | |
| lcl\|CP010341.1_prot_AJQ90619.1_883 | RM25_RS04405 | Anaerobic glycerol-3-phosphate dehydrogenase subuni | 241310000 | 198240000 | 599689984 | 367769984 | 582720000 | 842329984 | 13 | 13 | 39,2 | 39,2 | 39,2 | 45,912 | 0 | 169,44 | 3150900000 | 79 | AJQ90619.1_883 | RM25_0897 | CytoplasmicMembrane | 0 | CYT | 0.065502 | OTHER | |
| lcl\|CP010341.1_prot_AJQ90620.1_884 | RM25_RS04410 | Glycerol-3-phosphate dehydrogenase, anaerobic, C sub | 59406000 | 59479000 | 202460000 | 71896000 | 105250000 | 143300000 | 8 | 8 | 19,9 | 19,9 | 19,9 | 50,664 | 0 | 118,35 | 715220000 | 23 | AJQ90620.1_884 | RM25_0898 | CytoplasmicMembrane | 0 | CYT | 0.189830 | OTHER | |
| lcl\|CP010341.1_prot_AJQ90623.1_887 | RM25_RS04425 | putative phosphoketolase | 7449300 | 2590200 | 43679000 | 2928600 | 6698200 | 8671300 | 5 | 5 | 5,9 | 5,9 | 5,9 | 96,719 | 0 | 32,225 | 73626000 | 4 | AJQ90623.1_887 | RM25_0901 | Unknown | 0 | CYT | 0.312381 | OTHER | G |
| lcl\|CP010341.1_prot_AJQ90625.1_889 | RM25_RS04435 | Acyl-CoA thioesterase II | 5057200 | 6490500 | 18462000 | 8396600 | 10869000 | 11908000 | 1 | 1 | 4,1 | 4,1 | 4,1 | 32,142 | 0 | 25,63 | 61183000 | 4 | AJQ90625.1_889 | RM25_0903 | CytoplasmicMembrane | 0 | CYT | 0.129761 | OTHER | I |
| lcl\|CP010341.1_prot_AJQ90626.1_890 | RM25_RS04440 | Solute-binding lipoprotein | 91060000 | 194700000 | 248020000 | 70297000 | 145850000 | 257700000 | 8 | 8 | 22,2 | 22,2 | 22,2 | 48,757 | 0 | 81,743 | 1056500000 | 32 | AJQ90626.1_890 | RM25_0904 | CytoplasmicMembrane | 1 | SPI | 0.927617 | LIPO(Sec/SPII) | G |
| lcl\|CP010341.1_prot_AJQ90629.1_893 | RM25_RS04450 | RNA polymerase sigma factor SigB | 9075300 | 2124700 | 5351800 | 4013200 | 4444600 | 6259700 | 1 | 1 | 5,2 | 5,2 | 5,2 | 34,422 | 0,0051125 | 6,5953 | 31269000 | 1 | AJQ90629.1_893 | RM25_0907 | Cytoplasmic | 0 | CYT | 0.051572 | OTHER | K |
| lcl\|CP010341.1_prot_AJQ90630.1_894 | RM25_RS04455 | RNA polymerase sigma factor | 95677000 | 121160000 | 238970000 | 192850000 | 217120000 | 275340000 | 11 | 11 | 21,8 | 21,8 | 21,8 | 55,66 | 0 | 78,439 | 1313900000 | 35 | AJQ90630.1_894 | RM25_0908 | Cytoplasmic | 0 | CYT | 0.597669 | OTHER | K |
| lcl\|CP010341.1_prot_AJQ90631.1_895 | RM25_RS04460 | L-lactate dehydrogenase | 4051900 | 0 | 12701000 | 0 | 0 | 5750900 | 2 | 2 | 7,8 | 7,8 | 7,8 | 34 | 0 | 14,547 | 22504000 | 3 | AJQ90631.1_895 | RM25_0909 | Cytoplasmic | 0 | CYT | 0.052421 | OTHER | C |
| lcl\|CP010341.1_prot_AJQ90633.1_897 | RM25_RS04470 | DNA gyrase, B subunit | 2807900 | 2403400 | 28686000 | 10915000 | 20758000 | 22873000 | 4 | 4 | 5,8 | 5,8 | 5,8 | 76,753 | 0 | 23,032 | 95859000 | 4 | AJQ90633.1_897 | RM25_0911 | Cytoplasmic | 0 | CYT | 0.083295 | OTHER | L |
| lcl\|CP010341.1_prot_AJQ90634.1_898 | RM25_RS04475 | 2-dehydropantoate 2-reductase (Precursor) | 48516000 | 35741000 | 127790000 | 55788000 | 50436000 | 47591000 | 8 | 8 | 25,9 | 25,9 | 25,9 | 32,273 | 0 | 51,446 | 405410000 | 18 | AJQ90634.1_898 | RM25_0912 | Cytoplasmic | 0 | CYT | 0.050905 | OTHER | H |
| lcl\|CP010341.1_prot_AJQ90639.1_903 | RM25_RS04500 | Hypothetical protein | 17406000 | 25415000 | 53063000 | 30510000 | 29685000 | 44444000 | 4 | 4 | 16,1 | 16,1 | 16,1 | 46,58 | 0 | 71,939 | 228310000 | 11 | AJQ90639.1_903 | RM25_0917 | Cytoplasmic | 0 | CYT | 0.864957 | OTHER | S |
| lcl\|CP010341.1_prot_AJQ90640.1_904 | RM25_RS04505 | PF13834 domain protein | 184470000 | 229810000 | 281080000 | 179110000 | 178480000 | 172690000 | 3 | 3 | 33,9 | 33,9 | 33,9 | 13,827 | 0 | 60,726 | 1390500000 | 12 | AJQ90640.1_904 | RM25_0918 | Cytoplasmic | 0 | CYT | 0.103263 | OTHER | S |
| lcl\|CP010341.1_prot_AJQ90642.1_906 | RM25_RS04515 | Hypothetical protein | 12822000 | 22615000 | 31528000 | 30153000 | 31231000 | 34442000 | 2 | 2 | 13,2 | 13,2 | 13,2 | 29,649 | 0 | 28,27 | 205720000 | 4 | AJQ90642.1_906 | RM25_0920 | Cytoplasmic | 0 | CYT | 0.195488 | OTHER | S |
| lcl\|CP010341.1_prot_AJQ90644.1_908 | RM25_RS04525 | Trk system potassium uptake protein ceoC | 6488500 | 7052300 | 13626000 | 7703500 | 7440000 | 0 | 1 | 1 | 6,8 | 6,8 | 6,8 | 24,018 | 0,00453 | 9,5677 | 42310000 | 3 | AJQ90644.1_908 | RM25_0922 | CytoplasmicMembrane | 0 | CYT | 0.047288 | OTHER | P |
| lcl\|CP010341.1_prot_AJQ90645.1_909 | RM25_RS04530 | Trk family potassium (K+) transporter, NAD+ binding pr | 0 | 13766000 | 20338000 | 0 | 5416000 | 14239000 | 1 | 1 | 5 | 5 | 5 | 23,805 | 0,0042599 | 7,4291 | 60459000 | 1 | AJQ90645.1_909 | RM25_0923 | CytoplasmicMembrane | 0 | CYT | 0.039759 | OTHER | P |
| lcl\|CP010341.1_prot_AJQ90649.1_913 | RM25_RS04550 | Putative glutathione S-transferase with thioredoxin-like | 71749000 | 83895000 | 290240000 | 265950000 | 277670016 | 382140000 | 11 | 11 | 40,9 | 40,9 | 40,9 | 39,332 | 0 | 104,26 | 1526400000 | 40 | AJQ90649.1_913 | RM25_0927 | Cytoplasmic | 0 | CYT | 0.259982 | OTHER | O |
| lcl\|CP010341.1_prot_AJQ90653.1_917 | RM25_RS04565 | Putative glutathione S-transferase | 1116000 | 1042200 | 2396000 | 2507800 | 6420500 | 11152000 | 2 | 2 | 6,7 | 6,7 | 6,7 | 41,181 | 0 | 14,084 | 24635000 | 3 | AJQ90653.1_917 | RM25_0931 | Cytoplasmic | 0 | CYT | 0.210769 | OTHER | O |
| lcl\|CP010341.1_prot_AJQ90654.1_918 | RM25_RS04570 | Alpha-ketoglutarate transporter, MFS super | 14655000 | 11744000 | 16367000 | 23562000 | 35804000 | 55952000 | 2 | 2 | 6,5 | 6,5 | 6,5 | 50,3 | 0 | 17,736 | 179690000 | 12 | AJQ90654.1_918 | RM25_0932 | CytoplasmicMembrane | 10 | CYT | 0.963217 | OTHER | |
| lcl\|CP010341.1_prot_AJQ90655.1_919 | RM25_RS04575 | Aconitate hydratase 1 | 167490000 | 165650000 | 537150016 | 196610000 | 166460000 | 229610000 | 18 | 18 | 24,9 | 24,9 | 24,9 | 96,389 | 0 | 161,19 | 1585400000 | 49 | AJQ90655.1_919 | RM25_0933 | Cytoplasmic | 0 | CYT | 0.152075 | OTHER | C |
| lcl\|CP010341.1_prot_AJQ90659.1_923 | RM25_RS04595 | 3'-5' exonuclease | 13779000 | 12068000 | 25719000 | 0 | 15992000 | 26091000 | 2 | 2 | 4,8 | 4,8 | 4,8 | 47,372 | 0 | 11,049 | 97839000 | 2 | AJQ90659.1_923 | RM25_0937 | Cytoplasmic | 0 | CYT | 0.116860 | OTHER | J |
| lcl\|CP010341.1_prot_AJQ90661.1_925 | RM25_RS04605 | YbaK/prolyl-tRNA synthetase-associated domain protei | 4376800 | 5314000 | 6750300 | 4502000 | 3562200 | 4434400 | 1 | 1 | 13,2 | 13,2 | 13,2 | 17,848 | 0,0011669 | 10,731 | 31009000 | 1 | AJQ90661.1_925 | RM25_0939 | Cytoplasmic | 0 | CYT | 0.041574 | OTHER | S |
| lcl\|CP010341.1_prot_AJQ90665.1_929 | RM25_RS04625 | Cobalamin biosynthesis protein CbiX | 142930000 | 297329984 | 657800000 | 51536000 | 224400000 | 161190000 | 1 | 1 | 2,5 | 2,5 | 2,5 | 29,502 | 0,0070423 | 6,2429 | 1549100000 | 21 | AJQ90665.1_929 | RM25_0943 | Unknown | 0 | CYT | 0.041012 | OTHER | S |
| lcl\|CP010341.1_prot_AJQ90666.1_930 | RM25_RS04650 | Threonine--tRNA ligase | 86167000 | 79303000 | 314160000 | 75274000 | 127740000 | 223400000 | 14 | 14 | 30,1 | 30,1 | 30,1 | 75,831 | 0 | 281,83 | 983680000 | 49 | AJQ90666.1_930 | RM25_0947 | Cytoplasmic | 0 | CYT | 0.136611 | OTHER | J |
| lcl\|CP010341.1_prot_AJQ90667.1_931 | RM25_RS04655 | Histidine triad domain protein | 1421300 | 1338900 | 0 | 6010700 | 4204500 | 0 | 2 | 2 | 6,5 | 6,5 | 6,5 | 22,077 | 0 | 11,353 | 12975000 | 1 | AJQ90667.1_931 | RM25_0948 | Cytoplasmic | 0 | CYT | 0.215086 | OTHER | FG |
| lcl\|CP010341.1_prot_AJQ90670.1_934 | RM25_RS04670 | Phosphatidylinositol alpha-mannosyltransferase | 11106000 | 10232000 | 24217000 | 18852000 | 27164000 | 32746000 | 4 | 4 | 16 | 16 | 16 | 41,055 | 0 | 26,338 | 131710000 | 5 | AJQ90670.1_934 | RM25_0951 | Unknown | 0 | CYT | 0.052099 | OTHER | M |
| lcl\|CP010341.1_prot_AJQ90672.1_936 | RM25_RS13200 | Hypothetical protein | 691209984 | 1326800000 | 1565600000 | 328060000 | 633660032 | 946019968 | 7 | 7 | 44,5 | 44,5 | 44,5 | 28,05 | 0 | 290,69 | 5799100000 | 53 | AJQ90672.1_936 | RM25_0953 | Cytoplasmic | 1 | CYT | 0.300504 | OTHER | S |

| | | | | | | | | | | | | | | | | | | | | | | | | | | |
|---|---|---|---|---|---|---|---|---|---|---|---|---|---|---|---|---|---|---|---|---|---|---|---|---|---|---|
| lcl\|CP010341.1_prot_AJQ90674.1_938 | RM25_RS04690 | PF05949 family protein | 2935699968 | 5024900096 | 6190700032 | 1720000000 | 2646200064 | 4143000064 | 10 | 10 | 68,9 | 68,9 | 68,9 | 25,862 | 0 | 323,31 | 2,4066E+10 | 106 | AJQ90674.1_938 | RM25_0955 | Cytoplasmic | 1 | TMH | 0.198122 | OTHER | S |
| lcl\|CP010341.1_prot_AJQ90675.1_939 | RM25_RS04695 | Glycine cleavage system H protein | 37201000 | 68597000 | 32075000 | 63616000 | 42644000 | 55385000 | 2 | 2 | 23,6 | 23,6 | 23,6 | 13,3 | 0 | 17,564 | 347500000 | 10 | AJQ90675.1_939 | RM25_0956 | Unknown | 0 | CYT | 0.822280 | OTHER | E |
| lcl\|CP010341.1_prot_AJQ90676.1_940 | RM25_RS04700 | Oxoglutarate dehydrogenase inhibitor | 42115000 | 59743000 | 109460000 | 46270000 | 40198000 | 62057000 | 4 | 4 | 25,9 | 25,9 | 25,9 | 17,331 | 0 | 34,027 | 401640000 | 15 | AJQ90676.1_940 | RM25_0957 | Cytoplasmic | 0 | CYT | 0.150930 | OTHER | T |
| lcl\|CP010341.1_prot_AJQ90678.1_942 | RM25_RS04710 | Protein of hypothetical function | 3415600 | 3129800 | 8170800 | 8166200 | 4632300 | 5701200 | 1 | 1 | 5,9 | 5,9 | 5,9 | 16,56 | 0,0041408 | 6,8043 | 39089000 | 1 | AJQ90678.1_942 | RM25_0959 | Cytoplasmic | 0 | CYT | 0.029262 | OTHER | |
| lcl\|CP010341.1_prot_AJQ90680.1_944 | RM25_RS04720 | Glycine dehydrogenase | 5453700 | 2796400 | 46303000 | 20567000 | 12844000 | 19138000 | 4 | 4 | 6,4 | 6,4 | 6,4 | 103,5 | 0 | 25,588 | 107100000 | 3 | AJQ90680.1_944 | RM25_0961 | Cytoplasmic | 0 | CYT | 0.086590 | OTHER | E |
| lcl\|CP010341.1_prot_AJQ90682.1_946 | RM25_RS04730 | YhgE/Pip C-terminal domain protein | 812940032 | 881520000 | 1545900032 | 113400000 | 156040000 | 447430016 | 14 | 14 | 30,7 | 30,7 | 30,7 | 77,294 | 0 | 323,31 | 4071700000 | 58 | AJQ90682.1_946 | RM25_0963 | CytoplasmicMembrane | 6 | CYT | 0.829268 | OTHER | S |
| lcl\|CP010341.1_prot_AJQ90683.1_947 | RM25_RS04735 | Hypothetical protein | 5464100 | 2751400 | 5078600 | 0 | 0 | 4698100 | 2 | 2 | 14,1 | 14,1 | 14,1 | 28,959 | 0 | 13,939 | 17992000 | 1 | AJQ90683.1_947 | RM25_0964 | Cytoplasmic | 0 | CYT | 0.046228 | OTHER | |
| lcl\|CP010341.1_prot_AJQ90686.1_950 | RM25_RS04750 | Cobinamide kinase/cobinamide phosphate guanylyltrar | 17327000 | 7734300 | 19628000 | 43844000 | 39986000 | 70140000 | 4 | 4 | 25,1 | 25,1 | 25,1 | 23,919 | 0 | 32,314 | 221870000 | 10 | AJQ90686.1_950 | RM25_0967 | Unknown | 0 | CYT | 0.103898 | OTHER | H |
| lcl\|CP010341.1_prot_AJQ90687.1_951 | RM25_RS04755 | Cobyrinic acid a,c-diamide synthase | 6276400 | 5037000 | 16840000 | 4637100 | 16037000 | 21054000 | 5 | 5 | 7,2 | 7,2 | 7,2 | 90,565 | 0 | 31,919 | 74436000 | 6 | AJQ90687.1_951 | RM25_0968 | Cytoplasmic | 0 | CYT | 0.082910 | OTHER | H |
| lcl\|CP010341.1_prot_AJQ90689.1_953 | RM25_RS04765 | Adenosylcobyric acid synthase (glutamine-hydrolyzing) | 16879000 | 25799000 | 64721000 | 18328000 | 14495000 | 25787000 | 5 | 5 | 11,9 | 11,9 | 11,9 | 51,726 | 0 | 29,848 | 174300000 | 7 | AJQ90689.1_953 | RM25_0970 | Cytoplasmic | 0 | CYT | 0.081341 | OTHER | H |
| lcl\|CP010341.1_prot_AJQ90695.1_959 | RM25_RS04795 | Trehalose-6-phosphate phosphatase | 181660000 | 272140000 | 191610000 | 155170000 | 129530000 | 317769984 | 2 | 2 | 8,4 | 8,4 | 8,4 | 29,958 | 0 | 14,269 | 1414200000 | 7 | AJQ90695.1_959 | RM25_0976 | Cytoplasmic | 0 | CYT | 0.066376 | OTHER | G |
| lcl\|CP010341.1_prot_AJQ90703.1_967 | RM25_RS11835 | Hypothetical protein | 0 | 0 | 0 | 22483000 | 4756000 | 0 | 1 | 1 | 8 | 8 | 8 | 20,372 | 0,0044248 | 8,3908 | 40788000 | 2 | AJQ90703.1_967 | RM25_0985 | Unknown | 0 | CYT | 0.258589 | OTHER | |
| lcl\|CP010341.1_prot_AJQ90704.1_968 | RM25_RS04860 | Phosphofructokinase (Precursor) | 170730000 | 172690000 | 491550016 | 174500000 | 169730000 | 208180000 | 9 | 9 | 23,5 | 23,5 | 23,5 | 43,245 | 0 | 78,76 | 1476900000 | 40 | AJQ90704.1_968 | RM25_0987 | Cytoplasmic | 0 | SPI | 0.082195 | SP(Sec/SPI) | G |
| lcl\|CP010341.1_prot_AJQ90705.1_969 | RM25_RS11840 | Hypothetical protein | 139790000 | 209060000 | 317160000 | 135220000 | 246900000 | 351609984 | 4 | 4 | 27,9 | 27,9 | 27,9 | 14,583 | 0 | 70,089 | 1517200000 | 18 | AJQ90705.1_969 | RM25_0988 | Cytoplasmic | 1 | TMH | 0.451581 | OTHER | S |
| lcl\|CP010341.1_prot_AJQ90706.1_970 | RM25_RS04870 | Sua5/YciO/YrdC/YwlC family protein | 19063000 | 1922200 | 58420000 | 31511000 | 41707000 | 7776800 | 2 | 2 | 16,6 | 16,6 | 16,6 | 22,663 | 0 | 33,412 | 201600000 | 5 | AJQ90706.1_970 | RM25_0989 | Cytoplasmic | 0 | CYT | 0.080897 | OTHER | J |
| lcl\|CP010341.1_prot_AJQ90708.1_972 | RM25_RS04880 | 1D-myo-inositol 2-acetamido-2-deoxy-alpha-D-glucopyr | 0 | 5154400 | 15506000 | 37454000 | 49522000 | 12343000 | 1 | 1 | 3 | 3 | 3 | 28,886 | 0,0043526 | 7,8403 | 119980000 | 3 | AJQ90708.1_972 | RM25_0991 | Cytoplasmic | 0 | CYT | 0.131530 | OTHER | S |
| lcl\|CP010341.1_prot_AJQ90709.1_973 | RM25_RS04885 | Proteasome assembly chaperones 2 | 12049000 | 16495000 | 59117000 | 22248000 | 47016000 | 72864000 | 4 | 4 | 16,8 | 16,8 | 16,8 | 38,285 | 0 | 28,17 | 258990000 | 14 | AJQ90709.1_973 | RM25_0992 | Cytoplasmic | 0 | CYT | 0.510918 | OTHER | S |
| lcl\|CP010341.1_prot_AJQ90710.1_974 | RM25_RS04890 | ABC-type uncharacterized transport system, ATPase coi | 1443700 | 0 | 4683900 | 0 | 1170400 | 0 | 1 | 1 | 4,6 | 4,6 | 4,6 | 28,668 | 0,008008 | 6,1719 | 7298100 | 0 | AJQ90710.1_974 | RM25_0993 | CytoplasmicMembrane | 0 | CYT | 0.042275 | OTHER | S |
| lcl\|CP010341.1_prot_AJQ90712.1_976 | RM25_RS04900 | ABC transporter substrate binding protein | 3652300 | 2286600 | 10965000 | 5424600 | 0 | 9608800 | 3 | 3 | 11,7 | 11,7 | 11,7 | 33,504 | 0 | 24,44 | 31937000 | 6 | AJQ90712.1_976 | RM25_0995 | Unknown | 0 | SPII | 0.793800 | LIPO(Sec/SPII) | |
| lcl\|CP010341.1_prot_AJQ90713.1_977 | RM25_RS04905 | Phosphoglycerate mutase | 10778000 | 15913000 | 40556000 | 24113000 | 17700000 | 27540000 | 4 | 4 | 27,1 | 27,1 | 27,1 | 22,905 | 0 | 36,408 | 161880000 | 9 | AJQ90713.1_977 | RM25_0996 | Unknown | 0 | CYT | 0.191908 | OTHER | |
| lcl\|CP010341.1_prot_AJQ90714.1_978 | RM25_RS04910 | Glutamine--tRNA ligase | 36611000 | 47434000 | 77134000 | 8185200 | 30720000 | 63030000 | 6 | 6 | 15,8 | 15,8 | 15,8 | 64,243 | 0 | 147,14 | 263110000 | 10 | AJQ90714.1_978 | RM25_0997 | Cytoplasmic | 0 | CYT | 0.186540 | OTHER | |
| lcl\|CP010341.1_prot_AJQ90717.1_981 | RM25_RS04925 | Anthranilate synthase component I | 13183000 | 5720900 | 29980000 | 19297000 | 4662200 | 37906000 | 4 | 4 | 10,2 | 10,2 | 10,2 | 55,276 | 0 | 39,991 | 122950000 | 2 | AJQ90717.1_981 | RM25_1000 | Cytoplasmic | 0 | CYT | 0.072665 | OTHER | E |
| lcl\|CP010341.1_prot_AJQ90718.1_982 | RM25_RS04930 | Indole-3-glycerol phosphate synthase | 21310000 | 39228000 | 95894000 | 22210000 | 45927000 | 63040000 | 6 | 6 | 28,3 | 28,3 | 28,3 | 30,352 | 0 | 49,862 | 302600000 | 10 | AJQ90718.1_982 | RM25_1001 | Cytoplasmic | 0 | CYT | 0.042930 | OTHER | E |
| lcl\|CP010341.1_prot_AJQ90719.1_983 | RM25_RS04935 | Tryptophan synthase beta chain | 29339000 | 20484000 | 72212000 | 12455000 | 16367000 | 29579000 | 3 | 3 | 10,8 | 10,8 | 10,8 | 44,74 | 0 | 34,684 | 187840000 | 8 | AJQ90719.1_983 | RM25_1002 | Cytoplasmic | 0 | CYT | 0.084128 | OTHER | E |
| lcl\|CP010341.1_prot_AJQ90720.1_984 | RM25_RS04940 | Tryptophan synthase alpha chain | 25740000 | 23584000 | 95454000 | 29365000 | 51854000 | 73379000 | 5 | 5 | 19,8 | 19,8 | 19,8 | 31,267 | 0 | 33,98 | 312750000 | 7 | AJQ90720.1_984 | RM25_1003 | Cytoplasmic | 0 | CYT | 0.076670 | OTHER | E |
| lcl\|CP010341.1_prot_AJQ90722.1_986 | RM25_RS04950 | Ferredoxin-dependent glutamate synthase NADPH large | 22558000 | 2646700 | 19273000 | 10303000 | 23664000 | 24566000 | 8 | 8 | 8,1 | 8,1 | 8,1 | 164,11 | 0 | 48,053 | 103010000 | 9 | AJQ90722.1_986 | RM25_1005 | CytoplasmicMembrane | 0 | CYT | 0.089727 | OTHER | E |
| lcl\|CP010341.1_prot_AJQ90725.1_989 | RM25_RS04965 | DNA polymerase III, alpha subunit | 4610900 | 6500300 | 17929000 | 13691000 | 9679300 | 18695000 | 2 | 2 | 2,2 | 2,2 | 2,2 | 131,75 | 0 | 11,866 | 86876000 | 0 | AJQ90725.1_989 | RM25_1008 | Cytoplasmic | 0 | CYT | 0.100809 | OTHER | L |
| lcl\|CP010341.1_prot_AJQ90726.1_990 | RM25_RS11845 | Putative membrane protein | 4320800 | 7976100 | 7534500 | 0 | 9068900 | 17102000 | 1 | 1 | 7,1 | 7,1 | 7,1 | 25,004 | 0,0045662 | 10,121 | 48251000 | 3 | AJQ90726.1_990 | RM25_1009 | CytoplasmicMembrane | 4 | CYT | 0.950105 | OTHER | - |
| lcl\|CP010341.1_prot_AJQ90730.1_994 | RM25_RS04985 | Histidinol dehydrogenase | 66805000 | 54522000 | 180190000 | 91719000 | 97969000 | 115280000 | 8 | 8 | 18,8 | 18,8 | 18,8 | 46,278 | 0 | 85,495 | 650930000 | 30 | AJQ90730.1_994 | RM25_1013 | Cytoplasmic | 0 | CYT | 0.038840 | OTHER | E |
| lcl\|CP010341.1_prot_AJQ90733.1_997 | RM25_RS05000 | Imidazole glycerol phosphate synthase, glutamine amic | 5668800 | 20335000 | 15564000 | 8478200 | 22870000 | 37880000 | 2 | 2 | 16,4 | 16,4 | 16,4 | 23,619 | 0 | 17,752 | 117420000 | 4 | AJQ90733.1_997 | RM25_1016 | Cytoplasmic | 0 | CYT | 0.060795 | OTHER | E |
| lcl\|CP010341.1_prot_AJQ90734.1_998 | RM25_RS05005 | Phosphoribosyl isomerase A | 44827000 | 72855000 | 119070000 | 72831000 | 60172000 | 84437000 | 7 | 7 | 42,3 | 42,3 | 42,3 | 25,174 | 0 | 64,953 | 508230000 | 28 | AJQ90734.1_998 | RM25_1017 | Cytoplasmic | 0 | CYT | 0.041965 | OTHER | E |
| lcl\|CP010341.1_prot_AJQ90735.1_999 | RM25_RS05010 | Mycothiol acetyltransferase | 13214000 | 11438000 | 25739000 | 0 | 0 | 5199000 | 2 | 2 | 8,7 | 8,7 | 8,7 | 38,345 | 0 | 15,574 | 55590000 | 3 | AJQ90735.1_999 | RM25_1018 | CytoplasmicMembrane | 0 | CYT | 0.140523 | OTHER | |
| lcl\|CP010341.1_prot_AJQ90736.1_1000 | RM25_RS05015 | putative transcriptional regulatory protein | 121360000 | 172670000 | 159070000 | 93430000 | 143530000 | 99395000 | 5 | 5 | 36,8 | 36,8 | 36,8 | 28,994 | 0 | 133,93 | 907170000 | 28 | AJQ90736.1_1000 | RM25_1019 | Cytoplasmic | 0 | CYT | 0.317459 | OTHER | K |
| lcl\|CP010341.1_prot_AJQ90740.1_1004 | RM25_RS11860 | Preprotein translocase, YajC subunit | 32030000 | 26245000 | 57085000 | 33019000 | 45860000 | 52316000 | 2 | 2 | 18,6 | 18,6 | 18,6 | 17,869 | 0 | 36,008 | 262160000 | 11 | AJQ90740.1_1004 | RM25_1023 | Cytoplasmic | 1 | TMH | 0.954134 | OTHER | U |
| lcl\|CP010341.1_prot_AJQ90741.1_1005 | RM25_RS05040 | Protein translocase subunit SecD | 649710016 | 1366800000 | 1447699968 | 286880000 | 726220032 | 975960000 | 10 | 10 | 27,1 | 27,1 | 27,1 | 59,862 | 0 | 246,71 | 5607700000 | 46 | AJQ90741.1_1005 | RM25_1024 | CytoplasmicMembrane | 6 | TMH | 0.923646 | OTHER | U |
| lcl\|CP010341.1_prot_AJQ90742.1_1006 | RM25_RS05045 | Protein translocase subunit SecF | 35184000 | 35016000 | 98717000 | 57993000 | 58683000 | 103310000 | 3 | 3 | 15,7 | 15,7 | 15,7 | 44,402 | 0 | 72,115 | 422300000 | 12 | AJQ90742.1_1006 | RM25_1025 | CytoplasmicMembrane | 6 | CYT | 0.919590 | OTHER | U |
| lcl\|CP010341.1_prot_AJQ90743.1_1007 | RM25_RS05050 | Adenine phosphoribosyltransferase | 14438000 | 16736000 | 27002000 | 8446100 | 6575100 | 12165000 | 2 | 2 | 20,7 | 20,7 | 20,7 | 19,052 | 0 | 25,014 | 91844000 | 10 | AJQ90743.1_1007 | RM25_1026 | Cytoplasmic | 0 | CYT | 0.042359 | OTHER | |
| lcl\|CP010341.1_prot_AJQ90744.1_1008 | RM25_RS05055 | GTP diphosphokinase | 7536700 | 7632900 | 30999000 | 8466900 | 7769400 | 9649800 | 3 | 3 | 4,3 | 4,3 | 4,3 | 87,15 | 0 | 18,854 | 74225000 | 8 | AJQ90744.1_1008 | RM25_1027 | Cytoplasmic | 0 | CYT | 0.088551 | OTHER | KT |
| lcl\|CP010341.1_prot_AJQ90745.1_1009 | RM25_RS05060 | Conserved alanine and arginine rich protein | 121990000 | 106620000 | 313849984 | 202620000 | 219890000 | 323140000 | 7 | 7 | 18,7 | 18,7 | 18,7 | 43,103 | 0 | 52,163 | 1458200000 | 29 | AJQ90745.1_1009 | RM25_1028 | Cytoplasmic | 0 | CYT | 0.070755 | OTHER | D |
| lcl\|CP010341.1_prot_AJQ90746.1_1010 | RM25_RS05065 | Metallo-beta-lactamase family protein | 7496900 | 9563900 | 21508000 | 0 | 11157000 | 0 | 1 | 1 | 3,8 | 3,8 | 3,8 | 25,568 | 0,0041973 | 7,1455 | 59320000 | 5 | AJQ90746.1_1010 | RM25_1029 | Cytoplasmic | 0 | CYT | 0.089356 | OTHER | S |
| lcl\|CP010341.1_prot_AJQ90747.1_1011 | RM25_RS05070 | Histidine--tRNA ligase | 91385000 | 68159000 | 185660000 | 106170000 | 93977000 | 186790000 | 12 | 12 | 36,3 | 36,3 | 36,3 | 48,897 | 0 | 142,03 | 812820000 | 45 | AJQ90747.1_1011 | RM25_1030 | Cytoplasmic | 0 | CYT | 0.074074 | OTHER | J |
| lcl\|CP010341.1_prot_AJQ90749.1_1013 | RM25_RS05080 | Hypothetical protein | 85719000 | 64982000 | 310990016 | 156380000 | 200340000 | 255280000 | 6 | 6 | 19,9 | 19,9 | 19,9 | 39,422 | 0 | 102,56 | 1175600000 | 24 | AJQ90749.1_1013 | RM25_1032 | Cytoplasmic | 0 | CYT | 0.085503 | OTHER | |
| lcl\|CP010341.1_prot_AJQ90750.1_1014 | RM25_RS05085 | Aspartate--tRNA ligase | 615129984 | 1179299968 | 1569299968 | 198400000 | 498750016 | 988710016 | 11 | 11 | 22,3 | 22,3 | 22,3 | 67,366 | 0 | 159,48 | 5228100000 | 50 | AJQ90750.1_1014 | RM25_1033 | Cytoplasmic | 0 | CYT | 0.119452 | OTHER | J |
| lcl\|CP010341.1_prot_AJQ90751.1_1015 | RM25_RS05090 | Recombination factor protein RarA | 0 | 1263300 | 4532600 | 0 | 0 | 0 | 1 | 1 | 3,4 | 3,4 | 3,4 | 50,213 | 0,0070493 | 6,2601 | 8578300 | 1 | AJQ90751.1_1015 | RM25_1034 | Unknown | 0 | CYT | 0.071135 | OTHER | L |
| lcl\|CP010341.1_prot_AJQ90752.1_1016 | RM25_RS05095 | Lactate/malate dehydrogenase, NAD binding domain pı | 11188000 | 13368000 | 29950000 | 0 | 9295300 | 11221000 | 2 | 2 | 9,8 | 9,8 | 9,8 | 34,973 | 0 | 22,15 | 76968000 | 4 | AJQ90752.1_1016 | RM25_1035 | Cytoplasmic | 0 | CYT | 0.068835 | OTHER | |
| lcl\|CP010341.1_prot_AJQ90753.1_1017 | RM25_RS05100 | Alanine--tRNA ligase | 227530000 | 261190000 | 651369984 | 321780000 | 302520000 | 460729984 | 28 | 28 | 40,6 | 40,6 | 40,6 | 97,098 | 0 | 282,33 | 2402600000 | 86 | AJQ90753.1_1017 | RM25_1036 | Cytoplasmic | 0 | CYT | 0.095415 | OTHER | J |
| lcl\|CP010341.1_prot_AJQ90754.1_1018 | RM25_RS05105 | Putative crossover junction endodeoxyribonuclease | 1760900 | 0 | 6308600 | 6860500 | 9647500 | 7308200 | 1 | 1 | 13,8 | 13,8 | 13,8 | 19,575 | 0 | 12,84 | 36181000 | 4 | AJQ90754.1_1018 | RM25_1037 | Cytoplasmic | 0 | CYT | 0.087395 | OTHER | L |
| lcl\|CP010341.1_prot_AJQ90755.1_1019 | RM25_RS05110 | Amino deoxychorismate lyase | 738700032 | 1287800064 | 1208400000 | 195380000 | 622300032 | 758110016 | 14 | 14 | 51 | 51 | 51 | 43,845 | 0 | 323,31 | 4945900000 | 72 | AJQ90755.1_1019 | RM25_1038 | Unknown | 1 | CYT | 0.922792 | OTHER | S |
| lcl\|CP010341.1_prot_AJQ90756.1_1020 | RM25_RS05115 | Chorismate synthase | 20297000 | 11953000 | 60742000 | 29154000 | 43264000 | 50392000 | 5 | 5 | 18,2 | 18,2 | 18,2 | 42,724 | 0 | 35,246 | 243850000 | 11 | AJQ90756.1_1020 | RM25_1039 | Cytoplasmic | 0 | CYT | 0.056728 | OTHER | E |
| lcl\|CP010341.1_prot_AJQ90757.1_1021 | RM25_RS05120 | 3-dehydroquinate synthase | 8768300 | 11015000 | 24327000 | 26771000 | 25444000 | 33154000 | 4 | 4 | 8,7 | 8,7 | 8,7 | 58,38 | 0 | 21,523 | 154560000 | 5 | AJQ90757.1_1021 | RM25_1040 | Cytoplasmic | 0 | CYT | 0.054230 | OTHER | E |
| lcl\|CP010341.1_prot_AJQ90758.1_1022 | RM25_RS05125 | Translation elongation factor P | 141380000 | 210610000 | 630870016 | 374169984 | 448140000 | 453720000 | 4 | 4 | 23,8 | 23,8 | 23,8 | 21,312 | 0 | 106,35 | 2512700000 | 32 | AJQ90758.1_1022 | RM25_1041 | Cytoplasmic | 0 | CYT | 0.862099 | OTHER | |
| lcl\|CP010341.1_prot_AJQ90759.1_1023 | RM25_RS05130 | N utilization substance protein B-like protein | 0 | 0 | 13199000 | 12736000 | 16112000 | 4271200 | 2 | 2 | 13,5 | 13,5 | 13,5 | 22,711 | 0 | 15,027 | 46318000 | 8 | AJQ90759.1_1023 | RM25_1042 | Cytoplasmic | 0 | CYT | 0.090847 | OTHER | |
| lcl\|CP010341.1_prot_AJQ90760.1_1024 | RM25_RS05135 | Carbamoyl-phosphate synthase small chain | 42544000 | 28806000 | 124500000 | 41863000 | 78118000 | 95933000 | 5 | 5 | 18,8 | 18,8 | 18,8 | 42,508 | 0 | 63,423 | 437180000 | 13 | AJQ90760.1_1024 | RM25_1043 | Unknown | 0 | CYT | 0.121468 | OTHER | F |
| lcl\|CP010341.1_prot_AJQ90761.1_1025 | RM25_RS05140 | Carbamoyl-phosphate synthase large chain | 184100000 | 233760000 | 456569984 | 159910000 | 179690000 | 456460000 | 22 | 22 | 24,7 | 24,7 | 24,7 | 120,75 | 0 | 209,21 | 1762500000 | 61 | AJQ90761.1_1025 | RM25_1044 | Cytoplasmic | 0 | SPI | 0.067376 | OTHER | F |
| lcl\|CP010341.1_prot_AJQ90762.1_1026 | RM25_RS05145 | Dihydroorotate dehydrogenase (quinone) | 6888500 | 2006200 | 12770000 | 2474800 | 3282200 | 21607000 | 2 | 2 | 6 | 6 | 6 | 38,321 | 0 | 11,194 | 54198000 | 1 | AJQ90762.1_1026 | RM25_1045 | CytoplasmicMembrane | 0 | CYT | 0.069922 | OTHER | F |
| lcl\|CP010341.1_prot_AJQ90763.1_1027 | RM25_RS05150 | Orotidine 5'-phosphate decarboxylase | 5006300 | 8570200 | 22022000 | 0 | 15599000 | 17044000 | 2 | 2 | 7,1 | 7,1 | 7,1 | 28,041 | 0 | 13,677 | 71681000 | 5 | AJQ90763.1_1027 | RM25_1046 | Cytoplasmic | 0 | CYT | 0.058039 | OTHER | F |
| lcl\|CP010341.1_prot_AJQ90765.1_1029 | RM25_RS05160 | Guanylate kinase | 5928400 | 10328000 | 19335000 | 9705100 | 11830000 | 0 | 1 | 1 | 7,8 | 7,8 | 7,8 | 23,31 | 0,0045249 | 9,5645 | 69000000 | 3 | AJQ90765.1_1029 | RM25_1048 | Cytoplasmic | 0 | CYT | 0.089671 | OTHER | F |
| lcl\|CP010341.1_prot_AJQ90766.1_1030 | RM25_RS05165 | DNA-directed RNA polymerase, omega subunit | 35020000 | 23268000 | 72203000 | 60621000 | 35468000 | 112500000 | 3 | 3 | 34,4 | 34,4 | 34,4 | 13,46 | 0 | 33,101 | 373550000 | 14 | AJQ90766.1_1030 | RM25_1049 | Cytoplasmic | 0 | CYT | 0.316019 | OTHER | K |
| lcl\|CP010341.1_prot_AJQ90767.1_1031 | RM25_RS05170 | S-adenosylmethionine synthase | 33912000 | 49656000 | 162390000 | 17430000 | 44466000 | 99305000 | 9 | 9 | 28 | 28 | 28 | 42,781 | 0 | 70,255 | 434900000 | 22 | AJQ90767.1_1031 | RM25_1050 | Cytoplasmic | 0 | CYT | 0.089125 | OTHER | H |
| lcl\|CP010341.1_prot_AJQ90769.1_1033 | RM25_RS05180 | Methionyl-tRNA formyltransferase | 9410400 | 13121000 | 6402300 | 0 | 22223000 | 24497000 | 2 | 2 | 9,5 | 9,5 | 9,5 | 33,349 | 0 | 15,294 | 75654000 | 6 | AJQ90769.1_1033 | RM25_1052 | Cytoplasmic | 0 | CYT | 0.049906 | OTHER | J |
| lcl\|CP010341.1_prot_AJQ90770.1_1034 | RM25_RS05185 | Sun protein | 3506600 | 4149000 | 7893200 | 6600300 | 7979600 | 7740100 | 1 | 1 | 2,9 | 2,9 | 2,9 | 53,559 | 0,0043478 | 7,6584 | 41453000 | 3 | AJQ90770.1_1034 | RM25_1053 | Cytoplasmic | 0 | CYT | 0.108629 | OTHER | JK |
| lcl\|CP010341.1_prot_AJQ90772.1_1036 | RM25_RS05195 | Aminopeptidase C | 15735000 | 2608600 | 66168000 | 17454000 | 23422000 | 21446000 | 5 | 5 | 13,3 | 13,3 | 13,3 | 50,013 | 0 | 44,117 | 154110000 | 14 | AJQ90772.1_1036 | RM25_1055 | Cytoplasmic | 0 | CYT | 0.164960 | OTHER | E |
| lcl\|CP010341.1_prot_AJQ90774.1_1038 | RM25_RS05205 | Methionine synthase | 14727000 | 20727000 | 47212000 | 14539000 | 19783000 | 36124000 | 6 | 6 | 5,8 | 5,8 | 5,8 | 124,89 | 0 | 46,916 | 162620000 | 10 | AJQ90774.1_1038 | RM25_1057 | Cytoplasmic | 0 | CYT | 0.061351 | OTHER | |
| lcl\|CP010341.1_prot_AJQ90775.1_1039 | RM25_RS05210 | Signal peptide peptidase SppA | 280580000 | 501630016 | 607699968 | 205120000 | 154300000 | 412689984 | 12 | 12 | 34,9 | 34,9 | 34,9 | 43,195 | 0 | 103,91 | 2311500000 | 43 | AJQ90775.1_1039 | RM25_1058 | Unknown | 1 | CYT | 0.478456 | OTHER | |
| lcl\|CP010341.1_prot_AJQ90776.1_1040 | RM25_RS05215 | Hydrolase | 2100400 | 2607000 | 3278500 | 4040800 | 3724400 | 2882100 | 1 | 1 | 9,1 | 9,1 | 9,1 | 24,775 | 0,0045872 | 10,224 | 18633000 | 1 | AJQ90776.1_1040 | RM25_1059 | Cytoplasmic | 0 | CYT | 0.070475 | OTHER | S |
| lcl\|CP010341.1_prot_AJQ90779.1_1043 | RM25_RS05230 | Proteasome subunit alpha | 1227800 | 616570 | 3022600 | 0 | 0 | 2177400 | 1 | 1 | 5 | 5 | 5 | 25,983 | 0,0041885 | 7,0132 | 7044400 | 2 | AJQ90779.1_1043 | RM25_1062 | Cytoplasmic | 0 | CYT | 0.067696 | OTHER | O |
| lcl\|CP010341.1_prot_AJQ90780.1_1044 | RM25_RS05235 | Prephenate dehydrogenase | 3181000 | 3945400 | 7296000 | 5006900 | 4385900 | 8215800 | 1 | 1 | 4,7 | 4,7 | 4,7 | 38,625 | 0,0045506 | 9,9953 | 32031000 | 2 | AJQ90780.1_1044 | RM25_1063 | Unknown | 0 | CYT | 0.048050 | OTHER | E |
| lcl\|CP010341.1_prot_AJQ90781.1_1045 | RM25_RS05240 | Cytidylate kinase | 16835000 | 30914000 | 30446000 | 17064000 | 26863000 | 18281000 | 3 | 3 | 17 | 17 | 17 | 24,303 | 0 | 21,729 | 154810000 | 10 | AJQ90781.1_1045 | RM25_1064 | Cytoplasmic | 0 | CYT | 0.054611 | OTHER | |
| lcl\|CP010341.1_prot_AJQ90782.1_1046 | RM25_RS05245 | GTPase Der | 47457000 | 40966000 | 113230000 | 51663000 | 85361000 | 95640000 | 7 | 7 | 21,6 | 21,6 | 21,6 | 48,832 | 0 | 103,72 | 476790000 | 26 | AJQ90782.1_1046 | RM25_1065 | CytoplasmicMembrane | 0 | CYT | 0.054668 | OTHER | S |
| lcl\|CP010341.1_prot_AJQ90790.1_1054 | RM25_RS05290 | S-ribosylhomocysteine lyase | 12855000 | 15370000 | 38508000 | 19136000 | 38182000 | 34422000 | 2 | 2 | 16,2 | 16,2 | 16,2 | 17,213 | 0 | 11,58 | 169950000 | 6 | AJQ90790.1_1054 | RM25_1073 | Cytoplasmic | 0 | CYT | 0.130950 | OTHER | |
| lcl\|CP010341.1_prot_AJQ90791.1_1055 | RM25_RS05295 | Antiseptic resistance protein | 1547500 | 1080400 | 0 | 0 | 1823700 | 4852600 | 1 | 1 | 3 | 3 | 3 | 54,511 | 0,0042689 | 7,4895 | 9304300 | 1 | AJQ90791.1_1055 | RM25_1074 | CytoplasmicMembrane | 13 | TMH | 0.967351 | OTHER | |
| lcl\|CP010341.1_prot_AJQ90793.1_1057 | RM25_RS05305 | Putative sulfonate/nitrate/taurine transport system sub | 17417000 | 18753000 | 33316000 | 0 | 22102000 | 42684000 | 2 | 2 | 7,5 | 7,5 | 7,5 | 35,362 | 0 | 19,584 | 134270000 | 10 | AJQ90793.1_1057 | RM25_1076 | Unknown | 0 | SPII | 0.181707 | LIPO(Sec/SPII) | P |
| lcl\|CP010341.1_prot_AJQ90796.1_1060 | RM25_RS05320 | AsnC-family transcriptional regulatory protein | 1604200 | 1646200 | 5096100 | 5477700 | 3803700 | 5200700 | 1 | 1 | 8,8 | 8,8 | 8,8 | 14,862 | 0,0042781 | 7,5298 | 23997000 | 2 | AJQ90796.1_1060 | RM25_1079 | Cytoplasmic | 0 | CYT | 0.037438 | OTHER | K |
| lcl\|CP010341.1_prot_AJQ90797.1_1061 | RM25_RS05325 | Apolipoprotein N-acyltransferase | 8713200 | 15059000 | 20141000 | 0 | 9466000 | 24402000 | 2 | 2 | 4,6 | 4,6 | 4,6 | 57,416 | 0 | 11,175 | 80112000 | 2 | AJQ90797.1_1061 | RM25_1080 | CytoplasmicMembrane | 7 | TMH | 0.815818 | OTHER | M |
| lcl\|CP010341.1_prot_AJQ90798.1_1062 | RM25_RS05330 | Polyprenol-phosphate-mannose synthase domain 1 | 148330000 | 134120000 | 264830000 | 166820000 | 138640000 | 277060000 | 8 | 8 | 33,2 | 33,2 | 33,2 | 29,729 | 0 | 65,164 | 1286700000 | 39 | AJQ90798.1_1062 | RM25_1081 | Cytoplasmic | 0 | CYT | 0.067621 | OTHER | S |
| lcl\|CP010341.1_prot_AJQ90799.1_1063 | RM25_RS05335 | Putative membrane-associated GTPase | 11309000 | 12347000 | 28824000 | 21797000 | 12600000 | 43066000 | 2 | 2 | 22,3 | 22,3 | 22,3 | 12,676 | 0 | 15,032 | 150310000 | 8 | AJQ90799.1_1063 | RM25_1082 | Cytoplasmic | 0 | CYT | 0.121415 | OTHER | K |
| lcl\|CP010341.1_prot_AJQ90803.1_1067 | RM25_RS05355 | UDP-N-acetylglucosamine transferase | 25428000 | 19190000 | 79167000 | 27329000 | 29741000 | 43983000 | 6 | 6 | 12,3 | 12,3 | 12,3 | 53,33 | 0 | 45,189 | 253350000 | 16 | AJQ90803.1_1067 | RM25_1086 | Cytoplasmic | 0 | CYT | 0.057491 | OTHER | M |
| lcl\|CP010341.1_prot_AJQ90804.1_1068 | RM25_RS05360 | Dihydrolipoyl dehydrogenase | 77408000 | 59376000 | 241380000 | 127360000 | 84007000 | 130670000 | 5 | 5 | 13,7 | 13,7 | 13,7 | 49,617 | 0 | 48,732 | 820450000 | 19 | AJQ90804.1_1068 | RM25_1087 | Cytoplasmic | 0 | CYT | 0.061230 | OTHER | C |
| lcl\|CP010341.1_prot_AJQ90805.1_1069 | RM25_RS05365 | Receptor ligand binding region family protein | 179450000 | 301560000 | 460340000 | 153740000 | 250380000 | 385080000 | 11 | 11 | 44,4 | 44,4 | 44,4 | 40,807 | 0 | 145 | 1850700000 | 41 | AJQ90805.1_1069 | RM25_1088 | Unknown | 0 | SPII | 0.839962 | LIPO(Sec/SPII) | E |
| lcl\|CP010341.1_prot_AJQ90810.1_1074 | RM25_RS05390 | AcuB family protein | 9635100 | 9759600 | 26266000 | 26195000 | 20403000 | 36657000 | 2 | 2 | 12 | 12 | 12 | 23,411 | 0 | 15,084 | 148230000 | 9 | AJQ90810.1_1074 | RM25_1093 | Unknown | 0 | CYT | 0.044737 | OTHER | |
| lcl\|CP010341.1_prot_AJQ90812.1_1076 | RM25_RS05400 | Dipeptidase pepE | 2599100 | 1138800 | 7719700 | 0 | 3094500 | 0 | 1 | 1 | 4,2 | 4,2 | 4,2 | 40,341 | 0,0051387 | 6,6629 | 14552000 | 1 | AJQ90812.1_1076 | RM25_1095 | Cytoplasmic | 0 | CYT | 0.061908 | OTHER | E |
| lcl\|CP010341.1_prot_AJQ90813.1_1077 | RM25_RS05405 | Putative esterase | 526609984 | 812769984 | 1166200064 | 313710016 | 625590016 | 643480000 | 7 | 7 | 20,3 | 20,3 | 20,3 | 46,942 | 0 | 194,1 | 4335000000 | 60 | AJQ90813.1_1077 | RM25_1096 | CytoplasmicMembrane | 4 | TMH | 0.844777 | OTHER | |
| lcl\|CP010341.1_prot_AJQ90815.1_1079 | RM25_RS05415 | Aspartyl/glutamyl-tRNA(Asn/Gln) amidotransferase sub | 36479000 | 43082000 | 165390000 | 91570000 | 78560000 | 106760000 | 9 | 9 | 22,9 | 22,9 | 22,9 | 54,833 | 0 | 78,805 | 567230000 | 26 | AJQ90815.1_1079 | RM25_1098 | Cytoplasmic | 0 | CYT | 0.076056 | OTHER | J |
| lcl\|CP010341.1_prot_AJQ90816.1_1080 | RM25_RS05420 | Glutamyl-tRNA(Gln) amidotransferase subunit A | 43542000 | 30653000 | 132820000 | 55515000 | 50255000 | 87138000 | 6 | 6 | 14,5 | 14,5 | 14,5 | 53,047 | 0 | 47,036 | 434050000 | 21 | AJQ90816.1_1080 | RM25_1099 | Cytoplasmic | 0 | CYT | 0.067721 | OTHER | J |
| lcl\|CP010341.1_prot_AJQ90822.1_1086 | RM25_RS05450 | Glycogen debranching enzyme GlgX | 4682600 | 4164500 | 12638000 | 5958400 | 9194300 | 14496000 | 1 | 1 | 1,9 | 1,9 | 1,9 | 80,886 | 0,0042735 | 7,5015 | 56666000 | 2 | AJQ90822.1_1086 | RM25_1105 | Cytoplasmic | 0 | CYT | 0.392176 | OTHER | G |
| lcl\|CP010341.1_prot_AJQ90823.1_1087 | RM25_RS05455 | Hypothetical protein | 93509000 | 74923000 | 149720000 | 18491000 | 32543000 | 88477000 | 1 | 1 | 18,1 | 18,1 | 18,1 | 11,071 | 0 | 46,262 | 478040000 | 7 | AJQ90823.1_1087 | RM25_1106 | Unknown | 0 | SPII | 0.050425 | LIPO(Sec/SPII) | |
| lcl\|CP010341.1_prot_AJQ90827.1_1091 | RM25_RS05475 | Alpha-glucan phosphorylase | 73900000 | 39958000 | 180330000 | 116600000 | 128570000 | 214310000 | 12 | 12 | 21 | 21 | 21 | 95,47 | 0 | 129,23 | 811970000 | 34 | AJQ90827.1_1091 | RM25_1110 | Cytoplasmic | 0 | CYT | 0.129895 | OTHER | G |
| lcl\|CP010341.1_prot_AJQ90828.1_1092 | RM25_RS05480 | Alpha-1,4-glucan:maltose-1-phosphate maltosyltransfe | 6884000 | 0 | 16269000 | 13088000 | 10737000 | 16357000 | 2 | 2 | 4 | 4 | 4 | 78,328 | 0 | 11,251 | 76372000 | 3 | AJQ90828.1_1092 | RM25_1111 | Cytoplasmic | 0 | CYT | 0.261427 | OTHER | G |
| lcl\|CP010341.1_prot_AJQ90829.1_1093 | RM25_RS05485 | Maltose alpha-D-glucosyltransferase | 12942000 | 12754000 | 24319000 | 0 | 0 | 0 | 2 | 2 | 6,3 | 6,3 | 6,3 | 68,428 | 0 | 18,108 | 50015000 | 5 | AJQ90829.1_1093 | RM25_1112 | Cytoplasmic | 0 | CYT | 0.300070 | OTHER | G |
| lcl\|CP010341.1_prot_AJQ90830.1_1094 | RM25_RS05490 | Maltokinase | 1478500 | 1568400 | 3862600 | 1054500 | 900390 | 0 | 1 | 1 | 3,1 | 3,1 | 3,1 | 49,26 | 0,0043197 | 7,5824 | 13522000 | 2 | AJQ90830.1_1094 | RM25_1113 | Cytoplasmic | 0 | CYT | 0.119727 | OTHER | G |
| lcl\|CP010341.1_prot_AJQ90831.1_1095 | RM25_RS05495 | 1,4-alpha-glucan branching enzyme GlgB | 18625000 | 4558000 | 42018000 | 10279000 | 28135000 | 34604000 | 5 | 5 | 9,6 | 9,6 | 9,6 | 72,667 | 0 | 35,487 | 139780000 | 10 | AJQ90831.1_1095 | RM25_1114 | Cytoplasmic | 0 | CYT | 0.282269 | OTHER | G |
| lcl\|CP010341.1_prot_AJQ90833.1_1097 | RM25_RS05505 | Phosphoglucomutase | 113690000 | 146280000 | 309969984 | 188140000 | 149280000 | 172550000 | 13 | 13 | 33,1 | 33,1 | 33,1 | 58,707 | 0 | 117,23 | 1228300000 | 52 | AJQ90833.1_1097 | RM25_1116 | Cytoplasmic | 0 | CYT | 0.460182 | OTHER | G |
| lcl\|CP010341.1_prot_AJQ90834.1_1098 | RM25_RS05510 | Thioredoxin TrxA | 84081000 | 78347000 | 236910000 | 135620000 | 156410000 | 194210000 | 7 | 7 | 33 | 33 | 33 | 33,905 | 0 | 84,382 | 1010200000 | 39 | AJQ90834.1_1098 | RM25_1117 | Unknown | 0 | CYT | 0.100219 | OTHER | O |
| lcl\|CP010341.1_prot_AJQ90835.1_1099 | RM25_RS05515 | Methylmalonyl-CoA epimerase | 197100000 | 62782000 | 652449984 | 394420000 | 368529984 | 675150016 | 6 | 6 | 50,7 | 50,7 | 50,7 | 16,717 | 0 | 49,514 | 2540000000 | 36 | AJQ90835.1_1099 | RM25_1118 | Cytoplasmic | 0 | CYT | 0.103911 | OTHER | E |
| lcl\|CP010341.1_prot_AJQ90837.1_1101 | RM25_RS12740 | Hypothetical protein | 193900000 | 212690000 | 526040000 | 363680000 | 346960000 | 423060000 | 19 | 19 | 37,5 | 37,5 | 37,5 | 64,112 | 0 | 244,95 | 2307000000 | 75 | AJQ90837.1_1101 | RM25_1120 | Cytoplasmic | 0 | CYT | 0.504288 | OTHER | |
| lcl\|CP010341.1_prot_AJQ90838.1_1102 | RM25_RS05530 | Endonuclease NucS | 3734800 | 3169300 | 10600000 | 5958400 | 5013800 | 10668000 | 1 | 1 | 6,5 | 6,5 | 6,5 | 23,688 | 0,004315 | 7,582 | 42603000 | 3 | AJQ90838.1_1102 | RM25_1121 | Cytoplasmic | 0 | CYT | 0.067786 | OTHER | L |
| lcl\|CP010341.1_prot_AJQ90841.1_1105 | RM25_RS05545 | ATP:cob(I)alamin adenosyltransferase | 18533000 | 17973000 | 96721000 | 16556000 | 5932400 | 18771000 | 5 | 5 | 36,1 | 36,1 | 36,1 | 22,197 | 0 | 57,557 | 179460000 | 11 | AJQ90841.1_1105 | RM25_1124 | Cytoplasmic | 0 | CYT | 0.077854 | OTHER | S |
| lcl\|CP010341.1_prot_AJQ90844.1_1108 | RM25_RS05560 | ATP synthase F1, epsilon subunit | 27692000 | 24666000 | 65613000 | 34330000 | 46653000 | 62557000 | 3 | 3 | 23,9 | 23,9 | 23,9 | 16,667 | 0 | 31,967 | 283470000 | 12 | AJQ90844.1_1108 | RM25_1127 | Cytoplasmic | 0 | CYT | 0.054111 | OTHER | |
| lcl\|CP010341.1_prot_AJQ90845.1_1109 | RM25_RS05565 | ATP synthase subunit beta | 315380000 | 331600000 | 707689984 | 217340000 | 167650000 | 297340000 | 11 | 11 | 30,4 | 30,4 | 30,4 | 52,497 | 0 | 136,51 | 2203300000 | 42 | AJQ90845.1_1109 | RM25_1128 | CytoplasmicMembrane | 0 | CYT | 0.107652 | OTHER | C |

| | | | | | | | | | | | | | | | | | | | | | | | | | | |
|---|---|---|---|---|---|---|---|---|---|---|---|---|---|---|---|---|---|---|---|---|---|---|---|---|---|---|
| lcl\|CP010341.1_prot_AJQ90846.1_1110 | RM25_RS05570 | ATP synthase gamma chain | 5734500 | 4838700 | 22441000 | 0 | 3762000 | 14517000 | 3 | 3 | 9 | 9 | 9 | 34,683 | 0 | 17,952 | 51293000 | 4 | AJQ90846.1_1110 | RM25_1129 | Cytoplasmic | 0 | CYT | 0.056581 | OTHER | C |
| lcl\|CP010341.1_prot_AJQ90847.1_1111 | RM25_RS05575 | ATP synthase subunit alpha | 158800000 | 108110000 | 635920000 | 253130000 | 293960000 | 410620000 | 19 | 19 | 47,9 | 47,9 | 47,9 | 58,988 | 0 | 244,04 | 2027000000 | 71 | AJQ90847.1_1111 | RM25_1130 | Cytoplasmic | 0 | CYT | 0.069777 | OTHER | C |
| lcl\|CP010341.1_prot_AJQ90848.1_1112 | RM25_RS05580 | ATP synthase F1, delta subunit | 34549000 | 18290000 | 80972000 | 44186000 | 66420000 | 118880000 | 5 | 5 | 23,2 | 23,2 | 23,2 | 27,779 | 0 | 37,162 | 413750000 | 15 | AJQ90848.1_1112 | RM25_1131 | Cytoplasmic | 0 | CYT | 0.035751 | OTHER | |
| lcl\|CP010341.1_prot_AJQ90849.1_1113 | RM25_RS05585 | ATP synthase subunit b | 73575000 | 54602000 | 178920000 | 139310000 | 151180000 | 229350000 | 4 | 4 | 32,6 | 32,6 | 32,6 | 20,192 | 0 | 72,239 | 919820000 | 31 | AJQ90849.1_1113 | RM25_1132 | CytoplasmicMembrane | 1 | CYT | 0.697238 | OTHER | C |
| lcl\|CP010341.1_prot_AJQ90856.1_1120 | RM25_RS05620 | Release factor glutamine methyltransferase | 13347000 | 13584000 | 26099000 | 9510300 | 15704000 | 13672000 | 2 | 2 | 5,8 | 5,8 | 5,8 | 31,291 | 0 | 11,425 | 91915000 | 5 | AJQ90856.1_1120 | RM25_1139 | Cytoplasmic | 0 | CYT | 0.099476 | OTHER | J |
| lcl\|CP010341.1_prot_AJQ90857.1_1121 | RM25_RS05625 | Peptide chain release factor 1 | 18011000 | 20418000 | 49637000 | 0 | 74277000 | 16885000 | 2 | 2 | 6,4 | 6,4 | 6,4 | 39,329 | 0,0045767 | 10,133 | 179230000 | 1 | AJQ90857.1_1121 | RM25_1140 | Cytoplasmic | 0 | CYT | 0.101060 | OTHER | J |
| lcl\|CP010341.1_prot_AJQ90858.1_1122 | RM25_RS05630 | 50S ribosomal protein L31 | 8018900 | 35577000 | 117280000 | 80357000 | 112570000 | 101640000 | 1 | 1 | 14,9 | 14,9 | 14,9 | 8,2172 | 0 | 11,227 | 503320000 | 7 | AJQ90858.1_1122 | RM25_1141 | Cytoplasmic | 0 | CYT | 0.933518 | OTHER | |
| lcl\|CP010341.1_prot_AJQ90859.1_1123 | RM25_RS05635 | Transcription termination factor Rho | 80927000 | 94167000 | 205000000 | 129370000 | 133110000 | 215440000 | 14 | 14 | 22,4 | 22,4 | 22,4 | 72,445 | 0 | 90,664 | 933800000 | 33 | AJQ90859.1_1123 | RM25_1142 | Cytoplasmic | 0 | CYT | 0.153585 | OTHER | K |
| lcl\|CP010341.1_prot_AJQ90860.1_1124 | RM25_RS05640 | Phosphoserine phosphatase/homoserine phosphotrans | 97619000 | 148300000 | 179200000 | 95962000 | 151180000 | 162170000 | 5 | 5 | 27,1 | 27,1 | 27,1 | 23,257 | 0 | 71,294 | 914280000 | 33 | AJQ90860.1_1124 | RM25_1143 | Cytoplasmic | 0 | CYT | 0.078938 | OTHER | |
| lcl\|CP010341.1_prot_AJQ90861.1_1125 | RM25_RS05645 | Homoserine dehydrogenase | 11764000 | 9292300 | 48897000 | 11383000 | 11092000 | 33867000 | 3 | 3 | 9,1 | 9,1 | 9,1 | 46,551 | 0 | 20,556 | 140200000 | 5 | AJQ90861.1_1125 | RM25_1144 | Cytoplasmic | 0 | CYT | 0.043431 | OTHER | E |
| lcl\|CP010341.1_prot_AJQ90862.1_1126 | RM25_RS05650 | Diaminopimelate decarboxylase | 11358000 | 5659200 | 34152000 | 8845200 | 11774000 | 26476000 | 2 | 2 | 6 | 6 | 6 | 50,013 | 0 | 12,762 | 105780000 | 4 | AJQ90862.1_1126 | RM25_1145 | Cytoplasmic | 0 | CYT | 0.084626 | OTHER | E |
| lcl\|CP010341.1_prot_AJQ90866.1_1130 | RM25_RS05675 | 2-oxoglutarate decarboxylase | 30036000 | 23837000 | 147450000 | 116330000 | 92969000 | 120030000 | 11 | 11 | 10,1 | 10,1 | 10,1 | 139,56 | 0 | 76,561 | 555450000 | 24 | AJQ90866.1_1130 | RM25_1150 | Cytoplasmic | 0 | CYT | 0.190770 | OTHER | C |
| lcl\|CP010341.1_prot_AJQ90870.1_1134 | RM25_RS05695 | RNA polymerase sigma factor | 12627000 | 18156000 | 33772000 | 54553000 | 46006000 | 57745000 | 5 | 5 | 32,3 | 32,3 | 32,3 | 22,137 | 0 | 36,21 | 286440000 | 15 | AJQ90870.1_1134 | RM25_1154 | Cytoplasmic | 0 | CYT | 0.068797 | OTHER | K |
| lcl\|CP010341.1_prot_AJQ90873.1_1137 | RM25_RS05710 | Putative ribosome biogenesis GTPase RsgA | 3634500 | 0 | 11008000 | 5260800 | 0 | 8423800 | 1 | 1 | 3,9 | 3,9 | 3,9 | 38,331 | 0,0041841 | 7,0095 | 31393000 | 1 | AJQ90873.1_1137 | RM25_1157 | Cytoplasmic | 0 | CYT | 0.092456 | OTHER | S |
| lcl\|CP010341.1_prot_AJQ90874.1_1138 | RM25_RS05715 | Histidinol-phosphate phosphatase HisN | 10515000 | 7608800 | 41689000 | 21855000 | 19965000 | 22540000 | 3 | 3 | 13,7 | 13,7 | 13,7 | 29,041 | 0 | 21,265 | 134970000 | 10 | AJQ90874.1_1138 | RM25_1158 | Unknown | 0 | CYT | 0.087975 | OTHER | G |
| lcl\|CP010341.1_prot_AJQ90875.1_1139 | RM25_RS05720 | PF14262 domain protein | 4085300 | 14108000 | 55798000 | 14811000 | 14077000 | 23988000 | 1 | 1 | 1,6 | 1,6 | 1,6 | 56,222 | 0,0051282 | 6,6305 | 134580000 | 3 | AJQ90875.1_1139 | RM25_1159 | Extracellular | 0 | SPII | 0.867182 | LIPO(Sec/SPII) | |
| lcl\|CP010341.1_prot_AJQ90881.1_1145 | RM25_RS05745 | Phosphoglycerate dehydrogenase | 57768000 | 29337000 | 159890000 | 149230000 | 146000000 | 231580000 | 7 | 7 | 24,1 | 24,1 | 24,1 | 43,642 | 0 | 81,725 | 882170000 | 26 | AJQ90881.1_1145 | RM25_1165 | Cytoplasmic | 0 | CYT | 0.069174 | OTHER | EH |
| lcl\|CP010341.1_prot_AJQ90886.1_1150 | RM25_RS05770 | ABC-type dipeptide transport system, periplasmic comp | 112490000 | 88943000 | 230450000 | 86694000 | 111890000 | 188040000 | 8 | 8 | 19,7 | 19,7 | 19,7 | 54,972 | 0 | 103,15 | 887610000 | 36 | AJQ90886.1_1150 | RM25_1170 | Cellwall | 0 | SPII | 0.856347 | LIPO(Sec/SPII) | |
| lcl\|CP010341.1_prot_AJQ90889.1_1153 | RM25_RS05790 | Prolyl aminopeptidase | 2017400 | 0 | 3556300 | 0 | 0 | 0 | 1 | 1 | 3,2 | 3,2 | 3,2 | 44,399 | 0,0042328 | 7,3499 | 5573600 | 1 | AJQ90889.1_1153 | RM25_1174 | Cytoplasmic | 0 | CYT | 0.158976 | OTHER | S |
| lcl\|CP010341.1_prot_AJQ90890.1_1154 | RM25_RS05795 | Peptidase M16 inactive domain protein | 3545900 | 3017900 | 22263000 | 2200000 | 9182100 | 15774000 | 3 | 3 | 10,8 | 10,8 | 10,8 | 46,316 | 0 | 16,601 | 75238000 | 0 | AJQ90890.1_1154 | RM25_1175 | Cytoplasmic | 0 | CYT | 0.143828 | OTHER | |
| lcl\|CP010341.1_prot_AJQ90891.1_1155 | RM25_RS05800 | Peptidase M16 inactive domain protein | 1893100 | 6376000 | 20403000 | 13774000 | 8427300 | 14889000 | 2 | 2 | 5,5 | 5,5 | 5,5 | 48,504 | 0 | 15,69 | 65762000 | 5 | AJQ90891.1_1155 | RM25_1176 | Cytoplasmic | 0 | CYT | 0.086710 | OTHER | |
| lcl\|CP010341.1_prot_AJQ90892.1_1156 | RM25_RS05805 | UPF0182 protein | 187460000 | 294870016 | 439990016 | 249670000 | 299230016 | 524350016 | 18 | 18 | 20,8 | 20,8 | 20,8 | 107,23 | 0 | 323,31 | 2174200000 | 61 | AJQ90892.1_1156 | RM25_1177 | CytoplasmicMembrane | 7 | TMH | 0.854075 | OTHER | T |
| lcl\|CP010341.1_prot_AJQ90893.1_1157 | RM25_RS05810 | Hypothetical protein | 50534000 | 81227000 | 144810000 | 93378000 | 85062000 | 86882000 | 5 | 5 | 36,7 | 36,7 | 36,7 | 21,513 | 0 | 55,43 | 611340000 | 25 | AJQ90893.1_1157 | RM25_1178 | Unknown | 0 | CYT | 0.095661 | OTHER | - |
| lcl\|CP010341.1_prot_AJQ90894.1_1158 | RM25_RS05815 | putative secreted protein containing a PDZ domain | 739360000 | 1240899968 | 1185900032 | 485920000 | 777420032 | 1281299968 | 9 | 9 | 36,9 | 36,9 | 36,9 | 36,324 | 0 | 323,31 | 6191300000 | 49 | AJQ90894.1_1158 | RM25_1179 | Unknown | 1 | SPI | 0.463151 | OTHER | T |
| lcl\|CP010341.1_prot_AJQ90895.1_1159 | RM25_RS05820 | Putative hydrolase | 2040200 | 3690400 | 7890000 | 4103900 | 0 | 6201200 | 1 | 1 | 3,6 | 3,6 | 3,6 | 50,68 | 0,0098717 | 5,9587 | 23926000 | 0 | AJQ90895.1_1159 | RM25_1180 | Cytoplasmic | 0 | CYT | 0.103928 | OTHER | S |
| lcl\|CP010341.1_prot_AJQ90898.1_1162 | RM25_RS05835 | Hypothetical protein | 62833000 | 69447000 | 99801000 | 51616000 | 67203000 | 63195000 | 1 | 1 | 15,1 | 15,1 | 15,1 | 5,7808 | 0,0050968 | 6,4835 | 438250000 | 7 | AJQ90898.1_1162 | RM25_1183 | Unknown | 0 | CYT | 0.083024 | OTHER | |
| lcl\|CP010341.1_prot_AJQ90905.1_1169 | RM25_RS05870 | UvrD/REP helicase | 36031000 | 1181000 | 97881000 | 1613800 | 4430800 | 72069000 | 2 | 2 | 2,9 | 2,9 | 2,9 | 120,61 | 0,0011641 | 10,69 | 213210000 | 1 | AJQ90905.1_1169 | RM25_1190 | Cytoplasmic | 0 | CYT | 0.081737 | OTHER | L |
| lcl\|CP010341.1_prot_AJQ90907.1_1171 | RM25_RS05880 | Cold shock-like protein CspA | 1294400000 | 2179599872 | 2795000064 | 1884700032 | 1067900032 | 2146000000 | 2 | 2 | 54,4 | 54,4 | 54,4 | 7,3951 | 0 | 60,425 | 1,2835E+10 | 32 | AJQ90907.1_1171 | RM25_1192 | Cytoplasmic | 0 | CYT | 0.734862 | OTHER | |
| lcl\|CP010341.1_prot_AJQ90908.1_1172 | RM25_RS05885 | Serine/threonine phosphatase stp | 11983000 | 16233000 | 38422000 | 0 | 20635000 | 30330000 | 2 | 2 | 9,7 | 9,7 | 9,7 | 29,913 | 0 | 18,397 | 117600000 | 9 | AJQ90908.1_1172 | RM25_1193 | Unknown | 0 | CYT | 0.390157 | OTHER | T |
| lcl\|CP010341.1_prot_AJQ90909.1_1173 | RM25_RS05890 | Hypothetical protein | 27873000 | 26857000 | 87720000 | 73285000 | 60506000 | 52689000 | 5 | 5 | 56,4 | 56,4 | 56,4 | 10,07 | 0 | 36,662 | 363250000 | 14 | AJQ90909.1_1173 | RM25_1194 | Cytoplasmic | 0 | CYT | 0.060349 | OTHER | S |
| lcl\|CP010341.1_prot_AJQ90910.1_1174 | RM25_RS05895 | Hypothetical protein | 17423000 | 19213000 | 37252000 | 31415000 | 6023200 | 48518000 | 2 | 2 | 11,2 | 11,2 | 11,2 | 26,859 | 0 | 13,971 | 159840000 | 7 | AJQ90910.1_1174 | RM25_1195 | Cytoplasmic | 0 | CYT | 0.064181 | OTHER | S |
| lcl\|CP010341.1_prot_AJQ90911.1_1175 | RM25_RS05900 | DEAD-box ATP-dependent RNA helicase CshA | 67967000 | 72391000 | 193930000 | 94743000 | 73484000 | 216610000 | 9 | 9 | 20,2 | 20,2 | 20,2 | 55,495 | 0 | 117,66 | 802120000 | 29 | AJQ90911.1_1175 | RM25_1196 | Cytoplasmic | 0 | CYT | 0.164976 | OTHER | L |
| lcl\|CP010341.1_prot_AJQ90912.1_1176 | RM25_RS05905 | Tryptophan--tRNA ligase | 25979000 | 13033000 | 94898000 | 11655000 | 11788000 | 66043000 | 4 | 4 | 11,5 | 11,5 | 11,5 | 42,611 | 0 | 28,062 | 250220000 | 11 | AJQ90912.1_1176 | RM25_1197 | Cytoplasmic | 0 | CYT | 0.097646 | OTHER | J |
| lcl\|CP010341.1_prot_AJQ90913.1_1177 | RM25_RS05910 | PHP domain protein | 26058000 | 23546000 | 88374000 | 21054000 | 31351000 | 77164000 | 5 | 5 | 24,6 | 24,6 | 24,6 | 31,166 | 0 | 40,512 | 282320000 | 13 | AJQ90913.1_1177 | RM25_1198 | Cytoplasmic | 0 | CYT | 0.069029 | OTHER | S |
| lcl\|CP010341.1_prot_AJQ90914.1_1178 | RM25_RS05915 | Hypothetical protein | 956750 | 22787000 | 37099000 | 19433000 | 24916000 | 32932000 | 1 | 1 | 7,6 | 7,6 | 7,6 | 17,206 | 0,0043956 | 8,0735 | 138120000 | 3 | AJQ90914.1_1178 | RM25_1199 | Unknown | 0 | CYT | 0.746589 | OTHER | |
| lcl\|CP010341.1_prot_AJQ90915.1_1179 | RM25_RS05920 | Hypothetical protein | 5087200 | 6426000 | 7026000 | 0 | 4534700 | 9108900 | 1 | 1 | 23,9 | 23,9 | 23,9 | 7,24 | 0,0044643 | 8,5211 | 32183000 | 3 | AJQ90915.1_1179 | RM25_1200 | Unknown | 0 | CYT | 0.235022 | OTHER | |
| lcl\|CP010341.1_prot_AJQ90918.1_1182 | RM25_RS06075 | Protein translocase subunit SecA | 206040000 | 235020000 | 434460000 | 245580000 | 307310016 | 459110016 | 22 | 22 | 27,4 | 27,4 | 27,4 | 106,53 | 0 | 263,22 | 2179000000 | 103 | AJQ90918.1_1182 | RM25_1231 | CytoplasmicMembrane | 0 | CYT | 0.082400 | OTHER | I |
| lcl\|CP010341.1_prot_AJQ90919.1_1183 | RM25_RS06075 | S30AE family protein | 272169984 | 349460000 | 1036099968 | 551059968 | 566700032 | 585870016 | 8 | 8 | 47 | 47 | 47 | 22,024 | 0 | 72,639 | 3729600000 | 52 | AJQ90919.1_1183 | RM25_1231 | CytoplasmicMembrane | 0 | CYT | 0.082400 | OTHER | I |
| lcl\|CP010341.1_prot_AJQ90925.1_1189 | RM25_RS05975 | MutG family lantibiotic protection ABC superfamily ATP | 107560000 | 198480000 | 185850000 | 42093000 | 59704000 | 86772000 | 2 | 2 | 6,6 | 6,6 | 6,6 | 67,487 | 0 | 62,651 | 708420000 | 7 | AJQ90925.1_1189 | RM25_1210 | CytoplasmicMembrane | 4 | SPI | 0.803167 | SP(Sec/SPI) | |
| lcl\|CP010341.1_prot_AJQ90926.1_1190 | RM25_RS05980 | Hypothetical protein | 945110016 | 1261299968 | 1771699968 | 1112199936 | 1600300032 | 2244499968 | 10 | 10 | 23,2 | 23,2 | 23,2 | 45,741 | 0 | 323,31 | 9811800000 | 75 | AJQ90926.1_1190 | RM25_1211 | Extracellular | 0 | SPI | 0.298566 | SP(Sec/SPI) | M |
| lcl\|CP010341.1_prot_AJQ90927.1_1191 | RM25_RS05985 | Cell division protein FtsX | 18075000 | 57768000 | 40701000 | 2005600 | 9845000 | 20819000 | 1 | 1 | 4,3 | 4,3 | 4,3 | 33,638 | 0 | 55,804 | 154700000 | 5 | AJQ90927.1_1191 | RM25_1212 | CytoplasmicMembrane | 4 | SPI | 0.851227 | SP(Sec/SPI) | D |
| lcl\|CP010341.1_prot_AJQ90928.1_1192 | RM25_RS05990 | Cell division ATP-binding protein FtsE | 2280400 | 1463900 | 3643500 | 2431600 | 3530700 | 4451700 | 2 | 2 | 8,8 | 8,8 | 8,8 | 25,583 | 0,0023148 | 10,565 | 19264000 | 1 | AJQ90928.1_1192 | RM25_1213 | CytoplasmicMembrane | 0 | CYT | 0.050443 | OTHER | D |
| lcl\|CP010341.1_prot_AJQ90929.1_1193 | RM25_RS05995 | Peptide chain release factor 2 | 24740000 | 38402000 | 85541000 | 82411000 | 70526000 | 90540000 | 6 | 6 | 22,5 | 22,5 | 22,5 | 41,071 | 0 | 70,129 | 456590000 | 18 | AJQ90929.1_1193 | RM25_1214 | Cytoplasmic | 0 | CYT | 0.135406 | OTHER | J |
| lcl\|CP010341.1_prot_AJQ90930.1_1194 | RM25_RS06000 | GatB/YqeY domain protein | 17189000 | 18820000 | 16107000 | 33634000 | 38263000 | 42514000 | 2 | 2 | 13,7 | 13,7 | 13,7 | 16,553 | 0 | 19,334 | 191690000 | 13 | AJQ90930.1_1194 | RM25_1215 | Cytoplasmic | 0 | CYT | 0.033844 | OTHER | S |
| lcl\|CP010341.1_prot_AJQ90932.1_1196 | RM25_RS06015 | 4-aminobutyrate transaminase | 14160000 | 11830000 | 107500000 | 15235000 | 8392700 | 17656000 | 3 | 3 | 7,9 | 7,9 | 7,9 | 46,697 | 0 | 47,561 | 178300000 | 4 | AJQ90932.1_1196 | RM25_1218 | Cytoplasmic | 0 | CYT | 0.071154 | OTHER | E |
| lcl\|CP010341.1_prot_AJQ90935.1_1199 | RM25_RS06030 | Putative heat shock protein Hsp18 | 1000400000 | 832030016 | 2391000064 | 1876300032 | 1355500032 | 3184900096 | 11 | 11 | 74,7 | 74,7 | 74,7 | 16,901 | 0 | 323,31 | 1,2241E+10 | 105 | AJQ90935.1_1199 | RM25_1221 | Cellwall | 0 | CYT | 0.828807 | OTHER | O |
| lcl\|CP010341.1_prot_AJQ90937.1_1201 | RM25_RS06040 | Hypothetical protein | 25454000 | 32112000 | 50319000 | 46065000 | 46708000 | 62097000 | 2 | 2 | 18,4 | 18,4 | 18,4 | 15,463 | 0 | 19,003 | 297510000 | 6 | AJQ90937.1_1201 | RM25_1223 | Cytoplasmic | 0 | CYT | 0.086227 | OTHER | S |
| lcl\|CP010341.1_prot_AJQ90939.1_1203 | RM25_RS06045 | Pyruvate dehydrogenase E1 component | 80014000 | 59791000 | 243610000 | 109470000 | 89915000 | 192950000 | 11 | 11 | 16,3 | 16,3 | 16,3 | 102,14 | 0 | 94,288 | 825850000 | 21 | AJQ90939.1_1203 | RM25_1225 | Unknown | 0 | CYT | 0.291158 | OTHER | C |
| lcl\|CP010341.1_prot_AJQ90940.1_1204 | RM25_RS06050 | Aromatic-amino-acid transaminase | 49431000 | 64335000 | 88573000 | 95668000 | 81470000 | 136350000 | 8 | 8 | 23,4 | 23,4 | 23,4 | 43,099 | 0 | 57,345 | 583260000 | 22 | AJQ90940.1_1204 | RM25_1226 | Cytoplasmic | 0 | CYT | 0.087244 | OTHER | |
| lcl\|CP010341.1_prot_AJQ90942.1_1206 | RM25_RS06075 | Malonyl-CoA--acyl carrier protein transacylase | 146320000 | 252890000 | 498070016 | 247150000 | 192190000 | 281340000 | 5 | 5 | 22,6 | 22,6 | 22,6 | 32,968 | 0 | 108,5 | 1801200000 | 48 | AJQ90942.1_1206 | RM25_1231 | CytoplasmicMembrane | 0 | CYT | 0.082400 | OTHER | I |
| lcl\|CP010341.1_prot_AJQ90943.1_1207 | RM25_RS06075 | 3-oxoacyl-acyl-carrier-protein synthase 3 | 195100000 | 310500000 | 814099968 | 246800000 | 383409984 | 467609984 | 10 | 10 | 45,8 | 45,8 | 45,8 | 35,24 | 0 | 182,37 | 2586000000 | 59 | AJQ90943.1_1207 | RM25_1231 | CytoplasmicMembrane | 0 | CYT | 0.082400 | OTHER | I |
| lcl\|CP010341.1_prot_AJQ90944.1_1208 | RM25_RS06075 | Acyl carrier protein | 1467000064 | 2567300096 | 3161700096 | 2576300032 | 2353700096 | 3344300032 | 3 | 3 | 48,2 | 48,2 | 48,2 | 9,1852 | 0 | 95,133 | 1,6948E+10 | 53 | AJQ90944.1_1208 | RM25_1231 | CytoplasmicMembrane | 0 | CYT | 0.082400 | OTHER | I |
| lcl\|CP010341.1_prot_AJQ90945.1_1209 | RM25_RS06075 | 3-oxoacyl- | 62236000 | 63892000 | 276740000 | 82384000 | 51733000 | 114960000 | 4 | 4 | 15,3 | 15,3 | 15,3 | 43,747 | 0 | 203,88 | 713320000 | 15 | AJQ90945.1_1209 | RM25_1231 | CytoplasmicMembrane | 0 | CYT | 0.082400 | OTHER | I |
| lcl\|CP010341.1_prot_AJQ90948.1_1212 | RM25_RS06090 | Pyridoxal phosphate enzyme, YggS family | 3689000 | 4480100 | 11889000 | 5930600 | 8258200 | 9449700 | 2 | 2 | 12,4 | 12,4 | 12,4 | 26,996 | 0 | 12,835 | 45985000 | 2 | AJQ90948.1_1212 | RM25_1234 | Cytoplasmic | 0 | CYT | 0.045866 | OTHER | S |
| lcl\|CP010341.1_prot_AJQ90949.1_1213 | RM25_RS06095 | UPF0353 protein | 8085800 | 9384200 | 14412000 | 3251700 | 5457600 | 10822000 | 2 | 2 | 11,8 | 11,8 | 11,8 | 34,303 | 0 | 33,712 | 55570000 | 7 | AJQ90949.1_1213 | RM25_1235 | CytoplasmicMembrane | 3 | TMH | 0.865391 | OTHER | |
| lcl\|CP010341.1_prot_AJQ90950.1_1214 | RM25_RS06100 | UPF0353 protein | 12254000 | 19637000 | 23869000 | 17015000 | 14666000 | 23716000 | 2 | 2 | 7,1 | 7,1 | 7,1 | 34,453 | 0 | 31,479 | 128330000 | 6 | AJQ90950.1_1214 | RM25_1236 | CytoplasmicMembrane | 2 | TMH | 0.630060 | OTHER | |
| lcl\|CP010341.1_prot_AJQ90952.1_1216 | RM25_RS06110 | Transcriptional regulator moxR1 | 40502000 | 27940000 | 81834000 | 55530000 | 69840000 | 100640000 | 8 | 8 | 32,3 | 32,3 | 32,3 | 37,443 | 0 | 60,451 | 419130000 | 17 | AJQ90952.1_1216 | RM25_1238 | Cytoplasmic | 0 | CYT | 0.067248 | OTHER | S |
| lcl\|CP010341.1_prot_AJQ90955.1_1219 | RM25_RS06125 | Dihydrolipoyl dehydrogenase | 13326000 | 15255000 | 36009000 | 16834000 | 14004000 | 18425000 | 2 | 2 | 5 | 5 | 5 | 49,863 | 0 | 17,752 | 130760000 | 3 | AJQ90955.1_1219 | RM25_1241 | Cytoplasmic | 0 | CYT | 0.055086 | OTHER | C |
| lcl\|CP010341.1_prot_AJQ90960.1_1224 | RM25_RS06150 | Succinate dehydrogenase/fumarate reductase iron-sulf | 1315400 | 1136600 | 13560000 | 0 | 7603700 | 0 | 2 | 2 | 6,3 | 6,3 | 6,3 | 27,282 | 0 | 10,976 | 23616000 | 2 | AJQ90960.1_1224 | RM25_1246 | Cytoplasmic | 0 | CYT | 0.083911 | OTHER | C |
| lcl\|CP010341.1_prot_AJQ90961.1_1225 | RM25_RS06155 | Succinate dehydrogenase flavoprotein subunit | 64227000 | 22343000 | 259410000 | 83557000 | 134330000 | 147990000 | 14 | 14 | 22,8 | 22,8 | 22,8 | 76,466 | 0 | 93,436 | 763120000 | 31 | AJQ90961.1_1225 | RM25_1247 | Cytoplasmic | 0 | CYT | 0.159137 | OTHER | C |
| lcl\|CP010341.1_prot_AJQ90962.1_1226 | RM25_RS06160 | Succinate dehydrogenase cytochrome B subunit, b558 f | 13351000 | 15159000 | 28015000 | 20352000 | 12309000 | 33647000 | 1 | 1 | 6,4 | 6,4 | 6,4 | 27,474 | 0,0045819 | 10,135 | 122830000 | 6 | AJQ90962.1_1226 | RM25_1248 | CytoplasmicMembrane | 5 | CYT | 0.947575 | OTHER | C |
| lcl\|CP010341.1_prot_AJQ90967.1_1231 | RM25_RS06185 | Glycine--tRNA ligase | 33331000 | 15982000 | 102940000 | 53452000 | 24533000 | 60736000 | 4 | 4 | 9,7 | 9,7 | 9,7 | 52,98 | 0 | 29,368 | 329570000 | 8 | AJQ90967.1_1231 | RM25_1253 | Cytoplasmic | 0 | CYT | 0.176735 | OTHER | J |
| lcl\|CP010341.1_prot_AJQ90968.1_1232 | RM25_RS06190 | Metal transport system ABC transporter substrate-bind | 26040000 | 23201000 | 48932000 | 24836000 | 25568000 | 56567000 | 3 | 3 | 11,4 | 11,4 | 11,4 | 33,558 | 0 | 28,104 | 221470000 | 10 | AJQ90968.1_1232 | RM25_1254 | CytoplasmicMembrane | 1 | SPII | 0.929883 | LIPO(Sec/SPII) | P |
| lcl\|CP010341.1_prot_AJQ90973.1_1237 | RM25_RS06215 | Ethanolamine utilization protein EutE | 9365100 | 11000000 | 24691000 | 0 | 0 | 0 | 1 | 1 | 2,6 | 2,6 | 2,6 | 50,118 | 0,0043384 | 7,6249 | 45056000 | 3 | AJQ90973.1_1237 | RM25_1259 | Cytoplasmic | 0 | CYT | 0.069359 | OTHER | C |
| lcl\|CP010341.1_prot_AJQ90978.1_1242 | RM25_RS06240 | Propanediol utilization protein PduJ | 54571000 | 62965000 | 124970000 | 21093000 | 0 | 23066000 | 3 | 3 | 46,7 | 46,7 | 46,7 | 9,2237 | 0 | 36,787 | 292640000 | 13 | AJQ90978.1_1242 | RM25_1264 | Unknown | 0 | CYT | 0.019046 | OTHER | |
| lcl\|CP010341.1_prot_AJQ90982.1_1246 | RM25_RS06260 | Propanediol dehydratase small subunit | 4142800 | 3674000 | 11851000 | 0 | 1327100 | 0 | 1 | 1 | 7,9 | 7,9 | 7,9 | 19,469 | 0,0070707 | 6,2832 | 20995000 | 2 | AJQ90982.1_1246 | RM25_1268 | Cytoplasmic | 0 | CYT | 0.097673 | OTHER | |
| lcl\|CP010341.1_prot_AJQ90983.1_1247 | RM25_RS06265 | Propanediol dehydratase medium subunit | 11026000 | 11627000 | 27566000 | 0 | 0 | 3566500 | 2 | 2 | 9,3 | 9,3 | 9,3 | 24,234 | 0 | 12,423 | 53785000 | 4 | AJQ90983.1_1247 | RM25_1269 | Cytoplasmic | 0 | CYT | 0.086353 | OTHER | |
| lcl\|CP010341.1_prot_AJQ90984.1_1248 | RM25_RS06270 | Glycerol dehydratase large subunit | 3672800 | 3174900 | 8704800 | 5127200 | 7651000 | 8296800 | 2 | 2 | 4,3 | 4,3 | 4,3 | 60,572 | 0 | 10,853 | 41078000 | 3 | AJQ90984.1_1248 | RM25_1270 | Cytoplasmic | 0 | CYT | 0.080512 | OTHER | |
| lcl\|CP010341.1_prot_AJQ90985.1_1249 | RM25_RS06275 | Propanediol utilization protein PduB | 20766000 | 675390 | 46354000 | 5099300 | 3620900 | 0 | 2 | 2 | 12,3 | 12,3 | 12,3 | 27,383 | 0 | 25,558 | 76516000 | 4 | AJQ90985.1_1249 | RM25_1271 | Unknown | 0 | CYT | 0.056528 | OTHER | |
| lcl\|CP010341.1_prot_AJQ90986.1_1250 | RM25_RS06280 | Propanediol utilization protein PduA | 29002000 | 63841000 | 112320000 | 29339000 | 3249700 | 0 | 5 | 5 | 75,5 | 75,5 | 75,5 | 9,5489 | 0 | 39,692 | 260440000 | 5 | AJQ90986.1_1250 | RM25_1272 | Unknown | 0 | CYT | 0.028571 | OTHER | CQ |
| lcl\|CP010341.1_prot_AJQ90991.1_1255 | RM25_RS06300 | Ketol-acid reductoisomerase | 231620000 | 257870000 | 637249984 | 175240000 | 211570000 | 280550016 | 13 | 13 | 43,4 | 43,4 | 43,4 | 37,187 | 0 | 141,82 | 1894100000 | 50 | AJQ90991.1_1255 | RM25_1277 | Cytoplasmic | 0 | CYT | 0.086735 | OTHER | EH |
| lcl\|CP010341.1_prot_AJQ90995.1_1259 | RM25_RS06320 | Imidazole glycerol phosphate synthase subunit HisF | 5089900 | 8221700 | 16245000 | 12863000 | 16195000 | 13277000 | 1 | 1 | 5,9 | 5,9 | 5,9 | 26,841 | 0,004158 | 6,8922 | 71891000 | 2 | AJQ90995.1_1259 | RM25_1281 | Cytoplasmic | 0 | CYT | 0.045692 | OTHER | E |
| lcl\|CP010341.1_prot_AJQ90996.1_1260 | RM25_RS06325 | Voltage-dependent potassium channel beta subunit | 9811500 | 4354000 | 32095000 | 15784000 | 7612900 | 27175000 | 2 | 2 | 7,4 | 7,4 | 7,4 | 36,991 | 0 | 14,887 | 108870000 | 3 | AJQ90996.1_1260 | RM25_1282 | Cytoplasmic | 0 | CYT | 0.111994 | OTHER | C |
| lcl\|CP010341.1_prot_AJQ90997.1_1261 | RM25_RS06330 | Antibiotic biosynthesis monooxygenase | 60348000 | 100180000 | 117800000 | 86438000 | 129340000 | 97454000 | 4 | 4 | 37,6 | 37,6 | 37,6 | 14,058 | 0 | 40,474 | 604300000 | 16 | AJQ90997.1_1261 | RM25_1283 | Cytoplasmic | 0 | CYT | 0.094410 | OTHER | |
| lcl\|CP010341.1_prot_AJQ91002.1_1266 | RM25_RS12290 | Hypothetical protein | 4702200 | 4224100 | 15367000 | 2451200 | 3122800 | 2936300 | 1 | 1 | 8,8 | 8,8 | 8,8 | 10,414 | 0,0041929 | 7,1164 | 34465000 | 6 | AJQ91002.1_1266 | RM25_1288 | CytoplasmicMembrane | 1 | SPI | 0.974848 | SP(Sec/SPI) | U |
| lcl\|CP010341.1_prot_AJQ91003.1_1267 | RM25_RS06360 | Protein pafC | 3718100 | 5080300 | 17280000 | 13867000 | 10133000 | 0 | 2 | 2 | 8,8 | 8,8 | 8,8 | 34,124 | 0 | 13,286 | 50078000 | 1 | AJQ91003.1_1267 | RM25_1289 | Cytoplasmic | 0 | CYT | 0.063572 | OTHER | K |
| lcl\|CP010341.1_prot_AJQ91004.1_1268 | RM25_RS06365 | Protein pafB | 2766000 | 0 | 6254800 | 4507800 | 3771500 | 0 | 2 | 2 | 10,2 | 10,2 | 10,2 | 35,622 | 0 | 16,838 | 20701000 | 2 | AJQ91004.1_1268 | RM25_1290 | Cytoplasmic | 0 | CYT | 0.059013 | OTHER | K |
| lcl\|CP010341.1_prot_AJQ91006.1_1270 | RM25_RS06375 | Peptidyl-prolyl cis-trans isomerase | 613529984 | 633960000 | 1083600000 | 239970000 | 554120000 | 728220032 | 4 | 4 | 18,2 | 18,2 | 18,2 | 35,697 | 0 | 135,23 | 3943600000 | 26 | AJQ91006.1_1270 | RM25_1292 | Extracellular | 0 | SPI | 0.846925 | LIPO(Sec/SPII) | O |
| lcl\|CP010341.1_prot_AJQ91007.1_1271 | RM25_1293 | Peptidyl-prolyl cis-trans isomerase | 46904000 | 22812000 | 141010000 | 2792900 | 89730000 | 78999000 | 3 | 3 | 24,3 | 24,3 | 24,3 | 13,655 | 0 | 67,362 | 383250000 | 11 | AJQ91007.1_1271 | RM25_1293 | Unknown | 0 | CYT | 0.698586 | OTHER | |
| lcl\|CP010341.1_prot_AJQ91008.1_1272 | RM25_RS06385 | Pseudouridylate synthase | 7803200 | 2334300 | 36847000 | 19883000 | 4453500 | 12527000 | 4 | 4 | 16,5 | 16,5 | 16,5 | 29,09 | 0 | 37,629 | 90659000 | 6 | AJQ91008.1_1272 | RM25_1294 | Cytoplasmic | 0 | CYT | 0.087731 | OTHER | J |
| lcl\|CP010341.1_prot_AJQ91012.1_1276 | RM25_RS06405 | Initiation inhibitor protein | 25445000 | 22568000 | 79622000 | 48238000 | 17576000 | 41119000 | 4 | 4 | 20,8 | 20,8 | 20,8 | 32,259 | 0 | 34,092 | 278480000 | 11 | AJQ91012.1_1276 | RM25_1298 | CytoplasmicMembrane | 0 | CYT | 0.056441 | OTHER | D |
| lcl\|CP010341.1_prot_AJQ91014.1_1278 | RM25_RS06415 | ADP-ribose pyrophosphatase (ADPRase) | 5972800 | 7837300 | 14552000 | 18938000 | 14681000 | 19154000 | 2 | 2 | 17,3 | 17,3 | 17,3 | 24,44 | 0 | 14,383 | 84951000 | 3 | AJQ91014.1_1278 | RM25_1300 | Cytoplasmic | 0 | CYT | 0.070747 | OTHER | L |
| lcl\|CP010341.1_prot_AJQ91015.1_1279 | RM25_RS06420 | CTP synthase | 95978000 | 110100000 | 175050000 | 153670000 | 183010000 | 364110016 | 13 | 13 | 32,7 | 32,7 | 32,7 | 63,876 | 0 | 148,17 | 1208200000 | 48 | AJQ91015.1_1279 | RM25_1301 | Cytoplasmic | 0 | CYT | 0.093592 | OTHER | F |
| lcl\|CP010341.1_prot_AJQ91022.1_1286 | RM25_RS06455 | HAD-superfamily hydrolase, subfamily IIA | 2578800 | 3531300 | 4737800 | 0 | 2682000 | 3922700 | 2 | 2 | 5,2 | 5,2 | 5,2 | 36,818 | 0 | 11,737 | 17453000 | 4 | AJQ91022.1_1286 | RM25_1308 | Cytoplasmic | 0 | CYT | 0.062583 | OTHER | G |
| lcl\|CP010341.1_prot_AJQ91023.1_1287 | RM25_RS12295 | Hypothetical protein | 3797800 | 4655500 | 9021700 | 5852500 | 5773400 | 0 | 1 | 1 | 3 | 3 | 3 | 39,038 | 0,0051546 | 6,6891 | 33386000 | 2 | AJQ91023.1_1287 | RM25_1309 | Cytoplasmic | 0 | CYT | 0.725284 | OTHER | S |
| lcl\|CP010341.1_prot_AJQ91026.1_1290 | RM25_RS06480 | Tyrosine--tRNA ligase | 81205000 | 64303000 | 195090000 | 79221000 | 119840000 | 146550000 | 8 | 8 | 28,7 | 28,7 | 28,7 | 45,985 | 0 | 123,49 | 769200000 | 32 | AJQ91026.1_1290 | RM25_1315 | Cytoplasmic | 0 | CYT | 0.073900 | OTHER | J |
| lcl\|CP010341.1_prot_AJQ91027.1_1291 | RM25_RS06485 | Argininosuccinate lyase | 3802500 | 0 | 16448000 | 0 | 0 | 0 | 1 | 1 | 3,6 | 3,6 | 3,6 | 50,937 | 0,0045147 | 9,339 | 20250000 | 2 | AJQ91027.1_1291 | RM25_1316 | Cytoplasmic | 0 | CYT | 0.073832 | OTHER | E |
| lcl\|CP010341.1_prot_AJQ91031.1_1295 | RM25_RS06505 | Arginine biosynthesis bifunctional protein ArgJ | 11143000 | 15436000 | 55233000 | 5001200 | 4271300 | 11521000 | 3 | 3 | 8,6 | 8,6 | 8,6 | 39,52 | 0 | 21,042 | 102610000 | 7 | AJQ91031.1_1295 | RM25_1320 | Cytoplasmic | 0 | CYT | 0.065459 | OTHER | E |
| lcl\|CP010341.1_prot_AJQ91033.1_1297 | RM25_RS06515 | Aminotransferase AlaT | 6767000 | 10514000 | 24824000 | 29485000 | 25730000 | 31842000 | 2 | 2 | 6,7 | 6,7 | 6,7 | 44,568 | 0 | 12,12 | 146540000 | 5 | AJQ91033.1_1297 | RM25_1322 | Cytoplasmic | 0 | CYT | 0.098123 | OTHER | E |
| lcl\|CP010341.1_prot_AJQ91034.1_1298 | RM25_RS06520 | Phenylalanine--tRNA ligase beta subunit | 95269000 | 65039000 | 230150000 | 166670000 | 118770000 | 240240000 | 16 | 16 | 27,3 | 27,3 | 27,3 | 90,117 | 0 | 171,86 | 1012800000 | 56 | AJQ91034.1_1298 | RM25_1323 | Cytoplasmic | 0 | CYT | 0.083500 | OTHER | J |
| lcl\|CP010341.1_prot_AJQ91035.1_1299 | RM25_RS06525 | Phenylalanine--tRNA ligase alpha subunit | 27187000 | 18591000 | 91034000 | 66942000 | 56657000 | 97782000 | 9 | 9 | 25,5 | 25,5 | 25,5 | 39,779 | 0 | 81,62 | 385380000 | 19 | AJQ91035.1_1299 | RM25_1324 | Cytoplasmic | 0 | CYT | 0.102721 | OTHER | J |
| lcl\|CP010341.1_prot_AJQ91038.1_1302 | RM25_RS06535 | 50S ribosomal protein L20 | 140000000 | 115460000 | 344950016 | 222350000 | 248450000 | 136970000 | 4 | 4 | 35,8 | 35,8 | 35,8 | 13,93 | 0 | 92,002 | 1389300000 | 27 | AJQ91038.1_1302 | RM25_1327 | Cytoplasmic | 0 | CYT | 0.375369 | OTHER | J |
| lcl\|CP010341.1_prot_AJQ91039.1_1303 | RM25_RS06540 | 50S ribosomal protein L35 | 28770000 | 38472000 | 371640000 | 68899000 | 311360000 | 455009984 | 2 | 2 | 20,3 | 20,3 | 20,3 | 7,1424 | 0 | 47,067 | 1542800000 | 11 | AJQ91039.1_1303 | RM25_1328 | Cytoplasmic | 0 | CYT | 0.865452 | OTHER | J |
| lcl\|CP010341.1_prot_AJQ91040.1_1304 | RM25_RS06545 | Translation initiation factor IF-3 | 113720000 | 149300000 | 331400000 | 262720000 | 209810000 | 393560000 | 9 | 9 | 39,3 | 39,3 | 39,3 | 27,638 | 0 | 176,21 | 1689700000 | 48 | AJQ91040.1_1304 | RM25_1329 | Cytoplasmic | 0 | CYT | 0.102758 | OTHER | J |
| lcl\|CP010341.1_prot_AJQ91042.1_1306 | RM25_RS06555 | ATP phosphoribosyltransferase | 35523000 | 20066000 | 76232000 | 58440000 | 44156000 | 89374000 | 5 | 5 | 23,3 | 23,3 | 23,3 | 30,81 | 0 | 43,853 | 365430000 | 12 | AJQ91042.1_1306 | RM25_1331 | Cytoplasmic | 0 | CYT | 0.044567 | OTHER | E |
| lcl\|CP010341.1_prot_AJQ91043.1_1307 | RM25_RS06560 | Phosphoribosyl-ATP diphosphatase | 47994000 | 69021000 | 103520000 | 51615000 | 37474000 | 62674000 | 3 | 3 | 34,1 | 34,1 | 34,1 | 9,8672 | 0 | 35,683 | 380600000 | 18 | AJQ91043.1_1307 | RM25_1332 | Cytoplasmic | 0 | CYT | 0.028290 | OTHER | |
| lcl\|CP010341.1_prot_AJQ91044.1_1308 | RM25_RS06565 | ATP-dependent zinc metalloprotease FtsH | 27085000 | 26031000 | 65641000 | 27751000 | 34547000 | 53131000 | 9 | 9 | 18 | 18 | 18 | 77,504 | 0 | 96,621 | 253680000 | 21 | AJQ91044.1_1308 | RM25_1333 | CytoplasmicMembrane | 2 | CYT | 0.613950 | OTHER | O |
| lcl\|CP010341.1_prot_AJQ91045.1_1309 | RM25_RS06570 | L-cysteine:1D-myo-inositol 2-amino-2-deoxy-alpha-D-gl | 24490000 | 26325000 | 69362000 | 38407000 | 40150000 | 54517000 | 3 | 3 | 8,9 | 8,9 | 8,9 | 46,019 | 0 | 24,051 | 274660000 | 8 | AJQ91045.1_1309 | RM25_1334 | Cytoplasmic | 0 | CYT | 0.109855 | OTHER | J |
| lcl\|CP010341.1_prot_AJQ91051.1_1315 | RM25_RS06610 | Hypothetical protein | 46970000 | 61432000 | 122740000 | 66833000 | 53424000 | 57071000 | 4 | 4 | 44,5 | 44,5 | 44,5 | 13,8 | 0 | 37,812 | 459050000 | 12 | AJQ91051.1_1315 | RM25_1341 | CytoplasmicMembrane | 0 | CYT | 0.084444 | OTHER | S |

| | | | | | | | | | | | | | | | | | | | | | | | | |
|---|---|---|---|---|---|---|---|---|---|---|---|---|---|---|---|---|---|---|---|---|---|---|---|---|
| lcl\|CP010341.1_prot_AJQ91052.1_1316 | RM25_RS06615 | Pyridoxal-phosphate dependent TrpB-like enzyme | 5650300 | 3559800 | 15835000 | 0 | 14747000 | 0 | 1 | 1 | 2,7 | 2,7 | 2,7 | 46,941 | 0,0042827 | 7,5305 | 39792000 | 2 | AJQ91052.1_1316 | RM25_1342 | Cytoplasmic | 0 | CYT | 0.162059 | OTHER | E |
| lcl\|CP010341.1_prot_AJQ91056.1_1320 | RM25_RS06635 | Protein often found in Actinomycetes clustered with si | 9426400 | 9645100 | 21035000 | 14538000 | 12989000 | 23843000 | 3 | 3 | 25 | 25 | 25 | 12,294 | 0 | 21,041 | 99208000 | 7 | AJQ91056.1_1320 | RM25_1346 | Cytoplasmic | 0 | CYT | 0.073258 | OTHER | S |
| lcl\|CP010341.1_prot_AJQ91059.1_1323 | RM25_RS06650 | 50S ribosomal protein L19 | 193630000 | 224800000 | 718409984 | 525710016 | 526729984 | 883379968 | 5 | 5 | 39 | 39 | 39 | 13,255 | 0 | 45,124 | 3456200000 | 35 | AJQ91059.1_1323 | RM25_1349 | Cytoplasmic | 0 | CYT | 0.049147 | OTHER | J |
| lcl\|CP010341.1_prot_AJQ91060.1_1324 | RM25_RS06655 | Succinate dehydrogenase/fumarate reductase iron-sulf | 114870000 | 84133000 | 249760000 | 350880000 | 297510016 | 411220000 | 6 | 6 | 32,7 | 32,7 | 32,7 | 27,133 | 0 | 60,026 | 1793600000 | 30 | AJQ91060.1_1324 | RM25_1350 | Cytoplasmic | 0 | CYT | 0.106597 | OTHER | C |
| lcl\|CP010341.1_prot_AJQ91061.1_1325 | RM25_RS06660 | Succinate dehydrogenase or fumarate reductase, flavop | 246510000 | 168350000 | 597740032 | 677769984 | 972569984 | 1196000000 | 18 | 18 | 28,1 | 28,1 | 28,1 | 74,809 | 0 | 323,31 | 4302900000 | 84 | AJQ91061.1_1325 | RM25_1351 | Cytoplasmic | 0 | CYT | 0.170794 | OTHER | C |
| lcl\|CP010341.1_prot_AJQ91066.1_1330 | RM25_RS06685 | UPF0109 protein | 20340000 | 29409000 | 56726000 | 36814000 | 35086000 | 83051000 | 3 | 3 | 52,4 | 52,4 | 52,4 | 9,5538 | 0 | 41,038 | 281140000 | 14 | AJQ91066.1_1330 | RM25_1356 | Cytoplasmic | 0 | CYT | 0.047103 | OTHER | S |
| lcl\|CP010341.1_prot_AJQ91067.1_1331 | RM25_RS06690 | 30S ribosomal protein S16 | 357120000 | 445340000 | 1140000000 | 934350016 | 912289984 | 1358099968 | 6 | 6 | 46 | 46 | 46 | 17,753 | 0 | 229,49 | 5872900000 | 52 | AJQ91067.1_1331 | RM25_1357 | Cytoplasmic | 0 | CYT | 0.879160 | OTHER | J |
| lcl\|CP010341.1_prot_AJQ91069.1_1333 | RM25_RS06700 | Signal recognition particle protein | 205530000 | 222310000 | 514590016 | 282529984 | 366129984 | 417310016 | 19 | 19 | 48 | 48 | 48 | 56,292 | 0 | 184,88 | 2239500000 | 67 | AJQ91069.1_1333 | RM25_1359 | CytoplasmicMembrane | 0 | CYT | 0.075722 | OTHER | U |
| lcl\|CP010341.1_prot_AJQ91070.1_1334 | RM25_RS06705 | Signal recognition particle receptor FtsY | 131590000 | 92882000 | 237520000 | 111690000 | 150850000 | 176180000 | 9 | 9 | 28,4 | 28,4 | 28,4 | 39,979 | 0 | 85,589 | 957110000 | 36 | AJQ91070.1_1334 | RM25_1360 | CytoplasmicMembrane | 1 | SPII | 0.388480 | LIPO(Sec/SPII) | U |
| lcl\|CP010341.1_prot_AJQ91076.1_1340 | RM25_RS06735 | Ribonuclease III | 3418200 | 4550900 | 11207000 | 5692800 | 6221600 | 8117200 | 3 | 3 | 17,5 | 17,5 | 17,5 | 25,997 | 0 | 20,641 | 39208000 | 4 | AJQ91076.1_1340 | RM25_1366 | Cytoplasmic | 0 | CYT | 0.085617 | OTHER | K |
| lcl\|CP010341.1_prot_AJQ91077.1_1341 | RM25_RS12305 | Hypothetical protein | 45185000 | 53120000 | 127340000 | 95060000 | 92265000 | 99519000 | 1 | 1 | 12,9 | 12,9 | 12,9 | 7,9221 | 0,0041365 | 6,8015 | 578770000 | 5 | AJQ91077.1_1341 | RM25_1367 | Unknown | 0 | SPI | 0.800557 | OTHER | J |
| lcl\|CP010341.1_prot_AJQ91081.1_1345 | RM25_RS06755 | Pyridoxamine 5'-phosphate oxidase family protein | 105470000 | 165690000 | 303180000 | 182840000 | 142700000 | 209120000 | 4 | 4 | 44,3 | 44,3 | 44,3 | 15,18 | 0 | 85,975 | 1266500000 | 16 | AJQ91081.1_1345 | RM25_1371 | Unknown | 0 | CYT | 0.512455 | OTHER | S |
| lcl\|CP010341.1_prot_AJQ91083.1_1347 | RM25_RS06765 | 50S ribosomal protein L28 | 2031000 | 4447300 | 40843000 | 25468000 | 32606000 | 4735400 | 2 | 2 | 27,9 | 27,9 | 27,9 | 6,6758 | 0 | 15,94 | 132080000 | 8 | AJQ91083.1_1347 | RM25_1373 | Cytoplasmic | 0 | CYT | 0.801050 | OTHER | |
| lcl\|CP010341.1_prot_AJQ91084.1_1348 | RM25_RS06770 | Metallo-beta-lactamase family protein | 56020000 | 47666000 | 135440000 | 88502000 | 109240000 | 168770000 | 7 | 7 | 14,9 | 14,9 | 14,9 | 63,381 | 0 | 61,463 | 668940000 | 17 | AJQ91084.1_1348 | RM25_1374 | Cytoplasmic | 0 | CYT | 0.046565 | OTHER | S |
| lcl\|CP010341.1_prot_AJQ91085.1_1349 | RM25_RS06775 | Putative GCN5-related N-acetyltransferase | 15232000 | 22889000 | 33774000 | 2431300 | 4443800 | 8278100 | 3 | 3 | 19,1 | 19,1 | 19,1 | 17,643 | 0 | 23,396 | 87048000 | 9 | AJQ91085.1_1349 | RM25_1375 | Cytoplasmic | 0 | CYT | 0.097150 | OTHER | K |
| lcl\|CP010341.1_prot_AJQ91087.1_1351 | RM25_RS06785 | Dihydrodipicolinate reductase | 8870000 | 6694300 | 26526000 | 0 | 6166700 | 3218000 | 2 | 2 | 9,3 | 9,3 | 9,3 | 26,23 | 0,0011628 | 10,656 | 51475000 | 1 | AJQ91087.1_1351 | RM25_1377 | Cytoplasmic | 0 | SPI | 0.044147 | OTHER | E |
| lcl\|CP010341.1_prot_AJQ91088.1_1352 | RM25_RS06790 | Polyribonucleotide nucleotidyltransferase | 398249984 | 299080000 | 1073900032 | 769750016 | 737529984 | 1269600000 | 26 | 26 | 43,8 | 43,8 | 43,8 | 79,493 | 0 | 308,68 | 5052500000 | 128 | AJQ91088.1_1352 | RM25_1378 | Cytoplasmic | 0 | CYT | 0.063865 | OTHER | J |
| lcl\|CP010341.1_prot_AJQ91089.1_1353 | RM25_RS06795 | 30S ribosomal protein S15 | 149450000 | 173660000 | 390260000 | 412200000 | 365750016 | 525300000 | 5 | 5 | 64,4 | 64,4 | 64,4 | 10,163 | 0 | 87,408 | 2347800000 | 25 | AJQ91089.1_1353 | RM25_1379 | Cytoplasmic | 0 | CYT | 0.468293 | OTHER | J |
| lcl\|CP010341.1_prot_AJQ91091.1_1355 | RM25_RS06805 | Glycerol kinase | 54547000 | 55049000 | 181970000 | 69984000 | 65639000 | 99848000 | 8 | 8 | 22 | 22 | 22 | 55,743 | 0 | 57,842 | 552650000 | 19 | AJQ91091.1_1355 | RM25_1381 | Cytoplasmic | 0 | CYT | 0.089731 | OTHER | C |
| lcl\|CP010341.1_prot_AJQ91094.1_1358 | RM25_RS06820 | Ribosome-binding factor A | 32005000 | 29601000 | 74820000 | 22978000 | 40927000 | 39686000 | 6 | 6 | 49,3 | 49,3 | 49,3 | 16,28 | 0 | 58,343 | 265360000 | 9 | AJQ91094.1_1358 | RM25_1384 | Cytoplasmic | 0 | CYT | 0.092623 | OTHER | J |
| lcl\|CP010341.1_prot_AJQ91095.1_1359 | RM25_RS06825 | Translation initiation factor IF-2 | 260460000 | 302700000 | 740350016 | 374320000 | 389689984 | 765800000 | 20 | 20 | 27,9 | 27,9 | 27,9 | 101,91 | 0 | 244,53 | 3146300000 | 93 | AJQ91095.1_1359 | RM25_1385 | Cytoplasmic | 0 | CYT | 0.166711 | OTHER | J |
| lcl\|CP010341.1_prot_AJQ91097.1_1361 | RM25_RS06835 | Transcription termination factor NusA | 24985000 | 17021000 | 86278000 | 26391000 | 18348000 | 17215000 | 5 | 5 | 19,3 | 19,3 | 19,3 | 38,234 | 0 | 34,148 | 196620000 | 5 | AJQ91097.1_1361 | RM25_1387 | Cytoplasmic | 0 | CYT | 0.078389 | OTHER | K |
| lcl\|CP010341.1_prot_AJQ91098.1_1362 | RM25_RS06840 | Ribosome maturation factor RimP | 26194000 | 51231000 | 116700000 | 67235000 | 72831000 | 118770000 | 6 | 6 | 58 | 58 | 58 | 17,212 | 0 | 84,764 | 534970000 | 17 | AJQ91098.1_1362 | RM25_1388 | Cytoplasmic | 0 | CYT | 0.093418 | OTHER | S |
| lcl\|CP010341.1_prot_AJQ91099.1_1363 | RM25_RS06845 | Hypothetical protein | 80460000 | 129290000 | 187930000 | 23599000 | 53736000 | 52921000 | 3 | 3 | 12,5 | 12,5 | 12,5 | 33,422 | 0 | 70,957 | 558680000 | 12 | AJQ91099.1_1363 | RM25_1389 | Unknown | 0 | SPII | 0.185645 | LIPO(Sec/SPII) | |
| lcl\|CP010341.1_prot_AJQ91100.1_1364 | RM25_RS06850 | Proline--tRNA ligase | 88781000 | 139620000 | 239810000 | 67310000 | 82878000 | 194260000 | 7 | 7 | 15,2 | 15,2 | 15,2 | 66,084 | 0 | 60,772 | 833260000 | 17 | AJQ91100.1_1364 | RM25_1390 | Cytoplasmic | 0 | CYT | 0.063078 | OTHER | J |
| lcl\|CP010341.1_prot_AJQ91102.1_1366 | RM25_RS06860 | 4-hydroxy-3-methylbut-2-en-1-yl diphosphate synthase | 31471000 | 40458000 | 89794000 | 64637000 | 60875000 | 104260000 | 8 | 8 | 28,6 | 28,6 | 28,6 | 41,12 | 0 | 58,342 | 432150000 | 21 | AJQ91102.1_1366 | RM25_1392 | Cytoplasmic | 0 | CYT | 0.043282 | OTHER | I |
| lcl\|CP010341.1_prot_AJQ91103.1_1367 | RM25_RS06865 | 1-deoxy-D-xylulose 5-phosphate reductoisomerase | 0 | 4891400 | 17840000 | 0 | 7099800 | 10523000 | 1 | 1 | 3,8 | 3,8 | 3,8 | 42,164 | 0,0051493 | 6,6798 | 40354000 | 2 | AJQ91103.1_1367 | RM25_1393 | Cytoplasmic | 0 | CYT | 0.061421 | OTHER | I |
| lcl\|CP010341.1_prot_AJQ91106.1_1370 | RM25_RS06880 | Ribosome-recycling factor | 127760000 | 246880000 | 431889984 | 220540000 | 216170000 | 279489984 | 8 | 8 | 38,4 | 38,4 | 38,4 | 20,51 | 0 | 100,29 | 1691900000 | 29 | AJQ91106.1_1370 | RM25_1396 | Cytoplasmic | 0 | CYT | 0.065169 | OTHER | J |
| lcl\|CP010341.1_prot_AJQ91107.1_1371 | RM25_RS06885 | UMP kinase | 12149000 | 11703000 | 48208000 | 49219000 | 12737000 | 23532000 | 3 | 3 | 14,9 | 14,9 | 14,9 | 27,017 | 0 | 27,855 | 189160000 | 6 | AJQ91107.1_1371 | RM25_1397 | Cytoplasmic | 0 | CYT | 0.068569 | OTHER | F |
| lcl\|CP010341.1_prot_AJQ91108.1_1372 | RM25_RS06890 | Translation elongation factor Ts | 559590016 | 639630016 | 1591000064 | 785769984 | 1016400000 | 1676499968 | 9 | 9 | 40,7 | 40,7 | 40,7 | 28,943 | 0 | 284,51 | 7045200000 | 92 | AJQ91108.1_1372 | RM25_1398 | Cytoplasmic | 0 | CYT | 0.041287 | OTHER | J |
| lcl\|CP010341.1_prot_AJQ91109.1_1373 | RM25_RS06895 | 30S ribosomal protein S2 | 497480000 | 503150016 | 1314300032 | 688400000 | 965000000 | 1362599936 | 14 | 14 | 47,1 | 47,1 | 47,1 | 35,602 | 0 | 323,31 | 5955400000 | 100 | AJQ91109.1_1373 | RM25_1399 | Cytoplasmic | 0 | CYT | 0.096870 | OTHER | J |
| lcl\|CP010341.1_prot_AJQ91112.1_1376 | RM25_RS06915 | Peptide deformylase | 19344000 | 20368000 | 67638000 | 0 | 28454000 | 34269000 | 2 | 2 | 12,1 | 12,1 | 12,1 | 22,715 | 0 | 16,769 | 188300000 | 5 | AJQ91112.1_1376 | RM25_1402 | Unknown | 0 | CYT | 0.128958 | OTHER | J |
| lcl\|CP010341.1_prot_AJQ91115.1_1379 | RM25_RS06930 | ABC-type molybdenum transport system, ATPase comp | 43203000 | 16416000 | 85890000 | 49801000 | 59951000 | 71571000 | 6 | 6 | 34,8 | 34,8 | 34,8 | 28,891 | 0 | 48,345 | 349750000 | 8 | AJQ91115.1_1379 | RM25_1405 | CytoplasmicMembrane | 0 | CYT | 0.045073 | OTHER | P |
| lcl\|CP010341.1_prot_AJQ91117.1_1381 | RM25_RS06940 | Enoyl-acyl-carrier-protein reductase (NADH) | 30060000 | 26317000 | 71144000 | 43612000 | 48737000 | 59818000 | 3 | 3 | 21 | 21 | 21 | 26,855 | 0 | 34,733 | 294660000 | 7 | AJQ91117.1_1381 | RM25_1407 | CytoplasmicMembrane | 0 | CYT | 0.045006 | OTHER | I |
| lcl\|CP010341.1_prot_AJQ91118.1_1382 | RM25_RS06945 | 3-oxoacyl-acyl-carrier protein reductase fabG1 | 1994100 | 1043800 | 3090700 | 7476600 | 9598200 | 7048400 | 1 | 1 | 6,9 | 6,9 | 6,9 | 24,295 | 0,0044593 | 8,504 | 31440000 | 2 | AJQ91118.1_1382 | RM25_1408 | Cytoplasmic | 0 | CYT | 0.044338 | OTHER | IQ |
| lcl\|CP010341.1_prot_AJQ91119.1_1383 | RM25_RS06950 | Short-chain dehydrogenase/reductase SDR | 9518500 | 7635700 | 19356000 | 0 | 6448900 | 9922000 | 2 | 2 | 7,3 | 7,3 | 7,3 | 25,302 | 0 | 11,628 | 52882000 | 2 | AJQ91119.1_1383 | RM25_1409 | Cytoplasmic | 0 | SPI | 0.028921 | SP(Sec/SPI) | IQ |
| lcl\|CP010341.1_prot_AJQ91121.1_1385 | RM25_RS06960 | Macrolide ABC transporter ATP-binding protein | 16380000 | 17020000 | 57732000 | 29079000 | 20723000 | 45425000 | 5 | 5 | 13 | 13 | 13 | 57,747 | 0 | 37,527 | 201230000 | 13 | AJQ91121.1_1385 | RM25_1411 | Cytoplasmic | 0 | CYT | 0.063686 | OTHER | S |
| lcl\|CP010341.1_prot_AJQ91122.1_1386 | RM25_RS06965 | Putative aromatic ring hydroxylating enzyme | 33592000 | 44557000 | 58061000 | 77877000 | 63707000 | 75914000 | 2 | 2 | 34,9 | 34,9 | 34,9 | 16,409 | 0 | 31,185 | 432370000 | 8 | AJQ91122.1_1386 | RM25_1412 | Cytoplasmic | 0 | CYT | 0.540416 | OTHER | S |
| lcl\|CP010341.1_prot_AJQ91123.1_1387 | RM25_RS06970 | SUF system FeS assembly protein, NifU family | 27072000 | 34859000 | 59401000 | 71662000 | 69918000 | 79794000 | 3 | 3 | 32 | 32 | 32 | 16,64 | 0 | 29,46 | 385630000 | 6 | AJQ91123.1_1387 | RM25_1413 | Cytoplasmic | 0 | CYT | 0.093552 | OTHER | C |
| lcl\|CP010341.1_prot_AJQ91124.1_1388 | RM25_RS06975 | Cysteine desulfurase, SufS family protein | 41577000 | 8171700 | 200480000 | 90999000 | 74156000 | 84314000 | 5 | 5 | 15,6 | 15,6 | 15,6 | 45,203 | 0 | 42,74 | 526830000 | 12 | AJQ91124.1_1388 | RM25_1414 | Cytoplasmic | 0 | CYT | 0.067734 | OTHER | E |
| lcl\|CP010341.1_prot_AJQ91125.1_1389 | RM25_RS06980 | FeS assembly ATPase SufC | 241010000 | 214030000 | 838830016 | 458350016 | 545900032 | 625040000 | 8 | 8 | 45,8 | 45,8 | 45,8 | 26,923 | 0 | 239,64 | 3391400000 | 56 | AJQ91125.1_1389 | RM25_1415 | Cytoplasmic | 0 | CYT | 0.079178 | OTHER | O |
| lcl\|CP010341.1_prot_AJQ91126.1_1390 | RM25_RS06985 | Dioxygenase | 249630000 | 345449984 | 338000000 | 312550016 | 326440000 | 488950016 | 3 | 3 | 59,1 | 59,1 | 59,1 | 12,379 | 0 | 180,01 | 2225400000 | 29 | AJQ91126.1_1390 | RM25_1416 | Cytoplasmic | 0 | CYT | 0.230596 | OTHER | |
| lcl\|CP010341.1_prot_AJQ91127.1_1391 | RM25_RS06990 | FeS assembly protein SufD | 153610000 | 77639000 | 555630016 | 308840000 | 205160000 | 395049984 | 11 | 11 | 30,9 | 30,9 | 30,9 | 44,269 | 0 | 216,38 | 1835400000 | 46 | AJQ91127.1_1391 | RM25_1417 | Cytoplasmic | 0 | CYT | 0.070762 | OTHER | |
| lcl\|CP010341.1_prot_AJQ91128.1_1392 | RM25_RS06995 | FeS assembly protein SufB | 117020000 | 68253000 | 335470016 | 219160000 | 243290000 | 303180000 | 13 | 13 | 39,6 | 39,6 | 39,6 | 53,956 | 0 | 158,28 | 1390800000 | 42 | AJQ91128.1_1392 | RM25_1418 | Unknown | 0 | CYT | 0.250184 | OTHER | O |
| lcl\|CP010341.1_prot_AJQ91133.1_1397 | RM25_RS07020 | Glutamine amidotransferase subunit PdxT | 3685400 | 5602500 | 8965100 | 0 | 0 | 0 | 1 | 1 | 5,8 | 5,8 | 5,8 | 21,169 | 0,0042918 | 7,5317 | 18253000 | 2 | AJQ91133.1_1397 | RM25_1423 | Cytoplasmic | 0 | CYT | 0.024016 | OTHER | |
| lcl\|CP010341.1_prot_AJQ91134.1_1398 | RM25_RS07025 | Pyridoxal biosynthesis lyase PdxS | 80964000 | 33075000 | 229110000 | 126790000 | 223680000 | 377880000 | 8 | 8 | 27 | 27 | 27 | 31,349 | 0 | 74,946 | 1192900000 | 28 | AJQ91134.1_1398 | RM25_1424 | Cytoplasmic | 0 | CYT | 0.037839 | OTHER | H |
| lcl\|CP010341.1_prot_AJQ91136.1_1400 | RM25_RS07035 | Glucose-6-phosphate 1-dehydrogenase Zwf | 3210900 | 1546700 | 8928300 | 6342800 | 4400300 | 17754000 | 2 | 2 | 4,5 | 4,5 | 4,5 | 59,604 | 0 | 11,185 | 54874000 | 3 | AJQ91136.1_1400 | RM25_1426 | Cytoplasmic | 0 | CYT | 0.203076 | OTHER | G |
| lcl\|CP010341.1_prot_AJQ91137.1_1401 | RM25_RS07040 | Putative OpcA protein | 8800800 | 2746400 | 42566000 | 7432700 | 14213000 | 36211000 | 3 | 3 | 12 | 12 | 12 | 32,237 | 0 | 24,989 | 119350000 | 5 | AJQ91137.1_1401 | RM25_1427 | CytoplasmicMembrane | 0 | CYT | 0.045389 | TAT(Tat/SPI) | G |
| lcl\|CP010341.1_prot_AJQ91139.1_1403 | RM25_RS07050 | Electron transport protein | 23560000 | 32856000 | 56123000 | 26693000 | 43999000 | 53610000 | 2 | 2 | 21,2 | 21,2 | 21,2 | 12,989 | 0 | 17,166 | 255380000 | 14 | AJQ91139.1_1403 | RM25_1429 | Unknown | 0 | CYT | 0.497848 | OTHER | |
| lcl\|CP010341.1_prot_AJQ91141.1_1405 | RM25_RS07060 | Triosephosphate isomerase 2 | 48820000 | 22472000 | 217250000 | 136170000 | 107220000 | 146840000 | 4 | 4 | 18,7 | 18,7 | 18,7 | 27,619 | 0 | 189,23 | 749940000 | 30 | AJQ91141.1_1405 | RM25_1431 | Cytoplasmic | 0 | CYT | 0.048658 | OTHER | G |
| lcl\|CP010341.1_prot_AJQ91142.1_1406 | RM25_RS07065 | Phosphoglycerate kinase | 811009984 | 838070016 | 1930000000 | 661249984 | 701960000 | 1054099968 | 19 | 19 | 62,3 | 62,3 | 62,3 | 42,436 | 0 | 323,31 | 6375000000 | 98 | AJQ91142.1_1406 | RM25_1432 | Cytoplasmic | 0 | CYT | 0.045628 | OTHER | G |
| lcl\|CP010341.1_prot_AJQ91143.1_1407 | RM25_RS07070 | Glyceraldehyde-3-phosphate dehydrogenase, type I | 823960000 | 157820000 | 3060600064 | 1689500032 | 1272400000 | 1875399936 | 14 | 14 | 57,5 | 57,5 | 57,5 | 36,112 | 0 | 323,31 | 9712800000 | 111 | AJQ91143.1_1407 | RM25_1433 | Cytoplasmic | 0 | CYT | 0.068292 | OTHER | G |
| lcl\|CP010341.1_prot_AJQ91145.1_1409 | RM25_RS07080 | ABC transporter, ATP-binding protein | 1455100 | 2395200 | 8481200 | 0 | 0 | 3232600 | 1 | 1 | 4,1 | 4,1 | 4,1 | 24,983 | 0,0050865 | 6,4453 | 15564000 | 1 | AJQ91145.1_1409 | RM25_1435 | CytoplasmicMembrane | 0 | CYT | 0.058895 | OTHER | |
| lcl\|CP010341.1_prot_AJQ91146.1_1410 | RM25_RS07085 | Putative sporulation transcription regulator WhiA | 5075300 | 4616600 | 12471000 | 10086000 | 6120500 | 11524000 | 1 | 1 | 4,3 | 4,3 | 4,3 | 35,419 | 0,004415 | 8,2923 | 59824000 | 2 | AJQ91146.1_1410 | RM25_1436 | Cytoplasmic | 0 | CYT | 0.033961 | OTHER | K |
| lcl\|CP010341.1_prot_AJQ91148.1_1412 | RM25_RS07095 | UPF0042 nucleotide-binding protein | 2473800 | 0 | 10299000 | 9417600 | 7234900 | 13964000 | 2 | 2 | 7,1 | 7,1 | 7,1 | 32,524 | 0 | 11,107 | 52612000 | 2 | AJQ91148.1_1412 | RM25_1438 | Cytoplasmic | 0 | CYT | 0.066565 | OTHER | S |
| lcl\|CP010341.1_prot_AJQ91153.1_1417 | RM25_RS07120 | ABC transporter (Permease) (Na+ exclusion) | 194240000 | 349240000 | 519740000 | 211970000 | 64456000 | 364030016 | 3 | 3 | 16,2 | 16,2 | 16,2 | 41,493 | 0 | 202,88 | 1936000000 | 25 | AJQ91153.1_1417 | RM25_1443 | CytoplasmicMembrane | 6 | SPI | 0.894005 | OTHER | CP |
| lcl\|CP010341.1_prot_AJQ91155.1_1419 | RM25_RS07130 | UvrABC system protein A | 5505700 | 5606300 | 15639000 | 10638000 | 13086000 | 24439000 | 3 | 3 | 4,1 | 4,1 | 4,1 | 111,75 | 0 | 15,686 | 85079000 | 3 | AJQ91155.1_1419 | RM25_1445 | Cytoplasmic | 0 | CYT | 0.130224 | OTHER | L |
| lcl\|CP010341.1_prot_AJQ91157.1_1421 | RM25_RS07140 | Metallo-beta-lactamase domain protein | 12258000 | 28309000 | 41978000 | 24702000 | 18538000 | 15470000 | 2 | 2 | 14,2 | 14,2 | 14,2 | 20,992 | 0 | 16,425 | 170170000 | 9 | AJQ91157.1_1421 | RM25_1447 | Unknown | 0 | CYT | 0.132074 | OTHER | S |
| lcl\|CP010341.1_prot_AJQ91159.1_1423 | RM25_RS07150 | UvrABC system protein B | 1167600 | 1702300 | 4758000 | 3456600 | 5165900 | 0 | 2 | 2 | 3,2 | 3,2 | 3,2 | 78,388 | 0 | 11,727 | 20229000 | 3 | AJQ91159.1_1423 | RM25_1449 | Cytoplasmic | 0 | CYT | 0.089946 | OTHER | L |
| lcl\|CP010341.1_prot_AJQ91161.1_1425 | RM25_RS07160 | Phosphorylase, Pnp/Udp family | 53641000 | 60257000 | 124660000 | 87939000 | 121170000 | 133390000 | 6 | 6 | 32,9 | 32,9 | 32,9 | 28,263 | 0 | 62,046 | 637750000 | 17 | AJQ91161.1_1425 | RM25_1451 | Cytoplasmic | 0 | CYT | 0.130119 | OTHER | |
| lcl\|CP010341.1_prot_AJQ91162.1_1426 | RM25_RS07165 | 30S ribosomal protein S1 | 492830016 | 402830016 | 1040700032 | 360169984 | 413420000 | 958710016 | 16 | 16 | 50,4 | 50,4 | 50,4 | 53,615 | 0 | 323,31 | 4009700000 | 100 | AJQ91162.1_1426 | RM25_1452 | Cytoplasmic | 0 | CYT | 0.099100 | OTHER | J |
| lcl\|CP010341.1_prot_AJQ91163.1_1427 | RM25_RS07170 | Succinate-semialdehyde dehydrogenase | 73380000 | 49928000 | 275560000 | 137990000 | 129060000 | 221930000 | 9 | 9 | 25,6 | 25,6 | 25,6 | 52,247 | 0 | 104,13 | 955810000 | 34 | AJQ91163.1_1427 | RM25_1453 | Cytoplasmic | 0 | CYT | 0.072095 | OTHER | C |
| lcl\|CP010341.1_prot_AJQ91165.1_1429 | RM25_RS07180 | DNA-directed DNA polymerase | 11771000 | 8227300 | 31043000 | 30954000 | 27334000 | 37168000 | 5 | 5 | 7,5 | 7,5 | 7,5 | 98,65 | 0 | 36,485 | 169590000 | 7 | AJQ91165.1_1429 | RM25_1455 | Cytoplasmic | 0 | CYT | 0.100163 | OTHER | L |
| lcl\|CP010341.1_prot_AJQ91166.1_1430 | RM25_RS07185 | Putative transcriptional regulatory protein pdtaR | 128370000 | 196430000 | 390280000 | 224350000 | 286580000 | 325049984 | 7 | 7 | 53,1 | 53,1 | 53,1 | 22,601 | 0 | 137,75 | 1757800000 | 54 | AJQ91166.1_1430 | RM25_1456 | Cytoplasmic | 0 | CYT | 0.031189 | OTHER | |
| lcl\|CP010341.1_prot_AJQ91167.1_1431 | RM25_RS07195 | Pyruvate kinase | 171140000 | 153020000 | 454740000 | 194810000 | 280310016 | 499040000 | 11 | 11 | 31,7 | 31,7 | 31,7 | 51,555 | 0 | 119,95 | 1878100000 | 44 | AJQ91167.1_1431 | RM25_1458 | Cytoplasmic | 0 | CYT | 0.070797 | OTHER | G |
| lcl\|CP010341.1_prot_AJQ91168.1_1432 | RM25_RS07200 | Pseudouridine synthase (Precursor) | 4424100 | 4264500 | 16130000 | 6296200 | 9196100 | 12999000 | 3 | 3 | 9,2 | 9,2 | 9,2 | 32,844 | 0 | 17,798 | 61384000 | 5 | AJQ91168.1_1432 | RM25_1459 | Cytoplasmic | 0 | CYT | 0.064924 | OTHER | J |
| lcl\|CP010341.1_prot_AJQ91170.1_1434 | RM25_RS07210 | C4-type zinc finger protein, DksA/TraR family | 144870000 | 307440000 | 262080000 | 274820000 | 312350016 | 346120000 | 5 | 5 | 57 | 57 | 57 | 15,752 | 0 | 140,14 | 1896700000 | 31 | AJQ91170.1_1434 | RM25_1461 | Cytoplasmic | 0 | CYT | 0.066587 | OTHER | T |
| lcl\|CP010341.1_prot_AJQ91171.1_1435 | RM25_RS07215 | DivIVA domain protein | 643320000 | 808920000 | 1913500032 | 945960000 | 1429799936 | 1553699968 | 18 | 18 | 58,4 | 58,4 | 58,4 | 36,527 | 0 | 323,31 | 8024300000 | 130 | AJQ91171.1_1435 | RM25_1462 | Cytoplasmic | 0 | CYT | 0.487521 | OTHER | |
| lcl\|CP010341.1_prot_AJQ91173.1_1437 | RM25_RS07225 | Cell division protein sepF | 13117000 | 19726000 | 37852000 | 34827000 | 30754000 | 38277000 | 4 | 4 | 30,6 | 30,6 | 30,6 | 18,854 | 0 | 125,37 | 198140000 | 15 | AJQ91173.1_1437 | RM25_1464 | Cytoplasmic | 0 | CYT | 0.107297 | OTHER | D |
| lcl\|CP010341.1_prot_AJQ91175.1_1439 | RM25_RS07235 | Cell division protein FtsZ | 138370000 | 241740000 | 490710016 | 252400000 | 265880000 | 405880000 | 14 | 14 | 48,4 | 48,4 | 48,4 | 42,787 | 0 | 306,2 | 2003100000 | 51 | AJQ91175.1_1439 | RM25_1466 | Cytoplasmic | 0 | CYT | 0.092938 | OTHER | D |
| lcl\|CP010341.1_prot_AJQ91176.1_1440 | RM25_RS07240 | POTRA domain protein, FtsQ-type | 860329984 | 1327900032 | 1580499968 | 486569984 | 801790016 | 1282400000 | 5 | 5 | 43,6 | 43,6 | 43,6 | 25,981 | 0 | 191,38 | 6834000000 | 46 | AJQ91176.1_1440 | RM25_1467 | CytoplasmicMembrane | 1 | TMH | 0.635591 | OTHER | |
| lcl\|CP010341.1_prot_AJQ91177.1_1441 | RM25_RS07245 | UDP-N-acetylmuramate--L-alanine ligase | 16823000 | 14576000 | 36475000 | 5533300 | 17580000 | 19403000 | 4 | 4 | 10,8 | 10,8 | 10,8 | 51,208 | 0 | 25,654 | 113480000 | 9 | AJQ91177.1_1441 | RM25_1468 | Cytoplasmic | 0 | CYT | 0.067919 | OTHER | M |
| lcl\|CP010341.1_prot_AJQ91178.1_1442 | RM25_RS07250 | UDP-N-acetylglucosamine--N-acetylmuramyl-(pentapep | 8390700 | 6559000 | 27649000 | 18426000 | 20228000 | 29772000 | 3 | 3 | 8,7 | 8,7 | 8,7 | 38,059 | 0 | 22,662 | 125120000 | 10 | AJQ91178.1_1442 | RM25_1469 | CytoplasmicMembrane | 0 | CYT | 0.039555 | OTHER | M |
| lcl\|CP010341.1_prot_AJQ91179.1_1443 | RM25_RS07255 | Cell division protein FtsW | 1325300 | 1348300 | 0 | 0 | 1382100 | 1713900 | 1 | 1 | 3,3 | 3,3 | 3,3 | 44,21 | 0,004171 | 6,9043 | 5769600 | 1 | AJQ91179.1_1443 | RM25_1470 | CytoplasmicMembrane | 10 | CYT | 0.967586 | OTHER | |
| lcl\|CP010341.1_prot_AJQ91180.1_1444 | RM25_RS07260 | UDP-N-acetylmuramoyl-L-alanine--D-glutamate ligase | 30935000 | 33949000 | 76838000 | 81535000 | 55387000 | 81919000 | 5 | 5 | 11,8 | 11,8 | 11,8 | 50,178 | 0 | 32,757 | 461940000 | 15 | AJQ91180.1_1444 | RM25_1471 | Cytoplasmic | 0 | CYT | 0.054970 | OTHER | M |
| lcl\|CP010341.1_prot_AJQ91183.1_1447 | RM25_RS07275 | UDP-N-acetylmuramoyl-L-alanyl-D-glutamate--2,6-diam | 15828000 | 25899000 | 44522000 | 9998200 | 33503000 | 46897000 | 6 | 6 | 17,4 | 17,4 | 17,4 | 55,429 | 0 | 44,679 | 185770000 | 8 | AJQ91183.1_1447 | RM25_1474 | Cytoplasmic | 0 | CYT | 0.083079 | OTHER | M |
| lcl\|CP010341.1_prot_AJQ91184.1_1448 | RM25_RS07280 | Penicillin binding protein transpeptidase domain protei | 2240900096 | 4623200256 | 5744600064 | 1993900032 | 3280100096 | 4116900096 | 18 | 18 | 40,9 | 40,9 | 40,9 | 71,889 | 0 | 323,31 | 2,4216E+10 | 143 | AJQ91184.1_1448 | RM25_1475 | CytoplasmicMembrane | 1 | CYT | 0.913901 | OTHER | M |
| lcl\|CP010341.1_prot_AJQ91185.1_1449 | RM25_RS12605 | Hypothetical protein | 36760000 | 73043000 | 55406000 | 0 | 0 | 0 | 1 | 1 | 14 | 14 | 14 | 17,738 | 0 | 20,943 | 177010000 | 1 | AJQ91185.1_1449 | RM25_1476 | Unknown | 1 | CYT | 0.927041 | OTHER | |
| lcl\|CP010341.1_prot_AJQ91186.1_1450 | RM25_RS07290 | Ribosomal RNA small subunit methyltransferase H | 5171500 | 8817000 | 12469000 | 6699300 | 8327800 | 10300000 | 1 | 1 | 2 | 2 | 2 | 37,054 | 0,0045558 | 10,023 | 51785000 | 6 | AJQ91186.1_1450 | RM25_1477 | Cytoplasmic | 0 | CYT | 0.045109 | OTHER | M |
| lcl\|CP010341.1_prot_AJQ91190.1_1454 | RM25_RS07310 | DNA polymerase III, alpha subunit | 5850000 | 15227000 | 58676000 | 29093000 | 30034000 | 86150000 | 2 | 2 | 1,6 | 1,6 | 1,6 | 130,54 | 0 | 11,039 | 252080000 | 3 | AJQ91190.1_1454 | RM25_1481 | Cytoplasmic | 0 | CYT | 0.070506 | OTHER | |
| lcl\|CP010341.1_prot_AJQ91192.1_1456 | RM25_RS07320 | DNA or RNA helicase | 18525000 | 17166000 | 32030000 | 12828000 | 25349000 | 44784000 | 4 | 4 | 6,9 | 6,9 | 6,9 | 81,178 | 0 | 30,298 | 160550000 | 11 | AJQ91192.1_1456 | RM25_1483 | Cytoplasmic | 0 | CYT | 0.062244 | OTHER | L |
| lcl\|CP010341.1_prot_AJQ91194.1_1458 | RM25_RS07330 | Putative GERANYLGERANYL PYROPHOSPHATE SYNTHET | 4980800 | 9012300 | 13919000 | 3399800 | 0 | 8258100 | 3 | 3 | 12,1 | 12,1 | 12,1 | 37,626 | 0 | 27,529 | 48199000 | 7 | AJQ91194.1_1458 | RM25_1485 | Cytoplasmic | 0 | CYT | 0.053074 | OTHER | H |
| lcl\|CP010341.1_prot_AJQ91195.1_1459 | RM25_1486 | Hypothetical protein | 257140000 | 495560000 | 591720000 | 399849984 | 521140000 | 828460032 | 7 | 7 | 13,6 | 13,6 | 13,6 | 79,632 | 0 | 159,21 | 3447500000 | 41 | AJQ91195.1_1459 | RM25_1486 | CytoplasmicMembrane | 1 | CYT | 0.456671 | OTHER | KLT |
| lcl\|CP010341.1_prot_AJQ91198.1_1462 | RM25_RS07350 | 3-deoxy-7-phosphoheptulonate synthase | 24525000 | 11965000 | 73883000 | 45406000 | 46417000 | 67300000 | 4 | 4 | 11,2 | 11,2 | 11,2 | 49,549 | 0 | 27,356 | 286270000 | 9 | AJQ91198.1_1462 | RM25_1489 | Cytoplasmic | 0 | CYT | 0.095401 | OTHER | E |
| lcl\|CP010341.1_prot_AJQ91199.1_1463 | RM25_RS11895 | Acyltransferase | 13116000 | 1248500 | 21909000 | 16564000 | 20945000 | 21966000 | 2 | 2 | 6,1 | 6,1 | 6,1 | 34,225 | 0 | 12,5 | 108300000 | 5 | AJQ91199.1_1463 | RM25_1490 | Cytoplasmic | 0 | CYT | 0.096073 | OTHER | |
| lcl\|CP010341.1_prot_AJQ91201.1_1465 | RM25_RS11900 | Peptidase M23B family / metalloendopeptidase | 1080199936 | 1515299968 | 2184999936 | 727840000 | 1193799936 | 1855900032 | 9 | 9 | 37,3 | 37,3 | 37,3 | 39,029 | 0 | 323,31 | 9437200000 | 53 | AJQ91201.1_1465 | RM25_1492 | Extracellular | 1 | SPI | 0.714273 | SP(Sec/SPI) | D |
| lcl\|CP010341.1_prot_AJQ91202.1_1466 | RM25_RS07370 | Transcriptional regulator, AsnC family | 26020000 | 33975000 | 59293000 | 17056000 | 0 | 52471000 | 2 | 2 | 33,7 | 33,7 | 33,7 | 9,7419 | 0 | 21,937 | 201940000 | 7 | AJQ91202.1_1466 | RM25_1493 | Cytoplasmic | 0 | CYT | 0.024414 | OTHER | K |
| lcl\|CP010341.1_prot_AJQ91203.1_1467 | RM25_RS07375 | Peptidyl-prolyl cis-trans isomerase | 741870016 | 1201900032 | 1766099968 | 1607299968 | 1188700032 | 1409600000 | 10 | 10 | 75,3 | 75,3 | 75,3 | 18,771 | 0 | 323,31 | 8890600000 | 80 | AJQ91203.1_1467 | RM25_1494 | Cytoplasmic | 0 | CYT | 0.189089 | OTHER | O |
| lcl\|CP010341.1_prot_AJQ91204.1_1468 | RM25_RS11905 | Putative regulatory protein | 357500000 | 368880000 | 894720000 | 999219968 | 1506200064 | 1179800064 | 15 | 15 | 50,9 | 50,9 | 50,9 | 35,943 | 0 | 270,28 | 5968200000 | 74 | AJQ91204.1_1468 | RM25_1495 | Unknown | 0 | CYT | 0.761253 | OTHER | J |
| lcl\|CP010341.1_prot_AJQ91209.1_1473 | RM25_RS07405 | Anthranilate phosphoribosyltransferase | 7333100 | 12506000 | 29938000 | 0 | 5697200 | 11003000 | 3 | 3 | 9,7 | 9,7 | 9,7 | 36,561 | 0 | 24,312 | 71629000 | 6 | AJQ91209.1_1473 | RM25_1500 | Cytoplasmic | 0 | CYT | 0.055495 | OTHER | E |
| lcl\|CP010341.1_prot_AJQ91210.1_1474 | RM25_RS07410 | Response regulator | 70039000 | 71190000 | 158290000 | 126630000 | 176450000 | 214230000 | 7 | 7 | 47,7 | 47,7 | 47,7 | 18,78 | 0 | 69,687 | 905680000 | 35 | AJQ91210.1_1474 | RM25_1501 | Unknown | 0 | CYT | 0.254348 | OTHER | T |
| lcl\|CP010341.1_prot_AJQ91211.1_1475 | RM25_RS07415 | IMP dehydrogenase family protein | 32049000 | 22577000 | 107990000 | 33625000 | 50290000 | 99208000 | 7 | 7 | 21,4 | 21,4 | 21,4 | 51,624 | 0 | 78,447 | 372880000 | 14 | AJQ91211.1_1475 | RM25_1502 | Cytoplasmic | 0 | CYT | 0.051079 | OTHER | F |
| lcl\|CP010341.1_prot_AJQ91212.1_1476 | RM25_RS07420 | Iron-sulfur cluster assembly accessory protein | 169520000 | 276480000 | 378500000 | 404510016 | 530609984 | 604510016 | 4 | 4 | 39,7 | 39,7 | 39,7 | 12,441 | 0 | 49,256 | 2762100000 | 25 | AJQ91212.1_1476 | RM25_1503 | Cytoplasmic | 0 | CYT | 0.117597 | OTHER | S |
| lcl\|CP010341.1_prot_AJQ91214.1_1478 | RM25_RS07430 | Hypothetical protein | 15602000 | 29338000 | 35353000 | 55312000 | 0 | 0 | 1 | 1 | 12,5 | 12,5 | 12,5 | 10,473 | 0,005144 | 6,6651 | 135600000 | 2 | AJQ91214.1_1478 | RM25_1505 | Cytoplasmic | 0 | CYT | 0.107181 | OTHER | - |
| lcl\|CP010341.1_prot_AJQ91215.1_1479 | RM25_RS07435 | PspA/IM30 family protein | 36297000 | 24595000 | 70766000 | 29235000 | 28565000 | 39356000 | 6 | 6 | 35,2 | 35,2 | 35,2 | 28,742 | 0 | 50,108 | 246740000 | 8 | AJQ91215.1_1479 | RM25_1506 | Cytoplasmic | 0 | CYT | 0.065365 | OTHER | KT |
| lcl\|CP010341.1_prot_AJQ91216.1_1480 | RM25_RS07440 | Hypothetical protein | 15670000 | 23329000 | 40439000 | 25355000 | 34742000 | 40777000 | 1 | 1 | 6,5 | 6,5 | 6,5 | 21,459 | 0,0044944 | 9,194 | 204240000 | 7 | AJQ91216.1_1480 | RM25_1507 | CytoplasmicMembrane | 2 | CYT | 0.908663 | OTHER | S |
| lcl\|CP010341.1_prot_AJQ91219.1_1483 | RM25_RS07455 | Hypothetical protein | 318980000 | 189110000 | 765760000 | 483560000 | 386630016 | 628310016 | 14 | 13 | 32,3 | 32,3 | 30,9 | 59,587 | 0 | 323,31 | 3007100000 | 62 | AJQ91219.1_1483 | RM25_1510 | Cytoplasmic | 0 | CYT | 0.320528 | OTHER | C |
| lcl\|CP010341.1_prot_AJQ91220.1_1484 | RM25_RS07460 | Hypothetical protein | 13014000 | 7397400 | 9702600 | 23094000 | 18268000 | 33984000 | 2 | 2 | 10,2 | 10,2 | 10,2 | 28 | 0 | 12,504 | 109620000 | 1 | AJQ91220.1_1484 | RM25_1511 | CytoplasmicMembrane | 2 | CYT | 0.760872 | OTHER | S |
| lcl\|CP010341.1_prot_AJQ91222.1_1486 | RM25_RS07470 | Glutamate--ammonia ligase | 126240000 | 78830000 | 353400000 | 152450000 | 72111000 | 120660000 | 12 | 12 | 36,8 | 36,8 | 36,8 | 53,426 | 0 | 92,444 | 954420000 | 22 | AJQ91222.1_1486 | RM25_1513 | Cytoplasmic | 0 | CYT | 0.283197 | OTHER | E |

| | | | | | | | | | | | | | | | | | | | | | | | | | | |
|---|---|---|---|---|---|---|---|---|---|---|---|---|---|---|---|---|---|---|---|---|---|---|---|---|---|---|
| lcl\|CP010341.1_prot_AJQ91223.1_1487 | RM25_RS07475 | Hypothetical protein | 9308400 | 15921000 | 26722000 | 6003100 | 7086500 | 19186000 | 2 | 2 | 13,8 | 13,8 | 13,8 | 19,346 | 0 | 20,724 | 84227000 | 8 | AJQ91223.1_1487 | RM25_1514 | Unknown | 0 | SPII | 0.426757 | LIPO(Sec/SPII) | |
| lcl\|CP010341.1_prot_AJQ91229.1_1493 | RM25_RS11920 | Hypothetical protein | 14928000 | 51635000 | 28071000 | 864540 | 13538000 | 13522000 | 4 | 4 | 15 | 15 | 15 | 31,344 | 0 | 27,89 | 122560000 | 10 | AJQ91229.1_1493 | RM25_1520 | Unknown | 1 | TMH | 0.955144 | OTHER | |
| lcl\|CP010341.1_prot_AJQ91231.1_1495 | RM25_RS07515 | Methionyl aminopeptidase | 55105000 | 70564000 | 142980000 | 98794000 | 105150000 | 147590000 | 4 | 4 | 12,6 | 12,6 | 12,6 | 33,254 | 0 | 31,472 | 659050000 | 27 | AJQ91231.1_1495 | RM25_1522 | Cytoplasmic | 0 | CYT | 0.174698 | OTHER | J |
| lcl\|CP010341.1_prot_AJQ91232.1_1496 | RM25_RS11930 | S-layer domain protein | 210180000 | 281720000 | 453169984 | 131880000 | 135680000 | 233270000 | 4 | 4 | 24,7 | 24,7 | 24,7 | 23,35 | 0 | 61,992 | 1556700000 | 19 | AJQ91232.1_1496 | RM25_1523 | Unknown | 0 | SPI | 0.431193 | SP(Sec/SPI) | |
| lcl\|CP010341.1_prot_AJQ91234.1_1498 | RM25_RS07525 | Zn-ribbon protein, possibly nucleic acid-binding | 56010000 | 71564000 | 205520000 | 121160000 | 162670000 | 170220000 | 10 | 10 | 50,2 | 50,2 | 50,2 | 27,727 | 0 | 119,49 | 890620000 | 43 | AJQ91234.1_1498 | RM25_1525 | Cytoplasmic | 0 | CYT | 0.057214 | OTHER | S |
| lcl\|CP010341.1_prot_AJQ91235.1_1499 | RM25_RS07530 | Xaa-Pro aminopeptidase | 11557000 | 7314600 | 27523000 | 4409700 | 5483500 | 9630600 | 2 | 2 | 6 | 6 | 6 | 53,445 | 0 | 18,69 | 82998000 | 2 | AJQ91235.1_1499 | RM25_1526 | Cytoplasmic | 0 | CYT | 0.177440 | OTHER | E |
| lcl\|CP010341.1_prot_AJQ91236.1_1500 | RM25_RS12870 | Hypothetical protein | 0 | 0 | 0 | 0 | 34390000 | 26380000 | 1 | 1 | 5,8 | 5,8 | 5,8 | 29,728 | 0,0045045 | 9,2092 | 60770000 | 2 | AJQ91236.1_1500 | RM25_1527 | CytoplasmicMembrane | 2 | CYT | 0.964822 | OTHER | |
| lcl\|CP010341.1_prot_AJQ91237.1_1501 | RM25_RS07540 | Citrate lyase beta subunit | 28659000 | 22575000 | 64860000 | 31011000 | 25219000 | 32874000 | 4 | 4 | 18,1 | 18,1 | 18,1 | 31,11 | 0 | 32,276 | 224060000 | 8 | AJQ91237.1_1501 | RM25_1528 | Cytoplasmic | 0 | CYT | 0.050778 | OTHER | G |
| lcl\|CP010341.1_prot_AJQ91238.1_1502 | RM25_RS07545 | MgtE intracellular region | 4086500 | 4064100 | 8864600 | 14531000 | 11449000 | 7937600 | 2 | 2 | 6,4 | 6,4 | 6,4 | 46,935 | 0 | 12,474 | 68960000 | 4 | AJQ91238.1_1502 | RM25_1529 | Cytoplasmic | 0 | CYT | 0.073664 | OTHER | P |
| lcl\|CP010341.1_prot_AJQ91240.1_1504 | RM25_RS07555 | Putative ATP-binding protein Mrp | 44174000 | 47201000 | 112530000 | 82615000 | 61256000 | 137930000 | 3 | 3 | 14,5 | 14,5 | 14,5 | 40,371 | 0 | 44,934 | 556440000 | 20 | AJQ91240.1_1504 | RM25_1531 | Cytoplasmic | 0 | CYT | 0.072128 | OTHER | D |
| lcl\|CP010341.1_prot_AJQ91242.1_1506 | RM25_RS07565 | Dihydrodipicolinate synthase | 29013000 | 46527000 | 128810000 | 29410000 | 18955000 | 38533000 | 4 | 4 | 23,5 | 23,5 | 23,5 | 30,779 | 0 | 60,446 | 309280000 | 8 | AJQ91242.1_1506 | RM25_1533 | Cytoplasmic | 0 | CYT | 0.054845 | OTHER | E |
| lcl\|CP010341.1_prot_AJQ91243.1_1507 | RM25_RS07570 | Putative O-methyltransferase | 24676000 | 29214000 | 63120000 | 28524000 | 34576000 | 52986000 | 4 | 4 | 30,8 | 30,8 | 30,8 | 23,723 | 0 | 30,546 | 257960000 | 11 | AJQ91243.1_1507 | RM25_1534 | CytoplasmicMembrane | 0 | CYT | 0.081803 | OTHER | S |
| lcl\|CP010341.1_prot_AJQ91244.1_1508 | RM25_RS07575 | Glucose-1-phosphate adenylyltransferase | 6718400 | 1692400 | 17343000 | 0 | 2541300 | 2442800 | 2 | 2 | 6,8 | 6,8 | 6,8 | 43,095 | 0 | 16,64 | 30738000 | 3 | AJQ91244.1_1508 | RM25_1535 | CytoplasmicMembrane | 0 | CYT | 0.141306 | OTHER | G |
| lcl\|CP010341.1_prot_AJQ91246.1_1510 | RM25_RS12310 | Hypothetical protein | 91144000 | 59720000 | 147810000 | 75892000 | 106070000 | 93473000 | 1 | 1 | 18,2 | 18,2 | 18,2 | 5,7257 | 0 | 30,02 | 621310000 | 15 | AJQ91246.1_1510 | RM25_1537 | Unknown | 0 | CYT | 0.169584 | OTHER | |
| lcl\|CP010341.1_prot_AJQ91249.1_1513 | RM25_RS07595 | TIGR00730 family protein | 10019000 | 9436700 | 27516000 | 16898000 | 9906600 | 22039000 | 2 | 2 | 8,3 | 8,3 | 8,3 | 30,446 | 0 | 22,141 | 97048000 | 5 | AJQ91249.1_1513 | RM25_1540 | Cytoplasmic | 0 | CYT | 0.084357 | OTHER | S |
| lcl\|CP010341.1_prot_AJQ91250.1_1514 | RM25_RS07600 | Succinyl-diaminopimelate desuccinylase | 2368600 | 2924200 | 7434700 | 1939500 | 2092500 | 3118500 | 1 | 1 | 1,9 | 1,9 | 1,9 | 39,24 | 0,0070281 | 6,2282 | 19878000 | 2 | AJQ91250.1_1514 | RM25_1541 | Cytoplasmic | 0 | CYT | 0.097939 | OTHER | E |
| lcl\|CP010341.1_prot_AJQ91251.1_1515 | RM25_RS07605 | 2,3,4,5-tetrahydropyridine-2,6-dicarboxylate N-succinyl | 16633000 | 7959400 | 39382000 | 7594300 | 20195000 | 32138000 | 2 | 2 | 9,6 | 9,6 | 9,6 | 32,641 | 0 | 33,535 | 128060000 | 5 | AJQ91251.1_1515 | RM25_1542 | Cytoplasmic | 0 | CYT | 0.081592 | OTHER | E |
| lcl\|CP010341.1_prot_AJQ91252.1_1516 | RM25_RS07610 | Conserved protein, putative lipoprotein | 288720000 | 414729984 | 572700032 | 201480000 | 301060000 | 324140000 | 7 | 7 | 33,8 | 33,8 | 33,8 | 29,675 | 0 | 245,82 | 2266500000 | 48 | AJQ91252.1_1516 | RM25_1543 | Unknown | 1 | SPI | 0.534482 | LIPO(Sec/SPII) | M |
| lcl\|CP010341.1_prot_AJQ91254.1_1518 | RM25_RS07620 | Ferredoxin fdxC | 160380000 | 283100000 | 355550016 | 236300000 | 133210000 | 207540000 | 2 | 2 | 32,1 | 32,1 | 32,1 | 11,702 | 0 | 104,06 | 1564100000 | 11 | AJQ91254.1_1518 | RM25_1545 | Cytoplasmic | 0 | CYT | 0.411644 | OTHER | C |
| lcl\|CP010341.1_prot_AJQ91255.1_1519 | RM25_RS07625 | Glyoxylate reductase | 11800000 | 8955000 | 20273000 | 6456800 | 9259900 | 16316000 | 2 | 2 | 10,9 | 10,9 | 10,9 | 34,083 | 0 | 22,672 | 78204000 | 5 | AJQ91255.1_1519 | RM25_1546 | Cytoplasmic | 0 | CYT | 0.051171 | OTHER | CH |
| lcl\|CP010341.1_prot_AJQ91256.1_1520 | RM25_RS07630 | Fumarate hydratase class II | 284000000 | 200620000 | 982950016 | 505590016 | 348560000 | 755590016 | 13 | 13 | 34,4 | 34,4 | 34,4 | 51,182 | 0 | 228,95 | 3341700000 | 52 | AJQ91256.1_1520 | RM25_1547 | Cytoplasmic | 0 | CYT | 0.165926 | OTHER | C |
| lcl\|CP010341.1_prot_AJQ91258.1_1522 | RM25_RS07640 | Aspartate ammonia-lyase | 5198500 | 3905300 | 17201000 | 13456000 | 7541100 | 12602000 | 2 | 2 | 12,7 | 3,5 | 3,5 | 53,038 | 0 | 12,879 | 59903000 | 6 | AJQ91258.1_1522 | RM25_1549 | Cytoplasmic | 0 | CYT | 0.065463 | OTHER | E |
| lcl\|CP010341.1_prot_AJQ91259.1_1523 | RM25_RS07645 | Aspartate ammonia-lyase | 206680000 | 95780000 | 681369984 | 320510016 | 250340000 | 406920000 | 9 | 6 | 28,8 | 28,8 | 19,6 | 53,047 | 0 | 175,59 | 2164500000 | 52 | AJQ91259.1_1523 | RM25_1550 | Cytoplasmic | 0 | CYT | 0.064424 | OTHER | E |
| lcl\|CP010341.1_prot_AJQ91261.1_1525 | RM25_RS07655 | Aspartate racemase | 21086000 | 24588000 | 84525000 | 32445000 | 34443000 | 72073000 | 3 | 3 | 14,6 | 14,6 | 14,6 | 25,992 | 0 | 29,027 | 285250000 | 17 | AJQ91261.1_1525 | RM25_1552 | Cytoplasmic | 0 | CYT | 0.077248 | LIPO(Sec/SPII) | |
| lcl\|CP010341.1_prot_AJQ91265.1_1529 | RM25_RS07675 | Purine phosphoribosyltransferase | 46395000 | 82034000 | 133500000 | 52591000 | 53743000 | 62664000 | 5 | 5 | 39,3 | 39,3 | 39,3 | 18,258 | 0 | 47,989 | 480590000 | 22 | AJQ91265.1_1529 | RM25_1556 | Unknown | 0 | CYT | 0.069809 | OTHER | F |
| lcl\|CP010341.1_prot_AJQ91270.1_1534 | RM25_RS07695 | Cysteine synthase | 1876300032 | 757990016 | 4212900096 | 1792999936 | 2206700032 | 3335699968 | 17 | 17 | 81,4 | 81,4 | 81,4 | 33,625 | 0 | 323,31 | 1,5113E+10 | 144 | AJQ91270.1_1534 | RM25_1561 | Cytoplasmic | 0 | CYT | 0.101893 | OTHER | |
| lcl\|CP010341.1_prot_AJQ91275.1_1539 | RM25_RS07720 | Peptidyl-tRNA hydrolase PTH2 | 3356200 | 2301800 | 2903200 | 0 | 5733700 | 6074300 | 2 | 2 | 11,4 | 11,4 | 11,4 | 25,846 | 0 | 13,781 | 20369000 | 4 | AJQ91275.1_1539 | RM25_1566 | Cytoplasmic | 0 | CYT | 0.128637 | OTHER | S |
| lcl\|CP010341.1_prot_AJQ91278.1_1542 | RM25_RS07735 | Glycerate kinase | 0 | 2712800 | 7715900 | 4069900 | 0 | 0 | 1 | 1 | 2,7 | 2,7 | 2,7 | 36,499 | 0,008016 | 6,1896 | 14499000 | 2 | AJQ91278.1_1542 | RM25_1569 | Unknown | 0 | CYT | 0.044293 | OTHER | |
| lcl\|CP010341.1_prot_AJQ91285.1_1549 | RM25_RS07770 | Thiamine biosynthesis protein | 2932500 | 4298700 | 7429800 | 0 | 0 | 4026800 | 1 | 1 | 16,9 | 16,9 | 16,9 | 7,8737 | 0,0042463 | 7,3814 | 18688000 | 1 | AJQ91285.1_1549 | RM25_1576 | Unknown | 0 | CYT | 0.324105 | OTHER | |
| lcl\|CP010341.1_prot_AJQ91286.1_1550 | RM25_RS07775 | Glycine oxidase ThiO | 2132900 | 1638600 | 3944400 | 0 | 0 | 1906000 | 1 | 1 | 4,8 | 4,8 | 4,8 | 44,55 | 0 | 11,991 | 9621900 | 1 | AJQ91286.1_1550 | RM25_1577 | Cytoplasmic | 0 | CYT | 0.071626 | LIPO(Sec/SPII) | C |
| lcl\|CP010341.1_prot_AJQ91296.1_1560 | RM25_RS07815 | FrnE protein | 0 | 3132100 | 0 | 139440000 | 369030016 | 505480000 | 2 | 2 | 13,4 | 13,4 | 13,4 | 25,237 | 0 | 11,32 | 1020400000 | 1 | AJQ91296.1_1560 | RM25_1587 | Cytoplasmic | 0 | CYT | 0.078827 | OTHER | Q |
| lcl\|CP010341.1_prot_AJQ91299.1_1563 | RM25_RS07830 | Hypothetical protein | 82626000 | 169000000 | 217210000 | 51997000 | 131590000 | 181070000 | 6 | 6 | 36,7 | 36,7 | 36,7 | 27,068 | 0 | 78,12 | 882770000 | 24 | AJQ91299.1_1563 | RM25_1590 | Unknown | 1 | SPI | 0.477699 | SP(Sec/SPI) | |
| lcl\|CP010341.1_prot_AJQ91317.1_1581 | RM25_RS07905 | GTP-binding protein YchF | 28876000 | 35620000 | 61518000 | 17191000 | 28289000 | 41852000 | 5 | 5 | 20,1 | 20,1 | 20,1 | 39,174 | 0 | 40,61 | 213350000 | 16 | AJQ91317.1_1581 | RM25_1608 | Cytoplasmic | 0 | CYT | 0.065107 | OTHER | J |
| lcl\|CP010341.1_prot_AJQ91321.1_1585 | RM25_RS07925 | Exodeoxyribonuclease 7 large subunit | 3013100 | 0 | 6079000 | 9427200 | 7688900 | 7017500 | 1 | 1 | 7,6 | 7,6 | 7,6 | 41,073 | 0 | 18,675 | 42402000 | 3 | AJQ91321.1_1585 | RM25_1612 | Cytoplasmic | 0 | CYT | 0.051011 | OTHER | L |
| lcl\|CP010341.1_prot_AJQ91322.1_1586 | RM25_RS07930 | Exodeoxyribonuclease 7 small subunit | 12095000 | 16062000 | 33055000 | 19946000 | 17544000 | 27639000 | 2 | 2 | 28,8 | 28,8 | 28,8 | 8,6464 | 0 | 14,094 | 143990000 | 9 | AJQ91322.1_1586 | RM25_1613 | Cytoplasmic | 0 | CYT | 0.039672 | OTHER | |
| lcl\|CP010341.1_prot_AJQ91324.1_1588 | RM25_RS07940 | Isoprenyl transferase 1 | 2829400 | 0 | 2760000 | 788080 | 0 | 0 | 2 | 2 | 7,1 | 7,1 | 7,1 | 30,405 | 0 | 11,322 | 6377500 | 3 | AJQ91324.1_1588 | RM25_1615 | Cytoplasmic | 0 | CYT | 0.094434 | OTHER | I |
| lcl\|CP010341.1_prot_AJQ91325.1_1589 | RM25_RS07945 | Hypothetical protein | 3768600 | 5862200 | 12412000 | 9804200 | 5923100 | 8403800 | 1 | 1 | 5,8 | 5,8 | 5,8 | 18,898 | 0,0050607 | 6,3395 | 54596000 | 1 | AJQ91325.1_1589 | RM25_1616 | Cytoplasmic | 0 | CYT | 0.064694 | OTHER | |
| lcl\|CP010341.1_prot_AJQ91326.1_1590 | RM25_RS07950 | YyaL | 8646300 | 10815000 | 50972000 | 2786700 | 3273000 | 14998000 | 4 | 4 | 7,3 | 7,3 | 7,3 | 96,673 | 0 | 23,677 | 91491000 | 4 | AJQ91326.1_1590 | RM25_1617 | Cytoplasmic | 0 | CYT | 0.127179 | OTHER | O |
| lcl\|CP010341.1_prot_AJQ91328.1_1592 | RM25_RS07960 | Transcription elongation factor | 42256000 | 50510000 | 75846000 | 103970000 | 80782000 | 108610000 | 5 | 5 | 40,1 | 40,1 | 40,1 | 18,124 | 0 | 67,986 | 573350000 | 33 | AJQ91328.1_1592 | RM25_1619 | Cytoplasmic | 0 | CYT | 0.792401 | OTHER | K |
| lcl\|CP010341.1_prot_AJQ91329.1_1593 | RM25_RS07965 | Putative membrane protein | 17260000 | 0 | 3965100 | 0 | 4117400 | 3874800 | 2 | 2 | 15,3 | 15,3 | 15,3 | 15,134 | 0 | 11,767 | 30694000 | 1 | AJQ91329.1_1593 | RM25_1620 | Unknown | 0 | CYT | 0.033149 | OTHER | |
| lcl\|CP010341.1_prot_AJQ91330.1_1594 | RM25_RS07970 | Hypothetical protein | 12092000 | 7990700 | 32965000 | 11219000 | 9738300 | 14262000 | 5 | 5 | 18,8 | 18,8 | 18,8 | 33,529 | 0 | 28,512 | 94718000 | 3 | AJQ91330.1_1594 | RM25_1621 | Cytoplasmic | 0 | CYT | 0.098551 | OTHER | - |
| lcl\|CP010341.1_prot_AJQ91333.1_1597 | RM25_RS07985 | Peptide methionine sulfoxide reductase MsrA | 7081800 | 2005200 | 23932000 | 9988700 | 10779000 | 10812000 | 3 | 3 | 20,9 | 20,9 | 20,9 | 23,484 | 0 | 19,595 | 87154000 | 2 | AJQ91333.1_1597 | RM25_1624 | Cytoplasmic | 0 | CYT | 0.503537 | OTHER | O |
| lcl\|CP010341.1_prot_AJQ91336.1_1600 | RM25_RS08000 | Putative hydrolase | 0 | 11430000 | 15366000 | 4992700 | 6611300 | 7154500 | 1 | 1 | 7,8 | 7,8 | 7,8 | 22,949 | 0,0044199 | 8,3558 | 45554000 | 2 | AJQ91336.1_1600 | RM25_1627 | Cytoplasmic | 0 | CYT | 0.040098 | OTHER | |
| lcl\|CP010341.1_prot_AJQ91339.1_1603 | RM25_RS08015 | Hypothetical protein | 21489000 | 27710000 | 53096000 | 14236000 | 15673000 | 19016000 | 3 | 3 | 31,3 | 31,3 | 31,3 | 14,903 | 0 | 20,113 | 159990000 | 5 | AJQ91339.1_1603 | RM25_1630 | Cytoplasmic | 0 | CYT | 0.093249 | OTHER | |
| lcl\|CP010341.1_prot_AJQ91341.1_1605 | RM25_RS08020 | Putative 3-methyladenine DNA glycosylase | 6449400 | 8920000 | 22707000 | 18471000 | 12864000 | 7746400 | 4 | 4 | 24,1 | 24,1 | 24,1 | 22,875 | 0 | 28,891 | 85526000 | 5 | AJQ91341.1_1605 | RM25_1632 | Unknown | 0 | CYT | 0.104437 | OTHER | L |
| lcl\|CP010341.1_prot_AJQ91342.1_1606 | RM25_RS11945 | Hypothetical protein | 167890000 | 274569984 | 365060000 | 123620000 | 229800000 | 402929984 | 8 | 8 | 29,1 | 29,1 | 29,1 | 33,469 | 0 | 178,19 | 1692500000 | 45 | AJQ91342.1_1606 | RM25_1633 | Unknown | 0 | SPII | 0.557017 | LIPO(Sec/SPII) | |
| lcl\|CP010341.1_prot_AJQ91346.1_1610 | RM25_RS08045 | putative nucleoside-diphosphate-sugar epimerases | 36692000 | 47225000 | 109630000 | 42232000 | 38738000 | 68217000 | 5 | 5 | 24,5 | 24,5 | 24,5 | 27,757 | 0 | 44,222 | 370080000 | 25 | AJQ91346.1_1610 | RM25_1637 | Cytoplasmic | 0 | SPI | 0.073311 | OTHER | |
| lcl\|CP010341.1_prot_AJQ91356.1_1620 | RM25_RS08100 | Hypothetical protein | 48773000 | 103760000 | 156630000 | 5282000 | 61175000 | 67280000 | 2 | 2 | 15,5 | 15,5 | 15,5 | 22,898 | 0 | 38,972 | 450560000 | 8 | AJQ91356.1_1620 | RM25_1647 | Unknown | 1 | CYT | 0.935971 | OTHER | D |
| lcl\|CP010341.1_prot_AJQ91357.1_1621 | RM25_RS08105 | Phosphopyruvate hydratase | 997100032 | 552110016 | 2649100032 | 974929984 | 930940032 | 1470000000 | 15 | 15 | 52,4 | 52,4 | 52,4 | 45,95 | 0 | 323,31 | 8076400000 | 130 | AJQ91357.1_1621 | RM25_1648 | Cytoplasmic | 0 | CYT | 0.084493 | OTHER | G |
| lcl\|CP010341.1_prot_AJQ91359.1_1623 | RM25_RS08115 | Hypothetical protein | 54268000 | 79972000 | 114640000 | 55879000 | 68035000 | 143350000 | 3 | 3 | 24,9 | 24,9 | 24,9 | 18,267 | 0 | 104,59 | 551750000 | 21 | AJQ91359.1_1623 | RM25_1650 | Unknown | 0 | SPI | 0.073784 | LIPO(Sec/SPII) | |
| lcl\|CP010341.1_prot_AJQ91363.1_1627 | RM25_RS08135 | Aminoacyl-tRNA hydrolase | 1851500 | 1712900 | 3843000 | 1612100 | 3843900 | 5459700 | 1 | 1 | 6,6 | 6,6 | 6,6 | 21,208 | 0,0089463 | 6,0404 | 18323000 | 0 | AJQ91363.1_1627 | RM25_1654 | Cytoplasmic | 0 | CYT | 0.088332 | OTHER | J |
| lcl\|CP010341.1_prot_AJQ91364.1_1628 | RM25_RS08140 | 50S ribosomal protein L25 | 256600000 | 258520000 | 593449984 | 432100000 | 371249984 | 552409984 | 5 | 5 | 43,8 | 43,8 | 43,8 | 23,477 | 0 | 104 | 2961600000 | 27 | AJQ91364.1_1628 | RM25_1655 | Cytoplasmic | 0 | CYT | 0.069680 | OTHER | J |
| lcl\|CP010341.1_prot_AJQ91365.1_1629 | RM25_RS08145 | Ribose-phosphate diphosphokinase | 7513600 | 5210900 | 23282000 | 9489200 | 10139000 | 18082000 | 2 | 2 | 5,4 | 5,4 | 5,4 | 36,342 | 0 | 19,756 | 73716000 | 7 | AJQ91365.1_1629 | RM25_1656 | Cytoplasmic | 0 | CYT | 0.056729 | OTHER | F |
| lcl\|CP010341.1_prot_AJQ91366.1_1630 | RM25_RS08150 | Bifunctional protein GlmU | 59632000 | 58208000 | 146140000 | 98203000 | 89095000 | 138930000 | 7 | 7 | 15,9 | 15,9 | 15,9 | 54,519 | 0 | 45,737 | 653060000 | 27 | AJQ91366.1_1630 | RM25_1657 | Cytoplasmic | 0 | CYT | 0.103556 | OTHER | M |
| lcl\|CP010341.1_prot_AJQ91368.1_1632 | RM25_RS08160 | Transcriptional regulator, TetR family | 15762000 | 4551900 | 16031000 | 10133000 | 5434400 | 10095000 | 3 | 3 | 22,9 | 22,9 | 22,9 | 26,992 | 0 | 22,143 | 66468000 | 5 | AJQ91368.1_1632 | RM25_1659 | Unknown | 1 | CYT | 0.375391 | OTHER | |
| lcl\|CP010341.1_prot_AJQ91371.1_1635 | RM25_RS08175 | Hydrolase, TatD family | 2449500 | 2474300 | 15241000 | 2785300 | 0 | 0 | 4 | 4 | 13,8 | 13,8 | 13,8 | 34,534 | 0 | 24,162 | 24999000 | 2 | AJQ91371.1_1635 | RM25_1662 | Cytoplasmic | 0 | CYT | 0.066879 | OTHER | L |
| lcl\|CP010341.1_prot_AJQ91374.1_1638 | RM25_RS08190 | Dolichyl-phosphate-mannose-protein mannosyltransfer | 0 | 0 | 0 | 0 | 8468700 | 22782000 | 1 | 1 | 1,7 | 1,7 | 1,7 | 58,362 | 0 | 18,084 | 31251000 | 1 | AJQ91374.1_1638 | RM25_1665 | CytoplasmicMembrane | 11 | TMH | 0.951139 | OTHER | O |
| lcl\|CP010341.1_prot_AJQ91377.1_1641 | RM25_RS08205 | Hypothetical protein | 32078000 | 56537000 | 64743000 | 41256000 | 39000000 | 45859000 | 2 | 2 | 23,7 | 23,7 | 23,7 | 18,495 | 0 | 29,03 | 279470000 | 4 | AJQ91377.1_1641 | RM25_1668 | Unknown | 0 | CYT | 0.302196 | OTHER | |
| lcl\|CP010341.1_prot_AJQ91378.1_1642 | RM25_RS08210 | Galactokinase | 67187000 | 72384000 | 147200000 | 99067000 | 113940000 | 147330000 | 11 | 11 | 33,4 | 33,4 | 33,4 | 41,944 | 0 | 110,05 | 718940000 | 46 | AJQ91378.1_1642 | RM25_1669 | Cytoplasmic | 0 | CYT | 0.070045 | OTHER | G |
| lcl\|CP010341.1_prot_AJQ91384.1_1648 | RM25_RS08240 | Glutamate dehydrogenase | 6225000 | 3035200 | 29426000 | 7003300 | 18132000 | 27729000 | 3 | 3 | 6,8 | 6,8 | 6,8 | 48,914 | 0 | 18,262 | 101780000 | 4 | AJQ91384.1_1648 | RM25_1675 | Cytoplasmic | 0 | CYT | 0.095862 | OTHER | E |
| lcl\|CP010341.1_prot_AJQ91400.1_1664 | RM25_RS08325 | Heat shock protein transcriptional repressor HspR1 (Hs | 10232000 | 9433700 | 26039000 | 14132000 | 28078000 | 26502000 | 2 | 2 | 17,3 | 17,3 | 17,3 | 15,846 | 0 | 16,283 | 124710000 | 8 | AJQ91400.1_1664 | RM25_1691 | Unknown | 0 | CYT | 0.351926 | OTHER | |
| lcl\|CP010341.1_prot_AJQ91401.1_1665 | RM25_RS08330 | Chaperone protein DnaJ | 9850400 | 9103500 | 24681000 | 17178000 | 19661000 | 25948000 | 2 | 2 | 7,4 | 7,4 | 7,4 | 38,36 | 0 | 13,68 | 131000000 | 1 | AJQ91401.1_1665 | RM25_1692 | Cytoplasmic | 0 | CYT | 0.135969 | OTHER | O |
| lcl\|CP010341.1_prot_AJQ91402.1_1666 | RM25_RS08335 | Protein GrpE | 9531600 | 12001000 | 31173000 | 13085000 | 17134000 | 22576000 | 2 | 2 | 18,5 | 18,5 | 18,5 | 20,886 | 0 | 38,865 | 116350000 | 7 | AJQ91402.1_1666 | RM25_1693 | Cytoplasmic | 0 | CYT | 0.174429 | OTHER | |
| lcl\|CP010341.1_prot_AJQ91403.1_1667 | RM25_RS08340 | Chaperone protein DnaK | 164930000 | 100450000 | 532500000 | 167620000 | 209410000 | 322900000 | 13 | 13 | 29,1 | 26,4 | 26,4 | 65,435 | 0 | 100,22 | 1624700000 | 52 | AJQ91403.1_1667 | RM25_1694 | Cytoplasmic | 0 | CYT | 0.122656 | OTHER | O |
| lcl\|CP010341.1_prot_AJQ91404.1_1668 | RM25_RS08345 | Putative SERINE PROTEASE PEPD | 9625500 | 11036000 | 16393000 | 4674600 | 7261200 | 21037000 | 2 | 2 | 6,3 | 6,3 | 6,3 | 36,01 | 0 | 20,906 | 73021000 | 2 | AJQ91404.1_1668 | RM25_1695 | Unknown | 0 | SPII | 0.452606 | LIPO(Sec/SPII) | |
| lcl\|CP010341.1_prot_AJQ91409.1_1673 | RM25_RS08375 | Hypothetical protein | 10423000 | 11604000 | 22084000 | 3546800 | 6334500 | 8978000 | 3 | 3 | 17,3 | 17,3 | 17,3 | 26,729 | 0 | 33,658 | 71244000 | 8 | AJQ91409.1_1673 | RM25_1700 | Unknown | 0 | CYT | 0.035102 | OTHER | |
| lcl\|CP010341.1_prot_AJQ91410.1_1674 | RM25_RS08380 | ATP-dependent chaperone protein ClpB | 1115200000 | 854520000 | 2300100096 | 1188899968 | 1178400000 | 2248999936 | 29 | 26 | 39,5 | 39,5 | 34,6 | 99,81 | 0 | 323,31 | 9786800000 | 151 | AJQ91410.1_1674 | RM25_1701 | Cytoplasmic | 0 | CYT | 0.047893 | OTHER | O |
| lcl\|CP010341.1_prot_AJQ91417.1_1681 | RM25_RS08415 | Hypothetical protein | 28109000 | 24559000 | 58450000 | 24699000 | 27323000 | 51760000 | 7 | 7 | 16,1 | 16,1 | 16,1 | 60,344 | 0 | 42,982 | 219720000 | 7 | AJQ91417.1_1681 | RM25_1708 | Cytoplasmic | 0 | CYT | 0.045515 | OTHER | |
| lcl\|CP010341.1_prot_AJQ91436.1_1700 | RM25_RS08520 | UTP--glucose-1-phosphate uridylyltransferase | 57215000 | 84215000 | 146520000 | 58758000 | 78492000 | 92828000 | 9 | 9 | 20,8 | 20,8 | 20,8 | 49,907 | 0 | 116,24 | 573360000 | 44 | AJQ91436.1_1700 | RM25_1727 | Cytoplasmic | 0 | CYT | 0.154169 | OTHER | G |
| lcl\|CP010341.1_prot_AJQ91437.1_1701 | RM25_RS08530 | Hypothetical protein | 18408000 | 22904000 | 32752000 | 31511000 | 0 | 0 | 1 | 1 | 15,5 | 15,5 | 15,5 | 11,876 | 0 | 25,535 | 105570000 | 5 | AJQ91437.1_1701 | RM25_1729 | Unknown | 0 | CYT | 0.053782 | OTHER | |
| lcl\|CP010341.1_prot_AJQ91438.1_1702 | RM25_RS08535 | Isochorismatase hydrolase | 7041400 | 11020000 | 22156000 | 15739000 | 9537400 | 17114000 | 1 | 1 | 4,9 | 4,9 | 4,9 | 22,624 | 0,0042283 | 7,2754 | 93038000 | 7 | AJQ91438.1_1702 | RM25_1730 | Unknown | 0 | CYT | 0.077785 | OTHER | Q |
| lcl\|CP010341.1_prot_AJQ91440.1_1704 | RM25_RS08545 | Hypothetical protein | 3834900 | 0 | 140690000 | 0 | 0 | 0 | 1 | 1 | 12,6 | 12,6 | 12,6 | 13,819 | 0,0043057 | 7,5503 | 144530000 | 2 | AJQ91440.1_1704 | RM25_1732 | Cytoplasmic | 0 | CYT | 0.067880 | OTHER | |
| lcl\|CP010341.1_prot_AJQ91441.1_1705 | RM25_RS08550 | Thioredoxin | 100540000 | 122200000 | 150390000 | 214400000 | 178960000 | 254030000 | 3 | 3 | 55 | 55 | 55 | 12,225 | 0 | 208,54 | 1171600000 | 20 | AJQ91441.1_1705 | RM25_1733 | Cytoplasmic | 0 | CYT | 0.116867 | OTHER | O |
| lcl\|CP010341.1_prot_AJQ91442.1_1706 | RM25_RS08555 | Iron-dependent repressor and activator IdeR | 9272700 | 4383500 | 28973000 | 6098600 | 27212000 | 30912000 | 2 | 2 | 12 | 12 | 12 | 24,745 | 0 | 14,232 | 106850000 | 6 | AJQ91442.1_1706 | RM25_1734 | Cytoplasmic | 0 | CYT | 0.068879 | OTHER | K |
| lcl\|CP010341.1_prot_AJQ91448.1_1712 | RM25_RS12620 | Cold shock protein B cspB | 28759000 | 47731000 | 94392000 | 53240000 | 43216000 | 79457000 | 5 | 5 | 49,2 | 49,2 | 49,2 | 14,191 | 0 | 39,289 | 398870000 | 23 | AJQ91448.1_1712 | RM25_1740 | Cytoplasmic | 0 | CYT | 0.064793 | OTHER | K |
| lcl\|CP010341.1_prot_AJQ91453.1_1717 | RM25_RS11970 | Hypothetical protein | 65138000 | 93726000 | 122920000 | 60344000 | 83904000 | 116470000 | 3 | 3 | 14,2 | 14,2 | 14,2 | 32,881 | 0 | 68,266 | 558460000 | 13 | AJQ91453.1_1717 | RM25_1745 | Extracellular | 1 | SPI | 0.942354 | SP(Sec/SPI) | |
| lcl\|CP010341.1_prot_AJQ91461.1_1725 | RM25_RS08655 | 60 kDa chaperonin 1 | 3171399936 | 2739699968 | 7704600064 | 5896399872 | 6837700096 | 11256999936 | 25 | 25 | 72,2 | 72,2 | 72,2 | 56,538 | 0 | 323,31 | 4,1936E+10 | 303 | AJQ91461.1_1725 | RM25_1756 | Cytoplasmic | 0 | CYT | 0.047034 | OTHER | O |
| lcl\|CP010341.1_prot_AJQ91463.1_1727 | RM25_RS08665 | Mannose-6-phosphate isomerase, class I | 40048000 | 54366000 | 93124000 | 56110000 | 60071000 | 64701000 | 4 | 4 | 16,3 | 16,3 | 16,3 | 42,933 | 0 | 49,245 | 392810000 | 18 | AJQ91463.1_1727 | RM25_1758 | Cytoplasmic | 0 | CYT | 0.132548 | OTHER | G |
| lcl\|CP010341.1_prot_AJQ91464.1_1728 | RM25_RS08675 | Conserved transmembrane transport protein MmpL13 | 1931100 | 9971300 | 13477000 | 35251000 | 32766000 | 34159000 | 2 | 2 | 3,6 | 3,6 | 3,6 | 82,021 | 0 | 12,777 | 160540000 | 2 | AJQ91464.1_1728 | RM25_1760 | CytoplasmicMembrane | 12 | TMH | 0.879539 | OTHER | S |
| lcl\|CP010341.1_prot_AJQ91469.1_1733 | RM25_RS08700 | Amidohydrolase | 3875600 | 2041000 | 8833900 | 4772900 | 8064900 | 6506000 | 1 | 1 | 2,4 | 2,4 | 2,4 | 44,499 | 0,0042061 | 7,1598 | 35608000 | 3 | AJQ91469.1_1733 | RM25_1765 | Cytoplasmic | 0 | CYT | 0.098332 | OTHER | |
| lcl\|CP010341.1_prot_AJQ91470.1_1734 | RM25_RS08705 | L-serine dehydratase 1 | 8521000 | 0 | 11431000 | 6934400 | 8833400 | 11405000 | 2 | 2 | 8,4 | 8,4 | 8,4 | 48,652 | 0 | 13,065 | 51155000 | 2 | AJQ91470.1_1734 | RM25_1766 | Cytoplasmic | 0 | CYT | 0.062628 | OTHER | E |
| lcl\|CP010341.1_prot_AJQ91471.1_1735 | RM25_RS08710 | ATP binding protein of ABC transporter for sugars | 410240000 | 501169984 | 766220032 | 511940000 | 490350016 | 803299968 | 7 | 7 | 22,2 | 22,2 | 22,2 | 40,071 | 0 | 109,22 | 3852700000 | 36 | AJQ91471.1_1735 | RM25_1767 | CytoplasmicMembrane | 0 | CYT | 0.076953 | OTHER | P |
| lcl\|CP010341.1_prot_AJQ91472.1_1736 | RM25_RS08715 | Cystathionine gamma-synthase | 41256000 | 30850000 | 129240000 | 29988000 | 25248000 | 43539000 | 7 | 7 | 23 | 23 | 23 | 41,703 | 0 | 53,337 | 311900000 | 14 | AJQ91472.1_1736 | RM25_1768 | Cytoplasmic | 0 | CYT | 0.083786 | OTHER | E |
| lcl\|CP010341.1_prot_AJQ91473.1_1737 | RM25_RS08720 | Lipopolysaccharide kinase, Kdo/WaaP family | 1405300 | 1871900 | 5462300 | 0 | 3005900 | 3500800 | 1 | 1 | 2,8 | 2,8 | 2,8 | 52,3 | 0,0070352 | 6,241 | 21087000 | 2 | AJQ91473.1_1737 | RM25_1769 | Cytoplasmic | 0 | CYT | 0.115098 | OTHER | T |
| lcl\|CP010341.1_prot_AJQ91474.1_1738 | RM25_RS08725 | Hypothetical protein | 6443500 | 15337000 | 12939000 | 9353600 | 19241000 | 17074000 | 3 | 3 | 29,3 | 29,3 | 29,3 | 14,032 | 0 | 28,208 | 87516000 | 8 | AJQ91474.1_1738 | RM25_1770 | Cytoplasmic | 0 | CYT | 0.212423 | OTHER | |
| lcl\|CP010341.1_prot_AJQ91475.1_1739 | RM25_RS08730 | Protein of hypothetical function DUF162 | 369870016 | 269260000 | 629470016 | 702329984 | 896870016 | 1255200000 | 9 | 9 | 58,3 | 58,3 | 58,3 | 23,184 | 0 | 323,31 | 4614800000 | 58 | AJQ91475.1_1739 | RM25_1771 | Cytoplasmic | 0 | CYT | 0.096012 | OTHER | S |
| lcl\|CP010341.1_prot_AJQ91476.1_1740 | RM25_RS08735 | Iron-sulfur cluster-binding protein | 528040000 | 496350016 | 975120000 | 763449984 | 849030016 | 1859000064 | 15 | 15 | 33,4 | 33,4 | 33,4 | 57,312 | 0 | 202,66 | 6102800000 | 95 | AJQ91476.1_1740 | RM25_1772 | Cytoplasmic | 0 | CYT | 0.109371 | OTHER | C |
| lcl\|CP010341.1_prot_AJQ91477.1_1741 | RM25_RS08740 | Lactate utilization protein A | 178970000 | 201820000 | 397689984 | 159150000 | 196940000 | 448150016 | 6 | 6 | 26,4 | 26,4 | 26,4 | 28,925 | 0 | 68,38 | 1681700000 | 53 | AJQ91477.1_1741 | RM25_1773 | CytoplasmicMembrane | 0 | CYT | 0.091107 | OTHER | C |
| lcl\|CP010341.1_prot_AJQ91478.1_1742 | RM25_RS08745 | L-lactate permease | 119180000 | 98065000 | 180260000 | 118560000 | 150800000 | 188480000 | 4 | 4 | 10,2 | 10,2 | 10,2 | 59,708 | 0 | 29,867 | 947880000 | 15 | AJQ91478.1_1742 | RM25_1774 | CytoplasmicMembrane | 13 | TMH | 0.958075 | OTHER | C |
| lcl\|CP010341.1_prot_AJQ91480.1_1744 | RM25_RS08755 | Cysteine--tRNA ligase | 17909000 | 21038000 | 74004000 | 12493000 | 14016000 | 26590000 | 5 | 5 | 12 | 12 | 12 | 51,793 | 0 | 33,817 | 170290000 | 8 | AJQ91480.1_1744 | RM25_1776 | Cytoplasmic | 0 | CYT | 0.154515 | OTHER | J |
| lcl\|CP010341.1_prot_AJQ91481.1_1745 | RM25_RS08760 | RNA methyltransferase, TrmH family, group 3 | 474690 | 652150 | 7527600 | 956640 | 5157200 | 1147900 | 2 | 2 | 8,5 | 8,5 | 8,5 | 32,289 | 0 | 11,591 | 15916000 | 1 | AJQ91481.1_1745 | RM25_1777 | Cytoplasmic | 0 | CYT | 0.064752 | OTHER | J |
| lcl\|CP010341.1_prot_AJQ91485.1_1749 | RM25_RS08780 | Glycine hydroxymethyltransferase | 72432000 | 43186000 | 256810000 | 26188000 | 60332000 | 101650000 | 11 | 11 | 29,3 | 29,3 | 29,3 | 51,95 | 0 | 116,67 | 576300000 | 28 | AJQ91485.1_1749 | RM25_1781 | Cytoplasmic | 0 | CYT | 0.108544 | OTHER | E |
| lcl\|CP010341.1_prot_AJQ91488.1_1752 | RM25_RS08795 | Transcriptional regulator, CarD family | 23048000 | 30817000 | 64466000 | 40192000 | 30875000 | 63193000 | 2 | 2 | 12,4 | 12,4 | 12,4 | 18,2 | 0 | 13,14 | 285730000 | 6 | AJQ91488.1_1752 | RM25_1784 | Cytoplasmic | 0 | CYT | 0.034394 | OTHER | K |
| lcl\|CP010341.1_prot_AJQ91492.1_1756 | RM25_RS08815 | Phosphate-transport system transcriptional regulator p | 4836800 | 8970700 | 19372000 | 7458400 | 4772000 | 16645000 | 3 | 3 | 15,1 | 15,1 | 15,1 | 25,166 | 0 | 21,43 | 62055000 | 5 | AJQ91492.1_1756 | RM25_1788 | Cytoplasmic | 0 | CYT | 0.043297 | OTHER | P |
| lcl\|CP010341.1_prot_AJQ91493.1_1757 | RM25_RS08820 | 2,3-bisphosphoglycerate-dependent phosphoglycerate | 201270000 | 180010000 | 506950016 | 268729984 | 350880000 | 543740032 | 12 | 12 | 70,3 | 70,3 | 70,3 | 27,984 | 0 | 182,67 | 2175200000 | 43 | AJQ91493.1_1757 | RM25_1789 | Cytoplasmic | 0 | CYT | 0.306372 | OTHER | G |
| lcl\|CP010341.1_prot_AJQ91495.1_1759 | RM25_RS08830 | Short-chain dehydrogenase/reductase yusR | 10838000 | 2409800 | 40810000 | 18685000 | 11004000 | 18534000 | 2 | 2 | 7,5 | 7,5 | 7,5 | 25,454 | 0 | 15,58 | 109250000 | 2 | AJQ91495.1_1759 | RM25_1791 | Cytoplasmic | 0 | CYT | 0.054630 | OTHER | S |
| lcl\|CP010341.1_prot_AJQ91496.1_1760 | RM25_RS08835 | Biotin-requiring enzyme | 903560000 | 1208600064 | 1646700032 | 1432800000 | 865350016 | 2006400000 | 5 | 5 | 54,5 | 54,5 | 54,5 | 12,367 | 0 | 226,95 | 9396000000 | 50 | AJQ91496.1_1760 | RM25_1792 | Unknown | 0 | CYT | 0.265380 | OTHER | |
| lcl\|CP010341.1_prot_AJQ91497.1_1761 | RM25_RS08840 | Hypothetical protein | 25034000 | 27831000 | 61369000 | 84112000 | 58506000 | 84878000 | 2 | 2 | 18,3 | 18,3 | 18,3 | 8,8387 | 0 | 20,649 | 424430000 | 13 | AJQ91497.1_1761 | RM25_1793 | CytoplasmicMembrane | 0 | CYT | 0.036916 | OTHER | |
| lcl\|CP010341.1_prot_AJQ91498.1_1762 | RM25_RS08845 | Carboxyl transferase | 573329984 | 380969984 | 1414400000 | 892259968 | 865160000 | 1739399936 | 14 | 14 | 34,7 | 34,7 | 34,7 | 56,378 | 0 | 234,06 | 6552200000 | 119 | AJQ91498.1_1762 | RM25_1794 | Cytoplasmic | 0 | CYT | 0.081675 | OTHER | |

| | | | | | | | | | | | | | | | | | | | | | | | | | | |
|---|---|---|---|---|---|---|---|---|---|---|---|---|---|---|---|---|---|---|---|---|---|---|---|---|---|---|
| lcl\|CP010341.1_prot_AJQ91499.1_1763 | RM25_RS08850 | Oxaloacetate decarboxylase alpha subunit | 1246000000 | 527870016 | 3104100096 | 1476899968 | 1320300032 | 1868700032 | 20 | 20 | 47,3 | 47,3 | 47,3 | 55,649 | 0 | 246,25 | 1,0364E+10 | 187 | AJQ91499.1_1763 | RM25_1795 | Cytoplasmic | 0 | CYT | 0.089447 | OTHER | C |
| lcl\|CP010341.1_prot_AJQ91500.1_1764 | RM25_RS08855 | Glutamate-1-semialdehyde 2,1-aminomutase 2 | 112810000 | 133260000 | 357089984 | 132470000 | 129590000 | 202420000 | 9 | 9 | 27,7 | 27,7 | 27,7 | 45,932 | 0 | 86,665 | 1135300000 | 39 | AJQ91500.1_1764 | RM25_1796 | Cytoplasmic | 0 | CYT | 0.043163 | OTHER | H |
| lcl\|CP010341.1_prot_AJQ91504.1_1768 | RM25_RS08875 | Ferrochelatase | 12203000 | 20481000 | 35226000 | 5071700 | 16596000 | 13849000 | 5 | 5 | 18,5 | 18,5 | 18,5 | 37,455 | 0 | 33,577 | 105740000 | 12 | AJQ91504.1_1768 | RM25_1800 | Cytoplasmic | 0 | CYT | 0.082869 | OTHER | H |
| lcl\|CP010341.1_prot_AJQ91505.1_1769 | RM25_RS08880 | Protoporphyrinogen oxidase | 13494000 | 17873000 | 24010000 | 5690000 | 14728000 | 28394000 | 5 | 5 | 14,4 | 14,4 | 14,4 | 55,155 | 0 | 34,136 | 113990000 | 9 | AJQ91505.1_1769 | RM25_1801 | Cytoplasmic | 0 | CYT | 0.088631 | OTHER | |
| lcl\|CP010341.1_prot_AJQ91506.1_1770 | RM25_RS08885 | Coenzyme PQQ synthesis protein | 35115000 | 51614000 | 61279000 | 37154000 | 24469000 | 53734000 | 7 | 7 | 22 | 22 | 22 | 40,664 | 0 | 54,328 | 276910000 | 14 | AJQ91506.1_1770 | RM25_1802 | Cytoplasmic | 0 | CYT | 0.100483 | OTHER | |
| lcl\|CP010341.1_prot_AJQ91507.1_1771 | RM25_RS12355 | Uroporphyrinogen decarboxylase | 22824000 | 7926600 | 79696000 | 13114000 | 21294000 | 39366000 | 4 | 4 | 9,8 | 9,8 | 9,8 | 66,07 | 0 | 39,694 | 192930000 | 9 | AJQ91507.1_1771 | RM25_1803 | Cytoplasmic | 0 | CYT | 0.055389 | OTHER | H |
| lcl\|CP010341.1_prot_AJQ91508.1_1772 | RM25_RS08895 | Hydroxymethylbilane synthase | 15637000 | 17827000 | 36042000 | 15742000 | 3619300 | 6784200 | 3 | 3 | 13,9 | 13,9 | 13,9 | 32,825 | 0 | 24,338 | 97192000 | 5 | AJQ91508.1_1772 | RM25_1804 | Cytoplasmic | 0 | CYT | 0.045502 | OTHER | H |
| lcl\|CP010341.1_prot_AJQ91509.1_1773 | RM25_RS08900 | Glutamyl-tRNA reductase | 21327000 | 21195000 | 62269000 | 10287000 | 35077000 | 64050000 | 6 | 6 | 16,5 | 16,5 | 16,5 | 44,023 | 0 | 36,096 | 222870000 | 9 | AJQ91509.1_1773 | RM25_1805 | Cytoplasmic | 0 | CYT | 0.033857 | OTHER | H |
| lcl\|CP010341.1_prot_AJQ91510.1_1774 | RM25_RS08905 | Aspartate--ammonia ligase | 14142000 | 10969000 | 57884000 | 6878400 | 25268000 | 56914000 | 6 | 6 | 10,4 | 10,4 | 10,4 | 72,574 | 0 | 36,351 | 172050000 | 9 | AJQ91510.1_1774 | RM25_1806 | Unknown | 0 | CYT | 0.131987 | OTHER | E |
| lcl\|CP010341.1_prot_AJQ91511.1_1775 | RM25_RS08910 | Dihydroxy-acid dehydratase | 8394400 | 18294000 | 42063000 | 14684000 | 20437000 | 20793000 | 2 | 2 | 4,2 | 4,2 | 4,2 | 65,034 | 0 | 14,074 | 131010000 | 6 | AJQ91511.1_1775 | RM25_1807 | CytoplasmicMembrane | 0 | CYT | 0.084665 | OTHER | EG |
| lcl\|CP010341.1_prot_AJQ91513.1_1777 | RM25_RS08920 | Inosose dehydratase | 96862000 | 36546000 | 113750000 | 72882000 | 135340000 | 256890000 | 5 | 5 | 22,5 | 22,5 | 22,5 | 34,517 | 0 | 89,272 | 777730000 | 20 | AJQ91513.1_1777 | RM25_1809 | Cytoplasmic | 0 | CYT | 0.091070 | OTHER | |
| lcl\|CP010341.1_prot_AJQ91515.1_1779 | RM25_RS08930 | Inositol 2-dehydrogenase | 30661000 | 2734900 | 52413000 | 10035000 | 13528000 | 47058000 | 4 | 4 | 13,1 | 13,1 | 13,1 | 34,756 | 0 | 32,567 | 160630000 | 8 | AJQ91515.1_1779 | RM25_1811 | Cytoplasmic | 0 | CYT | 0.050394 | OTHER | S |
| lcl\|CP010341.1_prot_AJQ91516.1_1780 | RM25_RS08935 | Transaldolase | 59078000 | 15361000 | 182640000 | 58926000 | 79765000 | 145150000 | 7 | 7 | 25,7 | 25,7 | 25,7 | 39,396 | 0 | 68,203 | 575450000 | 27 | AJQ91516.1_1780 | RM25_1812 | Cytoplasmic | 0 | CYT | 0.083965 | OTHER | H |
| lcl\|CP010341.1_prot_AJQ91517.1_1781 | RM25_RS08940 | UbiC transcription regulator-associated | 9731200 | 10060000 | 48557000 | 8638500 | 12767000 | 32311000 | 4 | 4 | 24,8 | 24,8 | 24,8 | 27,4 | 0 | 30,333 | 129150000 | 7 | AJQ91517.1_1781 | RM25_1813 | CytoplasmicMembrane | 0 | CYT | 0.078578 | OTHER | K |
| lcl\|CP010341.1_prot_AJQ91518.1_1782 | RM25_RS08945 | Putative 5-dehydro-2-deoxygluconokinase | 15921000 | 7334300 | 24213000 | 2998200 | 10617000 | 47085000 | 3 | 3 | 11,2 | 11,2 | 11,2 | 35,635 | 0 | 19,955 | 108170000 | 7 | AJQ91518.1_1782 | RM25_1814 | Cytoplasmic | 0 | CYT | 0.071665 | OTHER | G |
| lcl\|CP010341.1_prot_AJQ91520.1_1784 | RM25_RS08955 | Myo-inositol catabolism protein | 11671000 | 0 | 18846000 | 4095800 | 6840400 | 17275000 | 4 | 4 | 13,8 | 13,8 | 13,8 | 31,858 | 0 | 24,498 | 60965000 | 7 | AJQ91520.1_1784 | RM25_1816 | Cytoplasmic | 0 | CYT | 0.559072 | OTHER | G |
| lcl\|CP010341.1_prot_AJQ91521.1_1785 | RM25_RS08960 | 3D-(3,5/4)-trihydroxycyclohexane-1,2-dione hydrolase | 42876000 | 20607000 | 117120000 | 44931000 | 116770000 | 187100000 | 12 | 12 | 21,4 | 21,4 | 21,4 | 69,85 | 0 | 81,797 | 569450000 | 25 | AJQ91521.1_1785 | RM25_1817 | Cytoplasmic | 0 | CYT | 0.127557 | OTHER | E |
| lcl\|CP010341.1_prot_AJQ91522.1_1786 | RM25_RS08965 | Methylmalonate-semialdehyde dehydrogenase (acylati | 195690000 | 107780000 | 479590016 | 275710016 | 334720000 | 761510016 | 11 | 11 | 33,7 | 33,7 | 33,7 | 52,826 | 0 | 287,44 | 2379600000 | 56 | AJQ91522.1_1786 | RM25_1818 | Cytoplasmic | 0 | CYT | 0.076124 | OTHER | C |
| lcl\|CP010341.1_prot_AJQ91527.1_1791 | RM25_RS08985 | Hypothetical protein | 6144800 | 3694500 | 9225200 | 4425500 | 10578000 | 17480000 | 1 | 1 | 15,4 | 15,4 | 15,4 | 9,0004 | 0 | 12,458 | 56896000 | 7 | AJQ91527.1_1791 | RM25_1823 | CytoplasmicMembrane | 2 | SPI | 0.955043 | OTHER | |
| lcl\|CP010341.1_prot_AJQ91528.1_1792 | RM25_RS08990 | Fructose-bisphosphate aldolase, class II | 54282000 | 10902000 | 347790016 | 134380000 | 166870000 | 229760000 | 10 | 10 | 30,9 | 30,9 | 30,9 | 36,874 | 0 | 79,578 | 1009800000 | 28 | AJQ91528.1_1792 | RM25_1824 | Cytoplasmic | 0 | CYT | 0.148189 | OTHER | G |
| lcl\|CP010341.1_prot_AJQ91533.1_1797 | RM25_RS09015 | Orotate phosphoribosyltransferase | 51886000 | 66406000 | 119580000 | 66338000 | 58403000 | 78408000 | 6 | 6 | 35,7 | 35,7 | 35,7 | 19,322 | 0 | 42,941 | 482670000 | 17 | AJQ91533.1_1797 | RM25_1829 | Unknown | 0 | CYT | 0.035408 | OTHER | |
| lcl\|CP010341.1_prot_AJQ91534.1_1798 | RM25_RS09020 | Peptidase C60 family protein | 246600000 | 423760000 | 510089984 | 73530000 | 220720000 | 283400000 | 4 | 4 | 33,7 | 33,7 | 33,7 | 21,037 | 0 | 77,042 | 1793800000 | 11 | AJQ91534.1_1798 | RM25_1830 | Unknown | 1 | SPI | 0.263717 | SP(Sec/SPI) | |
| lcl\|CP010341.1_prot_AJQ91537.1_1801 | RM25_RS09035 | ATP-dependent chaperone protein ClpB | 184900000 | 167620000 | 505400000 | 432489984 | 462630016 | 610289984 | 27 | 27 | 41,7 | 38 | 38 | 94,419 | 0 | 323,31 | 2667000000 | 105 | AJQ91537.1_1801 | RM25_1833 | Cytoplasmic | 0 | CYT | 0.046730 | OTHER | O |
| lcl\|CP010341.1_prot_AJQ91538.1_1802 | RM25_RS09040 | Glutamate-1-semialdehyde 2,1-aminomutase 2 | 35750000 | 82491000 | 196710000 | 34807000 | 43028000 | 58467000 | 8 | 8 | 22,6 | 22,6 | 22,6 | 49,456 | 0 | 105,58 | 463200000 | 16 | AJQ91538.1_1802 | RM25_1834 | Cytoplasmic | 0 | CYT | 0.089179 | OTHER | H |
| lcl\|CP010341.1_prot_AJQ91540.1_1804 | RM25_RS09050 | Adenylosuccinate synthase | 946000 | 1804800 | 0 | 1276100 | 0 | 0 | 2 | 2 | 4,4 | 4,4 | 4,4 | 46,564 | 0,0023202 | 10,649 | 4026900 | 1 | AJQ91540.1_1804 | RM25_1836 | Cytoplasmic | 0 | CYT | 0.099666 | OTHER | F |
| lcl\|CP010341.1_prot_AJQ91542.1_1806 | RM25_RS09060 | Putative aldehyde dehydrogenase AldA | 22152000 | 2750600 | 249780000 | 102930000 | 22929000 | 22355000 | 10 | 10 | 28,7 | 28,7 | 28,7 | 53,436 | 0 | 144,94 | 432690000 | 15 | AJQ91542.1_1806 | RM25_1838 | Cytoplasmic | 0 | CYT | 0.084271 | OTHER | C |
| lcl\|CP010341.1_prot_AJQ91553.1_1817 | RM25_RS09120 | Glycosyl transferase family 51 | 189930000 | 254540000 | 463180000 | 259710000 | 338640000 | 536569984 | 13 | 13 | 24,2 | 24,2 | 24,2 | 77,967 | 0 | 274,24 | 2233600000 | 62 | AJQ91553.1_1817 | RM25_1849 | CytoplasmicMembrane | 1 | TMH | 0.903416 | SP(Sec/SPI) | M |
| lcl\|CP010341.1_prot_AJQ91556.1_1820 | RM25_RS09135 | Endoribonuclease L-PSP | 33529000 | 38527000 | 66611000 | 16876000 | 23148000 | 31629000 | 3 | 3 | 31,8 | 31,8 | 31,8 | 15,249 | 0 | 73,669 | 212230000 | 18 | AJQ91556.1_1820 | RM25_1852 | Cytoplasmic | 0 | CYT | 0.024034 | OTHER | J |
| lcl\|CP010341.1_prot_AJQ91560.1_1824 | RM25_RS09155 | Phosphoenolpyruvate-protein phosphotransferase | 6182800 | 0 | 8039000 | 4300200 | 2285900 | 4821900 | 2 | 2 | 3,6 | 3,6 | 3,6 | 60,131 | 0 | 12,569 | 25630000 | 2 | AJQ91560.1_1824 | RM25_1856 | Cytoplasmic | 0 | SPI | 0.045443 | SP(Sec/SPI) | G |
| lcl\|CP010341.1_prot_AJQ91561.1_1825 | RM25_RS09160 | Phosphocarrier, HPr family | 308929984 | 411260000 | 437500000 | 327800000 | 242320000 | 413089984 | 3 | 3 | 46,6 | 46,6 | 46,6 | 9,027 | 0 | 109,21 | 2399400000 | 15 | AJQ91561.1_1825 | RM25_1857 | Cytoplasmic | 0 | CYT | 0.063825 | OTHER | |
| lcl\|CP010341.1_prot_AJQ91567.1_1831 | RM25_1863 | Hypothetical protein | 0 | 0 | 0 | 9291800 | 0 | 11115000 | 1 | 1 | 19 | 19 | 19 | 11,931 | 0 | 20,9 | 27539000 | 1 | AJQ91567.1_1831 | RM25_1863 | Unknown | 0 | CYT | 0.143159 | OTHER | |
| lcl\|CP010341.1_prot_AJQ91569.1_1833 | RM25_RS09195 | Thioredoxin | 52559000 | 85540000 | 85296000 | 84287000 | 86999000 | 95258000 | 3 | 3 | 33,8 | 33,8 | 33,8 | 15,499 | 0 | 66,044 | 604120000 | 15 | AJQ91569.1_1833 | RM25_1865 | Cytoplasmic | 0 | CYT | 0.147748 | OTHER | CO |
| lcl\|CP010341.1_prot_AJQ91570.1_1834 | RM25_RS09200 | FHA domain protein | 22149000 | 33670000 | 57825000 | 36501000 | 32592000 | 42120000 | 5 | 5 | 26,5 | 26,5 | 26,5 | 26,018 | 0 | 49,206 | 254270000 | 17 | AJQ91570.1_1834 | RM25_1866 | Cytoplasmic | 0 | CYT | 0.097523 | OTHER | T |
| lcl\|CP010341.1_prot_AJQ91571.1_1835 | RM25_RS09205 | FHA domain protein | 3868400 | 4601200 | 8870800 | 0 | 5281800 | 8146900 | 1 | 1 | 6,4 | 6,4 | 6,4 | 17,303 | 0,0042194 | 7,2543 | 30769000 | 4 | AJQ91571.1_1835 | RM25_1867 | Unknown | 1 | CYT | 0.802670 | OTHER | |
| lcl\|CP010341.1_prot_AJQ91572.1_1836 | RM25_1868 | Putative SERINE/THREONINE PHOSPHATASE PPP | 12748000 | 24045000 | 40187000 | 2010300 | 20640000 | 28342000 | 2 | 2 | 4,5 | 4,5 | 4,5 | 52,955 | 0 | 14,028 | 129580000 | 4 | AJQ91572.1_1836 | RM25_1868 | Unknown | 1 | CYT | 0.627097 | OTHER | T |
| lcl\|CP010341.1_prot_AJQ91574.1_1838 | RM25_RS09220 | Penicillin-binding protein pbpA | 143770000 | 279860000 | 349780000 | 127200000 | 230160000 | 265800000 | 9 | 9 | 26,2 | 26,2 | 26,2 | 49,704 | 0 | 190,45 | 1478900000 | 39 | AJQ91574.1_1838 | RM25_1870 | CytoplasmicMembrane | 1 | SPI | 0.924512 | SP(Sec/SPI) | M |
| lcl\|CP010341.1_prot_AJQ91575.1_1839 | RM25_RS09225 | Non-specific serine/threonine protein kinase | 5864000 | 4573600 | 26068000 | 14282000 | 14826000 | 21839000 | 2 | 2 | 4,5 | 4,5 | 4,5 | 62,562 | 0 | 12,159 | 87454000 | 6 | AJQ91575.1_1839 | RM25_1871 | CytoplasmicMembrane | 1 | CYT | 0.758223 | OTHER | KLT |
| lcl\|CP010341.1_prot_AJQ91576.1_1840 | RM25_RS11995 | Hypothetical protein | 18076000 | 8434000 | 31507000 | 26759000 | 29682000 | 53212000 | 4 | 4 | 9,2 | 9,2 | 9,2 | 60,011 | 0 | 96,805 | 174890000 | 11 | AJQ91576.1_1840 | RM25_1872 | CytoplasmicMembrane | 1 | CYT | 0.956603 | OTHER | KLT |
| lcl\|CP010341.1_prot_AJQ91577.1_1841 | RM25_RS09235 | Galactokinase | 30208000 | 65937000 | 127690000 | 91373000 | 49798000 | 46887000 | 8 | 8 | 25,4 | 25,4 | 25,4 | 40,505 | 0 | 59,233 | 474610000 | 24 | AJQ91577.1_1841 | RM25_1873 | CytoplasmicMembrane | 0 | CYT | 0.067043 | OTHER | G |
| lcl\|CP010341.1_prot_AJQ91579.1_1843 | RM25_RS09245 | Ferrous iron transport protein B | 1886400 | 1614000 | 8057300 | 11720000 | 12020000 | 23408000 | 3 | 3 | 6 | 6 | 6 | 69,798 | 0 | 28,798 | 68873000 | 5 | AJQ91579.1_1843 | RM25_1875 | CytoplasmicMembrane | 9 | CYT | 0.917375 | OTHER | |
| lcl\|CP010341.1_prot_AJQ91580.1_1844 | RM25_RS09250 | Ferrous iron transport protein A | 1081600 | 1772700 | 10853000 | 17414000 | 24370000 | 24818000 | 2 | 2 | 24,4 | 24,4 | 24,4 | 12,85 | 0 | 13,107 | 92272000 | 4 | AJQ91580.1_1844 | RM25_1876 | Cytoplasmic | 0 | CYT | 0.039254 | OTHER | P |
| lcl\|CP010341.1_prot_AJQ91583.1_1847 | RM25_RS12000 | Hypothetical protein | 14862000 | 13890000 | 44083000 | 13550000 | 21616000 | 28044000 | 4 | 4 | 8,7 | 8,7 | 8,7 | 65,032 | 0 | 57,884 | 142430000 | 9 | AJQ91583.1_1847 | RM25_1879 | Cytoplasmic | 2 | CYT | 0.965492 | OTHER | |
| lcl\|CP010341.1_prot_AJQ91585.1_1849 | RM25_RS09280 | Putative tRNA adenosine deaminase-associated protein | 12268000 | 32059000 | 42419000 | 13499000 | 15485000 | 17655000 | 1 | 1 | 7,9 | 7,9 | 7,9 | 26,348 | 0 | 28,745 | 146630000 | 2 | AJQ91585.1_1849 | RM25_1881 | Cytoplasmic | 0 | CYT | 0.837763 | OTHER | S |
| lcl\|CP010341.1_prot_AJQ91586.1_1850 | RM25_RS09285 | Uracil phosphoribosyltransferase | 13822000 | 23369000 | 31275000 | 7200900 | 11951000 | 0 | 2 | 2 | 11,7 | 11,7 | 11,7 | 22,817 | 0 | 29,425 | 103380000 | 2 | AJQ91586.1_1850 | RM25_1882 | Cytoplasmic | 0 | CYT | 0.045438 | OTHER | F |
| lcl\|CP010341.1_prot_AJQ91590.1_1854 | RM25_RS09305 | Putative surface-anchored fimbrial subunit | 144550000 | 187990000 | 170330000 | 73149000 | 78809000 | 79788000 | 6 | 6 | 26,6 | 26,6 | 26,6 | 39,502 | 0 | 113,63 | 779750000 | 15 | AJQ91590.1_1854 | RM25_1886 | Unknown | 1 | SPI | 0.922509 | SP(Sec/SPI) | |
| lcl\|CP010341.1_prot_AJQ91591.1_1855 | RM25_RS09310 | Surface-anchored fimbrial subunit | 15446000 | 19665000 | 29940000 | 0 | 3672900 | 15204000 | 3 | 3 | 4,3 | 4,3 | 4,3 | 98,147 | 0 | 22,941 | 83928000 | 5 | AJQ91591.1_1855 | RM25_1887 | Unknown | 1 | CYT | 0.926733 | SP(Sec/SPI) | |
| lcl\|CP010341.1_prot_AJQ91593.1_1857 | RM25_RS09320 | GTP-binding translation elongation factor typA | 44516000 | 50157000 | 137030000 | 74769000 | 84691000 | 150690000 | 8 | 8 | 13,2 | 13,2 | 13,2 | 68,864 | 0 | 56,244 | 583260000 | 21 | AJQ91593.1_1857 | RM25_1889 | CytoplasmicMembrane | 0 | CYT | 0.079980 | OTHER | T |
| lcl\|CP010341.1_prot_AJQ91595.1_1859 | RM25_RS09330 | UPF0678 fatty acid-binding protein-like protein | 19061000 | 29531000 | 45233000 | 36149000 | 32297000 | 25499000 | 2 | 2 | 12,6 | 12,6 | 12,6 | 18,883 | 0 | 17,735 | 214110000 | 10 | AJQ91595.1_1859 | RM25_1891 | Unknown | 0 | CYT | 0.790482 | OTHER | S |
| lcl\|CP010341.1_prot_AJQ91598.1_1862 | RM25_RS09345 | Hypothetical protein | 89188000 | 166960000 | 180090000 | 39679000 | 70826000 | 183090000 | 3 | 3 | 27,2 | 27,2 | 27,2 | 18,361 | 0 | 33,389 | 763310000 | 10 | AJQ91598.1_1862 | RM25_1894 | Unknown | 0 | SPII | 0.550873 | LIPO(Sec/SPII) | |
| lcl\|CP010341.1_prot_AJQ91599.1_1863 | RM25_RS09350 | Inositol-3-phosphate synthase | 160080000 | 116070000 | 558240000 | 271569984 | 158190000 | 222440000 | 10 | 10 | 39 | 39 | 39 | 39,23 | 0 | 180 | 1667700000 | 41 | AJQ91599.1_1863 | RM25_1895 | Cytoplasmic | 0 | CYT | 0.054503 | OTHER | I |
| lcl\|CP010341.1_prot_AJQ91604.1_1868 | RM25_RS09370 | Mycothiol acetyltransferase | 19238000 | 32381000 | 72554000 | 18147000 | 5162600 | 24638000 | 4 | 4 | 15,7 | 15,7 | 15,7 | 34,301 | 0 | 26,801 | 210920000 | 6 | AJQ91604.1_1868 | RM25_1900 | Cytoplasmic | 0 | CYT | 0.142919 | OTHER | K |
| lcl\|CP010341.1_prot_AJQ91605.1_1869 | RM25_RS09375 | Polyphosphate kinase | 9925000 | 0 | 18228000 | 0 | 0 | 0 | 3 | 3 | 5,9 | 5,9 | 5,9 | 82,373 | 0 | 17,61 | 28154000 | 2 | AJQ91605.1_1869 | RM25_1901 | CytoplasmicMembrane | 0 | CYT | 0.062033 | OTHER | P |
| lcl\|CP010341.1_prot_AJQ91615.1_1879 | RM25_RS09425 | putative B-hexosaminidase | 436150016 | 888649984 | 1107500032 | 252050000 | 324609984 | 814800000 | 7 | 7 | 27,9 | 27,9 | 27,9 | 41,267 | 0 | 126,05 | 4024400000 | 42 | AJQ91615.1_1879 | RM25_1911 | Unknown | 1 | SPII | 0.305952 | LIPO(Sec/SPII) | G |
| lcl\|CP010341.1_prot_AJQ91616.1_1880 | RM25_RS09430 | Mycothione reductase | 18237000 | 11577000 | 33750000 | 13460000 | 6550000 | 14380000 | 3 | 3 | 9,9 | 9,9 | 9,9 | 53,733 | 0 | 25,544 | 101310000 | 8 | AJQ91616.1_1880 | RM25_1912 | Cytoplasmic | 0 | CYT | 0.083596 | OTHER | C |
| lcl\|CP010341.1_prot_AJQ91617.1_1881 | RM25_RS09435 | Phosphoglycerate mutase | 5753800 | 3286500 | 12644000 | 13268000 | 15152000 | 26879000 | 2 | 2 | 14,6 | 14,6 | 14,6 | 24,288 | 0 | 12,528 | 76984000 | 5 | AJQ91617.1_1881 | RM25_1913 | CytoplasmicMembrane | 0 | CYT | 0.164491 | OTHER | G |
| lcl\|CP010341.1_prot_AJQ91618.1_1882 | RM25_RS09440 | Uroporphyrin-III C-methyltransferase/uroporphyrinoge | 17097000 | 13603000 | 37680000 | 24597000 | 17477000 | 47272000 | 4 | 4 | 9,3 | 9,3 | 9,3 | 57,187 | 0 | 29,955 | 180840000 | 17 | AJQ91618.1_1882 | RM25_1914 | Cytoplasmic | 0 | CYT | 0.058666 | OTHER | H |
| lcl\|CP010341.1_prot_AJQ91619.1_1883 | RM25_RS09445 | Pyrroline-5-carboxylate reductase | 7001900 | 0 | 22612000 | 11106000 | 14791000 | 25549000 | 2 | 2 | 10,1 | 10,1 | 10,1 | 28,964 | 0 | 12,989 | 92100000 | 2 | AJQ91619.1_1883 | RM25_1915 | Cytoplasmic | 1 | CYT | 0.105414 | TAT(Tat/SPI) | E |
| lcl\|CP010341.1_prot_AJQ91620.1_1884 | RM25_RS09450 | Aspartate-semialdehyde dehydrogenase | 44381000 | 38920000 | 119290000 | 26792000 | 32053000 | 50335000 | 5 | 5 | 19,8 | 19,8 | 19,8 | 36,567 | 0 | 38,489 | 329270000 | 13 | AJQ91620.1_1884 | RM25_1916 | Cytoplasmic | 0 | CYT | 0.085147 | OTHER | E |
| lcl\|CP010341.1_prot_AJQ91624.1_1888 | RM25_RS09470 | Hypothetical protein | 54987000 | 75004000 | 163260000 | 103730000 | 66734000 | 126050000 | 5 | 5 | 25,5 | 25,5 | 25,5 | 23,209 | 0 | 50,895 | 666520000 | 18 | AJQ91624.1_1888 | RM25_1920 | Cytoplasmic | 0 | CYT | 0.113277 | OTHER | S |
| lcl\|CP010341.1_prot_AJQ91625.1_1889 | RM25_RS09475 | DNA repair protein radA | 1312600 | 1119900 | 4669800 | 2905500 | 4943300 | 4031500 | 1 | 1 | 2,4 | 2,4 | 2,4 | 49,134 | 0,0080241 | 6,2188 | 20591000 | 2 | AJQ91625.1_1889 | RM25_1921 | Cytoplasmic | 0 | CYT | 0.051596 | OTHER | O |
| lcl\|CP010341.1_prot_AJQ91632.1_1896 | RM25_RS09510 | 3-dehydroquinate dehydratase | 13312000 | 13161000 | 24022000 | 19931000 | 22582000 | 25364000 | 2 | 2 | 12,4 | 12,4 | 12,4 | 25,953 | 0 | 15,002 | 124590000 | 4 | AJQ91632.1_1896 | RM25_1928 | Cytoplasmic | 0 | CYT | 0.061238 | SP(Sec/SPI) | |
| lcl\|CP010341.1_prot_AJQ91634.1_1898 | RM25_RS09520 | ATP-dependent protease ATP-binding subunit clpC1 | 420060000 | 480860000 | 1175500032 | 882160000 | 970249984 | 1223000064 | 26 | 26 | 43,9 | 42,4 | 42,4 | 94,357 | 0 | 323,31 | 5841700000 | 140 | AJQ91634.1_1898 | RM25_1930 | Cytoplasmic | 0 | CYT | 0.068499 | OTHER | O |
| lcl\|CP010341.1_prot_AJQ91635.1_1899 | RM25_RS09525 | Hypothetical protein | 23214000 | 19489000 | 67240000 | 19981000 | 20356000 | 22681000 | 2 | 2 | 16,1 | 16,1 | 16,1 | 12,989 | 0 | 16,328 | 191870000 | 8 | AJQ91635.1_1899 | RM25_1931 | Unknown | 0 | CYT | 0.078370 | OTHER | |
| lcl\|CP010341.1_prot_AJQ91636.1_1900 | RM25_RS09530 | 2-amino-4-hydroxy-6-hydroxymethyldihydropteridine d | 17850000 | 22736000 | 64232000 | 31116000 | 11386000 | 44476000 | 5 | 5 | 30,9 | 30,9 | 30,9 | 19,83 | 0 | 29,428 | 191800000 | 6 | AJQ91636.1_1900 | RM25_1932 | Cytoplasmic | 0 | CYT | 0.135337 | OTHER | H |
| lcl\|CP010341.1_prot_AJQ91637.1_1901 | RM25_RS09535 | Dihydroneopterin aldolase | 3772400 | 0 | 10629000 | 2766200 | 1720800 | 0 | 1 | 1 | 8,1 | 8,1 | 8,1 | 13,459 | 0,0043668 | 7,906 | 18889000 | 1 | AJQ91637.1_1901 | RM25_1933 | Cytoplasmic | 0 | CYT | 0.038156 | OTHER | |
| lcl\|CP010341.1_prot_AJQ91640.1_1904 | RM25_RS09550 | GTP cyclohydrolase 1 | 765030 | 0 | 14303000 | 0 | 4794500 | 1615200 | 2 | 2 | 8,2 | 8,2 | 8,2 | 25,719 | 0 | 11,025 | 21477000 | 2 | AJQ91640.1_1904 | RM25_1936 | Cytoplasmic | 0 | CYT | 0.088948 | OTHER | H |
| lcl\|CP010341.1_prot_AJQ91642.1_1906 | RM25_RS09560 | Hypoxanthine phosphoribosyltransferase | 20938000 | 30258000 | 55014000 | 55577000 | 70073000 | 61086000 | 4 | 4 | 27,9 | 27,9 | 27,9 | 20,03 | 0 | 31,487 | 338860000 | 15 | AJQ91642.1_1906 | RM25_1938 | Cytoplasmic | 0 | CYT | 0.038314 | OTHER | F |
| lcl\|CP010341.1_prot_AJQ91646.1_1910 | RM25_RS09575 | D-alanyl-D-alanine carboxypeptidase, serine-type, PBP4 | 430809984 | 1080400000 | 1296600064 | 385310016 | 660750016 | 1195900032 | 8 | 8 | 24,3 | 24,3 | 24,3 | 45,932 | 0 | 157,1 | 5360400000 | 73 | AJQ91646.1_1910 | RM25_1942 | Unknown | 1 | SPI | 0.412781 | SP(Sec/SPI) | M |
| lcl\|CP010341.1_prot_AJQ91647.1_1911 | RM25_RS09580 | Inorganic diphosphatase | 32333000 | 27620000 | 103610000 | 35365000 | 38688000 | 47546000 | 4 | 4 | 24,3 | 24,3 | 24,3 | 21,173 | 0 | 26,785 | 306810000 | 13 | AJQ91647.1_1911 | RM25_1943 | Cytoplasmic | 0 | CYT | 0.125637 | OTHER | C |
| lcl\|CP010341.1_prot_AJQ91649.1_1913 | RM25_RS09590 | Phosphoglycerate mutase family protein | 5341100 | 9399800 | 0 | 8245100 | 12542000 | 10259000 | 2 | 2 | 15,6 | 15,6 | 15,6 | 19,219 | 0 | 13,597 | 62670000 | 5 | AJQ91649.1_1913 | RM25_1945 | Unknown | 0 | CYT | 0.099794 | OTHER | |
| lcl\|CP010341.1_prot_AJQ91651.1_1915 | RM25_RS09600 | Hypothetical protein | 83974000 | 86753000 | 112230000 | 13251000 | 29368000 | 45450000 | 5 | 5 | 20,6 | 20,6 | 20,6 | 37,068 | 0 | 250,76 | 387040000 | 16 | AJQ91651.1_1915 | RM25_1947 | Unknown | 1 | TMH | 0.490259 | OTHER | |
| lcl\|CP010341.1_prot_AJQ91654.1_1918 | RM25_RS09615 | Putative flavin-containing monoamine oxidase AofH | 21165000 | 21278000 | 50273000 | 13440000 | 45947000 | 28139000 | 4 | 4 | 12,9 | 12,9 | 12,9 | 49,022 | 0 | 30,637 | 184480000 | 9 | AJQ91654.1_1918 | RM25_1950 | Cytoplasmic | 0 | CYT | 0.082482 | OTHER | E |
| lcl\|CP010341.1_prot_AJQ91656.1_1920 | RM25_RS09625 | Antimicrobial peptide ABC transporter, permease prote | 4895200 | 4931700 | 6950800 | 0 | 0 | 2701700 | 1 | 1 | 2,2 | 2,2 | 2,2 | 49,602 | 0,0011614 | 10,654 | 20222000 | 4 | AJQ91656.1_1920 | RM25_1952 | CytoplasmicMembrane | 4 | SPI | 0.907309 | SP(Sec/SPI) | |
| lcl\|CP010341.1_prot_AJQ91659.1_1923 | RM25_RS12010 | Hypothetical protein | 62239000 | 60799000 | 159970000 | 71197000 | 61193000 | 118390000 | 4 | 4 | 7,7 | 7,7 | 7,7 | 56,964 | 0 | 45,473 | 593990000 | 20 | AJQ91659.1_1923 | RM25_1955 | CytoplasmicMembrane | 4 | CYT | 0.941164 | OTHER | |
| lcl\|CP010341.1_prot_AJQ91660.1_1924 | RM25_RS09650 | DNA polymerase III, delta' subunit | 0 | 1239200 | 5405600 | 5071000 | 8177200 | 5698000 | 2 | 2 | 5,7 | 5,7 | 5,7 | 48,834 | 0 | 12,763 | 32942000 | 0 | AJQ91660.1_1924 | RM25_1956 | Cytoplasmic | 0 | CYT | 0.043440 | OTHER | L |
| lcl\|CP010341.1_prot_AJQ91663.1_1927 | RM25_RS09665 | DNA topoisomerase | 66043000 | 73409000 | 143100000 | 104810000 | 102230000 | 184860000 | 16 | 16 | 21,2 | 21,2 | 21,2 | 101,75 | 0 | 213,52 | 751070000 | 50 | AJQ91663.1_1927 | RM25_1959 | Cytoplasmic | 0 | CYT | 0.262338 | OTHER | L |
| lcl\|CP010341.1_prot_AJQ91664.1_1928 | RM25_RS09670 | RRNA or tRNA methyltransferase | 4570100 | 0 | 5737000 | 5675600 | 4375100 | 12451000 | 3 | 3 | 7,3 | 7,3 | 7,3 | 54,028 | 0 | 17,375 | 39399000 | 2 | AJQ91664.1_1928 | RM25_1960 | Cytoplasmic | 0 | CYT | 0.064999 | OTHER | J |
| lcl\|CP010341.1_prot_AJQ91667.1_1931 | RM25_RS09690 | Dimethylsulfoxide reductase, chain B | 104710000 | 94978000 | 280169984 | 0 | 21067000 | 13851000 | 5 | 5 | 31,3 | 31,3 | 31,3 | 23,523 | 0 | 40,049 | 520440000 | 13 | AJQ91667.1_1931 | RM25_1963 | CytoplasmicMembrane | 0 | CYT | 0.658943 | OTHER | C |
| lcl\|CP010341.1_prot_AJQ91668.1_1932 | RM25_RS09695 | Anaerobic dimethyl sulfoxide reductase chain A | 480520000 | 363169984 | 955920000 | 15467000 | 51008000 | 82338000 | 17 | 17 | 26,3 | 26,3 | 26,3 | 94,702 | 0 | 323,31 | 1965300000 | 56 | AJQ91668.1_1932 | RM25_1964 | Cytoplasmic | 0 | CYT | 0.424969 | TAT(Tat/SPI) | C |
| lcl\|CP010341.1_prot_AJQ91670.1_1934 | RM25_1966 | Cytoplasmic chaperone TorD | 1347800 | 1721600 | 3938600 | 0 | 0 | 0 | 1 | 1 | 3,7 | 3,7 | 3,7 | 31,033 | 0,0044793 | 8,7888 | 7008000 | 2 | AJQ91670.1_1934 | RM25_1966 | Cytoplasmic | 0 | CYT | 0.195394 | OTHER | |
| lcl\|CP010341.1_prot_AJQ91682.1_1946 | RM25_RS09765 | Hypothetical protein | 639240000 | 904440000 | 1240600064 | 206960000 | 358060000 | 644649984 | 5 | 5 | 33,8 | 33,8 | 33,8 | 16,434 | 0 | 55,82 | 4160200000 | 21 | AJQ91682.1_1946 | RM25_1978 | CytoplasmicMembrane | 2 | CYT | 0.874483 | OTHER | S |
| lcl\|CP010341.1_prot_AJQ91684.1_1948 | RM25_RS12015 | Hypothetical protein | 59588000 | 63048000 | 161770000 | 41619000 | 34073000 | 90323000 | 3 | 3 | 20,1 | 20,1 | 20,1 | 27,742 | 0 | 59,363 | 495900000 | 19 | AJQ91684.1_1948 | RM25_1980 | Unknown | 0 | CYT | 0.094704 | OTHER | |
| lcl\|CP010341.1_prot_AJQ91686.1_1950 | RM25_RS09785 | Heavy metal transport/detoxification protein | 60372000 | 134360000 | 134830000 | 35065000 | 30486000 | 66641000 | 2 | 2 | 50 | 50 | 50 | 7,4384 | 0 | 21,3 | 493240000 | 6 | AJQ91686.1_1950 | RM25_1982 | Unknown | 0 | CYT | 0.674618 | OTHER | P |
| lcl\|CP010341.1_prot_AJQ91688.1_1952 | RM25_RS09795 | Replicative DNA helicase | 4108400 | 4939700 | 8121000 | 5321300 | 3940200 | 17237000 | 2 | 2 | 6 | 6 | 6 | 49,877 | 0 | 22,645 | 48685000 | 9 | AJQ91688.1_1952 | RM25_1984 | Cytoplasmic | 0 | CYT | 0.077288 | OTHER | L |
| lcl\|CP010341.1_prot_AJQ91691.1_1955 | RM25_RS09810 | NADH:flavin oxidoreductase/NADH oxidase | 4521400 | 5923000 | 38468000 | 11266000 | 14614000 | 16423000 | 1 | 1 | 3 | 3 | 3 | 39,164 | 0,0044395 | 8,4755 | 102950000 | 4 | AJQ91691.1_1955 | RM25_1987 | Unknown | 0 | CYT | 0.182725 | OTHER | C |
| lcl\|CP010341.1_prot_AJQ91698.1_1962 | RM25_RS09850 | Hypothetical protein | 16932000 | 14038000 | 30881000 | 17916000 | 24305000 | 42345000 | 1 | 1 | 11,3 | 11,3 | 11,3 | 15,348 | 0 | 43,529 | 162480000 | 7 | AJQ91698.1_1962 | RM25_1994 | CytoplasmicMembrane | 2 | CYT | 0.953696 | OTHER | |
| lcl\|CP010341.1_prot_AJQ91712.1_1976 | RM25_RS13015 | Hypothetical protein | 13114000 | 8271200 | 37403000 | 18103000 | 18268000 | 35584000 | 2 | 2 | 10 | 10 | 10 | 26,482 | 0 | 11,527 | 149380000 | 2 | AJQ91712.1_1976 | RM25_2008 | Cytoplasmic | 0 | CYT | 0.697644 | OTHER | |
| lcl\|CP010341.1_prot_AJQ91719.1_1983 | RM25_RS09955 | 6,7-dimethyl-8-ribityllumazine synthase | 2441800 | 2396800 | 5353300 | 30283000 | 29201000 | 20413000 | 3 | 3 | 27,7 | 27,7 | 27,7 | 15,277 | 0 | 21,742 | 99730000 | 8 | AJQ91719.1_1983 | RM25_2015 | Unknown | 0 | CYT | 0.024044 | OTHER | H |
| lcl\|CP010341.1_prot_AJQ91721.1_1985 | RM25_RS09965 | Riboflavin synthase, alpha subunit | 3122900 | 4692300 | 12098000 | 21562000 | 38642000 | 37843000 | 1 | 1 | 5 | 5 | 5 | 22,44 | 0,0044994 | 9,2027 | 128200000 | 7 | AJQ91721.1_1985 | RM25_2017 | Cytoplasmic | 0 | CYT | 0.043145 | OTHER | H |
| lcl\|CP010341.1_prot_AJQ91724.1_1988 | RM25_RS12030 | Hypothetical protein | 7370300 | 8450800 | 17510000 | 10011000 | 8223400 | 19050000 | 2 | 2 | 19,4 | 19,4 | 19,4 | 17,209 | 0 | 26,096 | 74020000 | 4 | AJQ91724.1_1988 | RM25_2020 | Cytoplasmic | 0 | CYT | 0.100618 | OTHER | |
| lcl\|CP010341.1_prot_AJQ91727.1_1991 | RM25_RS10000 | NAD(P) transhydrogenase subunit beta | 14921000 | 6305300 | 34170000 | 23847000 | 14048000 | 28765000 | 2 | 2 | 9,4 | 9,4 | 9,4 | 49,345 | 0 | 17,055 | 131750000 | 6 | AJQ91727.1_1991 | RM25_2023 | CytoplasmicMembrane | 8 | TMH | 0.928151 | OTHER | |
| lcl\|CP010341.1_prot_AJQ91728.1_1992 | RM25_RS10005 | NAD(P)(+) transhydrogenase (AB-specific), alpha subuni | 319230016 | 303089984 | 586099968 | 464700000 | 627600000 | 1012400000 | 14 | 14 | 40,2 | 40,2 | 40,2 | 54,277 | 0 | 316,45 | 3638200000 | 59 | AJQ91728.1_1992 | RM25_2024 | CytoplasmicMembrane | 3 | CYT | 0.534334 | OTHER | C |
| lcl\|CP010341.1_prot_AJQ91735.1_1999 | RM25_RS10040 | SubI | 1146300032 | 867729984 | 1555699968 | 498849984 | 1144000000 | 1831100032 | 12 | 12 | 51,6 | 51,6 | 51,6 | 37,833 | 0 | 288,44 | 7501600000 | 73 | AJQ91735.1_1999 | RM25_2031 | CytoplasmicMembrane | 0 | SPI | 0.592031 | TAT(Tat/SPI) | P |
| lcl\|CP010341.1_prot_AJQ91736.1_2000 | RM25_RS10045 | Ferredoxin--nitrite reductase | 260250000 | 129130000 | 496689984 | 193980000 | 195730000 | 353900000 | 20 | 20 | 40,3 | 40,3 | 40,3 | 66,165 | 0 | 323,31 | 1757000000 | 61 | AJQ91736.1_2000 | RM25_2032 | Cytoplasmic | 0 | SPI | 0.245003 | OTHER | C |
| lcl\|CP010341.1_prot_AJQ91737.1_2001 | RM25_RS10050 | Phosphoadenosine phosphosulfate reductase | 8459100 | 10675000 | 26240000 | 21849000 | 34177000 | 48556000 | 2 | 2 | 13,3 | 13,3 | 13,3 | 27,004 | 0 | 125,02 | 170630000 | 7 | AJQ91737.1_2001 | RM25_2033 | Cytoplasmic | 0 | CYT | 0.175242 | OTHER | EH |
| lcl\|CP010341.1_prot_AJQ91738.1_2002 | RM25_RS10055 | Sulfate adenylyltransferase subunit 2 | 19266000 | 13029000 | 18545000 | 50766000 | 66127000 | 60177000 | 4 | 4 | 18,6 | 18,6 | 18,6 | 39,392 | 0 | 26,215 | 267390000 | 7 | AJQ91738.1_2002 | RM25_2034 | Cytoplasmic | 0 | CYT | 0.283592 | OTHER | EH |
| lcl\|CP010341.1_prot_AJQ91739.1_2003 | RM25_RS10060 | Sulfate adenylyltransferase subunit 1 | 106900000 | 61618000 | 154480000 | 122540000 | 162970000 | 285470016 | 11 | 11 | 32,3 | 32,3 | 32,3 | 48,108 | 0 | 112,28 | 1007000000 | 40 | AJQ91739.1_2003 | RM25_2035 | Cytoplasmic | 0 | CYT | 0.058680 | OTHER | P |
| lcl\|CP010341.1_prot_AJQ91744.1_2008 | RM25_RS10085 | L-aspartate oxidase | 3523600 | 2365000 | 14456000 | 4148900 | 3690700 | 14194000 | 2 | 2 | 5,5 | 5,5 | 5,5 | 54,741 | 0 | 12,817 | 45386000 | 2 | AJQ91744.1_2008 | RM25_2040 | Cytoplasmic | 1 | TMH | 0.156413 | TAT(Tat/SPI) | H |
| lcl\|CP010341.1_prot_AJQ91747.1_2011 | RM25_RS10100 | Molybdate ABC transporter, periplasmic molybdate-bin | 132370000 | 214510000 | 259610000 | 484980000 | 797160000 | 1292300032 | 4 | 4 | 22,5 | 22,5 | 22,5 | 26,185 | 0 | 146,72 | 3629100000 | 35 | AJQ91747.1_2011 | RM25_2043 | Unknown | 0 | SPII | 0.782155 | LIPO(Sec/SPII) | P |

| | | | | | | | | | | | | | | | | | | | | | | | | | | |
|---|---|---|---|---|---|---|---|---|---|---|---|---|---|---|---|---|---|---|---|---|---|---|---|---|---|---|
| lcl\|CP010341.1_prot_AJQ91748.1_2012 | RM25_RS10105 | Molybdenum cofactor synthesis domain protein | 12424000 | 6151300 | 23451000 | 9175900 | 3104700 | 3798000 | 3 | 3 | 19 | 19 | 19 | 17,716 | 0 | 22,238 | 59091000 | 6 | AJQ91748.1_2012 | RM25_2044 | Cytoplasmic | 0 | CYT | 0.050342 | OTHER | |
| lcl\|CP010341.1_prot_AJQ91752.1_2016 | RM25_RS10125 | Nitrate reductase, alpha subunit | 1199600 | 2296500 | 2217400 | 0 | 0 | 0 | 1 | 1 | 1 | 1 | 1 | 126,75 | 0,008991 | 6,125 | 5713400 | 0 | AJQ91752.1_2016 | RM25_2048 | CytoplasmicMembrane | 0 | CYT | 0.398803 | OTHER | C |
| lcl\|CP010341.1_prot_AJQ91764.1_2028 | RM25_RS10185 | Histidine triad protein | 44504000 | 73400000 | 89869000 | 79339000 | 86372000 | 92277000 | 4 | 4 | 38,9 | 38,9 | 38,9 | 15,918 | 0 | 31,867 | 526780000 | 22 | AJQ91764.1_2028 | RM25_2060 | Cytoplasmic | 0 | CYT | 0.053880 | OTHER | |
| lcl\|CP010341.1_prot_AJQ91772.1_2036 | RM25_RS10215 | Lysine-arginine-ornithine-binding periplasmic protein | 8705600 | 26973000 | 37791000 | 0 | 5418400 | 12192000 | 3 | 3 | 11,9 | 11,9 | 11,9 | 30,121 | 0 | 26,687 | 97579000 | 5 | AJQ91772.1_2036 | RM25_2068 | Unknown | 0 | SPII | 0.866149 | LIPO(Sec/SPII) | |
| lcl\|CP010341.1_prot_AJQ91782.1_2046 | RM25_RS10265 | Two-component system response regulator | 13212000 | 15878000 | 37389000 | 12051000 | 21026000 | 22579000 | 1 | 1 | 6,8 | 6,8 | 6,8 | 23,637 | 0,0043764 | 7,9359 | 132230000 | 1 | AJQ91782.1_2046 | RM25_2078 | Cytoplasmic | 0 | CYT | 0.034196 | OTHER | T |
| lcl\|CP010341.1_prot_AJQ91793.1_2057 | RM25_RS12050 | Hypothetical protein | 34146000 | 54575000 | 75159000 | 196770000 | 173010000 | 252410000 | 1 | 1 | 2 | 2 | 2 | 35,618 | 0,0089731 | 6,0892 | 970880000 | 9 | AJQ91793.1_2057 | RM25_2089 | Unknown | 0 | SPI | 0.070062 | OTHER | D |
| lcl\|CP010341.1_prot_AJQ91799.1_2063 | RM25_RS10350 | ABC-type Fe3+-siderophores transport systems | 5969200 | 9242900 | 22015000 | 24962000 | 19704000 | 23683000 | 2 | 2 | 9,3 | 9,3 | 9,3 | 29,871 | 0 | 14,465 | 120640000 | 7 | AJQ91799.1_2063 | RM25_2095 | Cytoplasmic | 0 | CYT | 0.104304 | OTHER | P |
| lcl\|CP010341.1_prot_AJQ91800.1_2064 | RM25_RS10355 | Hypothetical protein | 12788000 | 21817000 | 52728000 | 0 | 19706000 | 29330000 | 2 | 2 | 23,9 | 23,9 | 23,9 | 21,722 | 0 | 26,52 | 155460000 | 3 | AJQ91800.1_2064 | RM25_2096 | Cytoplasmic | 0 | CYT | 0.221009 | OTHER | |
| lcl\|CP010341.1_prot_AJQ91805.1_2069 | RM25_RS10380 | Hypothetical protein | 69706 | 71303 | 0 | 0 | 49156 | 2065900 | 2 | 2 | 14,7 | 14,7 | 14,7 | 19,875 | 0,004561 | 10,115 | 3343900 | 0 | AJQ91805.1_2069 | RM25_2101 | Unknown | 0 | CYT | 0.149479 | OTHER | |
| lcl\|CP010341.1_prot_AJQ91809.1_2073 | RM25_RS10400 | Ribose-phosphate diphosphokinase | 11770000 | 8438500 | 26209000 | 15043000 | 15308000 | 30217000 | 2 | 2 | 7,4 | 7,4 | 7,4 | 35,278 | 0 | 11,809 | 115560000 | 3 | AJQ91809.1_2073 | RM25_2105 | Cytoplasmic | 0 | CYT | 0.052782 | OTHER | EF |
| lcl\|CP010341.1_prot_AJQ91811.1_2075 | RM25_RS10410 | 1D-myo-inositol 2-acetamido-2-deoxy-alpha-D-glucopyr | 83254000 | 111610000 | 255140000 | 104430000 | 130240000 | 165660000 | 6 | 6 | 30,5 | 30,5 | 30,5 | 29,259 | 0 | 154,71 | 906300000 | 34 | AJQ91811.1_2075 | RM25_2107 | Cytoplasmic | 0 | CYT | 0.121015 | OTHER | |
| lcl\|CP010341.1_prot_AJQ91813.1_2077 | RM25_RS10420 | Hypothetical protein | 595240000 | 940550016 | 1094499968 | 207880000 | 595080000 | 725520000 | 6 | 6 | 35,8 | 35,8 | 35,8 | 29,995 | 0 | 243,79 | 4356700000 | 64 | AJQ91813.1_2077 | RM25_2109 | Cytoplasmic | 0 | SPII | 0.714845 | LIPO(Sec/SPII) | |
| lcl\|CP010341.1_prot_AJQ91817.1_2081 | RM25_RS10440 | Aminoglycoside adenyltransferase | 5159100 | 7965600 | 12939000 | 0 | 0 | 4268100 | 1 | 1 | 5,4 | 5,4 | 5,4 | 30,784 | 0,0044444 | 8,4827 | 30332000 | 3 | AJQ91817.1_2081 | RM25_2113 | Cytoplasmic | 0 | CYT | 0.085054 | OTHER | |
| lcl\|CP010341.1_prot_AJQ91818.1_2082 | RM25_RS10445 | Aldose 1-epimerase | 42841000 | 48037000 | 152410000 | 43068000 | 51785000 | 82953000 | 5 | 5 | 21,8 | 21,8 | 21,8 | 32,217 | 0 | 39,302 | 455020000 | 18 | AJQ91818.1_2082 | RM25_2114 | Cytoplasmic | 0 | CYT | 0.162927 | OTHER | |
| lcl\|CP010341.1_prot_AJQ91821.1_2085 | RM25_RS10460 | Transketolase | 194080000 | 135500000 | 582460032 | 107430000 | 102490000 | 230350000 | 14 | 14 | 26,5 | 26,5 | 26,5 | 74,04 | 0 | 130,26 | 1441000000 | 42 | AJQ91821.1_2085 | RM25_2117 | Cytoplasmic | 0 | CYT | 0.099607 | OTHER | G |
| lcl\|CP010341.1_prot_AJQ91822.1_2086 | RM25_RS10465 | Ribulokinase | 0 | 0 | 15143000 | 5794300 | 5314300 | 6314200 | 1 | 1 | 2,9 | 2,9 | 2,9 | 62,864 | 0 | 20,293 | 32566000 | 2 | AJQ91822.1_2086 | RM25_2118 | Cytoplasmic | 0 | CYT | 0.073534 | OTHER | G |
| lcl\|CP010341.1_prot_AJQ91823.1_2087 | RM25_RS10470 | Sn-glycerol-1-phosphate dehydrogenase | 125910000 | 96794000 | 227080000 | 151250000 | 122310000 | 169880000 | 12 | 12 | 42,1 | 42,1 | 42,1 | 49,718 | 0 | 162,92 | 1021300000 | 32 | AJQ91823.1_2087 | RM25_2119 | Cytoplasmic | 0 | CYT | 0.101491 | OTHER | C |
| lcl\|CP010341.1_prot_AJQ91824.1_2088 | RM25_RS10475 | FGGY-family pentulose kinase | 24406000 | 25428000 | 59008000 | 2588300 | 31500000 | 50672000 | 3 | 3 | 5,5 | 5,5 | 5,5 | 56,508 | 0 | 18,005 | 196930000 | 6 | AJQ91824.1_2088 | RM25_2120 | Cytoplasmic | 0 | CYT | 0.180471 | OTHER | G |
| lcl\|CP010341.1_prot_AJQ91825.1_2089 | RM25_RS10480 | HAD-superfamily hydrolase, subfamily IIA | 26873000 | 35183000 | 67755000 | 51249000 | 41153000 | 47721000 | 5 | 5 | 20,7 | 20,7 | 20,7 | 32,348 | 0 | 40,4 | 303980000 | 13 | AJQ91825.1_2089 | RM25_2121 | Cytoplasmic | 0 | CYT | 0.113414 | OTHER | G |
| lcl\|CP010341.1_prot_AJQ91827.1_2091 | RM25_RS10490 | Ribose 5-phosphate isomerase | 132770000 | 164810000 | 377640000 | 198670000 | 201820000 | 295929984 | 6 | 6 | 45,7 | 45,7 | 45,7 | 17,161 | 0 | 51,525 | 1518100000 | 30 | AJQ91827.1_2091 | RM25_2123 | Cytoplasmic | 0 | CYT | 0.062567 | OTHER | |
| lcl\|CP010341.1_prot_AJQ91828.1_2092 | RM25_RS10495 | Serine 3-dehydrogenase | 45664000 | 42864000 | 123320000 | 31088000 | 69170000 | 57376000 | 5 | 5 | 33,5 | 33,5 | 33,5 | 23,93 | 0 | 76,33 | 397250000 | 15 | AJQ91828.1_2092 | RM25_2124 | Cytoplasmic | 0 | CYT | 0.044410 | OTHER | S |
| lcl\|CP010341.1_prot_AJQ91830.1_2094 | RM25_RS10505 | Triose-phosphate isomerase | 24775000 | 22297000 | 52868000 | 8707600 | 23054000 | 28977000 | 3 | 3 | 18,8 | 18,8 | 18,8 | 26,382 | 0 | 23,081 | 177500000 | 11 | AJQ91830.1_2094 | RM25_2126 | Cytoplasmic | 0 | CYT | 0.081204 | OTHER | G |
| lcl\|CP010341.1_prot_AJQ91831.1_2095 | RM25_RS10510 | DAK1 domain protein | 24056000 | 13325000 | 31122000 | 6260400 | 9962100 | 12939000 | 4 | 4 | 7,5 | 7,5 | 7,5 | 58,242 | 0 | 23,643 | 113120000 | 5 | AJQ91831.1_2095 | RM25_2127 | Cytoplasmic | 0 | CYT | 0.078968 | OTHER | G |
| lcl\|CP010341.1_prot_AJQ91832.1_2096 | RM25_RS10515 | Transcriptional regulator | 8872000 | 11582000 | 23679000 | 15900000 | 16373000 | 28743000 | 4 | 4 | 18,1 | 18,1 | 18,1 | 30,327 | 0 | 27,707 | 111820000 | 7 | AJQ91832.1_2096 | RM25_2128 | Cytoplasmic | 0 | CYT | 0.064355 | OTHER | K |
| lcl\|CP010341.1_prot_AJQ91845.1_2109 | RM25_RS10570 | Glycogen debranching enzyme GlgX | 18808000 | 13613000 | 60856000 | 31467000 | 30250000 | 42520000 | 3 | 3 | 4,8 | 4,8 | 4,8 | 79,701 | 0 | 21,441 | 205450000 | 10 | AJQ91845.1_2109 | RM25_2141 | Cytoplasmic | 0 | CYT | 0.246574 | OTHER | G |
| lcl\|CP010341.1_prot_AJQ91851.1_2115 | RM25_RS10600 | Nucleotidyl transferase, PF08843 family | 1716400 | 4120700 | 8344400 | 0 | 0 | 0 | 2 | 2 | 9,7 | 9,7 | 9,7 | 29,108 | 0 | 12,459 | 14181000 | 1 | AJQ91851.1_2115 | RM25_2147 | Cytoplasmic | 0 | CYT | 0.081683 | OTHER | |
| lcl\|CP010341.1_prot_AJQ91853.1_2117 | RM25_RS10610 | Hsp20/alpha crystallin family protein | 723889984 | 572640000 | 1709200000 | 631800000 | 724369984 | 1015100032 | 10 | 10 | 63,8 | 63,8 | 63,8 | 17,337 | 0 | 323,31 | 5872100000 | 64 | AJQ91853.1_2117 | RM25_2149 | Cellwall | 0 | CYT | 0.103267 | OTHER | O |
| lcl\|CP010341.1_prot_AJQ91859.1_2123 | RM25_RS10640 | Hypothetical protein | 36986000 | 25834000 | 60193000 | 47911000 | 43834000 | 47860000 | 3 | 3 | 18,9 | 18,9 | 18,9 | 17,011 | 0 | 20,108 | 284010000 | 5 | AJQ91859.1_2123 | RM25_2155 | CytoplasmicMembrane | 2 | CYT | 0.905080 | OTHER | |
| lcl\|CP010341.1_prot_AJQ91873.1_2137 | RM25_RS10705 | Nitric oxide reductase | 166080000 | 263240000 | 249740000 | 116500000 | 289129984 | 308169984 | 8 | 8 | 12,2 | 12,2 | 12,2 | 90,092 | 0 | 79,499 | 1446000000 | 19 | AJQ91873.1_2137 | RM25_2169 | CytoplasmicMembrane | 14 | CYT | 0.951724 | OTHER | |
| lcl\|CP010341.1_prot_AJQ91891.1_2155 | RM25_RS10795 | Ornithine carbamoyltransferase | 4166600 | 2469700 | 20604000 | 2512900 | 0 | 2900800 | 2 | 2 | 7,2 | 7,2 | 7,2 | 36,663 | 0 | 18,57 | 35887000 | 2 | AJQ91891.1_2155 | RM25_2187 | Cytoplasmic | 0 | CYT | 0.106910 | OTHER | E |
| lcl\|CP010341.1_prot_AJQ91894.1_2158 | RM25_RS10810 | Cyanophycinase | 20338000 | 16416000 | 38318000 | 18923000 | 16846000 | 26499000 | 3 | 3 | 21,1 | 21,1 | 21,1 | 30,854 | 0 | 25,074 | 157490000 | 2 | AJQ91894.1_2158 | RM25_2190 | CytoplasmicMembrane | 0 | CYT | 0.287540 | OTHER | PQ |
| lcl\|CP010341.1_prot_AJQ91900.1_2164 | RM25_RS10840 | Glycerol-3-phosphate dehydrogenase NAD(P)+ | 5899100 | 8452300 | 11199000 | 0 | 2447500 | 3028200 | 2 | 2 | 7,4 | 7,4 | 7,4 | 36,011 | 0 | 11,1 | 31026000 | 1 | AJQ91900.1_2164 | RM25_2196 | Unknown | 0 | SPI | 0.049419 | OTHER | |
| lcl\|CP010341.1_prot_AJQ91903.1_2167 | RM25_RS10855 | Aliphatic compound ABC transporter, periplasmic subst | 24885000 | 15310000 | 21247000 | 26086000 | 31024000 | 54060000 | 3 | 3 | 12,6 | 12,6 | 12,6 | 34,78 | 0 | 16,285 | 191670000 | 4 | AJQ91903.1_2167 | RM25_2199 | Unknown | 0 | SPII | 0.557823 | LIPO(Sec/SPII) | |
| lcl\|CP010341.1_prot_AJQ91913.1_2177 | RM25_RS10905 | Inosose dehydratase | 12421000 | 4345500 | 15559000 | 0 | 10585000 | 17349000 | 2 | 2 | 8,5 | 8,5 | 8,5 | 35,742 | 0,0045924 | 10,232 | 67361000 | 1 | AJQ91913.1_2177 | RM25_2209 | Cytoplasmic | 0 | CYT | 0.368678 | OTHER | |
| lcl\|CP010341.1_prot_AJQ91919.1_2183 | RM25_RS10935 | Glycerophosphodiester phosphodiesterase | 2918300 | 1327200 | 7793900 | 0 | 0 | 3515400 | 1 | 1 | 4,8 | 4,8 | 4,8 | 29,555 | 0,0042508 | 7,3827 | 15555000 | 2 | AJQ91919.1_2183 | RM25_2215 | Cytoplasmic | 0 | CYT | 0.075840 | OTHER | C |
| lcl\|CP010341.1_prot_AJQ91920.1_2184 | RM25_RS10940 | Inner membrane protein yihN | 2928700 | 2157600 | 5975200 | 4524100 | 8616700 | 13016000 | 1 | 1 | 4,2 | 4,2 | 4,2 | 48,531 | 0 | 15,321 | 40505000 | 5 | AJQ91920.1_2184 | RM25_2216 | CytoplasmicMembrane | 12 | TMH | 0.966902 | OTHER | |
| lcl\|CP010341.1_prot_AJQ91925.1_2189 | RM25_RS10965 | Sulfurtransferase | 41989000 | 52003000 | 101430000 | 55283000 | 47192000 | 65923000 | 6 | 6 | 25,4 | 25,4 | 25,4 | 29,004 | 0 | 38,887 | 394670000 | 19 | AJQ91925.1_2189 | RM25_2221 | Cytoplasmic | 0 | CYT | 0.751594 | OTHER | P |
| lcl\|CP010341.1_prot_AJQ91926.1_2190 | RM25_RS10970 | Glutamate decarboxylase | 8361600 | 3926500 | 31777000 | 13690000 | 6348800 | 29056000 | 3 | 3 | 7,2 | 7,2 | 7,2 | 52,976 | 0 | 19,004 | 106420000 | 4 | AJQ91926.1_2190 | RM25_2222 | Cytoplasmic | 0 | CYT | 0.190525 | OTHER | E |
| lcl\|CP010341.1_prot_AJQ91930.1_2194 | RM25_RS12100 | Conserved transmembrane protein | 21808000 | 45596000 | 0 | 8984300 | 0 | 0 | 1 | 1 | 3,5 | 3,5 | 3,5 | 55,13 | 0,0023121 | 10,479 | 76389000 | 2 | AJQ91930.1_2194 | RM25_2226 | CytoplasmicMembrane | 9 | TMH | 0.922888 | OTHER | M |
| lcl\|CP010341.1_prot_AJQ91936.1_2200 | RM25_RS11025 | Putative alcohol dehydrogenase AdhA | 16589000 | 12183000 | 50579000 | 25492000 | 23312000 | 31892000 | 3 | 3 | 8 | 8 | 8 | 35,92 | 0 | 19,973 | 177040000 | 8 | AJQ91936.1_2200 | RM25_2232 | Cytoplasmic | 0 | CYT | 0.072622 | OTHER | |
| lcl\|CP010341.1_prot_AJQ91937.1_2201 | RM25_RS11030 | Methionine aminotransferase | 3570600 | 4373000 | 6975700 | 0 | 0 | 0 | 1 | 1 | 5,6 | 5,6 | 5,6 | 43,488 | 0,0044743 | 8,6361 | 14919000 | 0 | AJQ91937.1_2201 | RM25_2233 | Cytoplasmic | 0 | CYT | 0.067682 | OTHER | E |
| lcl\|CP010341.1_prot_AJQ91940.1_2204 | RM25_RS11040 | Hypothetical protein | 21837000 | 26729000 | 42643000 | 19793000 | 21580000 | 10267000 | 2 | 2 | 9,2 | 9,2 | 9,2 | 29,24 | 0 | 14,456 | 142850000 | 5 | AJQ91940.1_2204 | RM25_2236 | Cytoplasmic | 0 | CYT | 0.316482 | OTHER | |
| lcl\|CP010341.1_prot_AJQ91948.1_2212 | RM25_RS11075 | 50S ribosomal protein L9 | 547969984 | 679950016 | 1457600000 | 570240000 | 929560000 | 1398400000 | 7 | 7 | 60,7 | 60,7 | 60,7 | 16,069 | 0 | 266,64 | 6129400000 | 51 | AJQ91948.1_2212 | RM25_2244 | Cytoplasmic | 0 | CYT | 0.057220 | OTHER | J |
| lcl\|CP010341.1_prot_AJQ91949.1_2213 | RM25_RS11080 | Hypothetical protein | 81810000 | 70395000 | 200500000 | 130210000 | 184310000 | 282710016 | 2 | 2 | 26,6 | 26,6 | 26,6 | 8,9764 | 0 | 62,431 | 1065200000 | 20 | AJQ91949.1_2213 | RM25_2245 | Cytoplasmic | 0 | CYT | 0.623945 | OTHER | J |
| lcl\|CP010341.1_prot_AJQ91950.1_2214 | RM25_RS11085 | Single-stranded DNA-binding protein 2 | 28037000 | 20797000 | 71378000 | 42450000 | 27763000 | 73333000 | 3 | 3 | 28,2 | 28,2 | 28,2 | 19,651 | 0 | 49,714 | 292320000 | 7 | AJQ91950.1_2214 | RM25_2246 | Cytoplasmic | 0 | CYT | 0.852414 | OTHER | L |
| lcl\|CP010341.1_prot_AJQ91951.1_2215 | RM25_RS11090 | 30S ribosomal protein S6 | 151040000 | 189550000 | 558860032 | 453020000 | 520780000 | 674700032 | 5 | 5 | 48,4 | 48,4 | 48,4 | 10,886 | 0 | 77,522 | 2829200000 | 36 | AJQ91951.1_2215 | RM25_2247 | Cytoplasmic | 0 | CYT | 0.357149 | OTHER | J |
| lcl\|CP010341.1_prot_AJQ91957.1_2221 | RM25_RS12825 | Hypothetical protein | 3949600000 | 5196499968 | 7267700224 | 1498000000 | 2857400064 | 4338800128 | 32 | 32 | 53,4 | 53,4 | 53,4 | 113,89 | 0 | 323,31 | 2,6233E+10 | 206 | AJQ91957.1_2221 | RM25_2253 | Cellwall | 1 | SPI | 0.881818 | SP(Sec/SPI) | |
| lcl\|CP010341.1_prot_AJQ91960.1_2224 | RM25_RS11135 | DNA-3-methyladenine glycosylase I | 3306800 | 0 | 4840300 | 5964800 | 5754400 | 19796000 | 1 | 1 | 4,3 | 4,3 | 4,3 | 22,599 | 0,0089552 | 6,0483 | 39662000 | 1 | AJQ91960.1_2224 | RM25_2256 | Cytoplasmic | 0 | CYT | 0.121471 | OTHER | L |
| lcl\|CP010341.1_prot_AJQ91965.1_2229 | RM25_RS11160 | Bifunctional penicillin-binding protein ponA1 | 3920300032 | 6255699968 | 9544200192 | 3356000000 | 5430700032 | 8431300096 | 25 | 25 | 57,4 | 57,4 | 57,4 | 75,486 | 0 | 323,31 | 3,9443E+10 | 245 | AJQ91965.1_2229 | RM25_2261 | CytoplasmicMembrane | 1 | CYT | 0.912201 | OTHER | M |
| lcl\|CP010341.1_prot_AJQ91969.1_2233 | RM25_RS11185 | Ferritin-like protein | 122210000 | 39090000 | 403910016 | 232020000 | 241860000 | 304870016 | 7 | 7 | 40,7 | 40,7 | 40,7 | 19,394 | 0 | 173,52 | 1466500000 | 44 | AJQ91969.1_2233 | RM25_2266 | Cytoplasmic | 0 | CYT | 0.082425 | OTHER | P |
| lcl\|CP010341.1_prot_AJQ91976.1_2240 | RM25_RS11220 | Putative nitrite reductase, copper-dependent | 704020 | 687970 | 2392500 | 0 | 1907800 | 1661800 | 1 | 1 | 2,1 | 2,1 | 2,1 | 97,789 | 0,0041494 | 6,8538 | 8514000 | 3 | AJQ91976.1_2240 | RM25_2273 | CytoplasmicMembrane | 11 | TMH | 0.861011 | OTHER | |
| lcl\|CP010341.1_prot_AJQ91981.1_2245 | RM25_RS11245 | Transmembrane protein | 13447000 | 8569800 | 24885000 | 22686000 | 25281000 | 29899000 | 1 | 1 | 11,5 | 11,5 | 11,5 | 13,199 | 0 | 12,936 | 139010000 | 6 | AJQ91981.1_2245 | RM25_2278 | CytoplasmicMembrane | 4 | TMH | 0.972411 | OTHER | |
| lcl\|CP010341.1_prot_AJQ91984.1_2248 | RM25_RS11255 | Hypothetical protein | 6867400 | 7562800 | 9327500 | 7275400 | 14032000 | 13199000 | 2 | 2 | 10,7 | 10,7 | 10,7 | 34,95 | 0 | 10,857 | 63098000 | 0 | AJQ91984.1_2248 | RM25_2281 | Cytoplasmic | 0 | CYT | 0.057064 | OTHER | |
| lcl\|CP010341.1_prot_AJQ91999.1_2263 | RM25_RS11325 | 2,3-diaminopropionate biosynthesis protein SbnA | 6738200 | 12338000 | 7278700 | 0 | 0 | 10591000 | 1 | 1 | 4,6 | 4,6 | 4,6 | 40,85 | 0 | 19,454 | 36946000 | 3 | AJQ91999.1_2263 | RM25_2296 | Cytoplasmic | 0 | CYT | 0.107169 | OTHER | E |
| lcl\|CP010341.1_prot_AJQ92000.1_2264 | RM25_RS11330 | Pyridoxal-phosphate dependent enzyme | 0 | 7402300 | 2543400 | 0 | 5997500 | 11863000 | 2 | 2 | 3,8 | 3,8 | 3,8 | 70,839 | 0,0045977 | 10,271 | 29421000 | 2 | AJQ92000.1_2264 | RM25_2297 | Cytoplasmic | 0 | CYT | 0.118957 | OTHER | |
| lcl\|CP010341.1_prot_AJQ92001.1_2265 | RM25_RS11335 | Decarboxylase | 3316300 | 3639800 | 8480400 | 2612100 | 4360200 | 6270200 | 1 | 1 | 3,7 | 3,7 | 3,7 | 60,565 | 0,0046083 | 10,31 | 31091000 | 4 | AJQ92001.1_2265 | RM25_2298 | Cytoplasmic | 0 | CYT | 0.101388 | OTHER | E |
| lcl\|CP010341.1_prot_AJQ92010.1_2274 | RM25_RS11385 | Cytochrome P450 | 67290000 | 89346000 | 168690000 | 44049000 | 62793000 | 98136000 | 8 | 8 | 27,7 | 27,7 | 27,7 | 44,283 | 0 | 94,449 | 558210000 | 26 | AJQ92010.1_2274 | RM25_2307 | CytoplasmicMembrane | 0 | CYT | 0.119302 | OTHER | |
| lcl\|CP010341.1_prot_AJQ92011.1_2275 | RM25_RS11390 | Putative glyoxal reductase | 125240000 | 209710000 | 296569984 | 218830000 | 205050000 | 214950000 | 9 | 9 | 40,1 | 40,1 | 40,1 | 31,411 | 0 | 254,73 | 1418400000 | 43 | AJQ92011.1_2275 | RM25_2308 | Cytoplasmic | 0 | CYT | 0.203071 | OTHER | |
| lcl\|CP010341.1_prot_AJQ92015.1_2279 | RM25_RS11410 | Xanthine permease | 3010200 | 2908000 | 5032500 | 3957100 | 6054300 | 19915000 | 2 | 2 | 3,4 | 3,4 | 3,4 | 68,401 | 0 | 12,351 | 40878000 | 5 | AJQ92015.1_2279 | RM25_2312 | CytoplasmicMembrane | 12 | CYT | 0.948355 | OTHER | F |
| lcl\|CP010341.1_prot_AJQ92016.1_2280 | RM25_RS11415 | Putative xylan esterase | 18046000 | 12191000 | 40517000 | 12148000 | 8547500 | 19753000 | 3 | 3 | 15,3 | 15,3 | 15,3 | 35,23 | 0 | 31,615 | 116900000 | 4 | AJQ92016.1_2280 | RM25_2313 | CytoplasmicMembrane | 0 | CYT | 0.057841 | OTHER | |
| lcl\|CP010341.1_prot_AJQ92018.1_2282 | RM25_RS11425 | Catalase | 30520000 | 17034000 | 115770000 | 44299000 | 20617000 | 22714000 | 6 | 6 | 13,1 | 13,1 | 13,1 | 53,589 | 0 | 48,808 | 265350000 | 13 | AJQ92018.1_2282 | RM25_2315 | Cytoplasmic | 0 | CYT | 0.361989 | OTHER | P |
| lcl\|CP010341.1_prot_AJQ92019.1_2283 | RM25_RS11430 | Dyp-type peroxidase family protein | 9926200 | 13329000 | 26729000 | 15138000 | 10635000 | 6112700 | 2 | 2 | 8,5 | 8,5 | 8,5 | 32,948 | 0 | 14,922 | 95248000 | 6 | AJQ92019.1_2283 | RM25_2316 | Unknown | 0 | CYT | 0.202532 | OTHER | |
| lcl\|CP010341.1_prot_AJQ92021.1_2285 | RM25_RS12125 | Hypothetical protein | 9515600 | 13905000 | 25211000 | 30093000 | 25231000 | 44641000 | 1 | 1 | 10,8 | 10,8 | 10,8 | 25,277 | 0 | 34,733 | 171880000 | 6 | AJQ92021.1_2285 | RM25_2318 | Unknown | 2 | TMH | 0.964759 | OTHER | |
| lcl\|CP010341.1_prot_AJQ92026.1_2290 | RM25_RS11465 | Fructose-bisphosphate aldolase class 1 | 275540000 | 335120000 | 516129984 | 280169984 | 315220000 | 337020000 | 12 | 12 | 61 | 61 | 61 | 32,33 | 0 | 304,53 | 2248800000 | 65 | AJQ92026.1_2290 | RM25_2323 | Unknown | 0 | CYT | 0.072291 | OTHER | |
| lcl\|CP010341.1_prot_AJQ92027.1_2291 | RM25_RS11470 | YceI family protein | 69323000 | 81310000 | 135950000 | 161230000 | 164470000 | 143170000 | 5 | 5 | 32,8 | 32,8 | 32,8 | 19,955 | 0 | 43,945 | 842240000 | 23 | AJQ92027.1_2291 | RM25_2324 | Cytoplasmic | 0 | CYT | 0.446175 | OTHER | S |
| lcl\|CP010341.1_prot_AJQ92028.1_2292 | RM25_RS11475 | Citrate (Si)-synthase | 22123000 | 11672000 | 80762000 | 51078000 | 45182000 | 55101000 | 5 | 5 | 12,9 | 12,9 | 12,9 | 47,554 | 0 | 35,756 | 294370000 | 8 | AJQ92028.1_2292 | RM25_2325 | Cytoplasmic | 0 | CYT | 0.118794 | OTHER | C |
| lcl\|CP010341.1_prot_AJQ92029.1_2293 | RM25_RS11480 | L-glyceraldehyde 3-phosphate reductase | 0 | 0 | 0 | 0 | 4691000 | 11138000 | 1 | 1 | 2,8 | 2,8 | 2,8 | 38,79 | 0,0041322 | 6,7952 | 15829000 | 1 | AJQ92029.1_2293 | RM25_2326 | Cytoplasmic | 0 | CYT | 0.143980 | OTHER | C |
| lcl\|CP010341.1_prot_AJQ92032.1_2296 | RM25_RS11495 | Membrane protein YdfJ | 1909200000 | 2438599936 | 3756400128 | 1518300032 | 1812899968 | 2734899968 | 20 | 20 | 35,5 | 35,5 | 35,5 | 87,682 | 0 | 323,31 | 1,5173E+10 | 148 | AJQ92032.1_2296 | RM25_2329 | CytoplasmicMembrane | 12 | TMH | 0.896021 | SP(Sec/SPI) | P |
| lcl\|CP010341.1_prot_AJQ92034.1_2298 | RM25_RS11505 | Putative NAD(P) binding enzyme | 123790000 | 129710000 | 290169984 | 90765000 | 100640000 | 133760000 | 8 | 8 | 55 | 55 | 55 | 23,456 | 0 | 103,6 | 952870000 | 32 | AJQ92034.1_2298 | RM25_2331 | Cytoplasmic | 0 | CYT | 0.108450 | OTHER | |
| lcl\|CP010341.1_prot_AJQ92040.1_2304 | RM25_RS12135 | MscS transporter, small conductance mechanosensitive | 9789200 | 17031000 | 26387000 | 18830000 | 35672000 | 25709000 | 5 | 5 | 15,2 | 15,2 | 15,2 | 62,228 | 0 | 163,04 | 139320000 | 17 | AJQ92040.1_2304 | RM25_2337 | Unknown | 2 | CYT | 0.805368 | OTHER | M |
| lcl\|CP010341.1_prot_AJQ92041.1_2305 | RM25_RS11540 | CCA tRNA nucleotidyltransferase | 16398000 | 6774000 | 35315000 | 20810000 | 6359800 | 26026000 | 4 | 4 | 11,1 | 11,1 | 11,1 | 55,274 | 0 | 29,641 | 125230000 | 5 | AJQ92041.1_2305 | RM25_2338 | Cytoplasmic | 0 | CYT | 0.067919 | OTHER | J |
| lcl\|CP010341.1_prot_AJQ92043.1_2307 | RM25_RS11550 | Hypothetical protein | 10357000 | 10673000 | 9206500 | 1091300 | 2255500 | 2607000 | 4 | 4 | 5,7 | 5,7 | 5,7 | 72,563 | 0 | 26,562 | 37381000 | 6 | AJQ92043.1_2307 | RM25_2340 | Unknown | 2 | TMH | 0.789696 | SP(Sec/SPI) | |
| lcl\|CP010341.1_prot_AJQ92044.1_2308 | RM25_RS12140 | Hypothetical protein | 100010000 | 146810000 | 161500000 | 16334000 | 40109000 | 52988000 | 3 | 3 | 5,5 | 5,5 | 5,5 | 73,273 | 0 | 58,08 | 535220000 | 5 | AJQ92044.1_2308 | RM25_2341 | Cytoplasmic | 1 | CYT | 0.582451 | OTHER | |
| lcl\|CP010341.1_prot_AJQ92045.1_2309 | RM25_RS11560 | Phosphoglycerate mutase family protein | 26850000 | 26204000 | 75114000 | 25694000 | 23908000 | 34411000 | 3 | 3 | 15,5 | 15,5 | 15,5 | 26,155 | 0 | 21,237 | 231310000 | 6 | AJQ92045.1_2309 | RM25_2342 | Unknown | 0 | CYT | 0.545056 | OTHER | |
| lcl\|CP010341.1_prot_AJQ92046.1_2310 | RM25_RS11565 | Putative exported solute-binding protein | 5317500 | 12548000 | 20679000 | 18871000 | 17216000 | 31281000 | 2 | 2 | 5,7 | 5,7 | 5,7 | 46,173 | 0 | 12,459 | 117830000 | 5 | AJQ92046.1_2310 | RM25_2343 | CytoplasmicMembrane | 0 | SPII | 0.422605 | LIPO(Sec/SPII) | |
| lcl\|CP010341.1_prot_AJQ92047.1_2311 | RM25_RS11570 | Thioredoxin-disulfide reductase | 151370000 | 163870000 | 237340000 | 250710000 | 231210000 | 301060000 | 8 | 8 | 36 | 36 | 36 | 34,621 | 0 | 112,71 | 1546700000 | 57 | AJQ92047.1_2311 | RM25_2344 | Cytoplasmic | 0 | CYT | 0.066632 | OTHER | C |
| lcl\|CP010341.1_prot_AJQ92048.1_2312 | RM25_RS11575 | Thioredoxin | 56832000 | 119520000 | 137540000 | 152520000 | 141760000 | 160010000 | 2 | 2 | 33,6 | 33,6 | 33,6 | 11,847 | 0 | 57,034 | 900610000 | 8 | AJQ92048.1_2312 | RM25_2345 | Cytoplasmic | 0 | CYT | 0.774800 | OTHER | O |
| lcl\|CP010341.1_prot_AJQ92049.1_2313 | RM25_RS11580 | OsmC family protein | 42175000 | 55976000 | 122160000 | 97741000 | 89639000 | 81376000 | 5 | 5 | 44,6 | 44,6 | 44,6 | 15,735 | 0 | 39,736 | 571140000 | 21 | AJQ92049.1_2313 | RM25_2346 | Unknown | 0 | CYT | 0.320057 | OTHER | |
| lcl\|CP010341.1_prot_AJQ92050.1_2314 | RM25_RS11585 | Chromosome partitioning protein ParB | 14185000 | 25267000 | 65882000 | 32116000 | 27458000 | 34703000 | 6 | 6 | 22,9 | 22,9 | 22,9 | 36,187 | 0 | 43,071 | 212250000 | 11 | AJQ92050.1_2314 | RM25_2347 | Cytoplasmic | 0 | CYT | 0.076717 | OTHER | K |
| lcl\|CP010341.1_prot_AJQ92051.1_2315 | RM25_2348 | Chromosome partitioning protein | 27810000 | 37629000 | 73119000 | 28690000 | 23658000 | 42151000 | 8 | 8 | 20 | 20 | 20 | 59,86 | 0 | 57,698 | 251870000 | 18 | AJQ92051.1_2315 | RM25_2348 | Cytoplasmic | 0 | CYT | 0.720355 | OTHER | D |
| lcl\|CP010341.1_prot_AJQ92052.1_2316 | RM25_RS11600 | 16S rRNA (guanine(527)-N(7))-methyltransferase | 6631000 | 7566800 | 16541000 | 5177200 | 16758000 | 23384000 | 3 | 3 | 16 | 16 | 16 | 27,831 | 0 | 24,72 | 76059000 | 7 | AJQ92052.1_2316 | RM25_2349 | Cytoplasmic | 0 | CYT | 0.055610 | OTHER | J |
| lcl\|CP010341.1_prot_AJQ92053.1_2317 | RM25_2350 | Hypothetical protein | 44967000 | 65210000 | 109330000 | 67219000 | 71803000 | 134720000 | 5 | 5 | 21,3 | 21,3 | 21,3 | 33,821 | 0 | 156,1 | 540910000 | 23 | AJQ92053.1_2317 | RM25_2350 | Cytoplasmic | 0 | CYT | 0.457795 | OTHER | S |
