## Supplemental Table 4 for "Aerobic Adaptation and Metabolic Dynamics of *Propionibacterium freudenreichii* DSM 20271: Insights from Comparative Transcriptomics and Surfaceome Analysis"

**Table S4.** Proteins uniquely identified in at least two out of three replica samples in aerobic or anaerobic growth conditions. The colour gradient from pink to red indicates the log-2 transformed raw intensities for each biological replicate (grey colour indicates zero). The number of transmembrane spanning domains (TMD) were predicted using TMHMM 2.0, subcellular localization of proteins were done with prediction tool PSORTb 3.0.3 PSORTb v3.0.2, and the presence of possible classical and non-classical signal peptide sequences were analysed with LipoP 1.0, SignalP 5.0, and SecretomeP 2.0. The COG categorization was accomplished with EggNOG 5.0.0.

**Uniquelly identified in Aerobic condition (O+)**

| Protein IDs | NCBI-LocusTag | Protein name (21 log2 25) | Anaerobic 1 | Anaerobic 2 | Anaerobic 3 | Aerobic 1 | Aerobic 3 | Aerobic 4 | Mol. weight [kDa] | No.Peptides matched | Sequence coverage [%] | No. of TMHMM | PsortB | LipoP | SecP Score | SignalP | COG |
|---|---|---|---|---|---|---|---|---|---|---|---|---|---|---|---|---|---|
| lcl\|CP010341.1_prot_AJQ89873.1_137 | RM25_RS00680 | Hypothetical protein | | | | | | | 36,978 | 1 | 33 | 0 | Cytoplasmic | CYT | 0.086450 | OTHER | X |
| lcl\|CP010341.1_prot_AJQ90098.1_362 | RM25_RS01775 | Integral membrane protein MviN | | | | | | | 68,944 | 1 | 28 | 14 | CytoplasmicMembrane | CYT | 0.962179 | OTHER | KLT |
| lcl\|CP010341.1_prot_AJQ90550.1_814 | RM25_RS04060 | 30S ribosomal protein S20 | | | | | | | 93,957 | 1 | 128 | 0 | Cytoplasmic | CYT | 0.860127 | OTHER | J |
| lcl\|CP010341.1_prot_AJQ90703.1_967 | RM25_RS11835 | Hypothetical protein | | | | | | | 20,372 | 1 | 8 | 0 | Unknown | CYT | 0.258589 | OTHER | X |
| lcl\|CP010341.1_prot_AJQ91236.1_1500 | RM25_RS12870 | Hypothetical protein | | | | | | | 29,728 | 1 | 58 | 2 | CytoplasmicMembrane | CYT | 0.964822 | OTHER | X |
| lcl\|CP010341.1_prot_AJQ91374.1_1638 | RM25_RS08190 | mannosyltransferase | | | | | | | 58,362 | 1 | 17 | 11 | CytoplasmicMembrane | TMH | 0.951139 | OTHER | O |
| lcl\|CP010341.1_prot_AJQ91567.1_1831 | NA | Hypothetical protein | | | | | | | 11,931 | 1 | 19 | 0 | Unknown | CYT | 0.143159 | OTHER | X |
| lcl\|CP010341.1_prot_AJQ92029.1_2293 | RM25_RS11480 | L-glyceraldehyde 3-phosphate reductase | | | | | | | 3,879 | 1 | 28 | 0 | Cytoplasmic | CYT | 0.143980 | OTHER | C |

**Uniquelly identified in Anaerobic condition (N+)**

| Protein IDs | NCBI-LocusTag | Protein name (20 log2 27) | Anaerobic 1 | Anaerobic 2 | Anaerobic 3 | Aerobic 1 | Aerobic 3 | Aerobic 4 | Mol. weight [kDa] | Unique peptides | Sequence coverage [%] | No. of TMHMM | PsortB | LipoP | SecP Score | SignalP | COG |
|---|---|---|---|---|---|---|---|---|---|---|---|---|---|---|---|---|---|
| lcl\|CP010341.1_prot_AJQ89897.1_161 | RM25_RS00815 | 3-carboxymuconate cyclase (Precursor) | | | | | | | 38,268 | 3 | 152 | 0 | Cytoplasmic | CYT | 0.162919 | OTHER | X |
| lcl\|CP010341.1_prot_AJQ89936.1_200 | RM25_RS01010 | containing protein | | | | | | | 26,381 | 1 | 112 | 0 | Unknown | CYT | 0.487471 | OTHER | X |
| lcl\|CP010341.1_prot_AJQ89980.1_244 | RM25_RS01220 | Thermostable beta-glucosidase B | | | | | | | 85,968 | 1 | 18 | 0 | Cytoplasmic | CYT | 0.063195 | OTHER | X |
| lcl\|CP010341.1_prot_AJQ90154.1_418 | RM25_RS02060 | Cobalt ABC transporter, ATPase subunit | | | | | | | 31,244 | 1 | 61 | 0 | CytoplasmicMembrane | CYT | 0.067773 | OTHER | P |
| lcl\|CP010341.1_prot_AJQ90430.1_694 | RM25_RS03435 | Hypothetical protein | | | | | | | 11,266 | 1 | 1 | 9 | CytoplasmicMembrane | CYT | 0.822891 | OTHER | S |
| lcl\|CP010341.1_prot_AJQ90568.1_832 | RM25_RS04150 | CBS domain protein | | | | | | | 47,868 | 1 | 28 | 3 | CytoplasmicMembrane | CYT | 0.360026 | SP(Sec/SPI | S |
| lcl\|CP010341.1_prot_AJQ90751.1_1015 | RM25_RS05090 | Recombination factor protein RarA | | | | | | | 50,213 | 1 | 34 | 0 | Unknown | CYT | 0.071135 | OTHER | L |
| lcl\|CP010341.1_prot_AJQ90829.1_1093 | RM25_RS05485 | Maltose alpha-D-glucosyltransferase | | | | | | | 68,428 | 2 | 63 | 0 | Cytoplasmic | CYT | 0.300070 | OTHER | G |
| lcl\|CP010341.1_prot_AJQ90889.1_1153 | RM25_RS05790 | Prolyl aminopeptidase | | | | | | | 44,399 | 1 | 32 | 0 | Cytoplasmic | CYT | 0.158976 | OTHER | S |
| lcl\|CP010341.1_prot_AJQ90973.1_1237 | RM25_RS06215 | Ethanolamine utilization protein EutE | | | | | | | 50,118 | 1 | 26 | 0 | Cytoplasmic | CYT | 0.069359 | OTHER | C |
| lcl\|CP010341.1_prot_AJQ91027.1_1291 | RM25_RS06485 | Argininosuccinate lyase | | | | | | | 50,937 | 1 | 36 | 0 | Cytoplasmic | CYT | 0.073832 | OTHER | E |
| lcl\|CP010341.1_prot_AJQ91133.1_1397 | RM25_RS07020 | Glutamine amidotransferase subunit PdxT | | | | | | | 21,169 | 1 | 58 | 0 | Cytoplasmic | CYT | 0.024016 | OTHER | X |
| lcl\|CP010341.1_prot_AJQ91185.1_1449 | RM25_RS12605 | Hypothetical protein | | | | | | | 17,738 | 1 | 14 | 1 | Unknown | CYT | 0.927041 | OTHER | X |
| lcl\|CP010341.1_prot_AJQ91440.1_1704 | RM25_RS08545 | Hypothetical protein | | | | | | | 13,819 | 1 | 126 | 0 | Cytoplasmic | CYT | 0.067880 | OTHER | X |
| lcl\|CP010341.1_prot_AJQ91605.1_1869 | RM25_RS09375 | Polyphosphate kinase | | | | | | | 82,373 | 3 | 59 | 0 | CytoplasmicMembrane | CYT | 0.062033 | OTHER | P |
| lcl\|CP010341.1_prot_AJQ91670.1_1934 | NA | Cytoplasmic chaperone TorD | | | | | | | 31,033 | 1 | 37 | 0 | Cytoplasmic | CYT | 0.195394 | OTHER | X |
| lcl\|CP010341.1_prot_AJQ91752.1_2016 | RM25_RS10125 | Nitrate reductase, alpha subunit | | | | | | | 12,675 | 1 | 1 | 0 | CytoplasmicMembrane | CYT | 0.398803 | OTHER | C |
| lcl\|CP010341.1_prot_AJQ91851.1_2115 | RM25_RS10600 | Nucleotidyl transferase, PF08843 family | | | | | | | 29,108 | 2 | 97 | 0 | Cytoplasmic | CYT | 0.081683 | OTHER | X |
| lcl\|CP010341.1_prot_AJQ91937.1_2201 | RM25_RS11030 | Methionine aminotransferase | | | | | | | 43,488 | 1 | 56 | 0 | Cytoplasmic | CYT | 0.067682 | OTHER | E |
