## Supplemental Table 5 for "Aerobic Adaptation and Metabolic Dynamics of *Propionibacterium freudenreichii* DSM 20271: Insights from Comparative Transcriptomics and Surfaceome Analysis"

**Table_S5.** List of proteins showing statistically significant protein abundance change between the aerobic and anaerobic surfaceomes. For this purpose, the LFQ intensities after log2 conversion and missing value imputation were subjected to two-sample t-test with a permutation-based false discovery rate of 0.05 to confirm the statistical significance. The number of transmembrane spanning domains (TMD) were predicted using TMHMM 2.0, subcellular localization of proteins were done with prediction tool PSORTb 3.0.3 PSORTb v3.0.2, and the presence of possible classical and non-classical signal peptide sequences were analysed with LipoP 1.0, SignalP 5.0, and SecretomeP 2.0. The COG categorization was accomplished with EggNOG 5.0.0.

**Abundance increased under Aerobic (O+) condition**

| | | | | | | | | | Log2 Transfomed LFQ intensities | | | | | | | | | | | | | | | | | | | | | |
|---|---|---|---|---|---|---|---|---|---|---|---|---|---|---|---|---|---|---|---|---|---|---|---|---|---|---|---|---|---|---|
| Protein IDs | NCBI-LocusTag | Protein name | No. of TMHMM | PsortB | LipoP | SecP Score | SignalP | COG | Anaerobic 1 | Anaerobic 2 | Anaerobic 3 | Aerobic 1 | Aerobic 3 | Aerobic 4 | Student's T-test Sign Anaerobic_Aerobic | Razor + unique peptides | Unique peptides | Sequence coverage [%] | Unique + razor sequence coverage [%] | Unique sequence coverage [%] | Mol. weight [kDa] | Q-value | Score | Intensity | MS/MS count | -Log Student's T-test p-value Anaerobic_Aerobic | Student's T-test q-value Anaerobic_Aerobic | Student's T-test Difference Anaerobic_Aerobic | Student's T-test Test statistic Anaerobic_Aerobic | LocusTag |
| lcl\|CP010341.1_prot_AJQ89910.1_174 | RM25_RS00880 | Cytochrome bd-type menaquinol ox | 9 | CytoplasmicMembrane | TMH | 0,949 | OTHER | C | 26,311 | 26,792 | 26,154 | 29,073 | 29,817 | 29,732 | + | 9 | 9 | 26,8 | 26,8 | 26,8 | 55,025 | 0 | 134,84 | 5967700000 | 41 | 3,297434504 | 0,034333333 | -3,12153244 | -10,28427926 | RM25_0178 |
| lcl\|CP010341.1_prot_AJQ91497.1_1761 | RM25_RS08840 | Hypothetical protein | 0 | CytoplasmicMembrane | CYT | 0,037 | OTHER | X | 21,643 | 23,262 | 22,237 | 26,027 | 25,173 | 25,276 | + | 2 | 2 | 18,3 | 18,3 | 18,3 | 88,387 | 0 | 20,649 | 424430000 | 13 | 2,334804885 | 0,03952 | -3,111777623 | -5,718856121 | RM25_1793 |
| lcl\|CP010341.1_prot_AJQ89886.1_150 | RM25_RS00745 | putative thiol peroxidase | 0 | Unknown | CYT | 0,060 | OTHER | O | 24,504 | 24,718 | 25,743 | 27,776 | 27,446 | 27,158 | + | 4 | 4 | 45,5 | 45,5 | 45,5 | 17,439 | 0 | 96,766 | 1572600000 | 25 | 2,3718069 | 0,043911111 | -2,471943537 | -5,854198855 | RM25_0151 |
| lcl\|CP010341.1_prot_AJQ89975.1_239 | RM25_RS01195 | Putative iron-siderophore transport | 0 | Unknown | SPII | 0,817 | LIPO(Sec/SPII) | P | 24,049 | 22,448 | 24,539 | 26,039 | 26,025 | 26,112 | + | 2 | 2 | 1,3 | 1,3 | 1,3 | 34,454 | 0 | 44,363 | 498810000 | 14 | 1,706418398 | 0,044818898 | -2,38034757 | -3,766735155 | RM25_0243 |
| lcl\|CP010341.1_prot_AJQ90937.1_1201 | RM25_RS06040 | Hypothetical protein | 0 | Cytoplasmic | CYT | 0,086 | OTHER | S | 22,342 | 22,699 | 22,898 | 24,964 | 24,812 | 24,834 | + | 2 | 2 | 18,4 | 18,4 | 18,4 | 15,463 | 0 | 19,003 | 297510000 | 6 | 3,709742343 | 0,016 | -2,223818461 | -13,11632658 | RM25_1223 |
| lcl\|CP010341.1_prot_AJQ90670.1_934 | RM25_RS04670 | Phosphatidylinositol alpha-mannosy | 0 | Unknown | CYT | 0,052 | OTHER | M | 22,340 | 21,987 | 22,007 | 24,008 | 23,972 | 23,711 | + | 4 | 4 | 1,6 | 1,6 | 1,6 | 41,055 | 0 | 26,338 | 131710000 | 5 | 3,567180953 | 0,020888889 | -1,785416285 | -12,06208795 | RM25_0951 |
| lcl\|CP010341.1_prot_AJQ91719.1_1983 | RM25_RS09955 | 6,7-dimethyl-8-ribityllumazine syntl | 0 | Unknown | CYT | 0,024 | OTHER | H | 21,890 | 22,962 | 21,787 | 24,391 | 24,033 | 23,539 | + | 3 | 3 | 27,7 | 27,7 | 27,7 | 15,277 | 0 | 21,742 | 99730000 | 8 | 1,772960502 | 0,045614679 | -1,774866104 | -3,946528587 | RM25_2015 |
| lcl\|CP010341.1_prot_AJQ91747.1_2011 | RM25_RS10100 | Molybdate ABC transporter, peripla | 0 | Unknown | SPII | 0,782 | LIPO(Sec/SPII) | P | 26,993 | 27,486 | 26,650 | 28,304 | 28,801 | 28,975 | + | 4 | 4 | 22,5 | 22,5 | 22,5 | 26,185 | 0 | 146,72 | 3629100000 | 35 | 2,195933711 | 0,043672131 | -1,650386175 | -5,233389156 | RM25_2043 |
| lcl\|CP010341.1_prot_AJQ90232.1_496 | RM25_RS02465 | Alkyl hydroperoxide reductase, F su | 0 | CytoplasmicMembrane | CYT | 0,064 | OTHER | O | 22,924 | 22,127 | 22,849 | 23,751 | 24,361 | 24,477 | + | 5 | 5 | 14,7 | 14,7 | 14,7 | 58,196 | 0 | 47,793 | 147230000 | 8 | 1,999514787 | 0,039170732 | -1,562941869 | -4,60261137 | RM25_0502 |
| lcl\|CP010341.1_prot_AJQ91033.1_1297 | RM25_RS06515 | Aminotransferase AlaT | 0 | Cytoplasmic | CYT | 0,098 | OTHER | E | 22,193 | 22,493 | 23,095 | 24,272 | 24,080 | 23,950 | + | 2 | 2 | 6,7 | 6,7 | 6,7 | 44,568 | 0 | 12,12 | 146540000 | 5 | 2,233224909 | 0,042690909 | -1,507084529 | -5,360369476 | RM25_1322 |
| lcl\|CP010341.1_prot_AJQ90363.1_627 | RM25_RS03115 | Iron compound ABC transporter, irc | 1 | Unknown | SPII | 0,916 | LIPO(Sec/SPII) | X | 28,144 | 28,424 | 27,866 | 29,170 | 29,438 | 30,284 | + | 16 | 16 | 56,2 | 56,2 | 56,2 | 35,557 | 0 | 255,25 | 7123700000 | 97 | 1,789188229 | 0,046 | -1,486309687 | -3,991217126 | RM25_0636 |
| lcl\|CP010341.1_prot_AJQ89861.1_125 | RM25_RS00630 | Putative secreted peptidoglycan-bin | 0 | Extracellular | SPI | 0,875 | SP(Sec/SPI) | X | 31,557 | 31,268 | 30,627 | 32,769 | 32,993 | 32,129 | + | 4 | 4 | 34,7 | 34,7 | 34,7 | 20,027 | 0 | 322,7 | 5,4687E+10 | 129 | 1,762508619 | 0,045357143 | -1,479375203 | -3,917923152 | RM25_0126 |
| lcl\|CP010341.1_prot_AJQ91061.1_1325 | RM25_RS06660 | Succinate dehydrogenase or fumara | 0 | Cytoplasmic | CYT | 0,171 | OTHER | C | 27,574 | 27,025 | 27,771 | 28,924 | 29,048 | 28,825 | + | 18 | 18 | 28,1 | 28,1 | 28,1 | 74,809 | 0 | 323,31 | 4302900000 | 84 | 2,502812623 | 0,041529412 | -1,475554784 | -6,355031796 | RM25_1351 |
| lcl\|CP010341.1_prot_AJQ90231.1_495 | RM25_RS02460 | Alkyl hydroperoxide reductase subu | 0 | Cytoplasmic | CYT | 0,183 | OTHER | X | 26,373 | 26,219 | 26,904 | 28,135 | 27,721 | 28,009 | + | 5 | 5 | 43,3 | 43,3 | 43,3 | 20,761 | 0 | 143,75 | 2080400000 | 27 | 2,422515157 | 0,036571429 | -1,456131617 | -6,043974828 | RM25_0501 |
| lcl\|CP010341.1_prot_AJQ90271.1_535 | RM25_RS02675 | 30S ribosomal protein S14 type Z | 0 | Cytoplasmic | CYT | 0,318 | OTHER | X | 25,793 | 25,266 | 26,249 | 26,946 | 27,207 | 27,313 | + | 1 | 1 | 21,3 | 21,3 | 21,3 | 69,383 | 0,00454 | 9,6632 | 1196700000 | 10 | 1,983381119 | 0,042285714 | -1,385424932 | -4,553484498 | RM25_0544 |
| lcl\|CP010341.1_prot_AJQ90272.1_536 | RM25_RS02680 | 30S ribosomal protein S8 | 0 | Cytoplasmic | CYT | 0,068 | OTHER | J | 26,822 | 26,498 | 27,126 | 28,058 | 28,447 | 28,066 | + | 8 | 8 | 65,2 | 65,2 | 65,2 | 14,765 | 0 | 199,73 | 3170800000 | 70 | 2,459043289 | 0,040421053 | -1,374928792 | -6,183845445 | RM25_0545 |
| lcl\|CP010341.1_prot_AJQ90262.1_526 | RM25_RS02630 | 30S ribosomal protein S19 | 0 | Cytoplasmic | CYT | 0,203 | OTHER | J | 26,560 | 26,840 | 26,725 | 28,140 | 28,011 | 28,051 | + | 6 | 6 | 47,3 | 47,3 | 47,3 | 10,612 | 0 | 70,916 | 2220100000 | 29 | 3,954654544 | 0,024 | -1,358671824 | -15,13801619 | RM25_0535 |
| lcl\|CP010341.1_prot_AJQ91799.1_2063 | RM25_RS10350 | ABC-type Fe3+-siderophores transp | 0 | Cytoplasmic | CYT | 0,104 | OTHER | P | 22,908 | 22,417 | 22,003 | 23,981 | 23,634 | 23,610 | + | 2 | 2 | 9,3 | 9,3 | 9,3 | 29,871 | 0 | 14,465 | 120640000 | 7 | 1,970928573 | 0,041409091 | -1,299201965 | -4,515830932 | RM25_2095 |
| lcl\|CP010341.1_prot_AJQ90509.1_773 | RM25_RS03855 | ATP-dependent Clp protease protec | 0 | Cytoplasmic | CYT | 0,170 | OTHER | O | 24,840 | 23,988 | 24,664 | 25,783 | 25,914 | 25,654 | + | 6 | 6 | 40,3 | 40,3 | 40,3 | 22,645 | 0 | 63,213 | 590380000 | 19 | 2,049334839 | 0,040106667 | -1,285877864 | -4,756798319 | RM25_0785 |
| lcl\|CP010341.1_prot_AJQ90276.1_540 | RM25_RS02700 | Hypothetical protein | 0 | Cytoplasmic | CYT | 0,216 | OTHER | X | 27,385 | 27,798 | 27,532 | 29,221 | 28,875 | 28,473 | + | 3 | 3 | 5,5 | 5,5 | 5,5 | 66,828 | 0 | 39,117 | 4813300000 | 50 | 2,183463314 | 0,042967742 | -1,284446081 | -5,191461988 | RM25_0549 |
| lcl\|CP010341.1_prot_AJQ90250.1_514 | RM25_RS02570 | 30S ribosomal protein S7 | 0 | Unknown | CYT | 0,092 | OTHER | J | 27,786 | 27,645 | 28,314 | 29,074 | 29,161 | 29,240 | + | 10 | 10 | 63,5 | 63,5 | 63,5 | 17,455 | 0 | 203,19 | 5139300000 | 59 | 2,396780548 | 0,042976744 | -1,243258794 | -5,947033265 | RM25_0523 |
| lcl\|CP010341.1_prot_AJQ91134.1_1398 | RM25_RS07025 | Pyridoxal biosynthesis lyase PdxS | 0 | Cytoplasmic | CYT | 0,038 | OTHER | H | 25,718 | 25,561 | 26,022 | 26,792 | 27,144 | 26,997 | + | 8 | 8 | 2,7 | 2,7 | 2,7 | 31,349 | 0 | 74,946 | 1192900000 | 28 | 2,690564161 | 0,038714286 | -1,210542043 | -7,136329072 | RM25_1424 |
| lcl\|CP010341.1_prot_AJQ91060.1_1324 | RM25_RS06655 | Succinate dehydrogenase/fumarate | 0 | Cytoplasmic | CYT | 0,107 | OTHER | C | 26,656 | 26,194 | 26,520 | 27,996 | 27,586 | 27,331 | + | 6 | 6 | 32,7 | 32,7 | 32,7 | 27,133 | 0 | 60,026 | 1793600000 | 30 | 2,118148172 | 0,042985075 | -1,180932999 | -4,976139054 | RM25_1350 |
| lcl\|CP010341.1_prot_AJQ90585.1_849 | RM25_RS04225 | 2-isopropylmalate synthase/homoc | 0 | Cytoplasmic | CYT | 0,098 | OTHER | E | 21,987 | 22,109 | 21,704 | 22,847 | 23,478 | 22,914 | + | 2 | 2 | 4,1 | 4,1 | 4,1 | 57,583 | 0 | 13,267 | 69762000 | 5 | 2,098039979 | 0,044235294 | -1,146244049 | -4,911262785 | RM25_0861 |
| lcl\|CP010341.1_prot_AJQ90275.1_539 | RM25_RS02695 | 30S ribosomal protein S5 | 0 | Cytoplasmic | CYT | 0,047 | OTHER | J | 27,241 | 26,830 | 27,412 | 28,260 | 28,366 | 28,206 | + | 5 | 5 | 23,6 | 23,6 | 23,6 | 21,586 | 0 | 48,232 | 2763500000 | 17 | 2,470935672 | 0,042666667 | -1,116245906 | -6,229966742 | RM25_0548 |
| lcl\|CP010341.1_prot_AJQ90649.1_913 | RM25_RS04550 | Putative glutathione S-transferase v | 0 | Cytoplasmic | CYT | 0,260 | OTHER | O | 26,271 | 25,989 | 26,768 | 27,585 | 27,332 | 27,330 | + | 11 | 11 | 40,9 | 40,9 | 40,9 | 39,332 | 0 | 104,26 | 1526400000 | 40 | 1,937328794 | 0,041098901 | -1,073200226 | -4,415365269 | RM25_0927 |
| lcl\|CP010341.1_prot_AJQ90590.1_854 | RM25_RS04260 | 3-isopropylmalate dehydratase sma | 0 | Cytoplasmic | CYT | 0,248 | OTHER | E | 23,826 | 24,128 | 24,101 | 25,397 | 25,019 | 24,806 | + | 4 | 4 | 22,7 | 22,7 | 22,7 | 22,153 | 0 | 25,865 | 307930000 | 17 | 2,22729172 | 0,041928571 | -1,055456161 | -5,340004147 | RM25_0868 |
| lcl\|CP010341.1_prot_AJQ91015.1_1279 | RM25_RS06420 | CTP synthase | 0 | Cytoplasmic | CYT | 0,094 | OTHER | F | 26,274 | 25,564 | 25,722 | 26,929 | 26,745 | 27,036 | + | 13 | 13 | 32,7 | 32,7 | 32,7 | 63,876 | 0 | 148,17 | 1208200000 | 48 | 1,978593578 | 0,042372093 | -1,050053279 | -4,538980987 | RM25_1301 |
| lcl\|CP010341.1_prot_AJQ90293.1_557 | RM25_RS02780 | 30S ribosomal protein S11 | 0 | Cytoplasmic | CYT | 0,273 | OTHER | J | 25,517 | 25,491 | 25,654 | 26,537 | 26,728 | 26,511 | + | 4 | 4 | 45,2 | 45,2 | 45,2 | 14,237 | 0 | 32,709 | 810590000 | 18 | 3,589834127 | 0,026857143 | -1,038036346 | -12,22400129 | RM25_0566 |
| lcl\|CP010341.1_prot_AJQ89830.1_94 | RM25_RS00465 | Putative helicase | 0 | Cytoplasmic | CYT | 0,083 | OTHER | X | 24,026 | 23,664 | 24,073 | 24,929 | 24,632 | 25,270 | + | 5 | 5 | 6,4 | 6,4 | 6,4 | 12,474 | 0 | 35,792 | 254380000 | 8 | 1,981100095 | 0,041788235 | -1,022888184 | -4,546570067 | RM25_0095 |
| lcl\|CP010341.1_prot_AJQ91475.1_1739 | RM25_RS08730 | Protein of hypothetical function DUI | 0 | Cytoplasmic | CYT | 0,096 | OTHER | S | 28,353 | 27,939 | 27,881 | 28,895 | 29,169 | 29,135 | + | 9 | 9 | 58,3 | 58,3 | 58,3 | 23,184 | 0 | 323,31 | 4614800000 | 58 | 2,377015351 | 0,044909091 | -1,008555094 | -5,873460329 | RM25_1771 |
| lcl\|CP010341.1_prot_AJQ90244.1_508 | RM25_RS02540 | 50S ribosomal protein L10 | 0 | Cytoplasmic | CYT | 0,076 | OTHER | J | 28,389 | 28,571 | 28,603 | 29,647 | 29,425 | 29,480 | + | 9 | 9 | 58,6 | 58,6 | 58,6 | 21,881 | 0 | 232,1 | 6765300000 | 57 | 3,336382656 | 0,037454545 | -0,996184667 | -10,5246925 | RM25_0517 |
| lcl\|CP010341.1_prot_AJQ90525.1_789 | RM25_RS03935 | Integral membrane protein TerC | 9 | CytoplasmicMembrane | TMH | 0,964 | OTHER | P | 21,607 | 21,576 | 21,056 | 22,287 | 22,157 | 22,766 | + | 3 | 3 | 8,6 | 8,6 | 8,6 | 46,008 | 0 | 26,792 | 55233000 | 6 | 1,736335076 | 0,044677966 | -0,989890416 | -3,846891041 | RM25_0801 |
| lcl\|CP010341.1_prot_AJQ91580.1_1844 | RM25_RS09250 | Ferrous iron transport protein A | 0 | Cytoplasmic | CYT | 0,039 | OTHER | P | 22,833 | 22,338 | 22,958 | 23,784 | 23,709 | 23,559 | + | 2 | 2 | 24,4 | 24,4 | 24,4 | 1,285 | 0 | 13,107 | 92272000 | 4 | 2,083692184 | 0,041777778 | -0,974457423 | -4,865369438 | RM25_1876 |
| lcl\|CP010341.1_prot_AJQ90277.1_541 | RM25_RS02705 | 50S ribosomal protein L15 | 0 | Cytoplasmic | CYT | 0,065 | OTHER | J | 27,632 | 27,333 | 27,656 | 28,594 | 28,576 | 28,319 | + | 5 | 5 | 41,5 | 41,5 | 41,5 | 15,464 | 0 | 45,04 | 3344000000 | 21 | 2,659526669 | 0,03737931 | -0,956205368 | -7,001690161 | RM25_0550 |
| lcl\|CP010341.1_prot_AJQ91204.1_1468 | RM25_RS11905 | Putative regulatory protein | 0 | Unknown | CYT | 0,761 | OTHER | J | 28,431 | 28,439 | 28,674 | 29,540 | 29,811 | 28,979 | + | 15 | 15 | 50,9 | 50,9 | 50,9 | 35,943 | 0 | 270,28 | 5968200000 | 74 | 1,644827077 | 0,050289855 | -0,928223292 | -3,605081577 | RM25_1495 |
| lcl\|CP010341.1_prot_AJQ91579.1_1843 | RM25_RS09245 | Ferrous iron transport protein B | 9 | CytoplasmicMembrane | CYT | 0,917 | OTHER | X | 22,097 | 22,497 | 22,011 | 22,940 | 22,999 | 23,443 | + | 3 | 3 | 0,6 | 0,6 | 0,6 | 69,798 | 0 | 28,798 | 68873000 | 5 | 1,87758565 | 0,045583333 | -0,925839742 | -4,240707447 | RM25_1875 |
| lcl\|CP010341.1_prot_AJQ90274.1_538 | RM25_RS02690 | 50S ribosomal protein L18 | 0 | Cytoplasmic | CYT | 0,303 | OTHER | J | 27,856 | 27,814 | 28,241 | 28,995 | 28,800 | 28,891 | + | 8 | 8 | 55,9 | 55,9 | 55,9 | 14,023 | 0 | 95,318 | 5347400000 | 52 | 2,484190318 | 0,043885714 | -0,924957911 | -6,28171365 | RM25_0547 |
| lcl\|CP010341.1_prot_AJQ91123.1_1387 | RM25_RS06970 | SUF system FeS assembly protein, N | 0 | Cytoplasmic | CYT | 0,094 | OTHER | C | 24,498 | 24,787 | 24,519 | 25,327 | 25,745 | 25,449 | + | 3 | 3 | 3,2 | 3,2 | 3,2 | 1,664 | 0 | 29,46 | 385630000 | 6 | 2,367458192 | 0,042956522 | -0,905517578 | -5,838156879 | RM25_1413 |
| lcl\|CP010341.1_prot_AJQ90259.1_523 | RM25_RS02615 | 50S ribosomal protein L4 | 0 | Cytoplasmic | CYT | 0,311 | OTHER | J | 27,933 | 27,920 | 28,571 | 29,185 | 29,040 | 28,915 | + | 9 | 9 | 44,6 | 44,6 | 44,6 | 24,839 | 0 | 255,39 | 5405400000 | 78 | 1,777701593 | 0,045570093 | -0,905157725 | -3,959550011 | RM25_0532 |
| lcl\|CP010341.1_prot_AJQ90618.1_882 | RM25_RS04400 | Glycerol-3-phosphate dehydrogena: | 0 | Cytoplasmic | CYT | 0,101 | OTHER | X | 26,685 | 26,067 | 26,622 | 27,336 | 27,403 | 27,273 | + | 12 | 12 | 31,9 | 31,9 | 31,9 | 5,883 | 0 | 258,58 | 1661400000 | 38 | 1,931324252 | 0,042 | -0,879077276 | -4,397583122 | RM25_0896 |
| lcl\|CP010341.1_prot_AJQ90257.1_521 | RM25_RS02605 | 30S ribosomal protein S10 | 0 | Cytoplasmic | CYT | 0,068 | OTHER | J | 28,763 | 28,561 | 28,857 | 29,585 | 29,695 | 29,522 | + | 6 | 6 | 57,3 | 57,3 | 57,3 | 11,744 | 0 | 49,602 | 7288700000 | 43 | 3,00911771 | 0,0308 | -0,873648961 | -8,657630465 | RM25_0530 |
| lcl\|CP010341.1_prot_AJQ90248.1_512 | RM25_RS02560 | DNA-directed RNA polymerase, beta | 0 | Cytoplasmic | CYT | 0,078 | OTHER | K | 28,358 | 28,049 | 28,312 | 28,885 | 29,028 | 29,304 | + | 39 | 39 | 32,3 | 32,3 | 32,3 | 14,361 | 0 | 323,31 | 5253800000 | 165 | 2,225496185 | 0,041192982 | -0,83224233 | -5,333853256 | RM25_0521 |
| lcl\|CP010341.1_prot_AJQ91210.1_1474 | RM25_RS07410 | Response regulator | 0 | Unknown | CYT | 0,254 | OTHER | T | 25,823 | 25,710 | 25,572 | 26,542 | 26,486 | 26,535 | + | 7 | 7 | 47,7 | 47,7 | 47,7 | 1,878 | 0 | 69,687 | 905680000 | 35 | 3,404198233 | 0,0284 | -0,818848928 | -10,95598508 | RM25_1501 |
| lcl\|CP010341.1_prot_AJQ90334.1_598 | RM25_RS02990 | 60 kDa chaperonin 1 | 0 | Cytoplasmic | CYT | 0,044 | OTHER | O | 31,342 | 31,046 | 31,104 | 31,888 | 32,023 | 32,015 | + | 26 | 26 | 7,4 | 7,4 | 7,4 | 56,146 | 0 | 323,31 | 3,7612E+10 | 234 | 2,889449383 | 0,028615385 | -0,811395645 | -8,054914602 | RM25_0607 |
| lcl\|CP010341.1_prot_AJQ90240.1_504 | RM25_RS02520 | 50S ribosomal protein L11 | 0 | Unknown | SPI | 0,140 | SP(Sec/SPI) | J | 28,320 | 28,174 | 28,484 | 29,195 | 29,094 | 29,072 | + | 5 | 5 | 3,8 | 3,8 | 3,8 | 14,871 | 0 | 86,225 | 5530400000 | 40 | 2,914765989 | 0,025666667 | -0,79434967 | -8,179145631 | RM25_0513 |
| lcl\|CP010341.1_prot_AJQ90810.1_1074 | RM25_RS05390 | AcuB family protein | 0 | Unknown | CYT | 0,045 | OTHER | X | 23,062 | 23,431 | 23,203 | 24,172 | 23,786 | 24,014 | + | 2 | 2 | 1,2 | 1,2 | 1,2 | 23,411 | 0 | 15,084 | 148230000 | 9 | 2,089926979 | 0,042971429 | -0,758556366 | -4,885271826 | RM25_1093 |
| lcl\|CP010341.1_prot_AJQ90273.1_537 | RM25_RS02685 | 50S ribosomal protein L6 | 0 | Cytoplasmic | CYT | 0,127 | OTHER | J | 27,882 | 28,176 | 28,173 | 28,939 | 28,865 | 28,685 | + | 7 | 7 | 45,6 | 45,6 | 45,6 | 19,743 | 0 | 74,195 | 4570800000 | 56 | 2,438947245 | 0,037463415 | -0,752593358 | -6,106562687 | RM25_0546 |
| lcl\|CP010341.1_prot_AJQ90448.1_712 | RM25_RS03525 | Naphthoate synthase | 0 | Unknown | CYT | 0,215 | OTHER | H | 22,954 | 23,333 | 23,300 | 23,741 | 24,138 | 23,957 | + | 3 | 3 | 11,1 | 11,1 | 11,1 | 39,348 | 0 | 27,53 | 148900000 | 11 | 1,961653623 | 0,041555556 | -0,74978892 | -4,487933969 | RM25_0721 |
| lcl\|CP010341.1_prot_AJQ91461.1_1725 | RM25_RS08655 | 60 kDa chaperonin 1 | 0 | Cytoplasmic | CYT | 0,047 | OTHER | O | 31,282 | 31,094 | 31,356 | 31,966 | 31,956 | 32,059 | + | 25 | 25 | 72,2 | 72,2 | 72,2 | 56,538 | 0 | 323,31 | 4,1936E+10 | 303 | 3,047733156 | 0,032421053 | -0,749621073 | -8,860734573 | RM25_1756 |
| lcl\|CP010341.1_prot_AJQ91951.1_2215 | RM25_RS11090 | 30S ribosomal protein S6 | 0 | Cytoplasmic | CYT | 0,357 | OTHER | J | 27,106 | 26,926 | 27,471 | 28,030 | 28,019 | 27,691 | + | 5 | 5 | 48,4 | 48,4 | 48,4 | 10,886 | 0 | 77,522 | 2829200000 | 36 | 1,726474296 | 0,044363636 | -0,745400111 | -3,82035022 | RM25_2247 |
| lcl\|CP010341.1_prot_AJQ90268.1_532 | RM25_RS02660 | 50S ribosomal protein L14 | 0 | Cytoplasmic | CYT | 0,040 | OTHER | J | 26,632 | 26,509 | 26,619 | 27,362 | 27,322 | 27,233 | + | 3 | 3 | 26,8 | 26,8 | 26,8 | 13,344 | 0 | 36,727 | 1549600000 | 25 | 3,718782453 | 0,0192 | -0,719022751 | -13,18607926 | RM25_0541 |
| lcl\|CP010341.1_prot_AJQ90619.1_883 | RM25_RS04405 | Anaerobic glycerol-3-phosphate deh | 0 | CytoplasmicMembrane | CYT | 0,066 | OTHER | X | 27,802 | 27,349 | 27,812 | 28,153 | 28,484 | 28,468 | + | 13 | 13 | 39,2 | 39,2 | 39,2 | 45,912 | 0 | 169,44 | 3150900000 | 79 | 1,727082239 | 0,045109244 | -0,71396637 | -3,821983077 | RM25_0897 |
| lcl\|CP010341.1_prot_AJQ91537.1_1801 | RM25_RS09035 | ATP-dependent chaperone protein ( | 0 | Cytoplasmic | CYT | 0,047 | OTHER | O | 27,424 | 27,105 | 27,232 | 28,048 | 27,843 | 27,986 | + | 27 | 27 | 41,7 | 3,8 | 3,8 | 94,419 | 0 | 323,31 | 2667000000 | 105 | 2,503009926 | 0,042787879 | -0,705193202 | -6,355812477 | RM25_1833 |
| lcl\|CP010341.1_prot_AJQ90510.1_774 | RM25_RS03860 | ATP-dependent Clp protease protec | 0 | Cytoplasmic | CYT | 0,196 | OTHER | O | 25,419 | 24,882 | 24,973 | 25,844 | 25,848 | 25,676 | + | 4 | 4 | 27,5 | 27,5 | 27,5 | 24,806 | 0 | 58,87 | 530780000 | 19 | 1,783824443 | 0,045561905 | -0,697654724 | -3,976408857 | RM25_0786 |
| lcl\|CP010341.1_prot_AJQ90247.1_511 | RM25_RS02555 | DNA-directed RNA polymerase, beta | 0 | Cytoplasmic | CYT | 0,136 | OTHER | K | 28,444 | 28,015 | 28,227 | 28,803 | 28,843 | 29,116 | + | 34 | 34 | 38,9 | 38,9 | 38,9 | 12,872 | 0 | 323,31 | 4789200000 | 142 | 1,92553677 | 0,042255319 | -0,692023595 | -4,380492567 | RM25_0520 |
| lcl\|CP010341.1_prot_AJQ90292.1_556 | RM25_RS02775 | 30S ribosomal protein S13 | 0 | Cytoplasmic | CYT | 0,091 | OTHER | J | 28,250 | 28,296 | 28,424 | 29,228 | 28,946 | 28,866 | + | 6 | 6 | 46,8 | 46,8 | 46,8 | 14,026 | 0 | 124,71 | 5208400000 | 45 | 2,326844686 | 0,038745098 | -0,690237045 | -5,69007892 | RM25_0565 |
| lcl\|CP010341.1_prot_AJQ90261.1_525 | RM25_RS02625 | 50S ribosomal protein L2 | 0 | Cytoplasmic | CYT | 0,824 | OTHER | J | 28,660 | 28,737 | 29,120 | 29,325 | 29,666 | 29,529 | + | 12 | 12 | 53,2 | 53,2 | 53,2 | 30,369 | 0 | 186,99 | 7555400000 | 103 | 1,739961499 | 0,045843478 | -0,668240865 | -3,856681892 | RM25_0534 |
| lcl\|CP010341.1_prot_AJQ90210.1_474 | RM25_RS02350 | NADH-quinone oxidoreductase subi | 0 | Cytoplasmic | CYT | 0,138 | OTHER | C | 25,522 | 25,447 | 25,547 | 25,983 | 26,049 | 26,484 | + | 7 | 7 | 2,1 | 2,1 | 2,1 | 50,313 | 0 | 79,511 | 727010000 | 26 | 1,851589701 | 0,047393939 | -0,667116165 | -4,166253478 | RM25_0479 |
| lcl\|CP010341.1_prot_AJQ91067.1_1331 | RM25_RS06690 | 30S ribosomal protein S16 | 0 | Cytoplasmic | CYT | 0,879 | OTHER | J | 28,381 | 28,464 | 28,607 | 29,228 | 28,992 | 29,224 | + | 6 | 6 | 4,6 | 4,6 | 4,6 | 17,753 | 0 | 229,49 | 5872900000 | 52 | 2,53879963 | 0,041548387 | -0,663860957 | -6,498784167 | RM25_1357 |
| lcl\|CP010341.1_prot_AJQ90849.1_1113 | RM25_RS05585 | ATP synthase subunit b | 1 | CytoplasmicMembrane | CYT | 0,697 | OTHER | C | 26,119 | 25,585 | 26,057 | 26,635 | 26,530 | 26,573 | + | 4 | 4 | 32,6 | 32,6 | 32,6 | 20,192 | 0 | 72,239 | 919820000 | 31 | 1,736856728 | 0,045059829 | -0,65877978 | -3,84829843 | RM25_1132 |
| lcl\|CP010341.1_prot_AJQ90266.1_530 | RM25_RS02650 | 50S ribosomal protein L29 | 0 | Cytoplasmic | CYT | 0,051 | OTHER | J | 27,687 | 27,870 | 27,758 | 28,417 | 28,598 | 28,252 | + | 4 | 4 | 48,8 | 48,8 | 48,8 | 93,074 | 0 | 96,847 | 3063100000 | 28 | 2,343036183 | 0,040326531 | -0,65052859 | -5,748738797 | RM25_0539 |
| lcl\|CP010341.1_prot_AJQ90295.1_559 | RM25_RS02790 | DNA-directed RNA polymerase, alpł | 0 | Cytoplasmic | CYT | 0,076 | OTHER | K | 28,245 | 28,077 | 28,280 | 29,014 | 28,530 | 28,990 | + | 13 | 13 | 47,2 | 47,2 | 47,2 | 36,975 | 0 | 323,31 | 4527400000 | 63 | 1,719737286 | 0,04516129 | -0,643993378 | -3,802285731 | RM25_0568 |
| lcl\|CP010341.1_prot_AJQ90173.1_437 | RM25_RS02150 | UPF0237 protein | 0 | Cytoplasmic | CYT | 0,023 | OTHER | X | 23,860 | 23,630 | 24,090 | 24,684 | 24,493 | 24,321 | + | 3 | 3 | 38,2 | 38,2 | 38,2 | 98,172 | 0 | 23,188 | 201890000 | 12 | 1,711633935 | 0,045536 | -0,639289856 | -3,78063079 | RM25_0441 |
| lcl\|CP010341.1_prot_AJQ90258.1_522 | RM25_RS02610 | 50S ribosomal protein L3 | 0 | Cytoplasmic | CYT | 0,345 | OTHER | J | 27,935 | 27,977 | 27,938 | 28,773 | 28,360 | 28,628 | + | 6 | 6 | 38,8 | 38,8 | 38,8 | 22,967 | 0 | 323,31 | 4151100000 | 45 | 2,197421452 | 0,0444 | -0,6373113 | -5,238408931 | RM25_0531 |
| lcl\|CP010341.1_prot_AJQ90528.1_792 | RM25_RS03950 | Hypothetical protein | 0 | Cytoplasmic | CYT | 0,457 | OTHER | J | 27,518 | 27,515 | 27,746 | 28,241 | 28,199 | 28,217 | + | 20 | 20 | 34,3 | 34,3 | 34,3 | 10,118 | 0 | 323,31 | 3035200000 | 83 | 2,89440206 | 0,02976 | -0,62583669 | -8,079082677 | RM25_0804 |
| lcl\|CP010341.1_prot_AJQ90312.1_576 | RM25_RS02875 | 30S ribosomal protein S9 | 0 | Cytoplasmic | CYT | 0,819 | OTHER | J | 28,909 | 28,962 | 28,925 | 29,837 | 29,315 | 29,470 | + | 8 | 8 | 6,5 | 6,5 | 6,5 | 19,467 | 0 | 166,76 | 7181900000 | 50 | 1,759365105 | 0,044955752 | -0,608418147 | -3,90934674 | RM25_0585 |
| lcl\|CP010341.1_prot_AJQ91476.1_1740 | RM25_RS08735 | Iron-sulfur cluster-binding protein | 0 | Cytoplasmic | CYT | 0,109 | OTHER | C | 28,823 | 28,673 | 28,548 | 29,319 | 29,037 | 29,510 | + | 15 | 15 | 33,4 | 33,4 | 33,4 | 57,312 | 0 | 202,66 | 6102800000 | 95 | 1,726969721 | 0,044733333 | -0,607461294 | -3,821680833 | RM25_1772 |
| lcl\|CP010341.1_prot_AJQ91949.1_2213 | RM25_RS11080 | Hypothetical protein | 0 | Cytoplasmic | CYT | 0,624 | OTHER | J | 26,235 | 25,969 | 26,320 | 26,654 | 26,864 | 26,817 | + | 2 | 2 | 26,6 | 26,6 | 26,6 | 89,764 | 0 | 62,431 | 1065200000 | 20 | 2,092113359 | 0,043594203 | -0,603222529 | -4,892265773 | RM25_2245 |
| lcl\|CP010341.1_prot_AJQ90263.1_527 | RM25_RS02635 | 50S ribosomal protein L22 | 0 | Cytoplasmic | CYT | 0,091 | OTHER | J | 28,045 | 27,985 | 28,377 | 28,706 | 28,767 | 28,696 | + | 5 | 5 | 34,8 | 34,8 | 34,8 | 17,268 | 0 | 84,978 | 4350000000 | 37 | 2,047310359 | 0,039578947 | -0,587465286 | -4,75045851 | RM25_0536 |
| lcl\|CP010341.1_prot_AJQ89803.1_67 | RM25_RS00340 | Bifunctional UDP-galactofuranosyl t | 0 | Cytoplasmic | CYT | 0,349 | OTHER | S | 26,246 | 25,874 | 26,029 | 26,480 | 26,662 | 26,736 | + | 9 | 9 | 21,3 | 21,3 | 21,3 | 75,349 | 0 | 110,49 | 981220000 | 34 | 1,919467475 | 0,041810526 | -0,576452255 | -4,362621032 | RM25_0068 |
| lcl\|CP010341.1_prot_AJQ90294.1_558 | RM25_RS02785 | 30S ribosomal protein S4 | 0 | Cytoplasmic | CYT | 0,104 | OTHER | J | 28,885 | 28,779 | 29,135 | 29,512 | 29,349 | 29,652 | + | 12 | 12 | 49,3 | 49,3 | 49,3 | 23,303 | 0 | 138,37 | 6877400000 | 58 | 1,850090274 | 0,04692 | -0,571011225 | -4,16198699 | RM25_0567 |
| lcl\|CP010341.1_prot_AJQ90241.1_505 | RM25_RS02525 | 50S ribosomal protein L1 | 0 | Cytoplasmic | CYT | 0,155 | OTHER | J | 28,908 | 28,803 | 28,899 | 29,536 | 29,418 | 29,333 | + | 11 | 11 | 64,4 | 64,4 | 64,4 | 25,411 | 0 | 323,31 | 7884600000 | 109 | 2,926130587 | 0,026782609 | -0,558855057 | -8,235475183 | RM25_0514 |
| lcl\|CP010341.1_prot_AJQ91125.1_1389 | RM25_RS06980 | FeS assembly ATPase SufC | 0 | Cytoplasmic | CYT | 0,079 | OTHER | O | 27,754 | 27,583 | 27,953 | 28,335 | 28,477 | 28,124 | + | 8 | 8 | 45,8 | 45,8 | 45,8 | 26,923 | 0 | 239,64 | 3391400000 | 56 | 1,682066816 | 0,047877863 | -0,549134572 | -3,702287533 | RM25_1415 |
| lcl\|CP010341.1_prot_AJQ90859.1_1123 | RM25_RS05635 | Transcription termination factor Rh | 0 | Cytoplasmic | CYT | 0,154 | OTHER | K | 26,039 | 26,096 | 25,908 | 26,388 | 26,731 | 26,533 | + | 14 | 14 | 22,4 | 22,4 | 22,4 | 72,445 | 0 | 90,664 | 933800000 | 33 | 2,034952179 | 0,039948718 | -0,536281586 | -4,711895561 | RM25_1142 |
| lcl\|CP010341.1_prot_AJQ91238.1_1502 | RM25_RS07545 | MgtE intracellular region | 0 | Cytoplasmic | CYT | 0,074 | OTHER | P | 22,149 | 21,917 | 22,054 | 22,444 | 22,554 | 22,730 | + | 2 | 2 | 6,4 | 6,4 | 6,4 | 46,935 | 0 | 12,474 | 68960000 | 4 | 2,127587088 | 0,045 | -0,536040624 | -5,006819067 | RM25_1529 |
| lcl\|CP010341.1_prot_AJQ90091.1_355 | RM25_RS12205 | Hypothetical protein | 0 | Cytoplasmic | CYT | 0,858 | OTHER | L | 24,314 | 24,422 | 24,328 | 24,695 | 25,073 | 24,855 | + | 9 | 9 | 12,3 | 12,3 | 12,3 | 11,686 | 0 | 98,352 | 291220000 | 25 | 1,97438487 | 0,041885057 | -0,519639969 | -4,526259009 | RM25_0359 |
| lcl\|CP010341.1_prot_AJQ91089.1_1353 | RM25_RS06795 | 30S ribosomal protein S15 | 0 | Cytoplasmic | CYT | 0,468 | OTHER | J | 27,340 | 27,559 | 27,409 | 27,829 | 28,151 | 27,841 | + | 5 | 5 | 64,4 | 64,4 | 64,4 | 10,163 | 0 | 87,408 | 2347800000 | 25 | 1,823689517 | 0,046455446 | -0,504184723 | -4,087358912 | RM25_1379 |
| lcl\|CP010341.1_prot_AJQ91066.1_1330 | RM25_RS06685 | UPF0109 protein | 0 | Cytoplasmic | CYT | 0,047 | OTHER | S | 24,096 | 24,299 | 24,278 | 24,855 | 24,566 | 24,718 | + | 3 | 3 | 52,4 | 52,4 | 52,4 | 95,538 | 0 | 41,038 | 281140000 | 14 | 2,0113938 | 0,039654321 | -0,488576889 | -4,639032315 | RM25_1356 |
| lcl\|CP010341.1_prot_AJQ90742.1_1006 | RM25_RS05045 | Protein translocase subunit SecF | 6 | CytoplasmicMembrane | CYT | 0,920 | OTHER | U | 24,987 | 25,007 | 25,010 | 25,438 | 25,287 | 25,606 | + | 3 | 3 | 15,7 | 15,7 | 15,7 | 44,402 | 0 | 72,115 | 422300000 | 12 | 2,060677484 | 0,040648649 | -0,442232768 | -4,792436706 | RM25_1025 |
| lcl\|CP010341.1_prot_AJQ90265.1_529 | RM25_RS02645 | 50S ribosomal protein L16 | 0 | Cytoplasmic | CYT | 0,080 | OTHER | J | 27,310 | 27,186 | 27,589 | 27,793 | 27,799 | 27,818 | + | 4 | 4 | 46,8 | 46,8 | 46,8 | 15,732 | 0 | 61,346 | 2308000000 | 29 | 1,678820471 | 0,048454545 | -0,441445669 | -3,693749177 | RM25_0538 |
| lcl\|CP010341.1_prot_AJQ90319.1_583 | RM25_RS13145 | Hypothetical protein | 0 | Cytoplasmic | CYT | 0,790 | OTHER | L | 24,400 | 24,523 | 24,577 | 24,895 | 24,731 | 25,052 | + | 4 | 4 | 0,5 | 0,5 | 0,5 | 11,691 | 0 | 39,17 | 302670000 | 13 | 1,677433801 | 0,048656716 | -0,392719269 | -3,690105825 | RM25_0592 |
| lcl\|CP010341.1_prot_AJQ90270.1_534 | RM25_RS02670 | 50S ribosomal protein L5 | 0 | Cytoplasmic | CYT | 0,101 | OTHER | J | 28,007 | 27,951 | 27,983 | 28,476 | 28,248 | 28,342 | + | 11 | 11 | 52,7 | 52,7 | 52,7 | 24,709 | 0 | 207,14 | 4039800000 | 77 | 2,275045005 | 0,038888889 | -0,374914805 | -5,505686478 | RM25_0543 |
| lcl\|CP010341.1_prot_AJQ91084.1_1348 | RM25_RS06770 | Metallo-beta-lactamase family prot | 0 | Cytoplasmic | CYT | 0,047 | OTHER | S | 25,594 | 25,602 | 25,713 | 25,985 | 26,102 | 25,930 | + | 7 | 7 | 14,9 | 14,9 | 14,9 | 63,381 | 0 | 61,463 | 668940000 | 17 | 2,361200514 | 0,042042553 | -0,3695666 | -5,815136655 | RM25_1374 |
| lcl\|CP010341.1_prot_AJQ90834.1_1098 | RM25_RS05510 | Thioredoxin TrxA | 0 | Unknown | CYT | 0,100 | OTHER | O | 26,152 | 25,973 | 26,217 | 26,554 | 26,525 | 26,369 | + | 7 | 7 | 3,3 | 3,3 | 3,3 | 33,905 | 0 | 84,382 | 1010200000 | 39 | 1,782465788 | 0,045132075 | -0,368717829 | -3,972663758 | RM25_1117 |
| lcl\|CP010341.1_prot_AJQ90209.1_473 | RM25_RS02345 | NADH-quinone oxidoreductase subi | 0 | Cytoplasmic | CYT | 0,144 | OTHER | C | 27,245 | 27,064 | 27,272 | 27,416 | 27,614 | 27,563 | + | 10 | 10 | 52,2 | 52,2 | 52,2 | 28,677 | 0 | 146,21 | 1986100000 | 52 | 1,723122854 | 0,044520325 | -0,337105433 | -3,811356797 | RM25_0478 |
| lcl\|CP010341.1_prot_AJQ89751.1_15 | RM25_RS00075 | DNA gyrase, A subunit | 0 | Cytoplasmic | CYT | 0,111 | OTHER | L | 24,909 | 24,802 | 24,901 | 25,039 | 25,261 | 25,303 | + | 7 | 7 | 10,5 | 10,5 | 10,5 | 10,064 | 0 | 50,533 | 392330000 | 19 | 1,691156618 | 0,0463125 | -0,330416361 | -3,726261399 | RM25_0015 |
| lcl\|CP010341.1_prot_AJQ89855.1_119 | RM25_RS12905 | Hypothetical protein | 0 | Cytoplasmic | SPI | 0,065 | OTHER | H | 23,425 | 23,521 | 23,543 | 23,635 | 23,712 | 23,756 | + | 2 | 2 | 3,7 | 3,7 | 3,7 | 53,443 | 0 | 13,324 | 134040000 | 7 | 1,809937811 | 0,046901961 | -0,204601288 | -4,048852113 | RM25_0120 |

**Abundance increased under Anaerobic (N+) condition**

| | | | | | | | | | Log2 Transfomed LFQ intensities | | | | | | | | | | | | | | | | | | | | | |
|---|---|---|---|---|---|---|---|---|---|---|---|---|---|---|---|---|---|---|---|---|---|---|---|---|---|---|---|---|---|---|
| Protein IDs | NCBI-LocusTag | Description | No. of TMHMM | PsortB | LipoP | SecP Score | SignalP | COG | Anaerobic 1 | Anaerobic 2 | Anaerobic 3 | Aerobic 1 | Aerobic 3 | Aerobic 4 | Student's T-test Significant Anaerobic_Aerobic | Razor + unique peptides | Unique peptides | Sequence coverage [%] | Unique + razor sequence coverage [%] | Unique sequence coverage [%] | Mol. weight [kDa] | Q-value | Score | Intensity | MS/MS count | -Log Student's T-test p-value Anaerobic_Aerobic | Student's T-test q-value Anaerobic_Aerobic | Student's T-test Difference Anaerobic_Aerobic | Student's T-test Test statistic Anaerobic_Aerobic | LocusTag |
| lcl\|CP010341.1_prot_AJQ90153.1_417 | RM25_RS02055 | Multifunctional siroheme synthase | 0 | Unknown | CYT | 0,055 | OTHER | H | 25,879 | 25,904 | 25,880 | 25,722 | 25,626 | 25,598 | + | 6 | 6 | 42,8 | 42,8 | 42,8 | 27,095 | 0 | 58,271 | 633970000 | 26 | 2,463437689 | 0,041513514 | 0,238755544 | 6,200854199 | RM25_0421 |
| lcl\|CP010341.1_prot_AJQ89917.1_181 | RM25_RS00915 | TRNA uridine 5-carboxymethylamin | 0 | Cytoplasmic | CYT | 0,185 | OTHER | C | 28,721 | 28,636 | 28,638 | 28,424 | 28,239 | 28,429 | + | 22 | 22 | 45,3 | 45,3 | 45,3 | 59,855 | 0 | 213,31 | 4368100000 | 114 | 1,930621299 | 0,041548387 | 0,301124573 | 4,395504736 | RM25_0185 |
| lcl\|CP010341.1_prot_AJQ90252.1_516 | RM25_RS02580 | Translation elongation factor Tu | 0 | Cytoplasmic | CYT | 0,102 | OTHER | J | 31,302 | 31,144 | 31,302 | 30,833 | 31,011 | 30,958 | + | 15 | 15 | 55,8 | 55,8 | 55,8 | 43,679 | 0 | 323,31 | 2,9737E+10 | 227 | 1,868567934 | 0,045113402 | 0,315003077 | 4,21477584 | RM25_0525 |
| lcl\|CP010341.1_prot_AJQ91500.1_1764 | RM25_RS08855 | Glutamate-1-semialdehyde 2,1-ami | 0 | Cytoplasmic | CYT | 0,043 | OTHER | H | 26,769 | 26,777 | 26,829 | 26,348 | 26,544 | 26,453 | + | 9 | 9 | 27,7 | 27,7 | 27,7 | 45,932 | 0 | 86,665 | 1135300000 | 39 | 2,345229072 | 0,041166667 | 0,343056361 | 5,7567214 | RM25_1796 |
| lcl\|CP010341.1_prot_AJQ89802.1_66 | RM25_RS00335 | UDP-galactopyranose mutase | 0 | Cytoplasmic | CYT | 0,245 | OTHER | M | 25,627 | 25,692 | 25,947 | 25,255 | 25,315 | 25,450 | + | 9 | 9 | 28,1 | 28,1 | 28,1 | 45,347 | 0 | 223,77 | 536070000 | 31 | 1,666334658 | 0,050488889 | 0,415435155 | 3,661025176 | RM25_0067 |
| lcl\|CP010341.1_prot_AJQ89967.1_231 | RM25_RS01155 | Arginine--tRNA ligase | 0 | Cytoplasmic | CYT | 0,089 | OTHER | J | 25,224 | 25,481 | 25,592 | 24,986 | 24,884 | 25,088 | + | 7 | 7 | 21,3 | 21,3 | 21,3 | 5,838 | 0 | 69,123 | 405990000 | 16 | 1,645924662 | 0,050656934 | 0,446309408 | 3,607923597 | RM25_0235 |
| lcl\|CP010341.1_prot_AJQ91142.1_1406 | RM25_RS07065 | Phosphoglycerate kinase | 0 | Cytoplasmic | CYT | 0,046 | OTHER | G | 29,470 | 29,513 | 29,447 | 28,944 | 28,968 | 28,867 | + | 19 | 19 | 62,3 | 62,3 | 62,3 | 42,436 | 0 | 323,31 | 6375000000 | 98 | 3,976240375 | 0,032 | 0,55076472 | 15,33002179 | RM25_1432 |
| lcl\|CP010341.1_prot_AJQ92010.1_2274 | RM25_RS11385 | Cytochrome P450 | 0 | CytoplasmicMembrane | CYT | 0,119 | OTHER | X | 25,903 | 25,937 | 25,788 | 25,188 | 25,529 | 25,254 | + | 8 | 8 | 27,7 | 27,7 | 27,7 | 44,283 | 0 | 94,449 | 558210000 | 26 | 2,07714005 | 0,041205479 | 0,551988602 | 4,844520716 | RM25_2307 |
| lcl\|CP010341.1_prot_AJQ90375.1_639 | RM25_RS03175 | Malate dehydrogenase | 0 | Cytoplasmic | CYT | 0,115 | OTHER | C | 29,994 | 29,773 | 30,181 | 29,476 | 29,281 | 29,365 | + | 8 | 8 | 34,5 | 34,5 | 34,5 | 34,909 | 0 | 166,35 | 9365000000 | 61 | 2,017974205 | 0,040658228 | 0,608963013 | 4,65929958 | RM25_0648 |
| lcl\|CP010341.1_prot_AJQ89881.1_145 | RM25_RS00720 | PTS system mannose-specific EIIBC/ | 8 | CytoplasmicMembrane | CYT | 0,829 | OTHER | X | 22,771 | 22,255 | 22,438 | 22,010 | 21,869 | 21,748 | + | 3 | 3 | 3,5 | 3,5 | 3,5 | 10,199 | 0 | 23,255 | 49508000 | 8 | 1,652629953 | 0,050117647 | 0,612315496 | 3,625315932 | RM25_0146 |

| | | | | | | | | | | | | | | | | | | | | | | | | | | | | | | |
|---|---|---|---|---|---|---|---|---|---|---|---|---|---|---|---|---|---|---|---|---|---|---|---|---|---|---|---|---|---|---|
| lcl\|CP010341.1_prot_AJQ90718.1_982 | RM25_RS04930 | Indole-3-glycerol phosphate syntha | 0 | Cytoplasmic | CYT | 0,043 | OTHER | E | 25,013 | 25,152 | 25,085 | 24,527 | 24,367 | 24,319 | + | 6 | 6 | 28,3 | 28,3 | 28,3 | 30,352 | 0 | 49,862 | 302600000 | 10 | 3,095130657 | 0,02575 | 0,678885778 | 9,116017874 | RM25_1001 |
| lcl\|CP010341.1_prot_AJQ90179.1_443 | RM25_RS02195 | Cell-wall peptidases, NlpC/P60 fami | 0 | Extracellular | SPI | 0,884 | SP(Sec/SPI) | M | 33,428 | 33,504 | 33,067 | 32,546 | 32,704 | 32,622 | + | 22 | 22 | 61,2 | 61,2 | 61,2 | 58,794 | 0 | 323,31 | 9,055E+10 | 250 | 2,121819656 | 0,043636364 | 0,70923233 | 4,988055449 | RM25_0447 |
| lcl\|CP010341.1_prot_AJQ89750.1_14 | RM25_RS00070 | DNA gyrase, B subunit | 0 | Cytoplasmic | CYT | 0,100 | OTHER | L | 25,435 | 25,580 | 25,640 | 24,610 | 24,752 | 25,112 | + | 12 | 12 | 19,6 | 19,6 | 19,6 | 75,053 | 0 | 111,6 | 443410000 | 18 | 1,966108511 | 0,042022472 | 0,726942698 | 4,501317534 | RM25_0014 |
| lcl\|CP010341.1_prot_AJQ90752.1_1016 | RM25_RS05095 | Lactate/malate dehydrogenase, NA | 0 | Cytoplasmic | CYT | 0,069 | OTHER | X | 23,677 | 23,261 | 23,304 | 22,625 | 22,770 | 22,606 | + | 2 | 2 | 9,8 | 9,8 | 9,8 | 34,973 | 0 | 22,15 | 76968000 | 4 | 2,20601771 | 0,045931034 | 0,746759415 | 5,26748812 | RM25_1035 |
| lcl\|CP010341.1_prot_AJQ90095.1_359 | RM25_RS01760 | Isoleucine--tRNA ligase | 0 | Cytoplasmic | CYT | 0,203 | OTHER | J | 23,041 | 23,012 | 23,044 | 22,634 | 21,978 | 22,124 | + | 2 | 2 | 2,1 | 2,1 | 2,1 | 12,202 | 0 | 15,605 | 50034000 | 4 | 1,774287672 | 0,046037037 | 0,786958694 | 3,950170781 | RM25_0363 |
| lcl\|CP010341.1_prot_AJQ90829.1_1093 | RM25_RS05485 | Maltose alpha-D-glucosyltransferas | 0 | Cytoplasmic | CYT | 0,300 | OTHER | G | 23,637 | 23,534 | 23,218 | 22,961 | 22,604 | 22,417 | + | 2 | 2 | 6,3 | 6,3 | 6,3 | 68,428 | 0 | 18,108 | 50015000 | 5 | 1,772351805 | 0,044792793 | 0,802618663 | 3,944858868 | RM25_1112 |
| lcl\|CP010341.1_prot_AJQ91620.1_1884 | RM25_RS09450 | Aspartate-semialdehyde dehydroge | 0 | Cytoplasmic | CYT | 0,085 | OTHER | E | 25,289 | 25,032 | 25,555 | 24,502 | 24,340 | 24,561 | + | 5 | 5 | 19,8 | 19,8 | 19,8 | 36,567 | 0 | 38,489 | 329270000 | 13 | 2,127444916 | 0,044307692 | 0,824794134 | 5,006355871 | RM25_1916 |
| lcl\|CP010341.1_prot_AJQ89885.1_149 | RM25_RS00740 | Putative periplasmic or exported pr | 0 | Unknown | SPII | 0,834 | LIPO(Sec/SPII) | X | 25,909 | 26,311 | 26,344 | 25,070 | 25,657 | 25,351 | + | 3 | 3 | 14,9 | 14,9 | 14,9 | 22,112 | 0 | 59,766 | 595480000 | 10 | 1,707892087 | 0,045174603 | 0,828356425 | 3,77065814 | RM25_0150 |
| lcl\|CP010341.1_prot_AJQ90775.1_1039 | RM25_RS05210 | Signal peptide peptidase SppA | 1 | Unknown | CYT | 0,478 | OTHER | X | 28,019 | 28,383 | 27,906 | 27,094 | 27,206 | 27,459 | + | 12 | 12 | 34,9 | 34,9 | 34,9 | 43,195 | 0 | 103,91 | 2311500000 | 43 | 2,03766625 | 0,040467532 | 0,849553426 | 4,72034445 | RM25_1058 |
| lcl\|CP010341.1_prot_AJQ90735.1_999 | RM25_RS05010 | Mycothiol acetyltransferase | 0 | CytoplasmicMembrane | CYT | 0,141 | OTHER | X | 23,602 | 23,460 | 23,296 | 22,659 | 22,592 | 22,489 | + | 2 | 2 | 8,7 | 8,7 | 8,7 | 38,345 | 0 | 15,574 | 55590000 | 3 | 3,002776376 | 0,029333333 | 0,872595469 | 8,624688258 | RM25_1018 |
| lcl\|CP010341.1_prot_AJQ90743.1_1007 | RM25_RS05050 | Adenine phosphoribosyltransferase | 0 | Cytoplasmic | CYT | 0,042 | OTHER | X | 23,682 | 23,798 | 23,294 | 22,738 | 22,310 | 22,602 | + | 2 | 2 | 20,7 | 20,7 | 20,7 | 19,052 | 0 | 25,014 | 91844000 | 10 | 2,204264085 | 0,045152542 | 1,04123497 | 5,261545654 | RM25_1026 |
| lcl\|CP010341.1_prot_AJQ89813.1_77 | RM25_RS00390 | Acetate kinase | 0 | Cytoplasmic | CYT | 0,061 | OTHER | C | 27,442 | 27,239 | 27,586 | 26,593 | 26,489 | 26,002 | + | 8 | 8 | 26,8 | 26,8 | 26,8 | 42,431 | 0 | 69,26 | 1622700000 | 38 | 2,156981569 | 0,042285714 | 1,061317444 | 5,103302383 | RM25_0078 |
| lcl\|CP010341.1_prot_AJQ90089.1_353 | RM25_RS01730 | TQXA domain-containing protein | 2 | Unknown | SPI | 0,911 | SP(Sec/SPI) | X | 25,372 | 25,542 | 24,980 | 24,159 | 23,963 | 24,536 | + | 5 | 5 | 12,4 | 12,4 | 12,4 | 47,513 | 0 | 32,787 | 303320000 | 13 | 1,98571836 | 0,042795181 | 1,079085668 | 4,560577347 | RM25_0357 |
| lcl\|CP010341.1_prot_AJQ91223.1_1487 | RM25_RS07475 | Hypothetical protein | 0 | Unknown | SPII | 0,427 | LIPO(Sec/SPII) | X | 23,187 | 23,865 | 23,319 | 22,568 | 21,943 | 22,537 | + | 2 | 2 | 13,8 | 13,8 | 13,8 | 19,346 | 0 | 20,724 | 84227000 | 8 | 1,724433383 | 0,044885246 | 1,107899984 | 3,814871899 | RM25_1514 |
| lcl\|CP010341.1_prot_AJQ91232.1_1496 | RM25_RS11930 | S-layer domain protein | 0 | Unknown | SPI | 0,431 | SP(Sec/SPI) | X | 27,514 | 28,013 | 27,451 | 26,532 | 26,184 | 26,788 | + | 4 | 4 | 24,7 | 24,7 | 24,7 | 2,335 | 0 | 61,992 | 1556700000 | 19 | 2,012095344 | 0,04015 | 1,157848358 | 4,641189902 | RM25_1523 |
| lcl\|CP010341.1_prot_AJQ90382.1_646 | RM25_RS03210 | Basic membrane protein | 0 | Unknown | CYT | 0,686 | OTHER | X | 26,294 | 26,686 | 25,949 | 24,668 | 25,472 | 25,293 | + | 2 | 2 | 9,9 | 9,9 | 9,9 | 31,261 | 0 | 28,311 | 611590000 | 16 | 1,644193958 | 0,049928058 | 1,165904363 | 3,603442847 | RM25_0655 |
| lcl\|CP010341.1_prot_AJQ89745.1_9 | RM25_RS00040 | DNA polymerase III, beta subunit | 0 | Cytoplasmic | CYT | 0,045 | OTHER | L | 25,427 | 24,810 | 25,614 | 24,139 | 23,741 | 24,445 | + | 7 | 7 | 18,1 | 18,1 | 18,1 | 41,525 | 0 | 50,476 | 304520000 | 14 | 1,683780438 | 0,048246154 | 1,175278982 | 3,706799628 | RM25_0009 |
| lcl\|CP010341.1_prot_AJQ89841.1_105 | RM25_RS00525 | Arginine/ornithine binding protein | 0 | Unknown | SPII | 0,803 | LIPO(Sec/SPII) | E | 28,007 | 28,196 | 27,756 | 26,772 | 26,572 | 26,982 | + | 5 | 5 | 18,2 | 18,2 | 18,2 | 31,534 | 0 | 57,177 | 1895100000 | 22 | 2,649149478 | 0,036133333 | 1,211041133 | 6,95717363 | RM25_0106 |
| lcl\|CP010341.1_prot_AJQ90176.1_440 | RM25_RS02165 | Phosphoribosylformylglycinamidine | 0 | Cytoplasmic | CYT | 0,053 | OTHER | F | 23,724 | 24,367 | 23,974 | 22,930 | 22,775 | 22,696 | + | 4 | 4 | 19,5 | 19,5 | 19,5 | 36,474 | 0 | 32,997 | 154360000 | 8 | 2,44241664 | 0,0384 | 1,221396764 | 6,119846492 | RM25_0444 |
| lcl\|CP010341.1_prot_AJQ91242.1_1506 | RM25_RS07565 | Dihydrodipicolinate synthase | 0 | Cytoplasmic | CYT | 0,055 | OTHER | E | 25,101 | 24,876 | 25,693 | 24,305 | 23,663 | 23,998 | + | 4 | 4 | 23,5 | 23,5 | 23,5 | 30,779 | 0 | 60,446 | 309280000 | 8 | 1,803338647 | 0,046446602 | 1,234202703 | 4,030461385 | RM25_1533 |
| lcl\|CP010341.1_prot_AJQ91472.1_1736 | RM25_RS08715 | Cystathionine gamma-synthase | 0 | Cytoplasmic | CYT | 0,084 | OTHER | E | 25,314 | 24,719 | 25,771 | 24,134 | 23,810 | 24,068 | + | 7 | 7 | 2,3 | 2,3 | 2,3 | 41,703 | 0 | 53,337 | 311900000 | 14 | 1,772386959 | 0,0452 | 1,264155706 | 3,944955285 | RM25_1768 |
| lcl\|CP010341.1_prot_AJQ90105.1_369 | RM25_RS01810 | Hypothetical protein | 0 | Unknown | SPII | 0,760 | LIPO(Sec/SPII) | X | 25,604 | 25,856 | 24,766 | 24,099 | 24,139 | 24,132 | + | 3 | 3 | 13,4 | 13,4 | 13,4 | 26,104 | 0 | 21,07 | 310860000 | 6 | 1,755988468 | 0,045403509 | 1,285439809 | 3,900148136 | RM25_0373 |
| lcl\|CP010341.1_prot_AJQ91339.1_1603 | RM25_RS08015 | Hypothetical protein | 0 | Cytoplasmic | CYT | 0,093 | OTHER | X | 24,423 | 24,496 | 24,415 | 23,630 | 22,416 | 23,282 | + | 3 | 3 | 31,3 | 31,3 | 31,3 | 14,903 | 0 | 20,113 | 159990000 | 5 | 1,677519526 | 0,049022556 | 1,335414251 | 3,690330994 | RM25_1630 |
| lcl\|CP010341.1_prot_AJQ90719.1_983 | RM25_RS04935 | Tryptophan synthase beta chain | 0 | Cytoplasmic | CYT | 0,084 | OTHER | E | 24,907 | 24,181 | 24,906 | 23,113 | 23,296 | 23,353 | + | 3 | 3 | 10,8 | 10,8 | 10,8 | 4,474 | 0 | 34,684 | 187840000 | 8 | 2,298331618 | 0,037283019 | 1,410654704 | 5,587969533 | RM25_1002 |
| lcl\|CP010341.1_prot_AJQ89842.1_106 | RM25_RS00530 | Arginine/ornithine binding protein | 0 | Unknown | SPII | 0,832 | LIPO(Sec/SPII) | E | 27,754 | 27,596 | 26,802 | 25,674 | 25,950 | 25,983 | + | 5 | 5 | 20,8 | 20,8 | 20,8 | 31,488 | 0 | 83,192 | 1207300000 | 29 | 2,088647652 | 0,042366197 | 1,51495107 | 4,881182966 | RM25_0107 |
| lcl\|CP010341.1_prot_AJQ90133.1_397 | RM25_RS01955 | Carboxylic ester hydrolase | 0 | Unknown | SPI | 0,780 | SP(Sec/SPI) | X | 25,353 | 25,610 | 25,147 | 23,987 | 23,592 | 23,726 | + | 4 | 4 | 13,3 | 13,3 | 13,3 | 46,469 | 0 | 30,176 | 267650000 | 11 | 3,081609836 | 0,024235294 | 1,602043152 | 9,042511956 | RM25_0401 |
| lcl\|CP010341.1_prot_AJQ91112.1_1376 | RM25_RS06915 | Peptide deformylase | 0 | Unknown | CYT | 0,129 | OTHER | J | 24,164 | 24,033 | 24,707 | 22,111 | 22,410 | 23,374 | + | 2 | 2 | 12,1 | 12,1 | 12,1 | 22,715 | 0 | 16,769 | 188300000 | 5 | 1,738734296 | 0,045448276 | 1,669389725 | 3,853366789 | RM25_1402 |
| lcl\|CP010341.1_prot_AJQ89896.1_160 | RM25_RS00810 | Shikimate dehydrogenase | 0 | Cytoplasmic | CYT | 0,041 | OTHER | E | 24,128 | 24,173 | 24,455 | 21,898 | 22,357 | 23,405 | + | 3 | 3 | 12,5 | 12,5 | 12,5 | 29,183 | 0 | 33,145 | 124600000 | 4 | 1,685568959 | 0,047658915 | 1,698677699 | 3,711512643 | RM25_0164 |
| lcl\|CP010341.1_prot_AJQ91504.1_1768 | RM25_RS08875 | Ferrochelatase | 0 | Cytoplasmic | CYT | 0,083 | OTHER | H | 23,490 | 24,001 | 24,053 | 22,803 | 21,960 | 21,502 | + | 5 | 5 | 18,5 | 18,5 | 18,5 | 37,455 | 0 | 33,577 | 105740000 | 12 | 1,854831666 | 0,046897959 | 1,759581248 | 4,175488558 | RM25_1800 |
| lcl\|CP010341.1_prot_AJQ91085.1_1349 | RM25_RS06775 | Putative GCN5-related N-acetyltran | 0 | Cytoplasmic | CYT | 0,097 | OTHER | K | 23,932 | 24,325 | 23,702 | 21,969 | 22,306 | 22,030 | + | 3 | 3 | 19,1 | 19,1 | 19,1 | 17,643 | 0 | 23,396 | 87048000 | 9 | 3,072349898 | 0,034222222 | 1,884539922 | 8,992486276 | RM25_1375 |
| lcl\|CP010341.1_prot_AJQ90682.1_946 | RM25_RS04730 | YhgE/Pip C-terminal domain protein | 6 | CytoplasmicMembrane | CYT | 0,829 | OTHER | S | 29,400 | 29,522 | 28,948 | 26,761 | 26,955 | 27,707 | + | 14 | 14 | 30,7 | 30,7 | 30,7 | 77,294 | 0 | 323,31 | 4071700000 | 58 | 2,50769186 | 0,044125 | 2,148890813 | 6,374361749 | RM25_0963 |
| lcl\|CP010341.1_prot_AJQ90978.1_1242 | RM25_RS06240 | Propanediol utilization protein PduJ | 0 | Unknown | CYT | 0,019 | OTHER | C | 25,686 | 25,832 | 25,630 | 23,971 | 23,190 | 23,187 | + | 3 | 3 | 46,7 | 46,7 | 46,7 | 92,237 | 0 | 36,787 | 292640000 | 13 | 2,973339805 | 0,028 | 2,266682307 | 8,473265957 | RM25_1264 |
| lcl\|CP010341.1_prot_AJQ91604.1_1868 | RM25_RS09370 | Mycothiol acetyltransferase | 0 | Cytoplasmic | CYT | 0,143 | OTHER | K | 24,400 | 24,698 | 24,255 | 22,704 | 22,206 | 21,491 | + | 4 | 4 | 15,7 | 15,7 | 15,7 | 34,301 | 0 | 26,801 | 210920000 | 6 | 2,456233048 | 0,039384615 | 2,317504247 | 6,172988909 | RM25_1900 |
| lcl\|CP010341.1_prot_AJQ91377.1_1641 | RM25_RS08205 | Hypothetical protein | 0 | Unknown | CYT | 0,302 | OTHER | X | 24,975 | 25,433 | 25,004 | 22,955 | 22,306 | 22,351 | + | 2 | 2 | 23,7 | 23,7 | 23,7 | 18,495 | 0 | 29,03 | 279470000 | 4 | 3,274006727 | 0,031692308 | 2,60031573 | 10,14216389 | RM25_1668 |
| lcl\|CP010341.1_prot_AJQ91651.1_1915 | RM25_RS09600 | Hypothetical protein | 1 | Unknown | TMH | 0,490 | OTHER | X | 25,847 | 26,482 | 25,544 | 23,207 | 22,963 | 22,830 | + | 5 | 5 | 20,6 | 20,6 | 20,6 | 37,068 | 0 | 250,76 | 387040000 | 16 | 3,240452987 | 0,029428571 | 2,957466761 | 9,941827487 | RM25_1947 |
| lcl\|CP010341.1_prot_AJQ90986.1_1250 | RM25_RS06280 | Propanediol utilization protein Pdu | 0 | Unknown | CYT | 0,029 | OTHER | C | 25,558 | 25,466 | 25,238 | 23,136 | 21,753 | 22,058 | + | 5 | 5 | 75,5 | 75,5 | 75,5 | 95,489 | 0 | 39,692 | 260440000 | 5 | 2,70901036 | 0,040148148 | 3,104549408 | 7,217421008 | RM25_1272 |
| lcl\|CP010341.1_prot_AJQ91667.1_1931 | RM25_RS09690 | Dimethylsulfoxide reductase, chain | 0 | CytoplasmicMembrane | CYT | 0,659 | OTHER | C | 26,647 | 26,541 | 26,655 | 23,092 | 22,961 | 22,956 | + | 5 | 5 | 31,3 | 31,3 | 31,3 | 23,523 | 0 | 40,049 | 520440000 | 13 | 6,405594506 | 0 | 3,611536662 | 62,48145308 | RM25_1963 |
| lcl\|CP010341.1_prot_AJQ90249.1_513 | RM25_RS02565 | 30S ribosomal protein S12 | 0 | Cytoplasmic | CYT | 0,710 | OTHER | J | 26,556 | 26,370 | 26,757 | 22,654 | 22,903 | 22,867 | + | 2 | 2 | 13,8 | 13,8 | 13,8 | 13,611 | 0 | 67,025 | 771680000 | 11 | 4,987032174 | 0 | 3,752916336 | 27,56421206 | RM25_0522 |
| lcl\|CP010341.1_prot_AJQ90072.1_336 | RM25_RS01660 | Hypothetical protein | 0 | Unknown | CYT | 0,438 | OTHER | X | 25,228 | 25,670 | 25,177 | 21,905 | 21,885 | 20,822 | + | 3 | 3 | 3,1 | 3,1 | 3,1 | 18,363 | 0 | 71,528 | 468780000 | 19 | 3,214108156 | 0,027466667 | 3,821423213 | 9,787133774 | RM25_0340 |
| lcl\|CP010341.1_prot_AJQ90246.1_510 | RM25_RS02550 | Hypothetical protein | 1 | Unknown | TMH | 0,932 | OTHER | X | 26,625 | 27,188 | 26,175 | 22,473 | 22,977 | 22,612 | + | 2 | 2 | 8,7 | 8,7 | 8,7 | 22,183 | 0 | 18,02 | 811460000 | 11 | 3,569320606 | 0,0235 | 3,975711187 | 12,07729298 | RM25_0519 |
| lcl\|CP010341.1_prot_AJQ91668.1_1932 | RM25_RS09695 | Anaerobic dimethyl sulfoxide reduc | 0 | Cytoplasmic | CYT | 0,425 | TAT(Tat/SPI) | C | 28,654 | 28,458 | 28,588 | 22,324 | 24,880 | 24,665 | + | 17 | 17 | 26,3 | 26,3 | 26,3 | 94,702 | 0 | 323,31 | 1965300000 | 56 | 2,306763822 | 0,038 | 4,610172272 | 5,618009619 | RM25_1964 |
