## Supplemental experiment for "Aerobic Adaptation and Metabolic Dynamics of *Propionibacterium freudenreichii* DSM 20271: Insights from Comparative Transcriptomics and Surfaceome Analysis"

**Supplemental material.** Results and Materials & methods for validation of RNAseq data with ddPCR.

To validate the differential expression findings obtained from RNA-seq data, the same samples were subjected to ddPCR analysis. This analysis included genes selected from both the logarithmic and stationary growth phases under different atmospheric conditions. Ten specific genes were chosen for ddPCR analysis.

The results of the ddPCR analysis demonstrated a strong Pearson correlation (>0.96) between the log2 fold change values obtained from ddPCR and those derived from RNA-seq analysis. This correlation was observed for both sampling points: I and III. These findings provide additional support and validation for the differential expression results obtained from the RNA-seq data.

ddPCR was conducted on the same samples utilized for RNA-seq to validate the differential expression results obtained from the RNA-seq data. Genes were selected from sampling point I and sampling point III under different atmospheric conditions. The ddPCR results for the selected 10 genes revealed a robust Pearson correlation (> 0.96) between the ddPCR log2 fold change and the RNA-seq log2 fold change results across the various growth phases. The ddPCR data were normalized using *dnaN* as a reference.

sampling point I

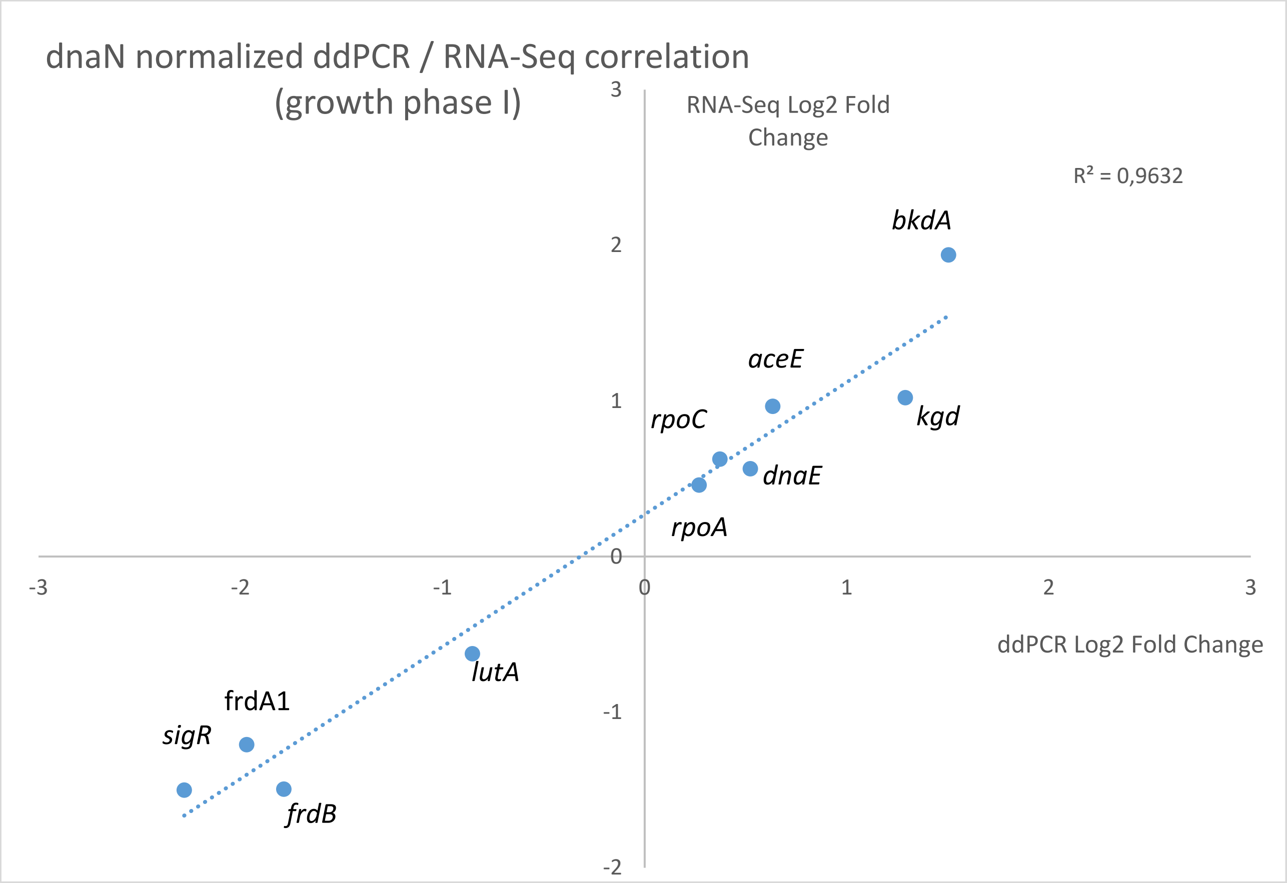


**Figure.** Comparison of relative expression changes (log2 fold change) for 10 genes obtained using droplet digital reverse transcription versus RNA sequencing from sampling point I.

sampling point III

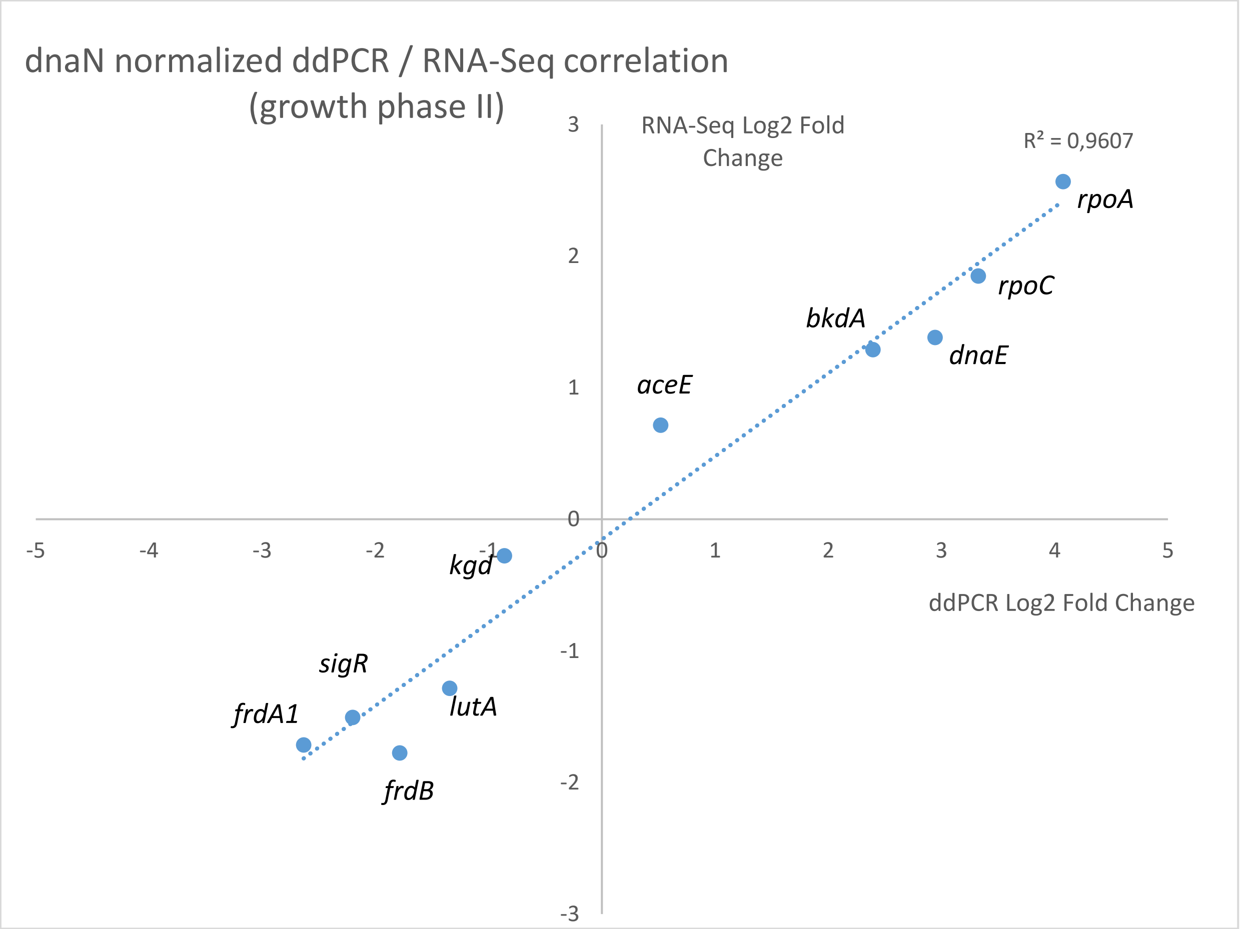


**Figure.** Comparison of relative expression changes (log2 fold change) for 10 genes obtained using droplet digital reverse transcription versus RNA sequencing from sampling point III.

**Table.** Primers used in ddPCR

| NCBI locus tag | Old locus tag | Product feature/name | Gene name | Forward primer (5'-3') | Reverse primer (5'-3') | Amplicon (bp) |
|---|---|---|---|---|---|---|
| RM25_RS06155 | RM25_1247 | Succinate dehydrogenase flavoprotein subunit | *frdA1* | gatgcttgacctgatcgtca | cggttgcattcgagttcttg | 167 |
| RM25_RS08740 | RM25_1773 | Lactate utilization protein A | *lutA* | tggttcgctcgtatgtgaag | ggtgatgtcgaagctgatct | 170 |
| RM25_RS06150 | RM25_1246 | Succinate dehydrogenase/fumarate reductase iron-sulfur subunit | *frdB* | attggatatctggcgccagg | gccctgttcgatgatctggt | 133 |
| RM25_RS06045 | RM25_1225 | Pyruvate dehydrogenase E1 component | *aceE* | gactaccagacgatgaaggc | gatcttcaggtagtcgtggc | 147 |
| RM25_RS05695 | RM25_1154 | RNA polymerase sigma factor | *sigR* | atgtcgaggactggcagttg | caggactgcctcacggaatt | 149 |
| RM25_RS02560 | RM25_0521 | DNA-directed RNA polymerase, beta' subunit | *rpoC* | cccgagacgatcaactaccg | acattggaccgtgtgacctc | 170 |
| RM25_RS02790 | RM25_0568 | DNA-directed RNA polymerase, alpha subunit | *rpoA* | aaacccgcactgacttcgat | caactcaaccagcgtcttgc | 104 |
| RM25_RS05675 | RM25_1150 | 2-oxoglutarate decarboxylase | *kgd* | cttccacaaggacgtcgtca | catcctccaccgagatgtcg | 176 |
| RM25_RS04965 | RM25_1008 | DNA polymerase III, alpha subunit | *dnaE* | gtgaagcgcgttgtcgatac | acgaggttgagcagcttctc | 107 |
| RM25_RS00040 | RM25_0009 | DNA polymerase III, beta subunit | *dnaN* | acgccctatgtgcagttctc | acatgcttgtagtcggcctc | 101 |
| RM25_RS01015 | RM25_0205 | Pyruvate dehydrogenase E1 component, alpha subunit | *bkdA* | tgatgaagccccaggacatg | tcatgtaatagggcgccacc | 156 |

**Materials and methods**

Gene expression levels of genes *frdA, lutA, frdB, ace, sigR, rpoC, rpoA, kgd, dnaE* and *bkdA* were verified with droplet digital PCR (ddPCR). Primers were designed with Primer3Plus (1).

dd-PCR mixes included 1x Q200 ddPCR EvaGreen Supermix (Bio-Rad), 200 nm forward and reverse primers and 5µL of 1:200 diluted complementary DNA (cDNA) in a total volume of 22µl. cDNA was synthesized using 400 ng of purified single stranded RNA with iScript™ Advanced cDNA Synthesis Kit (Bio-Rad). Reaction contained 1 x iScript™ Advanced Reaction Mix and iScript™ Advanced Reverse Transcriptase in reaction volume of 22 µL and reaction time 30 min at + 42 °C. Droplets were generated using an automated droplet generator (Bio-Rad) and amplification was performed in a Veriti thermal cycler (Applied Biosystems) with reaction volume of 40 µL. Thermal cycling conditions with ramp rates of 50 % were 95 °C for 5 min, 40 cycles of +95 °C for 30 s and +60 °C for 1 min, after cycles +4 °C for 5 min and +90 °C for 5 min, and then +4 °C until fluorescence measurement by a QX dropletdroplet reader (Bio-Rad).

Results were analyzed using QuantaSoft software (version 1.7.4.0917, Bio-Rad) using mainly automatic thresholds separation of PCR positive and PCR negative droplets.  The concentration of the target genes was normalized by a gene dnaN which expression was found to be mostly constant across growth conditions and growth phases.


1. Untergasser A, Nijveen H, Rao X, Bisseling T, Geurts R, Leunissen JA. 2007. Primer3Plus, an enhanced web interface to Primer3. Nucleic Acids Res 35:W71-4.
